# Deep indel mutagenesis reveals the regulatory and modulatory architecture of alternative exon splicing

**DOI:** 10.1101/2024.04.21.590414

**Authors:** Pablo Baeza-Centurion, Belén Miñana, Andre J. Faure, Mike Thompson, Sophie Bonnal, Gioia Quarantani, Ben Lehner, Juan Valcárcel

**Author notes:** These authors contributed equally to this work (Pablo Baeza-Centurion, Belén Miñana). These authors jointly supervised this work (Ben Lehner, Juan Valcárcel). Corresponding author: (BL), (JV).

## Abstract

Altered splicing is a frequent mechanism by which genetic variants cause disease and antisense oligonucleotides (AONs) that target pre-mRNA splicing have been approved as therapeutics for multiple pathologies including patient-customized treatments for rare diseases. However, the regulatory architecture of human exons remains poorly understood and AON discovery is currently slow and expensive, limiting the wider adoption of the approach. Here we show that that systematic deletion scans –which can be made experimentally at very low cost – provide an efficient strategy to chart the regulatory landscape of human exons and to rapidly identify effective splicing-modulating oligonucleotides in a fully quantitative manner. Our results suggest a mechanism for the evolutionary origins of unusually short microexons and the repression of transmembrane domain-encoding exons, and reveal a checkerboard architecture of sequential enhancers and silencers in a model alternative exon. Accurate prediction of the effects of deletions using deep learning provides a resource, DANGO, that maps the splicing regulatory landscape of all human exons and predicts effective splicing-altering antisense oligonucleotides genome-wide.

## Introduction

Pre-mRNA splicing is the process by which introns are removed from transcripts and exons are joined together to form mature mRNAs that are then exported to the cytoplasm and translated^1^. Altered splicing is an important mechanism by which genetic variants cause disease^2–4^. Multiplex assays of variant effects (MAVEs) have revealed that random nucleotide substitutions in exons frequently affect splicing, with 60-70% of substitutions in over 90% of positions in alternatively-spliced exons^5–7^ and 5% of substitutions in constitutively-spliced exons^8^ altering exon inclusion. Comprehensive testing has also shown that ∼10% of disease-causing missense variants, as well as 3% of common exonic substitutions, affect splicing^9,10^.

The impact of genetic variation beyond substitutions on splicing have been far less studied. Insertions and deletions (indels^11,12^) are abundant variants evident in 24% of Mendelian diseases^13^, and disease-causing indels are enriched close to splice sites^14^. However, the effects of indels on splicing have not been systematically tested.

The frequent disruption of splicing in human disease has led to extensive efforts to therapeutically modulate splicing^15^. In particular, antisense oligonucleotides (AONs) that modulate splicing have been approved as therapies for spinal muscular atrophy^16^ and Duchenne muscular dystrophy^17^. Indeed AONs – because of their programmable sequence specificity – may represent a general strategy to therapeutically modulate many splicing changes^18^. However, predicting the impact of AONs on splicing is very challenging: AONs need to be long to achieve sequence specificity (typically 18 to 21 nts) whereas splicing regulatory elements are short and very poorly mapped genome-wide. Parameters that influence AON efficacy include length, proximity to splice sites and binding energy^19–23^. In practice, however, identifying effective AONs requires laborious testing of many different designs.

Here we use deep indel mutagenesis (DIM) to comprehensively quantify for the first time the impact of all possible deletions and all small insertions on the splicing of a human exon, FAS exon 6. Exon 6 encodes a transmembrane helix of the FAS/CD95 receptor. Inclusion of this exon generates a pro-apoptotic receptor whereas exon skipping produces an anti-apototic soluble inhibitor^24–27^. A variant in exon 6 that causes exon 6 skipping causes autoimmune lymphoproliferative syndrome (ALPS)^28^. Our data show that insertions and deletions frequently disrupt splicing, with deletions in particular representing an efficient experimental strategy to map the regulatory architecture of the exon. The deletion scan also defines the minimum length of an exon and reveals a novel repressive regulatory mechanism whereby cryptic sequences within an exon are recognised by a core component of the spliceosome (U2 snRNP) that normally recognises intronic branch points. We show that this recognition of cryptic exonic elements can be a mechanism for the evolutionary birth of unusually short microexons^29^. Using our data, we show that deep learning methods accurately predict the effects of indels on splicing and that deletions accurately predict the splice-altering effects of AONs. Finally, we provide a genome-wide resource, DANGO, that charts splicing regulatory landscapes and splice-altering AONs across the human exome.

## Results

### Deep indel mutagenesis of a human alternatively spliced exon

To quantify in parallel how single nucleotide (nt) substitutions, deletions and insertions affect alternative splicing of a human exon, we designed a library containing all single-nt substitutions in the 63 nt-long FAS exon 6 (n = 189), as well as all deletions ranging in length from 1 to 60 nts (n = 2010), all possible 1-, 2-, and 3-nt insertions (n = 5208), and a random selection of 585 4-nt-long insertions (**figure 1A**). We cloned this library into a plasmid minigene vector spanning FAS exons 5–7, transfected it into HEK293 cells, isolated RNA 48 h post-transfection and quantified the inclusion of each variant by counting how often it is present in the final exon inclusion product compared to every other variant in the library by deep sequencing of reverse transcription (RT)-PCR products (**figure 1B**). The resulting enrichment scores allow the percent-spliced-in (PSI) value of each variant to be measured. Our results were highly reproducible across nine experimental replicates (Pearson’s r between 0.97 and 0.98 for all pairs of replicates, **figure S1**) and our estimated PSI values were very well correlated with PSI values determined by RT-PCR for 40 individual, independently transfected, mutant minigenes, which included indel and substitution mutants (Spearman’s rho = 0.95, **figure 1C**).

**Figure 1.**
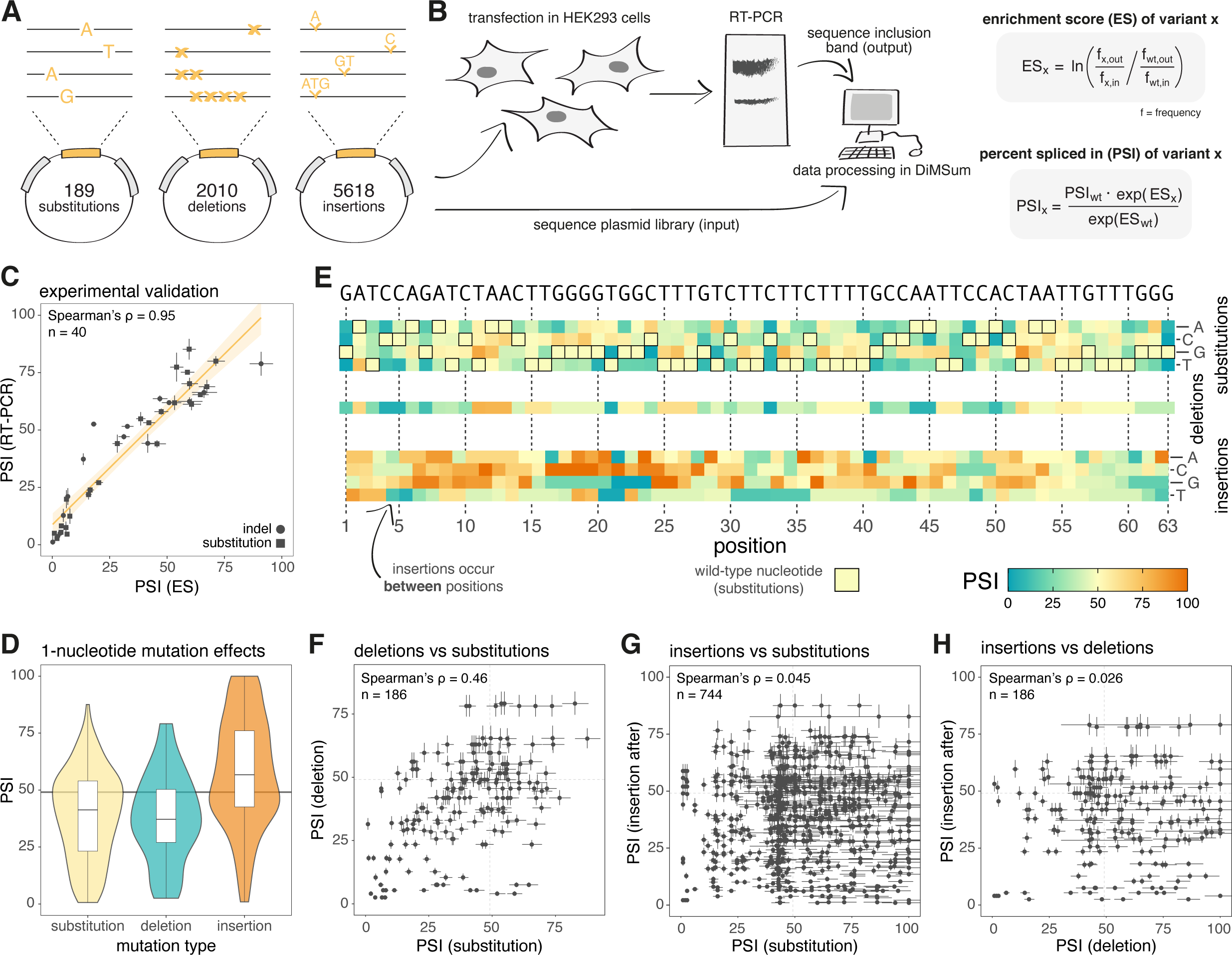
Deep indel mutagenesis of FAS exon 6. A. Plasmid library design. B. Experimental protocol of our massively-parallel splicing assay (MPSA). C. Correlation between PSI values from individual transfections and MPSA-derived PSI values. D. Distribution of PSI values for variants with 1-nucleotide (nt) mutations. E. Heatmaps displaying inclusion levels of 1-nt substitutions, deletions, and insertions. F. Correlation between PSI values for 1-nt deletions and substitutions at the same position. G. Correlation between PSI values for 1-nt insertions after a given position and substitutions at that position. H. Correlation between PSI values for 1-nt insertions after a given position and 1-nt deletions at the same position

### The effects of single-nt substitutions, insertions and deletions on exon inclusion

We quantified the effects of all single-nt substitutions (n = 189), insertions (n = 187) and deletions (n = 63) on the inclusion of FAS exon 6 (**figure 1D-E**). The results revealed that nearly the entire range of inclusion values can be obtained as a consequence of at least one of the three classes of variant. Regardless of the type of mutation, just under two thirds affected splicing by more than 10 PSI units (**figure 1D-E**): 61.9% of substitutions changed the inclusion of FAS exon 6 by more than 10 PSI units (mean absolute ΔPSI = 17.5 PSI units; consistent with previous data^5^), similar to 58.7% of single-nt deletions (mean absolute ΔPSI = 15.5 PSI units) and 61.3% of single-nt insertions (mean absolute ΔPSI = 19.7 PSI units).

Substitutions have a mode near the wild type (WT) PSI (49.1%) and they more often promote skipping than inclusion, with 47.6% decreasing and 14.3% increasing inclusion by more than 10 PSI units. Similarly, deletions tend to promote skipping, with 15.9% of single-nt deletions promoting inclusion and 42.9% promoting skipping by more than 10 PSI units. In contrast, insertions more frequently promote inclusion, with 41.1% increasing the inclusion of FAS exon 6 and only 20.2% decreasing inclusion by more than 10 PSI units (**figure 1E**).

We next compared the effects of different mutation types in the same position along the exon. The effects of single-nt substitutions and deletions in the same positions correlate moderately well (Spearman’s rho = 0.46, **figure 1F**), consistent with some of these variants disrupting existing regulatory elements and with positive regulatory elements covering a larger proportion of the exon than negative elements (**figure 1D-E**). However, the effects of substitutions and deletions correlate poorly with the effects of insertions before or after the substituted/deleted position (**figure 1G-H**, **figure S2, figure S3**). This suggests that insertions often affect splicing by a different mechanism, for example by the creation of new regulatory sequences (see below).

### The effects of short multi-nt deletions and insertions on exon inclusion

We next analysed the influence of short multi-nt deletions (ranging from 2 to 9 nts) and insertions (either 2,3 or 4 nts) on exon inclusion. Short multi-nt deletions had a similar effect on exon inclusion as single-nt deletions. Thus, 64.5% of 2-nt deletions affected splicing by more than 10 PSI units (mean absolute ΔPSI = 17.0), as well as 62.3% of 3-nt deletions (mean absolute ΔPSI = 17.9) and 73.3% of 4-nt deletions (mean absolute ΔPSI = 20.8).

Considering all short deletions up to 10 nts long, 66.0% altered inclusion by more than 10 PSI units (mean absolute ΔPSI = 18.7). Short multi-nt deletions longer than 2nt more frequently promote skipping, with 30.6% of 2-nt deletions, 39.3% of 3-nt deletions, 43.3% of 4-nt deletions and 45.8% of all deletions spanning 10 or fewer nts decreasing exon inclusion by more than 10 PSI units. In contrast, 33.9% of 2-nt deletions, 23.0% of 3-nt deletions, 30% of 4-nt deletions and 20.2% of all deletions up to 10 nts long increase inclusion by more than 10 PSI units.

Longer multi-nt deletions are particularly effective at revealing splicing regulatory elements (**figure 2A**). Thus, while 1-4 nt deletions at the 5’ end of the exon showed multiple, seemingly contradictory effects, longer deletions delineated ∼8 5’ terminal nts whose collective deletion strongly promotes exon skipping. Similarly, deletions that cover exon positions 9 to 18 consistently increase exon inclusion. Interestingly, deletions that partially overlap this region (on either side) have the inverse effect and promote exon skipping. Systematic deletion scans thus delineate consistent discrete regulatory elements (enhancers and silencers), often adjacent to each other, and reveal patterns such as alternating elements with antagonistic effects (**figure 2A**).

**Figure 2.**
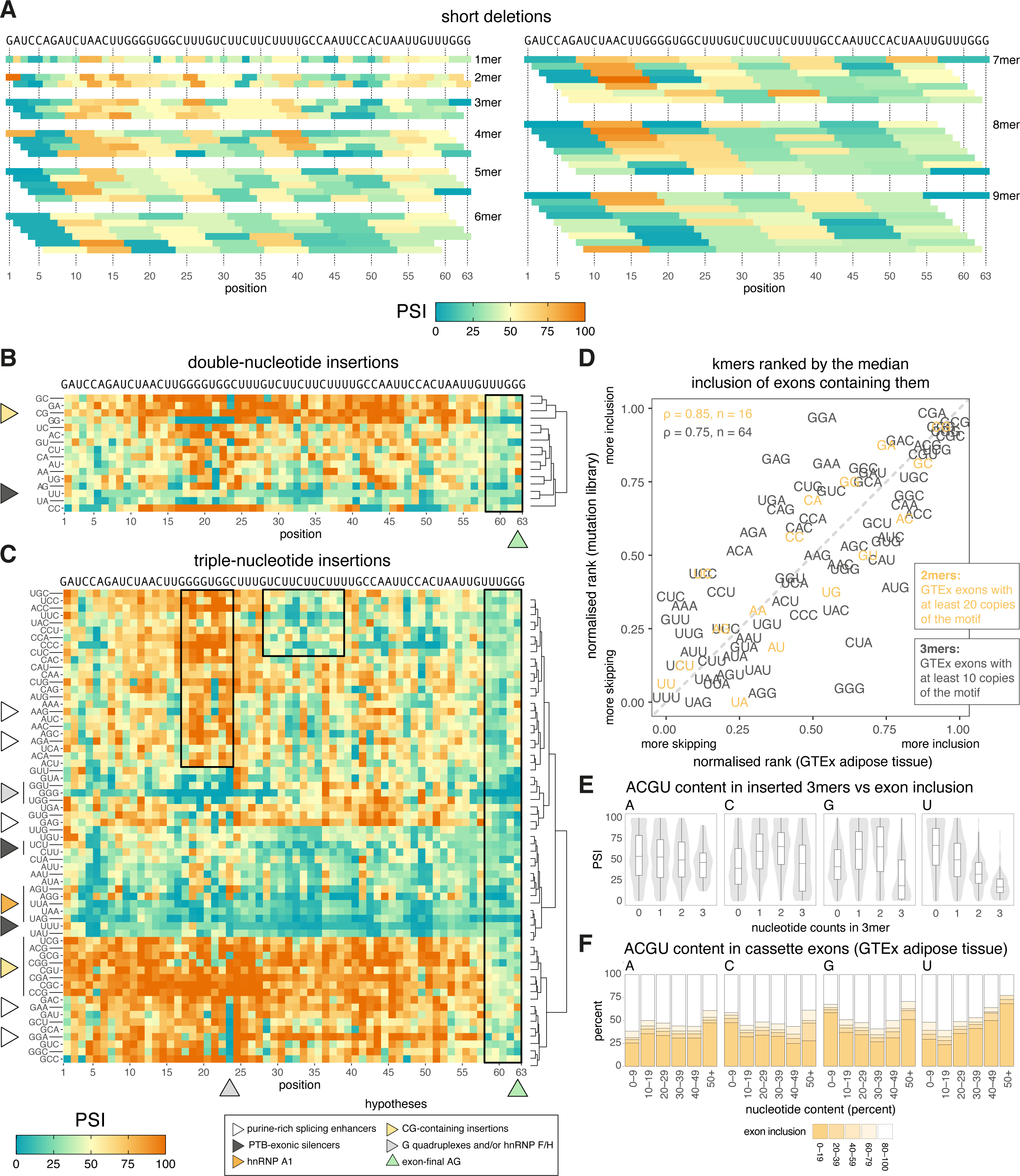
The effects of short indels on the inclusion of FAS exon 6. A. Heatmap showing inclusion values for short deletions (1-9 nt in length). B. Heatmap showing inclusion values for 2-nt insertions. C. Heatmap showing inclusion values for 3-nt insertions. D. Correlation between 2 (or 3) nt sequences ranked by the median PSI of exon variants in our library, with this sequence inserted, and their rank based on median PSI of exons containing 30 (or 20) such 2mers (or 3mers) in their sequence (GTEx adipose tissue). E. Distribution of PSI values in exons with a 3-nt insertion compared to the frequency of each nucleotide in the insertion. F. Distribution of PSI values in cassette exons (GTEx adipose tissue) relative to the percentage of each nucleotide present in the exon sequence.

Short multi-nt insertions had a stronger effect on FAS exon 6 inclusion compared to single-nt insertions. Thus, 74.5% of 2-nt insertions and 77.5% of 3-nt insertions changed inclusion by more than 10 PSI units (mean absolute ΔPSI = 23.9 and ΔPSI = 24.2, respectively). Our library also contained a random selection of 585 4-nt insertions, of which 88.6% changed splicing by more than 10 PSI units (mean absolute ΔPSI = 31.2). Similar to single-nt insertions, but in contrast with short multi-nt deletions, short multi-nt insertions tended to promote inclusion rather than skipping: 44.6% of 2-nt insertions increased exon inclusion by more than 10 PSI units (compared to 30.0% which decreased inclusion by this same amount), as well as 42.4% of 3-nt insertions (35.1% for skipping) and 57.5% of 4-nt insertions (31.2% for skipping).

In contrast to the regulatory landscape emerging from deletion analyses (**figure 2A**), double or triple nt insertions tended to show autonomous effects that were strongly influenced by the nature of the inserted nts (**figure 2B-C**, **figure S4**). Thus, insertion of GC, CG or GA dinucleotides resulted in general increases in exon inclusion, almost independently of the position of the insertion, with the prominent exception of the exon 3’ end. In contrast, insertion of GG or CC showed markedly different effects depending on the site of insertion.

The effects of triple nt insertions were even more idiosyncratic (**figure 2C**). For example, all CG-containing triplets enhanced exon inclusion in nearly all positions (as was the case of GC, CG or GA dinucleotides), with the notable exception of the five 3’ nts of the exon. Almost any insertion in this region (except those positioning an A at the 3’ end of the exon) led to enhanced exon skipping (black vertical rectangles on the right of panels **2B-C**), suggesting that it harbours a strong enhancer sequence very sensitive to insertions or deletions, possibly important for activation of the adjacent 5’ splice site. Effects similar to those of CG-containing triplets were observed upon insertion of a variety of GC / GA / GG-containing triplets (lower rows of **figure 2C**); some of these effects might be related to enrichment in purine residues which, together with other purines present in the insertion site, could function as purine-rich exonic enhancers, a well-known class of exonic regulatory elements^30^. Indeed, these effects are also observed for other purine-rich triplets such as GAG or AAG, albeit not for all (e.g. AGG or GGG). Triplets containing AU/UA dinucleotides (e.g. UAG, UAA, UUA, AUU, CUA) promote skipping when inserted at most exonic positions (cluster of blue colour in mid-low rows of panel **2C**), which might be explained by enhanced binding of hnRNP proteins such as hnRNP A1^31^, which are known to mediate effects of exonic silencers^32^. Pyrimidine-rich triplets (e.g. UCC, UUC, CUU), which could provide/reinforce binding sites for other repressive hnRNPs such as PTB/hnRNP I, have however very regional effects, promoting skipping mainly in already pyrimidine-rich regions like the previously described PTB silencer located in positions 28-39 (middle upper black rectangle in panel **2C**). A variety of cytosine-containing triplets systematically promote inclusion when introduced between exon positions 17-24 (upper left black rectangle in panel **2C**), which is a G-rich sequence, while insertion of 3 additional Gs in this region strongly inhibits inclusion, suggesting that a G-rich silencer in this region, possibly forming G-quadruplexes recognized by hnRNP F/H factors^33^, is disrupted by C-containing triplets. The latter effects could be in part linked to the creation of CG dinucleotides, which as discussed above display strong enhancing effects.

We next evaluated the extent to which the effects of 2– and 3-nt insertions in FAS exon 6 predict the association between 2– and 3-nt kmer content and exon inclusion transcriptome-wide. Strikingly, there is a strong positive correlation (Spearman’s rho between 0.72 and 0.87 for 2-mers, between 0.69 and 0.79 for 3-mers) between the ΔPSI induced by kmer insertions in our library and the PSI of exons containing at least 20 (for 2-mers) or 10 (for 3-mers) such kmers in the GTEx database, either in adipose tissue (**figure 2D)** or in all GTEx tissues (**figure S5**). Furthermore, the relationship between the content of each individual nucleotide in the inserted 3-mers and the PSI of the mutated exon (**figure 2E**) resembles the relationship between the content of each individual nucleotide in cassette exons and their PSI (**figure 2F**). For example, increasing the number of uridines in an inserted triplet correlates with increased exon skipping (**figure 2E**), similarly to how a larger uridine content is associated with increased skipping in exons throughout the genome (results for GTEx adipose tissue are shown in **figure 2F**, results for all GTEx tissues are shown in **figures S6-S9**). These results argue that the effects of systematic analysis of insertion mutations observed in our experiment for one individual model exon have captured sequence features relevant for the inclusion of alternatively-spliced exons genome-wide.

### Systematic deletion scans reveal the checkerboard regulatory landscape of a human exon

Deletions are more likely to cause loss-of-function molecular effects than insertions that are more likely to create new regulatory elements. Plotting the effects on inclusion of all possible deletions from size 1 to 60 nts provides a comprehensive deletion map of FAS exon 6 (**figure 5B**, bottom triangle). The map reveals several regions whose deletion leads to enhanced inclusion (exonic silencers, red areas) or enhanced skipping (exonic enhancers, more green areas), arranged in an alternating ‘checkerboard’ pattern and covering most of the exon length. These regulatory elements can also be visualised by plotting the effects on exon inclusion of 1 to 6 nt-long deletions versus the position of the deletion along the exon (**figure 5C**, left panel) and using local polynomial regression (LOESS) to identify sequence ‘blocks’ where deletions promote more inclusion or more skipping. These regulatory blocks recapitulate well-known regulatory elements in FAS exon 6, namely EWS binding exonic enhancers at positions 15-23 and 55-63^34^, a PTB-binding silencer in positions 25-40^35^, and an SRSF6 enhancer at positions 40-45^36^, as well as other elements that have not been fully characterised to date, in particular a very active silencer located between positions 8 and 13. Interestingly, deletions affecting nt C33 show different effects compared to deletions affecting only its flanking nts, and these differential effects are consistent across a wide range of deletion mutant lengths (blue square in the lower middle part of panel **5B**). Mutation of C33 has been reported to cause FAS exon 6 skipping in patients with ALPS^28^ and our deletion scans further emphasise the particular effects of deletions containing this nucleotide, demarcating a splicing enhancer embedded within two silencers previously associated with PTB-mediated repression^35^. Finally, a general observation is that deletions above a certain length tend to induce strong skipping (lower left triangle in panel **5B**), defining the exon length below which exon definition is likely failing (40 – 50 nts)^37,38^ (but see below).

Collectively, our results suggest that systematic deletion mutagenesis is a particularly informative experimental design to rapidly identify splicing regulatory elements throughout an exon.

### Small insertions create novel microexons by activating cryptic splice sites within FAS exon 6

We next analysed the relationship between exon length and inclusion in our library. Specifically, we asked, at 1 nt resolution, how short an exon can be while still being recognised by the splicing machinery. There is no clear dependence of exon inclusion on length for exons longer than 50 nts (up to 13 nt-long deletions). Then, inclusion gradually decreases with increasing deletion length (i.e. shorter exons are less included), with almost no exons shorter than 30 nts showing detectable levels of inclusion in our library (**figure 3A**). Consistent with this – and previous large deletions in constitutive exons^37–39^ – exons shorter than 30 nts are more likely to be skipped genome-wide compared to longer exons (results for GTEx adipose tissue are shown in **figure 3B**, results for all GTEx tissues are shown in **figure S10**).

**Figure 3.**
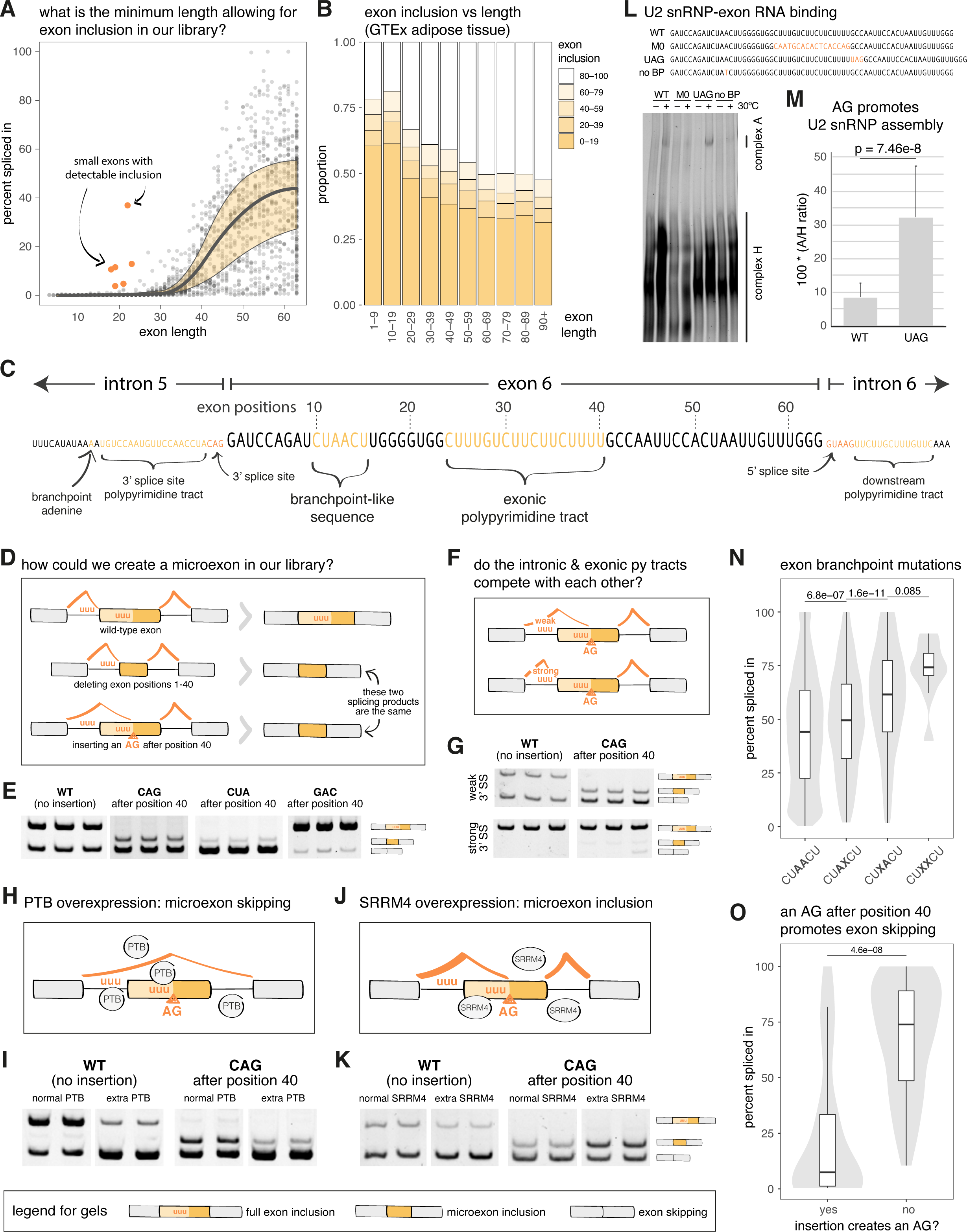
Deep indel mutagenesis of FAS exon 6 reveals the origin of novel microexons. A. Relationship between exon length and inclusion (black curve represents a constrained B-spline fit to rolling median PSI values, and the yellow shaded area indicates the rolling interquartile range of PSI values). B. Distribution of cassette exon inclusion values versus exon length (GTEx adipose tissue). C. Sequence of FAS exon 6 and surrounding intronic sequences. D. Hypothetical mechanisms explaining how our experimental assay could result in the detection of a microexon in our mutant library. E. RT-PCR analysis of FAS exon 6 inclusion for the WT exonic sequence, two variants with insertions that introduce an AG after exon position 40 and one variant with an insertion after exon position 40 that does not introduce an AG dinucleotide. F. Illustration showing the impact of a weak or strong 3’ splice site before FAS exon 6 on the recognition, by the splicing machinery, of the exonic 3’ splice-site-like sequence in the central region of the exon (followed by an AG insertion). G. RT-PCR analysis of FAS exon 6 inclusion in the presence of a weak or strong 3’ splice site, for the WT exon and an exon with an insertion introducing an AG dinucleotide after exon position 40. H-I. RT-PCR analysis of FAS exon 6 inclusion upon overexpression of PTB (WT sequence and variant with an insertion introducing an AG dinucleotide after exon position 40). J-K. RT-PCR analysis of FAS exon 6 inclusion upon overexpression of SRRM4 (WT sequence and variant with an insertion introducing an AG dinucleotide after exon position 40). L. Spliceosome assembly assays using the indicated fluorescently-labeled RNAs (wild type or mutant Fas exon 6 sequences) and HeLa nuclear extracts. The position of complexes assembling U2 snRNP (A complex) and hnRNP proteins (H complex) are indicated. M. Quantification of the ratio between A and H complexes for wild type (WT) and 3’ ss-containing (AUG) mutant RNAs as in L. P value corresponds to a 2-tailed t-test from 20 replicates with WT RNA, and 19 replicates with UAG-containing RNA. N. Distributions of PSI values for double-nucleotide substitutions targeting (i) neither of the putative exonic branchpoint adenines; (ii) the second putative exonic branchpoint adenine as well as another nucleotide in the exon; (iii) the first putative exonic branchpoint adenine as well as another nucleotide in the exon; (iv) both putative exonic branchpoint adenines. P values correspond to 2-tailed Wilcoxon tests with 16230 data points in the CTAACT group, 544 data points in the CTAXCT group, 549 data points in the CTXACT group, and 9 datapoints in the CTXXCT group. Boxplot boxes represent the median, the interquartile range, and the boxplot whiskers extend up to 1.5 times the interquartile range. O. Distribution of PSI values for insertions that either introduce (left) or do not introduce (right) an AG dinucleotide after exonic position 40. P value corresponds to a 2-tailed Wilcoxon test with 60 data points in the “no” group and 29 data points in the “yes” group.Boxplot boxes represent the median, the interquartile range, and the boxplot whiskers extend up to 1.5 times the interquartile range.

Exons shorter than 27 nts are detected in multicellular animal but are categorised as a special class – microexons – whose recognition requires a dedicated set of regulatory sequences and factors (such as SRRM3/4) that enable their inclusion in specific tissues (e.g. the brain or endocrine pancreas^40,41^). However, we observed that a group of very short exons (microexons, length-wise) from our library were detectably included (**figure 3A**).

These microexons appeared to correspond to large deletions at the 5’ or 3’ ends of FAS exon 6 (**figure S11A**). However, these deletion mutants showed no evidence of exon inclusion when tested individually (**figure S11B**). We realised that these apparently contradictory results could be explained if the clones detected by deep sequencing of the exon inclusion amplicon product were not the result of splicing of very short exons flanked by FAS exon 6 splice sites, but rather result from the use of cryptic splice sites within exon 6 that have been activated by another mutation that is no longer present in the spliced-in exon sequence. The central part of FAS exon 6 (positions 24 – 40) contains a pyrimidine-rich tract that resembles the polypyrimidine (Py)-tracts that precede 3’ splice sites, and nts in positions 10-15 contain a sequence that – strikingly – matches a branch point sequence^42^ (**figure 3C**). We reasoned that, if an AG-containing kmer were to be introduced after the pyrimidine (Py)-rich segment, this would result in a 3’ splice site-like sequence arrangement that, if recognised by the spliceosome, could create a novel microexon spanning from this new 3’ splice site to the 3’ end of exon 6 (**figure 3D**). To test this possibility, we inserted AG-creating triplets (like CAG or CUA –as the next nt is a G-) after the Py-tract in the FAS exon 6 minigene and observed that exons corresponding to the final part of the exon were included to some extent in the mature RNA (**figure 3E**). In contrast, insertion of GAC, that does not create a 3’ splice site, does not result in activation of a shorter exon, but rather enhances the inclusion of its full-length version (**figure 3E**).

Interestingly, the exonic Py-tract is longer and more uridine-rich than the Py-tract associated with the natural 3’ splice site of intron 5 (**figure 3C**). To assess whether the interplay between these 3’ splice sites plays a role in regulation, we strengthened the Py-tract of the natural 3’ splice site (**figure 3F**). In the presence of this mutation, inclusion of the full-length exon was enhanced in the wild type minigene and activation of the cryptic 3’ splice site in the AG-containing construct was greatly reduced (**figure 3G**).

Previous work showed that the pyrimidine-rich sequence within FAS exon 6 functions as a silencer when bound by PTB^43^ (**figure 3H**). As expected, overexpression of PTB led to skipping of the wild type exon (**figure 3I**, left panel) and also to reduced inclusion of the shorter exon in the AG-containing construct (**figure 3I**, right panel), most likely due to direct competition between PTB and the Py-tract-binding splicing factor U2AF^43,44^. Finally, we tested whether over-expression of SRRM4, which triggers inclusion of microexons in neurons^40^, has any effect on inclusion of the shorter version of FAS exon 6 (**figure 3J**). Surprisingly, SRRM4 overexpression enhanced inclusion of the shorter exon (**figure 3K**) despite this exon not being flanked by *cis*-acting sequences typically required for the inclusion of microexons (e.g. intronic UGC motifs^45^). Interestingly, SRRM4 reduced, rather than enhanced, inclusion of the wild type full length exon 6 (**figure 3K**). These results show that the short exon activated by a cryptic 3’ splice site in exon 6 is not only recognized by the splicing machinery but can be subject to splicing regulation by mechanisms similar to those operating on natural microexons.

Our results thus far do not account for the detection of short exons spanning the first third of FAS exon 6 (**figure 3C**). However, inserting sequences that mimic a 5’ splice site (e.g. UAA) after exon positions 18 or 19 induced the accumulation of spliced products containing these sequences (**figure S11C**). Interestingly, the Py-tract in the central region of FAS exon 6 is similar to a Py-tract found downstream of FAS exon 6 5’ splice site (**figure 3C**), which is recognized by the protein TIA1 and enhances 5’ splice site recognition by U1 snRNP^35,46^. The Py-tract in the central region of FAS exon 6 could therefore enhance recognition of upstream 5’ splice sites generated by exonic mutations.

These findings reveal that FAS exon 6 contains 3’ and 5’-like sequences that can be activated by simple mutations to function as *bona fide* splice sites, promoting the inclusion of very short exons even in the absence of regulatory sequence elements known to be involved in the activation of microexons. The evolutionary birth of new microexons is therefore likely to be simpler and more frequent than previously appreciated.

### Exonic binding of U2 snRNP promotes FAS exon 6 skipping

The presence of a relatively strong Py-tract preceded by a near-consensus branch point sequence within exon 6 (**figure 3C**), and the inclusion of a shorter version of the exon when a mutation creates a functional 3’ splice site AG downstream of the Py-tract, opened the possibility that 3’ splice site recognizing factors assemble on FAS exon 6 (**figure 3L-M**).

To directly assess whether U2 snRNP, the key ribonucleoprotein complex involved 3’ splice site recognition, can assemble on FAS exon 6 sequences, we incubated *in vitro* transcribed FAS exon 6 (wild type and mutants, all lacking the flanking splice sites) with HeLa nuclear extracts and measured the interaction by native gel electrophoresis. The results indicated that U2 snRNP can indeed assemble (complex A) on FAS exon 6, an interaction that was decreased upon mutation of the Py-tract or branch point sequences and enhanced upon introduction of an AG dinucleotide (**figure 3L-M** and **figure S12**).

It is conceivable that U2 snRNP assembly on the wild type exon, in the absence of a 3’ splice site, competes with recognition of the 3’ splice site of intron 5 by the splicing machinery and this contributes to modulate the levels of exon 6 inclusion. To test this possible mechanism, we took advantage of our saturation mutagenesis results. We observed that exonic variants with an intact branchpoint-like sequence at positions 10-15 (CUAACU) displayed an average inclusion of 45%, whereas variants harbouring mutations at either of the two adenosines that could serve as branch sites in this sequence increased the levels of exon inclusion, and mutation of both adenosines further increased exon inclusion to an average of 75% (**figure 3N**). Also consistent with our model, insertion of AG-containing sequences after position 40 reduced full length exon inclusion, compared to insertion of non-AG-containing sequences, from an average of 75% to 25% inclusion (**figure 3O**).

Collectively, our results reveal a novel mechanism of exon skipping based upon assembly of U2 snRNP on exonic sequences that resemble (but cannot be active as) 3’ splice sites. This illustrates the value of saturation mutagenesis approaches to discover and test mechanistic hypotheses.

### Cryptic 3’ splice sites regulate alternative exons encoding one-pass transmembrane helices

It has previously been reported that the Py-tract binding protein U2AF2 binds to an exonic polypyrimidine tract in IL7R exon 6^47^, promoting exon skipping. Like FAS exon 6, IL7R exon 6 encodes a one-pass transmembrane helix. Interestingly, transmembrane helices are enriched in nonpolar amino acid residues that are encoded by codons with the highest number of pyrimidines (**figure 4A**). Therefore transmembrane-encoding exons are expected to be rich in pyrimidines, allowing regulation by mechanisms similar to those described for IL7R exon 6 or FAS exon 6 (**figure 3**).

**Figure 4.**
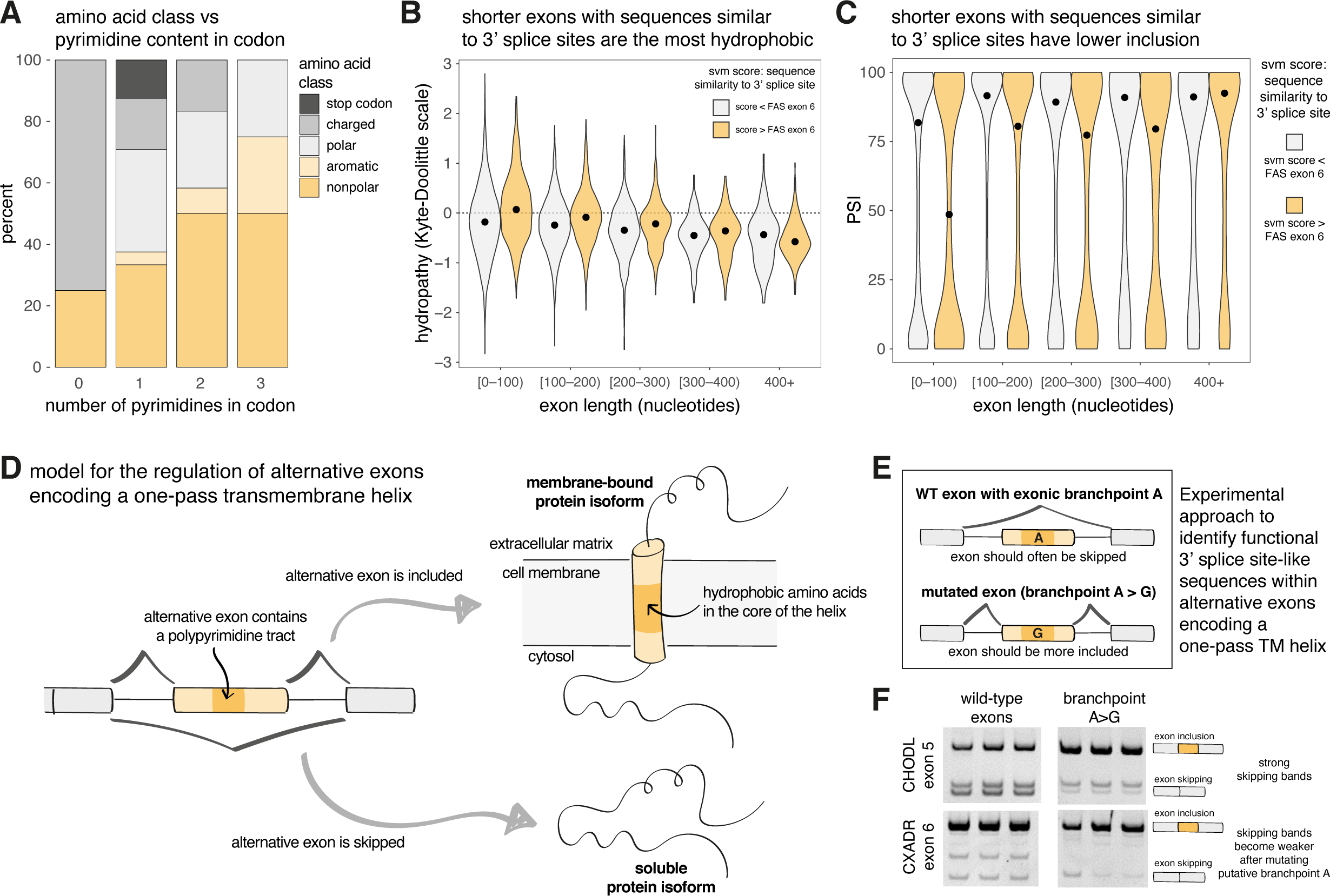
The inclusion of short alternative exons encoding one-pass transmembrane helices is regulated by exonic 3’ splice site-like sequences. A. Amino acid category encoded by each codon, categorized by the number of pyrimidines in the codon. B. Hydropathy score of exons categorized into different length groups, divided by whether they have an exonic sequence more or less similar to a 3’ splice site than FAS exon 6. C. Inclusion of exons categorized into different length groups, divided by whether they have an exonic sequence more or less similar to a 3’ splice site than FAS exon 6. D. Model for the regulation of alternative exons encoding a one-pass transmembrane helix. E. Hypothesis suggesting that 3’ splice site-like sequences in alternative exons encoding transmembrane helices promote exon skipping. F. RT-PCR analysis of the inclusion of two alternative exons (CHODL exon 5 and CXADR exon 6) that each encode a one-pass transmembrane helix, including wild-type (WT) sequences and sequence variants with the putative branchpoint adenine mutated.

To investigate whether alternative exons, and particularly those coding for transmembrane domains, are generally regulated by this type of mechanism, we first used SVM-BPfinder^48^ to scan exons throughout the genome harbouring a branchpoint motif followed by a polypyrimidine tract. For each input sequence, this tool returns a score (‘SVM score’) that reflects how strong a 3’ splice site is predicted to be. Interestingly, 23% of all exons had an SVM score greater than that of FAS exon 6 (1.19), suggesting that a significant proportion of exons across the genome may have cryptic 3’ splice sites or at least sequence elements that resemble 3’ splice site regions. We found that shorter exons (<100 nts) with SVM scores greater than 1.19 encoded the most hydrophobic amino acid sequences (**figure 4B**), consistent with transmembrane domains. These exons had a lower average PSI compared to other exons (**figure 4C**). These results are compatible with the existence of a category of (relatively) short exons containing 3’ splice site-like sequences and, in particular, those encoding individual transmembrane helices (**figure 4D**), whose inclusion is decreased by exonic 3’ splice site-like sequences.

To experimentally validate this hypothesis, we built minigenes containing one-pass transmembrane domain-encoding exon 5 of CHODL^49^ and exon 6 of CXADR^50^, and mutated their exonic putative branchpoint adenosines (**figure 4E-F**). In both cases, the mutations reduced exon skipping, suggesting that the levels of exon skipping were regulated by the recognition of 3’ splice site-like sequences. Cryptic 3’ splice sites in exons may therefore be a widespread mechanism to regulate the inclusion of alternative exons encoding transmembrane helices and thus modulate the balance between soluble and membrane-bound protein isoforms.

### Deep learning variant effect predictors accurately predict the effects of different types of genetic variation on alternative splicing

Multiple computational models have been used to predict the effects of genetic variation on splicing. Our deep indel mutagenesis dataset provides a unique opportunity to test the performance of these models for indel mutations, as well as independent evaluation of their accuracy for predicting the effects of substitutions.

We evaluated the performance of five different models: SMS score (an additive model using 7-mer sequences as input features with parameters learnt in a saturation mutagenesis assay^7^), HAL (an additive model using hexamers as input features with parameters learned from millions of random 50-nt-long exonic and intronic sequences^51^), MMSplice (a modular neural network where modules were trained to predict the effects of mutations on different splicing-relevant sequence regions^52^), SpliceAI (a deep learning model trained on human sequencing data^53^) and Pangolin (a deep learning model based on the SpliceAI architecture but also trained with data from three additional mammalian species^54^).

All models predicted the effects of single-nt substitutions at least moderately well (**figure 5A**), with Pangolin showing the best performance (rho = 0.82), followed by SpliceAI (rho = 0.79), MMsplice (rho = 0.74), HAL (rho = 0.69) and SMS scores (rho = 0.50). These models had a similar range and order of performance when predicting the effects of all insertions (1-, 2-, 3– and 4-nts long) in our library (rho = 0.80 for Pangolin, rho = 0.78 for SpliceAI, rho = 0.66 for MMsplice, rho = 0.67 for HAL and rho = 0.54 for SMS scores).

Interestingly, the predictive performance for deletions was substantially worse for most methods (rho = 0.18 for MMsplice, rho = –0.41 for HAL, and rho = 0.17 for SMS scores) except SpliceAI (rho = 0.90) and Pangolin (rho = 0.89) which remained highly predictive.

**Figure 5.**
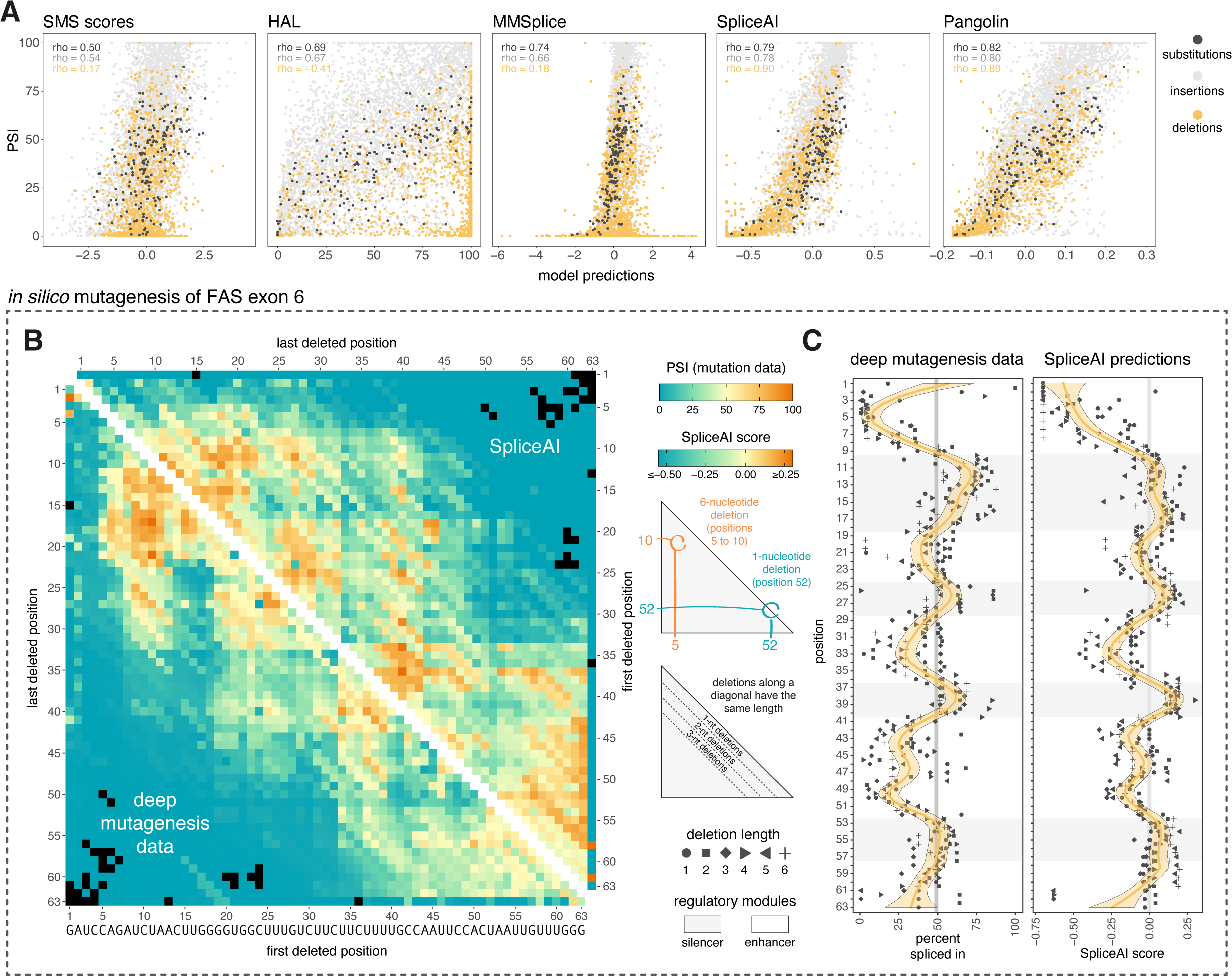
Deep learning predicts the inclusion of variants in our deep indel mutagenesis library. A. Correlations between the inclusion levels of all variants in our library and their inclusion levels according to five different predictors (Substitutions n = 189, Deletion n = 1985, Insertion n = 5744). B. Lower triangle: Heatmap displaying inclusion levels of all deletion variants in our mutant library. Upper triangle: Heatmap showing inclusion levels of all deletion variants in our mutant library as predicted by SpliceAI. C. Left: Inclusion levels of all 1-6 nt long deletion variants along the sequence of FAS exon 6. The yellow line represents a loess fit with a 95% confidence band. Right: Inclusion levels of the same variants as predicted by SpliceAI.

Considering all variants of all types in the library, SpliceAI performed best, with a Spearman correlation of 0.84. Indeed, SpliceAI predictions closely replicated our comprehensive deletion maps (**figure 5B-C**), recapitulating a similar regulatory architecture as that uncovered by our experimental dataset. This opens the possibility that this type of model is used to perform *in silico* deletion mutagenesis experiments and build regulatory maps for other exons as well.

### *In silico* deletion mutagenesis reveals the regulatory architecture of exons genome-wide

To test the utility of *in silico* deletion mutagenesis, we used SpliceAI to predict the effects of all 4 nt deletions in 18,551 exons expressed in at least 80% of GTEx tissues with a length between 50 and 200 nts. The impact of a 4 nt deletion is predicted to depend on each individual exon, although highly-included exons (PSI > 90%) are predicted to be more robust to PSI changes compared to exons included at lower levels (**figure 6A**), an expected consequence of the scaling law that the effects of splicing mutations follow (Baeza et al, 2019).

**Figure 6.**
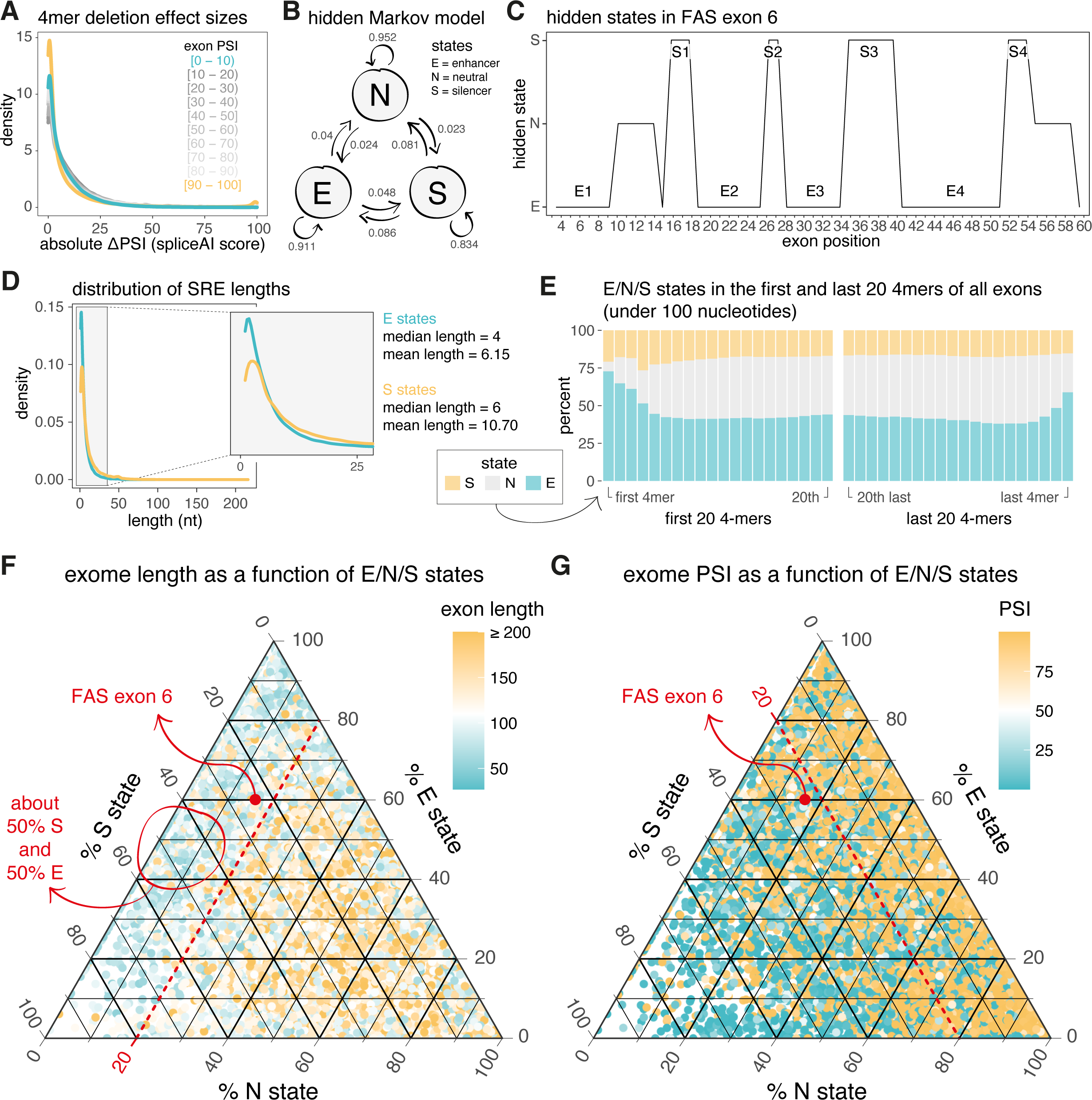
Architecture of regulatory elements in alternatively spliced exons across the transcriptome. A. Distribution of absolute effect sizes of all 4mer deletions in the exome, as predicted by SpliceAI and split by exon PSI groups. B. Hidden Markov model (HMM) with three states (enhancer, silencer, neutral) used to model the splicing regulatory architecture of exons across the genome. C. Predicted regulatory architecture of FAS exon 6 based on our HMM. D. Distribution of exonic splicing enhancer and silencer lengths across the exome, as predicted by our HMM. E. Distribution of the three states of our model in the first and last 20 4mers of all exons under 100 nucleotides long. F. Ternary plot illustrating the relative composition of E/N/S states along the sequences of all 18,551 exons in our dataset. The colour of each point corresponds to the exon’s length. G. Ternary plot illustrating the relative composition of E/N/S states along the sequences of all 18,551 exons in our. The colour of each point corresponds to the exon’s PSI value.

To systematically analyse the regulatory architecture of exons throughout the genome, we used SpliceAI predictions to train a hidden Markov model with 3 states (**figure 6B**): E (enhancer – corresponding to regions of the exon that promote skipping upon deletion), S (silencer – regions that promote inclusion upon deletion) and N (neutral – which have no consistent effect upon deletion). Our model captured the regulatory architecture of FAS exon 6 as uncovered by our deep indel mutagenesis experiment: regions of the exon corresponding to inferred enhancers were predicted to be in state E, and regions corresponding to inferred silencers in state S (**figure 6C**). This suggests that the model can accurately detect splicing regulatory elements along an exon sequence.

We first used our model to study the distribution of splicing regulatory element (SRE) lengths (i.e. stretches of nts in the same E or S states within an exon) throughout the transcriptome. This revealed that most exonic SREs are short, with a median length of 5 nts and a mean of 8.57 nts (compatible with the average binding site of various RNA-binding protein domains^55–57^) and similar to what we find in FAS exon 6. Enhancers were predicted to be slightly shorter than silencers (median lengths = 4 vs 6 nts; mean lengths = 6.15 vs 10.70 nts; **figure 6D**). The model also correctly interpreted exonic sequences that are part of splice site sequences and their immediate neighbourhood as inclusion-promoting (i.e. belonging to the E state, **figure 6E**, **figure S13**).

We next used our model to gain a comprehensive overview of the regulatory architecture of the entire exome. To do this, we used ternary plots to visualise all 18,551 exons in our dataset based on their predicted E/N/S states. This revealed two strong trends. First, for exons shorter than 100 nts, the percentage of nts in the N state tends to be below 20%, irrespective of the proportion of nts in the S or E states, while exons longer than 150 nts tend to have a higher than 20% percentage of nts in the N state (**figure 6F**). This suggests that the inclusion of short exons, such as FAS exon 6, may require a higher density of SREs compared to longer exons. Indeed, as exon length increases, the absolute number of nucleotides in the E or S states rises at a slower rate than expected if the proportion of nucleotides in these states remained constant regardless of exon length (**figure S14**).

Second, across highly-included exons, consistently fewer than 20% of nts are in the S state, regardless of the proportion of nts in the N or E states (**figure 6G**). This suggests that maintaining exon inclusion relies more on a low proportion of silencers than on a high proportion of enhancers, as also evidenced by the high inclusion values of exons whose nts predominantly fall into the N state (bottom right-hand corner in **figure 6G**). Interestingly, FAS exon 6, which has intermediate inclusion levels, approximately aligns with this boundary, with 23% of its nts predicted to be in the S state.

Our deletion analysis revealed not only that nearly the entire sequence of FAS exon 6 is covered by SREs (as is apparently typical of short exons across the genome), but also that its enhancers and silencers alternate in a ‘checkerboard’ pattern along the exon. We used our genome-wide *in silico* deletion mutagenesis to evaluate if this ‘checkerboard’ pattern is likely to be common in additional exons

We first hypothesised that exons encompassed entirely by SREs arranged in an alternating checkerboard pattern would distribute approximately 50% of their nts in the S state and the remaining 50% in the E state. Our ternary plots suggest that exons meeting this criterion are shorter than 100 nts (**figure 6F**) and display relatively low levels of inclusion (**figure 6G**). We next explicitly evaluated this hypothesis by counting the number of times the sequence of each exon in our dataset transitions from the E to the S state directly, and vice versa – without passing through the N state. For example, in the case of FAS exon 6, we counted 7 such transitions (**figure 6C**), equivalent to 9.5 E/S state transitions per 100 nts of exon sequence. The sequences of short (≤100 nt long) highly included (≥ 90%) exons were predicted to have very few E/S state transitions, with an average of 1.5 E/S state transitions per 100 nts (**figure S15A**, results for longer exons shown in **figure S14B-C**). In contrast, short alternatively spliced exons included at lower levels (< 90%) had many more state changes (two-tailed Wilcoxon rank sum test p value < 2.2e-16), with an average of 3.7 E/S state transitions per 100 nts (**figure S15A**, results for longer exons shown in **figure S15B-C**).

Repeating this analysis for E/N state transitions (i.e. where the sequence transitions from the E to the N state and vice versa, without passing through the S state) reveals that short highly-included exons have significantly more E/N state transitions per 100 nts compared to short exons with a PSI below 90% (median 2.4 vs 1.4, two-tailed Wilcoxon rank sum test p value < 2.2e-16, **figure S16A**), in agreement with previous findings that constitutive exons are sustained by strong enhancers^58,59^. Interestingly, this result did not hold true for exons longer than 100 nts (median 1.7 vs 1.8, Wilcoxon rank sum test p value 0.57, **figure S16B-C**).

Alternative exons are therefore predicted to have a high density of splicing regulatory elements, which suggests that their precise inclusion levels are tightly regulated and therefore sensitive to mutation. The alternating pattern of enhancers and silencers further suggests that some of these regulatory domains likely act by modulating the function of a neighbouring domain (e.g. a silencer protein binding to its site might sterically prevent a neighbouring enhancer from being bound by an enhancer protein). On the other hand, the lower density of enhancer-silencer alternations in constitutive exons suggests that their high inclusion levels have not been achieved by fine-tuning binding of splicing regulatory machinery. Their high inclusion levels might therefore simply be a function of their stronger splice sites^60–62^.

### Deletion scans accurately predict the effects of antisense oligonucleotides on exon inclusion

Antisense oligonucleotides (AONs) are an increasingly appealing therapeutic strategy to clinically modulate alternative splicing, as recently illustrated by the clinical success of Nusinersen for the treatment of Spinal Muscular Atrophy^16^ and Eteplirsen for Duchenne muscular dystrophy^17^. AONs base-pair to splice sites or regulatory sequences, competing with the binding of splicing factors and regulators^63^.

Clinically-used AONs are typically longer than individual regulatory elements (18-21 nt AONs versus 5-10 nt regulatory motifs), making the prediction of AON effects challenging. We reasoned that deletion mutagenesis might provide a rapid method to predict the effects of AONs binding to different regions of a transcript, since deleting (a set of) regulatory motif(s) will inhibit the assembly of cognate *trans*-acting regulatory factors, which is also the mechanism of action of AONs, as they typically compete with the binding of *trans*-acting factors to the same sequences (**figure 7A**).

**Figure 7.**
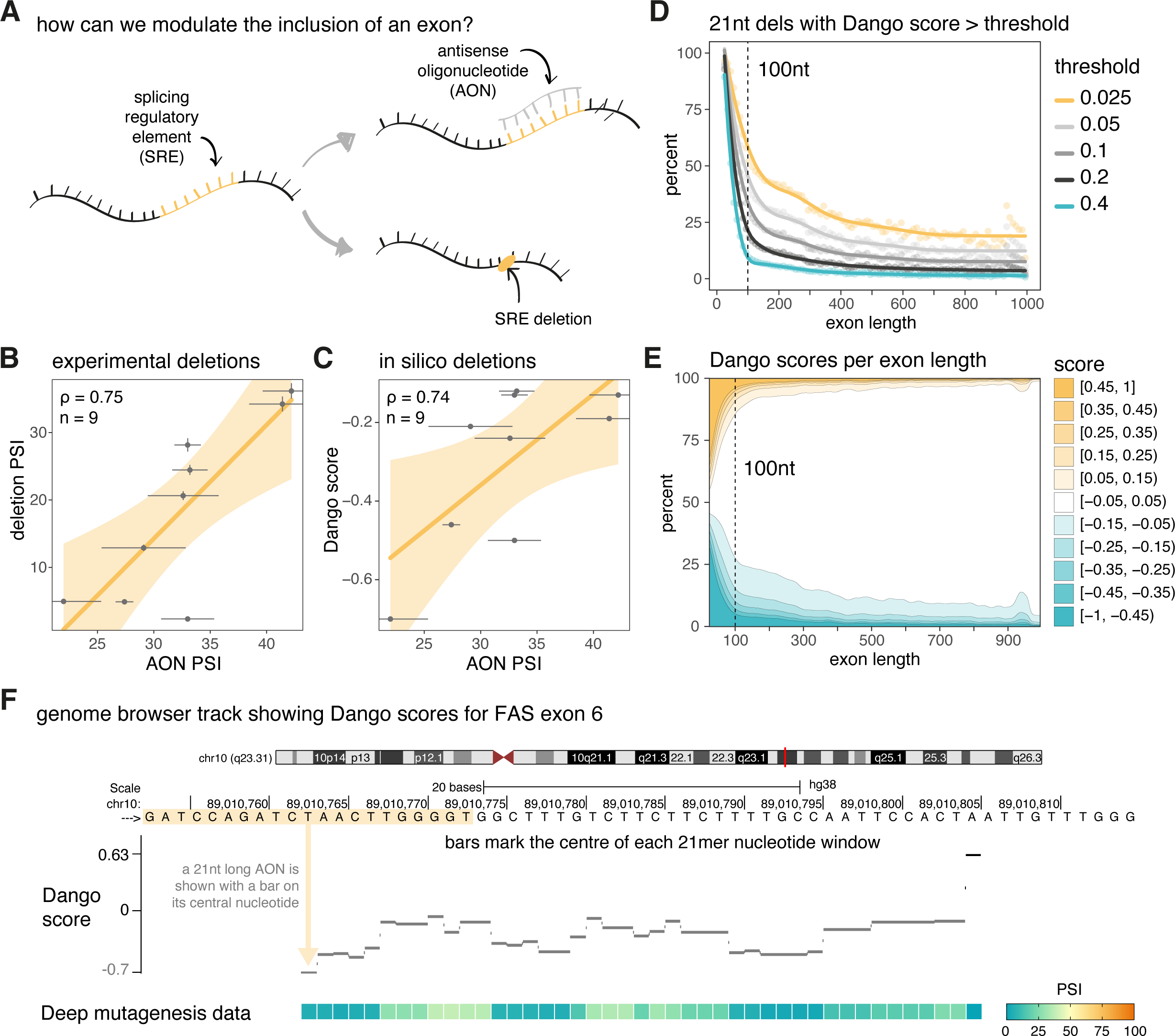
Design of splicing-modulating antisense oligonucleotides. A. The activity of a splicing regulatory element (SRE) could be modulated by using an antisense oligonucleotide (AON) to basepair with this region (therefore sterically blocking any proteins that might bind to the SRE) or alternatively by deleting the SRE altogether. B. Correlation between the PSI values of nine FAS exon 6 variants with 21-nt deletions and the PSI values of WT FAS exon 6 with a 21-nt AON base pairing to the corresponding regions. Horizontal error bars represent the standard deviation of three replicates. Vertical error bars represent the standard error of the mean in our deep insertion mutagenesis library. C. Correlation between the Dango scores of the same nine FAS exon 6 variants as in panel B, and the PSI values of the WT FAS exon 6 with a 21-nt AON base pairing to the corresponding regions. Horizontal error bars represent the standard deviation of three replicates. D. Percentage of 21-nt deletions with Dango scores above the indicated thresholds as a function of exon length. E. Distribution of Dango scores as a function of exon length. F. Custom genome browser track displaying the Dango scores for FAS exon 6. The corresponding PSI values as measured experimentally in our deep mutagenesis assay are shown in the heatmap below, using the same colour scale as figure 5B.

We compared the effects on splicing of an array of partially overlapping AONs collectively covering the entire length of FAS exon 6 (AON walk) with the effects of deletions of the same length (deletion walk). Changes in exon inclusion correlated well for 21-nt AONs and 21-nt deletions spaced every 5nt along the exon (Spearman rho = 0.75, n=9, **figure 7B**). These AONs modulate exon 6 inclusion over a wide dynamic range (from 20% to 50% inclusion, compared with the approximately 50% inclusion level of the WT exon). The correlation between the AON effects and the SpliceAI-predicted effects of 21-nt deletions was similarly strong (rho = 0.74, **figure 7C),** suggesting that *in silico* deletion mutagenesis could be an efficient, affordable strategy to identify regions in an exon that can be targeted by AONs to achieve a range of desired splicing outcomes for therapeutic or biotechnological applications.

### DANGO: a genome-wide resource for AON discovery

We used SpliceAI to predict the effects of all possible 21-nt deletions across the exome to generate a resource we refer to as DANGO (**D**eletion/**AN**tisense oli**GO** – with each DANGO score corresponding to the SpliceAI predictions for a particular 21-nt deletion). 12.4% of all 21nt deletions had an absolute DANGO score greater than 0.1 (mean absolute DANGO score across the exome = 0.05). Short exons (≤100 nt) were most vulnerable to these deletions (**figure 7D**), with 44.8% of all 21nt deletions in these exons having an absolute DANGO score > 0.1, compared to 10.2% in longer exons. Varying the DANGO score threshold (0.025, 0.05, 0.2, or 0.4) altered the proportion of deletions considered impactful, but it did not change the fundamental finding that short exons are more sensitive to the effects of 21nt deletions.

Regardless of the length of the exon, the proportion of negative DANGO scores is greater than the proportion of positive scores (**figure 7E**). This suggests that AONs targeting exonic regions are more likely to reduce recognition of the exon, rather than increase inclusion.

To visualise our results on an exon-by-exon basis, we generated a custom genome browser track (**supplementary data 1**) displaying the DANGO scores for all exons in the genome. This track allows users to interactively explore exonic regions of interest for sequences that may be susceptible to splicing changes upon 21-nt deletions. Visualising FAS exon 6 reveals that DANGO scores cluster around the identified regulatory domains of this exon (**figure 7F**), suggesting that these scores can accurately reflect the regulatory architecture of exonic sequences. Interestingly, since 21-nt deletions push FAS exon 6 below the length threshold for exon definition (**figure 3A**), nearly all 21mer deletions in this exon are predicted to promote skipping (**figure 7F**) and do, in, fact, promote skipping as demonstrated in our experimental assay (**figure 5B**), but deletions spanning silencer regions promote less skipping than deletions spanning enhancer regions.

DANGO is therefore a genome-wide resource that accurately predicts the effects of 21-nt deletions that can be used for AON selection.

## Discussion

Here, by comprehensively quantifying the effects of substitutions, insertions and deletions on splicing we have shown that experimental and *in silico* deletion scanning are particularly effective strategies for revealing the splicing regulatory landscape of exons and we have provided a first overview of the exonic splicing regulatory landscape of the human genome. Moreover, we have shown that deletion scanning accurately predicts the effects of AONs on exon inclusion and have provided a resource, DANGO, to facilitate AON design genome-wide. Finally, by analysing the effects of insertions, we have discovered a novel regulatory mechanism whereby the inclusion of short exons encoding one-pass transmembrane domains is repressed by the binding of a core component of the spliceosome that normally recognizes introns, U2 snRNP, to exons. This recognition of cryptic intron-like sequences in exons provides a simple mechanism for the evolutionary birth of short microexons.

The FAS exon 6 regulatory landscape has a checkerboard organisation of alternating silencers and enhancers that cover the entire exon. *In silico* deletion scans indicate that this architecture is not unusual, but typical of short alternatively spliced exons. The dense splicing regulatory landscape of human exons (Figure 6) is remarkable and a valuable resource to be further explored in future work.

Collectively, our results highlight the power of deep indel mutagenesis for charting regulatory landscapes, generating novel mechanistic hypotheses, and deriving biological insights. One shortcoming of our study is that we have only considered exonic sequences. In future work it will be important to extend deletion scanning to introns and to quantify the effects of inhibiting combinations of intronic and exonic regulatory sequences, potentially allowing finer control of desired splicing changes using AONs. Indeed, we envisage that both *in silico* and high-throughput experimental deletion scans will play an increasingly important role in accelerating the discovery of AONs to effectively modulate splicing for many different therapeutic goals.

## Methods

### Indel library construction

A sequence library was designed to include: 63-nt-long wild-type sequence of FAS exon 6, all possible 189 single-nt substitutions, all 2010 possible deletions ranging in length from 1 to 60 nts, all 5208 possible 1-, 2-, and 3-nt-long insertions as well as 400 randomly-selected 4-nt insertions (full library design along with measured PSI values available in **Supplementary Table 1**). The library was synthesised and purified by Twist Bioscience.

### Indel library amplification

Oligo libraries were resuspended in 10 mM Tris buffer, pH 8.0 to a concentration of 20 ng/ul. 20 ng of template ssDNA Twist library were PCR amplified with Pfx Accuprime polymerase (Thermo scientific, 12344024) in a total reaction volume of 50 ul in quadruplicate, for 12 cycles (as recommended by Twist Bioscience for a 100-150 nt oligo pool) using the following flanking intronic primers: FAS_i5_GC_F (5’-tgtccaatgttccaacctacag-3’) and FAS_i6_GC_R (5’-ctacttcccaagttatttcaatctg-3’). PCR reactions were combined and cleaned-up with the Quiaquick PCR purification kit, eluted with 50 ul elution buffer and dsDNA measured with a NanoDrop spectrophotometer.

### Indel library subcloning

The amplified library was recombined with pCMV FAS wt minigene exon 5-6-7 ^46^. We used a vector:insert ratio of 1:8, using 150 ng of vector backbone and 20 ng of dsDNA amplified libraries and incubated at 50°C for two hours for DNA assembly, using a Gibson master mix developed at the CRG Protein Technologies Unit, which contains a mix of T5 exonuclease (T5E4111K 1000U from Epicentre Biotech-Ecogen), Phusion polymerase (F530s 100U from VITRO and a Taq DNA ligase (Protein Technologies Unit CRG, homemade). After transformation into Stellar competent cells (Clontech, 636766), combining five replicates in order to maximise the number of individual transformants amplified, cells were grown for 18 hours in LB medium containing ampicillin. We obtained approximately 4.29 million clones. After bacterial transformation, the final plasmid library was purified using the Quiagen plasmid maxi kit (50912163, Quiagen) and quantified with a NanoDrop spectrophotometer.

### Transfection of Hek293 cell line to generate output libraries

750,000 Hek293 cells were plated on 100×20 mm petri dishes and transfected with 80 ng cloned libraries in 8 ml OPTIMEM Reduced Serum Medium with no phenol Red (Life technologies, 11058021) using Lipofectamine 2000 (Life technologies, 11668019) in nine biological replicates. 48h post transfection, cells were collected and RNA was prepared using Maxwell simplyRNA Tissue Kit (Promega, AS1280). cDNA was prepared with 400 ng total RNA using specific vector backbone PT2 primer (5’-AAGCTTGCATCGAATCAGTAG –3’) and Superscript III reverse transcriptase (Thermo Fisher, 18080085). PCR amplification of cDNA samples was performed with GoTaq flexi (PROMEGA, M7806) using a distinct 8-mer barcoded oligos to distinguish the nine experimental replicates. PCR products were run on a 2% agarose gel and the smear corresponding to sizes of the amplification product expected from exon inclusion (full length and insertion and deletion mutants) was excised, purified using the Quiaquick Gel extraction kit (Quiagen, 50928704) and quantified with a NanoDrop spectrophotometer.

### Input indel library

20 ng of the plasmid library were amplified in triplicates using GoTaq flexi DNA polymerase (M7806, Promega) for 25 cycles with three different pairs of barcoded intronic primers FAS_i5_TR_F and PT2 (“Indel library amplification primers” in **Supplementary Table 2**). Since the insertions and deletions result in a library with exons of different length, this resulted in a PCR smear (corresponding to exons ranging in length from 3 to 67 nts), which was gel-purified and sequenced. Each pair of primers had a distinct 8-mer barcode sequence to discriminate between technical replicates (“Primers used for amplifying technical replicates (input library)” in **Supplementary Table 2**).

### Sequencing

Equimolar quantities of three independent amplifications of the input library and equimolar quantities of the purified inclusion smear (output library) of each of the nine replicates were pooled and sequenced at the CRG Genomics Core Facility where Illumina Ampliseq PCR-free libraries were prepared and run on a single lane of an Illumina HiSeq2500. In total, 424 million paired-end reads were obtained (188 and 236 million for input and output respectively). The median sequencing coverage for all exon variants in the input was 2114 reads. In the output, the median sequencing coverage was between 278 and 468 reads. Raw sequencing data has been submitted to GEO with accession number GSE244179.

### Data processing and calculation of PSI values

FastQ files from paired-end sequencing were processed with DiMSum v1.2.7^64^ using default settings with minor adjustments (https://github.com/lehner-lab/DiMSum). First, DiMSum was run in default paired-end mode to demultiplex reads into input and replicate output samples (Stage 0 only). Second, DiMSum Stages 1-5 were run in single-end mode (’--paired’ = F) using only demultiplexed forward reads that have full coverage of the exon sequence. Reverse reads, originally intended to cover a unique molecular identifier (UMI) and a 3’ portion of the exon sequence, were discarded. The final stage estimates an enrichment score (ES) and associated error for each mutant variant based on its frequency in the input and output libraries, and relative to the wild type sequence in both libraries. Experimental design files and command-line options required for running DiMSum on this dataset are available on GitHub (https://github.com/lehner-lab/fas-indel-library).

The PSI of the wild type FAS exon 6 sequence has been previously experimentally shown to be 49.1%^5^. Therefore, the PSI of a variant of interest is estimated as follows:

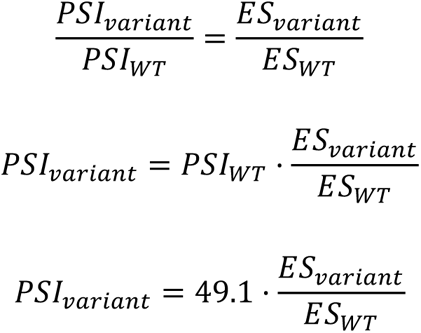

### Experimental validation of estimated PSI values

To confirm the accuracy of our PSI estimates on single-nt substitutions, we took the previously experimentally-determined values of 25 exon variants^5^ and plotted them against the experimentally-determined values (Figure 1C). To validate our estimates of indel PSI values, 15 individual clones from the indel library were Sanger sequenced. They covered a wide range of estimated PSI and were therefore good for validation and checking correlation.

Individual mutants were transfected into Hek293 cells in triplicates to quantify the ratio between exon 6 inclusion and skipping. For RT-PCR, minigene-specific primers were used (“Primers used for amplifying biological replicates (Output library)” in **Supplementary Table 2**). To avoid amplification of endogenous FAS RNAs, these primers (PT1 and PT2) are complementary to a plasmid backbone sequence distinct from endogenous DNA. RT-PCR products were fractionated by electrophoresis using 6% polyacrylamide gels in 1 x TBE and Sybr safe staining (ThermoFisher Scientific, S33102). The bands corresponding to exon inclusion or skipping were quantified using ImageJ v1.47 (NIH, USA). PSI measurements are shown in **Supplementary Table 3**.

Under the particular experimental conditions in which these indel mutants were tested, the wild-type exon was included with a PSI of 49.9% (compared to 49.1% in the experiment done to validate the single nt substitutions^5^). To visualise these results in the same plot as the single-nt substitutions (Figure 1C), we used the splicing scaling law^65^ to adjust all experimentally-determined PSI values to what their values should have been if the wild type had a PSI of 49.1%.

### Estimating PSI values in the GTEx dataset

We estimated the PSI of exons in the GTEx dataset (GTEx Consortium, 2017) from the proportion of reads supporting exon inclusion in the GTEx junction read counts file (**GTEx_Analysis_2016-01-15_v7_STARv2.4.2a_junctions.gct.gz**; available for download at https://www.gtexportal.org/home/datasets). To do this, we used the *quantifySplicing* function from the Psichomics package in R^66^. The *minReads* argument was set to 10 (such that a splicing event requires at least 10 reads for it to be quantified) and the *eventType* argument was set to ‘SE’ (instructing the *quantifySplicing* function to quantify alternative exon events). All estimates were based on the Psichomics hg19/GRCh37 alternative splicing annotations.

### Experimentally validating microexon inclusion

We initially cloned the microexon sequences observed in the output (i.e. those sequences corresponding to large deletions, **figure S11A**) into our plasmid vector backbone (pCMV_FAS_exon4_exon6) and transfected them. No inclusion band was found in the polyacrylamide gels (i.e. these sequences were 100% skipped, **figure S11B**).

Since the nt composition of the PTB binding domain in the central region of FAS exon 6 is very similar to the polypyrimidine tract of a 3’ splice site, we reasoned that an AG-containing insertion right after nt 40 (e.g. CAG, AG, TA, A, CTA) could create a new 3’ splice site in this region of the exon. Such a splice site would produce the microexons detected in the deep mutagenesis experiment. We introduced these insertions (as well as the non-AG-containing GAC insertion as a negative control) into the vector backbone using site-directed mutagenesis (Agilent, 200523) using the relevant mutagenesis primers. These minigene constructs were transfected into Hek293 cells in triplicate. RT-PCR products were fractionated by electrophoresis on 6% native acrylamide gels.

### Experimentally testing exon inclusion with different splice sites

To test the effects of mutations in the presence of different 3’ splice site strengths, we used partially complementary oligonucleotides in combination with TaqPlus precision (Agilent,600212) and PCR “around the world” (primers pointing in opposite directions from the mutagenesis site to amplify the full length of the plasmid) to replace the naturally weak 3’ splice site of FAS exon 6 (UUUCAUAUAAAAUGUCCAAUGUUCCAACCUACAG) with a strong 3’ splice site sequence (UACUAACGGCUUUUUUUUCCUUUUUCAG).

### PTB/SRRM4 overexpression experiments

We overexpressed the SRRM4 (or PTBP1) protein by co-transfecting minigenes containing a CAG or AG insertion after position 40 along with 1000 ng of pcDNA5_SRRM4_flag (or pcDNA5_PTBP1_T7) in lipofectamine 2000 for 24 hours. RT-PCR was then used to analyse the splicing ratios as described above.

### In vitro transcription experiments

T7 promoter-containing transcription templates were generated by PCR using Gotaq flexi enzyme (Promega, M7806): FAS exon 6 WT/noBP templates were generated from FAS WT minigene, Fas M0 template was generated from Fas M0 minigene (Izquierdo et al, 2007), FAS exon 6 UAG mutant template was generated from a ssDNA oligonucleotide. PCR products were purified on agarose gel.

Cy5-CTP/Cy5-UTP labelled RNA were transcribed directly from the PCR templates using Megascript T7 Transcription kit (Ambion) according to the manufacturer’s instructions. A complex formation 15 ng/ul fluorescently labelled RNA were incubated with 3 ul of HeLa cell nuclear extracts (CILBIOTECH, CC-01-20-50) supplemented with 3 mM MgCl2, 24.9 mM KCl, 3.33% PVA, 13.3 mM HEPES pH 8, 0.13 mM EDTA,13.3 % glycerol, 0.03 % NP-40, 0.66 mM DTT, 2 mM ATP and 22 mM creatine phosphate in a final volume of 9 ul. The mixture was incubated for 18 min at 30°C. 1 microliter of heparin (10 ug/ul stock) was added and incubated for 10 min at room temperature. 3 ul of 50% glycerol were added and 9 ul loaded on a composite gel (4% acrylamide, 0.05% bis-acrylamide, 0.5% agarose, 50mM Tris, 50mM glycine).

The gel was run for 6 hours at 200 Volts in 50mM Tris / 50mM glycine buffer. After electrophoresis, fluorescence was detected using a Typhoon PhosphorImager. The inactivation of U1 snRNP and U2 snRNP was performed as described in Dönmez et al^67^ using 2′-O-methylated oligoribonucleotide complementary to U1 snRNA (5’-CUGCCAGGUAAGUAU-3’) or U2 snRNA (5’-CAGAUACUACACUUG-3’).

### Scanning the exome for sequences similar to 3’ splice sites

To identify exonic sequences similar to 3’ splice sites in our GTEx dataset (see **Estimating PSI values in the GTEx dataset** section), we used SVM-BPfinder, a support vector machine that scores how closely a nt sequence resembles a 3’ splice site preceded by a branchpoint^48^. SVM-BPfinder was run at each position of each exon with the *--species* argument set to Hsap, the *--max-len* argument set to 1000, and the *--min-dist* argument set to 15. The final score assigned to each exon was the maximum score across all of its positions. This corresponds to the sequence within the exon that most closely resembles a 3’ splice site.

### Measuring hydropathy in exons throughout the genome

To measure hydropathy, the *hydrophobicity* function from the *Peptides* package^68^ in R was used with the *scale* argument set to “KyleDoolittle”.

### Preparation of minigene constructs carrying a transmembrane domain (CHODL exon 5 and CXADR exon 6)

Genomic DNA sequences were amplified from commercial genomic DNA (PROMEGA, G304A), and branch site mutations were produced with the help of Taq Plus precision (Agilent, 600212). Sequences of genomic regions were cloned into the pCMV_FAS567 minigene replacing FAS exon 6. The amplified genomic sequences were:

**CHODL exon 5 (103 bp) GRCh37/hg19 chr21:19635108-19635210**

GTATAATTCCCAATCTAATTTATGTTGTTATACCAACAATACCCCTGCTCTTACTGATACTGGTTGCTTTTGGAACCTGTTGTTTCCAGATGCTGCATAAAAG

**CXADR exon 6 (139 bp) GRCh37/hg19 chr21:18933656-18933794**

CTTCAAATAAAGCTGGACTAATTGCAGGAGCCATTATAGGAACTTTGCTTGCTCTAGCGCTCATTGGTCTTATCATCTTTTGCTGTCGTAAAAAGCGCAGAGAAGAAAAATATGAAAAGGAAGTTCATCACGATATCAG

Minigene constructs with the wild type exons contained the amplified genomic sequences above. In the case of exons whose putative branchpoint adenines were mutated and substituted with guanines, the minigene constructs contained the following exonic sequences:

**CHODL exon 5**

GTATAATTCCCAATCTAATTTATGTTGTTATACCAACAATACCCCTGCTCTT**<u>G</u>**CTG**<u>G</u>**T**<u>G</u>**C TGGTTGCTTTTGGAACCTGTTGTTTCCAGATGCTGCATAAAAG

**CXADR exon 6**

CTTCAAATAAAGCTGGACTAATTGCAGGAGCCATTATAGGAACTTTGCTTGCTCTAGCG CTC**<u>G</u>**TTGGTCTT**<u>G</u>**TC**<u>G</u>**TCTTTTGCTGTCGTAAAAAGCGCAGAGAAGAAAAATATGAAAA GGAAGTTCATCACGATATCAG

Sequences were all confirmed by Sanger sequencing of acrylamide purified PCR bands by crush and soak method. All minigene constructs were confirmed by Sanger sequencing.

### Predicting FAS exon 6 mutation effects using SMS Scores

We downloaded Supplementary Table 7 from Ke et al.^7^, which lists the SMS scores for all possible 7-mers. To calculate the total SMS score for each exon in our indel library, we performed a sliding window analysis by adding the SMS scores of consecutive 7-mers along its sequence. The final SMS score for each variant was obtained by subtracting the total SMS score for the wild type exon from that of the variant:

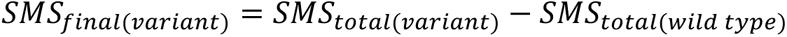

### Predicting FAS exon 6 mutation effects using HAL

To predict the effects of exon variants on inclusion with HAL^51^, we uploaded a file containing the sequences in our library to http://splicing.cs.washington.edu/SE using 49.1% as the wild type levels of inclusion. An output file was returned that contains the predicted PSI values for each sequence in the input file.

### Predicting FAS exon 6 mutation effects using MMSplice

We converted our indel library design file to VCF format, and used this new file as input for MMSplice^52^. We ran the algorithm online, on the Google Colab notebook provided for this purpose (available at https://colab.research.google.com/drive/1Kw5rHMXaxXXsmE3WecxbXyGQJma80Eq6).

This returned a CSV file with multiple columns containing different metrics for each exon variant. We selected *delta_logit_psi* as the predictor for the mutation effects.

### Predicting FAS exon 6 mutation effects using SpliceAI

We converted our indel library design file to VCF format, and used this new file as input for SpliceAI^53^. We ran SpliceAI using GRCh38 as both the genome reference and gene annotation files. All other parameters were set to the default configuration (parameter “D”, the maximum distance between the variant and gained/lost splice site was left to its default of 50, which means that the algorithm could be capturing information about splice site gain and loss beyond the boundaries of the exon). As SpliceAI outputs four scores for each mutant sequence (corresponding to splice site acceptor loss, splice site acceptor gain, splice site donor loss, and splice site donor gain), we selected the score associated with the highest absolute value in each case. If this score corresponded to a splice site loss, the score was multiplied by minus 1.

### Predicting FAS exon 6 mutation effects using Pangolin

We converted our indel library design file to VCF format, and used this new file as input for the Google Colab Notebook made available by the authors of Pangolin^54^ at https://colab.research.google.com/github/tkzeng/Pangolin/blob/main/PangolinColab.ipynb. Pangolin was used with the default options chosen for the Colab Notebook, including GRCh37 as the genome reference. Like SpliceAI, Pangolin outputs four scores for each mutant sequence (corresponding to splice site acceptor loss, splice site acceptor gain, splice site donor loss, and splice site donor gain). The Pangolin score selected for each mutation corresponded to that with the highest absolute value out of these four. If this score corresponded to a splice site loss, the score was multiplied by minus 1.

### *In silico* 4mer deletions in exons genome-wide

The SpliceAI developers created a file with annotations for all possible substitutions, 1 base insertions, and 1-4 base deletions across the genome. This file is available for download at https://basespace.illumina.com/s/otSPW8hnhaZR. We downloaded the file and extracted 4mer deletion data for all exons we calculated PSI values for (see **Estimating PSI values in the GTEx dataset** section). For each 4mer deletion in these exons, we computed its SpliceAI score by taking the maximum value among the acceptor gain, acceptor loss, donor gain, and donor loss scores. If the maximum value was the acceptor loss or the donor loss score, we multiplied the value by –1.

### Hidden Markov model

We used the *depmixS4* package in R to build a hidden Markov model that predicts the locations of exonic splicing enhancers (which promote exon inclusion) and silencers (which promote skipping) in each exon based on SpliceAI scores for 4mer deletions (see section above titled ***in silico* 4mer deletions in exons genome-wide**). Exons with lengths between 50 and 200 nts were used as input for the model, with each exon as a separate time series during training. The model has three hidden states: E (enhancer), N (neutral), and S (silencer), with mean scores of –0.15, 0, and 0.15, respectively. The standard deviations for the states were fixed at 0.1, 0.025, and 0.1 to account for the variability of positive and negative values in the dataset. For example, although breaking a splice site has a much stronger effect than breaking a weak enhancer (resulting in a much more negative spliceAI score), both sequence elements should be classified together in the E state.

### AON Walk

100,000 HEK293 cells in a 6-well plate were transfected with Antisense oligonucleotide harbouring 2’-O Me phosphorothioate modifications at each nucleotide position (Integrated technologies) using 3 ul of Lipofectamine 2000 (11668027, ThermoFisher Scientific) in one ml OPTIMEM I Reduced Serum Medium with no phenol red (11058021, ThermoFisher Scientific) to a final concentration of 2.5 nMolar (exact sequences shown in **Supplementary Table 4**). Six hours post-transfection, the cell culture medium was replaced with DMEM Glutamax (61965059, ThermoFisher Scientific) containing 10% FBS and Pen/Strep antibiotics. 24 hours post-transfection, total RNA was isolated using the automated Maxwell LEV 16 simplyRNA tissue kit (AS1280, Promega). cDNA was synthesised with 400 ng total RNA using Superscript III (18080085, Life Technologies) with a mix of random primers and oligodT. Effects on endogenous FAS exon 6 inclusion were determined by PCR using GoTaq flexi DNA polymerase (M7806, Promega) and the following primers:

FAS_e5_for 5’-TGTGAACATGGAATCATCAAGG-3’ FAS_e7_endo_R 5’-AAAGTTGGAGATTCATGAGAACC-3’

### Exome-wide 21mer deletion scan

After validating the ability of SpliceAI to predict AON effects, we characterised the AON targetability across the genome by performing an exon-wide scan of deletions using SpliceAI scores as a proxy for targetability. Specifically, for all exons in the genome, we produced SpliceAI scores for each length-21 deletion within each of the exons’ boundaries. To compile the list and sequences of each exon, we used R package *biomaRt*^69^ with the *hsapiens_gene_ensembl* dataset. We limited the analyses to canonical exons only (referring to the *transcript_is_canonical* attribute of each exon). After compiling the list of all exons and each of their length-21 deletions, we passed each sequence to SpliceAI using the parameters as described under **Predicting FAS exon 6 mutation effects using SpliceAI**. The resulting scores are referred to as the “DANGO scores” of each exon.

## Statistical tests

All statistical tests were performed in R 3.6.2 using custom code (see **Code availability**

section).

## Code availability

All scripts used in this study have been made available at the following GitHub repository: https://github.com/lehner-lab/fas-indel-library.

## Data availability

DNA sequencing data have been deposited in the Gene Expression Omnibus under the accession number GSE244179.

## Availability of biological materials

All deep mutagenesis libraries used in this study are available upon request.

## Author contributions

PB-C, AJF, MT and GQ performed computational analyses; BM and SB performed experiments; PB-C, BL and JV wrote the manuscript with input from all authors.

## Supporting information

Baeza Minana et al Supplementary Table 1

Baeza Minana et al Supplementary Tables 2_4

## Acknowledgements

The authors would like to thank Manuel Irimia for insightful comments and discussion. This work was funded by European Research Council (ERC) Advanced (670146l, 883742) grants, the Spanish Ministry of Science and Innovation (PID2020-114630GB-I00, LCF/PR/HR21/52410004, EMBL Partnership, Severo Ochoa Centre of Excellence), the Bettencourt Schueller Foundation, the AXA Research Fund, Agencia de Gestio d’Ajuts Universitaris i de Recerca (AGAUR, 2017 SGR 1322), and the CERCA Program/Generalitat de Catalunya. GQ was supported by PRE2022-102744, financed by MCIN/AEI/10.13039/501100011033 and FSE+. The Genotype-Tissue Expression (GTEx) data used for the analyses described in this manuscript were obtained from the GTEx Portal on May 8, 2018 and dbGaP accession number phs000424.v7.p2 on May 8, 2018. The GTEx Project was supported by the Common Fund of the Office of the Director of the NIH and by NCI, NHGRI,NHLBI, NIDA, NIMH, and NINDS.

## Figure legends

**Figure S1.**
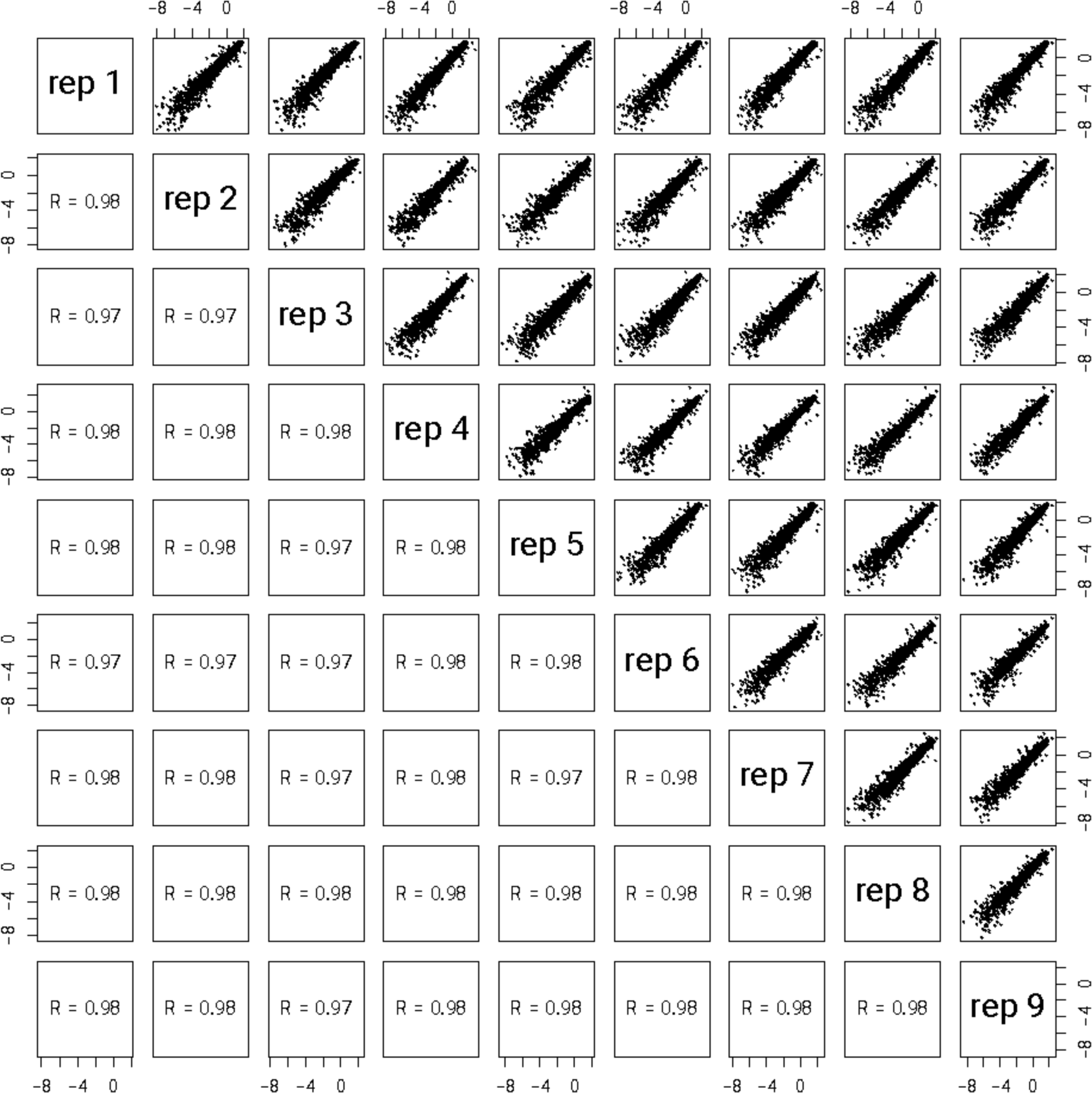
Pairwise correlations of enrichment scores for all exon variants in our library across nine experimental replicates.

**Figure S2.**
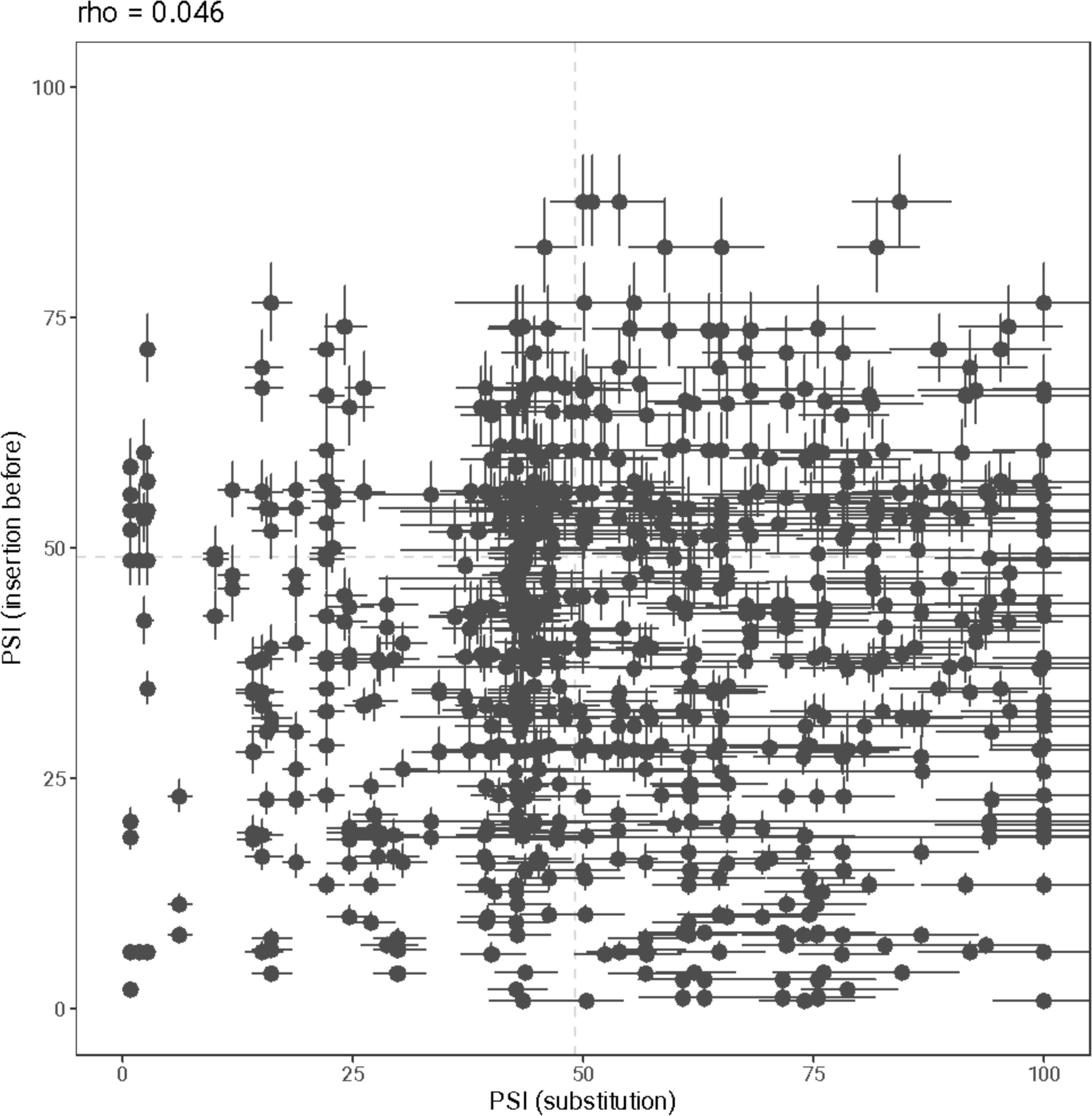
Correlation between PSI values for 1-nt insertions before a given position and substitutions at that position.

**Figure S3.**
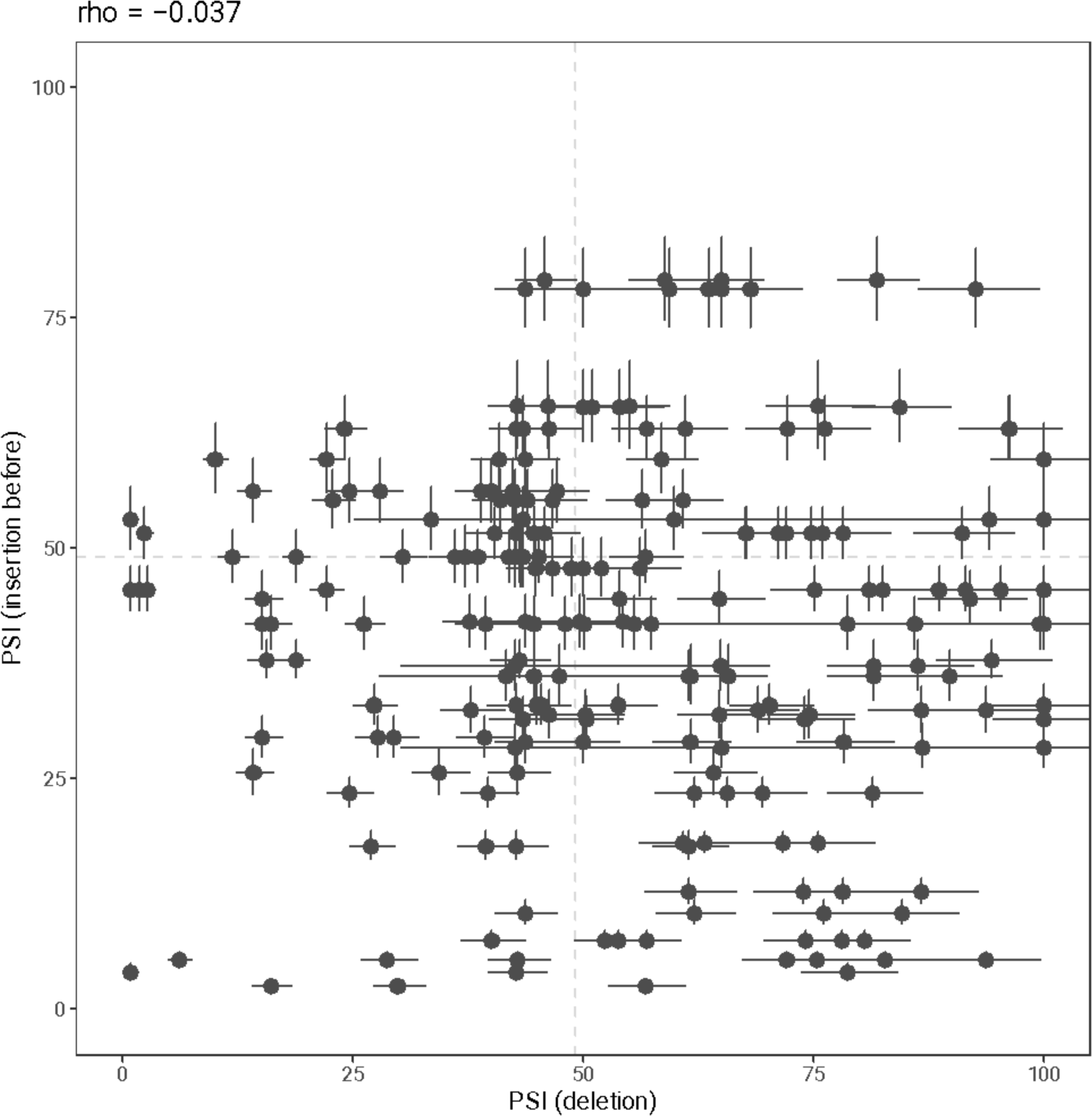
Correlation between PSI values for 1-nt insertions before a given position and deletions at that position.

**Figure S4.**
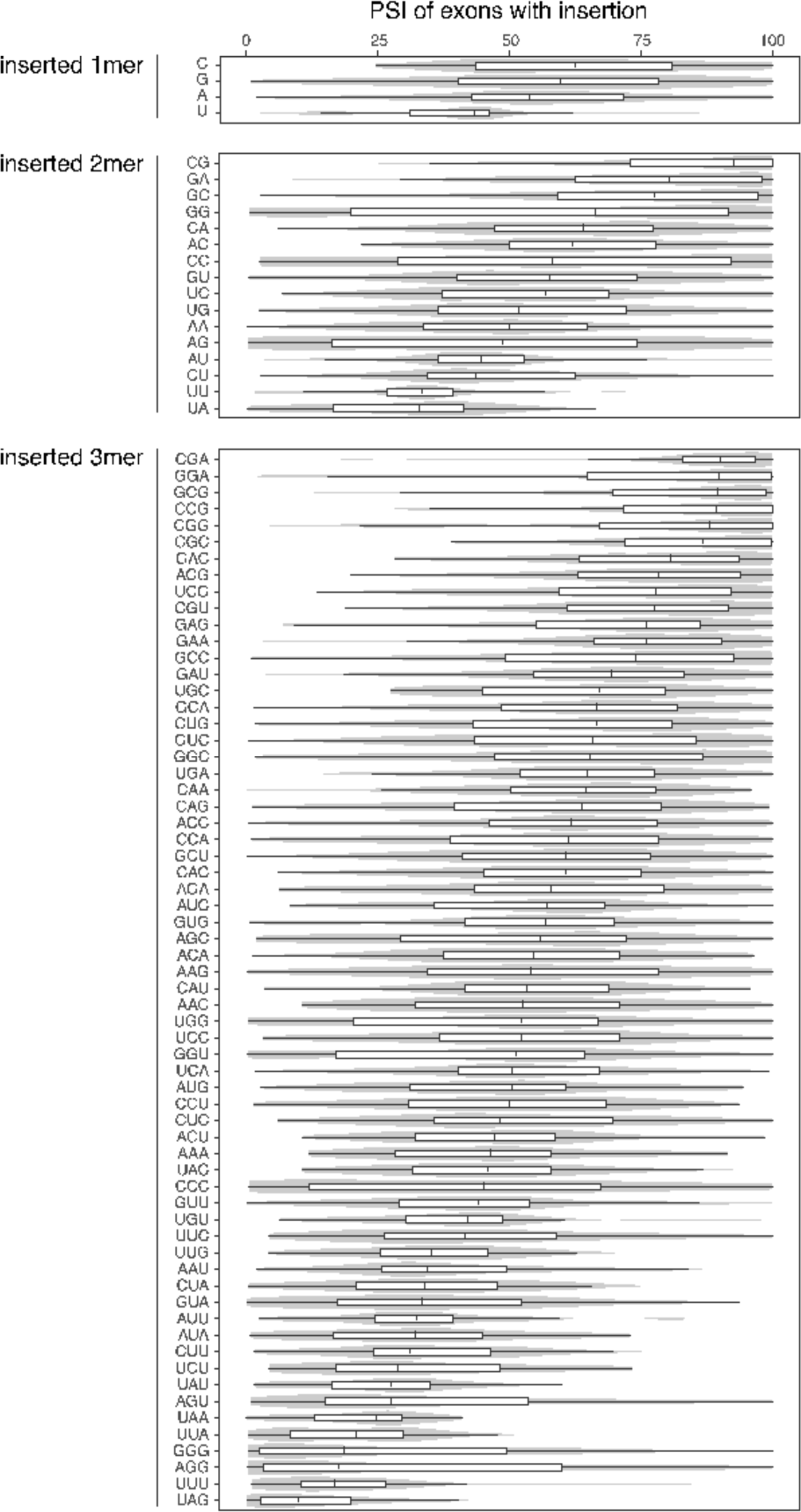
Distribution of inclusion values for exons with each possible 1-, 2-, and 3-nt insertion.

**Figure S5.**
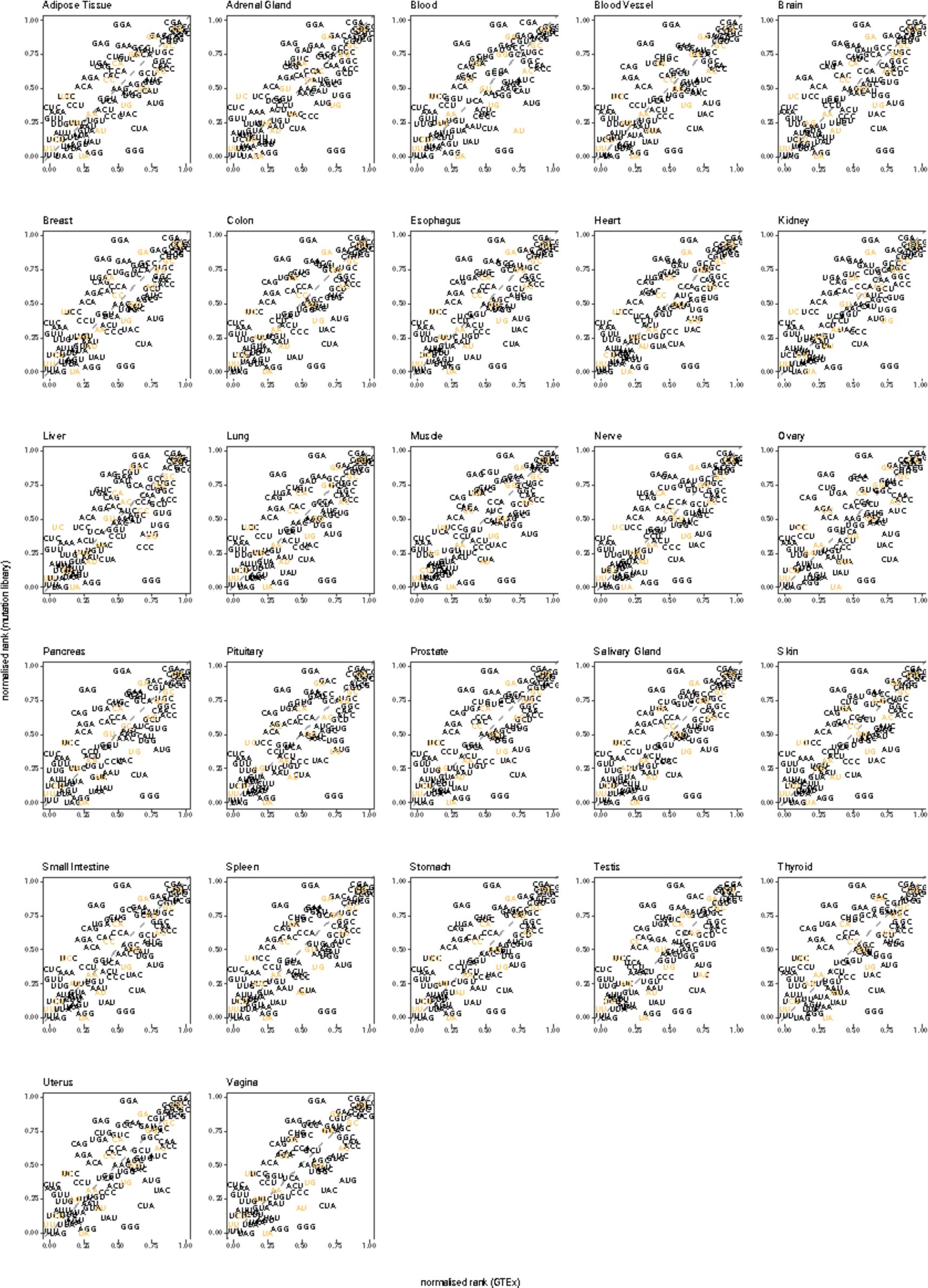
Correlation between 2 (or 3) nt sequences ranked by the median PSI of exon variants in our library, with this sequence inserted, and their rank based on median PSI of exons containing 30 (or 20) such 2mers (or 3mers) in their sequence (all GTEx tissues).

**Figure S6.**
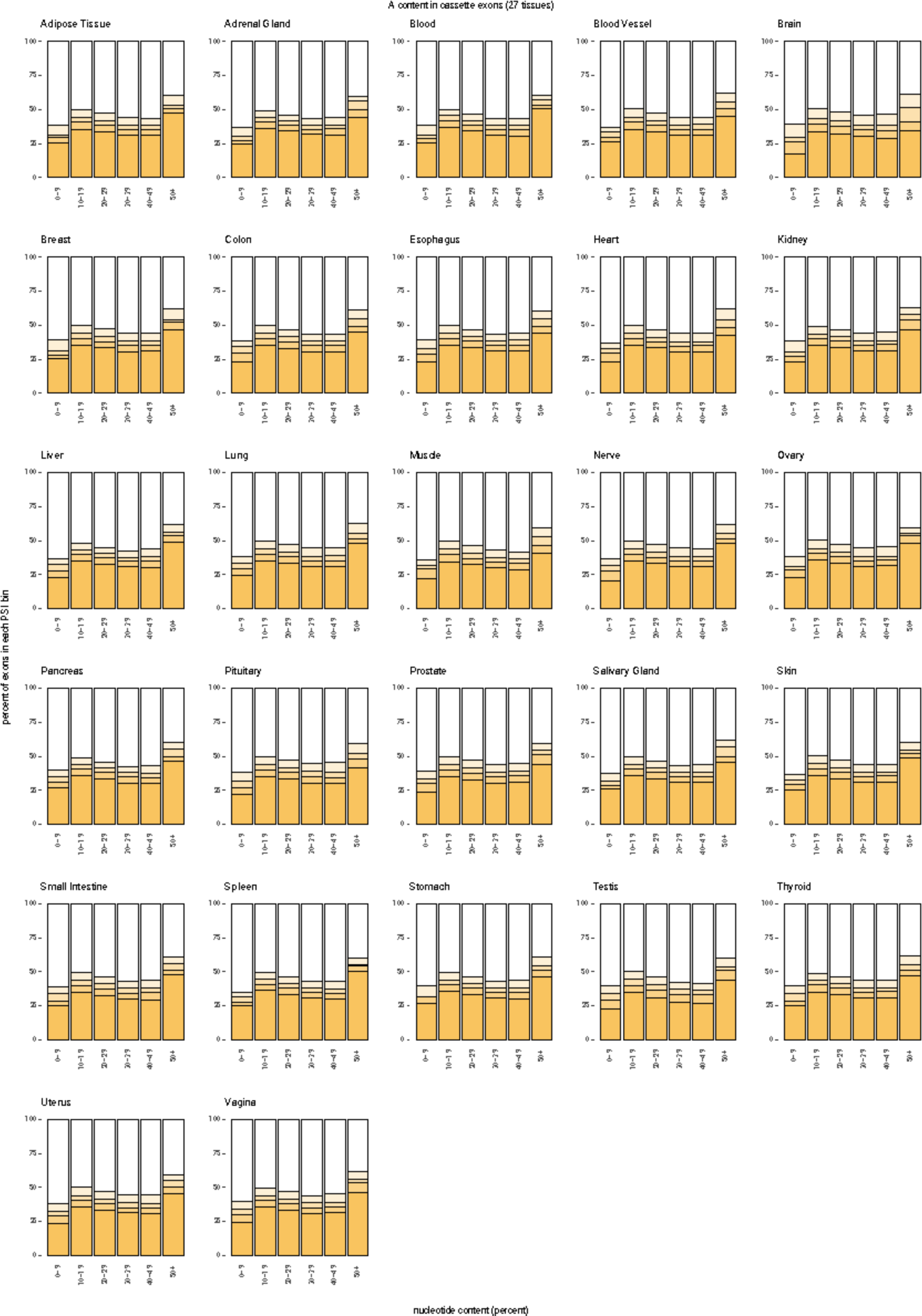
Distribution of PSI values in cassette exons (all GTEx tissues) relative to the percentage of each adenines present in the exon sequence.

**Figure S7.**
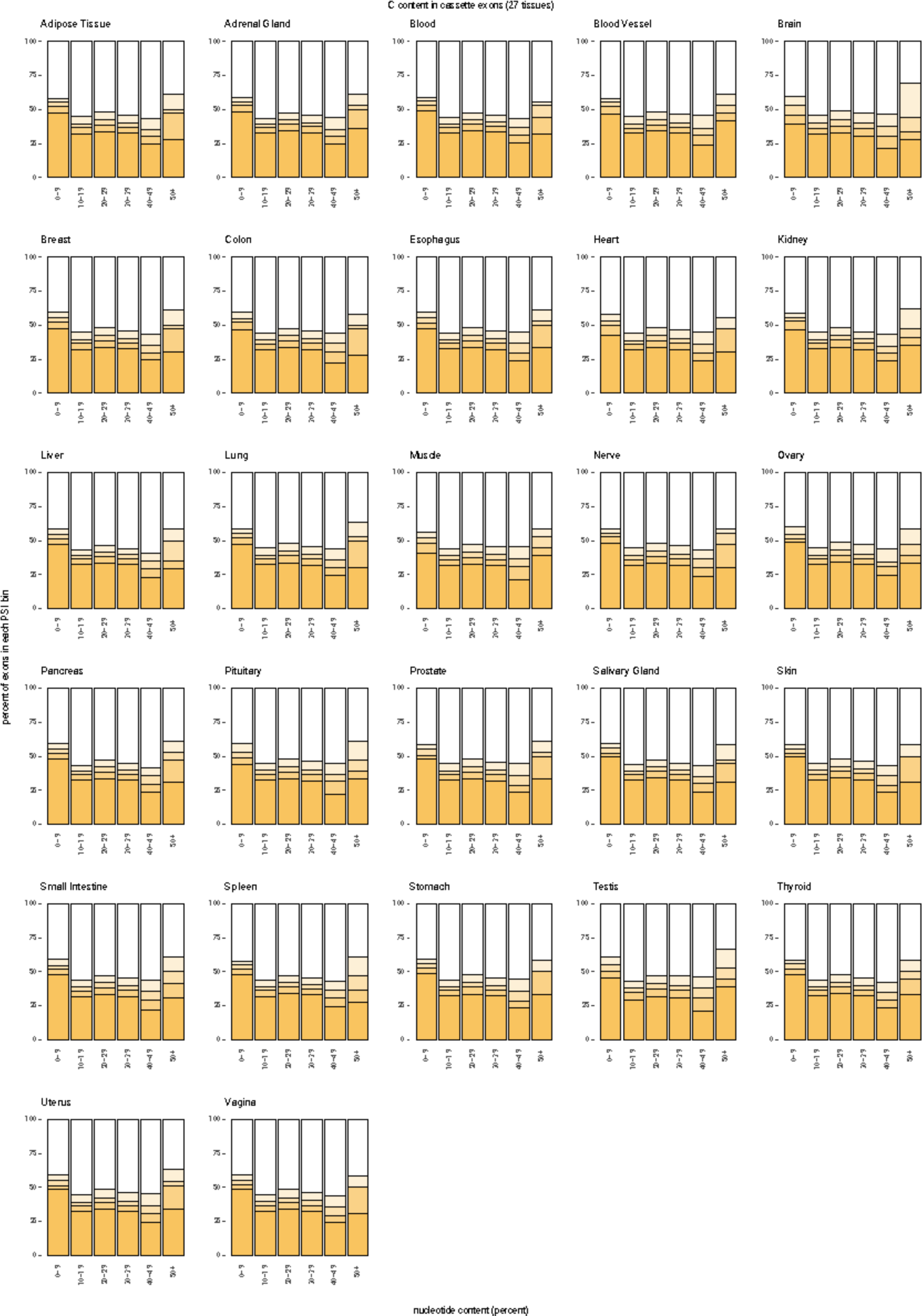
Distribution of PSI values in cassette exons (all GTEx tissues) relative to the percentage of each cytosines present in the exon sequence.

**Figure S8.**
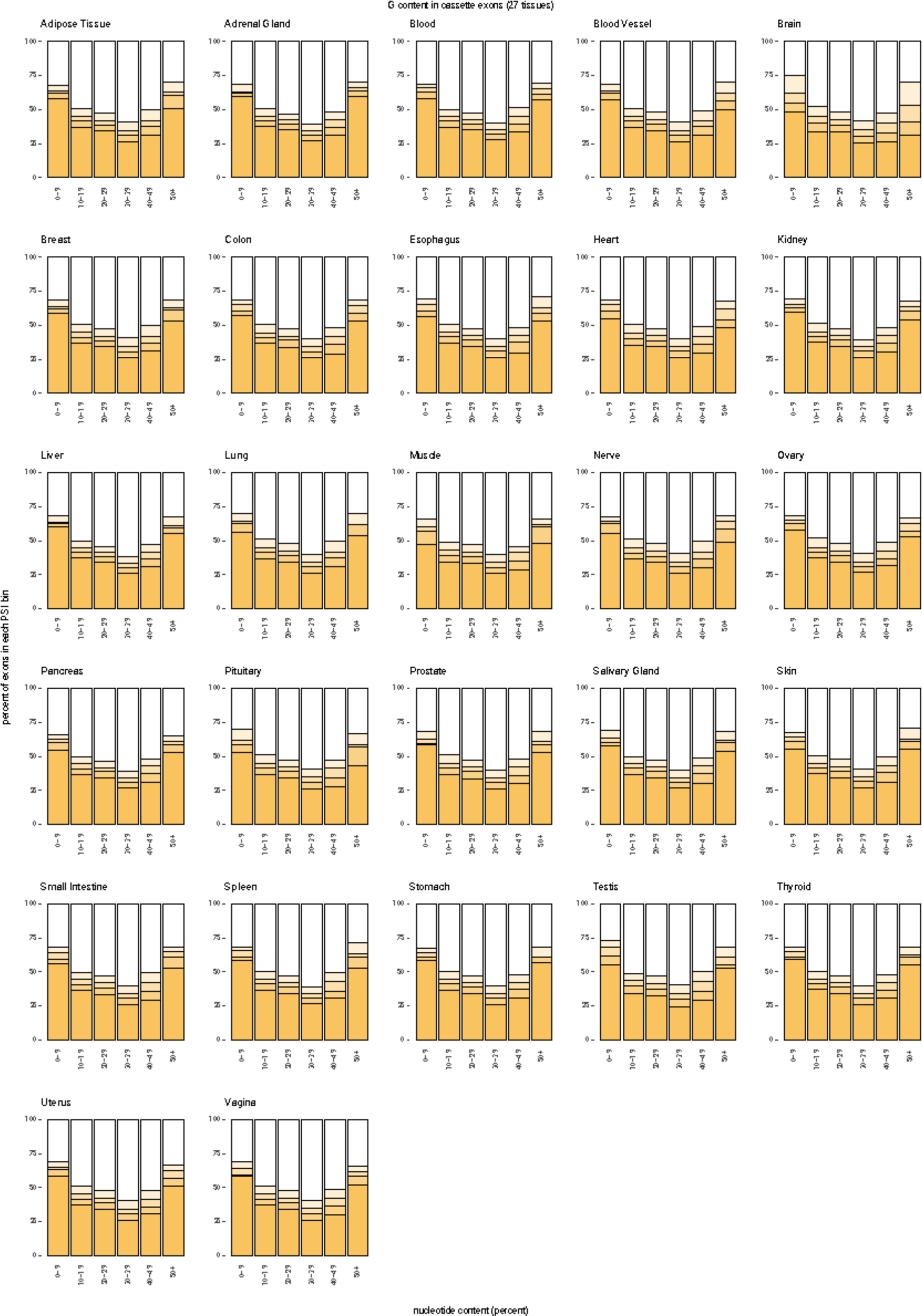
Distribution of PSI values in cassette exons (all GTEx tissues) relative to the percentage of each guanines present in the exon sequence.

**Figure S9.**
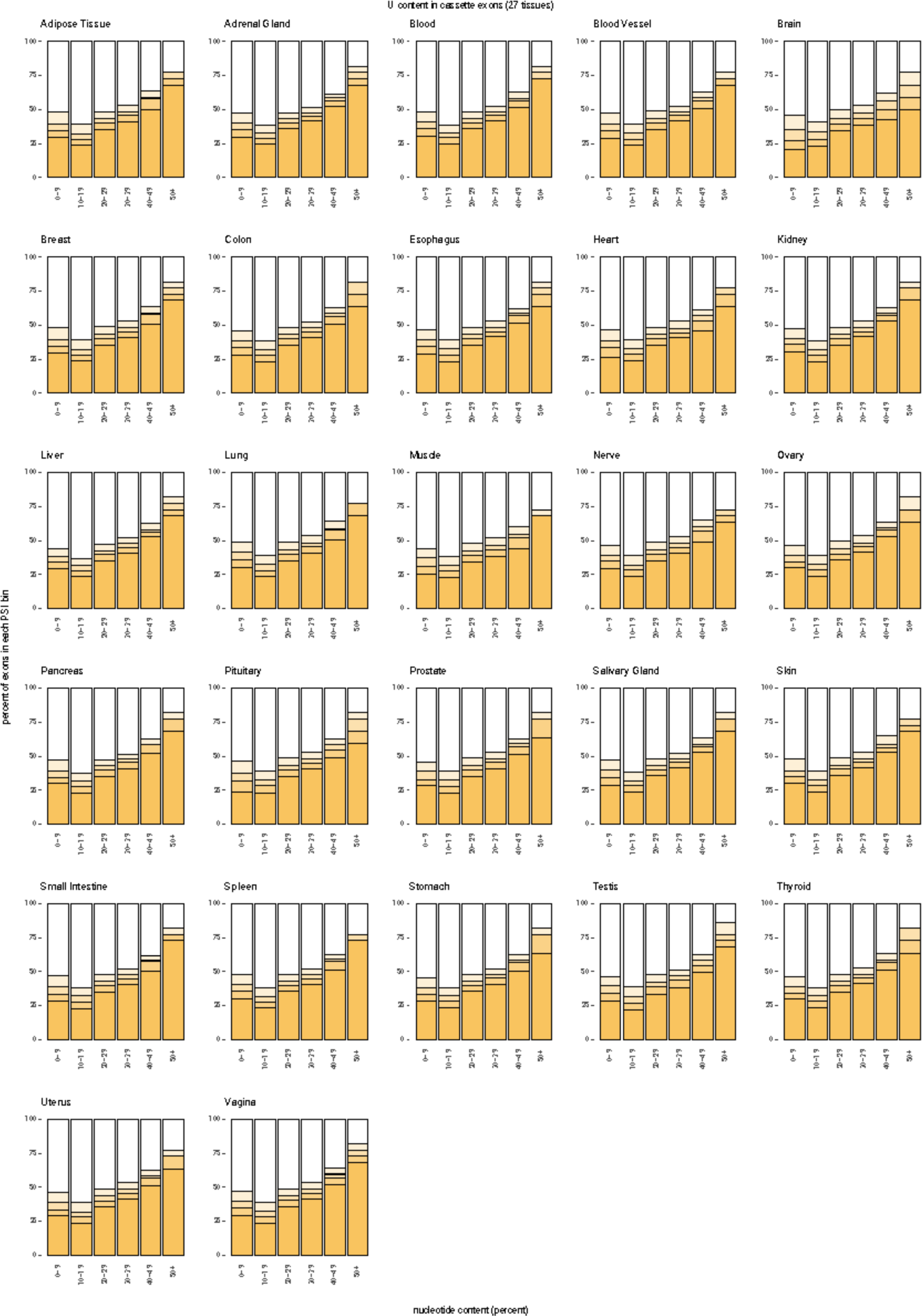
Distribution of PSI values in cassette exons (all GTEx tissues) relative to the percentage of each uracils present in the exon sequence.

**Figure S10.**
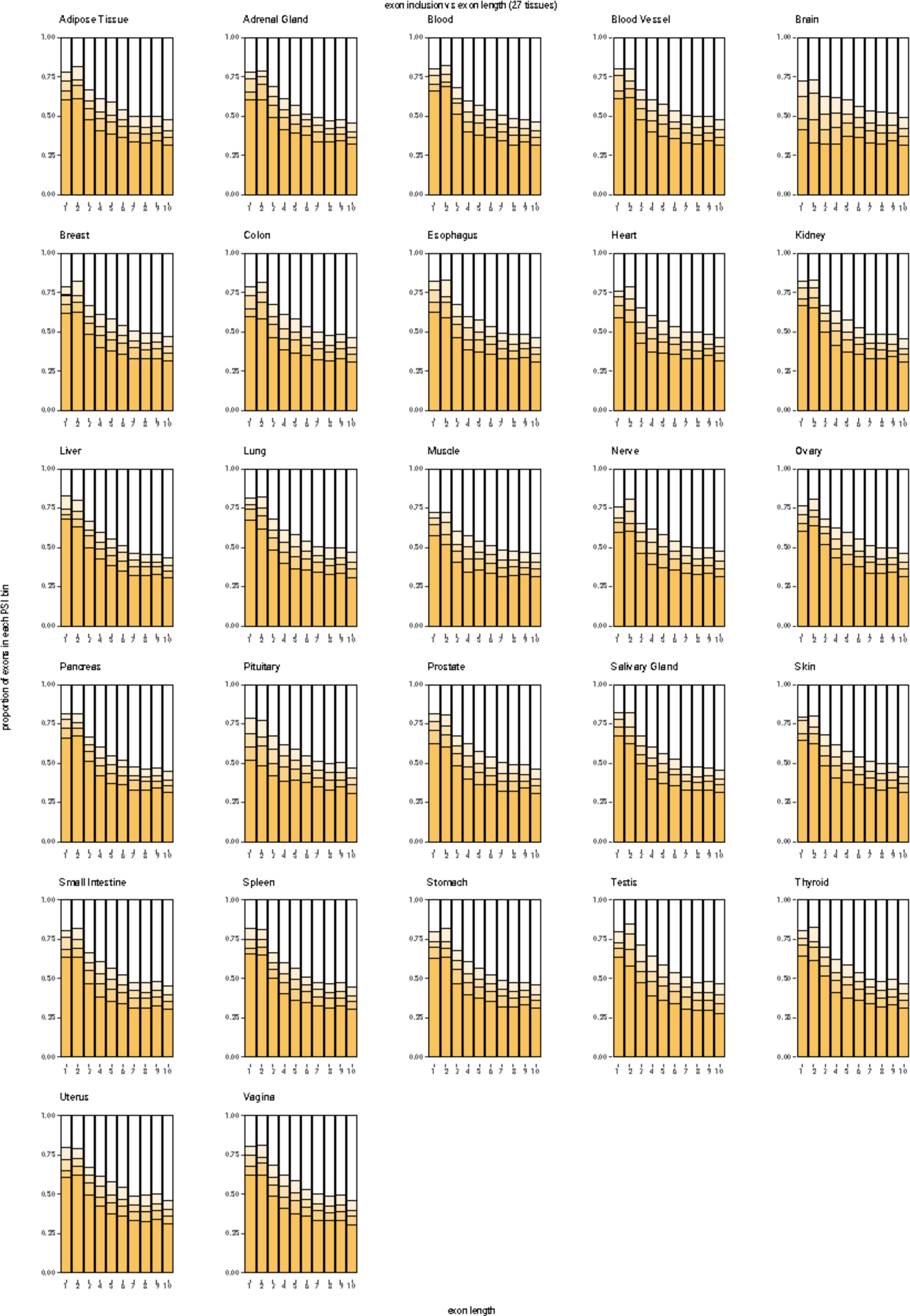
Distribution of cassette exon inclusion values versus exon length (all GTEx tissues).

**Figure S11.**
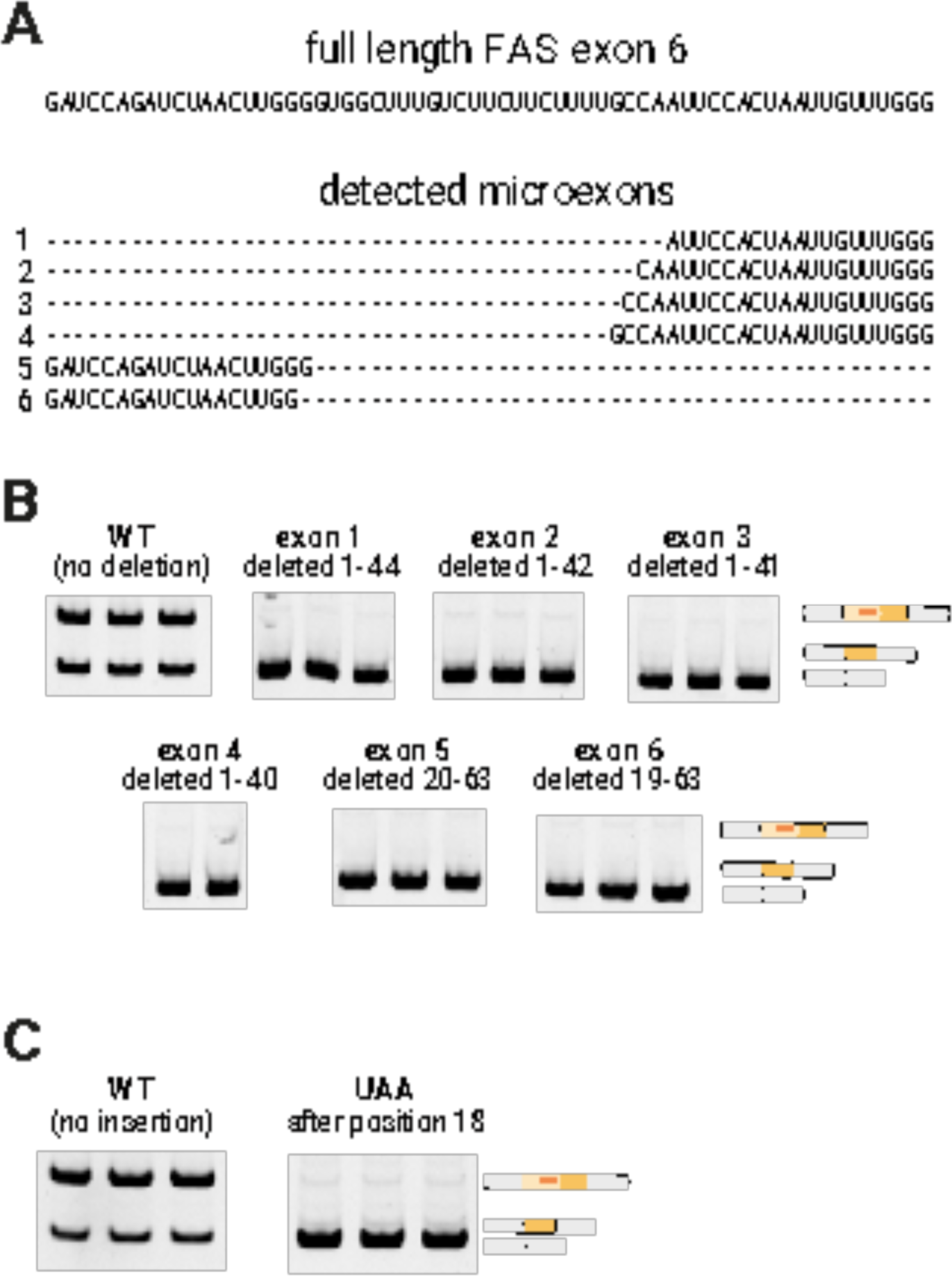
A. Sequences of microexons in our library with detectable levels of inclusion. B. RT-PCR analysis of the inclusion of exons with a sequence corresponding to the detectable microexons. C. RT-PCR analysis of the inclusion of FAS exon 6 with a UAA insertion after exon position 18.

**Figure S12.**
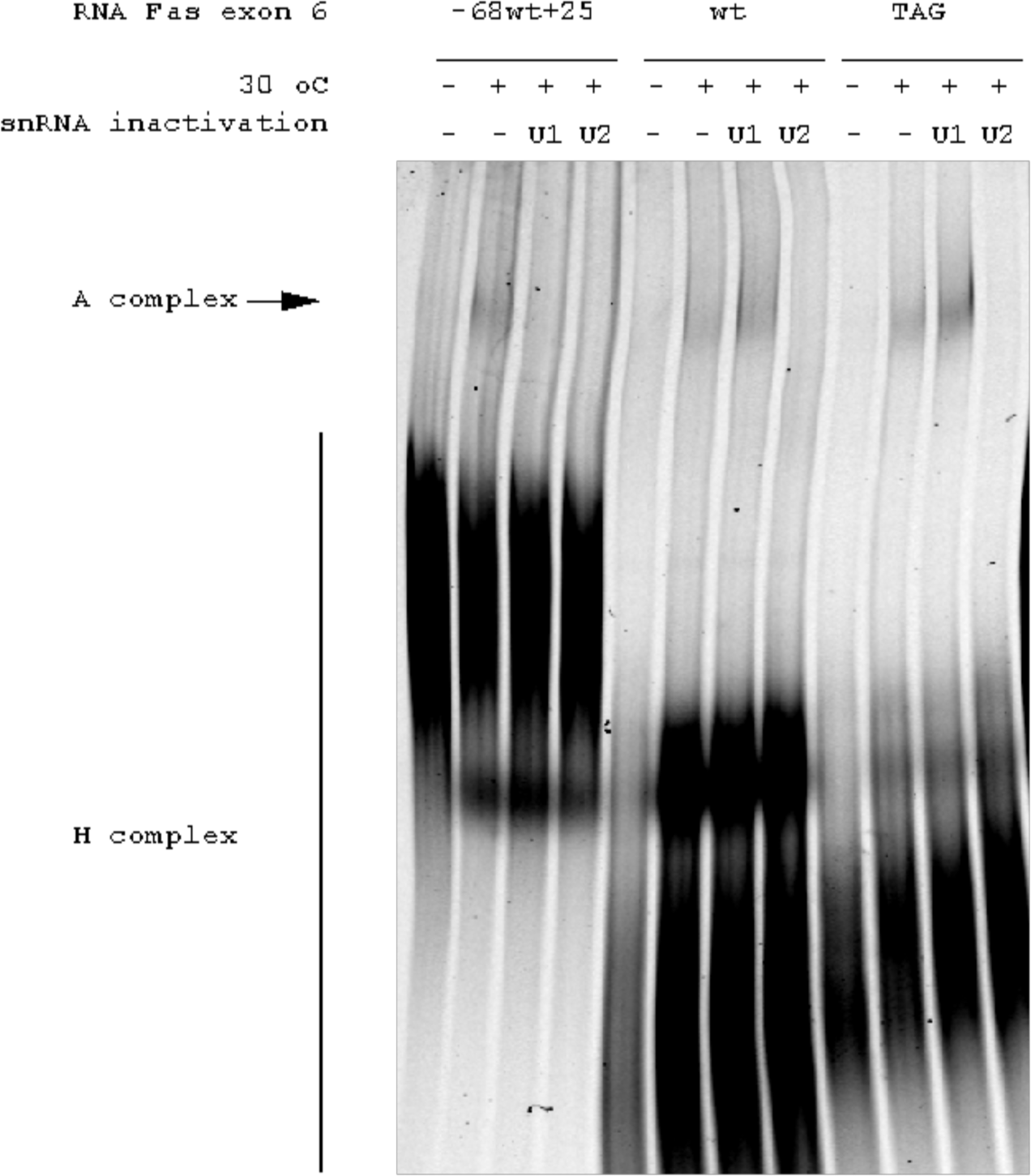
Validation of spliceosome assembly assays and enhanced A (U2 snRNP-containing) complex formation on Fas exon 6 upon inclusion of a 3’ splice site (TAG). Fluorescently-labeled RNAs corresponding to Fas exon 6 (wt), Fas exon 6 with a mutation that creates a 3’ splice site after the internal polypyrimidine tract (TAG) (see Figure 3) or Fas exon 6 flanked by intronic sequences (–68wt+25, which includes the 3’ 68 nucleotides of intron 5 and the 5’ 25 nucleotides of intron 6) were incubated with HeLa nuclear extracts under in vitro splicing conditions at 30 C (+) or on ice as a control (-) and the ribonucleoprotein complexes formed were fractionated by electrophoresis on a composite agarose-polyacrylamide gel. The electrophoretic positions of U2 snRNP-containing complexes (A complex) and hnRNP-containing complexes (H complex) are indicated. Controls of U1 / U2 snRNP inactivation by RNAse H-mediated digestion of the 5’ end of U1 snRNA or of the branch point recognition sequence of U2 snRNA are included. Inactivation of U2 snRNP (but not of U1 snRNP) reduces A complex formation on WT and TAG RNAs, while complexes formed on the –68wt+25 RNA are sensitive to inactivation of either U1 or U2 snRNPs because of exon definition-mediated effects (Izquierdo et al Mol Cell 19: 475-484, 2005). Note the significant increase in A/H complex ratio upon inclusion of a 3’ splice site (TAG) in Fas exon 6 compared to the wild type sequence (WT).

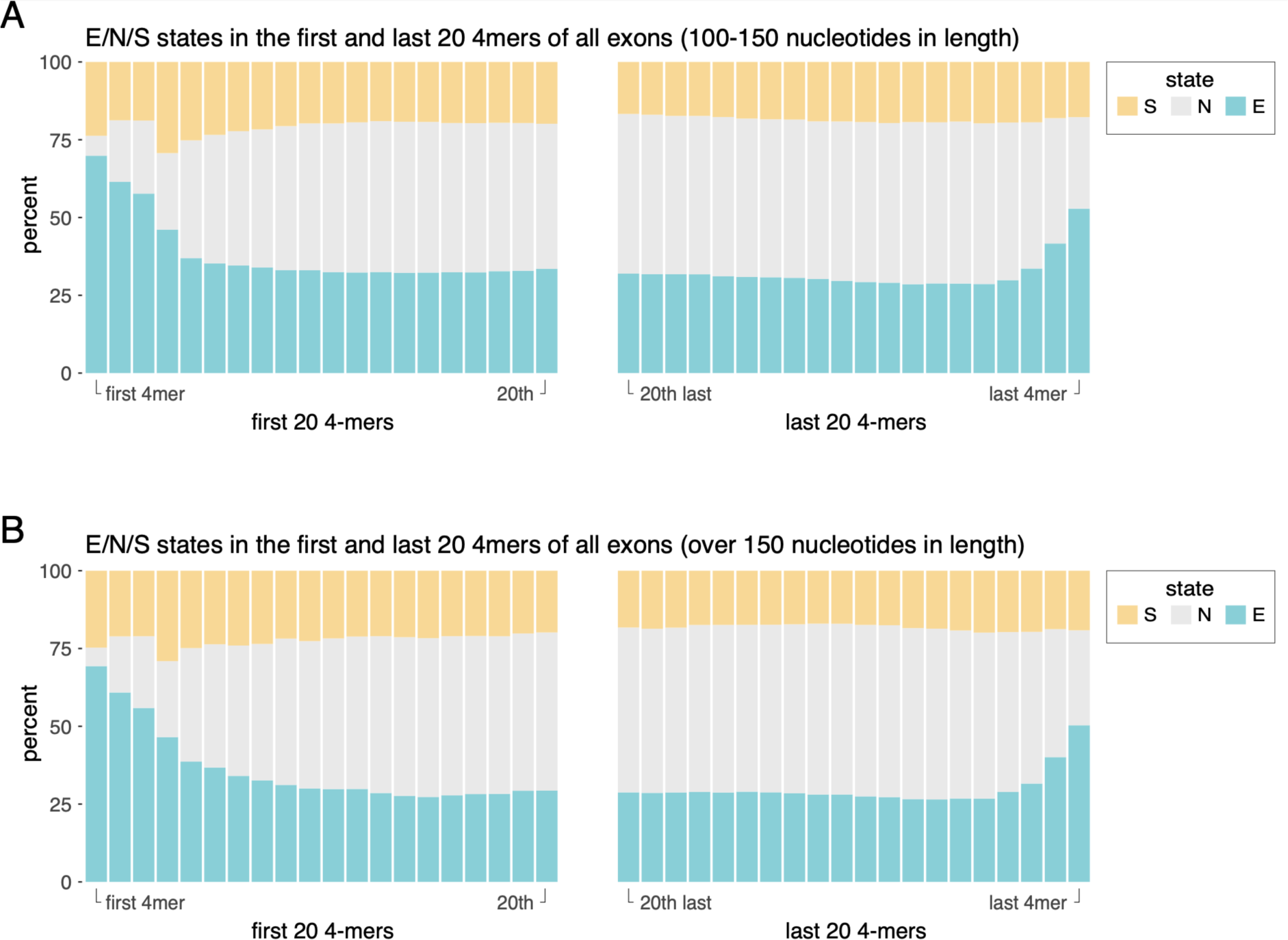
Figure S13

**Figure S14.**
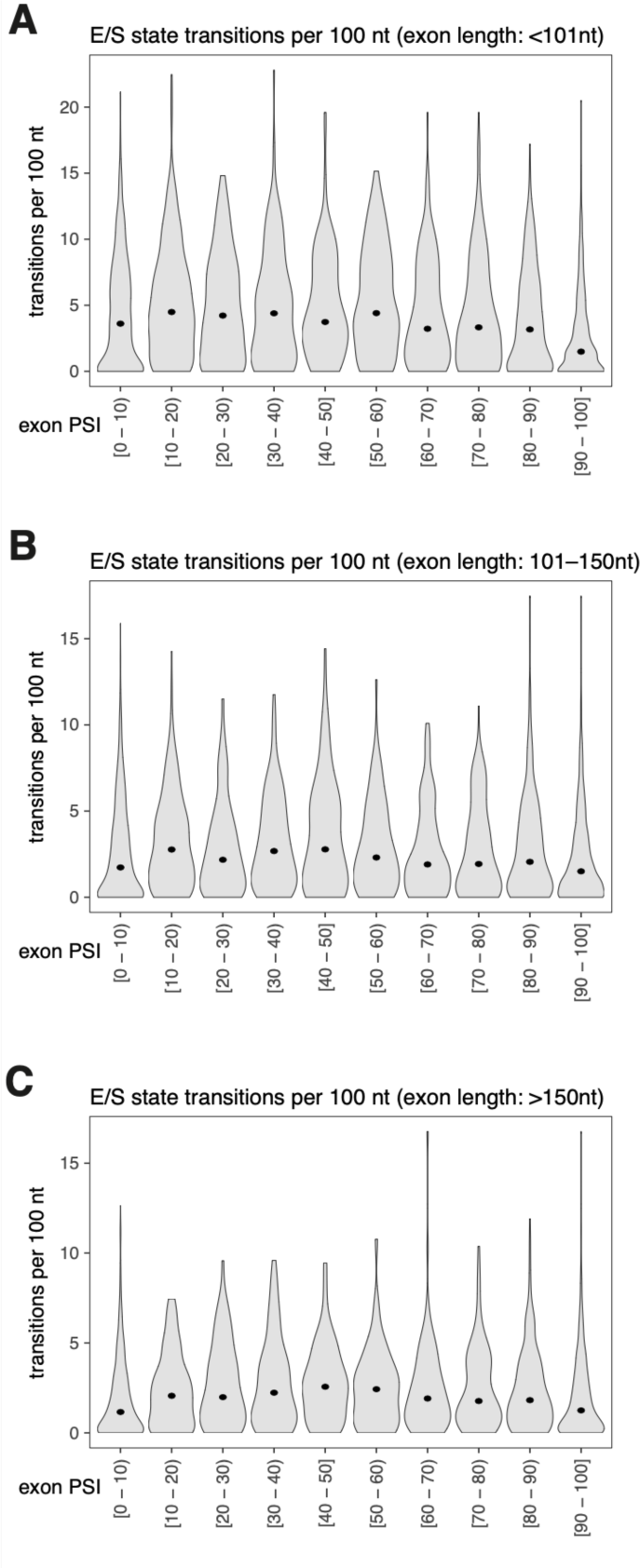
A. Distribution of the number of enhancer-silencer (“E”/“S”) state transitions per 100 nucleotides in all exons shorter than 101 nt, divided into groups based on exon inclusion levels. B. Distribution of the number of enhancer-silencer (“E”/“S”) state transitions per 100 nucleotides in all exons with a length between 101 and 150 nt, divided into groups based on exon inclusion levels. C. Distribution of the number of enhancer-silencer (“E”/“S”) state transitions per 100 nucleotides in all exons longer than 150 nt, divided into groups based on exon inclusion levels.

**Figure S15.**
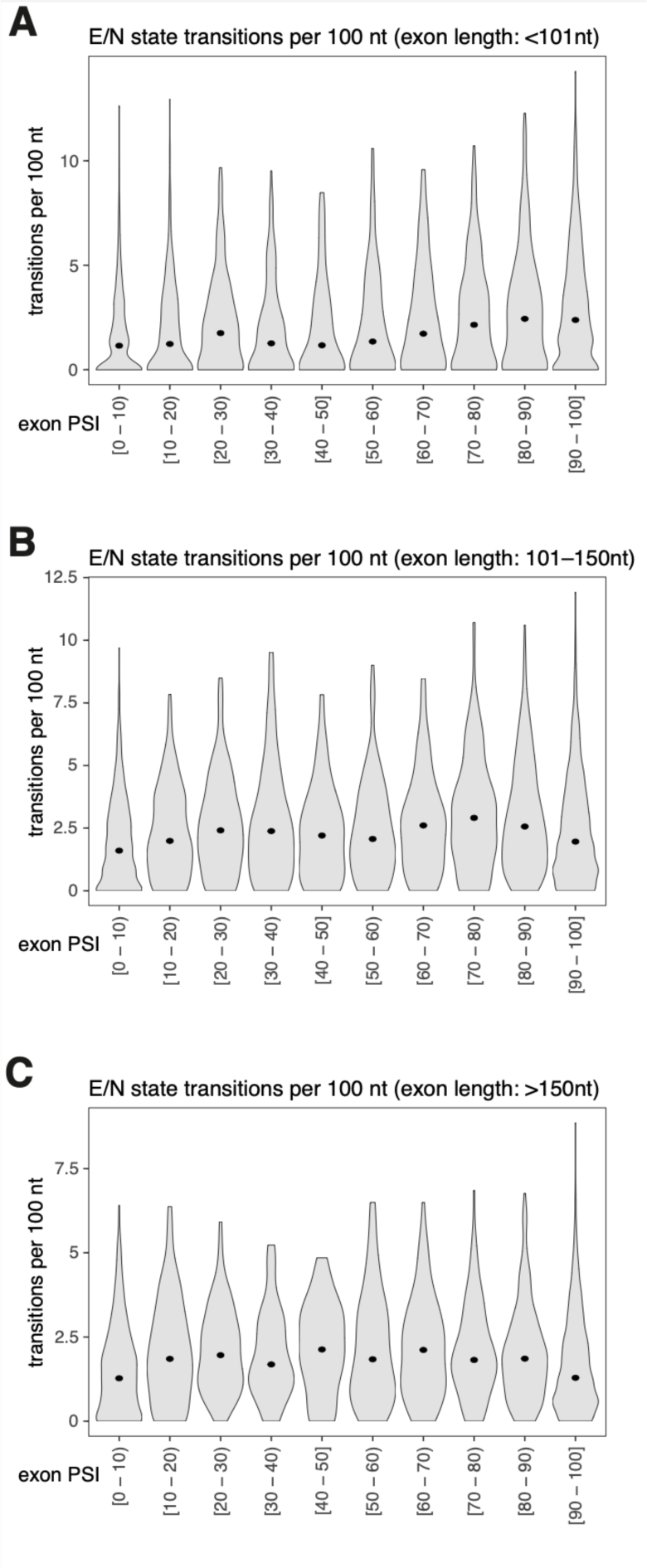
A. Distribution of the number of enhancer-neutral (“E”/“N”) state transitions per 100 nucleotides in all exons shorter than 101 nt, divided into groups based on exon inclusion levels. B. Distribution of the number of enhancer-neutral (“E”/“N”) state transitions per 100 nucleotides in all exons with a length between 101 and 150 nt, divided into groups based on exon inclusion levels. C. Distribution of the number of enhancer-neutral (“E”/“N”) state transitions per 100 nucleotides in all exons longer than 150 nt, divided into groups based on exon inclusion levels.

## References

1. Braunschweig, U., Gueroussov, S., Plocik, A. M., Graveley, B. R. & Blencowe, B. J. Dynamic integration of splicing within gene regulatory pathways. Cell 152, 1252–1269 (2013).

2. Anna, A. & Monika, G. Splicing mutations in human genetic disorders: examples, detection, and confirmation. J. Appl. Genet. 59, 253–268 (2018).

3. Scotti, M. M. & Swanson, M. S. RNA mis-splicing in disease. Nat. Rev. Genet. 17, 19– 32 (2016).

4. Sterne-Weiler, T. & Sanford, J. R. Exon identity crisis: disease-causing mutations that disrupt the splicing code. Genome Biol. 15, 201 (2014).

5. Julien, P., Miñana, B., Baeza-Centurion, P., Valcárcel, J. & Lehner, B. The complete local genotype-phenotype landscape for the alternative splicing of a human exon. Nat. Commun. 7, 11558 (2016).

6. Braun, S. et al. Decoding a cancer-relevant splicing decision in the RON proto-oncogene using high-throughput mutagenesis. Nat. Commun. 9, 1–18 (2018).

7. Ke, S. et al. Saturation mutagenesis reveals manifold determinants of exon definition. Genome Res. 28, 11–24 (2018).

8. Baeza-Centurion, P., Miñana, B., Valcárcel, J. & Lehner, B. Mutations primarily alter the inclusion of alternatively spliced exons. Elife 9, (2020).

9. Soemedi, R. et al. Pathogenic variants that alter protein code often disrupt splicing. Nat. Genet. 49, 848–855 (2017).

10. Cheung, R. et al. A Multiplexed Assay for Exon Recognition Reveals that an Unappreciated Fraction of Rare Genetic Variants Cause Large-Effect Splicing Disruptions. Mol. Cell 73, 183–194.e8 (2019).

11. Montgomery, S. B. et al. The origin, evolution, and functional impact of short insertion-deletion variants identified in 179 human genomes. Genome Res. 23, 749–761 (2013).

12. Mullaney, J. M., Mills, R. E., Pittard, W. S. & Devine, S. E. Small insertions and deletions (INDELs) in human genomes. Hum. Mol. Genet. 19, R131–6 (2010).

13. Stenson, P. D. et al. The Human Gene Mutation Database: building a comprehensive mutation repository for clinical and molecular genetics, diagnostic testing and personalized genomic medicine. Hum. Genet. 133, 1–9 (2014).

14. Zhang, X. et al. Impact of human pathogenic micro-insertions and micro-deletions on post-transcriptional regulation. Hum. Mol. Genet. 23, 3024–3034 (2014).

15. Rogalska, M. E., Vivori, C. & Valcárcel, J. Regulation of pre-mRNA splicing: roles in physiology and disease, and therapeutic prospects. Nat. Rev. Genet. 24, 251–269 (2023).

16. Qiu, J. et al. History of development of the life-saving drug ‘Nusinersen’ in spinal muscular atrophy. Front. Cell. Neurosci. 16, 942976 (2022).

17. Sheikh, O. & Yokota, T. Pharmacology and toxicology of eteplirsen and SRP-5051 for DMD exon 51 skipping: an update. Arch. Toxicol. 96, 1–9 (2022).

18. Kim, J. et al. A framework for individualized splice-switching oligonucleotide therapy. Nature 619, 828–836 (2023).

19. Popplewell, L. J., Trollet, C., Dickson, G. & Graham, I. R. Design of phosphorodiamidate morpholino oligomers (PMOs) for the induction of exon skipping of the human DMD gene. Mol. Ther. 17, 554–561 (2009).

20. Harding, P. L., Fall, A. M., Honeyman, K., Fletcher, S. & Wilton, S. D. The influence of antisense oligonucleotide length on dystrophin exon skipping. Mol. Ther. 15, 157–166 (2007).

21. Pramono, Z. A. D. et al. A prospective study in the rational design of efficient antisense oligonucleotides for exon skipping in the DMD gene. Hum. Gene Ther. 23, 781–790 (2012).

22. Wee, K. B. et al. Dynamics of co-transcriptional pre-mRNA folding influences the induction of dystrophin exon skipping by antisense oligonucleotides. PLoS One 3, e1844 (2008).

23. Aartsma-Rus, A., Houlleberghs, H., van Deutekom, J. C. T., van Ommen, G.-J. B. & ’t Hoen, P. A. C. Exonic sequences provide better targets for antisense oligonucleotides than splice site sequences in the modulation of Duchenne muscular dystrophy splicing. Oligonucleotides 20, 69–77 (2010).

24. Papoff, G. et al. An N-terminal domain shared by Fas/Apo-1 (CD95) soluble variants prevents cell death in vitro. J. Immunol. 156, 4622–4630 (1996).

25. Liu, C., Cheng, J. & Mountz, J. D. Differential expression of human Fas mRNA species upon peripheral blood mononuclear cell activation. Biochem. J 310 **(Pt** **3**), 957–963 (1995).

26. Cheng, J. et al. Protection from Fas-mediated apoptosis by a soluble form of the Fas molecule. Science 263, 1759–1762 (1994).

27. Cascino, I., Fiucci, G., Papoff, G. & Ruberti, G. Three functional soluble forms of the human apoptosis-inducing Fas molecule are produced by alternative splicing. J. Immunol. 154, 2706–2713 (1995).

28. Agrebi, N. et al. Rare splicing defects of FAS underly severe recessive autoimmune lymphoproliferative syndrome. Clin. Immunol. 183, 17–23 (2017).

29. Ustianenko, D., Weyn-Vanhentenryck, S. M. & Zhang, C. Microexons: discovery, regulation, and function. Wiley Interdiscip. Rev. RNA 8, (2017).

30. Liu, H. X., Zhang, M. & Krainer, A. R. Identification of functional exonic splicing enhancer motifs recognized by individual SR proteins. Genes Dev. 12, 1998–2012 (1998).

31. Jain, N., Lin, H.-C., Morgan, C. E., Harris, M. E. & Tolbert, B. S. Rules of RNA specificity of hnRNP A1 revealed by global and quantitative analysis of its affinity distribution. Proc. Natl. Acad. Sci. U. S. A. 114, 2206–2211 (2017).

32. Yu, Y. et al. Dynamic regulation of alternative splicing by silencers that modulate 5’ splice site competition. Cell 135, 1224–1236 (2008).

33. Vo, T. et al. HNRNPH1 destabilizes the G-quadruplex structures formed by G-rich RNA sequences that regulate the alternative splicing of an oncogenic fusion transcript. Nucleic Acids Res. 50, 6474–6496 (2022).

34. Paronetto, M. P. et al. Regulation of FAS exon definition and apoptosis by the Ewing sarcoma protein. Cell Rep. 7, 1211–1226 (2014).

35. Izquierdo, J. M. et al. Regulation of Fas alternative splicing by antagonistic effects of TIA-1 and PTB on exon definition. Mol. Cell 19, 475–484 (2005).

36. Choi, N. et al. SRSF6 Regulates the Alternative Splicing of the Apoptotic Fas Gene by Targeting a Novel RNA Sequence. Cancers 14, (2022).

37. Berget, S. M. Exon recognition in vertebrate splicing. J. Biol. Chem. 270, 2411–2414 (1995).

38. Ule, J. & Blencowe, B. J. Alternative Splicing Regulatory Networks: Functions, Mechanisms, and Evolution. Mol. Cell 76, 329–345 (2019).

39. Dominski, Z. & Kole, R. Selection of splice sites in pre-mRNAs with short internal exons. Mol. Cell. Biol. 11, 6075–6083 (1991).

40. Irimia, M. et al. A highly conserved program of neuronal microexons is misregulated in autistic brains. Cell 159, 1511–1523 (2014).

41. Juan-Mateu, J. et al. Pancreatic microexons regulate islet function and glucose homeostasis. Nat Metab 5, 219–236 (2023).

42. Gao, K., Masuda, A., Matsuura, T. & Ohno, K. Human branch point consensus sequence is yUnAy. Nucleic Acids Res. 36, 2257–2267 (2008).

43. Izquierdo, J. M. & Valcárcel, J. Fas-activated serine/threonine kinase (FAST K) synergizes with TIA-1/TIAR proteins to regulate Fas alternative splicing. J. Biol. Chem. 282, 1539–1543 (2007).

44. Singh, R., Valcárcel, J. & Green, M. R. Distinct binding specificities and functions of higher eukaryotic polypyrimidine tract-binding proteins. Science 268, 1173–1176 (1995).

45. Gonatopoulos-Pournatzis, T. et al. Genome-wide CRISPR-Cas9 Interrogation of Splicing Networks Reveals a Mechanism for Recognition of Autism-Misregulated Neuronal Microexons. Mol. Cell 72, 510–524.e12 (2018).

46. Förch, P. et al. The apoptosis-promoting factor TIA-1 is a regulator of alternative pre-mRNA splicing. Mol. Cell 6, 1089–1098 (2000).

47. Schott, G. et al. U2AF2 binds IL7R exon 6 ectopically and represses its inclusion. RNA 27, 571–583 (2021).

48. Corvelo, A., Hallegger, M., Smith, C. W. J. & Eyras, E. Genome-wide association between branch point properties and alternative splicing. PLoS Comput. Biol. 6, e1001016 (2010).

49. Weng, L. et al. A novel alternative spliced chondrolectin isoform lacking the transmembrane domain is expressed during T cell maturation. J. Biol. Chem. 278, 19164–19170 (2003).

50. Excoffon, K. J. D. A., Bowers, J. R. & Sharma, P. 1. Alternative splicing of viral receptors: A review of the diverse morphologies and physiologies of adenoviral receptors. Recent Res Dev Virol 9, 1–24 (2014).

51. Rosenberg, A. B., Patwardhan, R. P., Shendure, J. & Seelig, G. Learning the sequence determinants of alternative splicing from millions of random sequences. Cell 163, 698– 711 (2015).

52. Cheng, J. et al. MMSplice: modular modeling improves the predictions of genetic variant effects on splicing. Genome Biol. 20, 48 (2019).

53. Jaganathan, K. et al. Predicting Splicing from Primary Sequence with Deep Learning. Cell 176, 535–548.e24 (2019).

54. Zeng, T. & Li, Y. I. Predicting RNA splicing from DNA sequence using Pangolin. Genome Biol. 23, 103 (2022).

55. Hall, K. B. RNA-protein interactions. Curr. Opin. Struct. Biol. 12, 283–288 (2002).

56. Ray, D. et al. A compendium of RNA-binding motifs for decoding gene regulation. Nature 499, 172–177 (2013).

57. Pan, X., Rijnbeek, P., Yan, J. & Shen, H.-B. Prediction of RNA-protein sequence and structure binding preferences using deep convolutional and recurrent neural networks. BMC Genomics 19, 511 (2018).

58. Schaal, T. D. & Maniatis, T. Multiple distinct splicing enhancers in the protein-coding sequences of a constitutively spliced pre-mRNA. Mol. Cell. Biol. 19, 261–273 (1999).

59. Fairbrother, W. G., Yeh, R.-F., Sharp, P. A. & Burge, C. B. Predictive identification of exonic splicing enhancers in human genes. Science 297, 1007–1013 (2002).

60. Senapathy, P., Shapiro, M. B. & Harris, N. L. Splice junctions, branch point sites, and exons: sequence statistics, identification, and applications to genome project. Methods Enzymol. 183, 252–278 (1990).

61. Sorek, R. & Ast, G. Intronic sequences flanking alternatively spliced exons are conserved between human and mouse. Genome Res. 13, 1631–1637 (2003).

62. Zheng, C. L., Fu, X.-D. & Gribskov, M. Characteristics and regulatory elements defining constitutive splicing and different modes of alternative splicing in human and mouse. RNA 11, 1777–1787 (2005).

63. Havens, M. A. & Hastings, M. L. Splice-switching antisense oligonucleotides as therapeutic drugs. Nucleic Acids Res. 44, 6549–6563 (2016).

64. Faure, A. J., Schmiedel, J. M., Baeza-Centurion, P. & Lehner, B. DiMSum: an error model and pipeline for analyzing deep mutational scanning data and diagnosing common experimental pathologies. Genome Biol. 21, 207 (2020).

65. Baeza-Centurion, P., Miñana, B., Schmiedel, J. M., Valcárcel, J. & Lehner, B. Combinatorial Genetics Reveals a Scaling Law for the Effects of Mutations on Splicing. Cell 176, 549–563.e23 (2019).

66. Saraiva-Agostinho, N. & Barbosa-Morais, N. L. psichomics: graphical application for alternative splicing quantification and analysis. Nucleic Acids Res. 47, e7 (2019).

67. Dönmez, G., Hartmuth, K., Kastner, B., Will, C. L. & Lührmann, R. The 5’ end of U2 snRNA is in close proximity to U1 and functional sites of the pre-mRNA in early spliceosomal complexes. Mol. Cell 25, 399–411 (2007).

68. Osorio, D., Rondon-Villarreal, P. & Torres, R. Peptides: A Package for Data Mining of Antimicrobial Peptides. R J. 7, 4–14 (2015).

69. Durinck, S., Spellman, P. T., Birney, E. & Huber, W. Mapping identifiers for the integration of genomic datasets with the R/Bioconductor package biomaRt. Nat. Protoc. 4, 1184–1191 (2009).

