## Supplementary material for "Deep indel mutagenesis reveals the regulatory and modulatory architecture of alternative exon splicing": Baeza Minana et al Supplementary Table 1

| sequence | id | es | standard.error | psi |
|---|---|---|---|---|
| AACTTGGGGTGGCTTTGTCTTCTTCTTTTGCCAATTCCACTAATTGTTTGGG | Del_k11_1-11 | -4.060476526335533 | 0.3787933194834319 | 0.846523351764335 |
| AATCCAGATCTAACTTGGGGTGGCTTTGTCTTCTTCTTTTGCCAATTCCACTAATTGTTTGGG | G1A | -1.7826685651407692 | 0.04393235340578865 | 8.258066412720028 |
| AATTCCACTAATTGTTTGGG | Del_k43_1-43 | -4.127877339140709 | 0.09765651935818316 | 0.7913473306722291 |
| AATTGTTTGGG | Del_k52_1-52 | -6.846380662814996 | 0.6423745979493324 | 0.05220789094369205 |
| ACTAATTGTTTGGG | Del_k49_1-49 | -5.596628398009806 | 0.2262792916551992 | 0.18217830691004733 |
| ACTTGGGGTGGCTTTGTCTTCTTCTTTTGCCAATTCCACTAATTGTTTGGG | Del_k12_1-12 | -5.307860854252772 | 0.43302189927650503 | 0.24316821813212 |
| AGATCTAACTTGGGGTGGCTTTGTCTTCTTCTTTTGCCAATTCCACTAATTGTTTGGG | Del_k5_1-5 | -4.252389121246509 | 0.3161561182396023 | 0.6987026040441703 |
| ATCCAGATCTAACTTGGGGTGGCTTTGTCTTCTTCTTTTGCCAATTCCACTAATTGTTTGGG | Del_k1_1-1 | -0.9994971490403872 | 0.033973010186080316 | 18.071965782406128 |
| ATCTAACTTGGGGTGGCTTTGTCTTCTTCTTTTGCCAATTCCACTAATTGTTTGGG | Del_k7_1-7 | -1.9453761145056088 | 0.07573908089064428 | 7.0180325855889585 |
| ATTCCACTAATTGTTTGGG | Del_k44_1-44 | -2.5570045637430296 | 0.04286682303885742 | 3.8070494671739015 |
| ATTGTTTGGG | Del_k53_1-53 | -5.6892314390018 | 0.18222847586794835 | 0.1660655973038775 |
| CAATTCCACTAATTGTTTGGG | Del_k42_1-42 | -2.336634909063891 | 0.03913429960336808 | 4.745629673262323 |
| CACTAATTGTTTGGG | Del_k48_1-48 | -6.010504304325917 | 0.18240278725126652 | 0.12043497846043554 |
| CAGATCTAACTTGGGGTGGCTTTGTCTTCTTCTTTTGCCAATTCCACTAATTGTTTGGG | Del_k4_1-4 | 0.48963834882314883 | 0.21968836150954718 | 80.1177464825123 |
| CATCCAGATCTAACTTGGGGTGGCTTTGTCTTCTTCTTTTGCCAATTCCACTAATTGTTTGGG | G1C | -2.734503282287977 | 0.09260965373834638 | 3.187878831474206 |
| CCAATTCCACTAATTGTTTGGG | Del_k41_1-41 | -0.2851079160708989 | 0.02290333809563668 | 36.9199154198815 |
| CCACTAATTGTTTGGG | Del_k47_1-47 | -6.223151527283445 | 0.37294035316889207 | 0.09736460891923224 |
| CCAGATCTAACTTGGGGTGGCTTTGTCTTCTTCTTTTGCCAATTCCACTAATTGTTTGGG | Del_k3_1-3 | -3.092097280392142 | 0.13473360315273747 | 2.229465240856452 |
| CTAACTTGGGGTGGCTTTGTCTTCTTCTTTTGCCAATTCCACTAATTGTTTGGG | Del_k9_1-9 | -4.8826670753822 | 0.5718348376910144 | 0.3720198594254272 |
| CTAATTGTTTGGG | Del_k50_1-50 | -4.599586337559192 | 0.11373908789110301 | 0.4937493383794371 |
| CTTCTTCTTTTGCCAATTCCACTAATTGTTTGGG | Del_k29_1-29 | -4.919020532966754 | 0.2020526880559575 | 0.35873852511347254 |
| CTTCTTTTGCCAATTCCACTAATTGTTTGGG | Del_k32_1-32 | -5.289407094437264 | 0.24150068739879266 | 0.24769724629726014 |
| CTTGGGGTGGCTTTGTCTTCTTCTTTTGCCAATTCCACTAATTGTTTGGG | Del_k13_1-13 | -4.426482030997082 | 0.483137360099877 | 0.5870630841455571 |
| CTTTGTCTTCTTCTTTTGCCAATTCCACTAATTGTTTGGG | Del_k23_1-23 | -4.68531698466534 | 0.2633165134699821 | 0.45318359444036516 |
| CTTTTGCCAATTCCACTAATTGTTTGGG | Del_k35_1-35 | -4.530414794182313 | 0.1423869504603592 | 0.5291116772019473 |
| GAAAATCCAGATCTAACTTGGGGTGGCTTTGTCTTCTTCTTTTGCCAATTCCACTAATTGTTTGGG | Ins_k3_1_AAA;Ins_k3_2_AAA | 0.09635684362696817 | 0.03915340271701162 | 54.06655990750198 |
| GAAACTCCAGATCTAACTTGGGGTGGCTTTGTCTTCTTCTTTTGCCAATTCCACTAATTGTTTGGG | Ins_k3_2_AAC | 0.39223366013783223 | 0.0338159835136011 | 72.68192251392976 |
| GAAACTTGGGGTGGCTTTGTCTTCTTCTTTTGCCAATTCCACTAATTGTTTGGG | Del_k9_3-11 | -1.5049105669055918 | 0.0679001490520234 | 10.902024085300566 |
| GAAAGTCCAGATCTAACTTGGGGTGGCTTTGTCTTCTTCTTTTGCCAATTCCACTAATTGTTTGGG | Ins_k3_2_AAG | -0.3582695069070333 | 0.04732987345465192 | 34.31523854267865 |
| GAAATCCAGATCTAACTTGGGGTGGCTTTGTCTTCTTCTTTTGCCAATTCCACTAATTGTTTGGG | Ins_k2_1_AA;Ins_k2_2_AA | -0.04249302255217191 | 0.04423322722578412 | 47.057300195790056 |
| GAAATTCCACTAATTGTTTGGG | Del_k41_3-43 | -5.95159670168647 | 0.2973580482839166 | 0.12774263964186777 |
| GAAATTCCAGATCTAACTTGGGGTGGCTTTGTCTTCTTCTTTTGCCAATTCCACTAATTGTTTGGG | Ins_k3_2_AAT | 0.15996320334824554 | 0.040098479822945836 | 57.61726360430681 |
| GAAATTGTTTGGG | Del_k50_3-52 | -6.003470406374626 | 0.3345886242811709 | 0.12128509211002547 |
| GAACATCCAGATCTAACTTGGGGTGGCTTTGTCTTCTTCTTTTGCCAATTCCACTAATTGTTTGGG | Ins_k3_1_AAC;Ins_k3_2_ACA | -0.5722358478671377 | 0.09060111021003624 | 27.70528493669677 |
| GAACCAGATCTAACTTGGGGTGGCTTTGTCTTCTTCTTTTGCCAATTCCACTAATTGTTTGGG | T3A | -0.5639331305088365 | 0.04560833476761379 | 27.93627166870884 |
| GAACCTCCAGATCTAACTTGGGGTGGCTTTGTCTTCTTCTTTTGCCAATTCCACTAATTGTTTGGG | Ins_k3_2_ACC | 0.13784055000235507 | 0.04165108653422855 | 56.35661274024162 |
| GAACGTCCAGATCTAACTTGGGGTGGCTTTGTCTTCTTCTTTTGCCAATTCCACTAATTGTTTGGG | Ins_k3_2_ACG | 0.4005379423604043 | 0.03754305791112969 | 73.28800677532726 |
| GAACTAATTGTTTGGG | Del_k47_3-49 | -5.2110557038402074 | 0.6606634796606345 | 0.26788522127332415 |
| GAACTATCCAGATCTAACTTGGGGTGGCTTTGTCTTCTTCTTTTGCCAATTCCACTAATTGTTTGGG | Ins_k4_1_AACT;Ins_k4_2_ACTA | -0.3917866086385149 | 0.05841175236225112 | 33.184152451457145 |
| GAACTCCAGATCTAACTTGGGGTGGCTTTGTCTTCTTCTTTTGCCAATTCCACTAATTGTTTGGG | Ins_k2_2_AC | 0.014626127615474438 | 0.0523416160578550 4 | 49.82342038903893 |
| GAACTTCCAGATCTAACTTGGGGTGGCTTTGTCTTCTTCTTTTGCCAATTCCACTAATTGTTTGGG | Ins_k3_2_ACT | 0.15163708597296877 | 0.03396077407086373 | 57.13952710964929 |
| GAACTTGGGGTGGCTTTGTCTTCTTCTTTTGCCAATTCCACTAATTGTTTGGG | Del_k10_2-11;Del_k10_3-12 | -2.6008188039262405 | 0.15200037963526308 | 3.643847871556039 |
| GAAGATCCAGATCTAACTTGGGGTGGCTTTGTCTTCTTCTTTTGCCAATTCCACTAATTGTTTGGG | Ins_k3_1_AAG;Ins_k3_2_AGA | 0.5255075916391676 | 0.03558551629578763 | 83.0436710322543 |
| GAAGATCTAACTTGGGGTGGCTTTGTCTTCTTCTTTTGCCAATTCCACTAATTGTTTGGG | Del_k3_3-5 | -2.4193806572973386 | 0.09577974318690309 | 4.368756336890245 |
| GAAGCTCCAGATCTAACTTGGGGTGGCTTTGTCTTCTTCTTTTGCCAATTCCACTAATTGTTTGGG | Ins_k3_2_AGC | 0.4268572485813218 | 0.03369866084055943 | 75.24250394978446 |
| GAAGGTCCAGATCTAACTTGGGGTGGCTTTGTCTTCTTCTTTTGCCAATTCCACTAATTGTTTGGG | Ins_k3_2_AGG | -0.16718088065686193 | 0.04035373625811326 | 41.54088629425198 |
| GAAGTCCAGATCTAACTTGGGGTGGCTTTGTCTTCTTCTTTTGCCAATTCCACTAATTGTTTGGG | Ins_k2_2_AG | -0.4816900630379076 | 0.048989536468241515 | 30.330959961574152 |
| GAAGTTCCAGATCTAACTTGGGGTGGCTTTGTCTTCTTCTTTTGCCAATTCCACTAATTGTTTGGG | Ins_k3_2_AGT | 0.556135574198583 | 0.03258729819059649 | 85.62648238850907 |
| GAATAGTCCAGATCTAACTTGGGGTGGCTTTGTCTTCTTCTTTTGCCAATTCCACTAATTGTTTGGG | Ins_k4_2_ATAG | -2.1643979879581803 | 0.13070960709177437 | 5.637614679267494 |
| GAATATCCAGATCTAACTTGGGGTGGCTTTGTCTTCTTCTTTTGCCAATTCCACTAATTGTTTGGG | Ins_k3_1_AAT;Ins_k3_2_ATA | 0.11313468571985175 | 0.03841441717924935 | 54.9813326094715 |
| GAATCCAGATCTAACTTGGGGTGGCTTTGTCTTCTTCTTTTGCCAATTCCACTAATTGTTTGGG | Ins_k1_1_A;Ins_k1_2_A | 0.3775537840533878 | 0.036889048242167974 | 71.6227541487438 |
| GAATCTAACTTGGGGTGGCTTTGTCTTCTTCTTTTGCCAATTCCACTAATTGTTTGGG | Del_k5_3-7 | -1.9888036224563301 | 0.07405785011706124 | 6.7197799760791135 |
| GAATCTCCAGATCTAACTTGGGGTGGCTTTGTCTTCTTCTTTTGCCAATTCCACTAATTGTTTGGG | Ins_k3_2_ATC | 0.3269985192689526 | 0.03822014646208385 | 68.0918517037593 |
| GAATGTCCAGATCTAACTTGGGGTGGCTTTGTCTTCTTCTTTTGCCAATTCCACTAATTGTTTGGG | Ins_k3_2_ATG | -0.05239347527646332 | 0.04393805272399936 | 46.59371028159407 |
| GAATTCCACTAATTGTTTGGG | Del_k42_2-43;Del_k42_3-44 | -3.2249267193285696 | 0.05318140810860312 | 1.9521519251548203 |
| GAATTCCAGATCTAACTTGGGGTGGCTTTGTCTTCTTCTTTTGCCAATTCCACTAATTGTTTGGG | Ins_k2_2_AT | 0.1924779849200262 | 0.04239251939719238 | 59.52146594777845 |
| GAATTGTTTGGG | Del_k51_2-52;Del_k51_3-53 | -6.17149445546217 | 0.3740589897632382 | 0.10252635201042823 |
| GAATTTCCAGATCTAACTTGGGGTGGCTTTGTCTTCTTCTTTTGCCAATTCCACTAATTGTTTGGG | Ins_k3_2_ATT | -0.016010123294416756 | 0.043980438965113035 | 48.320162253009784 |
| GACAATCCAGATCTAACTTGGGGTGGCTTTGTCTTCTTCTTTTGCCAATTCCACTAATTGTTTGGG | Ins_k3_1_ACA;Ins_k3_2_CAA | -0.126097250572477 | 0.03951845541611664 | 43.28307946812363 |
| GACAATTCCACTAATTGTTTGGG | Del_k40_3-42 | -5.035904590915741 | 0.139568862955193 | 0.3191654827137867 |
| GACACTAATTGTTTGGG | Del_k46_3-48 | -6.497736829007486 | 0.5764113876279124 | 0.07398611827845544 |
| GACACTCCAGATCTAACTTGGGGTGGCTTTGTCTTCTTCTTTTGCCAATTCCACTAATTGTTTGGG | Ins_k3_2_CAC | -2.089791433536692 | 0.09598650276009252 | 6.074305132868789 |
| GACAGATCTAACTTGGGGTGGCTTTGTCTTCTTCTTTTGCCAATTCCACTAATTGTTTGGG | Del_k2_3-4 | -2.432172768655204 | 0.11859213145288595 | 4.313226647547303 |
| GACAGTCCAGATCTAACTTGGGGTGGCTTTGTCTTCTTCTTTTGCCAATTCCACTAATTGTTTGGG | Ins_k3_2_CAG | -0.40021307853945975 | 0.049717881324567594 | 32.905702014371734 |
| GACATCCAGATCTAACTTGGGGTGGCTTTGTCTTCTTCTTTTGCCAATTCCACTAATTGTTTGGG | Ins_k2_1_AC;Ins_k2_2_CA | -0.2251452902515802 | 0.05248191217422721 | 39.2014503387266 |
| GACATTCCAGATCTAACTTGGGGTGGCTTTGTCTTCTTCTTTTGCCAATTCCACTAATTGTTTGGG | Ins_k3_2_CAT | -0.44197623919969575 | 0.04995673497908426 | 31.559756977040898 |
| GACCAATTCCACTAATTGTTTGGG | Del_k39_3-41 | -5.63384617203692 | 0.2592544558470218 | 0.17552265830892913 |
| GACCACTAATTGTTTGGG | Del_k45_3-47 | -7.119538394007085 | 0.6461801125721695 | 0.039728782292136035 |
| GACCAGATCTAACTTGGGGTGGCTTTGTCTTCTTCTTTTGCCAATTCCACTAATTGTTTGGG | Del_k1_3-3 | -0.6502098991991593 | 0.05045640651213793 | 25.627067973347483 |
| GACCATCCAGATCTAACTTGGGGTGGCTTTGTCTTCTTCTTTTGCCAATTCCACTAATTGTTTGGG | Ins_k3_1_ACC;Ins_k3_2_CCA | 0.3081210319994885 | 0.03804111303603793 | 66.8185052447228 |
| GACCCAGATCTAACTTGGGGTGGCTTTGTCTTCTTCTTTTGCCAATTCCACTAATTGTTTGGG | T3C | -0.34754200470748176 | 0.029792886773413164 | 34.68533690591117 |
| GACCCTCCAGATCTAACTTGGGGTGGCTTTGTCTTCTTCTTTTGCCAATTCCACTAATTGTTTGGG | Ins_k3_2_CCC | 0.04582361837077509 | 0.04229712035869662 | 51.40228635934978 |
| GACCGTCCAGATCTAACTTGGGGTGGCTTTGTCTTCTTCTTTTGCCAATTCCACTAATTGTTTGGG | Ins_k3_2_CCG | 0.8290676143089943 | 0.035808107358028 | 100 |

| | | | | |
|---|---|---|---|---|
| **GACCTCCAGATCTAACTTGGGGTGGCTTTGTCTTCTTCTTTTGCCAATTCCACTAATTGTTTGGG** | Ins_k2_2_CC | 0.141699592695113 | 0.05004476461315015 | 56.574515492416126 |
| **GACCTTCCAGATCTAACTTGGGGTGGCTTTGTCTTCTTCTTTTGCCAATTCCACTAATTGTTTGGG** | Ins_k3_2_CCT | 0.36724017019320887 | 0.03437777535325566 | 70.88786093550901 |
| **GACGATCCAGATCTAACTTGGGGTGGCTTTGTCTTCTTCTTTTGCCAATTCCACTAATTGTTTGGG** | Ins_k3_1_ACG;Ins_k3_2_CGA | 0.5583905860505392 | 0.03459931630025922 | 85.81978899356427 |
| **GACGCTCCAGATCTAACTTGGGGTGGCTTTGTCTTCTTCTTTTGCCAATTCCACTAATTGTTTGGG** | Ins_k3_2_CGC | 0.33203818673762664 | 0.03193160737121883 | 68.43587815499103 |
| **GACGGTCCAGATCTAACTTGGGGTGGCTTTGTCTTCTTCTTTTGCCAATTCCACTAATTGTTTGGG** | Ins_k3_2_CGG | 0.47983091638637276 | 0.035205156254163456 | 79.33583762341485 |
| **GACGTCCAGATCTAACTTGGGGTGGCTTTGTCTTCTTCTTTTGCCAATTCCACTAATTGTTTGGG** | Ins_k2_2_CG | 0.1004306600697612 | 0.037313844883631764 | 54.28726640189474 |
| **GACGTTCCAGATCTAACTTGGGGTGGCTTTGTCTTCTTCTTTTGCCAATTCCACTAATTGTTTGGG** | Ins_k3_2_CGT | 0.6435900298912789 | 0.03287223229217832 | 93.452105526062 |
| **GACTAACTTGGGGTGGCTTTGTCTTCTTCTTTTGCCAATTCCACTAATTGTTTGGG** | Del_k7_3-9 | -3.001370871646456 | 0.15024199664674723 | 2.441196195309315 |
| **GACTAATTGTTTGGG** | Del_k48_2-49;Del_k48_3-50 | -4.729671597844334 | 0.12242193757309104 | 0.43352207425961775 |
| **GACTATCCAGATCTAACTTGGGGTGGCTTTGTCTTCTTCTTTTGCCAATTCCACTAATTGTTTGGG** | Ins_k3_1_ACT;Ins_k3_2_CTA | -0.17258948337902735 | 0.03833517304274221 | 41.31681464705032 |
| **GACTCCAGATCTAACTTGGGGTGGCTTTGTCTTCTTCTTTTGCCAATTCCACTAATTGTTTGGG** | Ins_k1_2_C | 0.2516728896156841 | 0.035780630001115886 | 63.15120463687102 |
| **GACTCTCCAGATCTAACTTGGGGTGGCTTTGTCTTCTTCTTTTGCCAATTCCACTAATTGTTTGGG** | Ins_k3_2_CTC | 0.2575019276404106 | 0.036198013455699694 | 63.520390363486854 |
| **GACTGTCCAGATCTAACTTGGGGTGGCTTTGTCTTCTTCTTTTGCCAATTCCACTAATTGTTTGGG** | Ins_k3_2_CTG | -0.017020597654959557 | 0.03670251046264081 | 48.27136062850345 |
| **GACTTCCAGATCTAACTTGGGGTGGCTTTGTCTTCTTCTTTTGCCAATTCCACTAATTGTTTGGG** | Ins_k2_2_CT | -0.05000267873184694 | 0.042950968864900775 | 46.70523963200467 |
| **GACTTCTTCTTTTGCCAATTCCACTAATTGTTTGGG** | Del_k27_3-29 | -3.0587711444182193 | 0.07707401846106336 | 2.305016628177026 |
| **GACTTCTTTTGCCAATTCCACTAATTGTTTGGG** | Del_k30_3-32 | -4.494559647005613 | 0.13978011201620807 | 0.5484272666261497 |
| **GACTTGGGGTGGCTTTGTCTTCTTCTTTTGCCAATTCCACTAATTGTTTGGG** | Del_k11_2-12;Del_k11_3-13 | -2.0381741830650286 | 0.08350188556301595 | 6.3960771155257525 |
| **GACTTTCCAGATCTAACTTGGGGTGGCTTTGTCTTCTTCTTTTGCCAATTCCACTAATTGTTTGGG** | Ins_k3_2_CTT | -0.4701155980760513 | 0.04694461300594615 | 30.68406414887739 |
| **GACTTTGTCTTCTTCTTTTGCCAATTCCACTAATTGTTTGGG** | Del_k21_3-23 | -2.6397114156740114 | 0.07481782998399997 | 3.5048496339583624 |
| **GACTTTTGCCAATTCCACTAATTGTTTGGG** | Del_k33_3-35 | -4.893516280043292 | 0.18181214636934293 | 0.36800555521183276 |
| **GAG** | Del_k60_3-62 | -5.603578043252746 | 0.11415662991876216 | 0.18091662151703197 |
| **GAGAATCCAGATCTAACTTGGGGTGGCTTTGTCTTCTTCTTTTGCCAATTCCACTAATTGTTTGGG** | Ins_k3_1_AGA;Ins_k3_2_GAA | 0.40645137223120087 | 0.03365631691510489 | 73.72267418474122 |
| **GAGACTCCAGATCTAACTTGGGGTGGCTTTGTCTTCTTCTTTTGCCAATTCCACTAATTGTTTGGG** | Ins_k3_2_GAC | 0.33479063735002246 | 0.0349267353271695 | 68.62450400230755 |
| **GAGAGTCCAGATCTAACTTGGGGTGGCTTTGTCTTCTTCTTTTGCCAATTCCACTAATTGTTTGGG** | Ins_k3_2_GAG | -0.03242786137746634 | 0.03953236169303045 | 47.53333115207258 |
| **GAGATCCAGATCTAACTTGGGGTGGCTTTGTCTTCTTCTTTTGCCAATTCCACTAATTGTTTGGG** | Ins_k2_1_AG;Ins_k2_2_GA | 0.41450996398379325 | 0.03596560121346356 | 74.31917537056681 |
| **GAGATCTAACTTGGGGTGGCTTTGTCTTCTTCTTTTGCCAATTCCACTAATTGTTTGGG** | Del_k4_2-5;Del_k4_3-6 | -2.5690103368973043 | 0.10126171470264472 | 3.7616161715274505 |
| **GAGATTCCAGATCTAACTTGGGGTGGCTTTGTCTTCTTCTTTTGCCAATTCCACTAATTGTTTGGG** | Ins_k3_2_GAT | 0.6084069898126576 | 0.036306215935866006 | 90.2213436233437 |
| **GAGCATCCAGATCTAACTTGGGGTGGCTTTGTCTTCTTCTTTTGCCAATTCCACTAATTGTTTGGG** | Ins_k3_1_AGC;Ins_k3_2_GCA | 0.2883774435824945 | 0.032590224922451824 | 65.51220612826916 |
| **GAGCCAATTCCACTAATTGTTTGGG** | Del_k38_3-40 | -5.659961574910675 | 0.21442502125523646 | 0.17099815019515172 |
| **GAGCCAGATCTAACTTGGGGTGGCTTTGTCTTCTTCTTTTGCCAATTCCACTAATTGTTTGGG** | T3G | -0.3602077653172824 | 0.03644033407191426 | 34.24879115987559 |
| **GAGCCTCCAGATCTAACTTGGGGTGGCTTTGTCTTCTTCTTTTGCCAATTCCACTAATTGTTTGGG** | Ins_k3_2_GCC | 0.4518694373598582 | 0.0374353684569154 | 77.14821734263795 |
| **GAGCGTCCAGATCTAACTTGGGGTGGCTTTGTCTTCTTCTTTTGCCAATTCCACTAATTGTTTGGG** | Ins_k3_2_GCG | 0.6471489865187607 | 0.03232909451336061 | 93.78529005934658 |
| **GAGCTCCAGATCTAACTTGGGGTGGCTTTGTCTTCTTCTTTTGCCAATTCCACTAATTGTTTGGG** | Ins_k2_2_GC | 0.1986744847306403 | 0.03915410970453119 | 59.89143577543666 |
| **GAGCTTCCAGATCTAACTTGGGGTGGCTTTGTCTTCTTCTTTTGCCAATTCCACTAATTGTTTGGG** | Ins_k3_2_GCT | 0.2209248652188711 | 0.034947993492770495 | 61.23897909054651 |
| **GAGCTTTGTCTTCTTCTTTTGCCAATTCCACTAATTGTTTGGG** | Del_k20_3-22 | -2.5255019566152344 | 0.06842777369621708 | 3.9288905301904906 |
| **GAGGATCCAGATCTAACTTGGGGTGGCTTTGTCTTCTTCTTTTGCCAATTCCACTAATTGTTTGGG** | Ins_k3_1_AGG;Ins_k3_2_GGA | 0.5779580030465159 | 0.03642781482178226 | 87.51559778236746 |
| **GAGGCTCCAGATCTAACTTGGGGTGGCTTTGTCTTCTTCTTTTGCCAATTCCACTAATTGTTTGGG** | Ins_k3_2_GGC | 0.3735066236753516 | 0.0356471932360508 | 71.33347115817519 |
| **GAGGCTTTGTCTTCTTCTTTTGCCAATTCCACTAATTGTTTGGG** | Del_k19_3-21 | -2.3340562891278607 | 0.07321448656329096 | 4.757882639629037 |
| **GAGGGGTGGCTTTGTCTTCTTCTTTTGCCAATTCCACTAATTGTTTGGG** | Del_k14_3-16 | -2.429921792789557 | 0.08943051420199812 | 4.322946552165211 |
| **GAGGGTCCAGATCTAACTTGGGGTGGCTTTGTCTTCTTCTTTTGCCAATTCCACTAATTGTTTGGG** | Ins_k3_2_GGG | -1.6953256995643198 | 0.0839088002693778 | 9.011786532275659 |
| **GAGGGTGGCTTTGTCTTCTTCTTTTGCCAATTCCACTAATTGTTTGGG** | Del_k15_3-17 | -3.318289322704298 | 0.12224338307031295 | 1.7781432681297749 |
| **GAGGTCCAGATCTAACTTGGGGTGGCTTTGTCTTCTTCTTTTGCCAATTCCACTAATTGTTTGGG** | Ins_k2_2_GG | -0.3129687795071228 | 0.053617420777235116 | 35.90549167678055 |
| **GAGGTGGCTTTGTCTTCTTCTTTTGCCAATTCCACTAATTGTTTGGG** | Del_k16_3-18 | -0.6970249155097493 | 0.03667753707337642 | 24.45498594570099 |
| **GAGGTTCCAGATCTAACTTGGGGTGGCTTTGTCTTCTTCTTTTGCCAATTCCACTAATTGTTTGGG** | Ins_k3_2_GGT | 0.36575007383868574 | 0.03462859734924432 | 70.78230985252141 |
| **GAGTATCCAGATCTAACTTGGGGTGGCTTTGTCTTCTTCTTTTGCCAATTCCACTAATTGTTTGGG** | Ins_k3_1_AGT;Ins_k3_2_GTA | 0.4029258472174544 | 0.032313571486007955 | 73.46322067647527 |
| **GAGTCCAGATCTAACTTGGGGTGGCTTTGTCTTCTTCTTTTGCCAATTCCACTAATTGTTTGGG** | Ins_k1_2_G | 0.2145477708224518 | 0.042080116448526377 | 60.849694910389246 |
| **GAGTCGTCCAGATCTAACTTGGGGTGGCTTTGTCTTCTTCTTTTGCCAATTCCACTAATTGTTTGGG** | Ins_k4_2_GTCG | 0.8927415811002957 | 0.032162392983047 | 100 |
| **GAGTCTCCAGATCTAACTTGGGGTGGCTTTGTCTTCTTCTTTTGCCAATTCCACTAATTGTTTGGG** | Ins_k3_2_GTC | 0.39979908690337407 | 0.03755558578014805 | 73.23387753089273 |
| **GAGTCTTCTTCTTTTGCCAATTCCACTAATTGTTTGGG** | Del_k25_3-27 | -2.0710410758930005 | 0.052940786265920185 | 6.189275021444298 |
| **GAGTGGCTTTGTCTTCTTCTTTTGCCAATTCCACTAATTGTTTGGG** | Del_k17_3-19 | -1.6789301773554437 | 0.04982691303189422 | 9.160757368741091 |
| **GAGTGTCCAGATCTAACTTGGGGTGGCTTTGTCTTCTTCTTTTGCCAATTCCACTAATTGTTTGGG** | Ins_k3_2_GTG | 0.19479450411449328 | 0.03253966684506783 | 59.659508393385764 |
| **GAGTTCCAGATCTAACTTGGGGTGGCTTTGTCTTCTTCTTTTGCCAATTCCACTAATTGTTTGGG** | Ins_k2_2_GT | 0.5803329489805995 | 0.038180258674166176 | 87.72368960108946 |
| **GAGTTTCCAGATCTAACTTGGGGTGGCTTTGTCTTCTTCTTTTGCCAATTCCACTAATTGTTTGGG** | Ins_k3_2_GTT | 0.41659288802909256 | 0.033937169468069316 | 74.47413789953549 |
| **GATAAACCAGATCTAACTTGGGGTGGCTTTGTCTTCTTCTTTTGCCAATTCCACTAATTGTTTGGG** | Ins_k3_3_AAA | -0.9233895097519496 | 0.055997744129395004 | 19.50107368939519 |
| **GATAACCAGATCTAACTTGGGGTGGCTTTGTCTTCTTCTTTTGCCAATTCCACTAATTGTTTGGG** | Ins_k2_3_AA | -0.8103075623529329 | 0.06685734863315189 | 21.835814144289035 |
| **GATAACCCAGATCTAACTTGGGGTGGCTTTGTCTTCTTCTTTTGCCAATTCCACTAATTGTTTGGG** | Ins_k3_3_AAC | -0.7318381814991158 | 0.055971979584989937 | 23.61827680746986 |
| **GATAACTTGGGGTGGCTTTGTCTTCTTCTTTTGCCAATTCCACTAATTGTTTGGG** | Del_k8_3-10;Del_k8_4-11 | -2.4615360739438716 | 0.11207665623961266 | 4.188417429750552 |
| **GATAAGCCAGATCTAACTTGGGGTGGCTTTGTCTTCTTCTTTTGCCAATTCCACTAATTGTTTGGG** | Ins_k3_3_AAG | -1.6013406267209493 | 0.076792380072251 | 9.899838145619002 |
| **GATAATCCAGATCTAACTTGGGGTGGCTTTGTCTTCTTCTTTTGCCAATTCCACTAATTGTTTGGG** | Ins_k3_1_ATA;Ins_k3_2_TAA;Ins_ | -0.2924150990141443 | 0.04210860417489794 | 36.65111811523235 |
| **GATAATTCCACTAATTGTTTGGG** | Del_k40_4-43 | -6.539153085113779 | 0.40933532106364845 | 0.07098447766258249 |
| **GATAATTGTTTGGG** | Del_k49_3-51;Del_k49_4-52 | -6.133299111470242 | 0.35210686990916884 | 0.10651812967079459 |
| **GATACACCAGATCTAACTTGGGGTGGCTTTGTCTTCTTCTTTTGCCAATTCCACTAATTGTTTGGG** | Ins_k3_3_ACA | -8.095929769791343e-5 | 0.4275991064871687 | 49.096025059389405 |
| **GATACAGATCTAACTTGGGGTGGCTTTGTCTTCTTCTTTTGCCAATTCCACTAATTGTTTGGG** | C4A | -1.8120410407356797 | 0.056660898118058835 | 8.019034227088833 |
| **GATACCAGATCTAACTTGGGGTGGCTTTGTCTTCTTCTTTTGCCAATTCCACTAATTGTTTGGG** | Ins_k1_3_A | -0.06186109915475002 | 0.03846124605927273 | 46.15466021753833 |
| **GATACCCAGATCTAACTTGGGGTGGCTTTGTCTTCTTCTTTTGCCAATTCCACTAATTGTTTGGG** | Ins_k2_3_AC | -0.8049967197152581 | 0.054264599759629814 | 21.95208920304956 |
| **GATACCCCAGATCTAACTTGGGGTGGCTTTGTCTTCTTCTTTTGCCAATTCCACTAATTGTTTGGG** | Ins_k3_3_ACC | -2.9559771850542753 | 0.15888416434350458 | 2.554564731741539 |
| **GATACGCCAGATCTAACTTGGGGTGGCTTTGTCTTCTTCTTTTGCCAATTCCACTAATTGTTTGGG** | Ins_k3_3_ACG | 0.449733663888253 | 0.03520482522129408 | 76.98362205834395 |
| **GATACTAATTGTTTGGG** | Del_k46_4-49 | -7.301261291829325 | 1.3384452204562938 | 0.03312714450396789 |
| **GATACTCCAGATCTAACTTGGGGTGGCTTTGTCTTCTTCTTTTGCCAATTCCACTAATTGTTTGGG** | Ins_k3_2_TAC;Ins_k3_3_ACT | 0.1534800211171542 | 0.03639347895506075 | 57.24492864654787 |
| **GATACTTCCAGATCTAACTTGGGGTGGCTTTGTCTTCTTCTTTTGCCAATTCCACTAATTGTTTGGG** | Ins_k4_2_TACT;Ins_k4_3_ACTT | 0.3799485373453698 | 0.03800091097984096 | 71.79447851169638 |
| **GATACTTGGGGTGGCTTTGTCTTCTTCTTTTGCCAATTCCACTAATTGTTTGGG** | Del_k9_4-12 | -1.9599613441609736 | 0.07633859158991667 | 6.916415821912519 |
| **GATAGACCAGATCTAACTTGGGGTGGCTTTGTCTTCTTCTTTTGCCAATTCCACTAATTGTTTGGG** | Ins_k3_3_AGA | -1.1813690424040046 | 0.05482467430796354 | 15.066744926547594 |
| **GATAGATCTAACTTGGGGTGGCTTTGTCTTCTTCTTTTGCCAATTCCACTAATTGTTTGGG** | Del_k2_4-5 | -4.343448663137837 | 0.27367256406219365 | 0.6378898698436974 |
| **GATAGCCAGATCTAACTTGGGGTGGCTTTGTCTTCTTCTTTTGCCAATTCCACTAATTGTTTGGG** | Ins_k2_3_AG | -2.5621393496263565 | 0.11272941123158065 | 3.7875511859034354 |
| **GATAGCCCAGATCTAACTTGGGGTGGCTTTGTCTTCTTCTTTTGCCAATTCCACTAATTGTTTGGG** | Ins_k3_3_AGC | -1.6510924809158813 | 0.07641284763626978 | 9.419354425213744 |

| | | | | |
|---|---|---|---|---|
| **GATAGGCCAGATCTAACTTGGGGTGGCTTTGTCTTCTTCTTTTGCCAATTCCACTAATTGTTTGGG** | Ins_k3_3_AGG | -3.0563404601135287 | 0.16421402755782347 | 2.3106262107174156 |
| **GATAGTCCAGATCTAACTTGGGGTGGCTTTGTCTTCTTCTTTTGCCAATTCCACTAATTGTTTGGG** | Ins_k3_2_TAG;Ins_k3_3_AGT | -1.832371914506219 | 0.08372829662752924 | 7.857646391339872 |
| **GATATACCAGATCTAACTTGGGGTGGCTTTGTCTTCTTCTTTTGCCAATTCCACTAATTGTTTGGG** | Ins_k3_3_ATA | -0.5423655624086646 | 0.05001347018686494 | 28.54533349248624 |
| **GATATCCAGATCTAACTTGGGGTGGCTTTGTCTTCTTCTTTTGCCAATTCCACTAATTGTTTGGG** | Ins_k2_1_AT;Ins_k2_2_TA;Ins_k2_ | -0.004249382694154091 | 0.04166117338204072 | 48.891797988026596 |
| **GATATCCCAGATCTAACTTGGGGTGGCTTTGTCTTCTTCTTTTGCCAATTCCACTAATTGTTTGGG** | Ins_k3_3_ATC | -0.5817322511944659 | 0.05030895528011745 | 27.44342968605193 |
| **GATATCTAACTTGGGGTGGCTTTGTCTTCTTCTTTTGCCAATTCCACTAATTGTTTGGG** | Del_k4_4-7 | -2.926734154654495 | 0.16539856080613435 | 2.6303709454624244 |
| **GATATGCCAGATCTAACTTGGGGTGGCTTTGTCTTCTTCTTTTGCCAATTCCACTAATTGTTTGGG** | Ins_k3_3_ATG | -0.1277059383874572 | 0.038913540420596825 | 43.213506481200454 |
| **GATATTCCACTAATTGTTTGGG** | Del_k41_4-44 | -6.266022071500408 | 0.3641834872226705 | 0.09327874256422779 |
| **GATATTCCAGATCTAACTTGGGGTGGCTTTGTCTTCTTCTTTTGCCAATTCCACTAATTGTTTGGG** | Ins_k3_2_TAT;Ins_k3_3_ATT | -0.09466187599295106 | 0.04265566187554307 | 44.66531094334581 |
| **GATATTGTTTGGG** | Del_k50_4-53 | -5.77580641818197 | 0.5062531947724284 | 0.1522932434669133 |
| **GATC** | Del_k59_5-63 | -7.32071562852895 | 0.5011626300856216 | 0.03248890626116566 |
| **GATCAAACAGATCTAACTTGGGGTGGCTTTGTCTTCTTCTTTTGCCAATTCCACTAATTGTTTGGG** | Ins_k3_4_AAA | 0.33157164303184383 | 0.3905130607871113 | 68.40395727361025 |
| **GATCAACAGATCTAACTTGGGGTGGCTTTGTCTTCTTCTTTTGCCAATTCCACTAATTGTTTGGG** | Ins_k2_4_AA | -0.11696611759608011 | 0.04000794879003268 | 43.68011294648114 |
| **GATCAACCAGATCTAACTTGGGGTGGCTTTGTCTTCTTCTTTTGCCAATTCCACTAATTGTTTGGG** | Ins_k3_3_CAA;Ins_k3_4_AAC | 0.2699937440014708 | 0.03479876629221172 | 64.31885215109935 |
| **GATCAACGCAGATCTAACTTGGGGTGGCTTTGTCTTCTTCTTTTGCCAATTCCACTAATTGTTTGGG** | Ins_k4_4_AACG | 1.044648761904658 | 0.029832792222595432 | 100 |
| **GATCAACTTGGGGTGGCTTTGTCTTCTTCTTTTGCCAATTCCACTAATTGTTTGGG** | Del_k7_5-11 | -0.05785087663510813 | 0.03463294971757522 | 46.34012229888845 |
| **GATCAAGATCTAACTTGGGGTGGCTTTGTCTTCTTCTTTTGCCAATTCCACTAATTGTTTGGG** | C5A | -0.17122492812743473 | 0.03138615415113304 | 41.3732322071408 |
| **GATCAAGCAGATCTAACTTGGGGTGGCTTTGTCTTCTTCTTTTGCCAATTCCACTAATTGTTTGGG** | Ins_k3_4_AAG | -0.1420340776140906 | 0.03644780074972771 | 42.598752004891914 |
| **GATCAATCAGATCTAACTTGGGGTGGCTTTGTCTTCTTCTTTTGCCAATTCCACTAATTGTTTGGG** | Ins_k3_4_AAT | -1.943295162310629 | 0.09127251419443719 | 7.032651981759048 |
| **GATCAATTCCACTAATTGTTTGGG** | Del_k39_4-42;Del_k39_5-43 | -5.93274188735467 | 0.33930691900101856 | 0.13017405329000117 |
| **GATCAATTGTTTGGG** | Del_k48_5-52 | -5.861843364055686 | 0.33050462742232706 | 0.13973823881234876 |
| **GATCACACAGATCTAACTTGGGGTGGCTTTGTCTTCTTCTTTTGCCAATTCCACTAATTGTTTGGG** | Ins_k3_4_ACA | -0.7606545937426143 | 0.05477625777458948 | 22.947395427258854 |
| **GATCACAGATCTAACTTGGGGTGGCTTTGTCTTCTTCTTTTGCCAATTCCACTAATTGTTTGGG** | Ins_k1_4_A | -0.3564166436534213 | 0.046259773137361807 | 34.378878927471526 |
| **GATCACCAGATCTAACTTGGGGTGGCTTTGTCTTCTTCTTTTGCCAATTCCACTAATTGTTTGGG** | Ins_k2_3_CA;Ins_k2_4_AC | 0.02215165716980838 | 0.04020924061161922 | 50.199782397156135 |
| **GATCACCCAGATCTAACTTGGGGTGGCTTTGTCTTCTTCTTTTGCCAATTCCACTAATTGTTTGGG** | Ins_k3_3_CAC;Ins_k3_4_ACC | -0.33073316562883687 | 0.04438600075377801 | 35.273284668998 |
| **GATCACGCAGATCTAACTTGGGGTGGCTTTGTCTTCTTCTTTTGCCAATTCCACTAATTGTTTGGG** | Ins_k3_4_ACG | 0.15734058788861194 | 0.036404518655436015 | 57.46635365391877 |
| **GATCACTAATTGTTTGGG** | Del_k45_4-48;Del_k45_5-49 | -5.880480323142302 | 0.3387779865132613 | 0.1371580609890391 |
| **GATCACTCAGATCTAACTTGGGGTGGCTTTGTCTTCTTCTTTTGCCAATTCCACTAATTGTTTGGG** | Ins_k3_4_ACT | -1.5166339544496386 | 0.0749034577146949 | 10.774961688058156 |
| **GATCACTTGGGGTGGCTTTGTCTTCTTCTTTTGCCAATTCCACTAATTGTTTGGG** | Del_k8_5-12 | -0.9143912770796698 | 0.0515431977054347 | 19.67734074446485 |
| **GATCAGACAGATCTAACTTGGGGTGGCTTTGTCTTCTTCTTTTGCCAATTCCACTAATTGTTTGGG** | Ins_k3_4_AGA | -1.5034111576611369 | 0.07756563721731663 | 10.918382942243129 |
| **GATCAGACCAGATCTAACTTGGGGTGGCTTTGTCTTCTTCTTTTGCCAATTCCACTAATTGTTTGGG** | Ins_k4_3_CAGA;Ins_k4_4_AGAC | 0.33061596858590436 | 0.038139984649230324 | 68.3386165868281 |
| **GATCAGATCTAACTTGGGGTGGCTTTGTCTTCTTCTTTTGCCAATTCCACTAATTGTTTGGG** | Del_k1_4-4;Del_k1_5-5 | -2.2166875764537703 | 0.07639028001150426 | 5.350400716303649 |
| **GATCAGCAGATCTAACTTGGGGTGGCTTTGTCTTCTTCTTTTGCCAATTCCACTAATTGTTTGGG** | Ins_k2_4_AG | -0.20290629432089913 | 0.04171997115964564 | 40.08301748439717 |
| **GATCAGCCAGATCTAACTTGGGGTGGCTTTGTCTTCTTCTTTTGCCAATTCCACTAATTGTTTGGG** | Ins_k3_3_CAG;Ins_k3_4_AGC | -0.06167160299734853 | 0.036964516732090544 | 46.16340717702715 |
| **GATCAGGCAGATCTAACTTGGGGTGGCTTTGTCTTCTTCTTTTGCCAATTCCACTAATTGTTTGGG** | Ins_k3_4_AGG | -1.5945612943313707 | 0.08042041349468577 | 9.967180449023347 |
| **GATCAGTCAGATCTAACTTGGGGTGGCTTTGTCTTCTTCTTTTGCCAATTCCACTAATTGTTTGGG** | Ins_k3_4_AGT | -3.0589368586772205 | 0.14760043199361506 | 2.3046346857020334 |
| **GATCATACAGATCTAACTTGGGGTGGCTTTGTCTTCTTCTTTTGCCAATTCCACTAATTGTTTGGG** | Ins_k3_4_ATA | -1.4740568154024594 | 0.07824142081905644 | 11.243635321995292 |
| **GATCATAGCAGATCTAACTTGGGGTGGCTTTGTCTTCTTCTTTTGCCAATTCCACTAATTGTTTGGG** | Ins_k4_4_ATAG | -1.9501993057787879 | 0.0947123729012538 | 6.984264771841734 |
| **GATCATCAGATCTAACTTGGGGTGGCTTTGTCTTCTTCTTTTGCCAATTCCACTAATTGTTTGGG** | Ins_k2_4_AT | -0.07494354842603757 | 0.04348056683458944 | 45.55477674499555 |
| **GATCATCCAGATCTAACTTGGGGTGGCTTTGTCTTCTTCTTTTGCCAATTCCACTAATTGTTTGGG** | Ins_k3_1_ATC;Ins_k3_2_TCA;Ins_ | 0.1513911853468266 | 0.03874167510489104 | 57.12547819154524 |
| **GATCATCTAACTTGGGGTGGCTTTGTCTTCTTCTTTTGCCAATTCCACTAATTGTTTGGG** | Del_k3_5-7 | -0.3918083012063748 | 0.04192399652195153 | 33.18343260978586 |
| **GATCATGCAGATCTAACTTGGGGTGGCTTTGTCTTCTTCTTTTGCCAATTCCACTAATTGTTTGGG** | Ins_k3_4_ATG | -0.4375088309100238 | 0.04279722579709338 | 31.701062697141058 |
| **GATCATTCAGATCTAACTTGGGGTGGCTTTGTCTTCTTCTTTTGCCAATTCCACTAATTGTTTGGG** | Ins_k3_4_ATT | -2.1716137809459966 | 0.1135797645708193 | 5.5970812351411166 |
| **GATCATTCCACTAATTGTTTGGG** | Del_k40_5-44 | -6.4118313955085835 | 0.5410600861934826 | 0.08062291531907291 |
| **GATCATTGCAGATCTAACTTGGGGTGGCTTTGTCTTCTTCTTTTGCCAATTCCACTAATTGTTTGGG** | Ins_k4_4_ATTG | -0.5017417751534199 | 0.07415943187437823 | 29.728829334048477 |
| **GATCATTGTTTGGG** | Del_k49_5-53 | -5.830235129219527 | 0.3889952637926539 | 0.1442256640740224 |
| **GATCC** | Del_k58_6-63 | -6.9325814947934346 | 0.42641998525342223 | 0.04789603953175231 |
| **GATCCA** | Del_k57_7-63 | -7.033527393473887 | 0.5676921539196547 | 0.043297154646993956 |
| **GATCCAAAAGATCTAACTTGGGGTGGCTTTGTCTTCTTCTTTTGCCAATTCCACTAATTGTTTGGG** | Ins_k3_5_AAA;Ins_k3_6_AAA | -0.7934282696493626 | 0.06207437501211275 | 22.20751544508607 |
| **GATCCAAACGATCTAACTTGGGGTGGCTTTGTCTTCTTCTTTTGCCAATTCCACTAATTGTTTGGG** | Ins_k3_6_AAC | -0.11612881160228927 | 0.03830899879434499 | 43.71670188279087 |
| **GATCCAAACTTGGGGTGGCTTTGTCTTCTTCTTTTGCCAATTCCACTAATTGTTTGGG** | Del_k5_7-11 | 0.03276363496931456 | 0.03474155173164098 | 50.735337999661525 |
| **GATCCAAAGATCTAACTTGGGGTGGCTTTGTCTTCTTCTTTTGCCAATTCCACTAATTGTTTGGG** | Ins_k2_5_AA;Ins_k2_6_AA | -0.6018387420289105 | 0.04908687135962151 | 26.897148916906726 |
| **GATCCAAAGGATCTAACTTGGGGTGGCTTTGTCTTCTTCTTTTGCCAATTCCACTAATTGTTTGGG** | Ins_k3_6_AAG | 0.23164583200160327 | 0.03522822876387283 | 61.89905213946083 |
| **GATCCAAATCTAACTTGGGGTGGCTTTGTCTTCTTCTTTTGCCAATTCCACTAATTGTTTGGG** | G7A | -1.057118987146654 | 0.04433133946692309 | 17.060059805475138 |
| **GATCCAAATGATCTAACTTGGGGTGGCTTTGTCTTCTTCTTTTGCCAATTCCACTAATTGTTTGGG** | Ins_k3_6_AAT | -0.7211135012608543 | 0.05640973353452054 | 23.872938415264354 |
| **GATCCAAATTCCACTAATTGTTTGGG** | Del_k37_7-43 | -5.81401719199489 | 0.29491626258839243 | 0.14658377701803238 |
| **GATCCAAATTGTTTGGG** | Del_k46_7-52 | -6.491462815872274 | 0.6094063160215726 | 0.07445176737317238 |
| **GATCCAACAGATCTAACTTGGGGTGGCTTTGTCTTCTTCTTTTGCCAATTCCACTAATTGTTTGGG** | Ins_k3_4_CAA;Ins_k3_5_AAC;Ins | 0.2391791609921261 | 0.038569877634696104 | 62.36711890027386 |
| **GATCCAACCGATCTAACTTGGGGTGGCTTTGTCTTCTTCTTTTGCCAATTCCACTAATTGTTTGGG** | Ins_k3_6_ACC | 0.23796180494386798 | 0.03508127719907357 | 62.29124210477664 |
| **GATCCAACGATCTAACTTGGGGTGGCTTTGTCTTCTTCTTTTGCCAATTCCACTAATTGTTTGGG** | Ins_k2_6_AC | 0.15266159306329086 | 0.04160296461635101 | 57.19809695780016 |
| **GATCCAACGGATCTAACTTGGGGTGGCTTTGTCTTCTTCTTTTGCCAATTCCACTAATTGTTTGGG** | Ins_k3_6_ACG | 0.48056914234506176 | 0.03174697679266523 | 79.39442702164932 |
| **GATCCAACTAATTGTTTGGG** | Del_k43_7-49 | -5.984853453940676 | 0.34782124505286466 | 0.12356420009714851 |
| **GATCCAACTGATCTAACTTGGGGTGGCTTTGTCTTCTTCTTTTGCCAATTCCACTAATTGTTTGGG** | Ins_k3_6_ACT | 0.11551748960498343 | 0.03359634157551968 | 55.11249855172081 |
| **GATCCAACTTGGGGTGGCTTTGTCTTCTTCTTTTGCCAATTCCACTAATTGTTTGGG** | Del_k6_6-11;Del_k6_7-12 | 0.028973922132113805 | 0.03665267084446091 | 50.54342950666542 |
| **GATCCAAGAGATCTAACTTGGGGTGGCTTTGTCTTCTTCTTTTGCCAATTCCACTAATTGTTTGGG** | Ins_k3_5_AAG;Ins_k3_6_AGA | -0.09163818600027052 | 0.041599804794011784 | 44.80056938380932 |
| **GATCCAAGATCTAACTTGGGGTGGCTTTGTCTTCTTCTTTTGCCAATTCCACTAATTGTTTGGG** | Ins_k1_5_A;Ins_k1_6_A | 0.38395180550649155 | 0.03574623265714463 | 72.08246712487711 |
| **GATCCAAGCGATCTAACTTGGGGTGGCTTTGTCTTCTTCTTTTGCCAATTCCACTAATTGTTTGGG** | Ins_k3_6_AGC | -0.20737798603659774 | 0.04335784155362193 | 39.9041787410408 |
| **GATCCAAGGAGATCTAACTTGGGGTGGCTTTGTCTTCTTCTTTTGCCAATTCCACTAATTGTTTGGG** | Ins_k4_5_AAGG;Ins_k4_6_AGGA | 0.6269609978850053 | 0.02985642906599123 | 91.9109370562209 |
| **GATCCAAGGATCTAACTTGGGGTGGCTTTGTCTTCTTCTTTTGCCAATTCCACTAATTGTTTGGG** | Ins_k2_6_AG | 0.39238313154026894 | 0.03784939046204201 | 72.69278719477903 |
| **GATCCAAGGGATCTAACTTGGGGTGGCTTTGTCTTCTTCTTTTGCCAATTCCACTAATTGTTTGGG** | Ins_k3_6_AGG | -0.4341249712636017 | 0.046385425807971364 | 31.80851634544959 |
| **GATCCAAGTGATCTAACTTGGGGTGGCTTTGTCTTCTTCTTTTGCCAATTCCACTAATTGTTTGGG** | Ins_k3_6_AGT | -0.9857099011963371 | 0.05982239527525284 | 18.322854008526274 |
| **GATCCAATAGATCTAACTTGGGGTGGCTTTGTCTTCTTCTTTTGCCAATTCCACTAATTGTTTGGG** | Ins_k3_5_AAT;Ins_k3_6_ATA | -2.6487391275062113 | 0.10293954841147127 | 3.4733512545192764 |
| **GATCCAATCGATCTAACTTGGGGTGGCTTTGTCTTCTTCTTTTGCCAATTCCACTAATTGTTTGGG** | Ins_k3_6_ATC | 0.18615409595417343 | 0.48416432547295807 | 59.14624647963428 |
| **GATCCAATCTAACTTGGGGTGGCTTTGTCTTCTTCTTTTGCCAATTCCACTAATTGTTTGGG** | Del_k1_7-7 | -1.3533518354018659 | 0.057278483484095914 | 12.68610372353109 |
| **GATCCAATGAGATCTAACTTGGGGTGGCTTTGTCTTCTTCTTTTGCCAATTCCACTAATTGTTTGGG** | Ins_k4_5_AATG;Ins_k4_6_ATGA | -0.464909333507604 | 0.05781083237183981 | 30.84423007619071 |

| | | | | |
|---|---|---|---|---|
| GATCCAATGATCTAACTTGGGGTGGCTTTGTCTTCTTCTTTTGCCAATTCCACTAATTGTTTGGG | Ins_k2_6_AT | -0.8938933383297063 | 0.07017779327722269 | 20.08484793020221 |
| GATCCAATGGATCTAACTTGGGGTGGCTTTGTCTTCTTCTTTTGCCAATTCCACTAATTGTTTGGG | Ins_k3_6_ATG | -0.2408955160080078 | 0.03967094530812021 | 38.58885556271528 |
| GATCCAATTCCACTAATTGTTTGGG | Del_k38_4-41;Del_k38_5-42;Del_ | -5.825679347547483 | 0.2838818898139689 | 0.1448842236989423 |
| GATCCAATTGATCTAACTTGGGGTGGCTTTGTCTTCTTCTTTTGCCAATTCCACTAATTGTTTGGG | Ins_k3_6_ATT | -0.7212337536030256 | 0.05656060972333116 | 23.870067811107305 |
| GATCCAATTGTTTGGG | Del_k47_6-52;Del_k47_7-53 | -6.11684384374125 | 0.5882368409275315 | 0.10828541470875189 |
| GATCCACAAGATCTAACTTGGGGTGGCTTTGTCTTCTTCTTTTGCCAATTCCACTAATTGTTTGGG | Ins_k3_5_ACA;Ins_k3_6_CAA | 0.22273486595926706 | 0.03514969055943724 | 61.34992206118261 |
| GATCCACAATTCCACTAATTGTTTGGG | Del_k36_7-42 | -4.583861607582973 | 0.1303094672107807 | 0.5015747786215836 |
| GATCCACACGATCTAACTTGGGGTGGCTTTGTCTTCTTCTTTTGCCAATTCCACTAATTGTTTGGG | Ins_k3_6_CAC | -0.010592833097778476 | 0.03607402690731692 | 48.58263690305323 |
| GATCCACACTAATTGTTTGGG | Del_k42_7-48 | -6.571749963085389 | 0.4642527070102976 | 0.06870791136140847 |
| GATCCACAGATCTAACTTGGGGTGGCTTTGTCTTCTTCTTTTGCCAATTCCACTAATTGTTTGGG | Ins_k2_4_CA;Ins_k2_5_AC;Ins_k | -0.19439182425264218 | 0.04594220732778164 | 40.42576020269529 |
| GATCCACAGCGATCTAACTTGGGGTGGCTTTGTCTTCTTCTTTTGCCAATTCCACTAATTGTTTGGG | Ins_k4_6_CAGC | 0.5124214245598109 | 0.03790666580594192 | 81.96402728464803 |
| GATCCACAGGATCTAACTTGGGGTGGCTTTGTCTTCTTCTTTTGCCAATTCCACTAATTGTTTGGG | Ins_k3_6_CAG | 0.19630818106837433 | 0.03461966858030144 | 59.749881997285975 |
| GATCCACATCTAACTTGGGGTGGCTTTGTCTTCTTCTTTTGCCAATTCCACTAATTGTTTGGG | G7C | -0.5819639370010341 | 0.04014129753226339 | 27.437072169411742 |
| GATCCACATGATCTAACTTGGGGTGGCTTTGTCTTCTTCTTTTGCCAATTCCACTAATTGTTTGGG | Ins_k3_6_CAT | -0.17792299850921567 | 0.04211634854966824 | 41.09703740461173 |
| GATCCACCAATTCCACTAATTGTTTGGG | Del_k35_7-41 | -4.004785767927794 | 0.09734045588340555 | 0.8950043206944955 |
| GATCCACCACTAATTGTTTGGG | Del_k41_7-47 | -5.233265288347862 | 0.19218125445194778 | 0.2620011846811754 |
| GATCCACCAGATCTAACTTGGGGTGGCTTTGTCTTCTTCTTTTGCCAATTCCACTAATTGTTTGGG | Ins_k3_3_CCA;Ins_k3_4_CAC;Ins | 0.2531686221134717 | 0.0342564337683328 | 63.245732622587624 |
| GATCCACCCGATCTAACTTGGGGTGGCTTTGTCTTCTTCTTTTGCCAATTCCACTAATTGTTTGGG | Ins_k3_6_CCC | -2.672865949307024 | 0.12633128919462436 | 3.3905531714303345 |
| GATCCACCGATCTAACTTGGGGTGGCTTTGTCTTCTTCTTTTGCCAATTCCACTAATTGTTTGGG | Ins_k2_6_CC | 0.12877278174772444 | 0.03838644326639669 | 55.847893993900385 |
| GATCCACCGGATCTAACTTGGGGTGGCTTTGTCTTCTTCTTTTGCCAATTCCACTAATTGTTTGGG | Ins_k3_6_CCG | 0.6482480989596642 | 0.02962589774394466 | 93.88842730775806 |
| GATCCACCTGATCTAACTTGGGGTGGCTTTGTCTTCTTCTTTTGCCAATTCCACTAATTGTTTGGG | Ins_k3_6_CCT | -0.20526608137708396 | 0.04339997380686759 | 39.98854161387433 |
| GATCCACGAGATCTAACTTGGGGTGGCTTTGTCTTCTTCTTTTGCCAATTCCACTAATTGTTTGGG | Ins_k3_5_ACG;Ins_k3_6_CGA | 0.45803131826550425 | 0.03400768111849054 | 77.62506309458325 |
| GATCCACGATCTAACTTGGGGTGGCTTTGTCTTCTTCTTTTGCCAATTCCACTAATTGTTTGGG | Ins_k1_6_C | 0.5222102300512266 | 0.03520195773011066 | 82.77029697551822 |
| GATCCACGCCGATCTAACTTGGGGTGGCTTTGTCTTCTTCTTTTGCCAATTCCACTAATTGTTTGGG | Ins_k4_6_CGCC | 0.8765203937200274 | 0.03324810507199159 | 100 |
| GATCCACGCGATCTAACTTGGGGTGGCTTTGTCTTCTTCTTTTGCCAATTCCACTAATTGTTTGGG | Ins_k3_6_CGC | 0.4335177916083548 | 0.033704902457806205 | 75.74533258176578 |
| GATCCACGGATCTAACTTGGGGTGGCTTTGTCTTCTTCTTTTGCCAATTCCACTAATTGTTTGGG | Ins_k2_6_CG | 0.5871439024903458 | 0.03523794813103431 | 88.32321091094857 |
| GATCCACGGGATCTAACTTGGGGTGGCTTTGTCTTCTTCTTTTGCCAATTCCACTAATTGTTTGGG | Ins_k3_6_CGG | 0.2835386392712713 | 0.036111254724143775 | 65.1959710995558 |
| GATCCACGTGATCTAACTTGGGGTGGCTTTGTCTTCTTCTTTTGCCAATTCCACTAATTGTTTGGG | Ins_k3_6_CGT | 0.23282099359143837 | 0.03405181854717363 | 61.97183628617283 |
| GATCCACTAACTTGGGGTGGCTTTGTCTTCTTCTTTTGCCAATTCCACTAATTGTTTGGG | Del_k3_7-9 | -0.8410442692037569 | 0.049354925953317905 | 21.174862893559734 |
| GATCCACTAATTGTTTGGG | Del_k44_3-46;Del_k44_4-47;Del_ | -6.458051787342631 | 0.4464436340230626 | 0.07698129932439096 |
| GATCCACTAGATCTAACTTGGGGTGGCTTTGTCTTCTTCTTTTGCCAATTCCACTAATTGTTTGGG | Ins_k3_5_ACT;Ins_k3_6_CTA | -1.375689877146486 | 0.0772213263451364 | 12.405862680024592 |
| GATCCACTCCGATCTAACTTGGGGTGGCTTTGTCTTCTTCTTTTGCCAATTCCACTAATTGTTTGGG | Ins_k4_6_CTCC | -2.267986959001489 | 0.15883473156665234 | 5.082849735190968 |
| GATCCACTCGATCTAACTTGGGGTGGCTTTGTCTTCTTCTTTTGCCAATTCCACTAATTGTTTGGG | Ins_k3_6_CTC | -0.28199723679830546 | 0.03938388253045092 | 37.034940245442925 |
| GATCCACTGATCTAACTTGGGGTGGCTTTGTCTTCTTCTTTTGCCAATTCCACTAATTGTTTGGG | Ins_k2_6_CT | -0.1612128944719084 | 0.048111178499662385 | 41.78954298157376 |
| GATCCACTGGATCTAACTTGGGGTGGCTTTGTCTTCTTCTTTTGCCAATTCCACTAATTGTTTGGG | Ins_k3_6_CTG | 0.14441041719157455 | 0.0386258845700784 | 56.72808713372772 |
| GATCCACTTCTTCTTTTGCCAATTCCACTAATTGTTTGGG | Del_k23_7-29 | -1.4457946906950765 | 0.05464483600872766 | 11.565937362985766 |
| GATCCACTTCTTTTGCCAATTCCACTAATTGTTTGGG | Del_k26_7-32 | -2.3923883083512214 | 0.062483345290840854 | 4.48828525839628 |
| GATCCACTTGATCTAACTTGGGGTGGCTTTGTCTTCTTCTTTTGCCAATTCCACTAATTGTTTGGG | Ins_k3_6_CTT | -0.24639016551231213 | 0.04307372394400427 | 38.377404782583376 |
| GATCCACTTGGGGTGGCTTTGTCTTCTTCTTTTGCCAATTCCACTAATTGTTTGGG | Del_k7_6-12;Del_k7_7-13 | 0.05541135352740301 | 0.037697124937595826 | 51.8974880050913 |
| GATCCACTTTGTCTTCTTCTTTTGCCAATTCCACTAATTGTTTGGG | Del_k17_7-23 | -0.9073665760248066 | 0.03801965437120362 | 19.816054822795728 |
| GATCCACTTTTGCCAATTCCACTAATTGTTTGGG | Del_k29_7-35 | -3.224764063552896 | 0.07747224972787957 | 1.9524694797657853 |
| GATCCAG | Del_k56_7-62;Del_k56_8-63 | -5.01995895092229 | 0.10953288140068358 | 0.3242955731883222 |
| GATCCAGA | Del_k55_9-63 | -6.0473331046175645 | 0.25200810706447135 | 0.11608018579653318 |
| GATCCAGAAAATCTAACTTGGGGTGGCTTTGTCTTCTTCTTTTGCCAATTCCACTAATTGTTTGGG | Ins_k3_7_AAA;Ins_k3_8_AAA | 0.31346235668285927 | 0.035976964849159285 | 67.17635943300638 |
| GATCCAGAAACTCTAACTTGGGGTGGCTTTGTCTTCTTCTTTTGCCAATTCCACTAATTGTTTGGG | Ins_k3_8_AAC | 0.07257552214208507 | 0.039412630990387644 | 52.79595389199841 |
| GATCCAGAAACTTGGGGTGGCTTTGTCTTCTTCTTTTGCCAATTCCACTAATTGTTTGGG | Del_k3_9-11 | 0.43581729939027425 | 0.031853713590668166 | 75.91970997773267 |
| GATCCAGAAAGTCTAACTTGGGGTGGCTTTGTCTTCTTCTTTTGCCAATTCCACTAATTGTTTGGG | Ins_k3_8_AAG | 0.11390064529668953 | 0.039309223478320436 | 55.02346222046077 |
| GATCCAGAAATCTAACTTGGGGTGGCTTTGTCTTCTTCTTTTGCCAATTCCACTAATTGTTTGGG | Ins_k2_7_AA;Ins_k2_8_AA | 0.2526282545611337 | 0.040547592953775205 | 63.21156591297654 |
| GATCCAGAAATTCCACTAATTGTTTGGG | Del_k35_9-43 | -3.308275313099529 | 0.06799753038223708 | 1.796039066695289 |
| GATCCAGAAATTCTAACTTGGGGTGGCTTTGTCTTCTTCTTTTGCCAATTCCACTAATTGTTTGGG | Ins_k3_8_AAT | 0.017170557859275015 | 0.0422750832359313 | 49.950354025135695 |
| GATCCAGAAATTGTTTGGG | Del_k44_9-52 | -5.7321026374634 | 0.3449402131627217 | 0.15909661772514952 |
| GATCCAGAACATCTAACTTGGGGTGGCTTTGTCTTCTTCTTTTGCCAATTCCACTAATTGTTTGGG | Ins_k3_7_AAC;Ins_k3_8_ACA | 0.6052595502672695 | 0.03765052014495382 | 89.93782381337088 |
| GATCCAGAACCATCTAACTTGGGGTGGCTTTGTCTTCTTCTTTTGCCAATTCCACTAATTGTTTGGG | Ins_k4_7_AACC;Ins_k4_8_ACCA | 0.7542918492833464 | 0.032490147054246715 | 100 |
| GATCCAGAACCTCTAACTTGGGGTGGCTTTGTCTTCTTCTTTTGCCAATTCCACTAATTGTTTGGG | Ins_k3_8_ACC | 0.4443326511434569 | 0.03921910727686584 | 76.56895335795654 |
| GATCCAGAACGTCTAACTTGGGGTGGCTTTGTCTTCTTCTTTTGCCAATTCCACTAATTGTTTGGG | Ins_k3_8_ACG | 0.6907431786242029 | 0.029385606155236263 | 97.96421054262277 |
| GATCCAGAACTAACTTGGGGTGGCTTTGTCTTCTTCTTTTGCCAATTCCACTAATTGTTTGGG | T9A | 0.013183389758906983 | 0.039770563468725074 | 49.75159008291046 |
| GATCCAGAACTAATTGTTTGGG | Del_k41_9-49 | -5.522868202407782 | 0.26976761332486693 | 0.19612380370422886 |
| GATCCAGAACTCTAACTTGGGGTGGCTTTGTCTTCTTCTTTTGCCAATTCCACTAATTGTTTGGG | Ins_k2_8_AC | -0.0521306835873544 | 0.045072400193230236 | 46.60595633043028 |
| GATCCAGAACTTCTAACTTGGGGTGGCTTTGTCTTCTTCTTTTGCCAATTCCACTAATTGTTTGGG | Ins_k3_8_ACT | 0.28479632799056287 | 0.03455192657652171 | 65.27801892144586 |
| GATCCAGAACTTGGGGTGGCTTTGTCTTCTTCTTTTGCCAATTCCACTAATTGTTTGGG | Del_k4_8-11;Del_k4_9-12 | 0.4236517370816 | 0.03260618973945646 | 75.001699395185 |
| GATCCAGAAGAATCTAACTTGGGGTGGCTTTGTCTTCTTCTTTTGCCAATTCCACTAATTGTTTGGG | Ins_k4_7_AAGA;Ins_k4_8_AGAA | 0.9997381845144964 | 0.03079412891649509 | 100 |
| GATCCAGAAGATCTAACTTGGGGTGGCTTTGTCTTCTTCTTTTGCCAATTCCACTAATTGTTTGGG | Ins_k3_5_AGA;Ins_k3_6_GAA;Ins | 0.7746904806960544 | 0.031694926036689705 | 100 |
| GATCCAGAAGCTCTAACTTGGGGTGGCTTTGTCTTCTTCTTTTGCCAATTCCACTAATTGTTTGGG | Ins_k3_8_AGC | 0.5792921508589124 | 0.033366899253454645 | 87.63243444705941 |
| GATCCAGAAGGTCTAACTTGGGGTGGCTTTGTCTTCTTCTTTTGCCAATTCCACTAATTGTTTGGG | Ins_k3_8_AGG | 0.27599241151740117 | 0.03461442140250897 | 64.70583910274347 |
| GATCCAGAAGTCTAACTTGGGGTGGCTTTGTCTTCTTCTTTTGCCAATTCCACTAATTGTTTGGG | Ins_k2_8_AG | 0.5025097905979434 | 0.036185636440931916 | 81.15564267234822 |
| GATCCAGAAGTTCTAACTTGGGGTGGCTTTGTCTTCTTCTTTTGCCAATTCCACTAATTGTTTGGG | Ins_k3_8_AGT | 0.7148933004297504 | 0.031197534663659494 | 100 |
| GATCCAGAATATCTAACTTGGGGTGGCTTTGTCTTCTTCTTTTGCCAATTCCACTAATTGTTTGGG | Ins_k3_7_AAT;Ins_k3_8_ATA | 0.06777432985443022 | 0.037180646197972886 | 52.543077904236796 |
| GATCCAGAATCTAACTTGGGGTGGCTTTGTCTTCTTCTTTTGCCAATTCCACTAATTGTTTGGG | Ins_k1_7_A;Ins_k1_8_A | 0.568233899042025 | 0.03540976109457892 | 86.66871128702482 |
| GATCCAGAATCTCTAACTTGGGGTGGCTTTGTCTTCTTCTTTTGCCAATTCCACTAATTGTTTGGG | Ins_k3_8_ATC | 0.12261937040005222 | 0.039525759780756106 | 55.505294089354564 |
| GATCCAGAATGTCTAACTTGGGGTGGCTTTGTCTTCTTCTTTTGCCAATTCCACTAATTGTTTGGG | Ins_k3_8_ATG | 0.0034430648208342296 | 0.03821932529927861 | 49.26934584977667 |
| GATCCAGAATGTTCTAACTTGGGGTGGCTTTGTCTTCTTCTTTTGCCAATTCCACTAATTGTTTGGG | Ins_k4_8_ATGT | 0.24374991257298828 | 0.04256083876709804 | 62.65283598105264 |
| GATCCAGAATTCCACTAATTGTTTGGG | Del_k36_8-43;Del_k36_9-44 | -3.703097072382989 | 0.07934015259675016 | 1.2101713515303345 |
| GATCCAGAATTCTAACTTGGGGTGGCTTTGTCTTCTTCTTTTGCCAATTCCACTAATTGTTTGGG | Ins_k2_8_AT | 0.06598419372254138 | 0.04360683635290756 | 52.44910278122732 |
| GATCCAGAATTGTTTGGG | Del_k45_8-52;Del_k45_9-53 | -5.568060537256938 | 0.3707612838439226 | 0.18745780433274567 |
| GATCCAGAATTTCTAACTTGGGGTGGCTTTGTCTTCTTCTTTTGCCAATTCCACTAATTGTTTGGG | Ins_k3_8_ATT | 0.14928393358952186 | 0.043730456750329376 | 57.00522717132406 |

| | | | | |
|---|---|---|---|---|
| **GATCCAGACAATCTAACTTGGGGTGGCTTTGTCTTCTTCTTTTGCCAATTCCACTAATTGTTTGGG** | Ins_k3_7_ACA;Ins_k3_8_CAA | -0.39449346875381286 | 0.18314897473970942 | 33.09444905477176 |
| **GATCCAGACAATTCCACTAATTGTTTGGG** | Del_k34_9-42 | -3.9518560394406443 | 0.12460339249842736 | 0.9436527740121785 |
| **GATCCAGACACTAATTGTTTGGG** | Del_k40_9-48 | -5.403043181705276 | 0.2032737109008087 | 0.22109028158929286 |
| **GATCCAGACACTCTAACTTGGGGTGGCTTTGTCTTCTTCTTTTGCCAATTCCACTAATTGTTTGGG** | Ins_k3_8_CAC | -0.3221337445608875 | 0.04155917668088781 | 35.5779224733636 |
| **GATCCAGACAGTCTAACTTGGGGTGGCTTTGTCTTCTTCTTTTGCCAATTCCACTAATTGTTTGGG** | Ins_k3_8_CAG | -0.08035401351689475 | 0.03574342284552025 | 45.30896978032371 |
| **GATCCAGACATATCTAACTTGGGGTGGCTTTGTCTTCTTCTTTTGCCAATTCCACTAATTGTTTGGG** | Ins_k4_7_ACAT;Ins_k4_8_CATA | -0.16149641278226962 | 0.03876806942874626 | 41.77769656037487 |
| **GATCCAGACATCTAACTTGGGGTGGCTTTGTCTTCTTCTTTTGCCAATTCCACTAATTGTTTGGG** | Ins_k2_7_AC;Ins_k2_8_CA | 0.17381781651973693 | 0.03927187170839986 | 58.42108395580326 |
| **GATCCAGACATTCTAACTTGGGGTGGCTTTGTCTTCTTCTTTTGCCAATTCCACTAATTGTTTGGG** | Ins_k3_8_CAT | -0.10673135840056669 | 0.042309854403834155 | 44.12946396137291 |
| **GATCCAGACCAATTCCACTAATTGTTTGGG** | Del_k33_9-41 | -3.1956266224637115 | 0.06923032850125009 | 2.010196367025633 |
| **GATCCAGACCACTAATTGTTTGGG** | Del_k39_9-47 | -5.094566836532408 | 0.16876156441518694 | 0.30098110155619895 |
| **GATCCAGACCATCTAACTTGGGGTGGCTTTGTCTTCTTCTTTTGCCAATTCCACTAATTGTTTGGG** | Ins_k3_7_ACC;Ins_k3_8_CCA | 0.39974984897237476 | 0.032302541507921376 | 73.23027173505574 |
| **GATCCAGACCCTCTAACTTGGGGTGGCTTTGTCTTCTTCTTTTGCCAATTCCACTAATTGTTTGGG** | Ins_k3_8_CCC | 0.26756076565777864 | 0.03461048493207763 | 64.16255598641459 |
| **GATCCAGACCGTCTAACTTGGGGTGGCTTTGTCTTCTTCTTTTGCCAATTCCACTAATTGTTTGGG** | Ins_k3_8_CCG | 0.7389896895205295 | 0.029755184689043198 | 100 |
| **GATCCAGACCTAACTTGGGGTGGCTTTGTCTTCTTCTTTTGCCAATTCCACTAATTGTTTGGG** | T9C | -0.07271675932851873 | 0.027136041092153954 | 45.65633065279488 |
| **GATCCAGACCTCTAACTTGGGGTGGCTTTGTCTTCTTCTTTTGCCAATTCCACTAATTGTTTGGG** | Ins_k2_8_CC | 0.26957562570595706 | 0.048609348567000193 | 64.29196488368945 |
| **GATCCAGACCTTCTAACTTGGGGTGGCTTTGTCTTCTTCTTTTGCCAATTCCACTAATTGTTTGGG** | Ins_k3_8_CCT | 0.40715336948537195 | 0.031930993886037005 | 73.77444546911272 |
| **GATCCAGACGATCTAACTTGGGGTGGCTTTGTCTTCTTCTTTTGCCAATTCCACTAATTGTTTGGG** | Ins_k3_6_GAC;Ins_k3_7_ACG;Ins | 0.6565812578211969 | 0.029697759098417254 | 94.67408344002797 |
| **GATCCAGACGCTCTAACTTGGGGTGGCTTTGTCTTCTTCTTTTGCCAATTCCACTAATTGTTTGGG** | Ins_k3_8_CGC | 0.6284328151381686 | 0.031282171271110985 | 92.04631275884395 |
| **GATCCAGACGGTCTAACTTGGGGTGGCTTTGTCTTCTTCTTTTGCCAATTCCACTAATTGTTTGGG** | Ins_k3_8_CGG | 0.5845821601548902 | 0.03174361505534465 | 88.09723916662539 |
| **GATCCAGACGTCTAACTTGGGGTGGCTTTGTCTTCTTCTTTTGCCAATTCCACTAATTGTTTGGG** | Ins_k2_8_CG | 0.7576259192665518 | 0.03486592537988289 | 100 |
| **GATCCAGACGTTCTAACTTGGGGTGGCTTTGTCTTCTTCTTTTGCCAATTCCACTAATTGTTTGGG** | Ins_k3_8_CGT | 0.9166115336901357 | 0.033563589872203085 | 100 |
| **GATCCAGACTAACTTGGGGTGGCTTTGTCTTCTTCTTTTGCCAATTCCACTAATTGTTTGGG** | Del_k1_9-9 | -0.2780388368087761 | 0.038631688335210294 | 37.1818298846957 |
| **GATCCAGACTAATTGTTTGGG** | Del_k42_8-49;Del_k42_9-50 | -5.728376498300383 | 0.2580087221298507 | 0.15969053969393937 |
| **GATCCAGACTATCTAACTTGGGGTGGCTTTGTCTTCTTCTTTTGCCAATTCCACTAATTGTTTGGG** | Ins_k3_7_ACT;Ins_k3_8_CTA | -0.2067500098477904 | 0.038715677757554606 | 39.92924548495948 |
| **GATCCAGACTCTAACTTGGGGTGGCTTTGTCTTCTTCTTTTGCCAATTCCACTAATTGTTTGGG** | Ins_k1_8_C | 0.40828629435718056 | 0.03873068163505953 | 73.85807373661582 |
| **GATCCAGACTCTCTAACTTGGGGTGGCTTTGTCTTCTTCTTTTGCCAATTCCACTAATTGTTTGGG** | Ins_k3_8_CTC | -0.05207127014437585 | 0.04390446181876975 | 46.60872543301932 |
| **GATCCAGACTGTCTAACTTGGGGTGGCTTTGTCTTCTTCTTTTGCCAATTCCACTAATTGTTTGGG** | Ins_k3_8_CTG | 0.17784486546570522 | 0.03665546168889959 | 58.65682286785463 |
| **GATCCAGACTTCTAACTTGGGGTGGCTTTGTCTTCTTCTTTTGCCAATTCCACTAATTGTTTGGG** | Ins_k2_8_CT | 0.013132313185630018 | 0.04457764780237916 | 49.749049007069225 |
| **GATCCAGACTTCTTCTTTTGCCAATTCCACTAATTGTTTGGG** | Del_k21_9-29 | -0.27041004204764435 | 0.030930261360343443 | 37.46656715388405 |
| **GATCCAGACTTCTTTTGCCAATTCCACTAATTGTTTGGG** | Del_k24_9-32 | -0.979801465650737 | 0.03423852962247079 | 18.431433863110087 |
| **GATCCAGACTTGGGGTGGCTTTGTCTTCTTCTTTTGCCAATTCCACTAATTGTTTGGG** | Del_k5_8-12;Del_k5_9-13 | 0.5030682952664303 | 0.030058262336623627 | 81.20098113734844 |
| **GATCCAGACTTTCTAACTTGGGGTGGCTTTGTCTTCTTCTTTTGCCAATTCCACTAATTGTTTGGG** | Ins_k3_8_CTT | -0.610456660648239 | 0.046516863152031164 | 26.66634741915004 |
| **GATCCAGACTTTGTCTTCTTCTTTTGCCAATTCCACTAATTGTTTGGG** | Del_k15_9-23 | -0.10263767314475258 | 0.029317618237151986 | 44.3104863689173 |
| **GATCCAGACTTTTGCCAATTCCACTAATTGTTTGGG** | Del_k27_9-35 | -1.7650215976925376 | 0.04156756293584239 | 8.405089683907521 |
| **GATCCAGAG** | Del_k54_9-62 | -6.267363158812893 | 0.20439862634476386 | 0.09315373147018259 |
| **GATCCAGAGAATCTAACTTGGGGTGGCTTTGTCTTCTTCTTTTGCCAATTCCACTAATTGTTTGGG** | Ins_k3_7_AGA;Ins_k3_8_GAA | 0.36465555044060394 | 0.03296711013808743 | 70.70487934070104 |
| **GATCCAGAGACTCTAACTTGGGGTGGCTTTGTCTTCTTCTTTTGCCAATTCCACTAATTGTTTGGG** | Ins_k3_8_GAC | 0.3499376212652261 | 0.03613885773977987 | 69.67187045724732 |
| **GATCCAGAGAGTCTAACTTGGGGTGGCTTTGTCTTCTTCTTTTGCCAATTCCACTAATTGTTTGGG** | Ins_k3_8_GAG | 0.1781617701488467 | 0.03364353582233773 | 58.675414435441475 |
| **GATCCAGAGATCTAACTTGGGGTGGCTTTGTCTTCTTCTTTTGCCAATTCCACTAATTGTTTGGG** | Ins_k2_5_AG;Ins_k2_6_GA;Ins_k | 0.06999996102221706 | 0.03679839954215799 | 52.660149647006115 |
| **GATCCAGAGATTCTAACTTGGGGTGGCTTTGTCTTCTTCTTTTGCCAATTCCACTAATTGTTTGGG** | Ins_k3_8_GAT | 0.34069045380867014 | 0.03624390943496659 | 69.0305726679217 |
| **GATCCAGAGCATCTAACTTGGGGTGGCTTTGTCTTCTTCTTTTGCCAATTCCACTAATTGTTTGGG** | Ins_k3_7_AGC;Ins_k3_8_GCA | 0.3340519614275186 | 0.03555276560932984 | 68.5738314511081 |
| **GATCCAGAGCCAATTCCACTAATTGTTTGGG** | Del_k32_9-40 | -1.3997968623782673 | 0.26211412572401255 | 12.110370751595998 |
| **GATCCAGAGCCCTCTAACTTGGGGTGGCTTTGTCTTCTTCTTTTGCCAATTCCACTAATTGTTTGGG** | Ins_k4_8_GCCC | -0.26648521413797244 | 0.058437903927474714 | 37.613905932867986 |
| **GATCCAGAGCCTCTAACTTGGGGTGGCTTTGTCTTCTTCTTTTGCCAATTCCACTAATTGTTTGGG** | Ins_k3_8_GCC | 0.2952567831638243 | 0.03461730348143649 | 65.9644405946699 |
| **GATCCAGAGCGTCTAACTTGGGGTGGCTTTGTCTTCTTCTTTTGCCAATTCCACTAATTGTTTGGG** | Ins_k3_8_GCG | 0.6078354863630102 | 0.03064674544498172 | 90.1697965453006 |
| **GATCCAGAGCTAACTTGGGGTGGCTTTGTCTTCTTCTTTTGCCAATTCCACTAATTGTTTGGG** | T9G | 0.06730860533652949 | 0.03242254228125803 | 52.518613002005424 |
| **GATCCAGAGCTCTAACTTGGGGTGGCTTTGTCTTCTTCTTTTGCCAATTCCACTAATTGTTTGGG** | Ins_k2_8_GC | 0.05091510063606286 | 0.04400282892785677 | 51.66466757658784 |
| **GATCCAGAGCTTCTAACTTGGGGTGGCTTTGTCTTCTTCTTTTGCCAATTCCACTAATTGTTTGGG** | Ins_k3_8_GCT | 0.3030851804534005 | 0.03855990741711384 | 66.48286300334702 |
| **GATCCAGAGCTTTGTCTTCTTCTTTTGCCAATTCCACTAATTGTTTGGG** | Del_k14_9-22 | -0.03384610602967253 | 0.030293117581097147 | 47.46596504146944 |
| **GATCCAGAGG** | Del_k53_9-61 | -7.075570423525976 | 0.5299047574730378 | 0.04151454676423188 |
| **GATCCAGAGGATCTAACTTGGGGTGGCTTTGTCTTCTTCTTTTGCCAATTCCACTAATTGTTTGGG** | Ins_k3_6_GAG;Ins_k3_7_AGG;In | 0.6405725354238068 | 0.0316664992821723 | 93.17053934051177 |
| **GATCCAGAGGCTCTAACTTGGGGTGGCTTTGTCTTCTTCTTTTGCCAATTCCACTAATTGTTTGGG** | Ins_k3_8_GGC | 0.5070164360558398 | 0.036453839272323436 | 81.52220774979418 |
| **GATCCAGAGGCTTTGTCTTCTTCTTTTGCCAATTCCACTAATTGTTTGGG** | Del_k13_9-21 | 0.4533484594437489 | 0.02633678171104449 | 77.26240568254309 |
| **GATCCAGAGGG** | Del_k52_9-60 | -6.996961979009241 | 0.39498112490901016 | 0.04490963388051931 |
| **GATCCAGAGGGGTGGCTTTGTCTTCTTCTTTTGCCAATTCCACTAATTGTTTGGG** | Del_k8_9-16 | 0.5183245958415779 | 0.0299321321999639 | 82.44930590884644 |
| **GATCCAGAGGGTCTAACTTGGGGTGGCTTTGTCTTCTTCTTTTGCCAATTCCACTAATTGTTTGGG** | Ins_k3_8_GGG | -0.9901536663373396 | 0.05278392445195586 | 18.24161219206596 |
| **GATCCAGAGGGTGGCTTTGTCTTCTTCTTTTGCCAATTCCACTAATTGTTTGGG** | Del_k9_9-17 | 0.5370317273273224 | 0.028577725004193335 | 84.00621314566614 |
| **GATCCAGAGGTCTAACTTGGGGTGGCTTTGTCTTCTTCTTTTGCCAATTCCACTAATTGTTTGGG** | Ins_k2_8_GG | 0.2677984201879492 | 0.04268449955467905 | 64.17780632059582 |
| **GATCCAGAGGTGGCTTTGTCTTCTTCTTTTGCCAATTCCACTAATTGTTTGGG** | Del_k10_9-18 | 0.5093391691831184 | 0.029350243203874225 | 81.71178216249685 |
| **GATCCAGAGGTTCTAACTTGGGGTGGCTTTGTCTTCTTCTTTTGCCAATTCCACTAATTGTTTGGG** | Ins_k3_8_GGT | 0.29755608909594233 | 0.040051504352368296 | 66.11628752861728 |
| **GATCCAGAGTATCTAACTTGGGGTGGCTTTGTCTTCTTCTTTTGCCAATTCCACTAATTGTTTGGG** | Ins_k3_7_AGT;Ins_k3_8_GTA | -0.14572877104490362 | 0.0395291425473796 | 42.441653070608794 |
| **GATCCAGAGTCTAACTTGGGGTGGCTTTGTCTTCTTCTTTTGCCAATTCCACTAATTGTTTGGG** | Ins_k1_8_G | 0.464655568482536 | 0.039091076778789535 | 78.14097782320465 |
| **GATCCAGAGTCTCTAACTTGGGGTGGCTTTGTCTTCTTCTTTTGCCAATTCCACTAATTGTTTGGG** | Ins_k3_8_GTC | 0.08486329061196918 | 0.03780549889822522 | 53.44870053626367 |
| **GATCCAGAGTCTTCTTCTTTTGCCAATTCCACTAATTGTTTGGG** | Del_k19_9-27 | 0.15364023965331253 | 0.027918255253013072 | 57.254101079995415 |
| **GATCCAGAGTGGCTTTGTCTTCTTCTTTTGCCAATTCCACTAATTGTTTGGG** | Del_k11_9-19 | 0.4518503322406755 | 0.02783325949636479 | 77.14674343083055 |
| **GATCCAGAGTGTCTAACTTGGGGTGGCTTTGTCTTCTTCTTTTGCCAATTCCACTAATTGTTTGGG** | Ins_k3_8_GTG | 0.8099554611624324 | 0.23185686767114724 | 100 |
| **GATCCAGAGTTCTAACTTGGGGTGGCTTTGTCTTCTTCTTTTGCCAATTCCACTAATTGTTTGGG** | Ins_k2_8_GT | -0.14145276467369955 | 0.042779996373166866 | 42.62352240965774 |
| **GATCCAGAGTTTCTAACTTGGGGTGGCTTTGTCTTCTTCTTTTGCCAATTCCACTAATTGTTTGGG** | Ins_k3_8_GTT | -0.0926402430728922 | 0.047591675034857624 | 44.75569914142724 |
| **GATCCAGAGTTTGGG** | Del_k48_9-56 | -6.823181143397824 | 0.6490947766345301 | 0.05343324781000633 |
| **GATCCAGAT** | Del_k54_10-63 | -6.44925317748744 | 0.38913367505701596 | 0.07766161627609046 |
| **GATCCAGATAAACTAACTTGGGGTGGCTTTGTCTTCTTCTTTTGCCAATTCCACTAATTGTTTGGG** | Ins_k3_9_AAA | -0.7507626107237833 | 0.04883079731622049 | 23.17551710084803 |
| **GATCCAGATAAAGCTAACTTGGGGTGGCTTTGTCTTCTTCTTTTGCCAATTCCACTAATTGTTTGGG** | Ins_k4_9_AAAG | -0.85361611593529 | 0.05796317345122374 | 20.910322129910547 |
| **GATCCAGATAACCTAACTTGGGGTGGCTTTGTCTTCTTCTTTTGCCAATTCCACTAATTGTTTGGG** | Ins_k3_9_AAC | 0.003076960721323907 | 0.036369150750364274 | 49.25131144171767 |
| **GATCCAGATAACTAACTTGGGGTGGCTTTGTCTTCTTCTTTTGCCAATTCCACTAATTGTTTGGG** | Ins_k2_9_AA | -0.8434935827964611 | 0.055046829186969605 | 21.12306247767873 |
| **GATCCAGATAACTCTAACTTGGGGTGGCTTTGTCTTCTTCTTTTGCCAATTCCACTAATTGTTTGGG** | Ins_k4_8_TAAC;Ins_k4_9_AACT | -0.43237681395064675 | 0.04901175786353047 | 31.86417126851982 |

| | | | | |
|---|---|---|---|---|
| **GATCCAGATAACTTGGGGTGGCTTTGTCTTCTTCTTTTGCCAATTCCACTAATTGTTTGGG** | Del_k2_9-10;Del_k2_10-11 | -0.013193777415571373 | 0.034673551960435464 | 48.456440355895325 |
| **GATCCAGATAAGCTAACTTGGGGTGGCTTTGTCTTCTTCTTTTGCCAATTCCACTAATTGTTTGGG** | Ins_k3_9_AAG | -0.5695772725116436 | 0.053282400070480344 | 27.77903952206634 |
| **GATCCAGATAATCTAACTTGGGGTGGCTTTGTCTTCTTCTTTTGCCAATTCCACTAATTGTTTGGG** | Ins_k3_7_ATA;Ins_k3_8_TAA;Ins_ | -0.5733039481246639 | 0.04687914535603349 | 27.675708712719082 |
| **GATCCAGATAATTCCACTAATTGTTTGGG** | Del_k34_10-43 | -3.5149446251329093 | 0.08348669680467341 | 1.4606980087648014 |
| **GATCCAGATAATTGTTTGGG** | Del_k43_9-51;Del_k43_10-52 | -6.242602162064536 | 0.5181770558944043 | 0.09548910447567634 |
| **GATCCAGATACACTAACTTGGGGTGGCTTTGTCTTCTTCTTTTGCCAATTCCACTAATTGTTTGGG** | Ins_k3_9_ACA | -0.3615343769016385 | 0.04290524482571241 | 34.203386440644934 |
| **GATCCAGATACCCTAACTTGGGGTGGCTTTGTCTTCTTCTTTTGCCAATTCCACTAATTGTTTGGG** | Ins_k3_9_ACC | 0.07585028852028797 | 0.03534962113504435 | 52.96913171039306 |
| **GATCCAGATACCTAACTTGGGGTGGCTTTGTCTTCTTCTTTTGCCAATTCCACTAATTGTTTGGG** | Ins_k2_9_AC | 0.1920072884830213 | 0.04519875338323133 | 59.49345599844034 |
| **GATCCAGATACGCTAACTTGGGGTGGCTTTGTCTTCTTCTTTTGCCAATTCCACTAATTGTTTGGG** | Ins_k3_9_ACG | 0.3193622461602075 | 0.03236536262026862 | 67.57386399258527 |
| **GATCCAGATACTAACTTGGGGTGGCTTTGTCTTCTTCTTTTGCCAATTCCACTAATTGTTTGGG** | Ins_k1_9_A | -0.16483456093185578 | 0.20605590141264613 | 41.63846893042267 |
| **GATCCAGATACTAATTGTTTGGG** | Del_k40_10-49 | -5.77241496910999 | 0.3287985753806142 | 0.1528106150699621 |
| **GATCCAGATACTCTAACTTGGGGTGGCTTTGTCTTCTTCTTTTGCCAATTCCACTAATTGTTTGGG** | Ins_k3_8_TAC;Ins_k3_9_ACT | -0.4459340364168849 | 0.03975560988728176 | 31.435096711452932 |
| **GATCCAGATACTTGGGGTGGCTTTGTCTTCTTCTTTTGCCAATTCCACTAATTGTTTGGG** | Del_k3_10-12 | 0.2653981645107437 | 0.029537647007136056 | 64.02394790026543 |
| **GATCCAGATAGACTAACTTGGGGTGGCTTTGTCTTCTTCTTTTGCCAATTCCACTAATTGTTTGGG** | Ins_k3_9_AGA | -0.6188853068984966 | 0.3412740522994148 | 26.442530770676743 |
| **GATCCAGATAGCCTAACTTGGGGTGGCTTTGTCTTCTTCTTTTGCCAATTCCACTAATTGTTTGGG** | Ins_k3_9_AGC | -0.38119205626053826 | 0.04407814330477116 | 33.537592657137054 |
| **GATCCAGATAGCTAACTTGGGGTGGCTTTGTCTTCTTCTTTTGCCAATTCCACTAATTGTTTGGG** | Ins_k2_9_AG | -1.636694613180563 | 0.07498166198717282 | 9.555954056356514 |
| **GATCCAGATAGGCTAACTTGGGGTGGCTTTGTCTTCTTCTTTTGCCAATTCCACTAATTGTTTGGG** | Ins_k3_9_AGG | -2.9521294581196678 | 0.12939311611544865 | 2.5644129337118304 |
| **GATCCAGATAGTCTAACTTGGGGTGGCTTTGTCTTCTTCTTTTGCCAATTCCACTAATTGTTTGGG** | Ins_k3_8_TAG;Ins_k3_9_AGT | -1.983365258848737 | 0.08195385279567467 | 6.756424134766598 |
| **GATCCAGATATAACTTGGGGTGGCTTTGTCTTCTTCTTTTGCCAATTCCACTAATTGTTTGGG** | C10A | -0.6428386497817586 | 0.03922153784599732 | 25.81666942450508 |
| **GATCCAGATATACTAACTTGGGGTGGCTTTGTCTTCTTCTTTTGCCAATTCCACTAATTGTTTGGG** | Ins_k3_9_ATA | -1.0919244499780643 | 0.05726024681780223 | 16.47649112802334 |
| **GATCCAGATATCCTAACTTGGGGTGGCTTTGTCTTCTTCTTTTGCCAATTCCACTAATTGTTTGGG** | Ins_k3_9_ATC | -0.13769426592122597 | 0.04160925531509721 | 42.7840242995954 |
| **GATCCAGATATCTAACTTGGGGTGGCTTTGTCTTCTTCTTTTGCCAATTCCACTAATTGTTTGGG** | Ins_k2_7_AT;Ins_k2_8_TA;Ins_k2_ | -0.3431193078178472 | 0.04750022876205568 | 34.839079365105604 |
| **GATCCAGATATGCTAACTTGGGGTGGCTTTGTCTTCTTCTTTTGCCAATTCCACTAATTGTTTGGG** | Ins_k3_9_ATG | -0.22140050048798837 | 0.03828148719576255 | 39.34852674190105 |
| **GATCCAGATATTCCACTAATTGTTTGGG** | Del_k35_10-44 | -3.6500870342197924 | 0.0772320880344277 | 1.2760533578953055 |
| **GATCCAGATATTCTAACTTGGGGTGGCTTTGTCTTCTTCTTTTGCCAATTCCACTAATTGTTTGGG** | Ins_k3_8_TAT;Ins_k3_9_ATT | -0.2692516711920892 | 0.04704779275665464 | 37.50999247979301 |
| **GATCCAGATATTGTTTGGG** | Del_k44_10-53 | -6.3297002107392375 | 0.5292393575202691 | 0.08752409291333696 |
| **GATCCAGATC** | Del_k53_11-63 | -6.790687544691106 | 0.3314572507019462 | 0.05519800266876294 |
| **GATCCAGATCAAACTTGGGGTGGCTTTGTCTTCTTCTTTTGCCAATTCCACTAATTGTTTGGG** | T11A | 0.11074415971796436 | 0.030640061543692598 | 54.85005527770428 |
| **GATCCAGATCAAAGTAACTTGGGGTGGCTTTGTCTTCTTCTTTTGCCAATTCCACTAATTGTTTGGG** | Ins_k4_10_AAAG | 0.04262379629401414 | 0.040201850654325286 | 51.23807105864819 |
| **GATCCAGATCAAATAACTTGGGGTGGCTTTGTCTTCTTCTTTTGCCAATTCCACTAATTGTTTGGG** | Ins_k3_10_AAA | -0.04328331181282796 | 0.035616565643702344 | 47.02012600792536 |
| **GATCCAGATCAACTAACTTGGGGTGGCTTTGTCTTCTTCTTTTGCCAATTCCACTAATTGTTTGGG** | Ins_k3_9_CAA;Ins_k3_10_AAC | 0.3122527222767091 | 0.03136914554731981 | 67.09514972428748 |
| **GATCCAGATCAACTTGGGGTGGCTTTGTCTTCTTCTTTTGCCAATTCCACTAATTGTTTGGG** | Del_k1_11-11 | 0.4771384557994598 | 0.029422920257632004 | 79.12251631596037 |
| **GATCCAGATCAAGTAACTTGGGGTGGCTTTGTCTTCTTCTTTTGCCAATTCCACTAATTGTTTGGG** | Ins_k3_10_AAG | 0.2590389668352646 | 0.03217032565017931 | 63.6180987647329 |
| **GATCCAGATCAATAACTTGGGGTGGCTTTGTCTTCTTCTTTTGCCAATTCCACTAATTGTTTGGG** | Ins_k2_10_AA | 0.00653115136339133 | 0.04456455479639578 | 49.42172901876534 |
| **GATCCAGATCAATTAACTTGGGGTGGCTTTGTCTTCTTCTTTTGCCAATTCCACTAATTGTTTGGG** | Ins_k3_10_AAT | -0.2320734788157176 | 0.03884527866795749 | 38.93079396062409 |
| **GATCCAGATCAATTCCACTAATTGTTTGGG** | Del_k33_10-42;Del_k33_11-43 | -3.1939253264478586 | 0.06753042415965202 | 2.0136192169107785 |
| **GATCCAGATCAATTGTTTGGG** | Del_k42_11-52 | -5.702096040446004 | 0.21316282483247323 | 0.16394291260222865 |
| **GATCCAGATCACATAACTTGGGGTGGCTTTGTCTTCTTCTTTTGCCAATTCCACTAATTGTTTGGG** | Ins_k3_10_ACA | 0.19015804303283634 | 0.036187151386251164 | 59.38353965808908 |
| **GATCCAGATCACCTAACTTGGGGTGGCTTTGTCTTCTTCTTTTGCCAATTCCACTAATTGTTTGGG** | Ins_k3_9_CAC;Ins_k3_10_ACC | 0.4374275426183789 | 0.03158884697817855 | 76.0420576549272 |
| **GATCCAGATCACGTAACTTGGGGTGGCTTTGTCTTCTTCTTTTGCCAATTCCACTAATTGTTTGGG** | Ins_k3_10_ACG | 0.2378104800686458 | 0.02983486749863206 | 62.28181660351209 |
| **GATCCAGATCACTAACTTGGGGTGGCTTTGTCTTCTTCTTTTGCCAATTCCACTAATTGTTTGGG** | Ins_k2_9_CA;Ins_k2_10_AC | -0.4085671548796968 | 0.0468275214394142 | 32.63195033105075 |
| **GATCCAGATCACTAATTGTTTGGG** | Del_k39_10-48;Del_k39_11-49 | -5.487347463183731 | 0.2020588314152536 | 0.203215471199515 |
| **GATCCAGATCACTTAACTTGGGGTGGCTTTGTCTTCTTCTTTTGCCAATTCCACTAATTGTTTGGG** | Ins_k3_10_ACT | -0.3047626025884213 | 0.03804936120287742 | 36.20135076956861 |
| **GATCCAGATCACTTGGGGTGGCTTTGTCTTCTTCTTTTGCCAATTCCACTAATTGTTTGGG** | Del_k2_11-12 | 0.5456712844210194 | 0.0293173708211047 | 84.73513386243859 |
| **GATCCAGATCAGACTAACTTGGGGTGGCTTTGTCTTCTTCTTTTGCCAATTCCACTAATTGTTTGGG** | Ins_k4_9_CAGA;Ins_k4_10_AGA( | 0.27159574438537926 | 0.03664996899604124 | 64.42197355514811 |
| **GATCCAGATCAGAGTAACTTGGGGTGGCTTTGTCTTCTTCTTTTGCCAATTCCACTAATTGTTTGGG** | Ins_k4_10_AGAG | 0.24416070864536255 | 0.03559797284529546 | 62.67857880716068 |
| **GATCCAGATCAGATAACTTGGGGTGGCTTTGTCTTCTTCTTTTGCCAATTCCACTAATTGTTTGGG** | Ins_k3_10_AGA | 0.06923606496945311 | 0.04066708362812445 | 52.61993812721883 |
| **GATCCAGATCAGCTAACTTGGGGTGGCTTTGTCTTCTTCTTTTGCCAATTCCACTAATTGTTTGGG** | Ins_k3_9_CAG;Ins_k3_10_AGC | 0.19822194621702283 | 0.03392068111736674 | 59.86433872580416 |
| **GATCCAGATCAGGTAACTTGGGGTGGCTTTGTCTTCTTCTTTTGCCAATTCCACTAATTGTTTGGG** | Ins_k3_10_AGG | -4.465759263269186 | 0.3869239998928535 | 0.5644518315336446 |
| **GATCCAGATCAGTAACTTGGGGTGGCTTTGTCTTCTTCTTTTGCCAATTCCACTAATTGTTTGGG** | Ins_k2_10_AG | -0.4123943357994034 | 0.04562679672309202 | 32.50730063399063 |
| **GATCCAGATCAGTGTAACTTGGGGTGGCTTTGTCTTCTTCTTTTGCCAATTCCACTAATTGTTTGGG** | Ins_k4_10_AGTG | -0.04700980407287725 | 0.03987095317429681 | 46.84523194536789 |
| **GATCCAGATCAGTTAACTTGGGGTGGCTTTGTCTTCTTCTTTTGCCAATTCCACTAATTGTTTGGG** | Ins_k3_10_AGT | -0.528661676125253 | 0.0405305881238682 | 28.93920813927003 |
| **GATCCAGATCATAACTTGGGGTGGCTTTGTCTTCTTCTTTTGCCAATTCCACTAATTGTTTGGG** | Ins_k1_10_A | 0.27986569837743014 | 0.03996444766477636 | 64.95694937623168 |
| **GATCCAGATCATATAACTTGGGGTGGCTTTGTCTTCTTCTTTTGCCAATTCCACTAATTGTTTGGG** | Ins_k3_10_ATA | -0.1957341222535991 | 0.039289753429814354 | 40.37153318813669 |
| **GATCCAGATCATCTAACTTGGGGTGGCTTTGTCTTCTTCTTTTGCCAATTCCACTAATTGTTTGGG** | Ins_k3_7_ATC;Ins_k3_8_TCA;Ins_ | 0.3550459692287463 | 0.031995858518109224 | 70.02868921606392 |
| **GATCCAGATCATGTAACTTGGGGTGGCTTTGTCTTCTTCTTTTGCCAATTCCACTAATTGTTTGGG** | Ins_k3_10_ATG | 0.016013728824397777 | 0.04034981110641661 | 49.89260341552842 |
| **GATCCAGATCATTAACTTGGGGTGGCTTTGTCTTCTTCTTTTGCCAATTCCACTAATTGTTTGGG** | Ins_k2_10_AT | -0.31245807416966165 | 0.04538027724191919 | 35.92383348625539 |
| **GATCCAGATCATTCCACTAATTGTTTGGG** | Del_k34_11-44 | -3.3374439506377063 | 0.06220329248633248 | 1.7444077227834227 |
| **GATCCAGATCATTGTTTGGG** | Del_k43_11-53 | -6.025484335171702 | 0.2690305459481467 | 0.11864430442996915 |
| **GATCCAGATCATTTAACTTGGGGTGGCTTTGTCTTCTTCTTTTGCCAATTCCACTAATTGTTTGGG** | Ins_k3_10_ATT | -0.3462662352826033 | 0.04388870376059675 | 34.72961563694733 |
| **GATCCAGATCCAACTTGGGGTGGCTTTGTCTTCTTCTTTTGCCAATTCCACTAATTGTTTGGG** | T11C | 0.08839801409994008 | 0.028053037374350283 | 53.637961208466294 |
| **GATCCAGATCCAATAACTTGGGGTGGCTTTGTCTTCTTCTTTTGCCAATTCCACTAATTGTTTGGG** | Ins_k3_10_CAA | 0.4451340606121599 | 0.032802176564092386 | 76.63034103723444 |
| **GATCCAGATCCAATTCCACTAATTGTTTGGG** | Del_k32_10-41;Del_k32_11-42 | -2.8952789846886637 | 0.06487213950266804 | 2.714424743562366 |
| **GATCCAGATCCACTAACTTGGGGTGGCTTTGTCTTCTTCTTTTGCCAATTCCACTAATTGTTTGGG** | Ins_k3_9_CCA;Ins_k3_10_CAC | 0.2852459750787685 | 0.03750103757994076 | 65.30737759260357 |
| **GATCCAGATCCACTAATTGTTTGGG** | Del_k38_9-46;Del_k38_10-47;De | -4.5719674169716775 | 0.10404440948504662 | 0.50757622506553 |
| **GATCCAGATCCAGTAACTTGGGGTGGCTTTGTCTTCTTCTTTTGCCAATTCCACTAATTGTTTGGG** | Ins_k3_10_CAG | 0.5745427871100977 | 0.03026166392650056 | 87.21722291544907 |
| **GATCCAGATCCATAACTTGGGGTGGCTTTGTCTTCTTCTTTTGCCAATTCCACTAATTGTTTGGG** | Ins_k2_10_CA | 0.4353585379618692 | 0.03669202357191372 | 75.88488893102645 |
| **GATCCAGATCCATTAACTTGGGGTGGCTTTGTCTTCTTCTTTTGCCAATTCCACTAATTGTTTGGG** | Ins_k3_10_CAT | 0.3504136436667439 | 0.03148630718023934 | 69.7050437233231 |
| **GATCCAGATCCCAATTCCACTAATTGTTTGGG** | Del_k31_11-41 | -3.024005140315187 | 0.06628818944701532 | 2.38656213816069 |
| **GATCCAGATCCCACTAATTGTTTGGG** | Del_k37_11-47 | -4.363350909806266 | 0.10831872294148812 | 0.6253199282553034 |
| **GATCCAGATCCCATAACTTGGGGTGGCTTTGTCTTCTTCTTTTGCCAATTCCACTAATTGTTTGGG** | Ins_k3_10_CCA | 0.257140696423774 | 0.03342492956540849 | 63.49744895941974 |
| **GATCCAGATCCCCTAACTTGGGGTGGCTTTGTCTTCTTCTTTTGCCAATTCCACTAATTGTTTGGG** | Ins_k3_9_CCC;Ins_k3_10_CCC | 0.31620027480780377 | 0.034566510102497124 | 67.36053481857776 |
| **GATCCAGATCCCGTAACTTGGGGTGGCTTTGTCTTCTTCTTTTGCCAATTCCACTAATTGTTTGGG** | Ins_k3_10_CCG | 0.5893672577057605 | 0.03292239105788716 | 88.51980324888483 |
| **GATCCAGATCCCTAACTTGGGGTGGCTTTGTCTTCTTCTTTTGCCAATTCCACTAATTGTTTGGG** | Ins_k2_9_CC;Ins_k2_10_CC | 0.09769475039719622 | 0.045101475726806534 | 54.13894433505571 |
| **GATCCAGATCCCTTAACTTGGGGTGGCTTTGTCTTCTTCTTTTGCCAATTCCACTAATTGTTTGGG** | Ins_k3_10_CCT | 0.16143900935810818 | 0.034043448046111355 | 57.70235828438131 |

| | | | | |
|---|---|---|---|---|
| GATCCAGATCCGATAACTTGGGGTGGCTTTGTCTTCTTCTTTTGCCAATTCCACTAATTGTTTGGG | Ins_k3_10_CGA | 0.5350395668326124 | 0.030029669480097106 | 83.83902587375937 |
| GATCCAGATCCGCTAACTTGGGGTGGCTTTGTCTTCTTCTTTTGCCAATTCCACTAATTGTTTGGG | Ins_k3_9_CCG;Ins_k3_10_CGC | 0.47060423590157563 | 0.03254093823456023 | 78.60719783112975 |
| GATCCAGATCCGGTAACTTGGGGTGGCTTTGTCTTCTTCTTTTGCCAATTCCACTAATTGTTTGGG | Ins_k3_10_CGG | 0.47347381705869207 | 0.031118071813343178 | 78.83309151996616 |
| GATCCAGATCCGTAACTTGGGGTGGCTTTGTCTTCTTCTTTTGCCAATTCCACTAATTGTTTGGG | Ins_k2_10_CG | 0.4998779126254036 | 0.0403474622618292 | 80.94233175133854 |
| GATCCAGATCCGTTAACTTGGGGTGGCTTTGTCTTCTTCTTTTGCCAATTCCACTAATTGTTTGGG | Ins_k3_10_CGT | 0.4838145908641942 | 0.029400751383685905 | 79.65251612817208 |
| GATCCAGATCCTAACTTGGGGTGGCTTTGTCTTCTTCTTTTGCCAATTCCACTAATTGTTTGGG | Ins_k1_9_C;Ins_k1_10_C | 0.507343393826436 | 0.03297070481918305 | 81.54886642698747 |
| GATCCAGATCCTAATTGTTTGGG | Del_k40_11-50 | -5.387034589069279 | 0.2786626155665169 | 0.22465810758444024 |
| GATCCAGATCCTACTAACTTGGGGTGGCTTTGTCTTCTTCTTTTGCCAATTCCACTAATTGTTTGGG | Ins_k4_9_CCTA;Ins_k4_10_CTAC | -0.13505821299107995 | 0.041243090829337595 | 42.89695403119513 |
| GATCCAGATCCTATAACTTGGGGTGGCTTTGTCTTCTTCTTTTGCCAATTCCACTAATTGTTTGGG | Ins_k3_10_CTA | -0.055502194595683174 | 0.04113626648861833 | 46.449088425137546 |
| GATCCAGATCCTCTAACTTGGGGTGGCTTTGTCTTCTTCTTTTGCCAATTCCACTAATTGTTTGGG | Ins_k3_8_TCC;Ins_k3_9_CCT;Ins | 0.12703843805964643 | 0.0354042173861616 | 55.75111849670394 |
| GATCCAGATCCTGTAACTTGGGGTGGCTTTGTCTTCTTCTTTTGCCAATTCCACTAATTGTTTGGG | Ins_k3_10_CTG | -1.5921631378432204 | 0.08482399261531787 | 9.991111991808541 |
| GATCCAGATCCTTAACTTGGGGTGGCTTTGTCTTCTTCTTTTGCCAATTCCACTAATTGTTTGGG | Ins_k2_10_CT | -0.16649345886307232 | 0.04584612285255547 | 41.56945222211978 |
| GATCCAGATCCTTCTTCTTTTGCCAATTCCACTAATTGTTTGGG | Del_k19_11-29 | -0.0560754500575158 | 0.029015058696973815 | 46.42246886213461 |
| GATCCAGATCCTTCTTTTGCCAATTCCACTAATTGTTTGGG | Del_k22_11-32 | -0.6828864168435669 | 0.03492704322642776 | 24.80319853293672 |
| GATCCAGATCCTTGGGGTGGCTTTGTCTTCTTCTTTTGCCAATTCCACTAATTGTTTGGG | Del_k3_11-13 | 0.40887467771503894 | 0.03169764467661932 | 73.90154338520074 |
| GATCCAGATCCTTTAACTTGGGGTGGCTTTGTCTTCTTCTTTTGCCAATTCCACTAATTGTTTGGG | Ins_k3_10_CTT | -0.11261732869584928 | 0.03850212203421623 | 43.87048217459545 |
| GATCCAGATCCTTTGTCTTCTTCTTTTGCCAATTCCACTAATTGTTTGGG | Del_k13_11-23 | 0.011253888821036173 | 0.030450745500594394 | 49.65568689558362 |
| GATCCAGATCCTTTTGCCAATTCCACTAATTGTTTGGG | Del_k25_11-35 | -1.4489438103072616 | 0.04212225043664598 | 11.529572132086285 |
| GATCCAGATCG | Del_k52_11-62 | -7.095480650059895 | 0.46261626338548567 | 0.04069615693230947 |
| GATCCAGATCGAACTTGGGGTGGCTTTGTCTTCTTCTTTTGCCAATTCCACTAATTGTTTGGG | T11G | 0.52121231368444 | 0.03158912379791573 | 82.6877403406422 |
| GATCCAGATCGAATAACTTGGGGTGGCTTTGTCTTCTTCTTTTGCCAATTCCACTAATTGTTTGGG | Ins_k3_10_GAA | 0.6249283454285646 | 0.02717812055836868 | 91.72430380876327 |
| GATCCAGATCGACTAACTTGGGGTGGCTTTGTCTTCTTCTTTTGCCAATTCCACTAATTGTTTGGG | Ins_k3_9_CGA;Ins_k3_10_GAC | 0.6064913777079297 | 0.02961505883651852 | 90.04867995649924 |
| GATCCAGATCGAGATAACTTGGGGTGGCTTTGTCTTCTTCTTTTGCCAATTCCACTAATTGTTTGGG | Ins_k4_10_GAGA | 0.9003857621312317 | 0.031055211801878053 | 100 |
| GATCCAGATCGAGTAACTTGGGGTGGCTTTGTCTTCTTCTTTTGCCAATTCCACTAATTGTTTGGG | Ins_k3_10_GAG | 0.5601059530139243 | 0.029759758979946878 | 85.96712775830476 |
| GATCCAGATCGATAACTTGGGGTGGCTTTGTCTTCTTCTTTTGCCAATTCCACTAATTGTTTGGG | Ins_k2_10_GA | 0.4858613685338786 | 0.03501931590082046 | 79.81571407750288 |
| GATCCAGATCGATCTAACTTGGGGTGGCTTTGTCTTCTTCTTTTGCCAATTCCACTAATTGTTTGGG | Ins_k4_6_GATC;Ins_k4_7_ATCG; | 0.6854712471462692 | 0.032884856554871916 | 97.44910892060885 |
| GATCCAGATCGATTAACTTGGGGTGGCTTTGTCTTCTTCTTTTGCCAATTCCACTAATTGTTTGGG | Ins_k3_10_GAT | 0.2064611222623145 | 0.1545444008964287 | 60.359609059393236 |
| GATCCAGATCGCATAACTTGGGGTGGCTTTGTCTTCTTCTTTTGCCAATTCCACTAATTGTTTGGG | Ins_k3_10_GCA | 0.662377467100808 | 0.15225720840107781 | 95.22442765534421 |
| GATCCAGATCGCCAATTCCACTAATTGTTTGGG | Del_k30_11-40 | -1.192165185268083 | 0.030836565300170544 | 14.904957109468844 |
| GATCCAGATCGCCTAACTTGGGGTGGCTTTGTCTTCTTCTTTTGCCAATTCCACTAATTGTTTGGG | Ins_k3_9_CGC;Ins_k3_10_GCC | 0.6761652155015534 | 0.030825414310432925 | 96.54645102505363 |
| GATCCAGATCGCGTAACTTGGGGTGGCTTTGTCTTCTTCTTTTGCCAATTCCACTAATTGTTTGGG | Ins_k3_10_GCG | 0.5581279090266107 | 0.027799636282997996 | 85.79724906728725 |
| GATCCAGATCGCTAACTTGGGGTGGCTTTGTCTTCTTCTTTTGCCAATTCCACTAATTGTTTGGG | Ins_k2_9_CG;Ins_k2_10_GC | 0.39301586609265415 | 0.03338192830097477 | 72.73879698740411 |
| GATCCAGATCGCTTAACTTGGGGTGGCTTTGTCTTCTTCTTTTGCCAATTCCACTAATTGTTTGGG | Ins_k3_10_GCT | 0.4450110574087862 | 0.031162019434711488 | 76.62091583948789 |
| GATCCAGATCGCTTTGTCTTCTTCTTTTGCCAATTCCACTAATTGTTTGGG | Del_k12_11-22 | 0.21511371586676697 | 0.027670355234119808 | 60.88414224040029 |
| GATCCAGATCGG | Del_k51_11-61 | -5.822846047153791 | 0.29313308502329494 | 0.1452953063122531 |
| GATCCAGATCGGATAACTTGGGGTGGCTTTGTCTTCTTCTTTTGCCAATTCCACTAATTGTTTGGG | Ins_k3_10_GGA | 0.6180608454959391 | 0.02772116764448418 | 91.0965451918891 |
| GATCCAGATCGGCTAACTTGGGGTGGCTTTGTCTTCTTCTTTTGCCAATTCCACTAATTGTTTGGG | Ins_k3_9_CGG;Ins_k3_10_GGC | 0.5845470457482727 | 0.029121012860439718 | 88.09414573865966 |
| GATCCAGATCGGCTTTGTCTTCTTCTTTTGCCAATTCCACTAATTGTTTGGG | Del_k11_11-21 | 0.70803397709618765 | 0.026932239783670573 | 99.67281899823162 |
| GATCCAGATCGGG | Del_k50_11-60 | -7.089952434751152 | 0.3967997859883931 | 0.040921757058555545 |
| GATCCAGATCGGGCTAACTTGGGGTGGCTTTGTCTTCTTCTTTTGCCAATTCCACTAATTGTTTGGG | Ins_k4_9_CGGG;Ins_k4_10_GGC | 0.2567806454934455 | 0.0349949947166726 | 63.474590759153486 |
| GATCCAGATCGGGGTGGCTTTGTCTTCTTCTTTTGCCAATTCCACTAATTGTTTGGG | Del_k6_11-16 | 0.5789542380021206 | 0.027109530398723648 | 87.60282732338337 |
| GATCCAGATCGGGTAACTTGGGGTGGCTTTGTCTTCTTCTTTTGCCAATTCCACTAATTGTTTGGG | Ins_k3_10_GGG | -0.8275296175163454 | 0.04520833058891198 | 21.462976280921318 |
| GATCCAGATCGGGTGGCTTTGTCTTCTTCTTTTGCCAATTCCACTAATTGTTTGGG | Del_k7_11-17 | 0.771529429117179 | 0.02646498354314802 | 100 |
| GATCCAGATCGGGTTAACTTGGGGTGGCTTTGTCTTCTTCTTTTGCCAATTCCACTAATTGTTTGGG | Ins_k4_10_GGGT | -0.5708428946434145 | 0.043022775656190046 | 27.74390399368904 |
| GATCCAGATCGGTAACTTGGGGTGGCTTTGTCTTCTTCTTTTGCCAATTCCACTAATTGTTTGGG | Ins_k2_10_GG | -0.2736188706376592 | 0.04666673866272346 | 37.34653604465094 |
| GATCCAGATCGGTGGCTTTGTCTTCTTCTTTTGCCAATTCCACTAATTGTTTGGG | Del_k8_11-18 | 0.5847279356566639 | 0.025370336434010453 | 88.11008252197031 |
| GATCCAGATCGGTTAACTTGGGGTGGCTTTGTCTTCTTCTTTTGCCAATTCCACTAATTGTTTGGG | Ins_k3_10_GGT | 0.13451809019704808 | 0.033472500481317814 | 56.1696808684218 |
| GATCCAGATCGTAACTTGGGGTGGCTTTGTCTTCTTCTTTTGCCAATTCCACTAATTGTTTGGG | Ins_k1_10_G | 0.564190792921883 | 0.034746943256564314 | 86.31900791030154 |
| GATCCAGATCGTATAACTTGGGGTGGCTTTGTCTTCTTCTTTTGCCAATTCCACTAATTGTTTGGG | Ins_k3_10_GTA | 0.1955563741656462 | 0.034930728142989434 | 59.70497850505593 |
| GATCCAGATCGTCTAACTTGGGGTGGCTTTGTCTTCTTCTTTTGCCAATTCCACTAATTGTTTGGG | Ins_k3_8_TCG;Ins_k3_9_CGT;Ins | 0.7504459368722269 | 0.029065416953853826 | 100 |
| GATCCAGATCGTCTTCTTCTTTTGCCAATTCCACTAATTGTTTGGG | Del_k17_11-27 | 0.4487968464857408 | 0.024877259531463882 | 76.91153623239141 |
| GATCCAGATCGTGGCTTTGTCTTCTTCTTTTGCCAATTCCACTAATTGTTTGGG | Del_k9_11-19 | 0.5160623980984018 | 0.025536224843319875 | 82.26300008485258 |
| GATCCAGATCGTGTAACTTGGGGTGGCTTTGTCTTCTTCTTTTGCCAATTCCACTAATTGTTTGGG | Ins_k3_10_GTG | 0.5040836152690653 | 0.032596949252262045 | 81.28346798591829 |
| GATCCAGATCGTTAACTTGGGGTGGCTTTGTCTTCTTCTTTTGCCAATTCCACTAATTGTTTGGG | Ins_k2_10_GT | 0.1953638207528422 | 0.033857690800154126 | 59.693483214447625 |
| GATCCAGATCGTTTAACTTGGGGTGGCTTTGTCTTCTTCTTTTGCCAATTCCACTAATTGTTTGGG | Ins_k3_10_GTT | 0.4055987196388715 | 0.02974741268946727 | 73.65984114666728 |
| GATCCAGATCGTTTGGG | Del_k46_11-56 | -6.953869528166113 | 0.36282284975548595 | 0.046887203212840046 |
| GATCCAGATCT | Del_k52_12-63 | -7.000953323782486 | 0.4818096587886961 | 0.04473074129654423 |
| GATCCAGATCTAA | Del_k50_14-63 | -6.712766373458638 | 0.6527739930316279 | 0.059671107413274437 |
| GATCCAGATCTAAAAACTTGGGGTGGCTTTGTCTTCTTCTTTTGCCAATTCCACTAATTGTTTGGG | Ins_k3_11_AAA;Ins_k3_12_AAA;I | -0.057261018735464964 | 0.037167836712118034 | 46.36746444929582 |
| GATCCAGATCTAAAACCTTGGGGTGGCTTTGTCTTCTTCTTTTGCCAATTCCACTAATTGTTTGGG | Ins_k3_13_AAC | 0.27172557315106183 | 0.03661284261065715 | 64.43033792341369 |
| GATCCAGATCTAAAACTTGGGGTGGCTTTGTCTTCTTCTTTTGCCAATTCCACTAATTGTTTGGG | Ins_k2_11_AA;Ins_k2_12_AA;Ins_ | 0.07114179407895582 | 0.04019154471914361 | 52.7203130884123 |
| GATCCAGATCTAAAAGCTTGGGGTGGCTTTGTCTTCTTCTTTTGCCAATTCCACTAATTGTTTGGG | Ins_k3_13_AAG | 0.32874244667864316 | 0.03869831682276199 | 68.21070255382803 |
| GATCCAGATCTAAAATCTTGGGGTGGCTTTGTCTTCTTCTTTTGCCAATTCCACTAATTGTTTGGG | Ins_k3_13_AAT | -0.29504114118840735 | 0.038712321692414335 | 36.55499699762057 |
| GATCCAGATCTAAAATTCCACTAATTGTTTGGG | Del_k30_14-43 | -3.4034608264034465 | 0.08714389085365407 | 1.6329663514160835 |
| GATCCAGATCTAAAATTGTTTGGG | Del_k39_14-52 | -5.707642010060498 | 0.2137370809239406 | 0.1630362067958709 |
| GATCCAGATCTAAACACTTGGGGTGGCTTTGTCTTCTTCTTTTGCCAATTCCACTAATTGTTTGGG | Ins_k3_12_AAC;Ins_k3_13_ACA | 0.12523261785860773 | 0.03507921415603416 | 55.650532847824465 |
| GATCCAGATCTAAACCCTTGGGGTGGCTTTGTCTTCTTCTTTTGCCAATTCCACTAATTGTTTGGG | Ins_k3_13_ACC | 0.43540507497293657 | 0.03393861077615245 | 75.88842046911546 |
| GATCCAGATCTAAACCTTGGGGTGGCTTTGTCTTCTTCTTTTGCCAATTCCACTAATTGTTTGGG | Ins_k2_13_AC | 0.2267486110114727 | 0.034672515998350174 | 61.596659847344625 |
| GATCCAGATCTAAACGCTTGGGGTGGCTTTGTCTTCTTCTTTTGCCAATTCCACTAATTGTTTGGG | Ins_k3_13_ACG | 0.48843615794285755 | 0.02974818625175796 | 80.02148753075181 |
| GATCCAGATCTAAACTAATTGTTTGGG | Del_k36_14-49 | -5.919618083715781 | 0.3000316960557941 | 0.1318936914191857 |
| GATCCAGATCTAAACTCTTGGGGTGGCTTTGTCTTCTTCTTTTGCCAATTCCACTAATTGTTTGGG | Ins_k3_13_ACT | -0.20491745997058997 | 0.03906361685034905 | 40.00248490581921 |
| GATCCAGATCTAAACTTGGGGTGGCTTTGTCTTCTTCTTTTGCCAATTCCACTAATTGTTTGGG | Ins_k1_11_A;Ins_k1_12_A;Ins_k1 | 0.28012648820831754 | 0.03504434179197522 | 64.97389169717096 |
| GATCCAGATCTAAAGACTTGGGGTGGCTTTGTCTTCTTCTTTTGCCAATTCCACTAATTGTTTGGG | Ins_k3_12_AAG;Ins_k3_13_AGA | 0.5430465681713081 | 0.03165352967955229 | 84.51301980037685 |
| GATCCAGATCTAAAGCCTTGGGGTGGCTTTGTCTTCTTCTTTTGCCAATTCCACTAATTGTTTGGG | Ins_k3_13_AGC | 0.4464399084611774 | 0.03488691336234682 | 76.73047396820445 |
| GATCCAGATCTAAAGCTTGGGGTGGCTTTGTCTTCTTCTTTTGCCAATTCCACTAATTGTTTGGG | Ins_k2_13_AG | 0.36865561556231774 | 0.035127713160709347 | 70.98826987491637 |

| | | | | |
|---|---|---|---|---|
| **GATCCAGATCTAAAGGCTTGGGGTGGCTTTGTCTTCTTCTTTTGCCAATTCCACTAATTGTTTGGG** | Ins_k3_13_AGG | 0.4125413355919172 | 0.03152205400187754 | 74.17301244931431 |
| **GATCCAGATCTAAAGTCTTGGGGTGGCTTTGTCTTCTTCTTTTGCCAATTCCACTAATTGTTTGGG** | Ins_k3_13_AGT | 0.2126006205891643 | 0.03468290815117763 | 60.73132669050087 |
| **GATCCAGATCTAAATACTTGGGGTGGCTTTGTCTTCTTCTTTTGCCAATTCCACTAATTGTTTGGG** | Ins_k3_12_AAT;Ins_k3_13_ATA | -0.08544919699348597 | 0.03814367006071171 | 45.07869939924658 |
| **GATCCAGATCTAAATCCTTGGGGTGGCTTTGTCTTCTTCTTTTGCCAATTCCACTAATTGTTTGGG** | Ins_k3_13_ATC | 0.32149236713334206 | 0.03267498725592506 | 67.7179579117668 |
| **GATCCAGATCTAAATCTTGGGGTGGCTTTGTCTTCTTCTTTTGCCAATTCCACTAATTGTTTGGG** | Ins_k2_13_AT | 0.013836268704437549 | 0.040150041945865814 | 49.78408245422066 |
| **GATCCAGATCTAAATGCTTGGGGTGGCTTTGTCTTCTTCTTTTGCCAATTCCACTAATTGTTTGGG** | Ins_k3_13_ATG | 0.2583787967044252 | 0.03457415327019445 | 63.576113856263724 |
| **GATCCAGATCTAAATTCCACTAATTGTTTGGG** | Del_k31_13-43;Del_k31_14-44 | -3.5849361596466434 | 0.08559671633992755 | 1.3619573257034592 |
| **GATCCAGATCTAAATTCTTGGGGTGGCTTTGTCTTCTTCTTTTGCCAATTCCACTAATTGTTTGGG** | Ins_k3_13_ATT | -0.4583247713455596 | 0.0478414022883763 | 31.04799594476852 |
| **GATCCAGATCTAAATTGGGGTGGCTTTGTCTTCTTCTTTTGCCAATTCCACTAATTGTTTGGG** | C14A | -0.5589818702372247 | 0.041965026146049064 | 28.074934415156825 |
| **GATCCAGATCTAAATTGTTTGGG** | Del_k40_13-52;Del_k40_14-53 | -5.913272839467213 | 0.2407663003389237 | 0.13273324989100252 |
| **GATCCAGATCTAAC** | Del_k49_15-63 | -5.794839530934444 | 0.18168732892258463 | 0.14942203964182352 |
| **GATCCAGATCTAACAAAGTTGGGGTGGCTTTGTCTTCTTCTTTTGCCAATTCCACTAATTGTTTGGG** | Ins_k4_14_AAAG | 0.3511677243255925 | 0.03583617970071784 | 69.75762677205735 |
| **GATCCAGATCTAACAAATTGGGGTGGCTTTGTCTTCTTCTTTTGCCAATTCCACTAATTGTTTGGG** | Ins_k3_14_AAA | -0.18191924081624467 | 0.037434560447323136 | 40.93313140721332 |
| **GATCCAGATCTAACAACTTGGGGTGGCTTTGTCTTCTTCTTTTGCCAATTCCACTAATTGTTTGGG** | Ins_k3_11_AAC;Ins_k3_12_ACA; | -0.6061722782807613 | 0.16248269853335842 | 26.7808413405699 |
| **GATCCAGATCTAACAAGTTGGGGTGGCTTTGTCTTCTTCTTTTGCCAATTCCACTAATTGTTTGGG** | Ins_k3_14_AAG | 0.34869369646522386 | 0.03597446424117185 | 69.58525777072573 |
| **GATCCAGATCTAACAATTCCACTAATTGTTTGGG** | Del_k29_14-42;Del_k29_15-43 | -3.0159766551977847 | 0.07729330095596028 | 2.4057997378276106 |
| **GATCCAGATCTAACAATTGGGGTGGCTTTGTCTTCTTCTTTTGCCAATTCCACTAATTGTTTGGG** | Ins_k2_14_AA | 0.014177365266751063 | 0.044856863241522986 | 49.801066530044 |
| **GATCCAGATCTAACAATTGTTTGGG** | Del_k38_15-52 | -5.727645607587059 | 0.1888290788608216 | 0.1598072986902427 |
| **GATCCAGATCTAACAATTTGGGGTGGCTTTGTCTTCTTCTTTTGCCAATTCCACTAATTGTTTGGG** | Ins_k3_14_AAT | -0.30494582928534 | 0.036030523413884415 | 36.19471832328228 |
| **GATCCAGATCTAACACATTGGGGTGGCTTTGTCTTCTTCTTTTGCCAATTCCACTAATTGTTTGGG** | Ins_k3_14_ACA | -0.10832201319528395 | 0.03847850402000553 | 44.0593250161479 |
| **GATCCAGATCTAACACATTTGGGGTGGCTTTGTCTTCTTCTTTTGCCAATTCCACTAATTGTTTGGG** | Ins_k4_14_ACAT | -0.17218666087109508 | 0.04090070332041986 | 41.33346134255294 |
| **GATCCAGATCTAACACCTTGGGGTGGCTTTGTCTTCTTCTTTTGCCAATTCCACTAATTGTTTGGG** | Ins_k3_13_CAC;Ins_k3_14_ACC | 0.2938979379023351 | 0.0345368991479901 | 65.87486599993085 |
| **GATCCAGATCTAACACGTTGGGGTGGCTTTGTCTTCTTCTTTTGCCAATTCCACTAATTGTTTGGG** | Ins_k3_14_ACG | 0.49094683909244385 | 0.030012494006476556 | 80.22264839077833 |
| **GATCCAGATCTAACACTAATTGTTTGGG** | Del_k35_14-48;Del_k35_15-49 | -5.221839547705332 | 0.19002542672189318 | 0.2650119093917329 |
| **GATCCAGATCTAACACTATTGGGGTGGCTTTGTCTTCTTCTTTTGCCAATTCCACTAATTGTTTGGG** | Ins_k4_14_ACTA | -0.4140349982432765 | 0.042391823627882005 | 32.45401085391282 |
| **GATCCAGATCTAACACTCTTGGGGTGGCTTTGTCTTCTTCTTTTGCCAATTCCACTAATTGTTTGGG** | Ins_k4_13_CACT;Ins_k4_14_ACT | -0.33583751602978895 | 0.24232424389602825 | 35.09369619542567 |
| **GATCCAGATCTAACACTTGGGGTGGCTTTGTCTTCTTCTTTTGCCAATTCCACTAATTGTTTGGG** | Ins_k2_12_AC;Ins_k2_13_CA;Ins | 0.0017959079774577669 | 0.03735068980576301 | 49.18825830987305 |
| **GATCCAGATCTAACACTTTGGGGTGGCTTTGTCTTCTTCTTTTGCCAATTCCACTAATTGTTTGGG** | Ins_k3_14_ACT | -0.2806587375964946 | 0.03699910421600052 | 37.08454467373854 |
| **GATCCAGATCTAACAGATTGGGGTGGCTTTGTCTTCTTCTTTTGCCAATTCCACTAATTGTTTGGG** | Ins_k3_14_AGA | 0.4059813941579711 | 0.03207744948321757 | 73.6880342850088 |
| **GATCCAGATCTAACAGCTTGGGGTGGCTTTGTCTTCTTCTTTTGCCAATTCCACTAATTGTTTGGG** | Ins_k3_13_CAG;Ins_k3_14_AGC | 0.40899008253666636 | 0.03359140561727408 | 73.9100724717725 |
| **GATCCAGATCTAACAGGTTGGGGTGGCTTTGTCTTCTTCTTTTGCCAATTCCACTAATTGTTTGGG** | Ins_k3_14_AGG | -1.1836792848620454 | 0.0565342044144219 | 15.03197726903664 |
| **GATCCAGATCTAACAGTTGGGGTGGCTTTGTCTTCTTCTTTTGCCAATTCCACTAATTGTTTGGG** | Ins_k2_14_AG | 0.3574017724034745 | 0.037905224836762764 | 70.19385750003737 |
| **GATCCAGATCTAACAGTTTGGGGTGGCTTTGTCTTCTTCTTTTGCCAATTCCACTAATTGTTTGGG** | Ins_k3_14_AGT | -0.017004699451384275 | 0.03852138014395911 | 48.27212806252198 |
| **GATCCAGATCTAACATATTGGGGTGGCTTTGTCTTCTTCTTTTGCCAATTCCACTAATTGTTTGGG** | Ins_k3_14_ATA | -0.4995024341560028 | 0.04520616816578014 | 29.795476915861418 |
| **GATCCAGATCTAACATATTTGGGGTGGCTTTGTCTTCTTCTTTTGCCAATTCCACTAATTGTTTGGG** | Ins_k4_14_ATAT | -0.4538360515558385 | 0.04254420740269385 | 31.187674954035508 |
| **GATCCAGATCTAACATCTTGGGGTGGCTTTGTCTTCTTCTTTTGCCAATTCCACTAATTGTTTGGG** | Ins_k3_13_CAT;Ins_k3_14_ATC | 0.339889814215227 | 0.03271530810217646 | 68.97532617750072 |
| **GATCCAGATCTAACATGGGGTGGCTTTGTCTTCTTCTTTTGCCAATTCCACTAATTGTTTGGG** | T15A | -0.14110125431545623 | 0.03171029166996075 | 42.63850765286936 |
| **GATCCAGATCTAACATGGTTGGGGTGGCTTTGTCTTCTTCTTTTGCCAATTCCACTAATTGTTTGGG** | Ins_k4_14_ATGG | -0.013490489359452296 | 0.04110026203152395 | 48.44206488407634 |
| **GATCCAGATCTAACATGTTGGGGTGGCTTTGTCTTCTTCTTTTGCCAATTCCACTAATTGTTTGGG** | Ins_k3_14_ATG | 0.20154773363400036 | 0.03265890408912054 | 60.063766233191465 |
| **GATCCAGATCTAACATTCCACTAATTGTTTGGG** | Del_k30_15-44 | -3.4429666824353053 | 0.08338246953230483 | 1.5697122971817967 |
| **GATCCAGATCTAACATTGGGGTGGCTTTGTCTTCTTCTTTTGCCAATTCCACTAATTGTTTGGG** | Ins_k1_14_A | 0.01738861347859888 | 0.05097960034877407 | 49.961247168130456 |
| **GATCCAGATCTAACATTGTTTGGG** | Del_k39_15-53 | -5.708022331198471 | 0.2142415241388485 | 0.1629742124697951 |
| **GATCCAGATCTAACATTTGGGGTGGCTTTGTCTTCTTCTTTTGCCAATTCCACTAATTGTTTGGG** | Ins_k2_14_AT | -0.06796479157402868 | 0.044509154295633506 | 45.87380436865331 |
| **GATCCAGATCTAACATTTTGGGGTGGCTTTGTCTTCTTCTTTTGCCAATTCCACTAATTGTTTGGG** | Ins_k3_14_ATT | -0.5188303829411983 | 0.051234165299639735 | 29.225121123296947 |
| **GATCCAGATCTAACCAAGTTGGGGTGGCTTTGTCTTCTTCTTTTGCCAATTCCACTAATTGTTTGGG** | Ins_k4_14_CAAG | 0.6084073237166041 | 0.029103976674750093 | 90.22137374861141 |
| **GATCCAGATCTAACCAATTCCACTAATTGTTTGGG** | Del_k28_14-41;Del_k28_15-42 | -2.8768804467606923 | 0.05291543639591047 | 2.7648284455483334 |
| **GATCCAGATCTAACCAATTGGGGTGGCTTTGTCTTCTTCTTTTGCCAATTCCACTAATTGTTTGGG** | Ins_k3_14_CAA | 0.13349452174871712 | 0.03700775885008357 | 56.11221676956836 |
| **GATCCAGATCTAACCACTAATTGTTTGGG** | Del_k34_14-47;Del_k34_15-48 | -3.402464853389539 | 0.3988484888710526 | 1.6345935520241492 |
| **GATCCAGATCTAACCACTTGGGGTGGCTTTGTCTTCTTCTTTTGCCAATTCCACTAATTGTTTGGG** | Ins_k3_12_ACC;Ins_k3_13_CCA; | 0.4064710104971023 | 0.03196909887112382 | 73.72412198443593 |
| **GATCCAGATCTAACCAGTTGGGGTGGCTTTGTCTTCTTCTTTTGCCAATTCCACTAATTGTTTGGG** | Ins_k3_14_CAG | 0.5543074348283619 | 0.03227349491891335 | 85.47008824362484 |
| **GATCCAGATCTAACCAGTTTGGGGTGGCTTTGTCTTCTTCTTTTGCCAATTCCACTAATTGTTTGGG** | Ins_k4_14_CAGT | 0.6855126976789001 | 0.03174881479696669 | 97.45314832179513 |
| **GATCCAGATCTAACCATATTGGGGTGGCTTTGTCTTCTTCTTTTGCCAATTCCACTAATTGTTTGGG** | Ins_k4_14_CATA | 0.2824106051185802 | 0.03633682789433307 | 65.12246928161213 |
| **GATCCAGATCTAACCATTGGGGTGGCTTTGTCTTCTTCTTTTGCCAATTCCACTAATTGTTTGGG** | Ins_k2_14_CA | 0.37312552878592997 | 0.033715778041574204 | 71.30629151621238 |
| **GATCCAGATCTAACCATTTGGGGTGGCTTTGTCTTCTTCTTTTGCCAATTCCACTAATTGTTTGGG** | Ins_k3_14_CAT | 0.3288006995785221 | 0.034243650568873636 | 68.21467614078993 |
| **GATCCAGATCTAACCCAATTCCACTAATTGTTTGGG** | Del_k27_15-41 | -3.0327694412916353 | 0.0656392164691603 | 2.3657369815563776 |
| **GATCCAGATCTAACCCACTAATTGTTTGGG** | Del_k33_15-47 | -4.274406775947776 | 0.10434338211333205 | 0.6834869327469258 |
| **GATCCAGATCTAACCCATTGGGGTGGCTTTGTCTTCTTCTTTTGCCAATTCCACTAATTGTTTGGG** | Ins_k3_14_CCA | 0.4091492464012523 | 0.03345826028305644 | 73.92183722077556 |
| **GATCCAGATCTAACCCCTTGGGGTGGCTTTGTCTTCTTCTTTTGCCAATTCCACTAATTGTTTGGG** | Ins_k3_13_CCC;Ins_k3_14_CCC | 0.25119052812065423 | 0.03335434662287719 | 63.120750272986186 |
| **GATCCAGATCTAACCCGTTGGGGTGGCTTTGTCTTCTTCTTTTGCCAATTCCACTAATTGTTTGGG** | Ins_k3_14_CCG | 0.6216325926799993 | 0.02923490424343484 | 91.42250078962186 |
| **GATCCAGATCTAACCCGTTTGGGGTGGCTTTGTCTTCTTCTTTTGCCAATTCCACTAATTGTTTGGG** | Ins_k4_14_CCGT | 0.7028881488602889 | 0.03291615471776972 | 99.16123717684043 |
| **GATCCAGATCTAACCCTTGGGGTGGCTTTGTCTTCTTCTTTTGCCAATTCCACTAATTGTTTGGG** | Ins_k2_13_CC;Ins_k2_14_CC | 0.5368928895204503 | 0.036802987941585576 | 83.99455071688111 |
| **GATCCAGATCTAACCCTTTGGGGTGGCTTTGTCTTCTTCTTTTGCCAATTCCACTAATTGTTTGGG** | Ins_k3_14_CCT | 0.1473473625982692 | 0.03703051257023925 | 56.89493932662244 |
| **GATCCAGATCTAACCGATTGGGGTGGCTTTGTCTTCTTCTTTTGCCAATTCCACTAATTGTTTGGG** | Ins_k3_14_CGA | 0.6777813364429468 | 0.02868747969684537 | 96.70260791659555 |
| **GATCCAGATCTAACCGCTTGGGGTGGCTTTGTCTTCTTCTTTTGCCAATTCCACTAATTGTTTGGG** | Ins_k3_13_CCG;Ins_k3_14_CGC | 1.4782710882472387 | 0.16571149013963557 | 100 |
| **GATCCAGATCTAACCGGATTGGGGTGGCTTTGTCTTCTTCTTTTGCCAATTCCACTAATTGTTTGGG** | Ins_k4_14_CGGA | 1.1315265619571466 | 0.029101451278266348 | 100 |
| **GATCCAGATCTAACCGGTTGGGGTGGCTTTGTCTTCTTCTTTTGCCAATTCCACTAATTGTTTGGG** | Ins_k3_14_CGG | 0.5935508646534498 | 0.0342728166187618 | 88.8909110560871 |
| **GATCCAGATCTAACCGTTGGGGTGGCTTTGTCTTCTTCTTTTGCCAATTCCACTAATTGTTTGGG** | Ins_k2_14_CG | 0.6343475280049278 | 0.03074805647987035 | 92.59235351451602 |
| **GATCCAGATCTAACCGTTTGGGGTGGCTTTGTCTTCTTCTTTTGCCAATTCCACTAATTGTTTGGG** | Ins_k3_14_CGT | 0.45582461897870297 | 0.03230170483772346 | 77.45395678273651 |
| **GATCCAGATCTAACCTAATTGTTTGGG** | Del_k36_15-50 | -5.249605673063253 | 0.17403878923194202 | 0.2577547730128879 |
| **GATCCAGATCTAACCTATTGGGGTGGCTTTGTCTTCTTCTTTTGCCAATTCCACTAATTGTTTGGG** | Ins_k3_14_CTA | -0.03179095267669385 | 0.036848159095094805 | 47.563615187319066 |
| **GATCCAGATCTAACCTCCTTGGGGTGGCTTTGTCTTCTTCTTTTGCCAATTCCACTAATTGTTTGGG** | Ins_k4_13_CCTC;Ins_k4_14_CT | 0.5744034724348435 | 0.031232416811321873 | 87.2050731227039 |
| **GATCCAGATCTAACCTCTTGGGGTGGCTTTGTCTTCTTCTTTTGCCAATTCCACTAATTGTTTGGG** | Ins_k3_13_CCT;Ins_k3_14_CTC | 0.13792946387634775 | 0.03410666814740604 | 56.361623847780656 |
| **GATCCAGATCTAACCTGGGGTGGCTTTGTCTTCTTCTTTTGCCAATTCCACTAATTGTTTGGG** | T15C | 0.006724352167455372 | 0.027400990216917966 | 49.43127825898074 |
| **GATCCAGATCTAACCTGTTGGGGTGGCTTTGTCTTCTTCTTTTGCCAATTCCACTAATTGTTTGGG** | Ins_k3_14_CTG | 0.48816268634186555 | 0.03588043162065745 | 79.99960691844241 |
| **GATCCAGATCTAACCTTCTTCTTTTGCCAATTCCACTAATTGTTTGGG** | Del_k15_15-29 | -1.4916264175493086 | 0.0524453456770952 | 11.047814407799722 |

| | | | | |
|---|---|---|---|---|
| GATCCAGATCTAACCTTCTTTTGCCAATTCCACTAATTGTTTGGG | Del_k18_15-32 | -1.4707752339589237 | 0.04947052437048746 | 11.280592833402055 |
| GATCCAGATCTAACCTTGGGGTGGCTTTGTCTTCTTCTTTTGCCAATTCCACTAATTGTTTGGG | Ins_k1_13_C;Ins_k1_14_C | 0.3279600674010491 | 0.040481718290173316 | 68.15735678467311 |
| GATCCAGATCTAACCTTTGGGGTGGCTTTGTCTTCTTCTTTTGCCAATTCCACTAATTGTTTGGG | Ins_k2_14_CT | 0.23541975582508856 | 0.039619054024222236 | 62.13309580067497 |
| GATCCAGATCTAACCTTTGTCTTCTTCTTTTGCCAATTCCACTAATTGTTTGGG | Del_k9_15-23 | -2.672230572104126 | 0.09847557549382567 | 3.3927081361559064 |
| GATCCAGATCTAACCTTTTGCCAATTCCACTAATTGTTTGGG | Del_k21_15-35 | -2.052104528524877 | 0.05822456636746403 | 6.307595273883 |
| GATCCAGATCTAACCTTTTGGGGTGGCTTTGTCTTCTTCTTTTGCCAATTCCACTAATTGTTTGGG | Ins_k3_14_CTT | -0.16665930452344013 | 0.03627791780089898 | 41.56255868051265 |
| GATCCAGATCTAACG | Del_k48_15-62 | -6.5358373815938515 | 0.32119652670987814 | 0.07122023177440989 |
| GATCCAGATCTAACGAATTGGGGTGGCTTTGTCTTCTTCTTTTGCCAATTCCACTAATTGTTTGGG | Ins_k3_14_GAA | 0.5056721433275715 | 0.0313187034213593 | 81.41269166605366 |
| GATCCAGATCTAACGACTTGGGGTGGCTTTGTCTTCTTCTTTTGCCAATTCCACTAATTGTTTGGG | Ins_k3_12_ACG;Ins_k3_13_CGA | 0.6196813130493151 | 0.03119150159858866 | 91.24428385812563 |
| GATCCAGATCTAACGAGGTTGGGGTGGCTTTGTCTTCTTCTTTTGCCAATTCCACTAATTGTTTGGG | Ins_k4_14_GAGG | 0.4407729927300683 | 0.03586358731677415 | 76.29687857287249 |
| GATCCAGATCTAACGAGTTGGGGTGGCTTTGTCTTCTTCTTTTGCCAATTCCACTAATTGTTTGGG | Ins_k3_14_GAG | 0.6251081246344372 | 0.030681699410160194 | 91.74079541364063 |
| GATCCAGATCTAACGATTGGGGTGGCTTTGTCTTCTTCTTTTGCCAATTCCACTAATTGTTTGGG | Ins_k2_14_GA | 0.6628726017628401 | 0.0334600897661956 | 95.27158824460626 |
| GATCCAGATCTAACGATTTGGGGTGGCTTTGTCTTCTTCTTTTGCCAATTCCACTAATTGTTTGGG | Ins_k3_14_GAT | 0.41617927245741315 | 0.03336390885342603 | 74.443340605972 |
| GATCCAGATCTAACGCATTGGGGTGGCTTTGTCTTCTTCTTTTGCCAATTCCACTAATTGTTTGGG | Ins_k3_14_GCA | 0.40738478036896786 | 0.03255311178511302 | 73.79151965422547 |
| GATCCAGATCTAACGCCAATTCCACTAATTGTTTGGG | Del_k26_15-40 | -0.9559180012368682 | 0.029749423685668058 | 18.87693928745625 |
| GATCCAGATCTAACGCCTTGGGGTGGCTTTGTCTTCTTCTTTTGCCAATTCCACTAATTGTTTGGG | Ins_k3_13_CGC;Ins_k3_14_GCC | 0.725737621213316 | 0.027898741378882812 | 100 |
| GATCCAGATCTAACGCGTTGGGGTGGCTTTGTCTTCTTCTTTTGCCAATTCCACTAATTGTTTGGG | Ins_k3_14_GCG | 0.6864643569799138 | 0.03268205133937863 | 97.54593466029694 |
| GATCCAGATCTAACGCTTGGGGTGGCTTTGTCTTCTTCTTTTGCCAATTCCACTAATTGTTTGGG | Ins_k2_13_CG;Ins_k2_14_GC | 0.6478553985240543 | 0.03257608043132392 | 93.85156451994573 |
| GATCCAGATCTAACGCTTTGGGGTGGCTTTGTCTTCTTCTTTTGCCAATTCCACTAATTGTTTGGG | Ins_k3_14_GCT | 0.40329047377122906 | 0.03423947443933857 | 73.49001220161362 |
| GATCCAGATCTAACGCTTTGTCTTCTTCTTTTGCCAATTCCACTAATTGTTTGGG | Del_k8_15-22 | -1.3765551999341228 | 0.048565868142706393 | 12.395132247660662 |
| GATCCAGATCTAACGG | Del_k47_15-61 | -6.5908435344431435 | 0.3290990375007953 | 0.06740847685280871 |
| GATCCAGATCTAACGGATTGGGGTGGCTTTGTCTTCTTCTTTTGCCAATTCCACTAATTGTTTGGG | Ins_k3_14_GGA | 0.6157928213210693 | 0.028465781064252096 | 90.89017014540899 |
| GATCCAGATCTAACGGCTTGGGGTGGCTTTGTCTTCTTCTTTTGCCAATTCCACTAATTGTTTGGG | Ins_k3_13_CGG;Ins_k3_14_GGC | 0.7356748327956919 | 0.03217060503210766 | 100 |
| GATCCAGATCTAACGGCTTTGTCTTCTTCTTTTGCCAATTCCACTAATTGTTTGGG | Del_k7_15-21 | 0.3022825785442681 | 0.0301791665848431 | 66.42952513797545 |
| GATCCAGATCTAACGGG | Del_k46_15-60 | -6.277771993121939 | 0.20645977755230407 | 0.09218913856685187 |
| GATCCAGATCTAACGGGGTGGCTTTGTCTTCTTCTTTTGCCAATTCCACTAATTGTTTGGG | Del_k2_15-16 | 0.5331323242364301 | 0.0308443790072998 | 83.6792769008383 |
| GATCCAGATCTAACGGGTGGCTTTGTCTTCTTCTTTTGCCAATTCCACTAATTGTTTGGG | Del_k3_15-17 | 0.561332343318298 | 0.027066735996199454 | 86.07262168542489 |
| GATCCAGATCTAACGGGTTGGGGTGGCTTTGTCTTCTTCTTTTGCCAATTCCACTAATTGTTTGGG | Ins_k3_14_GGG | -1.946309819345728 | 0.09682183046939102 | 7.011482872820775 |
| GATCCAGATCTAACGGTGGCTTTGTCTTCTTCTTTTGCCAATTCCACTAATTGTTTGGG | Del_k4_15-18 | 0.4989414476254963 | 0.02726410222905773 | 80.86656757143079 |
| GATCCAGATCTAACGGTTGGGGTGGCTTTGTCTTCTTCTTTTGCCAATTCCACTAATTGTTTGGG | Ins_k2_14_GG | 0.3828122583401318 | 0.036727776431464874 | 72.00037253793148 |
| GATCCAGATCTAACGGTTTGGGGTGGCTTTGTCTTCTTCTTTTGCCAATTCCACTAATTGTTTGGG | Ins_k3_14_GGT | 0.08558619265798195 | 0.03578514310959693 | 53.487352680411846 |
| GATCCAGATCTAACGTATTGGGGTGGCTTTGTCTTCTTCTTTTGCCAATTCCACTAATTGTTTGGG | Ins_k3_14_GTA | 0.2178183436536827 | 0.03137796537956797 | 61.049034068280505 |
| GATCCAGATCTAACGTCTTCTTCTTTTGCCAATTCCACTAATTGTTTGGG | Del_k13_15-27 | -0.05039765154362408 | 0.02720935660233946 | 46.68679597479384 |
| GATCCAGATCTAACGTCTTGGGGTGGCTTTGTCTTCTTCTTTTGCCAATTCCACTAATTGTTTGGG | Ins_k3_13_CGT;Ins_k3_14_GTC | 0.6264072404982737 | 0.031037986007357053 | 91.8600547854215 |
| GATCCAGATCTAACGTGGCTTTGTCTTCTTCTTTTGCCAATTCCACTAATTGTTTGGG | Del_k5_15-19 | 0.2734684201630406 | 0.026438750405336945 | 64.54272805629903 |
| GATCCAGATCTAACGTGGGGTGGCTTTGTCTTCTTCTTTTGCCAATTCCACTAATTGTTTGGG | T15G | -0.0060952945005163794 | 0.0354847567745124 | 48.80163128637854 |
| GATCCAGATCTAACGTGTTGGGGTGGCTTTGTCTTCTTCTTTTGCCAATTCCACTAATTGTTTGGG | Ins_k3_14_GTG | 0.42484232101171004 | 0.03196367292293224 | 75.09104839140112 |
| GATCCAGATCTAACGTTGGGGTGGCTTTGTCTTCTTCTTTTGCCAATTCCACTAATTGTTTGGG | Ins_k1_14_G | 0.6346739305325011 | 0.03621283131050161 | 92.62258082560413 |
| GATCCAGATCTAACGTTTGGG | Del_k42_15-56 | -6.511598929163329 | 0.27109774443437323 | 0.07296759106960625 |
| GATCCAGATCTAACGTTTGGGGTGGCTTTGTCTTCTTCTTTTGCCAATTCCACTAATTGTTTGGG | Ins_k2_14_GT | 0.523774180036841 | 0.03603918642311975 | 82.89984685864538 |
| GATCCAGATCTAACGTTTTGGGGTGGCTTTGTCTTCTTCTTTTGCCAATTCCACTAATTGTTTGGG | Ins_k3_14_GTT | 0.05323509679777605 | 0.0370995511012373 | 51.78466855414112 |
| GATCCAGATCTAACT | Del_k48_16-63 | -3.9251893554176633 | 0.06487573852582043 | 0.969155388191124 |
| GATCCAGATCTAACTAAATGGGGTGGCTTTGTCTTCTTCTTTTGCCAATTCCACTAATTGTTTGGG | Ins_k3_15_AAA | -0.4235253466671698 | 0.03992297506535715 | 32.147467883726385 |
| GATCCAGATCTAACTAACTGGGGTGGCTTTGTCTTCTTCTTTTGCCAATTCCACTAATTGTTTGGG | Ins_k3_15_AAC | -0.45761638669871074 | 0.04233031130909719 | 31.069997660329918 |
| GATCCAGATCTAACTAAGTGGGGTGGCTTTGTCTTCTTCTTTTGCCAATTCCACTAATTGTTTGGG | Ins_k3_15_AAG | -1.8397711103279624 | 0.07702625779605075 | 7.799720693061251 |
| GATCCAGATCTAACTAATGGGGTGGCTTTGTCTTCTTCTTTTGCCAATTCCACTAATTGTTTGGG | Ins_k2_15_AA | -0.35989070796689626 | 0.04557896406398428 | 34.2596517124727 |
| GATCCAGATCTAACTAATGTGGGGTGGCTTTGTCTTCTTCTTTTGCCAATTCCACTAATTGTTTGGG | Ins_k4_15_AATG | -1.5839683295293827 | 0.06867365388235813 | 10.073323633671025 |
| GATCCAGATCTAACTAATTCCACTAATTGTTTGGG | Del_k28_16-43 | -3.6319402136321006 | 0.08061836597188567 | 1.2994210526648267 |
| GATCCAGATCTAACTAATTGGGGTGGCTTTGTCTTCTTCTTTTGCCAATTCCACTAATTGTTTGGG | Ins_k3_14_TAA;Ins_k3_15_AAT | -0.7562167630219303 | 0.04736153099228558 | 23.04945838520556 |
| GATCCAGATCTAACTAATTGTTTGGG | Del_k37_13-49;Del_k37_14-50;D | -6.325686219961868 | 0.2241420811189508 | 0.0878761198588868 |
| GATCCAGATCTAACTACATGGGGTGGCTTTGTCTTCTTCTTTTGCCAATTCCACTAATTGTTTGGG | Ins_k3_15_ACA | 0.22433089008294682 | 0.033435828385066424 | 61.44791619647844 |
| GATCCAGATCTAACTACCTGGGGTGGCTTTGTCTTCTTCTTTTGCCAATTCCACTAATTGTTTGGG | Ins_k3_15_ACC | 0.5517076131492729 | 0.031027012923278214 | 85.24816985441191 |
| GATCCAGATCTAACTACGATGGGGTGGCTTTGTCTTCTTCTTTTGCCAATTCCACTAATTGTTTGGG | Ins_k4_15_ACGA | 0.8081249478851413 | 0.02856068928228479 | 100 |
| GATCCAGATCTAACTACGGTGGGGTGGCTTTGTCTTCTTCTTTTGCCAATTCCACTAATTGTTTGGG | Ins_k4_15_ACGG | 0.35395194441858135 | 0.03629902103721185 | 69.95211798568839 |
| GATCCAGATCTAACTACGTGGGGTGGCTTTGTCTTCTTCTTTTGCCAATTCCACTAATTGTTTGGG | Ins_k3_15_ACG | 0.3639224071126984 | 0.031092400601573787 | 70.65306152742507 |
| GATCCAGATCTAACTACTAATTGTTTGGG | Del_k34_16-49 | -3.8350917984450716 | 0.07843304183442874 | 1.0605283595626223 |
| GATCCAGATCTAACTACTGGGGTGGCTTTGTCTTCTTCTTTTGCCAATTCCACTAATTGTTTGGG | Ins_k2_15_AC | 0.5589063510056802 | 0.03345977345615328 | 85.86406324975816 |
| GATCCAGATCTAACTACTTGGGGTGGCTTTGTCTTCTTCTTTTGCCAATTCCACTAATTGTTTGGG | Ins_k3_12_ACT;Ins_k3_13_CTA;I | 0.28058678919158775 | 0.0343083912345189 | 65.00380612769875 |
| GATCCAGATCTAACTAGAGTGGGGTGGCTTTGTCTTCTTCTTTTGCCAATTCCACTAATTGTTTGGG | Ins_k4_15_AGAG | -0.8453763998916746 | 0.054491557502521396 | 21.0833290316861 |
| GATCCAGATCTAACTAGATGGGGTGGCTTTGTCTTCTTCTTTTGCCAATTCCACTAATTGTTTGGG | Ins_k3_15_AGA | 0.02861942104969306 | 0.03522353291348667 | 50.5255149817429 |
| GATCCAGATCTAACTAGCTGGGGTGGCTTTGTCTTCTTCTTTTGCCAATTCCACTAATTGTTTGGG | Ins_k3_15_AGC | 0.0011323991042476722 | 0.03533011921560081 | 49.15563228905081 |
| GATCCAGATCTAACTAGGGGTGGCTTTGTCTTCTTCTTTTGCCAATTCCACTAATTGTTTGGG | T16A | -0.7505379088947864 | 0.04054116506551539 | 23.180725267048857 |
| GATCCAGATCTAACTAGGTGGGGTGGCTTTGTCTTCTTCTTTTGCCAATTCCACTAATTGTTTGGG | Ins_k3_15_AGG | -2.7074940040541526 | 0.13189971291424055 | 3.275154458075796 |
| GATCCAGATCTAACTAGTGGGGTGGCTTTGTCTTCTTCTTTTGCCAATTCCACTAATTGTTTGGG | Ins_k2_15_AG | -2.154807164631647 | 0.100797435557134 | 5.691944161419326 |
| GATCCAGATCTAACTAGTTGGGGTGGCTTTGTCTTCTTCTTTTGCCAATTCCACTAATTGTTTGGG | Ins_k3_14_TAG;Ins_k3_15_AGT | -0.8417691352166701 | 0.04783542096163825 | 21.159519516738108 |
| GATCCAGATCTAACTATATGGGGTGGCTTTGTCTTCTTCTTTTGCCAATTCCACTAATTGTTTGGG | Ins_k3_15_ATA | -0.6381529189732709 | 0.03844742875703316 | 25.937923247331923 |
| GATCCAGATCTAACTATCTGGGGTGGCTTTGTCTTCTTCTTTTGCCAATTCCACTAATTGTTTGGG | Ins_k3_15_ATC | -0.6077367181199753 | 0.046728004861801654 | 26.738977081045224 |
| GATCCAGATCTAACTATGGGGTGGCTTTGTCTTCTTCTTTTGCCAATTCCACTAATTGTTTGGG | Ins_k1_15_A | 0.010534242100446067 | 0.04053290466917062 | 49.619965198393345 |
| GATCCAGATCTAACTATGTGGGGTGGCTTTGTCTTCTTCTTTTGCCAATTCCACTAATTGTTTGGG | Ins_k3_15_ATG | -1.024596393946286 | 0.05028462334335915 | 17.624018176594856 |
| GATCCAGATCTAACTATTCCACTAATTGTTTGGG | Del_k29_16-44 | -3.328267785641711 | 0.07498293939652736 | 1.760488362333835 |
| GATCCAGATCTAACTATTGGGGTGGCTTTGTCTTCTTCTTTTGCCAATTCCACTAATTGTTTGGG | Ins_k2_14_TA;Ins_k2_15_AT | -0.32157899450872307 | 0.043658076111151974 | 35.59766480323456 |
| GATCCAGATCTAACTATTGTTTGGG | Del_k38_16-53 | -5.199067186669726 | 0.13182458548669929 | 0.27111609584970636 |
| GATCCAGATCTAACTATTTGGGGTGGCTTTGTCTTCTTCTTTTGCCAATTCCACTAATTGTTTGGG | Ins_k3_14_TAT;Ins_k3_15_ATT | -0.5871603491088214 | 0.04805286714687656 | 27.294867632070233 |
| GATCCAGATCTAACTCAATGGGGTGGCTTTGTCTTCTTCTTTTGCCAATTCCACTAATTGTTTGGG | Ins_k3_15_CAA | 0.08066159556389962 | 0.03371258779086512 | 53.22459653393187 |
| GATCCAGATCTAACTCAATTCCACTAATTGTTTGGG | Del_k27_16-42 | -2.7283526541644116 | 0.06422021721693484 | 3.207546711574504 |

| | | | | |
|---|---|---|---|---|
| **GATCCAGATCTAACTCACTAATTGTTTGGG** | Del_k33_16-48 | -4.357270314857267 | 0.11512637138623372 | 0.6291338290945098 |
| **GATCCAGATCTAACTCACTGGGGTGGCTTTGTCTTCTTCTTTTGCCAATTCCACTAATTGTTTGGG** | Ins_k3_15_CAC | 0.19540539453848382 | 0.0345616358114529 | 59.6959649501102 |
| **GATCCAGATCTAACTCAGTGGGGTGGCTTTGTCTTCTTCTTTTGCCAATTCCACTAATTGTTTGGG** | Ins_k3_15_CAG | -0.528515022633574 | 0.042977464778999004 | 28.94345248640664 |
| **GATCCAGATCTAACTCATATGGGGTGGCTTTGTCTTCTTCTTTTGCCAATTCCACTAATTGTTTGGG** | Ins_k4_15_CATA | 0.23386767802088526 | 0.03587274265130198 | 62.03673520069447 |
| **GATCCAGATCTAACTCATGGGGTGGCTTTGTCTTCTTCTTTTGCCAATTCCACTAATTGTTTGGG** | Ins_k2_15_CA | 0.3298472267049447 | 0.033388817909936294 | 68.28610201783141 |
| **GATCCAGATCTAACTCATTGGGGTGGCTTTGTCTTCTTCTTTTGCCAATTCCACTAATTGTTTGGG** | Ins_k3_14_TCA;Ins_k3_15_CAT | 0.05476961108778999 | 0.0370868620416243 | 51.86419386880188 |
| **GATCCAGATCTAACTCCAATTCCACTAATTGTTTGGG** | Del_k26_16-41 | -1.2781563966238167 | 0.03498013638230656 | 13.676822860435001 |
| **GATCCAGATCTAACTCCACTAATTGTTTGGG** | Del_k32_15-46;Del_k32_16-47 | -2.715001512566193 | 0.05130995889357057 | 3.2506582758077642 |
| **GATCCAGATCTAACTCCATGGGGTGGCTTTGTCTTCTTCTTTTGCCAATTCCACTAATTGTTTGGG** | Ins_k3_15_CCA | 0.4930974424535419 | 0.03018531657881033 | 80.39536113978525 |
| **GATCCAGATCTAACTCCCTGGGGTGGCTTTGTCTTCTTCTTTTGCCAATTCCACTAATTGTTTGGG** | Ins_k3_15_CCC | 0.339038963208246 | 0.031390765960578056 | 68.91666341194139 |
| **GATCCAGATCTAACTCCGTGGGGTGGCTTTGTCTTCTTCTTTTGCCAATTCCACTAATTGTTTGGG** | Ins_k3_15_CCG | 0.6553081505088942 | 0.03224031742400326 | 94.5536298635487 |
| **GATCCAGATCTAACTCCTGGGGTGGCTTTGTCTTCTTCTTTTGCCAATTCCACTAATTGTTTGGG** | Ins_k2_15_CC | 0.6165714139800331 | 0.0335121102015681 | 90.96096412091468 |
| **GATCCAGATCTAACTCCTTGGGGTGGCTTTGTCTTCTTCTTTTGCCAATTCCACTAATTGTTTGGG** | Ins_k3_13_CTC;Ins_k3_14_TCC; | 0.3430624094282553 | 0.03373809241351262 | 69.19450446529952 |
| **GATCCAGATCTAACTCGAGTGGGGTGGCTTTGTCTTCTTCTTTTGCCAATTCCACTAATTGTTTGGG** | Ins_k4_15_CGAG | 0.45156087307801607 | 0.03453695150366181 | 77.12441583069528 |
| **GATCCAGATCTAACTCGATGGGGTGGCTTTGTCTTCTTCTTTTGCCAATTCCACTAATTGTTTGGG** | Ins_k3_15_CGA | 0.7114858984306496 | 0.028589593003380206 | 100 |
| **GATCCAGATCTAACTCGCTGGGGTGGCTTTGTCTTCTTCTTTTGCCAATTCCACTAATTGTTTGGG** | Ins_k3_15_CGC | 0.6845768062515372 | 0.02925079143669153 | 97.36198542165366 |
| **GATCCAGATCTAACTCGGGGTGGCTTTGTCTTCTTCTTTTGCCAATTCCACTAATTGTTTGGG** | T16C | 0.07247171614114245 | 0.02783555296621515 | 52.790473639605395 |
| **GATCCAGATCTAACTCGGTGGGGTGGCTTTGTCTTCTTCTTTTGCCAATTCCACTAATTGTTTGGG** | Ins_k3_15_CGG | -0.06458899965877729 | 0.042532806413726504 | 46.02892646921543 |
| **GATCCAGATCTAACTCGTGGGGTGGCTTTGTCTTCTTCTTTTGCCAATTCCACTAATTGTTTGGG** | Ins_k2_15_CG | 0.41686163365864015 | 0.03394102947780946 | 74.49415518826926 |
| **GATCCAGATCTAACTCGTTGGGGTGGCTTTGTCTTCTTCTTTTGCCAATTCCACTAATTGTTTGGG** | Ins_k3_14_TCG;Ins_k3_15_CGT | 0.45990373809674545 | 0.02944889423154369 | 77.77054596212133 |
| **GATCCAGATCTAACTCTAATTGTTTGGG** | Del_k35_16-50 | -4.735300985664551 | 0.13559115995087476 | 0.43108846666221934 |
| **GATCCAGATCTAACTCTATGGGGTGGCTTTGTCTTCTTCTTTTGCCAATTCCACTAATTGTTTGGG** | Ins_k3_15_CTA | -0.30209042839990863 | 0.043945532008676985 | 36.298216448027986 |
| **GATCCAGATCTAACTCTCTGGGGTGGCTTTGTCTTCTTCTTTTGCCAATTCCACTAATTGTTTGGG** | Ins_k3_15_CTC | 1.8828423670458339e-4 | 0.04072047190690887 | 49.10924562639774 |
| **GATCCAGATCTAACTCTGGGGTGGCTTTGTCTTCTTCTTTTGCCAATTCCACTAATTGTTTGGG** | Ins_k1_15_C | 0.10148125932848795 | 0.03461611022334155 | 54.34433053425369 |
| **GATCCAGATCTAACTCTGTGGGGTGGCTTTGTCTTCTTCTTTTGCCAATTCCACTAATTGTTTGGG** | Ins_k3_15_CTG | -0.4947536491549939 | 0.04162218602442691 | 29.937305720456155 |
| **GATCCAGATCTAACTCTTCTTCTTTTGCCAATTCCACTAATTGTTTGGG** | Del_k14_15-28;Del_k14_16-29 | -1.5078917849904059 | 0.043939709945905 | 10.869571172577416 |
| **GATCCAGATCTAACTCTTCTTTTGCCAATTCCACTAATTGTTTGGG** | Del_k17_15-31;Del_k17_16-32 | -1.1309453324733039 | 0.036491493714999455 | 15.845946120270781 |
| **GATCCAGATCTAACTCTTGGGGTGGCTTTGTCTTCTTCTTTTGCCAATTCCACTAATTGTTTGGG** | Ins_k2_13_CT;Ins_k2_14_TC;Ins_ | -0.037189809011402786 | 0.03787061524189841 | 47.307518000176195 |
| **GATCCAGATCTAACTCTTTGGGGTGGCTTTGTCTTCTTCTTTTGCCAATTCCACTAATTGTTTGGG** | Ins_k3_14_TCT;Ins_k3_15_CTT | -0.4410018538725953 | 0.04727490764577536 | 31.590523327872393 |
| **GATCCAGATCTAACTCTTTGTCTTCTTCTTTTGCCAATTCCACTAATTGTTTGGG** | Del_k8_16-23 | -3.1544344324495146 | 0.12985280229969307 | 2.0947298650239854 |
| **GATCCAGATCTAACTCTTTTGCCAATTCCACTAATTGTTTGGG** | Del_k20_15-34;Del_k20_16-35 | -2.1583243845263445 | 0.0582343211761494 | 5.671959507987829 |
| **GATCCAGATCTAACTG** | Del_k47_16-62 | -6.828240984612011 | 0.3572479552414217 | 0.05316356690716665 |
| **GATCCAGATCTAACTGAATGGGGTGGCTTTGTCTTCTTCTTTTGCCAATTCCACTAATTGTTTGGG** | Ins_k3_15_GAA | 0.590527113465259 | 0.02999534294179586 | 88.62253301685267 |
| **GATCCAGATCTAACTGACTGGGGTGGCTTTGTCTTCTTCTTTTGCCAATTCCACTAATTGTTTGGG** | Ins_k3_15_GAC | 0.7087237336301928 | 0.02755257750602659 | 99.7415926922241 |
| **GATCCAGATCTAACTGAGTGGGGTGGCTTTGTCTTCTTCTTTTGCCAATTCCACTAATTGTTTGGG** | Ins_k3_15_GAG | 0.0652414562279322 | 0.0394217187564056 | 52.41016132946146 |
| **GATCCAGATCTAACTGATGGGGTGGCTTTGTCTTCTTCTTTTGCCAATTCCACTAATTGTTTGGG** | Ins_k2_15_GA | 0.679215760163955 | 0.03068164644078994 | 96.84141996512074 |
| **GATCCAGATCTAACTGATTGGGGTGGCTTTGTCTTCTTCTTTTGCCAATTCCACTAATTGTTTGGG** | Ins_k3_14_TGA;Ins_k3_15_GAT | 0.4953982672612778 | 0.032678812733901386 | 80.58054974269253 |
| **GATCCAGATCTAACTGCATGGGGTGGCTTTGTCTTCTTCTTTTGCCAATTCCACTAATTGTTTGGG** | Ins_k3_15_GCA | 0.4916219793252059 | 0.03262725099793634 | 80.27682821571749 |
| **GATCCAGATCTAACTGCCAATTCCACTAATTGTTTGGG** | Del_k25_15-39;Del_k25_16-40 | -0.995881147945729 | 0.032947911712803556 | 18.13743232266479 |
| **GATCCAGATCTAACTGCCTGGGGTGGCTTTGTCTTCTTCTTTTGCCAATTCCACTAATTGTTTGGG** | Ins_k3_15_GCC | 0.666713348578581 | 0.0308305474823347 | 95.63820588588159 |
| **GATCCAGATCTAACTGCGTGGGGTGGCTTTGTCTTCTTCTTTTGCCAATTCCACTAATTGTTTGGG** | Ins_k3_15_GCG | 0.6860365058685696 | 0.02843849316632297 | 97.50420845068452 |
| **GATCCAGATCTAACTGCTGGGGTGGCTTTGTCTTCTTCTTTTGCCAATTCCACTAATTGTTTGGG** | Ins_k2_15_GC | 0.7433215322946616 | 0.03330044468743442 | 100 |
| **GATCCAGATCTAACTGCTTGGGGTGGCTTTGTCTTCTTCTTTTGCCAATTCCACTAATTGTTTGGG** | Ins_k3_13_CTG;Ins_k3_14_TGC; | 0.35362556726872885 | 0.03296262512954824 | 69.92929093811006 |
| **GATCCAGATCTAACTGCTTTGGGGTGGCTTTGTCTTCTTCTTTTGCCAATTCCACTAATTGTTTGGG** | Ins_k4_14_TGCT;Ins_k4_15_GCT | 0.4768683303591952 | 0.030554648745349907 | 79.1011461978419 |
| **GATCCAGATCTAACTGCTTTGTCTTCTTCTTTTGCCAATTCCACTAATTGTTTGGG** | Del_k7_16-22 | -0.679923492475476 | 0.03464814904220378 | 24.876797514541288 |
| **GATCCAGATCTAACTGG** | Del_k46_16-61 | -6.703397154991188 | 0.3701072771658197 | 0.060232806285621256 |
| **GATCCAGATCTAACTGGATGGGGTGGCTTTGTCTTCTTCTTTTGCCAATTCCACTAATTGTTTGGG** | Ins_k3_15_GGA | 0.4657695368337229 | 0.02794202728012492 | 78.22807290100405 |
| **GATCCAGATCTAACTGGCTGGGGTGGCTTTGTCTTCTTCTTTTGCCAATTCCACTAATTGTTTGGG** | Ins_k3_15_GGC | 0.5655810087240019 | 0.028282226949333855 | 86.43909341216202 |
| **GATCCAGATCTAACTGGCTTTGTCTTCTTCTTTTGCCAATTCCACTAATTGTTTGGG** | Del_k6_15-20;Del_k6_16-21 | -0.24491501216656847 | 0.030038255541185114 | 38.43405911629064 |
| **GATCCAGATCTAACTGGG** | Del_k45_15-59;Del_k45_16-60 | -6.263516833543835 | 0.2938784524032928 | 0.09351272097406281 |
| **GATCCAGATCTAACTGGGGGTGGCTTTGTCTTCTTCTTTTGCCAATTCCACTAATTGTTTGGG** | T16G | -0.5393720164557199 | 0.04131737106181104 | 28.630913289656903 |
| **GATCCAGATCTAACTGGGGTGGCTTTGTCTTCTTCTTTTGCCAATTCCACTAATTGTTTGGG** | Del_k1_15-15;Del_k1_16-16 | 0.1945133937334914 | 0.0327355769705106 | 59.642739843268096 |
| **GATCCAGATCTAACTGGGTGGCTTTGTCTTCTTCTTTTGCCAATTCCACTAATTGTTTGGG** | Del_k2_16-17 | 0.4334153878032949 | 0.030371769439401536 | 75.73757636863353 |
| **GATCCAGATCTAACTGGGTGGGGTGGCTTTGTCTTCTTCTTTTGCCAATTCCACTAATTGTTTGGG** | Ins_k3_15_GGG | -4.240347125084339 | 0.22414228600659378 | 0.7071672413982243 |
| **GATCCAGATCTAACTGGTGGCTTTGTCTTCTTCTTTTGCCAATTCCACTAATTGTTTGGG** | Del_k3_16-18 | 0.43965374026419 | 0.028893969450289783 | 76.21153087511134 |
| **GATCCAGATCTAACTGGTGGGGTGGCTTTGTCTTCTTCTTTTGCCAATTCCACTAATTGTTTGGG** | Ins_k2_15_GG | -0.38077625163224543 | 0.05125294626074375 | 33.55154064300327 |
| **GATCCAGATCTAACTGGTTGGGGTGGCTTTGTCTTCTTCTTTTGCCAATTCCACTAATTGTTTGGG** | Ins_k3_14_TGG;Ins_k3_15_GGT | 0.12316777358375519 | 0.03549241690420862 | 55.53574171737052 |
| **GATCCAGATCTAACTGTATGGGGTGGCTTTGTCTTCTTCTTTTGCCAATTCCACTAATTGTTTGGG** | Ins_k3_15_GTA | 0.020729465314110906 | 0.03333416988678022 | 50.12843941919925 |
| **GATCCAGATCTAACTGTCTGGGGTGGCTTTGTCTTCTTCTTTTGCCAATTCCACTAATTGTTTGGG** | Ins_k3_15_GTC | 0.1927126925391771 | 0.0332283646481857 | 59.535437728917024 |
| **GATCCAGATCTAACTGTCTTCTTCTTTTGCCAATTCCACTAATTGTTTGGG** | Del_k12_15-26;Del_k12_16-27 | -1.0305411343109872 | 0.040855076894342646 | 17.51955876402766 |
| **GATCCAGATCTAACTGTCTTGGGGTGGCTTTGTCTTCTTCTTTTGCCAATTCCACTAATTGTTTGGG** | Ins_k4_13_CTGT;Ins_k4_14_TGT | 0.26641274742778764 | 0.03383101310754799 | 64.08893846767148 |
| **GATCCAGATCTAACTGTGGCTTTGTCTTCTTCTTTTGCCAATTCCACTAATTGTTTGGG** | Del_k4_16-19 | 0.24589564217083204 | 0.030635482927854184 | 62.78741636052205 |
| **GATCCAGATCTAACTGTGGGGTGGCTTTGTCTTCTTCTTTTGCCAATTCCACTAATTGTTTGGG** | Ins_k1_15_G | -0.2637315491473775 | 0.0417375093864129 | 37.71762476671686 |
| **GATCCAGATCTAACTGTGTGGGGTGGCTTTGTCTTCTTCTTTTGCCAATTCCACTAATTGTTTGGG** | Ins_k3_15_GTG | -0.1331885914929728 | 0.035800373437939174 | 42.977230118220554 |
| **GATCCAGATCTAACTGTTGGGGTGGCTTTGTCTTCTTCTTTTGCCAATTCCACTAATTGTTTGGG** | Ins_k2_14_TG;Ins_k2_15_GT | 0.41673023908374107 | 0.034601155860715126 | 74.48436770344112 |
| **GATCCAGATCTAACTGTTTGGG** | Del_k41_15-55;Del_k41_16-56 | -5.614728165029033 | 0.204409777543629 | 0.17891058372199434 |
| **GATCCAGATCTAACTGTTTGGGGTGGCTTTGTCTTCTTCTTTTGCCAATTCCACTAATTGTTTGGG** | Ins_k3_14_TGT;Ins_k3_15_GTT | 0.02937460540909065 | 0.035959267832401935 | 50.56368547147198 |
| **GATCCAGATCTAACTT** | Del_k47_17-63 | -5.9566349585118115 | 0.18590830088409144 | 0.12710065800798562 |
| **GATCCAGATCTAACTTAAAGGGGTGGCTTTGTCTTCTTCTTTTGCCAATTCCACTAATTGTTTGGG** | Ins_k3_16_AAA | -0.7863924698626049 | 0.048977054475717645 | 22.364314032426687 |
| **GATCCAGATCTAACTTAAATGGGGTGGCTTTGTCTTCTTCTTTTGCCAATTCCACTAATTGTTTGGG** | Ins_k4_15_TAAA;Ins_k4_16_AAA | -1.0738926096208001 | 0.05670823922552677 | 16.776287401848833 |
| **GATCCAGATCTAACTTAACGGGGTGGCTTTGTCTTCTTCTTTTGCCAATTCCACTAATTGTTTGGG** | Ins_k3_16_AAC | -0.16922073277435729 | 0.033797039665120335 | 41.456235396391314 |
| **GATCCAGATCTAACTTAAGGGGGTGGCTTTGTCTTCTTCTTTTGCCAATTCCACTAATTGTTTGGG** | Ins_k3_16_AAG | -2.1578970409954015 | 0.10463203440353504 | 5.674383901178815 |
| **GATCCAGATCTAACTTAAGGGGTGGCTTTGTCTTCTTCTTTTGCCAATTCCACTAATTGTTTGGG** | Ins_k2_16_AA | -0.6177041591732539 | 0.05397521315917939 | 26.473781758129345 |
| **GATCCAGATCTAACTTAATGGGGTGGCTTTGTCTTCTTCTTTTGCCAATTCCACTAATTGTTTGGG** | Ins_k3_15_TAA;Ins_k3_16_AAT | -1.0007262137260222 | 0.05745956385957502 | 18.04976781162831 |
| **GATCCAGATCTAACTTAATTCCACTAATTGTTTGGG** | Del_k27_17-43 | -4.087171147955431 | 0.13016706391780355 | 0.8242246826230776 |

| | | | | |
|---|---|---|---|---|
| **GATCCAGATCTAACTTAATTGTTTGGG** | Del_k36_16-51;Del_k36_17-52 | -5.715931601126295 | 0.28678008242313724 | 0.16169028957162307 |
| **GATCCAGATCTAACTTACAGGGGTGGCTTTGTCTTCTTCTTTTGCCAATTCCACTAATTGTTTGGG** | Ins_k3_16_ACA | -0.2687882418708228 | 0.04462882362433581 | 37.527379738720256 |
| **GATCCAGATCTAACTTACCGGGGTGGCTTTGTCTTCTTCTTTTGCCAATTCCACTAATTGTTTGGG** | Ins_k3_16_ACC | 0.5163830194808992 | 0.0313911196097981 | 82.28937959035893 |
| **GATCCAGATCTAACTTACGGGGGTGGCTTTGTCTTCTTCTTTTGCCAATTCCACTAATTGTTTGGG** | Ins_k3_16_ACG | -0.009192214322147452 | 0.03650933986207996 | 48.65073033180495 |
| **GATCCAGATCTAACTTACGGGGTGGCTTTGTCTTCTTCTTTTGCCAATTCCACTAATTGTTTGGG** | Ins_k2_16_AC | 0.48574260595294927 | 0.0329774748023737 | 79.80623552016041 |
| **GATCCAGATCTAACTTACTAATTGTTTGGG** | Del_k33_17-49 | -5.217034835342197 | 0.20219916298229088 | 0.2662882792286898 |
| **GATCCAGATCTAACTTACTGGGGTGGCTTTGTCTTCTTCTTTTGCCAATTCCACTAATTGTTTGGG** | Ins_k3_15_TAC;Ins_k3_16_ACT | -0.047449416324171545 | 0.03752652276743331 | 46.82464273345647 |
| **GATCCAGATCTAACTTAGAGGGGTGGCTTTGTCTTCTTCTTTTGCCAATTCCACTAATTGTTTGGG** | Ins_k3_16_AGA | -0.3232648751819527 | 0.04083259800014028 | 35.5377019474427 |
| **GATCCAGATCTAACTTAGCGGGGTGGCTTTGTCTTCTTCTTTTGCCAATTCCACTAATTGTTTGGG** | Ins_k3_16_AGC | -0.552242997316519 | 0.04399678688189565 | 28.264766739999683 |
| **GATCCAGATCTAACTTAGGAGGGGTGGCTTTGTCTTCTTCTTTTGCCAATTCCACTAATTGTTTGGG** | Ins_k4_16_AGGA | -0.35088363068292816 | 0.04748040515948221 | 34.56962492389611 |
| **GATCCAGATCTAACTTAGGGGGGTGGCTTTGTCTTCTTCTTTTGCCAATTCCACTAATTGTTTGGG** | Ins_k3_16_AGG | -4.950451283282819 | 0.5195199306335014 | 0.34763845958642364 |
| **GATCCAGATCTAACTTAGGGGGTGGCTTTGTCTTCTTCTTTTGCCAATTCCACTAATTGTTTGGG** | Ins_k2_16_AG | -2.997882043116987 | 0.17418771782413014 | 2.4497279845623954 |
| **GATCCAGATCTAACTTAGGGGTGGCTTTGTCTTCTTCTTTTGCCAATTCCACTAATTGTTTGGG** | Ins_k1_16_A | -1.5860539335158403 | 0.06809308738424436 | 10.052336562710558 |
| **GATCCAGATCTAACTTAGGGTGGCTTTGTCTTCTTCTTTTGCCAATTCCACTAATTGTTTGGG** | G17A | -1.2891786333232915 | 0.04314099816124745 | 13.52690143411681 |
| **GATCCAGATCTAACTTAGGTGGGGTGGCTTTGTCTTCTTCTTTTGCCAATTCCACTAATTGTTTGGG** | Ins_k4_15_TAGG;Ins_k4_16_AG | -3.8479593647652597 | 0.24540567861444995 | 1.0469693632879613 |
| **GATCCAGATCTAACTTAGTGGGGTGGCTTTGTCTTCTTCTTTTGCCAATTCCACTAATTGTTTGGG** | Ins_k3_15_TAG;Ins_k3_16_AGT | -3.4863066487906678 | 0.21570149100331665 | 1.5031341866738561 |
| **GATCCAGATCTAACTTATAGGGGTGGCTTTGTCTTCTTCTTTTGCCAATTCCACTAATTGTTTGGG** | Ins_k3_16_ATA | -1.9845478932590959 | 0.08714935465894112 | 6.7484384780807805 |
| **GATCCAGATCTAACTTATCGGGGTGGCTTTGTCTTCTTCTTTTGCCAATTCCACTAATTGTTTGGG** | Ins_k3_16_ATC | 0.2125834135017669 | 0.03164384876533184 | 60.730281690245484 |
| **GATCCAGATCTAACTTATGGGGGTGGCTTTGTCTTCTTCTTTTGCCAATTCCACTAATTGTTTGGG** | Ins_k3_16_ATG | -1.561814394604871 | 0.06696343522323893 | 10.298977723822013 |
| **GATCCAGATCTAACTTATGGGGTGGCTTTGTCTTCTTCTTTTGCCAATTCCACTAATTGTTTGGG** | Ins_k2_15_TA;Ins_k2_16_AT | -0.2754323824511523 | 0.04007691387656493 | 37.27886903634581 |
| **GATCCAGATCTAACTTATTCCACTAATTGTTTGGG** | Del_k28_17-44 | -1.0711155248251094 | 0.40369356540398904 | 16.822941325487953 |
| **GATCCAGATCTAACTTATTGGGGTGGCTTTGTCTTCTTCTTTTGCCAATTCCACTAATTGTTTGGG** | Ins_k3_14_TTA;Ins_k3_15_TAT;In | -0.5651113835438294 | 0.04103182153870171 | 27.90337505590155 |
| **GATCCAGATCTAACTTATTGTTTGGG** | Del_k37_17-53 | -5.783031169558261 | 0.20507968833536286 | 0.1511969277196262 |
| **GATCCAGATCTAACTTCAAGGGGTGGCTTTGTCTTCTTCTTTTGCCAATTCCACTAATTGTTTGGG** | Ins_k3_16_CAA | 0.46895838997083295 | 0.02992254089361689 | 78.4779289020032 |
| **GATCCAGATCTAACTTCAATTCCACTAATTGTTTGGG** | Del_k26_17-42 | -1.0912270872246166 | 0.035024814187536925 | 16.48798522655582 |
| **GATCCAGATCTAACTTCACGGGGTGGCTTTGTCTTCTTCTTTTGCCAATTCCACTAATTGTTTGGG** | Ins_k3_16_CAC | 0.6290871165179883 | 0.02907047184962373 | 92.1065584955744 |
| **GATCCAGATCTAACTTCACTAATTGTTTGGG** | Del_k32_17-48 | -2.808677308907055 | 0.0577571146746585 | 2.9599776744542496 |
| **GATCCAGATCTAACTTCAGGGGGTGGCTTTGTCTTCTTCTTTTGCCAATTCCACTAATTGTTTGGG** | Ins_k3_16_CAG | 0.10590122521715628 | 0.03824182969862542 | 54.58506224251396 |
| **GATCCAGATCTAACTTCAGGGGTGGCTTTGTCTTCTTCTTTTGCCAATTCCACTAATTGTTTGGG** | Ins_k2_16_CA | 0.6215058449246103 | 0.030281565920290764 | 91.41091392717465 |
| **GATCCAGATCTAACTTCATGGGGTGGCTTTGTCTTCTTCTTTTGCCAATTCCACTAATTGTTTGGG** | Ins_k3_15_TCA;Ins_k3_16_CAT | 0.5353209051818961 | 0.03048093848290979 | 83.86261632519796 |
| **GATCCAGATCTAACTTCCAATTCCACTAATTGTTTGGG** | Del_k25_17-41 | -1.6427789129772563 | 0.04205724349711745 | 9.497989283377393 |
| **GATCCAGATCTAACTTCCACTAATTGTTTGGG** | Del_k31_15-45;Del_k31_16-46;D | -2.884488892572885 | 0.054937094627891495 | 2.7438722213818787 |
| **GATCCAGATCTAACTTCCAGGGGTGGCTTTGTCTTCTTCTTTTGCCAATTCCACTAATTGTTTGGG** | Ins_k3_16_CCA | 0.599469037416676 | 0.029338459491975364 | 89.41854259207679 |
| **GATCCAGATCTAACTTCCATGGGGTGGCTTTGTCTTCTTCTTTTGCCAATTCCACTAATTGTTTGGG** | Ins_k4_15_TCCA;Ins_k4_16_CC | 0.7867802138372584 | 0.032337828422153325 | 100 |
| **GATCCAGATCTAACTTCCCGGGGTGGCTTTGTCTTCTTCTTTTGCCAATTCCACTAATTGTTTGGG** | Ins_k3_16_CCC | 0.7816651509472239 | 0.031202176952280447 | 100 |
| **GATCCAGATCTAACTTCCGGGGGTGGCTTTGTCTTCTTCTTTTGCCAATTCCACTAATTGTTTGGG** | Ins_k3_16_CCG | 0.6193576515744765 | 0.03168588577457795 | 91.21475637735327 |
| **GATCCAGATCTAACTTCCGGGGTGGCTTTGTCTTCTTCTTTTGCCAATTCCACTAATTGTTTGGG** | Ins_k2_16_CC | 0.9684223759316113 | 0.030591583589713353 | 100 |
| **GATCCAGATCTAACTTCCTCGGGGTGGCTTTGTCTTCTTCTTTTGCCAATTCCACTAATTGTTTGGG** | Ins_k4_16_CCTC | 0.8057612008682609 | 0.029761484563289014 | 100 |
| **GATCCAGATCTAACTTCCTGGGGTGGCTTTGTCTTCTTCTTTTGCCAATTCCACTAATTGTTTGGG** | Ins_k3_15_TCC;Ins_k3_16_CCT | 0.47965095529273505 | 0.031079288073088827 | 79.32156154391946 |
| **GATCCAGATCTAACTTCGAGGGGTGGCTTTGTCTTCTTCTTTTGCCAATTCCACTAATTGTTTGGG** | Ins_k3_16_CGA | 0.570740814861874 | 0.02912773312228638 | 86.8862550184153 |
| **GATCCAGATCTAACTTCGCGGGGTGGCTTTGTCTTCTTCTTTTGCCAATTCCACTAATTGTTTGGG** | Ins_k3_16_CGC | 0.8026531334332494 | 0.02896211110214489 | 100 |
| **GATCCAGATCTAACTTCGGGGGGTGGCTTTGTCTTCTTCTTTTGCCAATTCCACTAATTGTTTGGG** | Ins_k3_16_CGG | -1.1421707658826143 | 0.06810931580907396 | 15.669063158663377 |
| **GATCCAGATCTAACTTCGGGGGTGGCTTTGTCTTCTTCTTTTGCCAATTCCACTAATTGTTTGGG** | Ins_k2_16_CG | 0.7176434322918916 | 0.033564802472155125 | 100 |
| **GATCCAGATCTAACTTCGGGGTGGCTTTGTCTTCTTCTTTTGCCAATTCCACTAATTGTTTGGG** | Ins_k1_16_C | 0.7332758967939335 | 0.028720272552075 | 100 |
| **GATCCAGATCTAACTTCGGGTGGCTTTGTCTTCTTCTTTTGCCAATTCCACTAATTGTTTGGG** | G17C | 0.3048408546420497 | 0.028013770887289992 | 66.59968777301754 |
| **GATCCAGATCTAACTTCGTGGGGTGGCTTTGTCTTCTTCTTTTGCCAATTCCACTAATTGTTTGGG** | Ins_k3_15_TCG;Ins_k3_16_CGT | 0.5815996813910747 | 0.03043843109482674 | 87.83488245279972 |
| **GATCCAGATCTAACTTCTAATTGTTTGGG** | Del_k34_17-50 | -3.556121762567278 | 0.07463882104161271 | 1.4017721705498698 |
| **GATCCAGATCTAACTTCTAGGGGTGGCTTTGTCTTCTTCTTTTGCCAATTCCACTAATTGTTTGGG** | Ins_k3_16_CTA | -0.349859506655291 | 0.04445903169264147 | 34.60504664242765 |
| **GATCCAGATCTAACTTCTCGGGGTGGCTTTGTCTTCTTCTTTTGCCAATTCCACTAATTGTTTGGG** | Ins_k3_16_CTC | 0.5618563912312953 | 0.03684067638380377 | 86.11773968414933 |
| **GATCCAGATCTAACTTCTGGGGGTGGCTTTGTCTTCTTCTTTTGCCAATTCCACTAATTGTTTGGG** | Ins_k3_16_CTG | -0.13667932672215588 | 0.03807281359880389 | 42.82746952635344 |
| **GATCCAGATCTAACTTCTGGGGTGGCTTTGTCTTCTTCTTTTGCCAATTCCACTAATTGTTTGGG** | Ins_k2_15_TC;Ins_k2_16_CT | 0.3625472730827819 | 0.03442618402672181 | 70.55597086983435 |
| **GATCCAGATCTAACTTCTTCTTCTTTTGCCAATTCCACTAATTGTTTGGG** | Del_k13_16-28;Del_k13_17-29 | -1.0333093668493252 | 0.04688049309054871 | 17.471127616666518 |
| **GATCCAGATCTAACTTCTTCTTTTGCCAATTCCACTAATTGTTTGGG** | Del_k16_14-29;Del_k16_15-30;D | -0.917353488856443 | 0.034863504521826315 | 19.61913854016242 |
| **GATCCAGATCTAACTTCTTGGGGTGGCTTTGTCTTCTTCTTTTGCCAATTCCACTAATTGTTTGGG** | Ins_k3_13_CTT;Ins_k3_14_TTC;I | -0.05948640435177477 | 0.03589186986680252 | 46.264393689475575 |
| **GATCCAGATCTAACTTCTTTGTCTTCTTCTTTTGCCAATTCCACTAATTGTTTGGG** | Del_k7_17-23 | -1.839395774125144 | 0.07500971068543653 | 7.802648760079631 |
| **GATCCAGATCTAACTTCTTTTGCCAATTCCACTAATTGTTTGGG** | Del_k19_14-32;Del_k19_15-33;D | -0.9795702743414668 | 0.038025489307685005 | 18.435695543049327 |
| **GATCCAGATCTAACTTG** | Del_k46_17-62;Del_k46_18-63 | -3.798444188246886 | 0.05671293643564615 | 1.1001151394497213 |
| **GATCCAGATCTAACTTGAAAGGGTGGCTTTGTCTTCTTCTTTTGCCAATTCCACTAATTGTTTGGG** | Ins_k3_17_AAA | 0.07327640747677899 | 0.03538428611191954 | 52.83297077259151 |
| **GATCCAGATCTAACTTGAACGGGTGGCTTTGTCTTCTTCTTTTGCCAATTCCACTAATTGTTTGGG** | Ins_k3_17_AAC | 0.49751062784134015 | 0.028632117841458863 | 80.75094482405568 |
| **GATCCAGATCTAACTTGAAGGGGTGGCTTTGTCTTCTTCTTTTGCCAATTCCACTAATTGTTTGGG** | Ins_k3_16_GAA;Ins_k3_17_AAG | 0.22910024118149908 | 0.034642676385856545 | 61.741682864432235 |
| **GATCCAGATCTAACTTGAAGGGTGGCTTTGTCTTCTTCTTTTGCCAATTCCACTAATTGTTTGGG** | Ins_k2_17_AA | 0.3612958650014376 | 0.030812965747480804 | 70.46773178077069 |
| **GATCCAGATCTAACTTGAATGGGTGGCTTTGTCTTCTTCTTTTGCCAATTCCACTAATTGTTTGGG** | Ins_k3_17_AAT | 0.11517880435232475 | 0.031211897982118337 | 55.09383592178133 |
| **GATCCAGATCTAACTTGAATTCCACTAATTGTTTGGG** | Del_k26_18-43 | -1.6298893756667576 | 0.034173060378645696 | 9.621206370302197 |
| **GATCCAGATCTAACTTGAATTGTTTGGG** | Del_k35_18-52 | -4.14984392839694 | 0.10924173304493176 | 0.7741536633815533 |
| **GATCCAGATCTAACTTGACAGGGTGGCTTTGTCTTCTTCTTTTGCCAATTCCACTAATTGTTTGGG** | Ins_k3_17_ACA | -0.4419666906914561 | 0.03478784678684341 | 31.560058327079158 |
| **GATCCAGATCTAACTTGACCGGGTGGCTTTGTCTTCTTCTTTTGCCAATTCCACTAATTGTTTGGG** | Ins_k3_17_ACC | 0.50942786277551 | 0.027505809306528906 | 81.71902979540211 |
| **GATCCAGATCTAACTTGACGGGGTGGCTTTGTCTTCTTCTTTTGCCAATTCCACTAATTGTTTGGG** | Ins_k3_16_GAC;Ins_k3_17_ACG | 0.1371295701233597 | 0.129106486548893 | 56.31655856307925 |
| **GATCCAGATCTAACTTGACGGGTGGCTTTGTCTTCTTCTTTTGCCAATTCCACTAATTGTTTGGG** | Ins_k2_17_AC | 0.570124013236172 | 0.02964607249214328 | 86.83267995935907 |
| **GATCCAGATCTAACTTGACTAATTGTTTGGG** | Del_k32_18-49 | -4.373000584332791 | 0.12482137114755609 | 0.6193148147657404 |
| **GATCCAGATCTAACTTGACTGGGTGGCTTTGTCTTCTTCTTTTGCCAATTCCACTAATTGTTTGGG** | Ins_k3_17_ACT | 0.25837640806437434 | 0.03188023858679463 | 63.57596199599327 |
| **GATCCAGATCTAACTTGAGAGGGTGGCTTTGTCTTCTTCTTTTGCCAATTCCACTAATTGTTTGGG** | Ins_k3_17_AGA | 0.18780426699301267 | 0.03165072191081953 | 59.24392847646592 |
| **GATCCAGATCTAACTTGAGCGGGTGGCTTTGTCTTCTTCTTTTGCCAATTCCACTAATTGTTTGGG** | Ins_k3_17_AGC | 0.3693512877862166 | 0.03270225745260266 | 71.03767162421101 |
| **GATCCAGATCTAACTTGAGGGGGTGGCTTTGTCTTCTTCTTTTGCCAATTCCACTAATTGTTTGGG** | Ins_k3_16_GAG;Ins_k3_17_AGG | -0.5982991148683093 | 0.04256217632624728 | 26.99252349139422 |
| **GATCCAGATCTAACTTGAGGGGTGGCTTTGTCTTCTTCTTTTGCCAATTCCACTAATTGTTTGGG** | Ins_k2_16_GA;Ins_k2_17_AG | 0.03583543289686908 | 0.03520626166869136 | 50.8914263189338 |
| **GATCCAGATCTAACTTGAGGGTGGCTTTGTCTTCTTCTTTTGCCAATTCCACTAATTGTTTGGG** | Ins_k1_17_A | 0.1747895766900126 | 0.03371358822557695 | 58.47788283126861 |

| | | | | |
|---|---|---|---|---|
| **GATCCAGATCTAACTTGAGGTGGCTTTGTCTTCTTCTTTTGCCAATTCCACTAATTGTTTGGG** | G18A | -0.2545530923150184 | 0.029209312560891817 | 38.0654079724752 |
| **GATCCAGATCTAACTTGAGTCGGGTGGCTTTGTCTTCTTCTTTTGCCAATTCCACTAATTGTTTGGG** | Ins_k4_17_AGTC | 0.413769172114715 | 0.03147777668370404 | 74.26414071686449 |
| **GATCCAGATCTAACTTGAGTGGGTGGCTTTGTCTTCTTCTTTTGCCAATTCCACTAATTGTTTGGG** | Ins_k3_17_AGT | -0.7069418743323628 | 0.04722718335301406 | 24.2136654176802 |
| **GATCCAGATCTAACTTGATAGGGTGGCTTTGTCTTCTTCTTTTGCCAATTCCACTAATTGTTTGGG** | Ins_k3_17_ATA | -2.2324899269348686 | 0.08151794620237754 | 5.266516339716293 |
| **GATCCAGATCTAACTTGATCGGGTGGCTTTGTCTTCTTCTTTTGCCAATTCCACTAATTGTTTGGG** | Ins_k3_17_ATC | 0.2961839249767294 | 0.032606139254019305 | 66.02562734572479 |
| **GATCCAGATCTAACTTGATGGGGTGGCTTTGTCTTCTTCTTTTGCCAATTCCACTAATTGTTTGGG** | Ins_k3_15_TGA;Ins_k3_16_GAT;I | 0.1832873551687001 | 0.03217721535419654 | 58.976932328389736 |
| **GATCCAGATCTAACTTGATGGGTGGCTTTGTCTTCTTCTTTTGCCAATTCCACTAATTGTTTGGG** | Ins_k2_17_AT | 0.29518707635858055 | 0.031383762689865605 | 65.95984258451452 |
| **GATCCAGATCTAACTTGATTCCACTAATTGTTTGGG** | Del_k27_18-44 | -2.7876147347956315 | 0.0430114285226817 | 3.0229836281036735 |
| **GATCCAGATCTAACTTGATTGGGTGGCTTTGTCTTCTTCTTTTGCCAATTCCACTAATTGTTTGGG** | Ins_k3_17_ATT | -0.08681655554331191 | 0.034895004852568516 | 45.017102776139716 |
| **GATCCAGATCTAACTTGATTGTTTGGG** | Del_k36_18-53 | -4.6401695737461415 | 0.11984107466038929 | 0.47411254965396815 |
| **GATCCAGATCTAACTTGCAAGGGTGGCTTTGTCTTCTTCTTTTGCCAATTCCACTAATTGTTTGGG** | Ins_k3_17_CAA | 0.32921287076209116 | 0.03027431453467686 | 68.24279805971584 |
| **GATCCAGATCTAACTTGCAATTCCACTAATTGTTTGGG** | Del_k25_18-42 | -2.0034009099071994 | 0.03997526696554987 | 6.622401873351032 |
| **GATCCAGATCTAACTTGCACGGGTGGCTTTGTCTTCTTCTTTTGCCAATTCCACTAATTGTTTGGG** | Ins_k3_17_CAC | 0.5157350997359382 | 0.031049975226945914 | 82.23607994533256 |
| **GATCCAGATCTAACTTGCACTAATTGTTTGGG** | Del_k31_18-48 | -4.227173774632696 | 0.10898962354285466 | 0.716544633523369 |
| **GATCCAGATCTAACTTGCAGGGGTGGCTTTGTCTTCTTCTTTTGCCAATTCCACTAATTGTTTGGG** | Ins_k3_16_GCA;Ins_k3_17_CAG | 0.18306881548872356 | 0.03256486815543576 | 58.964044936727205 |
| **GATCCAGATCTAACTTGCAGGGTGGCTTTGTCTTCTTCTTTTGCCAATTCCACTAATTGTTTGGG** | Ins_k2_17_CA | 0.11628353784253564 | 0.03506690573183144 | 55.15473355906452 |
| **GATCCAGATCTAACTTGCATGGGTGGCTTTGTCTTCTTCTTTTGCCAATTCCACTAATTGTTTGGG** | Ins_k3_17_CAT | 0.25646141133605604 | 0.030236908474753833 | 63.45433073567468 |
| **GATCCAGATCTAACTTGCCAATTCCACTAATTGTTTGGG** | Del_k24_15-38;Del_k24_16-39;D | -1.9048981913321639 | 0.04238853022719052 | 7.307935726075888 |
| **GATCCAGATCTAACTTGCCACTAATTGTTTGGG** | Del_k30_18-47 | -3.3018146511825033 | 0.06615404987134317 | 1.807680232220393 |
| **GATCCAGATCTAACTTGCCAGGGTGGCTTTGTCTTCTTCTTTTGCCAATTCCACTAATTGTTTGGG** | Ins_k3_17_CCA | 0.38810543673683195 | 0.029265234626339145 | 72.38249378017517 |
| **GATCCAGATCTAACTTGCCCGGGTGGCTTTGTCTTCTTCTTTTGCCAATTCCACTAATTGTTTGGG** | Ins_k3_17_CCC | 0.6244986813524112 | 0.029914813129521465 | 91.68490163596172 |
| **GATCCAGATCTAACTTGCCGGGGTGGCTTTGTCTTCTTCTTTTGCCAATTCCACTAATTGTTTGGG** | Ins_k3_16_GCC;Ins_k3_17_CCG | 0.9029163022779532 | 0.029090493181845204 | 100 |
| **GATCCAGATCTAACTTGCCGGGTGGCTTTGTCTTCTTCTTTTGCCAATTCCACTAATTGTTTGGG** | Ins_k2_17_CC | 0.7664418961423078 | 0.032178753998106054 | 100 |
| **GATCCAGATCTAACTTGCCTGGGTGGCTTTGTCTTCTTCTTTTGCCAATTCCACTAATTGTTTGGG** | Ins_k3_17_CCT | 0.4399363784652406 | 0.027945452720990603 | 76.2330742094384 |
| **GATCCAGATCTAACTTGCGAGGGTGGCTTTGTCTTCTTCTTTTGCCAATTCCACTAATTGTTTGGG** | Ins_k3_17_CGA | 0.5847361774744126 | 0.028941621285350834 | 88.11080871220486 |
| **GATCCAGATCTAACTTGCGCGGGTGGCTTTGTCTTCTTCTTTTGCCAATTCCACTAATTGTTTGGG** | Ins_k3_17_CGC | 0.7079653840124447 | 0.031223016314319466 | 99.66598236668669 |
| **GATCCAGATCTAACTTGCGGGGGGTGGCTTTGTCTTCTTCTTTTGCCAATTCCACTAATTGTTTGGG** | Ins_k4_16_GCGG;Ins_k4_17_CG | -1.596679534501638 | 0.0993844092790737 | 9.946089912308965 |
| **GATCCAGATCTAACTTGCGGGGGTGGCTTTGTCTTCTTCTTTTGCCAATTCCACTAATTGTTTGGG** | Ins_k3_16_GCG;Ins_k3_17_CGG | 0.3918003646817906 | 0.03332441434302453 | 72.65043658901482 |
| **GATCCAGATCTAACTTGCGGGGTGGCTTTGTCTTCTTCTTTTGCCAATTCCACTAATTGTTTGGG** | Ins_k2_16_GC;Ins_k2_17_CG | 0.6850294503091567 | 0.03166640236114898 | 97.40606572138306 |
| **GATCCAGATCTAACTTGCGGGTGGCTTTGTCTTCTTCTTTTGCCAATTCCACTAATTGTTTGGG** | Ins_k1_17_C | 0.7327886738843534 | 0.029680554578868285 | 100 |
| **GATCCAGATCTAACTTGCGGTGGCTTTGTCTTCTTCTTTTGCCAATTCCACTAATTGTTTGGG** | G18C | 0.21103588401455342 | 0.027487264393156456 | 60.636372470947556 |
| **GATCCAGATCTAACTTGCGTGGGTGGCTTTGTCTTCTTCTTTTGCCAATTCCACTAATTGTTTGGG** | Ins_k3_17_CGT | 0.2757201193408937 | 0.030513863891554596 | 64.6882226075072 |
| **GATCCAGATCTAACTTGCTAATTGTTTGGG** | Del_k33_18-50 | -4.692404280107382 | 0.11361545785657445 | 0.4499831032271201 |
| **GATCCAGATCTAACTTGCTAGGGTGGCTTTGTCTTCTTCTTTTGCCAATTCCACTAATTGTTTGGG** | Ins_k3_17_CTA | -2.1508750913399064 | 0.07009944996402516 | 5.714369362909071 |
| **GATCCAGATCTAACTTGCTCGGGTGGCTTTGTCTTCTTCTTTTGCCAATTCCACTAATTGTTTGGG** | Ins_k3_17_CTC | 0.5238296706231242 | 0.028707663769063103 | 82.90444714738554 |
| **GATCCAGATCTAACTTGCTGGGGTGGCTTTGTCTTCTTCTTTTGCCAATTCCACTAATTGTTTGGG** | Ins_k3_15_TGC;Ins_k3_16_GCT; | 0.5710918480443572 | 0.03309278439689003 | 86.91676033090368 |
| **GATCCAGATCTAACTTGCTGGGTGGCTTTGTCTTCTTCTTTTGCCAATTCCACTAATTGTTTGGG** | Ins_k2_17_CT | 0.6774531235645074 | 0.029033737615386895 | 96.67087408330991 |
| **GATCCAGATCTAACTTGCTTAGGGTGGCTTTGTCTTCTTCTTTTGCCAATTCCACTAATTGTTTGGG** | Ins_k4_17_CTTA | -1.486929976799019 | 0.06616237232667578 | 11.099821842912903 |
| **GATCCAGATCTAACTTGCTTCTTCTTTTGCCAATTCCACTAATTGTTTGGG** | Del_k12_18-29 | -1.4291789140257118 | 0.04966413770098885 | 11.759719860195075 |
| **GATCCAGATCTAACTTGCTTCTTTTGCCAATTCCACTAATTGTTTGGG** | Del_k15_18-32 | -1.5609697822267257 | 0.04420252354966229 | 10.307680042415466 |
| **GATCCAGATCTAACTTGCTTGGGTGGCTTTGTCTTCTTCTTTTGCCAATTCCACTAATTGTTTGGG** | Ins_k3_17_CTT | 0.12220836930652557 | 0.03298485863230273 | 55.482486040174955 |
| **GATCCAGATCTAACTTGCTTTGTCTTCTTCTTTTGCCAATTCCACTAATTGTTTGGG** | Del_k6_17-22;Del_k6_18-23 | -2.6387828570472465 | 0.08038836560299359 | 3.508105603766968 |
| **GATCCAGATCTAACTTGCTTTTGCCAATTCCACTAATTGTTTGGG** | Del_k18_18-35 | -1.7030170025341207 | 0.051893143530915654 | 8.942740020952316 |
| **GATCCAGATCTAACTTGG** | Del_k45_17-61;Del_k45_18-62;D | -1.5310288370982053 | 0.025543407691743694 | 10.620968395805033 |
| **GATCCAGATCTAACTTGGAAAGGTGGCTTTGTCTTCTTCTTTTGCCAATTCCACTAATTGTTTGGG** | Ins_k3_18_AAA | 0.46995809783150366 | 0.030082547390335505 | 78.55642313352845 |
| **GATCCAGATCTAACTTGGAACAGGTGGCTTTGTCTTCTTCTTTTGCCAATTCCACTAATTGTTTGGG** | Ins_k4_18_AACA | 0.4406190156639471 | 0.03688235669246289 | 76.28513150776814 |
| **GATCCAGATCTAACTTGGAACGGTGGCTTTGTCTTCTTCTTTTGCCAATTCCACTAATTGTTTGGG** | Ins_k3_18_AAC | 0.6334818474726442 | 0.03032859748035593 | 92.51223280111468 |
| **GATCCAGATCTAACTTGGAAGGGTGGCTTTGTCTTCTTCTTTTGCCAATTCCACTAATTGTTTGGG** | Ins_k3_17_GAA;Ins_k3_18_AAG | 0.6868597776778467 | 0.031943234795949405 | 97.58451396888671 |
| **GATCCAGATCTAACTTGGAAGGTGGCTTTGTCTTCTTCTTTTGCCAATTCCACTAATTGTTTGGG** | Ins_k2_18_AA | 0.7010938682813381 | 0.028622245871717612 | 98.98347362132859 |
| **GATCCAGATCTAACTTGGAATGGTGGCTTTGTCTTCTTCTTTTGCCAATTCCACTAATTGTTTGGG** | Ins_k3_18_AAT | 0.4840359462020112 | 0.027454024228951927 | 79.67014958934598 |
| **GATCCAGATCTAACTTGGAATTCCACTAATTGTTTGGG** | Del_k25_19-43 | -0.41795902622794806 | 0.02602357163815502 | 32.326909943962555 |
| **GATCCAGATCTAACTTGGAATTGTTTGGG** | Del_k34_19-52 | -2.37053026804063 | 0.04493188805446496 | 4.587470426222204 |
| **GATCCAGATCTAACTTGGACAGGTGGCTTTGTCTTCTTCTTTTGCCAATTCCACTAATTGTTTGGG** | Ins_k3_18_ACA | 0.3837686288755914 | 0.05242918814504225 | 72.06926451064432 |
| **GATCCAGATCTAACTTGGACCGGTGGCTTTGTCTTCTTCTTTTGCCAATTCCACTAATTGTTTGGG** | Ins_k3_18_ACC | 0.7434272260228318 | 0.028068868277745834 | 100 |
| **GATCCAGATCTAACTTGGACGGGTGGCTTTGTCTTCTTCTTTTGCCAATTCCACTAATTGTTTGGG** | Ins_k3_17_GAC;Ins_k3_18_ACG | 0.5944290218550277 | 0.029403265636553832 | 88.96900553438148 |
| **GATCCAGATCTAACTTGGACGGTGGCTTTGTCTTCTTCTTTTGCCAATTCCACTAATTGTTTGGG** | Ins_k2_18_AC | 0.9093113207416794 | 0.028014964963486405 | 100 |
| **GATCCAGATCTAACTTGGACTAATTGTTTGGG** | Del_k31_19-49 | -1.8719424412601169 | 0.04118874865652235 | 7.552786692765037 |
| **GATCCAGATCTAACTTGGACTGGTGGCTTTGTCTTCTTCTTTTGCCAATTCCACTAATTGTTTGGG** | Ins_k3_18_ACT | 0.6511148446479511 | 0.026493643059392606 | 94.15796771923343 |
| **GATCCAGATCTAACTTGGAGAGGTGGCTTTGTCTTCTTCTTTTGCCAATTCCACTAATTGTTTGGG** | Ins_k3_18_AGA | 0.5835034112774751 | 0.028650215010997983 | 88.00225560968475 |
| **GATCCAGATCTAACTTGGAGCGGTGGCTTTGTCTTCTTCTTTTGCCAATTCCACTAATTGTTTGGG** | Ins_k3_18_AGC | 0.8307859855328306 | 0.027083364312911112 | 100 |
| **GATCCAGATCTAACTTGGAGGGGTGGCTTTGTCTTCTTCTTTTGCCAATTCCACTAATTGTTTGGG** | Ins_k3_16_GGA;Ins_k3_17_GAG | 0.5259389483781081 | 0.02981374749241233 | 83.07950020640271 |
| **GATCCAGATCTAACTTGGAGGGTGGCTTTGTCTTCTTCTTTTGCCAATTCCACTAATTGTTTGGG** | Ins_k2_17_GA;Ins_k2_18_AG | 0.6130481975121049 | 0.03426178549316582 | 90.6410528434487 |
| **GATCCAGATCTAACTTGGAGGTGGCTTTGTCTTCTTCTTTTGCCAATTCCACTAATTGTTTGGG** | Ins_k1_18_A | 0.6219762266420061 | 0.027009323911180598 | 91.45392206418411 |
| **GATCCAGATCTAACTTGGAGTGGCTTTGTCTTCTTCTTTTGCCAATTCCACTAATTGTTTGGG** | G19A | 0.1535021517002343 | 0.025957967968055242 | 57.24619552421551 |
| **GATCCAGATCTAACTTGGAGTGGTGGCTTTGTCTTCTTCTTTTGCCAATTCCACTAATTGTTTGGG** | Ins_k3_18_AGT | 0.5245906842714352 | 0.0281444158000948 | 82.96756257597559 |
| **GATCCAGATCTAACTTGGATAGGTGGCTTTGTCTTCTTCTTTTGCCAATTCCACTAATTGTTTGGG** | Ins_k3_18_ATA | 0.1851284026812403 | 0.03128672464415457 | 59.0856116741728 |
| **GATCCAGATCTAACTTGGATCGGTGGCTTTGTCTTCTTCTTTTGCCAATTCCACTAATTGTTTGGG** | Ins_k3_18_ATC | 0.6306289269454393 | 0.026747926451999568 | 92.24867888109188 |
| **GATCCAGATCTAACTTGGATGGGTGGCTTTGTCTTCTTCTTTTGCCAATTCCACTAATTGTTTGGG** | Ins_k3_17_GAT;Ins_k3_18_ATG | 0.6000531098049005 | 0.02923824624879162 | 89.47078474891559 |
| **GATCCAGATCTAACTTGGATGGTGGCTTTGTCTTCTTCTTTTGCCAATTCCACTAATTGTTTGGG** | Ins_k2_18_AT | 0.7142366629486272 | 0.029067359263101725 | 100 |
| **GATCCAGATCTAACTTGGATTCCACTAATTGTTTGGG** | Del_k26_19-44 | -0.6187596954396614 | 0.026858689732479117 | 26.445852464159156 |
| **GATCCAGATCTAACTTGGATTGGTGGCTTTGTCTTCTTCTTTTGCCAATTCCACTAATTGTTTGGG** | Ins_k3_18_ATT | 0.497756478438099 | 0.028538775177726478 | 80.77079993262466 |
| **GATCCAGATCTAACTTGGATTGTTTGGG** | Del_k35_19-53 | -2.7873587632911563 | 0.05734738615189074 | 3.023757524814491 |
| **GATCCAGATCTAACTTGGCAAGGTGGCTTTGTCTTCTTCTTTTGCCAATTCCACTAATTGTTTGGG** | Ins_k3_18_CAA | 0.29657847172275 | 0.028117967923618297 | 66.05168268183405 |
| **GATCCAGATCTAACTTGGCAATTCCACTAATTGTTTGGG** | Del_k24_19-42 | -1.0204515606146454 | 0.03091720112992059 | 17.69721839111907 |
| **GATCCAGATCTAACTTGGCACGGTGGCTTTGTCTTCTTCTTTTGCCAATTCCACTAATTGTTTGGG** | Ins_k3_18_CAC | 0.6540952335010004 | 0.030889252326352 | 94.439013681738885 |

| | | | | |
|---|---|---|---|---|
| **GATCCAGATCTAACTTGGCACTAATTGTTTGGG** | Del_k30_19-48 | -2.5550756093689246 | 0.05331886493511009 | 3.8144001792108586 |
| **GATCCAGATCTAACTTGGCAGGGTGGCTTTGTCTTCTTCTTTTGCCAATTCCACTAATTGTTTGGG** | Ins_k3_17_GCA;Ins_k3_18_CAG | 0.28871288992606403 | 0.031273816109344633 | 65.53418564454215 |
| **GATCCAGATCTAACTTGGCAGGTGGCTTTGTCTTCTTCTTTTGCCAATTCCACTAATTGTTTGGG** | Ins_k2_18_CA | 0.36386224975798587 | 0.03133426174990908 | 70.64881135398211 |
| **GATCCAGATCTAACTTGGCATGGTGGCTTTGTCTTCTTCTTTTGCCAATTCCACTAATTGTTTGGG** | Ins_k3_18_CAT | 0.48986208244654605 | 0.02807790248666762 | 80.13567352159718 |
| **GATCCAGATCTAACTTGGCCAATTCCACTAATTGTTTGGG** | Del_k23_18-40;Del_k23_19-41 | -0.5910558962186453 | 0.03080316634675442 | 27.18874602441621 |
| **GATCCAGATCTAACTTGGCCACGGTGGCTTTGTCTTCTTCTTTTGCCAATTCCACTAATTGTTTGGG** | Ins_k4_18_CCAC | 0.9317578411197525 | 0.02694373646228926 | 100 |
| **GATCCAGATCTAACTTGGCCACTAATTGTTTGGG** | Del_k29_19-47 | -1.6287012369693075 | 0.036893124454354304 | 9.632644491598601 |
| **GATCCAGATCTAACTTGGCCAGGTGGCTTTGTCTTCTTCTTTTGCCAATTCCACTAATTGTTTGGG** | Ins_k3_18_CCA | 0.47127212561031384 | 0.027450809865130398 | 78.65971630591606 |
| **GATCCAGATCTAACTTGGCCCGGTGGCTTTGTCTTCTTCTTTTGCCAATTCCACTAATTGTTTGGG** | Ins_k3_18_CCC | 0.7872773712636207 | 0.02964724495774437 | 100 |
| **GATCCAGATCTAACTTGGCCGGGTGGCTTTGTCTTCTTCTTTTGCCAATTCCACTAATTGTTTGGG** | Ins_k3_17_GCC;Ins_k3_18_CCG | 0.7898277105292297 | 0.028355046483391915 | 100 |
| **GATCCAGATCTAACTTGGCCGGTGGCTTTGTCTTCTTCTTTTGCCAATTCCACTAATTGTTTGGG** | Ins_k2_18_CC | 0.8637195514567204 | 0.028596525178074574 | 100 |
| **GATCCAGATCTAACTTGGCCTGGTGGCTTTGTCTTCTTCTTTTGCCAATTCCACTAATTGTTTGGG** | Ins_k3_18_CCT | 0.5983187991651162 | 0.028938761764262997 | 89.3157490938289 |
| **GATCCAGATCTAACTTGGCGAGGTGGCTTTGTCTTCTTCTTTTGCCAATTCCACTAATTGTTTGGG** | Ins_k3_18_CGA | 0.6113298792608854 | 0.027599610265054957 | 90.48543640561037 |
| **GATCCAGATCTAACTTGGCGCGGTGGCTTTGTCTTCTTCTTTTGCCAATTCCACTAATTGTTTGGG** | Ins_k3_18_CGC | 0.8242884479761114 | 0.028926256081559455 | 100 |
| **GATCCAGATCTAACTTGGCGGGGTGGCTTTGTCTTCTTCTTTTGCCAATTCCACTAATTGTTTGGG** | Ins_k3_16_GGC;Ins_k3_17_GCG | 0.6733026360007139 | 0.03093641022945957 | 96.27047432454768 |
| **GATCCAGATCTAACTTGGCGGGTGGCTTTGTCTTCTTCTTTTGCCAATTCCACTAATTGTTTGGG** | Ins_k2_17_GC;Ins_k2_18_CG | 0.7520121668096931 | 0.03249765944698628 | 100 |
| **GATCCAGATCTAACTTGGCGGTGGCTTTGTCTTCTTCTTTTGCCAATTCCACTAATTGTTTGGG** | Ins_k1_18_C | 0.8275440913162333 | 0.02768557860551124 | 100 |
| **GATCCAGATCTAACTTGGCGTGGCTTTGTCTTCTTCTTTTGCCAATTCCACTAATTGTTTGGG** | G19C | 0.3773107737357376 | 0.02662477397561625 | 71.6053511951414 |
| **GATCCAGATCTAACTTGGCGTGGTGGCTTTGTCTTCTTCTTTTGCCAATTCCACTAATTGTTTGGG** | Ins_k3_18_CGT | 0.6996566506108559 | 0.024695094847267934 | 98.84131500486227 |
| **GATCCAGATCTAACTTGGCTAATTGTTTGGG** | Del_k32_19-50 | -3.065753548746799 | 0.05345011374586097 | 2.288978128900039 |
| **GATCCAGATCTAACTTGGCTAGGTGGCTTTGTCTTCTTCTTTTGCCAATTCCACTAATTGTTTGGG** | Ins_k3_18_CTA | -0.43838782211772254 | 0.038141384517192074 | 31.673209984693546 |
| **GATCCAGATCTAACTTGGCTCGGTGGCTTTGTCTTCTTCTTTTGCCAATTCCACTAATTGTTTGGG** | Ins_k3_18_CTC | 0.673327263552761 | 0.033275599000786374 | 96.27284525985974 |
| **GATCCAGATCTAACTTGGCTGGGTGGCTTTGTCTTCTTCTTTTGCCAATTCCACTAATTGTTTGGG** | Ins_k3_17_GCT;Ins_k3_18_CTG | 0.5033157434121753 | 0.029260287630087208 | 81.2210766557604 |
| **GATCCAGATCTAACTTGGCTGGTGGCTTTGTCTTCTTCTTTTGCCAATTCCACTAATTGTTTGGG** | Ins_k2_18_CT | 0.824184522048832 | 0.02912876477024814 | 100 |
| **GATCCAGATCTAACTTGGCTTCTTCTTTTGCCAATTCCACTAATTGTTTGGG** | Del_k11_19-29 | -0.5251804448972447 | 0.03398541916152518 | 29.04012777473138 |
| **GATCCAGATCTAACTTGGCTTCTTTTGCCAATTCCACTAATTGTTTGGG** | Del_k14_19-32 | -2.0885613992329928 | 0.053416492204589584 | 6.08178133360242 |
| **GATCCAGATCTAACTTGGCTTGGTGGCTTTGTCTTCTTCTTTTGCCAATTCCACTAATTGTTTGGG** | Ins_k3_18_CTT | 0.6001670283699896 | 0.028371405148173625 | 89.48097771290448 |
| **GATCCAGATCTAACTTGGCTTTGTCTTCTTCTTTTGCCAATTCCACTAATTGTTTGGG** | Del_k5_16-20;Del_k5_17-21;Del_ | -0.14797956294749232 | 0.031722409882701065 | 42.34623316700179 |
| **GATCCAGATCTAACTTGGCTTTTGCCAATTCCACTAATTGTTTGGG** | Del_k17_19-35 | -2.0423319750462254 | 0.05401680862033424 | 6.369538766083166 |
| **GATCCAGATCTAACTTGGG** | Del_k44_15-58;Del_k44_16-59;D | -1.4512514074462737 | 0.023607245277471867 | 11.502997198333434 |
| **GATCCAGATCTAACTTGGGAAAGTGGCTTTGTCTTCTTCTTTTGCCAATTCCACTAATTGTTTGGG** | Ins_k3_19_AAA | 0.607503226297649 | 0.02885502549142857 | 90.13984169948193 |
| **GATCCAGATCTAACTTGGGAACGTGGCTTTGTCTTCTTCTTTTGCCAATTCCACTAATTGTTTGGG** | Ins_k3_19_AAC | 0.8575013316334661 | 0.028118336001637798 | 100 |
| **GATCCAGATCTAACTTGGGAAGGTGGCTTTGTCTTCTTCTTTTGCCAATTCCACTAATTGTTTGGG** | Ins_k3_18_GAA;Ins_k3_19_AAG | 0.5558118222962076 | 0.029034503105296167 | 85.59876513894032 |
| **GATCCAGATCTAACTTGGGAAGTGGCTTTGTCTTCTTCTTTTGCCAATTCCACTAATTGTTTGGG** | Ins_k2_19_AA | 0.6437907920917465 | 0.031667426410186904 | 93.47086905984689 |
| **GATCCAGATCTAACTTGGGAATGTGGCTTTGTCTTCTTCTTTTGCCAATTCCACTAATTGTTTGGG** | Ins_k3_19_AAT | 0.2903712411715049 | 0.030471643966961743 | 65.64295450648619 |
| **GATCCAGATCTAACTTGGGAATTCCACTAATTGTTTGGG** | Del_k24_20-43 | -0.533446814281518 | 0.02870414337254738 | 28.801060820795918 |
| **GATCCAGATCTAACTTGGGAATTGTTTGGG** | Del_k33_20-52 | -2.754710458195697 | 0.062349960017883246 | 3.124107294458288 |
| **GATCCAGATCTAACTTGGGACAGTGGCTTTGTCTTCTTCTTTTGCCAATTCCACTAATTGTTTGGG** | Ins_k3_19_ACA | 0.5909100759023378 | 0.027168001626474928 | 88.65647861760704 |
| **GATCCAGATCTAACTTGGGACCGTGGCTTTGTCTTCTTCTTTTGCCAATTCCACTAATTGTTTGGG** | Ins_k3_19_ACC | 0.6356400105906315 | 0.02738417714708509 | 92.71210489061647 |
| **GATCCAGATCTAACTTGGGACGGTGGCTTTGTCTTCTTCTTTTGCCAATTCCACTAATTGTTTGGG** | Ins_k3_18_GAC;Ins_k3_19_ACG | 0.7815952431285358 | 0.028540999582125012 | 100 |
| **GATCCAGATCTAACTTGGGACGTGGCTTTGTCTTCTTCTTTTGCCAATTCCACTAATTGTTTGGG** | Ins_k2_19_AC | 0.7981986150429141 | 0.03034029369384951 | 100 |
| **GATCCAGATCTAACTTGGGACTAATTGTTTGGG** | Del_k30_20-49 | -1.9705610993647438 | 0.03802473147932521 | 6.843490684405206 |
| **GATCCAGATCTAACTTGGGACTGTGGCTTTGTCTTCTTCTTTTGCCAATTCCACTAATTGTTTGGG** | Ins_k3_19_ACT | 0.6951015898283588 | 0.029827143780941135 | 98.3921106604352 |
| **GATCCAGATCTAACTTGGGAGAGGTGGCTTTGTCTTCTTCTTTTGCCAATTCCACTAATTGTTTGGG** | Ins_k4_18_GAGA;Ins_k4_19_AG/ | 0.8208004998963863 | 0.027212310096372865 | 100 |
| **GATCCAGATCTAACTTGGGAGAGTGGCTTTGTCTTCTTCTTTTGCCAATTCCACTAATTGTTTGGG** | Ins_k3_19_AGA | 0.9309555822530784 | 0.029224919503031733 | 100 |
| **GATCCAGATCTAACTTGGGAGCGTGGCTTTGTCTTCTTCTTTTGCCAATTCCACTAATTGTTTGGG** | Ins_k3_19_AGC | 0.7231538561282824 | 0.02728684994112931 | 100 |
| **GATCCAGATCTAACTTGGGAGGGTGGCTTTGTCTTCTTCTTTTGCCAATTCCACTAATTGTTTGGG** | Ins_k3_17_GGA;Ins_k3_18_GAG | -0.764251652209993 | 0.047365939973993465 | 22.8650005826108 |
| **GATCCAGATCTAACTTGGGAGGTGGCTTTGTCTTCTTCTTTTGCCAATTCCACTAATTGTTTGGG** | Ins_k2_18_GA;Ins_k2_19_AG | 0.6924041505264422 | 0.032038142080638446 | 98.12706155179454 |
| **GATCCAGATCTAACTTGGGAGTGGCTTTGTCTTCTTCTTTTGCCAATTCCACTAATTGTTTGGG** | Ins_k1_19_A | 0.5189468933940997 | 0.032061024035314745 | 82.50062987785451 |
| **GATCCAGATCTAACTTGGGAGTGTGGCTTTGTCTTCTTCTTTTGCCAATTCCACTAATTGTTTGGG** | Ins_k3_19_AGT | 0.4851412002314739 | 0.028199666164621804 | 79.7582540231288 |
| **GATCCAGATCTAACTTGGGATAGTGGCTTTGTCTTCTTCTTTTGCCAATTCCACTAATTGTTTGGG** | Ins_k3_19_ATA | 0.33482455012691714 | 0.035663240027037885 | 68.62683128926344 |
| **GATCCAGATCTAACTTGGGATCGTGGCTTTGTCTTCTTCTTTTGCCAATTCCACTAATTGTTTGGG** | Ins_k3_19_ATC | 0.746772609993894 | 0.026327988680669122 | 100 |
| **GATCCAGATCTAACTTGGGATGGCTTTGTCTTCTTCTTTTGCCAATTCCACTAATTGTTTGGG** | G20A | 0.09755336755184774 | 0.02699484418141696 | 54.13129055813059 |
| **GATCCAGATCTAACTTGGGATGGTGGCTTTGTCTTCTTCTTTTGCCAATTCCACTAATTGTTTGGG** | Ins_k3_18_GAT;Ins_k3_19_ATG | 0.5825109256485701 | 0.027847820285200674 | 87.91495796367634 |
| **GATCCAGATCTAACTTGGGATGTGGCTTTGTCTTCTTCTTTTGCCAATTCCACTAATTGTTTGGG** | Ins_k2_19_AT | 0.6291383414702463 | 0.02923846201765585 | 92.11127677048165 |
| **GATCCAGATCTAACTTGGGATTCCACTAATTGTTTGGG** | Del_k25_20-44 | -0.5781569043982226 | 0.0274471865624967 | 27.541725079808447 |
| **GATCCAGATCTAACTTGGGATTGTGGCTTTGTCTTCTTCTTTTGCCAATTCCACTAATTGTTTGGG** | Ins_k3_19_ATT | 0.5288422255680196 | 0.027889092506049075 | 83.32105350271748 |
| **GATCCAGATCTAACTTGGGATTGTTTGGG** | Del_k34_20-53 | -2.7055984294980706 | 0.058644065684870066 | 3.2813686454007174 |
| **GATCCAGATCTAACTTGGGCAAGTGGCTTTGTCTTCTTCTTTTGCCAATTCCACTAATTGTTTGGG** | Ins_k3_19_CAA | 0.28176906471049157 | 0.03089405034637367 | 65.08070398458878 |
| **GATCCAGATCTAACTTGGGCAATTCCACTAATTGTTTGGG** | Del_k23_20-42 | -0.8572229535074318 | 0.03227161757352623 | 20.83503784511645 |
| **GATCCAGATCTAACTTGGGCACAGTGGCTTTGTCTTCTTCTTTTGCCAATTCCACTAATTGTTTGGG** | Ins_k4_19_CACA | 0.6425022264851894 | 0.02897456684022891 | 93.35050327901868 |
| **GATCCAGATCTAACTTGGGCACGTGGCTTTGTCTTCTTCTTTTGCCAATTCCACTAATTGTTTGGG** | Ins_k3_19_CAC | 0.7793407007879136 | 0.027617355618081645 | 100 |
| **GATCCAGATCTAACTTGGGCACTAATTGTTTGGG** | Del_k29_20-48 | -2.4003831113798872 | 0.0459957147361854 | 4.452545358912278 |
| **GATCCAGATCTAACTTGGGCAGCGTGGCTTTGTCTTCTTCTTTTGCCAATTCCACTAATTGTTTGGG** | Ins_k4_19_CAGC | 0.9907373095420196 | 0.025989991974253095 | 100 |
| **GATCCAGATCTAACTTGGGCAGGTGGCTTTGTCTTCTTCTTTTGCCAATTCCACTAATTGTTTGGG** | Ins_k3_18_GCA;Ins_k3_19_CAG | 0.2681636348446761 | 0.02942806187830806 | 64.20124927669568 |
| **GATCCAGATCTAACTTGGGCAGTGGCTTTGTCTTCTTCTTTTGCCAATTCCACTAATTGTTTGGG** | Ins_k2_19_CA | 0.6224794494451071 | 0.029775250684498093 | 91.49995534472663 |
| **GATCCAGATCTAACTTGGGCATCGTGGCTTTGTCTTCTTCTTTTGCCAATTCCACTAATTGTTTGGG** | Ins_k4_19_CATC | 0.721803032146344 | 0.029649467878335934 | 100 |
| **GATCCAGATCTAACTTGGGCATGTGGCTTTGTCTTCTTCTTTTGCCAATTCCACTAATTGTTTGGG** | Ins_k3_19_CAT | 0.3182127813011758 | 0.03177127156117713 | 67.49623483507852 |
| **GATCCAGATCTAACTTGGGCCAATTCCACTAATTGTTTGGG** | Del_k22_19-40;Del_k22_20-41 | -0.3989147183023447 | 0.030719719973735782 | 32.94845321666651 |
| **GATCCAGATCTAACTTGGGCCACTAATTGTTTGGG** | Del_k28_20-47 | -1.5829433412637075 | 0.032658938493714346 | 10.083653965521929 |
| **GATCCAGATCTAACTTGGGCCAGTGGCTTTGTCTTCTTCTTTTGCCAATTCCACTAATTGTTTGGG** | Ins_k3_19_CCA | 0.7342976078335053 | 0.028427800749676446 | 100 |
| **GATCCAGATCTAACTTGGGCCCGTGGCTTTGTCTTCTTCTTTTGCCAATTCCACTAATTGTTTGGG** | Ins_k3_19_CCC | 0.7667717178532825 | 0.028467384899715245 | 100 |
| **GATCCAGATCTAACTTGGGCCGGTGGCTTTGTCTTCTTCTTTTGCCAATTCCACTAATTGTTTGGG** | Ins_k3_18_GCC;Ins_k3_19_CCG | 0.9157374146150872 | 0.02730126326384951 | 100 |
| **GATCCAGATCTAACTTGGGCCGTGGCTTTGTCTTCTTCTTTTGCCAATTCCACTAATTGTTTGGG** | Ins_k2_19_CC | 0.977938307951973 | 0.02874736616811971 | 100 |
| **GATCCAGATCTAACTTGGGCCTGGTGGCTTTGTCTTCTTCTTTTGCCAATTCCACTAATTGTTTGGG** | Ins_k4_18_GCCT;Ins_k4_19_CCT | 0.8571844857196567 | 0.028392580843824178 | 100 |

| | | | | |
|---|---|---|---|---|
| **GATCCAGATCTAACTTGGGCCTGTGGCTTTGTCTTCTTCTTTTGCCAATTCCACTAATTGTTTGGG** | Ins_k3_19_CCT | 0.1911209605770734 | 0.03432661768588978 | 59.440748649615166 |
| **GATCCAGATCTAACTTGGGCGAGTGGCTTTGTCTTCTTCTTTTGCCAATTCCACTAATTGTTTGGG** | Ins_k3_19_CGA | 0.7246238908066298 | 0.028442413054814663 | 100 |
| **GATCCAGATCTAACTTGGGCGCGTGGCTTTGTCTTCTTCTTTTGCCAATTCCACTAATTGTTTGGG** | Ins_k3_19_CGC | 0.7338734723226827 | 0.13658490872402393 | 100 |
| **GATCCAGATCTAACTTGGGCGGGTGGCTTTGTCTTCTTCTTTTGCCAATTCCACTAATTGTTTGGG** | Ins_k3_17_GGC;Ins_k3_18_GCG | -0.46532665383719046 | 0.045718299906081644 | 30.831360837413722 |
| **GATCCAGATCTAACTTGGGCGGTGGCTTTGTCTTCTTCTTTTGCCAATTCCACTAATTGTTTGGG** | Ins_k2_18_GC;Ins_k2_19_CG | 0.8017688758825008 | 0.029907572119315454 | 100 |
| **GATCCAGATCTAACTTGGGCGTGGCTTTGTCTTCTTCTTTTGCCAATTCCACTAATTGTTTGGG** | Ins_k1_19_C | 0.7257041561403108 | 0.028675846628722888 | 100 |
| **GATCCAGATCTAACTTGGGCGTGTGGCTTTGTCTTCTTCTTTTGCCAATTCCACTAATTGTTTGGG** | Ins_k3_19_CGT | 0.8469300988940711 | 0.027590020173816166 | 100 |
| **GATCCAGATCTAACTTGGGCTAATTGTTTGGG** | Del_k31_20-50 | -2.06902868886798 | 0.4240328801636371 | 6.2017427789609085 |
| **GATCCAGATCTAACTTGGGCTAGTGGCTTTGTCTTCTTCTTTTGCCAATTCCACTAATTGTTTGGG** | Ins_k3_19_CTA | 0.12609762375870193 | 0.034314820860735513 | 55.6986917129296 |
| **GATCCAGATCTAACTTGGGCTCGGTGGCTTTGTCTTCTTCTTTTGCCAATTCCACTAATTGTTTGGG** | Ins_k4_18_GCTC;Ins_k4_19_CT( | 0.8586598892140493 | 0.037045564295004774 | 100 |
| **GATCCAGATCTAACTTGGGCTCGTGGCTTTGTCTTCTTCTTTTGCCAATTCCACTAATTGTTTGGG** | Ins_k3_19_CTC | 0.825960779887041 | 0.028176396715157054 | 100 |
| **GATCCAGATCTAACTTGGGCTGGCTTTGTCTTCTTCTTTTGCCAATTCCACTAATTGTTTGGG** | G20C | -0.008065303153512948 | 0.02915095777742211 | 48.705586286266694 |
| **GATCCAGATCTAACTTGGGCTGGTGGCTTTGTCTTCTTCTTTTGCCAATTCCACTAATTGTTTGGG** | Ins_k3_18_GCT;Ins_k3_19_CTG | 0.5473695862656647 | 0.028396442001396738 | 84.87916196356083 |
| **GATCCAGATCTAACTTGGGCTGTGGCTTTGTCTTCTTCTTTTGCCAATTCCACTAATTGTTTGGG** | Ins_k2_19_CT | 0.5460382526981269 | 0.030416148438075895 | 84.76623467467928 |
| **GATCCAGATCTAACTTGGGCTTCTTCTTTTGCCAATTCCACTAATTGTTTGGG** | Del_k10_20-29 | 0.21496469894159254 | 0.02815274557447712 | 60.87507014869613 |
| **GATCCAGATCTAACTTGGGCTTCTTTTGCCAATTCCACTAATTGTTTGGG** | Del_k13_20-32 | -0.3187536849951264 | 0.03177678968715301 | 35.69838143509757 |
| **GATCCAGATCTAACTTGGGCTTGTGGCTTTGTCTTCTTCTTTTGCCAATTCCACTAATTGTTTGGG** | Ins_k3_19_CTT | 0.23627358260389886 | 0.028573307076927738 | 62.18616935630719 |
| **GATCCAGATCTAACTTGGGCTTTGTCTTCTTCTTTTGCCAATTCCACTAATTGTTTGGG** | Del_k4_18-21;Del_k4_19-22;Del_ | 0.10471448147157347 | 0.03979754934630046 | 54.52032218382339 |
| **GATCCAGATCTAACTTGGGCTTTTGCCAATTCCACTAATTGTTTGGG** | Del_k16_20-35 | -0.894445578169806 | 0.03451384887718027 | 20.073759339055513 |
| **GATCCAGATCTAACTTGGGG** | Del_k43_18-60;Del_k43_19-61;D | -5.6833564042794045 | 0.14168657855217034 | 0.16704411003784742 |
| **GATCCAGATCTAACTTGGGGAAATGGCTTTGTCTTCTTCTTTTGCCAATTCCACTAATTGTTTGGG** | Ins_k3_20_AAA | 0.2941133981426096 | 0.03097379845479295 | 65.88906094355531 |
| **GATCCAGATCTAACTTGGGGAACTGGCTTTGTCTTCTTCTTTTGCCAATTCCACTAATTGTTTGGG** | Ins_k3_20_AAC | 0.21841435148458743 | 0.033452152434990466 | 61.08543061589066 |
| **GATCCAGATCTAACTTGGGGAAGTGGCTTTGTCTTCTTCTTTTGCCAATTCCACTAATTGTTTGGG** | Ins_k3_19_GAA;Ins_k3_20_AAG | 0.3929647981416158 | 0.030208306920056178 | 72.73508246092841 |
| **GATCCAGATCTAACTTGGGGAATGGCTTTGTCTTCTTCTTTTGCCAATTCCACTAATTGTTTGGG** | Ins_k2_20_AA | 0.4170852575484101 | 0.03145261615953736 | 74.51081572379505 |
| **GATCCAGATCTAACTTGGGGAATTCCACTAATTGTTTGGG** | Del_k23_21-43 | -0.4484139831855582 | 0.027309433367838008 | 31.35723593014147 |
| **GATCCAGATCTAACTTGGGGAATTGGCTTTGTCTTCTTCTTTTGCCAATTCCACTAATTGTTTGGG** | Ins_k3_20_AAT | -3.1550769096571445 | 0.12245853639763328 | 2.0933844810651085 |
| **GATCCAGATCTAACTTGGGGAATTGTTTGGG** | Del_k32_21-52 | -2.667487675340171 | 0.05847590636310231 | 3.4088376205995665 |
| **GATCCAGATCTAACTTGGGGACATGGCTTTGTCTTCTTCTTTTGCCAATTCCACTAATTGTTTGGG** | Ins_k3_20_ACA | 0.49499406156597114 | 0.027945000408743235 | 80.54798520738429 |
| **GATCCAGATCTAACTTGGGGACCTGGCTTTGTCTTCTTCTTTTGCCAATTCCACTAATTGTTTGGG** | Ins_k3_20_ACC | 0.797863157257729 | 0.02805552678316648 | 100 |
| **GATCCAGATCTAACTTGGGGACGTGGCTTTGTCTTCTTCTTTTGCCAATTCCACTAATTGTTTGGG** | Ins_k3_19_GAC;Ins_k3_20_ACG | 0.5920249253870584 | 0.026369428551327383 | 88.75537236268856 |
| **GATCCAGATCTAACTTGGGGACTAATTGTTTGGG** | Del_k29_21-49 | -1.861965574041454 | 0.0379130659755149 | 7.628516990121825 |
| **GATCCAGATCTAACTTGGGGACTGGCTTTGTCTTCTTCTTTTGCCAATTCCACTAATTGTTTGGG** | Ins_k2_20_AC | 0.6696137776991631 | 0.03262606641674257 | 95.91600039016286 |
| **GATCCAGATCTAACTTGGGGACTTGGCTTTGTCTTCTTCTTTTGCCAATTCCACTAATTGTTTGGG** | Ins_k3_20_ACT | 0.16987727877921632 | 0.03114398397182428 | 58.19132645099576 |
| **GATCCAGATCTAACTTGGGGAGATGGCTTTGTCTTCTTCTTTTGCCAATTCCACTAATTGTTTGGG** | Ins_k3_20_AGA | 0.5327051350558232 | 0.027953949571541355 | 83.64353765335869 |
| **GATCCAGATCTAACTTGGGGAGCTGGCTTTGTCTTCTTCTTTTGCCAATTCCACTAATTGTTTGGG** | Ins_k3_20_AGC | 0.6283093237572082 | 0.028842485392447093 | 92.03494653439887 |
| **GATCCAGATCTAACTTGGGGAGGCTTTGTCTTCTTCTTTTGCCAATTCCACTAATTGTTTGGG** | T21A | 0.05692997109577547 | 0.03064953392071855 | 51.97636031541034 |
| **GATCCAGATCTAACTTGGGGAGGTGGCTTTGTCTTCTTCTTTTGCCAATTCCACTAATTGTTTGGG** | Ins_k3_18_GGA;Ins_k3_19_GAG | 0.29709195603342176 | 0.03154740120449395 | 66.0856078938713 |
| **GATCCAGATCTAACTTGGGGAGTGGCTTTGTCTTCTTCTTTTGCCAATTCCACTAATTGTTTGGG** | Ins_k2_19_GA;Ins_k2_20_AG | 0.7207551852295726 | 0.032922674009596226 | 100 |
| **GATCCAGATCTAACTTGGGGAGTTGGCTTTGTCTTCTTCTTTTGCCAATTCCACTAATTGTTTGGG** | Ins_k3_20_AGT | -1.8625051221855495 | 0.07851606140578188 | 7.62440214811511 |
| **GATCCAGATCTAACTTGGGGATATGGCTTTGTCTTCTTCTTTTGCCAATTCCACTAATTGTTTGGG** | Ins_k3_20_ATA | 0.3962065763264111 | 0.031123748636299395 | 72.97125606891315 |
| **GATCCAGATCTAACTTGGGGATCTGGCTTTGTCTTCTTCTTTTGCCAATTCCACTAATTGTTTGGG** | Ins_k3_20_ATC | -0.7915648491766112 | 0.0501586857067085 | 22.248935963951315 |
| **GATCCAGATCTAACTTGGGGATGATGGCTTTGTCTTCTTCTTTTGCCAATTCCACTAATTGTTTGGG** | Ins_k4_20_ATGA | 0.7112419415118485 | 0.03137286992568259 | 99.99307927191663 |
| **GATCCAGATCTAACTTGGGGATGGCTTTGTCTTCTTCTTTTGCCAATTCCACTAATTGTTTGGG** | Ins_k1_20_A | 0.6628028852645054 | 0.02877066593298768 | 95.26494647460623 |
| **GATCCAGATCTAACTTGGGGATGTGGCTTTGTCTTCTTCTTTTGCCAATTCCACTAATTGTTTGGG** | Ins_k3_19_GAT;Ins_k3_20_ATG | 0.44096431549469206 | 0.030357515785523818 | 76.311477299103 |
| **GATCCAGATCTAACTTGGGGATTCCACTAATTGTTTGGG** | Del_k24_21-44 | -0.5566366539640791 | 0.026672428515566265 | 28.14085347522782 |
| **GATCCAGATCTAACTTGGGGATTGGCTTTGTCTTCTTCTTTTGCCAATTCCACTAATTGTTTGGG** | Ins_k2_20_AT | -2.6331946311055225 | 0.11254624233975458 | 3.5277645687176133 |
| **GATCCAGATCTAACTTGGGGATTGTTTGGG** | Del_k33_21-53 | -3.026945671500602 | 0.06478081840783684 | 2.379554685633133 |
| **GATCCAGATCTAACTTGGGGATTTGGCTTTGTCTTCTTCTTTTGCCAATTCCACTAATTGTTTGGG** | Ins_k3_20_ATT | -0.27128681662853593 | 0.035531043669139825 | 37.43373181686815 |
| **GATCCAGATCTAACTTGGGGCAATGGCTTTGTCTTCTTCTTTTGCCAATTCCACTAATTGTTTGGG** | Ins_k3_20_CAA | 0.11842303600196236 | 0.031145014850773596 | 55.27286333415489 |
| **GATCCAGATCTAACTTGGGGCAATTCCACTAATTGTTTGGG** | Del_k22_21-42 | -0.8365104360987319 | 0.03239711464651999 | 21.271084148672752 |
| **GATCCAGATCTAACTTGGGGCACTAATTGTTTGGG** | Del_k28_21-48 | -2.418593840948203 | 0.04862483762187051 | 4.372195098461007 |
| **GATCCAGATCTAACTTGGGGCACTGGCTTTGTCTTCTTCTTTTGCCAATTCCACTAATTGTTTGGG** | Ins_k3_20_CAC | 0.5138382138490241 | 0.030055443225549663 | 82.0802353423328 |
| **GATCCAGATCTAACTTGGGGCAGTGGCTTTGTCTTCTTCTTTTGCCAATTCCACTAATTGTTTGGG** | Ins_k3_19_GCA;Ins_k3_20_CAG | 0.6461371090470647 | 0.029598642689861537 | 93.69043883416117 |
| **GATCCAGATCTAACTTGGGGCATGGCTTTGTCTTCTTCTTTTGCCAATTCCACTAATTGTTTGGG** | Ins_k2_20_CA | 0.5120105141602371 | 0.02833846795329932 | 81.9303543322028 |
| **GATCCAGATCTAACTTGGGGCATTGGCTTTGTCTTCTTCTTTTGCCAATTCCACTAATTGTTTGGG** | Ins_k3_20_CAT | -0.23578939392033094 | 0.04159706169061288 | 38.78639888139006 |
| **GATCCAGATCTAACTTGGGGCCAATTCCACTAATTGTTTGGG** | Del_k21_20-40;Del_k21_21-41 | -0.36089751702229944 | 0.032093359845425044 | 34.22517614296363 |
| **GATCCAGATCTAACTTGGGGCCACTAATTGTTTGGG** | Del_k27_21-47 | -1.5048868137602978 | 0.03517157578681071 | 10.902283045738214 |
| **GATCCAGATCTAACTTGGGGCCATGGCTTTGTCTTCTTCTTTTGCCAATTCCACTAATTGTTTGGG** | Ins_k3_20_CCA | 0.6915357376651685 | 0.028352420838070297 | 98.04188373961078 |
| **GATCCAGATCTAACTTGGGGCCCTGGCTTTGTCTTCTTCTTTTGCCAATTCCACTAATTGTTTGGG** | Ins_k3_20_CCC | 0.7697031204782729 | 0.031460637958950946 | 100 |
| **GATCCAGATCTAACTTGGGGCCGGTGGCTTTGTCTTCTTCTTTTGCCAATTCCACTAATTGTTTGGG** | Ins_k4_18_GGCC;Ins_k4_19_GC | 1.0985404972474968 | 0.02978756216103087 | 100 |
| **GATCCAGATCTAACTTGGGGCCGTGGCTTTGTCTTCTTCTTTTGCCAATTCCACTAATTGTTTGGG** | Ins_k3_19_GCC;Ins_k3_20_CCG | 0.9199640432638068 | 0.02660839713379323 | 100 |
| **GATCCAGATCTAACTTGGGGCCTGGCTTTGTCTTCTTCTTTTGCCAATTCCACTAATTGTTTGGG** | Ins_k2_20_CC | 0.7581126943995391 | 0.03083010950923308 | 100 |
| **GATCCAGATCTAACTTGGGGCCTTGGCTTTGTCTTCTTCTTTTGCCAATTCCACTAATTGTTTGGG** | Ins_k3_20_CCT | 0.48059931278983314 | 0.03382457384945221 | 79.3968224229599 |
| **GATCCAGATCTAACTTGGGGCGATGGCTTTGTCTTCTTCTTTTGCCAATTCCACTAATTGTTTGGG** | Ins_k3_20_CGA | 0.7389330184397243 | 0.02851448689257883 | 100 |
| **GATCCAGATCTAACTTGGGGCGATTGGCTTTGTCTTCTTCTTTTGCCAATTCCACTAATTGTTTGGG** | Ins_k4_20_CGAT | 0.6915585036598613 | 0.028483877742151675 | 98.04411578602294 |
| **GATCCAGATCTAACTTGGGGCGCCTGGCTTTGTCTTCTTCTTTTGCCAATTCCACTAATTGTTTGGG** | Ins_k4_20_CGCC | 1.172086984089403 | 0.030603675407359237 | 100 |
| **GATCCAGATCTAACTTGGGGCGCTGGCTTTGTCTTCTTCTTTTGCCAATTCCACTAATTGTTTGGG** | Ins_k3_20_CGC | 0.7701774663016177 | 0.029379367556354594 | 100 |
| **GATCCAGATCTAACTTGGGGCGGCTTTGTCTTCTTCTTTTGCCAATTCCACTAATTGTTTGGG** | T21C | 0.18087221940532194 | 0.02538172206659015 | 58.834666894229514 |
| **GATCCAGATCTAACTTGGGGCGGTGGCTTTGTCTTCTTCTTTTGCCAATTCCACTAATTGTTTGGG** | Ins_k3_18_GGC;Ins_k3_19_GCG | 0.8051781544030554 | 0.029902563911513243 | 100 |
| **GATCCAGATCTAACTTGGGGCGTGGCTTTGTCTTCTTCTTTTGCCAATTCCACTAATTGTTTGGG** | Ins_k2_19_GC;Ins_k2_20_CG | 0.8505579141738739 | 0.03291788434518877 | 100 |
| **GATCCAGATCTAACTTGGGGCGTTGGCTTTGTCTTCTTCTTTTGCCAATTCCACTAATTGTTTGGG** | Ins_k3_20_CGT | 0.4537767982937979 | 0.03272257650053774 | 77.29550726138517 |
| **GATCCAGATCTAACTTGGGGCGTTTGGCTTTGTCTTCTTCTTTTGCCAATTCCACTAATTGTTTGGG** | Ins_k4_20_CGTT | 0.7108615672031673 | 0.029711962269106433 | 99.95505170632867 |
| **GATCCAGATCTAACTTGGGGCTAATTGTTTGGG** | Del_k30_21-50 | -2.8791766200732445 | 0.05877574859668164 | 2.758487203339581 |
| **GATCCAGATCTAACTTGGGGCTATGGCTTTGTCTTCTTCTTTTGCCAATTCCACTAATTGTTTGGG** | Ins_k3_20_CTA | 0.29057791735210725 | 0.03176882466843705 | 65.65652274367446 |
| **GATCCAGATCTAACTTGGGGCTCTGGCTTTGTCTTCTTCTTTTGCCAATTCCACTAATTGTTTGGG** | Ins_k3_20_CTC | 0.20486601090755685 | 0.03021264025992472 | 60.263405509699304 |

| | | | | |
|---|---|---|---|---|
| GATCCAGATCTAACTTGGGGCTGGCTTTGTCTTCTTCTTTTGCCAATTCCACTAATTGTTTGGG | Ins_k1_20_C | 0.5902537529422767 | 0.03165032312936409 | 88.5983104257801 |
| GATCCAGATCTAACTTGGGGCTGTGGCTTTGTCTTCTTCTTTTGCCAATTCCACTAATTGTTTGGG | Ins_k3_19_GCT;Ins_k3_20_CTG | 0.6076840833639698 | 0.029149555742120737 | 90.1561456011038 |
| GATCCAGATCTAACTTGGGGCTGTTGGCTTTGTCTTCTTCTTTTGCCAATTCCACTAATTGTTTGGG | Ins_k4_20_CTGT | 0.719474551922668 | 0.03055317634747797 | 100 |
| GATCCAGATCTAACTTGGGGCTTCTTCTTTTGCCAATTCCACTAATTGTTTGGG | Del_k9_21-29 | 0.1842330197299047 | 0.031510493641915174 | 59.032731102524565 |
| GATCCAGATCTAACTTGGGGCTTCTTTTGCCAATTCCACTAATTGTTTGGG | Del_k12_21-32 | -0.2014675356884041 | 0.033227403075032955 | 40.140728778175024 |
| GATCCAGATCTAACTTGGGGCTTGGCTTTGTCTTCTTCTTTTGCCAATTCCACTAATTGTTTGGG | Ins_k2_20_CT | 0.4324279941414788 | 0.032763427832747494 | 75.66283047365006 |
| GATCCAGATCTAACTTGGGGCTTTGGCTTTGTCTTCTTCTTTTGCCAATTCCACTAATTGTTTGGG | Ins_k3_20_CTT | 0.004433435749477521 | 0.030202259798945513 | 49.31816494809714 |
| GATCCAGATCTAACTTGGGGCTTTGTCTTCTTCTTTTGCCAATTCCACTAATTGTTTGGG | Del_k3_19-21;Del_k3_20-22;Del_ | 0.0218641531152034 | 0.03207103713856903 | 50.18535183069925 |
| GATCCAGATCTAACTTGGGGCTTTTGCCAATTCCACTAATTGTTTGGG | Del_k15_21-35 | -0.37340502206030546 | 0.030708632651636925 | 33.79977050726954 |
| GATCCAGATCTAACTTGGGGG | Del_k42_19-60;Del_k42_20-61;D | -6.11985076671535 | 0.21980109282440743 | 0.10796029785313963 |
| GATCCAGATCTAACTTGGGGGAATGGCTTTGTCTTCTTCTTTTGCCAATTCCACTAATTGTTTGGG | Ins_k3_20_GAA | 0.16324940051022419 | 0.032059610300518696 | 57.806916740569896 |
| GATCCAGATCTAACTTGGGGGACTGGCTTTGTCTTCTTCTTTTGCCAATTCCACTAATTGTTTGGG | Ins_k3_20_GAC | 0.4335943643122546 | 0.030678533655534638 | 75.75113282875682 |
| GATCCAGATCTAACTTGGGGGAGTGGCTTTGTCTTCTTCTTTTGCCAATTCCACTAATTGTTTGGG | Ins_k3_19_GGA;Ins_k3_20_GAG | 0.4679979020084102 | 0.03705572429334421 | 78.40258798379152 |
| GATCCAGATCTAACTTGGGGGATGGCTTTGTCTTCTTCTTTTGCCAATTCCACTAATTGTTTGGG | Ins_k2_20_GA | 0.6756825531896828 | 0.03357493484195315 | 96.49986293586083 |
| GATCCAGATCTAACTTGGGGGATTGGCTTTGTCTTCTTCTTTTGCCAATTCCACTAATTGTTTGGG | Ins_k3_20_GAT | -2.3123052940142426 | 0.0938967898972395 | 4.862505017332698 |
| GATCCAGATCTAACTTGGGGGCATGGCTTTGTCTTCTTCTTTTGCCAATTCCACTAATTGTTTGGG | Ins_k3_20_GCA | 0.47073169937659287 | 0.029951530532555445 | 78.61721801631693 |
| GATCCAGATCTAACTTGGGGGCCAATTCCACTAATTGTTTGGG | Del_k20_21-40 | -0.20132602945730377 | 0.02707560847983928 | 40.146409343326255 |
| GATCCAGATCTAACTTGGGGGCCTGGCTTTGTCTTCTTCTTTTGCCAATTCCACTAATTGTTTGGG | Ins_k3_20_GCC | 0.6130499472670656 | 0.030818429832052217 | 90.64121144321932 |
| GATCCAGATCTAACTTGGGGGCGTGGCTTTGTCTTCTTCTTTTGCCAATTCCACTAATTGTTTGGG | Ins_k3_19_GGC;Ins_k3_20_GCG | 0.6883151838603565 | 0.03337201200016879 | 97.72664247610017 |
| GATCCAGATCTAACTTGGGGGCTGGCTTTGTCTTCTTCTTTTGCCAATTCCACTAATTGTTTGGG | Ins_k2_20_GC | 0.5408942429298365 | 0.03343322698060301 | 84.33131590771016 |
| GATCCAGATCTAACTTGGGGGCTGTGGCTTTGTCTTCTTCTTTTGCCAATTCCACTAATTGTTTGGG | Ins_k4_19_GGCT;Ins_k4_20_GC | 0.6136488194322343 | 0.032448164219537896 | 90.69551019915374 |
| GATCCAGATCTAACTTGGGGGCTTGGCTTTGTCTTCTTCTTTTGCCAATTCCACTAATTGTTTGGG | Ins_k3_20_GCT | 0.1702258591666388 | 0.033052326830477576 | 58.21161434188968 |
| GATCCAGATCTAACTTGGGGGCTTTGTCTTCTTCTTTTGCCAATTCCACTAATTGTTTGGG | Del_k2_20-21;Del_k2_21-22 | 0.16229758034355057 | 0.03438747891694888 | 57.751921128561285 |
| GATCCAGATCTAACTTGGGGGGATGGCTTTGTCTTCTTCTTTTGCCAATTCCACTAATTGTTTGGG | Ins_k3_20_GGA | -0.8105103357635723 | 0.05306008854460986 | 21.831386870662765 |
| GATCCAGATCTAACTTGGGGGGCTGGCTTTGTCTTCTTCTTTTGCCAATTCCACTAATTGTTTGGG | Ins_k3_20_GGC | -0.8537086601714037 | 0.05730958129738915 | 20.908387089661936 |
| GATCCAGATCTAACTTGGGGGGCTTTGTCTTCTTCTTTTGCCAATTCCACTAATTGTTTGGG | Del_k1_21-21 | -2.5063650299575437 | 0.10934718226076458 | 4.004801454333607 |
| GATCCAGATCTAACTTGGGGGGGCTTTGTCTTCTTCTTTTGCCAATTCCACTAATTGTTTGGG | T21G | -3.1454066706323403 | 0.1368574202785684 | 2.11372620551803 |
| GATCCAGATCTAACTTGGGGGGGTGGCTTTGTCTTCTTCTTTTGCCAATTCCACTAATTGTTTGGG | Ins_k3_16_GGG;Ins_k3_17_GGG | -3.774598497399937 | 0.2917097050164517 | 1.1266634180110753 |
| GATCCAGATCTAACTTGGGGGGTGGCTTTGTCTTCTTCTTTTGCCAATTCCACTAATTGTTTGGG | Ins_k2_16_GG;Ins_k2_17_GG;Ins | -3.249152432065402 | 0.2001936685327121 | 1.90542789991807 |
| GATCCAGATCTAACTTGGGGGGTTGGCTTTGTCTTCTTCTTTTGCCAATTCCACTAATTGTTTGGG | Ins_k3_20_GGT | -4.277833329526272 | 0.35919806747846156 | 0.6811489360746853 |
| GATCCAGATCTAACTTGGGGGTATGGCTTTGTCTTCTTCTTTTGCCAATTCCACTAATTGTTTGGG | Ins_k3_20_GTA | -0.7456457866724114 | 0.04992734360649213 | 23.294406051679495 |
| GATCCAGATCTAACTTGGGGGTCTGGCTTTGTCTTCTTCTTTTGCCAATTCCACTAATTGTTTGGG | Ins_k3_20_GTC | -0.675725171137435 | 0.05164046134178484 | 24.981457849499776 |
| GATCCAGATCTAACTTGGGGGTCTTCTTCTTTTGCCAATTCCACTAATTGTTTGGG | Del_k7_21-27 | 0.12679929704407644 | 0.03259498974607528 | 55.73778771163974 |
| GATCCAGATCTAACTTGGGGGTGGCTTTGTCTTCTTCTTTTGCCAATTCCACTAATTGTTTGGG | Ins_k1_16_G;Ins_k1_17_G;Ins_k1 | -0.7979326870624996 | 0.04259107454383295 | 22.10770848068195 |
| GATCCAGATCTAACTTGGGGGTGTGGCTTTGTCTTCTTCTTTTGCCAATTCCACTAATTGTTTGGG | Ins_k3_19_GGT;Ins_k3_20_GTG | 0.03510777798275655 | 0.04156569509892787 | 50.85440839225813 |
| GATCCAGATCTAACTTGGGGGTTGGCTTTGTCTTCTTCTTTTGCCAATTCCACTAATTGTTTGGG | Ins_k2_20_GT | -2.750719928200041 | 0.14030596287268826 | 3.1365990461043998 |
| GATCCAGATCTAACTTGGGGGTTTGGCTTTGTCTTCTTCTTTTGCCAATTCCACTAATTGTTTGGG | Ins_k3_20_GTT | -0.5269950284113637 | 0.04776900676770644 | 28.987479819125806 |
| GATCCAGATCTAACTTGGGGGTTTGGG | Del_k36_21-56 | -5.355362739577107 | 0.29118031770984715 | 0.2318873223605649 |
| GATCCAGATCTAACTTGGGGT | Del_k42_22-63 | -6.014477044848083 | 0.297994473662743 | 0.11995747067650978 |
| GATCCAGATCTAACTTGGGGTAAAGGCTTTGTCTTCTTCTTTTGCCAATTCCACTAATTGTTTGGG | Ins_k3_21_AAA | -0.4060820872844446 | 0.03873611296515519 | 32.713143777204735 |
| GATCCAGATCTAACTTGGGGTAACGGCTTTGTCTTCTTCTTTTGCCAATTCCACTAATTGTTTGGG | Ins_k3_21_AAC | 0.10266007033029179 | 0.03339077128297173 | 54.408430002129414 |
| GATCCAGATCTAACTTGGGGTAAGGCTTTGTCTTCTTCTTTTGCCAATTCCACTAATTGTTTGGG | Ins_k2_21_AA | -5.243592498187242 | 0.5730100164502593 | 0.2593093668762388 |
| GATCCAGATCTAACTTGGGGTAAGGGCTTTGTCTTCTTCTTTTGCCAATTCCACTAATTGTTTGGG | Ins_k3_21_AAG | -4.976220804281248 | 0.4320370329898404 | 0.3387944256757678 |
| GATCCAGATCTAACTTGGGGTAATGGCTTTGTCTTCTTCTTTTGCCAATTCCACTAATTGTTTGGG | Ins_k3_20_TAA;Ins_k3_21_AAT | -0.6007350595644194 | 0.040021167954349963 | 26.926851216449634 |
| GATCCAGATCTAACTTGGGGTAATTCCACTAATTGTTTGGG | Del_k22_22-43 | -2.161447795521048 | 0.0529833314944849 | 5.654271285473479 |
| GATCCAGATCTAACTTGGGGTAATTGTTTGGG | Del_k31_21-51;Del_k31_22-52 | -4.094077988978421 | 0.10940035536299625 | 0.8185515081829077 |
| GATCCAGATCTAACTTGGGGTACAGGCTTTGTCTTCTTCTTTTGCCAATTCCACTAATTGTTTGGG | Ins_k3_21_ACA | -0.18782945466125378 | 0.037220149620209524 | 40.69192135089252 |
| GATCCAGATCTAACTTGGGGTACAGGGCTTTGTCTTCTTCTTTTGCCAATTCCACTAATTGTTTGGG | Ins_k4_21_ACAG | -1.6698249887115957 | 0.09313711387457276 | 9.244548681670807 |
| GATCCAGATCTAACTTGGGGTACCGGCTTTGTCTTCTTCTTTTGCCAATTCCACTAATTGTTTGGG | Ins_k3_21_ACC | 0.7154851911214977 | 0.028792998524721624 | 100 |
| GATCCAGATCTAACTTGGGGTACGAGGCTTTGTCTTCTTCTTTTGCCAATTCCACTAATTGTTTGGG | Ins_k4_21_ACGA | 0.5377029887349047 | 0.03057271115372451 | 84.06262220504212 |
| GATCCAGATCTAACTTGGGGTACGGCTTTGTCTTCTTCTTTTGCCAATTCCACTAATTGTTTGGG | Ins_k2_21_AC | 0.24345744788331192 | 0.0319117041519026 | 62.634514918082694 |
| GATCCAGATCTAACTTGGGGTACGGGCTTTGTCTTCTTCTTTTGCCAATTCCACTAATTGTTTGGG | Ins_k3_21_ACG | -0.4478504144976868 | 0.04050021543893158 | 31.3749128670661 |
| GATCCAGATCTAACTTGGGGTACTAATTGTTTGGG | Del_k28_22-49 | -3.8942717707935 | 0.10034290597778421 | 0.9995873491707578 |
| GATCCAGATCTAACTTGGGGTACTGGCTTTGTCTTCTTCTTTTGCCAATTCCACTAATTGTTTGGG | Ins_k3_20_TAC;Ins_k3_21_ACT | 0.0015135523303003717 | 0.03386094499458216 | 49.17437168794074 |
| GATCCAGATCTAACTTGGGGTAGAGGCTTTGTCTTCTTCTTTTGCCAATTCCACTAATTGTTTGGG | Ins_k3_21_AGA | 0.1324680572059561 | 0.033216012136980014 | 56.054649119269186 |
| GATCCAGATCTAACTTGGGGTAGCGGCTTTGTCTTCTTCTTTTGCCAATTCCACTAATTGTTTGGG | Ins_k3_21_AGC | 0.6048637174523157 | 0.030353177077524326 | 89.90223051636222 |
| GATCCAGATCTAACTTGGGGTAGCTTTGTCTTCTTCTTTTGCCAATTCCACTAATTGTTTGGG | G22A | -0.9692276191748778 | 0.03896774363747234 | 18.6273590309636 |
| GATCCAGATCTAACTTGGGGTAGGCGGCTTTGTCTTCTTCTTTTGCCAATTCCACTAATTGTTTGGG | Ins_k4_21_AGGC | 0.2531843817779623 | 0.03859878774911892 | 63.246729361968335 |
| GATCCAGATCTAACTTGGGGTAGGCTTTGTCTTCTTCTTTTGCCAATTCCACTAATTGTTTGGG | Ins_k1_21_A | -3.260877129690536 | 0.1565739464876279 | 1.8832177917781199 |
| GATCCAGATCTAACTTGGGGTAGGGCTTTGTCTTCTTCTTTTGCCAATTCCACTAATTGTTTGGG | Ins_k2_21_AG | -4.694170670773323 | 0.45983077430052927 | 0.4491889588648298 |
| GATCCAGATCTAACTTGGGGTAGGGGCTTTGTCTTCTTCTTTTGCCAATTCCACTAATTGTTTGGG | Ins_k3_21_AGG | -4.488548577289231 | 0.3382470410135953 | 0.5517338291929416 |
| GATCCAGATCTAACTTGGGGTAGTCGGCTTTGTCTTCTTCTTTTGCCAATTCCACTAATTGTTTGGG | Ins_k4_21_AGTC | 0.5565105808683231 | 0.03129904940277989 | 85.65859891209004 |
| GATCCAGATCTAACTTGGGGTAGTGGCTTTGTCTTCTTCTTTTGCCAATTCCACTAATTGTTTGGG | Ins_k3_19_GTA;Ins_k3_20_TAG;I | -0.42283841699182584 | 0.037717529332079384 | 32.169558519890984 |
| GATCCAGATCTAACTTGGGGTATAGGCTTTGTCTTCTTCTTTTGCCAATTCCACTAATTGTTTGGG | Ins_k3_21_ATA | -2.511036886318534 | 0.10320060248230743 | 3.986135234085683 |
| GATCCAGATCTAACTTGGGGTATCGGCTTTGTCTTCTTCTTTTGCCAATTCCACTAATTGTTTGGG | Ins_k3_21_ATC | 0.5095757440167457 | 0.02904981354383146 | 81.73111540055595 |
| GATCCAGATCTAACTTGGGGTATGGCTTTGTCTTCTTCTTTTGCCAATTCCACTAATTGTTTGGG | Ins_k2_20_TA;Ins_k2_21_AT | -0.19944561442408087 | 0.0389448060248364 | 40.22197227755848 |
| GATCCAGATCTAACTTGGGGTATGGGCTTTGTCTTCTTCTTTTGCCAATTCCACTAATTGTTTGGG | Ins_k3_21_ATG | -2.8459049038081146 | 0.1151705037764734 | 2.85181071459342 |
| GATCCAGATCTAACTTGGGGTATTAGGCTTTGTCTTCTTCTTTTGCCAATTCCACTAATTGTTTGGG | Ins_k4_21_ATTA | -2.998235516230067 | 0.16238375754755857 | 2.448862224605941 |
| GATCCAGATCTAACTTGGGGTATTCCACTAATTGTTTGGG | Del_k23_22-44 | -1.4878752698083502 | 0.039274506780661626 | 11.089334216640834 |
| GATCCAGATCTAACTTGGGGTATTGGCTTTGTCTTCTTCTTTTGCCAATTCCACTAATTGTTTGGG | Ins_k3_20_TAT;Ins_k3_21_ATT | -1.0907584401531942 | 0.049413296924036426 | 16.49571408345752 |
| GATCCAGATCTAACTTGGGGTATTGTTTGGG | Del_k32_22-53 | -4.092232759232605 | 0.1107362785476993 | 0.8200633181637029 |
| GATCCAGATCTAACTTGGGGTCAAGGCTTTGTCTTCTTCTTTTGCCAATTCCACTAATTGTTTGGG | Ins_k3_21_CAA | 0.5324309572065469 | 0.030077683461904644 | 83.62060759170016 |
| GATCCAGATCTAACTTGGGGTCAATTCCACTAATTGTTTGGG | Del_k21_22-42 | -0.9050700270274972 | 0.03525455500251154 | 19.861615659952587 |
| GATCCAGATCTAACTTGGGGTCACGGCTTTGTCTTCTTCTTTTGCCAATTCCACTAATTGTTTGGG | Ins_k3_21_CAC | 0.5868911900728262 | 0.03263547016724601 | 88.30089335887622 |
| GATCCAGATCTAACTTGGGGTCACTAATTGTTTGGG | Del_k27_22-48 | -2.6679909762782534 | 0.058059129340622524 | 3.4071223811043785 |

| | | | | |
|---|---|---|---|---|
| **GATCCAGATCTAACTTGGGGTCAGGCTTTGTCTTCTTCTTTTGCCAATTCCACTAATTGTTTGGG** | Ins_k2_21_CA | -0.1899860571958485 | 0.041820773149993 | 40.60425960991179 |
| **GATCCAGATCTAACTTGGGGTCAGGGCTTTGTCTTCTTCTTTTGCCAATTCCACTAATTGTTTGGG** | Ins_k3_21_CAG | -2.2340658158489424 | 0.1059052655629295 | 5.258223431068382 |
| **GATCCAGATCTAACTTGGGGTCATGGCTTTGTCTTCTTCTTTTGCCAATTCCACTAATTGTTTGGG** | Ins_k3_20_TCA;Ins_k3_21_CAT | 0.37525687311315825 | 0.029566104856202875 | 71.45843185025701 |
| **GATCCAGATCTAACTTGGGGTCCAATTCCACTAATTGTTTGGG** | Del_k20_22-41 | -0.8077748757387716 | 0.034977056056159916 | 21.891187510574 |
| **GATCCAGATCTAACTTGGGGTCCACTAATTGTTTGGG** | Del_k26_21-46;Del_k26_22-47 | -1.5734756317623755 | 0.03703052943026588 | 10.179576438513909 |
| **GATCCAGATCTAACTTGGGGTCCAGGCTTTGTCTTCTTCTTTTGCCAATTCCACTAATTGTTTGGG** | Ins_k3_21_CCA | 0.6465225316213492 | 0.02915885910977489 | 93.72655620406022 |
| **GATCCAGATCTAACTTGGGGTCCCGGCTTTGTCTTCTTCTTTTGCCAATTCCACTAATTGTTTGGG** | Ins_k3_21_CCC | 0.955761326586885 | 0.02687142258884978 | 100 |
| **GATCCAGATCTAACTTGGGGTCCGGCTTTGTCTTCTTCTTTTGCCAATTCCACTAATTGTTTGGG** | Ins_k2_21_CC | 1.1092246334591729 | 0.02937717741264309 | 100 |
| **GATCCAGATCTAACTTGGGGTCCGGGCTTTGTCTTCTTCTTTTGCCAATTCCACTAATTGTTTGGG** | Ins_k3_21_CCG | 0.5249686497498443 | 0.03278178351312069 | 82.99892737748976 |
| **GATCCAGATCTAACTTGGGGTCCGTGGCTTTGTCTTCTTCTTTTGCCAATTCCACTAATTGTTTGGG** | Ins_k4_19_GTCC;Ins_k4_20_TC( | 0.8069048994517745 | 0.03282224828334645 | 100 |
| **GATCCAGATCTAACTTGGGGTCCTGGCTTTGTCTTCTTCTTTTGCCAATTCCACTAATTGTTTGGG** | Ins_k3_20_TCC;Ins_k3_21_CCT | 0.26125665149311206 | 0.0360488757313752 | 63.75934020292322 |
| **GATCCAGATCTAACTTGGGGTCCTGGGCTTTGTCTTCTTCTTTTGCCAATTCCACTAATTGTTTGGG** | Ins_k4_21_CCTG | 0.0929524895390933 | 0.043744494427189876 | 53.882811143656156 |
| **GATCCAGATCTAACTTGGGGTCGAGGCTTTGTCTTCTTCTTTTGCCAATTCCACTAATTGTTTGGG** | Ins_k3_21_CGA | 0.5723590426929197 | 0.029220169287648746 | 87.02697059864833 |
| **GATCCAGATCTAACTTGGGGTCGCGGCTTTGTCTTCTTCTTTTGCCAATTCCACTAATTGTTTGGG** | Ins_k3_21_CGC | 0.8837969892058848 | 0.02833102607323364 | 100 |
| **GATCCAGATCTAACTTGGGGTCGCTTTGTCTTCTTCTTTTGCCAATTCCACTAATTGTTTGGG** | G22C | 0.12754597010615776 | 0.031682189778496274 | 55.77942115761604 |
| **GATCCAGATCTAACTTGGGGTCGGCTTTGTCTTCTTCTTTTGCCAATTCCACTAATTGTTTGGG** | Ins_k1_21_C | 0.834889838652809 | 0.03137457737835329 | 100 |
| **GATCCAGATCTAACTTGGGGTCGGGCTTTGTCTTCTTCTTTTGCCAATTCCACTAATTGTTTGGG** | Ins_k2_21_CG | -0.30690149635113273 | 0.05393695181946144 | 36.124002675372076 |
| **GATCCAGATCTAACTTGGGGTCGGGGCTTTGTCTTCTTCTTTTGCCAATTCCACTAATTGTTTGGG** | Ins_k3_21_CGG | -0.8113789556737496 | 0.06236979326556104 | 21.812431926872705 |
| **GATCCAGATCTAACTTGGGGTCGTGGCTTTGTCTTCTTCTTTTGCCAATTCCACTAATTGTTTGGG** | Ins_k3_19_GTC;Ins_k3_20_TCG; | 0.7011522165949202 | 0.030682869948104617 | 98.98924930858605 |
| **GATCCAGATCTAACTTGGGGTCTAATTGTTTGGG** | Del_k29_22-50 | -3.374749199746837 | 0.09378976648715924 | 1.6805310338626163 |
| **GATCCAGATCTAACTTGGGGTCTAGGCTTTGTCTTCTTCTTTTGCCAATTCCACTAATTGTTTGGG** | Ins_k3_21_CTA | -1.1636647891023255 | 0.054732619228361604 | 15.335865656535333 |
| **GATCCAGATCTAACTTGGGGTCTCGGCTTTGTCTTCTTCTTTTGCCAATTCCACTAATTGTTTGGG** | Ins_k3_21_CTC | 0.7682807838329353 | 0.033536150847155985 | 100 |
| **GATCCAGATCTAACTTGGGGTCTGGCTTTGTCTTCTTCTTTTGCCAATTCCACTAATTGTTTGGG** | Ins_k2_20_TC;Ins_k2_21_CT | -1.0124364056373052 | 0.057144622272444975 | 17.839634319629575 |
| **GATCCAGATCTAACTTGGGGTCTGGGCTTTGTCTTCTTCTTTTGCCAATTCCACTAATTGTTTGGG** | Ins_k3_21_CTG | -3.2407197703091413 | 0.19268963341633538 | 1.9215636670146101 |
| **GATCCAGATCTAACTTGGGGTCTTCTTCTTTTGCCAATTCCACTAATTGTTTGGG** | Del_k8_20-27;Del_k8_21-28;Del_ | 0.2577990125558077 | 0.02850930731823472 | 63.53926411669886 |
| **GATCCAGATCTAACTTGGGGTCTTCTTTTGCCAATTCCACTAATTGTTTGGG** | Del_k11_21-31;Del_k11_22-32 | 0.16525531700940232 | 0.03296010755706296 | 57.922988965301776 |
| **GATCCAGATCTAACTTGGGGTCTTGGCTTTGTCTTCTTCTTTTGCCAATTCCACTAATTGTTTGGG** | Ins_k3_20_TCT;Ins_k3_21_CTT | -0.10904702444993508 | 0.03726125900405792 | 44.027393086541764 |
| **GATCCAGATCTAACTTGGGGTCTTTGTCTTCTTCTTTTGCCAATTCCACTAATTGTTTGGG** | Del_k2_22-23 | -0.25663202297843224 | 0.03612294625122668 | 37.98635483009872 |
| **GATCCAGATCTAACTTGGGGTCTTTTGCCAATTCCACTAATTGTTTGGG** | Del_k14_21-34;Del_k14_22-35 | -0.5379957115465014 | 0.030423561114330085 | 28.67034528517298 |
| **GATCCAGATCTAACTTGGGGTG** | Del_k41_22-62;Del_k41_23-63 | -5.683555953485284 | 0.20876724766128854 | 0.16701077984396 |
| **GATCCAGATCTAACTTGGGGTGAAAGCTTTGTCTTCTTCTTTTGCCAATTCCACTAATTGTTTGGG** | Ins_k3_22_AAA | 0.4774925442556772 | 0.03338090382134574 | 79.15053764633676 |
| **GATCCAGATCTAACTTGGGGTGAACGCTTTGTCTTCTTCTTTTGCCAATTCCACTAATTGTTTGGG** | Ins_k3_22_AAC | 0.5957315096433624 | 0.028286069060364495 | 89.08496207722771 |
| **GATCCAGATCTAACTTGGGGTGAAGCTTTGTCTTCTTCTTTTGCCAATTCCACTAATTGTTTGGG** | Ins_k2_22_AA | 0.7783468440793424 | 0.029881028438182577 | 100 |
| **GATCCAGATCTAACTTGGGGTGAAGGCTTTGTCTTCTTCTTTTGCCAATTCCACTAATTGTTTGGG** | Ins_k3_21_GAA;Ins_k3_22_AAG | 0.6490368666896242 | 0.028666159644386045 | 93.96251268367098 |
| **GATCCAGATCTAACTTGGGGTGAATGCTTTGTCTTCTTCTTTTGCCAATTCCACTAATTGTTTGGG** | Ins_k3_22_AAT | 0.23416577754476353 | 0.03276069755472183 | 62.05523107858999 |
| **GATCCAGATCTAACTTGGGGTGAATTCCACTAATTGTTTGGG** | Del_k21_23-43 | -0.4442365355286405 | 0.02946769694584179 | 31.48850313194152 |
| **GATCCAGATCTAACTTGGGGTGAATTGTTTGGG** | Del_k30_23-52 | -2.652406746030544 | 0.060471689858938485 | 3.460635659346459 |
| **GATCCAGATCTAACTTGGGGTGACAGCTTTGTCTTCTTCTTTTGCCAATTCCACTAATTGTTTGGG** | Ins_k3_22_ACA | 0.6744327258126774 | 0.03051907364044364 | 96.37933010359316 |
| **GATCCAGATCTAACTTGGGGTGACCGCTTTGTCTTCTTCTTTTGCCAATTCCACTAATTGTTTGGG** | Ins_k3_22_ACC | 0.7213187472058975 | 0.028908263883655437 | 100 |
| **GATCCAGATCTAACTTGGGGTGACGCTTTGTCTTCTTCTTTTGCCAATTCCACTAATTGTTTGGG** | Ins_k2_22_AC | 0.8211643129483981 | 0.028500990522191554 | 100 |
| **GATCCAGATCTAACTTGGGGTGACGGCTTTGTCTTCTTCTTTTGCCAATTCCACTAATTGTTTGGG** | Ins_k3_21_GAC;Ins_k3_22_ACG | 0.8015574558710694 | 0.028642084528593414 | 100 |
| **GATCCAGATCTAACTTGGGGTGACTAATTGTTTGGG** | Del_k27_23-49 | -2.5600439490226536 | 0.05442515802321569 | 3.7954959437632176 |
| **GATCCAGATCTAACTTGGGGTGACTGCTTTGTCTTCTTCTTTTGCCAATTCCACTAATTGTTTGGG** | Ins_k3_22_ACT | 0.5436846236554802 | 0.02903586945138407 | 84.5669610030545 |
| **GATCCAGATCTAACTTGGGGTGACTTTGTCTTCTTCTTTTGCCAATTCCACTAATTGTTTGGG** | G23A | -0.002774037162532772 | 0.028038318496682313 | 48.96398351992861 |
| **GATCCAGATCTAACTTGGGGTGAGAGCTTTGTCTTCTTCTTTTGCCAATTCCACTAATTGTTTGGG** | Ins_k3_22_AGA | -0.09668446222555427 | 0.03449509762989545 | 44.57506279845055 |
| **GATCCAGATCTAACTTGGGGTGAGCAGCTTTGTCTTCTTCTTTTGCCAATTCCACTAATTGTTTGGG** | Ins_k4_22_AGCA | 0.48262132723448636 | 0.036064910702388764 | 79.55752636284885 |
| **GATCCAGATCTAACTTGGGGTGAGCGCTTTGTCTTCTTCTTTTGCCAATTCCACTAATTGTTTGGG** | Ins_k3_22_AGC | -0.33172101746398264 | 0.042829901301510466 | 35.23845709507973 |
| **GATCCAGATCTAACTTGGGGTGAGCTTTGTCTTCTTCTTTTGCCAATTCCACTAATTGTTTGGG** | Ins_k1_22_A | -0.13859584445214368 | 0.03813630201995923 | 42.74546852496161 |
| **GATCCAGATCTAACTTGGGGTGAGGCTTTGTCTTCTTCTTTTGCCAATTCCACTAATTGTTTGGG** | Ins_k2_21_GA;Ins_k2_22_AG | -0.025715392736759046 | 0.0441615667912544 | 47.85347040712608 |
| **GATCCAGATCTAACTTGGGGTGAGGGCTTTGTCTTCTTCTTTTGCCAATTCCACTAATTGTTTGGG** | Ins_k3_21_GAG;Ins_k3_22_AGG | -1.668935370709498 | 0.0817409085551505 | 9.252776457845554 |
| **GATCCAGATCTAACTTGGGGTGAGTGCTTTGTCTTCTTCTTTTGCCAATTCCACTAATTGTTTGGG** | Ins_k3_22_AGT | -3.8974917564321405 | 0.20999492547506982 | 0.9963738687188427 |
| **GATCCAGATCTAACTTGGGGTGATAGCTTTGTCTTCTTCTTTTGCCAATTCCACTAATTGTTTGGG** | Ins_k3_22_ATA | -0.22844670050560018 | 0.03652477558712263 | 39.07224366807484 |
| **GATCCAGATCTAACTTGGGGTGATATGCTTTGTCTTCTTCTTTTGCCAATTCCACTAATTGTTTGGG** | Ins_k4_22_ATAT | 0.36361578834947517 | 0.03182540737860342 | 70.63140129396828 |
| **GATCCAGATCTAACTTGGGGTGATCAGCTTTGTCTTCTTCTTTTGCCAATTCCACTAATTGTTTGGG** | Ins_k4_22_ATCA | 0.9161523639436135 | 0.02740952076435448 | 100 |
| **GATCCAGATCTAACTTGGGGTGATCGCTTTGTCTTCTTCTTTTGCCAATTCCACTAATTGTTTGGG** | Ins_k3_22_ATC | 0.7101771564931207 | 0.029283562927242993 | 99.88666480345145 |
| **GATCCAGATCTAACTTGGGGTGATGCTTTGTCTTCTTCTTTTGCCAATTCCACTAATTGTTTGGG** | Ins_k2_22_AT | 0.48228839647582644 | 0.03009065621132913 | 79.53104362394355 |
| **GATCCAGATCTAACTTGGGGTGATGGCTTTGTCTTCTTCTTTTGCCAATTCCACTAATTGTTTGGG** | Ins_k3_20_TGA;Ins_k3_21_GAT;I | 0.6058269282859421 | 0.029257515134710265 | 89.98886703668197 |
| **GATCCAGATCTAACTTGGGGTGATTCCACTAATTGTTTGGG** | Del_k22_23-44 | -0.5309333716282277 | 0.029963863625690295 | 28.873541685613993 |
| **GATCCAGATCTAACTTGGGGTGATTGCTTTGTCTTCTTCTTTTGCCAATTCCACTAATTGTTTGGG** | Ins_k3_22_ATT | 0.014634684056572866 | 0.031809633703991765 | 49.82384670202467 |
| **GATCCAGATCTAACTTGGGGTGATTGTTTGGG** | Del_k31_23-53 | -3.0163032827492815 | 0.06721970380831262 | 2.405014065667924 |
| **GATCCAGATCTAACTTGGGGTGCAAGCTTTGTCTTCTTCTTTTGCCAATTCCACTAATTGTTTGGG** | Ins_k3_22_CAA | 0.5186554196944861 | 0.028920601527186632 | 82.47658661820269 |
| **GATCCAGATCTAACTTGGGGTGCAATTCCACTAATTGTTTGGG** | Del_k20_23-42 | -0.5542783770929999 | 0.033561620684615885 | 28.207295712914046 |
| **GATCCAGATCTAACTTGGGGTGCACGCTTTGTCTTCTTCTTTTGCCAATTCCACTAATTGTTTGGG** | Ins_k3_22_CAC | 0.5899432031711618 | 0.032424737166658175 | 88.57080051257593 |
| **GATCCAGATCTAACTTGGGGTGCACTAATTGTTTGGG** | Del_k26_23-48 | -1.8162306757574527 | 0.04216554473695288 | 7.985507681486088 |
| **GATCCAGATCTAACTTGGGGTGCAGCTTTGTCTTCTTCTTTTGCCAATTCCACTAATTGTTTGGG** | Ins_k2_22_CA | 0.8444637836703817 | 0.03040721733465578 | 100 |
| **GATCCAGATCTAACTTGGGGTGCAGGCTTTGTCTTCTTCTTTTGCCAATTCCACTAATTGTTTGGG** | Ins_k3_21_GCA;Ins_k3_22_CAG | 0.35494721210067226 | 0.03353192047827384 | 70.02177372531678 |
| **GATCCAGATCTAACTTGGGGTGCATGCTTTGTCTTCTTCTTTTGCCAATTCCACTAATTGTTTGGG** | Ins_k3_22_CAT | -0.09033629916457786 | 0.04030998211107193 | 44.85893263825459 |
| **GATCCAGATCTAACTTGGGGTGCCAATTCCACTAATTGTTTGGG** | Del_k19_21-39;Del_k19_22-40;D | -0.1498777185144845 | 0.029729847311562706 | 42.26592966715342 |
| **GATCCAGATCTAACTTGGGGTGCCACTAATTGTTTGGG** | Del_k25_23-47 | -0.867465661381029 | 0.03164760549658333 | 20.622719850770817 |
| **GATCCAGATCTAACTTGGGGTGCCAGCTTTGTCTTCTTCTTTTGCCAATTCCACTAATTGTTTGGG** | Ins_k3_22_CCA | 0.7471577474053219 | 0.028612033642858786 | 100 |
| **GATCCAGATCTAACTTGGGGTGCCCGCTTTGTCTTCTTCTTTTGCCAATTCCACTAATTGTTTGGG** | Ins_k3_22_CCC | 0.7538990333958477 | 0.028789327023080808 | 100 |
| **GATCCAGATCTAACTTGGGGTGCCGCTTTGTCTTCTTCTTTTGCCAATTCCACTAATTGTTTGGG** | Ins_k2_22_CC | 0.9151332638595393 | 0.03076639686853427 | 100 |
| **GATCCAGATCTAACTTGGGGTGCCGGCTTTGTCTTCTTCTTTTGCCAATTCCACTAATTGTTTGGG** | Ins_k3_21_GCC;Ins_k3_22_CCG | 0.9096225469159418 | 0.02880502522975384 | 100 |
| **GATCCAGATCTAACTTGGGGTGCCTGCTTTGTCTTCTTCTTTTGCCAATTCCACTAATTGTTTGGG** | Ins_k3_22_CCT | 0.6441700279554412 | 0.0284427329612501 | 93.50632328793758 |
| **GATCCAGATCTAACTTGGGGTGCCTTTGTCTTCTTCTTTTGCCAATTCCACTAATTGTTTGGG** | G23C | -0.1081110511165132 | 0.030271522933267604 | 44.06862084344159 |

| | | | | |
|---|---|---|---|---|
| **GATCCAGATCTAACTTGGGGTGCGAGCTTTGTCTTCTTCTTTTGCCAATTCCACTAATTGTTTGGG** | Ins_k3_22_CGA | 0.5932623588063746 | 0.030341140571347132 | 88.86526920758506 |
| **GATCCAGATCTAACTTGGGGTGCGCGCTTTGTCTTCTTCTTTTGCCAATTCCACTAATTGTTTGGG** | Ins_k3_22_CGC | 0.623295530274274 | 0.030919044604237608 | 91.57465718137561 |
| **GATCCAGATCTAACTTGGGGTGCGCTTTGTCTTCTTCTTTTGCCAATTCCACTAATTGTTTGGG** | Ins_k1_22_C | 0.4716647554493918 | 0.032981054504993716 | 78.69060652148453 |
| **GATCCAGATCTAACTTGGGGTGCGGCTTTGTCTTCTTCTTTTGCCAATTCCACTAATTGTTTGGG** | Ins_k2_21_GC;Ins_k2_22_CG | 0.7907924316723522 | 0.03144623953576928 | 100 |
| **GATCCAGATCTAACTTGGGGTGCGGGCTTTGTCTTCTTCTTTTGCCAATTCCACTAATTGTTTGGG** | Ins_k3_21_GCG;Ins_k3_22_CGG | -0.5114815971078236 | 0.04929278294429215 | 29.440681362298005 |
| **GATCCAGATCTAACTTGGGGTGCGTCGCTTTGTCTTCTTCTTTTGCCAATTCCACTAATTGTTTGGG** | Ins_k4_22_CGTC | 0.8381970986384824 | 0.030489774609471554 | 100 |
| **GATCCAGATCTAACTTGGGGTGCGTGCTTTGTCTTCTTCTTTTGCCAATTCCACTAATTGTTTGGG** | Ins_k3_22_CGT | 0.16052352881510512 | 0.0341422670541227 | 57.64955707101605 |
| **GATCCAGATCTAACTTGGGGTGCTAATTGTTTGGG** | Del_k28_23-50 | -2.6241877643113596 | 0.05297608805135971 | 3.5596821974116524 |
| **GATCCAGATCTAACTTGGGGTGCTAGCTTTGTCTTCTTCTTTTGCCAATTCCACTAATTGTTTGGG** | Ins_k3_22_CTA | -0.5499099062130305 | 0.04598363006096984 | 28.33078800268539 |
| **GATCCAGATCTAACTTGGGGTGCTATGCTTTGTCTTCTTCTTTTGCCAATTCCACTAATTGTTTGGG** | Ins_k4_22_CTAT | 0.02530511683545067 | 0.036365368293074504 | 50.358335249790905 |
| **GATCCAGATCTAACTTGGGGTGCTCGCTTTGTCTTCTTCTTTTGCCAATTCCACTAATTGTTTGGG** | Ins_k3_22_CTC | 0.03997313564855176 | 0.166120301735635 | 51.10243616060107 |
| **GATCCAGATCTAACTTGGGGTGCTGCTTTGTCTTCTTCTTTTGCCAATTCCACTAATTGTTTGGG** | Ins_k2_22_CT | 0.7689812475121625 | 0.031099109360144154 | 100 |
| **GATCCAGATCTAACTTGGGGTGCTGGCTTTGTCTTCTTCTTTTGCCAATTCCACTAATTGTTTGGG** | Ins_k3_20_TGC;Ins_k3_21_GCT; | 0.7341169134399788 | 0.028884449789869093 | 100 |
| **GATCCAGATCTAACTTGGGGTGCTTCTTCTTTTGCCAATTCCACTAATTGTTTGGG** | Del_k7_23-29 | 0.22059590835842346 | 0.028373173274782476 | 61.21883742129963 |
| **GATCCAGATCTAACTTGGGGTGCTTCTTTTGCCAATTCCACTAATTGTTTGGG** | Del_k10_23-32 | 0.28028005337636375 | 0.031306545448439034 | 64.98387018992288 |
| **GATCCAGATCTAACTTGGGGTGCTTGCTTTGTCTTCTTCTTTTGCCAATTCCACTAATTGTTTGGG** | Ins_k3_22_CTT | 0.09547681220077298 | 0.03277815793269584 | 54.019000565604244 |
| **GATCCAGATCTAACTTGGGGTGCTTTGTCTTCTTCTTTTGCCAATTCCACTAATTGTTTGGG** | Del_k1_22-22;Del_k1_23-23 | 0.0792404605535878 | 0.033315934051270195 | 53.149010917791685 |
| **GATCCAGATCTAACTTGGGGTGCTTTTGCCAATTCCACTAATTGTTTGGG** | Del_k13_23-35 | -0.036196231074644425 | 0.02988932798228461 | 47.354545064962224 |
| **GATCCAGATCTAACTTGGGGTGG** | Del_k40_22-61;Del_k40_23-62;D | -6.11610158378963 | 0.4425627612739813 | 0.10836582047273229 |
| **GATCCAGATCTAACTTGGGGTGGAAACTTTGTCTTCTTCTTTTGCCAATTCCACTAATTGTTTGGG** | Ins_k3_23_AAA | 0.5152604755519059 | 0.031262416453925405 | 82.19705797410865 |
| **GATCCAGATCTAACTTGGGGTGGAACCTTTGTCTTCTTCTTTTGCCAATTCCACTAATTGTTTGGG** | Ins_k3_23_AAC | 0.7398766052550005 | 0.031215371861233788 | 100 |
| **GATCCAGATCTAACTTGGGGTGGAACTTTGTCTTCTTCTTTTGCCAATTCCACTAATTGTTTGGG** | Ins_k2_23_AA | 0.3747001489535756 | 0.03797219076095914 | 71.41866028676246 |
| **GATCCAGATCTAACTTGGGGTGGAAGCTTTGTCTTCTTCTTTTGCCAATTCCACTAATTGTTTGGG** | Ins_k3_22_GAA;Ins_k3_23_AAG | 0.6090519382069448 | 0.028921112679257302 | 90.2795505022429 |
| **GATCCAGATCTAACTTGGGGTGGAATCTTTGTCTTCTTCTTTTGCCAATTCCACTAATTGTTTGGG** | Ins_k3_23_AAT | 0.4397002795469158 | 0.03055506149273947 | 76.21507778762843 |
| **GATCCAGATCTAACTTGGGGTGGAATTCCACTAATTGTTTGGG** | Del_k20_24-43 | -0.47623505365635854 | 0.03255116569301419 | 30.49686773553629 |
| **GATCCAGATCTAACTTGGGGTGGAATTGTTTGGG** | Del_k29_24-52 | -2.6341865912403413 | 0.05719400219446304 | 3.524266901960218 |
| **GATCCAGATCTAACTTGGGGTGGACACTTTGTCTTCTTCTTTTGCCAATTCCACTAATTGTTTGGG** | Ins_k3_23_ACA | 0.4973750509067498 | 0.02997624533138773 | 80.73999760060343 |
| **GATCCAGATCTAACTTGGGGTGGACAGCTTTGTCTTCTTCTTTTGCCAATTCCACTAATTGTTTGGG** | Ins_k4_22_GACA;Ins_k4_23_AC/ | 0.9481840363184839 | 0.031524493749777 | 100 |
| **GATCCAGATCTAACTTGGGGTGGACCACTTTGTCTTCTTCTTTTGCCAATTCCACTAATTGTTTGGG** | Ins_k4_23_ACCA | 1.0657738075339542 | 0.02833970651202214 | 100 |
| **GATCCAGATCTAACTTGGGGTGGACCCTTTGTCTTCTTCTTTTGCCAATTCCACTAATTGTTTGGG** | Ins_k3_23_ACC | 0.7887488651066472 | 0.028996890287699407 | 100 |
| **GATCCAGATCTAACTTGGGGTGGACCTTTGTCTTCTTCTTTTGCCAATTCCACTAATTGTTTGGG** | Ins_k2_23_AC | 0.8757914888876817 | 0.03204219000446402 | 100 |
| **GATCCAGATCTAACTTGGGGTGGACGCTTTGTCTTCTTCTTTTGCCAATTCCACTAATTGTTTGGG** | Ins_k3_22_GAC;Ins_k3_23_ACG | 0.7644242799027781 | 0.028496837173263047 | 100 |
| **GATCCAGATCTAACTTGGGGTGGACTAATTGTTTGGG** | Del_k26_24-49 | -1.610213149811095 | 0.035229835641630215 | 9.812390119071102 |
| **GATCCAGATCTAACTTGGGGTGGACTCTTTGTCTTCTTCTTTTGCCAATTCCACTAATTGTTTGGG** | Ins_k3_23_ACT | 0.46022510575840525 | 0.030451595745163526 | 77.79554291701452 |
| **GATCCAGATCTAACTTGGGGTGGACTTTGTCTTCTTCTTTTGCCAATTCCACTAATTGTTTGGG** | Ins_k1_23_A | 1.2613707072827827 | 0.2884346935880907 | 100 |
| **GATCCAGATCTAACTTGGGGTGGAGACTTTGTCTTCTTCTTTTGCCAATTCCACTAATTGTTTGGG** | Ins_k3_23_AGA | 0.6590299631143448 | 0.02927742950843303 | 94.90619644145693 |
| **GATCCAGATCTAACTTGGGGTGGAGCCTTTGTCTTCTTCTTTTGCCAATTCCACTAATTGTTTGGG** | Ins_k3_23_AGC | 0.8174840822330061 | 0.02688514096523372 | 100 |
| **GATCCAGATCTAACTTGGGGTGGAGCTTTGTCTTCTTCTTTTGCCAATTCCACTAATTGTTTGGG** | Ins_k2_22_GA;Ins_k2_23_AG | 0.6106812378900519 | 0.030447361810753013 | 90.42676283920821 |
| **GATCCAGATCTAACTTGGGGTGGAGGACTTTGTCTTCTTCTTTTGCCAATTCCACTAATTGTTTGGG** | Ins_k4_23_AGGA | 0.9879149560935807 | 0.030635656638179465 | 100 |
| **GATCCAGATCTAACTTGGGGTGGAGGCTTTGTCTTCTTCTTTTGCCAATTCCACTAATTGTTTGGG** | Ins_k3_21_GGA;Ins_k3_22_GAG | 0.6331447812466451 | 0.030555967735858135 | 92.48105530668101 |
| **GATCCAGATCTAACTTGGGGTGGAGTCTTTGTCTTCTTCTTTTGCCAATTCCACTAATTGTTTGGG** | Ins_k3_23_AGT | 0.7177926759193314 | 0.02998237125559111 | 100 |
| **GATCCAGATCTAACTTGGGGTGGATACTTTGTCTTCTTCTTTTGCCAATTCCACTAATTGTTTGGG** | Ins_k3_23_ATA | 0.36889272942556095 | 0.03010164938816735 | 71.00510417357529 |
| **GATCCAGATCTAACTTGGGGTGGATCCTTTGTCTTCTTCTTTTGCCAATTCCACTAATTGTTTGGG** | Ins_k3_23_ATC | 0.28156899347131203 | 0.03209518468097616 | 65.06768450995054 |
| **GATCCAGATCTAACTTGGGGTGGATCTTTGTCTTCTTCTTTTGCCAATTCCACTAATTGTTTGGG** | Ins_k2_23_AT | 0.6673305729564565 | 0.030180373927985636 | 95.69725433920844 |
| **GATCCAGATCTAACTTGGGGTGGATGCTTTGTCTTCTTCTTTTGCCAATTCCACTAATTGTTTGGG** | Ins_k3_22_GAT;Ins_k3_23_ATG | 0.5773081642485172 | 0.029431410061667083 | 87.45874522599757 |
| **GATCCAGATCTAACTTGGGGTGGATTCCACTAATTGTTTGGG** | Del_k21_24-44 | -0.5649565743823959 | 0.02986630361221437 | 27.907695088376794 |
| **GATCCAGATCTAACTTGGGGTGGATTCTTTGTCTTCTTCTTTTGCCAATTCCACTAATTGTTTGGG** | Ins_k3_23_ATT | 0.45976016985400325 | 0.028138911927237862 | 77.75938138295913 |
| **GATCCAGATCTAACTTGGGGTGGATTGTTTGGG** | Del_k30_24-53 | -2.6835443053060475 | 0.06938587453771439 | 3.3545402591848488 |
| **GATCCAGATCTAACTTGGGGTGGATTTGTCTTCTTCTTTTGCCAATTCCACTAATTGTTTGGG** | C24A | 0.31512024294819274 | 0.02698987131439814 | 67.28782256773505 |
| **GATCCAGATCTAACTTGGGGTGGC** | Del_k39_25-63 | -6.1194304521088085 | 0.677181579507975 | 0.10800568468095903 |
| **GATCCAGATCTAACTTGGGGTGGCAAATTTGTCTTCTTCTTTTGCCAATTCCACTAATTGTTTGGG** | Ins_k3_24_AAA | -0.12142480777069098 | 0.036606290420302064 | 43.48579039002363 |
| **GATCCAGATCTAACTTGGGGTGGCAACTTTGTCTTCTTCTTTTGCCAATTCCACTAATTGTTTGGG** | Ins_k3_23_CAA;Ins_k3_24_AAC | 0.38134935836935724 | 0.03244282725913264 | 71.89512020064123 |
| **GATCCAGATCTAACTTGGGGTGGCAAGTTTGTCTTCTTCTTTTGCCAATTCCACTAATTGTTTGGG** | Ins_k3_24_AAG | 0.314658071456444 | 0.03163722776278516 | 67.25673123971855 |
| **GATCCAGATCTAACTTGGGGTGGCAATTCCACTAATTGTTTGGG** | Del_k19_24-42;Del_k19_25-43 | -0.6047543963529391 | 0.03267836671102257 | 26.818840344202762 |
| **GATCCAGATCTAACTTGGGGTGGCAATTGTTTGGG** | Del_k28_25-52 | -2.8975782500808247 | 0.07480093085728051 | 2.708190730261791 |
| **GATCCAGATCTAACTTGGGGTGGCAATTTGTCTTCTTCTTTTGCCAATTCCACTAATTGTTTGGG** | Ins_k2_24_AA | 0.06544680910791367 | 0.0350565470605094 | 52.42092501216955 |
| **GATCCAGATCTAACTTGGGGTGGCAATTTTGTCTTCTTCTTTTGCCAATTCCACTAATTGTTTGGG** | Ins_k3_24_AAT | 0.07620231173411529 | 0.040825422120882394 | 52.987781356733024 |
| **GATCCAGATCTAACTTGGGGTGGCACATTTGTCTTCTTCTTTTGCCAATTCCACTAATTGTTTGGG** | Ins_k3_24_ACA | 0.30309137846600354 | 0.035630040695930495 | 66.48327506624679 |
| **GATCCAGATCTAACTTGGGGTGGCACCTTTGTCTTCTTCTTTTGCCAATTCCACTAATTGTTTGGG** | Ins_k3_23_CAC;Ins_k3_24_ACC | 0.6683914005639362 | 0.028791550063969114 | 95.79882649431923 |
| **GATCCAGATCTAACTTGGGGTGGCACGTTTGTCTTCTTCTTTTGCCAATTCCACTAATTGTTTGGG** | Ins_k3_24_ACG | 0.6238255655868077 | 0.029498930525743063 | 91.62320784906242 |
| **GATCCAGATCTAACTTGGGGTGGCACTAATTGTTTGGG** | Del_k25_24-48;Del_k25_25-49 | -2.2254134729347346 | 0.05172824640550952 | 5.303916775494417 |
| **GATCCAGATCTAACTTGGGGTGGCACTTTGTCTTCTTCTTTTGCCAATTCCACTAATTGTTTGGG** | Ins_k2_23_CA;Ins_k2_24_AC | 0.3615199583106652 | 0.033735922992292985 | 70.483524897479 |
| **GATCCAGATCTAACTTGGGGTGGCACTTTTGTCTTCTTCTTTTGCCAATTCCACTAATTGTTTGGG** | Ins_k3_24_ACT | 0.2497846133350029 | 0.03833951163654432 | 63.03207022979499 |
| **GATCCAGATCTAACTTGGGGTGGCAGATTTGTCTTCTTCTTTTGCCAATTCCACTAATTGTTTGGG** | Ins_k3_24_AGA | 0.6985490701546615 | 0.02923835606739636 | 98.73190089974572 |
| **GATCCAGATCTAACTTGGGGTGGCAGCTTTGTCTTCTTCTTTTGCCAATTCCACTAATTGTTTGGG** | Ins_k3_22_GCA;Ins_k3_23_CAG; | 0.5781201311773296 | 0.030773842049494186 | 87.5297876729121 |
| **GATCCAGATCTAACTTGGGGTGGCAGGTTTGTCTTCTTCTTTTGCCAATTCCACTAATTGTTTGGG** | Ins_k3_24_AGG | -1.489157966513682 | 0.06737052108857401 | 11.075119082978409 |
| **GATCCAGATCTAACTTGGGGTGGCAGGTTTTGTCTTCTTCTTTTGCCAATTCCACTAATTGTTTGGG** | Ins_k4_24_AGGT | -0.008341942540310954 | 0.040024922233352196 | 48.692114266274416 |
| **GATCCAGATCTAACTTGGGGTGGCAGTTTGTCTTCTTCTTTTGCCAATTCCACTAATTGTTTGGG** | Ins_k2_24_AG | 0.7322683273080177 | 0.034209150961923784 | 100 |
| **GATCCAGATCTAACTTGGGGTGGCAGTTTTGTCTTCTTCTTTTGCCAATTCCACTAATTGTTTGGG** | Ins_k3_24_AGT | 0.5955438424796471 | 0.02883811076761749 | 89.06824532370692 |
| **GATCCAGATCTAACTTGGGGTGGCATAATTTGTCTTCTTCTTTTGCCAATTCCACTAATTGTTTGGG** | Ins_k4_24_ATAA | -0.12937021466122278 | 0.04213879837662459 | 43.14164708119126 |
| **GATCCAGATCTAACTTGGGGTGGCATATTTGTCTTCTTCTTTTGCCAATTCCACTAATTGTTTGGG** | Ins_k3_24_ATA | 0.049577037597067886 | 0.03742432753093383 | 51.59558322439822 |
| **GATCCAGATCTAACTTGGGGTGGCATCTTTGTCTTCTTCTTTTGCCAATTCCACTAATTGTTTGGG** | Ins_k3_23_CAT;Ins_k3_24_ATC | 0.23163062914883537 | 0.03309980639894585 | 61.8981111044379 |
| **GATCCAGATCTAACTTGGGGTGGCATGTTTGTCTTCTTCTTTTGCCAATTCCACTAATTGTTTGGG** | Ins_k3_24_ATG | 0.12303561590851142 | 0.03319097267056394 | 55.52840272781479 |
| **GATCCAGATCTAACTTGGGGTGGCATTCCACTAATTGTTTGGG** | Del_k20_25-44 | -0.7492444603670703 | 0.0340633669551505 | 23.21072774116203 |
| **GATCCAGATCTAACTTGGGGTGGCATTGTCTTCTTCTTTTGCCAATTCCACTAATTGTTTGGG** | T25A | -0.13583263551022173 | 0.030094154858238097 | 42.863746523971116 |

| | | | | |
|---|---|---|---|---|
| GATCCAGATCTAACTTGGGGTGGCATTGTTTGGG | Del_k29_25-53 | -2.4535686689528147 | 0.05729397050779571 | 4.221921540860733 |
| GATCCAGATCTAACTTGGGGTGGCATTTGTCTTCTTCTTTTGCCAATTCCACTAATTGTTTGGG | Ins_k1_24_A | 0.19808235914923555 | 0.03447035205232785 | 59.85598302148409 |
| GATCCAGATCTAACTTGGGGTGGCATTTTGTCTTCTTCTTTTGCCAATTCCACTAATTGTTTGGG | Ins_k2_24_AT | 0.15809875982730126 | 0.04035950611529503 | 57.509939551392314 |
| GATCCAGATCTAACTTGGGGTGGCATTTTTGTCTTCTTCTTTTGCCAATTCCACTAATTGTTTGGG | Ins_k3_24_ATT | -0.37434967354087534 | 0.037683973617395054 | 33.76785658015942 |
| GATCCAGATCTAACTTGGGGTGGCCAATTCCACTAATTGTTTGGG | Del_k18_23-40;Del_k18_24-41;D | -0.33030427766194354 | 0.029408446141269112 | 35.288416200979384 |
| GATCCAGATCTAACTTGGGGTGGCCAATTTGTCTTCTTCTTTTGCCAATTCCACTAATTGTTTGGG | Ins_k3_24_CAA | 0.5269703422828604 | 0.03047910227704425 | 83.1652321006021 |
| GATCCAGATCTAACTTGGGGTGGCCACTAATTGTTTGGG | Del_k24_24-47;Del_k24_25-48 | -1.2853998252363803 | 0.03851789030944605 | 13.578113698366067 |
| GATCCAGATCTAACTTGGGGTGGCCACTTTGTCTTCTTCTTTTGCCAATTCCACTAATTGTTTGGG | Ins_k3_23_CCA;Ins_k3_24_CAC | 0.46132285205980716 | 0.029293824769962576 | 77.88098957731428 |
| GATCCAGATCTAACTTGGGGTGGCCAGTTTGTCTTCTTCTTTTGCCAATTCCACTAATTGTTTGGG | Ins_k3_24_CAG | 0.6422008706003728 | 0.029112574235502092 | 93.32237579390942 |
| GATCCAGATCTAACTTGGGGTGGCCATTTGTCTTCTTCTTTTGCCAATTCCACTAATTGTTTGGG | Ins_k2_24_CA | 0.670752039284886 | 0.033622993286720464 | 96.02524004872205 |
| GATCCAGATCTAACTTGGGGTGGCCATTTTGTCTTCTTCTTTTGCCAATTCCACTAATTGTTTGGG | Ins_k3_24_CAT | 0.4150307616982112 | 0.03160449471824203 | 74.35789070779173 |
| GATCCAGATCTAACTTGGGGTGGCCCAATTCCACTAATTGTTTGGG | Del_k17_25-41 | -0.14536329586628233 | 0.029242925777911252 | 42.457167276201595 |
| GATCCAGATCTAACTTGGGGTGGCCCACTAATTGTTTGGG | Del_k23_25-47 | -0.8838138624051883 | 0.03423410037891097 | 20.288316376388682 |
| GATCCAGATCTAACTTGGGGTGGCCCAGTTTGTCTTCTTCTTTTGCCAATTCCACTAATTGTTTGGG | Ins_k4_24_CCAG | 0.9839824036282163 | 0.030180241194094428 | 100 |
| GATCCAGATCTAACTTGGGGTGGCCCATTTGTCTTCTTCTTTTGCCAATTCCACTAATTGTTTGGG | Ins_k3_24_CCA | 0.7181940433410725 | 0.031647073833828344 | 100 |
| GATCCAGATCTAACTTGGGGTGGCCCCTTTGTCTTCTTCTTTTGCCAATTCCACTAATTGTTTGGG | Ins_k3_23_CCC;Ins_k3_24_CCC | 0.4357074948025208 | 0.03561233190013962 | 75.91137410294323 |
| GATCCAGATCTAACTTGGGGTGGCCCGTTTGTCTTCTTCTTTTGCCAATTCCACTAATTGTTTGGG | Ins_k3_24_CCG | 0.9311950197895829 | 0.03315491764796676 | 100 |
| GATCCAGATCTAACTTGGGGTGGCCCTTTGTCTTCTTCTTTTGCCAATTCCACTAATTGTTTGGG | Ins_k2_23_CC;Ins_k2_24_CC | 0.669361686823031 | 0.03736510457030012 | 95.89182388905512 |
| GATCCAGATCTAACTTGGGGTGGCCCTTTTGTCTTCTTCTTTTGCCAATTCCACTAATTGTTTGGG | Ins_k3_24_CCT | 0.2794752688186667 | 0.033961889450344035 | 64.93159321336793 |
| GATCCAGATCTAACTTGGGGTGGCCGATTTGTCTTCTTCTTTTGCCAATTCCACTAATTGTTTGGG | Ins_k3_24_CGA | 0.857272888437445 | 0.0283866450333923 | 100 |
| GATCCAGATCTAACTTGGGGTGGCCGCTTTGTCTTCTTCTTTTGCCAATTCCACTAATTGTTTGGG | Ins_k3_22_GCC;Ins_k3_23_CCG | 0.9215045075266532 | 0.029980124653098305 | 100 |
| GATCCAGATCTAACTTGGGGTGGCCGGTTTGTCTTCTTCTTTTGCCAATTCCACTAATTGTTTGGG | Ins_k3_24_CGG | 0.6725506504481502 | 0.030199537557458744 | 96.19810753151874 |
| GATCCAGATCTAACTTGGGGTGGCCGTTTGTCTTCTTCTTTTGCCAATTCCACTAATTGTTTGGG | Ins_k2_24_CG | 1.1434208606719678 | 0.029330239043225053 | 100 |
| GATCCAGATCTAACTTGGGGTGGCCGTTTTGTCTTCTTCTTTTGCCAATTCCACTAATTGTTTGGG | Ins_k3_24_CGT | 0.8006368695243451 | 0.030617003538154056 | 100 |
| GATCCAGATCTAACTTGGGGTGGCCTAATTGTTTGGG | Del_k26_25-50 | -1.5499535932809723 | 0.039877293464197786 | 10.421859147960312 |
| GATCCAGATCTAACTTGGGGTGGCCTATTTGTCTTCTTCTTTTGCCAATTCCACTAATTGTTTGGG | Ins_k3_24_CTA | 0.2616022689889378 | 0.03471902102794833 | 63.781380354931336 |
| GATCCAGATCTAACTTGGGGTGGCCTCTTTGTCTTCTTCTTTTGCCAATTCCACTAATTGTTTGGG | Ins_k3_23_CCT;Ins_k3_24_CTC | 0.4149540286727271 | 0.03479574134929512 | 74.35218522077155 |
| GATCCAGATCTAACTTGGGGTGGCCTGGTTTGTCTTCTTCTTTTGCCAATTCCACTAATTGTTTGGG | Ins_k4_24_CTGG | 0.919179718504088 | 0.033259039232813155 | 100 |
| GATCCAGATCTAACTTGGGGTGGCCTGTTTGTCTTCTTCTTTTGCCAATTCCACTAATTGTTTGGG | Ins_k3_24_CTG | 0.8004884772555028 | 0.030051274912308034 | 100 |
| GATCCAGATCTAACTTGGGGTGGCCTGTTTTGTCTTCTTCTTTTGCCAATTCCACTAATTGTTTGGG | Ins_k4_24_CTGT | 0.7309639234252199 | 0.033790437603362346 | 100 |
| GATCCAGATCTAACTTGGGGTGGCCTTCTTCTTTTGCCAATTCCACTAATTGTTTGGG | Del_k5_25-29 | 0.2575707513322064 | 0.03132791121497853 | 63.52476222169797 |
| GATCCAGATCTAACTTGGGGTGGCCTTCTTTTGCCAATTCCACTAATTGTTTGGG | Del_k8_25-32 | 0.24523237082042748 | 0.029656667436274087 | 62.74578507403024 |
| GATCCAGATCTAACTTGGGGTGGCCTTGTCTTCTTCTTTTGCCAATTCCACTAATTGTTTGGG | T25C | -0.11973175135456399 | 0.02443409806006265 | 43.5594766463469 |
| GATCCAGATCTAACTTGGGGTGGCCTTTGTCTTCTTCTTTTGCCAATTCCACTAATTGTTTGGG | Ins_k1_23_C;Ins_k1_24_C | 0.6497285892626998 | 0.03285059202841425 | 94.02753115949837 |
| GATCCAGATCTAACTTGGGGTGGCCTTTTGCCAATTCCACTAATTGTTTGGG | Del_k11_25-35 | -0.4610578970728335 | 0.033340997547829224 | 30.963253726591773 |
| GATCCAGATCTAACTTGGGGTGGCCTTTTGTCTTCTTCTTTTGCCAATTCCACTAATTGTTTGGG | Ins_k2_24_CT | 0.4002275290331241 | 0.03521850658181497 | 73.26526073181456 |
| GATCCAGATCTAACTTGGGGTGGCCTTTTTGTCTTCTTCTTTTGCCAATTCCACTAATTGTTTGGG | Ins_k3_24_CTT | 0.03397807806358355 | 0.038981805841576955 | 50.79699060974642 |
| GATCCAGATCTAACTTGGGGTGGCG | Del_k38_25-62 | -5.731361786711214 | 0.21037600064522152 | 0.15921452824571727 |
| GATCCAGATCTAACTTGGGGTGGCGAATTTGTCTTCTTCTTTTGCCAATTCCACTAATTGTTTGGG | Ins_k3_24_GAA | 0.8676510897265928 | 0.028242718549062875 | 100 |
| GATCCAGATCTAACTTGGGGTGGCGACTTTGTCTTCTTCTTTTGCCAATTCCACTAATTGTTTGGG | Ins_k3_23_CGA;Ins_k3_24_GAC | 0.6515938192080182 | 0.028812892482120094 | 94.20307779282496 |
| GATCCAGATCTAACTTGGGGTGGCGAGGTTTGTCTTCTTCTTTTGCCAATTCCACTAATTGTTTGGG | Ins_k4_24_GAGG | 0.3549537642441173 | 0.034660948105473786 | 70.02223251952555 |
| GATCCAGATCTAACTTGGGGTGGCGAGTTTGTCTTCTTCTTTTGCCAATTCCACTAATTGTTTGGG | Ins_k3_24_GAG | 0.7221390669794476 | 0.02914873094149933 | 100 |
| GATCCAGATCTAACTTGGGGTGGCGATTTGTCTTCTTCTTTTGCCAATTCCACTAATTGTTTGGG | Ins_k2_24_GA | 0.9011455099404757 | 0.031313226416358836 | 100 |
| GATCCAGATCTAACTTGGGGTGGCGATTTTGTCTTCTTCTTTTGCCAATTCCACTAATTGTTTGGG | Ins_k3_24_GAT | 0.7796016286472656 | 0.02912356437437655 | 100 |
| GATCCAGATCTAACTTGGGGTGGCGCATTTGTCTTCTTCTTTTGCCAATTCCACTAATTGTTTGGG | Ins_k3_24_GCA | 0.7086834011816587 | 0.02743372336714128 | 99.73756995069418 |
| GATCCAGATCTAACTTGGGGTGGCGCCAATTCCACTAATTGTTTGGG | Del_k16_25-40 | 0.2115418225243887 | 0.02564036961341608 | 60.6670585088469 |
| GATCCAGATCTAACTTGGGGTGGCGCCTTTGTCTTCTTCTTTTGCCAATTCCACTAATTGTTTGGG | Ins_k3_23_CGC;Ins_k3_24_GCC | 1.0233023536168906 | 0.1293496772371782 | 100 |
| GATCCAGATCTAACTTGGGGTGGCGCGTTTGTCTTCTTCTTTTGCCAATTCCACTAATTGTTTGGG | Ins_k3_24_GCG | 0.9587997154316144 | 0.030512477313611085 | 100 |
| GATCCAGATCTAACTTGGGGTGGCGCTTTGTCTTCTTCTTTTGCCAATTCCACTAATTGTTTGGG | Ins_k2_22_GC;Ins_k2_23_CG;Ins | 0.8398328643774704 | 0.03065410287717327 | 100 |
| GATCCAGATCTAACTTGGGGTGGCGCTTTTGTCTTCTTCTTTTGCCAATTCCACTAATTGTTTGGG | Ins_k3_24_GCT | 0.7778445534986975 | 0.029472974516915905 | 100 |
| GATCCAGATCTAACTTGGGGTGGCGG | Del_k37_25-61 | -5.140153458644567 | 0.18909154458922908 | 0.2875684316768059 |
| GATCCAGATCTAACTTGGGGTGGCGGATTTGTCTTCTTCTTTTGCCAATTCCACTAATTGTTTGGG | Ins_k3_24_GGA | 0.8394292517370459 | 0.030833577698720402 | 100 |
| GATCCAGATCTAACTTGGGGTGGCGGCTTTGTCTTCTTCTTTTGCCAATTCCACTAATTGTTTGGG | Ins_k3_21_GGC;Ins_k3_22_GCG | 0.8027976765762728 | 0.0293394530891954 | 100 |
| GATCCAGATCTAACTTGGGGTGGCGGG | Del_k36_25-60 | -5.63784143812752 | 0.2639482324990865 | 0.17482279757991487 |
| GATCCAGATCTAACTTGGGGTGGCGGGGTTTGTCTTCTTCTTTTGCCAATTCCACTAATTGTTTGGG | Ins_k4_24_GGGG | -3.248639282493943 | 0.23965199941780554 | 1.9064059203422785 |
| GATCCAGATCTAACTTGGGGTGGCGGGTTTGTCTTCTTCTTTTGCCAATTCCACTAATTGTTTGGG | Ins_k3_24_GGG | -2.132663878567871 | 0.10417750522648299 | 5.819388318073613 |
| GATCCAGATCTAACTTGGGGTGGCGGTTTGTCTTCTTCTTTTGCCAATTCCACTAATTGTTTGGG | Ins_k2_24_GG | 0.8104062584510372 | 0.031011680009325318 | 100 |
| GATCCAGATCTAACTTGGGGTGGCGGTTTTGTCTTCTTCTTTTGCCAATTCCACTAATTGTTTGGG | Ins_k3_24_GGT | 0.8486099617487082 | 0.0320165004873038 | 100 |
| GATCCAGATCTAACTTGGGGTGGCGTATTTGTCTTCTTCTTTTGCCAATTCCACTAATTGTTTGGG | Ins_k3_24_GTA | 0.6443376692619811 | 0.029855984781109787 | 93.52200012414924 |
| GATCCAGATCTAACTTGGGGTGGCGTCTTCTTCTTTTGCCAATTCCACTAATTGTTTGGG | Del_k3_25-27 | 0.5548486507534831 | 0.02695850956125105 | 85.51635853648405 |
| GATCCAGATCTAACTTGGGGTGGCGTCTTTGTCTTCTTCTTTTGCCAATTCCACTAATTGTTTGGG | Ins_k3_23_CGT;Ins_k3_24_GTC | 0.7279716436973285 | 0.02919970243728186 | 100 |
| GATCCAGATCTAACTTGGGGTGGCGTGTTTGTCTTCTTCTTTTGCCAATTCCACTAATTGTTTGGG | Ins_k3_24_GTG | 0.7480440757954294 | 0.02734507469818888 | 100 |
| GATCCAGATCTAACTTGGGGTGGCGTTGTCTTCTTCTTTTGCCAATTCCACTAATTGTTTGGG | T25G | 0.2955045081751688 | 0.02762632849124599 | 65.98078366067408 |
| GATCCAGATCTAACTTGGGGTGGCGTTTGGG | Del_k32_25-56 | -2.388915060997139 | 0.052872840739184745 | 4.50390128676038 |
| GATCCAGATCTAACTTGGGGTGGCGTTTGTCTTCTTCTTTTGCCAATTCCACTAATTGTTTGGG | Ins_k1_24_G | 0.867809090232001 | 0.028451227751132936 | 100 |
| GATCCAGATCTAACTTGGGGTGGCGTTTTGTCTTCTTCTTTTGCCAATTCCACTAATTGTTTGGG | Ins_k2_24_GT | 0.8189353081404178 | 0.030155515402517243 | 100 |
| GATCCAGATCTAACTTGGGGTGGCGTTTTTGTCTTCTTCTTTTGCCAATTCCACTAATTGTTTGGG | Ins_k3_24_GTT | 0.8605817726169338 | 0.030682070404614716 | 100 |
| GATCCAGATCTAACTTGGGGTGGCT | Del_k38_26-63 | -5.919372564750763 | 0.7726628269867702 | 0.1319260777973677 |
| GATCCAGATCTAACTTGGGGTGGCTAAATTGTCTTCTTCTTTTGCCAATTCCACTAATTGTTTGGG | Ins_k3_25_AAA | -0.7101112239506862 | 0.0482852210089984 | 24.13704532849164 |
| GATCCAGATCTAACTTGGGGTGGCTAACCTTGTCTTCTTCTTTTGCCAATTCCACTAATTGTTTGGG | Ins_k4_25_AACC | -0.4132496720921684 | 0.04088894510986296 | 32.47950784776235 |
| GATCCAGATCTAACTTGGGGTGGCTAACTTGTCTTCTTCTTTTGCCAATTCCACTAATTGTTTGGG | Ins_k3_25_AAC | -0.7500389857285684 | 0.04320783161283263 | 23.192293553497404 |
| GATCCAGATCTAACTTGGGGTGGCTAAGTTGTCTTCTTCTTTTGCCAATTCCACTAATTGTTTGGG | Ins_k3_25_AAG | -0.2623518163729544 | 0.03620046481130365 | 37.76970092711224 |
| GATCCAGATCTAACTTGGGGTGGCTAAGTTTGTCTTCTTCTTTTGCCAATTCCACTAATTGTTTGGG | Ins_k4_24_TAAG;Ins_k4_25_AAG | -0.4748458200040994 | 0.045751546113441684 | 30.53926445313771 |
| GATCCAGATCTAACTTGGGGTGGCTAATTCCACTAATTGTTTGGG | Del_k18_26-43 | -1.3264184327606645 | 0.04997806746090098 | 13.032426553985221 |
| GATCCAGATCTAACTTGGGGTGGCTAATTGTCTTCTTCTTTTGCCAATTCCACTAATTGTTTGGG | Ins_k2_25_AA | -0.37959047262910933 | 0.04140865102961891 | 33.59134895268505 |

| | | | | |
|---|---|---|---|---|
| GATCCAGATCTAACTTGGGGTGGCTAATTGTTTGGG | Del_k27_24-50;Del_k27_25-51;D | -3.5871187294916296 | 0.10455865694569592 | 1.3589880002730188 |
| GATCCAGATCTAACTTGGGGTGGCTAATTTGTCTTCTTCTTTTGCCAATTCCACTAATTGTTTGGG | Ins_k3_24_TAA;Ins_k3_25_AAT | -0.7747916684706646 | 0.04808465596754187 | 22.625268712924655 |
| GATCCAGATCTAACTTGGGGTGGCTACATTGTCTTCTTCTTTTGCCAATTCCACTAATTGTTTGGG | Ins_k3_25_ACA | 0.3943438988239426 | 0.030767132183401383 | 72.83546066269336 |
| GATCCAGATCTAACTTGGGGTGGCTACCTTGTCTTCTTCTTTTGCCAATTCCACTAATTGTTTGGG | Ins_k3_25_ACC | 0.2260621678504372 | 0.032851978666366875 | 61.55439175043256 |
| GATCCAGATCTAACTTGGGGTGGCTACGTTGTCTTCTTCTTTTGCCAATTCCACTAATTGTTTGGG | Ins_k3_25_ACG | 0.6809678334543046 | 0.028014642112138777 | 97.01124195728671 |
| GATCCAGATCTAACTTGGGGTGGCTACTAATTGTTTGGG | Del_k24_26-49 | -1.70605823884768 | 0.04340578267661752 | 8.9155843495953 |
| GATCCAGATCTAACTTGGGGTGGCTACTTGTCTTCTTCTTTTGCCAATTCCACTAATTGTTTGGG | Ins_k2_25_AC | 0.32019154122774396 | 0.03474429612279992 | 67.62992590741047 |
| GATCCAGATCTAACTTGGGGTGGCTACTTTGTCTTCTTCTTTTGCCAATTCCACTAATTGTTTGGG | Ins_k3_23_CTA;Ins_k3_24_TAC;I | 0.18249821949140999 | 0.0331931407847821 | 58.93040988562623 |
| GATCCAGATCTAACTTGGGGTGGCTAGATTGTCTTCTTCTTTTGCCAATTCCACTAATTGTTTGGG | Ins_k3_25_AGA | -0.1347976622883545 | 0.03846320921038023 | 42.90813231890486 |
| GATCCAGATCTAACTTGGGGTGGCTAGCTTGTCTTCTTCTTTTGCCAATTCCACTAATTGTTTGGG | Ins_k3_25_AGC | -0.6015961096269707 | 0.036595094699167816 | 26.90367582854187 |
| GATCCAGATCTAACTTGGGGTGGCTAGGTTGTCTTCTTCTTTTGCCAATTCCACTAATTGTTTGGG | Ins_k3_25_AGG | -2.2674047853018093 | 0.08050590728188826 | 5.085809698148911 |
| GATCCAGATCTAACTTGGGGTGGCTAGTTGTCTTCTTCTTTTGCCAATTCCACTAATTGTTTGGG | Ins_k2_25_AG | -0.39042159411630384 | 0.041981811756006515 | 33.22948023096116 |
| GATCCAGATCTAACTTGGGGTGGCTAGTTTGTCTTCTTCTTTTGCCAATTCCACTAATTGTTTGGG | Ins_k3_24_TAG;Ins_k3_25_AGT | -1.1993791123547806 | 0.05487310045911772 | 14.797820737260476 |
| GATCCAGATCTAACTTGGGGTGGCTATATTGTCTTCTTCTTTTGCCAATTCCACTAATTGTTTGGG | Ins_k3_25_ATA | -0.24109227572139424 | 0.03757692725792782 | 38.581263577477756 |
| GATCCAGATCTAACTTGGGGTGGCTATCTTGTCTTCTTCTTTTGCCAATTCCACTAATTGTTTGGG | Ins_k3_25_ATC | -0.4605473888655759 | 0.04610377099470929 | 30.97906475722966 |
| GATCCAGATCTAACTTGGGGTGGCTATGGTTGTCTTCTTCTTTTGCCAATTCCACTAATTGTTTGGG | Ins_k4_25_ATGG | 0.2703914678246363 | 0.03290519593250397 | 64.34443837866735 |
| GATCCAGATCTAACTTGGGGTGGCTATGTCTTCTTCTTTTGCCAATTCCACTAATTGTTTGGG | T26A | -0.09008290581077016 | 0.03240966199041123 | 44.87030103392115 |
| GATCCAGATCTAACTTGGGGTGGCTATGTTGTCTTCTTCTTTTGCCAATTCCACTAATTGTTTGGG | Ins_k3_25_ATG | 0.14018726802447345 | 0.032991825468154753 | 56.48902112082584 |
| GATCCAGATCTAACTTGGGGTGGCTATGTTTGTCTTCTTCTTTTGCCAATTCCACTAATTGTTTGGG | Ins_k4_24_TATG;Ins_k4_25_ATG | -0.13861896860850548 | 0.039517137131656735 | 42.74448008349215 |
| GATCCAGATCTAACTTGGGGTGGCTATTCCACTAATTGTTTGGG | Del_k19_26-44 | -0.8782618991131818 | 0.03558040746612242 | 20.4012696301845 |
| GATCCAGATCTAACTTGGGGTGGCTATTGTCTTCTTCTTTTGCCAATTCCACTAATTGTTTGGG | Ins_k1_25_A | 0.025223969408675218 | 0.03818134090640748 | 50.35424896626663 |
| GATCCAGATCTAACTTGGGGTGGCTATTGTTTGGG | Del_k28_26-53 | -3.4625471554664466 | 0.08235687455273875 | 1.5392755433343082 |
| GATCCAGATCTAACTTGGGGTGGCTATTTGTCTTCTTCTTTTGCCAATTCCACTAATTGTTTGGG | Ins_k2_24_TA;Ins_k2_25_AT | 0.07246243802957053 | 0.03851607310535328 | 52.78998384597321 |
| GATCCAGATCTAACTTGGGGTGGCTATTTTGTCTTCTTCTTTTGCCAATTCCACTAATTGTTTGGG | Ins_k3_24_TAT;Ins_k3_25_ATT | -0.33115071351238523 | 0.03853542300172102 | 35.258559458092925 |
| GATCCAGATCTAACTTGGGGTGGCTCAATTCCACTAATTGTTTGGG | Del_k17_26-42 | -0.27933090454198994 | 0.02974235823860157 | 37.13381946507607 |
| GATCCAGATCTAACTTGGGGTGGCTCAATTGTCTTCTTCTTTTGCCAATTCCACTAATTGTTTGGG | Ins_k3_25_CAA | 0.3206432224072686 | 0.031794452522600256 | 67.66047997194397 |
| GATCCAGATCTAACTTGGGGTGGCTCACTAATTGTTTGGG | Del_k23_26-48 | -1.3993816105343133 | 0.04328885278024408 | 12.115400649646508 |
| GATCCAGATCTAACTTGGGGTGGCTCACTTGTCTTCTTCTTTTGCCAATTCCACTAATTGTTTGGG | Ins_k3_25_CAC | 0.264727193885543 | 0.035191422898639205 | 63.98100412053344 |
| GATCCAGATCTAACTTGGGGTGGCTCAGTTGTCTTCTTCTTTTGCCAATTCCACTAATTGTTTGGG | Ins_k3_25_CAG | 0.49204181719443096 | 0.02974691547928994 | 80.31053854416479 |
| GATCCAGATCTAACTTGGGGTGGCTCATTGTCTTCTTCTTTTGCCAATTCCACTAATTGTTTGGG | Ins_k2_25_CA | 0.5247285597481668 | 0.03451843804270965 | 82.97900255684729 |
| GATCCAGATCTAACTTGGGGTGGCTCATTTGTCTTCTTCTTTTGCCAATTCCACTAATTGTTTGGG | Ins_k3_24_TCA;Ins_k3_25_CAT | 0.36023004389621593 | 0.03269155408844256 | 70.39266579556339 |
| GATCCAGATCTAACTTGGGGTGGCTCCAATTCCACTAATTGTTTGGG | Del_k16_26-41 | -0.16109197784615714 | 0.02959640121304379 | 41.794596337614 |
| GATCCAGATCTAACTTGGGGTGGCTCCACTAATTGTTTGGG | Del_k22_25-46;Del_k22_26-47 | -0.6098873677265542 | 0.030393927246359177 | 26.68153270401042 |
| GATCCAGATCTAACTTGGGGTGGCTCCATTGTCTTCTTCTTTTGCCAATTCCACTAATTGTTTGGG | Ins_k3_25_CCA | 0.5756154560988274 | 0.02994357476037583 | 87.3108283205859 |
| GATCCAGATCTAACTTGGGGTGGCTCCCATTGTCTTCTTCTTTTGCCAATTCCACTAATTGTTTGGG | Ins_k4_25_CCCA | 0.27524670618236147 | 0.03600296538012526 | 64.6576055995418 |
| GATCCAGATCTAACTTGGGGTGGCTCCCTTGTCTTCTTCTTTTGCCAATTCCACTAATTGTTTGGG | Ins_k3_25_CCC | 0.21499674559595172 | 0.03190736604326671 | 60.87702102228759 |
| GATCCAGATCTAACTTGGGGTGGCTCCCTTTGTCTTCTTCTTTTGCCAATTCCACTAATTGTTTGGG | Ins_k4_23_CTCC;Ins_k4_24_TC( | 0.18681033554370308 | 0.03635507153060056 | 59.18507332660995 |
| GATCCAGATCTAACTTGGGGTGGCTCCGTTGTCTTCTTCTTTTGCCAATTCCACTAATTGTTTGGG | Ins_k3_25_CCG | 0.8678706145097406 | 0.028063221302225418 | 100 |
| GATCCAGATCTAACTTGGGGTGGCTCCTTGTCTTCTTCTTTTGCCAATTCCACTAATTGTTTGGG | Ins_k2_25_CC | 0.6381868930199333 | 0.032372461813073676 | 92.9485326706241 |
| GATCCAGATCTAACTTGGGGTGGCTCCTTTGTCTTCTTCTTTTGCCAATTCCACTAATTGTTTGGG | Ins_k3_23_CTC;Ins_k3_24_TCC; | 0.3798970955055532 | 0.029871888332456388 | 71.79078536662496 |
| GATCCAGATCTAACTTGGGGTGGCTCCTTTTGTCTTCTTCTTTTGCCAATTCCACTAATTGTTTGGG | Ins_k4_24_TCCT;Ins_k4_25_CCT | 0.4152771031241258 | 0.036919905883721654 | 74.37621039297274 |
| GATCCAGATCTAACTTGGGGTGGCTCGATTGTCTTCTTCTTTTGCCAATTCCACTAATTGTTTGGG | Ins_k3_25_CGA | 0.7043516249263622 | 0.026430066577637513 | 99.30646351584657 |
| GATCCAGATCTAACTTGGGGTGGCTCGCTTGTCTTCTTCTTTTGCCAATTCCACTAATTGTTTGGG | Ins_k3_25_CGC | 0.7086448038398523 | 0.03114405050407391 | 99.73372041990716 |
| GATCCAGATCTAACTTGGGGTGGCTCGGTTGTCTTCTTCTTTTGCCAATTCCACTAATTGTTTGGG | Ins_k3_25_CGG | 0.8515118706938647 | 0.03228622887274075 | 100 |
| GATCCAGATCTAACTTGGGGTGGCTCGTTGTCTTCTTCTTTTGCCAATTCCACTAATTGTTTGGG | Ins_k2_25_CG | 0.8531096054754808 | 0.03160164016746948 | 100 |
| GATCCAGATCTAACTTGGGGTGGCTCGTTTGTCTTCTTCTTTTGCCAATTCCACTAATTGTTTGGG | Ins_k3_24_TCG;Ins_k3_25_CGT | 0.5171347376983371 | 0.029869760737843348 | 82.35126127197836 |
| GATCCAGATCTAACTTGGGGTGGCTCTAATTGTTTGGG | Del_k25_26-50 | -2.248506950889232 | 0.058791243327421826 | 5.182834378345397 |
| GATCCAGATCTAACTTGGGGTGGCTCTATTGTCTTCTTCTTTTGCCAATTCCACTAATTGTTTGGG | Ins_k3_25_CTA | -0.2292943088635202 | 0.0365994267886115 | 39.03913973934436 |
| GATCCAGATCTAACTTGGGGTGGCTCTCTTGTCTTCTTCTTTTGCCAATTCCACTAATTGTTTGGG | Ins_k3_25_CTC | 0.3252199212831838 | 0.4159919728505182 | 67.97085131089706 |
| GATCCAGATCTAACTTGGGGTGGCTCTGTCTTCTTCTTTTGCCAATTCCACTAATTGTTTGGG | T26C | -0.15465408802214564 | 0.02683455841604483 | 42.06453332403732 |
| GATCCAGATCTAACTTGGGGTGGCTCTGTTGTCTTCTTCTTTTGCCAATTCCACTAATTGTTTGGG | Ins_k3_25_CTG | 0.303829263960701 | 0.03480803383935181 | 66.5323502142514 |
| GATCCAGATCTAACTTGGGGTGGCTCTTCTTCTTTTGCCAATTCCACTAATTGTTTGGG | Del_k4_25-28;Del_k4_26-29 | 0.2657503947381019 | 0.03069294176780715 | 64.04650304205887 |
| GATCCAGATCTAACTTGGGGTGGCTCTTCTTTTGCCAATTCCACTAATTGTTTGGG | Del_k7_25-31;Del_k7_26-32 | 0.13195490674285126 | 0.030237367579191335 | 56.02589202910256 |
| GATCCAGATCTAACTTGGGGTGGCTCTTGTCTTCTTCTTTTGCCAATTCCACTAATTGTTTGGG | Ins_k1_25_C | 0.41111804659769535 | 0.0358538024788406 | 74.0675179094445 |
| GATCCAGATCTAACTTGGGGTGGCTCTTTGTCTTCTTCTTTTGCCAATTCCACTAATTGTTTGGG | Ins_k2_23_CT;Ins_k2_24_TC;Ins_ | -0.11870945931088989 | 0.039633184041826745 | 43.604029922098576 |
| GATCCAGATCTAACTTGGGGTGGCTCTTTTGCCAATTCCACTAATTGTTTGGG | Del_k10_25-34;Del_k10_26-35 | -0.563317252866099 | 0.03509311183012884 | 27.95348229312643 |
| GATCCAGATCTAACTTGGGGTGGCTCTTTTGTCTTCTTCTTTTGCCAATTCCACTAATTGTTTGGG | Ins_k3_24_TCT;Ins_k3_25_CTT | -0.3625566094448166 | 0.04215229496284954 | 34.16844049042528 |
| GATCCAGATCTAACTTGGGGTGGCTG | Del_k37_26-62 | -5.092791328071305 | 0.1269846991690806 | 0.30151597074051373 |
| GATCCAGATCTAACTTGGGGTGGCTGAATTGTCTTCTTCTTTTGCCAATTCCACTAATTGTTTGGG | Ins_k3_25_GAA | 0.4441942707963715 | 0.03166908642541453 | 76.55835845269515 |
| GATCCAGATCTAACTTGGGGTGGCTGACTTGTCTTCTTCTTTTGCCAATTCCACTAATTGTTTGGG | Ins_k3_25_GAC | 0.2945548843304223 | 0.030330135701885367 | 65.91815647605512 |
| GATCCAGATCTAACTTGGGGTGGCTGAGTTGTCTTCTTCTTTTGCCAATTCCACTAATTGTTTGGG | Ins_k3_25_GAG | 0.4971449363473235 | 0.032377645734678696 | 80.72142028916437 |
| GATCCAGATCTAACTTGGGGTGGCTGATTGTCTTCTTCTTTTGCCAATTCCACTAATTGTTTGGG | Ins_k2_25_GA | 0.5595834484109317 | 0.03084885881066047 | 85.92222127128973 |
| GATCCAGATCTAACTTGGGGTGGCTGATTTGTCTTCTTCTTTTGCCAATTCCACTAATTGTTTGGG | Ins_k3_24_TGA;Ins_k3_25_GAT | 0.39770087515804575 | 0.03140477756521623 | 73.08037844201195 |
| GATCCAGATCTAACTTGGGGTGGCTGCATTGTCTTCTTCTTTTGCCAATTCCACTAATTGTTTGGG | Ins_k3_25_GCA | 0.5857351981573811 | 0.028914700337946443 | 88.19887721630529 |
| GATCCAGATCTAACTTGGGGTGGCTGCCAATTCCACTAATTGTTTGGG | Del_k15_25-39;Del_k15_26-40 | 0.04201853058108618 | 0.02751221959282538 | 51.20706779459268 |
| GATCCAGATCTAACTTGGGGTGGCTGCCTTGTCTTCTTCTTTTGCCAATTCCACTAATTGTTTGGG | Ins_k3_25_GCC | 0.8408540886872538 | 0.028658280135584108 | 100 |
| GATCCAGATCTAACTTGGGGTGGCTGCGTTGTCTTCTTCTTTTGCCAATTCCACTAATTGTTTGGG | Ins_k3_25_GCG | 1.0278065454221492 | 0.02733509487575845 | 100 |
| GATCCAGATCTAACTTGGGGTGGCTGCTATTGTCTTCTTCTTTTGCCAATTCCACTAATTGTTTGGG | Ins_k4_25_GCTA | 0.7061603959195835 | 0.03195633521186445 | 99.48624871260168 |
| GATCCAGATCTAACTTGGGGTGGCTGCTTGTCTTCTTCTTTTGCCAATTCCACTAATTGTTTGGG | Ins_k2_25_GC | 0.8244981613156475 | 0.030225484576507575 | 100 |
| GATCCAGATCTAACTTGGGGTGGCTGCTTTGTCTTCTTCTTTTGCCAATTCCACTAATTGTTTGGG | Ins_k3_22_GCT;Ins_k3_23_CTG; | 0.6205396665124372 | 0.030008415573718753 | 91.32263732784067 |
| GATCCAGATCTAACTTGGGGTGGCTGG | Del_k36_26-61 | -5.068876442512376 | 0.15468696684618813 | 0.3088136039029822 |
| GATCCAGATCTAACTTGGGGTGGCTGGACTTGTCTTCTTCTTTTGCCAATTCCACTAATTGTTTGGG | Ins_k4_25_GGAC | 0.9368094137769478 | 0.028148986120691263 | 100 |
| GATCCAGATCTAACTTGGGGTGGCTGGATTGTCTTCTTCTTTTGCCAATTCCACTAATTGTTTGGG | Ins_k3_25_GGA | 0.7193259955554356 | 0.029302029628984202 | 100 |
| GATCCAGATCTAACTTGGGGTGGCTGGCTTGTCTTCTTCTTTTGCCAATTCCACTAATTGTTTGGG | Ins_k3_25_GGC | 0.5050826523327336 | 0.030354737737461973 | 81.36471376011214 |
| GATCCAGATCTAACTTGGGGTGGCTGGG | Del_k35_25-59;Del_k35_26-60 | -4.438529891354761 | 0.11456990435994582 | 0.5800326658745283 |

| | | | | |
|---|---|---|---|---|
| **GATCCAGATCTAACTTGGGGTGGCTGGGTTGTCTTCTTCTTTTGCCAATTCCACTAATTGTTTGGG** | Ins_k3_25_GGG | -1.7106783501314817 | 0.07125315548162442 | 8.874488364852386 |
| **GATCCAGATCTAACTTGGGGTGGCTGGTTGTCTTCTTCTTTTGCCAATTCCACTAATTGTTTGGG** | Ins_k2_25_GG | 0.6590193360288237 | 0.03232444578909602 | 94.90518787054995 |
| **GATCCAGATCTAACTTGGGGTGGCTGGTTTGTCTTCTTCTTTTGCCAATTCCACTAATTGTTTGGG** | Ins_k3_24_TGG;Ins_k3_25_GGT | 0.31671717458548243 | 0.03355255743792843 | 67.39536246447555 |
| **GATCCAGATCTAACTTGGGGTGGCTGTACTTGTCTTCTTCTTTTGCCAATTCCACTAATTGTTTGGG** | Ins_k4_25_GTAC | 0.5404114270801299 | 0.0300536611465678 | 84.29060923946638 |
| **GATCCAGATCTAACTTGGGGTGGCTGTATTGTCTTCTTCTTTTGCCAATTCCACTAATTGTTTGGG** | Ins_k3_25_GTA | 0.4949063945628942 | 0.031559845940529394 | 80.54092411643413 |
| **GATCCAGATCTAACTTGGGGTGGCTGTCTTCTTCTTTTGCCAATTCCACTAATTGTTTGGG** | Del_k2_25-26;Del_k2_26-27 | 0.560137249970277 | 0.02801097147749652 | 85.9698183098528 |
| **GATCCAGATCTAACTTGGGGTGGCTGTCTTGTCTTCTTCTTTTGCCAATTCCACTAATTGTTTGGG** | Ins_k3_25_GTC | 0.602700502027316 | 0.02941190693411096 | 89.70796282176045 |
| **GATCCAGATCTAACTTGGGGTGGCTGTGTCTTCTTCTTTTGCCAATTCCACTAATTGTTTGGG** | T26G | 0.41128542099006027 | 0.029504222021455997 | 74.07991595277711 |
| **GATCCAGATCTAACTTGGGGTGGCTGTGTTGTCTTCTTCTTTTGCCAATTCCACTAATTGTTTGGG** | Ins_k3_25_GTG | 0.4792562750980296 | 0.030519808947373675 | 79.29026107180948 |
| **GATCCAGATCTAACTTGGGGTGGCTGTTGTCTTCTTCTTTTGCCAATTCCACTAATTGTTTGGG** | Ins_k1_25_G | 0.7429335652236164 | 0.027944998199282544 | 100 |
| **GATCCAGATCTAACTTGGGGTGGCTGTTTGGG** | Del_k31_25-55;Del_k31_26-56 | -1.9752989839377644 | 0.0447106035502759 | 6.81114370410461 |
| **GATCCAGATCTAACTTGGGGTGGCTGTTTGTCTTCTTCTTTTGCCAATTCCACTAATTGTTTGGG** | Ins_k2_24_TG;Ins_k2_25_GT | 0.7929524030456401 | 0.029243482200461607 | 100 |
| **GATCCAGATCTAACTTGGGGTGGCTGTTTTGTCTTCTTCTTTTGCCAATTCCACTAATTGTTTGGG** | Ins_k3_24_TGT;Ins_k3_25_GTT | 0.5373109352339267 | 0.030258718074823353 | 84.02967161932355 |
| **GATCCAGATCTAACTTGGGGTGGCTGTTTTTGTCTTCTTCTTTTGCCAATTCCACTAATTGTTTGGG** | Ins_k4_24_TGTT;Ins_k4_25_GTT | 0.7597039749709824 | 0.03370844904200543 | 100 |
| **GATCCAGATCTAACTTGGGGTGGCTT** | Del_k37_27-63 | -6.770904746253466 | 0.7714142711120225 | 0.05630084632955228 |
| **GATCCAGATCTAACTTGGGGTGGCTTAAAATGTCTTCTTCTTTTGCCAATTCCACTAATTGTTTGGG** | Ins_k4_26_AAAA | -0.15344110920710358 | 0.03972584680019933 | 42.115587669483965 |
| **GATCCAGATCTAACTTGGGGTGGCTTAAATGTCTTCTTCTTTTGCCAATTCCACTAATTGTTTGGG** | Ins_k3_26_AAA | -0.15951122596703837 | 0.03568065384999812 | 41.860715469514616 |
| **GATCCAGATCTAACTTGGGGTGGCTTAACTGTCTTCTTCTTTTGCCAATTCCACTAATTGTTTGGG** | Ins_k3_26_AAC | -0.4217548430006657 | 0.03732689220416811 | 32.20443550929703 |
| **GATCCAGATCTAACTTGGGGTGGCTTAAGATGTCTTCTTCTTTTGCCAATTCCACTAATTGTTTGGG** | Ins_k4_26_AAGA | 0.3388653542948219 | 0.1607807922832209 | 68.90469990340553 |
| **GATCCAGATCTAACTTGGGGTGGCTTAAGTGTCTTCTTCTTTTGCCAATTCCACTAATTGTTTGGG** | Ins_k3_26_AAG | 0.06876602323365488 | 0.036370584567078604 | 52.59521037215765 |
| **GATCCAGATCTAACTTGGGGTGGCTTAATGTCTTCTTCTTTTGCCAATTCCACTAATTGTTTGGG** | Ins_k2_26_AA | -0.14259507419456213 | 0.034989627896183075 | 42.57486095271068 |
| **GATCCAGATCTAACTTGGGGTGGCTTAATTCCACTAATTGTTTGGG** | Del_k17_27-43 | -1.3402150638903012 | 0.04353868227870467 | 12.853857629384166 |
| **GATCCAGATCTAACTTGGGGTGGCTTAATTGTCTTCTTCTTTTGCCAATTCCACTAATTGTTTGGG** | Ins_k3_25_TAA;Ins_k3_26_AAT | -0.3588496164073174 | 0.038272257231871526 | 34.29533771968186 |
| **GATCCAGATCTAACTTGGGGTGGCTTAATTGTTTGGG** | Del_k26_26-51;Del_k26_27-52 | -3.308341332420766 | 0.10212109129214114 | 1.7959204973291683 |
| **GATCCAGATCTAACTTGGGGTGGCTTACATGTCTTCTTCTTTTGCCAATTCCACTAATTGTTTGGG** | Ins_k3_26_ACA | 0.11681523194956234 | 0.029057888252995923 | 55.18406680333803 |
| **GATCCAGATCTAACTTGGGGTGGCTTACCTGTCTTCTTCTTTTGCCAATTCCACTAATTGTTTGGG** | Ins_k3_26_ACC | 0.4929918683736716 | 0.02781369616752399 | 80.3868739215301 |
| **GATCCAGATCTAACTTGGGGTGGCTTACGTGTCTTCTTCTTTTGCCAATTCCACTAATTGTTTGGG** | Ins_k3_26_ACG | 0.5979958879735388 | 0.030593118525154915 | 89.28691269491091 |
| **GATCCAGATCTAACTTGGGGTGGCTTACTAATTGTTTGGG** | Del_k23_27-49 | -2.7661211249394704 | 0.06918545941588458 | 3.088661760430609 |
| **GATCCAGATCTAACTTGGGGTGGCTTACTGTCTTCTTCTTTTGCCAATTCCACTAATTGTTTGGG** | Ins_k2_26_AC | 0.10746254095689423 | 0.033974862769804166 | 54.67035332517646 |
| **GATCCAGATCTAACTTGGGGTGGCTTACTTGTCTTCTTCTTTTGCCAATTCCACTAATTGTTTGGG** | Ins_k3_25_TAC;Ins_k3_26_ACT | -0.1386525493642865 | 0.03963706003113922 | 42.743044715645986 |
| **GATCCAGATCTAACTTGGGGTGGCTTAGATGTCTTCTTCTTTTGCCAATTCCACTAATTGTTTGGG** | Ins_k3_26_AGA | 0.4263403413428757 | 0.029074463114052557 | 75.20362060526091 |
| **GATCCAGATCTAACTTGGGGTGGCTTAGATTGTCTTCTTCTTTTGCCAATTCCACTAATTGTTTGGG** | Ins_k4_25_TAGA;Ins_k4_26_AGA | 0.25473138829160424 | 0.038178615551842876 | 63.34464818525805 |
| **GATCCAGATCTAACTTGGGGTGGCTTAGCCTGTCTTCTTCTTTTGCCAATTCCACTAATTGTTTGGG** | Ins_k4_26_AGCC | 0.3603101349332004 | 0.033698203491356604 | 70.39830384293857 |
| **GATCCAGATCTAACTTGGGGTGGCTTAGCGTGTCTTCTTCTTTTGCCAATTCCACTAATTGTTTGGG** | Ins_k4_26_AGCG | 0.43207368244121147 | 0.03234919482523675 | 75.63602699621114 |
| **GATCCAGATCTAACTTGGGGTGGCTTAGCTGTCTTCTTCTTTTGCCAATTCCACTAATTGTTTGGG** | Ins_k3_26_AGC | 0.36171728827211713 | 0.030463739401113576 | 70.49743478110328 |
| **GATCCAGATCTAACTTGGGGTGGCTTAGGTGTCTTCTTCTTTTGCCAATTCCACTAATTGTTTGGG** | Ins_k3_26_AGG | -0.42675553907984887 | 0.04024685423154018 | 32.04379291208153 |
| **GATCCAGATCTAACTTGGGGTGGCTTAGTCTTCTTCTTTTGCCAATTCCACTAATTGTTTGGG** | T27A | -0.418974031950633 | 0.03816593831988794 | 32.29411459192329 |
| **GATCCAGATCTAACTTGGGGTGGCTTAGTGTCTTCTTCTTTTGCCAATTCCACTAATTGTTTGGG** | Ins_k2_26_AG | -0.15960681978762487 | 0.036893066811539574 | 41.856714035049656 |
| **GATCCAGATCTAACTTGGGGTGGCTTAGTTGTCTTCTTCTTTTGCCAATTCCACTAATTGTTTGGG** | Ins_k3_25_TAG;Ins_k3_26_AGT | -0.8186696209768103 | 0.043662486026487514 | 21.653983086824187 |
| **GATCCAGATCTAACTTGGGGTGGCTTATATGTCTTCTTCTTTTGCCAATTCCACTAATTGTTTGGG** | Ins_k3_26_ATA | -0.2130214493969128 | 0.03768414796185078 | 39.67961522438539 |
| **GATCCAGATCTAACTTGGGGTGGCTTATCTGTCTTCTTCTTTTGCCAATTCCACTAATTGTTTGGG** | Ins_k3_26_ATC | -0.32581783048644936 | 0.036702142175788835 | 35.44709149417841 |
| **GATCCAGATCTAACTTGGGGTGGCTTATGTCTTCTTCTTTTGCCAATTCCACTAATTGTTTGGG** | Ins_k1_26_A | 0.21848312445179674 | 0.03630218760148929 | 61.0896317866692 |
| **GATCCAGATCTAACTTGGGGTGGCTTATGTGTCTTCTTCTTTTGCCAATTCCACTAATTGTTTGGG** | Ins_k3_26_ATG | 0.13223979068095792 | 0.03198909846459069 | 56.04185517957436 |
| **GATCCAGATCTAACTTGGGGTGGCTTATTCCACTAATTGTTTGGG** | Del_k18_27-44 | -1.1209964934059518 | 0.03840501705274017 | 16.004381706346056 |
| **GATCCAGATCTAACTTGGGGTGGCTTATTGTCTTCTTCTTTTGCCAATTCCACTAATTGTTTGGG** | Ins_k2_25_TA;Ins_k2_26_AT | -0.022182318049063454 | 0.043589615988656495 | 48.02283931245548 |
| **GATCCAGATCTAACTTGGGGTGGCTTATTGTTTGGG** | Del_k27_27-53 | -3.306802483646846 | 0.09009870030193773 | 1.7986862748954566 |
| **GATCCAGATCTAACTTGGGGTGGCTTATTTGTCTTCTTCTTTTGCCAATTCCACTAATTGTTTGGG** | Ins_k3_24_TTA;Ins_k3_25_TAT;In | -0.3064770085648903 | 0.042304252621892885 | 36.13934012834839 |
| **GATCCAGATCTAACTTGGGGTGGCTTCAATGTCTTCTTCTTTTGCCAATTCCACTAATTGTTTGGG** | Ins_k3_26_CAA | 0.5653392697864212 | 0.031335954417118124 | 86.41820024300314 |
| **GATCCAGATCTAACTTGGGGTGGCTTCAATTCCACTAATTGTTTGGG** | Del_k16_27-42 | -0.21407083715301162 | 0.03255906894538273 | 39.63799776224896 |
| **GATCCAGATCTAACTTGGGGTGGCTTCACTAATTGTTTGGG** | Del_k22_27-48 | -0.9057236921806285 | 0.03495794596352956 | 19.848637056203415 |
| **GATCCAGATCTAACTTGGGGTGGCTTCACTGTCTTCTTCTTTTGCCAATTCCACTAATTGTTTGGG** | Ins_k3_26_CAC | 0.4802599167509245 | 0.02883444942007592 | 79.36988002825727 |
| **GATCCAGATCTAACTTGGGGTGGCTTCAGTGTCTTCTTCTTTTGCCAATTCCACTAATTGTTTGGG** | Ins_k3_26_CAG | 0.6030826394653868 | 0.02989384412816344 | 89.74225014366513 |
| **GATCCAGATCTAACTTGGGGTGGCTTCATGTCTTCTTCTTTTGCCAATTCCACTAATTGTTTGGG** | Ins_k2_26_CA | 0.600774305188883 | 0.02911307046122133 | 89.53533393936911 |
| **GATCCAGATCTAACTTGGGGTGGCTTCATTGTCTTCTTCTTTTGCCAATTCCACTAATTGTTTGGG** | Ins_k3_25_TCA;Ins_k3_26_CAT | 0.40447330126387815 | 0.03334724888998584 | 73.5769896379916 |
| **GATCCAGATCTAACTTGGGGTGGCTTCCAATTCCACTAATTGTTTGGG** | Del_k15_27-41 | -0.1386553810382654 | 0.027895220528179308 | 42.74292368144985 |
| **GATCCAGATCTAACTTGGGGTGGCTTCCACTAATTGTTTGGG** | Del_k21_25-45;Del_k21_26-46;D | -0.8672347571596313 | 0.03125231569123172 | 20.627482273651676 |
| **GATCCAGATCTAACTTGGGGTGGCTTCCATGTCTTCTTCTTTTGCCAATTCCACTAATTGTTTGGG** | Ins_k3_26_CCA | 0.5185698328856372 | 0.031229593269853556 | 82.46952801241538 |
| **GATCCAGATCTAACTTGGGGTGGCTTCCCTGTCTTCTTCTTTTGCCAATTCCACTAATTGTTTGGG** | Ins_k3_26_CCC | 0.4256450901205535 | 0.030351674090432698 | 75.15135336764784 |
| **GATCCAGATCTAACTTGGGGTGGCTTCCGTGTCTTCTTCTTTTGCCAATTCCACTAATTGTTTGGG** | Ins_k3_26_CCG | 0.7171022947198427 | 0.02923482592178118 | 100 |
| **GATCCAGATCTAACTTGGGGTGGCTTCCTCTGTCTTCTTCTTTTGCCAATTCCACTAATTGTTTGGG** | Ins_k4_26_CCTC | 0.4375569777498338 | 0.03419613150415553 | 76.05190080566689 |
| **GATCCAGATCTAACTTGGGGTGGCTTCCTGTCTTCTTCTTTTGCCAATTCCACTAATTGTTTGGG** | Ins_k2_26_CC | 0.8550054194487671 | 0.027885806397642845 | 100 |
| **GATCCAGATCTAACTTGGGGTGGCTTCCTTGTCTTCTTCTTTTGCCAATTCCACTAATTGTTTGGG** | Ins_k3_25_TCC;Ins_k3_26_CCT | 0.38572894555517423 | 0.03358848584637461 | 72.21068165785906 |
| **GATCCAGATCTAACTTGGGGTGGCTTCGATGTCTTCTTCTTTTGCCAATTCCACTAATTGTTTGGG** | Ins_k3_26_CGA | 0.7571009228408714 | 0.028121818934637284 | 100 |
| **GATCCAGATCTAACTTGGGGTGGCTTCGCTGTCTTCTTCTTTTGCCAATTCCACTAATTGTTTGGG** | Ins_k3_26_CGC | 0.7700829122757935 | 0.027294855313799462 | 100 |
| **GATCCAGATCTAACTTGGGGTGGCTTCGGGTGTCTTCTTCTTTTGCCAATTCCACTAATTGTTTGGG** | Ins_k4_26_CGGG | 0.5902075608671926 | 0.03214497926418135 | 88.59421798049259 |
| **GATCCAGATCTAACTTGGGGTGGCTTCGGTGTCTTCTTCTTTTGCCAATTCCACTAATTGTTTGGG** | Ins_k3_26_CGG | 0.7242435742107565 | 0.02780219773294507 | 100 |
| **GATCCAGATCTAACTTGGGGTGGCTTCGTCTTCTTCTTTTGCCAATTCCACTAATTGTTTGGG** | T27C | 0.14071862607067898 | 0.02470425580067318 | 56.51904499272721 |
| **GATCCAGATCTAACTTGGGGTGGCTTCGTGTCTTCTTCTTTTGCCAATTCCACTAATTGTTTGGG** | Ins_k2_26_CG | 0.9029636298714245 | 0.02956203680898192 | 100 |
| **GATCCAGATCTAACTTGGGGTGGCTTCGTTGTCTTCTTCTTTTGCCAATTCCACTAATTGTTTGGG** | Ins_k3_25_TCG;Ins_k3_26_CGT | 0.8145202327497529 | 0.029999146997659376 | 100 |
| **GATCCAGATCTAACTTGGGGTGGCTTCTAATTGTTTGGG** | Del_k24_27-50 | -1.6225321113698936 | 0.04333151175303626 | 9.692253162962672 |
| **GATCCAGATCTAACTTGGGGTGGCTTCTATGTCTTCTTCTTTTGCCAATTCCACTAATTGTTTGGG** | Ins_k3_26_CTA | 0.11137038792488663 | 0.033506326681497514 | 54.88441468676311 |
| **GATCCAGATCTAACTTGGGGTGGCTTCTCTGTCTTCTTCTTTTGCCAATTCCACTAATTGTTTGGG** | Ins_k3_26_CTC | -0.23535529686242396 | 0.04193867143767111 | 38.8032395980196 |
| **GATCCAGATCTAACTTGGGGTGGCTTCTGTCTTCTTCTTTTGCCAATTCCACTAATTGTTTGGG** | Ins_k1_26_C | 0.3847680568125017 | 0.03276121083730096 | 72.14132855239865 |
| **GATCCAGATCTAACTTGGGGTGGCTTCTGTGTCTTCTTCTTTTGCCAATTCCACTAATTGTTTGGG** | Ins_k3_26_CTG | 0.4941571591146705 | 0.029883373953236012 | 80.48060260138391 |
| **GATCCAGATCTAACTTGGGGTGGCTTCTTCTTCTTTTGCCAATTCCACTAATTGTTTGGG** | Del_k3_26-28;Del_k3_27-29 | 0.27664746174248434 | 0.03058406119739204 | 64.74823856258367 |

| | | | | |
|---|---|---|---|---|
| GATCCAGATCTAACTTGGGGTGGCTTCTTCTTTTGCCAATTCCACTAATTGTTTGGG | Del_k6_24-29;Del_k6_25-30;Del_ | -0.14071487011126765 | 0.03170945862234485 | 42.65498568193667 |
| GATCCAGATCTAACTTGGGGTGGCTTCTTGTCTTCTTCTTTTGCCAATTCCACTAATTGTTTGGG | Ins_k2_25_TC;Ins_k2_26_CT | 0.24422580913827951 | 0.03553246998049573 | 62.68265934635748 |
| GATCCAGATCTAACTTGGGGTGGCTTCTTTGTCTTCTTCTTTTGCCAATTCCACTAATTGTTTGGG | Ins_k3_23_CTT;Ins_k3_24_TTC;I | -0.054479605118310626 | 0.037566404504576545 | 46.496611068130804 |
| GATCCAGATCTAACTTGGGGTGGCTTCTTTTGCCAATTCCACTAATTGTTTGGG | Del_k9_24-32;Del_k9_25-33;Del_ | -0.306917877105296 | 0.03286935620597661 | 36.123410941811386 |
| GATCCAGATCTAACTTGGGGTGGCTTG | Del_k36_27-62 | -4.706422662217077 | 0.1139305275261624 | 0.4437190764843404 |
| GATCCAGATCTAACTTGGGGTGGCTTGAATGTCTTCTTCTTTTGCCAATTCCACTAATTGTTTGGG | Ins_k3_26_GAA | 0.5818923492024328 | 0.02771293265696729 | 87.86059265769927 |
| GATCCAGATCTAACTTGGGGTGGCTTGACCTGTCTTCTTCTTTTGCCAATTCCACTAATTGTTTGGG | Ins_k4_26_GACC | 0.8312262686517644 | 0.02786124137089988 | 100 |
| GATCCAGATCTAACTTGGGGTGGCTTGACTGTCTTCTTCTTTTGCCAATTCCACTAATTGTTTGGG | Ins_k3_26_GAC | 0.5301760729963663 | 0.028702936275317996 | 83.4322652289082 |
| GATCCAGATCTAACTTGGGGTGGCTTGAGTGTCTTCTTCTTTTGCCAATTCCACTAATTGTTTGGG | Ins_k3_26_GAG | 0.5283825013566266 | 0.030200950074089742 | 83.2827576005547 |
| GATCCAGATCTAACTTGGGGTGGCTTGAGTTGTCTTCTTCTTTTGCCAATTCCACTAATTGTTTGGG | Ins_k4_25_TGAG;Ins_k4_26_GA | 0.7000066523842918 | 0.030388063428165326 | 98.87591569520099 |
| GATCCAGATCTAACTTGGGGTGGCTTGATGTCTTCTTCTTTTGCCAATTCCACTAATTGTTTGGG | Ins_k2_26_GA | 0.8381812371659241 | 0.02942064133182603 | 100 |
| GATCCAGATCTAACTTGGGGTGGCTTGATTGTCTTCTTCTTTTGCCAATTCCACTAATTGTTTGGG | Ins_k3_25_TGA;Ins_k3_26_GAT | -0.13074595653951532 | 0.1530466501269982 | 43.08233611824558 |
| GATCCAGATCTAACTTGGGGTGGCTTGCATGTCTTCTTCTTTTGCCAATTCCACTAATTGTTTGGG | Ins_k3_26_GCA | 0.2995570707553812 | 0.029186379184484215 | 66.2487174580956 |
| GATCCAGATCTAACTTGGGGTGGCTTGCCAATTCCACTAATTGTTTGGG | Del_k14_25-38;Del_k14_26-39;D | -0.17816864769342677 | 0.028446897980750552 | 41.08694319076825 |
| GATCCAGATCTAACTTGGGGTGGCTTGCCTGTCTTCTTCTTTTGCCAATTCCACTAATTGTTTGGG | Ins_k3_26_GCC | 0.6150789998031634 | 0.030106287832833218 | 90.82531393682561 |
| GATCCAGATCTAACTTGGGGTGGCTTGCGTGTCTTCTTCTTTTGCCAATTCCACTAATTGTTTGGG | Ins_k3_26_GCG | 0.8944106812820243 | 0.028119829555211487 | 100 |
| GATCCAGATCTAACTTGGGGTGGCTTGCTGTCTTCTTCTTTTGCCAATTCCACTAATTGTTTGGG | Ins_k2_26_GC | 0.8156626810975531 | 0.03242433161658924 | 100 |
| GATCCAGATCTAACTTGGGGTGGCTTGCTTGTCTTCTTCTTTTGCCAATTCCACTAATTGTTTGGG | Ins_k3_25_TGC;Ins_k3_26_GCT | 0.3541148964020384 | 0.03162899035884321 | 69.96351775084312 |
| GATCCAGATCTAACTTGGGGTGGCTTGCTTTGTCTTCTTCTTTTGCCAATTCCACTAATTGTTTGGG | Ins_k4_22_GCTT;Ins_k4_23_CTT | 0.20756722086547919 | 0.04126572951770235 | 60.426409675885076 |
| GATCCAGATCTAACTTGGGGTGGCTTGG | Del_k35_27-61 | -5.28643522727357 | 0.14196368095521003 | 0.2484344645247977 |
| GATCCAGATCTAACTTGGGGTGGCTTGGATGTCTTCTTCTTTTGCCAATTCCACTAATTGTTTGGG | Ins_k3_26_GGA | 0.7308555800082073 | 0.027469747687578195 | 100 |
| GATCCAGATCTAACTTGGGGTGGCTTGGCTGTCTTCTTCTTTTGCCAATTCCACTAATTGTTTGGG | Ins_k3_26_GGC | 0.21935822785561876 | 0.03295579246636638 | 61.14311492961102 |
| GATCCAGATCTAACTTGGGGTGGCTTGGG | Del_k34_25-58;Del_k34_26-59;D | -4.793565075131537 | 0.1523142431957375 | 0.40668919239120527 |
| GATCCAGATCTAACTTGGGGTGGCTTGGGTGTCTTCTTCTTTTGCCAATTCCACTAATTGTTTGGG | Ins_k3_26_GGG | -1.246081863922496 | 0.05788889821216468 | 14.122611577521488 |
| GATCCAGATCTAACTTGGGGTGGCTTGGTCTTCTTCTTTTGCCAATTCCACTAATTGTTTGGG | T27G | -0.03594469255866428 | 0.03199199643943184 | 47.36645805517791 |
| GATCCAGATCTAACTTGGGGTGGCTTGGTGTCTTCTTCTTTTGCCAATTCCACTAATTGTTTGGG | Ins_k2_26_GG | 0.6222375443077222 | 0.03523781000501007 | 91.47782371244395 |
| GATCCAGATCTAACTTGGGGTGGCTTGGTTGTCTTCTTCTTTTGCCAATTCCACTAATTGTTTGGG | Ins_k3_25_TGG;Ins_k3_26_GGT | 0.05820173745031598 | 0.032433595926074435 | 52.04250415252923 |
| GATCCAGATCTAACTTGGGGTGGCTTGTATGTCTTCTTCTTTTGCCAATTCCACTAATTGTTTGGG | Ins_k3_26_GTA | -1.0154352110229312 | 0.4789435327582425 | 17.786216862464574 |
| GATCCAGATCTAACTTGGGGTGGCTTGTCTGTCTTCTTCTTTTGCCAATTCCACTAATTGTTTGGG | Ins_k3_26_GTC | -0.035202314034220916 | 0.03321359269947135 | 47.40163495208785 |
| GATCCAGATCTAACTTGGGGTGGCTTGTCTTCTTCTTTTGCCAATTCCACTAATTGTTTGGG | Del_k1_25-25;Del_k1_26-26;Del_ | 0.24829648058280607 | 0.02800534656215936 | 62.938339900531155 |
| GATCCAGATCTAACTTGGGGTGGCTTGTGTCTTCTTCTTTTGCCAATTCCACTAATTGTTTGGG | Ins_k1_26_G | 0.43991190088175536 | 0.031736443035798076 | 76.23120823083752 |
| GATCCAGATCTAACTTGGGGTGGCTTGTGTGTCTTCTTCTTTTGCCAATTCCACTAATTGTTTGGG | Ins_k3_26_GTG | 0.276514857402423 | 0.03188425574685325 | 64.73965323437741 |
| GATCCAGATCTAACTTGGGGTGGCTTGTTGTCTTCTTCTTTTGCCAATTCCACTAATTGTTTGGG | Ins_k2_25_TG;Ins_k2_26_GT | 0.34090666929111957 | 0.033769636380978366 | 69.04549976017093 |
| GATCCAGATCTAACTTGGGGTGGCTTGTTTGGG | Del_k30_25-54;Del_k30_26-55;D | -3.1784912859571457 | 0.07545853235164844 | 2.0449385677905503 |
| GATCCAGATCTAACTTGGGGTGGCTTGTTTGTCTTCTTCTTTTGCCAATTCCACTAATTGTTTGGG | Ins_k3_24_TTG;Ins_k3_25_TGT;I | 0.005656759256416399 | 0.036404590026752136 | 49.3785339364699 |
| GATCCAGATCTAACTTGGGGTGGCTTT | Del_k36_28-63 | -5.786154930403209 | 0.23258816508746674 | 0.1507253615903016 |
| GATCCAGATCTAACTTGGGGTGGCTTTAAAGTCTTCTTCTTTTGCCAATTCCACTAATTGTTTGGG | Ins_k3_27_AAA | -0.0847916500186473 | 0.03612805442410025 | 45.10835050909666 |
| GATCCAGATCTAACTTGGGGTGGCTTTAACGTCTTCTTCTTTTGCCAATTCCACTAATTGTTTGGG | Ins_k3_27_AAC | 0.1975976307672216 | 0.03462439973306405 | 59.82697615847307 |
| GATCCAGATCTAACTTGGGGTGGCTTTAAGGTCTTCTTCTTTTGCCAATTCCACTAATTGTTTGGG | Ins_k3_27_AAG | -0.3509498347537217 | 0.039085299731155786 | 34.56733634975764 |
| GATCCAGATCTAACTTGGGGTGGCTTTAAGTCTTCTTCTTTTGCCAATTCCACTAATTGTTTGGG | Ins_k2_27_AA | -0.026145139349396995 | 0.03690988200083726 | 47.83290995852365 |
| GATCCAGATCTAACTTGGGGTGGCTTTAATGTCTTCTTCTTTTGCCAATTCCACTAATTGTTTGGG | Ins_k3_26_TAA;Ins_k3_27_AAT | -0.6279613594465462 | 0.05147752580662772 | 26.203622782621693 |
| GATCCAGATCTAACTTGGGGTGGCTTTAATTCCACTAATTGTTTGGG | Del_k16_28-43 | -1.5266988009138518 | 0.04685920408248602 | 10.667057284538854 |
| GATCCAGATCTAACTTGGGGTGGCTTTAATTGTTTGGG | Del_k25_27-51;Del_k25_28-52 | -3.637739885546973 | 0.11613642996107204 | 1.291906648482524 |
| GATCCAGATCTAACTTGGGGTGGCTTTACAGTCTTCTTCTTTTGCCAATTCCACTAATTGTTTGGG | Ins_k3_27_ACA | 0.1599673587113304 | 0.030179649572570244 | 57.61750302545449 |
| GATCCAGATCTAACTTGGGGTGGCTTTACCGTCTTCTTCTTTTGCCAATTCCACTAATTGTTTGGG | Ins_k3_27_ACC | 0.5041622727830293 | 0.031270325111722515 | 81.28986179289362 |
| GATCCAGATCTAACTTGGGGTGGCTTTACGGTCTTCTTCTTTTGCCAATTCCACTAATTGTTTGGG | Ins_k3_27_ACG | 0.46123328048737244 | 0.02969371107522226 | 77.87401396702792 |
| GATCCAGATCTAACTTGGGGTGGCTTTACGTCTTCTTCTTTTGCCAATTCCACTAATTGTTTGGG | Ins_k2_27_AC | 0.5527813289167067 | 0.03300971854833638 | 85.33975131596068 |
| GATCCAGATCTAACTTGGGGTGGCTTTACTAATTGTTTGGG | Del_k22_28-49 | -3.396482224207208 | 0.08763605443202448 | 1.6444020300980364 |
| GATCCAGATCTAACTTGGGGTGGCTTTACTGTCTTCTTCTTTTGCCAATTCCACTAATTGTTTGGG | Ins_k3_26_TAC;Ins_k3_27_ACT | -0.8476089250085467 | 0.05138663285805891 | 21.03631247242851 |
| GATCCAGATCTAACTTGGGGTGGCTTTAGAGTCTTCTTCTTTTGCCAATTCCACTAATTGTTTGGG | Ins_k3_27_AGA | 0.12906238444093798 | 0.033000717951417236 | 55.864070036611025 |
| GATCCAGATCTAACTTGGGGTGGCTTTAGCGTCTTCTTCTTTTGCCAATTCCACTAATTGTTTGGG | Ins_k3_27_AGC | 0.02084021629848376 | 0.03131657217283791 | 50.13399150065357 |
| GATCCAGATCTAACTTGGGGTGGCTTTAGGGTCTTCTTCTTTTGCCAATTCCACTAATTGTTTGGG | Ins_k3_27_AGG | -3.9658837434737504 | 0.21486172793589173 | 0.9305079039113793 |
| GATCCAGATCTAACTTGGGGTGGCTTTAGGTCTTCTTCTTTTGCCAATTCCACTAATTGTTTGGG | Ins_k2_27_AG | -2.065048117430251 | 0.09559106850292302 | 6.226478457537512 |
| GATCCAGATCTAACTTGGGGTGGCTTTAGTCTTCTTCTTTTGCCAATTCCACTAATTGTTTGGG | Ins_k1_27_A | -0.7118903991408948 | 0.0475129157786553 | 24.094139476110364 |
| GATCCAGATCTAACTTGGGGTGGCTTTAGTGTCTTCTTCTTTTGCCAATTCCACTAATTGTTTGGG | Ins_k3_26_TAG;Ins_k3_27_AGT | -0.6970714287932168 | 0.04396890958479952 | 24.453848490461098 |
| GATCCAGATCTAACTTGGGGTGGCTTTATAGTCTTCTTCTTTTGCCAATTCCACTAATTGTTTGGG | Ins_k3_27_ATA | -1.0814882793181884 | 0.0539511572397966 | 16.649342988242868 |
| GATCCAGATCTAACTTGGGGTGGCTTTATCGTCTTCTTCTTTTGCCAATTCCACTAATTGTTTGGG | Ins_k3_27_ATC | 0.44842720377366607 | 0.02915359003751668 | 76.8831116973334 |
| GATCCAGATCTAACTTGGGGTGGCTTTATCTTCTTCTTTTGCCAATTCCACTAATTGTTTGGG | G28A | -0.5397508942816935 | 0.0353543582877061 | 28.620067726175574 |
| GATCCAGATCTAACTTGGGGTGGCTTTATGGTCTTCTTCTTTTGCCAATTCCACTAATTGTTTGGG | Ins_k3_27_ATG | -0.19729363677709327 | 0.03670421043927055 | 40.30862226379455 4 |
| GATCCAGATCTAACTTGGGGTGGCTTTATGTCTTCTTCTTTTGCCAATTCCACTAATTGTTTGGG | Ins_k2_26_TA;Ins_k2_27_AT | -0.18082127948837262 | 0.044255685800993365 | 40.97809908439497 |
| GATCCAGATCTAACTTGGGGTGGCTTTATTCCACTAATTGTTTGGG | Del_k17_28-44 | -1.4208181075437798 | 0.04333210184508731 | 11.858452770462053 |
| GATCCAGATCTAACTTGGGGTGGCTTTATTGTCTTCTTCTTTTGCCAATTCCACTAATTGTTTGGG | Ins_k3_25_TTA;Ins_k3_26_TAT;In | -0.6173442117045924 | 0.04419127724286567 | 26.483312644065307 |
| GATCCAGATCTAACTTGGGGTGGCTTTATTGTTTGGG | Del_k26_28-53 | -3.1392694835606103 | 0.09130870579610051 | 2.1267384270363956 |
| GATCCAGATCTAACTTGGGGTGGCTTTCAAGTCTTCTTCTTTTGCCAATTCCACTAATTGTTTGGG | Ins_k3_27_CAA | 0.5448166689106952 | 0.029600842030762707 | 84.66274883785835 |
| GATCCAGATCTAACTTGGGGTGGCTTTCAATTCCACTAATTGTTTGGG | Del_k15_28-42 | -0.7901418873797857 | 0.03873624050370094 | 22.280617885584252 |
| GATCCAGATCTAACTTGGGGTGGCTTTCACGTCTTCTTCTTTTGCCAATTCCACTAATTGTTTGGG | Ins_k3_27_CAC | 0.48822331065305224 | 0.028066513085917517 | 80.00445698652159 |
| GATCCAGATCTAACTTGGGGTGGCTTTCACTAATTGTTTGGG | Del_k21_28-48 | -2.080306033865389 | 0.054414390041402694 | 6.132196471970735 |
| GATCCAGATCTAACTTGGGGTGGCTTTCAGGTCTTCTTCTTTTGCCAATTCCACTAATTGTTTGGG | Ins_k3_27_CAG | -0.5625600033053412 | 0.04477513430376971 | 27.97465807197727 |
| GATCCAGATCTAACTTGGGGTGGCTTTCAGTCTTCTTCTTTTGCCAATTCCACTAATTGTTTGGG | Ins_k2_27_CA | 0.35148285506817534 | 0.03210432926664851 | 69.77961300887046 |
| GATCCAGATCTAACTTGGGGTGGCTTTCATGTCTTCTTCTTTTGCCAATTCCACTAATTGTTTGGG | Ins_k3_26_TCA;Ins_k3_27_CAT | -0.07801989657669683 | 0.034857578491398616 | 45.41484973425754 |
| GATCCAGATCTAACTTGGGGTGGCTTTCCAATTCCACTAATTGTTTGGG | Del_k14_28-41 | -0.2411867290837566 | 0.030343556569249776 | 38.57761961950343 |
| GATCCAGATCTAACTTGGGGTGGCTTTCCACTAATTGTTTGGG | Del_k20_26-45;Del_k20_27-46;D | -0.9742413071599625 | 0.037250080159395016 | 18.53420099260825 |
| GATCCAGATCTAACTTGGGGTGGCTTTCCAGTCTTCTTCTTTTGCCAATTCCACTAATTGTTTGGG | Ins_k3_27_CCA | 0.7074092546793915 | 0.028044968754170386 | 99.61057059986761 |
| GATCCAGATCTAACTTGGGGTGGCTTTCCCGTCTTCTTCTTTTGCCAATTCCACTAATTGTTTGGG | Ins_k3_27_CCC | 0.5640580422579549 | 0.032718048255953806 | 86.30754976524683 |
| GATCCAGATCTAACTTGGGGTGGCTTTCCGGTCTTCTTCTTTTGCCAATTCCACTAATTGTTTGGG | Ins_k3_27_CCG | 0.8756757275016072 | 0.029859023568226572 | 100 |

| | | | | |
|---|---|---|---|---|
| **GATCCAGATCTAACTTGGGGTGGCTTTCCGTCTTCTTCTTTTGCCAATTCCACTAATTGTTTGGG** | Ins_k2_27_CC | 0.6339013605755375 | 0.03531889788842157 | 92.55105103676259 |
| **GATCCAGATCTAACTTGGGGTGGCTTTCCTGTCTTCTTCTTTTGCCAATTCCACTAATTGTTTGGG** | Ins_k3_26_TCC;Ins_k3_27_CCT | 0.2450637123075573 | 0.031871695312773174 | 62.73520335560431 |
| **GATCCAGATCTAACTTGGGGTGGCTTTCGAGTCTTCTTCTTTTGCCAATTCCACTAATTGTTTGGG** | Ins_k3_27_CGA | 0.6970050596341901 | 0.02857836599561299 | 98.57957543237406 |
| **GATCCAGATCTAACTTGGGGTGGCTTTCGCGTCTTCTTCTTTTGCCAATTCCACTAATTGTTTGGG** | Ins_k3_27_CGC | 0.7812420072136619 | 0.029226785638862888 | 100 |
| **GATCCAGATCTAACTTGGGGTGGCTTTCGGGTCTTCTTCTTTTGCCAATTCCACTAATTGTTTGGG** | Ins_k3_27_CGG | -0.5585468100094548 | 0.05299009810675072 | 28.087151359878785 |
| **GATCCAGATCTAACTTGGGGTGGCTTTCGGTCTTCTTCTTTTGCCAATTCCACTAATTGTTTGGG** | Ins_k2_27_CG | 0.5808751943459298 | 0.029829261671789828 | 87.77127026423652 |
| **GATCCAGATCTAACTTGGGGTGGCTTTCGGTGTCTTCTTCTTTTGCCAATTCCACTAATTGTTTGGG** | Ins_k4_26_TCGG;Ins_k4_27_CG | 0.801593515279448 | 0.030806228922565015 | 100 |
| **GATCCAGATCTAACTTGGGGTGGCTTTCGTCTTCTTCTTTTGCCAATTCCACTAATTGTTTGGG** | Ins_k1_27_C | 0.6725016662135077 | 0.029699944961150263 | 96.1933954562569 |
| **GATCCAGATCTAACTTGGGGTGGCTTTCGTGTCTTCTTCTTTTGCCAATTCCACTAATTGTTTGGG** | Ins_k3_26_TCG;Ins_k3_27_CGT | 0.6293962428238 | 0.030280535662530257 | 92.13503545700534 |
| **GATCCAGATCTAACTTGGGGTGGCTTTCTAATTGTTTGGG** | Del_k23_28-50 | -2.826182406767871 | 0.07651925326684511 | 2.908613851574794 |
| **GATCCAGATCTAACTTGGGGTGGCTTTCTAGTCTTCTTCTTTTGCCAATTCCACTAATTGTTTGGG** | Ins_k3_27_CTA | -0.9608529548323955 | 0.04907945552225914 | 18.78401195268364 |
| **GATCCAGATCTAACTTGGGGTGGCTTTCTCGTCTTCTTCTTTTGCCAATTCCACTAATTGTTTGGG** | Ins_k3_27_CTC | 0.5375167146730977 | 0.030647320583314566 | 84.04696497727083 |
| **GATCCAGATCTAACTTGGGGTGGCTTTCTCTTCTTCTTTTGCCAATTCCACTAATTGTTTGGG** | G28C | -1.241199419094781 | 0.04919105240641613 | 14.191733052955795 |
| **GATCCAGATCTAACTTGGGGTGGCTTTCTGGTCTTCTTCTTTTGCCAATTCCACTAATTGTTTGGG** | Ins_k3_27_CTG | 0.3087441772345155 | 0.03727207991502908 | 66.86015585371916 |
| **GATCCAGATCTAACTTGGGGTGGCTTTCTGTCTTCTTCTTTTGCCAATTCCACTAATTGTTTGGG** | Ins_k2_26_TC;Ins_k2_27_CT | -0.5693684061239995 | 0.04363302045889716 | 27.784842235678354 |
| **GATCCAGATCTAACTTGGGGTGGCTTTCTTCTTCTTTTGCCAATTCCACTAATTGTTTGGG** | Del_k2_27-28;Del_k2_28-29 | -0.0020302993762241206 | 0.0341897253333698 | 49.0004134301117 |
| **GATCCAGATCTAACTTGGGGTGGCTTTCTTCTTTTGCCAATTCCACTAATTGTTTGGG** | Del_k5_26-30;Del_k5_27-31;Del_ | -0.14526818979693806 | 0.03426287098994279 | 42.46120540251884 |
| **GATCCAGATCTAACTTGGGGTGGCTTTCTTGTCTTCTTCTTTTGCCAATTCCACTAATTGTTTGGG** | Ins_k3_25_TTC;Ins_k3_26_TCT;I | -0.6967076800407956 | 0.05119558747124409 | 24.462745165320495 |
| **GATCCAGATCTAACTTGGGGTGGCTTTCTTTTGCCAATTCCACTAATTGTTTGGG** | Del_k8_26-33;Del_k8_27-34;Del_ | -0.6116667209829021 | 0.0413814616602672 | 26.63409904504192 |
| **GATCCAGATCTAACTTGGGGTGGCTTTG** | Del_k35_28-62;Del_k35_29-63 | -4.438534488927894 | 0.10659596496212888 | 0.5800299991380572 |
| **GATCCAGATCTAACTTGGGGTGGCTTTGAAATCTTCTTCTTTTGCCAATTCCACTAATTGTTTGGG** | Ins_k3_28_AAA | 0.4854326983596471 | 0.031234671507176914 | 79.78150679378781 |
| **GATCCAGATCTAACTTGGGGTGGCTTTGAACCTCTTCTTCTTTTGCCAATTCCACTAATTGTTTGGG** | Ins_k4_28_AACC | 0.6765135564123697 | 0.031087046088799027 | 96.58008796196026 |
| **GATCCAGATCTAACTTGGGGTGGCTTTGAACTCTTCTTCTTTTGCCAATTCCACTAATTGTTTGGG** | Ins_k3_28_AAC | 0.4564844802217749 | 0.030018358862291886 | 77.5050825130289 |
| **GATCCAGATCTAACTTGGGGTGGCTTTGAAGTCTTCTTCTTTTGCCAATTCCACTAATTGTTTGGG** | Ins_k3_27_GAA;Ins_k3_28_AAG | 0.6422771359886568 | 0.02925483848316296 | 93.32949333254238 |
| **GATCCAGATCTAACTTGGGGTGGCTTTGAATCTTCTTCTTTTGCCAATTCCACTAATTGTTTGGG** | Ins_k2_28_AA | 0.5653629996188556 | 0.03207122731580451 | 86.42025095674563 |
| **GATCCAGATCTAACTTGGGGTGGCTTTGAATTCCACTAATTGTTTGGG** | Del_k15_29-43 | -0.37901423866051864 | 0.030383639148808806 | 33.61071100700332 |
| **GATCCAGATCTAACTTGGGGTGGCTTTGAATTCTTCTTCTTTTGCCAATTCCACTAATTGTTTGGG** | Ins_k3_28_AAT | 0.43995660008748994 | 0.031872100588662154 | 76.23461578145447 |
| **GATCCAGATCTAACTTGGGGTGGCTTTGAATTGTTTGGG** | Del_k24_29-52 | -2.456607644387915 | 0.060159718529663 | 4.209110700782691 |
| **GATCCAGATCTAACTTGGGGTGGCTTTGACATCTTCTTCTTTTGCCAATTCCACTAATTGTTTGGG** | Ins_k3_28_ACA | 0.6118770390264896 | 0.030787966117463015 | 90.53495994319272 |
| **GATCCAGATCTAACTTGGGGTGGCTTTGACCTCTTCTTCTTTTGCCAATTCCACTAATTGTTTGGG** | Ins_k3_28_ACC | 0.6071801361942777 | 0.029159535495188998 | 90.11072311292625 |
| **GATCCAGATCTAACTTGGGGTGGCTTTGACGTCTTCTTCTTTTGCCAATTCCACTAATTGTTTGGG** | Ins_k3_27_GAC;Ins_k3_28_ACG | 0.7145960430956053 | 0.03056532771865094 | 100 |
| **GATCCAGATCTAACTTGGGGTGGCTTTGACTAATTGTTTGGG** | Del_k21_29-49 | -2.2479724586951946 | 0.057289882178173546 | 5.185605303316529 |
| **GATCCAGATCTAACTTGGGGTGGCTTTGACTATCTTCTTCTTTTGCCAATTCCACTAATTGTTTGGG** | Ins_k4_28_ACTA | 0.23207252894529606 | 0.03220080825099042 | 61.925469911616716 |
| **GATCCAGATCTAACTTGGGGTGGCTTTGACTCTTCTTCTTTTGCCAATTCCACTAATTGTTTGGG** | Ins_k2_28_AC | 0.4282240089621887 | 0.030551823996168948 | 75.34541273294433 |
| **GATCCAGATCTAACTTGGGGTGGCTTTGACTTCTTCTTCTTTTGCCAATTCCACTAATTGTTTGGG** | Ins_k3_28_ACT | 0.37029003644335445 | 0.0315315217759999 | 71.10438945379131 |
| **GATCCAGATCTAACTTGGGGTGGCTTTGACTTCTTCTTTTGCCAATTCCACTAATTGTTTGGG** | T29A | 0.3490975561373123 | 0.029269748130135994 | 69.61336612564466 |
| **GATCCAGATCTAACTTGGGGTGGCTTTGAGATCTTCTTCTTTTGCCAATTCCACTAATTGTTTGGG** | Ins_k3_28_AGA | 0.6456530361209663 | 0.029215544672422197 | 93.64509680459713 |
| **GATCCAGATCTAACTTGGGGTGGCTTTGAGCATCTTCTTCTTTTGCCAATTCCACTAATTGTTTGGG** | Ins_k4_28_AGCA | 0.852000038419066 | 0.030655564926640136 | 100 |
| **GATCCAGATCTAACTTGGGGTGGCTTTGAGCCTCTTCTTCTTTTGCCAATTCCACTAATTGTTTGGG** | Ins_k4_28_AGCC | 0.8779722643307049 | 0.0342011459239658 | 100 |
| **GATCCAGATCTAACTTGGGGTGGCTTTGAGCTCTTCTTCTTTTGCCAATTCCACTAATTGTTTGGG** | Ins_k3_28_AGC | 0.4250127335885354 | 0.03134162542539336 | 75.10384594085473 |
| **GATCCAGATCTAACTTGGGGTGGCTTTGAGGTCTTCTTCTTTTGCCAATTCCACTAATTGTTTGGG** | Ins_k3_27_GAG;Ins_k3_28_AGG | 0.28673129492029703 | 0.0323712773902488 | 65.40445201177359 |
| **GATCCAGATCTAACTTGGGGTGGCTTTGAGTCTTCTTCTTTTGCCAATTCCACTAATTGTTTGGG** | Ins_k2_27_GA;Ins_k2_28_AG | 0.6901362178165285 | 0.03423578067686071 | 97.90476814769553 |
| **GATCCAGATCTAACTTGGGGTGGCTTTGAGTTCTTCTTCTTTTGCCAATTCCACTAATTGTTTGGG** | Ins_k3_28_AGT | 0.39327530303469205 | 0.03522675096572135 | 72.75767056660801 |
| **GATCCAGATCTAACTTGGGGTGGCTTTGATATCTTCTTCTTTTGCCAATTCCACTAATTGTTTGGG** | Ins_k3_28_ATA | 0.2662111473854968 | 0.03673457404079565 | 64.07601943724832 |
| **GATCCAGATCTAACTTGGGGTGGCTTTGATCTCTTCTTCTTTTGCCAATTCCACTAATTGTTTGGG** | Ins_k3_28_ATC | 0.2137894943459806 | 0.03266781837179211 | 60.80357150749398 |
| **GATCCAGATCTAACTTGGGGTGGCTTTGATCTTCTTCTTTTGCCAATTCCACTAATTGTTTGGG** | Ins_k1_28_A | 0.6730918619845327 | 0.02892464357673975 | 96.2501851483253 |
| **GATCCAGATCTAACTTGGGGTGGCTTTGATGTCTTCTTCTTTTGCCAATTCCACTAATTGTTTGGG** | Ins_k3_26_TGA;Ins_k3_27_GAT;I | 0.6530125390466848 | 0.030075253939794728 | 94.33682041737774 |
| **GATCCAGATCTAACTTGGGGTGGCTTTGATTCCACTAATTGTTTGGG** | Del_k16_29-44 | -0.4844059225386069 | 0.03385198457433293 | 30.248697093556515 |
| **GATCCAGATCTAACTTGGGGTGGCTTTGATTCTTCTTCTTTTGCCAATTCCACTAATTGTTTGGG** | Ins_k2_28_AT | 0.376824723674554 | 0.03495540154219357 | 71.57055586663272 |
| **GATCCAGATCTAACTTGGGGTGGCTTTGATTGTTTGGG** | Del_k25_29-53 | -2.6676367022836316 | 0.06746228056393308 | 3.408329649799822 |
| **GATCCAGATCTAACTTGGGGTGGCTTTGATTTCTTCTTCTTTTGCCAATTCCACTAATTGTTTGGG** | Ins_k3_28_ATT | 0.048288369212106 | 0.04219707688040152 | 51.529136450617095 |
| **GATCCAGATCTAACTTGGGGTGGCTTTGCAATCTTCTTCTTTTGCCAATTCCACTAATTGTTTGGG** | Ins_k3_28_CAA | 0.040899647949962925 | 0.03689631531575546 | 51.149805136917024 |
| **GATCCAGATCTAACTTGGGGTGGCTTTGCAATTCCACTAATTGTTTGGG** | Del_k14_29-42 | -0.6490087546441907 | 0.030790610859873172 | 25.65786828061107 |
| **GATCCAGATCTAACTTGGGGTGGCTTTGCACTAATTGTTTGGG** | Del_k20_29-48 | -1.6061808177810954 | 0.041075858297093384 | 9.852036814639023 |
| **GATCCAGATCTAACTTGGGGTGGCTTTGCACTCTTCTTCTTTTGCCAATTCCACTAATTGTTTGGG** | Ins_k3_28_CAC | 0.10123368641056496 | 0.03360308022091444 | 54.330878015079186 |
| **GATCCAGATCTAACTTGGGGTGGCTTTGCAGTCTTCTTCTTTTGCCAATTCCACTAATTGTTTGGG** | Ins_k3_27_GCA;Ins_k3_28_CAG | 0.5576965157097842 | 0.031135922754339147 | 85.76024468973199 |
| **GATCCAGATCTAACTTGGGGTGGCTTTGCATCTTCTTCTTTTGCCAATTCCACTAATTGTTTGGG** | Ins_k2_28_CA | 0.4390455569125642 | 0.03244785974645715 | 76.16519438280105 |
| **GATCCAGATCTAACTTGGGGTGGCTTTGCATTCTTCTTCTTTTGCCAATTCCACTAATTGTTTGGG** | Ins_k3_28_CAT | -0.010008080680455333 | 0.03588453278876664 | 48.61105402510289 |
| **GATCCAGATCTAACTTGGGGTGGCTTTGCCAATTCCACTAATTGTTTGGG** | Del_k13_25-37;Del_k13_26-38;D | -0.5847147755536958 | 0.03110618442811792 | 27.361700928102692 |
| **GATCCAGATCTAACTTGGGGTGGCTTTGCCACTAATTGTTTGGG** | Del_k19_29-47 | -2.930760161841661 | 0.08988800091655791 | 2.619802342043045 |
| **GATCCAGATCTAACTTGGGGTGGCTTTGCCAGTCTTCTTCTTTTGCCAATTCCACTAATTGTTTGGG** | Ins_k4_27_GCCA;Ins_k4_28_CC | 0.7320932218524305 | 0.028510602565767727 | 100 |
| **GATCCAGATCTAACTTGGGGTGGCTTTGCCATCTTCTTCTTTTGCCAATTCCACTAATTGTTTGGG** | Ins_k3_28_CCA | -0.2118252622438062 | 0.037743130466178186 | 39.72710786974017 |
| **GATCCAGATCTAACTTGGGGTGGCTTTGCCCTCTTCTTCTTTTGCCAATTCCACTAATTGTTTGGG** | Ins_k3_28_CCC | -0.047180593584772335 | 0.036856275016753946 | 46.837231954246064 |
| **GATCCAGATCTAACTTGGGGTGGCTTTGCCGTCTTCTTCTTTTGCCAATTCCACTAATTGTTTGGG** | Ins_k3_27_GCC;Ins_k3_28_CCG | 0.719862592725274 | 0.02849360308935486 | 100 |
| **GATCCAGATCTAACTTGGGGTGGCTTTGCCTCTTCTTCTTTTGCCAATTCCACTAATTGTTTGGG** | Ins_k2_28_CC | 0.24585776166926254 | 0.037797029966106 | 62.78503798674535 |
| **GATCCAGATCTAACTTGGGGTGGCTTTGCCTTCTTCTTCTTTTGCCAATTCCACTAATTGTTTGGG** | Ins_k3_28_CCT | 0.17414914566942974 | 0.03196565609189871 | 58.44044377093291 |
| **GATCCAGATCTAACTTGGGGTGGCTTTGCCTTCTTCTTTTGCCAATTCCACTAATTGTTTGGG** | T29C | -0.35504915316704944 | 0.0250614994148923 | 34.425923876765985 |
| **GATCCAGATCTAACTTGGGGTGGCTTTGCGATCTTCTTCTTTTGCCAATTCCACTAATTGTTTGGG** | Ins_k3_28_CGA | 0.6986462508174778 | 0.028516749857453918 | 98.74149619754742 |
| **GATCCAGATCTAACTTGGGGTGGCTTTGCGCTCTTCTTCTTTTGCCAATTCCACTAATTGTTTGGG** | Ins_k3_28_CGC | 0.2546596561184056 | 0.0312503574771027 | 63.340104498949415 |
| **GATCCAGATCTAACTTGGGGTGGCTTTGCGGTCTTCTTCTTTTGCCAATTCCACTAATTGTTTGGG** | Ins_k3_27_GCG;Ins_k3_28_CGG | 0.5581082575294798 | 0.029793918822888628 | 85.7955630394599 |
| **GATCCAGATCTAACTTGGGGTGGCTTTGCGTCTTCTTCTTTTGCCAATTCCACTAATTGTTTGGG** | Ins_k2_27_GC;Ins_k2_28_CG | 0.622208460351179 | 0.031682761429365436 | 91.47516321408354 |
| **GATCCAGATCTAACTTGGGGTGGCTTTGCGTTCTTCTTCTTTTGCCAATTCCACTAATTGTTTGGG** | Ins_k3_28_CGT | 0.2767772139983028 | 0.032350525093364065 | 64.75664033766029 |
| **GATCCAGATCTAACTTGGGGTGGCTTTGCTAATTGTTTGGG** | Del_k22_29-50 | -2.8136888881356734 | 0.06612506707150755 | 2.9451806210965352 |
| **GATCCAGATCTAACTTGGGGTGGCTTTGCTATCTTCTTCTTTTGCCAATTCCACTAATTGTTTGGG** | Ins_k3_28_CTA | -0.7746892587831707 | 0.049905185642564805 | 22.62758587827107 |
| **GATCCAGATCTAACTTGGGGTGGCTTTGCTCTCTTCTTCTTTTGCCAATTCCACTAATTGTTTGGG** | Ins_k3_28_CTC | -0.5811206425272641 | 0.03933569907289608 | 27.460219459367725 |

| | | | | |
|---|---|---|---|---|
| **GATCCAGATCTAACTTGGGGTGGCTTTGCTCTTCTTCTTTTGCCAATTCCACTAATTGTTTGGG** | Ins_k1_28_C | 0.14711432167537775 | 0.035451339886914505 | 56.881682022261124 |
| **GATCCAGATCTAACTTGGGGTGGCTTTGCTGATCTTCTTCTTTTGCCAATTCCACTAATTGTTTGGG** | Ins_k4_28_CTGA | 0.8994938188928585 | 0.029906831251831974 | 100 |
| **GATCCAGATCTAACTTGGGGTGGCTTTGCTGTCTTCTTCTTTTGCCAATTCCACTAATTGTTTGGG** | Ins_k3_26_TGC;Ins_k3_27_GCT; | 0.4869160799507475 | 0.029667657392660564 | 79.89994103214333 |
| **GATCCAGATCTAACTTGGGGTGGCTTTGCTTCTTCTTCTTTTGCCAATTCCACTAATTGTTTGGG** | Ins_k2_28_CT | -0.011052639951153909 | 0.03794469077496259 | 48.560303408590535 |
| **GATCCAGATCTAACTTGGGGTGGCTTTGCTTCTTCTTTTGCCAATTCCACTAATTGTTTGGG** | Del_k1_29-29 | -0.09798803821280944 | 0.03334465886824482 | 44.51699367393738 |
| **GATCCAGATCTAACTTGGGGTGGCTTTGCTTCTTTTGCCAATTCCACTAATTGTTTGGG** | Del_k4_29-32 | -0.18752092588263675 | 0.035918892710468144 | 40.70447791661783 |
| **GATCCAGATCTAACTTGGGGTGGCTTTGCTTTCTTCTTCTTTTGCCAATTCCACTAATTGTTTGGG** | Ins_k3_28_CTT | -1.1377605673656142 | 0.06182644388369886 | 15.738319442496628 |
| **GATCCAGATCTAACTTGGGGTGGCTTTGCTTTTGCCAATTCCACTAATTGTTTGGG** | Del_k7_29-35 | -0.23330230942043484 | 0.03522164508099139 | 38.882983990732626 |
| **GATCCAGATCTAACTTGGGGTGGCTTTGG** | Del_k34_28-61;Del_k34_29-62 | -4.37265366380775 | 0.10933831682542708 | 0.6195297050592163 |
| **GATCCAGATCTAACTTGGGGTGGCTTTGGAATCTTCTTCTTTTGCCAATTCCACTAATTGTTTGGG** | Ins_k3_28_GAA | 0.5169240514725135 | 0.028910127639542166 | 82.33391282315293 |
| **GATCCAGATCTAACTTGGGGTGGCTTTGGACTCTTCTTCTTTTGCCAATTCCACTAATTGTTTGGG** | Ins_k3_28_GAC | 0.5422626985383221 | 0.028173644080459345 | 84.4467985683606 |
| **GATCCAGATCTAACTTGGGGTGGCTTTGGAGTCTTCTTCTTTTGCCAATTCCACTAATTGTTTGGG** | Ins_k3_27_GGA;Ins_k3_28_GAG | 0.5752536803685586 | 0.02880680920091935 | 87.27924709491478 |
| **GATCCAGATCTAACTTGGGGTGGCTTTGGATCTTCTTCTTTTGCCAATTCCACTAATTGTTTGGG** | Ins_k2_28_GA | 0.7874044812460035 | 0.02888557027738655 | 100 |
| **GATCCAGATCTAACTTGGGGTGGCTTTGGATTCTTCTTCTTTTGCCAATTCCACTAATTGTTTGGG** | Ins_k3_28_GAT | 0.6895992722273363 | 0.030584889494045368 | 97.85221272524159 |
| **GATCCAGATCTAACTTGGGGTGGCTTTGGCATCTTCTTCTTTTGCCAATTCCACTAATTGTTTGGG** | Ins_k3_28_GCA | -2.7781470319752968 | 0.10038843786018567 | 3.0517402535116034 |
| **GATCCAGATCTAACTTGGGGTGGCTTTGGCCAATTCCACTAATTGTTTGGG** | Del_k12_29-40 | -0.47449931580296956 | 0.03207885453560753 | 30.54984827013277 |
| **GATCCAGATCTAACTTGGGGTGGCTTTGGCCTCTTCTTCTTTTGCCAATTCCACTAATTGTTTGGG** | Ins_k3_28_GCC | -0.25382800573531344 | 0.0370813089214524 | 38.09301869782055 |
| **GATCCAGATCTAACTTGGGGTGGCTTTGGCGTCTTCTTCTTTTGCCAATTCCACTAATTGTTTGGG** | Ins_k3_27_GGC;Ins_k3_28_GCG | -0.37655730315926605 | 0.04188086335273017 | 33.6933918852992 |
| **GATCCAGATCTAACTTGGGGTGGCTTTGGCTCTTCTTCTTTTGCCAATTCCACTAATTGTTTGGG** | Ins_k2_28_GC | -1.3692375715123006 | 0.0645515973321671 | 12.48616789598351 |
| **GATCCAGATCTAACTTGGGGTGGCTTTGGCTTCTTCTTCTTTTGCCAATTCCACTAATTGTTTGGG** | Ins_k3_28_GCT | -1.361844106681814 | 0.05991679124252312 | 12.578826049506523 |
| **GATCCAGATCTAACTTGGGGTGGCTTTGGCTTCTTCTTTTGCCAATTCCACTAATTGTTTGGG** | T29G | -2.0601854567533207 | 0.07590767951211323 | 6.256829442955131 |
| **GATCCAGATCTAACTTGGGGTGGCTTTGGG** | Del_k33_25-57;Del_k33_26-58;D | -4.506552325262248 | 0.12424101996304358 | 0.5418894362763463 |
| **GATCCAGATCTAACTTGGGGTGGCTTTGGGATCTTCTTCTTTTGCCAATTCCACTAATTGTTTGGG** | Ins_k3_28_GGA | -0.03779435345607851 | 0.0381173158689996 | 47.27892714606957 |
| **GATCCAGATCTAACTTGGGGTGGCTTTGGGCTCTTCTTCTTTTGCCAATTCCACTAATTGTTTGGG** | Ins_k3_28_GGC | -2.095605518894895 | 0.07680018541025034 | 6.039091072308217 |
| **GATCCAGATCTAACTTGGGGTGGCTTTGGGG** | Del_k32_29-60 | -5.229642556744697 | 0.22851124318736438 | 0.26295206600694 |
| **GATCCAGATCTAACTTGGGGTGGCTTTGGGGTCTTCTTCTTTTGCCAATTCCACTAATTGTTTGGG** | Ins_k3_27_GGG;Ins_k3_28_GGG | -3.484562025215677 | 0.1780335087957135 | 1.5057588788965324 |
| **GATCCAGATCTAACTTGGGGTGGCTTTGGGTCTTCTTCTTTTGCCAATTCCACTAATTGTTTGGG** | Ins_k2_27_GG;Ins_k2_28_GG | -2.117839076705617 | 0.10427104326681039 | 5.9063022459437065 |
| **GATCCAGATCTAACTTGGGGTGGCTTTGGGTTCTTCTTCTTTTGCCAATTCCACTAATTGTTTGGG** | Ins_k3_28_GGT | -3.100953535832517 | 0.1683932090578424 | 2.209807701765322 |
| **GATCCAGATCTAACTTGGGGTGGCTTTGGTATCTTCTTCTTTTGCCAATTCCACTAATTGTTTGGG** | Ins_k3_28_GTA | -0.9506898607376368 | 0.05158068421866796 | 18.975889014397563 |
| **GATCCAGATCTAACTTGGGGTGGCTTTGGTCTCTTCTTCTTTTGCCAATTCCACTAATTGTTTGGG** | Ins_k3_28_GTC | -0.4551666765281083 | 0.0396281601401813 | 31.14620345254139 |
| **GATCCAGATCTAACTTGGGGTGGCTTTGGTCTTCTTCTTTTGCCAATTCCACTAATTGTTTGGG** | Ins_k1_27_G;Ins_k1_28_G | -0.13759148989034276 | 0.03605410125634948 | 42.78842169776782[4] |
| **GATCCAGATCTAACTTGGGGTGGCTTTGGTGTCTTCTTCTTTTGCCAATTCCACTAATTGTTTGGG** | Ins_k3_26_TGG;Ins_k3_27_GGT; | 0.09095269429461161 | 0.032018016397059354 | 53.77516422594379 |
| **GATCCAGATCTAACTTGGGGTGGCTTTGGTTCTTCTTCTTTTGCCAATTCCACTAATTGTTTGGG** | Ins_k2_28_GT | -0.2617372877831895 | 0.03856072891037104 | 37.792918621396616 |
| **GATCCAGATCTAACTTGGGGTGGCTTTGGTTTCTTCTTCTTTTGCCAATTCCACTAATTGTTTGGG** | Ins_k3_28_GTT | -1.0323065935807711 | 0.04685491455142889 | 17.488655983432015 |
| **GATCCAGATCTAACTTGGGGTGGCTTTGGTTTGGG** | Del_k28_29-56 | -4.239427388475367 | 0.12906565882198456 | 0.7078179481923643 |
| **GATCCAGATCTAACTTGGGGTGGCTTTGT** | Del_k34_30-63 | -6.2459097314397045 | 0.8891867780577029 | 0.09517378938877537 |
| **GATCCAGATCTAACTTGGGGTGGCTTTGTAAACTTCTTCTTTTGCCAATTCCACTAATTGTTTGGG** | Ins_k3_29_AAA | -0.5971351266978572 | 0.04145373578491832 | 27.023960762185062 |
| **GATCCAGATCTAACTTGGGGTGGCTTTGTAACCTTCTTCTTTTGCCAATTCCACTAATTGTTTGGG** | Ins_k3_29_AAC | -0.4141649855817692 | 0.04851434646662615 | 32.44979251758961 |
| **GATCCAGATCTAACTTGGGGTGGCTTTGTAACTTCTTCTTTTGCCAATTCCACTAATTGTTTGGG** | Ins_k2_29_AA | -0.5587454148432427 | 0.044224738259632544 | 28.081573669747822 |
| **GATCCAGATCTAACTTGGGGTGGCTTTGTAAGCTTCTTCTTTTGCCAATTCCACTAATTGTTTGGG** | Ins_k3_29_AAG | -0.18675323543779904 | 0.037606137455702626 | 40.735738353010404 |
| **GATCCAGATCTAACTTGGGGTGGCTTTGTAATCTTCTTCTTTTGCCAATTCCACTAATTGTTTGGG** | Ins_k3_28_TAA;Ins_k3_29_AAT | -0.5818719488541825 | 0.05376852089577231 | 27.439596170922968 |
| **GATCCAGATCTAACTTGGGGTGGCTTTGTAATTCCACTAATTGTTTGGG** | Del_k14_30-43 | -1.6070332110804857 | 0.047474393654309305 | 9.843642582574713 |
| **GATCCAGATCTAACTTGGGGTGGCTTTGTAATTGTTTGGG** | Del_k23_29-51;Del_k23_30-52 | -3.7879592886056477 | 0.1405210635143154 | 1.111710417719914 |
| **GATCCAGATCTAACTTGGGGTGGCTTTGTACACCTTCTTCTTTTGCCAATTCCACTAATTGTTTGGG** | Ins_k4_29_ACAC | -0.02922559828761473 | 0.03988117197414657 | 47.68578935932173 |
| **GATCCAGATCTAACTTGGGGTGGCTTTGTACACTTCTTCTTTTGCCAATTCCACTAATTGTTTGGG** | Ins_k3_29_ACA | -0.2032408714502853 | 0.04243810687496067 | 40.06960886670359 |
| **GATCCAGATCTAACTTGGGGTGGCTTTGTACCCTTCTTCTTTTGCCAATTCCACTAATTGTTTGGG** | Ins_k3_29_ACC | 0.11559948544268872 | 0.03915087667241456 | 55.11701773248205[4] |
| **GATCCAGATCTAACTTGGGGTGGCTTTGTACCTTCTTCTTTTGCCAATTCCACTAATTGTTTGGG** | Ins_k2_29_AC | 0.3161234214001605 | 0.03619432046086182 | 67.35535813086186 |
| **GATCCAGATCTAACTTGGGGTGGCTTTGTACGCTTCTTCTTTTGCCAATTCCACTAATTGTTTGGG** | Ins_k3_29_ACG | 0.391767763002953 | 0.03456995632183861 | 72.64806810142228 |
| **GATCCAGATCTAACTTGGGGTGGCTTTGTACGGCTTCTTCTTTTGCCAATTCCACTAATTGTTTGGG** | Ins_k4_29_ACGG | 0.7301779170814799 | 0.03477524135284668 | 100 |
| **GATCCAGATCTAACTTGGGGTGGCTTTGTACTAATTGTTTGGG** | Del_k20_30-49 | -3.311844282565348 | 0.0890943857609292 | 1.7896404830734727 |
| **GATCCAGATCTAACTTGGGGTGGCTTTGTACTCTTCTTCTTTTGCCAATTCCACTAATTGTTTGGG** | Ins_k3_28_TAC;Ins_k3_29_ACT | -0.4071603772574679 | 0.04041545083482661 | 32.67788853338762 |
| **GATCCAGATCTAACTTGGGGTGGCTTTGTACTTCTTCTTTTGCCAATTCCACTAATTGTTTGGG** | Ins_k1_29_A | 0.023445348159427226 | 0.04060744369774782 | 50.26476742953614 |
| **GATCCAGATCTAACTTGGGGTGGCTTTGTAGACTTCTTCTTTTGCCAATTCCACTAATTGTTTGGG** | Ins_k3_29_AGA | 0.02693550978831627 | 0.03245278795663541 | 50.440506091879065 |
| **GATCCAGATCTAACTTGGGGTGGCTTTGTAGCCTTCTTCTTTTGCCAATTCCACTAATTGTTTGGG** | Ins_k3_29_AGC | 0.07457795852651983 | 0.032948402525985084 | 52.90178035103721 |
| **GATCCAGATCTAACTTGGGGTGGCTTTGTAGCTTCTTCTTTTGCCAATTCCACTAATTGTTTGGG** | Ins_k2_29_AG | -0.8208706383766521 | 0.05157939337056351 | 21.60637470593138 |
| **GATCCAGATCTAACTTGGGGTGGCTTTGTAGGCTTCTTCTTTTGCCAATTCCACTAATTGTTTGGG** | Ins_k3_29_AGG | -1.5392535548035435 | 0.06713041163725596 | 10.533972179065758 |
| **GATCCAGATCTAACTTGGGGTGGCTTTGTAGTCTTCTTCTTTTGCCAATTCCACTAATTGTTTGGG** | Ins_k3_27_GTA;Ins_k3_28_TAG;I | -0.9274629120551909 | 0.0506169365066714 | 19.421799538281697 |
| **GATCCAGATCTAACTTGGGGTGGCTTTGTATACTTCTTCTTTTGCCAATTCCACTAATTGTTTGGG** | Ins_k3_29_ATA | -0.5341128368588836 | 0.04690409229914047 | 28.781885050494008 |
| **GATCCAGATCTAACTTGGGGTGGCTTTGTATCCTTCTTCTTTTGCCAATTCCACTAATTGTTTGGG** | Ins_k3_29_ATC | -0.31605431456886013 | 0.03853197062364955 | 35.79487476724025 |
| **GATCCAGATCTAACTTGGGGTGGCTTTGTATCTTCTTCTTTTGCCAATTCCACTAATTGTTTGGG** | Ins_k2_28_TA;Ins_k2_29_AT | -0.3578215981811036 | 0.04362372403849934 | 34.330612080168045 |
| **GATCCAGATCTAACTTGGGGTGGCTTTGTATGCTTCTTCTTTTGCCAATTCCACTAATTGTTTGGG** | Ins_k3_29_ATG | -0.2810010803998864 | 0.0332751290219685 | 37.071851219632734 |
| **GATCCAGATCTAACTTGGGGTGGCTTTGTATTCCACTAATTGTTTGGG** | Del_k15_30-44 | -1.440098096459257 | 0.04383179498906852 | 11.632011836148255 |
| **GATCCAGATCTAACTTGGGGTGGCTTTGTATTCTTCTTCTTTTGCCAATTCCACTAATTGTTTGGG** | Ins_k3_28_TAT;Ins_k3_29_ATT | -0.8562160875653709 | 0.05058303406335988 | 20.856026499731843 |
| **GATCCAGATCTAACTTGGGGTGGCTTTGTATTCTTCTTTTGCCAATTCCACTAATTGTTTGGG** | C30A | -0.9555270558833153 | 0.04416793552622221 | 18.884320581907353 |
| **GATCCAGATCTAACTTGGGGTGGCTTTGTATTGTTTGGG** | Del_k24_30-53 | -3.6534977430064575 | 0.10438980691417893 | 1.2717085251867553 |
| **GATCCAGATCTAACTTGGGGTGGCTTTGTC** | Del_k33_31-63 | -5.863417256397815 | 0.41861158298223 | 0.13951847885300261 |
| **GATCCAGATCTAACTTGGGGTGGCTTTGTCAAATTCTTCTTTTGCCAATTCCACTAATTGTTTGGG** | Ins_k3_30_AAA | -0.16422754117702743 | 0.038170645738204216 | 41.66375197650212 |
| **GATCCAGATCTAACTTGGGGTGGCTTTGTCAACTTCTTCTTTTGCCAATTCCACTAATTGTTTGGG** | Ins_k3_29_CAA;Ins_k3_30_AAC | 0.3348066829891453 | 0.030432080986351123 | 68.62560513516792 |
| **GATCCAGATCTAACTTGGGGTGGCTTTGTCAAGTTCTTCTTTTGCCAATTCCACTAATTGTTTGGG** | Ins_k3_30_AAG | -0.007547256812675651 | 0.03375807776993257 | 48.73082457375743 |
| **GATCCAGATCTAACTTGGGGTGGCTTTGTCAATTCCACTAATTGTTTGGG** | Del_k13_30-42;Del_k13_31-43 | -0.5183211858108224 | 0.03166802680208165 | 29.240006260520772 |
| **GATCCAGATCTAACTTGGGGTGGCTTTGTCAATTCTTCTTTTGCCAATTCCACTAATTGTTTGGG** | Ins_k2_30_AA | 0.020570515394320865 | 0.04033409228346239 | 50.120472140990266 |
| **GATCCAGATCTAACTTGGGGTGGCTTTGTCAATTGTTTGGG** | Del_k22_31-52 | -2.0104386807726873 | 0.05487428700840056 | 6.575958546827879 |
| **GATCCAGATCTAACTTGGGGTGGCTTTGTCAATTTCTTCTTTTGCCAATTCCACTAATTGTTTGGG** | Ins_k3_30_AAT | -0.4783824643639626 | 0.039390208413127306 | 30.431448701119404 |
| **GATCCAGATCTAACTTGGGGTGGCTTTGTCACATTCTTCTTTTGCCAATTCCACTAATTGTTTGGG** | Ins_k3_30_ACA | -0.07357237340395184 | 0.03953263312523154 | 45.61728316082973 |
| **GATCCAGATCTAACTTGGGGTGGCTTTGTCACCCTTCTTCTTTTGCCAATTCCACTAATTGTTTGGG** | Ins_k4_29_CACC;Ins_k4_30_AC | -1.2745415275659155 | 0.06585077086387715 | 13.726352251400497 |

| | | | | |
|---|---|---|---|---|
| GATCCAGATCTAACTTGGGGTGGCTTTGTCACCTTCTTCTTTTGCCAATTCCACTAATTGTTTGGG | Ins_k3_29_CAC;Ins_k3_30_ACC | -0.606724054530729 | 0.046806603031093255 | 26.766068384422006 |
| GATCCAGATCTAACTTGGGGTGGCTTTGTCACGTTCTTCTTTTGCCAATTCCACTAATTGTTTGGG | Ins_k3_30_ACG | 0.5088527901168705 | 0.03053602274615455 | 81.6720489256778 |
| GATCCAGATCTAACTTGGGGTGGCTTTGTCACTAATTGTTTGGG | Del_k19_30-48;Del_k19_31-49 | -1.6527873479135589 | 0.05344983995300982 | 9.403403393516 |
| GATCCAGATCTAACTTGGGGTGGCTTTGTCACTATTCTTCTTTTGCCAATTCCACTAATTGTTTGGG | Ins_k4_30_ACTA | -0.2297340190351177 | 0.03983078333788428 | 39.021977605969376 |
| GATCCAGATCTAACTTGGGGTGGCTTTGTCACTGTTCTTCTTTTGCCAATTCCACTAATTGTTTGGG | Ins_k4_30_ACTG | 0.2258455439952597 | 0.037785384888600976 | 61.541059044931636 |
| GATCCAGATCTAACTTGGGGTGGCTTTGTCACTTCTTCTTTTGCCAATTCCACTAATTGTTTGGG | Ins_k2_29_CA;Ins_k2_30_AC | -0.0744112034280433 | 0.03617318755423458 | 45.57903405858741 |
| GATCCAGATCTAACTTGGGGTGGCTTTGTCACTTTCTTCTTTTGCCAATTCCACTAATTGTTTGGG | Ins_k3_30_ACT | -0.49825377750387906 | 0.04590096970354517 | 29.83270447369755 |
| GATCCAGATCTAACTTGGGGTGGCTTTGTCAGATTCTTCTTTTGCCAATTCCACTAATTGTTTGGG | Ins_k3_30_AGA | 0.3748683917400636 | 0.030168080260665166 | 71.4306769720083 |
| GATCCAGATCTAACTTGGGGTGGCTTTGTCAGCTTCTTCTTTTGCCAATTCCACTAATTGTTTGGG | Ins_k3_29_CAG;Ins_k3_30_AGC | 0.47383633989118357 | 0.029588214185061606 | 78.86167549645704 |
| GATCCAGATCTAACTTGGGGTGGCTTTGTCAGGTTCTTCTTTTGCCAATTCCACTAATTGTTTGGG | Ins_k3_30_AGG | -1.2499815082257089 | 0.0645410886322908 | 14.0676456591585 |
| GATCCAGATCTAACTTGGGGTGGCTTTGTCAGTTCTTCTTTTGCCAATTCCACTAATTGTTTGGG | Ins_k2_30_AG | 0.12034539216238804 | 0.03725529439229647 | 55.37921965810404 |
| GATCCAGATCTAACTTGGGGTGGCTTTGTCAGTTTCTTCTTTTGCCAATTCCACTAATTGTTTGGG | Ins_k3_30_AGT | -0.14317370336252344 | 0.03845417871595169 | 42.55023302224814 |
| GATCCAGATCTAACTTGGGGTGGCTTTGTCAGTTTTCTTCTTTTGCCAATTCCACTAATTGTTTGGG | Ins_k4_30_AGTT | -0.6223116062083235 | 0.04839396857046776 | 26.35208578003337 |
| GATCCAGATCTAACTTGGGGTGGCTTTGTCATATTCTTCTTTTGCCAATTCCACTAATTGTTTGGG | Ins_k3_30_ATA | -0.2599989519000512 | 0.04204482719039506 | 37.85867254260707 |
| GATCCAGATCTAACTTGGGGTGGCTTTGTCATCATTCTTCTTTTGCCAATTCCACTAATTGTTTGGG | Ins_k4_30_ATCA | 0.6797127843358322 | 0.0326791667664035 | 96.88956445517887 |
| GATCCAGATCTAACTTGGGGTGGCTTTGTCATCTTCTTCTTTTGCCAATTCCACTAATTGTTTGGG | Ins_k3_28_TCA;Ins_k3_29_CAT;I | 0.27169517021746337 | 0.0351815489256264 | 64.42837908190548 |
| GATCCAGATCTAACTTGGGGTGGCTTTGTCATCTTCTTTTGCCAATTCCACTAATTGTTTGGG | T31A | 0.13372472783399336 | 0.030100136402465363 | 56.12513563027018 |
| GATCCAGATCTAACTTGGGGTGGCTTTGTCATGTTCTTCTTTTGCCAATTCCACTAATTGTTTGGG | Ins_k3_30_ATG | 0.09267852428331746 | 0.037613642103978134 | 53.86805114747465 |
| GATCCAGATCTAACTTGGGGTGGCTTTGTCATTCCACTAATTGTTTGGG | Del_k14_31-44 | -2.155511144056675 | 0.06707073012631126 | 5.687938559937218 |
| GATCCAGATCTAACTTGGGGTGGCTTTGTCATTCTTCTTTTGCCAATTCCACTAATTGTTTGGG | Ins_k1_30_A | 0.0943881206716023 | 0.03535516644204297 | 53.96022253864978 |
| GATCCAGATCTAACTTGGGGTGGCTTTGTCATTGTTTGGG | Del_k23_31-53 | -1.7643806111856017 | 0.051006214137044974 | 8.410478960026607 |
| GATCCAGATCTAACTTGGGGTGGCTTTGTCATTTCTTCTTTTGCCAATTCCACTAATTGTTTGGG | Ins_k2_30_AT | -0.17455985965816256 | 0.04463696071730103 | 41.23548512673185 |
| GATCCAGATCTAACTTGGGGTGGCTTTGTCATTTTCTTCTTTTGCCAATTCCACTAATTGTTTGGG | Ins_k3_30_ATT | -0.6204790228426761 | 0.06056947997164069 | 26.40042245104593 |
| GATCCAGATCTAACTTGGGGTGGCTTTGTCCAATTCCACTAATTGTTTGGG | Del_k12_30-41;Del_k12_31-42 | -0.2605572766714401 | 0.03157047246793205 | 37.837541007593295 |
| GATCCAGATCTAACTTGGGGTGGCTTTGTCCAATTCTTCTTTTGCCAATTCCACTAATTGTTTGGG | Ins_k3_30_CAA | 0.2860525907210185 | 0.03532352121106175 | 65.36007679606772 |
| GATCCAGATCTAACTTGGGGTGGCTTTGTCCACTAATTGTTTGGG | Del_k18_29-46;Del_k18_30-47;D | -2.8309207030995176 | 0.09086689511490015 | 2.894864577024539 |
| GATCCAGATCTAACTTGGGGTGGCTTTGTCCACTTCTTCTTTTGCCAATTCCACTAATTGTTTGGG | Ins_k3_29_CCA;Ins_k3_30_CAC | -1.0057437252358223 | 0.053364065965166185 | 17.95942971961735 |
| GATCCAGATCTAACTTGGGGTGGCTTTGTCCAGTTCTTCTTTTGCCAATTCCACTAATTGTTTGGG | Ins_k3_30_CAG | 0.5722535232155341 | 0.028855458199926395 | 87.01778804267009 |
| GATCCAGATCTAACTTGGGGTGGCTTTGTCCATTCTTCTTTTGCCAATTCCACTAATTGTTTGGG | Ins_k2_30_CA | 0.21100949318317808 | 0.03746422790890972 | 60.63477224778216 |
| GATCCAGATCTAACTTGGGGTGGCTTTGTCCATTTCTTCTTTTGCCAATTCCACTAATTGTTTGGG | Ins_k3_30_CAT | 0.1214819233910156 | 0.037103163535401266 | 55.44219565097492 |
| GATCCAGATCTAACTTGGGGTGGCTTTGTCCCAATTCCACTAATTGTTTGGG | Del_k11_31-41 | -0.42655086190072544 | 0.03542945092584654 | 32.05035221647025 |
| GATCCAGATCTAACTTGGGGTGGCTTTGTCCCACTAATTGTTTGGG | Del_k17_31-47 | -1.8332728560387372 | 0.044921462559509975 | 7.850570299409262 |
| GATCCAGATCTAACTTGGGGTGGCTTTGTCCCATTCTTCTTTTGCCAATTCCACTAATTGTTTGGG | Ins_k3_30_CCA | -0.03964294770796206 | 0.04028940350380115 | 47.1916083265123 |
| GATCCAGATCTAACTTGGGGTGGCTTTGTCCCCTTCTTCTTTTGCCAATTCCACTAATTGTTTGGG | Ins_k3_29_CCC;Ins_k3_30_CCC | -0.07742729906585988 | 0.036603223477429274 | 45.44177043694798 |
| GATCCAGATCTAACTTGGGGTGGCTTTGTCCCGTTCTTCTTTTGCCAATTCCACTAATTGTTTGGG | Ins_k3_30_CCG | 0.6684013269437128 | 0.028752910390971206 | 95.79977743457285 |
| GATCCAGATCTAACTTGGGGTGGCTTTGTCCCTATTCTTCTTTTGCCAATTCCACTAATTGTTTGGG | Ins_k4_30_CCTA | -0.2758508910854716 | 0.04146829025335921 | 37.263270772008255 |
| GATCCAGATCTAACTTGGGGTGGCTTTGTCCCTTCTTCTTTTGCCAATTCCACTAATTGTTTGGG | Ins_k2_29_CC;Ins_k2_30_CC | 0.20132001378655717 | 0.03948636367712492 | 60.050090078732985 |
| GATCCAGATCTAACTTGGGGTGGCTTTGTCCCTTTCTTCTTTTGCCAATTCCACTAATTGTTTGGG | Ins_k3_30_CCT | -0.6064759191539774 | 0.044147476071270175 | 26.772710816961897 |
| GATCCAGATCTAACTTGGGGTGGCTTTGTCCGATTCTTCTTTTGCCAATTCCACTAATTGTTTGGG | Ins_k3_30_CGA | 0.6121741621409653 | 0.03135862725412397 | 90.56186396916604 |
| GATCCAGATCTAACTTGGGGTGGCTTTGTCCGCTTCTTCTTTTGCCAATTCCACTAATTGTTTGGG | Ins_k3_29_CCG;Ins_k3_30_CGC | 0.38452250969848245 | 0.0348648771769689 | 72.12361663201555 |
| GATCCAGATCTAACTTGGGGTGGCTTTGTCCGGTTCTTCTTTTGCCAATTCCACTAATTGTTTGGG | Ins_k3_30_CGG | 0.762097174710554 | 0.0335193892384398 | 100 |
| GATCCAGATCTAACTTGGGGTGGCTTTGTCCGTTCTTCTTTTGCCAATTCCACTAATTGTTTGGG | Ins_k2_30_CG | 0.6643208356685788 | 0.03595139283087124 | 95.40966374763512 |
| GATCCAGATCTAACTTGGGGTGGCTTTGTCCGTTTCTTCTTTTGCCAATTCCACTAATTGTTTGGG | Ins_k3_30_CGT | 0.33054477467754817 | 0.031317175347130834 | 68.33375146680713 |
| GATCCAGATCTAACTTGGGGTGGCTTTGTCCTAATTGTTTGGG | Del_k20_31-50 | -1.5107523869968809 | 0.0499615838632222 | 10.838522086184037 |
| GATCCAGATCTAACTTGGGGTGGCTTTGTCCTATTCTTCTTTTGCCAATTCCACTAATTGTTTGGG | Ins_k3_30_CTA | -0.24357587628642235 | 0.04414817206663188 | 38.48556202087003 |
| GATCCAGATCTAACTTGGGGTGGCTTTGTCCTCTTCTTCTTTTGCCAATTCCACTAATTGTTTGGG | Ins_k3_28_TCC;Ins_k3_29_CCT;I | 0.06688243954629636 | 0.04036573169643738 | 52.49623613425991 |
| GATCCAGATCTAACTTGGGGTGGCTTTGTCCTCTTCTTTTGCCAATTCCACTAATTGTTTGGG | T31C | -0.39753442100875824 | 0.026170536908917012 | 32.99396327895168 |
| GATCCAGATCTAACTTGGGGTGGCTTTGTCCTGTTCTTCTTTTGCCAATTCCACTAATTGTTTGGG | Ins_k3_30_CTG | 0.422592571622455 | 0.03395503824221745 | 74.92230224059283 |
| GATCCAGATCTAACTTGGGGTGGCTTTGTCCTTCTTCTTTTGCCAATTCCACTAATTGTTTGGG | Ins_k1_29_C;Ins_k1_30_C | 0.277037176095172 | 0.037622602245963775 | 64.77347679799462 |
| GATCCAGATCTAACTTGGGGTGGCTTTGTCCTTCTTTTGCCAATTCCACTAATTGTTTGGG | Del_k2_31-32 | 0.0741944145451352 | 0.03248223039615166 | 52.881494082165894 |
| GATCCAGATCTAACTTGGGGTGGCTTTGTCCTTTCTTCTTTTGCCAATTCCACTAATTGTTTGGG | Ins_k2_30_CT | -0.14886763346640985 | 0.04184252057947984 | 42.30864341938279 |
| GATCCAGATCTAACTTGGGGTGGCTTTGTCCTTTTCTTCTTTTGCCAATTCCACTAATTGTTTGGG | Ins_k3_30_CTT | -0.6668169328947298 | 0.05141940847615497 | 25.20499280072589 |
| GATCCAGATCTAACTTGGGGTGGCTTTGTCCTTTTGCCAATTCCACTAATTGTTTGGG | Del_k5_31-35 | -0.26778568837789796 | 0.03189234599571787 | 37.56502181029992 |
| GATCCAGATCTAACTTGGGGTGGCTTTGTCG | Del_k32_31-62 | -2.6996446431705476 | 0.05808828651255681 | 3.300963487044974 |
| GATCCAGATCTAACTTGGGGTGGCTTTGTCGAAGTTCTTCTTTTGCCAATTCCACTAATTGTTTGGG | Ins_k4_30_GAAG | 0.9317708505439948 | 0.029432676462372173 | 100 |
| GATCCAGATCTAACTTGGGGTGGCTTTGTCGAATTCTTCTTTTGCCAATTCCACTAATTGTTTGGG | Ins_k3_30_GAA | 0.520048363051707 | 0.03196202501861808 | 82.59155188312097 |
| GATCCAGATCTAACTTGGGGTGGCTTTGTCGAATTTCTTCTTTTGCCAATTCCACTAATTGTTTGGG | Ins_k4_30_GAAT | 0.8410930887728445 | 0.0300687195959441 | 100 |
| GATCCAGATCTAACTTGGGGTGGCTTTGTCGACTTCTTCTTTTGCCAATTCCACTAATTGTTTGGG | Ins_k3_29_CGA;Ins_k3_30_GAC | 0.7163038083038664 | 0.02809199564227204 | 100 |
| GATCCAGATCTAACTTGGGGTGGCTTTGTCGAGTTCTTCTTTTGCCAATTCCACTAATTGTTTGGG | Ins_k3_30_GAG | 0.6159522587329431 | 0.03113173585232752 | 90.90466259419031 |
| GATCCAGATCTAACTTGGGGTGGCTTTGTCGATTCTTCTTTTGCCAATTCCACTAATTGTTTGGG | Ins_k2_30_GA | 0.6573217277559309 | 0.031154546906474533 | 94.74421271351879 |
| GATCCAGATCTAACTTGGGGTGGCTTTGTCGATTTCTTCTTTTGCCAATTCCACTAATTGTTTGGG | Ins_k3_30_GAT | 0.5212429348557317 | 0.031123635164206714 | 82.69027237486962 |
| GATCCAGATCTAACTTGGGGTGGCTTTGTCGATTTTCTTCTTTTGCCAATTCCACTAATTGTTTGGG | Ins_k4_30_GATT | 0.812250911771808 | 0.034025599164572706 | 100 |
| GATCCAGATCTAACTTGGGGTGGCTTTGTCGCATTCTTCTTTTGCCAATTCCACTAATTGTTTGGG | Ins_k3_30_GCA | 0.5167458167429908 | 0.03114237444296763 | 82.31923936816884 |
| GATCCAGATCTAACTTGGGGTGGCTTTGTCGCCAATTCCACTAATTGTTTGGG | Del_k10_31-40 | 0.08034359400374302 | 0.028385760433260715 | 53.20767372007859 |
| GATCCAGATCTAACTTGGGGTGGCTTTGTCGCCTTCTTCTTTTGCCAATTCCACTAATTGTTTGGG | Ins_k3_29_CGC;Ins_k3_30_GCC | 0.6045841758566592 | 0.03126155851476422 | 89.87710261570096 |
| GATCCAGATCTAACTTGGGGTGGCTTTGTCGCGTTCTTCTTTTGCCAATTCCACTAATTGTTTGGG | Ins_k3_30_GCG | 0.7479888140321478 | 0.03201258716036241 | 100 |
| GATCCAGATCTAACTTGGGGTGGCTTTGTCGCTTCTTCTTTTGCCAATTCCACTAATTGTTTGGG | Ins_k2_29_CG;Ins_k2_30_GC | 0.6522410824584421 | 0.03332403144781766 | 94.2640717205916 |
| GATCCAGATCTAACTTGGGGTGGCTTTGTCGCTTTCTTCTTTTGCCAATTCCACTAATTGTTTGGG | Ins_k3_30_GCT | 0.16135670036533079 | 0.03664741723192376 | 57.6976090568448 |
| GATCCAGATCTAACTTGGGGTGGCTTTGTCGG | Del_k31_31-61 | -2.083527146473285 | 0.043033214435302045 | 6.112475754976977 |
| GATCCAGATCTAACTTGGGGTGGCTTTGTCGGATTCTTCTTTTGCCAATTCCACTAATTGTTTGGG | Ins_k3_30_GGA | 0.8153564865404959 | 0.02845616191480266 | 100 |
| GATCCAGATCTAACTTGGGGTGGCTTTGTCGGCTTCTTCTTTTGCCAATTCCACTAATTGTTTGGG | Ins_k3_29_CGG;Ins_k3_30_GGC | 0.6459252969430607 | 0.03134152876217863 | 93.67059616671939 |
| GATCCAGATCTAACTTGGGGTGGCTTTGTCGGG | Del_k30_31-60 | -3.1670946877002155 | 0.0666796642480448 | 2.068377217858453 |
| GATCCAGATCTAACTTGGGGTGGCTTTGTCGGGTTCTTCTTTTGCCAATTCCACTAATTGTTTGGG | Ins_k3_30_GGG | -0.10096748347708151 | 0.040171641402036384 | 44.384555122635625 |
| GATCCAGATCTAACTTGGGGTGGCTTTGTCGGTTCTTCTTTTGCCAATTCCACTAATTGTTTGGG | Ins_k2_30_GG | 0.6590265579730354 | 0.034232837067955504 | 94.90587327299713 |
| GATCCAGATCTAACTTGGGGTGGCTTTGTCGGTTTCTTCTTTTGCCAATTCCACTAATTGTTTGGG | Ins_k3_30_GGT | 0.4027207544446585 | 0.03421343988382179 | 73.44815544578599 |

| | | | | |
|---|---|---|---|---|
| **GATCCAGATCTAACTTGGGGTGGCTTTGTCGTATTCTTCTTTTGCCAATTCCACTAATTGTTTGGG** | Ins_k3_30_GTA | 0.13894068076795552 | 0.03584683757575601 | 56.41864650012951 |
| **GATCCAGATCTAACTTGGGGTGGCTTTGTCGTCCTTCTTCTTTTGCCAATTCCACTAATTGTTTGGG** | Ins_k4_29_CGTC;Ins_k4_30_GT( | 0.8727771198135377 | 0.03252760151191089 | 100 |
| **GATCCAGATCTAACTTGGGGTGGCTTTGTCGTCTTCTTCTTTTGCCAATTCCACTAATTGTTTGGG** | Ins_k3_27_GTC;Ins_k3_28_TCG; | 0.5272104803280075 | 0.027550773083994333 | 83.1852056349697 |
| **GATCCAGATCTAACTTGGGGTGGCTTTGTCGTCTTCTTTTGCCAATTCCACTAATTGTTTGGG** | T31G | 0.31688665964910423 | 0.0285750888784396 | 67.40678593979753 |
| **GATCCAGATCTAACTTGGGGTGGCTTTGTCGTGTTCTTCTTTTGCCAATTCCACTAATTGTTTGGG** | Ins_k3_30_GTG | 0.4991555446574889 | 0.031112570001226268 | 80.88388271702976 |
| **GATCCAGATCTAACTTGGGGTGGCTTTGTCGTTCTTCTTTTGCCAATTCCACTAATTGTTTGGG** | Ins_k1_30_G | 0.6279567340826617 | 0.03289710928773541 | 92.00250168274938 |
| **GATCCAGATCTAACTTGGGGTGGCTTTGTCGTTTCTTCTTTTGCCAATTCCACTAATTGTTTGGG** | Ins_k2_30_GT | 0.40009754351473403 | 0.03935864548772184 | 73.25573792784508 |
| **GATCCAGATCTAACTTGGGGTGGCTTTGTCGTTTGGG** | Del_k26_31-56 | -2.097465214680872 | 0.04853598955904996 | 6.02787063662219 |
| **GATCCAGATCTAACTTGGGGTGGCTTTGTCGTTTTCTTCTTTTGCCAATTCCACTAATTGTTTGGG** | Ins_k3_30_GTT | 0.1379672065145634 | 0.039317763115463776 | 56.36375112430305 |
| **GATCCAGATCTAACTTGGGGTGGCTTTGTCT** | Del_k32_32-63 | -6.618972791958213 | 1.1952672818445644 | 0.06553874679239309 |
| **GATCCAGATCTAACTTGGGGTGGCTTTGTCTAAATCTTCTTTTGCCAATTCCACTAATTGTTTGGG** | Ins_k3_31_AAA | -0.842728669891047 | 0.05517434121074461 | 21.13922596181049 |
| **GATCCAGATCTAACTTGGGGTGGCTTTGTCTAACTCTTCTTTTGCCAATTCCACTAATTGTTTGGG** | Ins_k3_31_AAC | -1.5287864662808959 | 0.07482842176827703 | 10.644811267676781 |
| **GATCCAGATCTAACTTGGGGTGGCTTTGTCTAAGTCTTCTTTTGCCAATTCCACTAATTGTTTGGG** | Ins_k3_31_AAG | -0.5842820450324743 | 0.04097756419800438 | 27.373543733393664 |
| **GATCCAGATCTAACTTGGGGTGGCTTTGTCTAATCTTCTTTTGCCAATTCCACTAATTGTTTGGG** | Ins_k2_31_AA | -0.4275375834069318 | 0.2807480308109979 | 32.01874304194752 |
| **GATCCAGATCTAACTTGGGGTGGCTTTGTCTAATTCCACTAATTGTTTGGG** | Del_k12_32-43 | -1.2918657009475552 | 0.04613337533036936 | 13.490602525868011 |
| **GATCCAGATCTAACTTGGGGTGGCTTTGTCTAATTCTTCTTTTGCCAATTCCACTAATTGTTTGGG** | Ins_k3_30_TAA;Ins_k3_31_AAT | -1.087293919465246 | 0.06347491313323834 | 16.552962938821327 |
| **GATCCAGATCTAACTTGGGGTGGCTTTGTCTAATTGTTTGGG** | Del_k21_30-50;Del_k21_31-51;D | -3.040322700697319 | 0.08125126916749184 | 2.347935271550611 |
| **GATCCAGATCTAACTTGGGGTGGCTTTGTCTACATCTTCTTTTGCCAATTCCACTAATTGTTTGGG** | Ins_k3_31_ACA | 0.35625422520125316 | 0.0305856066190093 | 70.11335293547845 |
| **GATCCAGATCTAACTTGGGGTGGCTTTGTCTACCTCTTCTTTTGCCAATTCCACTAATTGTTTGGG** | Ins_k3_31_ACC | -1.9890286021884724 | 0.09283650520233391 | 6.7182683318310765 |
| **GATCCAGATCTAACTTGGGGTGGCTTTGTCTACGTCTTCTTTTGCCAATTCCACTAATTGTTTGGG** | Ins_k3_31_ACG | 0.6797340686937106 | 0.02923669344289796 | 96.89162670929021 |
| **GATCCAGATCTAACTTGGGGTGGCTTTGTCTACTAATTGTTTGGG** | Del_k18_32-49 | -0.6102593019276119 | 0.03293025001886547 | 26.671610774727952 |
| **GATCCAGATCTAACTTGGGGTGGCTTTGTCTACTCTTCTTTTGCCAATTCCACTAATTGTTTGGG** | Ins_k2_31_AC | 0.19192549039414186 | 0.036368527945471504 | 59.4885897464666 |
| **GATCCAGATCTAACTTGGGGTGGCTTTGTCTACTTCTTCTTTTGCCAATTCCACTAATTGTTTGGG** | Ins_k3_29_CTA;Ins_k3_30_TAC;I | 0.20741486381600394 | 0.03543636279208775 | 60.41720398769002 |
| **GATCCAGATCTAACTTGGGGTGGCTTTGTCTACTTCTTTTGCCAATTCCACTAATTGTTTGGG** | T32A | -0.22567636964575932 | 0.032074738578935044 | 39.1806367835441 |
| **GATCCAGATCTAACTTGGGGTGGCTTTGTCTAGATCTTCTTTTGCCAATTCCACTAATTGTTTGGG** | Ins_k3_31_AGA | 0.12045773546589911 | 0.03400992178315296 | 55.38544149207045 |
| **GATCCAGATCTAACTTGGGGTGGCTTTGTCTAGCTCTTCTTTTGCCAATTCCACTAATTGTTTGGG** | Ins_k3_31_AGC | -1.2819843782500833 | 0.0614446788659626 | 13.6245683123536 |
| **GATCCAGATCTAACTTGGGGTGGCTTTGTCTAGGTCTTCTTTTGCCAATTCCACTAATTGTTTGGG** | Ins_k3_31_AGG | -2.7229397818528893 | 0.11587685123887012 | 3.224955826509545 |
| **GATCCAGATCTAACTTGGGGTGGCTTTGTCTAGTCTTCTTTTGCCAATTCCACTAATTGTTTGGG** | Ins_k2_31_AG | -0.8102101969051743 | 0.050914343791328545 | 21.83794030161582 |
| **GATCCAGATCTAACTTGGGGTGGCTTTGTCTAGTTCTTCTTTTGCCAATTCCACTAATTGTTTGGG** | Ins_k3_30_TAG;Ins_k3_31_AGT | -1.3785319553191728 | 0.06102395943896866 | 12.370654304665264 |
| **GATCCAGATCTAACTTGGGGTGGCTTTGTCTATATCTTCTTTTGCCAATTCCACTAATTGTTTGGG** | Ins_k3_31_ATA | -0.8095187100102649 | 0.057108139769067194 | 21.85304617330044 |
| **GATCCAGATCTAACTTGGGGTGGCTTTGTCTATCTCTTCTTTTGCCAATTCCACTAATTGTTTGGG** | Ins_k3_31_ATC | -1.7633438588233281 | 0.07304338150386011 | 8.419203065543341 |
| **GATCCAGATCTAACTTGGGGTGGCTTTGTCTATCTTCTTTTGCCAATTCCACTAATTGTTTGGG** | Ins_k1_31_A | -0.5705364384333254 | 0.04937522269139314 | 27.75240758828316 |
| **GATCCAGATCTAACTTGGGGTGGCTTTGTCTATGTCTTCTTTTGCCAATTCCACTAATTGTTTGGG** | Ins_k3_31_ATG | -0.4426650903329633 | 0.03792450294860231 | 31.538024488765572 |
| **GATCCAGATCTAACTTGGGGTGGCTTTGTCTATTCCACTAATTGTTTGGG** | Del_k13_32-44 | -1.1947448442653508 | 0.04038298166116605 | 14.866556953706102 |
| **GATCCAGATCTAACTTGGGGTGGCTTTGTCTATTCTTCTTTTGCCAATTCCACTAATTGTTTGGG** | Ins_k2_30_TA;Ins_k2_31_AT | -0.5996866988921894 | 0.057438237447007115 | 26.95509507057809 |
| **GATCCAGATCTAACTTGGGGTGGCTTTGTCTATTGTTTGGG** | Del_k22_32-53 | -2.575445775640794 | 0.07351932713512048 | 3.737486247680551 |
| **GATCCAGATCTAACTTGGGGTGGCTTTGTCTATTTCTTCTTTTGCCAATTCCACTAATTGTTTGGG** | Ins_k3_30_TAT;Ins_k3_31_ATT | -1.3394953447769553 | 0.06849300852862494 | 12.863112126319944 |
| **GATCCAGATCTAACTTGGGGTGGCTTTGTCTCAATCTTCTTTTGCCAATTCCACTAATTGTTTGGG** | Ins_k3_31_CAA | 6.450830124435925e-4 | 0.0314787739705726 | 49.13168379415095 |
| **GATCCAGATCTAACTTGGGGTGGCTTTGTCTCAATTCCACTAATTGTTTGGG** | Del_k11_32-42 | -0.30898247413146596 | 0.031541223900052644 | 36.048907591168486 |
| **GATCCAGATCTAACTTGGGGTGGCTTTGTCTCACTAATTGTTTGGG** | Del_k17_32-48 | -1.2359060723166135 | 0.0395955759389779 | 14.267053991440504 |
| **GATCCAGATCTAACTTGGGGTGGCTTTGTCTCACTCTTCTTTTGCCAATTCCACTAATTGTTTGGG** | Ins_k3_31_CAC | -0.6273304527506731 | 0.045745288016913604 | 26.220160039876003 |
| **GATCCAGATCTAACTTGGGGTGGCTTTGTCTCAGCTCTTCTTTTGCCAATTCCACTAATTGTTTGGG** | Ins_k4_31_CAGC | 0.4834129697742928 | 0.032701603181917886 | 79.62053242092689 |
| **GATCCAGATCTAACTTGGGGTGGCTTTGTCTCAGTCTTCTTTTGCCAATTCCACTAATTGTTTGGG** | Ins_k3_31_CAG | 0.13422409331710616 | 0.033784346972069626 | 56.15316958475101 |
| **GATCCAGATCTAACTTGGGGTGGCTTTGTCTCATCTCTTCTTTTGCCAATTCCACTAATTGTTTGGG** | Ins_k4_31_CATC | 0.3499450320218014 | 0.03456383373714206 | 69.6723867804326 |
| **GATCCAGATCTAACTTGGGGTGGCTTTGTCTCATCTTCTTTTGCCAATTCCACTAATTGTTTGGG** | Ins_k2_31_CA | 0.32383483615966824 | 0.03167098755856165 | 67.87677106552856 |
| **GATCCAGATCTAACTTGGGGTGGCTTTGTCTCATTCTTCTTTTGCCAATTCCACTAATTGTTTGGG** | Ins_k3_30_TCA;Ins_k3_31_CAT | -0.3447413987257399 | 0.04462647952926108 | 34.7826130203814 |
| **GATCCAGATCTAACTTGGGGTGGCTTTGTCTCCAATTCCACTAATTGTTTGGG** | Del_k10_32-41 | 0.31128297058313037 | 0.029106359102968358 | 67.03011562777066 |
| **GATCCAGATCTAACTTGGGGTGGCTTTGTCTCCACTAATTGTTTGGG** | Del_k16_31-46;Del_k16_32-47 | -0.39699933486095196 | 0.03256935971313589 | 33.01162261587541 |
| **GATCCAGATCTAACTTGGGGTGGCTTTGTCTCCATCTTCTTTTGCCAATTCCACTAATTGTTTGGG** | Ins_k3_31_CCA | 0.47999929085417237 | 0.029959351213955883 | 79.34919687749922 |
| **GATCCAGATCTAACTTGGGGTGGCTTTGTCTCCCTCTTCTTTTGCCAATTCCACTAATTGTTTGGG** | Ins_k3_31_CCC | -0.40436423692088364 | 0.04553301733831064 | 32.76938835919191 |
| **GATCCAGATCTAACTTGGGGTGGCTTTGTCTCCGTCTTCTTTTGCCAATTCCACTAATTGTTTGGG** | Ins_k3_31_CCG | 0.6429885485239233 | 0.03202208338004498 | 93.39591272700284 |
| **GATCCAGATCTAACTTGGGGTGGCTTTGTCTCCTCTTCTTTTGCCAATTCCACTAATTGTTTGGG** | Ins_k2_31_CC | 0.2946555320436848 | 0.03988325292159317 | 65.92479132165232 |
| **GATCCAGATCTAACTTGGGGTGGCTTTGTCTCCTTCTTCTTTTGCCAATTCCACTAATTGTTTGGG** | Ins_k3_29_CTC;Ins_k3_30_TCC; | 0.05444086771559442 | 0.035846207713278073 | 51.84714666104506 |
| **GATCCAGATCTAACTTGGGGTGGCTTTGTCTCCTTCTTTTGCCAATTCCACTAATTGTTTGGG** | T32C | -0.4411746964324227 | 0.027532444234576604 | 31.585063612803275 |
| **GATCCAGATCTAACTTGGGGTGGCTTTGTCTCGATCTTCTTTTGCCAATTCCACTAATTGTTTGGG** | Ins_k3_31_CGA | 0.5365917644031533 | 0.029784765570693966 | 83.96926165572101 |
| **GATCCAGATCTAACTTGGGGTGGCTTTGTCTCGCTCTTCTTTTGCCAATTCCACTAATTGTTTGGG** | Ins_k3_31_CGC | 0.2492491012873731 | 0.031920903009838276 | 62.99832483313956 |
| **GATCCAGATCTAACTTGGGGTGGCTTTGTCTCGGTCTTCTTTTGCCAATTCCACTAATTGTTTGGG** | Ins_k3_31_CGG | 0.7104224295362478 | 0.03149407756416867 | 99.91116731447549 |
| **GATCCAGATCTAACTTGGGGTGGCTTTGTCTCGTCTTCTTTTGCCAATTCCACTAATTGTTTGGG** | Ins_k2_31_CG | 0.6724737254104636 | 0.030824076674101336 | 96.1907077730885 |
| **GATCCAGATCTAACTTGGGGTGGCTTTGTCTCGTTCTTCTTTTGCCAATTCCACTAATTGTTTGGG** | Ins_k3_30_TCG;Ins_k3_31_CGT | 0.3006261421242058 | 0.035929778814127976 | 66.31957993692588 |
| **GATCCAGATCTAACTTGGGGTGGCTTTGTCTCTAATTGTTTGGG** | Del_k19_32-50 | -1.8850872571838784 | 0.05649064574437237 | 7.454156360914686 |
| **GATCCAGATCTAACTTGGGGTGGCTTTGTCTCTATCTTCTTTTGCCAATTCCACTAATTGTTTGGG** | Ins_k3_31_CTA | -1.2126570885905976 | 0.05866975851772639 | 14.602634333068304 |
| **GATCCAGATCTAACTTGGGGTGGCTTTGTCTCTATTCTTCTTTTGCCAATTCCACTAATTGTTTGGG** | Ins_k4_30_TCTA;Ins_k4_31_CTA | -1.1420955907148265 | 0.07339200373298066 | 15.670241127391852 |
| **GATCCAGATCTAACTTGGGGTGGCTTTGTCTCTCTCTTCTTTTGCCAATTCCACTAATTGTTTGGG** | Ins_k3_31_CTC | -1.353417547669217 | 0.058552773382955565 | 12.685270118280913 |
| **GATCCAGATCTAACTTGGGGTGGCTTTGTCTCTCTTCTTTTGCCAATTCCACTAATTGTTTGGG** | Ins_k1_31_C | -0.5120936388315668 | 0.04539375346415362 | 29.422667949971125 |
| **GATCCAGATCTAACTTGGGGTGGCTTTGTCTCTGTCTTCTTTTGCCAATTCCACTAATTGTTTGGG** | Ins_k3_31_CTG | -0.42785253515187005 | 0.04263396570952267 | 32.00866027083226 |
| **GATCCAGATCTAACTTGGGGTGGCTTTGTCTCTTCTTCTTTTGCCAATTCCACTAATTGTTTGGG** | Ins_k2_28_TC;Ins_k2_29_CT;Ins_ | -0.1674884993595771 | 0.04089069559610687 | 41.52810950599078 |
| **GATCCAGATCTAACTTGGGGTGGCTTTGTCTCTTCTTTTGCCAATTCCACTAATTGTTTGGG** | Del_k1_31-31;Del_k1_32-32 | -0.16132757609588882 | 0.03499289909134392 | 41.78475076371415 |
| **GATCCAGATCTAACTTGGGGTGGCTTTGTCTCTTTCTTCTTTTGCCAATTCCACTAATTGTTTGGG** | Ins_k3_30_TCT;Ins_k3_31_CTT | -1.8201260600314364 | 0.08926316150268745 | 7.954461567972874 |
| **GATCCAGATCTAACTTGGGGTGGCTTTGTCTCTTTTGCCAATTCCACTAATTGTTTGGG** | Del_k4_31-34;Del_k4_32-35 | -0.6119560321179426 | 0.03799239528885145 | 26.626394618158404 |
| **GATCCAGATCTAACTTGGGGTGGCTTTGTCTG** | Del_k31_32-62 | -2.234149648873605 | 0.042982497327313386 | 5.257782636770621 |
| **GATCCAGATCTAACTTGGGGTGGCTTTGTCTGAATCTTCTTTTGCCAATTCCACTAATTGTTTGGG** | Ins_k3_31_GAA | 0.37944815308862845 | 0.03085757807299871 | 71.75856267153806 |
| **GATCCAGATCTAACTTGGGGTGGCTTTGTCTGACTCTTCTTTTGCCAATTCCACTAATTGTTTGGG** | Ins_k3_31_GAC | 0.33646098930357915 | 0.03022587386880532 | 68.73922686373372 |
| **GATCCAGATCTAACTTGGGGTGGCTTTGTCTGAGTCTTCTTTTGCCAATTCCACTAATTGTTTGGG** | Ins_k3_31_GAG | 0.4903630067446001 | 0.029013928330843704 | 80.1758254833121 |
| **GATCCAGATCTAACTTGGGGTGGCTTTGTCTGATCTTCTTTTGCCAATTCCACTAATTGTTTGGG** | Ins_k2_31_GA | 0.43007025573425284 | 0.030764284967063314 | 75.48464744926521 |
| **GATCCAGATCTAACTTGGGGTGGCTTTGTCTGATGTCTTCTTTTGCCAATTCCACTAATTGTTTGGG** | Ins_k4_31_GATG | 0.505650383105311 | 0.028437921663640845 | 81.4109201270628 |

| | | | | |
|---|---|---|---|---|
| **GATCCAGATCTAACTTGGGGTGGCTTTGTCTGATTCTTCTTTTGCCAATTCCACTAATTGTTTGGG** | Ins_k3_30_TGA;Ins_k3_31_GAT | 0.13011797884616216 | 0.03541419977473173 | 55.92307097144446 |
| **GATCCAGATCTAACTTGGGGTGGCTTTGTCTGCATCTTCTTTTGCCAATTCCACTAATTGTTTGGG** | Ins_k3_31_GCA | 0.6472250489002329 | 0.028196410035143667 | 93.79242386315917 |
| **GATCCAGATCTAACTTGGGGTGGCTTTGTCTGCCAATTCCACTAATTGTTTGGG** | Del_k9_31-39;Del_k9_32-40 | -0.06841213140236335 | 0.18345736070523608 | 45.85328777816815 |
| **GATCCAGATCTAACTTGGGGTGGCTTTGTCTGCCTCTTCTTTTGCCAATTCCACTAATTGTTTGGG** | Ins_k3_31_GCC | -1.6254724293093106 | 0.06923050708643783 | 9.663796713120837 |
| **GATCCAGATCTAACTTGGGGTGGCTTTGTCTGCGTCTTCTTTTGCCAATTCCACTAATTGTTTGGG** | Ins_k3_31_GCG | 0.898670783189923 | 0.02872266944906916 | 100 |
| **GATCCAGATCTAACTTGGGGTGGCTTTGTCTGCTCTTCTTTTGCCAATTCCACTAATTGTTTGGG** | Ins_k2_31_GC | 0.39686315509687686 | 0.033579480334383155 | 73.01918317874187 |
| **GATCCAGATCTAACTTGGGGTGGCTTTGTCTGCTTCTTCTTTTGCCAATTCCACTAATTGTTTGGG** | Ins_k3_29_CTG;Ins_k3_30_TGC; | 0.303513074686653 | 0.03401512083203586 | 66.51131672419366 |
| **GATCCAGATCTAACTTGGGGTGGCTTTGTCTGCTTCTTTTGCCAATTCCACTAATTGTTTGGG** | T32G | 0.10009661396183855 | 0.03418217506416447 | 54.269134980377416 |
| **GATCCAGATCTAACTTGGGGTGGCTTTGTCTGG** | Del_k30_32-61 | -2.696182798230812 | 0.052211725804655766 | 3.312410713618231 |
| **GATCCAGATCTAACTTGGGGTGGCTTTGTCTGGATCTTCTTTTGCCAATTCCACTAATTGTTTGGG** | Ins_k3_31_GGA | 0.6718550175261062 | 0.03266857162565772 | 96.13121223086581 |
| **GATCCAGATCTAACTTGGGGTGGCTTTGTCTGGCTCTTCTTTTGCCAATTCCACTAATTGTTTGGG** | Ins_k3_31_GGC | 0.2568402798153604 | 0.03725689842967248 | 63.47837613620032 |
| **GATCCAGATCTAACTTGGGGTGGCTTTGTCTGGG** | Del_k29_31-59;Del_k29_32-60 | -2.964153282665094 | 0.06091205400942141 | 2.533763513414913 |
| **GATCCAGATCTAACTTGGGGTGGCTTTGTCTGGGTCTTCTTTTGCCAATTCCACTAATTGTTTGGG** | Ins_k3_31_GGG | -0.7820445534710472 | 0.057970930308701193 | 22.46176389841002 |
| **GATCCAGATCTAACTTGGGGTGGCTTTGTCTGGTCTTCTTTTGCCAATTCCACTAATTGTTTGGG** | Ins_k2_31_GG | 0.6930516219006093 | 0.03487143733239706 | 98.19061658799349 |
| **GATCCAGATCTAACTTGGGGTGGCTTTGTCTGGTTCTTCTTTTGCCAATTCCACTAATTGTTTGGG** | Ins_k3_30_TGG;Ins_k3_31_GGT | 0.20409367695750846 | 0.03223613786619084 | 60.2168800046049 |
| **GATCCAGATCTAACTTGGGGTGGCTTTGTCTGTATCTTCTTTTGCCAATTCCACTAATTGTTTGGG** | Ins_k3_31_GTA | -0.3590804478189422 | 0.042635588103661796 | 34.28742219207523 |
| **GATCCAGATCTAACTTGGGGTGGCTTTGTCTGTCTCTTCTTTTGCCAATTCCACTAATTGTTTGGG** | Ins_k3_31_GTC | -0.8577648572808259 | 0.049413459815441846 | 20.82375031814233 |
| **GATCCAGATCTAACTTGGGGTGGCTTTGTCTGTCTTCTTTTGCCAATTCCACTAATTGTTTGGG** | Ins_k1_31_G | -0.22402211441142866 | 0.04064386928958487 | 39.24550519669066 |
| **GATCCAGATCTAACTTGGGGTGGCTTTGTCTGTGTCTTCTTTTGCCAATTCCACTAATTGTTTGGG** | Ins_k3_31_GTG | 0.14304326689700497 | 0.03276044701742893 | 56.65058430377258 |
| **GATCCAGATCTAACTTGGGGTGGCTTTGTCTGTTCTTCTTTTGCCAATTCCACTAATTGTTTGGG** | Ins_k2_30_TG;Ins_k2_31_GT | -0.06697307378168627 | 0.0402061563408953 | 45.91932080264639 |
| **GATCCAGATCTAACTTGGGGTGGCTTTGTCTGTTTCTTCTTTTGCCAATTCCACTAATTGTTTGGG** | Ins_k3_30_TGT;Ins_k3_31_GTT | -0.6008026933730499 | 0.052092231150173314 | 26.92503011253223 |
| **GATCCAGATCTAACTTGGGGTGGCTTTGTCTGTTTGGG** | Del_k25_31-55;Del_k25_32-56 | -2.2936903242092948 | 0.05297835016134521 | 4.9538681239172675 |
| **GATCCAGATCTAACTTGGGGTGGCTTTGTCTT** | Del_k31_33-63 | -6.438571355635565 | 0.9472735350021984 | 0.07849563029028137 |
| **GATCCAGATCTAACTTGGGGTGGCTTTGTCTTAAACTTCTTTTGCCAATTCCACTAATTGTTTGGG** | Ins_k3_32_AAA | -1.410431055682579 | 0.06777715087146934 | 11.982269064296712 |
| **GATCCAGATCTAACTTGGGGTGGCTTTGTCTTAACCTTCTTTTGCCAATTCCACTAATTGTTTGGG** | Ins_k3_32_AAC | -1.3510643134265476 | 0.06742484117114167 | 12.715156681550873 |
| **GATCCAGATCTAACTTGGGGTGGCTTTGTCTTAACTTCTTTTGCCAATTCCACTAATTGTTTGGG** | Ins_k2_32_AA | -1.004034326957275 | 0.06785337904916426 | 17.99015579192941 |
| **GATCCAGATCTAACTTGGGGTGGCTTTGTCTTAAGCTTCTTTTGCCAATTCCACTAATTGTTTGGG** | Ins_k3_32_AAG | -0.7614574072401258 | 0.04773747408274025 | 22.928980341405918 |
| **GATCCAGATCTAACTTGGGGTGGCTTTGTCTTAATCTTCTTTTGCCAATTCCACTAATTGTTTGGG** | Ins_k3_31_TAA;Ins_k3_32_AAT | -1.2202304385583669 | 0.05896236105884703 | 14.49246118928707 |
| **GATCCAGATCTAACTTGGGGTGGCTTTGTCTTAATTCCACTAATTGTTTGGG** | Del_k11_33-43 | -1.284318147022021 | 0.04793978044425149 | 13.592808794395495 |
| **GATCCAGATCTAACTTGGGGTGGCTTTGTCTTAATTGTTTGGG** | Del_k20_32-51;Del_k20_33-52 | -2.9546073342496455 | 0.09601053760358592 | 2.5580665021987468 |
| **GATCCAGATCTAACTTGGGGTGGCTTTGTCTTACACTTCTTTTGCCAATTCCACTAATTGTTTGGG** | Ins_k3_32_ACA | -0.9894843648182815 | 0.053125148581458406 | 18.25382541752543 |
| **GATCCAGATCTAACTTGGGGTGGCTTTGTCTTACCCTTCTTTTGCCAATTCCACTAATTGTTTGGG** | Ins_k3_32_ACC | -2.01482951142308 | 0.0773815753920046 | 6.547147924055117 |
| **GATCCAGATCTAACTTGGGGTGGCTTTGTCTTACCTTCTTTTGCCAATTCCACTAATTGTTTGGG** | Ins_k2_32_AC | -0.5732114010921464 | 0.04371996387570494 | 27.678270135957494 |
| **GATCCAGATCTAACTTGGGGTGGCTTTGTCTTACGCTTCTTTTGCCAATTCCACTAATTGTTTGGG** | Ins_k3_32_ACG | -0.18772736119425965 | 0.03650381956279361 | 40.696075942296595 |
| **GATCCAGATCTAACTTGGGGTGGCTTTGTCTTACTAATTGTTTGGG** | Del_k17_33-49 | -2.8812880440843753 | 0.095471102255557 | 2.752669011720308 |
| **GATCCAGATCTAACTTGGGGTGGCTTTGTCTTACTCTTCTTTTGCCAATTCCACTAATTGTTTGGG** | Ins_k3_31_TAC;Ins_k3_32_ACT | -0.9431537874105249 | 0.04721110352119698 | 19.119432904749335 |
| **GATCCAGATCTAACTTGGGGTGGCTTTGTCTTACTTCTTTTGCCAATTCCACTAATTGTTTGGG** | Ins_k1_32_A | -0.6285597511813719 | 0.042482266820490354 | 26.18794744179102 |
| **GATCCAGATCTAACTTGGGGTGGCTTTGTCTTAGACTTCTTTTGCCAATTCCACTAATTGTTTGGG** | Ins_k3_32_AGA | -0.7376714498879982 | 0.04840377131312135 | 23.48090610956985 |
| **GATCCAGATCTAACTTGGGGTGGCTTTGTCTTAGCCCTTCTTTTGCCAATTCCACTAATTGTTTGGG** | Ins_k4_32_AGCC | -0.4668516543317724 | 0.05088292353275389 | 30.784378829785684 |
| **GATCCAGATCTAACTTGGGGTGGCTTTGTCTTAGCCTTCTTTTGCCAATTCCACTAATTGTTTGGG** | Ins_k3_32_AGC | -0.7587230661780413 | 0.05083949819664709 | 22.991761787692322 |
| **GATCCAGATCTAACTTGGGGTGGCTTTGTCTTAGCTTCTTTTGCCAATTCCACTAATTGTTTGGG** | Ins_k2_32_AG | -2.2872992479994894 | 0.09580996862262434 | 4.985630060998895 |
| **GATCCAGATCTAACTTGGGGTGGCTTTGTCTTAGGCTTCTTTTGCCAATTCCACTAATTGTTTGGG** | Ins_k3_32_AGG | -2.4151557902136904 | 0.09959944665595734 | 4.3872527967639865 |
| **GATCCAGATCTAACTTGGGGTGGCTTTGTCTTAGTCTTCTTTTGCCAATTCCACTAATTGTTTGGG** | Ins_k3_31_TAG;Ins_k3_32_AGT | -1.5621665014459936 | 0.06597951798451919 | 10.295352021663613 |
| **GATCCAGATCTAACTTGGGGTGGCTTTGTCTTATACTTCTTTTGCCAATTCCACTAATTGTTTGGG** | Ins_k3_32_ATA | -1.3761791069104068 | 0.06067547135826536 | 12.399794847157656 |
| **GATCCAGATCTAACTTGGGGTGGCTTTGTCTTATCCTTCTTTTGCCAATTCCACTAATTGTTTGGG** | Ins_k3_32_ATC | -0.8637567421115477 | 0.05481753141726385 | 20.699349873253908 |
| **GATCCAGATCTAACTTGGGGTGGCTTTGTCTTATCTTCTTTTGCCAATTCCACTAATTGTTTGGG** | Ins_k2_31_TA;Ins_k2_32_AT | -0.6100602118227493 | 0.05645478335695818 | 26.676921357138614 |
| **GATCCAGATCTAACTTGGGGTGGCTTTGTCTTATGCCTTCTTTTGCCAATTCCACTAATTGTTTGGG** | Ins_k4_32_ATGC | -0.2294861761935899 | 0.04420646879111605 | 39.03165012236373 |
| **GATCCAGATCTAACTTGGGGTGGCTTTGTCTTATGCTTCTTTTGCCAATTCCACTAATTGTTTGGG** | Ins_k3_32_ATG | -0.9851748538210765 | 0.058179389986609126 | 18.332660226632512 |
| **GATCCAGATCTAACTTGGGGTGGCTTTGTCTTATTCCACTAATTGTTTGGG** | Del_k12_33-44 | -1.1617183583417519 | 0.04437383047880718 | 15.365744926721172 |
| **GATCCAGATCTAACTTGGGGTGGCTTTGTCTTATTCTTCTTTTGCCAATTCCACTAATTGTTTGGG** | Ins_k3_30_TTA;Ins_k3_31_TAT;In | -0.9885390131102915 | 0.05697023200816253 | 18.27108986176141 |
| **GATCCAGATCTAACTTGGGGTGGCTTTGTCTTATTCTTTTGCCAATTCCACTAATTGTTTGGG** | C33A | -2.550886153227623 | 0.0890909415312736 | 3.8304139625663387 |
| **GATCCAGATCTAACTTGGGGTGGCTTTGTCTTATTGTTTGGG** | Del_k21_33-53 | -3.10865099375513 | 0.08965328671364552 | 2.192863098813144 |
| **GATCCAGATCTAACTTGGGGTGGCTTTGTCTTC** | Del_k30_34-63 | -6.220436354455035 | 0.48725472843637274 | 0.09762932987876617 |
| **GATCCAGATCTAACTTGGGGTGGCTTTGTCTTCAAATTCTTTTGCCAATTCCACTAATTGTTTGGG** | Ins_k3_33_AAA | 0.28382384187547305 | 0.03754265940402195 | 65.21456781208623 |
| **GATCCAGATCTAACTTGGGGTGGCTTTGTCTTCAACTTCTTTTGCCAATTCCACTAATTGTTTGGG** | Ins_k3_32_CAA;Ins_k3_33_AAC | 0.6692463275163216 | 0.03204761253890152 | 95.8807625127608 |
| **GATCCAGATCTAACTTGGGGTGGCTTTGTCTTCAAGTTCTTTTGCCAATTCCACTAATTGTTTGGG** | Ins_k3_33_AAG | 0.5389773352609366 | 0.029410445220865975 | 84.16981540175072 |
| **GATCCAGATCTAACTTGGGGTGGCTTTGTCTTCAATTCCACTAATTGTTTGGG** | Del_k10_33-42;Del_k10_34-43 | 0.31770947666570093 | 0.030147798566729753 | 67.46227221469799 |
| **GATCCAGATCTAACTTGGGGTGGCTTTGTCTTCAATTCTTTTGCCAATTCCACTAATTGTTTGGG** | Ins_k2_33_AA | 0.41823608732347606 | 0.03604112905590787 | 74.5966143494316 |
| **GATCCAGATCTAACTTGGGGTGGCTTTGTCTTCAATTGTTTGGG** | Del_k19_34-52 | -0.031630689540294576 | 0.0269526008259534 | 47.57123849231855 |
| **GATCCAGATCTAACTTGGGGTGGCTTTGTCTTCAATTTCTTTTGCCAATTCCACTAATTGTTTGGG** | Ins_k3_33_AAT | -0.016995252496632363 | 0.03781948890800565 | 48.272584089285594 |
| **GATCCAGATCTAACTTGGGGTGGCTTTGTCTTCACAATTCTTTTGCCAATTCCACTAATTGTTTGGG** | Ins_k4_33_ACAA | -0.006479115433204896 | 0.03593367730012151 | 48.78290379297338 |
| **GATCCAGATCTAACTTGGGGTGGCTTTGTCTTCACATTCTTTTGCCAATTCCACTAATTGTTTGGG** | Ins_k3_33_ACA | 0.1198206575917885 | 0.031597738134416464 | 55.35016788995676 |
| **GATCCAGATCTAACTTGGGGTGGCTTTGTCTTCACCTTCTTTTGCCAATTCCACTAATTGTTTGGG** | Ins_k3_32_CAC;Ins_k3_33_ACC | 0.3399528620794705 | 0.03240253254953 | 68.9796750615942 |
| **GATCCAGATCTAACTTGGGGTGGCTTTGTCTTCACGTTCTTTTGCCAATTCCACTAATTGTTTGGG** | Ins_k3_33_ACG | 0.48377237837882103 | 0.03179829957204715 | 79.64915386846525 |
| **GATCCAGATCTAACTTGGGGTGGCTTTGTCTTCACGTTTCTTTTGCCAATTCCACTAATTGTTTGGG** | Ins_k4_33_ACGT | 0.5769376803059749 | 0.036696751544674805 | 87.42634916673687 |
| **GATCCAGATCTAACTTGGGGTGGCTTTGTCTTCACTAATTGTTTGGG** | Del_k16_33-48;Del_k16_34-49 | -0.34838096964473214 | 0.030370272181566228 | 34.656249327845586 |
| **GATCCAGATCTAACTTGGGGTGGCTTTGTCTTCACTTCTTTTGCCAATTCCACTAATTGTTTGGG** | Ins_k2_32_CA;Ins_k2_33_AC | 0.5743833563900942 | 0.03222082958025357 | 87.20331891919447 |
| **GATCCAGATCTAACTTGGGGTGGCTTTGTCTTCACTTTCTTTTGCCAATTCCACTAATTGTTTGGG** | Ins_k3_33_ACT | -0.4040983289710266 | 0.03896342484754233 | 32.77810315868446 |
| **GATCCAGATCTAACTTGGGGTGGCTTTGTCTTCAGATTCTTTTGCCAATTCCACTAATTGTTTGGG** | Ins_k3_33_AGA | 0.3234276116337399 | 0.030283061698496174 | 67.84913560691002 |
| **GATCCAGATCTAACTTGGGGTGGCTTTGTCTTCAGCCTTCTTTTGCCAATTCCACTAATTGTTTGGG** | Ins_k4_32_CAGC;Ins_k4_33_AG | 0.5980255505419794 | 0.029010068577908332 | 89.28956121335031 |
| **GATCCAGATCTAACTTGGGGTGGCTTTGTCTTCAGCTTCTTTTGCCAATTCCACTAATTGTTTGGG** | Ins_k3_32_CAG;Ins_k3_33_AGC | 0.5236293670390688 | 0.029021807992339894 | 82.88784275250319 |
| **GATCCAGATCTAACTTGGGGTGGCTTTGTCTTCAGGTTCTTTTGCCAATTCCACTAATTGTTTGGG** | Ins_k3_33_AGG | -0.4095179555617343 | 0.04571674600339417 | 32.600938595745596 |
| **GATCCAGATCTAACTTGGGGTGGCTTTGTCTTCAGTTCTTTTGCCAATTCCACTAATTGTTTGGG** | Ins_k2_33_AG | 0.4590719602414217 | 0.03556330291860234 | 77.7058850396833 |
| **GATCCAGATCTAACTTGGGGTGGCTTTGTCTTCAGTTTCTTTTGCCAATTCCACTAATTGTTTGGG** | Ins_k3_33_AGT | 0.22559307303511822 | 0.038196297810819006 | 61.52552367586373 |
| **GATCCAGATCTAACTTGGGGTGGCTTTGTCTTCATATTCTTTTGCCAATTCCACTAATTGTTTGGG** | Ins_k3_33_ATA | 0.00961651819584275 | 0.044660479717953595 | 49.5744486591878 |

| | | | | |
|---|---|---|---|---|
| **GATCCAGATCTAACTTGGGGTGGCTTTGTCTTCATCATTCTTTTGCCAATTCCACTAATTGTTTGGG** | Ins_k4_33_ATCA | 0.3151760855974868 | 0.03570629939643299 | 67.29158020292962 |
| **GATCCAGATCTAACTTGGGGTGGCTTTGTCTTCATCTTCTTTTGCCAATTCCACTAATTGTTTGGG** | Ins_k3_31_TCA;Ins_k3_32_CAT;I | 0.5505324104450127 | 0.028873664527066908 | 85.148044819795075 |
| **GATCCAGATCTAACTTGGGGTGGCTTTGTCTTCATCTTTTGCCAATTCCACTAATTGTTTGGG** | T34A | 0.0547831045144378 | 0.03091297723710908 | 51.864893699219046 |
| **GATCCAGATCTAACTTGGGGTGGCTTTGTCTTCATGTTCTTTTGCCAATTCCACTAATTGTTTGGG** | Ins_k3_33_ATG | 0.02056860060879906 | 0.03219909887168637 | 50.120376171127745 |
| **GATCCAGATCTAACTTGGGGTGGCTTTGTCTTCATTCCACTAATTGTTTGGG** | Del_k11_34-44 | 0.0368488130595227 | 0.027298860229517757 | 50.94302482084799 |
| **GATCCAGATCTAACTTGGGGTGGCTTTGTCTTCATTCTTTTGCCAATTCCACTAATTGTTTGGG** | Ins_k1_33_A | 0.15589347806151377 | 0.035450480476479945 | 57.38325367072167 |
| **GATCCAGATCTAACTTGGGGTGGCTTTGTCTTCATTGTTTGGG** | Del_k20_34-53 | -0.19291968842084595 | 0.02850666653267236 | 40.48531623934931 |
| **GATCCAGATCTAACTTGGGGTGGCTTTGTCTTCATTTCTTTTGCCAATTCCACTAATTGTTTGGG** | Ins_k2_33_AT | -0.03001492506793324 | 0.04463942887472905 | 47.648164539832074 |
| **GATCCAGATCTAACTTGGGGTGGCTTTGTCTTCATTTTCTTTTGCCAATTCCACTAATTGTTTGGG** | Ins_k3_33_ATT | -0.6238485150914216 | 0.050211753959529655 | 26.311616132356455 |
| **GATCCAGATCTAACTTGGGGTGGCTTTGTCTTCCAATTCCACTAATTGTTTGGG** | Del_k9_33-41;Del_k9_34-42 | 0.12147029536490284 | 0.028792832884473218 | 55.44155097142432 |
| **GATCCAGATCTAACTTGGGGTGGCTTTGTCTTCCAATTCTTTTGCCAATTCCACTAATTGTTTGGG** | Ins_k3_33_CAA | -0.205189284361944 | 0.03864199334126253 | 39.99161273243494 |
| **GATCCAGATCTAACTTGGGGTGGCTTTGTCTTCCACTAATTGTTTGGG** | Del_k15_31-45;Del_k15_32-46;D | -0.5506618593229392 | 0.030043692849703454 | 28.309492586121394 |
| **GATCCAGATCTAACTTGGGGTGGCTTTGTCTTCCACTTCTTTTGCCAATTCCACTAATTGTTTGGG** | Ins_k3_32_CCA;Ins_k3_33_CAC | -0.03144975471527511 | 0.03444627783081824 | 47.579846564757666 |
| **GATCCAGATCTAACTTGGGGTGGCTTTGTCTTCCAGTTCTTTTGCCAATTCCACTAATTGTTTGGG** | Ins_k3_33_CAG | 0.26081095554165357 | 0.03489841468734346 | 63.73092925491967 |
| **GATCCAGATCTAACTTGGGGTGGCTTTGTCTTCCATCTTCTTTTGCCAATTCCACTAATTGTTTGGG** | Ins_k4_31_TCCA;Ins_k4_32_CC/ | 0.32503717998554404 | 0.03254431080715332 | 67.95843136417999 |
| **GATCCAGATCTAACTTGGGGTGGCTTTGTCTTCCATTCTTTTGCCAATTCCACTAATTGTTTGGG** | Ins_k2_33_CA | -0.05312766650192846 | 0.045830434288699935 | 46.55951414313194 |
| **GATCCAGATCTAACTTGGGGTGGCTTTGTCTTCCATTTCTTTTGCCAATTCCACTAATTGTTTGGG** | Ins_k3_33_CAT | -0.5545834017894032 | 0.051509605175167984 | 28.198693103173767 |
| **GATCCAGATCTAACTTGGGGTGGCTTTGTCTTCCCAATTCCACTAATTGTTTGGG** | Del_k8_34-41 | -0.6327112975733377 | 0.03572141088263603 | 26.079452330149817 |
| **GATCCAGATCTAACTTGGGGTGGCTTTGTCTTCCCACTAATTGTTTGGG** | Del_k14_34-47 | -1.2438581530136603 | 0.03883633867883583 | 14.154051126225955 |
| **GATCCAGATCTAACTTGGGGTGGCTTTGTCTTCCCATTCTTTTGCCAATTCCACTAATTGTTTGGG** | Ins_k3_33_CCA | -0.9879719535260841 | 0.055512187587146496 | 18.28145359653129 |
| **GATCCAGATCTAACTTGGGGTGGCTTTGTCTTCCCCTTCTTTTGCCAATTCCACTAATTGTTTGGG** | Ins_k3_32_CCC;Ins_k3_33_CCC | -2.3306245472176004 | 0.09748038336325308 | 4.774238513404587 |
| **GATCCAGATCTAACTTGGGGTGGCTTTGTCTTCCCGTTCTTTTGCCAATTCCACTAATTGTTTGGG** | Ins_k3_33_CCG | 0.2473860691282841 | 0.031451940814133654 | 62.881066190233994 |
| **GATCCAGATCTAACTTGGGGTGGCTTTGTCTTCCCTTCTTTTGCCAATTCCACTAATTGTTTGGG** | Ins_k2_32_CC;Ins_k2_33_CC | -0.5260580440212832 | 0.04389437843623282 | 29.014653363829645 |
| **GATCCAGATCTAACTTGGGGTGGCTTTGTCTTCCCTTTCTTTTGCCAATTCCACTAATTGTTTGGG** | Ins_k3_33_CCT | -2.1645496033231226 | 0.0835425982238193 | 5.636759995053759 |
| **GATCCAGATCTAACTTGGGGTGGCTTTGTCTTCCGATTCTTTTGCCAATTCCACTAATTGTTTGGG** | Ins_k3_33_CGA | 0.5791384986282702 | 0.03314905557464732 | 87.61897056243444 |
| **GATCCAGATCTAACTTGGGGTGGCTTTGTCTTCCGCTTCTTTTGCCAATTCCACTAATTGTTTGGG** | Ins_k3_32_CCG;Ins_k3_33_CGC | 0.4599132732636311 | 0.031962506842054206 | 77.7712875207913 |
| **GATCCAGATCTAACTTGGGGTGGCTTTGTCTTCCGGTTCTTTTGCCAATTCCACTAATTGTTTGGG** | Ins_k3_33_CGG | 0.7121437188057951 | 0.031220377980828204 | 100 |
| **GATCCAGATCTAACTTGGGGTGGCTTTGTCTTCCGTTCTTTTGCCAATTCCACTAATTGTTTGGG** | Ins_k2_33_CG | 0.3017635842506464 | 0.03044104858749313 | 66.39505753851824 |
| **GATCCAGATCTAACTTGGGGTGGCTTTGTCTTCCGTTTCTTTTGCCAATTCCACTAATTGTTTGGG** | Ins_k3_33_CGT | -0.13854309664735803 | 0.03906322228186303 | 42.74772331405789 |
| **GATCCAGATCTAACTTGGGGTGGCTTTGTCTTCCTAATTGTTTGGG** | Del_k17_34-50 | -0.4495291988693905 | 0.03096752605772955 | 31.322285341175384 |
| **GATCCAGATCTAACTTGGGGTGGCTTTGTCTTCCTATTCTTTTGCCAATTCCACTAATTGTTTGGG** | Ins_k3_33_CTA | -1.0434490316841252 | 0.05561561129334035 | 17.294871338255806 |
| **GATCCAGATCTAACTTGGGGTGGCTTTGTCTTCCTCTTCTTTTGCCAATTCCACTAATTGTTTGGG** | Ins_k3_31_TCC;Ins_k3_32_CCT;I | -0.22063658873949082 | 0.041663132013796095 | 39.37859702782528 |
| **GATCCAGATCTAACTTGGGGTGGCTTTGTCTTCCTCTTTTGCCAATTCCACTAATTGTTTGGG** | T34C | -0.46898748608688035 | 0.027314365688103297 | 30.718698741696826 |
| **GATCCAGATCTAACTTGGGGTGGCTTTGTCTTCCTGGTTCTTTTGCCAATTCCACTAATTGTTTGGG** | Ins_k4_33_CTGG | 0.5361543520507004 | 0.03713355947500811 | 83.93254049517655 |
| **GATCCAGATCTAACTTGGGGTGGCTTTGTCTTCCTGTTCTTTTGCCAATTCCACTAATTGTTTGGG** | Ins_k3_33_CTG | 0.24151818517621693 | 0.035225659961959935 | 62.51316783908709 |
| **GATCCAGATCTAACTTGGGGTGGCTTTGTCTTCCTTCTTTTGCCAATTCCACTAATTGTTTGGG** | Ins_k1_32_C;Ins_k1_33_C | -0.020975397247149363 | 0.037245826657345875 | 48.08083406668643 |
| **GATCCAGATCTAACTTGGGGTGGCTTTGTCTTCCTTTCTTTTGCCAATTCCACTAATTGTTTGGG** | Ins_k2_33_CT | -1.310698765280439 | 1.2495519688306294 | 13.238910644525179 |
| **GATCCAGATCTAACTTGGGGTGGCTTTGTCTTCCTTTTCTTTTGCCAATTCCACTAATTGTTTGGG** | Ins_k3_33_CTT | -2.1345523128410013 | 0.10339519170909939 | 5.808409155701174 |
| **GATCCAGATCTAACTTGGGGTGGCTTTGTCTTCCTTTTGCCAATTCCACTAATTGTTTGGG** | Del_k2_34-35 | 0.10012414535357095 | 0.03127519594222207 | 54.27062910575912 |
| **GATCCAGATCTAACTTGGGGTGGCTTTGTCTTCG** | Del_k29_34-62 | -1.9833165043323826 | 0.03720440461001534 | 6.7567535489877235 |
| **GATCCAGATCTAACTTGGGGTGGCTTTGTCTTCGAATTCTTTTGCCAATTCCACTAATTGTTTGGG** | Ins_k3_33_GAA | 0.5926584316705132 | 0.030581537753603266 | 88.81161726263839 |
| **GATCCAGATCTAACTTGGGGTGGCTTTGTCTTCGACTTCTTTTGCCAATTCCACTAATTGTTTGGG** | Ins_k3_32_CGA;Ins_k3_33_GAC | 0.6326190537918341 | 0.026679384562619528 | 92.43244825500672 |
| **GATCCAGATCTAACTTGGGGTGGCTTTGTCTTCGAGTTCTTTTGCCAATTCCACTAATTGTTTGGG** | Ins_k3_33_GAG | 0.5076637811068658 | 0.03447536559070658 | 81.57499783238558 |
| **GATCCAGATCTAACTTGGGGTGGCTTTGTCTTCGATTCTTTTGCCAATTCCACTAATTGTTTGGG** | Ins_k2_33_GA | 0.7174506386352495 | 0.03277861791251171 | 100 |
| **GATCCAGATCTAACTTGGGGTGGCTTTGTCTTCGATTTCTTTTGCCAATTCCACTAATTGTTTGGG** | Ins_k3_33_GAT | 0.20822104600500801 | 0.03130518443063208 | 60.46593090019973 |
| **GATCCAGATCTAACTTGGGGTGGCTTTGTCTTCGCAGTTCTTTTGCCAATTCCACTAATTGTTTGGG** | Ins_k4_33_GCAG | 0.5465546108298827 | 0.036491152834885306 | 84.81001571162633 |
| **GATCCAGATCTAACTTGGGGTGGCTTTGTCTTCGCATTCTTTTGCCAATTCCACTAATTGTTTGGG** | Ins_k3_33_GCA | 0.5494225489261555 | 0.03041249040459726 | 85.05359470441316 |
| **GATCCAGATCTAACTTGGGGTGGCTTTGTCTTCGCCAATTCCACTAATTGTTTGGG** | Del_k7_34-40 | 0.5591995798514797 | 0.028430669271961986 | 85.88924476171297 |
| **GATCCAGATCTAACTTGGGGTGGCTTTGTCTTCGCCTTCTTTTGCCAATTCCACTAATTGTTTGGG** | Ins_k3_32_CGC;Ins_k3_33_GCC | 0.5103897933107933 | 0.030296353952315756 | 81.79767564533397 |
| **GATCCAGATCTAACTTGGGGTGGCTTTGTCTTCGCGTTCTTTTGCCAATTCCACTAATTGTTTGGG** | Ins_k3_33_GCG | 0.6141935624701462 | 0.0309208887385305 | 90.74492940606699 |
| **GATCCAGATCTAACTTGGGGTGGCTTTGTCTTCGCTTCTTTTGCCAATTCCACTAATTGTTTGGG** | Ins_k2_32_CG;Ins_k2_33_GC | 0.7167782828088802 | 0.031023777531376782 | 100 |
| **GATCCAGATCTAACTTGGGGTGGCTTTGTCTTCGCTTTCTTTTGCCAATTCCACTAATTGTTTGGG** | Ins_k3_33_GCT | 0.13195002601409656 | 0.03466029262473458 | 56.02561858258763 |
| **GATCCAGATCTAACTTGGGGTGGCTTTGTCTTCGG** | Del_k28_34-61 | -2.0519353231120503 | 0.03817653977664221 | 6.308662643445069 |
| **GATCCAGATCTAACTTGGGGTGGCTTTGTCTTCGGATTCTTTTGCCAATTCCACTAATTGTTTGGG** | Ins_k3_33_GGA | 0.6833816349674526 | 0.028992707961357396 | 97.24569068242593 |
| **GATCCAGATCTAACTTGGGGTGGCTTTGTCTTCGGCTTCTTTTGCCAATTCCACTAATTGTTTGGG** | Ins_k3_32_CGG;Ins_k3_33_GGC | 0.5506883994564766 | 0.031240541037068397 | 85.16132801512391 |
| **GATCCAGATCTAACTTGGGGTGGCTTTGTCTTCGGG** | Del_k27_34-60 | -2.556432663543398 | 0.050368671606428106 | 3.8092273422284153 |
| **GATCCAGATCTAACTTGGGGTGGCTTTGTCTTCGGGTTCTTTTGCCAATTCCACTAATTGTTTGGG** | Ins_k3_33_GGG | 0.13914135179909748 | 0.03354633318935267 | 56.42996922413169 |
| **GATCCAGATCTAACTTGGGGTGGCTTTGTCTTCGGTTCTTTTGCCAATTCCACTAATTGTTTGGG** | Ins_k2_33_GG | 0.57400845011284 | 0.030720804461538402 | 87.17063197518738 |
| **GATCCAGATCTAACTTGGGGTGGCTTTGTCTTCGGTTTCTTTTGCCAATTCCACTAATTGTTTGGG** | Ins_k3_33_GGT | 0.422364914824368 | 0.03267787255831882 | 74.90524761053432 |
| **GATCCAGATCTAACTTGGGGTGGCTTTGTCTTCGTATTCTTTTGCCAATTCCACTAATTGTTTGGG** | Ins_k3_33_GTA | 0.1922919034029349 | 0.0328103253061819 | 59.51039113353645 |
| **GATCCAGATCTAACTTGGGGTGGCTTTGTCTTCGTCTTCTTTTGCCAATTCCACTAATTGTTTGGG** | Ins_k3_31_TCG;Ins_k3_32_CGT; | 0.7444976384788982 | 0.029134801944389013 | 100 |
| **GATCCAGATCTAACTTGGGGTGGCTTTGTCTTCGTCTTTTGCCAATTCCACTAATTGTTTGGG** | T34G | 0.444791167427129 | 0.028179836439497993 | 76.60406951994571 |
| **GATCCAGATCTAACTTGGGGTGGCTTTGTCTTCGTGTTCTTTTGCCAATTCCACTAATTGTTTGGG** | Ins_k3_33_GTG | 0.5074119663218895 | 0.029807479650193466 | 81.55445862799317 |
| **GATCCAGATCTAACTTGGGGTGGCTTTGTCTTCGTTCTTTTGCCAATTCCACTAATTGTTTGGG** | Ins_k1_33_G | 0.5605521085905758 | 0.029546123581052058 | 86.00549102912008 |
| **GATCCAGATCTAACTTGGGGTGGCTTTGTCTTCGTTTCTTTTGCCAATTCCACTAATTGTTTGGG** | Ins_k2_33_GT | 0.6110789843764013 | 0.03172675367993071 | 90.46273692020735 |
| **GATCCAGATCTAACTTGGGGTGGCTTTGTCTTCGTTTGGG** | Del_k23_34-56 | -0.37896815864210975 | 0.02898876280855036 | 33.61225982486976 |
| **GATCCAGATCTAACTTGGGGTGGCTTTGTCTTCGTTTTCTTTTGCCAATTCCACTAATTGTTTGGG** | Ins_k3_33_GTT | 0.3206455338509224 | 0.038263480763148956 | 67.66063636551178 |
| **GATCCAGATCTAACTTGGGGTGGCTTTGTCTTCT** | Del_k29_35-63 | -6.102806471260137 | 0.44853980541655897 | 0.10981617620794276 |
| **GATCCAGATCTAACTTGGGGTGGCTTTGTCTTCTAAATCTTTTGCCAATTCCACTAATTGTTTGGG** | Ins_k3_34_AAA | -0.5750022677881442 | 0.05142072077995198 | 27.628746402206946 |
| **GATCCAGATCTAACTTGGGGTGGCTTTGTCTTCTAACTCTTTTGCCAATTCCACTAATTGTTTGGG** | Ins_k3_34_AAC | -1.1470453654476167 | 0.056347718064129776 | 15.592868610039025 |
| **GATCCAGATCTAACTTGGGGTGGCTTTGTCTTCTAAGTCTTTTGCCAATTCCACTAATTGTTTGGG** | Ins_k3_34_AAG | -0.28092329704138463 | 0.038847263427960604 | 37.07473490487638 |
| **GATCCAGATCTAACTTGGGGTGGCTTTGTCTTCTAATCTTTTGCCAATTCCACTAATTGTTTGGG** | Ins_k2_34_AA | -0.4855624863153185 | 0.04549981279649366 | 30.213732769338293 |
| **GATCCAGATCTAACTTGGGGTGGCTTTGTCTTCTAATTCCACTAATTGTTTGGG** | Del_k9_35-43 | -0.1495974001353198 | 0.03559006904716281 | 42.277779244801216 |
| **GATCCAGATCTAACTTGGGGTGGCTTTGTCTTCTAATTCTTTTGCCAATTCCACTAATTGTTTGGG** | Ins_k3_33_TAA;Ins_k3_34_AAT | -0.9787574547563072 | 0.058267401582167 | 18.450686529110946 |
| **GATCCAGATCTAACTTGGGGTGGCTTTGTCTTCTAATTGTTTGGG** | Del_k18_33-50;Del_k18_34-51;D | -0.5396909114120725 | 0.034856247277715496 | 28.621784491454278 |

| | | | | |
|---|---|---|---|---|
| **GATCCAGATCTAACTTGGGGTGGCTTTGTCTTCTACACTCTTTTGCCAATTCCACTAATTGTTTGGG** | Ins_k4_34_ACAC | 0.062480076374368365 | 0.03920857715790982 | 52.26563660151634 |
| **GATCCAGATCTAACTTGGGGTGGCTTTGTCTTCTACATCTTTTGCCAATTCCACTAATTGTTTGGG** | Ins_k3_34_ACA | 0.4092946349983866 | 0.03414031646463278 | 73.93258539429867 |
| **GATCCAGATCTAACTTGGGGTGGCTTTGTCTTCTACCTCTTTTGCCAATTCCACTAATTGTTTGGG** | Ins_k3_34_ACC | -0.22194253036663175 | 0.038964695168465624 | 39.327204443909665 |
| **GATCCAGATCTAACTTGGGGTGGCTTTGTCTTCTACGTCTTTTGCCAATTCCACTAATTGTTTGGG** | Ins_k3_34_ACG | 0.5758644676617899 | 0.02822836648722906 | 87.33257243356496 |
| **GATCCAGATCTAACTTGGGGTGGCTTTGTCTTCTACTAATTGTTTGGG** | Del_k15_35-49 | -0.5278774199242672 | 0.030694769036095377 | 28.961912794674546 |
| **GATCCAGATCTAACTTGGGGTGGCTTTGTCTTCTACTCTTTTGCCAATTCCACTAATTGTTTGGG** | Ins_k2_34_AC | 0.20564892712056393 | 0.032996181774513085 | 60.31060518120696 |
| **GATCCAGATCTAACTTGGGGTGGCTTTGTCTTCTACTTCTTTTGCCAATTCCACTAATTGTTTGGG** | Ins_k3_32_CTA;Ins_k3_33_TAC;I | 0.1689112691203779 | 0.034460360051105664 | 58.135140210178434 |
| **GATCCAGATCTAACTTGGGGTGGCTTTGTCTTCTACTTTTGCCAATTCCACTAATTGTTTGGG** | T35A | -0.5567071554706873 | 0.06761751568340768 | 28.138869572595464 |
| **GATCCAGATCTAACTTGGGGTGGCTTTGTCTTCTAGATCTTTTGCCAATTCCACTAATTGTTTGGG** | Ins_k3_34_AGA | -0.5468181205798784 | 0.04758989563443821 | 28.4185162746295 |
| **GATCCAGATCTAACTTGGGGTGGCTTTGTCTTCTAGCTCTTTTGCCAATTCCACTAATTGTTTGGG** | Ins_k3_34_AGC | -3.193991062463348 | 0.14380697315091737 | 2.0134868539573003 |
| **GATCCAGATCTAACTTGGGGTGGCTTTGTCTTCTAGGTCTTTTGCCAATTCCACTAATTGTTTGGG** | Ins_k3_34_AGG | -4.88248089088458 | 0.35712051704661807 | 0.37208913020443146 |
| **GATCCAGATCTAACTTGGGGTGGCTTTGTCTTCTAGTCTTTTGCCAATTCCACTAATTGTTTGGG** | Ins_k2_34_AG | -1.2099781810036094 | 0.06030572313356221 | 14.641805886032023 |
| **GATCCAGATCTAACTTGGGGTGGCTTTGTCTTCTAGTTCTTTTGCCAATTCCACTAATTGTTTGGG** | Ins_k3_33_TAG;Ins_k3_34_AGT | -2.290734891641783 | 0.08616966634132785 | 4.968530603418018 |
| **GATCCAGATCTAACTTGGGGTGGCTTTGTCTTCTATATCTTTTGCCAATTCCACTAATTGTTTGGG** | Ins_k3_34_ATA | -0.7838107107298273 | 0.04925358295385206 | 22.42212790305937 |
| **GATCCAGATCTAACTTGGGGTGGCTTTGTCTTCTATATTCTTTTGCCAATTCCACTAATTGTTTGGG** | Ins_k4_33_TATA;Ins_k4_34_ATAT | -1.1483092110312474 | 0.08437782320628577 | 15.573174079954043 |
| **GATCCAGATCTAACTTGGGGTGGCTTTGTCTTCTATCTCTTTTGCCAATTCCACTAATTGTTTGGG** | Ins_k3_34_ATC | -1.6912262658715396 | 0.06641450928371156 | 9.048805580341494 |
| **GATCCAGATCTAACTTGGGGTGGCTTTGTCTTCTATCTTTTGCCAATTCCACTAATTGTTTGGG** | Ins_k1_34_A | -0.4974388393251999 | 0.04669186142424348 | 29.857026192548997 |
| **GATCCAGATCTAACTTGGGGTGGCTTTGTCTTCTATGTCTTTTGCCAATTCCACTAATTGTTTGGG** | Ins_k3_34_ATG | -0.5763336345531316 | 0.04137115829084824 | 27.591986883119596 |
| **GATCCAGATCTAACTTGGGGTGGCTTTGTCTTCTATTCCACTAATTGTTTGGG** | Del_k10_35-44 | -0.1101615628023 | 0.03347651470298458 | 43.978350203555905 |
| **GATCCAGATCTAACTTGGGGTGGCTTTGTCTTCTATTCTTTTGCCAATTCCACTAATTGTTTGGG** | Ins_k2_33_TA;Ins_k2_34_AT | -0.8204419124567182 | 0.05437888519586889 | 21.615639904777147 |
| **GATCCAGATCTAACTTGGGGTGGCTTTGTCTTCTATTGTTTGGG** | Del_k19_35-53 | -0.34777934313722025 | 0.028373963272928734 | 34.67710571934206 |
| **GATCCAGATCTAACTTGGGGTGGCTTTGTCTTCTATTTCTTTTGCCAATTCCACTAATTGTTTGGG** | Ins_k3_33_TAT;Ins_k3_34_ATT | -1.2298515908078569 | 0.37243759982111134 | 14.353695626438062 |
| **GATCCAGATCTAACTTGGGGTGGCTTTGTCTTCTCAATCTTTTGCCAATTCCACTAATTGTTTGGG** | Ins_k3_34_CAA | -0.27174925672413236 | 0.03629912601274278 | 37.41642496034884 |
| **GATCCAGATCTAACTTGGGGTGGCTTTGTCTTCTCAATTCCACTAATTGTTTGGG** | Del_k8_35-42 | 0.20342821802050415 | 0.029967323630859743 | 60.17682147377927 |
| **GATCCAGATCTAACTTGGGGTGGCTTTGTCTTCTCACTAATTGTTTGGG** | Del_k14_35-48 | -0.474434151109003 | 0.032743060523711426 | 30.551839106511434 |
| **GATCCAGATCTAACTTGGGGTGGCTTTGTCTTCTCACTCTTTTGCCAATTCCACTAATTGTTTGGG** | Ins_k3_34_CAC | -0.4687278672604997 | 0.045645264960212824 | 30.72667492955155 |
| **GATCCAGATCTAACTTGGGGTGGCTTTGTCTTCTCAGTCTTTTGCCAATTCCACTAATTGTTTGGG** | Ins_k3_34_CAG | -0.06282902422871117 | 0.035388505938507606 | 46.11000757832435 |
| **GATCCAGATCTAACTTGGGGTGGCTTTGTCTTCTCATCTTTTGCCAATTCCACTAATTGTTTGGG** | Ins_k2_34_CA | 0.7368155806664181 | 0.030394610844401277 | 100 |
| **GATCCAGATCTAACTTGGGGTGGCTTTGTCTTCTCATTCTTTTGCCAATTCCACTAATTGTTTGGG** | Ins_k3_33_TCA;Ins_k3_34_CAT | -0.6884260799890288 | 0.047856839494509945 | 24.66617704498563 |
| **GATCCAGATCTAACTTGGGGTGGCTTTGTCTTCTCCAATTCCACTAATTGTTTGGG** | Del_k7_35-41 | -0.0190419651991377 | 0.03095059861886953 | 48.173885017024226 |
| **GATCCAGATCTAACTTGGGGTGGCTTTGTCTTCTCCACTAATTGTTTGGG** | Del_k13_34-46;Del_k13_35-47 | -0.14954856992421173 | 0.029755426730070242 | 42.27984372809109 |
| **GATCCAGATCTAACTTGGGGTGGCTTTGTCTTCTCCATCTTTTGCCAATTCCACTAATTGTTTGGG** | Ins_k3_34_CCA | 0.20781224550536492 | 0.03268003001613867 | 60.441217449216076 |
| **GATCCAGATCTAACTTGGGGTGGCTTTGTCTTCTCCCTCTTTTGCCAATTCCACTAATTGTTTGGG** | Ins_k3_34_CCC | -0.5604077921478942 | 0.04369949427061941 | 28.034930279198846 |
| **GATCCAGATCTAACTTGGGGTGGCTTTGTCTTCTCCGTCTTTTGCCAATTCCACTAATTGTTTGGG** | Ins_k3_34_CCG | 0.6845768011682654 | 0.02824757989276981 | 97.36198492673623 |
| **GATCCAGATCTAACTTGGGGTGGCTTTGTCTTCTCCTCTTTTGCCAATTCCACTAATTGTTTGGG** | Ins_k2_34_CC | -0.04745774452556206 | 0.041578491708986365 | 46.8242527700256 |
| **GATCCAGATCTAACTTGGGGTGGCTTTGTCTTCTCCTTCTTTTGCCAATTCCACTAATTGTTTGGG** | Ins_k3_32_CTC;Ins_k3_33_TCC; | -0.10677111013606394 | 0.035254220268391596 | 44.12770977346012 |
| **GATCCAGATCTAACTTGGGGTGGCTTTGTCTTCTCCTTTTGCCAATTCCACTAATTGTTTGGG** | T35C | -0.2847480903015523 | 0.02579909793711315 | 36.93320254723372 |
| **GATCCAGATCTAACTTGGGGTGGCTTTGTCTTCTCGATCTTTTGCCAATTCCACTAATTGTTTGGG** | Ins_k3_34_CGA | 0.34726866667164147 | 0.03320154650094044 | 69.48616732539543 |
| **GATCCAGATCTAACTTGGGGTGGCTTTGTCTTCTCGCTCTTTTGCCAATTCCACTAATTGTTTGGG** | Ins_k3_34_CGC | 0.2755326854687963 | 0.03485473934836903 | 64.67609897968953 |
| **GATCCAGATCTAACTTGGGGTGGCTTTGTCTTCTCGGTCTTTTGCCAATTCCACTAATTGTTTGGG** | Ins_k3_34_CGG | 0.4944381754971052 | 0.03459263236380704 | 80.50322214726573 |
| **GATCCAGATCTAACTTGGGGTGGCTTTGTCTTCTCGTCTTTTGCCAATTCCACTAATTGTTTGGG** | Ins_k2_34_CG | 0.6975176239954652 | 0.03337112502280217 | 98.63011676122575 |
| **GATCCAGATCTAACTTGGGGTGGCTTTGTCTTCTCGTTCTTTTGCCAATTCCACTAATTGTTTGGG** | Ins_k3_33_TCG;Ins_k3_34_CGT | 0.18916226420489168 | 0.03501783235693082 | 59.324436218429874 |
| **GATCCAGATCTAACTTGGGGTGGCTTTGTCTTCTCTAATTGTTTGGG** | Del_k16_35-50 | -0.7571388083020928 | 0.034898194679164316 | 23.028215535830686 |
| **GATCCAGATCTAACTTGGGGTGGCTTTGTCTTCTCTATCTTTTGCCAATTCCACTAATTGTTTGGG** | Ins_k3_34_CTA | -1.596050869303546 | 0.071007080690678 | 9.952344638752347 |
| **GATCCAGATCTAACTTGGGGTGGCTTTGTCTTCTCTCTCTTTTGCCAATTCCACTAATTGTTTGGG** | Ins_k3_34_CTC | -1.7433432974186533 | 0.0719078696075592 | 8.589287071489247 |
| **GATCCAGATCTAACTTGGGGTGGCTTTGTCTTCTCTCTTTTGCCAATTCCACTAATTGTTTGGG** | Ins_k1_34_C | -0.4965596018134326 | 0.04851464188101576 | 29.883289153965826 |
| **GATCCAGATCTAACTTGGGGTGGCTTTGTCTTCTCTGTCTTTTGCCAATTCCACTAATTGTTTGGG** | Ins_k3_34_CTG | -0.6359470956183201 | 0.04485767508162605 | 25.995200873147745 |
| **GATCCAGATCTAACTTGGGGTGGCTTTGTCTTCTCTTCTTTTGCCAATTCCACTAATTGTTTGGG** | Ins_k2_31_TC;Ins_k2_32_CT;Ins_ | 0.3184863330915533 | 0.03576010168586049 | 67.51470107658295 |
| **GATCCAGATCTAACTTGGGGTGGCTTTGTCTTCTCTTTCTTTTGCCAATTCCACTAATTGTTTGGG** | Ins_k3_33_TCT;Ins_k3_34_CTT | -2.4021738554937846 | 0.11509290617882895 | 4.444579124391554 |
| **GATCCAGATCTAACTTGGGGTGGCTTTGTCTTCTCTTTTGCCAATTCCACTAATTGTTTGGG** | Del_k1_34-34;Del_k1_35-35 | -0.1601888412773328 | 0.035830159497609086 | 41.83235961607781 |
| **GATCCAGATCTAACTTGGGGTGGCTTTGTCTTCTG** | Del_k28_35-62 | -2.0758168279878007 | 0.0411965359746264 | 6.159787047967209 |
| **GATCCAGATCTAACTTGGGGTGGCTTTGTCTTCTGAACTCTTTTGCCAATTCCACTAATTGTTTGGG** | Ins_k4_34_GAAC | 0.4693065944776378 | 0.03248827905007108 | 78.50526002866617 |
| **GATCCAGATCTAACTTGGGGTGGCTTTGTCTTCTGAATCTTTTGCCAATTCCACTAATTGTTTGGG** | Ins_k3_34_GAA | 0.4511460700438821 | 0.03308303849667472 | 77.09243102315912 |
| **GATCCAGATCTAACTTGGGGTGGCTTTGTCTTCTGACTCTTTTGCCAATTCCACTAATTGTTTGGG** | Ins_k3_34_GAC | 0.23312213206378862 | 0.03484925853701432 | 61.99050120050164 |
| **GATCCAGATCTAACTTGGGGTGGCTTTGTCTTCTGAGTCTTTTGCCAATTCCACTAATTGTTTGGG** | Ins_k3_34_GAG | 0.4923803219871876 | 0.030575546871800984 | 80.33772864810147 |
| **GATCCAGATCTAACTTGGGGTGGCTTTGTCTTCTGATCTTTTGCCAATTCCACTAATTGTTTGGG** | Ins_k2_34_GA | 0.23424783407946104 | 0.03399464865101309 | 62.060323324735336 |
| **GATCCAGATCTAACTTGGGGTGGCTTTGTCTTCTGATTCTTTTGCCAATTCCACTAATTGTTTGGG** | Ins_k3_33_TGA;Ins_k3_34_GAT | 0.18760113891738517 | 0.03548268697211267 | 59.23189559343144 |
| **GATCCAGATCTAACTTGGGGTGGCTTTGTCTTCTGCAGTCTTTTGCCAATTCCACTAATTGTTTGGG** | Ins_k4_34_GCAG | 0.4974870213554272 | 0.02993056725715049 | 80.74903860051396 |
| **GATCCAGATCTAACTTGGGGTGGCTTTGTCTTCTGCATCTTTTGCCAATTCCACTAATTGTTTGGG** | Ins_k3_34_GCA | 0.37625620823564065 | 0.03145922024552345 | 71.52987846461208 |
| **GATCCAGATCTAACTTGGGGTGGCTTTGTCTTCTGCCAATTCCACTAATTGTTTGGG** | Del_k6_34-39;Del_k6_35-40 | 0.33786814964458944 | 0.02790308085419371 | 68.83602206486016 |
| **GATCCAGATCTAACTTGGGGTGGCTTTGTCTTCTGCCTCTTTTGCCAATTCCACTAATTGTTTGGG** | Ins_k3_34_GCC | 0.27277146550905007 | 0.031202303278651724 | 64.49776037363702 |
| **GATCCAGATCTAACTTGGGGTGGCTTTGTCTTCTGCGTCTTTTGCCAATTCCACTAATTGTTTGGG** | Ins_k3_34_GCG | 0.5624315984706423 | 0.031167933405464036 | 86.16728948078654 |
| **GATCCAGATCTAACTTGGGGTGGCTTTGTCTTCTGCTCTTTTGCCAATTCCACTAATTGTTTGGG** | Ins_k2_34_GC | 0.30149904174551045 | 0.03666754326914986 | 66.37749554671741 |
| **GATCCAGATCTAACTTGGGGTGGCTTTGTCTTCTGCTTCTTTTGCCAATTCCACTAATTGTTTGGG** | Ins_k3_32_CTG;Ins_k3_33_TGC; | -0.21001247871814605 | 0.11506498499832084 | 39.7991898311651 |
| **GATCCAGATCTAACTTGGGGTGGCTTTGTCTTCTGCTTTTGCCAATTCCACTAATTGTTTGGG** | T35G | 0.15316635604120088 | 0.028811108628597423 | 57.226975727405076 |
| **GATCCAGATCTAACTTGGGGTGGCTTTGTCTTCTGG** | Del_k27_35-61 | -2.247056626360034 | 0.043651213050850624 | 5.190356623705014 |
| **GATCCAGATCTAACTTGGGGTGGCTTTGTCTTCTGGATCTTTTGCCAATTCCACTAATTGTTTGGG** | Ins_k3_34_GGA | 0.7184901032295428 | 0.03150916077317477 | 100 |
| **GATCCAGATCTAACTTGGGGTGGCTTTGTCTTCTGGCTCTTTTGCCAATTCCACTAATTGTTTGGG** | Ins_k3_34_GGC | 0.2266257996006762 | 0.03552600287308291 | 61.58909553914961 |
| **GATCCAGATCTAACTTGGGGTGGCTTTGTCTTCTGGG** | Del_k26_34-59;Del_k26_35-60 | -2.2214122155033573 | 0.04463717008150927 | 5.325181626608589 |
| **GATCCAGATCTAACTTGGGGTGGCTTTGTCTTCTGGGTCTTTTGCCAATTCCACTAATTGTTTGGG** | Ins_k3_34_GGG | -0.12778313757668258 | 0.04108667578011693 | 43.21017056230329 |
| **GATCCAGATCTAACTTGGGGTGGCTTTGTCTTCTGGTCTTTTGCCAATTCCACTAATTGTTTGGG** | Ins_k2_34_GG | 0.5549129184409136 | 0.034306627292785614 | 85.52185465169408 |
| **GATCCAGATCTAACTTGGGGTGGCTTTGTCTTCTGGTTCTTTTGCCAATTCCACTAATTGTTTGGG** | Ins_k3_33_TGG;Ins_k3_34_GGT | 0.0746828139791641 | 0.03456723059879143 | 52.90732768199047 |
| **GATCCAGATCTAACTTGGGGTGGCTTTGTCTTCTGTATCTTTTGCCAATTCCACTAATTGTTTGGG** | Ins_k3_34_GTA | -0.45170298119719104 | 0.042452546647313905 | 31.2542714613018 |
| **GATCCAGATCTAACTTGGGGTGGCTTTGTCTTCTGTCTCTTTTGCCAATTCCACTAATTGTTTGGG** | Ins_k3_34_GTC | -0.6425002807715765 | 0.04270001697150923 | 25.8254064634727 |
| **GATCCAGATCTAACTTGGGGTGGCTTTGTCTTCTGTCTTTTGCCAATTCCACTAATTGTTTGGG** | Ins_k1_34_G | 0.14517215078665158 | 0.03763878355502689 | 56.77131528555947 |

| | | | | |
|---|---|---|---|---|
| GATCCAGATCTAACTTGGGGTGGCTTTGTCTTCTGTGTCTTTTGCCAATTCCACTAATTGTTTGGG | Ins_k3_34_GTG | 0.014052409471102312 | 0.03643767462107932 | 49.794843986931085 |
| GATCCAGATCTAACTTGGGGTGGCTTTGTCTTCTGTTCTTTTGCCAATTCCACTAATTGTTTGGG | Ins_k2_33_TG;Ins_k2_34_GT | -0.30125557675757786 | 0.03786456408446312 | 36.32853272668 |
| GATCCAGATCTAACTTGGGGTGGCTTTGTCTTCTGTTTCTTTTGCCAATTCCACTAATTGTTTGGG | Ins_k3_33_TGT;Ins_k3_34_GTT | -0.5067789015619967 | 0.0468972488154291 | 29.579457979739306 |
| GATCCAGATCTAACTTGGGGTGGCTTTGTCTTCTGTTTGGG | Del_k22_34-55;Del_k22_35-56 | -0.6730168569342082 | 0.029671134917159496 | 25.04920718844777 |
| GATCCAGATCTAACTTGGGGTGGCTTTGTCTTCTTAAACTTTTGCCAATTCCACTAATTGTTTGGG | Ins_k3_35_AAA | -0.7322860201054995 | 0.05948536993466346 | 23.607701999379618 |
| GATCCAGATCTAACTTGGGGTGGCTTTGTCTTCTTAACACTTTTGCCAATTCCACTAATTGTTTGGG | Ins_k4_35_AACA | -0.9795922239941975 | 0.05410328890735566 | 18.435290890375317 |
| GATCCAGATCTAACTTGGGGTGGCTTTGTCTTCTTAACCTTTTGCCAATTCCACTAATTGTTTGGG | Ins_k3_35_AAC | -0.588366913239672 | 0.04515572666302683 | 27.261954483734073 |
| GATCCAGATCTAACTTGGGGTGGCTTTGTCTTCTTAACTTTTGCCAATTCCACTAATTGTTTGGG | Ins_k2_35_AA | -0.748047682423235 | 0.051198699243895224 | 23.238522456916925 |
| GATCCAGATCTAACTTGGGGTGGCTTTGTCTTCTTAAGCTTTTGCCAATTCCACTAATTGTTTGGG | Ins_k3_35_AAG | -0.6303693417244141 | 0.043451168795245604 | 26.1406008316678 |
| GATCCAGATCTAACTTGGGGTGGCTTTGTCTTCTTAATCTTTTGCCAATTCCACTAATTGTTTGGG | Ins_k3_34_TAA;Ins_k3_35_AAT | -1.4356058897252366 | 0.058449742844739604 | 11.684382780533223 |
| GATCCAGATCTAACTTGGGGTGGCTTTGTCTTCTTAATTCCACTAATTGTTTGGG | Del_k8_36-43 | -0.4145953837034321 | 0.03369914918434082 | 32.43582919295158 |
| GATCCAGATCTAACTTGGGGTGGCTTTGTCTTCTTAATTGTTTGGG | Del_k17_35-51;Del_k17_36-52 | -0.7644658212765513 | 0.03609842943501086 | 22.860104131132108 |
| GATCCAGATCTAACTTGGGGTGGCTTTGTCTTCTTACACTTTTGCCAATTCCACTAATTGTTTGGG | Ins_k3_35_ACA | -0.370390500926973 | 0.042385504843241315 | 33.90181435916396 |
| GATCCAGATCTAACTTGGGGTGGCTTTGTCTTCTTACCCTTTTGCCAATTCCACTAATTGTTTGGG | Ins_k3_35_ACC | -0.6805973460773302 | 0.0486294394755694 | 24.860039841676734 |
| GATCCAGATCTAACTTGGGGTGGCTTTGTCTTCTTACCTTTTGCCAATTCCACTAATTGTTTGGG | Ins_k2_35_AC | -0.24566216404800573 | 0.03917997624863144 | 38.40535376167585 |
| GATCCAGATCTAACTTGGGGTGGCTTTGTCTTCTTACGCTTTTGCCAATTCCACTAATTGTTTGGG | Ins_k3_35_ACG | 0.17265980104221823 | 0.031206567006451035 | 58.35347059258537 |
| GATCCAGATCTAACTTGGGGTGGCTTTGTCTTCTTACTAATTGTTTGGG | Del_k14_36-49 | -1.1584400623255493 | 0.041156202227559044 | 15.416201046960817 |
| GATCCAGATCTAACTTGGGGTGGCTTTGTCTTCTTACTCTTTTGCCAATTCCACTAATTGTTTGGG | Ins_k3_34_TAC;Ins_k3_35_ACT | -1.3596678400761657 | 0.06359814865803219 | 12.606230737714187 |
| GATCCAGATCTAACTTGGGGTGGCTTTGTCTTCTTACTTTTGCCAATTCCACTAATTGTTTGGG | Ins_k1_35_A | 0.7281651020322302 | 0.5044270232323953 | 100 |
| GATCCAGATCTAACTTGGGGTGGCTTTGTCTTCTTAGACTTTTGCCAATTCCACTAATTGTTTGGG | Ins_k3_35_AGA | -0.7371992320540152 | 0.0483478908042052 | 23.49199683060491 |
| GATCCAGATCTAACTTGGGGTGGCTTTGTCTTCTTAGCCTTTTGCCAATTCCACTAATTGTTTGGG | Ins_k3_35_AGC | -0.9490876767262815 | 0.047942936451427554 | 19.00631624888311 |
| GATCCAGATCTAACTTGGGGTGGCTTTGTCTTCTTAGCTTTTGCCAATTCCACTAATTGTTTGGG | Ins_k2_35_AG | -2.417847797913067 | 0.10243562608021907 | 4.37545816120374 |
| GATCCAGATCTAACTTGGGGTGGCTTTGTCTTCTTAGGCTTTTGCCAATTCCACTAATTGTTTGGG | Ins_k3_35_AGG | -0.9245974668023693 | 0.05509160132836006 | 19.477531451809924 |
| GATCCAGATCTAACTTGGGGTGGCTTTGTCTTCTTAGTCTTTTGCCAATTCCACTAATTGTTTGGG | Ins_k3_34_TAG;Ins_k3_35_AGT | -1.6783301759001816 | 0.07248689247486549 | 9.166255485767804 |
| GATCCAGATCTAACTTGGGGTGGCTTTGTCTTCTTATACTTTTGCCAATTCCACTAATTGTTTGGG | Ins_k3_35_ATA | -0.9304179783668438 | 0.050792075714414735 | 19.364491548911932 |
| GATCCAGATCTAACTTGGGGTGGCTTTGTCTTCTTATCCTTTTGCCAATTCCACTAATTGTTTGGG | Ins_k3_35_ATC | -0.4055471684934019 | 0.040068782177725686 | 32.73064733359848 |
| GATCCAGATCTAACTTGGGGTGGCTTTGTCTTCTTATCTTTTGCCAATTCCACTAATTGTTTGGG | Ins_k2_34_TA;Ins_k2_35_AT | -1.0299838388518765 | 0.05489990670124395 | 17.5293250556747 |
| GATCCAGATCTAACTTGGGGTGGCTTTGTCTTCTTATGCTTTTGCCAATTCCACTAATTGTTTGGG | Ins_k3_35_ATG | -0.5285734393659727 | 0.03956936887667352 | 28.941761753872058 |
| GATCCAGATCTAACTTGGGGTGGCTTTGTCTTCTTATTCCACTAATTGTTTGGG | Del_k9_36-44 | -0.2658009491083787 | 0.034148634234097135 | 37.639652621098875 |
| GATCCAGATCTAACTTGGGGTGGCTTTGTCTTCTTATTCTTTTGCCAATTCCACTAATTGTTTGGG | Ins_k3_33_TTA;Ins_k3_34_TAT;In | -1.529931502692377 | 0.06427485112043024 | 10.632629546769905 |
| GATCCAGATCTAACTTGGGGTGGCTTTGTCTTCTTATTGTTTGGG | Del_k18_36-53 | -0.6605391744043927 | 0.034162963774560316 | 25.363721366787104 |
| GATCCAGATCTAACTTGGGGTGGCTTTGTCTTCTTATTTTGCCAATTCCACTAATTGTTTGGG | C36A | -0.6992701420309393 | 0.04025770777622703 | 24.400140555882093 |
| GATCCAGATCTAACTTGGGGTGGCTTTGTCTTCTTC | Del_k27_37-63 | -3.0305337088562574 | 0.3312326365551487 | 2.3710320534359726 |
| GATCCAGATCTAACTTGGGGTGGCTTTGTCTTCTTCAAATTTTGCCAATTCCACTAATTGTTTGGG | Ins_k3_36_AAA | 0.6085173941840629 | 0.03353628258390764 | 90.23130500395364 |
| GATCCAGATCTAACTTGGGGTGGCTTTGTCTTCTTCAACTTTTGCCAATTCCACTAATTGTTTGGG | Ins_k3_35_CAA;Ins_k3_36_AAC | 0.5216735226102405 | 0.03073384988745537 | 82.72588546030025 |
| GATCCAGATCTAACTTGGGGTGGCTTTGTCTTCTTCAAGTTTTGCCAATTCCACTAATTGTTTGGG | Ins_k3_36_AAG | 0.5938062206496781 | 0.03174307014942516 | 88.913612781623 |
| GATCCAGATCTAACTTGGGGTGGCTTTGTCTTCTTCAATTCCACTAATTGTTTGGG | Del_k7_36-42;Del_k7_37-43 | 0.2304525280828965 | 0.027726439963475433 | 61.825231811781585 |
| GATCCAGATCTAACTTGGGGTGGCTTTGTCTTCTTCAATTGTTTGGG | Del_k16_37-52 | 0.03957079939850583 | 0.027532812349857383 | 51.08187993360295 |
| GATCCAGATCTAACTTGGGGTGGCTTTGTCTTCTTCAATTTTGCCAATTCCACTAATTGTTTGGG | Ins_k2_36_AA | 0.7086441273112554 | 0.03409597112325124 | 99.73365294721603 |
| GATCCAGATCTAACTTGGGGTGGCTTTGTCTTCTTCAATTTTTGCCAATTCCACTAATTGTTTGGG | Ins_k3_36_AAT | 0.536864686231048 | 0.03274096459514493 | 83.99218182766441 |
| GATCCAGATCTAACTTGGGGTGGCTTTGTCTTCTTCACATTTTGCCAATTCCACTAATTGTTTGGG | Ins_k3_36_ACA | 0.2975388186594967 | 0.03318604176514449 | 66.11514568133563 |
| GATCCAGATCTAACTTGGGGTGGCTTTGTCTTCTTCACCTTTTGCCAATTCCACTAATTGTTTGGG | Ins_k3_35_CAC;Ins_k3_36_ACC | 0.28318426006828845 | 0.028693443912625648 | 65.17287109660194 |
| GATCCAGATCTAACTTGGGGTGGCTTTGTCTTCTTCACGTTTTGCCAATTCCACTAATTGTTTGGG | Ins_k3_36_ACG | 0.6873539238412931 | 0.031393744831303874 | 97.63274689815323 |
| GATCCAGATCTAACTTGGGGTGGCTTTGTCTTCTTCACTAATTGTTTGGG | Del_k13_36-48;Del_k13_37-49 | 4.586473443639605e-4 | 0.02754259931678145 | 49.12252474967173 |
| GATCCAGATCTAACTTGGGGTGGCTTTGTCTTCTTCACTTTTGCCAATTCCACTAATTGTTTGGG | Ins_k2_35_CA;Ins_k2_36_AC | 0.26280995182169375 | 0.03447635470727212 | 63.858454564267475 |
| GATCCAGATCTAACTTGGGGTGGCTTTGTCTTCTTCACTTTTTGCCAATTCCACTAATTGTTTGGG | Ins_k3_36_ACT | 0.11583878959081961 | 0.035919910849095606 | 55.130209041762605 |
| GATCCAGATCTAACTTGGGGTGGCTTTGTCTTCTTCAGATTTTGCCAATTCCACTAATTGTTTGGG | Ins_k3_36_AGA | 0.28935658212101756 | 0.03282460384455402 | 65.5763830679632 |
| GATCCAGATCTAACTTGGGGTGGCTTTGTCTTCTTCAGCTTTTGCCAATTCCACTAATTGTTTGGG | Ins_k3_35_CAG;Ins_k3_36_AGC | 0.3258105096597531 | 0.029991004307877496 | 68.01100596189472 |
| GATCCAGATCTAACTTGGGGTGGCTTTGTCTTCTTCAGGTTTTGCCAATTCCACTAATTGTTTGGG | Ins_k3_36_AGG | -0.6265835005647037 | 0.04922699225724811 | 26.23975256216198 |
| GATCCAGATCTAACTTGGGGTGGCTTTGTCTTCTTCAGTTTTGCCAATTCCACTAATTGTTTGGG | Ins_k2_36_AG | 0.4251974350878427 | 0.03064808711438642 | 75.11771901495311 |
| GATCCAGATCTAACTTGGGGTGGCTTTGTCTTCTTCAGTTTTTGCCAATTCCACTAATTGTTTGGG | Ins_k3_36_AGT | 0.6523386346254323 | 0.032075886406346496 | 94.27326783360034 |
| GATCCAGATCTAACTTGGGGTGGCTTTGTCTTCTTCATATTTTGCCAATTCCACTAATTGTTTGGG | Ins_k3_36_ATA | 0.20820390037152203 | 0.03328737342000299 | 60.46489418239774 |
| GATCCAGATCTAACTTGGGGTGGCTTTGTCTTCTTCATCTTTTGCCAATTCCACTAATTGTTTGGG | Ins_k3_34_TCA;Ins_k3_35_CAT;I | 0.328465579229158 | 0.03417237046910795 | 68.19181984470875 |
| GATCCAGATCTAACTTGGGGTGGCTTTGTCTTCTTCATGCTTTTGCCAATTCCACTAATTGTTTGGG | Ins_k4_35_CATG;Ins_k4_36_ATG | 0.40807014410417874 | 0.033839194370226706 | 73.84211102052613 |
| GATCCAGATCTAACTTGGGGTGGCTTTGTCTTCTTCATGTTTTGCCAATTCCACTAATTGTTTGGG | Ins_k3_36_ATG | 0.40808793393632237 | 0.031210121764604318 | 73.8434246709711 |
| GATCCAGATCTAACTTGGGGTGGCTTTGTCTTCTTCATTCCACTAATTGTTTGGG | Del_k8_37-44 | 0.15369495478615702 | 0.030389081563482797 | 57.257233831445575 |
| GATCCAGATCTAACTTGGGGTGGCTTTGTCTTCTTCATTGTTTGGG | Del_k17_37-53 | -0.038691107360576016 | 0.028226541852156208 | 47.23654858795956 |
| GATCCAGATCTAACTTGGGGTGGCTTTGTCTTCTTCATTTGCCAATTCCACTAATTGTTTGGG | T37A | 0.12186676872100366 | 0.030070935885688946 | 55.46353642724113 |
| GATCCAGATCTAACTTGGGGTGGCTTTGTCTTCTTCATTTTGCCAATTCCACTAATTGTTTGGG | Ins_k1_36_A | 0.4719966928131105 | 0.034598887796617975 | 78.71673120960287 |
| GATCCAGATCTAACTTGGGGTGGCTTTGTCTTCTTCATTTTTGCCAATTCCACTAATTGTTTGGG | Ins_k2_36_AT | 0.40702799369920384 | 0.03201214513635181 | 73.76519651982218 |
| GATCCAGATCTAACTTGGGGTGGCTTTGTCTTCTTCATTTTTTGCCAATTCCACTAATTGTTTGGG | Ins_k3_36_ATT | 0.09829044461520198 | 0.03615869864260116 | 54.171204198718215 |
| GATCCAGATCTAACTTGGGGTGGCTTTGTCTTCTTCCAATTCCACTAATTGTTTGGG | Del_k6_36-41;Del_k6_37-42 | 0.04397624870811423 | 0.029405354668077397 | 51.307414993161196 |
| GATCCAGATCTAACTTGGGGTGGCTTTGTCTTCTTCCAATTTTGCCAATTCCACTAATTGTTTGGG | Ins_k3_36_CAA | 0.33923713789468846 | 0.03167758330050255 | 68.93032230348228 |
| GATCCAGATCTAACTTGGGGTGGCTTTGTCTTCTTCCACTAATTGTTTGGG | Del_k12_34-45;Del_k12_35-46;D | -0.048091686161024993 | 0.02965278367904925 | 46.79457833356943 |
| GATCCAGATCTAACTTGGGGTGGCTTTGTCTTCTTCCACTTTTGCCAATTCCACTAATTGTTTGGG | Ins_k3_35_CCA;Ins_k3_36_CAC | 0.25145590458905154 | 0.03322256225223536 | 63.137503257601836 |
| GATCCAGATCTAACTTGGGGTGGCTTTGTCTTCTTCCAGTTTTGCCAATTCCACTAATTGTTTGGG | Ins_k3_36_CAG | 0.7048778428090485 | 0.030502081978589543 | 99.35873410446803 |
| GATCCAGATCTAACTTGGGGTGGCTTTGTCTTCTTCCATTTTGCCAATTCCACTAATTGTTTGGG | Ins_k2_36_CA | 0.5481445675710193 | 0.03326522242204498 | 84.94496722292614 |
| GATCCAGATCTAACTTGGGGTGGCTTTGTCTTCTTCCATTTTTGCCAATTCCACTAATTGTTTGGG | Ins_k3_36_CAT | 0.6683996893774773 | 0.16762813030491971 | 95.7996205562204 |
| GATCCAGATCTAACTTGGGGTGGCTTTGTCTTCTTCCCAATTCCACTAATTGTTTGGG | Del_k5_37-41 | -0.05388632778698832 | 0.032321304156568885 | 46.524204637971216 |
| GATCCAGATCTAACTTGGGGTGGCTTTGTCTTCTTCCCACTAATTGTTTGGG | Del_k11_37-47 | -0.323777273629536 | 0.03152022781491041 | 35.51949714858779 |
| GATCCAGATCTAACTTGGGGTGGCTTTGTCTTCTTCCCAGTTTTGCCAATTCCACTAATTGTTTGGG | Ins_k4_36_CCAG | 0.45730601616860156 | 0.0348800527933743 | 77.56878188645497 |
| GATCCAGATCTAACTTGGGGTGGCTTTGTCTTCTTCCCATTTTGCCAATTCCACTAATTGTTTGGG | Ins_k3_36_CCA | 0.014613366528420935 | 0.0362591969907232 | 49.822784592090784 |
| GATCCAGATCTAACTTGGGGTGGCTTTGTCTTCTTCCCCTTTTGCCAATTCCACTAATTGTTTGGG | Ins_k3_35_CCC;Ins_k3_36_CCC | -0.611111100934579 | 0.0420098025595092 | 26.648901596356694 |
| GATCCAGATCTAACTTGGGGTGGCTTTGTCTTCTTCCCGTTTTGCCAATTCCACTAATTGTTTGGG | Ins_k3_36_CCG | 0.6056196435356223 | 0.3496093367170438 | 89.97021564998751 |
| GATCCAGATCTAACTTGGGGTGGCTTTGTCTTCTTCCCTTTTGCCAATTCCACTAATTGTTTGGG | Ins_k2_35_CC;Ins_k2_36_CC | -0.16847151941095284 | 0.03901736673996201 | 41.48730659997379 |

| | | | | |
|---|---|---|---|---|
| **GATCCAGATCTAACTTGGGGTGGCTTTGTCTTCTTCCCTTTTTGCCAATTCCACTAATTGTTTGGG** | Ins_k3_36_CCT | -0.5116293002561544 | 0.04604513487547953 | 29.436333202098172 |
| **GATCCAGATCTAACTTGGGGTGGCTTTGTCTTCTTCCGATTTTGCCAATTCCACTAATTGTTTGGG** | Ins_k3_36_CGA | 0.524576334850251 | 0.03129520988756458 | 82.96637204801726 |
| **GATCCAGATCTAACTTGGGGTGGCTTTGTCTTCTTCCGATTTTTGCCAATTCCACTAATTGTTTGGG** | Ins_k4_36_CGAT | 0.6992122996860104 | 0.030077135654780002 | 98.79740453167909 |
| **GATCCAGATCTAACTTGGGGTGGCTTTGTCTTCTTCCGCTTTTGCCAATTCCACTAATTGTTTGGG** | Ins_k3_35_CCG;Ins_k3_36_CGC | 0.4596703424785007 | 0.02898586428240954 | 77.75239677551797 |
| **GATCCAGATCTAACTTGGGGTGGCTTTGTCTTCTTCCGGTTTTGCCAATTCCACTAATTGTTTGGG** | Ins_k3_36_CGG | 0.9334280677749376 | 0.03106063211886264 | 100 |
| **GATCCAGATCTAACTTGGGGTGGCTTTGTCTTCTTCCGTTTTGCCAATTCCACTAATTGTTTGGG** | Ins_k2_36_CG | 0.5308487991166406 | 0.033843463102323584 | 83.48841117630657 |
| **GATCCAGATCTAACTTGGGGTGGCTTTGTCTTCTTCCGTTTTTGCCAATTCCACTAATTGTTTGGG** | Ins_k3_36_CGT | 0.6318333577339619 | 0.031536728321999684 | 92.35985296744344 |
| **GATCCAGATCTAACTTGGGGTGGCTTTGTCTTCTTCCTAATTGTTTGGG** | Del_k14_37-50 | -0.35463639711362815 | 0.029301387776480666 | 34.44013631817457 |
| **GATCCAGATCTAACTTGGGGTGGCTTTGTCTTCTTCCTATTTTGCCAATTCCACTAATTGTTTGGG** | Ins_k3_36_CTA | -0.2363817084737705 | 0.03649774482122361 | 38.76343193535677 |
| **GATCCAGATCTAACTTGGGGTGGCTTTGTCTTCTTCCTCTTTTGCCAATTCCACTAATTGTTTGGG** | Ins_k3_34_TCC;Ins_k3_35_CCT;I | -0.3220509159868054 | 0.040969336797629125 | 35.5808694639967 |
| **GATCCAGATCTAACTTGGGGTGGCTTTGTCTTCTTCCTGTTTTGCCAATTCCACTAATTGTTTGGG** | Ins_k3_36_CTG | 0.5044811951289565 | 0.030703245160597973 | 81.315791080811607 |
| **GATCCAGATCTAACTTGGGGTGGCTTTGTCTTCTTCCTTTGCCAATTCCACTAATTGTTTGGG** | T37C | -0.2649349946357061 | 0.025218827952290423 | 37.67226096326815 |
| **GATCCAGATCTAACTTGGGGTGGCTTTGTCTTCTTCCTTTTGCCAATTCCACTAATTGTTTGGG** | Ins_k1_35_C;Ins_k1_36_C | 0.12398853686903577 | 0.0368546925328385 | 55.581342126203744 |
| **GATCCAGATCTAACTTGGGGTGGCTTTGTCTTCTTCCTTTTTGCCAATTCCACTAATTGTTTGGG** | Ins_k2_36_CT | 0.08417608803882803 | 0.03692268352572297 | 53.41198306933795 |
| **GATCCAGATCTAACTTGGGGTGGCTTTGTCTTCTTCCTTTTTTGCCAATTCCACTAATTGTTTGGG** | Ins_k3_36_CTT | -0.47876262474001685 | 0.0445090886284425 | 30.419882068864247 |
| **GATCCAGATCTAACTTGGGGTGGCTTTGTCTTCTTCG** | Del_k26_37-62 | -1.1990426027752386 | 0.030086820202132675 | 14.80280118363187 |
| **GATCCAGATCTAACTTGGGGTGGCTTTGTCTTCTTCGAACTTTTGCCAATTCCACTAATTGTTTGGG** | Ins_k4_35_CGAA;Ins_k4_36_GA | 0.7153760810775689 | 0.03028573034734721 | 100 |
| **GATCCAGATCTAACTTGGGGTGGCTTTGTCTTCTTCGAATTTTGCCAATTCCACTAATTGTTTGGG** | Ins_k3_36_GAA | 0.664441024478925 | 0.029781414840820447 | 95.42113161075709 |
| **GATCCAGATCTAACTTGGGGTGGCTTTGTCTTCTTCGACTTTTGCCAATTCCACTAATTGTTTGGG** | Ins_k3_35_CGA;Ins_k3_36_GAC | 0.5872155118631663 | 0.03099929468422903 | 88.32953590714921 |
| **GATCCAGATCTAACTTGGGGTGGCTTTGTCTTCTTCGAGTTTTGCCAATTCCACTAATTGTTTGGG** | Ins_k3_36_GAG | 1.1009056251513085 | 0.030654565854959782 | 100 |
| **GATCCAGATCTAACTTGGGGTGGCTTTGTCTTCTTCGATTTTGCCAATTCCACTAATTGTTTGGG** | Ins_k2_36_GA | 0.6912278436419887 | 0.02912692051516035 | 98.01170187623217 |
| **GATCCAGATCTAACTTGGGGTGGCTTTGTCTTCTTCGATTTTTGCCAATTCCACTAATTGTTTGGG** | Ins_k3_36_GAT | 0.5950421366844116 | 0.029980450758430338 | 89.02357047660848 |
| **GATCCAGATCTAACTTGGGGTGGCTTTGTCTTCTTCGCATTTTGCCAATTCCACTAATTGTTTGGG** | Ins_k3_36_GCA | 0.5988836735045413 | 0.029319676520497894 | 89.36621552085143 |
| **GATCCAGATCTAACTTGGGGTGGCTTTGTCTTCTTCGCCAATTCCACTAATTGTTTGGG** | Del_k4_37-40 | 0.5492002618439051 | 0.0286636573709853 | 85.03469049017056 |
| **GATCCAGATCTAACTTGGGGTGGCTTTGTCTTCTTCGCCTTTTGCCAATTCCACTAATTGTTTGGG** | Ins_k3_35_CGC;Ins_k3_36_GCC | 0.6382745109588195 | 0.029678127498476935 | 92.95667698626806 |
| **GATCCAGATCTAACTTGGGGTGGCTTTGTCTTCTTCGCGTTTTGCCAATTCCACTAATTGTTTGGG** | Ins_k3_36_GCG | 0.9218811532661235 | 0.031270442896463746 | 100 |
| **GATCCAGATCTAACTTGGGGTGGCTTTGTCTTCTTCGCTTTTGCCAATTCCACTAATTGTTTGGG** | Ins_k2_35_CG;Ins_k2_36_GC | 0.6401435407017262 | 0.0284182534253773 | 93.1305782430441 |
| **GATCCAGATCTAACTTGGGGTGGCTTTGTCTTCTTCGCTTTTTGCCAATTCCACTAATTGTTTGGG** | Ins_k3_36_GCT | 0.34231554261741426 | 0.0319360692431606 | 69.14284468031259 |
| **GATCCAGATCTAACTTGGGGTGGCTTTGTCTTCTTCGG** | Del_k25_37-61 | -0.9136856092476692 | 0.030985592811111982 | 19.691231311334217 |
| **GATCCAGATCTAACTTGGGGTGGCTTTGTCTTCTTCGGATTTTGCCAATTCCACTAATTGTTTGGG** | Ins_k3_36_GGA | 0.6908252389557696 | 0.030383181216821892 | 97.9722498480711 |
| **GATCCAGATCTAACTTGGGGTGGCTTTGTCTTCTTCGGCTTTTGCCAATTCCACTAATTGTTTGGG** | Ins_k3_35_CGG;Ins_k3_36_GGC | 0.5054390376859572 | 0.03009786110544705 | 81.39371612006664 |
| **GATCCAGATCTAACTTGGGGTGGCTTTGTCTTCTTCGGG** | Del_k24_37-60 | -1.298395201695575 | 0.03192513290947202 | 13.402802583420181 |
| **GATCCAGATCTAACTTGGGGTGGCTTTGTCTTCTTCGGGTTTTGCCAATTCCACTAATTGTTTGGG** | Ins_k3_36_GGG | 0.8158605828647041 | 0.03131442018909929 | 100 |
| **GATCCAGATCTAACTTGGGGTGGCTTTGTCTTCTTCGGTTTTGCCAATTCCACTAATTGTTTGGG** | Ins_k2_36_GG | 0.947582727171725 | 0.030537713111707426 | 100 |
| **GATCCAGATCTAACTTGGGGTGGCTTTGTCTTCTTCGGTTTTTGCCAATTCCACTAATTGTTTGGG** | Ins_k3_36_GGT | 0.8451419318478397 | 0.03265428271740331 | 100 |
| **GATCCAGATCTAACTTGGGGTGGCTTTGTCTTCTTCGTATTTTGCCAATTCCACTAATTGTTTGGG** | Ins_k3_36_GTA | 0.4968269233028766 | 0.031247002620034357 | 80.6957539058848 |
| **GATCCAGATCTAACTTGGGGTGGCTTTGTCTTCTTCGTCTTTTGCCAATTCCACTAATTGTTTGGG** | Ins_k3_34_TCG;Ins_k3_35_CGT; | 0.47050299075597346 | 0.030781659108799125 | 78.59923963681103 |
| **GATCCAGATCTAACTTGGGGTGGCTTTGTCTTCTTCGTGTTTTGCCAATTCCACTAATTGTTTGGG** | Ins_k3_36_GTG | 0.5419500495396425 | 0.027988497632512106 | 84.42040048822821 |
| **GATCCAGATCTAACTTGGGGTGGCTTTGTCTTCTTCGTTTGCCAATTCCACTAATTGTTTGGG** | T37G | 0.3720873840267368 | 0.0271519840043738 | 71.23230367506709 |
| **GATCCAGATCTAACTTGGGGTGGCTTTGTCTTCTTCGTTTGGG** | Del_k20_37-56 | -0.29955379093196227 | 0.028038769540507408 | 36.390408743670875 |
| **GATCCAGATCTAACTTGGGGTGGCTTTGTCTTCTTCGTTTTGCCAATTCCACTAATTGTTTGGG** | Ins_k1_36_G | 0.7069321937152924 | 0.02919251484659655 | 99.56306161826409 |
| **GATCCAGATCTAACTTGGGGTGGCTTTGTCTTCTTCGTTTTTGCCAATTCCACTAATTGTTTGGG** | Ins_k2_36_GT | 0.6235312016836143 | 0.033341823594618854 | 91.5962412531678 |
| **GATCCAGATCTAACTTGGGGTGGCTTTGTCTTCTTCGTTTTTTGCCAATTCCACTAATTGTTTGGG** | Ins_k3_36_GTT | 0.5603009906043017 | 0.03589530857777265 | 85.98389621494093 |
| **GATCCAGATCTAACTTGGGGTGGCTTTGTCTTCTTCT** | Del_k26_38-63 | -6.118758297695417 | 0.47796992127173443 | 0.10807830558208416 |
| **GATCCAGATCTAACTTGGGGTGGCTTTGTCTTCTTCTAAATTTGCCAATTCCACTAATTGTTTGGG** | Ins_k3_37_AAA | -0.05217680800732416 | 0.036945536532317094 | 46.603806707302844 |
| **GATCCAGATCTAACTTGGGGTGGCTTTGTCTTCTTCTAACTTTGCCAATTCCACTAATTGTTTGGG** | Ins_k3_37_AAC | -0.34949587921621683 | 0.04182085267265843 | 34.617632275019275 |
| **GATCCAGATCTAACTTGGGGTGGCTTTGTCTTCTTCTAAGTTTGCCAATTCCACTAATTGTTTGGG** | Ins_k3_37_AAG | 0.08176173127194941 | 0.03506030393314816 | 53.283183033766704 |
| **GATCCAGATCTAACTTGGGGTGGCTTTGTCTTCTTCTAATTCCACTAATTGTTTGGG** | Del_k6_38-43 | -0.23090021238413905 | 0.031233518973646793 | 38.976496959987564 |
| **GATCCAGATCTAACTTGGGGTGGCTTTGTCTTCTTCTAATTGTTTGGG** | Del_k15_36-50;Del_k15_37-51;D | -0.37740568424183174 | 0.030815784101893198 | 33.664819171005156 |
| **GATCCAGATCTAACTTGGGGTGGCTTTGTCTTCTTCTAATTTGCCAATTCCACTAATTGTTTGGG** | Ins_k2_37_AA | 0.896832051558581 | 0.515828531184252 | 100 |
| **GATCCAGATCTAACTTGGGGTGGCTTTGTCTTCTTCTAATTTTGCCAATTCCACTAATTGTTTGGG** | Ins_k3_36_TAA;Ins_k3_37_AAT | -0.18940192692449287 | 0.04086206947914086 | 40.62798471569757 |
| **GATCCAGATCTAACTTGGGGTGGCTTTGTCTTCTTCTACATTTGCCAATTCCACTAATTGTTTGGG** | Ins_k3_37_ACA | 0.43243115256098763 | 0.030242964646929507 | 75.66306944898732 |
| **GATCCAGATCTAACTTGGGGTGGCTTTGTCTTCTTCTACCTTTGCCAATTCCACTAATTGTTTGGG** | Ins_k3_37_ACC | 0.14030268050244432 | 0.03115246081134993 | 56.49554103496403 |
| **GATCCAGATCTAACTTGGGGTGGCTTTGTCTTCTTCTACGTTTGCCAATTCCACTAATTGTTTGGG** | Ins_k3_37_ACG | 0.6729752837897269 | 0.028395725697614435 | 96.23896512950847 |
| **GATCCAGATCTAACTTGGGGTGGCTTTGTCTTCTTCTACTAATTGTTTGGG** | Del_k12_38-49 | -0.03171521295838986 | 0.027447591979033462 | 47.56721777856278 |
| **GATCCAGATCTAACTTGGGGTGGCTTTGTCTTCTTCTACTTTGCCAATTCCACTAATTGTTTGGG** | Ins_k2_37_AC | 0.38684210482004167 | 0.03322025304990877 | 72.29110840275213 |
| **GATCCAGATCTAACTTGGGGTGGCTTTGTCTTCTTCTACTTTTGCCAATTCCACTAATTGTTTGGG** | Ins_k3_35_CTA;Ins_k3_36_TAC;I | 0.2011978396166656 | 0.03388530900266424 | 60.04275395697712 |
| **GATCCAGATCTAACTTGGGGTGGCTTTGTCTTCTTCTAGATTTGCCAATTCCACTAATTGTTTGGG** | Ins_k3_37_AGA | -0.43428799219525405 | 0.03861301771194266 | 31.80333131412628 |
| **GATCCAGATCTAACTTGGGGTGGCTTTGTCTTCTTCTAGCATTTGCCAATTCCACTAATTGTTTGGG** | Ins_k4_37_AGCA | -1.4528447577891377 | 0.0643093357239819 | 11.484683487756536 |
| **GATCCAGATCTAACTTGGGGTGGCTTTGTCTTCTTCTAGCTTTGCCAATTCCACTAATTGTTTGGG** | Ins_k3_37_AGC | -2.3251786865209154 | 0.09438108886504885 | 4.800309275718695 |
| **GATCCAGATCTAACTTGGGGTGGCTTTGTCTTCTTCTAGGTTTGCCAATTCCACTAATTGTTTGGG** | Ins_k3_37_AGG | -4.8462339788252375 | 0.37093430534845234 | 0.3858236249879253 |
| **GATCCAGATCTAACTTGGGGTGGCTTTGTCTTCTTCTAGTTTGCCAATTCCACTAATTGTTTGGG** | Ins_k2_37_AG | -0.6245154532334098 | 0.04505765863203052 | 26.29407376246847 |
| **GATCCAGATCTAACTTGGGGTGGCTTTGTCTTCTTCTAGTTTTGCCAATTCCACTAATTGTTTGGG** | Ins_k3_36_TAG;Ins_k3_37_AGT | -0.9851777015639327 | 0.05296048031984444 | 18.33260802000465 |
| **GATCCAGATCTAACTTGGGGTGGCTTTGTCTTCTTCTATATTTGCCAATTCCACTAATTGTTTGGG** | Ins_k3_37_ATA | -0.09617923568600362 | 0.03546972108338532 | 44.597588993115 |
| **GATCCAGATCTAACTTGGGGTGGCTTTGTCTTCTTCTATATTTTGCCAATTCCACTAATTGTTTGGG** | Ins_k4_36_TATA;Ins_k4_37_ATAT | -0.2869456973764616 | 0.050831052518271864 | 36.85212699874571 |
| **GATCCAGATCTAACTTGGGGTGGCTTTGTCTTCTTCTATCTTTGCCAATTCCACTAATTGTTTGGG** | Ins_k3_37_ATC | -0.8056438627000553 | 0.045373710793482305 | 21.937887658230846 |
| **GATCCAGATCTAACTTGGGGTGGCTTTGTCTTCTTCTATGTTTGCCAATTCCACTAATTGTTTGGG** | Ins_k3_37_ATG | 0.06439242600271433 | 0.03302561822567009 | 52.36568240303256 |
| **GATCCAGATCTAACTTGGGGTGGCTTTGTCTTCTTCTATTCCACTAATTGTTTGGG** | Del_k7_38-44 | 9.904564890517165e-5 | 0.03196859917046225 | 49.10486338220569 |
| **GATCCAGATCTAACTTGGGGTGGCTTTGTCTTCTTCTATTGCCAATTCCACTAATTGTTTGGG** | T38A | -0.13549794503298146 | 0.03254885749973498 | 42.87809501276887 |
| **GATCCAGATCTAACTTGGGGTGGCTTTGTCTTCTTCTATTGTTTGGG** | Del_k16_38-53 | -0.1574471658610253 | 0.031301889035593944 | 41.947207734233714 |
| **GATCCAGATCTAACTTGGGGTGGCTTTGTCTTCTTCTATTTGCCAATTCCACTAATTGTTTGGG** | Ins_k1_37_A | 0.229115394733167 | 0.03803691662696313 | 61.7426184773025 |
| **GATCCAGATCTAACTTGGGGTGGCTTTGTCTTCTTCTATTTTGCCAATTCCACTAATTGTTTGGG** | Ins_k2_36_TA;Ins_k2_37_AT | 0.07469703397741956 | 0.04232239517135973 | 52.90808002944699 |
| **GATCCAGATCTAACTTGGGGTGGCTTTGTCTTCTTCTATTTTTGCCAATTCCACTAATTGTTTGGG** | Ins_k3_36_TAT;Ins_k3_37_ATT | -0.24119628868975035 | 0.043257692579987 | 38.577250834422415 |
| **GATCCAGATCTAACTTGGGGTGGCTTTGTCTTCTTCTCAATTCCACTAATTGTTTGGG** | Del_k5_38-42 | 0.12190904718912443 | 0.03029297642185113 | 55.46588139016821 |
| **GATCCAGATCTAACTTGGGGTGGCTTTGTCTTCTTCTCAATTTGCCAATTCCACTAATTGTTTGGG** | Ins_k3_37_CAA | 0.19928277167904196 | 0.03266372203233547 | 59.92787803669217 |

| | | | | |
|---|---|---|---|---|
| **GATCCAGATCTAACTTGGGGTGGCTTTGTCTTCTTCTCACTAATTGTTTGGG** | Del_k11_38-48 | -0.2683178411645331 | 0.03308042891875959 | 37.545036797275365 |
| **GATCCAGATCTAACTTGGGGTGGCTTTGTCTTCTTCTCACTTTGCCAATTCCACTAATTGTTTGGG** | Ins_k3_37_CAC | -0.0738823798561331 | 0.03456247368796764 | 45.6031437004941 |
| **GATCCAGATCTAACTTGGGGTGGCTTTGTCTTCTTCTCAGTTTGCCAATTCCACTAATTGTTTGGG** | Ins_k3_37_CAG | 0.11856951910850944 | 0.032518477160026475 | 55.280960467916 |
| **GATCCAGATCTAACTTGGGGTGGCTTTGTCTTCTTCTCATTTGCCAATTCCACTAATTGTTTGGG** | Ins_k2_37_CA | 0.4221496826335076 | 0.03302378870694541 | 74.88912732484859 |
| **GATCCAGATCTAACTTGGGGTGGCTTTGTCTTCTTCTCATTTTGCCAATTCCACTAATTGTTTGGG** | Ins_k3_36_TCA;Ins_k3_37_CAT | 0.07786693765757735 | 0.03409146209696911 | 53.07605964598579 |
| **GATCCAGATCTAACTTGGGGTGGCTTTGTCTTCTTCTCCAATTCCACTAATTGTTTGGG** | Del_k4_38-41 | 0.5241974826236431 | 0.029140443171074528 | 82.93494600652208 |
| **GATCCAGATCTAACTTGGGGTGGCTTTGTCTTCTTCTCCACTAATTGTTTGGG** | Del_k10_37-46;Del_k10_38-47 | 0.4140656083704388 | 0.026964812079615357 | 74.28615856395596 |
| **GATCCAGATCTAACTTGGGGTGGCTTTGTCTTCTTCTCCATTTGCCAATTCCACTAATTGTTTGGG** | Ins_k3_37_CCA | 0.3747925226544899 | 0.029077740349049298 | 71.42525779744103 |
| **GATCCAGATCTAACTTGGGGTGGCTTTGTCTTCTTCTCCCTTTGCCAATTCCACTAATTGTTTGGG** | Ins_k3_37_CCC | -0.08615690927577357 | 0.033692543019541346 | 45.04680793633327 |
| **GATCCAGATCTAACTTGGGGTGGCTTTGTCTTCTTCTCCGTTTGCCAATTCCACTAATTGTTTGGG** | Ins_k3_37_CCG | 0.5154754182333554 | 0.02714809858649063 | 82.2147275290596 |
| **GATCCAGATCTAACTTGGGGTGGCTTTGTCTTCTTCTCCTTTGCCAATTCCACTAATTGTTTGGG** | Ins_k2_37_CC | 0.3152928588408457 | 0.032160794687942446 | 67.2994385178122 |
| **GATCCAGATCTAACTTGGGGTGGCTTTGTCTTCTTCTCCTTTTGCCAATTCCACTAATTGTTTGGG** | Ins_k3_35_CTC;Ins_k3_36_TCC; | -0.04670471161087664 | 0.03428353333553147 | 46.859526252946964 |
| **GATCCAGATCTAACTTGGGGTGGCTTTGTCTTCTTCTCGATTTGCCAATTCCACTAATTGTTTGGG** | Ins_k3_37_CGA | 0.5256181613454263 | 0.0312762664660262 | 83.05285365421753 |
| **GATCCAGATCTAACTTGGGGTGGCTTTGTCTTCTTCTCGCTTTGCCAATTCCACTAATTGTTTGGG** | Ins_k3_37_CGC | 0.22184558107698638 | 0.03213202047580429 | 61.29538875438103 |
| **GATCCAGATCTAACTTGGGGTGGCTTTGTCTTCTTCTCGGTTTGCCAATTCCACTAATTGTTTGGG** | Ins_k3_37_CGG | 0.44036943292732733 | 0.039650185508540034 | 76.26609443164463 |
| **GATCCAGATCTAACTTGGGGTGGCTTTGTCTTCTTCTCGTTTGCCAATTCCACTAATTGTTTGGG** | Ins_k2_37_CG | 0.5029996673231569 | 0.02990530398841866 | 81.19540867223681 |
| **GATCCAGATCTAACTTGGGGTGGCTTTGTCTTCTTCTCGTTTTGCCAATTCCACTAATTGTTTGGG** | Ins_k3_36_TCG;Ins_k3_37_CGT | 0.4260810325184059 | 0.031873062865119255 | 75.18412217097318 |
| **GATCCAGATCTAACTTGGGGTGGCTTTGTCTTCTTCTCGTTTTTGCCAATTCCACTAATTGTTTGGG** | Ins_k4_36_TCGT;Ins_k4_37_CGT | 0.5594102272224857 | 0.03385305034680232 | 85.90733901100583 |
| **GATCCAGATCTAACTTGGGGTGGCTTTGTCTTCTTCTCTAATTGTTTGGG** | Del_k13_38-50 | -0.33591954520826217 | 0.03207263393724002 | 35.09081760642293 |
| **GATCCAGATCTAACTTGGGGTGGCTTTGTCTTCTTCTCTATTTGCCAATTCCACTAATTGTTTGGG** | Ins_k3_37_CTA | -0.5606741345067512 | 0.04542162976484723 | 28.02746438402409 |
| **GATCCAGATCTAACTTGGGGTGGCTTTGTCTTCTTCTCTCTTTGCCAATTCCACTAATTGTTTGGG** | Ins_k3_37_CTC | -0.983139760275229 | 0.046585857939347206 | 18.37000689422944 |
| **GATCCAGATCTAACTTGGGGTGGCTTTGTCTTCTTCTCTGTTTGCCAATTCCACTAATTGTTTGGG** | Ins_k3_37_CTG | 0.06147321979057829 | 0.03239288297084926 | 52.213039084719604 |
| **GATCCAGATCTAACTTGGGGTGGCTTTGTCTTCTTCTCTTGCCAATTCCACTAATTGTTTGGG** | T38C | -0.11391975097661733 | 0.027449283817753557 | 43.81338147382736 |
| **GATCCAGATCTAACTTGGGGTGGCTTTGTCTTCTTCTCTTTGCCAATTCCACTAATTGTTTGGG** | Ins_k1_37_C | -0.03522636943050139 | 0.034427806456501085 | 47.40049470068939 |
| **GATCCAGATCTAACTTGGGGTGGCTTTGTCTTCTTCTCTTTTGCCAATTCCACTAATTGTTTGGG** | Ins_k2_34_TC;Ins_k2_35_CT;Ins_ | -0.4363677530437078 | 0.0420076616483469 | 31.737256724297875 |
| **GATCCAGATCTAACTTGGGGTGGCTTTGTCTTCTTCTCTTTTTGCCAATTCCACTAATTGTTTGGG** | Ins_k3_36_TCT;Ins_k3_37_CTT | -1.236101546995172 | 0.05690311209326948 | 14.264265416204355 |
| **GATCCAGATCTAACTTGGGGTGGCTTTGTCTTCTTCTG** | Del_k25_38-62 | -1.2937969290931624 | 0.1643930500431931 | 13.464574235941582 |
| **GATCCAGATCTAACTTGGGGTGGCTTTGTCTTCTTCTGAATTTGCCAATTCCACTAATTGTTTGGG** | Ins_k3_37_GAA | 0.4010118240897671 | 0.03183480327165532 | 73.32274485293958 |
| **GATCCAGATCTAACTTGGGGTGGCTTTGTCTTCTTCTGACATTTGCCAATTCCACTAATTGTTTGGG** | Ins_k4_37_GACA | -0.3201246030263328 | 0.038396135212201477 | 35.649475411038 |
| **GATCCAGATCTAACTTGGGGTGGCTTTGTCTTCTTCTGACTTTGCCAATTCCACTAATTGTTTGGG** | Ins_k3_37_GAC | -0.3753317869641691 | 0.03792131913820885 | 33.73470899494474 |
| **GATCCAGATCTAACTTGGGGTGGCTTTGTCTTCTTCTGAGTTTGCCAATTCCACTAATTGTTTGGG** | Ins_k3_37_GAG | 0.46540600185471326 | 0.02934636678765546 | 78.19963942875805 |
| **GATCCAGATCTAACTTGGGGTGGCTTTGTCTTCTTCTGATTTGCCAATTCCACTAATTGTTTGGG** | Ins_k2_37_GA | 0.4871488981697569 | 0.03297613210311228 | 79.91854535974257 |
| **GATCCAGATCTAACTTGGGGTGGCTTTGTCTTCTTCTGATTTTGCCAATTCCACTAATTGTTTGGG** | Ins_k3_36_TGA;Ins_k3_37_GAT | 0.32990070648012515 | 0.03186185833880092 | 68.28975404086908 |
| **GATCCAGATCTAACTTGGGGTGGCTTTGTCTTCTTCTGCATTTGCCAATTCCACTAATTGTTTGGG** | Ins_k3_37_GCA | 0.47200929686096504 | 0.03035701090587239 | 78.71772336530256 |
| **GATCCAGATCTAACTTGGGGTGGCTTTGTCTTCTTCTGCCAATTCCACTAATTGTTTGGG** | Del_k3_37-39;Del_k3_38-40 | 0.3138421986076263 | 0.02746036229306057 | 67.20188067738471 |
| **GATCCAGATCTAACTTGGGGTGGCTTTGTCTTCTTCTGCCTTTGCCAATTCCACTAATTGTTTGGG** | Ins_k3_37_GCC | 0.28231928203419243 | 0.028084061006928143 | 65.11652236840375 |
| **GATCCAGATCTAACTTGGGGTGGCTTTGTCTTCTTCTGCCTTTTGCCAATTCCACTAATTGTTTGGG** | Ins_k4_35_CTGC;Ins_k4_36_TG( | 0.4788358534353374 | 0.03064898847162749 | 79.2569327348827 |
| **GATCCAGATCTAACTTGGGGTGGCTTTGTCTTCTTCTGCGTTTGCCAATTCCACTAATTGTTTGGG** | Ins_k3_37_GCG | 0.6085970628213491 | 0.0265279203417398 | 90.23849389542464 |
| **GATCCAGATCTAACTTGGGGTGGCTTTGTCTTCTTCTGCTCTTTGCCAATTCCACTAATTGTTTGGG** | Ins_k4_37_GCTC | 0.3655880199388375 | 0.03287281724276567 | 70.77084023254297 |
| **GATCCAGATCTAACTTGGGGTGGCTTTGTCTTCTTCTGCTTTGCCAATTCCACTAATTGTTTGGG** | Ins_k2_37_GC | 0.3980680553013167 | 0.029372737817465685 | 73.10721703283585 |
| **GATCCAGATCTAACTTGGGGTGGCTTTGTCTTCTTCTGCTTTTGCCAATTCCACTAATTGTTTGGG** | Ins_k3_35_CTG;Ins_k3_36_TGC; | 0.30792032513952305 | 0.02877277534189397 | 66.80509565809058 |
| **GATCCAGATCTAACTTGGGGTGGCTTTGTCTTCTTCTGG** | Del_k24_38-61 | -1.0900977563238452 | 0.02790981299418935 | 16.506616136014376 |
| **GATCCAGATCTAACTTGGGGTGGCTTTGTCTTCTTCTGGATTTGCCAATTCCACTAATTGTTTGGG** | Ins_k3_37_GGA | 0.7192481683992216 | 0.027820123148167297 | 100 |
| **GATCCAGATCTAACTTGGGGTGGCTTTGTCTTCTTCTGGCTTTGCCAATTCCACTAATTGTTTGGG** | Ins_k3_37_GGC | 0.11546345449014705 | 0.03031726085115372 | 55.10952062198975 |
| **GATCCAGATCTAACTTGGGGTGGCTTTGTCTTCTTCTGGG** | Del_k23_37-59;Del_k23_38-60 | -1.0054082886103917 | 0.031087976790784136 | 17.965454980607273 |
| **GATCCAGATCTAACTTGGGGTGGCTTTGTCTTCTTCTGGGTTTGCCAATTCCACTAATTGTTTGGG** | Ins_k3_37_GGG | -0.026541221432968443 | 0.03466208913221556 | 47.81396795142645 |
| **GATCCAGATCTAACTTGGGGTGGCTTTGTCTTCTTCTGGTTTGCCAATTCCACTAATTGTTTGGG** | Ins_k2_37_GG | 0.6420343092002676 | 0.031313519898122394 | 93.3068331827715 |
| **GATCCAGATCTAACTTGGGGTGGCTTTGTCTTCTTCTGGTTTTGCCAATTCCACTAATTGTTTGGG** | Ins_k3_36_TGG;Ins_k3_37_GGT | 0.528926693591611 | 0.03156575038615558 | 83.32809176468027 |
| **GATCCAGATCTAACTTGGGGTGGCTTTGTCTTCTTCTGTATTTGCCAATTCCACTAATTGTTTGGG** | Ins_k3_37_GTA | 0.05404062942927135 | 0.03593143846659608 | 51.82639960007572 |
| **GATCCAGATCTAACTTGGGGTGGCTTTGTCTTCTTCTGTCTTTGCCAATTCCACTAATTGTTTGGG** | Ins_k3_37_GTC | -0.2877708937105234 | 0.035312844781102616 | 36.8217293024027 |
| **GATCCAGATCTAACTTGGGGTGGCTTTGTCTTCTTCTGTGTTTGCCAATTCCACTAATTGTTTGGG** | Ins_k3_37_GTG | 0.07380810278553063 | 0.03202092084636751 | 52.86106928456025 |
| **GATCCAGATCTAACTTGGGGTGGCTTTGTCTTCTTCTGTTGCCAATTCCACTAATTGTTTGGG** | T38G | 0.06945775173106039 | 0.030916192550184123 | 52.631604563997755 |
| **GATCCAGATCTAACTTGGGGTGGCTTTGTCTTCTTCTGTTTGCCAATTCCACTAATTGTTTGGG** | Ins_k1_37_G | 0.2917117623799236 | 0.03200205774599123 | 65.7310092856487 |
| **GATCCAGATCTAACTTGGGGTGGCTTTGTCTTCTTCTGTTTGGG** | Del_k19_37-55;Del_k19_38-56 | 0.04403649588373715 | 0.028007170020892588 | 51.31050621312075 |
| **GATCCAGATCTAACTTGGGGTGGCTTTGTCTTCTTCTGTTTTGCCAATTCCACTAATTGTTTGGG** | Ins_k2_36_TG;Ins_k2_37_GT | 0.377971495314233 | 0.03466732349016358 | 71.65267802902024 |
| **GATCCAGATCTAACTTGGGGTGGCTTTGTCTTCTTCTGTTTTTGCCAATTCCACTAATTGTTTGGG** | Ins_k3_36_TGT;Ins_k3_37_GTT | 0.08929194497167114 | 0.03588265329593488 | 53.68593127565479 |
| **GATCCAGATCTAACTTGGGGTGGCTTTGTCTTCTTCTT** | Del_k25_39-63 | -6.4801006779756065 | 0.5595860543842022 | 0.07530252266764287 |
| **GATCCAGATCTAACTTGGGGTGGCTTTGTCTTCTTCTTAAATTGCCAATTCCACTAATTGTTTGGG** | Ins_k3_38_AAA | -0.03348108282518769 | 0.03748934377888337 | 47.483294382746266 |
| **GATCCAGATCTAACTTGGGGTGGCTTTGTCTTCTTCTTAAATTTGCCAATTCCACTAATTGTTTGGG** | Ins_k4_37_TAAA;Ins_k4_38_AAA | -0.26539542510346836 | 0.04370271798254272 | 37.654919499106754 |
| **GATCCAGATCTAACTTGGGGTGGCTTTGTCTTCTTCTTAACTTGCCAATTCCACTAATTGTTTGGG** | Ins_k3_38_AAC | -0.1319418775903593 | 0.040294415113162424 | 43.0308438420479 |
| **GATCCAGATCTAACTTGGGGTGGCTTTGTCTTCTTCTTAAGTTGCCAATTCCACTAATTGTTTGGG** | Ins_k3_38_AAG | 0.041205918220736604 | 0.03808488089624243 | 51.165473200794935 |
| **GATCCAGATCTAACTTGGGGTGGCTTTGTCTTCTTCTTAATTCCACTAATTGTTTGGG** | Del_k5_39-43 | -0.36343427709673437 | 0.03662600634902096 | 34.1384651116247 |
| **GATCCAGATCTAACTTGGGGTGGCTTTGTCTTCTTCTTAATTGCCAATTCCACTAATTGTTTGGG** | Ins_k2_38_AA | 0.05362672639615518 | 0.04247467263630466 | 51.80495293481184 |
| **GATCCAGATCTAACTTGGGGTGGCTTTGTCTTCTTCTTAATTGTTTGGG** | Del_k14_38-51;Del_k14_39-52 | -0.4922527965081113 | 0.03245510836336275 | 30.012268206704174 |
| **GATCCAGATCTAACTTGGGGTGGCTTTGTCTTCTTCTTAATTTGCCAATTCCACTAATTGTTTGGG** | Ins_k3_37_TAA;Ins_k3_38_AAT | -0.40848812561331516 | 0.038357904419998395 | 32.634529312052166 |
| **GATCCAGATCTAACTTGGGGTGGCTTTGTCTTCTTCTTACATTGCCAATTCCACTAATTGTTTGGG** | Ins_k3_38_ACA | -0.04941229341640141 | 0.039030792218334893 | 46.73282186090806 |
| **GATCCAGATCTAACTTGGGGTGGCTTTGTCTTCTTCTTACCATTGCCAATTCCACTAATTGTTTGGG** | Ins_k4_38_ACCA | 0.43762928271511226 | 0.03231884981852361 | 76.05739993451877 |
| **GATCCAGATCTAACTTGGGGTGGCTTTGTCTTCTTCTTACCTTGCCAATTCCACTAATTGTTTGGG** | Ins_k3_38_ACC | 0.05242770208065317 | 0.03458438559913202 | 51.74287476063698 |
| **GATCCAGATCTAACTTGGGGTGGCTTTGTCTTCTTCTTACGTTGCCAATTCCACTAATTGTTTGGG** | Ins_k3_38_ACG | 0.4392608162739453 | 0.031297296392265514 | 76.18159141864777 |
| **GATCCAGATCTAACTTGGGGTGGCTTTGTCTTCTTCTTACTAATTGTTTGGG** | Del_k11_39-49 | -0.7333987456832369 | 0.04100831271433056 | 23.58144771514901 |
| **GATCCAGATCTAACTTGGGGTGGCTTTGTCTTCTTCTTACTTGCCAATTCCACTAATTGTTTGGG** | Ins_k2_38_AC | 0.1352556237463689 | 0.03716229832636448 | 56.2111231731834 |
| **GATCCAGATCTAACTTGGGGTGGCTTTGTCTTCTTCTTACTTTGCCAATTCCACTAATTGTTTGGG** | Ins_k3_37_TAC;Ins_k3_38_ACT | -0.46123845721754075 | 0.03766469697663973 | 30.95766350171939 |
| **GATCCAGATCTAACTTGGGGTGGCTTTGTCTTCTTCTTAGATTGCCAATTCCACTAATTGTTTGGG** | Ins_k3_38_AGA | -0.07484189832759225 | 0.035794247362046054 | 45.559407627897194 |
| **GATCCAGATCTAACTTGGGGTGGCTTTGTCTTCTTCTTAGCTTGCCAATTCCACTAATTGTTTGGG** | Ins_k3_38_AGC | -1.5363387935795667 | 0.06736742533867908 | 10.56472098364967 |
| **GATCCAGATCTAACTTGGGGTGGCTTTGTCTTCTTCTTAGGTTGCCAATTCCACTAATTGTTTGGG** | Ins_k3_38_AGG | -3.6701830488043536 | 0.2168244384145595 | 1.2506657205630205 |

| | | | | |
|---|---|---|---|---|
| **GATCCAGATCTAACTTGGGGTGGCTTTGTCTTCTTCTTAGTTGCCAATTCCACTAATTGTTTGGG** | Ins_k2_38_AG | -0.29671988746750344 | 0.04219963503526904 | 36.49368191306326 |
| **GATCCAGATCTAACTTGGGGTGGCTTTGTCTTCTTCTTAGTTTGCCAATTCCACTAATTGTTTGGG** | Ins_k3_37_TAG;Ins_k3_38_AGT | -1.15478294305491 | 0.04718992619073219 | 15.472683150833221 |
| **GATCCAGATCTAACTTGGGGTGGCTTTGTCTTCTTCTTATATTGCCAATTCCACTAATTGTTTGGG** | Ins_k3_38_ATA | -0.24300736054986422 | 0.037501078487566765 | 38.507447889149965 |
| **GATCCAGATCTAACTTGGGGTGGCTTTGTCTTCTTCTTATCTTGCCAATTCCACTAATTGTTTGGG** | Ins_k3_38_ATC | -0.298414348507122 | 0.03990652973167025 | 36.431897151587954 |
| **GATCCAGATCTAACTTGGGGTGGCTTTGTCTTCTTCTTATGCCAATTCCACTAATTGTTTGGG** | T39A | -0.15370876463312538 | 0.03122675589973606 | 42.104316712357836 |
| **GATCCAGATCTAACTTGGGGTGGCTTTGTCTTCTTCTTATGTTGCCAATTCCACTAATTGTTTGGG** | Ins_k3_38_ATG | 0.0908135083923951 | 0.03144154715931114 | 53.767680002055464 |
| **GATCCAGATCTAACTTGGGGTGGCTTTGTCTTCTTCTTATTCCACTAATTGTTTGGG** | Del_k6_39-44 | -0.3916619815127592 | 0.03298927812802971 | 33.188288354715326 |
| **GATCCAGATCTAACTTGGGGTGGCTTTGTCTTCTTCTTATTGCCAATTCCACTAATTGTTTGGG** | Ins_k1_38_A | 0.38367885511002486 | 0.03711564089829264 | 72.06279487179384 |
| **GATCCAGATCTAACTTGGGGTGGCTTTGTCTTCTTCTTATTGTTTGGG** | Del_k15_39-53 | -0.34741506691575214 | 0.031234810474764133 | 34.689740065441164 |
| **GATCCAGATCTAACTTGGGGTGGCTTTGTCTTCTTCTTATTTGCCAATTCCACTAATTGTTTGGG** | Ins_k2_37_TA;Ins_k2_38_AT | -0.016489551713218198 | 0.043042019554297276 | 48.29700174637076 |
| **GATCCAGATCTAACTTGGGGTGGCTTTGTCTTCTTCTTATTTTGCCAATTCCACTAATTGTTTGGG** | Ins_k3_36_TTA;Ins_k3_37_TAT;In | -0.36790686958320995 | 0.041630854674949695 | 33.986118614927456 |
| **GATCCAGATCTAACTTGGGGTGGCTTTGTCTTCTTCTTCAATTCCACTAATTGTTTGGG** | Del_k4_39-42 | 0.4670851887840157 | 0.02842009069663657 | 78.33106155143773 |
| **GATCCAGATCTAACTTGGGGTGGCTTTGTCTTCTTCTTCAATTGCCAATTCCACTAATTGTTTGGG** | Ins_k3_38_CAA | 0.530887713175642 | 0.029238490744871008 | 83.49166011247925 |
| **GATCCAGATCTAACTTGGGGTGGCTTTGTCTTCTTCTTCACCTTGCCAATTCCACTAATTGTTTGGG** | Ins_k4_38_CACC | 0.7167713442722776 | 0.028435282583214577 | 100 |
| **GATCCAGATCTAACTTGGGGTGGCTTTGTCTTCTTCTTCACTAATTGTTTGGG** | Del_k10_39-48 | 0.23937367397989806 | 0.027943503586767976 | 62.379251294825785 |
| **GATCCAGATCTAACTTGGGGTGGCTTTGTCTTCTTCTTCACTTGCCAATTCCACTAATTGTTTGGG** | Ins_k3_38_CAC | 0.43024254408680673 | 0.03005982587986988 | 75.49765369519753 |
| **GATCCAGATCTAACTTGGGGTGGCTTTGTCTTCTTCTTCAGATTGCCAATTCCACTAATTGTTTGGG** | Ins_k4_38_CAGA | 0.5895289174761542 | 0.02972072023060918 | 88.53411449669869 |
| **GATCCAGATCTAACTTGGGGTGGCTTTGTCTTCTTCTTCAGTTGCCAATTCCACTAATTGTTTGGG** | Ins_k3_38_CAG | 0.4579136075359269 | 0.029242416451599605 | 77.61592632953109 |
| **GATCCAGATCTAACTTGGGGTGGCTTTGTCTTCTTCTTCATTGCCAATTCCACTAATTGTTTGGG** | Ins_k2_38_CA | 0.4535005913774617 | 0.03250544736165004 | 77.27416065585362 |
| **GATCCAGATCTAACTTGGGGTGGCTTTGTCTTCTTCTTCATTTGCCAATTCCACTAATTGTTTGGG** | Ins_k3_37_TCA;Ins_k3_38_CAT | 0.5820443242866953 | 0.030060196389540803 | 87.87394629335601 |
| **GATCCAGATCTAACTTGGGGTGGCTTTGTCTTCTTCTTCCAATTCCACTAATTGTTTGGG** | Del_k3_39-41 | 0.22077016609618827 | 0.0287612092914189 | 61.22950620694949 |
| **GATCCAGATCTAACTTGGGGTGGCTTTGTCTTCTTCTTCCACTAATTGTTTGGG** | Del_k9_37-45;Del_k9_38-46;Del_ | 0.14657856561045124 | 0.029193410519711745 | 56.85121547818112 |
| **GATCCAGATCTAACTTGGGGTGGCTTTGTCTTCTTCTTCCATTGCCAATTCCACTAATTGTTTGGG** | Ins_k3_38_CCA | 0.3696479374793029 | 0.03106248564698268 | 71.05874805369952 |
| **GATCCAGATCTAACTTGGGGTGGCTTTGTCTTCTTCTTCCCTTGCCAATTCCACTAATTGTTTGGG** | Ins_k3_38_CCC | 0.06415060414611629 | 0.03321750608954023 | 52.35302076748364 |
| **GATCCAGATCTAACTTGGGGTGGCTTTGTCTTCTTCTTCCGTTGCCAATTCCACTAATTGTTTGGG** | Ins_k3_38_CCG | 0.5482398419766508 | 0.029416265258252108 | 84.95306068973368 |
| **GATCCAGATCTAACTTGGGGTGGCTTTGTCTTCTTCTTCCTTGCCAATTCCACTAATTGTTTGGG** | Ins_k2_38_CC | 0.4728125236416487 | 0.03789857681747968 | 78.78097694890938 |
| **GATCCAGATCTAACTTGGGGTGGCTTTGTCTTCTTCTTCCTTTGCCAATTCCACTAATTGTTTGGG** | Ins_k3_37_TCC;Ins_k3_38_CCT | 0.02453285912763359 | 0.032420253209993755 | 50.31946064977817 |
| **GATCCAGATCTAACTTGGGGTGGCTTTGTCTTCTTCTTCGAGTTGCCAATTCCACTAATTGTTTGGG** | Ins_k4_38_CGAG | 0.8364406891839198 | 0.02999836804508465 | 100 |
| **GATCCAGATCTAACTTGGGGTGGCTTTGTCTTCTTCTTCGATTGCCAATTCCACTAATTGTTTGGG** | Ins_k3_38_CGA | 0.5437848004215586 | 0.02737831419958628 | 84.57543307207001 |
| **GATCCAGATCTAACTTGGGGTGGCTTTGTCTTCTTCTTCGCTTGCCAATTCCACTAATTGTTTGGG** | Ins_k3_38_CGC | 0.6268267846905375 | 0.02851812970652653 | 91.89860222351923 |
| **GATCCAGATCTAACTTGGGGTGGCTTTGTCTTCTTCTTCGGTTGCCAATTCCACTAATTGTTTGGG** | Ins_k3_38_CGG | 0.4574204750820601 | 0.02843255989711457 | 77.5776608330756 |
| **GATCCAGATCTAACTTGGGGTGGCTTTGTCTTCTTCTTCGTTGCCAATTCCACTAATTGTTTGGG** | Ins_k2_38_CG | 0.7542645068656086 | 0.030483927233198198 | 100 |
| **GATCCAGATCTAACTTGGGGTGGCTTTGTCTTCTTCTTCGTTTGCCAATTCCACTAATTGTTTGGG** | Ins_k3_37_TCG;Ins_k3_38_CGT | 0.5078622620798257 | 0.03009296919711913 | 81.591190524242 |
| **GATCCAGATCTAACTTGGGGTGGCTTTGTCTTCTTCTTCTAATTGTTTGGG** | Del_k12_39-50 | 0.18084850494681526 | 0.030510852727390483 | 58.83327167850616 |
| **GATCCAGATCTAACTTGGGGTGGCTTTGTCTTCTTCTTCTATTGCCAATTCCACTAATTGTTTGGG** | Ins_k3_38_CTA | -0.595383229921266 | 0.22282708978247745 | 27.071345446353593 |
| **GATCCAGATCTAACTTGGGGTGGCTTTGTCTTCTTCTTCTCTTGCCAATTCCACTAATTGTTTGGG** | Ins_k3_38_CTC | -0.08608474061333504 | 0.03498618388871241 | 45.05005902152099 |
| **GATCCAGATCTAACTTGGGGTGGCTTTGTCTTCTTCTTCTGATTGCCAATTCCACTAATTGTTTGGG** | Ins_k4_38_CTGA | 0.7181251666138424 | 0.03090878031562588 | 100 |
| **GATCCAGATCTAACTTGGGGTGGCTTTGTCTTCTTCTTCTGCCAATTCCACTAATTGTTTGGG** | T39C | 0.08130072535851263 | 0.025581815044779917 | 53.258624832474354 |
| **GATCCAGATCTAACTTGGGGTGGCTTTGTCTTCTTCTTCTGTTGCCAATTCCACTAATTGTTTGGG** | Ins_k3_38_CTG | 0.4198874627107918 | 0.030632028081702876 | 74.71990313235837 |
| **GATCCAGATCTAACTTGGGGTGGCTTTGTCTTCTTCTTCTTGCCAATTCCACTAATTGTTTGGG** | Ins_k1_38_C | 0.4653842872580687 | 0.03292125182249161 | 78.19794137356645 |
| **GATCCAGATCTAACTTGGGGTGGCTTTGTCTTCTTCTTCTTTGCCAATTCCACTAATTGTTTGGG** | Ins_k2_37_TC;Ins_k2_38_CT | 0.26267142642635843 | 0.034342158777599864 | 63.849609159274216 |
| **GATCCAGATCTAACTTGGGGTGGCTTTGTCTTCTTCTTCTTTTGCCAATTCCACTAATTGTTTGGG** | Ins_k3_28_TCT;Ins_k3_29_CTT;Ir | -0.5740141151067709 | 0.029726354033815607 | 27.656061315477043 |
| **GATCCAGATCTAACTTGGGGTGGCTTTGTCTTCTTCTTG** | Del_k24_39-62 | -1.0843323615394533 | 0.032842008356597376 | 16.60205816069608 |
| **GATCCAGATCTAACTTGGGGTGGCTTTGTCTTCTTCTTGAATTGCCAATTCCACTAATTGTTTGGG** | Ins_k3_38_GAA | 0.608982833028521 | 0.029213257683382517 | 90.27331193335861 |
| **GATCCAGATCTAACTTGGGGTGGCTTTGTCTTCTTCTTGACTTGCCAATTCCACTAATTGTTTGGG** | Ins_k3_38_GAC | 0.2808883366863477 | 0.033107443111178046 | 65.02341081831054 |
| **GATCCAGATCTAACTTGGGGTGGCTTTGTCTTCTTCTTGAGTTGCCAATTCCACTAATTGTTTGGG** | Ins_k3_38_GAG | 0.4462077315873918 | 0.030388754166494086 | 76.7126609945947 |
| **GATCCAGATCTAACTTGGGGTGGCTTTGTCTTCTTCTTGATTGCCAATTCCACTAATTGTTTGGG** | Ins_k2_38_GA | 0.4556541521814632 | 0.03327006627560935 | 77.44075458009078 |
| **GATCCAGATCTAACTTGGGGTGGCTTTGTCTTCTTCTTGATTTGCCAATTCCACTAATTGTTTGGG** | Ins_k3_37_TGA;Ins_k3_38_GAT | 0.3229383657302677 | 0.031734590147194264 | 67.81594881417337 |
| **GATCCAGATCTAACTTGGGGTGGCTTTGTCTTCTTCTTGCATTGCCAATTCCACTAATTGTTTGGG** | Ins_k3_38_GCA | 0.10513748660626498 | 0.03264087434752612 | 54.54338943849181 |
| **GATCCAGATCTAACTTGGGGTGGCTTTGTCTTCTTCTTGCCAATTCCACTAATTGTTTGGG** | Del_k2_37-38;Del_k2_38-39;Del_ | 0.27173114280117183 | 0.030094821288256572 | 64.43069677885174 |
| **GATCCAGATCTAACTTGGGGTGGCTTTGTCTTCTTCTTGCCTTGCCAATTCCACTAATTGTTTGGG** | Ins_k3_38_GCC | 0.25400128888990126 | 0.0322727985539569 | 63.29841717418727 |
| **GATCCAGATCTAACTTGGGGTGGCTTTGTCTTCTTCTTGCGTTGCCAATTCCACTAATTGTTTGGG** | Ins_k3_38_GCG | 0.3481710867405863 | 0.030799242036139256 | 69.548901339263125 |
| **GATCCAGATCTAACTTGGGGTGGCTTTGTCTTCTTCTTGCTCTTGCCAATTCCACTAATTGTTTGGG** | Ins_k4_38_GCTC | -0.003973578373008602 | 0.03644456739420053 | 48.905284416902454 |
| **GATCCAGATCTAACTTGGGGTGGCTTTGTCTTCTTCTTGCTTGCCAATTCCACTAATTGTTTGGG** | Ins_k2_38_GC | 0.45567267854477134 | 0.03575056846375217 | 77.44218928893491 |
| **GATCCAGATCTAACTTGGGGTGGCTTTGTCTTCTTCTTGCTTTGCCAATTCCACTAATTGTTTGGG** | Ins_k3_37_TGC;Ins_k3_38_GCT | -0.07155165020487719 | 0.03393608991873515 | 45.70955626099025 |
| **GATCCAGATCTAACTTGGGGTGGCTTTGTCTTCTTCTTGG** | Del_k23_39-61 | -0.898305409912461 | 0.028077595456575757 | 19.996427345814915 |
| **GATCCAGATCTAACTTGGGGTGGCTTTGTCTTCTTCTTGGAATTGCCAATTCCACTAATTGTTTGGG** | Ins_k4_38_GGAA | 0.6244955739387024 | 0.0299302059771239 | 91.68461673348415 |
| **GATCCAGATCTAACTTGGGGTGGCTTTGTCTTCTTCTTGGATTGCCAATTCCACTAATTGTTTGGG** | Ins_k3_38_GGA | 0.6748427133384036 | 0.027723544007292465 | 96.41885252797037 |
| **GATCCAGATCTAACTTGGGGTGGCTTTGTCTTCTTCTTGGCTTGCCAATTCCACTAATTGTTTGGG** | Ins_k3_38_GGC | 0.36715175354391805 | 0.028827240345031237 | 70.88159354544459 |
| **GATCCAGATCTAACTTGGGGTGGCTTTGTCTTCTTCTTGGG** | Del_k22_37-58;Del_k22_38-59;D | -0.9263130216966995 | 0.031369426598386776 | 19.44414532345456 |
| **GATCCAGATCTAACTTGGGGTGGCTTTGTCTTCTTCTTGGGTTGCCAATTCCACTAATTGTTTGGG** | Ins_k3_38_GGG | 0.008752508965301886 | 0.03708794910083845 | 49.531634376578424 |
| **GATCCAGATCTAACTTGGGGTGGCTTTGTCTTCTTCTTGGTTGCCAATTCCACTAATTGTTTGGG** | Ins_k2_38_GG | 0.6235665010199654 | 0.0319130957969022 | 91.5994745967634 |
| **GATCCAGATCTAACTTGGGGTGGCTTTGTCTTCTTCTTGGTTTGCCAATTCCACTAATTGTTTGGG** | Ins_k3_37_TGG;Ins_k3_38_GGT | 0.3992322761109375 | 0.028916839055343084 | 73.19237954060164 |
| **GATCCAGATCTAACTTGGGGTGGCTTTGTCTTCTTCTTGTAATTGCCAATTCCACTAATTGTTTGGG** | Ins_k4_38_GTAA | -0.2220589966914424 | 0.03957319068323254 | 39.322624415657714 |
| **GATCCAGATCTAACTTGGGGTGGCTTTGTCTTCTTCTTGTATTGCCAATTCCACTAATTGTTTGGG** | Ins_k3_38_GTA | 0.050360567345388185 | 0.03403604295521875 | 51.636025740624724 |
| **GATCCAGATCTAACTTGGGGTGGCTTTGTCTTCTTCTTGTCTTGCCAATTCCACTAATTGTTTGGG** | Ins_k3_38_GTC | -0.011827892305374843 | 0.03241635540346867 | 48.522671508045875 |
| **GATCCAGATCTAACTTGGGGTGGCTTTGTCTTCTTCTTGTGCCAATTCCACTAATTGTTTGGG** | T39G | 0.20658256061467717 | 0.02931143647111763 | 60.36693947595451 |
| **GATCCAGATCTAACTTGGGGTGGCTTTGTCTTCTTCTTGTGTTGCCAATTCCACTAATTGTTTGGG** | Ins_k3_38_GTG | 0.19845902203383992 | 0.03120007581054174 | 59.87853279527461 |
| **GATCCAGATCTAACTTGGGGTGGCTTTGTCTTCTTCTTGTTGCCAATTCCACTAATTGTTTGGG** | Ins_k1_38_G | 0.3191353095797884 | 0.03582773029705046 | 67.5585307508676 |
| **GATCCAGATCTAACTTGGGGTGGCTTTGTCTTCTTCTTGTTTGCCAATTCCACTAATTGTTTGGG** | Ins_k2_37_TG;Ins_k2_38_GT | 0.2257138430135215 | 0.032863355304674116 | 61.53295456073419 |
| **GATCCAGATCTAACTTGGGGTGGCTTTGTCTTCTTCTTGTTTGGG** | Del_k18_37-54;Del_k18_38-55;D | -0.31192730683716596 | 0.0318254071980488 | 35.942905744545996 |
| **GATCCAGATCTAACTTGGGGTGGCTTTGTCTTCTTCTTGTTTTGCCAATTCCACTAATTGTTTGGG** | Ins_k3_36_TTG;Ins_k3_37_TGT;I | -0.11354153539301398 | 0.03453775027014041 | 43.82995551155323 |
| **GATCCAGATCTAACTTGGGGTGGCTTTGTCTTCTTCTTT** | Del_k24_40-63 | -4.820753213950046 | 0.18557844906622684 | 0.3957810284403398 |
| **GATCCAGATCTAACTTGGGGTGGCTTTGTCTTCTTCTTTAAATGCCAATTCCACTAATTGTTTGGG** | Ins_k3_39_AAA | -0.3740309798203866 | 0.0446494947722321 | 33.778619899015844 |
| **GATCCAGATCTAACTTGGGGTGGCTTTGTCTTCTTCTTTAACTGCCAATTCCACTAATTGTTTGGG** | Ins_k3_39_AAC | 0.10490518313522046 | 0.03544523325113776 | 54.5307202914031 |

| | | | | |
|---|---|---|---|---|
| **GATCCAGATCTAACTTGGGGTGGCTTTGTCTTCTTCTTTAAGTGCCAATTCCACTAATTGTTTGGG** | Ins_k3_39_AAG | 0.1274270322523019 | 0.03502525182520146 | 55.772787267492376 |
| **GATCCAGATCTAACTTGGGGTGGCTTTGTCTTCTTCTTTAATGCCAATTCCACTAATTGTTTGGG** | Ins_k2_39_AA | -0.19202356667611342 | 0.04354257420470352 | 40.5216122719769 |
| **GATCCAGATCTAACTTGGGGTGGCTTTGTCTTCTTCTTTAATTCCACTAATTGTTTGGG** | Del_k4_40-43 | -0.8137774008177646 | 0.04760327827403286 | 21.760178693744226 |
| **GATCCAGATCTAACTTGGGGTGGCTTTGTCTTCTTCTTTAATTGCCAATTCCACTAATTGTTTGGG** | Ins_k3_38_TAA;Ins_k3_39_AAT | -0.5084728828058918 | 0.05197428626428459 | 29.52939334896181 |
| **GATCCAGATCTAACTTGGGGTGGCTTTGTCTTCTTCTTTAATTGTTTGGG** | Del_k13_39-51;Del_k13_40-52 | -0.7548804749055247 | 0.04302422043682433 | 23.080279691021268 |
| **GATCCAGATCTAACTTGGGGTGGCTTTGTCTTCTTCTTTACATGCCAATTCCACTAATTGTTTGGG** | Ins_k3_39_ACA | -0.27725050105077836 | 0.0429999044914976 | 37.21115320753493 |
| **GATCCAGATCTAACTTGGGGTGGCTTTGTCTTCTTCTTTACCTGCCAATTCCACTAATTGTTTGGG** | Ins_k3_39_ACC | 0.3018224716714327 | 0.031532262560676365 | 66.3989674873319 |
| **GATCCAGATCTAACTTGGGGTGGCTTTGTCTTCTTCTTTACGTGCCAATTCCACTAATTGTTTGGG** | Ins_k3_39_ACG | 0.396201585364629 | 0.030592876215107614 | 72.97089187307175 |
| **GATCCAGATCTAACTTGGGGTGGCTTTGTCTTCTTCTTTACTAATTGTTTGGG** | Del_k10_40-49 | -1.5336947460461867 | 0.06665530206865176 | 10.592691569591555 |
| **GATCCAGATCTAACTTGGGGTGGCTTTGTCTTCTTCTTTACTGCCAATTCCACTAATTGTTTGGG** | Ins_k2_39_AC | 0.4325855634225184 | 0.03748377333611412 | 75.6747535507799 |
| **GATCCAGATCTAACTTGGGGTGGCTTTGTCTTCTTCTTTACTTGCCAATTCCACTAATTGTTTGGG** | Ins_k3_38_TAC;Ins_k3_39_ACT | -0.39652389652014736 | 0.04450205731164656 | 33.02732133855079 |
| **GATCCAGATCTAACTTGGGGTGGCTTTGTCTTCTTCTTTAGATGCCAATTCCACTAATTGTTTGGG** | Ins_k3_39_AGA | 0.14525146045015247 | 0.03452898354163234 | 56.775817978022424 |
| **GATCCAGATCTAACTTGGGGTGGCTTTGTCTTCTTCTTTAGCCAATTCCACTAATTGTTTGGG** | T40A | -1.3491094758837514 | 0.05470909800007062 | 12.740037057818173 |
| **GATCCAGATCTAACTTGGGGTGGCTTTGTCTTCTTCTTTAGCTGCCAATTCCACTAATTGTTTGGG** | Ins_k3_39_AGC | -0.3982863109859878 | 0.046873197172721974 | 32.9691647726956 |
| **GATCCAGATCTAACTTGGGGTGGCTTTGTCTTCTTCTTTAGGTGCCAATTCCACTAATTGTTTGGG** | Ins_k3_39_AGG | -3.536762158241382 | 0.16334805028949181 | 1.4291743166377842 |
| **GATCCAGATCTAACTTGGGGTGGCTTTGTCTTCTTCTTTAGTGCCAATTCCACTAATTGTTTGGG** | Ins_k2_39_AG | -0.33763841434312436 | 0.04879079987262242 | 35.03055289153907 |
| **GATCCAGATCTAACTTGGGGTGGCTTTGTCTTCTTCTTTAGTTGCCAATTCCACTAATTGTTTGGG** | Ins_k3_38_TAG;Ins_k3_39_AGT | -1.4982959099961668 | 0.059460827992975196 | 10.974376262852683 |
| **GATCCAGATCTAACTTGGGGTGGCTTTGTCTTCTTCTTTATATGCCAATTCCACTAATTGTTTGGG** | Ins_k3_39_ATA | -0.42127584574672194 | 0.048270640795998784 | 32.219865040527324 |
| **GATCCAGATCTAACTTGGGGTGGCTTTGTCTTCTTCTTTATCTGCCAATTCCACTAATTGTTTGGG** | Ins_k3_39_ATC | 0.08311428821771538 | 0.03504733245274704 | 53.3553003334545 |
| **GATCCAGATCTAACTTGGGGTGGCTTTGTCTTCTTCTTTATGCCAATTCCACTAATTGTTTGGG** | Ins_k1_39_A | -0.06959543331955587 | 0.04051748554039595 | 45.79906158413765 |
| **GATCCAGATCTAACTTGGGGTGGCTTTGTCTTCTTCTTTATGTGCCAATTCCACTAATTGTTTGGG** | Ins_k3_39_ATG | 0.1824656240483038 | 0.040290081375163204 | 58.92848905410894 |
| **GATCCAGATCTAACTTGGGGTGGCTTTGTCTTCTTCTTTATTCCACTAATTGTTTGGG** | Del_k5_40-44 | -0.8203291010499225 | 0.040643693795933786 | 21.618078533073554 |
| **GATCCAGATCTAACTTGGGGTGGCTTTGTCTTCTTCTTTATTGCCAATTCCACTAATTGTTTGGG** | Ins_k2_38_TA;Ins_k2_39_AT | -0.2232785330877124 | 0.041532202876791435 | 39.27469827376307 |
| **GATCCAGATCTAACTTGGGGTGGCTTTGTCTTCTTCTTTATTGTTTGGG** | Del_k14_40-53 | -0.4650338163257126 | 0.0330997539615237 | 30.84039073847918 |
| **GATCCAGATCTAACTTGGGGTGGCTTTGTCTTCTTCTTTATTTGCCAATTCCACTAATTGTTTGGG** | Ins_k3_37_TTA;Ins_k3_38_TAT;In | -0.6426722387677728 | 0.049883345790971076 | 25.82096596012726 |
| **GATCCAGATCTAACTTGGGGTGGCTTTGTCTTCTTCTTTCAATGCCAATTCCACTAATTGTTTGGG** | Ins_k3_39_CAA | 0.2592254320068157 | 0.03244868888387248 | 63.629962430478685 |
| **GATCCAGATCTAACTTGGGGTGGCTTTGTCTTCTTCTTTCAATTCCACTAATTGTTTGGG** | Del_k3_40-42 | -0.015915827196822274 | 0.038510854276059624 | 48.32471887057763 |
| **GATCCAGATCTAACTTGGGGTGGCTTTGTCTTCTTCTTTCACTAATTGTTTGGG** | Del_k9_40-48 | -0.804876029201089 | 0.04484264611315861 | 21.95473877186826 |
| **GATCCAGATCTAACTTGGGGTGGCTTTGTCTTCTTCTTTCACTGCCAATTCCACTAATTGTTTGGG** | Ins_k3_39_CAC | 0.357131269747057 | 0.03257501917459431 | 70.17487244298937 |
| **GATCCAGATCTAACTTGGGGTGGCTTTGTCTTCTTCTTTCAGTGCCAATTCCACTAATTGTTTGGG** | Ins_k3_39_CAG | -0.4051034498489255 | 0.03963621251063828 | 32.74517375464978 |
| **GATCCAGATCTAACTTGGGGTGGCTTTGTCTTCTTCTTTCATGCCAATTCCACTAATTGTTTGGG** | Ins_k2_39_CA | 0.10421312636174584 | 0.0329561971516318 | 54.49299499259296 |
| **GATCCAGATCTAACTTGGGGTGGCTTTGTCTTCTTCTTTCATTGCCAATTCCACTAATTGTTTGGG** | Ins_k3_38_TCA;Ins_k3_39_CAT | -0.17199485912774065 | 0.03600040713490051 | 41.34138993283172 |
| **GATCCAGATCTAACTTGGGGTGGCTTTGTCTTCTTCTTTCCAATTCCACTAATTGTTTGGG** | Del_k2_40-41 | -0.46641083845351367 | 0.03666050518718306 | 30.797952064205017 |
| **GATCCAGATCTAACTTGGGGTGGCTTTGTCTTCTTCTTTCCACTAATTGTTTGGG** | Del_k8_38-45;Del_k8_39-46;Del_ | -0.33140003589324846 | 0.03730867947261606 | 35.24976980587707 |
| **GATCCAGATCTAACTTGGGGTGGCTTTGTCTTCTTCTTTCCAGTGCCAATTCCACTAATTGTTTGGG** | Ins_k4_39_CCAG | 0.2480276752458367 | 0.03488470521321306 | 62.92142401250824 |
| **GATCCAGATCTAACTTGGGGTGGCTTTGTCTTCTTCTTTCCATGCCAATTCCACTAATTGTTTGGG** | Ins_k3_39_CCA | 0.1670822050077736 | 0.034408824807256544 | 58.0289044971213 |
| **GATCCAGATCTAACTTGGGGTGGCTTTGTCTTCTTCTTTCCCTGCCAATTCCACTAATTGTTTGGG** | Ins_k3_39_CCC | 0.2782369112472314 | 0.03748773667984572 | 64.85123444999769 |
| **GATCCAGATCTAACTTGGGGTGGCTTTGTCTTCTTCTTTCCGTGCCAATTCCACTAATTGTTTGGG** | Ins_k3_39_CCG | 0.6628579093051283 | 0.030308811488002702 | 95.27018848110787 |
| **GATCCAGATCTAACTTGGGGTGGCTTTGTCTTCTTCTTTCCTGCCAATTCCACTAATTGTTTGGG** | Ins_k2_39_CC | 0.7024331751065942 | 0.03136321521785358 | 99.11613167822817 |
| **GATCCAGATCTAACTTGGGGTGGCTTTGTCTTCTTCTTTCCTTGCCAATTCCACTAATTGTTTGGG** | Ins_k3_38_TCC;Ins_k3_39_CCT | -0.4726265129941551 | 0.04197275589027385 | 30.60711572036448 |
| **GATCCAGATCTAACTTGGGGTGGCTTTGTCTTCTTCTTTCGAATGCCAATTCCACTAATTGTTTGGG** | Ins_k4_39_CGAA | 0.7341238363100302 | 0.0297582328442884 | 100 |
| **GATCCAGATCTAACTTGGGGTGGCTTTGTCTTCTTCTTTCGATGCCAATTCCACTAATTGTTTGGG** | Ins_k3_39_CGA | 0.6101508331354941 | 0.029358275753614025 | 90.37881277185502 |
| **GATCCAGATCTAACTTGGGGTGGCTTTGTCTTCTTCTTTCGATTGCCAATTCCACTAATTGTTTGGG** | Ins_k4_38_TCGA;Ins_k4_39_CG/ | 0.46091327381414793 | 0.031494434585848066 | 77.84909774977821 |
| **GATCCAGATCTAACTTGGGGTGGCTTTGTCTTCTTCTTTCGCCAATTCCACTAATTGTTTGGG** | T40C | 0.11416791725017283 | 0.025183177836480317 | 55.03817041415212 |
| **GATCCAGATCTAACTTGGGGTGGCTTTGTCTTCTTCTTTCGCTGCCAATTCCACTAATTGTTTGGG** | Ins_k3_39_CGC | 0.5598673473042672 | 0.03245738021684373 | 85.94661795775387 |
| **GATCCAGATCTAACTTGGGGTGGCTTTGTCTTCTTCTTTCGGTGCCAATTCCACTAATTGTTTGGG** | Ins_k3_39_CGG | 0.789256148161176 | 0.031288605700980514 | 100 |
| **GATCCAGATCTAACTTGGGGTGGCTTTGTCTTCTTCTTTCGTGCCAATTCCACTAATTGTTTGGG** | Ins_k2_39_CG | 0.6323875741297451 | 0.029843051558085504 | 92.41105449932378 |
| **GATCCAGATCTAACTTGGGGTGGCTTTGTCTTCTTCTTTCGTTGCCAATTCCACTAATTGTTTGGG** | Ins_k3_38_TCG;Ins_k3_39_CGT | 0.21264790819151036 | 0.03342257431699546 | 60.73419859722962 |
| **GATCCAGATCTAACTTGGGGTGGCTTTGTCTTCTTCTTTCTAATTGTTTGGG** | Del_k11_40-50 | -1.6165315456011162 | 0.062150247508653354 | 9.750587008518636 |
| **GATCCAGATCTAACTTGGGGTGGCTTTGTCTTCTTCTTTCTATGCCAATTCCACTAATTGTTTGGG** | Ins_k3_39_CTA | -0.9687235300912759 | 0.05171846521212729 | 18.636751246365186 |
| **GATCCAGATCTAACTTGGGGTGGCTTTGTCTTCTTCTTTCTCTGCCAATTCCACTAATTGTTTGGG** | Ins_k3_39_CTC | -0.21455056572435993 | 0.04015646969876308 | 39.618986842616614 |
| **GATCCAGATCTAACTTGGGGTGGCTTTGTCTTCTTCTTTCTGCCAATTCCACTAATTGTTTGGG** | Ins_k1_39_C | 0.3708860879066477 | 0.03341626217402127 | 71.14678396257118 |
| **GATCCAGATCTAACTTGGGGTGGCTTTGTCTTCTTCTTTCTGTGCCAATTCCACTAATTGTTTGGG** | Ins_k3_39_CTG | -0.018311291145359232 | 0.03517669273816413 | 48.209097287655524 |
| **GATCCAGATCTAACTTGGGGTGGCTTTGTCTTCTTCTTTCTTGCCAATTCCACTAATTGTTTGGG** | Ins_k2_38_TC;Ins_k2_39_CT | -0.4688986843388507 | 0.0456846528543085 | 30.721426736965867 |
| **GATCCAGATCTAACTTGGGGTGGCTTTGTCTTCTTCTTTCTTTGCCAATTCCACTAATTGTTTGGG** | Ins_k3_37_TTC;Ins_k3_38_TCT;In | -1.3120901536260159 | 0.06603670568458472 | 13.220502987615609 |
| **GATCCAGATCTAACTTGGGGTGGCTTTGTCTTCTTCTTTG** | Del_k23_40-62 | -0.775770874058377 | 0.027916027922713783 | 22.603124766884694 |
| **GATCCAGATCTAACTTGGGGTGGCTTTGTCTTCTTCTTTGAATGCCAATTCCACTAATTGTTTGGG** | Ins_k3_39_GAA | 0.5174663958349313 | 0.0287378930576948 | 82.37857826753952 |
| **GATCCAGATCTAACTTGGGGTGGCTTTGTCTTCTTCTTTGACCTGCCAATTCCACTAATTGTTTGGG** | Ins_k4_39_GACC | 0.7700745867375716 | 0.030469940021073644 | 100 |
| **GATCCAGATCTAACTTGGGGTGGCTTTGTCTTCTTCTTTGACTGCCAATTCCACTAATTGTTTGGG** | Ins_k3_39_GAC | 0.6285113051761093 | 0.031755476047500966 | 92.05353776096634 |
| **GATCCAGATCTAACTTGGGGTGGCTTTGTCTTCTTCTTTGAGTGCCAATTCCACTAATTGTTTGGG** | Ins_k3_39_GAG | -0.21280364886568487 | 0.34225261612404934 | 39.68825840687088 |
| **GATCCAGATCTAACTTGGGGTGGCTTTGTCTTCTTCTTTGATGCCAATTCCACTAATTGTTTGGG** | Ins_k2_39_GA | 0.960997310603452 | 0.030910566677929536 | 100 |
| **GATCCAGATCTAACTTGGGGTGGCTTTGTCTTCTTCTTTGATTGCCAATTCCACTAATTGTTTGGG** | Ins_k3_38_TGA;Ins_k3_39_GAT | 0.3671373170546156 | 0.030430969144288656 | 70.8805702714639 |
| **GATCCAGATCTAACTTGGGGTGGCTTTGTCTTCTTCTTTGCATGCCAATTCCACTAATTGTTTGGG** | Ins_k3_39_GCA | -0.4060762398845036 | 0.04220444143722088 | 32.713335064598994 |
| **GATCCAGATCTAACTTGGGGTGGCTTTGTCTTCTTCTTTGCCAATTCCACTAATTGTTTGGG** | Del_k1_37-37;Del_k1_38-38;Del_ | 0.04936523867321618 | 0.028322187810769426 | 51.58465649257165 |
| **GATCCAGATCTAACTTGGGGTGGCTTTGTCTTCTTCTTTGCCTGCCAATTCCACTAATTGTTTGGG** | Ins_k3_39_GCC | 0.5670097656938156 | 0.030171443430420133 | 86.5626821374414 |
| **GATCCAGATCTAACTTGGGGTGGCTTTGTCTTCTTCTTTGCGTGCCAATTCCACTAATTGTTTGGG** | Ins_k3_39_GCG | 0.5637961746541391 | 0.028454857734041997 | 86.28495157299496 |
| **GATCCAGATCTAACTTGGGGTGGCTTTGTCTTCTTCTTTGCTGCCAATTCCACTAATTGTTTGGG** | Ins_k2_39_GC | 0.739028908663524 | 0.03250184451456147 | 100 |
| **GATCCAGATCTAACTTGGGGTGGCTTTGTCTTCTTCTTTGCTTGCCAATTCCACTAATTGTTTGGG** | Ins_k3_38_TGC;Ins_k3_39_GCT | -0.13465225808624254 | 0.038940211892108 | 42.914371795260855 |
| **GATCCAGATCTAACTTGGGGTGGCTTTGTCTTCTTCTTTGG** | Del_k22_40-61 | -0.6555831488774345 | 0.026842257219061125 | 25.489736626828215 |
| **GATCCAGATCTAACTTGGGGTGGCTTTGTCTTCTTCTTTGGATGCCAATTCCACTAATTGTTTGGG** | Ins_k3_39_GGA | 0.5471659649367222 | 0.03060580653115485 | 84.86188051529774 |
| **GATCCAGATCTAACTTGGGGTGGCTTTGTCTTCTTCTTTGGCCAATTCCACTAATTGTTTGGG** | T40G | 0.11169444616181523 | 0.029329975075864025 | 54.902203315541534 |
| **GATCCAGATCTAACTTGGGGTGGCTTTGTCTTCTTCTTTGGCGTGCCAATTCCACTAATTGTTTGGG** | Ins_k4_39_GGCG | 0.7497297910571248 | 0.028692101785183183 | 100 |
| **GATCCAGATCTAACTTGGGGTGGCTTTGTCTTCTTCTTTGGCTGCCAATTCCACTAATTGTTTGGG** | Ins_k3_39_GGC | 0.5320832233488735 | 0.030628699585311683 | 83.59153493030861 |
| **GATCCAGATCTAACTTGGGGTGGCTTTGTCTTCTTCTTTGGG** | Del_k21_37-57;Del_k21_38-58;D | -1.058655366282153 | 0.030656028727211692 | 17.033869210033654 |
| **GATCCAGATCTAACTTGGGGTGGCTTTGTCTTCTTCTTTGGGGTGCCAATTCCACTAATTGTTTGGG** | Ins_k4_39_GGGG | 0.10684209612742179 | 0.04018069719893354 | 54.636443907679585 |

| | | | | |
|---|---|---|---|---|
| **GATCCAGATCTAACTTGGGGTGGCTTTGTCTTCTTCTTTGGGTGCCAATTCCACTAATTGTTTGGG** | Ins_k3_39_GGG | 0.23136769607984753 | 0.03275739289100406 | 61.88183818355893 |
| **GATCCAGATCTAACTTGGGGTGGCTTTGTCTTCTTCTTTGGTGCCAATTCCACTAATTGTTTGGG** | Ins_k2_39_GG | 0.7562523069862153 | 0.033088240314656674 | 100 |
| **GATCCAGATCTAACTTGGGGTGGCTTTGTCTTCTTCTTTGGTTGCCAATTCCACTAATTGTTTGGG** | Ins_k3_38_TGG;Ins_k3_39_GGT | 0.24452898416085367 | 0.03220885762288283 | 62.70166604405224 |
| **GATCCAGATCTAACTTGGGGTGGCTTTGTCTTCTTCTTTGTATGCCAATTCCACTAATTGTTTGGG** | Ins_k3_39_GTA | -0.22322408898738083 | 0.040217147299256094 | 39.27683660758568 |
| **GATCCAGATCTAACTTGGGGTGGCTTTGTCTTCTTCTTTGTCTGCCAATTCCACTAATTGTTTGGG** | Ins_k3_39_GTC | 0.1687974511995609 | 0.033915166723733764 | 58.12852376593542 |
| **GATCCAGATCTAACTTGGGGTGGCTTTGTCTTCTTCTTTGTGCCAATTCCACTAATTGTTTGGG** | Ins_k1_39_G | 0.3207104557337018 | 0.03360709357796789 | 67.6650291640075 |
| **GATCCAGATCTAACTTGGGGTGGCTTTGTCTTCTTCTTTGTGTGCCAATTCCACTAATTGTTTGGG** | Ins_k3_39_GTG | 0.1743359760014147 | 0.03405780540625614 | 58.45136323845609 |
| **GATCCAGATCTAACTTGGGGTGGCTTTGTCTTCTTCTTTGTTGCCAATTCCACTAATTGTTTGGG** | Ins_k2_38_TG;Ins_k2_39_GT | 0.32239899157594665 | 0.033473484597889173 | 67.77938050701869 |
| **GATCCAGATCTAACTTGGGGTGGCTTTGTCTTCTTCTTTGTTTGCCAATTCCACTAATTGTTTGGG** | Ins_k3_37_TTG;Ins_k3_38_TGT;I | -0.25051173569544 | 0.03378247536308679 | 38.21955513298401 |
| **GATCCAGATCTAACTTGGGGTGGCTTTGTCTTCTTCTTTGTTTGGG** | Del_k17_38-54;Del_k17_39-55;D | -0.16809546867004505 | 0.02830428860525814 | 41.502910866173025 |
| **GATCCAGATCTAACTTGGGGTGGCTTTGTCTTCTTCTTTT** | Del_k23_41-63 | -6.173070676946207 | 0.6662367486920733 | 0.10236487506685543 |
| **GATCCAGATCTAACTTGGGGTGGCTTTGTCTTCTTCTTTTAAAGCCAATTCCACTAATTGTTTGGG** | Ins_k3_40_AAA | -0.7861991782014786 | 0.04991281180761414 | 22.368637285646905 |
| **GATCCAGATCTAACTTGGGGTGGCTTTGTCTTCTTCTTTTAACGCCAATTCCACTAATTGTTTGGG** | Ins_k3_40_AAC | 0.3543639790075833 | 0.03400373342616321 | 69.98094661666113 |
| **GATCCAGATCTAACTTGGGGTGGCTTTGTCTTCTTCTTTTAAGCCAATTCCACTAATTGTTTGGG** | Ins_k2_40_AA | -0.6452101289431044 | 0.04699282510876841 | 25.75551826895412 |
| **GATCCAGATCTAACTTGGGGTGGCTTTGTCTTCTTCTTTTAAGGCCAATTCCACTAATTGTTTGGG** | Ins_k3_40_AAG | -0.4931795913607518 | 0.05301153998112027 | 29.984465876532443 |
| **GATCCAGATCTAACTTGGGGTGGCTTTGTCTTCTTCTTTTAATGCCAATTCCACTAATTGTTTGGG** | Ins_k3_39_TAA;Ins_k3_40_AAT | -0.629725983717987 | 0.04985799252457358 | 26.157424007587856 |
| **GATCCAGATCTAACTTGGGGTGGCTTTGTCTTCTTCTTTTAATTCCACTAATTGTTTGGG** | Del_k3_41-43 | -1.3281333295401674 | 0.06699901821087446 | 13.010096440051418 |
| **GATCCAGATCTAACTTGGGGTGGCTTTGTCTTCTTCTTTTAATTGTTTGGG** | Del_k12_40-51;Del_k12_41-52 | -1.2457709772971883 | 0.04774707253157712 | 14.127002791124776 |
| **GATCCAGATCTAACTTGGGGTGGCTTTGTCTTCTTCTTTTACAGCCAATTCCACTAATTGTTTGGG** | Ins_k3_40_ACA | -3.5749211065092847 | 0.18151389294898454 | 1.3756659323206644 |
| **GATCCAGATCTAACTTGGGGTGGCTTTGTCTTCTTCTTTTACCAATTCCACTAATTGTTTGGG** | G41A | -0.708937223256993 | 0.031512277830985175 | 24.16539887673706 |
| **GATCCAGATCTAACTTGGGGTGGCTTTGTCTTCTTCTTTTACCGCCAATTCCACTAATTGTTTGGG** | Ins_k3_40_ACC | 0.6078759430013576 | 0.03154493573944453 | 90.17344458594252 |
| **GATCCAGATCTAACTTGGGGTGGCTTTGTCTTCTTCTTTTACGCCAATTCCACTAATTGTTTGGG** | Ins_k2_40_AC | 0.5746722883609454 | 0.03359446178383052 | 87.2285183862851 |
| **GATCCAGATCTAACTTGGGGTGGCTTTGTCTTCTTCTTTTACGGCCAATTCCACTAATTGTTTGGG** | Ins_k3_40_ACG | 0.6781685272197635 | 0.032600343481939666 | 96.74005752407842 |
| **GATCCAGATCTAACTTGGGGTGGCTTTGTCTTCTTCTTTTACTAATTGTTTGGG** | Del_k9_41-49 | -2.243450049957332 | 0.08458075298766385 | 5.209109838549208 |
| **GATCCAGATCTAACTTGGGGTGGCTTTGTCTTCTTCTTTTACTGCCAATTCCACTAATTGTTTGGG** | Ins_k3_39_TAC;Ins_k3_40_ACT | 0.1448949873390516 | 0.03505635933783745 | 56.755582532462896 |
| **GATCCAGATCTAACTTGGGGTGGCTTTGTCTTCTTCTTTTACTTGCCAATTCCACTAATTGTTTGGG** | Ins_k4_38_TTAC;Ins_k4_39_TAC | -0.6000657242416819 | 0.05446129097616258 | 26.944880342191148 |
| **GATCCAGATCTAACTTGGGGTGGCTTTGTCTTCTTCTTTTAGAGCCAATTCCACTAATTGTTTGGG** | Ins_k3_40_AGA | -0.3830981512069599 | 0.04510922457026766 | 33.47372770692374 |
| **GATCCAGATCTAACTTGGGGTGGCTTTGTCTTCTTCTTTTAGCCAATTCCACTAATTGTTTGGG** | Ins_k1_40_A | -3.0387596557953636 | 0.16884133709549093 | 2.3516080694331785 |
| **GATCCAGATCTAACTTGGGGTGGCTTTGTCTTCTTCTTTTAGCGCCAATTCCACTAATTGTTTGGG** | Ins_k3_40_AGC | -1.352638606561631 | 0.0666542226040673 | 12.69515504602535 |
| **GATCCAGATCTAACTTGGGGTGGCTTTGTCTTCTTCTTTTAGGCCAATTCCACTAATTGTTTGGG** | Ins_k2_40_AG | -3.731287617224346 | 0.24790715229275273 | 1.1765323405363264 |
| **GATCCAGATCTAACTTGGGGTGGCTTTGTCTTCTTCTTTTAGGGCCAATTCCACTAATTGTTTGGG** | Ins_k3_40_AGG | -4.514605704338755 | 0.3738813945076901 | 0.5375429207886738 |
| **GATCCAGATCTAACTTGGGGTGGCTTTGTCTTCTTCTTTTAGGGGCCAATTCCACTAATTGTTTGGG** | Ins_k4_40_AGGG | -3.930615445944292 | 0.3054741705238801 | 0.9639109047090328 |
| **GATCCAGATCTAACTTGGGGTGGCTTTGTCTTCTTCTTTTAGTGCCAATTCCACTAATTGTTTGGG** | Ins_k3_39_TAG;Ins_k3_40_AGT | -1.5416138873064622 | 0.07045432221693333 | 10.509137822346046 |
| **GATCCAGATCTAACTTGGGGTGGCTTTGTCTTCTTCTTTTATAGCCAATTCCACTAATTGTTTGGG** | Ins_k3_40_ATA | -3.9995651929711995 | 0.2543054007348001 | 0.899688975493134 |
| **GATCCAGATCTAACTTGGGGTGGCTTTGTCTTCTTCTTTTATCGCCAATTCCACTAATTGTTTGGG** | Ins_k3_40_ATC | 0.47553443413043306 | 0.03232279062138816 | 78.9957038174689 |
| **GATCCAGATCTAACTTGGGGTGGCTTTGTCTTCTTCTTTTATGCCAATTCCACTAATTGTTTGGG** | Ins_k2_39_TA;Ins_k2_40_AT | -0.1888658305184847 | 0.04024027187695322 | 40.64977107155876 |
| **GATCCAGATCTAACTTGGGGTGGCTTTGTCTTCTTCTTTTATGGCCAATTCCACTAATTGTTTGGG** | Ins_k3_40_ATG | 0.2182069579559709 | 0.03581989880392229 | 61.072763206502934 |
| **GATCCAGATCTAACTTGGGGTGGCTTTGTCTTCTTCTTTTATTCCACTAATTGTTTGGG** | Del_k4_41-44 | -1.1369860273989025 | 0.06355674837617717 | 15.750514121937375 |
| **GATCCAGATCTAACTTGGGGTGGCTTTGTCTTCTTCTTTTATTGCCAATTCCACTAATTGTTTGGG** | Ins_k3_38_TTA;Ins_k3_39_TAT;In | -0.7021081857653663 | 0.04770431785149753 | 24.330990062335143 |
| **GATCCAGATCTAACTTGGGGTGGCTTTGTCTTCTTCTTTTATTGTTTGGG** | Del_k13_41-53 | -0.9448498482750634 | 0.040710241534879385 | 19.087032667001978 |
| **GATCCAGATCTAACTTGGGGTGGCTTTGTCTTCTTCTTTTCAAGCCAATTCCACTAATTGTTTGGG** | Ins_k3_40_CAA | 0.4091353835700915 | 0.031310016823893554 | 73.92081246193011 |
| **GATCCAGATCTAACTTGGGGTGGCTTTGTCTTCTTCTTTTCAATTCCACTAATTGTTTGGG** | Del_k2_41-42 | -0.4721837622506027 | 0.04171131636961888 | 30.620670043979676 |
| **GATCCAGATCTAACTTGGGGTGGCTTTGTCTTCTTCTTTTCACGCCAATTCCACTAATTGTTTGGG** | Ins_k3_40_CAC | 0.5717308361851459 | 0.031919664121653375 | 86.97231685808157 |
| **GATCCAGATCTAACTTGGGGTGGCTTTGTCTTCTTCTTTTCACTAATTGTTTGGG** | Del_k8_41-48 | -1.2806830906279685 | 1.0664283141172173 | 13.642309335038254 |
| **GATCCAGATCTAACTTGGGGTGGCTTTGTCTTCTTCTTTTCAGCCAATTCCACTAATTGTTTGGG** | Ins_k2_40_CA | -2.0753081030548834 | 0.08455366728209036 | 6.162921482434854 |
| **GATCCAGATCTAACTTGGGGTGGCTTTGTCTTCTTCTTTTCAGGCCAATTCCACTAATTGTTTGGG** | Ins_k3_40_CAG | -3.677433399669323 | 0.2121698371853511 | 1.241630748212803 |
| **GATCCAGATCTAACTTGGGGTGGCTTTGTCTTCTTCTTTTCATGCCAATTCCACTAATTGTTTGGG** | Ins_k3_39_TCA;Ins_k3_40_CAT | -0.01866345169570005 | 0.039941954430954524 | 48.192122934447475 |
| **GATCCAGATCTAACTTGGGGTGGCTTTGTCTTCTTCTTTTCCAATTCCACTAATTGTTTGGG** | Del_k1_41-41 | -1.022041247793599 | 0.04624459439564769 | 17.669107699462515 |
| **GATCCAGATCTAACTTGGGGTGGCTTTGTCTTCTTCTTTTCCACTAATTGTTTGGG** | Del_k7_39-45;Del_k7_40-46;Del_ | -0.48039652692900753 | 0.03403180432112482 | 30.370219539868913 |
| **GATCCAGATCTAACTTGGGGTGGCTTTGTCTTCTTCTTTTCCAGCCAATTCCACTAATTGTTTGGG** | Ins_k3_40_CCA | -0.3992725786571877 | 0.039657583420157606 | 32.936664381011134 |
| **GATCCAGATCTAACTTGGGGTGGCTTTGTCTTCTTCTTTTCCCAATTCCACTAATTGTTTGGG** | G41C | -1.2894477270503042 | 0.04780998005704723 | 13.523261919502206 |
| **GATCCAGATCTAACTTGGGGTGGCTTTGTCTTCTTCTTTTCCCGCCAATTCCACTAATTGTTTGGG** | Ins_k3_40_CCC | 0.5927741388739483 | 0.028223287392388673 | 88.82189400103925 |
| **GATCCAGATCTAACTTGGGGTGGCTTTGTCTTCTTCTTTTCCGCCAATTCCACTAATTGTTTGGG** | Ins_k2_40_CC | 0.601654075633713 | 0.032436063684827446 | 89.61413914009728 |
| **GATCCAGATCTAACTTGGGGTGGCTTTGTCTTCTTCTTTTCCGGCCAATTCCACTAATTGTTTGGG** | Ins_k3_40_CCG | 0.7465229538647413 | 0.030017715334410687 | 100 |
| **GATCCAGATCTAACTTGGGGTGGCTTTGTCTTCTTCTTTTCCTGCCAATTCCACTAATTGTTTGGG** | Ins_k3_39_TCC;Ins_k3_40_CCT | 0.4285905443053031 | 0.03069530987226359 | 75.37303455152961 |
| **GATCCAGATCTAACTTGGGGTGGCTTTGTCTTCTTCTTTTCGAGCCAATTCCACTAATTGTTTGGG** | Ins_k3_40_CGA | 0.6068766309100668 | 0.030292433201406423 | 90.08337818217282 |
| **GATCCAGATCTAACTTGGGGTGGCTTTGTCTTCTTCTTTTCGCCAATTCCACTAATTGTTTGGG** | Ins_k1_40_C | 0.618442243002759 | 0.03041997440612216 | 91.13129581358419 |
| **GATCCAGATCTAACTTGGGGTGGCTTTGTCTTCTTCTTTTCGCGCCAATTCCACTAATTGTTTGGG** | Ins_k3_40_CGC | 0.5830194361318757 | 0.031757066523080135 | 87.95967501002008 |
| **GATCCAGATCTAACTTGGGGTGGCTTTGTCTTCTTCTTTTCGGCCAATTCCACTAATTGTTTGGG** | Ins_k2_40_CG | 0.849728130810167 | 0.029891827859596664 | 100 |
| **GATCCAGATCTAACTTGGGGTGGCTTTGTCTTCTTCTTTTCGGGCCAATTCCACTAATTGTTTGGG** | Ins_k3_40_CGG | 0.5090624977293372 | 0.031320920168029266 | 81.6891779720462 |
| **GATCCAGATCTAACTTGGGGTGGCTTTGTCTTCTTCTTTTCGTGCCAATTCCACTAATTGTTTGGG** | Ins_k3_39_TCG;Ins_k3_40_CGT | 0.5757808979900083 | 0.03175632738303194 | 87.32527438410291 |
| **GATCCAGATCTAACTTGGGGTGGCTTTGTCTTCTTCTTTTCTAATTGTTTGGG** | Del_k10_41-50 | -2.3261815179301486 | 0.08667925801401227 | 4.795497787761827 |
| **GATCCAGATCTAACTTGGGGTGGCTTTGTCTTCTTCTTTTCTAGCCAATTCCACTAATTGTTTGGG** | Ins_k3_40_CTA | -4.772413148233158 | 0.4082322578379511 | 0.4153830745010558 |
| **GATCCAGATCTAACTTGGGGTGGCTTTGTCTTCTTCTTTTCTCGCCAATTCCACTAATTGTTTGGG** | Ins_k3_40_CTC | 0.3212182619711236 | 0.032785958965087056 | 67.6993986136442 |
| **GATCCAGATCTAACTTGGGGTGGCTTTGTCTTCTTCTTTTCTGCCAATTCCACTAATTGTTTGGG** | Ins_k2_39_TC;Ins_k2_40_CT | 0.27840230537720545 | 0.0329092388519938 | 64.86196135055604 |
| **GATCCAGATCTAACTTGGGGTGGCTTTGTCTTCTTCTTTTCTGGCCAATTCCACTAATTGTTTGGG** | Ins_k3_40_CTG | 0.19242172972622096 | 0.035059620368758745 | 59.51811765035642 |
| **GATCCAGATCTAACTTGGGGTGGCTTTGTCTTCTTCTTTTCTTGCCAATTCCACTAATTGTTTGGG** | Ins_k3_38_TTC;Ins_k3_39_TCT;I | -1.1735753014448136 | 0.053408734668746266 | 15.184630019825287 |
| **GATCCAGATCTAACTTGGGGTGGCTTTGTCTTCTTCTTTTG** | Del_k22_41-62;Del_k22_42-63 | -0.6028602821540552 | 0.028298821145241657 | 26.869686429440748 |
| **GATCCAGATCTAACTTGGGGTGGCTTTGTCTTCTTCTTTTGAAACCAATTCCACTAATTGTTTGGG** | Ins_k3_41_AAA | 0.5125597554996191 | 0.03573022474712552 | 81.97536622981818 |
| **GATCCAGATCTAACTTGGGGTGGCTTTGTCTTCTTCTTTTGAACCAATTCCACTAATTGTTTGGG** | Ins_k2_41_AA | 0.7307072785099895 | 0.03398311836497851 | 100 |
| **GATCCAGATCTAACTTGGGGTGGCTTTGTCTTCTTCTTTTGAACCCAATTCCACTAATTGTTTGGG** | Ins_k3_41_AAC | 0.507704679748066 | 0.031788991858114485 | 81.57833420717897 |
| **GATCCAGATCTAACTTGGGGTGGCTTTGTCTTCTTCTTTTGAAGCCAATTCCACTAATTGTTTGGG** | Ins_k3_40_GAA;Ins_k3_41_AAG | 0.13771603551144174 | 0.035354660438112105 | 56.349595962151014 |
| **GATCCAGATCTAACTTGGGGTGGCTTTGTCTTCTTCTTTTGAATCCAATTCCACTAATTGTTTGGG** | Ins_k3_41_AAT | 0.5679623500922069 | 0.03169791002023084 | 86.6451796846283 |
| **GATCCAGATCTAACTTGGGGTGGCTTTGTCTTCTTCTTTTGAATTCCACTAATTGTTTGGG** | Del_k2_42-43 | 0.2830111033312709 | 0.03221847371402287 | 65.1615869518918 |
| **GATCCAGATCTAACTTGGGGTGGCTTTGTCTTCTTCTTTTGAATTGTTTGGG** | Del_k11_42-52 | -0.35164950236891346 | 0.0359889788051973 | 34.543159162953884 |

| | | | | |
|---|---|---|---|---|
| GATCCAGATCTAACTTGGGGTGGCTTTGTCTTCTTCTTTTGACAACCAATTCCACTAATTGTTTGGG | Ins_k4_41_ACAA | 0.547586396725751 | 0.031031496864195506 | 84.89756664880994 |
| GATCCAGATCTAACTTGGGGTGGCTTTGTCTTCTTCTTTTGACAATTCCACTAATTGTTTGGG | C42A | -0.9320543002351191 | 0.04130030102696883 | 19.332830918477793 |
| GATCCAGATCTAACTTGGGGTGGCTTTGTCTTCTTCTTTTGACACCAATTCCACTAATTGTTTGGG | Ins_k3_41_ACA | 0.6497785278250034 | 0.032588873949787774 | 94.03222687646914 |
| GATCCAGATCTAACTTGGGGTGGCTTTGTCTTCTTCTTTTGACAGCCAATTCCACTAATTGTTTGGG | Ins_k4_40_GACA;Ins_k4_41_AC | -0.29834378962942953 | 0.04495612827354951 | 36.43446783605442 |
| GATCCAGATCTAACTTGGGGTGGCTTTGTCTTCTTCTTTTGACCAATTCCACTAATTGTTTGGG | Ins_k1_41_A | 0.4200923763177229 | 0.03572267761367855 | 74.73521582605483 |
| GATCCAGATCTAACTTGGGGTGGCTTTGTCTTCTTCTTTTGACCACCAATTCCACTAATTGTTTGGG | Ins_k4_41_ACCA | 0.7585713148982418 | 0.033107668085414044 | 100 |
| GATCCAGATCTAACTTGGGGTGGCTTTGTCTTCTTCTTTTGACCCAATTCCACTAATTGTTTGGG | Ins_k2_41_AC | 0.3661047931055602 | 0.03608844601842187 | 70.80742215522547 |
| GATCCAGATCTAACTTGGGGTGGCTTTGTCTTCTTCTTTTGACCCCAATTCCACTAATTGTTTGGG | Ins_k3_41_ACC | -0.7601117145671943 | 0.05255296123062227 | 22.959856472481377 |
| GATCCAGATCTAACTTGGGGTGGCTTTGTCTTCTTCTTTTGACGCCAATTCCACTAATTGTTTGGG | Ins_k3_40_GAC;Ins_k3_41_ACG | 0.7501380546277641 | 0.028240887594465778 | 100 |
| GATCCAGATCTAACTTGGGGTGGCTTTGTCTTCTTCTTTTGACTAATTGTTTGGG | Del_k8_42-49 | -1.0694738699970443 | 0.050128168663372524 | 16.850581469917884 |
| GATCCAGATCTAACTTGGGGTGGCTTTGTCTTCTTCTTTTGACTACCAATTCCACTAATTGTTTGGG | Ins_k4_41_ACTA | 0.6297308767723802 | 0.03424412514538135 | 92.16587212693345 |
| GATCCAGATCTAACTTGGGGTGGCTTTGTCTTCTTCTTTTGACTCCAATTCCACTAATTGTTTGGG | Ins_k3_41_ACT | 0.653625890758128 | 0.030438371386763634 | 94.39469981603244 |
| GATCCAGATCTAACTTGGGGTGGCTTTGTCTTCTTCTTTTGACTCCCAATTCCACTAATTGTTTGGG | Ins_k4_41_ACTC | 0.4010612522442023 | 0.038548517990350355 | 73.3263691504662 |
| GATCCAGATCTAACTTGGGGTGGCTTTGTCTTCTTCTTTTGAGACCAATTCCACTAATTGTTTGGG | Ins_k3_41_AGA | 0.7685100184376108 | 0.029126770872418512 | 100 |
| GATCCAGATCTAACTTGGGGTGGCTTTGTCTTCTTCTTTTGAGCCAATTCCACTAATTGTTTGGG | Ins_k2_40_GA;Ins_k2_41_AG | 0.510977539545568 | 0.03305787734425109 | 81.84576605230126 |
| GATCCAGATCTAACTTGGGGTGGCTTTGTCTTCTTCTTTTGAGCCCAATTCCACTAATTGTTTGGG | Ins_k3_41_AGC | 0.40413651004517614 | 0.03519033109576698 | 73.55221372637192 |
| GATCCAGATCTAACTTGGGGTGGCTTTGTCTTCTTCTTTTGAGGCCAATTCCACTAATTGTTTGGG | Ins_k3_40_GAG;Ins_k3_41_AGG | 0.4067920580215829 | 0.031306467786357026 | 73.74779473112856 |
| GATCCAGATCTAACTTGGGGTGGCTTTGTCTTCTTCTTTTGAGTCCAATTCCACTAATTGTTTGGG | Ins_k3_41_AGT | 0.4445394650593839 | 0.030779149752695557 | 76.5847905206553 |
| GATCCAGATCTAACTTGGGGTGGCTTTGTCTTCTTCTTTTGATACCAATTCCACTAATTGTTTGGG | Ins_k3_41_ATA | 0.3255271439905004 | 0.031001518573176876 | 67.99173670792524 |
| GATCCAGATCTAACTTGGGGTGGCTTTGTCTTCTTCTTTTGATCCAATTCCACTAATTGTTTGGG | Ins_k2_41_AT | 0.5824500858747492 | 0.03505135452693825 | 87.9096094002227 |
| GATCCAGATCTAACTTGGGGTGGCTTTGTCTTCTTCTTTTGATCCCAATTCCACTAATTGTTTGGG | Ins_k3_41_ATC | 0.32187644259428105 | 0.033295476285091954 | 67.74397171297716 |
| GATCCAGATCTAACTTGGGGTGGCTTTGTCTTCTTCTTTTGATGCCAATTCCACTAATTGTTTGGG | Ins_k3_39_TGA;Ins_k3_40_GAT;I | 0.5765136982266601 | 0.02924369344365326 | 87.38928981823523 |
| GATCCAGATCTAACTTGGGGTGGCTTTGTCTTCTTCTTTTGATTCCAATTCCACTAATTGTTTGGG | Ins_k3_41_ATT | 0.1915220758116254 | 0.037876715937506515 | 59.46459602190413 |
| GATCCAGATCTAACTTGGGGTGGCTTTGTCTTCTTCTTTTGATTCCACTAATTGTTTGGG | Del_k3_42-44 | -0.11562938524782423 | 0.03517775385068941 | 43.73854060880516 |
| GATCCAGATCTAACTTGGGGTGGCTTTGTCTTCTTCTTTTGATTGTTTGGG | Del_k12_42-53 | -0.7041742002953958 | 0.03980812128327234 | 24.280773774988006 |
| GATCCAGATCTAACTTGGGGTGGCTTTGTCTTCTTCTTTTGC | Del_k21_43-63 | -4.468996872674614 | 0.13607629632739898 | 0.5626273121093196 |
| GATCCAGATCTAACTTGGGGTGGCTTTGTCTTCTTCTTTTGCAAACAATTCCACTAATTGTTTGGG | Ins_k3_42_AAA | 0.1853589479523048 | 0.039292100415276913 | 59.09923515288618 |
| GATCCAGATCTAACTTGGGGTGGCTTTGTCTTCTTCTTTTGCAAATTCCACTAATTGTTTGGG | C43A | -0.5471697784308674 | 0.035279587810407693 | 28.40852443722507 |
| GATCCAGATCTAACTTGGGGTGGCTTTGTCTTCTTCTTTTGCAACAATTCCACTAATTGTTTGGG | Ins_k2_42_AA | 0.7013800669091053 | 0.03172682774756496 | 99.01180660988847 |
| GATCCAGATCTAACTTGGGGTGGCTTTGTCTTCTTCTTTTGCAACCAATTCCACTAATTGTTTGGG | Ins_k3_41_CAA;Ins_k3_42_AAC | 0.5336747211966029 | 0.03437263716812654 | 83.72467659747447 |
| GATCCAGATCTAACTTGGGGTGGCTTTGTCTTCTTCTTTTGCAACTCAATTCCACTAATTGTTTGGG | Ins_k4_42_AACT | 0.639725911469387 | 0.03704123356417917 | 93.09169231166116 |
| GATCCAGATCTAACTTGGGGTGGCTTTGTCTTCTTCTTTTGCAAGCAATTCCACTAATTGTTTGGG | Ins_k3_42_AAG | 0.5364641767381906 | 0.0337728557061293 | 83.95854889712018 |
| GATCCAGATCTAACTTGGGGTGGCTTTGTCTTCTTCTTTTGCAATCAATTCCACTAATTGTTTGGG | Ins_k3_42_AAT | 0.2226124115356105 | 0.036778480764837584 | 61.342409951790124 |
| GATCCAGATCTAACTTGGGGTGGCTTTGTCTTCTTCTTTTGCAATTCCACTAATTGTTTGGG | Del_k1_42-42;Del_k1_43-43 | -0.3981433326430993 | 0.033622039648566394 | 32.97387898624841 |
| GATCCAGATCTAACTTGGGGTGGCTTTGTCTTCTTCTTTTGCAATTGTTTGGG | Del_k10_43-52 | -0.5240951096028323 | 0.037313132599158944 | 29.071663160497536 |
| GATCCAGATCTAACTTGGGGTGGCTTTGTCTTCTTCTTTTGCACAATTCCACTAATTGTTTGGG | Ins_k1_42_A | -0.21833013527836076 | 0.04204097401315899 | 39.469526751512404 |
| GATCCAGATCTAACTTGGGGTGGCTTTGTCTTCTTCTTTTGCACACAATTCCACTAATTGTTTGGG | Ins_k3_42_ACA | -0.448382311711954 | 0.04311556036635315 | 31.358229075738638 |
| GATCCAGATCTAACTTGGGGTGGCTTTGTCTTCTTCTTTTGCACCAATTCCACTAATTGTTTGGG | Ins_k2_41_CA;Ins_k2_42_AC | 0.5882564981914999 | 0.032268449713061746 | 88.42153362229065 |
| GATCCAGATCTAACTTGGGGTGGCTTTGTCTTCTTCTTTTGCACCCAATTCCACTAATTGTTTGGG | Ins_k3_41_CAC;Ins_k3_42_ACC | 0.13130319275490598 | 0.0395587443337147 | 55.9893910669658 |
| GATCCAGATCTAACTTGGGGTGGCTTTGTCTTCTTCTTTTGCACGCAATTCCACTAATTGTTTGGG | Ins_k3_42_ACG | 0.15743486062337297 | 0.034230524197583864 | 57.47177141960426 |
| GATCCAGATCTAACTTGGGGTGGCTTTGTCTTCTTCTTTTGCACTAATTGTTTGGG | Del_k7_42-48;Del_k7_43-49 | -0.8344423390727072 | 0.042396530762772625 | 21.31512033439799 |
| GATCCAGATCTAACTTGGGGTGGCTTTGTCTTCTTCTTTTGCACTCAATTCCACTAATTGTTTGGG | Ins_k3_42_ACT | -0.6269209292043576 | 0.042179232854902556 | 26.23090001178608 |
| GATCCAGATCTAACTTGGGGTGGCTTTGTCTTCTTCTTTTGCAGACAATTCCACTAATTGTTTGGG | Ins_k3_42_AGA | -2.0525376385358696 | 0.08705055233337873 | 6.304863982742056 |
| GATCCAGATCTAACTTGGGGTGGCTTTGTCTTCTTCTTTTGCAGCAATTCCACTAATTGTTTGGG | Ins_k2_42_AG | -2.2127716868472946 | 0.09493804393852963 | 5.371393370491835 |
| GATCCAGATCTAACTTGGGGTGGCTTTGTCTTCTTCTTTTGCAGCCAATTCCACTAATTGTTTGGG | Ins_k3_40_GCA;Ins_k3_41_CAG | -1.8871201604317511 | 0.07595185591666512 | 7.4390181746858675 |
| GATCCAGATCTAACTTGGGGTGGCTTTGTCTTCTTCTTTTGCAGCCCAATTCCACTAATTGTTTGGG | Ins_k4_41_CAGC;Ins_k4_42_AG | -1.402462272171329 | 0.08065279083787164 | 12.078134631117575 |
| GATCCAGATCTAACTTGGGGTGGCTTTGTCTTCTTCTTTTGCAGCTCAATTCCACTAATTGTTTGGG | Ins_k4_42_AGCT | -3.024474989676617 | 0.1572036052934745 | 2.3854410768496557 |
| GATCCAGATCTAACTTGGGGTGGCTTTGTCTTCTTCTTTTGCAGGCAATTCCACTAATTGTTTGGG | Ins_k3_42_AGG | -1.9891645645547216 | 0.5303602261975526 | 6.717354962264934 |
| GATCCAGATCTAACTTGGGGTGGCTTTGTCTTCTTCTTTTGCAGTCAATTCCACTAATTGTTTGGG | Ins_k3_42_AGT | -0.5935161015393614 | 0.04134756451876475 | 27.12193834078944 |
| GATCCAGATCTAACTTGGGGTGGCTTTGTCTTCTTCTTTTGCATACAATTCCACTAATTGTTTGGG | Ins_k3_42_ATA | -1.1664460960325647 | 0.04595098870888472 | 15.293271168733718 |
| GATCCAGATCTAACTTGGGGTGGCTTTGTCTTCTTCTTTTGCATCAATTCCACTAATTGTTTGGG | Ins_k2_42_AT | 0.38090338785256905 | 0.03502334503678502 | 71.86306424526758 |
| GATCCAGATCTAACTTGGGGTGGCTTTGTCTTCTTCTTTTGCATCCAATTCCACTAATTGTTTGGG | Ins_k3_41_CAT;Ins_k3_42_ATC | 0.2465298746085547 | 0.03225637711554971 | 62.82725080747261 |
| GATCCAGATCTAACTTGGGGTGGCTTTGTCTTCTTCTTTTGCATGCAATTCCACTAATTGTTTGGG | Ins_k3_42_ATG | -1.450802865189418 | 0.05588404018369024 | 11.508157935975309 |
| GATCCAGATCTAACTTGGGGTGGCTTTGTCTTCTTCTTTTGCATTCAATTCCACTAATTGTTTGGG | Ins_k3_42_ATT | -0.6602025793503175 | 0.051493760164969 | 25.372260106920336 |
| GATCCAGATCTAACTTGGGGTGGCTTTGTCTTCTTCTTTTGCATTCCACTAATTGTTTGGG | Del_k2_43-44 | -0.47611815241992617 | 0.03722977103093175 | 30.50043306547357 |
| GATCCAGATCTAACTTGGGGTGGCTTTGTCTTCTTCTTTTGCATTGTTTGGG | Del_k11_43-53 | -0.43906614889457896 | 0.03521758815632543 | 31.65173248345756 |
| GATCCAGATCTAACTTGGGGTGGCTTTGTCTTCTTCTTTTGCC | Del_k20_44-63 | -4.1102204619967715 | 0.1230970276757176 | 0.8054441398424741 |
| GATCCAGATCTAACTTGGGGTGGCTTTGTCTTCTTCTTTTGCCA | Del_k19_45-63 | -3.5726320051571645 | 0.10313482160941775 | 1.378818578052558 |
| GATCCAGATCTAACTTGGGGTGGCTTTGTCTTCTTCTTTTGCCAA | Del_k18_46-63 | -1.062146585128439 | 0.035983482026124704 | 16.974503933706508 |
| GATCCAGATCTAACTTGGGGTGGCTTTGTCTTCTTCTTTTGCCAAAAATTCCACTAATTGTTTGGG | Ins_k3_43_AAA;Ins_k3_44_AAA;I | 0.07125254859965946 | 0.036321491752945856 | 52.72615242478023 |
| GATCCAGATCTAACTTGGGGTGGCTTTGTCTTCTTCTTTTGCCAAAACGTTCCACTAATTGTTTGGG | Ins_k4_45_AACG | 0.7098611191101443 | 0.03545749724701792 | 99.8551018711087 |
| GATCCAGATCTAACTTGGGGTGGCTTTGTCTTCTTCTTTTGCCAAAACTTCCACTAATTGTTTGGG | Ins_k3_45_AAC | 0.3176967160820723 | 0.03779045693473869 | 67.46141136222411 |
| GATCCAGATCTAACTTGGGGTGGCTTTGTCTTCTTCTTTTGCCAAAAGTTCCACTAATTGTTTGGG | Ins_k3_45_AAG | 0.8766915476644226 | 0.22119946121103584 | 100 |
| GATCCAGATCTAACTTGGGGTGGCTTTGTCTTCTTCTTTTGCCAAAATTCCACTAATTGTTTGGG | Ins_k2_43_AA;Ins_k2_44_AA;Ins_ | 0.15295974405716017 | 0.03833662848651958 | 57.21515316979248 |
| GATCCAGATCTAACTTGGGGTGGCTTTGTCTTCTTCTTTTGCCAAAATTGTTTGGG | Del_k7_46-52 | -0.3545364138399602 | 0.03493883117158607 | 34.44357992789805 |
| GATCCAGATCTAACTTGGGGTGGCTTTGTCTTCTTCTTTTGCCAAAATTTCCACTAATTGTTTGGG | Ins_k3_45_AAT | -0.3402621085450837 | 0.046570744591062134 | 34.93876389880466 |
| GATCCAGATCTAACTTGGGGTGGCTTTGTCTTCTTCTTTTGCCAAACATTCCACTAATTGTTTGGG | Ins_k3_44_AAC;Ins_k3_45_ACA | -0.10633797452194327 | 0.04152500418378809 | 44.146827196051994 |
| GATCCAGATCTAACTTGGGGTGGCTTTGTCTTCTTCTTTTGCCAAACCTTCCACTAATTGTTTGGG | Ins_k3_45_ACC | 0.194860726993354 | 0.03391928087433546 | 59.663459348603354 |
| GATCCAGATCTAACTTGGGGTGGCTTTGTCTTCTTCTTTTGCCAAACGTTCCACTAATTGTTTGGG | Ins_k3_45_ACG | 0.37371736601869654 | 0.03499827473809904 | 71.3485057252003 |
| GATCCAGATCTAACTTGGGGTGGCTTTGTCTTCTTCTTTTGCCAAACTAATTGTTTGGG | Del_k4_46-49 | -0.08051340481044011 | 0.2957126542020267 | 45.301748500541066 |
| GATCCAGATCTAACTTGGGGTGGCTTTGTCTTCTTCTTTTGCCAAACTTCCACTAATTGTTTGGG | Ins_k2_45_AC | 0.461956397011944 | 0.03327644683705246 | 77.93034631833457 |
| GATCCAGATCTAACTTGGGGTGGCTTTGTCTTCTTCTTTTGCCAAACTTTCCACTAATTGTTTGGG | Ins_k3_45_ACT | -0.8306061427369108 | 0.048599366114735615 | 21.39704636261515 |
| GATCCAGATCTAACTTGGGGTGGCTTTGTCTTCTTCTTTTGCCAAAGATTCCACTAATTGTTTGGG | Ins_k3_44_AAG;Ins_k3_45_AGA | 0.5561998222245947 | 0.030621649747137346 | 85.63198389770552 |
| GATCCAGATCTAACTTGGGGTGGCTTTGTCTTCTTCTTTTGCCAAAGCTTCCACTAATTGTTTGGG | Ins_k3_45_AGC | -0.1461343565757059 | 0.03932255442775882 | 42.424442840566094 |
| GATCCAGATCTAACTTGGGGTGGCTTTGTCTTCTTCTTTTGCCAAAGCTTTCCACTAATTGTTTGGG | Ins_k4_45_AGCT | -0.34058237855439105 | 0.04508071998118708 | 34.92757585225912 |

| | | | | |
|---|---|---|---|---|
| GATCCAGATCTAACTTGGGGTGGCTTTGTCTTCTTCTTTTGCCAAAGGTTCCACTAATTGTTTGGG | Ins_k3_45_AGG | 0.2203661875083116 | 0.03911801083758398 | 61.20477579311078 |
| GATCCAGATCTAACTTGGGGTGGCTTTGTCTTCTTCTTTTGCCAAAGTTCCACTAATTGTTTGGG | Ins_k2_45_AG | 0.6431374072950957 | 0.03545077038004301 | 93.40981656263219 |
| GATCCAGATCTAACTTGGGGTGGCTTTGTCTTCTTCTTTTGCCAAAGTTTCCACTAATTGTTTGGG | Ins_k3_45_AGT | 0.20014654403863907 | 0.03772470082441223 | 59.979664443923326 |
| GATCCAGATCTAACTTGGGGTGGCTTTGTCTTCTTCTTTTGCCAAATATTCCACTAATTGTTTGGG | Ins_k3_44_AAT;Ins_k3_45_ATA | -0.10578024465149327 | 0.038897012439941175 | 44.17145606775511 |
| GATCCAGATCTAACTTGGGGTGGCTTTGTCTTCTTCTTTTGCCAAATCCACTAATTGTTTGGG | T46A | -0.11805127355698675 | 0.03513907661660226 | 43.63273892029647 |
| GATCCAGATCTAACTTGGGGTGGCTTTGTCTTCTTCTTTTGCCAAATCTTCCACTAATTGTTTGGG | Ins_k3_45_ATC | 0.17258842011421482 | 0.03327132803326089 | 58.349305416361084 |
| GATCCAGATCTAACTTGGGGTGGCTTTGTCTTCTTCTTTTGCCAAATGGTTCCACTAATTGTTTGGG | Ins_k4_45_ATGG | 0.6052542939302066 | 0.034235578185105746 | 89.93735107109666 |
| GATCCAGATCTAACTTGGGGTGGCTTTGTCTTCTTCTTTTGCCAAATGTTCCACTAATTGTTTGGG | Ins_k3_45_ATG | 0.0759474085017503 | 0.04085572102512454 | 52.97427632130106 |
| GATCCAGATCTAACTTGGGGTGGCTTTGTCTTCTTCTTTTGCCAAATTCCACTAATTGTTTGGG | Ins_k1_43_A;Ins_k1_44_A;Ins_k1 | 0.09185290560406234 | 0.03770506724216798 | 53.823595032656904 |
| GATCCAGATCTAACTTGGGGTGGCTTTGTCTTCTTCTTTTGCCAAATTGTTTGGG | Del_k8_45-52;Del_k8_46-53 | -0.3534088848777018 | 0.03958221657006349 | 34.4824379644947 |
| GATCCAGATCTAACTTGGGGTGGCTTTGTCTTCTTCTTTTGCCAAATTTCCACTAATTGTTTGGG | Ins_k2_45_AT | -0.11850377100715132 | 0.04609771103564904 | 43.6129996835053 |
| GATCCAGATCTAACTTGGGGTGGCTTTGTCTTCTTCTTTTGCCAAATTTTCCACTAATTGTTTGGG | Ins_k3_45_ATT | -0.7961739605612425 | 0.05483582243173721 | 22.146624104407486 |
| GATCCAGATCTAACTTGGGGTGGCTTTGTCTTCTTCTTTTGCCAACAATTCCACTAATTGTTTGGG | Ins_k3_42_CAA;Ins_k3_43_AAC; | 0.5850741824601586 | 0.031203835598684476 | 88.14059563862891 |
| GATCCAGATCTAACTTGGGGTGGCTTTGTCTTCTTCTTTTGCCAACACTAATTGTTTGGG | Del_k3_46-48 | -0.08285329858611379 | 0.0332596010272067 | 45.19587114039296 |
| GATCCAGATCTAACTTGGGGTGGCTTTGTCTTCTTCTTTTGCCAACACTTCCACTAATTGTTTGGG | Ins_k3_45_CAC | 0.45955006593630743 | 0.03405976012890819 | 77.74304554846448 |
| GATCCAGATCTAACTTGGGGTGGCTTTGTCTTCTTCTTTTGCCAACAGTTCCACTAATTGTTTGGG | Ins_k3_45_CAG | 0.34196490912019994 | 0.03486200558286285 | 69.118605132722 |
| GATCCAGATCTAACTTGGGGTGGCTTTGTCTTCTTCTTTTGCCAACATTCCACTAATTGTTTGGG | Ins_k2_44_AC;Ins_k2_45_CA | 0.5102533857249136 | 0.032475967450137595 | 81.7865185828399 |
| GATCCAGATCTAACTTGGGGTGGCTTTGTCTTCTTCTTTTGCCAACATTTCCACTAATTGTTTGGG | Ins_k3_45_CAT | 0.28493009148804294 | 0.0339866022944249 | 65.28675132159053 |
| GATCCAGATCTAACTTGGGGTGGCTTTGTCTTCTTCTTTTGCCAACCACTAATTGTTTGGG | Del_k2_46-47 | 0.32625800689459417 | 0.03239712352151791 | 68.04144750974767 |
| GATCCAGATCTAACTTGGGGTGGCTTTGTCTTCTTCTTTTGCCAACCATTCCACTAATTGTTTGGG | Ins_k3_44_ACC;Ins_k3_45_CCA | 0.46837349424133334 | 0.02968983949057077 | 78.43204091767967 |
| GATCCAGATCTAACTTGGGGTGGCTTTGTCTTCTTCTTTTGCCAACCCTTCCACTAATTGTTTGGG | Ins_k3_45_CCC | -0.9402376467534739 | 0.05999757092796938 | 19.17526923410896 |
| GATCCAGATCTAACTTGGGGTGGCTTTGTCTTCTTCTTTTGCCAACCGTTCCACTAATTGTTTGGG | Ins_k3_45_CCG | 0.7156486275161376 | 0.03252530333937811 | 100 |
| GATCCAGATCTAACTTGGGGTGGCTTTGTCTTCTTCTTTTGCCAACCTTCCACTAATTGTTTGGG | Ins_k2_45_CC | 0.5280229873152444 | 0.0330431833290895 | 83.25282166130995 |
| GATCCAGATCTAACTTGGGGTGGCTTTGTCTTCTTCTTTTGCCAACCTTTCCACTAATTGTTTGGG | Ins_k3_45_CCT | -0.1446089105330548 | 0.036764911458722946 | 42.489208424651835 |
| GATCCAGATCTAACTTGGGGTGGCTTTGTCTTCTTCTTTTGCCAACGAGTTCCACTAATTGTTTGGG | Ins_k4_45_CGAG | 0.9498408735688153 | 0.028587714529561638 | 100 |
| GATCCAGATCTAACTTGGGGTGGCTTTGTCTTCTTCTTTTGCCAACGATTCCACTAATTGTTTGGG | Ins_k3_44_ACG;Ins_k3_45_CGA | 0.800086439215957 | 0.030408562958641604 | 100 |
| GATCCAGATCTAACTTGGGGTGGCTTTGTCTTCTTCTTTTGCCAACGCTTCCACTAATTGTTTGGG | Ins_k3_45_CGC | 0.8499171813085906 | 0.028717819124824157 | 100 |
| GATCCAGATCTAACTTGGGGTGGCTTTGTCTTCTTCTTTTGCCAACGGTTCCACTAATTGTTTGGG | Ins_k3_45_CGG | 1.002219960773615 | 0.03270875784161952 | 100 |
| GATCCAGATCTAACTTGGGGTGGCTTTGTCTTCTTCTTTTGCCAACGTTCCACTAATTGTTTGGG | Ins_k2_45_CG | 0.7793100783032545 | 0.032448079548399164 | 100 |
| GATCCAGATCTAACTTGGGGTGGCTTTGTCTTCTTCTTTTGCCAACGTTTCCACTAATTGTTTGGG | Ins_k3_45_CGT | 0.6155627045721026 | 0.03225536265008204 | 90.86925720124376 |
| GATCCAGATCTAACTTGGGGTGGCTTTGTCTTCTTCTTTTGCCAACTAATTGTTTGGG | Del_k5_45-49;Del_k5_46-50 | 0.13678381254902028 | 0.03111577996588371 | 56.297090052281824 |
| GATCCAGATCTAACTTGGGGTGGCTTTGTCTTCTTCTTTTGCCAACTACTTCCACTAATTGTTTGGG | Ins_k4_45_CTAC | 0.5207602174242787 | 0.05041458532009129 | 82.65036597151585 |
| GATCCAGATCTAACTTGGGGTGGCTTTGTCTTCTTCTTTTGCCAACTATTCCACTAATTGTTTGGG | Ins_k3_44_ACT;Ins_k3_45_CTA | 0.1514876090468581 | 0.03368720495364993 | 57.13098670708999 |
| GATCCAGATCTAACTTGGGGTGGCTTTGTCTTCTTCTTTTGCCAACTCCACTAATTGTTTGGG | T46C | -0.2438395584622554 | 0.027615481109116477 | 38.47541540193828 |
| GATCCAGATCTAACTTGGGGTGGCTTTGTCTTCTTCTTTTGCCAACTCTTCCACTAATTGTTTGGG | Ins_k3_45_CTC | 0.5132764540174786 | 0.03123652955790466 | 82.0341389119274 |
| GATCCAGATCTAACTTGGGGTGGCTTTGTCTTCTTCTTTTGCCAACTGTTCCACTAATTGTTTGGG | Ins_k3_45_CTG | 0.4993899126405781 | 0.0320614104476226 | 80.9028415310693 |
| GATCCAGATCTAACTTGGGGTGGCTTTGTCTTCTTCTTTTGCCAACTTCCACTAATTGTTTGGG | Ins_k1_45_C | 0.41191288551501826 | 0.03214266862620123 | 74.12641305816588 |
| GATCCAGATCTAACTTGGGGTGGCTTTGTCTTCTTCTTTTGCCAACTTTCCACTAATTGTTTGGG | Ins_k2_45_CT | 0.1537112945952882 | 0.03625298381372516 | 57.258169411361344 |
| GATCCAGATCTAACTTGGGGTGGCTTTGTCTTCTTCTTTTGCCAACTTTTCCACTAATTGTTTGGG | Ins_k3_45_CTT | 0.02424779777524322 | 0.03621139301873438 | 50.30511856055859 |
| GATCCAGATCTAACTTGGGGTGGCTTTGTCTTCTTCTTTTGCCAAG | Del_k17_46-62 | -0.6737797898137563 | 0.025994863842211616 | 25.030103612976642 |
| GATCCAGATCTAACTTGGGGTGGCTTTGTCTTCTTCTTTTGCCAAGAATTCCACTAATTGTTTGGG | Ins_k3_43_AAG;Ins_k3_44_AGA; | 0.6843628905144834 | 0.03170600224479527 | 97.34116038826177 |
| GATCCAGATCTAACTTGGGGTGGCTTTGTCTTCTTCTTTTGCCAAGACTTCCACTAATTGTTTGGG | Ins_k3_45_GAC | 0.8679119870568945 | 0.02978975858075768 | 100 |
| GATCCAGATCTAACTTGGGGTGGCTTTGTCTTCTTCTTTTGCCAAGAGTTCCACTAATTGTTTGGG | Ins_k3_45_GAG | 1.0235760798916842 | 0.03255056778678385 | 100 |
| GATCCAGATCTAACTTGGGGTGGCTTTGTCTTCTTCTTTTGCCAAGATTCCACTAATTGTTTGGG | Ins_k2_44_AG;Ins_k2_45_GA | 0.7054799558057032 | 0.028294213612524852 | 99.41857730397963 |
| GATCCAGATCTAACTTGGGGTGGCTTTGTCTTCTTCTTTTGCCAAGATTTCCACTAATTGTTTGGG | Ins_k3_45_GAT | 0.7979742924270412 | 0.03255046921261504 | 100 |
| GATCCAGATCTAACTTGGGGTGGCTTTGTCTTCTTCTTTTGCCAAGCATTCCACTAATTGTTTGGG | Ins_k3_44_AGC;Ins_k3_45_GCA | 0.3355027103789998 | 0.03493485311392244 | 68.67338706282352 |
| GATCCAGATCTAACTTGGGGTGGCTTTGTCTTCTTCTTTTGCCAAGCCATTCCACTAATTGTTTGGG | Ins_k4_44_AGCC;Ins_k4_45_GC | -1.493544089948025 | 0.07157830269172635 | 11.026648620053596 |
| GATCCAGATCTAACTTGGGGTGGCTTTGTCTTCTTCTTTTGCCAAGCCTTCCACTAATTGTTTGGG | Ins_k3_45_GCC | 0.62932885180214 | 0.032828641702977876 | 92.12882659204844 |
| GATCCAGATCTAACTTGGGGTGGCTTTGTCTTCTTCTTTTGCCAAGCGTTCCACTAATTGTTTGGG | Ins_k3_45_GCG | 0.7016430214930285 | 0.03281737851361651 | 99.0378456416904 |
| GATCCAGATCTAACTTGGGGTGGCTTTGTCTTCTTCTTTTGCCAAGCTTCCACTAATTGTTTGGG | Ins_k2_45_GC | 0.761200117610483 | 0.032479069975812215 | 100 |
| GATCCAGATCTAACTTGGGGTGGCTTTGTCTTCTTCTTTTGCCAAGCTTTCCACTAATTGTTTGGG | Ins_k3_45_GCT | 0.1141305782857176 | 0.03756324157855411 | 55.03611538422993 |
| GATCCAGATCTAACTTGGGGTGGCTTTGTCTTCTTCTTTTGCCAAGG | Del_k16_46-61 | -0.39486506775529273 | 0.02959349820737929 | 33.08215347520251 |
| GATCCAGATCTAACTTGGGGTGGCTTTGTCTTCTTCTTTTGCCAAGGATTCCACTAATTGTTTGGG | Ins_k3_44_AGG;Ins_k3_45_GGA | 0.769384012325115 | 0.029768074502441958 | 100 |
| GATCCAGATCTAACTTGGGGTGGCTTTGTCTTCTTCTTTTGCCAAGGCTTCCACTAATTGTTTGGG | Ins_k3_45_GGC | 0.436544036862597 | 0.03156088366999067 | 75.97490372911584 |
| GATCCAGATCTAACTTGGGGTGGCTTTGTCTTCTTCTTTTGCCAAGGG | Del_k15_46-60 | -0.5396256411428056 | 0.030948933957481047 | 28.623652704003636 |
| GATCCAGATCTAACTTGGGGTGGCTTTGTCTTCTTCTTTTGCCAAGGGTTCCACTAATTGTTTGGG | Ins_k3_45_GGG | 0.20519756064764438 | 0.03674650859409475 | 60.283389138733284 |
| GATCCAGATCTAACTTGGGGTGGCTTTGTCTTCTTCTTTTGCCAAGGTTCCACTAATTGTTTGGG | Ins_k2_45_GG | 0.6660567921344889 | 0.031999501131880805 | 95.57543461420214 |
| GATCCAGATCTAACTTGGGGTGGCTTTGTCTTCTTCTTTTGCCAAGGTTTCCACTAATTGTTTGGG | Ins_k3_45_GGT | 0.123844183781469 | 0.03417783412359711 6 | 55.57331936692568 |
| GATCCAGATCTAACTTGGGGTGGCTTTGTCTTCTTCTTTTGCCAAGTATTCCACTAATTGTTTGGG | Ins_k3_44_AGT;Ins_k3_45_GTA | 0.2472277516652024 | 0.03202148843793895 | 62.871111807355994 |
| GATCCAGATCTAACTTGGGGTGGCTTTGTCTTCTTCTTTTGCCAAGTCCACTAATTGTTTGGG | T46G | 0.2844233621181602 | 0.033641352141091964 | 65.25367698780754 |
| GATCCAGATCTAACTTGGGGTGGCTTTGTCTTCTTCTTTTGCCAAGTCTTCCACTAATTGTTTGGG | Ins_k3_45_GTC | 0.6152264908073182 | 0.03171773770947331 | 90.83871084151801 |
| GATCCAGATCTAACTTGGGGTGGCTTTGTCTTCTTCTTTTGCCAAGTGTTCCACTAATTGTTTGGG | Ins_k3_45_GTG | 0.5483355185039289 | 0.030535073482333946 | 84.9611890924046 |
| GATCCAGATCTAACTTGGGGTGGCTTTGTCTTCTTCTTTTGCCAAGTTCCACTAATTGTTTGGG | Ins_k1_45_G | 0.4946684991295818 | 0.030322896110193613 | 80.52176607728744 |
| GATCCAGATCTAACTTGGGGTGGCTTTGTCTTCTTCTTTTGCCAAGTTCTTCCACTAATTGTTTGGG | Ins_k4_45_GTTC | 0.6683084146228615 | 0.031949653961413346 | 95.79087686840691 |
| GATCCAGATCTAACTTGGGGTGGCTTTGTCTTCTTCTTTTGCCAAGTTTCCACTAATTGTTTGGG | Ins_k2_45_GT | 0.5547642063843634 | 0.03888122030399131 | 85.50913746643167 |
| GATCCAGATCTAACTTGGGGTGGCTTTGTCTTCTTCTTTTGCCAAGTTTGGG | Del_k11_46-56 | 0.06125713916524578 | 0.02884418788086395 | 52.20175807743111 |
| GATCCAGATCTAACTTGGGGTGGCTTTGTCTTCTTCTTTTGCCAAGTTTTCCACTAATTGTTTGGG | Ins_k3_45_GTT | 0.20752469158605324 | 0.036928155050306305 | 60.4238398388703 |
| GATCCAGATCTAACTTGGGGTGGCTTTGTCTTCTTCTTTTGCCAAT | Del_k17_47-63 | -3.100538944349856 | 0.07836557208678978 | 2.2107240591606527 |
| GATCCAGATCTAACTTGGGGTGGCTTTGTCTTCTTCTTTTGCCAATAAATCCACTAATTGTTTGGG | Ins_k3_46_AAA | -0.3110540852086213 | 0.041597280253008816 | 35.97430557474507 |
| GATCCAGATCTAACTTGGGGTGGCTTTGTCTTCTTCTTTTGCCAATAACTCCACTAATTGTTTGGG | Ins_k3_46_AAC | 0.19025818613282064 | 0.03737122353440949 | 59.38948680761651 |
| GATCCAGATCTAACTTGGGGTGGCTTTGTCTTCTTCTTTTGCCAATAAGTCCACTAATTGTTTGGG | Ins_k3_46_AAG | 0.049430410820670345 | 0.03436602828480787 | 51.588018484963904 |
| GATCCAGATCTAACTTGGGGTGGCTTTGTCTTCTTCTTTTGCCAATAATCCACTAATTGTTTGGG | Ins_k2_46_AA | 0.013421632362407454 | 0.03680119468745272 | 49.763444443310874 |
| GATCCAGATCTAACTTGGGGTGGCTTTGTCTTCTTCTTTTGCCAATAATTCCACTAATTGTTTGGG | Ins_k3_43_AAT;Ins_k3_44_ATA;In | -0.4218915701908969 | 0.04536656271874601 | 32.200032588323225 |
| GATCCAGATCTAACTTGGGGTGGCTTTGTCTTCTTCTTTTGCCAATAATTGTTTGGG | Del_k6_46-51;Del_k6_47-52 | -0.9937952621655581 | 0.04506538832287368 | 18.17530441953808 |
| GATCCAGATCTAACTTGGGGTGGCTTTGTCTTCTTCTTTTGCCAATACATCCACTAATTGTTTGGG | Ins_k3_46_ACA | 0.1786926837745635 | 0.03143222210053633 | 58.70657428332182 |

| | | | | |
|---|---|---|---|---|
| GATCCAGATCTAACTTGGGGTGGCTTTGTCTTCTTCTTTTGCCAATACCACTAATTGTTTGGG | T47A | -0.2674316037166501 | 0.035718510880264794 | 37.57832536347544 |
| GATCCAGATCTAACTTGGGGTGGCTTTGTCTTCTTCTTTTGCCAATACCTCCACTAATTGTTTGGG | Ins_k3_46_ACC | 0.25361138768149105 | 0.03424962738620502 | 63.273741855621466 |
| GATCCAGATCTAACTTGGGGTGGCTTTGTCTTCTTCTTTTGCCAATACGCTCCACTAATTGTTTGGG | Ins_k4_46_ACGC | 0.708030414284911 | 0.030218372803813225 | 99.67246388342073 |
| GATCCAGATCTAACTTGGGGTGGCTTTGTCTTCTTCTTTTGCCAATACGTCCACTAATTGTTTGGG | Ins_k3_46_ACG | 0.44168278102766645 | 0.030563608879222087 | 76.36632416574511 |
| GATCCAGATCTAACTTGGGGTGGCTTTGTCTTCTTCTTTTGCCAATACTAATTGTTTGGG | Del_k3_47-49 | -1.4980576975778641 | 0.054476496790683726 | 10.976990806957694 |
| GATCCAGATCTAACTTGGGGTGGCTTTGTCTTCTTCTTTTGCCAATACTCCACTAATTGTTTGGG | Ins_k2_46_AC | 0.25251628197982545 | 0.03555919395971649 | 63.20448834702678 |
| GATCCAGATCTAACTTGGGGTGGCTTTGTCTTCTTCTTTTGCCAATACTTCCACTAATTGTTTGGG | Ins_k3_45_TAC;Ins_k3_46_ACT | 0.08708067320867985 | 0.03843937581497346 | 53.56734824972329 |
| GATCCAGATCTAACTTGGGGTGGCTTTGTCTTCTTCTTTTGCCAATAGATCCACTAATTGTTTGGG | Ins_k3_46_AGA | 0.20234180861417406 | 0.03444174153098686 | 60.111480308941935 |
| GATCCAGATCTAACTTGGGGTGGCTTTGTCTTCTTCTTTTGCCAATAGCTCCACTAATTGTTTGGG | Ins_k3_46_AGC | -0.2441945650604576 | 0.03993260786091835 | 38.461758799837376 |
| GATCCAGATCTAACTTGGGGTGGCTTTGTCTTCTTCTTTTGCCAATAGGTCCACTAATTGTTTGGG | Ins_k3_46_AGG | -2.5970420512155115 | 0.10734030356929276 | 3.6576358042995123 |
| GATCCAGATCTAACTTGGGGTGGCTTTGTCTTCTTCTTTTGCCAATAGTATCCACTAATTGTTTGGG | Ins_k4_46_AGTA | -0.6188664995176665 | 0.04551717958824698 | 26.44302809009968 |
| GATCCAGATCTAACTTGGGGTGGCTTTGTCTTCTTCTTTTGCCAATAGTCCACTAATTGTTTGGG | Ins_k2_46_AG | -0.20675054306086416 | 0.04174676783484272 | 39.92922419416944 |
| GATCCAGATCTAACTTGGGGTGGCTTTGTCTTCTTCTTTTGCCAATAGTTCCACTAATTGTTTGGG | Ins_k3_45_TAG;Ins_k3_46_AGT | -0.34516587429513007 | 0.04065103069203256 | 34.76785178402844 |
| GATCCAGATCTAACTTGGGGTGGCTTTGTCTTCTTCTTTTGCCAATATATCCACTAATTGTTTGGG | Ins_k3_46_ATA | -0.28951472582735116 | 0.04208075970847035 | 36.75757434224569 |
| GATCCAGATCTAACTTGGGGTGGCTTTGTCTTCTTCTTTTGCCAATATCCACTAATTGTTTGGG | Ins_k1_46_A | 0.14776602689983165 | 0.0317078116423438 | 56.9187641936207 |
| GATCCAGATCTAACTTGGGGTGGCTTTGTCTTCTTCTTTTGCCAATATCTCCACTAATTGTTTGGG | Ins_k3_46_ATC | 0.3466831540562655 | 0.03337605756926696 | 69.44549420630642 |
| GATCCAGATCTAACTTGGGGTGGCTTTGTCTTCTTCTTTTGCCAATATGTCCACTAATTGTTTGGG | Ins_k3_46_ATG | -0.1261359056323335 | 0.034698558494136236 | 43.28140639043268 |
| GATCCAGATCTAACTTGGGGTGGCTTTGTCTTCTTCTTTTGCCAATATTCCACTAATTGTTTGGG | Ins_k2_44_AT;Ins_k2_45_TA;Ins_ | 0.08240758985078345 | 0.038025962421940486 | 53.31760755012462 |
| GATCCAGATCTAACTTGGGGTGGCTTTGTCTTCTTCTTTTGCCAATATTGTTTGGG | Del_k7_47-53 | -0.38754761481053157 | 0.040084847687457625 | 33.32511843476678 |
| GATCCAGATCTAACTTGGGGTGGCTTTGTCTTCTTCTTTTGCCAATATTTCCACTAATTGTTTGGG | Ins_k3_45_TAT;Ins_k3_46_ATT | -0.6232584117608956 | 0.04955434377338089 | 26.327147286709945 |
| GATCCAGATCTAACTTGGGGTGGCTTTGTCTTCTTCTTTTGCCAATCAATCCACTAATTGTTTGGG | Ins_k3_46_CAA | 0.22745127429882017 | 0.034352152973117216 | 61.639956768639664 |
| GATCCAGATCTAACTTGGGGTGGCTTTGTCTTCTTCTTTTGCCAATCACTAATTGTTTGGG | Del_k2_47-48 | -0.3958987064276023 | 0.03835080830145913 | 33.047976148553474 |
| GATCCAGATCTAACTTGGGGTGGCTTTGTCTTCTTCTTTTGCCAATCACTCCACTAATTGTTTGGG | Ins_k3_46_CAC | 0.024890423072497336 | 0.03591615562962009 | 50.33745629173616 |
| GATCCAGATCTAACTTGGGGTGGCTTTGTCTTCTTCTTTTGCCAATCAGTCCACTAATTGTTTGGG | Ins_k3_46_CAG | 0.3476149755278636 | 0.03406755895353349 | 69.51023516773807 |
| GATCCAGATCTAACTTGGGGTGGCTTTGTCTTCTTCTTTTGCCAATCATCCACTAATTGTTTGGG | Ins_k2_46_CA | 0.7082891810807843 | 0.03247689698523576 | 99.69825914487127 |
| GATCCAGATCTAACTTGGGGTGGCTTTGTCTTCTTCTTTTGCCAATCATTCCACTAATTGTTTGGG | Ins_k3_44_ATC;Ins_k3_45_TCA;I | -0.03738416758185936 | 0.03394526684297767 | 47.2983242720747 |
| GATCCAGATCTAACTTGGGGTGGCTTTGTCTTCTTCTTTTGCCAATCCACTAATTGTTTGGG | Del_k1_46-46;Del_k1_47-47 | 0.1342461904821569 | 0.03214253052080143 | 56.15441042431692 |
| GATCCAGATCTAACTTGGGGTGGCTTTGTCTTCTTCTTTTGCCAATCCATCCACTAATTGTTTGGG | Ins_k3_46_CCA | 0.31297877839114563 | 0.03249584453096801 | 67.14388225712973 |
| GATCCAGATCTAACTTGGGGTGGCTTTGTCTTCTTCTTTTGCCAATCCCACTAATTGTTTGGG | T47C | -0.9843468837471274 | 0.03217573998057169 | 18.347845406245685 |
| GATCCAGATCTAACTTGGGGTGGCTTTGTCTTCTTCTTTTGCCAATCCCTCCACTAATTGTTTGGG | Ins_k3_46_CCC | -0.3340354397779847 | 0.04152919794584742 | 35.15699472918966 |
| GATCCAGATCTAACTTGGGGTGGCTTTGTCTTCTTCTTTTGCCAATCCGTCCACTAATTGTTTGGG | Ins_k3_46_CCG | 0.6653756826716837 | 0.028886871973031865 | 95.51035944544707 |
| GATCCAGATCTAACTTGGGGTGGCTTTGTCTTCTTCTTTTGCCAATCCTCCACTAATTGTTTGGG | Ins_k2_46_CC | 0.6991392762195066 | 0.032906053704317983 | 98.79019026612717 |
| GATCCAGATCTAACTTGGGGTGGCTTTGTCTTCTTCTTTTGCCAATCCTTCCACTAATTGTTTGGG | Ins_k3_45_TCC;Ins_k3_46_CCT | 0.23946571273373674 | 0.03734565045874469 | 62.384992867599905 |
| GATCCAGATCTAACTTGGGGTGGCTTTGTCTTCTTCTTTTGCCAATCGATCCACTAATTGTTTGGG | Ins_k3_46_CGA | 0.5932924305688518 | 0.02865252426274806 | 88.86794158303447 |
| GATCCAGATCTAACTTGGGGTGGCTTTGTCTTCTTCTTTTGCCAATCGCTCCACTAATTGTTTGGG | Ins_k3_46_CGC | 0.5808293065194325 | 0.029627584289644307 | 87.76724272382343 |
| GATCCAGATCTAACTTGGGGTGGCTTTGTCTTCTTCTTTTGCCAATCGGTCCACTAATTGTTTGGG | Ins_k3_46_CGG | 0.48828692263427054 | 0.03060215464717889 | 80.00954639040862 |
| GATCCAGATCTAACTTGGGGTGGCTTTGTCTTCTTCTTTTGCCAATCGTCCACTAATTGTTTGGG | Ins_k2_46_CG | 0.657797486590769 | 0.0281695933978685 | 94.78929883397666 |
| GATCCAGATCTAACTTGGGGTGGCTTTGTCTTCTTCTTTTGCCAATCGTTCCACTAATTGTTTGGG | Ins_k3_45_TCG;Ins_k3_46_CGT | 0.31720491095196746 | 0.03534425749503174 | 67.42824165121766 |
| GATCCAGATCTAACTTGGGGTGGCTTTGTCTTCTTCTTTTGCCAATCTAATTGTTTGGG | Del_k4_47-50 | -0.9532665572249377 | 0.05310406171762294 | 18.92705684768485 |
| GATCCAGATCTAACTTGGGGTGGCTTTGTCTTCTTCTTTTGCCAATCTATCCACTAATTGTTTGGG | Ins_k3_46_CTA | -0.03046447628000426 | 0.3919772832688597 | 47.62674906374752 |
| GATCCAGATCTAACTTGGGGTGGCTTTGTCTTCTTCTTTTGCCAATCTCCACTAATTGTTTGGG | Ins_k1_46_C | 0.4637690672298225 | 0.031621199827957365 | 78.07173644429533 |
| GATCCAGATCTAACTTGGGGTGGCTTTGTCTTCTTCTTTTGCCAATCTCTCCACTAATTGTTTGGG | Ins_k3_46_CTC | 0.32302866473992375 | 0.033305172538297165 | 67.82207280368112 |
| GATCCAGATCTAACTTGGGGTGGCTTTGTCTTCTTCTTTTGCCAATCTGTCCACTAATTGTTTGGG | Ins_k3_46_CTG | -0.41086575230151456 | 0.04296392105376307 | 32.55702875441002 |
| GATCCAGATCTAACTTGGGGTGGCTTTGTCTTCTTCTTTTGCCAATCTTCCACTAATTGTTTGGG | Ins_k2_45_TC;Ins_k2_46_CT | 0.3092392127818041 | 0.03622571553583878 | 66.89326220129738 |
| GATCCAGATCTAACTTGGGGTGGCTTTGTCTTCTTCTTTTGCCAATCTTTCCACTAATTGTTTGGG | Ins_k3_45_TCT;Ins_k3_46_CTT | -1.2606631471666936 | 0.061477722932209704 | 13.918179838578276 |
| GATCCAGATCTAACTTGGGGTGGCTTTGTCTTCTTCTTTTGCCAATG | Del_k16_47-62 | -0.4413328931083806 | 0.028758138996529358 | 31.580067355935846 |
| GATCCAGATCTAACTTGGGGTGGCTTTGTCTTCTTCTTTTGCCAATGAATCCACTAATTGTTTGGG | Ins_k3_46_GAA | 0.6305337682929051 | 0.030944668023627567 | 92.23990103876194 |
| GATCCAGATCTAACTTGGGGTGGCTTTGTCTTCTTCTTTTGCCAATGACGTCCACTAATTGTTTGGG | Ins_k4_46_GACG | 0.9729205513919932 | 0.02980313817700497 | 100 |
| GATCCAGATCTAACTTGGGGTGGCTTTGTCTTCTTCTTTTGCCAATGACTCCACTAATTGTTTGGG | Ins_k3_46_GAC | 0.5563434306782293 | 0.030195984947331035 | 85.64428225754781 |
| GATCCAGATCTAACTTGGGGTGGCTTTGTCTTCTTCTTTTGCCAATGAGTCCACTAATTGTTTGGG | Ins_k3_46_GAG | 0.44484405785782033 | 0.030929212205998154 | 76.60812124932325 |
| GATCCAGATCTAACTTGGGGTGGCTTTGTCTTCTTCTTTTGCCAATGATCCACTAATTGTTTGGG | Ins_k2_46_GA | 0.5002636903587288 | 0.03520921161947065 | 80.97356352448561 |
| GATCCAGATCTAACTTGGGGTGGCTTTGTCTTCTTCTTTTGCCAATGATTCCACTAATTGTTTGGG | Ins_k3_44_ATG;Ins_k3_45_TGA;I | 0.38074460833071555 | 0.03534052981244338 | 71.85165476810742 |
| GATCCAGATCTAACTTGGGGTGGCTTTGTCTTCTTCTTTTGCCAATGCATCCACTAATTGTTTGGG | Ins_k3_46_GCA | 0.46517297107058764 | 0.030837194903729753 | 78.18141862854975 |
| GATCCAGATCTAACTTGGGGTGGCTTTGTCTTCTTCTTTTGCCAATGCCACTAATTGTTTGGG | T47G | -0.944294193532821 | 0.047323418221358146 | 19.097641414346164 |
| GATCCAGATCTAACTTGGGGTGGCTTTGTCTTCTTCTTTTGCCAATGCCATCCACTAATTGTTTGGG | Ins_k4_46_GCCA | -1.5282497522720628 | 0.07135633639949687 | 10.650526020462282 |
| GATCCAGATCTAACTTGGGGTGGCTTTGTCTTCTTCTTTTGCCAATGCCTCCACTAATTGTTTGGG | Ins_k3_46_GCC | 0.37251605630760865 | 0.03216439014515784 | 71.2628455349119 |
| GATCCAGATCTAACTTGGGGTGGCTTTGTCTTCTTCTTTTGCCAATGCGTCCACTAATTGTTTGGG | Ins_k3_46_GCG | 0.6596128478094172 | 0.03041042896722482 | 94.96153193637002 |
| GATCCAGATCTAACTTGGGGTGGCTTTGTCTTCTTCTTTTGCCAATGCTCCACTAATTGTTTGGG | Ins_k2_46_GC | 0.5946721194083633 | 0.03302609103300202 | 88.9906363110375 |
| GATCCAGATCTAACTTGGGGTGGCTTTGTCTTCTTCTTTTGCCAATGCTTCCACTAATTGTTTGGG | Ins_k3_45_TGC;Ins_k3_46_GCT | 0.1391261465006195 | 0.033343664710097115 | 56.42911119612983 |
| GATCCAGATCTAACTTGGGGTGGCTTTGTCTTCTTCTTTTGCCAATGG | Del_k15_47-61 | -0.3573832500965691 | 0.027707353509819253 | 34.345664136993406 |
| GATCCAGATCTAACTTGGGGTGGCTTTGTCTTCTTCTTTTGCCAATGGATCCACTAATTGTTTGGG | Ins_k3_46_GGA | 0.735407486188393 | 0.029408121150116248 | 100 |
| GATCCAGATCTAACTTGGGGTGGCTTTGTCTTCTTCTTTTGCCAATGGCTCCACTAATTGTTTGGG | Ins_k3_46_GGC | 0.4103943031673525 | 0.03003031689342738 | 74.01393142372831 |
| GATCCAGATCTAACTTGGGGTGGCTTTGTCTTCTTCTTTTGCCAATGGG | Del_k14_46-59;Del_k14_47-60 | -0.32096908983579747 | 0.029010427294293623 | 35.619382607565235 |
| GATCCAGATCTAACTTGGGGTGGCTTTGTCTTCTTCTTTTGCCAATGGGTCCACTAATTGTTTGGG | Ins_k3_46_GGG | 0.3232198030348724 | 0.03085747358590218 | 67.83503743801629 |
| GATCCAGATCTAACTTGGGGTGGCTTTGTCTTCTTCTTTTGCCAATGGTCCACTAATTGTTTGGG | Ins_k2_46_GG | 0.717064232434089 | 0.03373365883132984 | 100 |
| GATCCAGATCTAACTTGGGGTGGCTTTGTCTTCTTCTTTTGCCAATGGTTCCACTAATTGTTTGGG | Ins_k3_45_TGG;Ins_k3_46_GGT | 0.49179452703790383 | 0.029225236171461964 | 80.29068099390697 |
| GATCCAGATCTAACTTGGGGTGGCTTTGTCTTCTTCTTTTGCCAATGTATCCACTAATTGTTTGGG | Ins_k3_46_GTA | 0.09818617745151292 | 0.036416952577166095 | 54.16555621535751 |
| GATCCAGATCTAACTTGGGGTGGCTTTGTCTTCTTCTTTTGCCAATGTCCACTAATTGTTTGGG | Ins_k1_46_G | 0.06376038999904465 | 0.033340810301054 | 52.33259586344066 |
| GATCCAGATCTAACTTGGGGTGGCTTTGTCTTCTTCTTTTGCCAATGTCTCCACTAATTGTTTGGG | Ins_k3_46_GTC | 0.48746998974175004 | 0.03068761735245374 | 79.94421065133744 |
| GATCCAGATCTAACTTGGGGTGGCTTTGTCTTCTTCTTTTGCCAATGTGTCCACTAATTGTTTGGG | Ins_k3_46_GTG | 0.16891284520643157 | 0.03739049804647873 | 58.135231836234354 |
| GATCCAGATCTAACTTGGGGTGGCTTTGTCTTCTTCTTTTGCCAATGTTCCACTAATTGTTTGGG | Ins_k2_45_TG;Ins_k2_46_GT | 0.32332098220222716 | 0.03965179800057636 | 67.84190127785467 |
| GATCCAGATCTAACTTGGGGTGGCTTTGTCTTCTTCTTTTGCCAATGTTTCCACTAATTGTTTGGG | Ins_k3_45_TGT;Ins_k3_46_GTT | -0.33242512195438695 | 0.04173694243506503 | 35.21365427211837 |
| GATCCAGATCTAACTTGGGGTGGCTTTGTCTTCTTCTTTTGCCAATGTTTGGG | Del_k10_46-55;Del_k10_47-56 | -0.008514650734661955 | 0.0319097481762489 | 48.683705465295674 |
| GATCCAGATCTAACTTGGGGTGGCTTTGTCTTCTTCTTTTGCCAATT | Del_k16_48-63 | -4.762263051631027 | 0.17534225945578716 | 0.4196207227218122 |
| GATCCAGATCTAACTTGGGGTGGCTTTGTCTTCTTCTTTTGCCAATTAAACCACTAATTGTTTGGG | Ins_k3_47_AAA | -0.4663949754749632 | 0.04528373524337084 | 30.798440615332936 |

| | | | | |
|---|---|---|---|---|
| GATCCAGATCTAACTTGGGGTGGCTTTGTCTTCTTCTTTTGCCAATTAACCACTAATTGTTTGGG | Ins_k2_47_AA | -0.3608106824832432 | 0.05133718612850544 | 34.22814819939483 |
| GATCCAGATCTAACTTGGGGTGGCTTTGTCTTCTTCTTTTGCCAATTAACCCACTAATTGTTTGGG | Ins_k3_47_AAC | -1.2192450205830845 | 0.05892107517370814 | 14.506749359802896 |
| GATCCAGATCTAACTTGGGGTGGCTTTGTCTTCTTCTTTTGCCAATTAAGCCACTAATTGTTTGGG | Ins_k3_47_AAG | -0.9381604432274746 | 0.059343106619966515 | 19.215141568113754 |
| GATCCAGATCTAACTTGGGGTGGCTTTGTCTTCTTCTTTTGCCAATTAATCCACTAATTGTTTGGG | Ins_k3_46_TAA;Ins_k3_47_AAT | -0.37025788046533625 | 0.04546240238132412 | 33.906310731583496 |
| GATCCAGATCTAACTTGGGGTGGCTTTGTCTTCTTCTTTTGCCAATTAATTGTTTGGG | Del_k5_47-51;Del_k5_48-52 | -1.125532867666663 | 0.043201680071455195 | 15.931944267004729 |
| GATCCAGATCTAACTTGGGGTGGCTTTGTCTTCTTCTTTTGCCAATTACACCACTAATTGTTTGGG | Ins_k3_47_ACA | 0.3494053146198832 | 0.035165659910690795 | 69.63479352663323 |
| GATCCAGATCTAACTTGGGGTGGCTTTGTCTTCTTCTTTTGCCAATTACACTAATTGTTTGGG | C48A | -1.5862754606347498 | 0.055285104716390955 | 10.050109944190812 |
| GATCCAGATCTAACTTGGGGTGGCTTTGTCTTCTTCTTTTGCCAATTACCACCACTAATTGTTTGGG | Ins_k4_47_ACCA | 0.49175585319447035 | 0.037520812572855305 | 80.2875759047243 |
| GATCCAGATCTAACTTGGGGTGGCTTTGTCTTCTTCTTTTGCCAATTACCACTAATTGTTTGGG | Ins_k1_47_A | -0.14824098938936703 | 0.04021705959280432 | 42.33516418886297 |
| GATCCAGATCTAACTTGGGGTGGCTTTGTCTTCTTCTTTTGCCAATTACCCACTAATTGTTTGGG | Ins_k2_47_AC | -0.757284224092597 | 0.05563899309585126 | 23.024867113127307 |
| GATCCAGATCTAACTTGGGGTGGCTTTGTCTTCTTCTTTTGCCAATTACCCCACTAATTGTTTGGG | Ins_k3_47_ACC | -4.446200472342686 | 0.2757681914469277 | 0.5756004987154322 |
| GATCCAGATCTAACTTGGGGTGGCTTTGTCTTCTTCTTTTGCCAATTACGCCACTAATTGTTTGGG | Ins_k3_47_ACG | 0.35497324946458586 | 0.031236172830803162 | 70.02359693145678 |
| GATCCAGATCTAACTTGGGGTGGCTTTGTCTTCTTCTTTTGCCAATTACTAATTGTTTGGG | Del_k2_48-49 | -2.362505131736084 | 0.34569253539871075 | 4.624433620644864 |
| GATCCAGATCTAACTTGGGGTGGCTTTGTCTTCTTCTTTTGCCAATTACTCCACTAATTGTTTGGG | Ins_k3_46_TAC;Ins_k3_47_ACT | 0.07266545199641095 | 0.03849461532508045 | 52.80070203793775 |
| GATCCAGATCTAACTTGGGGTGGCTTTGTCTTCTTCTTTTGCCAATTAGACCACTAATTGTTTGGG | Ins_k3_47_AGA | 0.22523907346042282 | 0.03604225418532719 | 61.50374752125171 |
| GATCCAGATCTAACTTGGGGTGGCTTTGTCTTCTTCTTTTGCCAATTAGCCACTAATTGTTTGGG | Ins_k2_47_AG | -1.2461650518911276 | 0.058610371228770775 | 14.121436795017143 |
| GATCCAGATCTAACTTGGGGTGGCTTTGTCTTCTTCTTTTGCCAATTAGCCCACTAATTGTTTGGG | Ins_k3_47_AGC | -1.2506237953544692 | 0.059176351736135155 | 14.05861309248084 |
| GATCCAGATCTAACTTGGGGTGGCTTTGTCTTCTTCTTTTGCCAATTAGGCCACTAATTGTTTGGG | Ins_k3_47_AGG | -1.7529633221442888 | 0.07007645825765373 | 8.5070540933133465 |
| GATCCAGATCTAACTTGGGGTGGCTTTGTCTTCTTCTTTTGCCAATTAGTCCACTAATTGTTTGGG | Ins_k3_46_TAG;Ins_k3_47_AGT | -0.5629421430950542 | 0.042621027397189065 | 27.96396988434531 |
| GATCCAGATCTAACTTGGGGTGGCTTTGTCTTCTTCTTTTGCCAATTATACCACTAATTGTTTGGG | Ins_k3_47_ATA | -0.5692454748845995 | 0.044289794036440205 | 27.788258070723035 |
| GATCCAGATCTAACTTGGGGTGGCTTTGTCTTCTTCTTTTGCCAATTATCCACTAATTGTTTGGG | Ins_k2_46_TA;Ins_k2_47_AT | 0.08777689541971989 | 0.0373379412679765 | 53.604656013101014 |
| GATCCAGATCTAACTTGGGGTGGCTTTGTCTTCTTCTTTTGCCAATTATCCCACTAATTGTTTGGG | Ins_k3_47_ATC | -1.4523296650154616 | 0.05687753406316106 | 11.490600689052021 |
| GATCCAGATCTAACTTGGGGTGGCTTTGTCTTCTTCTTTTGCCAATTATGCCACTAATTGTTTGGG | Ins_k3_47_ATG | -0.4600761470432788 | 0.040509378089585324 | 30.993666828442024 |
| GATCCAGATCTAACTTGGGGTGGCTTTGTCTTCTTCTTTTGCCAATTATTCCACTAATTGTTTGGG | Ins_k3_44_ATT;Ins_k3_45_TTA;In | -0.22184546449800613 | 0.039540139028289914 | 39.331021958441816 |
| GATCCAGATCTAACTTGGGGTGGCTTTGTCTTCTTCTTTTGCCAATTATTGTTTGGG | Del_k6_48-53 | -0.8205207985628914 | 0.0456521659890997 | 21.613934798368106 |
| GATCCAGATCTAACTTGGGGTGGCTTTGTCTTCTTCTTTTGCCAATTC | Del_k15_49-63 | -5.305954665148592 | 0.3477671353207762 | 0.24363218480362592 |
| GATCCAGATCTAACTTGGGGTGGCTTTGTCTTCTTCTTTTGCCAATTCAAAACACTAATTGTTTGGG | Ins_k4_48_AAAA | -0.8732595390836343 | 0.058957568088912296 | 20.503579808761145 |
| GATCCAGATCTAACTTGGGGTGGCTTTGTCTTCTTCTTTTGCCAATTCAAACACTAATTGTTTGGG | Ins_k3_48_AAA | -1.0093775349077285 | 0.05843456502710951 | 17.89428700000438 |
| GATCCAGATCTAACTTGGGGTGGCTTTGTCTTCTTCTTTTGCCAATTCAACACTAATTGTTTGGG | Ins_k2_48_AA | 0.3820178795693522 | 0.03335997805325691 | 71.94319968195717 |
| GATCCAGATCTAACTTGGGGTGGCTTTGTCTTCTTCTTTTGCCAATTCAACCACTAATTGTTTGGG | Ins_k3_47_CAA;Ins_k3_48_AAC | 0.4643936377892892 | 0.03241268681379395 | 78.12051298301434 |
| GATCCAGATCTAACTTGGGGTGGCTTTGTCTTCTTCTTTTGCCAATTCAACTAATTGTTTGGG | C49A | -0.03446363479775605 | 0.032333815662789075 | 47.4366624910666 |
| GATCCAGATCTAACTTGGGGTGGCTTTGTCTTCTTCTTTTGCCAATTCAAGACACTAATTGTTTGGG | Ins_k4_48_AAGA | 0.6311822845134945 | 0.03386764741843971 | 92.29973951178411 |
| GATCCAGATCTAACTTGGGGTGGCTTTGTCTTCTTCTTTTGCCAATTCAAGCACTAATTGTTTGGG | Ins_k3_48_AAG | 0.2277332362084895 | 0.03376795133654974 | 61.657339339058296 |
| GATCCAGATCTAACTTGGGGTGGCTTTGTCTTCTTCTTTTGCCAATTCAAGTCACTAATTGTTTGGG | Ins_k4_48_AAGT | 0.47345669273749813 | 0.03604415213408715 | 78.8317415683448 |
| GATCCAGATCTAACTTGGGGTGGCTTTGTCTTCTTCTTTTGCCAATTCAATCACTAATTGTTTGGG | Ins_k3_48_AAT | -0.6182388014320398 | 0.04475814756141267 | 26.459631538641286 |
| GATCCAGATCTAACTTGGGGTGGCTTTGTCTTCTTCTTTTGCCAATTCAATTGTTTGGG | Del_k4_49-52 | -0.46414400681243656 | 0.04178313277699981 | 30.86784502428234 |
| GATCCAGATCTAACTTGGGGTGGCTTTGTCTTCTTCTTTTGCCAATTCACACACTAATTGTTTGGG | Ins_k3_48_ACA | -0.6984574895067223 | 0.04368839516592782 | 24.419977450855768 |
| GATCCAGATCTAACTTGGGGTGGCTTTGTCTTCTTCTTTTGCCAATTCACACTAATTGTTTGGG | Ins_k1_48_A | -0.5625240156162313 | 0.042095013124896226 | 27.97566483339032 |
| GATCCAGATCTAACTTGGGGTGGCTTTGTCTTCTTCTTTTGCCAATTCACCACTAATTGTTTGGG | Ins_k2_47_CA;Ins_k2_48_AC | 0.46003495693841345 | 0.03280439454856278 | 77.78075159264924 |
| GATCCAGATCTAACTTGGGGTGGCTTTGTCTTCTTCTTTTGCCAATTCACCCACTAATTGTTTGGG | Ins_k3_47_CAC;Ins_k3_48_ACC | -0.07481183236609823 | 0.12937742561440654 | 45.560777435884816 |
| GATCCAGATCTAACTTGGGGTGGCTTTGTCTTCTTCTTTTGCCAATTCACGCACTAATTGTTTGGG | Ins_k3_48_ACG | 0.14503753281424148 | 0.03454067016855603 | 56.76367336058643 |
| GATCCAGATCTAACTTGGGGTGGCTTTGTCTTCTTCTTTTGCCAATTCACTAATTGTTTGGG | Del_k1_48-48;Del_k1_49-49 | -0.7393395992253013 | 0.03701060884122829 | 23.44176910386458 |
| GATCCAGATCTAACTTGGGGTGGCTTTGTCTTCTTCTTTTGCCAATTCACTCACTAATTGTTTGGG | Ins_k3_48_ACT | -0.6230715368492336 | 0.04998407299600641 | 26.332067629763483 |
| GATCCAGATCTAACTTGGGGTGGCTTTGTCTTCTTCTTTTGCCAATTCAGACACTAATTGTTTGGG | Ins_k3_48_AGA | -0.28560287305439824 | 0.043560311918654046 | 36.90164617153322 |
| GATCCAGATCTAACTTGGGGTGGCTTTGTCTTCTTCTTTTGCCAATTCAGCACTAATTGTTTGGG | Ins_k2_48_AG | 0.4709196305204968 | 0.033757911214892064 | 78.63199402842221 |
| GATCCAGATCTAACTTGGGGTGGCTTTGTCTTCTTCTTTTGCCAATTCAGCCACTAATTGTTTGGG | Ins_k3_47_CAG;Ins_k3_48_AGC | 0.08554436583397584 | 0.033658998789901806 | 53.485115521111695 |
| GATCCAGATCTAACTTGGGGTGGCTTTGTCTTCTTCTTTTGCCAATTCAGGCACTAATTGTTTGGG | Ins_k3_48_AGG | -1.354113659562217 | 0.06083404342487925 | 12.676442823634508 |
| GATCCAGATCTAACTTGGGGTGGCTTTGTCTTCTTCTTTTGCCAATTCAGTCACTAATTGTTTGGG | Ins_k3_48_AGT | -0.48214776278498556 | 0.04411127239136549 | 30.317080665388584 |
| GATCCAGATCTAACTTGGGGTGGCTTTGTCTTCTTCTTTTGCCAATTCATACACTAATTGTTTGGG | Ins_k3_48_ATA | -1.237217024808079 | 0.06152557521651185 | 14.248362815763098 |
| GATCCAGATCTAACTTGGGGTGGCTTTGTCTTCTTCTTTTGCCAATTCATCACTAATTGTTTGGG | Ins_k2_48_AT | 0.43820641123714377 | 0.032373313909717275 | 76.10130749826752 |
| GATCCAGATCTAACTTGGGGTGGCTTTGTCTTCTTCTTTTGCCAATTCATCCACTAATTGTTTGGG | Ins_k3_46_TCA;Ins_k3_47_CAT;I | 0.32522367123118956 | 0.03187833095565197 | 67.97110619853329 |
| GATCCAGATCTAACTTGGGGTGGCTTTGTCTTCTTCTTTTGCCAATTCATGCACTAATTGTTTGGG | Ins_k3_48_ATG | -0.8516637463044089 | 0.05499684484544903 | 20.951186686187807 |
| GATCCAGATCTAACTTGGGGTGGCTTTGTCTTCTTCTTTTGCCAATTCATTCACTAATTGTTTGGG | Ins_k3_48_ATT | -1.0534989234904102 | 0.0585888528680935 | 17.121930227817018 |
| GATCCAGATCTAACTTGGGGTGGCTTTGTCTTCTTCTTTTGCCAATTCATTGTTTGGG | Del_k5_49-53 | -0.2612903471168646 | 0.03387353054222079 | 37.8098135888704 |
| GATCCAGATCTAACTTGGGGTGGCTTTGTCTTCTTCTTTTGCCAATTCC | Del_k14_50-63 | -3.91191652710856 | 0.14474972473172346 | 0.9821045672768086 |
| GATCCAGATCTAACTTGGGGTGGCTTTGTCTTCTTCTTTTGCCAATTCCA | Del_k13_51-63 | -3.9352185213961843 | 0.13246738579997086 | 0.9594841462536334 |
| GATCCAGATCTAACTTGGGGTGGCTTTGTCTTCTTCTTTTGCCAATTCCAAAACTAATTGTTTGGG | Ins_k3_49_AAA;Ins_k3_50_AAA | -1.2058888375345338 | 0.0636012425046171 | 14.701803851841571 |
| GATCCAGATCTAACTTGGGGTGGCTTTGTCTTCTTCTTTTGCCAATTCCAAACCTAATTGTTTGGG | Ins_k3_50_AAC | -1.0003673478664117 | 0.07248300810007274 | 18.056246419475862 |
| GATCCAGATCTAACTTGGGGTGGCTTTGTCTTCTTCTTTTGCCAATTCCAAACTAATTGTTTGGG | Ins_k2_49_AA;Ins_k2_50_AA | -1.7364435897666946 | 0.0925014781468639 | 8.648755562902842 |
| GATCCAGATCTAACTTGGGGTGGCTTTGTCTTCTTCTTTTGCCAATTCCAAAGCTAATTGTTTGGG | Ins_k3_50_AAG | -0.4289582271113411 | 0.04706594721165338 | 31.973288111497144 |
| GATCCAGATCTAACTTGGGGTGGCTTTGTCTTCTTCTTTTGCCAATTCCAAATCTAATTGTTTGGG | Ins_k3_50_AAT | -1.5214072688025877 | 0.07467867341558641 | 10.723651964921308 |
| GATCCAGATCTAACTTGGGGTGGCTTTGTCTTCTTCTTTTGCCAATTCCAAATTGTTTGGG | Del_k2_51-52 | -0.4809511573448295 | 0.040249716725620235 | 30.353379962679437 |
| GATCCAGATCTAACTTGGGGTGGCTTTGTCTTCTTCTTTTGCCAATTCCAACACTAATTGTTTGGG | Ins_k3_48_CAA;Ins_k3_49_AAC; | 0.06292765395161926 | 0.03570774458438952 | 52.289034764375835 |
| GATCCAGATCTAACTTGGGGTGGCTTTGTCTTCTTCTTTTGCCAATTCCAACCCTAATTGTTTGGG | Ins_k3_50_ACC | -0.7644593022858888 | 0.0504605694322319 | 22.86025315642323 |
| GATCCAGATCTAACTTGGGGTGGCTTTGTCTTCTTCTTTTGCCAATTCCAACCTAATTGTTTGGG | Ins_k2_50_AC | 0.2823597556328348 | 0.04145240043994935 | 65.11915792172988 |
| GATCCAGATCTAACTTGGGGTGGCTTTGTCTTCTTCTTTTGCCAATTCCAACGCTAATTGTTTGGG | Ins_k3_50_ACG | 0.6247230135520045 | 0.033186071673167075 | 91.70547181880613 |
| GATCCAGATCTAACTTGGGGTGGCTTTGTCTTCTTCTTTTGCCAATTCCAACTAATTGTTTGGG | Ins_k1_49_A;Ins_k1_50_A | 0.29084866672195875 | 0.03293702149735752 | 65.67430161253411 |
| GATCCAGATCTAACTTGGGGTGGCTTTGTCTTCTTCTTTTGCCAATTCCAACTCCTAATTGTTTGGG | Ins_k4_50_ACTC | 0.235815834062987 | 0.03436468065542657 | 62.157710242075375 |
| GATCCAGATCTAACTTGGGGTGGCTTTGTCTTCTTCTTTTGCCAATTCCAACTCTAATTGTTTGGG | Ins_k3_50_ACT | 0.10485998455169931 | 0.035061145319437846 | 54.528255635787445 |
| GATCCAGATCTAACTTGGGGTGGCTTTGTCTTCTTCTTTTGCCAATTCCAAGACTAATTGTTTGGG | Ins_k3_49_AAG;Ins_k3_50_AGA | 0.029260020984033352 | 0.03316282196577636 | 50.55789199257102 |
| GATCCAGATCTAACTTGGGGTGGCTTTGTCTTCTTCTTTTGCCAATTCCAAGCCTAATTGTTTGGG | Ins_k3_50_AGC | 0.130879814528988 | 0.036830317598116495 | 55.965691395222144 |
| GATCCAGATCTAACTTGGGGTGGCTTTGTCTTCTTCTTTTGCCAATTCCAAGCTAATTGTTTGGG | Ins_k2_50_AG | -0.19727857973888369 | 0.04301619778404394 | 40.30922919682945 |
| GATCCAGATCTAACTTGGGGTGGCTTTGTCTTCTTCTTTTGCCAATTCCAAGGACTAATTGTTTGGG | Ins_k4_49_AAGG;Ins_k4_50_AG | 0.6114104543824961 | 0.03277446432096799 | 90.4927275743922 |
| GATCCAGATCTAACTTGGGGTGGCTTTGTCTTCTTCTTTTGCCAATTCCAAGGCTAATTGTTTGGG | Ins_k3_50_AGG | 0.04899339376761913 | 0.03816626459993589 | 51.56547856667592 |
| GATCCAGATCTAACTTGGGGTGGCTTTGTCTTCTTCTTTTGCCAATTCCAAGTCTAATTGTTTGGG | Ins_k3_50_AGT | -1.107723536810607 | 0.06210034572740696 | 16.218223185015624 |

| | | | | |
|---|---|---|---|---|
| GATCCAGATCTAACTTGGGGTGGCTTTGTCTTCTTCTTTTGCCAATTCCAATAATTGTTTGGG | C51A | -1.1823955305448 | 0.05603783241156366 | 15.0512870265929 |
| GATCCAGATCTAACTTGGGGTGGCTTTGTCTTCTTCTTTTGCCAATTCCAATACTAATTGTTTGGG | Ins_k3_49_AAT;Ins_k3_50_ATA | -1.7712555618614665 | 0.08319567546641853 | 8.35285563772874 |
| GATCCAGATCTAACTTGGGGTGGCTTTGTCTTCTTCTTTTGCCAATTCCAATCCTAATTGTTTGGG | Ins_k3_50_ATC | -0.6870285295994552 | 0.05748986533445057 | 24.7006733698856 |
| GATCCAGATCTAACTTGGGGTGGCTTTGTCTTCTTCTTTTGCCAATTCCAATCTAATTGTTTGGG | Ins_k2_50_AT | -1.087356652145096 | 0.07085314967090024 | 16.551924559667214 |
| GATCCAGATCTAACTTGGGGTGGCTTTGTCTTCTTCTTTTGCCAATTCCAATGCTAATTGTTTGGG | Ins_k3_50_ATG | -0.9913621250977167 | 0.05777275514838025 | 18.219581270420303 |
| GATCCAGATCTAACTTGGGGTGGCTTTGTCTTCTTCTTTTGCCAATTCCAATTCTAATTGTTTGGG | Ins_k3_50_ATT | -1.9335661198832603 | 0.0915044669186458 | 7.10140686854405 |
| GATCCAGATCTAACTTGGGGTGGCTTTGTCTTCTTCTTTTGCCAATTCCAATTGTTTGGG | Del_k3_50-52;Del_k3_51-53 | -0.5919114888011211 | 0.03643133018515486 | 27.165493483760034 |
| GATCCAGATCTAACTTGGGGTGGCTTTGTCTTCTTCTTTTGCCAATTCCAATTTCTAATTGTTTGGG | Ins_k4_50_ATTT | -2.6617423260851645 | 0.18330469928252496 | 3.4284789524106336 |
| GATCCAGATCTAACTTGGGGTGGCTTTGTCTTCTTCTTTTGCCAATTCCAC | Del_k12_52-63 | -1.393897913103543 | 0.040148182541765554 | 12.182020335218082 |
| GATCCAGATCTAACTTGGGGTGGCTTTGTCTTCTTCTTTTGCCAATTCCACAAATAATTGTTTGGG | Ins_k3_51_AAA | 0.2863648653270544 | 0.04242880423703539 | 65.38049027543941 |
| GATCCAGATCTAACTTGGGGTGGCTTTGTCTTCTTCTTTTGCCAATTCCACAAATTGTTTGGG | T52A | 0.13153791736696152 | 0.03174585957728562 | 56.00253469756967 |
| GATCCAGATCTAACTTGGGGTGGCTTTGTCTTCTTCTTTTGCCAATTCCACAACGTAATTGTTTGGG | Ins_k4_51_AACG | 0.7081736671248293 | 0.032436210537698074 | 99.68674326969058 |
| GATCCAGATCTAACTTGGGGTGGCTTTGTCTTCTTCTTTTGCCAATTCCACAACTAATTGTTTGGG | Ins_k3_49_ACA;Ins_k3_50_CAA; | 0.37139083484334423 | 0.035108439159111396 | 71.18270414837136 |
| GATCCAGATCTAACTTGGGGTGGCTTTGTCTTCTTCTTTTGCCAATTCCACAAGTAATTGTTTGGG | Ins_k3_51_AAG | 0.3165723054593612 | 0.03556100200724162 | 67.38559966439192 |
| GATCCAGATCTAACTTGGGGTGGCTTTGTCTTCTTCTTTTGCCAATTCCACAATAATTGTTTGGG | Ins_k2_51_AA | 0.3563276636411771 | 0.039874182917235025 | 70.11850213980833 |
| GATCCAGATCTAACTTGGGGTGGCTTTGTCTTCTTCTTTTGCCAATTCCACAATTAATTGTTTGGG | Ins_k3_51_AAT | 0.1766318822742856 | 0.04154162589118386 | 58.58571626193117 |
| GATCCAGATCTAACTTGGGGTGGCTTTGTCTTCTTCTTTTGCCAATTCCACAATTGTTTGGG | Del_k1_52-52 | 0.28468184075449066 | 0.03084276040014967 | 65.27054584927713 |
| GATCCAGATCTAACTTGGGGTGGCTTTGTCTTCTTCTTTTGCCAATTCCACACATAATTGTTTGGG | Ins_k3_51_ACA | 0.37448119510941824 | 0.03584072248861367 | 71.40302460835652 |
| GATCCAGATCTAACTTGGGGTGGCTTTGTCTTCTTCTTTTGCCAATTCCACACCTAATTGTTTGGG | Ins_k3_50_CAC;Ins_k3_51_ACC | 0.29585532508314094 | 0.0340361941665805 | 66.00393489586843 |
| GATCCAGATCTAACTTGGGGTGGCTTTGTCTTCTTCTTTTGCCAATTCCACACGTAATTGTTTGGG | Ins_k3_51_ACG | 0.39300358799895474 | 0.03232337304099971 | 72.73790389912183 |
| GATCCAGATCTAACTTGGGGTGGCTTTGTCTTCTTCTTTTGCCAATTCCACACTAATTGTTTGGG | Ins_k2_48_CA;Ins_k2_49_AC;Ins | 0.26876264859358256 | 0.0403652575298353 | 64.23971822827457 |
| GATCCAGATCTAACTTGGGGTGGCTTTGTCTTCTTCTTTTGCCAATTCCACACTTAATTGTTTGGG | Ins_k3_51_ACT | -0.028481099103500496 | 0.046455625224726446 | 47.72130460949053 |
| GATCCAGATCTAACTTGGGGTGGCTTTGTCTTCTTCTTTTGCCAATTCCACAGATAATTGTTTGGG | Ins_k3_51_AGA | -0.06440271830464939 | 0.03677914616334935 | 46.03750159863572 |
| GATCCAGATCTAACTTGGGGTGGCTTTGTCTTCTTCTTTTGCCAATTCCACAGCTAATTGTTTGGG | Ins_k3_50_CAG;Ins_k3_51_AGC | -0.12957458402380512 | 0.03962050517599205 | 43.132831151160204 |
| GATCCAGATCTAACTTGGGGTGGCTTTGTCTTCTTCTTTTGCCAATTCCACAGGTAATTGTTTGGG | Ins_k3_51_AGG | -1.6001913828384855 | 0.07513622785137969 | 9.91122201421296 |
| GATCCAGATCTAACTTGGGGTGGCTTTGTCTTCTTCTTTTGCCAATTCCACAGGTTAATTGTTTGGG | Ins_k4_51_AGGT | -3.553159449273151 | 0.22714868915502515 | 1.4059308154494499 |
| GATCCAGATCTAACTTGGGGTGGCTTTGTCTTCTTCTTTTGCCAATTCCACAGTAATTGTTTGGG | Ins_k2_51_AG | 0.08590301005759504 | 0.042934313738775036 | 53.504301089033845 |
| GATCCAGATCTAACTTGGGGTGGCTTTGTCTTCTTCTTTTGCCAATTCCACAGTATAATTGTTTGGG | Ins_k4_51_AGTA | -0.8103062204563861 | 0.04764822048938109 | 21.835843445712293 |
| GATCCAGATCTAACTTGGGGTGGCTTTGTCTTCTTCTTTTGCCAATTCCACAGTTAATTGTTTGGG | Ins_k3_51_AGT | -0.25823684833304905 | 0.04298569541560196 | 37.92544225483369 |
| GATCCAGATCTAACTTGGGGTGGCTTTGTCTTCTTCTTTTGCCAATTCCACATAATTGTTTGGG | Ins_k1_51_A | 0.43849232211004807 | 0.03894622334633857 | 76.12306880027168 |
| GATCCAGATCTAACTTGGGGTGGCTTTGTCTTCTTCTTTTGCCAATTCCACATATAATTGTTTGGG | Ins_k3_51_ATA | 0.07296484595762909 | 0.04230536189545721 | 52.81651261595367 |
| GATCCAGATCTAACTTGGGGTGGCTTTGTCTTCTTCTTTTGCCAATTCCACATCGTAATTGTTTGGG | Ins_k4_51_ATCG | 0.680115491086946 | 0.035617367016232124 | 96.92859039437462 |
| GATCCAGATCTAACTTGGGGTGGCTTTGTCTTCTTCTTTTGCCAATTCCACATCTAATTGTTTGGG | Ins_k3_50_CAT;Ins_k3_51_ATC | 0.392505185877716 | 0.03281867410028445 | 72.70166020624102 |
| GATCCAGATCTAACTTGGGGTGGCTTTGTCTTCTTCTTTTGCCAATTCCACATGTAATTGTTTGGG | Ins_k3_51_ATG | 0.03495939239511514 | 0.041431807524907376 | 50.84686289082037 |
| GATCCAGATCTAACTTGGGGTGGCTTTGTCTTCTTCTTTTGCCAATTCCACATTAATTGTTTGGG | Ins_k2_51_AT | -0.05497976906655733 | 0.03919221886747846 | 46.4733609544779 |
| GATCCAGATCTAACTTGGGGTGGCTTTGTCTTCTTCTTTTGCCAATTCCACATTGTTTGGG | Del_k2_52-53 | 0.09837689835383456 | 0.03029979976390572 | 54.175887704297786 |
| GATCCAGATCTAACTTGGGGTGGCTTTGTCTTCTTCTTTTGCCAATTCCACATTTAATTGTTTGGG | Ins_k3_51_ATT | -0.18584803038937253 | 0.04124105790219597 | 40.77262924341003 |
| GATCCAGATCTAACTTGGGGTGGCTTTGTCTTCTTCTTTTGCCAATTCCACCAATAATTGTTTGGG | Ins_k3_51_CAA | 0.4953621237301904 | 0.19104917051148393 | 80.57763732972063 |
| GATCCAGATCTAACTTGGGGTGGCTTTGTCTTCTTCTTTTGCCAATTCCACCAATTGTTTGGG | T52C | 0.08143733094430905 | 0.026913286253860837 | 53.26590075507282 |
| GATCCAGATCTAACTTGGGGTGGCTTTGTCTTCTTCTTTTGCCAATTCCACCACCTAATTGTTTGGG | Ins_k4_50_CCAC;Ins_k4_51_CA( | 0.4534153642108978 | 0.03889340143952335 | 77.26757507873133 |
| GATCCAGATCTAACTTGGGGTGGCTTTGTCTTCTTCTTTTGCCAATTCCACCACTAATTGTTTGGG | Ins_k3_47_CCA;Ins_k3_48_CAC; | 0.10131361860557742 | 0.03314794166660807 | 54.33522097498472 |
| GATCCAGATCTAACTTGGGGTGGCTTTGTCTTCTTCTTTTGCCAATTCCACCAGTAATTGTTTGGG | Ins_k3_51_CAG | 0.4670611606267505 | 0.03505893592008832 | 78.32917942298414 |
| GATCCAGATCTAACTTGGGGTGGCTTTGTCTTCTTCTTTTGCCAATTCCACCATAATTGTTTGGG | Ins_k2_51_CA | 0.4267655722100223 | 0.03391333095729981 | 75.23560630623516 |
| GATCCAGATCTAACTTGGGGTGGCTTTGTCTTCTTCTTTTGCCAATTCCACCATTAATTGTTTGGG | Ins_k3_51_CAT | 0.24958970366401345 | 0.035916962562031876 | 63.01978586693417 |
| GATCCAGATCTAACTTGGGGTGGCTTTGTCTTCTTCTTTTGCCAATTCCACCATTTAATTGTTTGGG | Ins_k4_51_CATT | 0.2832651735203735 | 0.03631116586127578 | 65.17814467193337 |
| GATCCAGATCTAACTTGGGGTGGCTTTGTCTTCTTCTTTTGCCAATTCCACCCATAATTGTTTGGG | Ins_k3_51_CCA | -1.5516252233595487 | 0.07197003389517621 | 10.404452207769713 |
| GATCCAGATCTAACTTGGGGTGGCTTTGTCTTCTTCTTTTGCCAATTCCACCCCTAATTGTTTGGG | Ins_k3_50_CCC;Ins_k3_51_CCC | -4.45262785789825 | 0.3242334527335101 | 0.5719127563506565 |
| GATCCAGATCTAACTTGGGGTGGCTTTGTCTTCTTCTTTTGCCAATTCCACCCGTAATTGTTTGGG | Ins_k3_51_CCG | -0.6161617272074734 | 0.05819505816729487 | 26.51464727340479 |
| GATCCAGATCTAACTTGGGGTGGCTTTGTCTTCTTCTTTTGCCAATTCCACCCTAATTGTTTGGG | Ins_k2_50_CC;Ins_k2_51_CC | -2.368214254978356 | 0.11924390687778318 | 4.598107380560083 |
| GATCCAGATCTAACTTGGGGTGGCTTTGTCTTCTTCTTTTGCCAATTCCACCCTTAATTGTTTGGG | Ins_k3_51_CCT | -2.509945973602714 | 0.11451254933199713 | 3.9904861324932264 |
| GATCCAGATCTAACTTGGGGTGGCTTTGTCTTCTTCTTTTGCCAATTCCACCGAGTAATTGTTTGGG | Ins_k4_51_CGAG | 0.8267446631051902 | 0.033530423990396685 | 100 |
| GATCCAGATCTAACTTGGGGTGGCTTTGTCTTCTTCTTTTGCCAATTCCACCGATAATTGTTTGGG | Ins_k3_51_CGA | 0.5100631041542434 | 0.031281122549727695 | 81.77095759615575 |
| GATCCAGATCTAACTTGGGGTGGCTTTGTCTTCTTCTTTTGCCAATTCCACCGCTAATTGTTTGGG | Ins_k3_50_CCG;Ins_k3_51_CGC | 0.5250581437799078 | 0.0346441509031166 | 83.00635561837838 |
| GATCCAGATCTAACTTGGGGTGGCTTTGTCTTCTTCTTTTGCCAATTCCACCGGTAATTGTTTGGG | Ins_k3_51_CGG | 0.6819299043478428 | 0.037226997921174634 | 97.10461855977321 |
| GATCCAGATCTAACTTGGGGTGGCTTTGTCTTCTTCTTTTGCCAATTCCACCGTAATTGTTTGGG | Ins_k2_51_CG | 0.6429537083161616 | 0.030386057483863752 | 93.39265885068254 |
| GATCCAGATCTAACTTGGGGTGGCTTTGTCTTCTTCTTTTGCCAATTCCACCGTTAATTGTTTGGG | Ins_k3_51_CGT | 0.3598314112743939 | 0.03643760356267171 | 70.36461057487487 |
| GATCCAGATCTAACTTGGGGTGGCTTTGTCTTCTTCTTTTGCCAATTCCACCTAATTGTTTGGG | Ins_k1_50_C;Ins_k1_51_C | 0.23413214907453284 | 0.03572770108129399 | 62.05314429118693 |
| GATCCAGATCTAACTTGGGGTGGCTTTGTCTTCTTCTTTTGCCAATTCCACCTATAATTGTTTGGG | Ins_k3_51_CTA | -0.4076791420884056 | 0.040219728207864854 | 32.660940790393795 |
| GATCCAGATCTAACTTGGGGTGGCTTTGTCTTCTTCTTTTGCCAATTCCACCTCTAATTGTTTGGG | Ins_k3_50_CCT;Ins_k3_51_CTC | -0.9373013206976931 | 0.05062097672435491 | 19.23165682244559 |
| GATCCAGATCTAACTTGGGGTGGCTTTGTCTTCTTCTTTTGCCAATTCCACCTGTAATTGTTTGGG | Ins_k3_51_CTG | 0.20763291146524732 | 0.03668181452652574 | 60.43037925335904 |
| GATCCAGATCTAACTTGGGGTGGCTTTGTCTTCTTCTTTTGCCAATTCCACCTTAATTGTTTGGG | Ins_k2_51_CT | 0.09086262709259274 | 0.03798457683009285 | 53.77032106547208 |
| GATCCAGATCTAACTTGGGGTGGCTTTGTCTTCTTCTTTTGCCAATTCCACCTTTAATTGTTTGGG | Ins_k3_51_CTT | -0.16585512900195537 | 0.0375376403776644 | 41.5959957156344 |
| GATCCAGATCTAACTTGGGGTGGCTTTGTCTTCTTCTTTTGCCAATTCCACCTTTTAATTGTTTGGG | Ins_k4_51_CTTT | -0.7465630739464536 | 0.05661864024388564 | 23.273048186598412 |
| GATCCAGATCTAACTTGGGGTGGCTTTGTCTTCTTCTTTTGCCAATTCCACG | Del_k11_52-62 | 0.06593916826903168 | 0.12261864823165167 | 52.44674128975243 |
| GATCCAGATCTAACTTGGGGTGGCTTTGTCTTCTTCTTTTGCCAATTCCACGAATAATTGTTTGGG | Ins_k3_51_GAA | 0.6516206329491894 | 0.030691163622555594 | 94.20560376363564 |
| GATCCAGATCTAACTTGGGGTGGCTTTGTCTTCTTCTTTTGCCAATTCCACGAATTGTTTGGG | T52G | 0.5784777824576048 | 0.028793008628830455 | 87.56109841236481 |
| GATCCAGATCTAACTTGGGGTGGCTTTGTCTTCTTCTTTTGCCAATTCCACGACTAATTGTTTGGG | Ins_k3_49_ACG;Ins_k3_50_CGA | 0.7207580908540427 | 0.02929605821028325 | 100 |
| GATCCAGATCTAACTTGGGGTGGCTTTGTCTTCTTCTTTTGCCAATTCCACGAGTAATTGTTTGGG | Ins_k3_51_GAG | 0.9167288321476985 | 0.03847634886396077 | 100 |
| GATCCAGATCTAACTTGGGGTGGCTTTGTCTTCTTCTTTTGCCAATTCCACGATAATTGTTTGGG | Ins_k2_51_GA | 0.8200941616473789 | 0.03577602919952366 | 100 |
| GATCCAGATCTAACTTGGGGTGGCTTTGTCTTCTTCTTTTGCCAATTCCACGATTAATTGTTTGGG | Ins_k3_51_GAT | 0.4899261081996733 | 0.034139649505102336 | 80.14080443270024 |
| GATCCAGATCTAACTTGGGGTGGCTTTGTCTTCTTCTTTTGCCAATTCCACGCATAATTGTTTGGG | Ins_k3_51_GCA | 0.654679353196963 | 0.031064368238425327 | 94.4941934839334 |
| GATCCAGATCTAACTTGGGGTGGCTTTGTCTTCTTCTTTTGCCAATTCCACGCCTAATTGTTTGGG | Ins_k3_50_CGC;Ins_k3_51_GCC | 0.7863050489248986 | 0.03126000678167561 | 100 |
| GATCCAGATCTAACTTGGGGTGGCTTTGTCTTCTTCTTTTGCCAATTCCACGCGTAATTGTTTGGG | Ins_k3_51_GCG | 0.8919533435377653 | 0.03361122662966691 | 100 |
| GATCCAGATCTAACTTGGGGTGGCTTTGTCTTCTTCTTTTGCCAATTCCACGCTAATTGTTTGGG | Ins_k2_50_CG;Ins_k2_51_GC | 0.6460181037292028 | 0.03179018727406683 | 93.6792898371152 |
| GATCCAGATCTAACTTGGGGTGGCTTTGTCTTCTTCTTTTGCCAATTCCACGCTTAATTGTTTGGG | Ins_k3_51_GCT | 0.5437645126915029 | 0.03203075168972075 | 84.57371724591968 |

| | | | | |
|---|---|---|---|---|
| **GATCCAGATCTAACTTGGGGTGGCTTTGTCTTCTTCTTTTGCCAATTCCACGG** | Del_k10_52-61 | -0.16628913463179107 | 0.028438472943945806 | 41.57794673627783 |
| **GATCCAGATCTAACTTGGGGTGGCTTTGTCTTCTTCTTTTGCCAATTCCACGGATAATTGTTTGGG** | Ins_k3_51_GGA | 0.8339515871934569 | 0.03445207329226869 | 100 |
| **GATCCAGATCTAACTTGGGGTGGCTTTGTCTTCTTCTTTTGCCAATTCCACGGCTAATTGTTTGGG** | Ins_k3_50_CGG;Ins_k3_51_GGC | 0.6518746651771967 | 0.032040959079109775 | 94.22953806296347 |
| **GATCCAGATCTAACTTGGGGTGGCTTTGTCTTCTTCTTTTGCCAATTCCACGGG** | Del_k9_52-60 | -0.19379407996626946 | 0.030855859335686683 | 40.44993169334423 |
| **GATCCAGATCTAACTTGGGGTGGCTTTGTCTTCTTCTTTTGCCAATTCCACGGGTAATTGTTTGGG** | Ins_k3_51_GGG | 0.48685328464591104 | 0.03654924193263954 | 79.89492384851924 |
| **GATCCAGATCTAACTTGGGGTGGCTTTGTCTTCTTCTTTTGCCAATTCCACGGTAATTGTTTGGG** | Ins_k2_51_GG | 0.5529229381372158 | 0.03297797093278615 | 85.3518370673297 |
| **GATCCAGATCTAACTTGGGGTGGCTTTGTCTTCTTCTTTTGCCAATTCCACGGTTAATTGTTTGGG** | Ins_k3_51_GGT | 0.48434654566352386 | 0.031049288535862706 | 79.69489893827533 |
| **GATCCAGATCTAACTTGGGGTGGCTTTGTCTTCTTCTTTTGCCAATTCCACGTAATTGTTTGGG** | Ins_k1_51_G | 0.5436461393310423 | 0.03608813775488515 | 84.56370656331339 |
| **GATCCAGATCTAACTTGGGGTGGCTTTGTCTTCTTCTTTTGCCAATTCCACGTATAATTGTTTGGG** | Ins_k3_51_GTA | -0.014219614373558676 | 0.03904086096636579 | 48.4067574361703 |
| **GATCCAGATCTAACTTGGGGTGGCTTTGTCTTCTTCTTTTGCCAATTCCACGTCTAATTGTTTGGG** | Ins_k3_50_CGT;Ins_k3_51_GTC | 0.5702650688163073 | 0.03077746686299139 | 86.84492905728695 |
| **GATCCAGATCTAACTTGGGGTGGCTTTGTCTTCTTCTTTTGCCAATTCCACGTGTAATTGTTTGGG** | Ins_k3_51_GTG | 1.019058190276467 | 0.20601111515990944 | 100 |
| **GATCCAGATCTAACTTGGGGTGGCTTTGTCTTCTTCTTTTGCCAATTCCACGTTAATTGTTTGGG** | Ins_k2_51_GT | 0.5966192404377919 | 0.03518786821055387 | 89.16408065418695 |
| **GATCCAGATCTAACTTGGGGTGGCTTTGTCTTCTTCTTTTGCCAATTCCACGTTTAATTGTTTGGG** | Ins_k3_51_GTT | 0.15230656869078008 | 0.036699157882325424 | 57.17779384358234 |
| **GATCCAGATCTAACTTGGGGTGGCTTTGTCTTCTTCTTTTGCCAATTCCACGTTTGGG** | Del_k5_52-56 | 0.2007490274635475 | 0.03275757046227472 | 60.01581208566644 |
| **GATCCAGATCTAACTTGGGGTGGCTTTGTCTTCTTCTTTTGCCAATTCCACT** | Del_k11_53-63 | -4.957287096412683 | 0.2611478017490324 | 0.34527017184831726 |
| **GATCCAGATCTAACTTGGGGTGGCTTTGTCTTCTTCTTTTGCCAATTCCACTA** | Del_k10_54-63 | -4.968024123972338 | 0.2894130143430394 | 0.3415828274808785 |
| **GATCCAGATCTAACTTGGGGTGGCTTTGTCTTCTTCTTTTGCCAATTCCACTAA** | Del_k9_55-63 | -3.856502679145198 | 0.15053198421455263 | 1.0380628745103837 |
| **GATCCAGATCTAACTTGGGGTGGCTTTGTCTTCTTCTTTTGCCAATTCCACTAAAAATTGTTTGGG** | Ins_k3_52_AAA;Ins_k3_53_AAA;I | 0.09373691811781482 | 0.03933579267113847 | 53.92509494276111 |
| **GATCCAGATCTAACTTGGGGTGGCTTTGTCTTCTTCTTTTGCCAATTCCACTAAAACTTGTTTGGG** | Ins_k3_54_AAC | 0.007916160720251919 | 0.03725804072165191 | 49.49022599941859 |
| **GATCCAGATCTAACTTGGGGTGGCTTTGTCTTCTTCTTTTGCCAATTCCACTAAAAGTTGTTTGGG** | Ins_k3_54_AAG | 0.17379006660544305 | 0.03784981780418318 | 58.41946279822414 |
| **GATCCAGATCTAACTTGGGGTGGCTTTGTCTTCTTCTTTTGCCAATTCCACTAAAATTGTTTGGG** | Ins_k2_52_AA;Ins_k2_53_AA;Ins_ | 0.10184351163527838 | 0.2388028649512735 | 54.36402045949581 |
| **GATCCAGATCTAACTTGGGGTGGCTTTGTCTTCTTCTTTTGCCAATTCCACTAAAATTTGTTTGGG** | Ins_k3_54_AAT | -0.11012029086588859 | 0.04986326884866193 | 43.980165312685266 |
| **GATCCAGATCTAACTTGGGGTGGCTTTGTCTTCTTCTTTTGCCAATTCCACTAAACATTGTTTGGG** | Ins_k3_53_AAC;Ins_k3_54_ACA | 0.05180343173244395 | 0.038239014636296496 | 51.71058329854266 |
| **GATCCAGATCTAACTTGGGGTGGCTTTGTCTTCTTCTTTTGCCAATTCCACTAAACCTTGTTTGGG** | Ins_k3_54_ACC | -0.059481870359395006 | 0.03924774780690166 | 46.26460345235955 |
| **GATCCAGATCTAACTTGGGGTGGCTTTGTCTTCTTCTTTTGCCAATTCCACTAAACGTTGTTTGGG** | Ins_k3_54_ACG | 0.4771284694502691 | 0.03947898335881494 | 79.12172617482891 |
| **GATCCAGATCTAACTTGGGGTGGCTTTGTCTTCTTCTTTTGCCAATTCCACTAAACTTGTTTGGG** | Ins_k2_54_AC | 0.05113669160629153 | 0.03929435603746053 | 51.67611726893004 |
| **GATCCAGATCTAACTTGGGGTGGCTTTGTCTTCTTCTTTTGCCAATTCCACTAAACTTTGTTTGGG** | Ins_k3_54_ACT | 0.33159233949572964 | 0.539309329035633 | 68.4053730082919 |
| **GATCCAGATCTAACTTGGGGTGGCTTTGTCTTCTTCTTTTGCCAATTCCACTAAAGATTGTTTGGG** | Ins_k3_53_AAG;Ins_k3_54_AGA | 0.5554236140623177 | 0.034973302556437366 | 85.56554144277602 |
| **GATCCAGATCTAACTTGGGGTGGCTTTGTCTTCTTCTTTTGCCAATTCCACTAAAGCTTGTTTGGG** | Ins_k3_54_AGC | 0.2986196537718887 | 0.03999648331110005 | 66.1866438841726 |
| **GATCCAGATCTAACTTGGGGTGGCTTTGTCTTCTTCTTTTGCCAATTCCACTAAAGGTTGTTTGGG** | Ins_k3_54_AGG | 0.2845243711072122 | 0.036007799561983016 | 65.26026852864882 |
| **GATCCAGATCTAACTTGGGGTGGCTTTGTCTTCTTCTTTTGCCAATTCCACTAAAGTTGTTTGGG** | Ins_k2_54_AG | 0.40969077037448476 | 0.03834198186521599 | 73.96187850845608 |
| **GATCCAGATCTAACTTGGGGTGGCTTTGTCTTCTTCTTTTGCCAATTCCACTAAAGTTTGTTTGGG** | Ins_k3_54_AGT | 0.22931129015556417 | 0.03463919893777076 | 61.75471475839338 |
| **GATCCAGATCTAACTTGGGGTGGCTTTGTCTTCTTCTTTTGCCAATTCCACTAAATATTGTTTGGG** | Ins_k3_53_AAT;Ins_k3_54_ATA | -0.05793600562090784 | 0.04036080752166488 | 46.336177579182696 |
| **GATCCAGATCTAACTTGGGGTGGCTTTGTCTTCTTCTTTTGCCAATTCCACTAAATCCTTGTTTGGG** | Ins_k4_54_ATCC | 0.1273551367404525 | 0.04319565731134761 | 55.76877759854484 |
| **GATCCAGATCTAACTTGGGGTGGCTTTGTCTTCTTCTTTTGCCAATTCCACTAAATCTTGTTTGGG** | Ins_k3_54_ATC | -0.17995625779469876 | 0.042881584057214914 | 41.013561364678104 |
| **GATCCAGATCTAACTTGGGGTGGCTTTGTCTTCTTCTTTTGCCAATTCCACTAAATGTTGTTTGGG** | Ins_k3_54_ATG | -0.3508153544929043 | 0.0507401907559765 | 34.571985286753836 |
| **GATCCAGATCTAACTTGGGGTGGCTTTGTCTTCTTCTTTTGCCAATTCCACTAAATGTTTGGG** | T55A | 0.11477477528976632 | 0.03829131720161636 | 55.07158090704022 |
| **GATCCAGATCTAACTTGGGGTGGCTTTGTCTTCTTCTTTTGCCAATTCCACTAAATTGTTTGGG** | Ins_k1_52_A;Ins_k1_53_A;Ins_k1 | 0.018691670495563528 | 0.037944961715692636 | 50.0263919562981 |
| **GATCCAGATCTAACTTGGGGTGGCTTTGTCTTCTTCTTTTGCCAATTCCACTAAATTTGTTTGGG** | Ins_k2_54_AT | -0.1413828159020375 | 0.04412464238713558 | 42.62650397697147 |
| **GATCCAGATCTAACTTGGGGTGGCTTTGTCTTCTTCTTTTGCCAATTCCACTAAATTTTGTTTGGG** | Ins_k3_54_ATT | -0.3368080869209347 | 0.057256206553544096 | 35.0596517993581 |
| **GATCCAGATCTAACTTGGGGTGGCTTTGTCTTCTTCTTTTGCCAATTCCACTAACAATTGTTTGGG** | Ins_k3_52_AAC;Ins_k3_53_ACA; | -0.18283652630326175 | 0.04169649843496018 | 40.89560125539864 |
| **GATCCAGATCTAACTTGGGGTGGCTTTGTCTTCTTCTTTTGCCAATTCCACTAACACTTGTTTGGG** | Ins_k3_54_CAC | -0.620713742863997 | 0.045160757668867395 | 26.394226470514123 |
| **GATCCAGATCTAACTTGGGGTGGCTTTGTCTTCTTCTTTTGCCAATTCCACTAACAGTTGTTTGGG** | Ins_k3_54_CAG | 0.05599615513555045 | 0.040756578231898696 | 51.92784661555239 |
| **GATCCAGATCTAACTTGGGGTGGCTTTGTCTTCTTCTTTTGCCAATTCCACTAACATTGTTTGGG** | Ins_k2_53_AC;Ins_k2_54_CA | -0.2400767392188231 | 0.04112588825766054 | 38.620464160398434 |
| **GATCCAGATCTAACTTGGGGTGGCTTTGTCTTCTTCTTTTGCCAATTCCACTAACATTTGTTTGGG** | Ins_k3_54_CAT | -0.3346799940598574 | 0.04527630507061266 | 35.13434143912102 |
| **GATCCAGATCTAACTTGGGGTGGCTTTGTCTTCTTCTTTTGCCAATTCCACTAACCATTGTTTGGG** | Ins_k3_53_ACC;Ins_k3_54_CCA | -0.06157169787993799 | 0.03396850261130969 | 46.168019368028155 |
| **GATCCAGATCTAACTTGGGGTGGCTTTGTCTTCTTCTTTTGCCAATTCCACTAACCCTTGTTTGGG** | Ins_k3_54_CCC | -0.9178396380964784 | 0.05965391295724363 | 19.60960302889955 |
| **GATCCAGATCTAACTTGGGGTGGCTTTGTCTTCTTCTTTTGCCAATTCCACTAACCGTTGTTTGGG** | Ins_k3_54_CCG | 0.2945741374438893 | 0.038555977452731385 | 65.91942561801872 |
| **GATCCAGATCTAACTTGGGGTGGCTTTGTCTTCTTCTTTTGCCAATTCCACTAACCTTGTTTGGG** | Ins_k2_54_CC | -0.2104864169569784 | 0.043710844365816855 | 39.780331942319854 |
| **GATCCAGATCTAACTTGGGGTGGCTTTGTCTTCTTCTTTTGCCAATTCCACTAACCTTTGTTTGGG** | Ins_k3_54_CCT | -0.5853595537850644 | 0.04195155671831465 | 27.344064385411443 |
| **GATCCAGATCTAACTTGGGGTGGCTTTGTCTTCTTCTTTTGCCAATTCCACTAACGACTTGTTTGGG** | Ins_k4_54_CGAC | 0.672477444319243 | 0.03301403971515966 | 96.1910654982213 |
| **GATCCAGATCTAACTTGGGGTGGCTTTGTCTTCTTCTTTTGCCAATTCCACTAACGATTGTTTGGG** | Ins_k3_53_ACG;Ins_k3_54_CGA | 0.1881901084304507 | 0.03734106995901233 | 59.26679164948489 |
| **GATCCAGATCTAACTTGGGGTGGCTTTGTCTTCTTCTTTTGCCAATTCCACTAACGCTTGTTTGGG** | Ins_k3_54_CGC | 0.3605405837671134 | 0.040565802275611504 | 70.41452891942181 |
| **GATCCAGATCTAACTTGGGGTGGCTTTGTCTTCTTCTTTTGCCAATTCCACTAACGGTTGTTTGGG** | Ins_k3_54_CGG | 0.5248969859208539 | 0.03769731466786468 | 82.99297956967568 |
| **GATCCAGATCTAACTTGGGGTGGCTTTGTCTTCTTCTTTTGCCAATTCCACTAACGTTGTTTGGG** | Ins_k2_54_CG | 0.26441639242664894 | 0.0451506719816261 | 63.961121820995174 |
| **GATCCAGATCTAACTTGGGGTGGCTTTGTCTTCTTCTTTTGCCAATTCCACTAACGTTTGTTTGGG** | Ins_k3_54_CGT | 0.18847883420302158 | 0.03958814201427938 | 59.28390597024553 |
| **GATCCAGATCTAACTTGGGGTGGCTTTGTCTTCTTCTTTTGCCAATTCCACTAACTAGTTGTTTGGG** | Ins_k4_54_CTAG | -1.188249848077764 | 0.07062811051091937 | 14.963429437101428 |
| **GATCCAGATCTAACTTGGGGTGGCTTTGTCTTCTTCTTTTGCCAATTCCACTAACTATTGTTTGGG** | Ins_k3_53_ACT;Ins_k3_54_CTA | -0.6281889512382036 | 0.05073970934739331 | 26.19765973176113 |
| **GATCCAGATCTAACTTGGGGTGGCTTTGTCTTCTTCTTTTGCCAATTCCACTAACTCTTGTTTGGG** | Ins_k3_54_CTC | -0.47191794621657046 | 0.04729327158281539 | 30.6288105909462 |
| **GATCCAGATCTAACTTGGGGTGGCTTTGTCTTCTTCTTTTGCCAATTCCACTAACTGTTGTTTGGG** | Ins_k3_54_CTG | -0.10588367798846804 | 0.03622773664649617 | 44.16688750293002 |
| **GATCCAGATCTAACTTGGGGTGGCTTTGTCTTCTTCTTTTGCCAATTCCACTAACTGTTTGGG** | T55C | 0.018058638512270297 | 0.028150298660944633 | 49.99473367159158 |
| **GATCCAGATCTAACTTGGGGTGGCTTTGTCTTCTTCTTTTGCCAATTCCACTAACTTGTTTGGG** | Ins_k1_54_C | -0.08960583742502282 | 0.03905981438801189 | 44.89171234292663 |
| **GATCCAGATCTAACTTGGGGTGGCTTTGTCTTCTTCTTTTGCCAATTCCACTAACTTTGTTTGGG** | Ins_k2_54_CT | -0.13907268274044982 | 0.0451482740999335 | 42.72509070776517 |
| **GATCCAGATCTAACTTGGGGTGGCTTTGTCTTCTTCTTTTGCCAATTCCACTAACTTTTGTTTGGG** | Ins_k3_54_CTT | -0.7471580694496965 | 0.04834517163461929 | 23.259204946321884 |
| **GATCCAGATCTAACTTGGGGTGGCTTTGTCTTCTTCTTTTGCCAATTCCACTAAG** | Del_k8_55-62 | 0.048879437641020096 | 0.026540334707646288 | 51.559602699274144 |
| **GATCCAGATCTAACTTGGGGTGGCTTTGTCTTCTTCTTTTGCCAATTCCACTAAGAATTGTTTGGG** | Ins_k3_52_AAG;Ins_k3_53_AGA; | 0.3019737114745632 | 0.03244220550485174 | 66.4090104135287 |
| **GATCCAGATCTAACTTGGGGTGGCTTTGTCTTCTTCTTTTGCCAATTCCACTAAGACTTGTTTGGG** | Ins_k3_54_GAC | 0.24398241351460126 | 0.03564724703581744 | 62.66740451794661 |
| **GATCCAGATCTAACTTGGGGTGGCTTTGTCTTCTTCTTTTGCCAATTCCACTAAGAGTTGTTTGGG** | Ins_k3_54_GAG | 0.33472143353727624 | 0.041377125521625556 | 68.6197550893061 |
| **GATCCAGATCTAACTTGGGGTGGCTTTGTCTTCTTCTTTTGCCAATTCCACTAAGATTGTTTGGG** | Ins_k2_53_AG;Ins_k2_54_GA | 0.38418259385019327 | 0.04259195045320664 | 72.09910483789606 |
| **GATCCAGATCTAACTTGGGGTGGCTTTGTCTTCTTCTTTTGCCAATTCCACTAAGATTTGTTTGGG** | Ins_k3_54_GAT | -0.05593037709302939 | 0.034993468774723153 | 46.42920399584231 |
| **GATCCAGATCTAACTTGGGGTGGCTTTGTCTTCTTCTTTTGCCAATTCCACTAAGCATTGTTTGGG** | Ins_k3_53_AGC;Ins_k3_54_GCA | 0.16956873763157543 | 0.03427836935190483 | 58.17337480190344 |
| **GATCCAGATCTAACTTGGGGTGGCTTTGTCTTCTTCTTTTGCCAATTCCACTAAGCCTTGTTTGGG** | Ins_k3_54_GCC | 0.09830982849996328 | 0.032705000074833755 | 54.17225425727486 |
| **GATCCAGATCTAACTTGGGGTGGCTTTGTCTTCTTCTTTTGCCAATTCCACTAAGCGCTTGTTTGGG** | Ins_k4_54_GCGC | 0.5806439163984838 | 0.03701528447102272 | 87.7509730522443 |
| **GATCCAGATCTAACTTGGGGTGGCTTTGTCTTCTTCTTTTGCCAATTCCACTAAGCGTTGTTTGGG** | Ins_k3_54_GCG | 0.5055539418163328 | 0.0364756556643741 | 81.403069131575 |
| **GATCCAGATCTAACTTGGGGTGGCTTTGTCTTCTTCTTTTGCCAATTCCACTAAGCTTGTTTGGG** | Ins_k2_54_GC | 0.22280095085050777 | 0.04104729459432012 | 61.35397649807668 |

| | | | | |
|---|---|---|---|---|
| GATCCAGATCTAACTTGGGGTGGCTTTGTCTTCTTCTTTTGCCAATTCCACTAAGCTTTGTTTGGG | Ins_k3_54_GCT | -0.18613094467649793 | 0.04318554914753706 | 40.76109571565022 |
| GATCCAGATCTAACTTGGGGTGGCTTTGTCTTCTTCTTTTGCCAATTCCACTAAGG | Del_k7_55-61 | -0.008084050488085692 | 0.029780954914826345 | 48.70467319490407 |
| GATCCAGATCTAACTTGGGGTGGCTTTGTCTTCTTCTTTTGCCAATTCCACTAAGGATTGTTTGGG | Ins_k3_53_AGG;Ins_k3_54_GGA | 0.5971196815379327 | 0.035608674480864 | 89.20871319184921 |
| GATCCAGATCTAACTTGGGGTGGCTTTGTCTTCTTCTTTTGCCAATTCCACTAAGGCTTGTTTGGG | Ins_k3_54_GGC | 0.12768980647687245 | 0.03632475061910636 | 55.78744484415106 |
| GATCCAGATCTAACTTGGGGTGGCTTTGTCTTCTTCTTTTGCCAATTCCACTAAGGG | Del_k6_55-60 | 0.0321665102841856 | 0.03213463039316865 | 50.705051720176016 |
| GATCCAGATCTAACTTGGGGTGGCTTTGTCTTCTTCTTTTGCCAATTCCACTAAGGGTTGTTTGGG | Ins_k3_54_GGG | -0.8901075816257982 | 0.06352279390152583 | 20.161028387254397 |
| GATCCAGATCTAACTTGGGGTGGCTTTGTCTTCTTCTTTTGCCAATTCCACTAAGGTTGTTTGGG | Ins_k2_54_GG | 0.3130165861279792 | 0.04688086234195725 | 67.14642086334928 |
| GATCCAGATCTAACTTGGGGTGGCTTTGTCTTCTTCTTTTGCCAATTCCACTAAGGTTTGTTTGGG | Ins_k3_54_GGT | 0.1881112779607278 | 0.04213887470268852 | 59.26211980460447 |
| GATCCAGATCTAACTTGGGGTGGCTTTGTCTTCTTCTTTTGCCAATTCCACTAAGTATTGTTTGGG | Ins_k3_53_AGT;Ins_k3_54_GTA | -0.32739407223048816 | 0.04343709135649774 | 35.391262320556656 |
| GATCCAGATCTAACTTGGGGTGGCTTTGTCTTCTTCTTTTGCCAATTCCACTAAGTCTTGTTTGGG | Ins_k3_54_GTC | -0.01791283205402472 | 0.03753300785848352 | 48.22831046833387 |
| GATCCAGATCTAACTTGGGGTGGCTTTGTCTTCTTCTTTTGCCAATTCCACTAAGTGTTGTTTGGG | Ins_k3_54_GTG | -0.16844924717456583 | 0.043239438752250364 | 41.48823062536347 |
| GATCCAGATCTAACTTGGGGTGGCTTTGTCTTCTTCTTTTGCCAATTCCACTAAGTGTTTGGG | T55G | 0.1322578709972465 | 0.03529751541783052 | 56.04286844320145 |
| GATCCAGATCTAACTTGGGGTGGCTTTGTCTTCTTCTTTTGCCAATTCCACTAAGTTGTTTGGG | Ins_k1_54_G | 0.1348764330522867 | 0.03810258375332209 | 56.189812479036014 |
| GATCCAGATCTAACTTGGGGTGGCTTTGTCTTCTTCTTTTGCCAATTCCACTAAGTTTGGG | Del_k2_55-56 | 0.1606223981155981 | 0.03412710626101506 | 57.655257124172884 |
| GATCCAGATCTAACTTGGGGTGGCTTTGTCTTCTTCTTTTGCCAATTCCACTAAGTTTGTTTGGG | Ins_k2_54_GT | -0.0516579007466188 | 0.046399778931508116 | 46.627996036446255 |
| GATCCAGATCTAACTTGGGGTGGCTTTGTCTTCTTCTTTTGCCAATTCCACTAAGTTTTGTTTGGG | Ins_k3_54_GTT | -0.5862934178602851 | 0.048930436330639364 | 27.318540665708777 |
| GATCCAGATCTAACTTGGGGTGGCTTTGTCTTCTTCTTTTGCCAATTCCACTAAT | Del_k8_56-63 | -3.945445674950469 | 0.15197887135600854 | 0.9497213623962567 |
| GATCCAGATCTAACTTGGGGTGGCTTTGTCTTCTTCTTTTGCCAATTCCACTAATAAATGTTTGGG | Ins_k3_55_AAA | -0.04376954030024527 | 0.051321015863060725 | 46.997269040483026 |
| GATCCAGATCTAACTTGGGGTGGCTTTGTCTTCTTCTTTTGCCAATTCCACTAATAACTGTTTGGG | Ins_k3_55_AAC | -0.23065580876980124 | 0.040686517967602816 | 38.986024120907665 |
| GATCCAGATCTAACTTGGGGTGGCTTTGTCTTCTTCTTTTGCCAATTCCACTAATAAGTGTTTGGG | Ins_k3_55_AAG | -0.046478138999926144 | 0.04600966041186235 | 46.8701445410192 |
| GATCCAGATCTAACTTGGGGTGGCTTTGTCTTCTTCTTTTGCCAATTCCACTAATAATGTTTGGG | Ins_k2_55_AA | -0.10822887008591009 | 0.048067586270221695 | 44.06342902980415 |
| GATCCAGATCTAACTTGGGGTGGCTTTGTCTTCTTCTTTTGCCAATTCCACTAATAATTGTTTGGG | Ins_k3_51_TAA;Ins_k3_52_AAT;In | -0.3315049459785428 | 0.04787497636570132 | 35.24607194349505 |
| GATCCAGATCTAACTTGGGGTGGCTTTGTCTTCTTCTTTTGCCAATTCCACTAATACATGTTTGGG | Ins_k3_55_ACA | 0.09623399352691853 | 0.037357088655468065 | 54.05991823318154 |
| GATCCAGATCTAACTTGGGGTGGCTTTGTCTTCTTCTTTTGCCAATTCCACTAATACCTGTTTGGG | Ins_k3_55_ACC | 0.23607257993616382 | 0.035836854590934 | 62.173671026511236 |
| GATCCAGATCTAACTTGGGGTGGCTTTGTCTTCTTCTTTTGCCAATTCCACTAATACGTGTTTGGG | Ins_k3_55_ACG | 0.30059995560441255 | 0.037016035959881975 | 66.31784328067178 |
| GATCCAGATCTAACTTGGGGTGGCTTTGTCTTCTTCTTTTGCCAATTCCACTAATACTGTTTGGG | Ins_k2_55_AC | 0.1457524765401429 | 0.0392200014737019 | 56.80427070339321 |
| GATCCAGATCTAACTTGGGGTGGCTTTGTCTTCTTCTTTTGCCAATTCCACTAATACTTGTTTGGG | Ins_k3_54_TAC;Ins_k3_55_ACT | -0.42977485408896254 | 0.04334716998277175 | 31.94718852012768 |
| GATCCAGATCTAACTTGGGGTGGCTTTGTCTTCTTCTTTTGCCAATTCCACTAATAGATGTTTGGG | Ins_k3_55_AGA | -0.0030107249843908113 | 0.0442545355193305 | 48.95239571272031 |
| GATCCAGATCTAACTTGGGGTGGCTTTGTCTTCTTCTTTTGCCAATTCCACTAATAGCTGTTTGGG | Ins_k3_55_AGC | 0.15756012674657582 | 0.03819672173872506 | 57.47897113653449 |
| GATCCAGATCTAACTTGGGGTGGCTTTGTCTTCTTCTTTTGCCAATTCCACTAATAGGTGTTTGGG | Ins_k3_55_AGG | -0.3713211802322621 | 0.059582084487922404 | 33.8702773198394 |
| GATCCAGATCTAACTTGGGGTGGCTTTGTCTTCTTCTTTTGCCAATTCCACTAATAGTGTTTGGG | Ins_k2_55_AG | -0.13357041748238968 | 0.050036474211537965 | 42.960823427258106 |
| GATCCAGATCTAACTTGGGGTGGCTTTGTCTTCTTCTTTTGCCAATTCCACTAATAGTTGTTTGGG | Ins_k3_54_TAG;Ins_k3_55_AGT | -0.744008203933866 | 0.053728566026021926 | 23.332583820029914 |
| GATCCAGATCTAACTTGGGGTGGCTTTGTCTTCTTCTTTTGCCAATTCCACTAATAGTTTGGG | T56A | -0.4150740127013289 | 0.03910628325736397 | 32.42030817922893 |
| GATCCAGATCTAACTTGGGGTGGCTTTGTCTTCTTCTTTTGCCAATTCCACTAATATAGTGTTTGGG | Ins_k4_55_ATAG | -0.9415442383492438 | 0.0847556138880719 | 19.150231349188072 |
| GATCCAGATCTAACTTGGGGTGGCTTTGTCTTCTTCTTTTGCCAATTCCACTAATATATGTTTGGG | Ins_k3_55_ATA | -0.19185356257516822 | 0.05064792302930892 | 40.52850169783878 |
| GATCCAGATCTAACTTGGGGTGGCTTTGTCTTCTTCTTTTGCCAATTCCACTAATATCTGTTTGGG | Ins_k3_55_ATC | -0.3086242155081136 | 0.042212396305578895 | 36.061824736876716 |
| GATCCAGATCTAACTTGGGGTGGCTTTGTCTTCTTCTTTTGCCAATTCCACTAATATGTGTTTGGG | Ins_k3_55_ATG | -0.4561972454134605 | 0.051692969700206814 | 31.114121678444867 |
| GATCCAGATCTAACTTGGGGTGGCTTTGTCTTCTTCTTTTGCCAATTCCACTAATATGTTTGGG | Ins_k1_55_A | 0.05639184273452347 | 0.04446211645954078 | 51.94839788617287 |
| GATCCAGATCTAACTTGGGGTGGCTTTGTCTTCTTCTTTTGCCAATTCCACTAATATTGTTTGGG | Ins_k2_53_AT;Ins_k2_54_TA;Ins_ | -0.28697140576991653 | 0.04456745152639714 | 36.85117960194325 |
| GATCCAGATCTAACTTGGGGTGGCTTTGTCTTCTTCTTTTGCCAATTCCACTAATATTTGTTTGGG | Ins_k3_54_TAT;Ins_k3_55_ATT | -0.44420160020532073 | 0.055787420374071035 | 31.489603212195007 |
| GATCCAGATCTAACTTGGGGTGGCTTTGTCTTCTTCTTTTGCCAATTCCACTAATCAATGTTTGGG | Ins_k3_55_CAA | 0.12778071235970068 | 0.03687308578546474 | 55.79251648159269 |
| GATCCAGATCTAACTTGGGGTGGCTTTGTCTTCTTCTTTTGCCAATTCCACTAATCACTGTTTGGG | Ins_k3_55_CAC | 0.11835643900780968 | 0.036618195252111636 | 55.26918245016788 |
| GATCCAGATCTAACTTGGGGTGGCTTTGTCTTCTTCTTTTGCCAATTCCACTAATCAGTGTTTGGG | Ins_k3_55_CAG | 0.12749995958500182 | 0.04060727252939066 | 55.77685477641952 |
| GATCCAGATCTAACTTGGGGTGGCTTTGTCTTCTTCTTTTGCCAATTCCACTAATCATGTTTGGG | Ins_k2_55_CA | 0.07363468255475902 | 0.038403303801861445 | 52.851902900567275 |
| GATCCAGATCTAACTTGGGGTGGCTTTGTCTTCTTCTTTTGCCAATTCCACTAATCATTGTTTGGG | Ins_k3_53_ATC;Ins_k3_54_TCA;I | 0.07980216219468228 | 0.04257556929392628 | 53.1788731905054 |
| GATCCAGATCTAACTTGGGGTGGCTTTGTCTTCTTCTTTTGCCAATTCCACTAATCCATGTTTGGG | Ins_k3_55_CCA | -0.21775340640680246 | 0.09104482973484818 | 39.49229653250066 |
| GATCCAGATCTAACTTGGGGTGGCTTTGTCTTCTTCTTTTGCCAATTCCACTAATCCCTGTTTGGG | Ins_k3_55_CCC | -0.19454567343962692 | 0.044589051490722155 | 40.419541210760556 |
| GATCCAGATCTAACTTGGGGTGGCTTTGTCTTCTTCTTTTGCCAATTCCACTAATCCGTGTTTGGG | Ins_k3_55_CCG | 0.4788860336575069 | 0.034105683824312116 | 79.26090996516413 |
| GATCCAGATCTAACTTGGGGTGGCTTTGTCTTCTTCTTTTGCCAATTCCACTAATCCTGTTTGGG | Ins_k2_55_CC | 0.4298446268166738 | 0.04030916233474552 | 75.46761785122453 |
| GATCCAGATCTAACTTGGGGTGGCTTTGTCTTCTTCTTTTGCCAATTCCACTAATCCTTGTTTGGG | Ins_k3_54_TCC;Ins_k3_55_CCT | -0.11505897251195235 | 0.03784707923640087 | 43.76349674638956 |
| GATCCAGATCTAACTTGGGGTGGCTTTGTCTTCTTCTTTTGCCAATTCCACTAATCGATGTTTGGG | Ins_k3_55_CGA | 0.46615471571252787 | 0.03648553492217493 | 78.25821050622703 |
| GATCCAGATCTAACTTGGGGTGGCTTTGTCTTCTTCTTTTGCCAATTCCACTAATCGCTGTTTGGG | Ins_k3_55_CGC | 0.6642500012855441 | 0.03124758494402149 | 95.40290570232182 |
| GATCCAGATCTAACTTGGGGTGGCTTTGTCTTCTTCTTTTGCCAATTCCACTAATCGGTGTTTGGG | Ins_k3_55_CGG | 0.7720600790666774 | 0.03688979160432656 | 100 |
| GATCCAGATCTAACTTGGGGTGGCTTTGTCTTCTTCTTTTGCCAATTCCACTAATCGTGTTTGGG | Ins_k2_55_CG | 0.6212968731573332 | 0.03717696753277886 | 91.39181362272456 |
| GATCCAGATCTAACTTGGGGTGGCTTTGTCTTCTTCTTTTGCCAATTCCACTAATCGTTGTTTGGG | Ins_k3_54_TCG;Ins_k3_55_CGT | 0.2034443500940476 | 0.040487985484872396 | 60.17779225851926 |
| GATCCAGATCTAACTTGGGGTGGCTTTGTCTTCTTCTTTTGCCAATTCCACTAATCGTTTGGG | T56C | 0.09909665767709153 | 0.028693432218518188 | 54.21489534093975 |
| GATCCAGATCTAACTTGGGGTGGCTTTGTCTTCTTCTTTTGCCAATTCCACTAATCTATGTTTGGG | Ins_k3_55_CTA | -0.5287490579674023 | 0.04077249981499989 | 28.936679488433175 |
| GATCCAGATCTAACTTGGGGTGGCTTTGTCTTCTTCTTTTGCCAATTCCACTAATCTCTGTTTGGG | Ins_k3_55_CTC | -0.07136454127135083 | 0.03593555718237121 | 45.71810972750425 |
| GATCCAGATCTAACTTGGGGTGGCTTTGTCTTCTTCTTTTGCCAATTCCACTAATCTGTGTTTGGG | Ins_k3_55_CTG | -0.149751034490413 | 0.040653763242539394 | 42.27128442437872 |
| GATCCAGATCTAACTTGGGGTGGCTTTGTCTTCTTCTTTTGCCAATTCCACTAATCTGTTTGGG | Ins_k1_55_C | -0.0077685551929217445 | 0.04115195185419234 | 48.72004171436987 |
| GATCCAGATCTAACTTGGGGTGGCTTTGTCTTCTTCTTTTGCCAATTCCACTAATCTTGTTTGGG | Ins_k2_54_TC;Ins_k2_55_CT | -0.14864384473425146 | 0.04701832596482104 | 42.31811267656997 |
| GATCCAGATCTAACTTGGGGTGGCTTTGTCTTCTTCTTTTGCCAATTCCACTAATCTTTGTTTGGG | Ins_k3_54_TCT;Ins_k3_55_CTT | -0.6312184498324802 | 0.05034639706534593 | 26.11841405637034 |
| GATCCAGATCTAACTTGGGGTGGCTTTGTCTTCTTCTTTTGCCAATTCCACTAATCTTTTGTTTGGG | Ins_k4_54_TCTT;Ins_k4_55_CTT | -1.2566744237787522 | 0.0719469647095993 | 13.973806473897401 |
| GATCCAGATCTAACTTGGGGTGGCTTTGTCTTCTTCTTTTGCCAATTCCACTAATG | Del_k7_56-62 | -0.18486175481897765 | 0.02776678818127867 | 40.81286212866816 |
| GATCCAGATCTAACTTGGGGTGGCTTTGTCTTCTTCTTTTGCCAATTCCACTAATGAATGTTTGGG | Ins_k3_55_GAA | 0.33023949197569014 | 0.03508912516583134 | 68.3128935384756 |
| GATCCAGATCTAACTTGGGGTGGCTTTGTCTTCTTCTTTTGCCAATTCCACTAATGACATGTTTGGG | Ins_k4_55_GACA | 0.4160025399263201 | 0.03523963311716458 | 74.43018520849127 |
| GATCCAGATCTAACTTGGGGTGGCTTTGTCTTCTTCTTTTGCCAATTCCACTAATGACTGTTTGGG | Ins_k3_55_GAC | 0.12966754707971984 | 0.033850192353149816 | 55.89788711603194 |
| GATCCAGATCTAACTTGGGGTGGCTTTGTCTTCTTCTTTTGCCAATTCCACTAATGAGTGTTTGGG | Ins_k3_55_GAG | 0.25358522768087455 | 0.044098075647524514 | 63.272086636145886 |
| GATCCAGATCTAACTTGGGGTGGCTTTGTCTTCTTCTTTTGCCAATTCCACTAATGATGTTTGGG | Ins_k2_55_GA | 0.4295561781359736 | 0.03401509494446788 | 75.44585245567049 |
| GATCCAGATCTAACTTGGGGTGGCTTTGTCTTCTTCTTTTGCCAATTCCACTAATGATTGTTTGGG | Ins_k3_53_ATG;Ins_k3_54_TGA;I | 0.062220181720156975 | 0.036498068900848835 | 52.252054806958995 |
| GATCCAGATCTAACTTGGGGTGGCTTTGTCTTCTTCTTTTGCCAATTCCACTAATGCATGTTTGGG | Ins_k3_55_GCA | -0.00935217559438628 | 0.03657221264153665 | 48.64294872148046 |
| GATCCAGATCTAACTTGGGGTGGCTTTGTCTTCTTCTTTTGCCAATTCCACTAATGCCTGTTTGGG | Ins_k3_55_GCC | 0.3948368757154789 | 0.0357868983563461 | 72.8713757136022 |
| GATCCAGATCTAACTTGGGGTGGCTTTGTCTTCTTCTTTTGCCAATTCCACTAATGCGTGTTTGGG | Ins_k3_55_GCG | 0.6753990037283681 | 0.038206394672090756 | 96.47250433065062 |
| GATCCAGATCTAACTTGGGGTGGCTTTGTCTTCTTCTTTTGCCAATTCCACTAATGCTGTGTTTGGG | Ins_k4_55_GCTG | 0.4650071312149098 | 0.035674447796961765 | 78.16845410841448 |
| GATCCAGATCTAACTTGGGGTGGCTTTGTCTTCTTCTTTTGCCAATTCCACTAATGCTGTTTGGG | Ins_k2_55_GC | 0.3786662104931242 | 0.036210568452345056 | 71.7024735268966 |

| | | | | |
|---|---|---|---|---|
| **GATCCAGATCTAACTTGGGGTGGCTTTGTCTTCTTCTTTTGCCAATTCCACTAATGCTTGTTTGGG** | Ins_k3_54_TGC;Ins_k3_55_GCT | -0.1481832295986818 | 0.039909535963194756 | 42.337609529705645 |
| **GATCCAGATCTAACTTGGGGTGGCTTTGTCTTCTTCTTTTGCCAATTCCACTAATGG** | Del_k6_56-61 | -0.032646797115201465 | 0.030226824035907894 | 47.52292554627084 |
| **GATCCAGATCTAACTTGGGGTGGCTTTGTCTTCTTCTTTTGCCAATTCCACTAATGGATGTTTGGG** | Ins_k3_55_GGA | 0.6915680910419946 | 0.20517023453148678 | 98.04505577693291 |
| **GATCCAGATCTAACTTGGGGTGGCTTTGTCTTCTTCTTTTGCCAATTCCACTAATGGCTGTTTGGG** | Ins_k3_55_GGC | 0.5185664364357399 | 0.03342995502944264 | 82.4692479092711 |
| **GATCCAGATCTAACTTGGGGTGGCTTTGTCTTCTTCTTTTGCCAATTCCACTAATGGG** | Del_k5_55-59;Del_k5_56-60 | 0.08963425807741376 | 0.029406909184231084 | 53.70431181929545 |
| **GATCCAGATCTAACTTGGGGTGGCTTTGTCTTCTTCTTTTGCCAATTCCACTAATGGGTGTTTGGG** | Ins_k3_55_GGG | 0.15046716891788614 | 0.04892741617719378 | 57.072717690723884 |
| **GATCCAGATCTAACTTGGGGTGGCTTTGTCTTCTTCTTTTGCCAATTCCACTAATGGTCTGTTTGGG** | Ins_k4_55_GGTC | 0.3271101513703776 | 0.04216700717514879 | 68.09945336454169 |
| **GATCCAGATCTAACTTGGGGTGGCTTTGTCTTCTTCTTTTGCCAATTCCACTAATGGTGTTTGGG** | Ins_k2_55_GG | 0.6536440300739483 | 0.0453635804524774 | 94.39641208683383 |
| **GATCCAGATCTAACTTGGGGTGGCTTTGTCTTCTTCTTTTGCCAATTCCACTAATGGTTGTTTGGG** | Ins_k3_54_TGG;Ins_k3_55_GGT | 0.004093802498146504 | 0.04037146608849303 | 49.30141770350769 |
| **GATCCAGATCTAACTTGGGGTGGCTTTGTCTTCTTCTTTTGCCAATTCCACTAATGGTTTGGG** | T56G | 0.21834249587666 | 0.0389984588024822 | 61.08104144283348 |
| **GATCCAGATCTAACTTGGGGTGGCTTTGTCTTCTTCTTTTGCCAATTCCACTAATGTACTGTTTGGG** | Ins_k4_55_GTAC | -0.2158119551410539 | 0.05004274852071624 | 39.5690433776144 |
| **GATCCAGATCTAACTTGGGGTGGCTTTGTCTTCTTCTTTTGCCAATTCCACTAATGTATGTTTGGG** | Ins_k3_55_GTA | -0.44327555669180985 | 0.04892300546468174 | 31.518777461219848 |
| **GATCCAGATCTAACTTGGGGTGGCTTTGTCTTCTTCTTTTGCCAATTCCACTAATGTCTGTTTGGG** | Ins_k3_55_GTC | 0.01422343120033076 | 0.03686655712021657 | 49.80336071550846 |
| **GATCCAGATCTAACTTGGGGTGGCTTTGTCTTCTTCTTTTGCCAATTCCACTAATGTGTGTTTGGG** | Ins_k3_55_GTG | -0.11671397408937767 | 0.039194602769215034 | 43.691127991960364 |
| **GATCCAGATCTAACTTGGGGTGGCTTTGTCTTCTTCTTTTGCCAATTCCACTAATGTGTTTGGG** | Ins_k1_55_G | 0.01882242623925423 | 0.039203069042789945 | 50.03293362205337 |
| **GATCCAGATCTAACTTGGGGTGGCTTTGTCTTCTTCTTTTGCCAATTCCACTAATGTTGTTTGGG** | Ins_k2_54_TG;Ins_k2_55_GT | -0.306301997306413 | 0.3152137507561203 | 36.14566547323192 |
| **GATCCAGATCTAACTTGGGGTGGCTTTGTCTTCTTCTTTTGCCAATTCCACTAATGTTTGGG** | Del_k1_55-55;Del_k1_56-56 | 0.11717382031630907 | 0.02972520138782461 | 55.20385871608624 |
| **GATCCAGATCTAACTTGGGGTGGCTTTGTCTTCTTCTTTTGCCAATTCCACTAATGTTTGTTTGGG** | Ins_k3_54_TGT;Ins_k3_55_GTT | -0.4794134083406206 | 0.05235837594412473 | 30.400091748787872 |
| **GATCCAGATCTAACTTGGGGTGGCTTTGTCTTCTTCTTTTGCCAATTCCACTAATT** | Del_k7_57-63 | -1.5012714829942229 | 0.04875143303029713 | 10.941769740776026 |
| **GATCCAGATCTAACTTGGGGTGGCTTTGTCTTCTTCTTTTGCCAATTCCACTAATTAAAGTTTGGG** | Ins_k3_56_AAA | 0.04874310333003429 | 0.042538422524526866 | 51.55257383551397 |
| **GATCCAGATCTAACTTGGGGTGGCTTTGTCTTCTTCTTTTGCCAATTCCACTAATTAACCGTTTGGG** | Ins_k4_56_AACC | 0.03335236913374506 | 0.038312227866107565 | 50.76521642084651 |
| **GATCCAGATCTAACTTGGGGTGGCTTTGTCTTCTTCTTTTGCCAATTCCACTAATTAACGTTTGGG** | Ins_k3_56_AAC | -0.08518299162439091 | 0.0381322702674897 | 45.09070118845797 |
| **GATCCAGATCTAACTTGGGGTGGCTTTGTCTTCTTCTTTTGCCAATTCCACTAATTAAGGTTTGGG** | Ins_k3_56_AAG | -1.0117603549904224 | 0.06717808498305668 | 17.851698893623535 |
| **GATCCAGATCTAACTTGGGGTGGCTTTGTCTTCTTCTTTTGCCAATTCCACTAATTAAGTTTGGG** | Ins_k2_56_AA | -0.011735312850539636 | 0.048213776300252295 | 48.52716391847048 |
| **GATCCAGATCTAACTTGGGGTGGCTTTGTCTTCTTCTTTTGCCAATTCCACTAATTAATGTTTGGG** | Ins_k3_55_TAA;Ins_k3_56_AAT | -0.644064587388962 | 0.05097997662521818 | 25.785039190872794 |
| **GATCCAGATCTAACTTGGGGTGGCTTTGTCTTCTTCTTTTGCCAATTCCACTAATTACAGTTTGGG** | Ins_k3_56_ACA | 0.054605967532486144 | 0.037079847182734306 | 51.85570732212749 |
| **GATCCAGATCTAACTTGGGGTGGCTTTGTCTTCTTCTTTTGCCAATTCCACTAATTACCGTTTGGG** | Ins_k3_56_ACC | 0.18418179711647187 | 0.03962258122337343 | 59.029707369201844 |
| **GATCCAGATCTAACTTGGGGTGGCTTTGTCTTCTTCTTTTGCCAATTCCACTAATTACGGTTTGGG** | Ins_k3_56_ACG | -0.03099559242898875 | 0.04154710139191914 | 47.601460444387655 |
| **GATCCAGATCTAACTTGGGGTGGCTTTGTCTTCTTCTTTTGCCAATTCCACTAATTACGTTTGGG** | Ins_k2_56_AC | 0.33265483151509334 | 0.04615821765835964 | 68.47809179591962 |
| **GATCCAGATCTAACTTGGGGTGGCTTTGTCTTCTTCTTTTGCCAATTCCACTAATTACTGTTTGGG** | Ins_k3_55_TAC;Ins_k3_56_ACT | -0.06551035294455887 | 0.044125745888233654 | 45.9865370973642 |
| **GATCCAGATCTAACTTGGGGTGGCTTTGTCTTCTTCTTTTGCCAATTCCACTAATTAGAGGTTTGGG** | Ins_k4_56_AGAG | -0.5863028843146348 | 0.07120651321544107 | 27.31828205721472 |
| **GATCCAGATCTAACTTGGGGTGGCTTTGTCTTCTTCTTTTGCCAATTCCACTAATTAGAGTTTGGG** | Ins_k3_56_AGA | -0.2256178102531542 | 0.0480497770542918 | 39.18293124501651 |
| **GATCCAGATCTAACTTGGGGTGGCTTTGTCTTCTTCTTTTGCCAATTCCACTAATTAGCGTTTGGG** | Ins_k3_56_AGC | -0.7258676486032936 | 0.048824540411223034 | 23.759712308598765 |
| **GATCCAGATCTAACTTGGGGTGGCTTTGTCTTCTTCTTTTGCCAATTCCACTAATTAGGGTTTGGG** | Ins_k3_56_AGG | -2.604481039944577 | 0.15024544803338355 | 3.6305276464271463 |
| **GATCCAGATCTAACTTGGGGTGGCTTTGTCTTCTTCTTTTGCCAATTCCACTAATTAGGTTTGGG** | Ins_k2_56_AG | -2.0237754264343497 | 0.12381055405651767 | 6.488838897825983 |
| **GATCCAGATCTAACTTGGGGTGGCTTTGTCTTCTTCTTTTGCCAATTCCACTAATTAGTGGTTTGGG** | Ins_k4_56_AGTG | -1.3032677123235548 | 0.07295492692094427 | 13.337656127675485 |
| **GATCCAGATCTAACTTGGGGTGGCTTTGTCTTCTTCTTTTGCCAATTCCACTAATTAGTGTTTGGG** | Ins_k3_55_TAG;Ins_k3_56_AGT | -0.9803675643344293 | 0.06902566145836382 | 18.421002805444193 |
| **GATCCAGATCTAACTTGGGGTGGCTTTGTCTTCTTCTTTTGCCAATTCCACTAATTAGTTTGGG** | Ins_k1_56_A | -0.7657126906937992 | 0.05167666631212599 | 22.831618329144092 |
| **GATCCAGATCTAACTTGGGGTGGCTTTGTCTTCTTCTTTTGCCAATTCCACTAATTATAGTTTGGG** | Ins_k3_56_ATA | -1.0637707407855153 | 0.07601492269830205 | 16.94695707336649 |
| **GATCCAGATCTAACTTGGGGTGGCTTTGTCTTCTTCTTTTGCCAATTCCACTAATTATCGTTTGGG** | Ins_k3_56_ATC | -0.08622193612717466 | 0.03883517090231284 | 45.04387877948546 |
| **GATCCAGATCTAACTTGGGGTGGCTTTGTCTTCTTCTTTTGCCAATTCCACTAATTATCTGTTTGGG** | Ins_k4_55_TATC;Ins_k4_56_ATC | -0.5890757395761668 | 0.050115297995575035 | 27.242637339470782 |
| **GATCCAGATCTAACTTGGGGTGGCTTTGTCTTCTTCTTTTGCCAATTCCACTAATTATGGTTTGGG** | Ins_k3_56_ATG | -0.6716746648569237 | 0.05351199704382594 | 25.08285060879382 |
| **GATCCAGATCTAACTTGGGGTGGCTTTGTCTTCTTCTTTTGCCAATTCCACTAATTATGTTTGGG** | Ins_k2_55_TA;Ins_k2_56_AT | -0.28630373259779074 | 0.04607457109645758 | 36.87579236164999 |
| **GATCCAGATCTAACTTGGGGTGGCTTTGTCTTCTTCTTTTGCCAATTCCACTAATTATTGTTTGGG** | Ins_k3_53_ATT;Ins_k3_54_TTA;In | -0.56672655269671 | 0.05076193264828169 | 27.858342762425565 |
| **GATCCAGATCTAACTTGGGGTGGCTTTGTCTTCTTCTTTTGCCAATTCCACTAATTATTTGGG** | G57A | -0.019868446255478678 | 0.03107116469864415 | 48.134086662205895 |
| **GATCCAGATCTAACTTGGGGTGGCTTTGTCTTCTTCTTTTGCCAATTCCACTAATTCAAGGTTTGGG** | Ins_k4_56_CAAG | 0.1663342929985273 | 0.04414889305565996 | 57.98552020840211 |
| **GATCCAGATCTAACTTGGGGTGGCTTTGTCTTCTTCTTTTGCCAATTCCACTAATTCAAGTTTGGG** | Ins_k3_56_CAA | 0.407854431596983 | 0.0366479305970709 | 73.82618407150409 |
| **GATCCAGATCTAACTTGGGGTGGCTTTGTCTTCTTCTTTTGCCAATTCCACTAATTCACGTTTGGG** | Ins_k3_56_CAC | 0.29437358703306227 | 0.03474936785189385 | 65.90620677569596 |
| **GATCCAGATCTAACTTGGGGTGGCTTTGTCTTCTTCTTTTGCCAATTCCACTAATTCAGGTTTGGG** | Ins_k3_56_CAG | -0.5164185788386154 | 0.05005484839977144 | 29.29569145702301 |
| **GATCCAGATCTAACTTGGGGTGGCTTTGTCTTCTTCTTTTGCCAATTCCACTAATTCAGTTTGGG** | Ins_k2_56_CA | -0.0075452476341549835 | 0.04152725278512925 | 48.73092248278182 |
| **GATCCAGATCTAACTTGGGGTGGCTTTGTCTTCTTCTTTTGCCAATTCCACTAATTCATGTTTGGG** | Ins_k3_55_TCA;Ins_k3_56_CAT | -0.00800348261600059 | 0.03848803374157091 | 48.70859738486367 |
| **GATCCAGATCTAACTTGGGGTGGCTTTGTCTTCTTCTTTTGCCAATTCCACTAATTCCAGTTTGGG** | Ins_k3_56_CCA | 0.5519720657712841 | 0.0357293383528501 | 85.2707169376378 |
| **GATCCAGATCTAACTTGGGGTGGCTTTGTCTTCTTCTTTTGCCAATTCCACTAATTCCCGTTTGGG** | Ins_k3_56_CCC | 0.1782851295997132 | 0.03690949508253379 | 58.68265304881211 |
| **GATCCAGATCTAACTTGGGGTGGCTTTGTCTTCTTCTTTTGCCAATTCCACTAATTCCGGTTTGGG** | Ins_k3_56_CCG | 0.4389298904041359 | 0.03689531815662603 | 76.15638513018058 |
| **GATCCAGATCTAACTTGGGGTGGCTTTGTCTTCTTCTTTTGCCAATTCCACTAATTCCGTTTGGG** | Ins_k2_56_CC | 0.31117988795213813 | 0.04088864356929342 | 67.02320634321576 |
| **GATCCAGATCTAACTTGGGGTGGCTTTGTCTTCTTCTTTTGCCAATTCCACTAATTCCTGTTTGGG** | Ins_k3_55_TCC;Ins_k3_56_CCT | 0.21993945963782535 | 0.032802683116598196 | 61.17866358127352 |
| **GATCCAGATCTAACTTGGGGTGGCTTTGTCTTCTTCTTTTGCCAATTCCACTAATTCGAGTTTGGG** | Ins_k3_56_CGA | 0.5420053284016177 | 0.03699883399539956 | 84.425067280881 |
| **GATCCAGATCTAACTTGGGGTGGCTTTGTCTTCTTCTTTTGCCAATTCCACTAATTCGCGTTTGGG** | Ins_k3_56_CGC | 0.3821495502780019 | 0.03267892674448847 | 71.95267311771514 |
| **GATCCAGATCTAACTTGGGGTGGCTTTGTCTTCTTCTTTTGCCAATTCCACTAATTCGGGTTTGGG** | Ins_k3_56_CGG | -0.5110169418703828 | 0.04864341609917384 | 29.454364307766777 |
| **GATCCAGATCTAACTTGGGGTGGCTTTGTCTTCTTCTTTTGCCAATTCCACTAATTCGGTTTGGG** | Ins_k2_56_CG | 0.028799413802064633 | 0.04899467762542802 | 50.534610026746 |
| **GATCCAGATCTAACTTGGGGTGGCTTTGTCTTCTTCTTTTGCCAATTCCACTAATTCGTGTTTGGG** | Ins_k3_55_TCG;Ins_k3_56_CGT | 0.46483393921866756 | 0.03509392382174956 | 78.15491713008626 |
| **GATCCAGATCTAACTTGGGGTGGCTTTGTCTTCTTCTTTTGCCAATTCCACTAATTCGTTTGGG** | Ins_k1_56_C | 0.13797925878467787 | 0.03634516237924848 | 56.3644304395499 |
| **GATCCAGATCTAACTTGGGGTGGCTTTGTCTTCTTCTTTTGCCAATTCCACTAATTCTAGTTTGGG** | Ins_k3_56_CTA | -0.9555665499832102 | 0.054627832622109976 | 18.88357477739138 |
| **GATCCAGATCTAACTTGGGGTGGCTTTGTCTTCTTCTTTTGCCAATTCCACTAATTCTCGTTTGGG** | Ins_k3_56_CTC | -0.06519478612550156 | 0.03666845703003344 | 46.00105121256161 |
| **GATCCAGATCTAACTTGGGGTGGCTTTGTCTTCTTCTTTTGCCAATTCCACTAATTCTGGTTTGGG** | Ins_k3_56_CTG | -0.24428529673809452 | 0.046770517094561266 | 38.45826925824498 |
| **GATCCAGATCTAACTTGGGGTGGCTTTGTCTTCTTCTTTTGCCAATTCCACTAATTCTGTTTGGG** | Ins_k2_55_TC;Ins_k2_56_CT | -0.06044250321358824 | 0.045136179180263136 | 46.22018149430928 |
| **GATCCAGATCTAACTTGGGGTGGCTTTGTCTTCTTCTTTTGCCAATTCCACTAATTCTTGTTTGGG** | Ins_k3_54_TTC;Ins_k3_55_TCT;Ir | -0.4892482980141699 | 0.04951034405303482 | 30.102575617637225 |
| **GATCCAGATCTAACTTGGGGTGGCTTTGTCTTCTTCTTTTGCCAATTCCACTAATTCTTTGGG** | G57C | -0.37278008090273407 | 0.036467991581572164 | 33.82089997662584 |
| **GATCCAGATCTAACTTGGGGTGGCTTTGTCTTCTTCTTTTGCCAATTCCACTAATTG** | Del_k6_57-62;Del_k6_58-63 | -0.2704464247422135 | 0.03395066801164145 | 37.46520404401169 |
| **GATCCAGATCTAACTTGGGGTGGCTTTGTCTTCTTCTTTTGCCAATTCCACTAATTGAAATTTGGG** | Ins_k3_57_AAA | 0.41664608980950435 | 0.038280108294195266 | 74.47810016166515 |
| **GATCCAGATCTAACTTGGGGTGGCTTTGTCTTCTTCTTTTGCCAATTCCACTAATTGAACTTTGGG** | Ins_k3_57_AAC | 0.32350371865757155 | 0.038325359018862144 | 67.85429959919617 |
| **GATCCAGATCTAACTTGGGGTGGCTTTGTCTTCTTCTTTTGCCAATTCCACTAATTGAAGTTTGGG** | Ins_k3_56_GAA;Ins_k3_57_AAG | 0.47712474464853094 | 0.03457942683924027 | 79.1214314626346 |
| **GATCCAGATCTAACTTGGGGTGGCTTTGTCTTCTTCTTTTGCCAATTCCACTAATTGAATTTGGG** | Ins_k2_57_AA | 0.2826961205210985 | 0.03765228838967745 | 65.14106540425546 |
| **GATCCAGATCTAACTTGGGGTGGCTTTGTCTTCTTCTTTTGCCAATTCCACTAATTGAATTTTGGG** | Ins_k3_57_AAT | 0.3565941640929283 | 0.04008969154413416 | 70.13719124252124 |
| **GATCCAGATCTAACTTGGGGTGGCTTTGTCTTCTTCTTTTGCCAATTCCACTAATTGACATTTGGG** | Ins_k3_57_ACA | 0.19117994509498715 | 0.04031451494655249 | 59.44425483692307 |

| | | | | |
|---|---|---|---|---|
| **GATCCAGATCTAACTTGGGGTGGCTTTGTCTTCTTCTTTTGCCAATTCCACTAATTGACCTTTGGG** | Ins_k3_57_ACC | 0.1038571074420066 | 0.036584819897203906 | 54.473597908451694 |
| **GATCCAGATCTAACTTGGGGTGGCTTTGTCTTCTTCTTTTGCCAATTCCACTAATTGACGTTTGGG** | Ins_k3_56_GAC;Ins_k3_57_ACG | 0.2851972718864966 | 0.031309071371259115 | 65.30419699228922 |
| **GATCCAGATCTAACTTGGGGTGGCTTTGTCTTCTTCTTTTGCCAATTCCACTAATTGACTCTTTGGG** | Ins_k4_57_ACTC | 0.06928799491118709 | 0.038540243457881825 | 52.62267074849164 |
| **GATCCAGATCTAACTTGGGGTGGCTTTGTCTTCTTCTTTTGCCAATTCCACTAATTGACTTTGGG** | Ins_k2_57_AC | 0.04063536817599911 | 0.03631691556478885 | 51.136289064068144 |
| **GATCCAGATCTAACTTGGGGTGGCTTTGTCTTCTTCTTTTGCCAATTCCACTAATTGACTTTTGGG** | Ins_k3_57_ACT | 0.24646040456698548 | 0.039534183968503986 | 62.8228863473487 |
| **GATCCAGATCTAACTTGGGGTGGCTTTGTCTTCTTCTTTTGCCAATTCCACTAATTGAGATTTGGG** | Ins_k3_57_AGA | 0.29651107886614037 | 0.042104650858265355 | 66.04723142024756 |
| **GATCCAGATCTAACTTGGGGTGGCTTTGTCTTCTTCTTTTGCCAATTCCACTAATTGAGCTTTGGG** | Ins_k3_57_AGC | 0.2628639231011687 | 0.03776939415868251 | 63.861901179773895 |
| **GATCCAGATCTAACTTGGGGTGGCTTTGTCTTCTTCTTTTGCCAATTCCACTAATTGAGGCTTTGGG** | Ins_k4_57_AGGC | -0.061544985655904605 | 0.04519941142414581 | 46.16925263497627 |
| **GATCCAGATCTAACTTGGGGTGGCTTTGTCTTCTTCTTTTGCCAATTCCACTAATTGAGGTTTGGG** | Ins_k3_56_GAG;Ins_k3_57_AGG | -0.34901993203879267 | 0.046635718657270175 | 34.634112360903764 |
| **GATCCAGATCTAACTTGGGGTGGCTTTGTCTTCTTCTTTTGCCAATTCCACTAATTGAGTTTGGG** | Ins_k2_56_GA;Ins_k2_57_AG | 0.17486278990255566 | 0.04734865767911163 | 58.48216434166306 |
| **GATCCAGATCTAACTTGGGGTGGCTTTGTCTTCTTCTTTTGCCAATTCCACTAATTGAGTTTTGGG** | Ins_k3_57_AGT | 0.08647192781840629 | 0.04658585007748442 | 53.53474929664499 |
| **GATCCAGATCTAACTTGGGGTGGCTTTGTCTTCTTCTTTTGCCAATTCCACTAATTGATATTTGGG** | Ins_k3_57_ATA | 0.17635928937921463 | 0.039852119538234934 | 58.56974838838883 |
| **GATCCAGATCTAACTTGGGGTGGCTTTGTCTTCTTCTTTTGCCAATTCCACTAATTGATCTTTGGG** | Ins_k3_57_ATC | 0.00733046753449762 | 0.040285346207511355 | 49.46124839810947 |
| **GATCCAGATCTAACTTGGGGTGGCTTTGTCTTCTTCTTTTGCCAATTCCACTAATTGATGTTTGGG** | Ins_k3_55_TGA;Ins_k3_56_GAT;I | 0.14248139488844747 | 0.03821327335799058 | 56.61876286680839 |
| **GATCCAGATCTAACTTGGGGTGGCTTTGTCTTCTTCTTTTGCCAATTCCACTAATTGATTGGG** | T58A | 0.053441275588819837 | 0.03278865536526578 | 51.79534655524907 |
| **GATCCAGATCTAACTTGGGGTGGCTTTGTCTTCTTCTTTTGCCAATTCCACTAATTGATTTGGG** | Ins_k1_57_A | 0.21402909916534452 | 0.03525245607736042 | 60.818142081781914 |
| **GATCCAGATCTAACTTGGGGTGGCTTTGTCTTCTTCTTTTGCCAATTCCACTAATTGATTTTGGG** | Ins_k2_57_AT | 0.03223495905312279 | 0.03711231021752991 | 50.70852253733039 |
| **GATCCAGATCTAACTTGGGGTGGCTTTGTCTTCTTCTTTTGCCAATTCCACTAATTGATTTTTGGG** | Ins_k3_57_ATT | -0.021137982906514696 | 0.039265521186899015 | 48.07301744802913 |
| **GATCCAGATCTAACTTGGGGTGGCTTTGTCTTCTTCTTTTGCCAATTCCACTAATTGCAATTTGGG** | Ins_k3_57_CAA | 0.2045144643610618 | 0.03842273940953768 | 60.24222384100173 |
| **GATCCAGATCTAACTTGGGGTGGCTTTGTCTTCTTCTTTTGCCAATTCCACTAATTGCACTTTGGG** | Ins_k3_57_CAC | 0.06000931313001279 | 0.036873466399799895 | 52.13665998860905 |
| **GATCCAGATCTAACTTGGGGTGGCTTTGTCTTCTTCTTTTGCCAATTCCACTAATTGCAGTTTGGG** | Ins_k3_56_GCA;Ins_k3_57_CAG | -0.5516426859109133 | 0.04618780157478332 | 28.281739495810232 |
| **GATCCAGATCTAACTTGGGGTGGCTTTGTCTTCTTCTTTTGCCAATTCCACTAATTGCATTTGGG** | Ins_k2_57_CA | 0.00897869022653619 | 0.03481861293457327 | 49.542838771177 |
| **GATCCAGATCTAACTTGGGGTGGCTTTGTCTTCTTCTTTTGCCAATTCCACTAATTGCATTTTGGG** | Ins_k3_57_CAT | 0.1581634979091575 | 0.040440291076360525 | 57.51366275508177 |
| **GATCCAGATCTAACTTGGGGTGGCTTTGTCTTCTTCTTTTGCCAATTCCACTAATTGCCATTTGGG** | Ins_k3_57_CCA | 0.23746001324159038 | 0.035359785873710955 | 62.25999271736126 |
| **GATCCAGATCTAACTTGGGGTGGCTTTGTCTTCTTCTTTTGCCAATTCCACTAATTGCCCTTTGGG** | Ins_k3_57_CCC | -1.7125065402767974 | 0.0687479432679127 | 8.85827893415448 |
| **GATCCAGATCTAACTTGGGGTGGCTTTGTCTTCTTCTTTTGCCAATTCCACTAATTGCCGTTTGGG** | Ins_k3_56_GCC;Ins_k3_57_CCG | 0.4639094174886569 | 0.03247431258288263 | 78.08269460168508 |
| **GATCCAGATCTAACTTGGGGTGGCTTTGTCTTCTTCTTTTGCCAATTCCACTAATTGCCTTTGGG** | Ins_k2_57_CC | -0.02270999627762471 | 0.039343098402423934 | 47.99750539034402 |
| **GATCCAGATCTAACTTGGGGTGGCTTTGTCTTCTTCTTTTGCCAATTCCACTAATTGCCTTTTGGG** | Ins_k3_57_CCT | -0.14744419426675665 | 0.0425639079246324 | 42.36891008370038 |
| **GATCCAGATCTAACTTGGGGTGGCTTTGTCTTCTTCTTTTGCCAATTCCACTAATTGCGATTTGGG** | Ins_k3_57_CGA | 0.6562626160480962 | 0.03835412532558502 | 94.64392112795545 |
| **GATCCAGATCTAACTTGGGGTGGCTTTGTCTTCTTCTTTTGCCAATTCCACTAATTGCGCTTTGGG** | Ins_k3_57_CGC | 0.16963728586074334 | 0.0353580142369067 | 58.17736262040822 |
| **GATCCAGATCTAACTTGGGGTGGCTTTGTCTTCTTCTTTTGCCAATTCCACTAATTGCGGTTTGGG** | Ins_k3_56_GCG;Ins_k3_57_CGG | 0.12159789819126954 | 0.043166938261734705 | 55.448625921408635 |
| **GATCCAGATCTAACTTGGGGTGGCTTTGTCTTCTTCTTTTGCCAATTCCACTAATTGCGTTTGGG** | Ins_k2_56_GC;Ins_k2_57_CG | 0.3762704497480812 | 0.04397609391802803 | 71.53089716552 |
| **GATCCAGATCTAACTTGGGGTGGCTTTGTCTTCTTCTTTTGCCAATTCCACTAATTGCGTTTTGGG** | Ins_k3_57_CGT | 0.19624283641078255 | 0.04608381525628521 | 59.74597778926666 |
| **GATCCAGATCTAACTTGGGGTGGCTTTGTCTTCTTCTTTTGCCAATTCCACTAATTGCTATTTGGG** | Ins_k3_57_CTA | -0.2992286427116459 | 0.038122583583157586 | 36.402242944140816 |
| **GATCCAGATCTAACTTGGGGTGGCTTTGTCTTCTTCTTTTGCCAATTCCACTAATTGCTCTTTGGG** | Ins_k3_57_CTC | -0.36935467449226134 | 0.038304298453719465 | 33.93694894818847 |
| **GATCCAGATCTAACTTGGGGTGGCTTTGTCTTCTTCTTTTGCCAATTCCACTAATTGCTGTTTGGG** | Ins_k3_55_TGC;Ins_k3_56_GCT; | 0.39889886073625735 | 0.03272469804820443 | 73.16798014374643 |
| **GATCCAGATCTAACTTGGGGTGGCTTTGTCTTCTTCTTTTGCCAATTCCACTAATTGCTTGGG** | T58C | -0.1433287259633384 | 0.027262826237025032 | 42.54363728571708 |
| **GATCCAGATCTAACTTGGGGTGGCTTTGTCTTCTTCTTTTGCCAATTCCACTAATTGCTTTGGG** | Ins_k1_57_C | -0.18002699949451112 | 0.0394036572691439 | 41.010660098253275 |
| **GATCCAGATCTAACTTGGGGTGGCTTTGTCTTCTTCTTTTGCCAATTCCACTAATTGCTTTTGGG** | Ins_k2_57_CT | -0.30803098972618337 | 0.041252275725757535 | 36.0832238876905 |
| **GATCCAGATCTAACTTGGGGTGGCTTTGTCTTCTTCTTTTGCCAATTCCACTAATTGCTTTTTGGG** | Ins_k3_57_CTT | -0.2940712333629058 | 0.04471000770638008 | 36.59046917486011 |
| **GATCCAGATCTAACTTGGGGTGGCTTTGTCTTCTTCTTTTGCCAATTCCACTAATTGG** | Del_k5_57-61;Del_k5_58-62 | -0.15103373388856117 | 0.029721174105738362 | 42.217097833272035 |
| **GATCCAGATCTAACTTGGGGTGGCTTTGTCTTCTTCTTTTGCCAATTCCACTAATTGGAATTTGGG** | Ins_k3_57_GAA | 0.1230963560866184 | 0.04150000913837438 | 55.53177563532094 |
| **GATCCAGATCTAACTTGGGGTGGCTTTGTCTTCTTCTTTTGCCAATTCCACTAATTGGACTTTGGG** | Ins_k3_57_GAC | 0.6100108518344323 | 0.03301202112178386 | 90.36616231348927 |
| **GATCCAGATCTAACTTGGGGTGGCTTTGTCTTCTTCTTTTGCCAATTCCACTAATTGGAGTTTGGG** | Ins_k3_56_GGA;Ins_k3_57_GAG | 0.7359963788502164 | 0.0333843462177553 | 100 |
| **GATCCAGATCTAACTTGGGGTGGCTTTGTCTTCTTCTTTTGCCAATTCCACTAATTGGATTTGGG** | Ins_k2_57_GA | 0.5100629402484169 | 0.036275056409154995 | 81.77094419342048 |
| **GATCCAGATCTAACTTGGGGTGGCTTTGTCTTCTTCTTTTGCCAATTCCACTAATTGGATTTTGGG** | Ins_k3_57_GAT | 0.6838038446078885 | 0.03423404297130974 | 97.28675741929895 |
| **GATCCAGATCTAACTTGGGGTGGCTTTGTCTTCTTCTTTTGCCAATTCCACTAATTGGCATTTGGG** | Ins_k3_57_GCA | 0.2618363475844091 | 0.033886702930634414 | 63.79631195837825 |
| **GATCCAGATCTAACTTGGGGTGGCTTTGTCTTCTTCTTTTGCCAATTCCACTAATTGGCCTTTGGG** | Ins_k3_57_GCC | -0.032926970051230686 | 0.035647297603386176 | 47.50961277371822 |
| **GATCCAGATCTAACTTGGGGTGGCTTTGTCTTCTTCTTTTGCCAATTCCACTAATTGGCGGTTTGGG** | Ins_k4_56_GGCG;Ins_k4_57_GC | 0.33753556808836405 | 0.044736942601404206 | 68.81313228008858 |
| **GATCCAGATCTAACTTGGGGTGGCTTTGTCTTCTTCTTTTGCCAATTCCACTAATTGGCGTTTGGG** | Ins_k3_56_GGC;Ins_k3_57_GCG | 0.2585688621051012 | 0.04052760654020139 | 63.588198624229 |
| **GATCCAGATCTAACTTGGGGTGGCTTTGTCTTCTTCTTTTGCCAATTCCACTAATTGGCTTTGGG** | Ins_k2_57_GC | 0.13893543568048583 | 0.03419683074521005 | 56.418350580169765 |
| **GATCCAGATCTAACTTGGGGTGGCTTTGTCTTCTTCTTTTGCCAATTCCACTAATTGGCTTTTGGG** | Ins_k3_57_GCT | 0.20908276161688266 | 0.034209404989319424 | 60.518057792896 |
| **GATCCAGATCTAACTTGGGGTGGCTTTGTCTTCTTCTTTTGCCAATTCCACTAATTGGG** | Del_k4_55-58;Del_k4_56-59;Del_ | 0.030073039628167653 | 0.030157142823835335 | 50.59901321526437 |
| **GATCCAGATCTAACTTGGGGTGGCTTTGTCTTCTTCTTTTGCCAATTCCACTAATTGGGATTTGGG** | Ins_k3_57_GGA | 0.42450437721019374 | 0.03999436792091135 | 75.06567612448951 |
| **GATCCAGATCTAACTTGGGGTGGCTTTGTCTTCTTCTTTTGCCAATTCCACTAATTGGGCTTTGGG** | Ins_k3_57_GGC | -0.34477079770483104 | 0.04105402767208961 | 34.78159046209963 |
| **GATCCAGATCTAACTTGGGGTGGCTTTGTCTTCTTCTTTTGCCAATTCCACTAATTGGGG** | Del_k3_58-60 | -0.600018387054966 | 0.03745763362954988 | 26.946155867212585 |
| **GATCCAGATCTAACTTGGGGTGGCTTTGTCTTCTTCTTTTGCCAATTCCACTAATTGGGGTTTGGG** | Ins_k3_56_GGG;Ins_k3_57_GGG | -1.7336137954557262 | 0.08136715149639896 | 8.673264423353597 |
| **GATCCAGATCTAACTTGGGGTGGCTTTGTCTTCTTCTTTTGCCAATTCCACTAATTGGGTTTGGG** | Ins_k2_56_GG;Ins_k2_57_GG | -1.1233709453788059 | 0.08530402219819917 | 15.96642515147074 |
| **GATCCAGATCTAACTTGGGGTGGCTTTGTCTTCTTCTTTTGCCAATTCCACTAATTGGGTTTTGGG** | Ins_k3_57_GGT | -1.1224105895626735 | 0.07900998441984476 | 15.981765965868917 |
| **GATCCAGATCTAACTTGGGGTGGCTTTGTCTTCTTCTTTTGCCAATTCCACTAATTGGTATTTGGG** | Ins_k3_57_GTA | 0.06499121053816637 | 0.04306993038518504 | 52.39704755339022 |
| **GATCCAGATCTAACTTGGGGTGGCTTTGTCTTCTTCTTTTGCCAATTCCACTAATTGGTCTTTGGG** | Ins_k3_57_GTC | 0.03654850968132306 | 0.033721999211162076 | 50.92772875524362 |
| **GATCCAGATCTAACTTGGGGTGGCTTTGTCTTCTTCTTTTGCCAATTCCACTAATTGGTGGTTTGGG** | Ins_k4_56_GGTG;Ins_k4_57_GT | -0.19654070485923542 | 0.05670482389827617 | 40.338983340539386 |
| **GATCCAGATCTAACTTGGGGTGGCTTTGTCTTCTTCTTTTGCCAATTCCACTAATTGGTGTTTGGG** | Ins_k3_55_TGG;Ins_k3_56_GGT; | 0.15826899921107396 | 0.03918982477809536 | 57.51973084147029 |
| **GATCCAGATCTAACTTGGGGTGGCTTTGTCTTCTTCTTTTGCCAATTCCACTAATTGGTTGGG** | T58G | -0.14068462455494152 | 0.03756842225922768 | 42.656275825219154 |
| **GATCCAGATCTAACTTGGGGTGGCTTTGTCTTCTTCTTTTGCCAATTCCACTAATTGGTTTGGG** | Ins_k1_56_G;Ins_k1_57_G | -0.1111227192171218 | 0.049645365809947516 | 43.93610043771381 |
| **GATCCAGATCTAACTTGGGGTGGCTTTGTCTTCTTCTTTTGCCAATTCCACTAATTGGTTTTGGG** | Ins_k2_57_GT | -0.08635593074198289 | 0.0430960220736368 | 45.03784354665231 |
| **GATCCAGATCTAACTTGGGGTGGCTTTGTCTTCTTCTTTTGCCAATTCCACTAATTGGTTTTTGGG** | Ins_k3_57_GTT | 0.11694185926754963 | 0.04630089536262372 | 55.191055056155484 |
| **GATCCAGATCTAACTTGGGGTGGCTTTGTCTTCTTCTTTTGCCAATTCCACTAATTGT** | Del_k5_59-63 | -4.119621623178892 | 0.2088709294514037 | 0.7979075116969384 |
| **GATCCAGATCTAACTTGGGGTGGCTTTGTCTTCTTCTTTTGCCAATTCCACTAATTGTAAATTGGG** | Ins_k3_58_AAA | -0.6533591100855711 | 0.0494170228058411 | 25.546489877202415 |
| **GATCCAGATCTAACTTGGGGTGGCTTTGTCTTCTTCTTTTGCCAATTCCACTAATTGTAACTTGGG** | Ins_k3_58_AAC | -0.6447266833260216 | 0.053376681658215616 | 25.76797267164757 |
| **GATCCAGATCTAACTTGGGGTGGCTTTGTCTTCTTCTTTTGCCAATTCCACTAATTGTAAGTTGGG** | Ins_k3_58_AAG | -0.42162124295945547 | 0.05114595551994493 | 32.20873831062904 |
| **GATCCAGATCTAACTTGGGGTGGCTTTGTCTTCTTCTTTTGCCAATTCCACTAATTGTAATTGGG** | Ins_k2_58_AA | -0.3613585940800115 | 0.04872510633001573 | 34.20939933688912 |
| **GATCCAGATCTAACTTGGGGTGGCTTTGTCTTCTTCTTTTGCCAATTCCACTAATTGTAATTTGGG** | Ins_k3_57_TAA;Ins_k3_58_AAT | -0.6486599501767883 | 0.04735654567545713 | 25.666819420699436 |
| **GATCCAGATCTAACTTGGGGTGGCTTTGTCTTCTTCTTTTGCCAATTCCACTAATTGTACATTGGG** | Ins_k3_58_ACA | -0.4498726788209596 | 0.045843192760793045 | 31.311528611589598 |
| **GATCCAGATCTAACTTGGGGTGGCTTTGTCTTCTTCTTTTGCCAATTCCACTAATTGTACCTTGGG** | Ins_k3_58_ACC | -0.4166614327473126 | 0.03851273462821305 | 32.368884358528746 |

| | | | | |
|---|---|---|---|---|
| GATCCAGATCTAACTTGGGGTGGCTTTGTCTTCTTCTTTTGCCAATTCCACTAATTGTACGTTGGG | Ins_k3_58_ACG | -0.23205539668044534 | 0.045744525303328384 | 38.93149791887126 |
| GATCCAGATCTAACTTGGGGTGGCTTTGTCTTCTTCTTTTGCCAATTCCACTAATTGTACGTTTGGG | Ins_k4_56_GTAC;Ins_k4_57_TAC | -0.13905338409298662 | 0.044875016444996266 | 42.72591525218484 |
| GATCCAGATCTAACTTGGGGTGGCTTTGTCTTCTTCTTTTGCCAATTCCACTAATTGTACTTGGG | Ins_k2_58_AC | -0.27445095406025216 | 0.036469703961083344 | 37.3154735362083 |
| GATCCAGATCTAACTTGGGGTGGCTTTGTCTTCTTCTTTTGCCAATTCCACTAATTGTACTTTGGG | Ins_k3_57_TAC;Ins_k3_58_ACT | -0.49019094488818316 | 0.04447421009852321 | 30.074212888950502 |
| GATCCAGATCTAACTTGGGGTGGCTTTGTCTTCTTCTTTTGCCAATTCCACTAATTGTAGATTGGG | Ins_k3_58_AGA | -0.9473090066036038 | 0.05407639829180973 | 19.040152298401836 |
| GATCCAGATCTAACTTGGGGTGGCTTTGTCTTCTTCTTTTGCCAATTCCACTAATTGTAGCATTGGG | Ins_k4_58_AGCA | -1.1944894887646704 | 0.06331547921974896 | 14.870353695539231 |
| GATCCAGATCTAACTTGGGGTGGCTTTGTCTTCTTCTTTTGCCAATTCCACTAATTGTAGCTTGGG | Ins_k3_58_AGC | -1.3844978693261827 | 0.06628445989647563 | 12.297071756696486 |
| GATCCAGATCTAACTTGGGGTGGCTTTGTCTTCTTCTTTTGCCAATTCCACTAATTGTAGGTTGGG | Ins_k3_58_AGG | -3.211825079361257 | 0.17761684039864384 | 1.977896597290999 |
| GATCCAGATCTAACTTGGGGTGGCTTTGTCTTCTTCTTTTGCCAATTCCACTAATTGTAGTTGGG | Ins_k2_58_AG | -1.3072985558146848 | 0.06354347226003533 | 13.28400233096082 |
| GATCCAGATCTAACTTGGGGTGGCTTTGTCTTCTTCTTTTGCCAATTCCACTAATTGTAGTTTGGG | Ins_k3_56_GTA;Ins_k3_57_TAG;I | -1.6360842037184837 | 0.07348496177660833 | 9.561788881766857 |
| GATCCAGATCTAACTTGGGGTGGCTTTGTCTTCTTCTTTTGCCAATTCCACTAATTGTATATTGGG | Ins_k3_58_ATA | -0.43758531090360003 | 0.04706568338600401 | 31.698638292780014 |
| GATCCAGATCTAACTTGGGGTGGCTTTGTCTTCTTCTTTTGCCAATTCCACTAATTGTATCTTGGG | Ins_k3_58_ATC | -0.6802748708169647 | 0.05100368657089031 | 24.868057882237753 |
| GATCCAGATCTAACTTGGGGTGGCTTTGTCTTCTTCTTTTGCCAATTCCACTAATTGTATGGG | T59A | -0.043239289866779496 | 0.03475123463721096 | 47.02219597093724 |
| GATCCAGATCTAACTTGGGGTGGCTTTGTCTTCTTCTTTTGCCAATTCCACTAATTGTATGTTGGG | Ins_k3_58_ATG | -0.6441572545485661 | 0.0487335256937115 | 25.782649875237954 |
| GATCCAGATCTAACTTGGGGTGGCTTTGTCTTCTTCTTTTGCCAATTCCACTAATTGTATTGGG | Ins_k1_58_A | -0.12264819385447329 | 0.039516059338943844 | 43.43262300796704 |
| GATCCAGATCTAACTTGGGGTGGCTTTGTCTTCTTCTTTTGCCAATTCCACTAATTGTATTTGGG | Ins_k2_57_TA;Ins_k2_58_AT | -0.1701746311072957 | 0.04839105624154015 | 41.416709217532 |
| GATCCAGATCTAACTTGGGGTGGCTTTGTCTTCTTCTTTTGCCAATTCCACTAATTGTATTTTGGG | Ins_k3_57_TAT;Ins_k3_58_ATT | -0.47313423599692495 | 0.04503390367215619 | 30.591579727988904 |
| GATCCAGATCTAACTTGGGGTGGCTTTGTCTTCTTCTTTTGCCAATTCCACTAATTGTCAATTGGG | Ins_k3_58_CAA | -0.21378843791647367 | 0.046396345790326844 | 39.64919308325556 |
| GATCCAGATCTAACTTGGGGTGGCTTTGTCTTCTTCTTTTGCCAATTCCACTAATTGTCACTTGGG | Ins_k3_58_CAC | -0.12434192924151666 | 0.03795714474177252 | 43.359121900710804 |
| GATCCAGATCTAACTTGGGGTGGCTTTGTCTTCTTCTTTTGCCAATTCCACTAATTGTCAGTTGGG | Ins_k3_58_CAG | -0.24330550875190887 | 0.04906596185076748 | 38.495968674135185 |
| GATCCAGATCTAACTTGGGGTGGCTTTGTCTTCTTCTTTTGCCAATTCCACTAATTGTCATTGGG | Ins_k2_58_CA | 0.1060461821232932 | 0.036311434744529296 | 54.592975297770074 |
| GATCCAGATCTAACTTGGGGTGGCTTTGTCTTCTTCTTTTGCCAATTCCACTAATTGTCATTTGGG | Ins_k3_57_TCA;Ins_k3_58_CAT | 0.022215956904616196 | 0.03762523654173841 | 50.20301033362829 |
| GATCCAGATCTAACTTGGGGTGGCTTTGTCTTCTTCTTTTGCCAATTCCACTAATTGTCCATTGGG | Ins_k3_58_CCA | -0.25984803375794 | 0.037696168302830055 | 37.86438653429165 |
| GATCCAGATCTAACTTGGGGTGGCTTTGTCTTCTTCTTTTGCCAATTCCACTAATTGTCCCTTGGG | Ins_k3_58_CCC | -0.7458508507430497 | 0.042961675495466915 | 23.289629695697677 |
| GATCCAGATCTAACTTGGGGTGGCTTTGTCTTCTTCTTTTGCCAATTCCACTAATTGTCCGTTGGG | Ins_k3_58_CCG | 0.09294929370925498 | 0.0414944857959738 | 53.88263894363569 |
| GATCCAGATCTAACTTGGGGTGGCTTTGTCTTCTTCTTTTGCCAATTCCACTAATTGTCCTTGGG | Ins_k2_58_CC | -0.4664208170601835 | 0.03825599316502943 | 30.797644745088444 |
| GATCCAGATCTAACTTGGGGTGGCTTTGTCTTCTTCTTTTGCCAATTCCACTAATTGTCCTTTGGG | Ins_k3_57_TCC;Ins_k3_58_CCT | -0.5487022655495791 | 0.03704710415044988 | 28.365022081377234 |
| GATCCAGATCTAACTTGGGGTGGCTTTGTCTTCTTCTTTTGCCAATTCCACTAATTGTCGATTGGG | Ins_k3_58_CGA | 0.12721068374901734 | 0.03787623573100336 | 55.76072221361829 |
| GATCCAGATCTAACTTGGGGTGGCTTTGTCTTCTTCTTTTGCCAATTCCACTAATTGTCGCTTGGG | Ins_k3_58_CGC | 0.0339929417530761 | 0.03428664888611773 | 50.7977456460533 |
| GATCCAGATCTAACTTGGGGTGGCTTTGTCTTCTTCTTTTGCCAATTCCACTAATTGTCGGTTGGG | Ins_k3_58_CGG | -0.07083866935443905 | 0.047009819074739195 | 45.74215792008461 |
| GATCCAGATCTAACTTGGGGTGGCTTTGTCTTCTTCTTTTGCCAATTCCACTAATTGTCGTCTTGGG | Ins_k4_58_CGTC | 0.40993681706802243 | 0.03348330655701883 | 73.98007882308272 |
| GATCCAGATCTAACTTGGGGTGGCTTTGTCTTCTTCTTTTGCCAATTCCACTAATTGTCGTTGGG | Ins_k2_58_CG | 0.14358639656192318 | 0.03469705223710742 | 56.681361273829054 |
| GATCCAGATCTAACTTGGGGTGGCTTTGTCTTCTTCTTTTGCCAATTCCACTAATTGTCGTTTGGG | Ins_k3_56_GTC;Ins_k3_57_TCG; | 0.004213591534869332 | 0.03767043098184858 | 49.3073238265808 |
| GATCCAGATCTAACTTGGGGTGGCTTTGTCTTCTTCTTTTGCCAATTCCACTAATTGTCTATTGGG | Ins_k3_58_CTA | -0.5737502867911408 | 0.041770919812837155 | 27.663358730140846 |
| GATCCAGATCTAACTTGGGGTGGCTTTGTCTTCTTCTTTTGCCAATTCCACTAATTGTCTCTTGGG | Ins_k3_58_CTC | -0.4852038515743285 | 0.04318211909999651 | 30.22457040682486 |
| GATCCAGATCTAACTTGGGGTGGCTTTGTCTTCTTCTTTTGCCAATTCCACTAATTGTCTGGG | T59C | 0.13669190470641868 | 0.026971119722391256 | 56.2919161459553 |
| GATCCAGATCTAACTTGGGGTGGCTTTGTCTTCTTCTTTTGCCAATTCCACTAATTGTCTGTTGGG | Ins_k3_58_CTG | -0.30178075209700356 | 0.04273569904551726 | 36.30945888616815 |
| GATCCAGATCTAACTTGGGGTGGCTTTGTCTTCTTCTTTTGCCAATTCCACTAATTGTCTGTTTGGG | Ins_k4_55_TGTC;Ins_k4_56_GTC | -0.4090087965477293 | 0.04833906579728737 | 32.61754188399489 |
| GATCCAGATCTAACTTGGGGTGGCTTTGTCTTCTTCTTTTGCCAATTCCACTAATTGTCTTGGG | Ins_k1_58_C | -0.27779424182862317 | 0.04213782831828949 | 37.190925485962254 |
| GATCCAGATCTAACTTGGGGTGGCTTTGTCTTCTTCTTTTGCCAATTCCACTAATTGTCTTTGGG | Ins_k2_57_TC;Ins_k2_58_CT | -0.2370077511876595 | 0.043385208606028125 | 38.73917196590921 |
| GATCCAGATCTAACTTGGGGTGGCTTTGTCTTCTTCTTTTGCCAATTCCACTAATTGTCTTTTGGG | Ins_k3_57_TCT;Ins_k3_58_CTT | -0.5401537281538824 | 0.045856273405787976 | 28.608540915321928 |
| GATCCAGATCTAACTTGGGGTGGCTTTGTCTTCTTCTTTTGCCAATTCCACTAATTGTG | Del_k4_59-62 | -0.5976438957221873 | 0.03456296973219816 | 27.010215304962475 |
| GATCCAGATCTAACTTGGGGTGGCTTTGTCTTCTTCTTTTGCCAATTCCACTAATTGTGAATTGGG | Ins_k3_58_GAA | -0.47231887191410526 | 0.04703264061265871 | 30.616533175025587 |
| GATCCAGATCTAACTTGGGGTGGCTTTGTCTTCTTCTTTTGCCAATTCCACTAATTGTGACTTGGG | Ins_k3_58_GAC | 0.18083512698313597 | 0.03352645237355907 | 58.83248461439917 |
| GATCCAGATCTAACTTGGGGTGGCTTTGTCTTCTTCTTTTGCCAATTCCACTAATTGTGAGTTGGG | Ins_k3_58_GAG | 0.09588750510706567 | 0.04676831810762862 | 54.0411903422216 |
| GATCCAGATCTAACTTGGGGTGGCTTTGTCTTCTTCTTTTGCCAATTCCACTAATTGTGATTGGG | Ins_k2_58_GA | 0.20600623368557291 | 0.03760472765029842 | 60.33215840670305 |
| GATCCAGATCTAACTTGGGGTGGCTTTGTCTTCTTCTTTTGCCAATTCCACTAATTGTGATTTGGG | Ins_k3_57_TGA;Ins_k3_58_GAT | 0.36040120016343724 | 0.03741241394429798 | 70.40471497259739 |
| GATCCAGATCTAACTTGGGGTGGCTTTGTCTTCTTCTTTTGCCAATTCCACTAATTGTGCATTGGG | Ins_k3_58_GCA | -0.0385761495236866 | 0.03273417178618762 | 47.241979111541994 |
| GATCCAGATCTAACTTGGGGTGGCTTTGTCTTCTTCTTTTGCCAATTCCACTAATTGTGCCGTTGGG | Ins_k4_58_GCCG | 0.31854176887833185 | 0.03770812782611085 | 67.51844391089878 |
| GATCCAGATCTAACTTGGGGTGGCTTTGTCTTCTTCTTTTGCCAATTCCACTAATTGTGCCTTGGG | Ins_k3_58_GCC | -0.02097743387490547 | 0.0339508502178748 | 48.08073614402495 |
| GATCCAGATCTAACTTGGGGTGGCTTTGTCTTCTTCTTTTGCCAATTCCACTAATTGTGCGTTGGG | Ins_k3_58_GCG | 0.19351908257811634 | 0.039554508858703846 | 59.58346587497221 |
| GATCCAGATCTAACTTGGGGTGGCTTTGTCTTCTTCTTTTGCCAATTCCACTAATTGTGCTTGGG | Ins_k2_58_GC | -0.08816485328233858 | 0.04121234253422461 | 44.9564472183137 |
| GATCCAGATCTAACTTGGGGTGGCTTTGTCTTCTTCTTTTGCCAATTCCACTAATTGTGCTTTGGG | Ins_k3_57_TGC;Ins_k3_58_GCT | -0.2340521758626809 | 0.034718025333363614 | 38.85383787507635 |
| GATCCAGATCTAACTTGGGGTGGCTTTGTCTTCTTCTTTTGCCAATTCCACTAATTGTGG | Del_k3_59-61 | -0.23034924221500952 | 0.033096374642931387 | 38.99797776420744 |
| GATCCAGATCTAACTTGGGGTGGCTTTGTCTTCTTCTTTTGCCAATTCCACTAATTGTGGATTGGG | Ins_k3_58_GGA | -1.8580999833660998 | 0.08200547254877305 | 7.6580627834458275 |
| GATCCAGATCTAACTTGGGGTGGCTTTGTCTTCTTCTTTTGCCAATTCCACTAATTGTGGCTTGGG | Ins_k3_58_GGC | -0.15547281498818744 | 0.0372165992406291 | 42.030108050663756 |
| GATCCAGATCTAACTTGGGGTGGCTTTGTCTTCTTCTTTTGCCAATTCCACTAATTGTGGG | Del_k2_58-59;Del_k2_59-60 | -0.09655884451840602 | 0.03357429455397338 | 44.58066256734298 |
| GATCCAGATCTAACTTGGGGTGGCTTTGTCTTCTTCTTTTGCCAATTCCACTAATTGTGGGTTGGG | Ins_k3_58_GGG | -3.2407003749539447 | 0.23841668222782644 | 1.9216009367858937 |
| GATCCAGATCTAACTTGGGGTGGCTTTGTCTTCTTCTTTTGCCAATTCCACTAATTGTGGTTGGG | Ins_k2_58_GG | -0.7861584524792299 | 0.05738705575529441 | 22.36954828310647 |
| GATCCAGATCTAACTTGGGGTGGCTTTGTCTTCTTCTTTTGCCAATTCCACTAATTGTGGTTTGGG | Ins_k3_56_GTG;Ins_k3_57_TGG; | -0.8782166515090484 | 0.057280223224140975 | 20.40219275964109 |
| GATCCAGATCTAACTTGGGGTGGCTTTGTCTTCTTCTTTTGCCAATTCCACTAATTGTGTATTGGG | Ins_k3_58_GTA | -0.29880199389155704 | 0.04949161489523486 | 36.41777723174862 |
| GATCCAGATCTAACTTGGGGTGGCTTTGTCTTCTTCTTTTGCCAATTCCACTAATTGTGTCTTGGG | Ins_k3_58_GTC | -0.4170742972870093 | 0.044174717516312155 | 32.35552315236148 |
| GATCCAGATCTAACTTGGGGTGGCTTTGTCTTCTTCTTTTGCCAATTCCACTAATTGTGTGGG | T59G | -0.07317154742617182 | 0.04140186395093471 | 45.63557141791575 |
| GATCCAGATCTAACTTGGGGTGGCTTTGTCTTCTTCTTTTGCCAATTCCACTAATTGTGTGTTGGG | Ins_k3_58_GTG | -0.19764913946058116 | 0.0574756890797753 | 40.2942949872556 |
| GATCCAGATCTAACTTGGGGTGGCTTTGTCTTCTTCTTTTGCCAATTCCACTAATTGTGTTGGG | Ins_k1_58_G | -0.12769298886497266 | 0.042044312923378395 | 43.21406607909752 |
| GATCCAGATCTAACTTGGGGTGGCTTTGTCTTCTTCTTTTGCCAATTCCACTAATTGTGTTTGGG | Ins_k2_55_TG;Ins_k2_56_GT;Ins_ | -0.070261908633789 | 0.04809771374802503 | 45.76854780964499 |
| GATCCAGATCTAACTTGGGGTGGCTTTGTCTTCTTCTTTTGCCAATTCCACTAATTGTGTTTTGGG | Ins_k3_57_TGT;Ins_k3_58_GTT | -0.09843586492169754 | 0.049230377712082364 | 44.49706223841847 |
| GATCCAGATCTAACTTGGGGTGGCTTTGTCTTCTTCTTTTGCCAATTCCACTAATTGTT | Del_k4_60-63 | -1.3058926681053995 | 0.05164912664512229 | 13.302691280772198 |
| GATCCAGATCTAACTTGGGGTGGCTTTGTCTTCTTCTTTTGCCAATTCCACTAATTGTTAAATGGG | Ins_k3_59_AAA | -0.2425150463396713 | 0.04777651396294022 | 38.52641032029822 |
| GATCCAGATCTAACTTGGGGTGGCTTTGTCTTCTTCTTTTGCCAATTCCACTAATTGTTAACTGGG | Ins_k3_59_AAC | -0.5629437783012189 | 0.04427162632255548 | 27.96392415752675 |
| GATCCAGATCTAACTTGGGGTGGCTTTGTCTTCTTCTTTTGCCAATTCCACTAATTGTTAAGTGGG | Ins_k3_59_AAG | -0.9537968367351979 | 0.06594907363847 | 18.917022877888414 |
| GATCCAGATCTAACTTGGGGTGGCTTTGTCTTCTTCTTTTGCCAATTCCACTAATTGTTAATGGG | Ins_k2_59_AA | -0.31515856059515357 | 0.04387516792179653 | 35.826952533299675 |
| GATCCAGATCTAACTTGGGGTGGCTTTGTCTTCTTCTTTTGCCAATTCCACTAATTGTTAATTGGG | Ins_k3_58_TAA;Ins_k3_59_AAT | -0.7063914755076539 | 0.05494514185636649 | 24.226996258971138 |
| GATCCAGATCTAACTTGGGGTGGCTTTGTCTTCTTCTTTTGCCAATTCCACTAATTGTTACATGGG | Ins_k3_59_ACA | -0.42263869079186445 | 0.04022506290856528 | 32.175984265242086 |
| GATCCAGATCTAACTTGGGGTGGCTTTGTCTTCTTCTTTTGCCAATTCCACTAATTGTTACCTGGG | Ins_k3_59_ACC | -0.20963438535872958 | 0.03469230391308479 | 39.814240485647375 |

| | | | | |
|---|---|---|---|---|
| **GATCCAGATCTAACTTGGGGTGGCTTTGTCTTCTTCTTTTGCCAATTCCACTAATTGTTACGTGGG** | Ins_k3_59_ACG | -0.33431278108282464 | 0.039355021478991135 | 35.147245594378596 |
| **GATCCAGATCTAACTTGGGGTGGCTTTGTCTTCTTCTTTTGCCAATTCCACTAATTGTTACTGGG** | Ins_k2_59_AC | -0.21058074710772012 | 0.03951036972053364 | 39.77657963459184 |
| **GATCCAGATCTAACTTGGGGTGGCTTTGTCTTCTTCTTTTGCCAATTCCACTAATTGTTACTTGGG** | Ins_k3_58_TAC;Ins_k3_59_ACT | -0.412971086510931 | 0.04476631042272571 | 32.48855743081758 |
| **GATCCAGATCTAACTTGGGGTGGCTTTGTCTTCTTCTTTTGCCAATTCCACTAATTGTTAGATGGG** | Ins_k3_59_AGA | -1.0803735713571918 | 0.06496676320760018 | 16.667912491280827 |
| **GATCCAGATCTAACTTGGGGTGGCTTTGTCTTCTTCTTTTGCCAATTCCACTAATTGTTAGCTGGG** | Ins_k3_59_AGC | -1.1747875496557743 | 0.06147754426040682 | 15.166233631996633 |
| **GATCCAGATCTAACTTGGGGTGGCTTTGTCTTCTTCTTTTGCCAATTCCACTAATTGTTAGGG** | T60A | -1.1236363177882454 | 0.05738034683217685 | 15.962188664906346 |
| **GATCCAGATCTAACTTGGGGTGGCTTTGTCTTCTTCTTTTGCCAATTCCACTAATTGTTAGGTGGG** | Ins_k3_59_AGG | -4.928974549695399 | 0.5086017417097787 | 0.35518534935680907 |
| **GATCCAGATCTAACTTGGGGTGGCTTTGTCTTCTTCTTTTGCCAATTCCACTAATTGTTAGTGGG** | Ins_k2_59_AG | -2.0737859939698877 | 0.09688639142874746 | 6.172309264014719 |
| **GATCCAGATCTAACTTGGGGTGGCTTTGTCTTCTTCTTTTGCCAATTCCACTAATTGTTAGTGTGGG** | Ins_k4_59_AGTG | -2.284328345212556 | 0.13264382447783873 | 5.00046390728876 |
| **GATCCAGATCTAACTTGGGGTGGCTTTGTCTTCTTCTTTTGCCAATTCCACTAATTGTTAGTTGGG** | Ins_k3_58_TAG;Ins_k3_59_AGT | -1.6410570886442608 | 0.07514532354324698 | 9.51435723976903 |
| **GATCCAGATCTAACTTGGGGTGGCTTTGTCTTCTTCTTTTGCCAATTCCACTAATTGTTATATGGG** | Ins_k3_59_ATA | -0.2987610833508606 | 0.04654928046139362 | 36.41926713318226 |
| **GATCCAGATCTAACTTGGGGTGGCTTTGTCTTCTTCTTTTGCCAATTCCACTAATTGTTATCATGGG** | Ins_k4_59_ATCA | -0.6440567594569416 | 0.04960557058978384 | 25.785241035196734 |
| **GATCCAGATCTAACTTGGGGTGGCTTTGTCTTCTTCTTTTGCCAATTCCACTAATTGTTATCTGGG** | Ins_k3_59_ATC | -0.7429036850200106 | 0.050840693647147925 | 23.358369337842873 |
| **GATCCAGATCTAACTTGGGGTGGCTTTGTCTTCTTCTTTTGCCAATTCCACTAATTGTTATGGG** | Ins_k1_59_A | -0.2404840841707701 | 0.0348444663249665 | 38.6047355129909 |
| **GATCCAGATCTAACTTGGGGTGGCTTTGTCTTCTTCTTTTGCCAATTCCACTAATTGTTATGTGGG** | Ins_k3_59_ATG | -0.7691378968757049 | 0.04997279659743213 | 22.753549106817566 |
| **GATCCAGATCTAACTTGGGGTGGCTTTGTCTTCTTCTTTTGCCAATTCCACTAATTGTTATTGGG** | Ins_k2_58_TA;Ins_k2_59_AT | -0.2622179829937556 | 0.04047083143040432 | 37.774756112087346 |
| **GATCCAGATCTAACTTGGGGTGGCTTTGTCTTCTTCTTTTGCCAATTCCACTAATTGTTATTTGGG** | Ins_k3_57_TTA;Ins_k3_58_TAT;In | -0.3205423061630871 | 0.04785383635823916 | 35.63458762288952 |
| **GATCCAGATCTAACTTGGGGTGGCTTTGTCTTCTTCTTTTGCCAATTCCACTAATTGTTCAATGGG** | Ins_k3_59_CAA | -0.13705052089430542 | 0.034644249610225865 | 42.81157516938618 |
| **GATCCAGATCTAACTTGGGGTGGCTTTGTCTTCTTCTTTTGCCAATTCCACTAATTGTTCACATGGG** | Ins_k4_59_CACA | -0.141039761039069 | 0.039484645721639886 | 42.641129715023965 |
| **GATCCAGATCTAACTTGGGGTGGCTTTGTCTTCTTCTTTTGCCAATTCCACTAATTGTTCACTGGG** | Ins_k3_59_CAC | -0.24453663048467345 | 0.037817338590188515 | 38.44860461192212 |
| **GATCCAGATCTAACTTGGGGTGGCTTTGTCTTCTTCTTTTGCCAATTCCACTAATTGTTCAGGTGGG** | Ins_k4_59_CAGG | -2.6648098528957957 | 0.17326383099173684 | 3.4179781153382165 |
| **GATCCAGATCTAACTTGGGGTGGCTTTGTCTTCTTCTTTTGCCAATTCCACTAATTGTTCAGTGGG** | Ins_k3_59_CAG | -0.6145088518650993 | 0.05358366907070563 | 26.55850891902519 |
| **GATCCAGATCTAACTTGGGGTGGCTTTGTCTTCTTCTTTTGCCAATTCCACTAATTGTTCATGGG** | Ins_k2_59_CA | -0.10666248751667828 | 0.043565679999141294 | 44.132503301221256 |
| **GATCCAGATCTAACTTGGGGTGGCTTTGTCTTCTTCTTTTGCCAATTCCACTAATTGTTCATTGGG** | Ins_k3_58_TCA;Ins_k3_59_CAT | -0.13577227609846693 | 0.0341683964558938 | 42.86633383258033 |
| **GATCCAGATCTAACTTGGGGTGGCTTTGTCTTCTTCTTTTGCCAATTCCACTAATTGTTCCATGGG** | Ins_k3_59_CCA | -0.20726869136530346 | 0.03528821791276313 | 39.90854029348245 |
| **GATCCAGATCTAACTTGGGGTGGCTTTGTCTTCTTCTTTTGCCAATTCCACTAATTGTTCCCTGGG** | Ins_k3_59_CCC | -0.6030401078329484 | 0.04574917499585036 | 26.864855004257105 |
| **GATCCAGATCTAACTTGGGGTGGCTTTGTCTTCTTCTTTTGCCAATTCCACTAATTGTTCCGTGGG** | Ins_k3_59_CCG | -0.09957110027418581 | 0.04082469810219296 | 44.446576262437944 |
| **GATCCAGATCTAACTTGGGGTGGCTTTGTCTTCTTCTTTTGCCAATTCCACTAATTGTTCCTGGG** | Ins_k2_59_CC | -0.10292693494297778 | 0.0368474472635434 | 44.29767089155337 |
| **GATCCAGATCTAACTTGGGGTGGCTTTGTCTTCTTCTTTTGCCAATTCCACTAATTGTTCCTTGGG** | Ins_k3_58_TCC;Ins_k3_59_CCT | -0.44808452536903737 | 0.03861386745375174 | 31.367568518605697 |
| **GATCCAGATCTAACTTGGGGTGGCTTTGTCTTCTTCTTTTGCCAATTCCACTAATTGTTCGATGGG** | Ins_k3_59_CGA | 0.005579890572663504 | 0.035609782547653464 | 49.374738419440625 |
| **GATCCAGATCTAACTTGGGGTGGCTTTGTCTTCTTCTTTTGCCAATTCCACTAATTGTTCGCTGGG** | Ins_k3_59_CGC | -0.09499834479617941 | 0.03246322375500966 | 44.650284987651304 |
| **GATCCAGATCTAACTTGGGGTGGCTTTGTCTTCTTCTTTTGCCAATTCCACTAATTGTTCGGG** | T60C | -0.2124277660254431 | 0.027904464270082918 | 39.703179346251964 |
| **GATCCAGATCTAACTTGGGGTGGCTTTGTCTTCTTCTTTTGCCAATTCCACTAATTGTTCGGTGGG** | Ins_k3_59_CGG | -0.7808787771267567 | 0.05404816582594687 | 22.487964560502558 |
| **GATCCAGATCTAACTTGGGGTGGCTTTGTCTTCTTCTTTTGCCAATTCCACTAATTGTTCGTGGG** | Ins_k2_59_CG | 0.10723439306090243 | 0.037336297234646196 | 54.65788182181991 |
| **GATCCAGATCTAACTTGGGGTGGCTTTGTCTTCTTCTTTTGCCAATTCCACTAATTGTTCGTTGGG** | Ins_k3_58_TCG;Ins_k3_59_CGT | -0.11994403426407357 | 0.03528358845526787 | 43.55023069532111 |
| **GATCCAGATCTAACTTGGGGTGGCTTTGTCTTCTTCTTTTGCCAATTCCACTAATTGTTCTATGGG** | Ins_k3_59_CTA | -0.4364528048120656 | 0.048941270853606354 | 31.734557529277925 |
| **GATCCAGATCTAACTTGGGGTGGCTTTGTCTTCTTCTTTTGCCAATTCCACTAATTGTTCTATTGGG** | Ins_k4_58_TCTA;Ins_k4_59_CTA | -0.9520727903158617 | 0.0622869864693331 | 18.94966483348363 |
| **GATCCAGATCTAACTTGGGGTGGCTTTGTCTTCTTCTTTTGCCAATTCCACTAATTGTTCTCTGGG** | Ins_k3_59_CTC | -0.7666110359724544 | 0.04841109899927375 | 22.81111686269061 |
| **GATCCAGATCTAACTTGGGGTGGCTTTGTCTTCTTCTTTTGCCAATTCCACTAATTGTTCTGGG** | Ins_k1_59_C | -0.13174369255250512 | 0.04140552333506379 | 43.03937275658734 |
| **GATCCAGATCTAACTTGGGGTGGCTTTGTCTTCTTCTTTTGCCAATTCCACTAATTGTTCTGTGGG** | Ins_k3_59_CTG | -0.6561353390959767 | 0.037456103231516304 | 25.475665328963732 |
| **GATCCAGATCTAACTTGGGGTGGCTTTGTCTTCTTCTTTTGCCAATTCCACTAATTGTTCTTGGG** | Ins_k2_58_TC;Ins_k2_59_CT | -0.4978207573723976 | 0.04218055931807167 | 29.845625432626967 |
| **GATCCAGATCTAACTTGGGGTGGCTTTGTCTTCTTCTTTTGCCAATTCCACTAATTGTTCTTTGGG** | Ins_k3_57_TTC;Ins_k3_58_TCT;I | -0.6189477688426048 | 0.044417456127632426 | 26.44087917037928 |
| **GATCCAGATCTAACTTGGGGTGGCTTTGTCTTCTTCTTTTGCCAATTCCACTAATTGTTG** | Del_k3_60-62 | -0.5482303844026247 | 0.03268246870735009 | 28.378410159072683 |
| **GATCCAGATCTAACTTGGGGTGGCTTTGTCTTCTTCTTTTGCCAATTCCACTAATTGTTGAATGGG** | Ins_k3_59_GAA | -0.20721012997508104 | 0.04024950608610494 | 39.910877461517025 |
| **GATCCAGATCTAACTTGGGGTGGCTTTGTCTTCTTCTTTTGCCAATTCCACTAATTGTTGACTGGG** | Ins_k3_59_GAC | -0.09003061355420161 | 0.035960185180758414 | 44.87264746456467 |
| **GATCCAGATCTAACTTGGGGTGGCTTTGTCTTCTTCTTTTGCCAATTCCACTAATTGTTGAGTGGG** | Ins_k3_59_GAG | -0.6337444979538303 | 0.057776144962513924 | 26.05252094523361 |
| **GATCCAGATCTAACTTGGGGTGGCTTTGTCTTCTTCTTTTGCCAATTCCACTAATTGTTGATCTGGG** | Ins_k4_59_GATC | -0.46768237963071524 | 0.04230358858731948 | 30.758816086758294 |
| **GATCCAGATCTAACTTGGGGTGGCTTTGTCTTCTTCTTTTGCCAATTCCACTAATTGTTGATGGG** | Ins_k2_59_GA | -0.020248545061485657 | 0.039665591388707594 | 48.1157944299854 |
| **GATCCAGATCTAACTTGGGGTGGCTTTGTCTTCTTCTTTTGCCAATTCCACTAATTGTTGATTGGG** | Ins_k3_58_TGA;Ins_k3_59_GAT | -0.0663057434169696 | 0.03862839534911542 | 45.94997438664127 |
| **GATCCAGATCTAACTTGGGGTGGCTTTGTCTTCTTCTTTTGCCAATTCCACTAATTGTTGCATGGG** | Ins_k3_59_GCA | -0.5768916504125169 | 0.04314958292916594 | 27.576594411867813 |
| **GATCCAGATCTAACTTGGGGTGGCTTTGTCTTCTTCTTTTGCCAATTCCACTAATTGTTGCCCTGGG** | Ins_k4_59_GCCC | -0.6167399565395321 | 0.05185734319107747 | 26.499320158342922 |
| **GATCCAGATCTAACTTGGGGTGGCTTTGTCTTCTTCTTTTGCCAATTCCACTAATTGTTGCCTGGG** | Ins_k3_59_GCC | 0.004785580661640143 | 0.037609116919004315 | 49.33553514719493 |
| **GATCCAGATCTAACTTGGGGTGGCTTTGTCTTCTTCTTTTGCCAATTCCACTAATTGTTGCGTGGG** | Ins_k3_59_GCG | -0.13429489056226723 | 0.038521644486961576 | 42.929710738709154 |
| **GATCCAGATCTAACTTGGGGTGGCTTTGTCTTCTTCTTTTGCCAATTCCACTAATTGTTGCTGGG** | Ins_k2_59_GC | 0.024679130894339596 | 0.03281678073778674 | 50.3268215045166 |
| **GATCCAGATCTAACTTGGGGTGGCTTTGTCTTCTTCTTTTGCCAATTCCACTAATTGTTGCTTGGG** | Ins_k3_58_TGC;Ins_k3_59_GCT | -0.518994840614107 | 0.04229225499915781 | 29.22031522308095 |
| **GATCCAGATCTAACTTGGGGTGGCTTTGTCTTCTTCTTTTGCCAATTCCACTAATTGTTGG** | Del_k2_60-61 | 0.0339816972764031 | 0.032878272985980075 | 50.79717445519872 |
| **GATCCAGATCTAACTTGGGGTGGCTTTGTCTTCTTCTTTTGCCAATTCCACTAATTGTTGGATGGG** | Ins_k3_59_GGA | 0.2720524043851035 | 0.0428857535721139 | 64.45139921182286 |
| **GATCCAGATCTAACTTGGGGTGGCTTTGTCTTCTTCTTTTGCCAATTCCACTAATTGTTGGCCTGGG** | Ins_k4_59_GGCC | -0.07069692580287934 | 0.04556319644585425 | 45.748642035534125 |
| **GATCCAGATCTAACTTGGGGTGGCTTTGTCTTCTTCTTTTGCCAATTCCACTAATTGTTGGCTGGG** | Ins_k3_59_GGC | 0.02480537432621477 | 0.03723839668696396 | 50.33317533623502 |
| **GATCCAGATCTAACTTGGGGTGGCTTTGTCTTCTTCTTTTGCCAATTCCACTAATTGTTGGG** | Del_k1_58-58;Del_k1_59-59;Del_ | -0.0013138002593555687 | 0.030606905785345865 | 49.03553476376032 |
| **GATCCAGATCTAACTTGGGGTGGCTTTGTCTTCTTCTTTTGCCAATTCCACTAATTGTTGGGG** | T60G | -0.6349068199317792 | 0.03876186159530187 | 26.02225711912029 |
| **GATCCAGATCTAACTTGGGGTGGCTTTGTCTTCTTCTTTTGCCAATTCCACTAATTGTTGGGTGGG** | Ins_k3_59_GGG | -4.0356626234708894 | 0.2783456475661162 | 0.8677916838668635 |
| **GATCCAGATCTAACTTGGGGTGGCTTTGTCTTCTTCTTTTGCCAATTCCACTAATTGTTGGTGGG** | Ins_k2_59_GG | -0.9535184300265769 | 0.0746888564815116 | 18.922290237164813 |
| **GATCCAGATCTAACTTGGGGTGGCTTTGTCTTCTTCTTTTGCCAATTCCACTAATTGTTGGTTGGG** | Ins_k3_58_TGG;Ins_k3_59_GGT | -0.545414011712067 | 0.05334309178711165 | 28.458446992317796 |
| **GATCCAGATCTAACTTGGGGTGGCTTTGTCTTCTTCTTTTGCCAATTCCACTAATTGTTGTATGGG** | Ins_k3_59_GTA | -0.4013904984294535 | 0.04382128359904511 | 32.866980986253445 |
| **GATCCAGATCTAACTTGGGGTGGCTTTGTCTTCTTCTTTTGCCAATTCCACTAATTGTTGTCGTGGG** | Ins_k4_59_GTCG | -0.24353494077063081 | 0.044243756435505675 | 38.48713747944777 |
| **GATCCAGATCTAACTTGGGGTGGCTTTGTCTTCTTCTTTTGCCAATTCCACTAATTGTTGTCTGGG** | Ins_k3_59_GTC | -0.5391259938243629 | 0.03726167839598201 | 28.63795800882722 |
| **GATCCAGATCTAACTTGGGGTGGCTTTGTCTTCTTCTTTTGCCAATTCCACTAATTGTTGTGGG** | Ins_k1_59_G | -0.3085068080730643 | 0.042275499053992256 | 36.066058911779216 |
| **GATCCAGATCTAACTTGGGGTGGCTTTGTCTTCTTCTTTTGCCAATTCCACTAATTGTTGTGTGGG** | Ins_k3_59_GTG | -0.7656116625104341 | 0.052491050189590646 | 22.833925082588763 |
| **GATCCAGATCTAACTTGGGGTGGCTTTGTCTTCTTCTTTTGCCAATTCCACTAATTGTTGTTGGG** | Ins_k2_58_TG;Ins_k2_59_GT | -0.3094735178252777 | 0.03703553730713016 | 36.03121034784251 |
| **GATCCAGATCTAACTTGGGGTGGCTTTGTCTTCTTCTTTTGCCAATTCCACTAATTGTTGTTTGGG** | Ins_k3_54_TTG;Ins_k3_55_TGT;I | -0.4539156746812192 | 0.04115235350647879 | 31.185191792742152 |
| **GATCCAGATCTAACTTGGGGTGGCTTTGTCTTCTTCTTTTGCCAATTCCACTAATTGTTT** | Del_k3_61-63 | -4.292104917687839 | 0.20017161829412455 | 0.6714968977654554 |
| **GATCCAGATCTAACTTGGGGTGGCTTTGTCTTCTTCTTTTGCCAATTCCACTAATTGTTTAAAGGG** | Ins_k3_60_AAA | -0.44093986687139924 | 0.04968176693310973 | 31.592481590372486 |
| **GATCCAGATCTAACTTGGGGTGGCTTTGTCTTCTTCTTTTGCCAATTCCACTAATTGTTTAACGGG** | Ins_k3_60_AAC | -0.4506460860762779 | 0.04130930505786317 | 31.28732141040545 |
| **GATCCAGATCTAACTTGGGGTGGCTTTGTCTTCTTCTTTTGCCAATTCCACTAATTGTTTAAGGG** | Ins_k2_60_AA | -0.8340157527321065 | 0.056192370831317145 | 21.324215013275875 |

| | | | | |
|---|---|---|---|---|
| GATCCAGATCTAACTTGGGGTGGCTTTGTCTTCTTCTTTTGCCAATTCCACTAATTGTTTAAGGGG | Ins_k3_60_AAG | -1.7276202128064515 | 0.1018922355748315 | 8.725404447199553 |
| GATCCAGATCTAACTTGGGGTGGCTTTGTCTTCTTCTTTTGCCAATTCCACTAATTGTTTAATGGG | Ins_k3_59_TAA;Ins_k3_60_AAT | -0.33675155326054085 | 0.04577545697371899 | 35.06163390583379 |
| GATCCAGATCTAACTTGGGGTGGCTTTGTCTTCTTCTTTTGCCAATTCCACTAATTGTTTACACGGG | Ins_k4_60_ACAC | -0.6751287395973576 | 0.04314051994153557 | 24.996362023096115 |
| GATCCAGATCTAACTTGGGGTGGCTTTGTCTTCTTCTTTTGCCAATTCCACTAATTGTTTACAGGG | Ins_k3_60_ACA | -1.0128857171337289 | 0.05342656103067238 | 17.83162056730811 |
| GATCCAGATCTAACTTGGGGTGGCTTTGTCTTCTTCTTTTGCCAATTCCACTAATTGTTTACAGGGG | Ins_k4_60_ACAG | -2.0678046190086476 | 0.10099250849092645 | 6.209338793449704 |
| GATCCAGATCTAACTTGGGGTGGCTTTGTCTTCTTCTTTTGCCAATTCCACTAATTGTTTACCGGG | Ins_k3_60_ACC | -0.5730769116306533 | 0.04143818754055911 | 27.68199282192854 |
| GATCCAGATCTAACTTGGGGTGGCTTTGTCTTCTTCTTTTGCCAATTCCACTAATTGTTTACGGG | Ins_k2_60_AC | -0.28287506515460065 | 0.03602764984180981 | 37.00244418978808 |
| GATCCAGATCTAACTTGGGGTGGCTTTGTCTTCTTCTTTTGCCAATTCCACTAATTGTTTACGGGG | Ins_k3_60_ACG | -0.902968505996153 | 0.05280167576998942 | 19.903399152044464 |
| GATCCAGATCTAACTTGGGGTGGCTTTGTCTTCTTCTTTTGCCAATTCCACTAATTGTTTACTGGG | Ins_k3_59_TAC;Ins_k3_60_ACT | -0.49253583968050774 | 0.04586462487906838 | 30.003774641179707 |
| GATCCAGATCTAACTTGGGGTGGCTTTGTCTTCTTCTTTTGCCAATTCCACTAATTGTTTAGAGGG | Ins_k3_60_AGA | -1.3648294462387578 | 0.07712399351339241 | 12.541329979619215 |
| GATCCAGATCTAACTTGGGGTGGCTTTGTCTTCTTCTTTTGCCAATTCCACTAATTGTTTAGCGGG | Ins_k3_60_AGC | -1.5127158542849488 | 0.06366627295290814 | 10.817261881305424 |
| GATCCAGATCTAACTTGGGGTGGCTTTGTCTTCTTCTTTTGCCAATTCCACTAATTGTTTAGG | G61A | -0.769162951606557 | 0.03860736243820979 | 22.752979029910357 |
| GATCCAGATCTAACTTGGGGTGGCTTTGTCTTCTTCTTTTGCCAATTCCACTAATTGTTTAGGG | Ins_k1_60_A | -1.416851704736924 | 0.06999484416775507 | 11.905581574939736 |
| GATCCAGATCTAACTTGGGGTGGCTTTGTCTTCTTCTTTTGCCAATTCCACTAATTGTTTAGGGG | Ins_k2_60_AG | -2.874320050631057 | 0.18602479993725424 | 2.7719165719328416 |
| GATCCAGATCTAACTTGGGGTGGCTTTGTCTTCTTCTTTTGCCAATTCCACTAATTGTTTAGGGGG | Ins_k3_60_AGG | -4.604432343632271 | 0.48305095130252484 | 0.4913624142825699 |
| GATCCAGATCTAACTTGGGGTGGCTTTGTCTTCTTCTTTTGCCAATTCCACTAATTGTTTAGTGGG | Ins_k3_59_TAG;Ins_k3_60_AGT | -2.538434638869759 | 0.1301300796316596 | 3.87840658738233 |
| GATCCAGATCTAACTTGGGGTGGCTTTGTCTTCTTCTTTTGCCAATTCCACTAATTGTTTATAAGGG | Ins_k4_60_ATAA | -0.7400725745376657 | 0.059601433375746006 | 23.424593161373192 |
| GATCCAGATCTAACTTGGGGTGGCTTTGTCTTCTTCTTTTGCCAATTCCACTAATTGTTTATAGGG | Ins_k3_60_ATA | -1.451357907488084 | 0.07972009118442573 | 11.50177219388354 |
| GATCCAGATCTAACTTGGGGTGGCTTTGTCTTCTTCTTTTGCCAATTCCACTAATTGTTTATCGGG | Ins_k3_60_ATC | -0.48186897833795744 | 0.046727002936355795 | 30.325533774200274 |
| GATCCAGATCTAACTTGGGGTGGCTTTGTCTTCTTCTTTTGCCAATTCCACTAATTGTTTATGGG | Ins_k2_59_TA;Ins_k2_60_AT | -0.29920841961272465 | 0.03907441398395801 | 36.402979117744664 |
| GATCCAGATCTAACTTGGGGTGGCTTTGTCTTCTTCTTTTGCCAATTCCACTAATTGTTTATGGGG | Ins_k3_60_ATG | -1.1385307604610402 | 0.07077026995788673 | 15.726202564295347 |
| GATCCAGATCTAACTTGGGGTGGCTTTGTCTTCTTCTTTTGCCAATTCCACTAATTGTTTATTGGG | Ins_k3_58_TTA;Ins_k3_59_TAT;In | -0.3635233382713783 | 0.05402679744510606 | 34.135424835208624 |
| GATCCAGATCTAACTTGGGGTGGCTTTGTCTTCTTCTTTTGCCAATTCCACTAATTGTTTCAACGGG | Ins_k4_60_CAAC | -0.27393203572390007 | 0.040682848356448906 | 37.334842244610236 |
| GATCCAGATCTAACTTGGGGTGGCTTTGTCTTCTTCTTTTGCCAATTCCACTAATTGTTTCAAGGG | Ins_k3_60_CAA | -0.3521773082764942 | 0.050154428816205944 | 34.52493189013546 |
| GATCCAGATCTAACTTGGGGTGGCTTTGTCTTCTTCTTTTGCCAATTCCACTAATTGTTTCACGGG | Ins_k3_60_CAC | -0.32840333227846186 | 0.04398095874605622 | 35.35556135226009 |
| GATCCAGATCTAACTTGGGGTGGCTTTGTCTTCTTCTTTTGCCAATTCCACTAATTGTTTCAGGG | Ins_k2_60_CA | -0.8992601980103843 | 0.05057888875505203 | 19.977344106658673 |
| GATCCAGATCTAACTTGGGGTGGCTTTGTCTTCTTCTTTTGCCAATTCCACTAATTGTTTCAGGGG | Ins_k3_60_CAG | -1.6401744515490395 | 0.07886238651001336 | 9.522758671568292 |
| GATCCAGATCTAACTTGGGGTGGCTTTGTCTTCTTCTTTTGCCAATTCCACTAATTGTTTCATAGGG | Ins_k4_60_CATA | -1.2195726900362907 | 0.06506826122014821 | 14.50199671986236 |
| GATCCAGATCTAACTTGGGGTGGCTTTGTCTTCTTCTTTTGCCAATTCCACTAATTGTTTCATGGG | Ins_k3_59_TCA;Ins_k3_60_CAT | -0.2856415964745456 | 0.042652706714273646 | 36.900217241251106 |
| GATCCAGATCTAACTTGGGGTGGCTTTGTCTTCTTCTTTTGCCAATTCCACTAATTGTTTCCAGGG | Ins_k3_60_CCA | -0.24576446619526054 | 0.04220925135500913 | 38.40142501248312 |
| GATCCAGATCTAACTTGGGGTGGCTTTGTCTTCTTCTTTTGCCAATTCCACTAATTGTTTCCCGGG | Ins_k3_60_CCC | -0.791347433674451 | 0.04399004277357168 | 22.253773753422607 |
| GATCCAGATCTAACTTGGGGTGGCTTTGTCTTCTTCTTTTGCCAATTCCACTAATTGTTTCCGGG | Ins_k2_60_CC | -0.679649296309518 | 0.03977238940505085 | 24.883619572289387 |
| GATCCAGATCTAACTTGGGGTGGCTTTGTCTTCTTCTTTTGCCAATTCCACTAATTGTTTCCGGGG | Ins_k3_60_CCG | -0.3419575304650444 | 0.046079126926638916 | 34.8795781392286 |
| GATCCAGATCTAACTTGGGGTGGCTTTGTCTTCTTCTTTTGCCAATTCCACTAATTGTTTCCGTGGG | Ins_k4_59_TCCG;Ins_k4_60_CC | -0.481268285363591 | 0.04489931189751503 | 30.343755581590273 |
| GATCCAGATCTAACTTGGGGTGGCTTTGTCTTCTTCTTTTGCCAATTCCACTAATTGTTTCCTGGG | Ins_k3_59_TCC;Ins_k3_60_CCT | -0.8162735006684922 | 0.045395563134379736 | 21.705930847141868 |
| GATCCAGATCTAACTTGGGGTGGCTTTGTCTTCTTCTTTTGCCAATTCCACTAATTGTTTCCTGGGG | Ins_k4_60_CCTG | -1.997467561485285 | 0.0772258635815491 | 6.661811691529562 |
| GATCCAGATCTAACTTGGGGTGGCTTTGTCTTCTTCTTTTGCCAATTCCACTAATTGTTTCGAGGG | Ins_k3_60_CGA | -0.137521650682758 | 0.04107158767483844 | 42.79141011158601 |
| GATCCAGATCTAACTTGGGGTGGCTTTGTCTTCTTCTTTTGCCAATTCCACTAATTGTTTCGCAGGG | Ins_k4_60_CGCA | -0.5413458653985946 | 0.05532090541481071 | 28.57445592920739 |
| GATCCAGATCTAACTTGGGGTGGCTTTGTCTTCTTCTTTTGCCAATTCCACTAATTGTTTCGCGGG | Ins_k3_60_CGC | -0.23087643522287965 | 0.03772020417283119 | 38.97742372145893 |
| GATCCAGATCTAACTTGGGGTGGCTTTGTCTTCTTCTTTTGCCAATTCCACTAATTGTTTCGCTGGG | Ins_k4_59_TCGC;Ins_k4_60_CG | -0.1598099849878789 | 0.04070177075084937 | 41.84821107114318 |
| GATCCAGATCTAACTTGGGGTGGCTTTGTCTTCTTCTTTTGCCAATTCCACTAATTGTTTCGG | G61C | 0.10076493670410636 | 0.03027501615021695 | 54.305416399986484 |
| GATCCAGATCTAACTTGGGGTGGCTTTGTCTTCTTCTTTTGCCAATTCCACTAATTGTTTCGGG | Ins_k1_60_C | -0.15771175883336946 | 0.03626305047220887 | 41.936110266078565 |
| GATCCAGATCTAACTTGGGGTGGCTTTGTCTTCTTCTTTTGCCAATTCCACTAATTGTTTCGGGG | Ins_k2_60_CG | -0.9101668615026522 | 0.054850508448700194 | 19.760641834550924 |
| GATCCAGATCTAACTTGGGGTGGCTTTGTCTTCTTCTTTTGCCAATTCCACTAATTGTTTCGGGGG | Ins_k3_60_CGG | -2.415306853415288 | 0.11126622521127617 | 4.386590094366523 |
| GATCCAGATCTAACTTGGGGTGGCTTTGTCTTCTTCTTTTGCCAATTCCACTAATTGTTTCGTGGG | Ins_k3_59_TCG;Ins_k3_60_CGT | -0.15320949159641173 | 0.038482508741922085 | 42.1253435110416 |
| GATCCAGATCTAACTTGGGGTGGCTTTGTCTTCTTCTTTTGCCAATTCCACTAATTGTTTCTAGGG | Ins_k3_60_CTA | -1.6476678211036708 | 0.07471675410179021 | 9.451667809363052 |
| GATCCAGATCTAACTTGGGGTGGCTTTGTCTTCTTCTTTTGCCAATTCCACTAATTGTTTCTCGGG | Ins_k3_60_CTC | -0.7074637922436053 | 0.04612396476041452 | 24.201031169308933 |
| GATCCAGATCTAACTTGGGGTGGCTTTGTCTTCTTCTTTTGCCAATTCCACTAATTGTTTCTGGG | Ins_k2_59_TC;Ins_k2_60_CT | -0.41069038998801083 | 0.04148003402792584 | 32.56273853091838 |
| GATCCAGATCTAACTTGGGGTGGCTTTGTCTTCTTCTTTTGCCAATTCCACTAATTGTTTCTGGGG | Ins_k3_60_CTG | -1.6524571477358958 | 0.07597737313893002 | 9.406508911680284 |
| GATCCAGATCTAACTTGGGGTGGCTTTGTCTTCTTCTTTTGCCAATTCCACTAATTGTTTCTTGGG | Ins_k3_58_TTC;Ins_k3_59_TCT;I | -0.704218659386161 | 0.04289701357617339 | 24.279694297859354 |
| GATCCAGATCTAACTTGGGGTGGCTTTGTCTTCTTCTTTTGCCAATTCCACTAATTGTTTG | Del_k2_61-62;Del_k2_62-63 | 0.2658859112933618 | 0.0280647931460419 | 64.05518299165273 |
| GATCCAGATCTAACTTGGGGTGGCTTTGTCTTCTTCTTTTGCCAATTCCACTAATTGTTTGAAAGG | Ins_k3_61_AAA | -0.04900984097452743 | 0.042470888069526466 | 46.7516333842995 |
| GATCCAGATCTAACTTGGGGTGGCTTTGTCTTCTTCTTTTGCCAATTCCACTAATTGTTTGAACGG | Ins_k3_61_AAC | -0.2144897017100369 | 0.041156503179656724 | 39.621398286583606 |
| GATCCAGATCTAACTTGGGGTGGCTTTGTCTTCTTCTTTTGCCAATTCCACTAATTGTTTGAAGCGG | Ins_k4_61_AAGC | 0.019851940543348934 | 0.03814295061054474 | 50.084469766940344 |
| GATCCAGATCTAACTTGGGGTGGCTTTGTCTTCTTCTTTTGCCAATTCCACTAATTGTTTGAAGG | Ins_k2_61_AA | 0.06635028396358444 | 0.03665825559889864 | 52.46830740100445 |
| GATCCAGATCTAACTTGGGGTGGCTTTGTCTTCTTCTTTTGCCAATTCCACTAATTGTTTGAAGGG | Ins_k3_60_GAA;Ins_k3_61_AAG | -0.14729296918194673 | 0.04067252975434833 | 42.375317810213375 |
| GATCCAGATCTAACTTGGGGTGGCTTTGTCTTCTTCTTTTGCCAATTCCACTAATTGTTTGAATGG | Ins_k3_61_AAT | -0.23682791875837045 | 0.04017718794265491 | 38.74613915175669 |
| GATCCAGATCTAACTTGGGGTGGCTTTGTCTTCTTCTTTTGCCAATTCCACTAATTGTTTGACAGG | Ins_k3_61_ACA | -0.26307024649896377 | 0.048096224339054715 | 37.7425757810436 |
| GATCCAGATCTAACTTGGGGTGGCTTTGTCTTCTTCTTTTGCCAATTCCACTAATTGTTTGACCGG | Ins_k3_61_ACC | 0.09946256474248327 | 0.034855328051110246 | 54.23473658399861 |
| GATCCAGATCTAACTTGGGGTGGCTTTGTCTTCTTCTTTTGCCAATTCCACTAATTGTTTGACGG | Ins_k2_61_AC | 0.09378804134248953 | 0.033634194302460126 | 53.92785183797556 |
| GATCCAGATCTAACTTGGGGTGGCTTTGTCTTCTTCTTTTGCCAATTCCACTAATTGTTTGACGGG | Ins_k3_60_GAC;Ins_k3_61_ACG | -0.09482066949424561 | 0.03685211624963886 | 44.65821894533121 |
| GATCCAGATCTAACTTGGGGTGGCTTTGTCTTCTTCTTTTGCCAATTCCACTAATTGTTTGACTCGG | Ins_k4_61_ACTC | -0.05074861281105425 | 0.03764835975915125 | 46.67041359266522 |
| GATCCAGATCTAACTTGGGGTGGCTTTGTCTTCTTCTTTTGCCAATTCCACTAATTGTTTGACTGG | Ins_k3_61_ACT | -0.10128659261790096 | 0.03810232175397121 | 44.370393864998334 |
| GATCCAGATCTAACTTGGGGTGGCTTTGTCTTCTTCTTTTGCCAATTCCACTAATTGTTTGAG | G62A | 0.34716792260241447 | 0.026900691504336924 | 69.47916735875256 |
| GATCCAGATCTAACTTGGGGTGGCTTTGTCTTCTTCTTTTGCCAATTCCACTAATTGTTTGAGAGG | Ins_k3_61_AGA | 0.00843149801391363 | 0.04407282809986239 | 49.51573673129902 |
| GATCCAGATCTAACTTGGGGTGGCTTTGTCTTCTTCTTTTGCCAATTCCACTAATTGTTTGAGCGG | Ins_k3_61_AGC | 0.12721490212160558 | 0.03507583477087872 | 55.7609574336165 |
| GATCCAGATCTAACTTGGGGTGGCTTTGTCTTCTTCTTTTGCCAATTCCACTAATTGTTTGAGG | Ins_k1_61_A | -0.08332167766581311 | 0.039974073139129096 | 45.17470729659976 |
| GATCCAGATCTAACTTGGGGTGGCTTTGTCTTCTTCTTTTGCCAATTCCACTAATTGTTTGAGGG | Ins_k2_60_GA;Ins_k2_61_AG | -0.3884647863310814 | 0.0448630897361215 | 33.294567597539995 |
| GATCCAGATCTAACTTGGGGTGGCTTTGTCTTCTTCTTTTGCCAATTCCACTAATTGTTTGAGGGG | Ins_k3_60_GAG;Ins_k3_61_AGG | -1.1380299767271391 | 0.06416783508155345 | 15.73407996300752 |
| GATCCAGATCTAACTTGGGGTGGCTTTGTCTTCTTCTTTTGCCAATTCCACTAATTGTTTGAGGGGG | Ins_k4_60_GAGG;Ins_k4_61_AG | -4.187975651045644 | 0.29532233675196407 | 0.7451895851646844 |
| GATCCAGATCTAACTTGGGGTGGCTTTGTCTTCTTCTTTTGCCAATTCCACTAATTGTTTGAGTGG | Ins_k3_61_AGT | -0.29459719569998777 | 0.046735575496778335 | 36.57122902641824 |
| GATCCAGATCTAACTTGGGGTGGCTTTGTCTTCTTCTTTTGCCAATTCCACTAATTGTTTGATAGG | Ins_k3_61_ATA | -1.2217222497548255 | 0.05796982959403053 | 14.470857291895282 |
| GATCCAGATCTAACTTGGGGTGGCTTTGTCTTCTTCTTTTGCCAATTCCACTAATTGTTTGATCGG | Ins_k3_61_ATC | 0.1740627355357416 | 0.038339974080805375 | 58.43539414254654 |
| GATCCAGATCTAACTTGGGGTGGCTTTGTCTTCTTCTTTTGCCAATTCCACTAATTGTTTGATGG | Ins_k2_61_AT | -0.03185701922079476 | 0.03739812728611965 | 47.560472927438965 |
| GATCCAGATCTAACTTGGGGTGGCTTTGTCTTCTTCTTTTGCCAATTCCACTAATTGTTTGATGGG | Ins_k3_59_TGA;Ins_k3_60_GAT;I | -0.04982684340901977 | 0.036262276899728454 | 46.71345278495312 |

| | | | | |
|---|---|---|---|---|
| **GATCCAGATCTAACTTGGGGTGGCTTTGTCTTCTTCTTTTGCCAATTCCACTAATTGTTTGATTGG** | Ins_k3_61_ATT | 0.013005118090237078 | 0.040158780798623436 | 49.74272157445279 |
| **GATCCAGATCTAACTTGGGGTGGCTTTGTCTTCTTCTTTTGCCAATTCCACTAATTGTTTGCAAGG** | Ins_k3_61_CAA | -0.524168076421014 | 0.041416018983889226 | 29.069541971126625 |
| **GATCCAGATCTAACTTGGGGTGGCTTTGTCTTCTTCTTTTGCCAATTCCACTAATTGTTTGCACAGG** | Ins_k4_61_CACA | -0.2565896438955618 | 0.043646808734083956 | 37.987964691089985 |
| **GATCCAGATCTAACTTGGGGTGGCTTTGTCTTCTTCTTTTGCCAATTCCACTAATTGTTTGCACGG** | Ins_k3_61_CAC | -0.08432783680938805 | 0.038070454970178316 | 45.12927721057581 |
| **GATCCAGATCTAACTTGGGGTGGCTTTGTCTTCTTCTTTTGCCAATTCCACTAATTGTTTGCAGG** | Ins_k2_61_CA | -0.443263244468144 | 0.04033066384400783 | 31.51916552984661 |
| **GATCCAGATCTAACTTGGGGTGGCTTTGTCTTCTTCTTTTGCCAATTCCACTAATTGTTTGCAGGG** | Ins_k3_60_GCA;Ins_k3_61_CAG | -0.3920277774162713 | 0.04709458585991844 | 33.176150434926605 |
| **GATCCAGATCTAACTTGGGGTGGCTTTGTCTTCTTCTTTTGCCAATTCCACTAATTGTTTGCATGG** | Ins_k3_61_CAT | -0.1430746890553819 | 0.04317712602404389 | 42.55444631267413 |
| **GATCCAGATCTAACTTGGGGTGGCTTTGTCTTCTTCTTTTGCCAATTCCACTAATTGTTTGCATGGG** | Ins_k4_59_TGCA;Ins_k4_60_GC/ | -0.6323250448986886 | 0.05415186210779861 | 26.08952753402768 |
| **GATCCAGATCTAACTTGGGGTGGCTTTGTCTTCTTCTTTTGCCAATTCCACTAATTGTTTGCCAGG** | Ins_k3_61_CCA | -0.5274418760809632 | 0.0391612424415264 | 28.974529724896264 |
| **GATCCAGATCTAACTTGGGGTGGCTTTGTCTTCTTCTTTTGCCAATTCCACTAATTGTTTGCCCGG** | Ins_k3_61_CCC | -0.2942652510432433 | 0.035026466806923715 | 36.58337066554876 |
| **GATCCAGATCTAACTTGGGGTGGCTTTGTCTTCTTCTTTTGCCAATTCCACTAATTGTTTGCCGG** | Ins_k2_61_CC | -0.08048530043278568 | 0.0371360961143836 | 45.303021695880425 |
| **GATCCAGATCTAACTTGGGGTGGCTTTGTCTTCTTCTTTTGCCAATTCCACTAATTGTTTGCCGGG** | Ins_k3_60_GCC;Ins_k3_61_CCG | -0.24201141657175296 | 0.04257994474368373 | 38.545818254182954 |
| **GATCCAGATCTAACTTGGGGTGGCTTTGTCTTCTTCTTTTGCCAATTCCACTAATTGTTTGCCTGG** | Ins_k3_61_CCT | -0.3886200427831301 | 0.036629316283378954 | 33.28939880235709 |
| **GATCCAGATCTAACTTGGGGTGGCTTTGTCTTCTTCTTTTGCCAATTCCACTAATTGTTTGCG** | G62C | -0.017573145242492316 | 0.033209808001083925 | 48.2446957721207 |
| **GATCCAGATCTAACTTGGGGTGGCTTTGTCTTCTTCTTTTGCCAATTCCACTAATTGTTTGCGAAGG** | Ins_k4_61_CGAA | 0.11377085830882383 | 0.04068053525184867 | 55.01632135444301 |
| **GATCCAGATCTAACTTGGGGTGGCTTTGTCTTCTTCTTTTGCCAATTCCACTAATTGTTTGCGAGG** | Ins_k3_61_CGA | -0.2898122031365841 | 0.042727402167520405 | 36.74664142416458 |
| **GATCCAGATCTAACTTGGGGTGGCTTTGTCTTCTTCTTTTGCCAATTCCACTAATTGTTTGCGCGG** | Ins_k3_61_CGC | -0.05823100870968756 | 0.0364609210472549 | 46.322510279721484 |
| **GATCCAGATCTAACTTGGGGTGGCTTTGTCTTCTTCTTTTGCCAATTCCACTAATTGTTTGCGG** | Ins_k1_61_C | 0.1444251127495575 | 0.036117637478188365 | 56.72892079074697 |
| **GATCCAGATCTAACTTGGGGTGGCTTTGTCTTCTTCTTTTGCCAATTCCACTAATTGTTTGCGGG** | Ins_k2_60_GC;Ins_k2_61_CG | -0.3401199225829669 | 0.04120712247951561 | 34.94373205375735 |
| **GATCCAGATCTAACTTGGGGTGGCTTTGTCTTCTTCTTTTGCCAATTCCACTAATTGTTTGCGGGG** | Ins_k3_60_GCG;Ins_k3_61_CGG | -0.3549588920375506 | 0.04163604027472284 | 34.42903133977859 |
| **GATCCAGATCTAACTTGGGGTGGCTTTGTCTTCTTCTTTTGCCAATTCCACTAATTGTTTGCGTGG** | Ins_k3_61_CGT | -0.30206584238271234 | 0.04478987721778239 | 36.299108887572494 |
| **GATCCAGATCTAACTTGGGGTGGCTTTGTCTTCTTCTTTTGCCAATTCCACTAATTGTTTGCTAGG** | Ins_k3_61_CTA | -1.6253705750087006 | 0.06498016859290877 | 9.664781062505545 |
| **GATCCAGATCTAACTTGGGGTGGCTTTGTCTTCTTCTTTTGCCAATTCCACTAATTGTTTGCTCGG** | Ins_k3_61_CTC | -0.22090888697758226 | 0.0346778031987647 | 39.367875764992824 |
| **GATCCAGATCTAACTTGGGGTGGCTTTGTCTTCTTCTTTTGCCAATTCCACTAATTGTTTGCTGG** | Ins_k2_61_CT | -0.4513712639007462 | 0.03965402278129506 | 31.264640763476656 |
| **GATCCAGATCTAACTTGGGGTGGCTTTGTCTTCTTCTTTTGCCAATTCCACTAATTGTTTGCTGGG** | Ins_k3_59_TGC;Ins_k3_60_GCT; | -0.09163843935883542 | 0.04260066212977113 | 44.80055803320279 |
| **GATCCAGATCTAACTTGGGGTGGCTTTGTCTTCTTCTTTTGCCAATTCCACTAATTGTTTGCTTGG** | Ins_k3_61_CTT | -0.43468856915654475 | 0.04000084643173883 | 31.79059418358039 |
| **GATCCAGATCTAACTTGGGGTGGCTTTGTCTTCTTCTTTTGCCAATTCCACTAATTGTTTGG** | Del_k1_61-61;Del_k1_62-62;Del_ | -0.2590928064594102 | 0.028019046980566883 | 37.892993553685706 |
| **GATCCAGATCTAACTTGGGGTGGCTTTGTCTTCTTCTTTTGCCAATTCCACTAATTGTTTGGA** | G63A | -1.8405175625284915 | 0.05859960527919116 | 7.793900746812479 |
| **GATCCAGATCTAACTTGGGGTGGCTTTGTCTTCTTCTTTTGCCAATTCCACTAATTGTTTGGAAAG** | Ins_k3_62_AAA | 0.6207792163036658 | 0.03311617805197093 | 91.34451626699003 |
| **GATCCAGATCTAACTTGGGGTGGCTTTGTCTTCTTCTTTTGCCAATTCCACTAATTGTTTGGAACG** | Ins_k3_62_AAC | -0.0579572337423121 | 0.03548029351419477 | 46.335193959619865 |
| **GATCCAGATCTAACTTGGGGTGGCTTTGTCTTCTTCTTTTGCCAATTCCACTAATTGTTTGGAAG** | Ins_k2_62_AA | 0.6974721004791411 | 0.038315430711267195 | 98.62562687369383 |
| **GATCCAGATCTAACTTGGGGTGGCTTTGTCTTCTTCTTTTGCCAATTCCACTAATTGTTTGGAAGG** | Ins_k3_61_GAA;Ins_k3_62_AAG | 0.04935857738455929 | 0.04021096097894131 | 51.584312873428964 |
| **GATCCAGATCTAACTTGGGGTGGCTTTGTCTTCTTCTTTTGCCAATTCCACTAATTGTTTGGAATG** | Ins_k3_62_AAT | -0.4632778803018369 | 0.05304420859850462 | 30.89459206467125 |
| **GATCCAGATCTAACTTGGGGTGGCTTTGTCTTCTTCTTTTGCCAATTCCACTAATTGTTTGGACAG** | Ins_k3_62_ACA | 0.3212576494553867 | 0.03402084410132072 | 67.70206517515595 |
| **GATCCAGATCTAACTTGGGGTGGCTTTGTCTTCTTCTTTTGCCAATTCCACTAATTGTTTGGACCG** | Ins_k3_62_ACC | 0.15435820525378663 | 0.036237702588851964 | 57.295222315106635 |
| **GATCCAGATCTAACTTGGGGTGGCTTTGTCTTCTTCTTTTGCCAATTCCACTAATTGTTTGGACG** | Ins_k2_62_AC | 0.08999266904972547 | 0.040335894030106414 | 53.723563483709725 |
| **GATCCAGATCTAACTTGGGGTGGCTTTGTCTTCTTCTTTTGCCAATTCCACTAATTGTTTGGACGAG** | Ins_k4_62_ACGA | 0.5889315850672752 | 0.034440516756818744 | 88.4812459924259 |
| **GATCCAGATCTAACTTGGGGTGGCTTTGTCTTCTTCTTTTGCCAATTCCACTAATTGTTTGGACGG** | Ins_k3_61_GAC;Ins_k3_62_ACG | 0.23732547021777783 | 0.03581815978139677 | 62.251616633162996 |
| **GATCCAGATCTAACTTGGGGTGGCTTTGTCTTCTTCTTTTGCCAATTCCACTAATTGTTTGGACTCG** | Ins_k4_62_ACTC | 0.1785642039819192 | 0.04152821022405342 | 58.69903215934668 |
| **GATCCAGATCTAACTTGGGGTGGCTTTGTCTTCTTCTTTTGCCAATTCCACTAATTGTTTGGACTG** | Ins_k3_62_ACT | -0.10857139286824563 | 0.043264233428801056 | 44.048338886000174 |
| **GATCCAGATCTAACTTGGGGTGGCTTTGTCTTCTTCTTTTGCCAATTCCACTAATTGTTTGGAG** | Ins_k1_62_A | 0.653055808433121 | 0.034169321402373484 | 94.3409024020274 |
| **GATCCAGATCTAACTTGGGGTGGCTTTGTCTTCTTCTTTTGCCAATTCCACTAATTGTTTGGAGAG** | Ins_k3_62_AGA | 0.5410734940692012 | 0.03384230124624488 | 84.34643374707524 |
| **GATCCAGATCTAACTTGGGGTGGCTTTGTCTTCTTCTTTTGCCAATTCCACTAATTGTTTGGAGCG** | Ins_k3_62_AGC | 0.043112915296075305 | 0.037829545416959744 | 51.26313870286274 |
| **GATCCAGATCTAACTTGGGGTGGCTTTGTCTTCTTCTTTTGCCAATTCCACTAATTGTTTGGAGG** | Ins_k2_61_GA;Ins_k2_62_AG | 0.2653065836665526 | 0.037745303789199575 | 64.01808480154612 |
| **GATCCAGATCTAACTTGGGGTGGCTTTGTCTTCTTCTTTTGCCAATTCCACTAATTGTTTGGAGGG** | Ins_k3_60_GGA;Ins_k3_61_GAG | -0.04761006084719932 | 0.04865704143189926 | 46.81712121522015 |
| **GATCCAGATCTAACTTGGGGTGGCTTTGTCTTCTTCTTTTGCCAATTCCACTAATTGTTTGGAGTG** | Ins_k3_62_AGT | -0.42666538009571875 | 0.055887711730507134 | 32.04668207813834 |
| **GATCCAGATCTAACTTGGGGTGGCTTTGTCTTCTTCTTTTGCCAATTCCACTAATTGTTTGGATAG** | Ins_k3_62_ATA | 0.20186897745286347 | 0.04300776477656904 | 60.083064446381414 |
| **GATCCAGATCTAACTTGGGGTGGCTTTGTCTTCTTCTTTTGCCAATTCCACTAATTGTTTGGATCG** | Ins_k3_62_ATC | 0.1872853620740803 | 0.03721311864801676 | 59.21319448526183 |
| **GATCCAGATCTAACTTGGGGTGGCTTTGTCTTCTTCTTTTGCCAATTCCACTAATTGTTTGGATG** | Ins_k2_62_AT | 0.04062674452648962 | 0.04654845817003828 | 51.13584808453548 |
| **GATCCAGATCTAACTTGGGGTGGCTTTGTCTTCTTCTTTTGCCAATTCCACTAATTGTTTGGATGG** | Ins_k3_61_GAT;Ins_k3_62_ATG | 0.12855685165619724 | 0.03617401863343285 | 55.835836054921465 |
| **GATCCAGATCTAACTTGGGGTGGCTTTGTCTTCTTCTTTTGCCAATTCCACTAATTGTTTGGATTG** | Ins_k3_62_ATT | -1.641377180369967 | 0.08243748070209994 | 9.511312260103564 |
| **GATCCAGATCTAACTTGGGGTGGCTTTGTCTTCTTCTTTTGCCAATTCCACTAATTGTTTGGC** | G63C | -3.99791669301029 | 0.1901103060056203 | 0.9011733358819344 |
| **GATCCAGATCTAACTTGGGGTGGCTTTGTCTTCTTCTTTTGCCAATTCCACTAATTGTTTGGCAAG** | Ins_k3_62_CAA | 0.3554890843017219 | 0.03619628768881389 | 70.05972685991233 |
| **GATCCAGATCTAACTTGGGGTGGCTTTGTCTTCTTCTTTTGCCAATTCCACTAATTGTTTGGCACG** | Ins_k3_62_CAC | -0.127331676591306 | 0.03584482329192941 | 43.22968267263051 |
| **GATCCAGATCTAACTTGGGGTGGCTTTGTCTTCTTCTTTTGCCAATTCCACTAATTGTTTGGCAG** | Ins_k2_62_CA | 0.4200162610289453 | 0.03398433642610601 | 74.7295275500055 |
| **GATCCAGATCTAACTTGGGGTGGCTTTGTCTTCTTCTTTTGCCAATTCCACTAATTGTTTGGCAGG** | Ins_k3_61_GCA;Ins_k3_62_CAG | -0.640448567356667 | 0.05189008073866573 | 25.878447189937944 |
| **GATCCAGATCTAACTTGGGGTGGCTTTGTCTTCTTCTTTTGCCAATTCCACTAATTGTTTGGCATG** | Ins_k3_62_CAT | -0.3217642984994624 | 0.04160356889696098 | 35.5910690250164 |
| **GATCCAGATCTAACTTGGGGTGGCTTTGTCTTCTTCTTTTGCCAATTCCACTAATTGTTTGGCCAG** | Ins_k3_62_CCA | 0.12760780819669126 | 0.03364434533937952 | 55.78287055716253 |
| **GATCCAGATCTAACTTGGGGTGGCTTTGTCTTCTTCTTTTGCCAATTCCACTAATTGTTTGGCCATG** | Ins_k4_62_CCAT | -0.41854770429372956 | 0.04637457841003263 | 32.307885401358924 |
| **GATCCAGATCTAACTTGGGGTGGCTTTGTCTTCTTCTTTTGCCAATTCCACTAATTGTTTGGCCCG** | Ins_k3_62_CCC | -0.17648360488614379 | 0.03459488651021621 | 41.15623481214383 |
| **GATCCAGATCTAACTTGGGGTGGCTTTGTCTTCTTCTTTTGCCAATTCCACTAATTGTTTGGCCCTG** | Ins_k4_62_CCCT | -0.33333659053174375 | 0.041015011993510306 | 35.18157275562357 |
| **GATCCAGATCTAACTTGGGGTGGCTTTGTCTTCTTCTTTTGCCAATTCCACTAATTGTTTGGCCG** | Ins_k2_62_CC | 0.12181046934495676 | 0.032129993838343 | 55.46041395264444 |
| **GATCCAGATCTAACTTGGGGTGGCTTTGTCTTCTTCTTTTGCCAATTCCACTAATTGTTTGGCCGG** | Ins_k3_61_GCC;Ins_k3_62_CCG | 0.006350654825155733 | 0.03681566443524681 | 49.41280937277066 |
| **GATCCAGATCTAACTTGGGGTGGCTTTGTCTTCTTCTTTTGCCAATTCCACTAATTGTTTGGCCTG** | Ins_k3_62_CCT | -0.2799315624394098 | 0.03984006380875278 | 37.111521440565674 |
| **GATCCAGATCTAACTTGGGGTGGCTTTGTCTTCTTCTTTTGCCAATTCCACTAATTGTTTGGCG** | Ins_k1_62_C | -0.13146092376043683 | 0.038250981551718216 | 43.05154466887046 |
| **GATCCAGATCTAACTTGGGGTGGCTTTGTCTTCTTCTTTTGCCAATTCCACTAATTGTTTGGCGAG** | Ins_k3_62_CGA | 0.44283087164318524 | 0.0323708454354804 | 76.4540499748145 |
| **GATCCAGATCTAACTTGGGGTGGCTTTGTCTTCTTCTTTTGCCAATTCCACTAATTGTTTGGCGCG** | Ins_k3_62_CGC | -0.20797026891453507 | 0.035362756960440334 | 39.880551177005856 |
| **GATCCAGATCTAACTTGGGGTGGCTTTGTCTTCTTCTTTTGCCAATTCCACTAATTGTTTGGCGG** | Ins_k2_61_GC;Ins_k2_62_CG | 0.08341208471064288 | 0.03542944104191828 | 53.37119172085517 |
| **GATCCAGATCTAACTTGGGGTGGCTTTGTCTTCTTCTTTTGCCAATTCCACTAATTGTTTGGCGGG** | Ins_k3_60_GGC;Ins_k3_61_GCG | -0.3587267290885895 | 0.04231652318104458 | 34.299552440741685 |
| **GATCCAGATCTAACTTGGGGTGGCTTTGTCTTCTTCTTTTGCCAATTCCACTAATTGTTTGGCGTG** | Ins_k3_62_CGT | -0.6910393985593677 | 0.048079175382609037 | 24.601800621144537 |
| **GATCCAGATCTAACTTGGGGTGGCTTTGTCTTCTTCTTTTGCCAATTCCACTAATTGTTTGGCGTTG** | Ins_k4_62_CGTT | -1.175589025339888 | 0.06652786691049747 | 15.15408313433563 |
| **GATCCAGATCTAACTTGGGGTGGCTTTGTCTTCTTCTTTTGCCAATTCCACTAATTGTTTGGCTAG** | Ins_k3_62_CTA | -0.004472160746721632 | 0.036459570670446494 | 48.88090718164566 |
| **GATCCAGATCTAACTTGGGGTGGCTTTGTCTTCTTCTTTTGCCAATTCCACTAATTGTTTGGCTCCG** | Ins_k4_62_CTCC | 0.07073863139279013 | 0.03492996026212926 | 52.69906250938018 |
| **GATCCAGATCTAACTTGGGGTGGCTTTGTCTTCTTCTTTTGCCAATTCCACTAATTGTTTGGCTCG** | Ins_k3_62_CTC | -0.02970876262399602 | 0.034916664687987355 | 47.662754851725516 |
| **GATCCAGATCTAACTTGGGGTGGCTTTGTCTTCTTCTTTTGCCAATTCCACTAATTGTTTGGCTG** | Ins_k2_62_CT | -0.11282628243140747 | 0.04382688187032461 | 43.86131623112675 |

| | | | | |
|---|---|---|---|---|
| GATCCAGATCTAACTTGGGGTGGCTTTGTCTTCTTCTTTTGCCAATTCCACTAATTGTTTGGCTGG | Ins_k3_61_GCT;Ins_k3_62_CTG | -0.32532355370395094 | 0.044020992934896454 | 35.46461649925585 |
| GATCCAGATCTAACTTGGGGTGGCTTTGTCTTCTTCTTTTGCCAATTCCACTAATTGTTTGGCTTG | Ins_k3_62_CTT | -0.771193575784318 | 0.04563174857825741 | 22.706823159049847 |
| GATCCAGATCTAACTTGGGGTGGCTTTGTCTTCTTCTTTTGCCAATTCCACTAATTGTTTGGCTTGG | Ins_k4_61_GCTT;Ins_k4_62_CTT | -0.3542100112137157 | 0.04432617233656313 | 34.454824237830096 |
| GATCCAGATCTAACTTGGGGTGGCTTTGTCTTCTTCTTTTGCCAATTCCACTAATTGTTTGGG | WT | 0 | 0.019335756304226577 | 49.1 |
| GATCCAGATCTAACTTGGGGTGGCTTTGTCTTCTTCTTTTGCCAATTCCACTAATTGTTTGGGAAG | Ins_k3_62_GAA | 0.5976575813977116 | 0.03523390004338329 | 89.25671145413814 |
| GATCCAGATCTAACTTGGGGTGGCTTTGTCTTCTTCTTTTGCCAATTCCACTAATTGTTTGGGACCG | Ins_k4_62_GACC | 0.05258863896841426 | 0.04167311748400608 | 51.75120276798859 |
| GATCCAGATCTAACTTGGGGTGGCTTTGTCTTCTTCTTTTGCCAATTCCACTAATTGTTTGGGACG | Ins_k3_62_GAC | -0.3729265332918809 | 0.04168573581821041 | 33.815947187703905 |
| GATCCAGATCTAACTTGGGGTGGCTTTGTCTTCTTCTTTTGCCAATTCCACTAATTGTTTGGGAG | Ins_k2_62_GA | 0.515523945723299 | 0.033364411936156774 | 82.21871730022897 |
| GATCCAGATCTAACTTGGGGTGGCTTTGTCTTCTTCTTTTGCCAATTCCACTAATTGTTTGGGAGG | Ins_k3_61_GGA;Ins_k3_62_GAG | -1.9436893455791284 | 0.1048794254157654 | 7.029880374311188 |
| GATCCAGATCTAACTTGGGGTGGCTTTGTCTTCTTCTTTTGCCAATTCCACTAATTGTTTGGGATG | Ins_k3_62_GAT | -0.6162352261437105 | 0.058909567570166035 | 26.51269854665104 |
| GATCCAGATCTAACTTGGGGTGGCTTTGTCTTCTTCTTTTGCCAATTCCACTAATTGTTTGGGCAG | Ins_k3_62_GCA | 0.3749326880579289 | 0.03829990767428634 | 71.43526984917119 |
| GATCCAGATCTAACTTGGGGTGGCTTTGTCTTCTTCTTTTGCCAATTCCACTAATTGTTTGGGCCG | Ins_k3_62_GCC | -0.042433561868139735 | 0.0408599768232211 | 47.06009833823731 |
| GATCCAGATCTAACTTGGGGTGGCTTTGTCTTCTTCTTTTGCCAATTCCACTAATTGTTTGGGCG | Ins_k2_62_GC | -0.7098181120878257 | 0.05075357524778893 | 24.144121219775 |
| GATCCAGATCTAACTTGGGGTGGCTTTGTCTTCTTCTTTTGCCAATTCCACTAATTGTTTGGGCGG | Ins_k3_61_GGC;Ins_k3_62_GCG | -0.9352777551469986 | 0.06740324538545465 | 19.27061274230987 |
| GATCCAGATCTAACTTGGGGTGGCTTTGTCTTCTTCTTTTGCCAATTCCACTAATTGTTTGGGCTG | Ins_k3_62_GCT | -0.6251402383280258 | 0.047651818343952344 | 26.277650748066485 |
| GATCCAGATCTAACTTGGGGTGGCTTTGTCTTCTTCTTTTGCCAATTCCACTAATTGTTTGGGG | Ins_k1_60_G;Ins_k1_61_G;Ins_k1 | -0.9592513558460166 | 0.03931371069770623 | 18.814120511669962 |
| GATCCAGATCTAACTTGGGGTGGCTTTGTCTTCTTCTTTTGCCAATTCCACTAATTGTTTGGGGAG | Ins_k3_62_GGA | 0.1278596542157543 | 0.0403049026377597 | 55.79692102024659 |
| GATCCAGATCTAACTTGGGGTGGCTTTGTCTTCTTCTTTTGCCAATTCCACTAATTGTTTGGGGCG | Ins_k3_62_GGC | -1.688483869124361 | 0.09468010104693363 | 9.07365505331393 |
| GATCCAGATCTAACTTGGGGTGGCTTTGTCTTCTTCTTTTGCCAATTCCACTAATTGTTTGGGGG | Ins_k2_60_GG;Ins_k2_61_GG;Ins | -2.8723060406386205 | 0.14117884755998386 | 2.777504865157159 |
| GATCCAGATCTAACTTGGGGTGGCTTTGTCTTCTTCTTTTGCCAATTCCACTAATTGTTTGGGGGG | Ins_k3_60_GGG;Ins_k3_61_GGG | -4.234070101802793 | 0.38055716972995507 | 0.7116201073875574 |
| GATCCAGATCTAACTTGGGGTGGCTTTGTCTTCTTCTTTTGCCAATTCCACTAATTGTTTGGGGTG | Ins_k3_62_GGT | -4.658367169398864 | 0.325342848812173 | 0.4655628689338704 |
| GATCCAGATCTAACTTGGGGTGGCTTTGTCTTCTTCTTTTGCCAATTCCACTAATTGTTTGGGTAG | Ins_k3_62_GTA | -1.964409395537037 | 0.10228930411619012 | 6.885719568864581 |
| GATCCAGATCTAACTTGGGGTGGCTTTGTCTTCTTCTTTTGCCAATTCCACTAATTGTTTGGGTCG | Ins_k3_62_GTC | -1.0914479430050352 | 0.06462524811298653 | 16.48434416180098 |
| GATCCAGATCTAACTTGGGGTGGCTTTGTCTTCTTCTTTTGCCAATTCCACTAATTGTTTGGGTG | Ins_k2_62_GT | -2.4363749473748904 | 0.13479151857833838 | 4.295139727167116 |
| GATCCAGATCTAACTTGGGGTGGCTTTGTCTTCTTCTTTTGCCAATTCCACTAATTGTTTGGGTGG | Ins_k3_61_GGT;Ins_k3_62_GTG | -3.6381632924025897 | 0.36069619767404854 | 1.2913597621366237 |
| GATCCAGATCTAACTTGGGGTGGCTTTGTCTTCTTCTTTTGCCAATTCCACTAATTGTTTGGGTTG | Ins_k3_62_GTT | -3.418096943556449 | 0.23766720944431133 | 1.6092401180983054 |
| GATCCAGATCTAACTTGGGGTGGCTTTGTCTTCTTCTTTTGCCAATTCCACTAATTGTTTGGT | G63T | -4.358128542147243 | 0.18449208676064047 | 0.6285941209026101 |
| GATCCAGATCTAACTTGGGGTGGCTTTGTCTTCTTCTTTTGCCAATTCCACTAATTGTTTGGTAAG | Ins_k3_62_TAA | -0.4265940890162163 | 0.037433036897180944 | 32.048966802137414 |
| GATCCAGATCTAACTTGGGGTGGCTTTGTCTTCTTCTTTTGCCAATTCCACTAATTGTTTGGTACG | Ins_k3_62_TAC | -1.4389591395808539 | 0.07094960604326403 | 11.645267743559176 |
| GATCCAGATCTAACTTGGGGTGGCTTTGTCTTCTTCTTTTGCCAATTCCACTAATTGTTTGGTAG | Ins_k2_62_TA | -1.2283259679423764 | 0.05324322261124448 | 14.375610665481586 |
| GATCCAGATCTAACTTGGGGTGGCTTTGTCTTCTTCTTTTGCCAATTCCACTAATTGTTTGGTAGG | Ins_k3_61_GTA;Ins_k3_62_TAG | -3.3286505428453883 | 0.22029336184402099 | 1.7598146516731912 |
| GATCCAGATCTAACTTGGGGTGGCTTTGTCTTCTTCTTTTGCCAATTCCACTAATTGTTTGGTATG | Ins_k3_62_TAT | -1.885854867348264 | 0.07432547335313747 | 7.448436670252322 |
| GATCCAGATCTAACTTGGGGTGGCTTTGTCTTCTTCTTTTGCCAATTCCACTAATTGTTTGGTCAG | Ins_k3_62_TCA | 0.04508571448597509 | 0.038404631090186686 | 51.36437040344364 |
| GATCCAGATCTAACTTGGGGTGGCTTTGTCTTCTTCTTTTGCCAATTCCACTAATTGTTTGGTCCG | Ins_k3_62_TCC | -0.16509909048348248 | 0.03749333376674672 | 41.62745578162196 |
| GATCCAGATCTAACTTGGGGTGGCTTTGTCTTCTTCTTTTGCCAATTCCACTAATTGTTTGGTCG | Ins_k2_62_TC | -0.44741958530118936 | 0.04477918903705124 | 31.388433007793353 |
| GATCCAGATCTAACTTGGGGTGGCTTTGTCTTCTTCTTTTGCCAATTCCACTAATTGTTTGGTCGG | Ins_k3_61_GTC;Ins_k3_62_TCG | -0.16771028392073795 | 0.037397822799951616 | 41.5189002337231 |
| GATCCAGATCTAACTTGGGGTGGCTTTGTCTTCTTCTTTTGCCAATTCCACTAATTGTTTGGTCTG | Ins_k3_62_TCT | -0.49051377554545617 | 0.0419035278721684 | 30.064505578029685 |
| GATCCAGATCTAACTTGGGGTGGCTTTGTCTTCTTCTTTTGCCAATTCCACTAATTGTTTGGTG | Ins_k1_62_T | -1.1444301351131616 | 0.07122867124787324 | 15.633700922708055 |
| GATCCAGATCTAACTTGGGGTGGCTTTGTCTTCTTCTTTTGCCAATTCCACTAATTGTTTGGTGAG | Ins_k3_62_TGA | 0.38517506195546547 | 0.03951953919455596 | 72.17069642017093 |
| GATCCAGATCTAACTTGGGGTGGCTTTGTCTTCTTCTTTTGCCAATTCCACTAATTGTTTGGTGCG | Ins_k3_62_TGC | -0.4439164712128152 | 0.03898094704283289 | 31.498583091184386 |
| GATCCAGATCTAACTTGGGGTGGCTTTGTCTTCTTCTTTTGCCAATTCCACTAATTGTTTGGTGCGG | Ins_k4_61_GTGC;Ins_k4_62_TG( | -0.18187834439444273 | 0.040031039467515 | 40.934805460052175 |
| GATCCAGATCTAACTTGGGGTGGCTTTGTCTTCTTCTTTTGCCAATTCCACTAATTGTTTGGTGG | Ins_k2_61_GT;Ins_k2_62_TG | -1.631696575755716 | 0.07281192064321436 | 9.603834627129771 |
| GATCCAGATCTAACTTGGGGTGGCTTTGTCTTCTTCTTTTGCCAATTCCACTAATTGTTTGGTGGG | Ins_k3_59_TGG;Ins_k3_60_GGT; | -1.4613158880875896 | 0.07232614594528797 | 11.387806147109263 |
| GATCCAGATCTAACTTGGGGTGGCTTTGTCTTCTTCTTTTGCCAATTCCACTAATTGTTTGGTGTG | Ins_k3_62_TGT | -1.3877668792592877 | 0.06288724146307798 | 12.256938141309819 |
| GATCCAGATCTAACTTGGGGTGGCTTTGTCTTCTTCTTTTGCCAATTCCACTAATTGTTTGGTTAG | Ins_k3_62_TTA | -0.4798440993801383 | 0.04935249759306754 | 30.38700152079545 |
| GATCCAGATCTAACTTGGGGTGGCTTTGTCTTCTTCTTTTGCCAATTCCACTAATTGTTTGGTTCG | Ins_k3_62_TTC | -0.40815196128381404 | 0.041553196017268264 | 32.645501720879835 |
| GATCCAGATCTAACTTGGGGTGGCTTTGTCTTCTTCTTTTGCCAATTCCACTAATTGTTTGGTTG | Ins_k2_62_TT | -1.0666294478351581 | 0.058251120885190674 | 16.898579868784456 |
| GATCCAGATCTAACTTGGGGTGGCTTTGTCTTCTTCTTTTGCCAATTCCACTAATTGTTTGGTTGG | Ins_k3_61_GTT;Ins_k3_62_TTG | -0.9245148531869898 | 0.050379564177817616 | 19.47914062757083 |
| GATCCAGATCTAACTTGGGGTGGCTTTGTCTTCTTCTTTTGCCAATTCCACTAATTGTTTGGTTTG | Ins_k3_62_TTT | -1.1038423562224053 | 0.05454393648377538 | 16.281291348312745 |
| GATCCAGATCTAACTTGGGGTGGCTTTGTCTTCTTCTTTTGCCAATTCCACTAATTGTTTGTAAGG | Ins_k3_61_TAA | -0.4789841231357824 | 0.050386677666311795 | 30.41314485995464 |
| GATCCAGATCTAACTTGGGGTGGCTTTGTCTTCTTCTTTTGCCAATTCCACTAATTGTTTGTACGG | Ins_k3_61_TAC | -0.16599031593726926 | 0.034074242687464375 | 41.590372860529115 |
| GATCCAGATCTAACTTGGGGTGGCTTTGTCTTCTTCTTTTGCCAATTCCACTAATTGTTTGTAGG | Ins_k2_61_TA | -1.3069283051710694 | 0.057750767738733264 | 13.288921652008343 |
| GATCCAGATCTAACTTGGGGTGGCTTTGTCTTCTTCTTTTGCCAATTCCACTAATTGTTTGTAGGG | Ins_k3_60_GTA;Ins_k3_61_TAG | -1.9336546752014185 | 0.08626033758130157 | 7.100778029043388 |
| GATCCAGATCTAACTTGGGGTGGCTTTGTCTTCTTCTTTTGCCAATTCCACTAATTGTTTGTATGG | Ins_k3_61_TAT | -0.5632015124007699 | 0.050403031717748584 | 27.956717829412234 |
| GATCCAGATCTAACTTGGGGTGGCTTTGTCTTCTTCTTTTGCCAATTCCACTAATTGTTTGTATTGG | Ins_k4_61_TATT | -0.6449210761757463 | 0.051844433573781414 | 25.762964048844296 |
| GATCCAGATCTAACTTGGGGTGGCTTTGTCTTCTTCTTTTGCCAATTCCACTAATTGTTTGTCAGG | Ins_k3_61_TCA | -0.6531538519714505 | 0.045359821780800225 | 25.551734039722053 |
| GATCCAGATCTAACTTGGGGTGGCTTTGTCTTCTTCTTTTGCCAATTCCACTAATTGTTTGTCCGG | Ins_k3_61_TCC | -0.23831543342333278 | 0.03776380815077697 | 38.688546547099385 |
| GATCCAGATCTAACTTGGGGTGGCTTTGTCTTCTTCTTTTGCCAATTCCACTAATTGTTTGTCGG | Ins_k2_61_TC | 0.03827993341060054 | 0.03697051025868201 | 51.015982613653485 |
| GATCCAGATCTAACTTGGGGTGGCTTTGTCTTCTTCTTTTGCCAATTCCACTAATTGTTTGTCGGG | Ins_k3_60_GTC;Ins_k3_61_TCG | -0.38547985471558655 | 0.045428518840539285 | 33.394098076923584 |
| GATCCAGATCTAACTTGGGGTGGCTTTGTCTTCTTCTTTTGCCAATTCCACTAATTGTTTGTCTGG | Ins_k3_61_TCT | -0.4811735015874173 | 0.047782275741130024 | 30.3466318136355 |
| GATCCAGATCTAACTTGGGGTGGCTTTGTCTTCTTCTTTTGCCAATTCCACTAATTGTTTGTG | G62T | -0.7972265804978653 | 0.03589841474990293 | 22.123324391368882 |
| GATCCAGATCTAACTTGGGGTGGCTTTGTCTTCTTCTTTTGCCAATTCCACTAATTGTTTGTGAGG | Ins_k3_61_TGA | -0.508356578312222 | 0.04709148511798894 | 29.53282794982952 |
| GATCCAGATCTAACTTGGGGTGGCTTTGTCTTCTTCTTTTGCCAATTCCACTAATTGTTTGTGCGG | Ins_k3_61_TGC | -0.028840180789145742 | 0.036806986513975466 | 47.704171839206275 |
| GATCCAGATCTAACTTGGGGTGGCTTTGTCTTCTTCTTTTGCCAATTCCACTAATTGTTTGTGG | Ins_k1_61_T | -0.4802778976612303 | 0.04165652070488585 | 30.373822550482192 |
| GATCCAGATCTAACTTGGGGTGGCTTTGTCTTCTTCTTTTGCCAATTCCACTAATTGTTTGTGGG | Ins_k2_59_TG;Ins_k2_60_GT;Ins_ | -0.21795100201034556 | 0.04348924936197247 | 39.484493799250295 |
| GATCCAGATCTAACTTGGGGTGGCTTTGTCTTCTTCTTTTGCCAATTCCACTAATTGTTTGTGGGG | Ins_k3_60_GTG;Ins_k3_61_TGG | -1.2974953906322833 | 0.07652182829720364 | 13.414868000947873 |
| GATCCAGATCTAACTTGGGGTGGCTTTGTCTTCTTCTTTTGCCAATTCCACTAATTGTTTGTGTGG | Ins_k3_61_TGT | -0.7821260609939512 | 0.05317438284610901 | 22.459933170284675 |
| GATCCAGATCTAACTTGGGGTGGCTTTGTCTTCTTCTTTTGCCAATTCCACTAATTGTTTGTGTTGG | Ins_k4_61_TGTT | -0.9384603980333528 | 0.055704043026803936 | 19.209378758389214 |
| GATCCAGATCTAACTTGGGGTGGCTTTGTCTTCTTCTTTTGCCAATTCCACTAATTGTTTGTTAGG | Ins_k3_61_TTA | -1.7731118418830178 | 0.08594788455574508 | 8.33736478081779 |
| GATCCAGATCTAACTTGGGGTGGCTTTGTCTTCTTCTTTTGCCAATTCCACTAATTGTTTGTTCGG | Ins_k3_61_TTC | -0.035637629281473515 | 0.03701258415633206 | 47.38100478828672 |
| GATCCAGATCTAACTTGGGGTGGCTTTGTCTTCTTCTTTTGCCAATTCCACTAATTGTTTGTTCTGG | Ins_k4_61_TTCT | -0.799539150776512 | 0.047354650064000504 | 22.072221760903386 |
| GATCCAGATCTAACTTGGGGTGGCTTTGTCTTCTTCTTTTGCCAATTCCACTAATTGTTTGTTGG | Ins_k2_61_TT | -0.36821352488526227 | 0.04507981592890555 | 33.97569818927906 |
| GATCCAGATCTAACTTGGGGTGGCTTTGTCTTCTTCTTTTGCCAATTCCACTAATTGTTTGTTGGG | Ins_k3_58_TTG;Ins_k3_59_TGT;Ii | -0.3887119799535004 | 0.05124763252579183 | 33.28633840991194 |
| GATCCAGATCTAACTTGGGGTGGCTTTGTCTTCTTCTTTTGCCAATTCCACTAATTGTTTGTTTGG | Ins_k3_61_TTT | -0.3260434159300828 | 0.04278246569467383 | 35.439096048180204 |
| GATCCAGATCTAACTTGGGGTGGCTTTGTCTTCTTCTTTTGCCAATTCCACTAATTGTTTTAAGGG | Ins_k3_60_TAA | -0.6832814745209137 | 0.054430054511484034 | 24.793401774203318 |

| | | | | |
|---|---|---|---|---|
| GATCCAGATCTAACTTGGGGTGGCTTTGTCTTCTTCTTTTGCCAATTCCACTAATTGTTTTACGGG | Ins_k3_60_TAC | -0.3471255757807209 | 0.03798456413052577 | 34.69978389139579 |
| GATCCAGATCTAACTTGGGGTGGCTTTGTCTTCTTCTTTTGCCAATTCCACTAATTGTTTTAGGG | Ins_k2_60_TA | -1.4808080967535686 | 0.07197850191272485 | 11.167982042170742 |
| GATCCAGATCTAACTTGGGGTGGCTTTGTCTTCTTCTTTTGCCAATTCCACTAATTGTTTTAGGGG | Ins_k3_60_TAG | -2.826430267530344 | 0.15347286617710143 | 2.9078930096657074 |
| GATCCAGATCTAACTTGGGGTGGCTTTGTCTTCTTCTTTTGCCAATTCCACTAATTGTTTTATGGG | Ins_k3_59_TTA;Ins_k3_60_TAT | -0.49428312075662256 | 0.05318176790781542 5 | 29.951395387502636 |
| GATCCAGATCTAACTTGGGGTGGCTTTGTCTTCTTCTTTTGCCAATTCCACTAATTGTTTTCAGGG | Ins_k3_60_TCA | -1.123885491215981 | 0.05708627771129729 | 15.958211807126764 |
| GATCCAGATCTAACTTGGGGTGGCTTTGTCTTCTTCTTTTGCCAATTCCACTAATTGTTTTCCGGG | Ins_k3_60_TCC | -1.004063877413387 | 0.05361066207459178 | 17.989624182474923 |
| GATCCAGATCTAACTTGGGGTGGCTTTGTCTTCTTCTTTTGCCAATTCCACTAATTGTTTTCGGG | Ins_k2_60_TC | -0.35808860898125494 | 0.03754734254238982 | 34.32144665965442 |
| GATCCAGATCTAACTTGGGGTGGCTTTGTCTTCTTCTTTTGCCAATTCCACTAATTGTTTTCGGGG | Ins_k3_60_TCG | -1.2858746812037931 | 0.0669047548152793 | 13.571667580660048 |
| GATCCAGATCTAACTTGGGGTGGCTTTGTCTTCTTCTTTTGCCAATTCCACTAATTGTTTTCTGGG | Ins_k3_59_TTC;Ins_k3_60_TCT | -0.8019233169380532 | 0.04889740516242934 | 22.019660598840563 |
| GATCCAGATCTAACTTGGGGTGGCTTTGTCTTCTTCTTTTGCCAATTCCACTAATTGTTTTGAGGG | Ins_k3_60_TGA | -0.6824279765321085 | 0.06338796089553099 | 24.814571925809016 |
| GATCCAGATCTAACTTGGGGTGGCTTTGTCTTCTTCTTTTGCCAATTCCACTAATTGTTTTGCGGG | Ins_k3_60_TGC | -0.533598897887277 | 0.044799521379372674 | 28.796680984675685 |
| GATCCAGATCTAACTTGGGGTGGCTTTGTCTTCTTCTTTTGCCAATTCCACTAATTGTTTTGG | G61T | -0.4885152852779492 | 0.033295220143743684 | 30.12464927810677 |
| GATCCAGATCTAACTTGGGGTGGCTTTGTCTTCTTCTTTTGCCAATTCCACTAATTGTTTTGGG | Ins_k1_57_T;Ins_k1_58_T;Ins_k1_ | -0.1418149302272454 | 0.03851950953189197 | 42.60808843306621 |
| GATCCAGATCTAACTTGGGGTGGCTTTGTCTTCTTCTTTTGCCAATTCCACTAATTGTTTTGGGG | Ins_k2_60_TG | -1.0646492540477221 | 0.0638278374408924 | 16.932075484617144 |
| GATCCAGATCTAACTTGGGGTGGCTTTGTCTTCTTCTTTTGCCAATTCCACTAATTGTTTTGGGGG | Ins_k3_60_TGG | -3.32852932319222 | 0.20746866505139225 | 1.7600279887249677 |
| GATCCAGATCTAACTTGGGGTGGCTTTGTCTTCTTCTTTTGCCAATTCCACTAATTGTTTTGTGGG | Ins_k3_59_TTG;Ins_k3_60_TGT | -0.8261226841474169 | 0.0598049955780484 | 21.493194510980924 |
| GATCCAGATCTAACTTGGGGTGGCTTTGTCTTCTTCTTTTGCCAATTCCACTAATTGTTTTTAGGG | Ins_k3_60_TTA | -1.711471897514438 | 0.09474120132715932 | 8.867448831306502 |
| GATCCAGATCTAACTTGGGGTGGCTTTGTCTTCTTCTTTTGCCAATTCCACTAATTGTTTTTCGGG | Ins_k3_60_TTC | -0.6264873723113525 | 0.047863359755365445 | 26.2422750649841 |
| GATCCAGATCTAACTTGGGGTGGCTTTGTCTTCTTCTTTTGCCAATTCCACTAATTGTTTTTGGG | Ins_k2_57_TT;Ins_k2_58_TT;Ins_ | -0.4180310176937978 | 0.04055372847589805 | 32.32458276609877 |
| GATCCAGATCTAACTTGGGGTGGCTTTGTCTTCTTCTTTTGCCAATTCCACTAATTGTTTTTGGGG | Ins_k3_60_TTG | -1.3241803783085766 | 0.0754820624107462 | 13.061626497599512 |
| GATCCAGATCTAACTTGGGGTGGCTTTGTCTTCTTCTTTTGCCAATTCCACTAATTGTTTTTTGGG | Ins_k3_57_TTT;Ins_k3_58_TTT;In | -0.5002459153172791 | 0.05032914564184142 | 29.773332772979806 |
| GATCCAGATCTAACTTGGGGTGGCTTTGTCTTCTTCTTTTGCCAATTCCACTAATTTAAGGTTTGGG | Ins_k4_56_TAAG | -1.2649399412715057 | 0.08414470309277855 | 13.858781756316295 |
| GATCCAGATCTAACTTGGGGTGGCTTTGTCTTCTTCTTTTGCCAATTCCACTAATTTAAGTTTGGG | Ins_k3_56_TAA | -0.5921764016081866 | 0.05684944633301444 | 27.158297949762574 |
| GATCCAGATCTAACTTGGGGTGGCTTTGTCTTCTTCTTTTGCCAATTCCACTAATTTACGTTTGGG | Ins_k3_56_TAC | -0.22784058622192127 | 0.04088394717076211 | 39.09593309158118 |
| GATCCAGATCTAACTTGGGGTGGCTTTGTCTTCTTCTTTTGCCAATTCCACTAATTTAGGTTTGGG | Ins_k3_56_TAG | -3.07276590718141 | 0.16060346361215352 | 2.2729831406502266 |
| GATCCAGATCTAACTTGGGGTGGCTTTGTCTTCTTCTTTTGCCAATTCCACTAATTTAGTTTGGG | Ins_k2_56_TA | -1.2495996712956368 | 0.07789797180381204 | 14.073018231448314 |
| GATCCAGATCTAACTTGGGGTGGCTTTGTCTTCTTCTTTTGCCAATTCCACTAATTTATGTTTGGG | Ins_k3_55_TTA;Ins_k3_56_TAT | -0.41616275422553056 | 0.05116286431565498 | 32.38503005135291 |
| GATCCAGATCTAACTTGGGGTGGCTTTGTCTTCTTCTTTTGCCAATTCCACTAATTTCAGTTTGGG | Ins_k3_56_TCA | -0.10897974563377438 | 0.039996438466305725 | 44.030355297073164 |
| GATCCAGATCTAACTTGGGGTGGCTTTGTCTTCTTCTTTTGCCAATTCCACTAATTTCCGTTTGGG | Ins_k3_56_TCC | 0.020813713667105382 | 0.03761515993309525 | 50.13266283556394 |
| GATCCAGATCTAACTTGGGGTGGCTTTGTCTTCTTCTTTTGCCAATTCCACTAATTTCGGTTTGGG | Ins_k3_56_TCG | -0.17694658238575323 | 0.04082250866266685 | 41.13718481165824 |
| GATCCAGATCTAACTTGGGGTGGCTTTGTCTTCTTCTTTTGCCAATTCCACTAATTTCGTTTGGG | Ins_k2_56_TC | 0.12340445177900217 | 0.042708118609228254 | 55.54888737207544 |
| GATCCAGATCTAACTTGGGGTGGCTTTGTCTTCTTCTTTTGCCAATTCCACTAATTTCTGTTTGGG | Ins_k3_55_TTC;Ins_k3_56_TCT | -0.5448520947261769 | 0.04526495473027343 | 28.474442770803762 |
| GATCCAGATCTAACTTGGGGTGGCTTTGTCTTCTTCTTTTGCCAATTCCACTAATTTGAGTTTGGG | Ins_k3_56_TGA | 0.036664600378823575 | 0.04383671695283566 | 50.933641333987985 |
| GATCCAGATCTAACTTGGGGTGGCTTTGTCTTCTTCTTTTGCCAATTCCACTAATTTGCAGTTTGGG | Ins_k4_56_TGCA | -1.1812296709302905 | 0.0755780369239559 | 15.068844947330176 |
| GATCCAGATCTAACTTGGGGTGGCTTTGTCTTCTTCTTTTGCCAATTCCACTAATTTGCGTTTGGG | Ins_k3_56_TGC | -0.004553862482849391 | 0.04076201571475372 | 48.87691368980525 |
| GATCCAGATCTAACTTGGGGTGGCTTTGTCTTCTTCTTTTGCCAATTCCACTAATTTGGG | Del_k3_55-57;Del_k3_56-58;Del_ | -0.35049162499018693 | 0.033071962173536584 | 34.58317907013988 |
| GATCCAGATCTAACTTGGGGTGGCTTTGTCTTCTTCTTTTGCCAATTCCACTAATTTGGGTTTGGG | Ins_k3_56_TGG | -1.5860912979035717 | 0.08043177299226228 | 10.051960970326556 |
| GATCCAGATCTAACTTGGGGTGGCTTTGTCTTCTTCTTTTGCCAATTCCACTAATTTGGTTTGGG | Ins_k2_56_TG | -0.4324349169044629 | 0.04882788196627043 | 31.862319919833144 |
| GATCCAGATCTAACTTGGGGTGGCTTTGTCTTCTTCTTTTGCCAATTCCACTAATTTGTGTTTGGG | Ins_k3_55_TTG;Ins_k3_56_TGT | -0.4746475714451825 | 0.05019244498290649 | 30.545319418480172 |
| GATCCAGATCTAACTTGGGGTGGCTTTGTCTTCTTCTTTTGCCAATTCCACTAATTTGTTTGGG | Ins_k1_54_T;Ins_k1_55_T;Ins_k1_ | -0.05101823196412769 | 0.03907389651890883 | 46.65783205146724 |
| GATCCAGATCTAACTTGGGGTGGCTTTGTCTTCTTCTTTTGCCAATTCCACTAATTTTAGTTTGGG | Ins_k3_56_TTA | -1.5815980106378325 | 0.07897727100504802 | 10.097228943391746 |
| GATCCAGATCTAACTTGGGGTGGCTTTGTCTTCTTCTTTTGCCAATTCCACTAATTTTCGTTTGGG | Ins_k3_56_TTC | -0.2479340139115382 | 0.041254875766327706 | 38.31820159977191 |
| GATCCAGATCTAACTTGGGGTGGCTTTGTCTTCTTCTTTTGCCAATTCCACTAATTTTGGG | Del_k2_56-57;Del_k2_57-58 | 0.23715448890020552 | 0.03510896544716171 | 62.2409736796291 |
| GATCCAGATCTAACTTGGGGTGGCTTTGTCTTCTTCTTTTGCCAATTCCACTAATTTTGGTTTGGG | Ins_k3_56_TTG | -0.6668793954479681 | 0.06279240129736832 | 25.203418481689706 |
| GATCCAGATCTAACTTGGGGTGGCTTTGTCTTCTTCTTTTGCCAATTCCACTAATTTTGTGTTTGGG | Ins_k4_55_TTTG;Ins_k4_56_TTG | -0.8046314649948296 | 0.06049676403087199 | 21.96010877175775 |
| GATCCAGATCTAACTTGGGGTGGCTTTGTCTTCTTCTTTTGCCAATTCCACTAATTTTGTTTGGG | Ins_k2_54_TT;Ins_k2_55_TT;Ins_ | -0.19932493031156373 | 0.04488512123223902 | 40.22682672350784 |
| GATCCAGATCTAACTTGGGGTGGCTTTGTCTTCTTCTTTTGCCAATTCCACTAATTTTTGGG | Del_k1_57-57 | -0.0016591047744426435 | 0.03506381357434407 | 49.01860549525132 |
| GATCCAGATCTAACTTGGGGTGGCTTTGTCTTCTTCTTTTGCCAATTCCACTAATTTTTGTTTGGG | Ins_k3_54_TTT;Ins_k3_55_TTT;In | -0.8768271212972363 | 0.05744542072197247 | 20.430561928211645 |
| GATCCAGATCTAACTTGGGGTGGCTTTGTCTTCTTCTTTTGCCAATTCCACTAATTTTTTGGG | G57T | -0.2503161201837696 | 0.03330940555250029 | 38.22703220210867 |
| GATCCAGATCTAACTTGGGGTGGCTTTGTCTTCTTCTTTTGCCAATTCCACTACAAATTGTTTGGG | Ins_k3_52_ACA;Ins_k3_53_CAA | 0.49405231698064433 | 0.03148040314022594 | 80.47216528556028 |
| GATCCAGATCTAACTTGGGGTGGCTTTGTCTTCTTCTTTTGCCAATTCCACTACAATTGTTTGGG | Ins_k2_52_AC;Ins_k2_53_CA | 0.45564161260557956 | 0.02986042780764983 | 77.43978351196064 |
| GATCCAGATCTAACTTGGGGTGGCTTTGTCTTCTTCTTTTGCCAATTCCACTACACATTGTTTGGG | Ins_k3_53_CAC | 0.2590352389072167 | 0.03349343618425541 | 63.61786160148023 |
| GATCCAGATCTAACTTGGGGTGGCTTTGTCTTCTTCTTTTGCCAATTCCACTACAGATTGTTTGGG | Ins_k3_53_CAG | 0.49773135108162214 | 0.03135367291929498 | 80.7687704014403 |
| GATCCAGATCTAACTTGGGGTGGCTTTGTCTTCTTCTTTTGCCAATTCCACTACATATTGTTTGGG | Ins_k3_53_CAT | 0.3972693228081745 | 0.03296905066512539 | 73.04884723713795 |
| GATCCAGATCTAACTTGGGGTGGCTTTGTCTTCTTCTTTTGCCAATTCCACTACATTGTTTGGG | Ins_k1_53_C | 0.5408860027123734 | 0.03252822915396044 | 84.33062100219124 |
| GATCCAGATCTAACTTGGGGTGGCTTTGTCTTCTTCTTTTGCCAATTCCACTACCAATTGTTTGGG | Ins_k3_52_ACC;Ins_k3_53_CCA | 0.16759282439214185 | 0.035255684947366195 | 58.05854274689965 |
| GATCCAGATCTAACTTGGGGTGGCTTTGTCTTCTTCTTTTGCCAATTCCACTACCATTGTTTGGG | Ins_k2_53_CC | 0.38942894037797315 | 0.029716067631207235 | 72.47835569705299 |
| GATCCAGATCTAACTTGGGGTGGCTTTGTCTTCTTCTTTTGCCAATTCCACTACCCATTGTTTGGG | Ins_k3_53_CCC | -0.017251318765534873 | 0.037359955693127396 | 48.26022469126757 |
| GATCCAGATCTAACTTGGGGTGGCTTTGTCTTCTTCTTTTGCCAATTCCACTACCGATTGTTTGGG | Ins_k3_53_CCG | 0.6984242287805922 | 0.03615585919807581 | 98.71957584292744 |
| GATCCAGATCTAACTTGGGGTGGCTTTGTCTTCTTCTTTTGCCAATTCCACTACCTATTGTTTGGG | Ins_k3_53_CCT | 0.018529193394586474 | 0.04211836167905053 | 50.01826447344351 |
| GATCCAGATCTAACTTGGGGTGGCTTTGTCTTCTTCTTTTGCCAATTCCACTACGAATTGTTTGGG | Ins_k3_52_ACG;Ins_k3_53_CGA | 0.5117172205666398 | 0.03383081849448403 | 81.90632820768244 |
| GATCCAGATCTAACTTGGGGTGGCTTTGTCTTCTTCTTTTGCCAATTCCACTACGATTGTTTGGG | Ins_k2_53_CG | 0.8886141345451333 | 0.029755780578590362 | 100 |
| GATCCAGATCTAACTTGGGGTGGCTTTGTCTTCTTCTTTTGCCAATTCCACTACGCATTGTTTGGG | Ins_k3_53_CGC | 0.8503700673811294 | 0.032790630518646155 | 100 |
| GATCCAGATCTAACTTGGGGTGGCTTTGTCTTCTTCTTTTGCCAATTCCACTACGGATTGTTTGGG | Ins_k3_53_CGG | 1.1102170451131617 | 0.03369043781799568 | 100 |
| GATCCAGATCTAACTTGGGGTGGCTTTGTCTTCTTCTTTTGCCAATTCCACTACGTATTGTTTGGG | Ins_k3_53_CGT | 0.6106074268664243 | 0.03483381990027694 | 90.4200885935992 |
| GATCCAGATCTAACTTGGGGTGGCTTTGTCTTCTTCTTTTGCCAATTCCACTACTAATTGTTTGGG | Ins_k3_49_ACT;Ins_k3_50_CTA;I | 0.1176420275600184 | 0.0376545807868815 | 55.22971161440265 |
| GATCCAGATCTAACTTGGGGTGGCTTTGTCTTCTTCTTTTGCCAATTCCACTACTATTGTTTGGG | Ins_k2_53_CT | 0.2770335381976059 | 0.03159427248573745 | 64.77324115914965 |
| GATCCAGATCTAACTTGGGGTGGCTTTGTCTTCTTCTTTTGCCAATTCCACTACTCATTGTTTGGG | Ins_k3_53_CTC | 0.3657085014816964 | 0.03290134528909123 | 70.77936732623199 |
| GATCCAGATCTAACTTGGGGTGGCTTTGTCTTCTTCTTTTGCCAATTCCACTACTGATTGTTTGGG | Ins_k3_53_CTG | 0.5614573442428674 | 0.03236010629826014 | 86.08338151519601 |
| GATCCAGATCTAACTTGGGGTGGCTTTGTCTTCTTCTTTTGCCAATTCCACTACTTATTGTTTGGG | Ins_k3_53_CTT | 0.06951642557111248 | 0.03907702246967841 | 52.634692752942684 |
| GATCCAGATCTAACTTGGGGTGGCTTTGTCTTCTTCTTTTGCCAATTCCACTACTTGTTTGGG | A54C | 0.2768401878137946 | 0.03642459882333731 | 64.76071843878623 |
| GATCCAGATCTAACTTGGGGTGGCTTTGTCTTCTTCTTTTGCCAATTCCACTAG | Del_k9_54-62 | -0.10576205219645618 | 0.02776120491954372 | 44.17225966229321 |
| GATCCAGATCTAACTTGGGGTGGCTTTGTCTTCTTCTTTTGCCAATTCCACTAGAAATTGTTTGGG | Ins_k3_52_AGA;Ins_k3_53_GAA | 0.06974588564789863 | 0.04014716997880971 | 52.64677169934838 |
| GATCCAGATCTAACTTGGGGTGGCTTTGTCTTCTTCTTTTGCCAATTCCACTAGAATTGTTTGGG | Ins_k2_52_AG;Ins_k2_53_GA | 0.013854957666583623 | 0.04052137865755905 | 49.785012875747405 |
| GATCCAGATCTAACTTGGGGTGGCTTTGTCTTCTTCTTTTGCCAATTCCACTAGACATTGTTTGGG | Ins_k3_53_GAC | -0.0405029793545818 | 0.042265361754644765 | 47.151039497635544 |

| | | | | |
|---|---|---|---|---|
| GATCCAGATCTAACTTGGGGTGGCTTTGTCTTCTTCTTTTGCCAATTCCACTAGAGATTGTTTGGG | Ins_k3_53_GAG | 0.37474374729295856 | 0.03527235853277652 | 71.42177408962975 |
| GATCCAGATCTAACTTGGGGTGGCTTTGTCTTCTTCTTTTGCCAATTCCACTAGATATTGTTTGGG | Ins_k3_53_GAT | -0.5573228555321421 | 0.05004755002209295 | 28.12154980130961 |
| GATCCAGATCTAACTTGGGGTGGCTTTGTCTTCTTCTTTTGCCAATTCCACTAGATTGTTTGGG | Ins_k1_53_G | 0.09439526934349707 | 0.04290907389467147 | 53.96060828395486 |
| GATCCAGATCTAACTTGGGGTGGCTTTGTCTTCTTCTTTTGCCAATTCCACTAGCAATTGTTTGGG | Ins_k3_52_AGC;Ins_k3_53_GCA | -0.32714706916847774 | 0.04110637505852528 | 35.40000515042667 |
| GATCCAGATCTAACTTGGGGTGGCTTTGTCTTCTTCTTTTGCCAATTCCACTAGCATTGTTTGGG | Ins_k2_53_GC | -0.3801843150289625 | 0.04386441144068908 | 33.5714069072108 |
| GATCCAGATCTAACTTGGGGTGGCTTTGTCTTCTTCTTTTGCCAATTCCACTAGCCATTGTTTGGG | Ins_k3_53_GCC | -0.033225815795601465 | 0.043447378371364184 | 47.495416849425524 |
| GATCCAGATCTAACTTGGGGTGGCTTTGTCTTCTTCTTTTGCCAATTCCACTAGCGATTGTTTGGG | Ins_k3_53_GCG | 0.14205626912234026 | 0.0407324038648017 | 56.5946978875524 |
| GATCCAGATCTAACTTGGGGTGGCTTTGTCTTCTTCTTTTGCCAATTCCACTAGCGTATTGTTTGGG | Ins_k4_53_GCGT | -0.14738087122173354 | 0.04399945360564839 | 42.37159309704858 |
| GATCCAGATCTAACTTGGGGTGGCTTTGTCTTCTTCTTTTGCCAATTCCACTAGCTATTGTTTGGG | Ins_k3_53_GCT | -1.0184222418181017 | 0.05353844089523242 | 17.73316815347378 |
| GATCCAGATCTAACTTGGGGTGGCTTTGTCTTCTTCTTTTGCCAATTCCACTAGG | Del_k8_54-61 | -0.49609826700601345 | 0.035136693757249024 | 29.897078535929094 |
| GATCCAGATCTAACTTGGGGTGGCTTTGTCTTCTTCTTTTGCCAATTCCACTAGGAATTGTTTGGG | Ins_k3_52_AGG;Ins_k3_53_GGA | -0.047683891782612786 | 0.21464394304761317 | 46.81366479096455 |
| GATCCAGATCTAACTTGGGGTGGCTTTGTCTTCTTCTTTTGCCAATTCCACTAGGATTGTTTGGG | Ins_k2_53_GG | 0.5812366574991663 | 0.039822290375728854 | 87.8030020789452 |
| GATCCAGATCTAACTTGGGGTGGCTTTGTCTTCTTCTTTTGCCAATTCCACTAGGCATTGTTTGGG | Ins_k3_53_GGC | -0.4337747254680572 | 0.04437752414251695 | 31.819659095798915 |
| GATCCAGATCTAACTTGGGGTGGCTTTGTCTTCTTCTTTTGCCAATTCCACTAGGG | Del_k7_54-60 | -0.5224684042802128 | 0.03415095096957126 | 29.118992674847746 |
| GATCCAGATCTAACTTGGGGTGGCTTTGTCTTCTTCTTTTGCCAATTCCACTAGGGATTGTTTGGG | Ins_k3_53_GGG | -1.2170390123896837 | 0.07379993385855256 | 14.538786692066568 |
| GATCCAGATCTAACTTGGGGTGGCTTTGTCTTCTTCTTTTGCCAATTCCACTAGGTATTGTTTGGG | Ins_k3_53_GGT | -2.2200767161165444 | 0.10318091300262629 | 5.332298154407119 |
| GATCCAGATCTAACTTGGGGTGGCTTTGTCTTCTTCTTTTGCCAATTCCACTAGTAATTGTTTGGG | Ins_k3_51_TAG;Ins_k3_52_AGT;I | -0.9697663667980696 | 0.06093250136540898 | 18.61732628836011 |
| GATCCAGATCTAACTTGGGGTGGCTTTGTCTTCTTCTTTTGCCAATTCCACTAGTATTGTTTGGG | Ins_k2_53_GT | -0.48282417995748395 | 0.05409725144459874 | 30.296580605484408 |
| GATCCAGATCTAACTTGGGGTGGCTTTGTCTTCTTCTTTTGCCAATTCCACTAGTCATTGTTTGGG | Ins_k3_53_GTC | -0.16536426123798892 | 0.04098301604441308 | 41.616418861163105 |
| GATCCAGATCTAACTTGGGGTGGCTTTGTCTTCTTCTTTTGCCAATTCCACTAGTGATTGTTTGGG | Ins_k3_53_GTG | 0.08025143214184405 | 0.04008226170550242 | 53.2027702277622 |
| GATCCAGATCTAACTTGGGGTGGCTTTGTCTTCTTCTTTTGCCAATTCCACTAGTGGATTGTTTGGG | Ins_k4_53_GTGG | 0.5397216753324376 | 0.040558464821662414 | 84.2324896907638 |
| GATCCAGATCTAACTTGGGGTGGCTTTGTCTTCTTCTTTTGCCAATTCCACTAGTTATTGTTTGGG | Ins_k3_53_GTT | -1.1818978065705523 | 0.05793962258284156 | 15.058780277619968 |
| GATCCAGATCTAACTTGGGGTGGCTTTGTCTTCTTCTTTTGCCAATTCCACTAGTTCATTGTTTGGG | Ins_k4_53_GTTC | -0.3121409316071006 | 0.04215360051222221 | 35.93522826965413 |
| GATCCAGATCTAACTTGGGGTGGCTTTGTCTTCTTCTTTTGCCAATTCCACTAGTTGTTTGGG | A54G | -0.09165827915547431 | 0.030735055962139303 | 44.799669208059186 |
| GATCCAGATCTAACTTGGGGTGGCTTTGTCTTCTTCTTTTGCCAATTCCACTAGTTTGGG | Del_k3_54-56 | -0.43399731210100523 | 0.03501470416646842 | 31.812577253210215 |
| GATCCAGATCTAACTTGGGGTGGCTTTGTCTTCTTCTTTTGCCAATTCCACTATAAATTGTTTGGG | Ins_k3_52_ATA;Ins_k3_53_TAA | -0.2526961868148507 | 0.052269675275560035 | 38.13615750517434 |
| GATCCAGATCTAACTTGGGGTGGCTTTGTCTTCTTCTTTTGCCAATTCCACTATAATTGTTTGGG | Ins_k2_51_TA;Ins_k2_52_AT;Ins_ | -0.26265263716206677 | 0.04309517668011215 | 37.75834072465301 |
| GATCCAGATCTAACTTGGGGTGGCTTTGTCTTCTTCTTTTGCCAATTCCACTATACATTGTTTGGG | Ins_k3_53_TAC | 0.0369373150701261 | 0.03652339876072704 | 50.94753358048505 |
| GATCCAGATCTAACTTGGGGTGGCTTTGTCTTCTTCTTTTGCCAATTCCACTATAGATTGTTTGGG | Ins_k3_53_TAG | -0.5592253130465195 | 0.05652051766829603 | 28.06810060610662 |
| GATCCAGATCTAACTTGGGGTGGCTTTGTCTTCTTCTTTTGCCAATTCCACTATATATTGTTTGGG | Ins_k3_53_TAT | -0.270254989681484 | 0.04503031904594921 | 37.472376884167744 |
| GATCCAGATCTAACTTGGGGTGGCTTTGTCTTCTTCTTTTGCCAATTCCACTATATTGTTTGGG | Ins_k1_53_T | 0.03765715447550744 | 0.03715340318224843 | 50.984220825641465 |
| GATCCAGATCTAACTTGGGGTGGCTTTGTCTTCTTCTTTTGCCAATTCCACTATCAATTGTTTGGG | Ins_k3_52_ATC;Ins_k3_53_TCA | 0.12188720142309228 | 0.04030349138539744 | 55.46466970873569 |
| GATCCAGATCTAACTTGGGGTGGCTTTGTCTTCTTCTTTTGCCAATTCCACTATCATTGTTTGGG | Ins_k2_53_TC | 0.11722791982767644 | 0.03622816801834149 | 55.20684529865397 |
| GATCCAGATCTAACTTGGGGTGGCTTTGTCTTCTTCTTTTGCCAATTCCACTATCCATTGTTTGGG | Ins_k3_53_TCC | 0.06049064067287224 | 0.035510938938853154 | 52.16176083943509 |
| GATCCAGATCTAACTTGGGGTGGCTTTGTCTTCTTCTTTTGCCAATTCCACTATCGATTGTTTGGG | Ins_k3_53_TCG | 0.28027742376282094 | 0.03557098748615286 | 64.98369930768246 |
| GATCCAGATCTAACTTGGGGTGGCTTTGTCTTCTTCTTTTGCCAATTCCACTATCTATTGTTTGGG | Ins_k3_53_TCT | -0.7287429146733142 | 0.04937389595636841 | 23.69149493251478 |
| GATCCAGATCTAACTTGGGGTGGCTTTGTCTTCTTCTTTTGCCAATTCCACTATGAATTGTTTGGG | Ins_k3_52_ATG;Ins_k3_53_TGA | 0.30068150767743584 | 0.033544269928906134 | 66.32325185880713 |
| GATCCAGATCTAACTTGGGGTGGCTTTGTCTTCTTCTTTTGCCAATTCCACTATGATTGTTTGGG | Ins_k2_53_TG | 0.4336642464996612 | 0.032605025539541 | 75.7564266685878 |
| GATCCAGATCTAACTTGGGGTGGCTTTGTCTTCTTCTTTTGCCAATTCCACTATGCAATTGTTTGGG | Ins_k4_52_ATGC;Ins_k4_53_TGC | -0.06997859635445358 | 0.03784351927763679 | 45.78151643824593 |
| GATCCAGATCTAACTTGGGGTGGCTTTGTCTTCTTCTTTTGCCAATTCCACTATGCATTGTTTGGG | Ins_k3_53_TGC | 0.02603687921087583 | 0.03577269744964823 | 50.39519907094944 |
| GATCCAGATCTAACTTGGGGTGGCTTTGTCTTCTTCTTTTGCCAATTCCACTATGGATTGTTTGGG | Ins_k3_53_TGG | 0.6766963979520946 | 0.03154936715004255 | 96.59774842843423 |
| GATCCAGATCTAACTTGGGGTGGCTTTGTCTTCTTCTTTTGCCAATTCCACTATGGG | Del_k6_54-59 | -0.08095065888690145 | 0.030298957607259216 | 45.281944456353436 |
| GATCCAGATCTAACTTGGGGTGGCTTTGTCTTCTTCTTTTGCCAATTCCACTATGGTATTGTTTGGG | Ins_k4_53_TGGT | -0.45081804511107626 | 0.046203635582778256 | 31.281941735369408 |
| GATCCAGATCTAACTTGGGGTGGCTTTGTCTTCTTCTTTTGCCAATTCCACTATGTATTGTTTGGG | Ins_k3_53_TGT | -0.37989567096293003 | 0.04283466548150088 | 33.581098493245044 |
| GATCCAGATCTAACTTGGGGTGGCTTTGTCTTCTTCTTTTGCCAATTCCACTATGTTTGGG | Del_k2_54-55 | 0.20454769410103602 | 0.032648603340911006 | 60.244225707696 |
| GATCCAGATCTAACTTGGGGTGGCTTTGTCTTCTTCTTTTGCCAATTCCACTATTAATTGTTTGGG | Ins_k3_51_TAT;Ins_k3_52_ATT;Ins | -0.32456086596648853 | 0.048352387493574914 | 35.49167524474867 |
| GATCCAGATCTAACTTGGGGTGGCTTTGTCTTCTTCTTTTGCCAATTCCACTATTATTGTTTGGG | Ins_k2_53_TT | -0.08524202565718397 | 0.04767089668717084 | 45.08803938109473 |
| GATCCAGATCTAACTTGGGGTGGCTTTGTCTTCTTCTTTTGCCAATTCCACTATTCATTGTTTGGG | Ins_k3_53_TTC | -0.04388553528655703 | 0.03916298153307261 | 46.99181790906206 |
| GATCCAGATCTAACTTGGGGTGGCTTTGTCTTCTTCTTTTGCCAATTCCACTATTGATTGTTTGGG | Ins_k3_53_TTG | 0.07423011727118067 | 0.035834837662983794 | 52.88338212936599 |
| GATCCAGATCTAACTTGGGGTGGCTTTGTCTTCTTCTTTTGCCAATTCCACTATTGGG | Del_k5_54-58 | 0.05868897073395074 | 0.029709423115450998 | 52.06786717106769 |
| GATCCAGATCTAACTTGGGGTGGCTTTGTCTTCTTCTTTTGCCAATTCCACTATTGTTTGGG | Del_k1_53-53;Del_k1_54-54 | -0.025829529039604116 | 0.03348023571999077 | 47.84800890061951 |
| GATCCAGATCTAACTTGGGGTGGCTTTGTCTTCTTCTTTTGCCAATTCCACTATTTATTGTTTGGG | Ins_k3_53_TTT | -0.5846371335660591 | 0.04883264755937605 | 27.363825427421972 |
| GATCCAGATCTAACTTGGGGTGGCTTTGTCTTCTTCTTTTGCCAATTCCACTATTTGGG | Del_k4_54-57 | 0.23203152150798378 | 0.030573685162223817 | 61.922930558857814 |
| GATCCAGATCTAACTTGGGGTGGCTTTGTCTTCTTCTTTTGCCAATTCCACTATTTGTTTGGG | A54T | 0.21161137653940276 | 0.03419578786682592 | 60.67127829309504 |
| GATCCAGATCTAACTTGGGGTGGCTTTGTCTTCTTCTTTTGCCAATTCCACTCAAAATTGTTTGGG | Ins_k3_52_CAA | 0.02815784902094376 | 0.04037835259881382 | 50.50219919865978 |
| GATCCAGATCTAACTTGGGGTGGCTTTGTCTTCTTCTTTTGCCAATTCCACTCAAATTGTTTGGG | Ins_k2_52_CA | 0.16152150117137865 | 0.03559518414159112 | 57.707118452880934 |
| GATCCAGATCTAACTTGGGGTGGCTTTGTCTTCTTCTTTTGCCAATTCCACTCAATTGTTTGGG | Ins_k1_52_C | 0.2273786591798016 | 0.034802491887405904 | 61.63548093835105 |
| GATCCAGATCTAACTTGGGGTGGCTTTGTCTTCTTCTTTTGCCAATTCCACTCACAATTGTTTGGG | Ins_k3_52_CAC | 0.1278314139684147 | 0.03854729473539729 | 55.79534532364532 |
| GATCCAGATCTAACTTGGGGTGGCTTTGTCTTCTTCTTTTGCCAATTCCACTCAGAATTGTTTGGG | Ins_k3_52_CAG | 0.39538896995800954 | 0.033821872641130984 | 72.91161868851415 |
| GATCCAGATCTAACTTGGGGTGGCTTTGTCTTCTTCTTTTGCCAATTCCACTCATAATTGTTTGGG | Ins_k3_51_TCA;Ins_k3_52_CAT | -0.19137508571230982 | 0.04193056422594847 | 40.547898288227316 |
| GATCCAGATCTAACTTGGGGTGGCTTTGTCTTCTTCTTTTGCCAATTCCACTCATTGTTTGGG | A53C | 0.04904074918466894 | 0.035626268026096315 | 51.56792052923844 |
| GATCCAGATCTAACTTGGGGTGGCTTTGTCTTCTTCTTTTGCCAATTCCACTCCAAATTGTTTGGG | Ins_k3_52_CCA | -1.2746709932049853 | 0.06496710461762112 | 13.724575275465316 |
| GATCCAGATCTAACTTGGGGTGGCTTTGTCTTCTTCTTTTGCCAATTCCACTCCAATTGTTTGGG | Ins_k2_52_CC | -0.9288973151507403 | 0.05086811291250302 | 19.393960819648147 |
| GATCCAGATCTAACTTGGGGTGGCTTTGTCTTCTTCTTTTGCCAATTCCACTCCCAATTGTTTGGG | Ins_k3_52_CCC | -1.3750572528457499 | 0.07132702934098648 | 12.413713413248049 |
| GATCCAGATCTAACTTGGGGTGGCTTTGTCTTCTTCTTTTGCCAATTCCACTCCGAATTGTTTGGG | Ins_k3_52_CCG | -0.06337005998975165 | 0.04096667340559051 | 46.085067162719376 |
| GATCCAGATCTAACTTGGGGTGGCTTTGTCTTCTTCTTTTGCCAATTCCACTCCTAATTGTTTGGG | Ins_k3_50_CTC;Ins_k3_51_TCC; | -0.9297906028863188 | 0.04586620440011394 | 19.376644167831515 |
| GATCCAGATCTAACTTGGGGTGGCTTTGTCTTCTTCTTTTGCCAATTCCACTCGAAATTGTTTGGG | Ins_k3_52_CGA | 0.5117895448136971 | 0.03369347742702624 | 81.91225223542212 |
| GATCCAGATCTAACTTGGGGTGGCTTTGTCTTCTTCTTTTGCCAATTCCACTCGAATTGTTTGGG | Ins_k2_52_CG | 0.6972682163265136 | 0.03375607161618011 | 98.60552072106394 |
| GATCCAGATCTAACTTGGGGTGGCTTTGTCTTCTTCTTTTGCCAATTCCACTCGCAATTGTTTGGG | Ins_k3_52_CGC | 0.5103800765659952 | 0.03766392381867838 | 81.7968808420561 |
| GATCCAGATCTAACTTGGGGTGGCTTTGTCTTCTTCTTTTGCCAATTCCACTCGGAATTGTTTGGG | Ins_k3_52_CGG | 0.4115586887370275 | 0.03064466294477464 | 74.10016237072557 |
| GATCCAGATCTAACTTGGGGTGGCTTTGTCTTCTTCTTTTGCCAATTCCACTCGTAATTGTTTGGG | Ins_k3_51_TCG;Ins_k3_52_CGT | 0.20916425656163296 | 0.03541638402912265 | 60.52298990964079 |
| GATCCAGATCTAACTTGGGGTGGCTTTGTCTTCTTCTTTTGCCAATTCCACTCTAAATTGTTTGGG | Ins_k3_52_CTA | -0.8644605888280931 | 0.06288207219449911 | 20.684785829839317 |
| GATCCAGATCTAACTTGGGGTGGCTTTGTCTTCTTCTTTTGCCAATTCCACTCTAATTGTTTGGG | Ins_k2_50_CT;Ins_k2_51_TC;Ins_ | -0.24464421688067478 | 0.31219339661234113 | 38.44446828763074 |
| GATCCAGATCTAACTTGGGGTGGCTTTGTCTTCTTCTTTTGCCAATTCCACTCTCAATTGTTTGGG | Ins_k3_52_CTC | -0.006565940098017475 | 0.040008468992061466 | 48.77866841757315 |
| GATCCAGATCTAACTTGGGGTGGCTTTGTCTTCTTCTTTTGCCAATTCCACTCTCCAATTGTTTGGG | Ins_k4_52_CTCC | 0.7497029966761941 | 0.03272837889772101 | 100 |

| | | | | |
|---|---|---|---|---|
| GATCCAGATCTAACTTGGGGTGGCTTTGTCTTCTTCTTTTGCCAATTCCACTCTGAATTGTTTGGG | Ins_k3_52_CTG | 0.3087716311035318 | 0.036523310353844714 | 66.86199144887736 |
| GATCCAGATCTAACTTGGGGTGGCTTTGTCTTCTTCTTTTGCCAATTCCACTCTTAATTGTTTGGG | Ins_k3_51_TCT;Ins_k3_52_CTT | -1.0892573829559085 | 0.06289774759424341 | 16.520493686955227 |
| GATCCAGATCTAACTTGGGGTGGCTTTGTCTTCTTCTTTTGCCAATTCCACTG | Del_k10_53-62 | -0.3008355296788714 | 0.029905885291632076 | 36.343795626069266 |
| GATCCAGATCTAACTTGGGGTGGCTTTGTCTTCTTCTTTTGCCAATTCCACTGAAAATTGTTTGGG | Ins_k3_52_GAA | 0.7666107291126221 | 0.034629492150829284 | 100 |
| GATCCAGATCTAACTTGGGGTGGCTTTGTCTTCTTCTTTTGCCAATTCCACTGAAATTGTTTGGG | Ins_k2_52_GA | 0.5940085592850701 | 0.03475579041677654 | 88.93160526092647 |
| GATCCAGATCTAACTTGGGGTGGCTTTGTCTTCTTCTTTTGCCAATTCCACTGAATTGTTTGGG | Ins_k1_52_G | 0.4661617269885421 | 0.03459657909730291 | 78.25875919806478 |
| GATCCAGATCTAACTTGGGGTGGCTTTGTCTTCTTCTTTTGCCAATTCCACTGACAATTGTTTGGG | Ins_k3_52_GAC | 0.3885873874209117 | 0.03320842724270537 | 72.41738698029334 |
| GATCCAGATCTAACTTGGGGTGGCTTTGTCTTCTTCTTTTGCCAATTCCACTGACTAATTGTTTGGG | Ins_k4_49_ACTG;Ins_k4_50_CT | 0.44246588429247113 | 0.034029944005008084 | 76.42615030548573 |
| GATCCAGATCTAACTTGGGGTGGCTTTGTCTTCTTCTTTTGCCAATTCCACTGAGAATTGTTTGGG | Ins_k3_52_GAG | 0.373772229821698 | 0.032246462849873064 | 71.35242028294563 |
| GATCCAGATCTAACTTGGGGTGGCTTTGTCTTCTTCTTTTGCCAATTCCACTGATAATTGTTTGGG | Ins_k3_51_TGA;Ins_k3_52_GAT | 0.35155944158300145 | 0.03425874244579536 | 69.78495739088791 |
| GATCCAGATCTAACTTGGGGTGGCTTTGTCTTCTTCTTTTGCCAATTCCACTGATTGTTTGGG | A53G | 0.3233562485701602 | 0.027529828578228267 | 67.84429385749497 |
| GATCCAGATCTAACTTGGGGTGGCTTTGTCTTCTTCTTTTGCCAATTCCACTGCAAAATTGTTTGGG | Ins_k4_52_GCAA | 0.8217958352827148 | 0.033087484024988636 | 100 |
| GATCCAGATCTAACTTGGGGTGGCTTTGTCTTCTTCTTTTGCCAATTCCACTGCAAATTGTTTGGG | Ins_k3_52_GCA | 0.5940391949679447 | 0.033945666603486316 | 88.93432978311637 |
| GATCCAGATCTAACTTGGGGTGGCTTTGTCTTCTTCTTTTGCCAATTCCACTGCAATTGTTTGGG | Ins_k2_52_GC | 0.6350517367405564 | 0.03318248254305265 | 92.65758082283796 |
| GATCCAGATCTAACTTGGGGTGGCTTTGTCTTCTTCTTTTGCCAATTCCACTGCCAATTGTTTGGG | Ins_k3_52_GCC | 0.3865348909665671 | 0.036250335172774636 | 72.26890298384856 |
| GATCCAGATCTAACTTGGGGTGGCTTTGTCTTCTTCTTTTGCCAATTCCACTGCGAATTGTTTGGG | Ins_k3_52_GCG | 0.4288043127075256 | 0.03099985036913948 | 75.38914864697722 |
| GATCCAGATCTAACTTGGGGTGGCTTTGTCTTCTTCTTTTGCCAATTCCACTGCTAATTGTTTGGG | Ins_k3_50_CTG;Ins_k3_51_TGC; | 0.3227519379155982 | 0.030634389618381986 | 67.80330721344582 |
| GATCCAGATCTAACTTGGGGTGGCTTTGTCTTCTTCTTTTGCCAATTCCACTGG | Del_k9_53-61 | -0.3744141536390018 | 0.030667394280151413 | 33.76567929565012 |
| GATCCAGATCTAACTTGGGGTGGCTTTGTCTTCTTCTTTTGCCAATTCCACTGGAAATTGTTTGGG | Ins_k3_52_GGA | 0.751751655035316 | 0.03174933799353776 | 100 |
| GATCCAGATCTAACTTGGGGTGGCTTTGTCTTCTTCTTTTGCCAATTCCACTGGAATTGTTTGGG | Ins_k2_52_GG | 0.9386857161471587 | 0.032592551003866176 | 100 |
| GATCCAGATCTAACTTGGGGTGGCTTTGTCTTCTTCTTTTGCCAATTCCACTGGCAATTGTTTGGG | Ins_k3_52_GGC | 0.2066106560569938 | 0.03628183853156072 | 60.36863553564606 |
| GATCCAGATCTAACTTGGGGTGGCTTTGTCTTCTTCTTTTGCCAATTCCACTGGCGAATTGTTTGGG | Ins_k4_52_GGCG | 0.9656602764833004 | 0.03297392843524956 | 100 |
| GATCCAGATCTAACTTGGGGTGGCTTTGTCTTCTTCTTTTGCCAATTCCACTGGG | Del_k8_52-59;Del_k8_53-60 | -0.16033632370967854 | 0.031876287825420936 | 41.826190532857744 |
| GATCCAGATCTAACTTGGGGTGGCTTTGTCTTCTTCTTTTGCCAATTCCACTGGGAATTGTTTGGG | Ins_k3_52_GGG | 0.26811247292093576 | 0.031079716483892586 | 64.19796470129948 |
| GATCCAGATCTAACTTGGGGTGGCTTTGTCTTCTTCTTTTGCCAATTCCACTGGTAATTGTTTGGG | Ins_k3_51_TGG;Ins_k3_52_GGT | 0.12868959388332435 | 0.034125089114325206 | 55.843248320102305 |
| GATCCAGATCTAACTTGGGGTGGCTTTGTCTTCTTCTTTTGCCAATTCCACTGTAAATTGTTTGGG | Ins_k3_52_GTA | -0.22310078398743743 | 0.040703318063824746 | 39.28167993651856 |
| GATCCAGATCTAACTTGGGGTGGCTTTGTCTTCTTCTTTTGCCAATTCCACTGTAATTGTTTGGG | Ins_k2_51_TG;Ins_k2_52_GT | 0.009358385415476511 | 0.03621651768458274 | 49.56165352041371 |
| GATCCAGATCTAACTTGGGGTGGCTTTGTCTTCTTCTTTTGCCAATTCCACTGTCAATTGTTTGGG | Ins_k3_52_GTC | 0.39594192275531503 | 0.03385792225565723 | 72.9519465206903 |
| GATCCAGATCTAACTTGGGGTGGCTTTGTCTTCTTCTTTTGCCAATTCCACTGTGAATTGTTTGGG | Ins_k3_52_GTG | 0.33631224192344944 | 0.03415325061206663 | 68.7290028442425 |
| GATCCAGATCTAACTTGGGGTGGCTTTGTCTTCTTCTTTTGCCAATTCCACTGTTAATTGTTTGGG | Ins_k3_51_TGT;Ins_k3_52_GTT | -0.09728172258014522 | 0.04129576109232457 | 44.54844782946172 |
| GATCCAGATCTAACTTGGGGTGGCTTTGTCTTCTTCTTTTGCCAATTCCACTGTTTGGG | Del_k4_52-55;Del_k4_53-56 | 0.15200707492352727 | 0.029156140074215003 | 57.16067201476941 |
| GATCCAGATCTAACTTGGGGTGGCTTTGTCTTCTTCTTTTGCCAATTCCACTTAAAATTGTTTGGG | Ins_k3_52_TAA | -0.4072850618002898 | 0.05017172489379673 | 32.673814359793965 |
| GATCCAGATCTAACTTGGGGTGGCTTTGTCTTCTTCTTTTGCCAATTCCACTTAAATTGTTTGGG | Ins_k2_52_TA | -0.14920339108418962 | 0.04409215254634405 | 42.294440354583834 |
| GATCCAGATCTAACTTGGGGTGGCTTTGTCTTCTTCTTTTGCCAATTCCACTTAATTGTTTGGG | Ins_k1_51_T;Ins_k1_52_T | -0.11713889123130633 | 0.039888742274118885 | 43.67256682648435 |
| GATCCAGATCTAACTTGGGGTGGCTTTGTCTTCTTCTTTTGCCAATTCCACTTACAATTGTTTGGG | Ins_k3_52_TAC | -0.06787480419823233 | 0.04109288953083444 | 45.8779326176686 |
| GATCCAGATCTAACTTGGGGTGGCTTTGTCTTCTTCTTTTGCCAATTCCACTTAGAATTGTTTGGG | Ins_k3_52_TAG | -0.7278246073407078 | 0.05848077813709068 | 23.713260998471114 |
| GATCCAGATCTAACTTGGGGTGGCTTTGTCTTCTTCTTTTGCCAATTCCACTTATAATTGTTTGGG | Ins_k3_51_TTA;Ins_k3_52_TAT | -0.7160581397961389 | 0.05729417938258111 | 23.993930319404104 |
| GATCCAGATCTAACTTGGGGTGGCTTTGTCTTCTTCTTTTGCCAATTCCACTTATTGTTTGGG | A53T | -0.230143816267869 | 0.034873641990893364 | 39.00598978363638 |
| GATCCAGATCTAACTTGGGGTGGCTTTGTCTTCTTCTTTTGCCAATTCCACTTCAAATTGTTTGGG | Ins_k3_52_TCA | 0.412880116061489 | 0.0354213603315242 | 74.19814507428231 |
| GATCCAGATCTAACTTGGGGTGGCTTTGTCTTCTTCTTTTGCCAATTCCACTTCAATTGTTTGGG | Ins_k2_52_TC | 0.6558577015224951 | 0.03389276605505767 | 94.60560618719244 |
| GATCCAGATCTAACTTGGGGTGGCTTTGTCTTCTTCTTTTGCCAATTCCACTTCCAAATTGTTTGGG | Ins_k4_52_TCCA | -0.011874213256981236 | 0.03853900126003275 | 48.52042394378219 |
| GATCCAGATCTAACTTGGGGTGGCTTTGTCTTCTTCTTTTGCCAATTCCACTTCCAATTGTTTGGG | Ins_k3_52_TCC | -0.021679898436880896 | 0.03492565394820547 | 48.046972990867765 |
| GATCCAGATCTAACTTGGGGTGGCTTTGTCTTCTTCTTTTGCCAATTCCACTTCGAATTGTTTGGG | Ins_k3_52_TCG | 0.6232984162073598 | 0.03235200739258967 | 91.57492146008994 |
| GATCCAGATCTAACTTGGGGTGGCTTTGTCTTCTTCTTTTGCCAATTCCACTTCTAATTGTTTGGG | Ins_k3_50_CTT;Ins_k3_51_TTC;I | -0.0022487036751850376 | 0.037603021272490045 | 48.98971269775322 |
| GATCCAGATCTAACTTGGGGTGGCTTTGTCTTCTTCTTTTGCCAATTCCACTTGAAATTGTTTGGG | Ins_k3_52_TGA | 0.5164271087913803 | 0.03561827661384748 | 82.29300775234599 |
| GATCCAGATCTAACTTGGGGTGGCTTTGTCTTCTTCTTTTGCCAATTCCACTTGAATTGTTTGGG | Ins_k2_52_TG | 0.4972941982925059 | 0.032188384677383546 | 80.73346982462081 |
| GATCCAGATCTAACTTGGGGTGGCTTTGTCTTCTTCTTTTGCCAATTCCACTTGCAATTGTTTGGG | Ins_k3_52_TGC | 0.1245347626717841 | 0.035515160530388445 | 55.61171038263201 |
| GATCCAGATCTAACTTGGGGTGGCTTTGTCTTCTTCTTTTGCCAATTCCACTTGGAATTGTTTGGG | Ins_k3_52_TGG | 0.14417734535285598 | 0.03821199673003774 | 56.71486695483801 |
| GATCCAGATCTAACTTGGGGTGGCTTTGTCTTCTTCTTTTGCCAATTCCACTTGGG | Del_k7_52-58;Del_k7_53-59 | 0.015457017671540063 | 0.028802277632973054 | 49.86483537686747 |
| GATCCAGATCTAACTTGGGGTGGCTTTGTCTTCTTCTTTTGCCAATTCCACTTGTAATTGTTTGGG | Ins_k3_51_TTG;Ins_k3_52_TGT | -0.41493598171892354 | 0.046952529565481696 | 32.424783495075815 |
| GATCCAGATCTAACTTGGGGTGGCTTTGTCTTCTTCTTTTGCCAATTCCACTTGTTTGGG | Del_k3_52-54;Del_k3_53-55 | 0.0353645056095091 | 0.03230422734665771 | 50.8674657998615 |
| GATCCAGATCTAACTTGGGGTGGCTTTGTCTTCTTCTTTTGCCAATTCCACTTTAAATTGTTTGGG | Ins_k3_52_TTA | -0.502913467594961 | 0.05066141143636653 | 29.694016688261627 |
| GATCCAGATCTAACTTGGGGTGGCTTTGTCTTCTTCTTTTGCCAATTCCACTTTAATTGTTTGGG | Ins_k2_51_TT;Ins_k2_52_TT | -0.3507138222427538 | 0.05399564048522336 | 34.57549563641601 |
| GATCCAGATCTAACTTGGGGTGGCTTTGTCTTCTTCTTTTGCCAATTCCACTTTCAATTGTTTGGG | Ins_k3_52_TTC | -0.3488643713241173 | 0.044060493846305226 | 34.639500487254296 |
| GATCCAGATCTAACTTGGGGTGGCTTTGTCTTCTTCTTTTGCCAATTCCACTTTGAATTGTTTGGG | Ins_k3_52_TTG | 0.16005977305660724 | 0.03114048416545562 | 57.622827955319224 |
| GATCCAGATCTAACTTGGGGTGGCTTTGTCTTCTTCTTTTGCCAATTCCACTTTGGG | Del_k6_52-57;Del_k6_53-58 | -0.25895240909371553 | 0.03148093613182803 | 37.898314003638774 |
| GATCCAGATCTAACTTGGGGTGGCTTTGTCTTCTTCTTTTGCCAATTCCACTTTGTTTGGG | Del_k2_53-54 | 0.019271698855876318 | 0.03421945629887656 | 50.055417099286174 |
| GATCCAGATCTAACTTGGGGTGGCTTTGTCTTCTTCTTTTGCCAATTCCACTTTTAATTGTTTGGG | Ins_k3_51_TTT;Ins_k3_52_TTT | -1.0577358957051146 | 0.062220343946029495 | 17.049538554231937 |
| GATCCAGATCTAACTTGGGGTGGCTTTGTCTTCTTCTTTTGCCAATTCCACTTTTGAATTGTTTGGG | Ins_k4_52_TTTG | -0.07851035366271508 | 0.044433964096425756 | 45.39258116073115 |
| GATCCAGATCTAACTTGGGGTGGCTTTGTCTTCTTCTTTTGCCAATTCCACTTTTGGG | Del_k5_53-57 | -0.06597082846890234 | 0.03089876233211564 | 45.96536629727327 |
| GATCCAGATCTAACTTGGGGTGGCTTTGTCTTCTTCTTTTGCCAATTCCAG | Del_k12_51-62 | -0.07058564914381715 | 0.026874623524264132 | 45.75373307482814 |
| GATCCAGATCTAACTTGGGGTGGCTTTGTCTTCTTCTTTTGCCAATTCCAGAACTAATTGTTTGGG | Ins_k3_49_AGA;Ins_k3_50_GAA | 0.25546525174084433 | 0.03327359370071822 | 63.39115156874584 |
| GATCCAGATCTAACTTGGGGTGGCTTTGTCTTCTTCTTTTGCCAATTCCAGACCTAATTGTTTGGG | Ins_k3_50_GAC | 0.43213859962054 | 0.03170831221218174 | 75.640937233117 |
| GATCCAGATCTAACTTGGGGTGGCTTTGTCTTCTTCTTTTGCCAATTCCAGACTAATTGTTTGGG | Ins_k2_49_AG;Ins_k2_50_GA | 0.3414783019156249 | 0.03504913690477197 | 69.08497970334584 |
| GATCCAGATCTAACTTGGGGTGGCTTTGTCTTCTTCTTTTGCCAATTCCAGAGCTAATTGTTTGGG | Ins_k3_50_GAG | 0.27407391907599277 | 0.03286495337509788 | 64.58182044198023 |
| GATCCAGATCTAACTTGGGGTGGCTTTGTCTTCTTCTTTTGCCAATTCCAGATCTAATTGTTTGGG | Ins_k3_50_GAT | 0.253505844381656 | 0.04102305965073289 | 63.2670640885161 |
| GATCCAGATCTAACTTGGGGTGGCTTTGTCTTCTTCTTTTGCCAATTCCAGCACTAATTGTTTGGG | Ins_k3_48_CAG;Ins_k3_49_AGC; | 0.4348497459554761 | 0.03155789936603793 | 75.846289126601 |
| GATCCAGATCTAACTTGGGGTGGCTTTGTCTTCTTCTTTTGCCAATTCCAGCCCTAATTGTTTGGG | Ins_k3_50_GCC | 0.29486086683679624 | 0.0359694934978147 | 65.93832936490737 |
| GATCCAGATCTAACTTGGGGTGGCTTTGTCTTCTTCTTTTGCCAATTCCAGCCTAATTGTTTGGG | Ins_k2_50_GC | 0.6013889965853145 | 0.034384386539603753 | 89.59038745754677 |
| GATCCAGATCTAACTTGGGGTGGCTTTGTCTTCTTCTTTTGCCAATTCCAGCGCTAATTGTTTGGG | Ins_k3_50_GCG | 0.7824634151380674 | 0.031197483762590194 | 100 |
| GATCCAGATCTAACTTGGGGTGGCTTTGTCTTCTTCTTTTGCCAATTCCAGCTAATTGTTTGGG | Ins_k1_50_G | 0.5058974901766919 | 0.03247019017164401 | 81.4310398268715 |
| GATCCAGATCTAACTTGGGGTGGCTTTGTCTTCTTCTTTTGCCAATTCCAGCTCTAATTGTTTGGG | Ins_k3_50_GCT | 0.24733668704442568 | 0.035942851490289314 | 62.8779610688196 |
| GATCCAGATCTAACTTGGGGTGGCTTTGTCTTCTTCTTTTGCCAATTCCAGG | Del_k11_51-61 | -0.38273964162801855 | 0.028424606823715188 | 33.48573051038099 |
| GATCCAGATCTAACTTGGGGTGGCTTTGTCTTCTTCTTTTGCCAATTCCAGGACTAATTGTTTGGG | Ins_k3_49_AGG;Ins_k3_50_GGA | 0.3935856893234888 | 0.03140898616141913 | 72.7802570550435 |
| GATCCAGATCTAACTTGGGGTGGCTTTGTCTTCTTCTTTTGCCAATTCCAGGCCTAATTGTTTGGG | Ins_k3_50_GGC | 0.12005416965113103 | 0.039962333509987034 | 55.36309433082687 |

| | | | | |
|---|---|---|---|---|
| **GATCCAGATCTAACTTGGGGTGGCTTTGTCTTCTTCTTTTGCCAATTCCAGGCTAATTGTTTGGG** | Ins_k2_50_GG | -0.1605892446599797 | 0.040623807498708704 | 41.81561315067786 |
| **GATCCAGATCTAACTTGGGGTGGCTTTGTCTTCTTCTTTTGCCAATTCCAGGG** | Del_k10_51-60 | -0.130384516748124 | 0.030990621809990602 | 43.09791060327456 |
| **GATCCAGATCTAACTTGGGGTGGCTTTGTCTTCTTCTTTTGCCAATTCCAGGGCTAATTGTTTGGG** | Ins_k3_50_GGG | -0.824292517847462 | 0.051724286834666275 | 21.532566649043638 |
| **GATCCAGATCTAACTTGGGGTGGCTTTGTCTTCTTCTTTTGCCAATTCCAGGTCTAATTGTTTGGG** | Ins_k3_50_GGT | -1.5937438106333628 | 0.07798138331551298 | 9.975331787894904 |
| **GATCCAGATCTAACTTGGGGTGGCTTTGTCTTCTTCTTTTGCCAATTCCAGTAATTGTTTGGG** | C51G | 0.037652880352254336 | 0.03326817843156377 | 50.984002913263375 |
| **GATCCAGATCTAACTTGGGGTGGCTTTGTCTTCTTCTTTTGCCAATTCCAGTACTAATTGTTTGGG** | Ins_k3_49_AGT;Ins_k3_50_GTA | 0.09020923329411679 | 0.035617900007109723 | 53.73519934655203 |
| **GATCCAGATCTAACTTGGGGTGGCTTTGTCTTCTTCTTTTGCCAATTCCAGTCCTAATTGTTTGGG** | Ins_k3_50_GTC | 0.34461095021042887 | 0.03399240349312677 | 69.3017379836759 |
| **GATCCAGATCTAACTTGGGGTGGCTTTGTCTTCTTCTTTTGCCAATTCCAGTCTAATTGTTTGGG** | Ins_k2_50_GT | 0.46770545832837374 | 0.035154616193377806 | 78.37966299473463 |
| **GATCCAGATCTAACTTGGGGTGGCTTTGTCTTCTTCTTTTGCCAATTCCAGTGCTAATTGTTTGGG** | Ins_k3_50_GTG | 0.30317266894636563 | 0.03314578747614226 | 66.48867974328438 |
| **GATCCAGATCTAACTTGGGGTGGCTTTGTCTTCTTCTTTTGCCAATTCCAGTTCTAATTGTTTGGG** | Ins_k3_50_GTT | 0.005788046365931443 | 0.03564117715756658 | 49.385017127032874 |
| **GATCCAGATCTAACTTGGGGTGGCTTTGTCTTCTTCTTTTGCCAATTCCAGTTTGGG** | Del_k6_51-56 | 0.1022519185919738 | 0.03279481064457463 | 54.38622763811985 |
| **GATCCAGATCTAACTTGGGGTGGCTTTGTCTTCTTCTTTTGCCAATTCCATAACTAATTGTTTGGG** | Ins_k3_49_ATA;Ins_k3_50_TAA | -1.7621350607740676 | 0.08301822410945826 | 8.429386335303631 |
| **GATCCAGATCTAACTTGGGGTGGCTTTGTCTTCTTCTTTTGCCAATTCCATAATTGTTTGGG** | Del_k1_51-51 | -0.5278370976163306 | 0.0420929283942583 | 28.963080629385427 |
| **GATCCAGATCTAACTTGGGGTGGCTTTGTCTTCTTCTTTTGCCAATTCCATACCTAATTGTTTGGG** | Ins_k3_50_TAC | -0.8883984271210357 | 0.05374771797924803 | 20.195516163816954 |
| **GATCCAGATCTAACTTGGGGTGGCTTTGTCTTCTTCTTTTGCCAATTCCATACTAATTGTTTGGG** | Ins_k2_49_AT;Ins_k2_50_TA | -1.1805861369091675 | 0.06585769082815736 | 15.078545382658113 |
| **GATCCAGATCTAACTTGGGGTGGCTTTGTCTTCTTCTTTTGCCAATTCCATAGCTAATTGTTTGGG** | Ins_k3_50_TAG | -3.053343782074494 | 0.14530300220640902 | 2.317560798713841 |
| **GATCCAGATCTAACTTGGGGTGGCTTTGTCTTCTTCTTTTGCCAATTCCATATCTAATTGTTTGGG** | Ins_k3_50_TAT | -1.6495577673917956 | 0.07751044007559986 | 9.433821534432775 |
| **GATCCAGATCTAACTTGGGGTGGCTTTGTCTTCTTCTTTTGCCAATTCCATCACTAATTGTTTGGG** | Ins_k3_48_CAT;Ins_k3_49_ATC;I | 0.15613686195236154 | 0.035162232190514536 | 57.39722152997657 |
| **GATCCAGATCTAACTTGGGGTGGCTTTGTCTTCTTCTTTTGCCAATTCCATCCCTAATTGTTTGGG** | Ins_k3_50_TCC | -1.4355679513680333 | 0.0652506856576658 | 11.684826075229735 |
| **GATCCAGATCTAACTTGGGGTGGCTTTGTCTTCTTCTTTTGCCAATTCCATCCTAATTGTTTGGG** | Ins_k2_50_TC | 0.08668666521048596 | 0.03933583987681515 | 53.546246443484065 |
| **GATCCAGATCTAACTTGGGGTGGCTTTGTCTTCTTCTTTTGCCAATTCCATCGCTAATTGTTTGGG** | Ins_k3_50_TCG | 0.6487085011733336 | 0.031631351230947 | 93.93166369983008 |
| **GATCCAGATCTAACTTGGGGTGGCTTTGTCTTCTTCTTTTGCCAATTCCATCTAATTGTTTGGG** | Ins_k1_50_T | 0.23462710943362497 | 0.03673621344169209 | 62.0838657400906 |
| **GATCCAGATCTAACTTGGGGTGGCTTTGTCTTCTTCTTTTGCCAATTCCATCTACTAATTGTTTGGG** | Ins_k4_49_ATCT;Ins_k4_50_TCT | 0.5293640256118568 | 0.033729208296985044 | 83.3645417771928 |
| **GATCCAGATCTAACTTGGGGTGGCTTTGTCTTCTTCTTTTGCCAATTCCATCTCTAATTGTTTGGG** | Ins_k3_50_TCT | -0.010658032533798378 | 0.03965546995949518 | 48.57946944578612 |
| **GATCCAGATCTAACTTGGGGTGGCTTTGTCTTCTTCTTTTGCCAATTCCATGACTAATTGTTTGGG** | Ins_k3_49_ATG;Ins_k3_50_TGA | -0.23203963022747753 | 0.04193984141983842 | 38.932111735341 |
| **GATCCAGATCTAACTTGGGGTGGCTTTGTCTTCTTCTTTTGCCAATTCCATGAGCTAATTGTTTGGG** | Ins_k4_50_TGAG | 0.21999306300549726 | 0.03936468579348333 | 61.18194305156569 |
| **GATCCAGATCTAACTTGGGGTGGCTTTGTCTTCTTCTTTTGCCAATTCCATGCCTAATTGTTTGGG** | Ins_k3_50_TGC | -0.31581281174528714 | 0.047302556036112886 | 35.803520374493246 |
| **GATCCAGATCTAACTTGGGGTGGCTTTGTCTTCTTCTTTTGCCAATTCCATGCTAATTGTTTGGG** | Ins_k2_50_TG | -0.44133477121695874 | 0.046773388895558904 | 31.580008045196138 |
| **GATCCAGATCTAACTTGGGGTGGCTTTGTCTTCTTCTTTTGCCAATTCCATGGCCTAATTGTTTGGG** | Ins_k4_50_TGGC | 0.24997803167671728 | 0.03730657946036176 | 63.044262967405196 |
| **GATCCAGATCTAACTTGGGGTGGCTTTGTCTTCTTCTTTTGCCAATTCCATGGCTAATTGTTTGGG** | Ins_k3_50_TGG | 0.1530760914352226 | 0.038505270762739255 | 57.221810390116794 |
| **GATCCAGATCTAACTTGGGGTGGCTTTGTCTTCTTCTTTTGCCAATTCCATGGG** | Del_k9_51-59 | -0.08756076343423125 | 0.031207773334995435 | 44.98361315619278 |
| **GATCCAGATCTAACTTGGGGTGGCTTTGTCTTCTTCTTTTGCCAATTCCATGTCTAATTGTTTGGG** | Ins_k3_50_TGT | -1.034909418036422 | 0.05676172370918261 | 17.443195270734467 |
| **GATCCAGATCTAACTTGGGGTGGCTTTGTCTTCTTCTTTTGCCAATTCCATGTTTGGG** | Del_k5_51-55 | -0.013140776598824622 | 0.033249519392836054 | 48.45900865487119 |
| **GATCCAGATCTAACTTGGGGTGGCTTTGTCTTCTTCTTTTGCCAATTCCATTAATTGTTTGGG** | C51T | -0.7526897314703915 | 0.04431393785193501 | 23.13089808795115 |
| **GATCCAGATCTAACTTGGGGTGGCTTTGTCTTCTTCTTTTGCCAATTCCATTACTAATTGTTTGGG** | Ins_k3_49_ATT;Ins_k3_50_TTA | -1.8990357413377288 | 0.07806980621006106 | 7.350903960332795 |
| **GATCCAGATCTAACTTGGGGTGGCTTTGTCTTCTTCTTTTGCCAATTCCATTCCTAATTGTTTGGG** | Ins_k3_50_TTC | -1.3365567207182898 | 0.06862760326350725 | 12.900967571302592 |
| **GATCCAGATCTAACTTGGGGTGGCTTTGTCTTCTTCTTTTGCCAATTCCATTCTAATTGTTTGGG** | Ins_k2_50_TT | -1.5079771109485915 | 0.07721551887095186 | 10.868643755568986 |
| **GATCCAGATCTAACTTGGGGTGGCTTTGTCTTCTTCTTTTGCCAATTCCATTGCTAATTGTTTGGG** | Ins_k3_50_TTG | -1.0947619830530888 | 0.06392022131231631 | 16.429804807801055 |
| **GATCCAGATCTAACTTGGGGTGGCTTTGTCTTCTTCTTTTGCCAATTCCATTGGG** | Del_k8_51-58 | -0.19378097878251582 | 0.03208104700756379 | 40.45046163880362 |
| **GATCCAGATCTAACTTGGGGTGGCTTTGTCTTCTTCTTTTGCCAATTCCATTGTTTGGG** | Del_k4_50-53;Del_k4_51-54 | -0.1409883172907695 | 0.038185734279663194 | 42.6433233909932 |
| **GATCCAGATCTAACTTGGGGTGGCTTTGTCTTCTTCTTTTGCCAATTCCATTTCTAATTGTTTGGG** | Ins_k3_50_TTT | -3.1677646785302604 | 0.16160150088634542 | 2.0669918882203127 |
| **GATCCAGATCTAACTTGGGGTGGCTTTGTCTTCTTCTTTTGCCAATTCCATTTGGG** | Del_k7_51-57 | -0.09197001643556779 | 0.03120009027707569 | 44.78570565762388 |
| **GATCCAGATCTAACTTGGGGTGGCTTTGTCTTCTTCTTTTGCCAATTCCCAAACTAATTGTTTGGG** | Ins_k3_49_CAA | -5.834768413273409 | 1.5097727242427577 | 0.14357332790066274 |
| **GATCCAGATCTAACTTGGGGTGGCTTTGTCTTCTTCTTTTGCCAATTCCCAACTAATTGTTTGGG** | Ins_k2_49_CA | -0.6723755180171184 | 0.0528847517688277 | 25.065277372526367 |
| **GATCCAGATCTAACTTGGGGTGGCTTTGTCTTCTTCTTTTGCCAATTCCCACACTAATTGTTTGGG** | Ins_k3_48_CCA;Ins_k3_49_CAC | -0.9189417084829401 | 0.05578471490056365 | 19.58800377024893 |
| **GATCCAGATCTAACTTGGGGTGGCTTTGTCTTCTTCTTTTGCCAATTCCCACTAATTGTTTGGG** | Ins_k1_47_C;Ins_k1_48_C;Ins_k | -0.6906368913741845 | 0.05249607313167936 | 24.61170501582429 |
| **GATCCAGATCTAACTTGGGGTGGCTTTGTCTTCTTCTTTTGCCAATTCCCAGACTAATTGTTTGGG** | Ins_k3_49_CAG | -1.691756864244393 | 0.08979741519766501 | 9.04400557237512 |
| **GATCCAGATCTAACTTGGGGTGGCTTTGTCTTCTTCTTTTGCCAATTCCCATACTAATTGTTTGGG** | Ins_k3_49_CAT | -2.631359411556896 | 0.11513301356222187 | 3.534244735662821 |
| **GATCCAGATCTAACTTGGGGTGGCTTTGTCTTCTTCTTTTGCCAATTCCCCAACTAATTGTTTGGG** | Ins_k3_49_CCA | -3.79647109419768 | 0.21993449021708741 | 1.1022879129231147 |
| **GATCCAGATCTAACTTGGGGTGGCTTTGTCTTCTTCTTTTGCCAATTCCCCACTAATTGTTTGGG** | Ins_k2_47_CC;Ins_k2_48_CC;Ins | -2.645335717634854 | 0.12199270667498621 | 3.4851926315652797 |
| **GATCCAGATCTAACTTGGGGTGGCTTTGTCTTCTTCTTTTGCCAATTCCCCCACTAATTGTTTGGG** | Ins_k3_47_CCC;Ins_k3_48_CCC | -2.2210631257309164 | 0.09896725799144132 | 5.327040917561675 |
| **GATCCAGATCTAACTTGGGGTGGCTTTGTCTTCTTCTTTTGCCAATTCCCCGACTAATTGTTTGGG** | Ins_k3_49_CCG | -1.0458346948338482 | 0.054870207983718056 | 17.253660777828067 |
| **GATCCAGATCTAACTTGGGGTGGCTTTGTCTTCTTCTTTTGCCAATTCCCCTAATTGTTTGGG** | A50C | -2.5104817636549717 | 0.08004865410307409 | 3.988348642393865 |
| **GATCCAGATCTAACTTGGGGTGGCTTTGTCTTCTTCTTTTGCCAATTCCCCTACTAATTGTTTGGG** | Ins_k3_49_CCT | -2.2415508455493147 | 0.10146566601678955 | 5.219012403437343 |
| **GATCCAGATCTAACTTGGGGTGGCTTTGTCTTCTTCTTTTGCCAATTCCCGAACTAATTGTTTGGG** | Ins_k3_49_CGA | -1.0048040308959987 | 0.5335926650153523 | 17.976314025872405 |
| **GATCCAGATCTAACTTGGGGTGGCTTTGTCTTCTTCTTTTGCCAATTCCCGACACTAATTGTTTGGG** | Ins_k4_48_CCGA;Ins_k4_49_CG | 0.42595099515902146 | 0.03406787568446881 | 75.1743460619019 |
| **GATCCAGATCTAACTTGGGGTGGCTTTGTCTTCTTCTTTTGCCAATTCCCGACTAATTGTTTGGG** | Ins_k2_49_CG | 0.47328626988578293 | 0.03386485739157508 | 78.81830798286853 |
| **GATCCAGATCTAACTTGGGGTGGCTTTGTCTTCTTCTTTTGCCAATTCCCGCAACTAATTGTTTGGG** | Ins_k4_49_CGCA | 0.5347775747491601 | 0.034084111057296834 | 83.81706358979396 |
| **GATCCAGATCTAACTTGGGGTGGCTTTGTCTTCTTCTTTTGCCAATTCCCGCACTAATTGTTTGGG** | Ins_k3_48_CCG;Ins_k3_49_CGC | 0.3761647471660118 | 0.032921209423081674 | 71.52333656458643 |
| **GATCCAGATCTAACTTGGGGTGGCTTTGTCTTCTTCTTTTGCCAATTCCCGGACTAATTGTTTGGG** | Ins_k3_49_CGG | 0.6589660914423432 | 0.03261267236132758 | 94.90013481759198 |
| **GATCCAGATCTAACTTGGGGTGGCTTTGTCTTCTTCTTTTGCCAATTCCCGTACTAATTGTTTGGG** | Ins_k3_49_CGT | -0.28662325958708507 | 0.04024967758645203 | 36.8640114330014 |
| **GATCCAGATCTAACTTGGGGTGGCTTTGTCTTCTTCTTTTGCCAATTCCCTAACTAATTGTTTGGG** | Ins_k3_49_CTA | -3.137484284150239 | 0.17923863345831464 | 2.130538470130492 |
| **GATCCAGATCTAACTTGGGGTGGCTTTGTCTTCTTCTTTTGCCAATTCCCTAATTGTTTGGG** | Del_k1_50-50 | -1.5500917296288965 | 0.068252134990553 | 10.42041960982759 |
| **GATCCAGATCTAACTTGGGGTGGCTTTGTCTTCTTCTTTTGCCAATTCCCTACTAATTGTTTGGG** | Ins_k2_49_CT | -1.0373950321109426 | 0.06260294557835998 | 17.39989205889626 |
| **GATCCAGATCTAACTTGGGGTGGCTTTGTCTTCTTCTTTTGCCAATTCCCTCAACTAATTGTTTGGG** | Ins_k4_49_CTCA | 0.12092384383276655 | 0.03730392359055149 | 55.411263127125125 |
| **GATCCAGATCTAACTTGGGGTGGCTTTGTCTTCTTCTTTTGCCAATTCCCTCACTAATTGTTTGGG** | Ins_k3_48_CCT;Ins_k3_49_CTC | -0.7043683802701197 | 0.32364050542395545 | 24.276059392684232 |
| **GATCCAGATCTAACTTGGGGTGGCTTTGTCTTCTTCTTTTGCCAATTCCCTGACTAATTGTTTGGG** | Ins_k3_49_CTG | -0.45341880870945533 | 0.03939295255340191 | 31.20069050343902 |
| **GATCCAGATCTAACTTGGGGTGGCTTTGTCTTCTTCTTTTGCCAATTCCCTTACTAATTGTTTGGG** | Ins_k3_49_CTT | -2.986135850601253 | 0.13991165923921783 | 2.478672622929985 |
| **GATCCAGATCTAACTTGGGGTGGCTTTGTCTTCTTCTTTTGCCAATTCCG** | Del_k13_50-62 | -0.12101804832883323 | 0.028313384232168146 | 43.503482243771764 |
| **GATCCAGATCTAACTTGGGGTGGCTTTGTCTTCTTCTTTTGCCAATTCCGAAACTAATTGTTTGGG** | Ins_k3_49_GAA | 0.30164802421009385 | 0.03964642041039129 | 66.3873853662834 |
| **GATCCAGATCTAACTTGGGGTGGCTTTGTCTTCTTCTTTTGCCAATTCCGAACTAATTGTTTGGG** | Ins_k2_49_GA | 0.20189208663644131 | 0.038919801617423526 | 60.08445293299095 |
| **GATCCAGATCTAACTTGGGGTGGCTTTGTCTTCTTCTTTTGCCAATTCCGACACTAATTGTTTGGG** | Ins_k3_48_CGA;Ins_k3_49_GAC | 0.28022736449622065 | 0.034673381651235215 | 64.98044635277508 |
| **GATCCAGATCTAACTTGGGGTGGCTTTGTCTTCTTCTTTTGCCAATTCCGACTAATTGTTTGGG** | Ins_k1_49_G | 0.34734609836025854 | 0.03387383226784872 | 69.49154796498027 |
| **GATCCAGATCTAACTTGGGGTGGCTTTGTCTTCTTCTTTTGCCAATTCCGAGACTAATTGTTTGGG** | Ins_k3_49_GAG | 0.6127152887880467 | 0.03402857316053914 | 90.61088266843518 |
| **GATCCAGATCTAACTTGGGGTGGCTTTGTCTTCTTCTTTTGCCAATTCCGATACTAATTGTTTGGG** | Ins_k3_49_GAT | -0.215794630374687 | 0.04449898275638563 | 39.56972890798459 |

| | | | | |
|---|---|---|---|---|
| **GATCCAGATCTAACTTGGGGTGGCTTTGTCTTCTTCTTTTGCCAATTCCGCAACTAATTGTTTGGG** | Ins_k3_49_GCA | 0.4884222481536009 | 0.03146166130436612 | 80.02037445646559 |
| **GATCCAGATCTAACTTGGGGTGGCTTTGTCTTCTTCTTTTGCCAATTCCGCACTAATTGTTTGGG** | Ins_k2_48_CG;Ins_k2_49_GC | 0.27041108547557535 | 0.034873829344381706 | 64.34570067778095 |
| **GATCCAGATCTAACTTGGGGTGGCTTTGTCTTCTTCTTTTGCCAATTCCGCCACTAATTGTTTGGG** | Ins_k3_47_CCG;Ins_k3_48_CGC | 0.4541919556889995 | 0.031119941039969568 | 77.32760372492169 |
| **GATCCAGATCTAACTTGGGGTGGCTTTGTCTTCTTCTTTTGCCAATTCCGCGACTAATTGTTTGGG** | Ins_k3_49_GCG | 0.6724521721942556 | 0.03186343336740563 | 96.18863457630877 |
| **GATCCAGATCTAACTTGGGGTGGCTTTGTCTTCTTCTTTTGCCAATTCCGCTAATTGTTTGGG** | A50G | -0.24267435772270574 | 0.026447385350365487 | 38.520273113463176 |
| **GATCCAGATCTAACTTGGGGTGGCTTTGTCTTCTTCTTTTGCCAATTCCGCTACTAATTGTTTGGG** | Ins_k3_49_GCT | 0.2418368406144904 | 0.03492061333098257 | 62.53309117415367 |
| **GATCCAGATCTAACTTGGGGTGGCTTTGTCTTCTTCTTTTGCCAATTCCGG** | Del_k12_50-61 | -0.1834935180428846 | 0.028497854459888002 | 40.868742007305116 |
| **GATCCAGATCTAACTTGGGGTGGCTTTGTCTTCTTCTTTTGCCAATTCCGGAACTAATTGTTTGGG** | Ins_k3_49_GGA | 0.6780875079677895 | 0.029357367369229986 | 96.73222003447995 |
| **GATCCAGATCTAACTTGGGGTGGCTTTGTCTTCTTCTTTTGCCAATTCCGGACTAATTGTTTGGG** | Ins_k2_49_GG | 0.8764894602865594 | 0.03309371885145979 | 100 |
| **GATCCAGATCTAACTTGGGGTGGCTTTGTCTTCTTCTTTTGCCAATTCCGGCACTAATTGTTTGGG** | Ins_k3_48_CGG;Ins_k3_49_GGC | 0.18133661953669 | 0.035225556178026404 | 58.86199606660865 |
| **GATCCAGATCTAACTTGGGGTGGCTTTGTCTTCTTCTTTTGCCAATTCCGGG** | Del_k11_50-60 | -0.16801172106748358 | 0.0320377893430629 | 41.50638678100513 |
| **GATCCAGATCTAACTTGGGGTGGCTTTGTCTTCTTCTTTTGCCAATTCCGGGACTAATTGTTTGGG** | Ins_k3_49_GGG | 0.7724082140632631 | 0.033182708011883597 | 100 |
| **GATCCAGATCTAACTTGGGGTGGCTTTGTCTTCTTCTTTTGCCAATTCCGGTACTAATTGTTTGGG** | Ins_k3_49_GGT | 0.13853519398231107 | 0.03561158378811367 | 56.395774122046696 |
| **GATCCAGATCTAACTTGGGGTGGCTTTGTCTTCTTCTTTTGCCAATTCCGTAACTAATTGTTTGGG** | Ins_k3_49_GTA | -1.1299478913091774 | 0.05709320109400949 | 15.861759404314855 |
| **GATCCAGATCTAACTTGGGGTGGCTTTGTCTTCTTCTTTTGCCAATTCCGTACTAATTGTTTGGG** | Ins_k2_49_GT | -0.7596989605691777 | 0.05377078824870191 | 22.969335201091276 |
| **GATCCAGATCTAACTTGGGGTGGCTTTGTCTTCTTCTTTTGCCAATTCCGTCACTAATTGTTTGGG** | Ins_k3_48_CGT;Ins_k3_49_GTC | 0.34371210225674365 | 0.031786058755746575 | 69.2394742453114 |
| **GATCCAGATCTAACTTGGGGTGGCTTTGTCTTCTTCTTTTGCCAATTCCGTGACTAATTGTTTGGG** | Ins_k3_49_GTG | 0.35591033905938196 | 0.03095054945949065 | 70.08924607028982 |
| **GATCCAGATCTAACTTGGGGTGGCTTTGTCTTCTTCTTTTGCCAATTCCGTTACTAATTGTTTGGG** | Ins_k3_49_GTT | -1.157675815545729 | 0.05636965966520496 | 15.427987332209621 |
| **GATCCAGATCTAACTTGGGGTGGCTTTGTCTTCTTCTTTTGCCAATTCCGTTTACTAATTGTTTGGG** | Ins_k4_49_GTTT | -1.3563066393035077 | 0.07152068927061507 | 12.648674100582877 |
| **GATCCAGATCTAACTTGGGGTGGCTTTGTCTTCTTCTTTTGCCAATTCCGTTTGGG** | Del_k7_50-56 | 0.4291449922851365 | 0.03185506214819518 | 75.41483656572092 |
| **GATCCAGATCTAACTTGGGGTGGCTTTGTCTTCTTCTTTTGCCAATTCCTAAACTAATTGTTTGGG** | Ins_k3_49_TAA | -2.798081015036602 | 0.1349424431617029 | 2.991509231531729 |
| **GATCCAGATCTAACTTGGGGTGGCTTTGTCTTCTTCTTTTGCCAATTCCTAACTAATTGTTTGGG** | Ins_k2_49_TA | -2.107943862304835 | 0.08953600362940703 | 5.965036487693057 |
| **GATCCAGATCTAACTTGGGGTGGCTTTGTCTTCTTCTTTTGCCAATTCCTAATTGTTTGGG** | Del_k2_49-50;Del_k2_50-51 | -0.9563420680432326 | 0.05278564364178663 | 18.868935901203844 |
| **GATCCAGATCTAACTTGGGGTGGCTTTGTCTTCTTCTTTTGCCAATTCCTACACTAATTGTTTGGG** | Ins_k3_48_CTA;Ins_k3_49_TAC | -0.10631943281815404 | 0.038776900757280616 | 44.147645761033864 |
| **GATCCAGATCTAACTTGGGGTGGCTTTGTCTTCTTCTTTTGCCAATTCCTACTAATTGTTTGGG** | Ins_k1_49_T | -0.2128157283901727 | 0.04045981756941954 | 39.687778994477114 |
| **GATCCAGATCTAACTTGGGGTGGCTTTGTCTTCTTCTTTTGCCAATTCCTAGACTAATTGTTTGGG** | Ins_k3_49_TAG | -1.3449534197947277 | 0.07035097428568757 | 12.793095547062979 |
| **GATCCAGATCTAACTTGGGGTGGCTTTGTCTTCTTCTTTTGCCAATTCCTATAACTAATTGTTTGGG** | Ins_k4_49_TATA | -3.038532410232167 | 0.16652826525571932 | 2.352142522657097 |
| **GATCCAGATCTAACTTGGGGTGGCTTTGTCTTCTTCTTTTGCCAATTCCTATACTAATTGTTTGGG** | Ins_k3_49_TAT | -2.7241969204576626 | 0.13257136884893278 | 3.220904157329825 |
| **GATCCAGATCTAACTTGGGGTGGCTTTGTCTTCTTCTTTTGCCAATTCCTCAACTAATTGTTTGGG** | Ins_k3_49_TCA | 0.24774585715107555 | 0.03552321792980899 | 62.90369411509367 |
| **GATCCAGATCTAACTTGGGGTGGCTTTGTCTTCTTCTTTTGCCAATTCCTCACTAATTGTTTGGG** | Ins_k2_48_CT;Ins_k2_49_TC | -0.0643518924881612 | 0.0399301162773928 | 46.039841551708065 |
| **GATCCAGATCTAACTTGGGGTGGCTTTGTCTTCTTCTTTTGCCAATTCCTCCACTAATTGTTTGGG** | Ins_k3_46_TCC;Ins_k3_47_CCT;I | 0.36885771331895356 | 0.0334930633079328 | 71.00261789480803 |
| **GATCCAGATCTAACTTGGGGTGGCTTTGTCTTCTTCTTTTGCCAATTCCTCGACTAATTGTTTGGG** | Ins_k3_49_TCG | 0.7269327853937078 | 0.030980796077196532 | 100 |
| **GATCCAGATCTAACTTGGGGTGGCTTTGTCTTCTTCTTTTGCCAATTCCTCTAATTGTTTGGG** | A50T | -0.43791584753995394 | 0.0392123505921966 | 31.68816246291988 |
| **GATCCAGATCTAACTTGGGGTGGCTTTGTCTTCTTCTTTTGCCAATTCCTCTACTAATTGTTTGGG** | Ins_k3_49_TCT | 0.18716171658459735 | 0.036927906143054814 | 59.205873493458995 |
| **GATCCAGATCTAACTTGGGGTGGCTTTGTCTTCTTCTTTTGCCAATTCCTGAACTAATTGTTTGGG** | Ins_k3_49_TGA | 0.1802875623032808 | 0.0403346473432578 | 58.80027884197262 |
| **GATCCAGATCTAACTTGGGGTGGCTTTGTCTTCTTCTTTTGCCAATTCCTGACTAATTGTTTGGG** | Ins_k2_49_TG | 0.45029551705512105 | 0.032847815477480155 | 77.02688770352611 |
| **GATCCAGATCTAACTTGGGGTGGCTTTGTCTTCTTCTTTTGCCAATTCCTGCACTAATTGTTTGGG** | Ins_k3_48_CTG;Ins_k3_49_TGC | 0.47560329066245033 | 0.03118426939583828 | 79.0011433749504 |
| **GATCCAGATCTAACTTGGGGTGGCTTTGTCTTCTTCTTTTGCCAATTCCTGCGACTAATTGTTTGGG** | Ins_k4_49_TGCG | 0.8680667714903025 | 0.0297774385150029 | 100 |
| **GATCCAGATCTAACTTGGGGTGGCTTTGTCTTCTTCTTTTGCCAATTCCTGGACTAATTGTTTGGG** | Ins_k3_49_TGG | 0.8514843260629876 | 0.028233948422210362 | 100 |
| **GATCCAGATCTAACTTGGGGTGGCTTTGTCTTCTTCTTTTGCCAATTCCTGGG** | Del_k10_50-59 | -0.0026368026914147524 | 0.027901238730086105 | 48.970703527408595 |
| **GATCCAGATCTAACTTGGGGTGGCTTTGTCTTCTTCTTTTGCCAATTCCTGTACTAATTGTTTGGG** | Ins_k3_49_TGT | -0.15298559474533663 | 0.03703452220126958 | 42.1347762987505 |
| **GATCCAGATCTAACTTGGGGTGGCTTTGTCTTCTTCTTTTGCCAATTCCTGTTTGGG** | Del_k6_50-55 | 0.07814078778326061 | 0.029309750098160215 | 53.09059652196242 |
| **GATCCAGATCTAACTTGGGGTGGCTTTGTCTTCTTCTTTTGCCAATTCCTTAACTAATTGTTTGGG** | Ins_k3_49_TTA | -3.1831186129469793 | 0.1635882487488675 | 2.0354978279040443 |
| **GATCCAGATCTAACTTGGGGTGGCTTTGTCTTCTTCTTTTGCCAATTCCTTACTAATTGTTTGGG** | Ins_k2_49_TT | -2.467404727524677 | 0.10902756400822955 | 4.163909044755479 |
| **GATCCAGATCTAACTTGGGGTGGCTTTGTCTTCTTCTTTTGCCAATTCCTTCACTAATTGTTTGGG** | Ins_k3_48_CTT;Ins_k3_49_TTC | -0.08056927982229178 | 0.03857617576829919 | 45.299217335521796 |
| **GATCCAGATCTAACTTGGGGTGGCTTTGTCTTCTTCTTTTGCCAATTCCTTGACTAATTGTTTGGG** | Ins_k3_49_TTG | -0.8137590371876067 | 0.051065426927708296 | 21.76057829328697 |
| **GATCCAGATCTAACTTGGGGTGGCTTTGTCTTCTTCTTTTGCCAATTCCTTGGG** | Del_k9_50-58 | -0.3335597505362067 | 0.031509221742508954 | 35.17372251165323 |
| **GATCCAGATCTAACTTGGGGTGGCTTTGTCTTCTTCTTTTGCCAATTCCTTGTTTGGG** | Del_k5_50-54 | -0.3270980080305668 | 0.03297711803100668 | 35.40174195756593 |
| **GATCCAGATCTAACTTGGGGTGGCTTTGTCTTCTTCTTTTGCCAATTCCTTTACTAATTGTTTGGG** | Ins_k3_49_TTT | -3.65763403057965 | 0.19601937843718156 | 1.266459236784611 |
| **GATCCAGATCTAACTTGGGGTGGCTTTGTCTTCTTCTTTTGCCAATTCCTTTGGG** | Del_k8_50-57 | -0.3415107703577151 | 0.034829155383931824 | 34.89516442470772 |
| **GATCCAGATCTAACTTGGGGTGGCTTTGTCTTCTTCTTTTGCCAATTCG** | Del_k14_49-62 | -0.22728365471578088 | 0.02768927803133935 | 39.11771291285321 |
| **GATCCAGATCTAACTTGGGGTGGCTTTGTCTTCTTCTTTTGCCAATTCGAACACTAATTGTTTGGG** | Ins_k3_48_GAA | 0.20414073066520955 | 0.035004644334077434 | 60.21971349873799 |
| **GATCCAGATCTAACTTGGGGTGGCTTTGTCTTCTTCTTTTGCCAATTCGACACTAATTGTTTGGG** | Ins_k2_48_GA | 0.6208961858458675 | 0.031155856964566093 | 91.35520141814685 |
| **GATCCAGATCTAACTTGGGGTGGCTTTGTCTTCTTCTTTTGCCAATTCGACCACTAATTGTTTGGG** | Ins_k3_47_CGA;Ins_k3_48_GAC | 0.653591355485073 | 0.030109334839498864 | 94.39143992559028 |
| **GATCCAGATCTAACTTGGGGTGGCTTTGTCTTCTTCTTTTGCCAATTCGACTAATTGTTTGGG** | C49G | 0.290193258940763 | 0.030025430270706296 | 65.63127226665702 |
| **GATCCAGATCTAACTTGGGGTGGCTTTGTCTTCTTCTTTTGCCAATTCGAGCACTAATTGTTTGGG** | Ins_k3_48_GAG | 0.2359112926508099 | 0.034394623408486556 | 62.16364401252756 |
| **GATCCAGATCTAACTTGGGGTGGCTTTGTCTTCTTCTTTTGCCAATTCGATCACTAATTGTTTGGG** | Ins_k3_48_GAT | 0.3743277969021006 | 0.03759740534315062 | 71.39207235243423 |
| **GATCCAGATCTAACTTGGGGTGGCTTTGTCTTCTTCTTTTGCCAATTCGATCCACTAATTGTTTGGG** | Ins_k4_46_TCGA;Ins_k4_47_CG/ | 0.6187260141932511 | 0.03194814506096816 5 | 91.15715991945879 |
| **GATCCAGATCTAACTTGGGGTGGCTTTGTCTTCTTCTTTTGCCAATTCGCACACTAATTGTTTGGG** | Ins_k3_48_GCA | -0.051683933322504094 | 0.03815713302919062 | 46.626782205400694 |
| **GATCCAGATCTAACTTGGGGTGGCTTTGTCTTCTTCTTTTGCCAATTCGCACTAATTGTTTGGG** | Ins_k1_48_G | -0.04021348236263389 | 0.03635134048964849 | 47.16469155775715 |
| **GATCCAGATCTAACTTGGGGTGGCTTTGTCTTCTTCTTTTGCCAATTCGCATCACTAATTGTTTGGG** | Ins_k4_48_GCAT | 0.4836815800601073 | 0.03463448082624957 | 79.64192218752423 |
| **GATCCAGATCTAACTTGGGGTGGCTTTGTCTTCTTCTTTTGCCAATTCGCCACTAATTGTTTGGG** | Ins_k2_47_CG;Ins_k2_48_GC | 0.41699303681543576 | 0.033413643455268004 | 74.50394459858946 |
| **GATCCAGATCTAACTTGGGGTGGCTTTGTCTTCTTCTTTTGCCAATTCGCCCACTAATTGTTTGGG** | Ins_k3_47_CGC;Ins_k3_48_GCC | 0.4281944980378314 | 0.033348479824097915 | 75.34318925297713 |
| **GATCCAGATCTAACTTGGGGTGGCTTTGTCTTCTTCTTTTGCCAATTCGCCCCACTAATTGTTTGGG** | Ins_k4_47_CGCC;Ins_k4_48_GC | -1.9606824655153428 | 0.08765902504355783 | 6.91143004465892 |
| **GATCCAGATCTAACTTGGGGTGGCTTTGTCTTCTTCTTTTGCCAATTCGCGCACTAATTGTTTGGG** | Ins_k3_48_GCG | 0.3619712304964683 | 0.031728058919153555 | 70.51533932976689 |
| **GATCCAGATCTAACTTGGGGTGGCTTTGTCTTCTTCTTTTGCCAATTCGCTCACTAATTGTTTGGG** | Ins_k3_48_GCT | -0.27186257320311075 | 0.04057021913331166 | 37.41218530303244 |
| **GATCCAGATCTAACTTGGGGTGGCTTTGTCTTCTTCTTTTGCCAATTCGG** | Del_k13_49-61 | -0.12319462624581379 | 0.027266367993324683 | 43.40889649897151 |
| **GATCCAGATCTAACTTGGGGTGGCTTTGTCTTCTTCTTTTGCCAATTCGGACACTAATTGTTTGGG** | Ins_k3_48_GGA | 0.622515498259891 | 0.029750929868804184 | 91.50325386912318 |
| **GATCCAGATCTAACTTGGGGTGGCTTTGTCTTCTTCTTTTGCCAATTCGGCACTAATTGTTTGGG** | Ins_k2_48_GG | -0.4757542317494422 | 0.050851402179728035 | 30.51153482348189 |
| **GATCCAGATCTAACTTGGGGTGGCTTTGTCTTCTTCTTTTGCCAATTCGGCCACTAATTGTTTGGG** | Ins_k3_47_CGG;Ins_k3_48_GGC | 0.19515187880599172 | 0.03650758748265206 | 59.68083300200355 |
| **GATCCAGATCTAACTTGGGGTGGCTTTGTCTTCTTCTTTTGCCAATTCGGG** | Del_k12_49-60 | -0.3856946684067518 | 0.027658250591819058 | 33.3869253378814 |
| **GATCCAGATCTAACTTGGGGTGGCTTTGTCTTCTTCTTTTGCCAATTCGGGCACTAATTGTTTGGG** | Ins_k3_48_GGG | -0.15604399764935073 | 0.04185778713854762 | 42.00610803654686 |
| **GATCCAGATCTAACTTGGGGTGGCTTTGTCTTCTTCTTTTGCCAATTCGGGCCACTAATTGTTTGGG** | Ins_k4_47_CGGG;Ins_k4_48_GG | 0.38133308345368744 | 0.03972477696921856 | 71.89395012314438 |
| **GATCCAGATCTAACTTGGGGTGGCTTTGTCTTCTTCTTTTGCCAATTCGGTCACTAATTGTTTGGG** | Ins_k3_48_GGT | -0.035604353979033054 | 0.038340063260505236 | 47.382581431782484 |
| **GATCCAGATCTAACTTGGGGTGGCTTTGTCTTCTTCTTTTGCCAATTCGTACACTAATTGTTTGGG** | Ins_k3_48_GTA | -0.2801116408292381 | 0.04359571896767454 | 37.10483905923488 |

| | | | | |
|---|---|---|---|---|
| **GATCCAGATCTAACTTGGGGTGGCTTTGTCTTCTTCTTTTGCCAATTCGTCACTAATTGTTTGGG** | Ins_k2_48_GT | 0.6241117548544217 | 0.030045388401889892 | 91.64943318033838 |
| **GATCCAGATCTAACTTGGGGTGGCTTTGTCTTCTTCTTTTGCCAATTCGTCCACTAATTGTTTGGG** | Ins_k3_46_TCG;Ins_k3_47_CGT; | 0.5655082413879224 | 0.02817223585675231 | 86.43280369844695 |
| **GATCCAGATCTAACTTGGGGTGGCTTTGTCTTCTTCTTTTGCCAATTCGTGCACTAATTGTTTGGG** | Ins_k3_48_GTG | 0.05906561847444236 | 0.03471107196427072 | 52.08748210931908 |
| **GATCCAGATCTAACTTGGGGTGGCTTTGTCTTCTTCTTTTGCCAATTCGTGCCACTAATTGTTTGGG** | Ins_k4_47_CGTG;Ins_k4_48_GT | 0.5171738749317014 | 0.033838241083976664 | 82.3544843355791 |
| **GATCCAGATCTAACTTGGGGTGGCTTTGTCTTCTTCTTTTGCCAATTCGTTCACTAATTGTTTGGG** | Ins_k3_48_GTT | -0.303420682101219 | 0.04057861918140675 | 36.24996271321742 |
| **GATCCAGATCTAACTTGGGGTGGCTTTGTCTTCTTCTTTTGCCAATTCGTTTGGG** | Del_k8_49-56 | -0.007643570646016062 | 0.03151867880125037 | 48.726131347255844 |
| **GATCCAGATCTAACTTGGGGTGGCTTTGTCTTCTTCTTTTGCCAATTCTAACACTAATTGTTTGGG** | Ins_k3_48_TAA | -2.938814187052982 | 0.12783334701063556 | 2.598787130055061 |
| **GATCCAGATCTAACTTGGGGTGGCTTTGTCTTCTTCTTTTGCCAATTCTAATTGTTTGGG** | Del_k3_48-50;Del_k3_49-51 | -2.1537210426737072 | 0.08659119479826699 | 5.698129665454221 |
| **GATCCAGATCTAACTTGGGGTGGCTTTGTCTTCTTCTTTTGCCAATTCTACACTAATTGTTTGGG** | Ins_k2_48_TA | 0.29907979285807684 | 0.034134425590073694 | 66.2171059538643 |
| **GATCCAGATCTAACTTGGGGTGGCTTTGTCTTCTTCTTTTGCCAATTCTACCACTAATTGTTTGGG** | Ins_k3_47_CTA;Ins_k3_48_TAC | 0.005829536220286924 | 0.03481943581428056 | 49.38706614670729 |
| **GATCCAGATCTAACTTGGGGTGGCTTTGTCTTCTTCTTTTGCCAATTCTACTAATTGTTTGGG** | C49T | -0.06059195959488217 | 0.030384588429701796 | 46.21327410942962 |
| **GATCCAGATCTAACTTGGGGTGGCTTTGTCTTCTTCTTTTGCCAATTCTAGCACTAATTGTTTGGG** | Ins_k3_48_TAG | -4.00879777118906 | 0.23046362424255273 | 0.8914207538886857 |
| **GATCCAGATCTAACTTGGGGTGGCTTTGTCTTCTTCTTTTGCCAATTCTATCACTAATTGTTTGGG** | Ins_k3_48_TAT | -2.442346312941079 | 0.1226096413838541 | 4.269568301846415 |
| **GATCCAGATCTAACTTGGGGTGGCTTTGTCTTCTTCTTTTGCCAATTCTCACACTAATTGTTTGGG** | Ins_k3_48_TCA | -1.1036194403887838 | 0.06570634306852896 | 16.28492111049679 |
| **GATCCAGATCTAACTTGGGGTGGCTTTGTCTTCTTCTTTTGCCAATTCTCACTAATTGTTTGGG** | Ins_k1_48_T | -1.243837456038054 | 0.06570181040346088 | 14.154344075308416 |
| **GATCCAGATCTAACTTGGGGTGGCTTTGTCTTCTTCTTTTGCCAATTCTCCACTAATTGTTTGGG** | Ins_k2_46_TC;Ins_k2_47_CT;Ins_ | 0.23020879315735177 | 0.035688211572657245 | 61.81016467977689 |
| **GATCCAGATCTAACTTGGGGTGGCTTTGTCTTCTTCTTTTGCCAATTCTCCCACTAATTGTTTGGG** | Ins_k3_47_CTC;Ins_k3_48_TCC | -0.304240708700739 | 0.041441997054871074 | 36.220248964258296 |
| **GATCCAGATCTAACTTGGGGTGGCTTTGTCTTCTTCTTTTGCCAATTCTCGCACTAATTGTTTGGG** | Ins_k3_48_TCG | -0.3318998509122627 | 0.04135620916721412 | 35.2321558437394 |
| **GATCCAGATCTAACTTGGGGTGGCTTTGTCTTCTTCTTTTGCCAATTCTCTCACTAATTGTTTGGG** | Ins_k3_48_TCT | -1.9976791586584344 | 0.08819294111037784 | 6.660402220132921 |
| **GATCCAGATCTAACTTGGGGTGGCTTTGTCTTCTTCTTTTGCCAATTCTGACACTAATTGTTTGGG** | Ins_k3_48_TGA | -0.5999170713890946 | 0.04801258254844312 | 26.94888607324094 |
| **GATCCAGATCTAACTTGGGGTGGCTTTGTCTTCTTCTTTTGCCAATTCTGCACTAATTGTTTGGG** | Ins_k2_48_TG | 0.11093114378931068 | 0.03702860558868645 | 54.860312323275856 |
| **GATCCAGATCTAACTTGGGGTGGCTTTGTCTTCTTCTTTTGCCAATTCTGCCACTAATTGTTTGGG** | Ins_k3_47_CTG;Ins_k3_48_TGC | -0.005695411102975223 | 0.03303567469325772 | 48.82115015087771 |
| **GATCCAGATCTAACTTGGGGTGGCTTTGTCTTCTTCTTTTGCCAATTCTGGCACTAATTGTTTGGG** | Ins_k3_48_TGG | -1.5566043339423732 | 0.06884882600387375 | 10.352776047093698 |
| **GATCCAGATCTAACTTGGGGTGGCTTTGTCTTCTTCTTTTGCCAATTCTGGG** | Del_k11_49-59 | -0.3204266610689794 | 0.03341517309807716 | 35.63870882642255 |
| **GATCCAGATCTAACTTGGGGTGGCTTTGTCTTCTTCTTTTGCCAATTCTGTCACTAATTGTTTGGG** | Ins_k3_48_TGT | -1.5666317730751218 | 0.07413395903132118 | 10.249482963485361 |
| **GATCCAGATCTAACTTGGGGTGGCTTTGTCTTCTTCTTTTGCCAATTCTGTTTGGG** | Del_k7_49-55 | -0.3413715424163465 | 0.03786074700349028 | 34.90002314484125 |
| **GATCCAGATCTAACTTGGGGTGGCTTTGTCTTCTTCTTTTGCCAATTCTTACACTAATTGTTTGGG** | Ins_k3_48_TTA | -2.1680359947265386 | 0.10889926467450861 | 5.617142260884768 |
| **GATCCAGATCTAACTTGGGGTGGCTTTGTCTTCTTCTTTTGCCAATTCTTCACTAATTGTTTGGG** | Ins_k2_48_TT | 0.3589342756766368 | 0.03367793815903952 | 70.30151228599846 |
| **GATCCAGATCTAACTTGGGGTGGCTTTGTCTTCTTCTTTTGCCAATTCTTCCACTAATTGTTTGGG** | Ins_k3_45_TTC;Ins_k3_46_TCT;I | 0.07823784975381332 | 0.034664247117978324 | 53.0957498499707 |
| **GATCCAGATCTAACTTGGGGTGGCTTTGTCTTCTTCTTTTGCCAATTCTTGCACTAATTGTTTGGG** | Ins_k3_48_TTG | -1.5533223342992275 | 0.07448184492078647 | 10.386809673010395 |
| **GATCCAGATCTAACTTGGGGTGGCTTTGTCTTCTTCTTTTGCCAATTCTTGGG** | Del_k10_49-58 | -0.20221563594295405 | 0.03069929114459483 | 40.11071071841731 |
| **GATCCAGATCTAACTTGGGGTGGCTTTGTCTTCTTCTTTTGCCAATTCTTGTTTGGG** | Del_k6_49-54 | -0.7221101472154534 | 0.04553759176099648 | 23.849157400363016 |
| **GATCCAGATCTAACTTGGGGTGGCTTTGTCTTCTTCTTTTGCCAATTCTTTCACTAATTGTTTGGG** | Ins_k3_48_TTT | -3.0897528558507914 | 0.13935008370763846 | 2.2346981856064203 |
| **GATCCAGATCTAACTTGGGGTGGCTTTGTCTTCTTCTTTTGCCAATTCTTTGGG** | Del_k9_49-57 | -0.3969207485060796 | 0.03571224225231925 | 33.01421698090473 |
| **GATCCAGATCTAACTTGGGGTGGCTTTGTCTTCTTCTTTTGCCAATTG** | Del_k15_48-62 | -0.44615794117159824 | 0.029085157946707844 | 31.428059031748738 |
| **GATCCAGATCTAACTTGGGGTGGCTTTGTCTTCTTCTTTTGCCAATTGAACCACTAATTGTTTGGG** | Ins_k3_47_GAA | 0.3560510057539126 | 0.0328310234216709 | 70.0991059863213 |
| **GATCCAGATCTAACTTGGGGTGGCTTTGTCTTCTTCTTTTGCCAATTGACCACTAATTGTTTGGG** | Ins_k2_47_GA | 0.3294258816107041 | 0.036163709667048044 | 68.25733606436364 |
| **GATCCAGATCTAACTTGGGGTGGCTTTGTCTTCTTCTTTTGCCAATTGACCCACTAATTGTTTGGG** | Ins_k3_47_GAC | 0.43267776418094384 | 0.03148418051205323 | 75.6817311421156 |
| **GATCCAGATCTAACTTGGGGTGGCTTTGTCTTCTTCTTTTGCCAATTGAGCCACTAATTGTTTGGG** | Ins_k3_47_GAG | -0.056460117427574175 | 0.03674779727217145 | 46.404615087228265 |
| **GATCCAGATCTAACTTGGGGTGGCTTTGTCTTCTTCTTTTGCCAATTGATCCACTAATTGTTTGGG** | Ins_k3_46_TGA;Ins_k3_47_GAT | 0.47540897774996993 | 0.03154895331393869 | 78.98579392403849 |
| **GATCCAGATCTAACTTGGGGTGGCTTTGTCTTCTTCTTTTGCCAATTGCACCACTAATTGTTTGGG** | Ins_k3_47_GCA | 0.2698515542893569 | 0.03088483774369031 | 64.30970732219419 |
| **GATCCAGATCTAACTTGGGGTGGCTTTGTCTTCTTCTTTTGCCAATTGCACTAATTGTTTGGG** | C48G | -0.913933576919282 | 0.04310345176496194 | 19.686349127891603 |
| **GATCCAGATCTAACTTGGGGTGGCTTTGTCTTCTTCTTTTGCCAATTGCCACTAATTGTTTGGG** | Ins_k1_47_G | -0.2313591189009253 | 0.03844208497534947 | 38.9586144950348 |
| **GATCCAGATCTAACTTGGGGTGGCTTTGTCTTCTTCTTTTGCCAATTGCCCACTAATTGTTTGGG** | Ins_k2_47_GC | -0.21345029432489787 | 0.04322065710183065 | 39.66260247082928 |
| **GATCCAGATCTAACTTGGGGTGGCTTTGTCTTCTTCTTTTGCCAATTGCCCCACTAATTGTTTGGG** | Ins_k3_47_GCC | -3.7647413840077606 | 0.18322391618060885 | 1.1378239821847997 |
| **GATCCAGATCTAACTTGGGGTGGCTTTGTCTTCTTCTTTTGCCAATTGCCCCCACTAATTGTTTGGG** | Ins_k4_47_GCCC | -2.388833682415174 | 0.1265547126370352 | 4.504267822774294 |
| **GATCCAGATCTAACTTGGGGTGGCTTTGTCTTCTTCTTTTGCCAATTGCGCCACTAATTGTTTGGG** | Ins_k3_47_GCG | 0.33972319406325224 | 0.032448396765271746 | 68.96383445557333 |
| **GATCCAGATCTAACTTGGGGTGGCTTTGTCTTCTTCTTTTGCCAATTGCTCCACTAATTGTTTGGG** | Ins_k3_46_TGC;Ins_k3_47_GCT | 0.24362100615175586 | 0.03202234722366006 | 62.64476014871077 |
| **GATCCAGATCTAACTTGGGGTGGCTTTGTCTTCTTCTTTTGCCAATTGG** | Del_k14_48-61 | -0.4362685083726877 | 0.02990973136676599 | 31.740406634203854 |
| **GATCCAGATCTAACTTGGGGTGGCTTTGTCTTCTTCTTTTGCCAATTGGACCACTAATTGTTTGGG** | Ins_k3_47_GGA | 0.6657723599348342 | 0.03190807190466495 | 95.54825374884179 |
| **GATCCAGATCTAACTTGGGGTGGCTTTGTCTTCTTCTTTTGCCAATTGGCCACTAATTGTTTGGG** | Ins_k2_47_GG | -0.30967535975474336 | 0.04645364793458384 | 36.02393847273426 |
| **GATCCAGATCTAACTTGGGGTGGCTTTGTCTTCTTCTTTTGCCAATTGGCCCACTAATTGTTTGGG** | Ins_k3_47_GGC | 0.14373810422895295 | 0.03651246032823769 | 56.689960923211856 |
| **GATCCAGATCTAACTTGGGGTGGCTTTGTCTTCTTCTTTTGCCAATTGGG** | Del_k13_46-58;Del_k13_47-59;D | -0.25604924479625396 | 0.028720393789363696 | 38.008498900828094 |
| **GATCCAGATCTAACTTGGGGTGGCTTTGTCTTCTTCTTTTGCCAATTGGGCCACTAATTGTTTGGG** | Ins_k3_47_GGG | -0.10464996498633017 | 0.039864013133834 | 44.2214103921895 |
| **GATCCAGATCTAACTTGGGGTGGCTTTGTCTTCTTCTTTTGCCAATTGGTCCACTAATTGTTTGGG** | Ins_k3_46_TGG;Ins_k3_47_GGT | 0.2755593923407865 | 0.035267655646005744 | 64.67782629905136 |
| **GATCCAGATCTAACTTGGGGTGGCTTTGTCTTCTTCTTTTGCCAATTGTACCACTAATTGTTTGGG** | Ins_k3_47_GTA | -0.23287371906227888 | 0.034444465396404875 | 38.89965243447937 |
| **GATCCAGATCTAACTTGGGGTGGCTTTGTCTTCTTCTTTTGCCAATTGTCCACTAATTGTTTGGG** | Ins_k2_46_TG;Ins_k2_47_GT | 0.06271865361604645 | 0.0359132635019011 | 52.27810748050609 |
| **GATCCAGATCTAACTTGGGGTGGCTTTGTCTTCTTCTTTTGCCAATTGTCCCACTAATTGTTTGGG** | Ins_k3_47_GTC | -0.4980036761641465 | 0.047933327985808016 | 29.840166606160032 |
| **GATCCAGATCTAACTTGGGGTGGCTTTGTCTTCTTCTTTTGCCAATTGTGCCACTAATTGTTTGGG** | Ins_k3_47_GTG | -0.08101364549925727 | 0.03553864822340491 | 45.27909238989319 |
| **GATCCAGATCTAACTTGGGGTGGCTTTGTCTTCTTCTTTTGCCAATTGTTCCACTAATTGTTTGGG** | Ins_k3_45_TTG;Ins_k3_46_TGT;I | 0.09341633107053704 | 0.03967114979705034 | 53.9078100266074 |
| **GATCCAGATCTAACTTGGGGTGGCTTTGTCTTCTTCTTTTGCCAATTGTTTGGG** | Del_k9_44-52;Del_k9_45-53;Del_ | -0.23999463589651682 | 0.03402919255511348 | 38.623635158987994 |
| **GATCCAGATCTAACTTGGGGTGGCTTTGTCTTCTTCTTTTGCCAATTTAACCACTAATTGTTTGGG** | Ins_k3_47_TAA | -0.838844132390084 | 0.05318783752604355 | 21.221501776084168 |
| **GATCCAGATCTAACTTGGGGTGGCTTTGTCTTCTTCTTTTGCCAATTTAATTGTTTGGG** | Del_k4_48-51 | -1.5978593662815261 | 0.05960211037471367 | 9.934362119116782 |
| **GATCCAGATCTAACTTGGGGTGGCTTTGTCTTCTTCTTTTGCCAATTTACCACTAATTGTTTGGG** | Ins_k2_47_TA | -0.45404024068071247 | 0.04352375810302765 | 31.181307420092505 |
| **GATCCAGATCTAACTTGGGGTGGCTTTGTCTTCTTCTTTTGCCAATTTACCCACTAATTGTTTGGG** | Ins_k3_47_TAC | -1.3474580226253705 | 0.0626148243395406 | 12.761094016034614 |
| **GATCCAGATCTAACTTGGGGTGGCTTTGTCTTCTTCTTTTGCCAATTTAGCCACTAATTGTTTGGG** | Ins_k3_47_TAG | -2.5009228931866367 | 0.11381637910960218 | 4.026655544123818 |
| **GATCCAGATCTAACTTGGGGTGGCTTTGTCTTCTTCTTTTGCCAATTTATACCACTAATTGTTTGGG** | Ins_k4_47_TATA | -0.8119923495335126 | 0.05709247797722877 | 21.799056417709238 |
| **GATCCAGATCTAACTTGGGGTGGCTTTGTCTTCTTCTTTTGCCAATTTATCCACTAATTGTTTGGG** | Ins_k3_46_TTA;Ins_k3_47_TAT | -0.4062589015919917 | 0.04053532876906601 | 32.707360136670346 |
| **GATCCAGATCTAACTTGGGGTGGCTTTGTCTTCTTCTTTTGCCAATTTCACCACTAATTGTTTGGG** | Ins_k3_47_TCA | 0.1545858881630333 | 0.033894062241946056 | 57.308268943200034 |
| **GATCCAGATCTAACTTGGGGTGGCTTTGTCTTCTTCTTTTGCCAATTTCACTAATTGTTTGGG** | C48T | -1.1311580249294624 | 0.041423896020573635 | 15.842576165464992 |
| **GATCCAGATCTAACTTGGGGTGGCTTTGTCTTCTTCTTTTGCCAATTTCCACTAATTGTTTGGG** | Ins_k1_45_T;Ins_k1_46_T;Ins_k1_ | -0.20309031511112424 | 0.04432952376019125 | 40.07564205448212 |
| **GATCCAGATCTAACTTGGGGTGGCTTTGTCTTCTTCTTTTGCCAATTTCCCACTAATTGTTTGGG** | Ins_k2_47_TC | -1.0146700784298035 | 0.0630902854984804 | 17.79983088429623 |
| **GATCCAGATCTAACTTGGGGTGGCTTTGTCTTCTTCTTTTGCCAATTTCCCCACTAATTGTTTGGG** | Ins_k3_47_TCC | -2.687210031434008 | 0.10811120361221402 | 3.342265944192765 |
| **GATCCAGATCTAACTTGGGGTGGCTTTGTCTTCTTCTTTTGCCAATTTCGCCACTAATTGTTTGGG** | Ins_k3_47_TCG | 0.2720046791924584 | 0.029622946464157803 | 64.44832332977835 |
| **GATCCAGATCTAACTTGGGGTGGCTTTGTCTTCTTCTTTTGCCAATTTCTCCACTAATTGTTTGGG** | Ins_k3_46_TTC;Ins_k3_47_TCT | 0.19350378938633783 | 0.03397424698459471 | 59.58255466056946 |

| | | | | |
|---|---|---|---|---|
| GATCCAGATCTAACTTGGGGTGGCTTTGTCTTCTTCTTTTGCCAATTTGACCACTAATTGTTTGGG | Ins_k3_47_TGA | 0.2597305300567194 | 0.03312826087537672 | 63.66210991855107 |
| GATCCAGATCTAACTTGGGGTGGCTTTGTCTTCTTCTTTTGCCAATTTGCCACTAATTGTTTGGG | Ins_k2_47_TG | -1.077302732741995 | 0.6198779846356436 | 16.719175630723353 |
| GATCCAGATCTAACTTGGGGTGGCTTTGTCTTCTTCTTTTGCCAATTTGCCCACTAATTGTTTGGG | Ins_k3_47_TGC | -0.5791343902384414 | 0.042055631899778186 | 27.514816587005633 |
| GATCCAGATCTAACTTGGGGTGGCTTTGTCTTCTTCTTTTGCCAATTTGGCCACTAATTGTTTGGG | Ins_k3_47_TGG | -0.8670030284564467 | 0.056454850588638666 | 20.632262807240945 |
| GATCCAGATCTAACTTGGGGTGGCTTTGTCTTCTTCTTTTGCCAATTTGGG | Del_k12_46-57;Del_k12_47-58;D | -0.3360005326796373 | 0.03466766597052583 | 35.087975804913214 |
| GATCCAGATCTAACTTGGGGTGGCTTTGTCTTCTTCTTTTGCCAATTTGTCCACTAATTGTTTGGG | Ins_k3_46_TTG;Ins_k3_47_TGT | -0.15932955928400971 | 0.03594489334553683 | 41.868320857645074 |
| GATCCAGATCTAACTTGGGGTGGCTTTGTCTTCTTCTTTTGCCAATTTGTTTGGG | Del_k8_47-54;Del_k8_48-55 | -0.41280217296115673 | 0.0362565811394169 | 32.49404565188466 |
| GATCCAGATCTAACTTGGGGTGGCTTTGTCTTCTTCTTTTGCCAATTTTACCACTAATTGTTTGGG | Ins_k3_47_TTA | -0.8430424102926838 | 0.05512748397458418 | 21.132594772857274 |
| GATCCAGATCTAACTTGGGGTGGCTTTGTCTTCTTCTTTTGCCAATTTTCCACTAATTGTTTGGG | Ins_k2_45_TT;Ins_k2_46_TT;Ins_I | -0.4611872951102679 | 0.04730504412896875 | 30.95924740153786 |
| GATCCAGATCTAACTTGGGGTGGCTTTGTCTTCTTCTTTTGCCAATTTTCCCACTAATTGTTTGGG | Ins_k3_47_TTC | -1.512031119697463 | 0.06451543898121961 | 10.824671371134924 |
| GATCCAGATCTAACTTGGGGTGGCTTTGTCTTCTTCTTTTGCCAATTTTGCCACTAATTGTTTGGG | Ins_k3_47_TTG | -0.6228171992576581 | 0.04857018526312679 | 26.338765716177114 |
| GATCCAGATCTAACTTGGGGTGGCTTTGTCTTCTTCTTTTGCCAATTTTGGG | Del_k11_47-57;Del_k11_48-58 | -0.2806300715199031 | 0.03219610751915883 | 37.085607757373666 |
| GATCCAGATCTAACTTGGGGTGGCTTTGTCTTCTTCTTTTGCCAATTTTGTTTGGG | Del_k7_48-54 | -0.531306931117139 | 0.03843669130572107 | 28.862757714491636 |
| GATCCAGATCTAACTTGGGGTGGCTTTGTCTTCTTCTTTTGCCAATTTTTCCACTAATTGTTTGGG | Ins_k3_45_TTT;Ins_k3_46_TTT;In | -1.3197755154837982 | 0.06072522473491357 | 13.119288073021874 |
| GATCCAGATCTAACTTGGGGTGGCTTTGTCTTCTTCTTTTGCCAATTTTTGGG | Del_k10_48-57 | -0.09275530557110656 | 0.03348765846723266 | 44.75054973513216 |
| GATCCAGATCTAACTTGGGGTGGCTTTGTCTTCTTCTTTTGCCACAAATTCCACTAATTGTTTGGG | Ins_k3_43_ACA;Ins_k3_44_CAA | 0.020062213288708164 | 0.03664495093226581 | 50.0950022732088 |
| GATCCAGATCTAACTTGGGGTGGCTTTGTCTTCTTCTTTTGCCACAATTCCACTAATTGTTTGGG | Ins_k2_42_CA;Ins_k2_43_AC;Ins | 0.17934553990832958 | 0.03374619101831431 | 58.74491374414941 |
| GATCCAGATCTAACTTGGGGTGGCTTTGTCTTCTTCTTTTGCCACACATTCCACTAATTGTTTGGG | Ins_k3_44_CAC | -0.07825866343044086 | 0.03613899775500747 | 45.40400746791165 |
| GATCCAGATCTAACTTGGGGTGGCTTTGTCTTCTTCTTTTGCCACACTAATTGTTTGGG | Del_k4_45-48 | -0.81200422233901 | 0.041787637665403894 | 21.798797603288794 |
| GATCCAGATCTAACTTGGGGTGGCTTTGTCTTCTTCTTTTGCCACAGATTCCACTAATTGTTTGGG | Ins_k3_44_CAG | 0.00822525313191158 | 0.035516678774786634 | 49.50552541707148 |
| GATCCAGATCTAACTTGGGGTGGCTTTGTCTTCTTCTTTTGCCACATATTCCACTAATTGTTTGGG | Ins_k3_44_CAT | -0.20210416649051277 | 0.03775519127490472 | 40.115182086583964 |
| GATCCAGATCTAACTTGGGGTGGCTTTGTCTTCTTCTTTTGCCACATTCCACTAATTGTTTGGG | Ins_k1_44_C | -0.08708333625666309 | 0.036478998790775534 | 45.00509468318312 |
| GATCCAGATCTAACTTGGGGTGGCTTTGTCTTCTTCTTTTGCCACCAATTCCACTAATTGTTTGGG | Ins_k3_41_CCA;Ins_k3_42_CAC; | -0.6858340904230186 | 0.04607225065238725 | 24.730194448899834 |
| GATCCAGATCTAACTTGGGGTGGCTTTGTCTTCTTCTTTTGCCACCACTAATTGTTTGGG | Del_k3_45-47 | -2.0276390170854244 | 0.07348522554847785 | 6.463817048740895 |
| GATCCAGATCTAACTTGGGGTGGCTTTGTCTTCTTCTTTTGCCACCATTCCACTAATTGTTTGGG | Ins_k2_44_CC | -0.5384500889961483 | 0.05557569857033926 | 28.65732108597709 |
| GATCCAGATCTAACTTGGGGTGGCTTTGTCTTCTTCTTTTGCCACCCATTCCACTAATTGTTTGGG | Ins_k3_44_CCC | -1.8822001400628265 | 0.07744578180091127 | 7.47570848014295 |
| GATCCAGATCTAACTTGGGGTGGCTTTGTCTTCTTCTTTTGCCACCGATTCCACTAATTGTTTGGG | Ins_k3_44_CCG | 0.007448337206701905 | 0.038470685247596385 | 49.467078722847454 |
| GATCCAGATCTAACTTGGGGTGGCTTTGTCTTCTTCTTTTGCCACCTATTCCACTAATTGTTTGGG | Ins_k3_44_CCT | -1.5215475493900301 | 0.06954627088945498 | 10.722147750232667 |
| GATCCAGATCTAACTTGGGGTGGCTTTGTCTTCTTCTTTTGCCACGAATTCCACTAATTGTTTGGG | Ins_k3_43_ACG;Ins_k3_44_CGA | 0.6878995820354885 | 0.029115248800160868 | 97.68603554389092 |
| GATCCAGATCTAACTTGGGGTGGCTTTGTCTTCTTCTTTTGCCACGATTCCACTAATTGTTTGGG | Ins_k2_44_CG | 0.7926620644285387 | 0.029449774206895273 | 100 |
| GATCCAGATCTAACTTGGGGTGGCTTTGTCTTCTTCTTTTGCCACGCATTCCACTAATTGTTTGGG | Ins_k3_44_CGC | 0.4774964852522279 | 0.03016256737232673 | 79.15084957894727 |
| GATCCAGATCTAACTTGGGGTGGCTTTGTCTTCTTCTTTTGCCACGGATTCCACTAATTGTTTGGG | Ins_k3_44_CGG | 0.8553109655598257 | 0.028606255068026897 | 100 |
| GATCCAGATCTAACTTGGGGTGGCTTTGTCTTCTTCTTTTGCCACGTATTCCACTAATTGTTTGGG | Ins_k3_44_CGT | 0.2597062011898694 | 0.032910948159654535 | 63.660561110395925 |
| GATCCAGATCTAACTTGGGGTGGCTTTGTCTTCTTCTTTTGCCACGTTATTCCACTAATTGTTTGGG | Ins_k4_44_CGTT | 0.30448642956258465 | 0.03416142923612943 | 66.57608735592315 |
| GATCCAGATCTAACTTGGGGTGGCTTTGTCTTCTTCTTTTGCCACTAATTCCACTAATTGTTTGGG | Ins_k3_43_ACT;Ins_k3_44_CTA | -0.8283873500002591 | 0.04895144732511835 | 21.44457468191649 |
| GATCCAGATCTAACTTGGGGTGGCTTTGTCTTCTTCTTTTGCCACTAATTGTTTGGG | Del_k6_42-47;Del_k6_43-48;Del_ | -0.8603400538299572 | 0.04177091896966555 | 20.770194056730002 |
| GATCCAGATCTAACTTGGGGTGGCTTTGTCTTCTTCTTTTGCCACTATTCCACTAATTGTTTGGG | Ins_k2_44_CT | -0.47893787185582304 | 0.04659025596336868 | 30.41455153936211 |
| GATCCAGATCTAACTTGGGGTGGCTTTGTCTTCTTCTTTTGCCACTCATTCCACTAATTGTTTGGG | Ins_k3_44_CTC | -2.0778549243512545 | 0.08133850283920446 | 6.147245593073166 |
| GATCCAGATCTAACTTGGGGTGGCTTTGTCTTCTTCTTTTGCCACTGATTCCACTAATTGTTTGGG | Ins_k3_44_CTG | 0.28334011170984474 | 0.03236219733621333 | 65.18302918710424 |
| GATCCAGATCTAACTTGGGGTGGCTTTGTCTTCTTCTTTTGCCACTTATTCCACTAATTGTTTGGG | Ins_k3_44_CTT | -0.7712778777605012 | 0.04543767284268495 | 22.70490900966912 |
| GATCCAGATCTAACTTGGGGTGGCTTTGTCTTCTTCTTTTGCCACTTCCACTAATTGTTTGGG | A45C | -2.100796132256316 | 0.08887014497292445 | 6.007825698927027 |
| GATCCAGATCTAACTTGGGGTGGCTTTGTCTTCTTCTTTTGCCAG | Del_k18_45-62 | -0.1043819225359785 | 0.024206314760345283 | 44.2332651961134 |
| GATCCAGATCTAACTTGGGGTGGCTTTGTCTTCTTCTTTTGCCAGAAATTCCACTAATTGTTTGGG | Ins_k3_43_AGA;Ins_k3_44_GAA | 0.495539506290122 | 0.02886637716132876 | 80.59193166504883 |
| GATCCAGATCTAACTTGGGGTGGCTTTGTCTTCTTCTTTTGCCAGAATTCCACTAATTGTTTGGG | Ins_k2_43_AG;Ins_k2_44_GA | 0.609860516233425 | 0.031500668702869765 | 90.35257808328943 |
| GATCCAGATCTAACTTGGGGTGGCTTTGTCTTCTTCTTTTGCCAGACATTCCACTAATTGTTTGGG | Ins_k3_44_GAC | 0.3328111068368844 | 0.031934121004646994 | 68.48879406797938 |
| GATCCAGATCTAACTTGGGGTGGCTTTGTCTTCTTCTTTTGCCAGAGATTCCACTAATTGTTTGGG | Ins_k3_44_GAG | 0.5627889581637253 | 0.029212144050415532 | 86.19808769960244 |
| GATCCAGATCTAACTTGGGGTGGCTTTGTCTTCTTCTTTTGCCAGATATTCCACTAATTGTTTGGG | Ins_k3_44_GAT | 0.10083281925947407 | 0.035304113347183023 | 54.30910291554566 |
| GATCCAGATCTAACTTGGGGTGGCTTTGTCTTCTTCTTTTGCCAGATTCCACTAATTGTTTGGG | Ins_k1_44_G | 0.35820168955645304 | 0.033211680024996346 | 70.25002923405695 |
| GATCCAGATCTAACTTGGGGTGGCTTTGTCTTCTTCTTTTGCCAGCAATTCCACTAATTGTTTGGG | Ins_k3_42_CAG;Ins_k3_43_AGC | 0.27740142829101133 | 0.03351207677505155 | 64.79707497673373 |
| GATCCAGATCTAACTTGGGGTGGCTTTGTCTTCTTCTTTTGCCAGCATTCCACTAATTGTTTGGG | Ins_k2_44_GC | 0.457162241351156 | 0.0335594029069167 | 77.55763025068119 |
| GATCCAGATCTAACTTGGGGTGGCTTTGTCTTCTTCTTTTGCCAGCCATTCCACTAATTGTTTGGG | Ins_k3_44_GCC | -0.6588578700288051 | 0.046173287598404174 | 25.406401371487664 |
| GATCCAGATCTAACTTGGGGTGGCTTTGTCTTCTTCTTTTGCCAGCGATTCCACTAATTGTTTGGG | Ins_k3_44_GCG | 0.6442890461614128 | 0.02833558324574222 | 93.5174529050827 |
| GATCCAGATCTAACTTGGGGTGGCTTTGTCTTCTTCTTTTGCCAGCTATTCCACTAATTGTTTGGG | Ins_k3_44_GCT | -0.017078743557688125 | 0.03485488286561593 | 48.26855392826362 |
| GATCCAGATCTAACTTGGGGTGGCTTTGTCTTCTTCTTTTGCCAGG | Del_k17_45-61 | -0.5561822100913021 | 0.027965754859744375 | 28.153644819918327 |
| GATCCAGATCTAACTTGGGGTGGCTTTGTCTTCTTCTTTTGCCAGGAATTCCACTAATTGTTTGGG | Ins_k3_43_AGG;Ins_k3_44_GGA | 0.10467292290784203 | 0.03531044750290942 | 54.51805644462001 |
| GATCCAGATCTAACTTGGGGTGGCTTTGTCTTCTTCTTTTGCCAGGATTCCACTAATTGTTTGGG | Ins_k2_44_GG | 0.4871540746981693 | 0.0352042137400507 | 79.91895906143407 |
| GATCCAGATCTAACTTGGGGTGGCTTTGTCTTCTTCTTTTGCCAGGCATTCCACTAATTGTTTGGG | Ins_k3_44_GGC | -1.004133597713603 | 0.047525562439591794 | 17.988369984198194 |
| GATCCAGATCTAACTTGGGGTGGCTTTGTCTTCTTCTTTTGCCAGGG | Del_k16_45-60 | -0.39119566591213373 | 0.0278317501074442 | 33.20376818029476 |
| GATCCAGATCTAACTTGGGGTGGCTTTGTCTTCTTCTTTTGCCAGGGATTCCACTAATTGTTTGGG | Ins_k3_44_GGG | -0.07126768686430707 | 0.0400255140914704 | 45.72253794235567 |
| GATCCAGATCTAACTTGGGGTGGCTTTGTCTTCTTCTTTTGCCAGGGTATTCCACTAATTGTTTGGG | Ins_k4_44_GGGT | -2.772805367558509 | 0.13833950924409433 | 3.068085241689143 |
| GATCCAGATCTAACTTGGGGTGGCTTTGTCTTCTTCTTTTGCCAGGTATTCCACTAATTGTTTGGG | Ins_k3_44_GGT | -2.7506019974436677 | 0.1437122704302234 | 3.136968969414586 |
| GATCCAGATCTAACTTGGGGTGGCTTTGTCTTCTTCTTTTGCCAGTAATTCCACTAATTGTTTGGG | Ins_k3_43_AGT;Ins_k3_44_GTA | 0.002388806629536888 | 0.034565945504584014 | 49.217430609176986 |
| GATCCAGATCTAACTTGGGGTGGCTTTGTCTTCTTCTTTTGCCAGTATTCCACTAATTGTTTGGG | Ins_k2_44_GT | 0.20033686523064131 | 0.037661239516632826 | 59.99108093152149 |
| GATCCAGATCTAACTTGGGGTGGCTTTGTCTTCTTCTTTTGCCAGTCATTCCACTAATTGTTTGGG | Ins_k3_44_GTC | 0.06409495354696726 | 0.03433240381964483 | 52.35010737157753 |
| GATCCAGATCTAACTTGGGGTGGCTTTGTCTTCTTCTTTTGCCAGTGATTCCACTAATTGTTTGGG | Ins_k3_44_GTG | 0.4016121332046787 | 0.03286088570355354 | 73.3667743793459 |
| GATCCAGATCTAACTTGGGGTGGCTTTGTCTTCTTCTTTTGCCAGTTATTCCACTAATTGTTTGGG | Ins_k3_44_GTT | 0.047595127903072854 | 0.038419820233411346 | 51.49342670377666 |
| GATCCAGATCTAACTTGGGGTGGCTTTGTCTTCTTCTTTTGCCAGTTCATTCCACTAATTGTTTGGG | Ins_k4_44_GTTC | 0.2097426410102028 | 0.04052320988002272 | 60.55800559107213 |
| GATCCAGATCTAACTTGGGGTGGCTTTGTCTTCTTCTTTTGCCAGTTCCACTAATTGTTTGGG | A45G | 0.2725650437108148 | 0.028004598690948967 | 64.48444800398747 |
| GATCCAGATCTAACTTGGGGTGGCTTTGTCTTCTTCTTTTGCCAGTTTGGG | Del_k12_45-56 | -0.08845786473756279 | 0.029771087931980418 | 44.943276393987986 |
| GATCCAGATCTAACTTGGGGTGGCTTTGTCTTCTTCTTTTGCCATAAATTCCACTAATTGTTTGGG | Ins_k3_43_ATA;Ins_k3_44_TAA | -0.5058044972321687 | 0.04213424372573889 | 29.60829437854306 |
| GATCCAGATCTAACTTGGGGTGGCTTTGTCTTCTTCTTTTGCCATAATTCCACTAATTGTTTGGG | Ins_k2_43_AT;Ins_k2_44_TA | -0.22563393589173836 | 0.0407390436521342 | 39.182299400323046 |
| GATCCAGATCTAACTTGGGGTGGCTTTGTCTTCTTCTTTTGCCATAATTGTTTGGG | Del_k7_45-51 | -0.9135503007550393 | 0.04132255311611895 | 19.693895882426464 |
| GATCCAGATCTAACTTGGGGTGGCTTTGTCTTCTTCTTTTGCCATACATTCCACTAATTGTTTGGG | Ins_k3_44_TAC | -0.1424765946685791 | 0.03680872514590062 | 42.579905500887094 |
| GATCCAGATCTAACTTGGGGTGGCTTTGTCTTCTTCTTTTGCCATAGATTCCACTAATTGTTTGGG | Ins_k3_44_TAG | -0.1994459889390643 | 0.03956882601487094 | 40.22195721383002 |
| GATCCAGATCTAACTTGGGGTGGCTTTGTCTTCTTCTTTTGCCATATATTCCACTAATTGTTTGGG | Ins_k3_44_TAT | -0.45289524616294363 | 0.043348325113300945 | 31.217030293489717 |

| | | | | |
|---|---|---|---|---|
| **GATCCAGATCTAACTTGGGGTGGCTTTGTCTTCTTCTTTTGCCATATTCCACTAATTGTTTGGG** | Ins_k1_44_T | -0.07853958796357152 | 0.03581373387036378 | 45.391254159753906 |
| **GATCCAGATCTAACTTGGGGTGGCTTTGTCTTCTTCTTTTGCCATCAATTCCACTAATTGTTTGGG** | Ins_k3_42_CAT;Ins_k3_43_ATC;Ir | 0.35822356815195555 | 0.03499525349202081 | 70.25156622284412 |
| **GATCCAGATCTAACTTGGGGTGGCTTTGTCTTCTTCTTTTGCCATCATTCCACTAATTGTTTGGG** | Ins_k2_44_TC | 0.345542398153046 | 0.03320215969906182 | 69.36631901720682 |
| **GATCCAGATCTAACTTGGGGTGGCTTTGTCTTCTTCTTTTGCCATCCACTAATTGTTTGGG** | Del_k2_45-46 | -1.3535458668803575 | 0.06277473393265448 | 12.683642458858449 |
| **GATCCAGATCTAACTTGGGGTGGCTTTGTCTTCTTCTTTTGCCATCCATTCCACTAATTGTTTGGG** | Ins_k3_44_TCC | -1.1231998568881019 | 0.05312788175411945 | 15.969157056743954 |
| **GATCCAGATCTAACTTGGGGTGGCTTTGTCTTCTTCTTTTGCCATCGAATTCCACTAATTGTTTGGG** | Ins_k4_43_ATCG;Ins_k4_44_TCG | 0.9529051044943502 | 0.034002703211353826 | 100 |
| **GATCCAGATCTAACTTGGGGTGGCTTTGTCTTCTTCTTTTGCCATCGATTCCACTAATTGTTTGGG** | Ins_k3_44_TCG | 0.6837028263896852 | 0.03167801753383026 | 97.2769301807831 |
| **GATCCAGATCTAACTTGGGGTGGCTTTGTCTTCTTCTTTTGCCATCTATTCCACTAATTGTTTGGG** | Ins_k3_44_TCT | -0.04299720364753812 | 0.03705132779792781 | 47.033580774576706 |
| **GATCCAGATCTAACTTGGGGTGGCTTTGTCTTCTTCTTTTGCCATGAATTCCACTAATTGTTTGGG** | Ins_k3_43_ATG;Ins_k3_44_TGA | 0.6466630653139767 | 0.029724916580732574 | 93.73972886868073 |
| **GATCCAGATCTAACTTGGGGTGGCTTTGTCTTCTTCTTTTGCCATGATTCCACTAATTGTTTGGG** | Ins_k2_44_TG | 0.534197082147295 | 0.03190132812269769 | 83.76842252372542 |
| **GATCCAGATCTAACTTGGGGTGGCTTTGTCTTCTTCTTTTGCCATGCATTCCACTAATTGTTTGGG** | Ins_k3_44_TGC | 0.10585462785679911 | 0.03265880627764775 | 54.58251878195826 |
| **GATCCAGATCTAACTTGGGGTGGCTTTGTCTTCTTCTTTTGCCATGCTATTCCACTAATTGTTTGGG** | Ins_k4_44_TGCT | -0.23390470084460296 | 0.037857593910560774 | 38.85956826805389 |
| **GATCCAGATCTAACTTGGGGTGGCTTTGTCTTCTTCTTTTGCCATGGATTCCACTAATTGTTTGGG** | Ins_k3_44_TGG | 0.7123965369931409 | 0.027837108945282935 | 100 |
| **GATCCAGATCTAACTTGGGGTGGCTTTGTCTTCTTCTTTTGCCATGGG** | Del_k15_45-59 | -0.479636939473953 | 0.028292317886330125 | 30.39329714125731 |
| **GATCCAGATCTAACTTGGGGTGGCTTTGTCTTCTTCTTTTGCCATGTATTCCACTAATTGTTTGGG** | Ins_k3_44_TGT | -0.31214521891050023 | 0.03827109523016658 | 35.93507420475807 |
| **GATCCAGATCTAACTTGGGGTGGCTTTGTCTTCTTCTTTTGCCATGTTTGGG** | Del_k11_45-55 | -0.15234054429113508 | 0.029806405626653838 | 42.16196412315622 |
| **GATCCAGATCTAACTTGGGGTGGCTTTGTCTTCTTCTTTTGCCATTAATTCCACTAATTGTTTGGG** | Ins_k3_43_ATT;Ins_k3_44_TTA | -0.6665497490900258 | 0.04716810558929303 | 25.2117280663368 |
| **GATCCAGATCTAACTTGGGGTGGCTTTGTCTTCTTCTTTTGCCATTATTCCACTAATTGTTTGGG** | Ins_k2_44_TT | -0.26246340432926585 | 0.0422836779652285 | 37.76548651851829 |
| **GATCCAGATCTAACTTGGGGTGGCTTTGTCTTCTTCTTTTGCCATTCATTCCACTAATTGTTTGGG** | Ins_k3_44_TTC | -0.2932812770393458 | 0.03839807781820215 | 36.61938546716708 |
| **GATCCAGATCTAACTTGGGGTGGCTTTGTCTTCTTCTTTTGCCATTCCACTAATTGTTTGGG** | Del_k1_44-44;Del_k1_45-45 | -1.8801059726383558 | 0.06418316219786106 | 7.491380269264602 |
| **GATCCAGATCTAACTTGGGGTGGCTTTGTCTTCTTCTTTTGCCATTGATTCCACTAATTGTTTGGG** | Ins_k3_44_TTG | -0.4460684571617356 | 0.031036781691011508 | 31.430871466325183 |
| **GATCCAGATCTAACTTGGGGTGGCTTTGTCTTCTTCTTTTGCCATTGGG** | Del_k14_45-58 | -0.31505550591957776 | 0.030924994416897316 | 35.83064485852229 |
| **GATCCAGATCTAACTTGGGGTGGCTTTGTCTTCTTCTTTTGCCATTGTATTCCACTAATTGTTTGGG** | Ins_k4_44_TTGT | -0.46485243710198615 | 0.043091568150656054 | 30.845985051941526 |
| **GATCCAGATCTAACTTGGGGTGGCTTTGTCTTCTTCTTTTGCCATTGTTTGGG** | Del_k10_44-53;Del_k10_45-54 | -0.08977166828698835 | 0.03061105062647508 | 44.88426852879781 |
| **GATCCAGATCTAACTTGGGGTGGCTTTGTCTTCTTCTTTTGCCATTTATTCCACTAATTGTTTGGG** | Ins_k3_44_TTT | -0.7500769108798104 | 0.047922554266373936 | 23.191413998935463 |
| **GATCCAGATCTAACTTGGGGTGGCTTTGTCTTCTTCTTTTGCCATTTCCACTAATTGTTTGGG** | A45T | -0.5648341877447094 | 0.03936128148017447 | 27.91111082636032 |
| **GATCCAGATCTAACTTGGGGTGGCTTTGTCTTCTTCTTTTGCCATTTGGG** | Del_k13_45-57 | -0.22798746979482504 | 0.028185531205871653 | 39.09019096296519 |
| **GATCCAGATCTAACTTGGGGTGGCTTTGTCTTCTTCTTTTGCCCAAAATTCCACTAATTGTTTGGG** | Ins_k3_43_CAA | -0.21397112909681631 | 0.04441723750956049 | 39.64195018699832 |
| **GATCCAGATCTAACTTGGGGTGGCTTTGTCTTCTTCTTTTGCCCAAATTCCACTAATTGTTTGGG** | Ins_k2_43_CA | -0.10634174113205258 | 0.04350354452853408 | 44.14666091247955 |
| **GATCCAGATCTAACTTGGGGTGGCTTTGTCTTCTTCTTTTGCCCAATTCCACTAATTGTTTGGG** | Ins_k1_41_C;Ins_k1_42_C;Ins_k1 | -0.1387984040522528 | 0.0389214940744321 | 42.73681089682316 |
| **GATCCAGATCTAACTTGGGGTGGCTTTGTCTTCTTCTTTTGCCCACAATTCCACTAATTGTTTGGG** | Ins_k3_42_CCA;Ins_k3_43_CAC | 0.07610110724864616 | 0.03666354259864829 | 52.982419026935155 |
| **GATCCAGATCTAACTTGGGGTGGCTTTGTCTTCTTCTTTTGCCCACCAATTCCACTAATTGTTTGGG** | Ins_k4_41_CCCA;Ins_k4_42_CC | 0.5563241758720254 | 0.031151088018974052 | 85.64263320936658 |
| **GATCCAGATCTAACTTGGGGTGGCTTTGTCTTCTTCTTTTGCCCACTAATTGTTTGGG** | Del_k5_43-47;Del_k5_44-48 | -0.55251355955022885 | 0.04333246578589488 | 28.25712039602782 |
| **GATCCAGATCTAACTTGGGGTGGCTTTGTCTTCTTCTTTTGCCCAGAATTCCACTAATTGTTTGGG** | Ins_k3_43_CAG | 0.5092739623436011 | 0.0325695078281743 | 81.7064541691434 |
| **GATCCAGATCTAACTTGGGGTGGCTTTGTCTTCTTCTTTTGCCCATAATTCCACTAATTGTTTGGG** | Ins_k3_43_CAT | -0.8973347364458788 | 0.05007283510054126 | 20.015846770701412 |
| **GATCCAGATCTAACTTGGGGTGGCTTTGTCTTCTTCTTTTGCCCATTCCACTAATTGTTTGGG** | A44C | -0.46816682661921016 | 0.04216461175752864 | 30.743918679728736 |
| **GATCCAGATCTAACTTGGGGTGGCTTTGTCTTCTTCTTTTGCCCCAAATTCCACTAATTGTTTGGG** | Ins_k3_43_CCA | -2.6597283561936735 | 0.11645672972249578 | 3.435390763547927 |
| **GATCCAGATCTAACTTGGGGTGGCTTTGTCTTCTTCTTTTGCCCCAATTCCACTAATTGTTTGGG** | Ins_k2_41_CC;Ins_k2_42_CC;Ins | -2.2224601677288964 | 0.09665487066834655 | 5.319604013717951 |
| **GATCCAGATCTAACTTGGGGTGGCTTTGTCTTCTTCTTTTGCCCCACTAATTGTTTGGG** | Del_k4_44-47 | -3.1520634962294003 | 0.13115904796145753 | 2.0997022281810023 |
| **GATCCAGATCTAACTTGGGGTGGCTTTGTCTTCTTCTTTTGCCCCCAATTCCACTAATTGTTTGGG** | Ins_k3_41_CCC;Ins_k3_42_CCC | -1.4147437619200416 | 0.06300730751821822 | 11.930704329463824 |
| **GATCCAGATCTAACTTGGGGTGGCTTTGTCTTCTTCTTTTGCCCCGAATTCCACTAATTGTTTGGG** | Ins_k3_43_CCG | 0.370481095271186 | 0.034171949002016686 | 71.11797587295537 |
| **GATCCAGATCTAACTTGGGGTGGCTTTGTCTTCTTCTTTTGCCCCTAAATTCCACTAATTGTTTGGG** | Ins_k4_43_CCTA | -2.347750674971376 | 0.11081256471032186 | 4.693170467013847 |
| **GATCCAGATCTAACTTGGGGTGGCTTTGTCTTCTTCTTTTGCCCCTAATTCCACTAATTGTTTGGG** | Ins_k3_43_CCT | -2.1892458116590934 | 0.09230295151723707 | 5.499258269985613 |
| **GATCCAGATCTAACTTGGGGTGGCTTTGTCTTCTTCTTTTGCCCGAAATTCCACTAATTGTTTGGG** | Ins_k3_43_CGA | 0.666661974710463 | 0.03094108849457275 | 95.63329270751093 |
| **GATCCAGATCTAACTTGGGGTGGCTTTGTCTTCTTCTTTTGCCCGAATTCCACTAATTGTTTGGG** | Ins_k2_43_CG | 0.7316942327454825 | 0.028519663459558062 | 100 |
| **GATCCAGATCTAACTTGGGGTGGCTTTGTCTTCTTCTTTTGCCCGCAATTCCACTAATTGTTTGGG** | Ins_k3_42_CCG;Ins_k3_43_CGC | 0.514538412590364 | 0.03170523355516216 | 82.13772794558366 |
| **GATCCAGATCTAACTTGGGGTGGCTTTGTCTTCTTCTTTTGCCCGGAATTCCACTAATTGTTTGGG** | Ins_k3_43_CGG | 0.9716223950919027 | 0.030655243491126496 | 100 |
| **GATCCAGATCTAACTTGGGGTGGCTTTGTCTTCTTCTTTTGCCCGTAATTCCACTAATTGTTTGGG** | Ins_k3_43_CGT | 0.04228861414245444 | 0.038600537870976386 | 51.22089984965113 |
| **GATCCAGATCTAACTTGGGGTGGCTTTGTCTTCTTCTTTTGCCCTAAATTCCACTAATTGTTTGGG** | Ins_k3_43_CTA | -1.280313928235784 | 0.07192536040755036 | 13.647346492294604 |
| **GATCCAGATCTAACTTGGGGTGGCTTTGTCTTCTTCTTTTGCCCTAATTCCACTAATTGTTTGGG** | Ins_k2_43_CT | -1.2224343995980265 | 0.06510803129280723 | 14.460555541774125 |
| **GATCCAGATCTAACTTGGGGTGGCTTTGTCTTCTTCTTTTGCCCTAATTGTTTGGG** | Del_k7_44-50 | -1.9246344290517685 | 0.07231067033312737 | 7.165118542097918 |
| **GATCCAGATCTAACTTGGGGTGGCTTTGTCTTCTTCTTTTGCCCTCAATTCCACTAATTGTTTGGG** | Ins_k3_42_CCT;Ins_k3_43_CTC | -0.17600224610467258 | 0.04311825793193825 | 41.17605049602705 |
| **GATCCAGATCTAACTTGGGGTGGCTTTGTCTTCTTCTTTTGCCCTGAATTCCACTAATTGTTTGGG** | Ins_k3_43_CTG | 0.3511441640484967 | 0.03633719115758936 | 69.75598328240166 |
| **GATCCAGATCTAACTTGGGGTGGCTTTGTCTTCTTCTTTTGCCCTTAATTCCACTAATTGTTTGGG** | Ins_k3_43_CTT | -1.9588445884049845 | 0.078711923129636 | 6.924144083579614 |
| **GATCCAGATCTAACTTGGGGTGGCTTTGTCTTCTTCTTTTGCCG** | Del_k19_44-62 | 0.0359250832815472 | 0.02756793321165913 | 50.895988959398295 |
| **GATCCAGATCTAACTTGGGGTGGCTTTGTCTTCTTCTTTTGCCGAAAATTCCACTAATTGTTTGGG** | Ins_k3_43_GAA | 0.9734211920260116 | 0.02760743959532871 | 100 |
| **GATCCAGATCTAACTTGGGGTGGCTTTGTCTTCTTCTTTTGCCGAAATTCCACTAATTGTTTGGG** | Ins_k2_43_GA | 0.7635098277589296 | 0.030422327991360004 | 100 |
| **GATCCAGATCTAACTTGGGGTGGCTTTGTCTTCTTCTTTTGCCGAATTCCACTAATTGTTTGGG** | Ins_k1_43_G | 0.716019820323911 | 0.028965879881503644 | 100 |
| **GATCCAGATCTAACTTGGGGTGGCTTTGTCTTCTTCTTTTGCCGACAATTCCACTAATTGTTTGGG** | Ins_k3_42_CGA;Ins_k3_43_GAC | 0.675408851544157 | 0.030257937828110183 | 96.47345437877989 |
| **GATCCAGATCTAACTTGGGGTGGCTTTGTCTTCTTCTTTTGCCGAGAATTCCACTAATTGTTTGGG** | Ins_k3_43_GAG | 0.7394058344228034 | 0.03267962316337701 | 100 |
| **GATCCAGATCTAACTTGGGGTGGCTTTGTCTTCTTCTTTTGCCGATAATTCCACTAATTGTTTGGG** | Ins_k3_43_GAT | 0.3338870315796618 | 0.029754552920427814 | 68.56252251213635 |
| **GATCCAGATCTAACTTGGGGTGGCTTTGTCTTCTTCTTTTGCCGATTCCACTAATTGTTTGGG** | A44G | 0.19369686989615809 | 0.024533424079830935 | 59.594060001292505 |
| **GATCCAGATCTAACTTGGGGTGGCTTTGTCTTCTTCTTTTGCCGCAAATTCCACTAATTGTTTGGG** | Ins_k3_43_GCA | 0.7140234788728286 | 0.029643034436867298 | 100 |
| **GATCCAGATCTAACTTGGGGTGGCTTTGTCTTCTTCTTTTGCCGCAATTCCACTAATTGTTTGGG** | Ins_k2_42_CG;Ins_k2_43_GC | 0.7050782551380133 | 0.031003837141365457 | 99.37864881528303 |
| **GATCCAGATCTAACTTGGGGTGGCTTTGTCTTCTTCTTTTGCCGCAGAATTCCACTAATTGTTTGGG** | Ins_k4_43_GCAG | 0.8734746979494155 | 0.03090124105065337 | 100 |
| **GATCCAGATCTAACTTGGGGTGGCTTTGTCTTCTTCTTTTGCCGCCAATTCCACTAATTGTTTGGG** | Ins_k3_40_GCC;Ins_k3_41_CCG | 0.32084370352427705 | 0.03183796113260187 | 67.6740459803649 |
| **GATCCAGATCTAACTTGGGGTGGCTTTGTCTTCTTCTTTTGCCGCGAATTCCACTAATTGTTTGGG** | Ins_k3_43_GCG | 0.7564489311924274 | 0.029935034934355995 | 100 |
| **GATCCAGATCTAACTTGGGGTGGCTTTGTCTTCTTCTTTTGCCGCGTAATTCCACTAATTGTTTGGG** | Ins_k4_43_GCGT | 0.7393778501509571 | 0.03149014359103693 | 100 |
| **GATCCAGATCTAACTTGGGGTGGCTTTGTCTTCTTCTTTTGCCGCTAATTCCACTAATTGTTTGGG** | Ins_k3_43_GCT | 0.42407044213293155 | 0.030918162659449223 | 75.03310956091886 |
| **GATCCAGATCTAACTTGGGGTGGCTTTGTCTTCTTCTTTTGCCGG** | Del_k18_44-61 | -0.2501969776441984 | 0.02814587198326144 | 38.23158693913158 |
| **GATCCAGATCTAACTTGGGGTGGCTTTGTCTTCTTCTTTTGCCGGAAATTCCACTAATTGTTTGGG** | Ins_k3_43_GGA | 0.7989739049876099 | 0.028336481305982132 | 100 |
| **GATCCAGATCTAACTTGGGGTGGCTTTGTCTTCTTCTTTTGCCGGAATTCCACTAATTGTTTGGG** | Ins_k2_43_GG | 0.9364581172297088 | 0.03016161668417414 | 100 |
| **GATCCAGATCTAACTTGGGGTGGCTTTGTCTTCTTCTTTTGCCGGCAATTCCACTAATTGTTTGGG** | Ins_k3_42_CGG;Ins_k3_43_GGC | 0.6677024833176493 | 0.030713840708228028 | 95.73285175874916 |
| **GATCCAGATCTAACTTGGGGTGGCTTTGTCTTCTTCTTTTGCCGGG** | Del_k17_44-60 | -0.23868381414127404 | 0.029065256123039403 | 38.67429705732604 |
| **GATCCAGATCTAACTTGGGGTGGCTTTGTCTTCTTCTTTTGCCGGGAATTCCACTAATTGTTTGGG** | Ins_k3_43_GGG | 0.6891825695320135 | 0.027989802043098356 | 97.81144593886044 |

| | | | | |
|---|---|---|---|---|
| GATCCAGATCTAACTTGGGGTGGCTTTGTCTTCTTCTTTTGCCGGTAATTCCACTAATTGTTTGGG | Ins_k3_43_GGT | 0.2984798642581019 | 0.03457020856719631 | 66.1773923320549 |
| GATCCAGATCTAACTTGGGGTGGCTTTGTCTTCTTCTTTTGCCGTAAATTCCACTAATTGTTTGGG | Ins_k3_43_GTA | 0.40868817148993575 | 0.02903561079678739 | 73.88776157255154 |
| GATCCAGATCTAACTTGGGGTGGCTTTGTCTTCTTCTTTTGCCGTAATTCCACTAATTGTTTGGG | Ins_k2_43_GT | 0.6324307734654275 | 0.03126752816380432 | 92.41504668171706 |
| GATCCAGATCTAACTTGGGGTGGCTTTGTCTTCTTCTTTTGCCGTCAATTCCACTAATTGTTTGGG | Ins_k3_42_CGT;Ins_k3_43_GTC | 0.7683262993246827 | 0.027434573374353916 | 100 |
| GATCCAGATCTAACTTGGGGTGGCTTTGTCTTCTTCTTTTGCCGTGAATTCCACTAATTGTTTGGG | Ins_k3_43_GTG | 0.7844791522491881 | 0.028916802949798332 | 100 |
| GATCCAGATCTAACTTGGGGTGGCTTTGTCTTCTTCTTTTGCCGTTAATTCCACTAATTGTTTGGG | Ins_k3_43_GTT | 0.4607641496777348 | 0.03316229219024668 | 77.83748943586716 |
| GATCCAGATCTAACTTGGGGTGGCTTTGTCTTCTTCTTTTGCCGTTTGGG | Del_k13_44-56 | 0.2020312838287276 | 0.027649084681762026 | 60.09281710226023 |
| GATCCAGATCTAACTTGGGGTGGCTTTGTCTTCTTCTTTTGCCTAAAATTCCACTAATTGTTTGGG | Ins_k3_43_TAA | -1.4741290125322029 | 0.05521437680547052 | 11.242823593099763 |
| GATCCAGATCTAACTTGGGGTGGCTTTGTCTTCTTCTTTTGCCTAAATTCCACTAATTGTTTGGG | Ins_k2_43_TA | -0.7598159320778557 | 0.05093924745760102 | 22.966648600430403 |
| GATCCAGATCTAACTTGGGGTGGCTTTGTCTTCTTCTTTTGCCTAATTCCACTAATTGTTTGGG | Ins_k1_43_T | -0.587318595364769 | 0.04441792506768716 | 27.29054866320021 |
| GATCCAGATCTAACTTGGGGTGGCTTTGTCTTCTTCTTTTGCCTAATTGTTTGGG | Del_k8_43-50;Del_k8_44-51 | -0.8579077615957292 | 0.04777261597094264 | 20.82077472698687 |
| GATCCAGATCTAACTTGGGGTGGCTTTGTCTTCTTCTTTTGCCTACAATTCCACTAATTGTTTGGG | Ins_k3_42_CTA;Ins_k3_43_TAC | -0.4163670658180788 | 0.033208257719384376 | 32.378414090170196 |
| GATCCAGATCTAACTTGGGGTGGCTTTGTCTTCTTCTTTTGCCTACGAATTCCACTAATTGTTTGGG | Ins_k4_43_TACG | 0.7687302722615232 | 0.02970790382660531 | 100 |
| GATCCAGATCTAACTTGGGGTGGCTTTGTCTTCTTCTTTTGCCTAGAATTCCACTAATTGTTTGGG | Ins_k3_43_TAG | -0.39368427898388203 | 0.034926988341336465 | 33.12123958223691 |
| GATCCAGATCTAACTTGGGGTGGCTTTGTCTTCTTCTTTTGCCTATAATTCCACTAATTGTTTGGG | Ins_k3_43_TAT | -2.821936367024597 | 0.09842037460149661 | 2.9209901982220163 |
| GATCCAGATCTAACTTGGGGTGGCTTTGTCTTCTTCTTTTGCCTATTCCACTAATTGTTTGGG | A44T | -0.5491519369499341 | 0.0392161212886939 | 28.352270009512686 |
| GATCCAGATCTAACTTGGGGTGGCTTTGTCTTCTTCTTTTGCCTCAAATTCCACTAATTGTTTGGG | Ins_k3_43_TCA | -0.049353378428061534 | 0.035183120814598526 | 46.73557520566893 |
| GATCCAGATCTAACTTGGGGTGGCTTTGTCTTCTTCTTTTGCCTCAATTCCACTAATTGTTTGGG | Ins_k2_42_CT;Ins_k2_43_TC | 0.18071982040741463 | 0.04250542832625573 | 58.825701233148855 |
| GATCCAGATCTAACTTGGGGTGGCTTTGTCTTCTTCTTTTGCCTCCAATTCCACTAATTGTTTGGG | Ins_k3_41_CCT;Ins_k3_42_CTC;I | 0.29843110776788895 | 0.031494791972613796 | 66.17416583333036 |
| GATCCAGATCTAACTTGGGGTGGCTTTGTCTTCTTCTTTTGCCTCCACTAATTGTTTGGG | Del_k3_44-46 | -0.13171055572249807 | 0.0329772959301313 | 43.04079896859593 |
| GATCCAGATCTAACTTGGGGTGGCTTTGTCTTCTTCTTTTGCCTCGAATTCCACTAATTGTTTGGG | Ins_k3_43_TCG | 0.611660030186288 | 0.030157318297536408 | 90.5153151881576 |
| GATCCAGATCTAACTTGGGGTGGCTTTGTCTTCTTCTTTTGCCTCTAATTCCACTAATTGTTTGGG | Ins_k3_43_TCT | -1.05158392011025 | 0.059354673976341296 | 17.154750197224445 |
| GATCCAGATCTAACTTGGGGTGGCTTTGTCTTCTTCTTTTGCCTGAAATTCCACTAATTGTTTGGG | Ins_k3_43_TGA | 0.5586513818358252 | 0.03201356801333268 | 85.84217335157417 |
| GATCCAGATCTAACTTGGGGTGGCTTTGTCTTCTTCTTTTGCCTGAATTCCACTAATTGTTTGGG | Ins_k2_43_TG | 0.7957719742792493 | 0.030191178560605783 | 100 |
| GATCCAGATCTAACTTGGGGTGGCTTTGTCTTCTTCTTTTGCCTGCAATTCCACTAATTGTTTGGG | Ins_k3_42_CTG;Ins_k3_43_TGC | 0.6261860984554892 | 0.028981895683719863 | 91.8397429112434 |
| GATCCAGATCTAACTTGGGGTGGCTTTGTCTTCTTCTTTTGCCTGGAATTCCACTAATTGTTTGGG | Ins_k3_43_TGG | 0.7264706599352095 | 0.029957240198374684 | 100 |
| GATCCAGATCTAACTTGGGGTGGCTTTGTCTTCTTCTTTTGCCTGGG | Del_k16_44-59 | -0.09625435024966633 | 0.027342186864195945 | 44.59423919048842 |
| GATCCAGATCTAACTTGGGGTGGCTTTGTCTTCTTCTTTTGCCTGTAATTCCACTAATTGTTTGGG | Ins_k3_43_TGT | -0.11381866210648289 | 0.03681669187897789 | 43.81781074292846 |
| GATCCAGATCTAACTTGGGGTGGCTTTGTCTTCTTCTTTTGCCTGTTTGGG | Del_k12_44-55 | -0.01204764225897359 | 0.02901081459969484 | 48.5120098247286 |
| GATCCAGATCTAACTTGGGGTGGCTTTGTCTTCTTCTTTTGCCTTAAATTCCACTAATTGTTTGGG | Ins_k3_43_TTA | -1.3782435459096187 | 0.07317401981451833 | 12.37422263231397 |
| GATCCAGATCTAACTTGGGGTGGCTTTGTCTTCTTCTTTTGCCTTAATTCCACTAATTGTTTGGG | Ins_k2_43_TT | -1.2014471069789199 | 0.06037222215159743 | 14.767250543919301 |
| GATCCAGATCTAACTTGGGGTGGCTTTGTCTTCTTCTTTTGCCTTCAATTCCACTAATTGTTTGGG | Ins_k3_42_CTT;Ins_k3_43_TTC | 0.18424482297780098 | 0.03242642639012106 | 59.0334278845959 |
| GATCCAGATCTAACTTGGGGTGGCTTTGTCTTCTTCTTTTGCCTTCCACTAATTGTTTGGG | Del_k2_44-45 | -0.2378856664004979 | 0.03369742519899957 | 38.70517718195998 |
| GATCCAGATCTAACTTGGGGTGGCTTTGTCTTCTTCTTTTGCCTTCTAATTCCACTAATTGTTTGGG | Ins_k4_43_TTCT | -0.935775170788546 | 0.063723180751543 | 19.261029621704274 |
| GATCCAGATCTAACTTGGGGTGGCTTTGTCTTCTTCTTTTGCCTTGAATTCCACTAATTGTTTGGG | Ins_k3_43_TTG | 0.17384079079591586 | 0.034315356763462875 | 58.42242615333869 |
| GATCCAGATCTAACTTGGGGTGGCTTTGTCTTCTTCTTTTGCCTTGGG | Del_k15_44-58 | -0.5042518157457383 | 0.028314472803167137 | 29.654302337679518 |
| GATCCAGATCTAACTTGGGGTGGCTTTGTCTTCTTCTTTTGCCTTGTTTGGG | Del_k11_44-54 | -0.7441176660489618 | 0.1746258215453797 | 23.330029925834204 |
| GATCCAGATCTAACTTGGGGTGGCTTTGTCTTCTTCTTTTGCCTTTAATTCCACTAATTGTTTGGG | Ins_k3_43_TTT | -2.2525163367853125 | 0.1212235932144365 | 5.162095997154751 |
| GATCCAGATCTAACTTGGGGTGGCTTTGTCTTCTTCTTTTGCCTTTGGG | Del_k14_44-57 | -0.46287012979124986 | 0.03060510071951709 | 30.90719191898394 |
| GATCCAGATCTAACTTGGGGTGGCTTTGTCTTCTTCTTTTGCG | Del_k20_43-62 | -0.4104652331373513 | 0.02816056096990196 | 32.57007108003071 |
| GATCCAGATCTAACTTGGGGTGGCTTTGTCTTCTTCTTTTGCGAACAATTCCACTAATTGTTTGGG | Ins_k3_42_GAA | 0.6457266783556088 | 0.029490668287176294 | 93.65199329272242 |
| GATCCAGATCTAACTTGGGGTGGCTTTGTCTTCTTCTTTTGCGAATTCCACTAATTGTTTGGG | C43G | 0.19621498598331033 | 0.030497084468108948 | 59.744313861416146 |
| GATCCAGATCTAACTTGGGGTGGCTTTGTCTTCTTCTTTTGCGACAATTCCACTAATTGTTTGGG | Ins_k2_42_GA | 0.7282332200788568 | 0.03228075489331552 | 100 |
| GATCCAGATCTAACTTGGGGTGGCTTTGTCTTCTTCTTTTGCGACCAATTCCACTAATTGTTTGGG | Ins_k3_41_CGA;Ins_k3_42_GAC | 0.6626316019601717 | 0.028861872962745047 | 95.2486305771472 |
| GATCCAGATCTAACTTGGGGTGGCTTTGTCTTCTTCTTTTGCGAGCAATTCCACTAATTGTTTGGG | Ins_k3_42_GAG | 0.6561289791783487 | 0.030835789687294717 | 94.63127405567168 |
| GATCCAGATCTAACTTGGGGTGGCTTTGTCTTCTTCTTTTGCGATCAATTCCACTAATTGTTTGGG | Ins_k3_42_GAT | 0.5395561689080997 | 0.03039441276818246 | 84.21854982618238 |
| GATCCAGATCTAACTTGGGGTGGCTTTGTCTTCTTCTTTTGCGCAATTCCACTAATTGTTTGGG | Ins_k1_42_G | 0.22485984547251392 | 0.03385304552774887 | 61.48042800081724 |
| GATCCAGATCTAACTTGGGGTGGCTTTGTCTTCTTCTTTTGCGCACAATTCCACTAATTGTTTGGG | Ins_k3_42_GCA | 0.2564081703229595 | 0.031881883343330995 | 63.45095245275334 |
| GATCCAGATCTAACTTGGGGTGGCTTTGTCTTCTTCTTTTGCGCCAATTCCACTAATTGTTTGGG | Ins_k2_40_GC;Ins_k2_41_CG;Ins | 0.5823353770617437 | 0.03164834056248565 | 87.89952597161675 |
| GATCCAGATCTAACTTGGGGTGGCTTTGTCTTCTTCTTTTGCGCCCAATTCCACTAATTGTTTGGG | Ins_k3_41_CGC;Ins_k3_42_GCC | 0.42477293442268005 | 0.031928223931786896 | 75.08583826044524 |
| GATCCAGATCTAACTTGGGGTGGCTTTGTCTTCTTCTTTTGCGCGCAATTCCACTAATTGTTTGGG | Ins_k3_42_GCG | 0.327509385059943 | 0.03192283220650512 | 68.12664638839024 |
| GATCCAGATCTAACTTGGGGTGGCTTTGTCTTCTTCTTTTGCGCTCAATTCCACTAATTGTTTGGG | Ins_k3_42_GCT | -0.03310640601805681 | 0.034449635984859776 | 47.50108860521071 |
| GATCCAGATCTAACTTGGGGTGGCTTTGTCTTCTTCTTTTGCGG | Del_k19_43-61 | -0.2983513343061889 | 0.02589635883773211 | 36.43419295080866 |
| GATCCAGATCTAACTTGGGGTGGCTTTGTCTTCTTCTTTTGCGGACAATTCCACTAATTGTTTGGG | Ins_k3_42_GGA | 0.5142718430957798 | 0.03418620687561224 | 82.11583545102414 |
| GATCCAGATCTAACTTGGGGTGGCTTTGTCTTCTTCTTTTGCGGCAATTCCACTAATTGTTTGGG | Ins_k2_42_GG | 0.7353859108222223 | 0.032511862852353815 | 100 |
| GATCCAGATCTAACTTGGGGTGGCTTTGTCTTCTTCTTTTGCGGCCAATTCCACTAATTGTTTGGG | Ins_k3_40_GCG;Ins_k3_41_CGG | 0.713925339940453 | 0.027329720721480343 | 100 |
| GATCCAGATCTAACTTGGGGTGGCTTTGTCTTCTTCTTTTGCGGCTCAATTCCACTAATTGTTTGGG | Ins_k4_42_GGCT | 1.1137806272528918 | 0.03351729722470549 | 100 |
| GATCCAGATCTAACTTGGGGTGGCTTTGTCTTCTTCTTTTGCGGG | Del_k18_43-60 | -0.3982782791301633 | 0.027137721892636125 | 32.969429577337145 |
| GATCCAGATCTAACTTGGGGTGGCTTTGTCTTCTTCTTTTGCGGGCAATTCCACTAATTGTTTGGG | Ins_k3_42_GGG | 0.3438784228180929 | 0.14187014450277047 | 69.25099115125799 |
| GATCCAGATCTAACTTGGGGTGGCTTTGTCTTCTTCTTTTGCGGTCAATTCCACTAATTGTTTGGG | Ins_k3_42_GGT | 0.5544942012355303 | 0.034348771245998375 | 85.48605267568988 |
| GATCCAGATCTAACTTGGGGTGGCTTTGTCTTCTTCTTTTGCGTACAATTCCACTAATTGTTTGGG | Ins_k3_42_GTA | -0.7256879978987999 | 0.038450035788962375 | 23.763981141091474 |
| GATCCAGATCTAACTTGGGGTGGCTTTGTCTTCTTCTTTTGCGTCAATTCCACTAATTGTTTGGG | Ins_k2_42_GT | 0.3360616816818952 | 0.032279321244433334 | 68.71178424592622 |
| GATCCAGATCTAACTTGGGGTGGCTTTGTCTTCTTCTTTTGCGTCCAATTCCACTAATTGTTTGGG | Ins_k3_41_CGT;Ins_k3_42_GTC | 0.4393066057552102 | 0.02964685825779797 | 76.18507981406606 |
| GATCCAGATCTAACTTGGGGTGGCTTTGTCTTCTTCTTTTGCGTGCAATTCCACTAATTGTTTGGG | Ins_k3_42_GTG | 0.5232558442876736 | 0.0315725570371057 | 82.85688803892586 |
| GATCCAGATCTAACTTGGGGTGGCTTTGTCTTCTTCTTTTGCGTTCAATTCCACTAATTGTTTGGG | Ins_k3_42_GTT | -0.2295698487845409 | 0.039540673138694284 | 39.028384379697385 |
| GATCCAGATCTAACTTGGGGTGGCTTTGTCTTCTTCTTTTGCGTTTGGG | Del_k14_43-56 | -0.18824286636521986 | 0.03240743007513292 | 40.67510231118267 |
| GATCCAGATCTAACTTGGGGTGGCTTTGTCTTCTTCTTTTGCTAACAATTCCACTAATTGTTTGGG | Ins_k3_42_TAA | -1.9151293950543717 | 0.08514918452918536 | 7.233547934093771 |
| GATCCAGATCTAACTTGGGGTGGCTTTGTCTTCTTCTTTTGCTAACCAATTCCACTAATTGTTTGGG | Ins_k4_41_CTAA;Ins_k4_42_TAA | -1.6221013905275716 | 0.07566997557642197 | 9.696428717593673 |
| GATCCAGATCTAACTTGGGGTGGCTTTGTCTTCTTCTTTTGCTAATTCCACTAATTGTTTGGG | C43T | -1.0997521575185027 | 0.043477338093621025 | 16.348021441722942 |
| GATCCAGATCTAACTTGGGGTGGCTTTGTCTTCTTCTTTTGCTAATTGTTTGGG | Del_k9_42-50;Del_k9_43-51 | -1.624764840546914 | 0.05993573010705339 | 9.670637126891677 |
| GATCCAGATCTAACTTGGGGTGGCTTTGTCTTCTTCTTTTGCTACAATTCCACTAATTGTTTGGG | Ins_k2_42_TA | -0.546164284363542 | 0.03512224147414422 | 28.437103405591923 |
| GATCCAGATCTAACTTGGGGTGGCTTTGTCTTCTTCTTTTGCTACCAATTCCACTAATTGTTTGGG | Ins_k3_41_CTA;Ins_k3_42_TAC | 0.7769232441022753 | 0.3183090838725596 | 100 |
| GATCCAGATCTAACTTGGGGTGGCTTTGTCTTCTTCTTTTGCTAGCAATTCCACTAATTGTTTGGG | Ins_k3_42_TAG | -2.8645561375385866 | 0.13380835367706762 | 2.7991138844700503 |
| GATCCAGATCTAACTTGGGGTGGCTTTGTCTTCTTCTTTTGCTATCAATTCCACTAATTGTTTGGG | Ins_k3_42_TAT | -2.331057448993045 | 0.09864085555578415 | 4.772172184366763 |
| GATCCAGATCTAACTTGGGGTGGCTTTGTCTTCTTCTTTTGCTCAATTCCACTAATTGTTTGGG | Ins_k1_42_T | -0.5978252440686707 | 0.046014027392356785 | 27.0053174911974 |

| | | | | |
|---|---|---|---|---|
| GATCCAGATCTAACTTGGGGTGGCTTTGTCTTCTTCTTTTGCTCACAATTCCACTAATTGTTTGGG | Ins_k3_42_TCA | -0.517526590002876 | 0.045970690641419946 | 29.26324948017922 |
| GATCCAGATCTAACTTGGGGTGGCTTTGTCTTCTTCTTTTGCTCACCAATTCCACTAATTGTTTGGG | Ins_k4_41_CTCA;Ins_k4_42_TCA | 0.4861831753899377 | 0.035015809013006945 | 79.84140345480144 |
| GATCCAGATCTAACTTGGGGTGGCTTTGTCTTCTTCTTTTGCTCCAATTCCACTAATTGTTTGGG | Ins_k2_41_CT;Ins_k2_42_TC | 0.39888909817221085 | 0.03320954448954785 | 73.16726584014086 |
| GATCCAGATCTAACTTGGGGTGGCTTTGTCTTCTTCTTTTGCTCCACTAATTGTTTGGG | Del_k4_43-46 | -0.029721711242168714 | 0.03689073322160889 | 47.6621376889076 |
| GATCCAGATCTAACTTGGGGTGGCTTTGTCTTCTTCTTTTGCTCCCAATTCCACTAATTGTTTGGG | Ins_k3_41_CTC;Ins_k3_42_TCC | -0.4084006084312777 | 0.04166878980497366 | 32.637385519076446 |
| GATCCAGATCTAACTTGGGGTGGCTTTGTCTTCTTCTTTTGCTCGCAATTCCACTAATTGTTTGGG | Ins_k3_42_TCG | -0.043482366305442666 | 0.037234710043132106 | 47.01076737207033 |
| GATCCAGATCTAACTTGGGGTGGCTTTGTCTTCTTCTTTTGCTCTACAATTCCACTAATTGTTTGGG | Ins_k4_42_TCTA | 0.10916090922116073 | 0.03240363779986534 | 54.763282609996736 |
| GATCCAGATCTAACTTGGGGTGGCTTTGTCTTCTTCTTTTGCTCTCAATTCCACTAATTGTTTGGG | Ins_k3_42_TCT | -1.567022022416819 | 0.07730923451224628 | 10.2454838898748 |
| GATCCAGATCTAACTTGGGGTGGCTTTGTCTTCTTCTTTTGCTGACAATTCCACTAATTGTTTGGG | Ins_k3_42_TGA | 0.3925840328147445 | 0.03424568762533103 | 72.7073927354584 |
| GATCCAGATCTAACTTGGGGTGGCTTTGTCTTCTTCTTTTGCTGCAATTCCACTAATTGTTTGGG | Ins_k2_42_TG | 0.6581153624781395 | 0.029325612019347008 | 94.81943485596035 |
| GATCCAGATCTAACTTGGGGTGGCTTTGTCTTCTTCTTTTGCTGCCAATTCCACTAATTGTTTGGG | Ins_k3_39_TGC;Ins_k3_40_GCT; | 0.6019460136666872 | 0.029861877432041106 | 89.6403047347848 |
| GATCCAGATCTAACTTGGGGTGGCTTTGTCTTCTTCTTTTGCTGGCAATTCCACTAATTGTTTGGG | Ins_k3_42_TGG | 0.48497355369144923 | 0.032056784519187705 | 79.7448839485581 |
| GATCCAGATCTAACTTGGGGTGGCTTTGTCTTCTTCTTTTGCTGGG | Del_k17_43-59 | -0.2587018065203709 | 0.02846681564542822 | 37.90781260879125 |
| GATCCAGATCTAACTTGGGGTGGCTTTGTCTTCTTCTTTTGCTGGGCAATTCCACTAATTGTTTGGG | Ins_k4_42_TGGG | 0.4227776179972573 | 0.03559912261025072 | 74.93616762384417 |
| GATCCAGATCTAACTTGGGGTGGCTTTGTCTTCTTCTTTTGCTGTCAATTCCACTAATTGTTTGGG | Ins_k3_42_TGT | 0.2100398897927972 | 0.034340944011539504 | 60.576009060132485 |
| GATCCAGATCTAACTTGGGGTGGCTTTGTCTTCTTCTTTTGCTGTTTGGG | Del_k13_43-55 | 0.07018293146026056 | 0.028888412966242092 | 52.66978577919123 |
| GATCCAGATCTAACTTGGGGTGGCTTTGTCTTCTTCTTTTGCTTACAATTCCACTAATTGTTTGGG | Ins_k3_42_TTA | -1.8765087070295572 | 0.06242430372152394 | 7.518377282470741 |
| GATCCAGATCTAACTTGGGGTGGCTTTGTCTTCTTCTTTTGCTTCAATTCCACTAATTGTTTGGG | Ins_k2_42_TT | 0.14397206425923303 | 0.03635443120072566 | 56.703225659834615 |
| GATCCAGATCTAACTTGGGGTGGCTTTGTCTTCTTCTTTTGCTTCCAATTCCACTAATTGTTTGGG | Ins_k3_41_CTT;Ins_k3_42_TTC | -0.4772925064896226 | 0.04007630096178467 | 30.464635781232868 |
| GATCCAGATCTAACTTGGGGTGGCTTTGTCTTCTTCTTTTGCTTCCACTAATTGTTTGGG | Del_k3_43-45 | -0.6005702829417233 | 0.04091527351157932 | 26.931288497622987 |
| GATCCAGATCTAACTTGGGGTGGCTTTGTCTTCTTCTTTTGCTTGCAATTCCACTAATTGTTTGGG | Ins_k3_42_TTG | -0.5233216391264144 | 0.04247806324167306 | 29.09415793205191 |
| GATCCAGATCTAACTTGGGGTGGCTTTGTCTTCTTCTTTTGCTTGGG | Del_k16_43-58 | -0.5452142758936968 | 0.029003159877789706 | 28.464131731221443 |
| GATCCAGATCTAACTTGGGGTGGCTTTGTCTTCTTCTTTTGCTTGTTTGGG | Del_k12_43-54 | -0.7874684210612233 | 0.038538005182238905 | 22.34026406255434 |
| GATCCAGATCTAACTTGGGGTGGCTTTGTCTTCTTCTTTTGCTTTCAATTCCACTAATTGTTTGGG | Ins_k3_42_TTT | -2.7144814499945062 | 0.12392762327950128 | 3.252349261181344 |
| GATCCAGATCTAACTTGGGGTGGCTTTGTCTTCTTCTTTTGCTTTGGG | Del_k15_43-57 | -0.8650269478486952 | 0.03585344450902567 | 20.673074131621387 |
| GATCCAGATCTAACTTGGGGTGGCTTTGTCTTCTTCTTTTGG | Del_k21_41-61;Del_k21_42-62 | -0.6975219629940713 | 0.03035835010429945 | 24.442833676835075 |
| GATCCAGATCTAACTTGGGGTGGCTTTGTCTTCTTCTTTTGGAACCAATTCCACTAATTGTTTGGG | Ins_k3_41_GAA | 0.7609707465110254 | 0.02835337879843167 | 100 |
| GATCCAGATCTAACTTGGGGTGGCTTTGTCTTCTTCTTTTGGACCAATTCCACTAATTGTTTGGG | Ins_k2_41_GA | 0.9200042045348866 | 0.02875382652528517 | 100 |
| GATCCAGATCTAACTTGGGGTGGCTTTGTCTTCTTCTTTTGGACCCAATTCCACTAATTGTTTGGG | Ins_k3_41_GAC | 0.7431012365227705 | 0.02889275046713013 | 100 |
| GATCCAGATCTAACTTGGGGTGGCTTTGTCTTCTTCTTTTGGACGCCAATTCCACTAATTGTTTGGG | Ins_k4_40_GGAC;Ins_k4_41_GA | 0.9947155543561568 | 0.02908636453065328 | 100 |
| GATCCAGATCTAACTTGGGGTGGCTTTGTCTTCTTCTTTTGGAGCCAATTCCACTAATTGTTTGGG | Ins_k3_40_GGA;Ins_k3_41_GAG | 0.8114203020984299 | 0.030024125838941724 | 100 |
| GATCCAGATCTAACTTGGGGTGGCTTTGTCTTCTTCTTTTGGATCCAATTCCACTAATTGTTTGGG | Ins_k3_41_GAT | 0.7741936503677098 | 0.029888450589338273 | 100 |
| GATCCAGATCTAACTTGGGGTGGCTTTGTCTTCTTCTTTTGGCAATTCCACTAATTGTTTGGG | C42G | -0.3842706310417462 | 0.0347636926936338 | 33.43450343551937 |
| GATCCAGATCTAACTTGGGGTGGCTTTGTCTTCTTCTTTTGGCACCAATTCCACTAATTGTTTGGG | Ins_k3_41_GCA | 0.57239940013938 | 0.033672723732520594 | 87.03048285582726 |
| GATCCAGATCTAACTTGGGGTGGCTTTGTCTTCTTCTTTTGGCCAATTCCACTAATTGTTTGGG | Ins_k1_40_G;Ins_k1_41_G | 0.4367319149410316 | 0.03186044136388102 | 75.98917908900933 |
| GATCCAGATCTAACTTGGGGTGGCTTTGTCTTCTTCTTTTGGCCCAATTCCACTAATTGTTTGGG | Ins_k2_41_GC | 0.5560820376914459 | 0.03469152579831241 | 85.62189836843086 |
| GATCCAGATCTAACTTGGGGTGGCTTTGTCTTCTTCTTTTGGCCCCAATTCCACTAATTGTTTGGG | Ins_k3_41_GCC | -0.4420121627157666 | 0.04240265724589178 | 31.558623259967618 |
| GATCCAGATCTAACTTGGGGTGGCTTTGTCTTCTTCTTTTGGCCGCCAATTCCACTAATTGTTTGGG | Ins_k4_40_GGCC;Ins_k4_41_GC | 0.6725435564951963 | 0.033781045373419645 | 96.1974251090902 |
| GATCCAGATCTAACTTGGGGTGGCTTTGTCTTCTTCTTTTGGCGCCAATTCCACTAATTGTTTGGG | Ins_k3_40_GGC;Ins_k3_41_GCG | 0.8169186113385398 | 0.030081391866729635 | 100 |
| GATCCAGATCTAACTTGGGGTGGCTTTGTCTTCTTCTTTTGGCTCCAATTCCACTAATTGTTTGGG | Ins_k3_41_GCT | 0.6654862534928006 | 0.029453141920040796 | 95.520920688188 |
| GATCCAGATCTAACTTGGGGTGGCTTTGTCTTCTTCTTTTGGG | Del_k20_38-57;Del_k20_39-58;D | -0.6204991144415448 | 0.028308932711914706 | 26.3998920296766 |
| GATCCAGATCTAACTTGGGGTGGCTTTGTCTTCTTCTTTTGGGACCAATTCCACTAATTGTTTGGG | Ins_k3_41_GGA | 0.7414805491139631 | 0.029283526284716872 | 100 |
| GATCCAGATCTAACTTGGGGTGGCTTTGTCTTCTTCTTTTGGGCCAATTCCACTAATTGTTTGGG | Ins_k2_40_GG;Ins_k2_41_GG | 0.42032990501498246 | 0.03684948053813794 | 74.75296969295236 |
| GATCCAGATCTAACTTGGGGTGGCTTTGTCTTCTTCTTTTGGGCCCAATTCCACTAATTGTTTGGG | Ins_k3_41_GGC | 0.3156340510454487 | 0.03432311730662522 | 67.32240447928936 |
| GATCCAGATCTAACTTGGGGTGGCTTTGTCTTCTTCTTTTGGGCTCCAATTCCACTAATTGTTTGGG | Ins_k4_41_GGCT | 0.36461332467219554 | 0.034688328211582915 | 70.7018938358737 |
| GATCCAGATCTAACTTGGGGTGGCTTTGTCTTCTTCTTTTGGGG | Del_k19_42-60 | -0.7147920743931311 | 0.029580651916926232 | 24.024327442776457 |
| GATCCAGATCTAACTTGGGGTGGCTTTGTCTTCTTCTTTTGGGGCCAATTCCACTAATTGTTTGGG | Ins_k3_40_GGG;Ins_k3_41_GGG | 0.03383618391391341 | 0.037612085244660656 | 50.78978332530589 |
| GATCCAGATCTAACTTGGGGTGGCTTTGTCTTCTTCTTTTGGGTCCAATTCCACTAATTGTTTGGG | Ins_k3_41_GGT | -0.07042307694722494 | 0.03737848334504345 | 45.761171964378335 |
| GATCCAGATCTAACTTGGGGTGGCTTTGTCTTCTTCTTTTGGTACCAATTCCACTAATTGTTTGGG | Ins_k3_41_GTA | 0.19585485041424122 | 0.034492064071744676 | 59.722801682827864 |
| GATCCAGATCTAACTTGGGGTGGCTTTGTCTTCTTCTTTTGGTCCAATTCCACTAATTGTTTGGG | Ins_k2_41_GT | 0.466387156849827 | 0.03643318898469613 | 78.27640304794572 |
| GATCCAGATCTAACTTGGGGTGGCTTTGTCTTCTTCTTTTGGTCCCAATTCCACTAATTGTTTGGG | Ins_k3_41_GTC | 0.13973990245146173 | 0.03202196061259061 | 56.46375552942252 |
| GATCCAGATCTAACTTGGGGTGGCTTTGTCTTCTTCTTTTGGTCTCCAATTCCACTAATTGTTTGGG | Ins_k4_41_GTCT | 0.8220973079988583 | 0.031487106667469754 | 100 |
| GATCCAGATCTAACTTGGGGTGGCTTTGTCTTCTTCTTTTGGTGCCAATTCCACTAATTGTTTGGG | Ins_k3_39_TGG;Ins_k3_40_GGT; | 0.6685690662951613 | 0.032514967942622194 | 95.81584817491874 |
| GATCCAGATCTAACTTGGGGTGGCTTTGTCTTCTTCTTTTGGTTCCAATTCCACTAATTGTTTGGG | Ins_k3_41_GTT | 0.13612686299979182 | 0.03503250725550307 | 56.26011785011459 |
| GATCCAGATCTAACTTGGGGTGGCTTTGTCTTCTTCTTTTGGTTTGGG | Del_k15_42-56 | -1.0875868077566067 | 0.04334481064290036 | 16.54811547970588 |
| GATCCAGATCTAACTTGGGGTGGCTTTGTCTTCTTCTTTTGTAACCAATTCCACTAATTGTTTGGG | Ins_k3_41_TAA | -0.6737784952206122 | 0.053325308709684015 | 25.030136016798153 |
| GATCCAGATCTAACTTGGGGTGGCTTTGTCTTCTTCTTTTGTAATTGTTTGGG | Del_k10_42-51 | -1.953571271159249 | 0.0795265689323944 | 6.960753734302429 |
| GATCCAGATCTAACTTGGGGTGGCTTTGTCTTCTTCTTTTGTACCAATTCCACTAATTGTTTGGG | Ins_k2_41_TA | -0.12122063488967877 | 0.041194245161465914 | 43.49466991557897 |
| GATCCAGATCTAACTTGGGGTGGCTTTGTCTTCTTCTTTTGTACCCAATTCCACTAATTGTTTGGG | Ins_k3_41_TAC | -0.6233653482405823 | 0.04713562755055782 | 26.324332104784414 |
| GATCCAGATCTAACTTGGGGTGGCTTTGTCTTCTTCTTTTGTACCCCAATTCCACTAATTGTTTGGG | Ins_k4_41_TACC | -2.6991567231628144 | 0.13972514365259808 | 3.3025744861624906 |
| GATCCAGATCTAACTTGGGGTGGCTTTGTCTTCTTCTTTTGTAGCCAATTCCACTAATTGTTTGGG | Ins_k3_40_GTA;Ins_k3_41_TAG | -3.6285959395098164 | 0.1915988143972291 | 1.3037739474449312 |
| GATCCAGATCTAACTTGGGGTGGCTTTGTCTTCTTCTTTTGTATCCAATTCCACTAATTGTTTGGG | Ins_k3_41_TAT | -0.9108204943923814 | 0.049242329789076236 | 19.74772984943448 |
| GATCCAGATCTAACTTGGGGTGGCTTTGTCTTCTTCTTTTGTCAATTCCACTAATTGTTTGGG | C42T | -0.84482149986849 | 0.03430728986408851 | 21.095031417981527 |
| GATCCAGATCTAACTTGGGGTGGCTTTGTCTTCTTCTTTTGTCACCAATTCCACTAATTGTTTGGG | Ins_k3_41_TCA | 0.32421513615460584 | 0.03360846299474783 | 67.9025895102872 |
| GATCCAGATCTAACTTGGGGTGGCTTTGTCTTCTTCTTTTGTCCAATTCCACTAATTGTTTGGG | Ins_k1_41_T | -0.19432475529960036 | 0.042055307102622744 | 40.428471607032506 |
| GATCCAGATCTAACTTGGGGTGGCTTTGTCTTCTTCTTTTGTCCACTAATTGTTTGGG | Del_k5_42-46 | -0.37676213648895285 | 0.03463324248720362 | 33.686491062434285 |
| GATCCAGATCTAACTTGGGGTGGCTTTGTCTTCTTCTTTTGTCCCAATTCCACTAATTGTTTGGG | Ins_k2_41_TC | -0.42772967965274417 | 0.04117063673909953 | 32.01259295233701 |
| GATCCAGATCTAACTTGGGGTGGCTTTGTCTTCTTCTTTTGTCCCCAATTCCACTAATTGTTTGGG | Ins_k3_41_TCC | -1.2130053074628733 | 0.05727212202717874 | 14.597550305437148 |
| GATCCAGATCTAACTTGGGGTGGCTTTGTCTTCTTCTTTTGTCGCCAATTCCACTAATTGTTTGGG | Ins_k3_40_GTC;Ins_k3_41_TCG | 0.5523375116942499 | 0.027845668007845742 | 85.30188446816766 |
| GATCCAGATCTAACTTGGGGTGGCTTTGTCTTCTTCTTTTGTCTCCAATTCCACTAATTGTTTGGG | Ins_k3_41_TCT | 0.12823897627685277 | 0.03293307183733623 | 55.818090038014844 |
| GATCCAGATCTAACTTGGGGTGGCTTTGTCTTCTTCTTTTGTGACCAATTCCACTAATTGTTTGGG | Ins_k3_41_TGA | 0.6138296430901973 | 0.029279206064642183 | 90.71191157590256 |
| GATCCAGATCTAACTTGGGGTGGCTTTGTCTTCTTCTTTTGTGCCAATTCCACTAATTGTTTGGG | Ins_k2_39_TG;Ins_k2_40_GT;Ins_ | 0.2523083496260413 | 0.03478189830136414 | 63.19134745525069 |
| GATCCAGATCTAACTTGGGGTGGCTTTGTCTTCTTCTTTTGTGCCCAATTCCACTAATTGTTTGGG | Ins_k3_41_TGC | 0.09456244654793922 | 0.03607401678534376 | 53.9696300216912 |
| GATCCAGATCTAACTTGGGGTGGCTTTGTCTTCTTCTTTTGTGGCCAATTCCACTAATTGTTTGGG | Ins_k3_40_GTG;Ins_k3_41_TGG | 0.280392606321341 | 0.03509311449042555 | 64.99118472751744 |
| GATCCAGATCTAACTTGGGGTGGCTTTGTCTTCTTCTTTTGTGGG | Del_k18_42-59 | -0.36358784974555314 | 0.029417516357796162 | 34.1332227796606 |

| | | | | |
|---|---|---|---|---|
| GATCCAGATCTAACTTGGGGTGGCTTTGTCTTCTTCTTTTGTGGTCCAATTCCACTAATTGTTTGGG | Ins_k4_41_TGGT | 0.31749566628234527 | 0.03809308425118759 | 67.44784962231867 |
| GATCCAGATCTAACTTGGGGTGGCTTTGTCTTCTTCTTTTGTGTCCAATTCCACTAATTGTTTGGG | Ins_k3_41_TGT | -0.13828499713623071 | 0.03683622297728004 | 42.758757904497 |
| GATCCAGATCTAACTTGGGGTGGCTTTGTCTTCTTCTTTTGTGTTTGGG | Del_k14_42-55 | -0.71308073223008 | 0.03684419029598079 | 24.065476487272896 |
| GATCCAGATCTAACTTGGGGTGGCTTTGTCTTCTTCTTTTGTTACCAATTCCACTAATTGTTTGGG | Ins_k3_41_TTA | -0.603074168863531 | 0.04733103052848322 | 26.863939975192707 |
| GATCCAGATCTAACTTGGGGTGGCTTTGTCTTCTTCTTTTGTTCACCAATTCCACTAATTGTTTGGG | Ins_k4_41_TTCA | 0.45164991791152054 | 0.03457222366738182 | 77.13128366723019 |
| GATCCAGATCTAACTTGGGGTGGCTTTGTCTTCTTCTTTTGTTCCAATTCCACTAATTGTTTGGG | Ins_k2_41_TT | -0.2918207063652873 | 0.040655959473276174 | 36.67290974616346 |
| GATCCAGATCTAACTTGGGGTGGCTTTGTCTTCTTCTTTTGTTCCACTAATTGTTTGGG | Del_k4_42-45 | -0.35656139277690946 | 0.03956506285608095 4 | 34.37390297502134 |
| GATCCAGATCTAACTTGGGGTGGCTTTGTCTTCTTCTTTTGTTCCCAATTCCACTAATTGTTTGGG | Ins_k3_41_TTC | -0.8167833838811582 | 0.05016777042536163 | 21.69486617847174 |
| GATCCAGATCTAACTTGGGGTGGCTTTGTCTTCTTCTTTTGTTGCCAATTCCACTAATTGTTTGGG | Ins_k3_38_TTG;Ins_k3_39_TGT;I | -0.09810699118376798 | 0.03637151292919365 | 44.51169856022272 |
| GATCCAGATCTAACTTGGGGTGGCTTTGTCTTCTTCTTTTGTTGGG | Del_k17_42-58 | -0.23477435673361735 | 0.029766722542670206 | 38.82578850617842 |
| GATCCAGATCTAACTTGGGGTGGCTTTGTCTTCTTCTTTTGTTGTTTGGG | Del_k13_42-54 | -0.8640319533424448 | 0.04093059174211268 | 20.693653963521633 |
| GATCCAGATCTAACTTGGGGTGGCTTTGTCTTCTTCTTTTGTTTCCAATTCCACTAATTGTTTGGG | Ins_k3_41_TTT | -1.9405063220433465 | 0.0803885671823782 | 7.052292298915644 |
| GATCCAGATCTAACTTGGGGTGGCTTTGTCTTCTTCTTTTGTTTGGG | Del_k16_39-54;Del_k16_40-55;D | -0.42051612250392445 | 0.031060666288417187 | 32.24435252155485 |
| GATCCAGATCTAACTTGGGGTGGCTTTGTCTTCTTCTTTTTAAGCCAATTCCACTAATTGTTTGGG | Ins_k3_40_TAA | -1.0024879043833046 | 0.06200497897448452 | 18.017997697082127 |
| GATCCAGATCTAACTTGGGGTGGCTTTGTCTTCTTCTTTTTAATTGTTTGGG | Del_k11_41-51 | -1.6675570712336325 | 0.0649379055283386 | 9.265538347620375 |
| GATCCAGATCTAACTTGGGGTGGCTTTGTCTTCTTCTTTTTACGCCAATTCCACTAATTGTTTGGG | Ins_k3_40_TAC | 0.41104278062416144 | 0.03199403026311205 | 74.06194335539156 |
| GATCCAGATCTAACTTGGGGTGGCTTTGTCTTCTTCTTTTTAGCCAATTCCACTAATTGTTTGGG | Ins_k2_40_TA | -4.010469475324493 | 0.24567456772714624 | 0.8899318070140728 |
| GATCCAGATCTAACTTGGGGTGGCTTTGTCTTCTTCTTTTTAGGCCAATTCCACTAATTGTTTGGG | Ins_k3_40_TAG | -4.6228098524726535 | 0.3563389088622098 | 0.48241486581775506 |
| GATCCAGATCTAACTTGGGGTGGCTTTGTCTTCTTCTTTTTATGCCAATTCCACTAATTGTTTGGG | Ins_k3_39_TTA;Ins_k3_40_TAT | -0.693006712372972 | 0.054397923083220986 | 24.55344873620339 |
| GATCCAGATCTAACTTGGGGTGGCTTTGTCTTCTTCTTTTTCAGCCAATTCCACTAATTGTTTGGG | Ins_k3_40_TCA | -3.2814604329731973 | 0.1764911209710844 | 1.84485115940167 |
| GATCCAGATCTAACTTGGGGTGGCTTTGTCTTCTTCTTTTTCCAATTCCACTAATTGTTTGGG | G41T | -1.649710323741835 | 0.05417562458225316 | 9.432382454825717 |
| GATCCAGATCTAACTTGGGGTGGCTTTGTCTTCTTCTTTTTCCACTAATTGTTTGGG | Del_k6_40-45;Del_k6_41-46 | -1.150074693346298 | 0.052194483762079864 | 15.545704172479743 |
| GATCCAGATCTAACTTGGGGTGGCTTTGTCTTCTTCTTTTTCCGCCAATTCCACTAATTGTTTGGG | Ins_k3_40_TCC | 0.3979273667672554 | 0.032106074558694526 | 73.09693240912348 |
| GATCCAGATCTAACTTGGGGTGGCTTTGTCTTCTTCTTTTTCGCCAATTCCACTAATTGTTTGGG | Ins_k2_40_TC | 0.6877222000827219 | 0.03033645127451631 | 97.6687093408714 |
| GATCCAGATCTAACTTGGGGTGGCTTTGTCTTCTTCTTTTTCGGCCAATTCCACTAATTGTTTGGG | Ins_k3_40_TCG | 0.7536869029400919 | 0.03253117964764434 | 100 |
| GATCCAGATCTAACTTGGGGTGGCTTTGTCTTCTTCTTTTTCTGCCAATTCCACTAATTGTTTGGG | Ins_k3_39_TTC;Ins_k3_40_TCT | -0.00636223334586572 | 0.03659537352080029 | 48.78860597383007 |
| GATCCAGATCTAACTTGGGGTGGCTTTGTCTTCTTCTTTTTGAGCCAATTCCACTAATTGTTTGGG | Ins_k3_40_TGA | 0.3458441955628411 | 0.0343373022501981 | 69.38725675193136 |
| GATCCAGATCTAACTTGGGGTGGCTTTGTCTTCTTCTTTTTGCCAATTCCACTAATTGTTTGGG | Ins_k1_36_T;Ins_k1_37_T;Ins_k1_ | -0.09529299538386882 | 0.041413073909872505 | 44.63713069299469 |
| GATCCAGATCTAACTTGGGGTGGCTTTGTCTTCTTCTTTTTGCGCCAATTCCACTAATTGTTTGGG | Ins_k3_40_TGC | 0.37160444821717575 | 0.036761036578915184 | 71.19791135013595 |
| GATCCAGATCTAACTTGGGGTGGCTTTGTCTTCTTCTTTTTGGCCAATTCCACTAATTGTTTGGG | Ins_k2_40_TG | -0.1211419329176115 | 0.04120941535436067 | 43.49809316658227 |
| GATCCAGATCTAACTTGGGGTGGCTTTGTCTTCTTCTTTTTGGG | Del_k19_39-57;Del_k19_40-58;D | -0.19457601591105983 | 0.026454063476955267 | 40.418314800592285 |
| GATCCAGATCTAACTTGGGGTGGCTTTGTCTTCTTCTTTTTGGGCCAATTCCACTAATTGTTTGGG | Ins_k3_40_TGG | 0.22063244227548529 | 0.03351934689650481 | 61.22107402608433 |
| GATCCAGATCTAACTTGGGGTGGCTTTGTCTTCTTCTTTTTGTGCCAATTCCACTAATTGTTTGGG | Ins_k3_39_TTG;Ins_k3_40_TGT | -0.09380580002045524 | 0.03501644452529312 | 44.70356421436534 |
| GATCCAGATCTAACTTGGGGTGGCTTTGTCTTCTTCTTTTTGTTTGGG | Del_k15_40-54;Del_k15_41-55 | -0.5660859841399262 | 0.03482469726361338 | 27.876193657593966 |
| GATCCAGATCTAACTTGGGGTGGCTTTGTCTTCTTCTTTTTTAGCCAATTCCACTAATTGTTTGGG | Ins_k3_40_TTA | -4.494182743268562 | 0.4640524656526145 | 0.5486340098711455 |
| GATCCAGATCTAACTTGGGGTGGCTTTGTCTTCTTCTTTTTTCCACTAATTGTTTGGG | Del_k5_41-45 | -1.4954371510915074 | 0.05973036623902601 | 11.005794245539121 |
| GATCCAGATCTAACTTGGGGTGGCTTTGTCTTCTTCTTTTTTCGCCAATTCCACTAATTGTTTGGG | Ins_k3_40_TTC | 0.45351682550108263 | 0.03627930492081691 | 77.27541514431314 |
| GATCCAGATCTAACTTGGGGTGGCTTTGTCTTCTTCTTTTTTGCCAATTCCACTAATTGTTTGGG | Ins_k2_36_TT;Ins_k2_37_TT;Ins_I | -0.33210159935085215 | 0.042993265163513705 | 35.22504852827879 |
| GATCCAGATCTAACTTGGGGTGGCTTTGTCTTCTTCTTTTTTGGCCAATTCCACTAATTGTTTGGG | Ins_k3_40_TTG | -0.2537498417915281 | 0.04280860896536224 | 38.095996314762274 |
| GATCCAGATCTAACTTGGGGTGGCTTTGTCTTCTTCTTTTTTGGG | Del_k18_40-57;Del_k18_41-58 | -0.14020575018148068 | 0.031177766595574922 | 42.676707714342726 |
| GATCCAGATCTAACTTGGGGTGGCTTTGTCTTCTTCTTTTTTGTTTGGG | Del_k14_41-54 | -0.6403843865508108 | 0.03844889459535329 | 25.880108142832984 |
| GATCCAGATCTAACTTGGGGTGGCTTTGTCTTCTTCTTTTTTTGCCAATTCCACTAATTGTTTGGG | Ins_k3_36_TTT;Ins_k3_37_TTT;In | -1.5444001172807849 | 0.25742769468395055 | 10.479897701308088 |
| GATCCAGATCTAACTTGGGGTGGCTTTGTCTTCTTCTTTTTTTGGG | Del_k17_41-57 | -0.14819737879970513 | 0.03295961939079312 | 42.33701049059554 |
| GATCCAGATCTAACTTGGGGTGGCTTTGTCTTCTTG | Del_k27_36-62 | -2.1624027415692586 | 0.03971207504271541 4 | 5.648874338760507 |
| GATCCAGATCTAACTTGGGGTGGCTTTGTCTTCTTGAACTTTTGCCAATTCCACTAATTGTTTGGG | Ins_k3_35_GAA | 0.5395450492462177 | 0.03220713876310131 | 84.21761334959079 |
| GATCCAGATCTAACTTGGGGTGGCTTTGTCTTCTTGACCTTTTGCCAATTCCACTAATTGTTTGGG | Ins_k3_35_GAC | 0.21129062006031807 | 0.03019972736403706 | 60.651820708228215 |
| GATCCAGATCTAACTTGGGGTGGCTTTGTCTTCTTGACTTTTGCCAATTCCACTAATTGTTTGGG | Ins_k2_35_GA | -0.008309967926552475 | 0.04144919907355599 | 48.69367120271225 |
| GATCCAGATCTAACTTGGGGTGGCTTTGTCTTCTTGAGCTTTTGCCAATTCCACTAATTGTTTGGG | Ins_k3_35_GAG | 0.4746221894032774 | 0.030987332745492816 | 78.9236732629355 |
| GATCCAGATCTAACTTGGGGTGGCTTTGTCTTCTTGATCTTTTGCCAATTCCACTAATTGTTTGGG | Ins_k3_34_TGA;Ins_k3_35_GAT | 0.15914797041094741 | 0.029877246359309276 | 57.570311254411045 |
| GATCCAGATCTAACTTGGGGTGGCTTTGTCTTCTTGCACTTTTGCCAATTCCACTAATTGTTTGGG | Ins_k3_35_GCA | -0.016252424266686494 | 0.036886959591547024 | 48.30845564903369 |
| GATCCAGATCTAACTTGGGGTGGCTTTGTCTTCTTGCCAATTCCACTAATTGTTTGGG | Del_k5_34-38;Del_k5_35-39;Del_ | 0.07949859759211086 | 0.02951182750772157 | 53.162732417007824 |
| GATCCAGATCTAACTTGGGGTGGCTTTGTCTTCTTGCCCTTTTGCCAATTCCACTAATTGTTTGGG | Ins_k3_35_GCC | 0.1338110609610772 | 0.03740010591450015 | 56.129981297904955 |
| GATCCAGATCTAACTTGGGGTGGCTTTGTCTTCTTGCCTTTTGCCAATTCCACTAATTGTTTGGG | Ins_k2_35_GC | 0.18385866363243797 | 0.037414244218092446 | 59.01063597566699 |
| GATCCAGATCTAACTTGGGGTGGCTTTGTCTTCTTGCGCTTTTGCCAATTCCACTAATTGTTTGGG | Ins_k3_35_GCG | 0.43343890964370724 | 0.03046073942944947 | 75.73935787677017 |
| GATCCAGATCTAACTTGGGGTGGCTTTGTCTTCTTGCTCTTTTGCCAATTCCACTAATTGTTTGGG | Ins_k3_34_TGC;Ins_k3_35_GCT | -0.48802660383869945 | 0.04123841400042605 | 30.13937423268585 |
| GATCCAGATCTAACTTGGGGTGGCTTTGTCTTCTTGCTTTTGCCAATTCCACTAATTGTTTGGG | Ins_k1_35_G | 0.02100909482866231 | 0.03595812899869614 | 50.1424587704001 |
| GATCCAGATCTAACTTGGGGTGGCTTTGTCTTCTTGG | Del_k26_36-61 | -1.8990832237172457 | 0.03746875514288184 | 7.350554930207615 |
| GATCCAGATCTAACTTGGGGTGGCTTTGTCTTCTTGGACTTTTGCCAATTCCACTAATTGTTTGGG | Ins_k3_35_GGA | 0.5028613023766084 | 0.03078870511052473 | 81.18417485105734 |
| GATCCAGATCTAACTTGGGGTGGCTTTGTCTTCTTGGCCTTTTGCCAATTCCACTAATTGTTTGGG | Ins_k3_35_GGC | -0.058081399605397155 | 0.03342809761429948 | 46.32944106743446 |
| GATCCAGATCTAACTTGGGGTGGCTTTGTCTTCTTGGCTTTTGCCAATTCCACTAATTGTTTGGG | Ins_k2_35_GG | 0.29053518821908764 | 0.03729759398143377 | 65.65371735731684 |
| GATCCAGATCTAACTTGGGGTGGCTTTGTCTTCTTGGG | Del_k25_34-58;Del_k25_35-59;D | -1.6616054555109023 | 0.03776421958554853 | 9.320847698066855 |
| GATCCAGATCTAACTTGGGGTGGCTTTGTCTTCTTGGGCTTTTGCCAATTCCACTAATTGTTTGGG | Ins_k3_35_GGG | 0.007462005214280754 | 0.036172398569979634 | 49.46775484387495 |
| GATCCAGATCTAACTTGGGGTGGCTTTGTCTTCTTGGTCTTTTGCCAATTCCACTAATTGTTTGGG | Ins_k3_34_TGG;Ins_k3_35_GGT | 0.008225262886534912 | 0.03200195882130019 | 49.50552589997923 |
| GATCCAGATCTAACTTGGGGTGGCTTTGTCTTCTTGTACTTTTGCCAATTCCACTAATTGTTTGGG | Ins_k3_35_GTA | -0.42867965729114216 | 0.04600426926078775 | 31.982196145314617 |
| GATCCAGATCTAACTTGGGGTGGCTTTGTCTTCTTGTATCTTTTGCCAATTCCACTAATTGTTTGGG | Ins_k4_34_TGTA;Ins_k4_35_GTA | -0.8771315332813648 | 0.06447124086291076 | 20.424343566838097 |
| GATCCAGATCTAACTTGGGGTGGCTTTGTCTTCTTGTCCTTTTGCCAATTCCACTAATTGTTTGGG | Ins_k3_35_GTC | -0.15371994767866148 | 0.03409822768232893 | 42.10384586049956 |
| GATCCAGATCTAACTTGGGGTGGCTTTGTCTTCTTGTCTTTTGCCAATTCCACTAATTGTTTGGG | Ins_k2_34_TG;Ins_k2_35_GT | -0.13535794728666356 | 0.039939850147841405 | 42.88409826964847 |
| GATCCAGATCTAACTTGGGGTGGCTTTGTCTTCTTGTGCTTTTGCCAATTCCACTAATTGTTTGGG | Ins_k3_35_GTG | -0.07384670962836659 | 0.03263874577509219 | 45.60477040402903 |
| GATCCAGATCTAACTTGGGGTGGCTTTGTCTTCTTGTTCTTTTGCCAATTCCACTAATTGTTTGGG | Ins_k3_33_TTG;Ins_k3_34_TGT;I | -1.0000726925005714 | 0.049451321747177375 | 18.061567573285057 |
| GATCCAGATCTAACTTGGGGTGGCTTTGTCTTCTTGTTTGGG | Del_k21_34-54;Del_k21_35-55;D | -1.3375039132485418 | 0.0364800344600396 | 12.8887536565636 |
| GATCCAGATCTAACTTGGGGTGGCTTTGTCTTCTTGTTTTGCCAATTCCACTAATTGTTTGGG | C36G | -0.3370897795093199 | 0.0359078142248961 | 35.04977714616846 |
| GATCCAGATCTAACTTGGGGTGGCTTTGTCTTCTTTAACTTTTGCCAATTCCACTAATTGTTTGGG | Ins_k3_35_TAA | -1.6132547847992285 | 0.06514689538372377 | 9.782589753859408 |
| GATCCAGATCTAACTTGGGGTGGCTTTGTCTTCTTTAATTGTTTGGG | Del_k16_36-51 | -1.0247064214171642 | 0.04881500485910747 | 17.622079157122837 |
| GATCCAGATCTAACTTGGGGTGGCTTTGTCTTCTTTACCTTTTGCCAATTCCACTAATTGTTTGGG | Ins_k3_35_TAC | -0.8851724550595921 | 0.05002361883026266 | 20.260771534136474 |
| GATCCAGATCTAACTTGGGGTGGCTTTGTCTTCTTTACTTTTGCCAATTCCACTAATTGTTTGGG | Ins_k2_35_TA | -0.9083337142897501 | 0.059711897402457394 | 19.79689922246653 |

| | | | | |
|---|---|---|---|---|
| **GATCCAGATCTAACTTGGGGTGGCTTTGTCTTCTTTAGACTTTTGCCAATTCCACTAATTGTTTGGG** | Ins_k4_35_TAGA | -1.2043246036119686 | 0.06587433153775417 | 14.72481890792242 |
| **GATCCAGATCTAACTTGGGGTGGCTTTGTCTTCTTTAGCTTTTGCCAATTCCACTAATTGTTTGGG** | Ins_k3_35_TAG | -3.4393779857304505 | 0.1580411090391624 | 1.575355638594034 |
| **GATCCAGATCTAACTTGGGGTGGCTTTGTCTTCTTTATCTTTTGCCAATTCCACTAATTGTTTGGG** | Ins_k3_34_TTA;Ins_k3_35_TAT | -1.6524810292514733 | 0.07219938477491628 | 9.40628427267355 |
| **GATCCAGATCTAACTTGGGGTGGCTTTGTCTTCTTTATGCTTTTGCCAATTCCACTAATTGTTTGGG** | Ins_k4_35_TATG | -1.092545926005657 | 0.05555792757327489 | 16.46625456498717 |
| **GATCCAGATCTAACTTGGGGTGGCTTTGTCTTCTTTCACTTTTGCCAATTCCACTAATTGTTTGGG** | Ins_k3_35_TCA | -0.616682857567697 | 0.0480046075512223 | 26.500833285476883 |
| **GATCCAGATCTAACTTGGGGTGGCTTTGTCTTCTTTCCACTAATTGTTTGGG** | Del_k11_35-45;Del_k11_36-46 | -0.3012911275870976 | 0.03424266006900646 | 36.327241240163175 |
| **GATCCAGATCTAACTTGGGGTGGCTTTGTCTTCTTTCCCTTTTGCCAATTCCACTAATTGTTTGGG** | Ins_k3_35_TCC | -1.2031013149223622 | 0.06460860021820351 | 14.742842634211632 |
| **GATCCAGATCTAACTTGGGGTGGCTTTGTCTTCTTTCCTTTTGCCAATTCCACTAATTGTTTGGG** | Ins_k2_35_TC | -0.9330755352650478 | 0.06137489851725838 | 19.313097632192783 |
| **GATCCAGATCTAACTTGGGGTGGCTTTGTCTTCTTTCGCCTTTTGCCAATTCCACTAATTGTTTGGG** | Ins_k4_35_TCGC | 0.6170012701146916 | 0.03223696216361285 | 91.0000726542758 |
| **GATCCAGATCTAACTTGGGGTGGCTTTGTCTTCTTTCGCTTTTGCCAATTCCACTAATTGTTTGGG** | Ins_k3_35_TCG | -0.12704378280232842 | 0.03956632467168252 | 43.24213002144266 |
| **GATCCAGATCTAACTTGGGGTGGCTTTGTCTTCTTTCTCTTTTGCCAATTCCACTAATTGTTTGGG** | Ins_k3_34_TTC;Ins_k3_35_TCT | -2.3139159187174934 | 0.09040800443335074 | 4.854679650188574 |
| **GATCCAGATCTAACTTGGGGTGGCTTTGTCTTCTTTCTTTTGCCAATTCCACTAATTGTTTGGG** | Ins_k1_33_T;Ins_k1_34_T;Ins_k1_ | -1.114466771833304 | 0.06791078950987636 | 16.109227798195814 |
| **GATCCAGATCTAACTTGGGGTGGCTTTGTCTTCTTTGACTTTTGCCAATTCCACTAATTGTTTGGG** | Ins_k3_35_TGA | -0.04444394538108076 | 0.03730037045415302 | 46.965584528755485 |
| **GATCCAGATCTAACTTGGGGTGGCTTTGTCTTCTTTGCCAATTCCACTAATTGTTTGGG** | Del_k4_34-37;Del_k4_35-38;Del_ | 0.22647044259592497 | 0.03135063713683472 | 61.579527984953835 |
| **GATCCAGATCTAACTTGGGGTGGCTTTGTCTTCTTTGCCTTTTGCCAATTCCACTAATTGTTTGGG** | Ins_k3_35_TGC | -0.16749329303833188 | 0.03622843556817748 | 41.52791043405166 |
| **GATCCAGATCTAACTTGGGGTGGCTTTGTCTTCTTTGCTTTTGCCAATTCCACTAATTGTTTGGG** | Ins_k2_35_TG | -0.3361893659568708 | 0.0398644276504446 | 35.08135065299474 |
| **GATCCAGATCTAACTTGGGGTGGCTTTGTCTTCTTTGGCTTTTGCCAATTCCACTAATTGTTTGGG** | Ins_k3_35_TGG | 0.008206697321357703 | 0.03482306156005429 | 49.504606810443235 |
| **GATCCAGATCTAACTTGGGGTGGCTTTGTCTTCTTTGGG** | Del_k24_34-57;Del_k24_35-58;D | -1.4682848339808112 | 0.03464158329690042 | 11.308721032242541 |
| **GATCCAGATCTAACTTGGGGTGGCTTTGTCTTCTTTGTCTTTTGCCAATTCCACTAATTGTTTGGG** | Ins_k3_34_TTG;Ins_k3_35_TGT | -1.068888035724264 | 0.05028162695434844 | 16.860456010198888 |
| **GATCCAGATCTAACTTGGGGTGGCTTTGTCTTCTTTGTTTGGG** | Del_k20_35-54;Del_k20_36-55 | -0.574260261011115 | 0.032884512337782626 | 27.64925472699525 |
| **GATCCAGATCTAACTTGGGGTGGCTTTGTCTTCTTTTACTTTTGCCAATTCCACTAATTGTTTGGG** | Ins_k3_35_TTA | -1.799717752902133 | 0.08974870057522148 | 8.118466501947205 |
| **GATCCAGATCTAACTTGGGGTGGCTTTGTCTTCTTTTCCACTAATTGTTTGGG** | Del_k10_36-45 | -0.3220238808075264 | 0.03793822495443614 | 35.58183141218473 |
| **GATCCAGATCTAACTTGGGGTGGCTTTGTCTTCTTTTCCTTTTGCCAATTCCACTAATTGTTTGGG** | Ins_k3_35_TTC | -1.3643663937001698 | 0.06021390451206142 | 12.547138619052376 |
| **GATCCAGATCTAACTTGGGGTGGCTTTGTCTTCTTTTCGCTTTTGCCAATTCCACTAATTGTTTGGG** | Ins_k4_35_TTCG | -0.03265345968038347 | 0.039588124709647676 | 47.52260892273652 |
| **GATCCAGATCTAACTTGGGGTGGCTTTGTCTTCTTTTCTTTTGCCAATTCCACTAATTGTTTGGG** | Ins_k2_33_TT;Ins_k2_34_TT;Ins_ | -2.1823640762696193 | 0.1100156918126246 | 5.537233227172481 |
| **GATCCAGATCTAACTTGGGGTGGCTTTGTCTTCTTTTGCCAATTCCACTAATTGTTTGGG** | Del_k3_29-31;Del_k3_30-32;Del_ | 0.05366383969344757 | 0.0300990647625484 | 51.80687562310976 |
| **GATCCAGATCTAACTTGGGGTGGCTTTGTCTTCTTTTGCTTTTGCCAATTCCACTAATTGTTTGGG** | Ins_k3_35_TTG | -1.081825484574774 | 0.050946229656770106 | 16.643729688738777 |
| **GATCCAGATCTAACTTGGGGTGGCTTTGTCTTCTTTTGGG** | Del_k23_35-57;Del_k23_36-58 | -1.162638983518998 | 0.03585320784967487 | 15.351605344700758 |
| **GATCCAGATCTAACTTGGGGTGGCTTTGTCTTCTTTTGTTTGGG** | Del_k19_36-54 | -0.45223140255719474 | 0.03630862058375232 | 31.23776039945337 |
| **GATCCAGATCTAACTTGGGGTGGCTTTGTCTTCTTTTTCTTTTGCCAATTCCACTAATTGTTTGGG** | Ins_k3_33_TTT;Ins_k3_34_TTT;In | -3.060201611176091 | 0.1950626094885193 | 2.301721735693006 |
| **GATCCAGATCTAACTTGGGGTGGCTTTGTCTTCTTTTTGCCAATTCCACTAATTGTTTGGG** | Del_k2_35-36;Del_k2_36-37 | -0.12977064754943352 | 0.03412764112181447 | 43.12437520519275 |
| **GATCCAGATCTAACTTGGGGTGGCTTTGTCTTCTTTTTGGG** | Del_k22_36-57 | -0.7925754095364507 | 0.032732285977213296 | 22.226463428056995 |
| **GATCCAGATCTAACTTGGGGTGGCTTTGTCTTCTTTTTTGCCAATTCCACTAATTGTTTGGG** | Del_k1_36-36 | -0.30599955492285325 | 0.04572491980468681 | 36.1565991077669 |
| **GATCCAGATCTAACTTGGGGTGGCTTTGTCTTCTTTTTTTGCCAATTCCACTAATTGTTTGGG** | C36T | -0.8837363049736552 | 0.044065325728730134 | 20.289889947117228 |
| **GATCCAGATCTAACTTGGGGTGGCTTTGTCTTG** | Del_k30_33-62 | -2.6153745703237035 | 0.049672351950210865 | 3.5911930186823335 |
| **GATCCAGATCTAACTTGGGGTGGCTTTGTCTTGAACTTCTTTTGCCAATTCCACTAATTGTTTGGG** | Ins_k3_32_GAA | 0.40705687036938 | 0.03184299380358939 | 73.76732664382786 |
| **GATCCAGATCTAACTTGGGGTGGCTTTGTCTTGACACTTCTTTTGCCAATTCCACTAATTGTTTGGG** | Ins_k4_32_GACA | 0.2249722715712933 | 0.038605062984074 | 61.48734039404808 |
| **GATCCAGATCTAACTTGGGGTGGCTTTGTCTTGACCTTCTTTTGCCAATTCCACTAATTGTTTGGG** | Ins_k3_32_GAC | 0.35335554942945036 | 0.03109028519071366 | 69.91041133109874 |
| **GATCCAGATCTAACTTGGGGTGGCTTTGTCTTGACTTCTTTTGCCAATTCCACTAATTGTTTGGG** | Ins_k2_32_GA | -0.5116730194113839 | 0.03495284698690798 | 29.435046298608892 |
| **GATCCAGATCTAACTTGGGGTGGCTTTGTCTTGAGCTTCTTTTGCCAATTCCACTAATTGTTTGGG** | Ins_k3_32_GAG | 0.31732716330830024 | 0.03400995307788677 | 67.4364854165425 |
| **GATCCAGATCTAACTTGGGGTGGCTTTGTCTTGATCTTCTTTTGCCAATTCCACTAATTGTTTGGG** | Ins_k3_31_TGA;Ins_k3_32_GAT | 0.39141037746273954 | 0.03435989171098912 | 72.62210937127209 |
| **GATCCAGATCTAACTTGGGGTGGCTTTGTCTTGCACTTCTTTTGCCAATTCCACTAATTGTTTGGG** | Ins_k3_32_GCA | -0.19137876665259107 | 0.03618082613890015 | 40.547749034109884 |
| **GATCCAGATCTAACTTGGGGTGGCTTTGTCTTGCCAATTCCACTAATTGTTTGGG** | Del_k8_31-38;Del_k8_32-39;Del_ | -0.1614793131474767 | 0.03150937907656141 | 41.77841094983643 |
| **GATCCAGATCTAACTTGGGGTGGCTTTGTCTTGCCCTTCTTTTGCCAATTCCACTAATTGTTTGGG** | Ins_k3_32_GCC | -0.7644144212305642 | 0.04491089320331169 | 22.86127917173403 |
| **GATCCAGATCTAACTTGGGGTGGCTTTGTCTTGCCTTCTTTTGCCAATTCCACTAATTGTTTGGG** | Ins_k2_32_GC | 0.07468128817570863 | 0.04139431014213546 | 52.90724695586866 |
| **GATCCAGATCTAACTTGGGGTGGCTTTGTCTTGCGCTTCTTTTGCCAATTCCACTAATTGTTTGGG** | Ins_k3_32_GCG | 0.35683288474540875 | 0.03197715648835905 | 70.15393643721018 |
| **GATCCAGATCTAACTTGGGGTGGCTTTGTCTTGCTCTTCTTTTGCCAATTCCACTAATTGTTTGGG** | Ins_k3_31_TGC;Ins_k3_32_GCT | -0.2718430458107732 | 0.03993274598604664 | 37.41291587258609 |
| **GATCCAGATCTAACTTGGGGTGGCTTTGTCTTGCTTCTTTTGCCAATTCCACTAATTGTTTGGG** | Ins_k1_32_G | -0.21991541244270457 | 0.04363447186994547 | 39.40700618137523 |
| **GATCCAGATCTAACTTGGGGTGGCTTTGTCTTGG** | Del_k29_33-61 | -2.714872329079759 | 0.052518181786979104 | 3.251078234302305 |
| **GATCCAGATCTAACTTGGGGTGGCTTTGTCTTGGACTTCTTTTGCCAATTCCACTAATTGTTTGGG** | Ins_k3_32_GGA | 0.6976619829208357 | 0.02819814791444902 | 98.64435592664115 |
| **GATCCAGATCTAACTTGGGGTGGCTTTGTCTTGGCCTTCTTTTGCCAATTCCACTAATTGTTTGGG** | Ins_k3_32_GGC | 0.13961768030907395 | 0.032380028917141014 | 56.45685482997218 |
| **GATCCAGATCTAACTTGGGGTGGCTTTGTCTTGGCTTCTTTTGCCAATTCCACTAATTGTTTGGG** | Ins_k2_32_GG | -0.27529803020488913 | 0.050258503791494887 | 37.28387787260566 |
| **GATCCAGATCTAACTTGGGGTGGCTTTGTCTTGGG** | Del_k28_31-58;Del_k28_32-59;D | -3.5763282024997913 | 0.0660248092917874 | 1.3737315995181367 |
| **GATCCAGATCTAACTTGGGGTGGCTTTGTCTTGGGCTTCTTTTGCCAATTCCACTAATTGTTTGGG** | Ins_k3_32_GGG | -0.8508240938110134 | 0.04975115859281414 | 20.968785789860128 |
| **GATCCAGATCTAACTTGGGGTGGCTTTGTCTTGGTCTTCTTTTGCCAATTCCACTAATTGTTTGGG** | Ins_k3_31_TGG;Ins_k3_32_GGT | 0.027874670825524195 | 0.0338904776193205 7 | 50.48790010171777 |
| **GATCCAGATCTAACTTGGGGTGGCTTTGTCTTGTACTTCTTTTGCCAATTCCACTAATTGTTTGGG** | Ins_k3_32_GTA | -0.9849052804045249 | 0.053373491969767095 | 18.337602890659767 |
| **GATCCAGATCTAACTTGGGGTGGCTTTGTCTTGTCCTTCTTTTGCCAATTCCACTAATTGTTTGGG** | Ins_k3_32_GTC | -0.27666598175181983 | 0.047489500248386066 | 37.23291020279358 |
| **GATCCAGATCTAACTTGGGGTGGCTTTGTCTTGTCTTCTTTTGCCAATTCCACTAATTGTTTGGG** | Ins_k2_31_TG;Ins_k2_32_GT | -0.05295476277357569 | 0.042868826582754774 | 46.56756515272232 |
| **GATCCAGATCTAACTTGGGGTGGCTTTGTCTTGTGCTTCTTTTGCCAATTCCACTAATTGTTTGGG** | Ins_k3_32_GTG | -0.1541359910294171 | 0.037476174542366916 | 42.086332478803676 |
| **GATCCAGATCTAACTTGGGGTGGCTTTGTCTTGTTCTTCTTTTGCCAATTCCACTAATTGTTTGGG** | Ins_k3_30_TTG;Ins_k3_31_TGT;I | -0.558061165176417 | 0.044351315359115524 | 28.100795052537727 |
| **GATCCAGATCTAACTTGGGGTGGCTTTGTCTTGTTCTTTTGCCAATTCCACTAATTGTTTGGG** | C33G | -1.8496310573121437 | 0.060314120821714834 | 7.723193756291461 |
| **GATCCAGATCTAACTTGGGGTGGCTTTGTCTTGTTTGGG** | Del_k24_31-54;Del_k24_32-55;D | -2.480354514861287 | 0.07066040568208179 | 4.110334943428917 |
| **GATCCAGATCTAACTTGGGGTGGCTTTGTCTTTAACTTCTTTTGCCAATTCCACTAATTGTTTGGG** | Ins_k3_32_TAA | -2.2479173968943664 | 0.10517967749441304 | 5.185890839943922 |
| **GATCCAGATCTAACTTGGGGTGGCTTTGTCTTTAATTGTTTGGG** | Del_k19_33-51 | -3.325534370245199 | 0.1453122218314034 | 1.7653070911222646 |
| **GATCCAGATCTAACTTGGGGTGGCTTTGTCTTTACCTTCTTTTGCCAATTCCACTAATTGTTTGGG** | Ins_k3_32_TAC | -1.5212162547142123 | 0.06437134632702061 | 10.72570052917153 |
| **GATCCAGATCTAACTTGGGGTGGCTTTGTCTTTACTTCTTTTGCCAATTCCACTAATTGTTTGGG** | Ins_k2_32_TA | -1.5960002628554018 | 0.08384922472698023 | 9.952848304309477 |
| **GATCCAGATCTAACTTGGGGTGGCTTTGTCTTTAGCCTTCTTTTGCCAATTCCACTAATTGTTTGGG** | Ins_k4_32_TAGC | -1.4829596138556487 | 0.07487208927117932 | 11.143979767722199 |
| **GATCCAGATCTAACTTGGGGTGGCTTTGTCTTTAGCTTCTTTTGCCAATTCCACTAATTGTTTGGG** | Ins_k3_32_TAG | -4.078807500124043 | 0.22317285526531813 | 0.8311471156461271 |
| **GATCCAGATCTAACTTGGGGTGGCTTTGTCTTTATCTTCTTTTGCCAATTCCACTAATTGTTTGGG** | Ins_k3_31_TTA;Ins_k3_32_TAT | -1.4096431433765368 | 0.06820795319945099 | 11.991713761854886 |
| **GATCCAGATCTAACTTGGGGTGGCTTTGTCTTTCACTTCTTTTGCCAATTCCACTAATTGTTTGGG** | Ins_k3_32_TCA | -1.4421499098251154 | 0.05557080663131723 | 11.608169587078212 |
| **GATCCAGATCTAACTTGGGGTGGCTTTGTCTTTCCACTAATTGTTTGGG** | Del_k14_32-45;Del_k14_33-46 | -1.2526925598041578 | 0.041296299008147415 | 14.029559196698584 |
| **GATCCAGATCTAACTTGGGGTGGCTTTGTCTTTCCCTTCTTTTGCCAATTCCACTAATTGTTTGGG** | Ins_k3_32_TCC | -2.1405321991242814 | 0.08421854522386696 | 5.773779174333118 |
| **GATCCAGATCTAACTTGGGGTGGCTTTGTCTTTCCTTCTTTTGCCAATTCCACTAATTGTTTGGG** | Ins_k2_32_TC | -1.9569787129299425 | 0.09442879187569475 | 6.937075734884549 |
| **GATCCAGATCTAACTTGGGGTGGCTTTGTCTTTCGCTTCTTTTGCCAATTCCACTAATTGTTTGGG** | Ins_k3_32_TCG | -0.5395360538460992 | 0.043729731544206334 | 28.62621713453974 |
| **GATCCAGATCTAACTTGGGGTGGCTTTGTCTTTCTCTTCTTTTGCCAATTCCACTAATTGTTTGGG** | Ins_k3_31_TTC;Ins_k3_32_TCT | -2.4056105056092183 | 0.09723799134864541 | 4.429330877482939 |

| | | | | |
|---|---|---|---|---|
| GATCCAGATCTAACTTGGGGTGGCTTTGTCTTTCTTCTTTTGCCAATTCCACTAATTGTTTGGG | Ins_k1_30_T;Ins_k1_31_T;Ins_k1_ | -1.1739424884524616 | 0.06927416390478644 | 15.179055444484517 |
| GATCCAGATCTAACTTGGGGTGGCTTTGTCTTTCTTTTGCCAATTCCACTAATTGTTTGGG | Del_k2_32-33;Del_k2_33-34 | -1.7384528428527861 | 0.0680427241921577 | 8.631395470347364 |
| GATCCAGATCTAACTTGGGGTGGCTTTGTCTTTGACTTCTTTTGCCAATTCCACTAATTGTTTGGG | Ins_k3_32_TGA | -0.013119046303288084 | 0.04223566068217491 | 48.46006169489206 |
| GATCCAGATCTAACTTGGGGTGGCTTTGTCTTTGCCAATTCCACTAATTGTTTGGG | Del_k7_31-37;Del_k7_32-38;Del_ | -0.32990474173220846 | 0.033601821702489945 | 35.302518008056836 |
| GATCCAGATCTAACTTGGGGTGGCTTTGTCTTTGCCCTTCTTTTGCCAATTCCACTAATTGTTTGGG | Ins_k4_32_TGCC | -0.6589788549171922 | 0.04740063552480064 | 25.403327766787026 |
| GATCCAGATCTAACTTGGGGTGGCTTTGTCTTTGCCTTCTTTTGCCAATTCCACTAATTGTTTGGG | Ins_k3_32_TGC | -0.4109478164045301 | 0.041208762408201996 | 32.55435710067338 |
| GATCCAGATCTAACTTGGGGTGGCTTTGTCTTTGCTTCTTTTGCCAATTCCACTAATTGTTTGGG | Ins_k2_32_TG | -0.8222993969467958 | 0.04775503568185122 | 21.57552645549931 |
| GATCCAGATCTAACTTGGGGTGGCTTTGTCTTTGGCTTCTTTTGCCAATTCCACTAATTGTTTGGG | Ins_k3_32_TGG | -1.1211843134422677 | 0.0554813009503387 | 16.0013760450633 |
| GATCCAGATCTAACTTGGGGTGGCTTTGTCTTTGGG | Del_k27_31-57;Del_k27_32-58;D | -3.059205112738794 | 0.06464572335146947 | 2.304016541000771 |
| GATCCAGATCTAACTTGGGGTGGCTTTGTCTTTGTCTTCTTTTGCCAATTCCACTAATTGTTTGGG | Ins_k3_31_TTG;Ins_k3_32_TGT | -0.7890343696063947 | 0.04731847536744112 | 22.305307735594823 |
| GATCCAGATCTAACTTGGGGTGGCTTTGTCTTTGTTTGGG | Del_k23_32-54;Del_k23_33-55 | -2.5170644468168026 | 0.07370421285186973 | 3.9621808286468263 |
| GATCCAGATCTAACTTGGGGTGGCTTTGTCTTTTACTTCTTTTGCCAATTCCACTAATTGTTTGGG | Ins_k3_32_TTA | -2.2436169819971994 | 0.10126719056273799 | 5.2082403437932525 |
| GATCCAGATCTAACTTGGGGTGGCTTTGTCTTTTCCACTAATTGTTTGGG | Del_k13_33-45 | -1.2510687273437073 | 0.04291691795066989 | 14.052359357138293 |
| GATCCAGATCTAACTTGGGGTGGCTTTGTCTTTTCCTTCTTTTGCCAATTCCACTAATTGTTTGGG | Ins_k3_32_TTC | -2.366674002209135 | 0.10035519031744011 | 4.605195085213299 |
| GATCCAGATCTAACTTGGGGTGGCTTTGTCTTTTCTTCTTTTGCCAATTCCACTAATTGTTTGGG | Ins_k2_30_TT;Ins_k2_31_TT;Ins_I | -1.4493738573684976 | 0.08993758166574062 | 11.524614939463032 |
| GATCCAGATCTAACTTGGGGTGGCTTTGTCTTTTCTTTTGCCAATTCCACTAATTGTTTGGG | Del_k1_33-33 | -2.973109183899654 | 0.12677316522733567 | 2.5111726892293245 |
| GATCCAGATCTAACTTGGGGTGGCTTTGTCTTTTGCCAATTCCACTAATTGTTTGGG | Del_k6_29-34;Del_k6_30-35;Del_ | -0.5035552923998508 | 0.0996158453823786 | 29.67496444654898 |
| GATCCAGATCTAACTTGGGGTGGCTTTGTCTTTTGCTTCTTTTGCCAATTCCACTAATTGTTTGGG | Ins_k3_32_TTG | -1.480537720025057 | 0.057183728363121356 | 11.171002012865882 |
| GATCCAGATCTAACTTGGGGTGGCTTTGTCTTTTGGG | Del_k26_32-57;Del_k26_33-58 | -3.0407438095209423 | 0.06393456421153094 | 2.3469467434440268 |
| GATCCAGATCTAACTTGGGGTGGCTTTGTCTTTTGTTTGGG | Del_k22_33-54 | -2.7530281007745048 | 0.06849141303695747 | 3.129367583151888 |
| GATCCAGATCTAACTTGGGGTGGCTTTGTCTTTTTCTTCTTTTGCCAATTCCACTAATTGTTTGGG | Ins_k3_30_TTT;Ins_k3_31_TTT;In | -2.213655214171596 | 0.11854710097231234 | 5.366649693572105 |
| GATCCAGATCTAACTTGGGGTGGCTTTGTCTTTTTCTTTTGCCAATTCCACTAATTGTTTGGG | C33T | -2.022505217265036 | 0.06756284231688138 | 6.4970863173572715 |
| GATCCAGATCTAACTTGGGGTGGCTTTGTCTTTTTGCCAATTCCACTAATTGTTTGGG | Del_k5_32-36;Del_k5_33-37 | -0.77996361451106 | 0.04209861141289786 | 22.508554124934246 |
| GATCCAGATCTAACTTGGGGTGGCTTTGTCTTTTTTGGG | Del_k25_33-57 | -2.4771763604312103 | 0.06003845971478935 | 4.123419003206608 |
| GATCCAGATCTAACTTGGGGTGGCTTTGTCTTTTTTGCCAATTCCACTAATTGTTTGGG | Del_k4_33-36 | -0.9919057834649341 | 0.06047253699336557 | 18.209678734657558 |
| GATCCAGATCTAACTTGGGGTGGCTTTGTG | Del_k33_30-62 | -3.896517176890128 | 0.0777608122243752 | 0.9973453876418293 |
| GATCCAGATCTAACTTGGGGTGGCTTTGTGAACTTCTTCTTTTGCCAATTCCACTAATTGTTTGGG | Ins_k3_29_GAA | 0.3140998668287731 | 0.03254715154006594 | 67.21919869749246 |
| GATCCAGATCTAACTTGGGGTGGCTTTGTGAATCTTCTTCTTTTGCCAATTCCACTAATTGTTTGGG | Ins_k4_28_TGAA;Ins_k4_29_GAA | 0.760436165055147 | 0.03304037260779251 | 100 |
| GATCCAGATCTAACTTGGGGTGGCTTTGTGACCCTTCTTCTTTTGCCAATTCCACTAATTGTTTGGG | Ins_k4_29_GACC | 0.7855297787924371 | 0.03239564494330014 | 100 |
| GATCCAGATCTAACTTGGGGTGGCTTTGTGACCTTCTTCTTTTGCCAATTCCACTAATTGTTTGGG | Ins_k3_29_GAC | 0.5751846577245221 | 0.028925047233506498 | 87.27322305841062 |
| GATCCAGATCTAACTTGGGGTGGCTTTGTGACTTCTTCTTTTGCCAATTCCACTAATTGTTTGGG | Ins_k2_29_GA | 0.7589095193538582 | 0.03185020766501737 | 100 |
| GATCCAGATCTAACTTGGGGTGGCTTTGTGAGCTTCTTCTTTTGCCAATTCCACTAATTGTTTGGG | Ins_k3_29_GAG | 0.502220801312675 | 0.03134690571035358 | 81.13219294969934 |
| GATCCAGATCTAACTTGGGGTGGCTTTGTGATCTTCTTCTTTTGCCAATTCCACTAATTGTTTGGG | Ins_k3_28_TGA;Ins_k3_29_GAT | 0.4804857315643396 | 0.03435346275883329 | 79.3878049466865 |
| GATCCAGATCTAACTTGGGGTGGCTTTGTGCACTTCTTCTTTTGCCAATTCCACTAATTGTTTGGG | Ins_k3_29_GCA | 0.27172548085756953 | 0.03252410260158563 | 64.43033197691305 |
| GATCCAGATCTAACTTGGGGTGGCTTTGTGCCAATTCCACTAATTGTTTGGG | Del_k11_29-39;Del_k11_30-40 | -0.5257978000549706 | 0.03546295456075104 | 29.022205234924055 |
| GATCCAGATCTAACTTGGGGTGGCTTTGTGCCCTTCTTCTTTTGCCAATTCCACTAATTGTTTGGG | Ins_k3_29_GCC | 0.2694485401444784 | 0.03633865605835631 | 64.28379482239305 |
| GATCCAGATCTAACTTGGGGTGGCTTTGTGCCTCTTCTTCTTTTGCCAATTCCACTAATTGTTTGGG | Ins_k4_28_TGCC;Ins_k4_29_GC | 0.5474354091664373 | 0.033704309814538806 | 84.88474914009649 |
| GATCCAGATCTAACTTGGGGTGGCTTTGTGCCTTCTTCTTTTGCCAATTCCACTAATTGTTTGGG | Ins_k2_29_GC | 0.5317428720391659 | 0.03642674816960196 | 83.56308928294588 |
| GATCCAGATCTAACTTGGGGTGGCTTTGTGCGACTTCTTCTTTTGCCAATTCCACTAATTGTTTGGG | Ins_k4_29_GCGA | 0.9803588433945123 | 0.028649160079841966 | 100 |
| GATCCAGATCTAACTTGGGGTGGCTTTGTGCGCTTCTTCTTTTGCCAATTCCACTAATTGTTTGGG | Ins_k3_29_GCG | 0.5697318108867921 | 0.03066310959894146 | 86.79863065582096 |
| GATCCAGATCTAACTTGGGGTGGCTTTGTGCTCTTCTTCTTTTGCCAATTCCACTAATTGTTTGGG | Ins_k3_28_TGC;Ins_k3_29_GCT | 0.05987441200148147 | 0.03128255915343254 | 52.129627168717064 |
| GATCCAGATCTAACTTGGGGTGGCTTTGTGCTTCTTCTTTTGCCAATTCCACTAATTGTTTGGG | Ins_k1_29_G | 0.4176193667176482 | 0.03235498032874542 | 74.5506232635202 |
| GATCCAGATCTAACTTGGGGTGGCTTTGTGG | Del_k32_30-61 | -3.6980329405816175 | 0.085905073868287593 | 1.2163153626673426 |
| GATCCAGATCTAACTTGGGGTGGCTTTGTGGACTTCTTCTTTTGCCAATTCCACTAATTGTTTGGG | Ins_k3_29_GGA | 0.6126033656109395 | 0.029788989587787525 | 90.60074177807736 |
| GATCCAGATCTAACTTGGGGTGGCTTTGTGGCCTTCTTCTTTTGCCAATTCCACTAATTGTTTGGG | Ins_k3_29_GGC | 0.4278232485165596 | 0.03056253372076505 | 75.3152233215264 |
| GATCCAGATCTAACTTGGGGTGGCTTTGTGGCTTCTTCTTTTGCCAATTCCACTAATTGTTTGGG | Ins_k2_29_GG | 0.36122504912605047 | 0.034565084181152984 | 70.46274172334782 |
| GATCCAGATCTAACTTGGGGTGGCTTTGTGGG | Del_k31_29-59;Del_k31_30-60 | -3.6286019602050548 | 0.09252751099836234 | 1.3037660978429642 |
| GATCCAGATCTAACTTGGGGTGGCTTTGTGGGCTTCTTCTTTTGCCAATTCCACTAATTGTTTGGG | Ins_k3_29_GGG | -2.376687419963839 | 0.11748402636676522 | 4.5593114523648035 |
| GATCCAGATCTAACTTGGGGTGGCTTTGTGGTCTTCTTCTTTTGCCAATTCCACTAATTGTTTGGG | Ins_k3_27_GTG;Ins_k3_28_TGG; | 0.07794661549468507 | 0.0342273072148322 | 53.08028880010327 |
| GATCCAGATCTAACTTGGGGTGGCTTTGTGTACTTCTTCTTTTGCCAATTCCACTAATTGTTTGGG | Ins_k3_29_GTA | -0.15025209207552193 | 0.040554465612355435 | 42.25010938208638 |
| GATCCAGATCTAACTTGGGGTGGCTTTGTGTCCTTCTTCTTTTGCCAATTCCACTAATTGTTTGGG | Ins_k3_29_GTC | 0.20757574467176454 | 0.03381349236354171 | 60.42692474109082 |
| GATCCAGATCTAACTTGGGGTGGCTTTGTGTCTTCTTCTTTTGCCAATTCCACTAATTGTTTGGG | Ins_k2_26_TG;Ins_k2_27_GT;Ins_ | 0.23169859137304594 | 0.03255196829462347 | 61.90231798069571 |
| GATCCAGATCTAACTTGGGGTGGCTTTGTGTGCTTCTTCTTTTGCCAATTCCACTAATTGTTTGGG | Ins_k3_29_GTG | 0.34724571839009744 | 0.03309813262663055 | 69.4845727555606 |
| GATCCAGATCTAACTTGGGGTGGCTTTGTGTTCTTCTTCTTTTGCCAATTCCACTAATTGTTTGGG | Ins_k3_28_TGT;Ins_k3_29_GTT | -0.07271615723369307 | 0.036029541953857616 | 45.656358142243604 |
| GATCCAGATCTAACTTGGGGTGGCTTTGTGTTCTTCTTTTGCCAATTCCACTAATTGTTTGGG | C30G | -0.2582666141813777 | 0.033605870583178925 | 37.924313388672644 |
| GATCCAGATCTAACTTGGGGTGGCTTTGTGTTTGGG | Del_k27_29-55;Del_k27_30-56 | -3.3544393934123273 | 0.08616125673709982 | 1.7150112517578793 |
| GATCCAGATCTAACTTGGGGTGGCTTTGTTAACTTCTTCTTTTGCCAATTCCACTAATTGTTTGGG | Ins_k3_29_TAA | -1.3613557701697852 | 0.07024104607389318 | 12.584970249641486 |
| GATCCAGATCTAACTTGGGGTGGCTTTGTTAATTGTTTGGG | Del_k22_30-51 | -4.21718021673888 | 0.14401125127028805 | 0.7237413643798059 |
| GATCCAGATCTAACTTGGGGTGGCTTTGTTACCTTCTTCTTTTGCCAATTCCACTAATTGTTTGGG | Ins_k3_29_TAC | -0.19100374839152567 | 0.03673942359342187 | 40.56295803209051 |
| GATCCAGATCTAACTTGGGGTGGCTTTGTTACTTCTTCTTTTGCCAATTCCACTAATTGTTTGGG | Ins_k2_29_TA | -0.06798573218361581 | 0.0468879923520107 | 45.87284375328372 |
| GATCCAGATCTAACTTGGGGTGGCTTTGTTAGCTTCTTCTTTTGCCAATTCCACTAATTGTTTGGG | Ins_k3_29_TAG | -1.630975890154285 | 0.0735702581337922 | 9.610758467120283 |
| GATCCAGATCTAACTTGGGGTGGCTTTGTTATCTTCTTCTTTTGCCAATTCCACTAATTGTTTGGG | Ins_k3_28_TTA;Ins_k3_29_TAT | -0.5765758916436905 | 0.045261836611086674 | 27.585303338255304 |
| GATCCAGATCTAACTTGGGGTGGCTTTGTTCACTTCTTCTTTTGCCAATTCCACTAATTGTTTGGG | Ins_k3_29_TCA | -0.34913239820638287 | 0.047935620223138464 | 34.63021741404762 |
| GATCCAGATCTAACTTGGGGTGGCTTTGTTCCACTAATTGTTTGGG | Del_k17_29-45;Del_k17_30-46 | -1.7530691259006983 | 0.04600769649125255 | 8.506154062648449 |
| GATCCAGATCTAACTTGGGGTGGCTTTGTTCCCTTCTTCTTTTGCCAATTCCACTAATTGTTTGGG | Ins_k3_29_TCC | -0.22782575068250038 | 0.038026831356725514 | 39.096513105140154 |
| GATCCAGATCTAACTTGGGGTGGCTTTGTTCCTTCTTCTTTTGCCAATTCCACTAATTGTTTGGG | Ins_k2_29_TC | 0.2328215260529315 | 0.036694790365242154 | 61.9718692837981 |
| GATCCAGATCTAACTTGGGGTGGCTTTGTTCGCTTCTTCTTTTGCCAATTCCACTAATTGTTTGGG | Ins_k3_29_TCG | 0.3175578186008694 | 0.03409566334680174 | 67.45204179282737 |
| GATCCAGATCTAACTTGGGGTGGCTTTGTTCGGCTTCTTCTTTTGCCAATTCCACTAATTGTTTGGG | Ins_k4_29_TCGG | 0.8115001355570186 | 0.03318542125192815 | 100 |
| GATCCAGATCTAACTTGGGGTGGCTTTGTTCGTCTTCTTCTTTTGCCAATTCCACTAATTGTTTGGG | Ins_k4_27_GTTC;Ins_k4_28_TTC | 0.7193201624135384 | 0.03135383924992616 | 100 |
| GATCCAGATCTAACTTGGGGTGGCTTTGTTCTCTTCTTCTTTTGCCAATTCCACTAATTGTTTGGG | Ins_k3_28_TTC;Ins_k3_29_TCT | -0.9852198683134141 | 0.04706745179662408 | 18.331835009812707 |
| GATCCAGATCTAACTTGGGGTGGCTTTGTTCTTCTTCTTTTGCCAATTCCACTAATTGTTTGGG | Ins_k1_28_T;Ins_k1_29_T | -0.057714255513458546 | 0.03695109975000746 | 46.34645377087086 |
| GATCCAGATCTAACTTGGGGTGGCTTTGTTCTTCTTTTGCCAATTCCACTAATTGTTTGGG | Del_k2_29-30;Del_k2_30-31 | 0.021079728919633278 | 0.031111148751893927 | 50.14600066248209 |
| GATCCAGATCTAACTTGGGGTGGCTTTGTTCTTTTGCCAATTCCACTAATTGTTTGGG | Del_k5_29-33;Del_k5_30-34 | -0.9144801598653043 | 0.04110243786691698 | 19.675591845330267 |
| GATCCAGATCTAACTTGGGGTGGCTTTGTTGAACTTCTTCTTTTGCCAATTCCACTAATTGTTTGGG | Ins_k4_29_TGAA | 0.5492203463054628 | 0.030659716558025688 | 85.03639838329379 |
| GATCCAGATCTAACTTGGGGTGGCTTTGTTGACTTCTTCTTTTGCCAATTCCACTAATTGTTTGGG | Ins_k3_29_TGA | 0.4598149751493983 | 0.032837612529509404 | 77.76364312560753 |

| | | | | |
|---|---|---|---|---|
| **GATCCAGATCTAACTTGGGGTGGCTTTGTTGAGCTTCTTCTTTTGCCAATTCCACTAATTGTTTGGG** | Ins_k4_29_TGAG | 0.6516037285401639 | 0.03141077212326453 | 94.2040112870371 |
| **GATCCAGATCTAACTTGGGGTGGCTTTGTTGCCAATTCCACTAATTGTTTGGG** | Del_k10_29-38;Del_k10_30-39 | -0.5118472440595095 | 0.038194085950316564 | 29.429918434738063 |
| **GATCCAGATCTAACTTGGGGTGGCTTTGTTGCCTTCTTCTTTTGCCAATTCCACTAATTGTTTGGG** | Ins_k3_29_TGC | 0.33941027084452213 | 0.03181849378448905 | 68.94225744666913 |
| **GATCCAGATCTAACTTGGGGTGGCTTTGTTGCTTCTTCTTTTGCCAATTCCACTAATTGTTTGGG** | Ins_k2_29_TG | -0.10584260408547402 | 0.439515643230369 | 44.16870164663958 |
| **GATCCAGATCTAACTTGGGGTGGCTTTGTTGGCTTCTTCTTTTGCCAATTCCACTAATTGTTTGGG** | Ins_k3_29_TGG | -0.20826800505683707 | 0.03626185419376286 | 39.86867906301195 |
| **GATCCAGATCTAACTTGGGGTGGCTTTGTTGGG** | Del_k30_29-58;Del_k30_30-59 | -3.212438575748115 | 0.08037500154520538 | 1.9766835370170766 |
| **GATCCAGATCTAACTTGGGGTGGCTTTGTTGTCTTCTTCTTTTGCCAATTCCACTAATTGTTTGGG** | Ins_k3_25_TTG;Ins_k3_26_TGT;I | -0.1489067157946239 | 0.03610077223105685 | 42.306989931405674 |
| **GATCCAGATCTAACTTGGGGTGGCTTTGTTGTTTGGG** | Del_k26_29-54;Del_k26_30-55 | -3.1767837205280163 | 0.09513832408599104 | 2.048433417186423 |
| **GATCCAGATCTAACTTGGGGTGGCTTTGTTTACTTCTTCTTTTGCCAATTCCACTAATTGTTTGGG** | Ins_k3_29_TTA | -0.865266860616382 | 0.0574068710416762 | 20.668114992094104 |
| **GATCCAGATCTAACTTGGGGTGGCTTTGTTTCCACTAATTGTTTGGG** | Del_k16_30-45 | -1.3194530699991613 | 0.044766305063334705 | 13.123519010309325 |
| **GATCCAGATCTAACTTGGGGTGGCTTTGTTTCCTTCTTCTTTTGCCAATTCCACTAATTGTTTGGG** | Ins_k3_29_TTC | -0.6252565232438759 | 0.04498957025068734 | 26.274595231319203 |
| **GATCCAGATCTAACTTGGGGTGGCTTTGTTTCTTCTTCTTTTGCCAATTCCACTAATTGTTTGGG** | Ins_k2_28_TT;Ins_k2_29_TT | -0.6277386738522248 | 0.051998862357691404 | 26.209458601686716 |
| **GATCCAGATCTAACTTGGGGTGGCTTTGTTTCTTCTTTTGCCAATTCCACTAATTGTTTGGG** | Del_k1_30-30 | -0.5109162199690834 | 0.0388960740669259 | 29.457331156752247 |
| **GATCCAGATCTAACTTGGGGTGGCTTTGTTTCTTTTGCCAATTCCACTAATTGTTTGGG** | Del_k4_30-33 | -1.4400782640604586 | 0.06023628007125542 | 11.632242529133407 |
| **GATCCAGATCTAACTTGGGGTGGCTTTGTTTGCCAATTCCACTAATTGTTTGGG** | Del_k9_29-37;Del_k9_30-38 | -0.42224623573153514 | 0.035499418372989006 | 32.18861437130047 |
| **GATCCAGATCTAACTTGGGGTGGCTTTGTTTGCTTCTTCTTTTGCCAATTCCACTAATTGTTTGGG** | Ins_k3_29_TTG | -0.37949731885578597 | 0.040288920511757156 | 33.59447825934212 |
| **GATCCAGATCTAACTTGGGGTGGCTTTGTTTGGG** | Del_k29_26-54;Del_k29_27-55;D | -2.9918397220535975 | 0.06372234754073908 | 2.4645748371169685 |
| **GATCCAGATCTAACTTGGGGTGGCTTTGTTTGTTTGGG** | Del_k25_30-54 | -3.767742564562003 | 0.12204295449571684 | 1.1344142860914064 |
| **GATCCAGATCTAACTTGGGGTGGCTTTGTTTTCTTCTTCTTTTGCCAATTCCACTAATTGTTTGGG** | Ins_k3_28_TTT;Ins_k3_29_TTT | -1.4477389972096635 | 0.07340497800302923 | 11.543471482979399 |
| **GATCCAGATCTAACTTGGGGTGGCTTTGTTTTCTTCTTTTGCCAATTCCACTAATTGTTTGGG** | C30T | -1.088792979728046 | 0.04708458649185452 | 16.52816763931553 |
| **GATCCAGATCTAACTTGGGGTGGCTTTGTTTTGCCAATTCCACTAATTGTTTGGG** | Del_k8_29-36;Del_k8_30-37 | -0.549708242059967 | 0.038066513327347104 | 28.336501883176645 |
| **GATCCAGATCTAACTTGGGGTGGCTTTGTTTTGGG** | Del_k28_30-57 | -3.4767618711707517 | 0.11076675098131192 | 1.5175499564339494 |
| **GATCCAGATCTAACTTGGGGTGGCTTTGTTTTTGCCAATTCCACTAATTGTTTGGG** | Del_k7_30-36 | -0.6717211462248102 | 0.04176483626665732 | 25.081684750682584 |
| **GATCCAGATCTAACTTGGGGTGGCTTTTAACGTCTTCTTCTTTTGCCAATTCCACTAATTGTTTGGG** | Ins_k4_27_TAAC | 0.05948177097295704 | 0.03656842177241679 | 52.10916295609661 |
| **GATCCAGATCTAACTTGGGGTGGCTTTTAAGTCTTCTTCTTTTGCCAATTCCACTAATTGTTTGGG** | Ins_k3_27_TAA | -0.5789962830185487 | 0.048617491704507486 | 27.51861684424576 |
| **GATCCAGATCTAACTTGGGGTGGCTTTTAATTGTTTGGG** | Del_k24_28-51 | -3.7833342011315954 | 0.14775667745377782 | 1.1168640845405888 |
| **GATCCAGATCTAACTTGGGGTGGCTTTTACGTCTTCTTCTTTTGCCAATTCCACTAATTGTTTGGG** | Ins_k3_27_TAC | 0.3527121820149586 | 0.03461442260539298 | 69.8654477161279 |
| **GATCCAGATCTAACTTGGGGTGGCTTTTAGGTCTTCTTCTTTTGCCAATTCCACTAATTGTTTGGG** | Ins_k3_27_TAG | -2.8396896537076386 | 0.13796079283400525 | 2.8695906274910676 |
| **GATCCAGATCTAACTTGGGGTGGCTTTTAGTCTTCTTCTTTTGCCAATTCCACTAATTGTTTGGG** | Ins_k2_27_TA | -1.3343858791894898 | 0.06544295088516346 | 12.92900394772296 |
| **GATCCAGATCTAACTTGGGGTGGCTTTTATGTCTTCTTCTTTTGCCAATTCCACTAATTGTTTGGG** | Ins_k3_26_TTA;Ins_k3_27_TAT | -0.5225546358516832 | 0.04727407476425474 | 29.116481806609375 |
| **GATCCAGATCTAACTTGGGGTGGCTTTTCAGTCTTCTTCTTTTGCCAATTCCACTAATTGTTTGGG** | Ins_k3_27_TCA | 0.0416259582778542 | 0.03846569864448495 | 51.186969263368276 |
| **GATCCAGATCTAACTTGGGGTGGCTTTTCCACTAATTGTTTGGG** | Del_k19_27-45;Del_k19_28-46 | -0.9505549690254496 | 0.03801300996949914 | 18.97844887720525 |
| **GATCCAGATCTAACTTGGGGTGGCTTTTCCGTCTTCTTCTTTTGCCAATTCCACTAATTGTTTGGG** | Ins_k3_27_TCC | 0.4334457353283885 | 0.030857132203649683 | 75.73987485150936 |
| **GATCCAGATCTAACTTGGGGTGGCTTTTCGGTCTTCTTCTTTTGCCAATTCCACTAATTGTTTGGG** | Ins_k3_27_TCG | -0.051131300876524066 | 0.48571260987248394 | 46.652556799381536 |
| **GATCCAGATCTAACTTGGGGTGGCTTTTCGTCTTCTTCTTTTGCCAATTCCACTAATTGTTTGGG** | Ins_k2_27_TC | 0.6405616440057871 | 0.029446287999825124 | 93.16952458674675 |
| **GATCCAGATCTAACTTGGGGTGGCTTTTCTGTCTTCTTCTTTTGCCAATTCCACTAATTGTTTGGG** | Ins_k3_26_TTC;Ins_k3_27_TCT | -1.9507689065540426 | 0.0825731617715627 | 6.980287662003062 |
| **GATCCAGATCTAACTTGGGGTGGCTTTTCTTCTTCTTTTGCCAATTCCACTAATTGTTTGGG** | Del_k1_28-28 | -0.431630195731993 | 0.04173720080773678 | 31.887970522684018 |
| **GATCCAGATCTAACTTGGGGTGGCTTTTCTTCTTTTGCCAATTCCACTAATTGTTTGGG** | Del_k4_27-30;Del_k4_28-31 | -0.3096261693050923 | 0.03996040820615867 | 36.02571055005021 |
| **GATCCAGATCTAACTTGGGGTGGCTTTTCTTTTGCCAATTCCACTAATTGTTTGGG** | Del_k7_27-33;Del_k7_28-34 | -0.809769397238291 | 0.04592183350382277 | 21.847568580341314 |
| **GATCCAGATCTAACTTGGGGTGGCTTTTGAGTCTTCTTCTTTTGCCAATTCCACTAATTGTTTGGG** | Ins_k3_27_TGA | 0.2397031496998644 | 0.03440359481541207 | 62.39980712969541 |
| **GATCCAGATCTAACTTGGGGTGGCTTTTGCCAATTCCACTAATTGTTTGGG** | Del_k12_24-35;Del_k12_25-36;D | -0.8326083419044332 | 0.034623102486620726 | 21.35424807384704 |
| **GATCCAGATCTAACTTGGGGTGGCTTTTGCGTCTTCTTCTTTTGCCAATTCCACTAATTGTTTGGG** | Ins_k3_27_TGC | 0.4120762121732351 | 0.03448252775081062 | 74.13852086623329 |
| **GATCCAGATCTAACTTGGGGTGGCTTTTGGG** | Del_k32_26-57;Del_k32_27-58;D | -4.12989003282431 | 0.11564366427090074 | 0.7897561926720851 |
| **GATCCAGATCTAACTTGGGGTGGCTTTTGGGTCTTCTTCTTTTGCCAATTCCACTAATTGTTTGGG** | Ins_k3_27_TGG | -2.763181345107455 | 0.11547561462172486 | 3.097755105649325 |
| **GATCCAGATCTAACTTGGGGTGGCTTTTGGTCTTCTTCTTTTGCCAATTCCACTAATTGTTTGGG** | Ins_k2_27_TG | 0.07637099636094002 | 0.04329656291178668 | 52.996720334770345 |
| **GATCCAGATCTAACTTGGGGTGGCTTTTGTAGTCTTCTTCTTTTGCCAATTCCACTAATTGTTTGGG** | Ins_k4_27_TGTA | -0.9828802157746711 | 0.06320023956256207 | 18.374775347280163 |
| **GATCCAGATCTAACTTGGGGTGGCTTTTGTCTTCTTCTTTTGCCAATTCCACTAATTGTTTGGG** | Ins_k1_24_T;Ins_k1_25_T;Ins_k1_ | -0.12166403891425331 | 0.04595956442095883 | 43.47538847894006 |
| **GATCCAGATCTAACTTGGGGTGGCTTTTGTGTCTTCTTCTTTTGCCAATTCCACTAATTGTTTGGG** | Ins_k3_26_TTG;Ins_k3_27_TGT | -0.07101961633199172 | 0.03878296253418686 | 45.733881763657564 |
| **GATCCAGATCTAACTTGGGGTGGCTTTTGTTTGGG** | Del_k28_27-54;Del_k28_28-55 | -3.6590042729432604 | 0.09320391919216185 | 1.2647250690729497 |
| **GATCCAGATCTAACTTGGGGTGGCTTTTTAGTCTTCTTCTTTTGCCAATTCCACTAATTGTTTGGG** | Ins_k3_27_TTA | -1.7720661547391856 | 0.08363638494639886 | 8.346087615865924 |
| **GATCCAGATCTAACTTGGGGTGGCTTTTTCCACTAATTGTTTGGG** | Del_k18_28-45 | -1.344208552520707 | 0.04635039752405137 | 12.802628255129793 |
| **GATCCAGATCTAACTTGGGGTGGCTTTTTCGTCTTCTTCTTTTGCCAATTCCACTAATTGTTTGGG** | Ins_k3_27_TTC | 0.6283112842672074 | 0.03415492896070911 | 92.0351269700087 |
| **GATCCAGATCTAACTTGGGGTGGCTTTTTCTTCTTCTTTTGCCAATTCCACTAATTGTTTGGG** | G28T | -1.5605111427310088 | 0.04608761079096197 | 10.312408635869101 |
| **GATCCAGATCTAACTTGGGGTGGCTTTTTCTTCTTTTGCCAATTCCACTAATTGTTTGGG** | Del_k3_28-30 | -0.8548224630150371 | 0.05110524224320392 | 20.885112232872668 |
| **GATCCAGATCTAACTTGGGGTGGCTTTTTCTTTTGCCAATTCCACTAATTGTTTGGG** | Del_k6_28-33 | -1.596232734608896 | 0.06122141146391978 | 9.95053481713254 |
| **GATCCAGATCTAACTTGGGGTGGCTTTTTGCCAATTCCACTAATTGTTTGGG** | Del_k11_26-36;Del_k11_27-37;D | -0.762336918526986 | 0.03911159313119689 | 22.908822910043035 |
| **GATCCAGATCTAACTTGGGGTGGCTTTTTGGG** | Del_k31_27-57;Del_k31_28-58 | -3.2685292869161473 | 0.08585221273913601 | 1.8688621091658717 |
| **GATCCAGATCTAACTTGGGGTGGCTTTTTGGTCTTCTTCTTTTGCCAATTCCACTAATTGTTTGGG** | Ins_k3_27_TTG | -0.20389066004719802 | 0.040114699962337644 | 40.04358054914705 |
| **GATCCAGATCTAACTTGGGGTGGCTTTTTGTCTTCTTCTTTTGCCAATTCCACTAATTGTTTGGG** | Ins_k2_24_TT;Ins_k2_25_TT;Ins_ | -0.3844950673673706 | 0.052952547334821916 | 33.42700036043202 |
| **GATCCAGATCTAACTTGGGGTGGCTTTTTGTTTGGG** | Del_k27_28-54 | -3.791784537856195 | 0.1211873657613687 | 1.1074659714856752 |
| **GATCCAGATCTAACTTGGGGTGGCTTTTTTGCCAATTCCACTAATTGTTTGGG** | Del_k10_27-36;Del_k10_28-37 | -0.9445804707881997 | 0.1763169038353242 | 19.092174976473608 |
| **GATCCAGATCTAACTTGGGGTGGCTTTTTTGGG** | Del_k30_28-57 | -3.3465490467761834 | 0.08187986090220818 | 1.728596811925028 |
| **GATCCAGATCTAACTTGGGGTGGCTTTTTTGTCTTCTTCTTTTGCCAATTCCACTAATTGTTTGGG** | Ins_k3_24_TTT;Ins_k3_25_TTT;In | -1.2556174858997897 | 0.06580454476720099 | 13.988583727216684 |
| **GATCCAGATCTAACTTGGGGTGGCTTTTTTTGCCAATTCCACTAATTGTTTGGG** | Del_k9_28-36 | -0.7319792794112342 | 0.047155238907352334 | 23.614944553017096 |
| **GATCCAGATCTAACTTGGGGTGGG** | Del_k39_21-59;Del_k39_22-60;D | -5.711457402942378 | 0.5763713588361297 | 0.16241534478239533 |
| **GATCCAGATCTAACTTGGGGTGGGAACTTTGTCTTCTTCTTTTGCCAATTCCACTAATTGTTTGGG** | Ins_k3_23_GAA | -2.726198439482753 | 0.12201590587999557 | 3.2144639036760574 |
| **GATCCAGATCTAACTTGGGGTGGGACCTTTGTCTTCTTCTTTTGCCAATTCCACTAATTGTTTGGG** | Ins_k3_23_GAC | -0.5526873939947858 | 0.04347404066529694 | 28.25220876211694 |
| **GATCCAGATCTAACTTGGGGTGGGACTCTTTGTCTTCTTCTTTTGCCAATTCCACTAATTGTTTGGG** | Ins_k4_23_GACT | -2.1376822171397403 | 0.09457156003402781 | 5.79025781171869 |
| **GATCCAGATCTAACTTGGGGTGGGACTTTGTCTTCTTCTTTTGCCAATTCCACTAATTGTTTGGG** | Ins_k2_23_GA | -1.7184695168128874 | 0.08134134179515924 | 8.805614399462009 |
| **GATCCAGATCTAACTTGGGGTGGGAGCTTTGTCTTCTTCTTTTGCCAATTCCACTAATTGTTTGGG** | Ins_k3_22_GGA;Ins_k3_23_GAG | -1.1515247717563213 | 0.053900027078861004 | 15.523178018782387 |
| **GATCCAGATCTAACTTGGGGTGGGATCTTTGTCTTCTTCTTTTGCCAATTCCACTAATTGTTTGGG** | Ins_k3_23_GAT | -2.6079293962349337 | 0.13154957424690755 | 3.618029854384613 |
| **GATCCAGATCTAACTTGGGGTGGGCACTTTGTCTTCTTCTTTTGCCAATTCCACTAATTGTTTGGG** | Ins_k3_23_GCA | -3.4899773460908716 | 0.1697447509925307 | 1.497626750322801 |
| **GATCCAGATCTAACTTGGGGTGGGCATCTTTGTCTTCTTCTTTTGCCAATTCCACTAATTGTTTGGG** | Ins_k4_23_GCAT | -3.087974594047784 | 0.15669859540734968 | 2.238675599424549 |
| **GATCCAGATCTAACTTGGGGTGGGCCAATTCCACTAATTGTTTGGG** | Del_k17_24-40 | -2.1279743866433636 | 0.05420197721261155 | 5.8467423807955665 |
| **GATCCAGATCTAACTTGGGGTGGGCCCTTTGTCTTCTTCTTTTGCCAATTCCACTAATTGTTTGGG** | Ins_k3_23_GCC | -1.8740356039002948 | 0.08282630731173836 | 7.536994015917168 |

| | | | | |
|---|---|---|---|---|
| **GATCCAGATCTAACTTGGGGTGGGCCTTTGTCTTCTTCTTTTGCCAATTCCACTAATTGTTTGGG** | Ins_k2_23_GC | -2.895354882674524 | 0.14146357360325998 | 2.7142187320095905 |
| **GATCCAGATCTAACTTGGGGTGGGCGCTTTGTCTTCTTCTTTTGCCAATTCCACTAATTGTTTGGG** | Ins_k3_22_GGC;Ins_k3_23_GCG | -1.3368599901432647 | 0.06463559982648456 | 12.897055695491769 |
| **GATCCAGATCTAACTTGGGGTGGGCTCTTTGTCTTCTTCTTTTGCCAATTCCACTAATTGTTTGGG** | Ins_k3_23_GCT | -5.348918222486026 | 0.5289172060874984 | 0.23338654988261912 |
| **GATCCAGATCTAACTTGGGGTGGGCTTTGTCTTCTTCTTTTGCCAATTCCACTAATTGTTTGGG** | Ins_k1_21_G;Ins_k1_22_G;Ins_k1 | -3.9732707538946057 | 0.20297771451480562 | 0.9236595578719172 |
| **GATCCAGATCTAACTTGGGGTGGGGACTTTGTCTTCTTCTTTTGCCAATTCCACTAATTGTTTGGG** | Ins_k3_23_GGA | -3.1468529514442065 | 0.16577867496826235 | 2.1106713734704274 |
| **GATCCAGATCTAACTTGGGGTGGGGCCTTTGTCTTCTTCTTTTGCCAATTCCACTAATTGTTTGGG** | Ins_k3_23_GGC | -3.287705840719585 | 0.1818279441489726 | 1.833365216213282 |
| **GATCCAGATCTAACTTGGGGTGGGGCTTTGTCTTCTTCTTTTGCCAATTCCACTAATTGTTTGGG** | Ins_k2_21_GG;Ins_k2_22_GG;Ins | -4.234600114428469 | 0.3816395023340766 | 0.7112430396801069 |
| **GATCCAGATCTAACTTGGGGTGGGGGCTTTGTCTTCTTCTTTTGCCAATTCCACTAATTGTTTGGG** | Ins_k3_21_GGG;Ins_k3_22_GGG | -4.64028343462736 | 0.3655291408173317 | 0.4740585698544199 |
| **GATCCAGATCTAACTTGGGGTGGGGTCTTTGTCTTCTTCTTTTGCCAATTCCACTAATTGTTTGGG** | Ins_k3_23_GGT | -4.85669392596675 | 0.5120850924908369 | 0.38180896344215987 |
| **GATCCAGATCTAACTTGGGGTGGGTACTTTGTCTTCTTCTTTTGCCAATTCCACTAATTGTTTGGG** | Ins_k3_23_GTA | -5.381917036689779 | 0.5743160452388083 | 0.22581075406626913 |
| **GATCCAGATCTAACTTGGGGTGGGTCACTTTGTCTTCTTCTTTTGCCAATTCCACTAATTGTTTGGG** | Ins_k4_23_GTCA | -3.9921223479941426 | 0.3899625276240798 | 0.9064102025635038 |
| **GATCCAGATCTAACTTGGGGTGGGTCCTTTGTCTTCTTCTTTTGCCAATTCCACTAATTGTTTGGG** | Ins_k3_23_GTC | -4.784383381201219 | 0.35046979739964146 | 0.41044048332925065 |
| **GATCCAGATCTAACTTGGGGTGGGTCTTCTTCTTTTGCCAATTCCACTAATTGTTTGGG** | Del_k4_24-27 | -4.333881181371789 | 0.3043269668851997 | 0.6440221580425526 |
| **GATCCAGATCTAACTTGGGGTGGGTCTTTGTCTTCTTCTTTTGCCAATTCCACTAATTGTTTGGG** | Ins_k2_23_GT | -4.5297022568862015 | 0.4645889786322448 | 0.5294888233551551 |
| **GATCCAGATCTAACTTGGGGTGGGTGCTTTGTCTTCTTCTTTTGCCAATTCCACTAATTGTTTGGG** | Ins_k3_22_GGT;Ins_k3_23_GTG | -3.8038231319065456 | 0.2680499656966805 | 1.0942135684336718 |
| **GATCCAGATCTAACTTGGGGTGGGTTCTTTGTCTTCTTCTTTTGCCAATTCCACTAATTGTTTGGG** | Ins_k3_23_GTT | -5.50260308950771 | 0.663554476224667 | 0.20013881970036443 |
| **GATCCAGATCTAACTTGGGGTGGGTTTGGG** | Del_k33_24-56 | -6.518568689069914 | 0.6091416721828643 | 0.0724607926620838 |
| **GATCCAGATCTAACTTGGGGTGGGTTTGTCTTCTTCTTTTGCCAATTCCACTAATTGTTTGGG** | C24G | -3.933970630876591 | 0.18869386678304076 | 0.9606822248034016 |
| **GATCCAGATCTAACTTGGGGTGGTAACTTTGTCTTCTTCTTTTGCCAATTCCACTAATTGTTTGGG** | Ins_k3_23_TAA | -1.4803506822737562 | 0.05743337241965026 | 11.17309160737258 |
| **GATCCAGATCTAACTTGGGGTGGTAATTGTTTGGG** | Del_k28_24-51 | -5.167291635437418 | 0.21149641645325085 | 0.2798692915671452 |
| **GATCCAGATCTAACTTGGGGTGGTACCTTTGTCTTCTTCTTTTGCCAATTCCACTAATTGTTTGGG** | Ins_k3_23_TAC | -0.016298075805070256 | 0.03831504822650235 | 48.30625034405453 |
| **GATCCAGATCTAACTTGGGGTGGTACTTTGTCTTCTTCTTTTGCCAATTCCACTAATTGTTTGGG** | Ins_k2_23_TA | -0.5815834193866988 | 0.0530905605692399 | 27.44751444526694 |
| **GATCCAGATCTAACTTGGGGTGGTAGCTTTGTCTTCTTCTTTTGCCAATTCCACTAATTGTTTGGG** | Ins_k3_22_GTA;Ins_k3_23_TAG | -1.4569120519276906 | 0.06618282461343139 | 11.438066768143425 |
| **GATCCAGATCTAACTTGGGGTGGTATCTTTGTCTTCTTCTTTTGCCAATTCCACTAATTGTTTGGG** | Ins_k3_23_TAT | -1.770020960961374 | 0.08300486153398819 | 8.363174449313764 |
| **GATCCAGATCTAACTTGGGGTGGTCACTTTGTCTTCTTCTTTTGCCAATTCCACTAATTGTTTGGG** | Ins_k3_23_TCA | -0.05574868329709237 | 0.037753250895638475 | 46.43764066058026 |
| **GATCCAGATCTAACTTGGGGTGGTCCACTAATTGTTTGGG** | Del_k23_24-46 | -1.2479335154490374 | 0.038822056205672674 | 14.096485617786481 |
| **GATCCAGATCTAACTTGGGGTGGTCCCTTTGTCTTCTTCTTTTGCCAATTCCACTAATTGTTTGGG** | Ins_k3_23_TCC | 4.5208346511971254e-4 | 0.03566967894648709 | 49.12220231640931 |
| **GATCCAGATCTAACTTGGGGTGGTCCTTTGTCTTCTTCTTTTGCCAATTCCACTAATTGTTTGGG** | Ins_k2_23_TC | 0.35558022551019475 | 0.04215541310823836 | 70.06611247907573 |
| **GATCCAGATCTAACTTGGGGTGGTCGCTTTGTCTTCTTCTTTTGCCAATTCCACTAATTGTTTGGG** | Ins_k3_22_GTC;Ins_k3_23_TCG | 0.2975627558399505 | 0.03300135283106558 | 66.1167283104503 |
| **GATCCAGATCTAACTTGGGGTGGTCTCTTTGTCTTCTTCTTTTGCCAATTCCACTAATTGTTTGGG** | Ins_k3_23_TCT | -1.0561792848674947 | 0.051612680923153874 | 17.076098717277674 |
| **GATCCAGATCTAACTTGGGGTGGTCTTCTTCTTTTGCCAATTCCACTAATTGTTTGGG** | Del_k5_23-27;Del_k5_24-28 | 0.06036762244676641 | 0.02944222511553049 | 52.15534438682446 |
| **GATCCAGATCTAACTTGGGGTGGTCTTCTTTTGCCAATTCCACTAATTGTTTGGG** | Del_k8_24-31 | 0.15401172814252895 | 0.032023996942797894 | 57.27537427063514 |
| **GATCCAGATCTAACTTGGGGTGGTCTTTGTCTTCTTCTTTTGCCAATTCCACTAATTGTTTGGG** | Ins_k1_23_T | -0.3811587334160096 | 0.04177332793367938 | 33.538710243743516 |
| **GATCCAGATCTAACTTGGGGTGGTCTTTTGCCAATTCCACTAATTGTTTGGG** | Del_k11_24-34 | -0.43126006901527486 | 0.03495690994738079 | 31.899775297012802 |
| **GATCCAGATCTAACTTGGGGTGGTGACTTTGTCTTCTTCTTTTGCCAATTCCACTAATTGTTTGGG** | Ins_k3_23_TGA | 0.5524904573211156 | 0.0313214913796472 | 85.31493201611772 |
| **GATCCAGATCTAACTTGGGGTGGTGCACTTTGTCTTCTTCTTTTGCCAATTCCACTAATTGTTTGGG** | Ins_k4_23_TGCA | 0.5584052312186145 | 0.033402744327856226 | 85.82104584800167 |
| **GATCCAGATCTAACTTGGGGTGGTGCCAATTCCACTAATTGTTTGGG** | Del_k16_24-39 | -0.166117610535923 | 0.029152356948773883 | 41.585078967657104 |
| **GATCCAGATCTAACTTGGGGTGGTGCCTTTGTCTTCTTCTTTTGCCAATTCCACTAATTGTTTGGG** | Ins_k3_23_TGC | 0.41293210674107805 | 0.033524138676435694 | 74.20200278655065 |
| **GATCCAGATCTAACTTGGGGTGGTGCTTTGTCTTCTTCTTTTGCCAATTCCACTAATTGTTTGGG** | Ins_k2_22_GT;Ins_k2_23_TG | 0.22425543370517875 | 0.035226880317890986 | 61.443279734228376 |
| **GATCCAGATCTAACTTGGGGTGGTGGCTTTGTCTTCTTCTTTTGCCAATTCCACTAATTGTTTGGG** | Ins_k3_18_GGT;Ins_k3_19_GTG; | -0.18167790539898504 | 0.03560289771834546 | 40.94301121368687 |
| **GATCCAGATCTAACTTGGGGTGGTGTCTTCTTCTTTTGCCAATTCCACTAATTGTTTGGG** | Del_k3_24-26 | 0.3668321862353934 | 0.0299334043352024 | 70.8589457243157 |
| **GATCCAGATCTAACTTGGGGTGGTGTCTTTGTCTTCTTCTTTTGCCAATTCCACTAATTGTTTGGG** | Ins_k3_23_TGT | -0.07790946840171564 | 0.03427718546105224 | 45.41986509014394 |
| **GATCCAGATCTAACTTGGGGTGGTGTTTGGG** | Del_k32_24-55 | -3.773130693648413 | 0.1167956855545367 | 1.1283183530655547 |
| **GATCCAGATCTAACTTGGGGTGGTTACTTTGTCTTCTTCTTTTGCCAATTCCACTAATTGTTTGGG** | Ins_k3_23_TTA | -1.2577859303626266 | 0.051021661872252974 | 13.958283124750455 |
| **GATCCAGATCTAACTTGGGGTGGTTCCACTAATTGTTTGGG** | Del_k22_24-45 | -1.5402123256775255 | 0.039364368724667906 | 10.523877353438827 |
| **GATCCAGATCTAACTTGGGGTGGTTCCTTTGTCTTCTTCTTTTGCCAATTCCACTAATTGTTTGGG** | Ins_k3_23_TTC | 0.028414204196122395 | 0.037428358886618364 | 50.51514735837522 |
| **GATCCAGATCTAACTTGGGGTGGTTCTTCTTTTGCCAATTCCACTAATTGTTTGGG** | Del_k7_24-30 | 0.007480833805995402 | 0.03171647624733931 | 49.46868626080254 |
| **GATCCAGATCTAACTTGGGGTGGTTCTTTGTCTTCTTCTTTTGCCAATTCCACTAATTGTTTGGG** | Ins_k2_23_TT | -1.122898637291984 | 0.06923118970390713 | 15.973968004322396 |
| **GATCCAGATCTAACTTGGGGTGGTTCTTTTGCCAATTCCACTAATTGTTTGGG** | Del_k10_24-33 | -0.7206446284232705 | 0.03822906473900309 | 23.884134412184892 |
| **GATCCAGATCTAACTTGGGGTGGTTGCCAATTCCACTAATTGTTTGGG** | Del_k15_24-38 | -0.5740625033620893 | 0.03257316424229003 | 27.654723119297756 |
| **GATCCAGATCTAACTTGGGGTGGTTGCTTTGTCTTCTTCTTTTGCCAATTCCACTAATTGTTTGGG** | Ins_k3_22_GTT;Ins_k3_23_TTG | -0.474321600376616 | 0.043189395532454614 | 30.555277931896168 |
| **GATCCAGATCTAACTTGGGGTGGTTGGG** | Del_k35_24-58 | -5.84808215019691 | 0.5895408737675388 | 0.1416744986881524 |
| **GATCCAGATCTAACTTGGGGTGGTTGTCTTCTTCTTTTGCCAATTCCACTAATTGTTTGGG** | Del_k2_24-25 | -0.0377125883575512 | 0.03755562288461561 | 47.282793070252616 |
| **GATCCAGATCTAACTTGGGGTGGTTGTTTGGG** | Del_k31_24-54 | -4.714235259983368 | 0.1982986504570231 | 0.4402659841928571 |
| **GATCCAGATCTAACTTGGGGTGGTTTCTTTGTCTTCTTCTTTTGCCAATTCCACTAATTGTTTGGG** | Ins_k3_23_TTT | -1.9773124600171623 | 0.08695608275236151 | 6.79744342640228 |
| **GATCCAGATCTAACTTGGGGTGGTTTGCCAATTCCACTAATTGTTTGGG** | Del_k14_24-37 | -0.5579132915042353 | 0.03196108247394369 | 28.104950727543258 |
| **GATCCAGATCTAACTTGGGGTGGTTTGGG** | Del_k34_23-56;Del_k34_24-57 | -5.173921287529607 | 0.30838648780707195 | 0.27801999241118236 |
| **GATCCAGATCTAACTTGGGGTGGTTTGTCTTCTTCTTTTGCCAATTCCACTAATTGTTTGGG** | Del_k1_24-24 | -0.4448040492239364 | 0.03706506762528567 | 31.470638044990018 |
| **GATCCAGATCTAACTTGGGGTGGTTTTGCCAATTCCACTAATTGTTTGGG** | Del_k13_24-36 | -0.4382248971985791 | 0.032902861359066955 | 31.67837076026737 |
| **GATCCAGATCTAACTTGGGGTGGTTTTGTCTTCTTCTTTTGCCAATTCCACTAATTGTTTGGG** | C24T | -0.9605307114979382 | 0.037439762883347674 | 18.79006595070763 |
| **GATCCAGATCTAACTTGGGGTGTAAGCTTTGTCTTCTTCTTTTGCCAATTCCACTAATTGTTTGGG** | Ins_k3_22_TAA | -0.5318771776050178 | 0.04244614992799554 | 28.846303520193587 |
| **GATCCAGATCTAACTTGGGGTGTAATTGTTTGGG** | Del_k29_23-51 | -3.1539027321097137 | 0.08498034795836787 | 2.095843929753061 |
| **GATCCAGATCTAACTTGGGGTGTACGCTTTGTCTTCTTCTTTTGCCAATTCCACTAATTGTTTGGG** | Ins_k3_22_TAC | 0.47008639904662974 | 0.03253950763882448 | 78.56650266466667 |
| **GATCCAGATCTAACTTGGGGTGTAGCTTTGTCTTCTTCTTTTGCCAATTCCACTAATTGTTTGGG** | Ins_k2_22_TA | -0.6990587394848995 | 0.04610253736887727 | 24.405299352991545 |
| **GATCCAGATCTAACTTGGGGTGTAGGCTTTGTCTTCTTCTTTTGCCAATTCCACTAATTGTTTGGG** | Ins_k3_21_GTA;Ins_k3_22_TAG | -1.5984230385039133 | 0.07263978301029324 | 9.928763973051042 |
| **GATCCAGATCTAACTTGGGGTGTATGCTTTGTCTTCTTCTTTTGCCAATTCCACTAATTGTTTGGG** | Ins_k3_22_TAT | -0.11996790188769829 | 0.035415930730791266 | 43.549191267210496 |
| **GATCCAGATCTAACTTGGGGTGTCAGCTTTGTCTTCTTCTTTTGCCAATTCCACTAATTGTTTGGG** | Ins_k3_22_TCA | 0.4655206242279728 | 0.027568735553794268 | 78.20860337074174 |
| **GATCCAGATCTAACTTGGGGTGTCCACTAATTGTTTGGG** | Del_k24_23-46 | -0.9910159220895085 | 0.0342434782847148 | 18.225890036258257 |
| **GATCCAGATCTAACTTGGGGTGTCCGCTTTGTCTTCTTCTTTTGCCAATTCCACTAATTGTTTGGG** | Ins_k3_22_TCC | 0.7586389597170007 | 0.029389127071249688 | 100 |
| **GATCCAGATCTAACTTGGGGTGTCGCTTTGTCTTCTTCTTTTGCCAATTCCACTAATTGTTTGGG** | Ins_k2_22_TC | 0.6489757071378877 | 0.03330388381684884 | 93.95676615424458 |
| **GATCCAGATCTAACTTGGGGTGTCGGCTTTGTCTTCTTCTTTTGCCAATTCCACTAATTGTTTGGG** | Ins_k3_21_GTC;Ins_k3_22_TCG | 0.6603713873455561 | 0.028454406986872155 | 95.03359133925395 |
| **GATCCAGATCTAACTTGGGGTGTCTGCTTTGTCTTCTTCTTTTGCCAATTCCACTAATTGTTTGGG** | Ins_k3_22_TCT | 0.377377362390538 | 0.0306300096546904 | 71.61011945790834 |
| **GATCCAGATCTAACTTGGGGTGTCTTCTTCTTTTGCCAATTCCACTAATTGTTTGGG** | Del_k6_21-26;Del_k6_22-27;Del_ | 0.030957373830593715 | 0.030022030786418123 | 50.64377944449601 |
| **GATCCAGATCTAACTTGGGGTGTCTTCTTTTGCCAATTCCACTAATTGTTTGGG** | Del_k9_23-31 | 0.24095775447990697 | 0.031433568546881174 | 62.47814335622086 |
| **GATCCAGATCTAACTTGGGGTGTCTTTGTCTTCTTCTTTTGCCAATTCCACTAATTGTTTGGG** | G23T | -0.8956073705009265 | 0.03414143710179448 | 20.050451341546704 |

| | | | | |
|---|---|---|---|---|
| **GATCCAGATCTAACTTGGGGTGTCTTTTGCCAATTCCACTAATTGTTTGGG** | Del_k12_23-34 | -0.1456224111007397 | 0.029089409611482035 | 42.446167402527394 |
| **GATCCAGATCTAACTTGGGGTGTGAGCTTTGTCTTCTTCTTTTGCCAATTCCACTAATTGTTTGGG** | Ins_k3_22_TGA | 0.4510421098777855 | 0.02892219218501011 | 77.08441689780727 |
| **GATCCAGATCTAACTTGGGGTGTGCCAATTCCACTAATTGTTTGGG** | Del_k17_23-39 | -0.08771447234304897 | 0.02608138244266818 | 44.976699305473694 |
| **GATCCAGATCTAACTTGGGGTGTGCGCTTTGTCTTCTTCTTTTGCCAATTCCACTAATTGTTTGGG** | Ins_k3_22_TGC | 0.5193857791295506 | 0.02959875165545616 | 82.53684617430022 |
| **GATCCAGATCTAACTTGGGGTGTGCTTTGTCTTCTTCTTTTGCCAATTCCACTAATTGTTTGGG** | Ins_k1_22_T | 0.471778362136555 | 0.034037936277944374 | 78.69954680843087 |
| **GATCCAGATCTAACTTGGGGTGTGGCTTTGTCTTCTTCTTTTGCCAATTCCACTAATTGTTTGGG** | Ins_k2_19_GT;Ins_k2_20_TG;Ins_ | 0.42864945812916155 | 0.03389307514687547 | 75.37747519601726 |
| **GATCCAGATCTAACTTGGGGTGTGGG** | Del_k37_23-59 | -5.806503776242154 | 0.31973926770689254 | 0.1476892696898553 |
| **GATCCAGATCTAACTTGGGGTGTGGGCTTTGTCTTCTTCTTTTGCCAATTCCACTAATTGTTTGGG** | Ins_k3_21_GTG;Ins_k3_22_TGG | -3.157131251973919 | 0.17402245134960334 | 2.0890883670956555 |
| **GATCCAGATCTAACTTGGGGTGTGTCTTCTTCTTTTGCCAATTCCACTAATTGTTTGGG** | Del_k4_23-26 | 0.15713763543598536 | 0.030042359389951947 | 57.45469189993202 |
| **GATCCAGATCTAACTTGGGGTGTGTGCTTTGTCTTCTTCTTTTGCCAATTCCACTAATTGTTTGGG** | Ins_k3_22_TGT | 0.012389585090766375 | 0.034302391222373524 | 49.71211270918998 |
| **GATCCAGATCTAACTTGGGGTGTGTTTGGG** | Del_k33_23-55 | -3.9214731532941465 | 0.09732854481074614 | 0.9727636658947971 |
| **GATCCAGATCTAACTTGGGGTGTTAGCTTTGTCTTCTTCTTTTGCCAATTCCACTAATTGTTTGGG** | Ins_k3_22_TTA | -1.4285248013411103 | 0.060617227025792884 | 11.767414558448499 |
| **GATCCAGATCTAACTTGGGGTGTTCCACTAATTGTTTGGG** | Del_k23_23-45 | -0.8425641807649071 | 0.03008917056344354 | 21.142703420610456 |
| **GATCCAGATCTAACTTGGGGTGTTCGCTTTGTCTTCTTCTTTTGCCAATTCCACTAATTGTTTGGG** | Ins_k3_22_TTC | 0.49891867020622294 | 0.030182075772577324 | 80.86472566069308 |
| **GATCCAGATCTAACTTGGGGTGTTCTTCTTTTGCCAATTCCACTAATTGTTTGGG** | Del_k8_23-30 | 0.2346235977176242 | 0.028187173726469113 | 62.083647719568695 |
| **GATCCAGATCTAACTTGGGGTGTTCTTTTGCCAATTCCACTAATTGTTTGGG** | Del_k11_23-33 | -0.2646983138096885 | 0.033164224336526785 | 37.68117832035298 |
| **GATCCAGATCTAACTTGGGGTGTTGAGCTTTGTCTTCTTCTTTTGCCAATTCCACTAATTGTTTGGG** | Ins_k4_22_TTGA | 0.6315208682999196 | 0.033357763318298296 | 92.33099599824523 |
| **GATCCAGATCTAACTTGGGGTGTTGCCAATTCCACTAATTGTTTGGG** | Del_k16_23-38 | -0.12812234140919004 | 0.026571286169043862 | 43.1955159924289 |
| **GATCCAGATCTAACTTGGGGTGTTGCTTTGTCTTCTTCTTTTGCCAATTCCACTAATTGTTTGGG** | Ins_k2_22_TT | 0.279464060148351 | 0.032274984880201356 | 64.93086542062534 |
| **GATCCAGATCTAACTTGGGGTGTTGGCTTTGTCTTCTTCTTTTGCCAATTCCACTAATTGTTTGGG** | Ins_k3_20_TGT;Ins_k3_21_GTT;Ir | 0.06047753162058078 | 0.033396213658946426 | 52.16107705266654 |
| **GATCCAGATCTAACTTGGGGTGTTGGG** | Del_k36_23-58 | -5.117553274161519 | 0.17154371946618627 | 0.29414152810507416 |
| **GATCCAGATCTAACTTGGGGTGTTGTCTTCTTCTTTTGCCAATTCCACTAATTGTTTGGG** | Del_k3_23-25 | 0.10461828037085763 | 0.028973939378607764 | 54.515077521093126 |
| **GATCCAGATCTAACTTGGGGTGTTGTTTGGG** | Del_k32_23-54 | -3.5080050786640755 | 0.08646863264318713 | 1.4708698436132128 |
| **GATCCAGATCTAACTTGGGGTGTTTGCCAATTCCACTAATTGTTTGGG** | Del_k15_23-37 | -0.31097676646941314 | 0.028986501843093426 | 35.97708717022934 |
| **GATCCAGATCTAACTTGGGGTGTTTGCTTTGTCTTCTTCTTTTGCCAATTCCACTAATTGTTTGGG** | Ins_k3_22_TTT | -0.25151332798212384 | 0.03662116139707078 | 38.18129388564603 |
| **GATCCAGATCTAACTTGGGGTGTTTGGG** | Del_k35_21-55;Del_k35_22-56;D | -4.222191998076039 | 0.11513310292845187 | 0.7201232052015918 |
| **GATCCAGATCTAACTTGGGGTGTTTGTCTTCTTCTTTTGCCAATTCCACTAATTGTTTGGG** | Del_k2_23-24 | 0.09969384367212764 | 0.032014867064313855 | 54.24728138644362 |
| **GATCCAGATCTAACTTGGGGTGTTTTGCCAATTCCACTAATTGTTTGGG** | Del_k14_23-36 | -0.28501973709884565 | 0.03016832216630638 | 36.923171123611716 |
| **GATCCAGATCTAACTTGGGGTTAAGGCTTTGTCTTCTTCTTTTGCCAATTCCACTAATTGTTTGGG** | Ins_k3_21_TAA | -0.8037206763069076 | 0.04783577804283728 | 21.980118901529263 |
| **GATCCAGATCTAACTTGGGGTTAATTGTTTGGG** | Del_k30_22-51 | -4.267375911304064 | 0.1169266170377205 | 0.688309369942769 |
| **GATCCAGATCTAACTTGGGGTTACGGCTTTGTCTTCTTCTTTTGCCAATTCCACTAATTGTTTGGG** | Ins_k3_21_TAC | 0.5682088578065014 | 0.030693476738395467 | 86.66654102258613 |
| **GATCCAGATCTAACTTGGGGTTAGGCTTTGTCTTCTTCTTTTGCCAATTCCACTAATTGTTTGGG** | Ins_k2_21_TA | -2.8801321808533014 | 0.1513261337947287 | 2.7558525601371113 |
| **GATCCAGATCTAACTTGGGGTTAGGGCTTTGTCTTCTTCTTTTGCCAATTCCACTAATTGTTTGGG** | Ins_k3_21_TAG | -5.333674587728423 | 0.43431163607856016 | 0.23697146334397315 |
| **GATCCAGATCTAACTTGGGGTTATGGCTTTGTCTTCTTCTTTTGCCAATTCCACTAATTGTTTGGG** | Ins_k3_20_TTA;Ins_k3_21_TAT | -0.05260011968730205 | 0.036662863594886914 | 46.584082946535894 |
| **GATCCAGATCTAACTTGGGGTTCAGGCTTTGTCTTCTTCTTTTGCCAATTCCACTAATTGTTTGGG** | Ins_k3_21_TCA | -0.1686293120982441 | 0.03941977238602289 | 41.48076072283626 |
| **GATCCAGATCTAACTTGGGGTTCCACTAATTGTTTGGG** | Del_k25_21-45;Del_k25_22-46 | -1.4385564021406385 | 0.04008058637376676 | 11.649958673423978 |
| **GATCCAGATCTAACTTGGGGTTCCCGGCTTTGTCTTCTTCTTTTGCCAATTCCACTAATTGTTTGGG** | Ins_k4_21_TCCC | 1.0181269422609813 | 0.030455229782516413 | 100 |
| **GATCCAGATCTAACTTGGGGTTCCGGCTTTGTCTTCTTCTTTTGCCAATTCCACTAATTGTTTGGG** | Ins_k3_21_TCC | 0.8822970260608896 | 0.028760135469184343 | 100 |
| **GATCCAGATCTAACTTGGGGTTCGGCTTTGTCTTCTTCTTTTGCCAATTCCACTAATTGTTTGGG** | Ins_k2_21_TC | 0.6937576640899217 | 0.03269575008487505 | 98.25996778545392 |
| **GATCCAGATCTAACTTGGGGTTCGGGCTTTGTCTTCTTCTTTTGCCAATTCCACTAATTGTTTGGG** | Ins_k3_21_TCG | -0.1992250672099492 | 0.040934549596288144 | 40.23084409978305 |
| **GATCCAGATCTAACTTGGGGTTCTGGCTTTGTCTTCTTCTTTTGCCAATTCCACTAATTGTTTGGG** | Ins_k3_20_TTC;Ins_k3_21_TCT | 0.012426047441659871 | 0.038935069226763507 | 49.71392536273384 |
| **GATCCAGATCTAACTTGGGGTTCTTCTTCTTTTGCCAATTCCACTAATTGTTTGGG** | Del_k7_22-28 | 0.125356813493831 | 0.0313987303833086 | 55.65744483031216 |
| **GATCCAGATCTAACTTGGGGTTCTTCTTTTGCCAATTCCACTAATTGTTTGGG** | Del_k10_21-30;Del_k10_22-31 | -0.08039306652868405 | 0.030373159822725038 | 45.30720036314347 |
| **GATCCAGATCTAACTTGGGGTTCTTTTGCCAATTCCACTAATTGTTTGGG** | Del_k13_21-33;Del_k13_22-34 | -0.49842213431819476 | 0.03493455556962959 | 29.827682357375547 |
| **GATCCAGATCTAACTTGGGGTTGAGGCTTTGTCTTCTTCTTTTGCCAATTCCACTAATTGTTTGGG** | Ins_k3_21_TGA | 0.06115105506498042 | 0.03565377116117167 | 52.19622059461783 |
| **GATCCAGATCTAACTTGGGGTTGCCAATTCCACTAATTGTTTGGG** | Del_k18_21-38;Del_k18_22-39 | -0.42566245169447353 | 0.027614418646453143 | 32.0788387284737 |
| **GATCCAGATCTAACTTGGGGTTGCGGCTTTGTCTTCTTCTTTTGCCAATTCCACTAATTGTTTGGG** | Ins_k3_21_TGC | 0.6886921867548699 | 0.03147322442740406 | 97.7634926490592 |
| **GATCCAGATCTAACTTGGGGTTGCTTTGTCTTCTTCTTTTGCCAATTCCACTAATTGTTTGGG** | G22T | -0.8775756723362191 | 0.03361390915984718 | 20.415274332340005 |
| **GATCCAGATCTAACTTGGGGTTGGCGGCTTTGTCTTCTTCTTTTGCCAATTCCACTAATTGTTTGGG** | Ins_k4_21_TGGC | 0.27603785402028913 | 0.03990100369727935 | 64.70877956483424 |
| **GATCCAGATCTAACTTGGGGTTGGCTTTGTCTTCTTCTTTTGCCAATTCCACTAATTGTTTGGG** | Ins_k1_20_T;Ins_k1_21_T | -2.8928477874751986 | 0.1347154274105156 | 2.721032074042295 |
| **GATCCAGATCTAACTTGGGGTTGGG** | Del_k38_21-58;Del_k38_22-59 | -6.163004918514301 | 0.37955676926718884 | 0.10340045839327383 |
| **GATCCAGATCTAACTTGGGGTTGGGCTTTGTCTTCTTCTTTTGCCAATTCCACTAATTGTTTGGG** | Ins_k2_21_TG | -2.9752551916874435 | 0.1600782110808933 | 2.5057894713614552 |
| **GATCCAGATCTAACTTGGGGTTGGGGCTTTGTCTTCTTCTTTTGCCAATTCCACTAATTGTTTGGG** | Ins_k3_21_TGG | -3.928411099612868 | 0.19982952952565658 | 0.9660380417877175 |
| **GATCCAGATCTAACTTGGGGTTGTCTTCTTCTTTTGCCAATTCCACTAATTGTTTGGG** | Del_k5_21-25;Del_k5_22-26 | 0.001655846569231012 | 0.03162955804971751 | 49.18136941559132 |
| **GATCCAGATCTAACTTGGGGTTGTGGCTTTGTCTTCTTCTTTTGCCAATTCCACTAATTGTTTGGG** | Ins_k3_19_GTT;Ins_k3_20_TTG;Ir | -0.09654286711805549 | 0.03630265009145736 | 44.58137485612696 |
| **GATCCAGATCTAACTTGGGGTTGTTTGGG** | Del_k34_21-54;Del_k34_22-55 | -4.560956356668136 | 0.12547057206006265 | 0.5131960608833385 |
| **GATCCAGATCTAACTTGGGGTTTAGGCTTTGTCTTCTTCTTTTGCCAATTCCACTAATTGTTTGGG** | Ins_k3_21_TTA | -3.1220620444372322 | 0.12862443133263735 | 2.1636508221504536 |
| **GATCCAGATCTAACTTGGGGTTTCCACTAATTGTTTGGG** | Del_k24_22-45 | -1.3857766984653934 | 0.037884482210392506 | 12.281355954062041 |
| **GATCCAGATCTAACTTGGGGTTTCGGCTTTGTCTTCTTCTTTTGCCAATTCCACTAATTGTTTGGG** | Ins_k3_21_TTC | 0.6793058679801913 | 0.029130744250957758 | 96.85014652715485 |
| **GATCCAGATCTAACTTGGGGTTTCTTCTTTTGCCAATTCCACTAATTGTTTGGG** | Del_k9_22-30 | -0.18648658287591408 | 0.03351831120145682 | 40.74660209035984 |
| **GATCCAGATCTAACTTGGGGTTTCTTTTGCCAATTCCACTAATTGTTTGGG** | Del_k12_22-33 | -0.8820104506423745 | 0.04081019544435814 | 20.324937576415756 |
| **GATCCAGATCTAACTTGGGGTTTGCCAATTCCACTAATTGTTTGGG** | Del_k17_21-37;Del_k17_22-38 | -0.5727173133500478 | 0.028145357168759457 | 27.69194900895762 |
| **GATCCAGATCTAACTTGGGGTTTGGCTTTGTCTTCTTCTTTTGCCAATTCCACTAATTGTTTGGG** | Ins_k2_20_TT;Ins_k2_21_TT | -0.23067308052892033 | 0.04116516202738623 | 38.98535076950504 |
| **GATCCAGATCTAACTTGGGGTTTGGG** | Del_k37_20-56;Del_k37_21-57;D | -5.567282476994087 | 0.27970507404968786 | 0.1876037145573711 |
| **GATCCAGATCTAACTTGGGGTTTGGGCTTTGTCTTCTTCTTTTGCCAATTCCACTAATTGTTTGGG** | Ins_k3_21_TTG | -2.441256180872875 | 0.09160407475588746 | 4.2742252330434765 |
| **GATCCAGATCTAACTTGGGGTTTGTCTTCTTCTTTTGCCAATTCCACTAATTGTTTGGG** | Del_k4_21-24;Del_k4_22-25 | -0.37511648241726586 | 0.040113460870584546 | 33.74197301313982 |
| **GATCCAGATCTAACTTGGGGTTTGTTTGGG** | Del_k33_22-54 | -4.527019912220899 | 0.14098037503342276 | 0.5309110014089661 |
| **GATCCAGATCTAACTTGGGGTTTTGCCAATTCCACTAATTGTTTGGG** | Del_k16_21-36;Del_k16_22-37 | -0.4516206031063864 | 0.032105268067124444 | 31.25684623456527 |
| **GATCCAGATCTAACTTGGGGTTTTGGCTTTGTCTTCTTCTTTTGCCAATTCCACTAATTGTTTGGG** | Ins_k3_20_TTT;Ins_k3_21_TTT | -0.6173616612743965 | 0.04869220568120014 | 26.482850525684576 |
| **GATCCAGATCTAACTTGGGGTTTTGGG** | Del_k36_22-57 | -5.495968062837253 | 0.22413841515482777 | 0.2014711612799608 |
| **GATCCAGATCTAACTTGGGGTTTTGTCTTCTTCTTTTGCCAATTCCACTAATTGTTTGGG** | Del_k3_22-24 | -0.22714423375048653 | 0.035028446332229894 | 39.123167122354374 |
| **GATCCAGATCTAACTTGGGGTTTTTGCCAATTCCACTAATTGTTTGGG** | Del_k15_22-36 | -0.48916642956182976 | 0.034074191319454605 | 30.105040169797658 |
| **GATCCAGATCTAACTTGGGTAAGTGGCTTTGTCTTCTTCTTTTGCCAATTCCACTAATTGTTTGGG** | Ins_k3_19_TAA | -6.151207362227298 | 0.7449997571845904 | 0.10462755526096928 |
| **GATCCAGATCTAACTTGGGTAATTGTTTGGG** | Del_k32_20-51 | -5.223614751216579 | 0.2655534703888271 | 0.26454187664498596 |
| **GATCCAGATCTAACTTGGGTACGTGGCTTTGTCTTCTTCTTTTGCCAATTCCACTAATTGTTTGGG** | Ins_k3_19_TAC | -0.8252885466443888 | 0.043416128755659804 | 21.511130269987074 |

| | | | | |
|---|---|---|---|---|
| GATCCAGATCTAACTTGGGTAGGTGGCTTTGTCTTCTTCTTTTGCCAATTCCACTAATTGTTTGGG | Ins_k3_18_GTA;Ins_k3_19_TAG | -4.299682622770718 | 0.33500223901771653 | 0.6664277229227664 |
| GATCCAGATCTAACTTGGGTAGTGGCTTTGTCTTCTTCTTTTGCCAATTCCACTAATTGTTTGGG | Ins_k2_19_TA | -0.437451567364197 | 0.04592363286034759 | 31.702878064374257 |
| GATCCAGATCTAACTTGGGTATGTGGCTTTGTCTTCTTCTTTTGCCAATTCCACTAATTGTTTGGG | Ins_k3_19_TAT | -1.778220610050599 | 0.07359194442728965 | 8.294879732558321 |
| GATCCAGATCTAACTTGGGTCAGTGGCTTTGTCTTCTTCTTTTGCCAATTCCACTAATTGTTTGGG | Ins_k3_19_TCA | -0.005999899954824306 | 0.031286516016750726 | 48.80628691788153 |
| GATCCAGATCTAACTTGGGTCCACTAATTGTTTGGG | Del_k27_20-46 | -5.0533379194238215 | 0.18628334030001245 | 0.3136495859245737 |
| GATCCAGATCTAACTTGGGTCCGTGGCTTTGTCTTCTTCTTTTGCCAATTCCACTAATTGTTTGGG | Ins_k3_19_TCC | -0.15084215324337658 | 0.034759534429771304 | 42.225186586912336 |
| GATCCAGATCTAACTTGGGTCGAGTGGCTTTGTCTTCTTCTTTTGCCAATTCCACTAATTGTTTGGG | Ins_k4_19_TCGA | 0.7921664189174793 | 0.031707181571786536 | 100 |
| GATCCAGATCTAACTTGGGTCGGGTGGCTTTGTCTTCTTCTTTTGCCAATTCCACTAATTGTTTGGG | Ins_k4_17_GGTC;Ins_k4_18_GT | -0.015859627130423548 | 0.047242705663834386 | 48.32743479930383 |
| GATCCAGATCTAACTTGGGTCGGTGGCTTTGTCTTCTTCTTTTGCCAATTCCACTAATTGTTTGGG | Ins_k3_18_GTC;Ins_k3_19_TCG | 0.6344189869303505 | 0.02833204320870222 | 92.59897030101195 |
| GATCCAGATCTAACTTGGGTCGTGGCTTTGTCTTCTTCTTTTGCCAATTCCACTAATTGTTTGGG | Ins_k2_19_TC | 0.7764327679635341 | 0.030390101672223245 | 100 |
| GATCCAGATCTAACTTGGGTCTCGTGGCTTTGTCTTCTTCTTTTGCCAATTCCACTAATTGTTTGGG | Ins_k4_19_TCTC | 0.6945947027488595 | 0.030698898604835472 | 98.34224960884573 |
| GATCCAGATCTAACTTGGGTCTGTGGCTTTGTCTTCTTCTTTTGCCAATTCCACTAATTGTTTGGG | Ins_k3_19_TCT | -0.5219129422963977 | 0.03903975883118488 | 29.135171661275574 |
| GATCCAGATCTAACTTGGGTCTTCTTCTTTTGCCAATTCCACTAATTGTTTGGG | Del_k9_19-27;Del_k9_20-28 | -0.22617452314906195 | 0.034357926399430985 | 39.16112367273454 |
| GATCCAGATCTAACTTGGGTCTTCTTTTGCCAATTCCACTAATTGTTTGGG | Del_k12_20-31 | -0.4520073590747963 | 0.033176327119633885 | 31.24475980013137 |
| GATCCAGATCTAACTTGGGTCTTTTGCCAATTCCACTAATTGTTTGGG | Del_k15_20-34 | -1.262033216840737 | 0.042137044102805436 | 13.899124019344368 |
| GATCCAGATCTAACTTGGGTGAGTGGCTTTGTCTTCTTCTTTTGCCAATTCCACTAATTGTTTGGG | Ins_k3_19_TGA | -2.8612607794576568 | 0.12370321992041361 | 2.8083531820645873 |
| GATCCAGATCTAACTTGGGTGCCAATTCCACTAATTGTTTGGG | Del_k20_20-39 | -0.2241664028950851 | 0.026741241599787153 | 39.23984293076521 |
| GATCCAGATCTAACTTGGGTGCGTGGCTTTGTCTTCTTCTTTTGCCAATTCCACTAATTGTTTGGG | Ins_k3_19_TGC | 0.3524278318574281 | 0.03069203096921103 | 69.84558428928199 |
| GATCCAGATCTAACTTGGGTGGCTTTGTCTTCTTCTTTTGCCAATTCCACTAATTGTTTGGG | Del_k1_17-17;Del_k1_18-18;Del_ | -0.07534345007235166 | 0.02793370743187835 | 45.536562956882406 |
| GATCCAGATCTAACTTGGGTGGG | Del_k40_20-59 | -6.324935020845301 | 0.6300357951070661 | 0.087942157122954015 |
| GATCCAGATCTAACTTGGGTGGGTGGCTTTGTCTTCTTCTTTTGCCAATTCCACTAATTGTTTGGG | Ins_k3_17_GGT;Ins_k3_18_GTG; | -4.154513633510725 | 0.3074468586951809 | 0.7705470215915211 |
| GATCCAGATCTAACTTGGGTGGTGGCTTTGTCTTCTTCTTTTGCCAATTCCACTAATTGTTTGGG | Ins_k2_18_GT;Ins_k2_19_TG | 0.3879527671967211 | 0.03308928622000001 | 72.3714440216399 |
| GATCCAGATCTAACTTGGGTGTCTTCTTCTTTTGCCAATTCCACTAATTGTTTGGG | Del_k7_20-26 | 0.36414253552018583 | 0.030439672852604103 | 70.66861598526943 |
| GATCCAGATCTAACTTGGGTGTGGCTTTGTCTTCTTCTTTTGCCAATTCCACTAATTGTTTGGG | Ins_k1_19_T | 0.425591232379386 | 0.034253515208870336 | 75.14730599450196 |
| GATCCAGATCTAACTTGGGTGTGTGGCTTTGTCTTCTTCTTTTGCCAATTCCACTAATTGTTTGGG | Ins_k3_19_TGT | 0.1151788823406289 | 0.032817457104134015 | 55.09384021845633 |
| GATCCAGATCTAACTTGGGTGTTTGGG | Del_k36_20-55 | -4.833736482213654 | 0.13113752261036415 | 0.3906757107437948 |
| GATCCAGATCTAACTTGGGTTAGTGGCTTTGTCTTCTTCTTTTGCCAATTCCACTAATTGTTTGGG | Ins_k3_19_TTA | -1.7510135471971773 | 0.07692163381311427 | 8.523657115076285 |
| GATCCAGATCTAACTTGGGTTCCACTAATTGTTTGGG | Del_k26_20-45 | -1.5747097778262626 | 0.0343845532793684 | 10.167021103472544 |
| GATCCAGATCTAACTTGGGTTCGTGGCTTTGTCTTCTTCTTTTGCCAATTCCACTAATTGTTTGGG | Ins_k3_19_TTC | 0.4239926483776093 | 0.030140368532677903 | 75.02727268059198 |
| GATCCAGATCTAACTTGGGTTCTTCTTTTGCCAATTCCACTAATTGTTTGGG | Del_k11_20-30 | -0.2446763838350663 | 0.034925792102986956 | 38.44323166606201 |
| GATCCAGATCTAACTTGGGTTCTTTTGCCAATTCCACTAATTGTTTGGG | Del_k14_20-33 | -1.1536991146106408 | 0.04467788968813758 | 15.489461975982753 |
| GATCCAGATCTAACTTGGGTTGCCAATTCCACTAATTGTTTGGG | Del_k19_20-38 | -0.34043261480318876 | 0.02992895621868284 | 34.932807128757176 |
| GATCCAGATCTAACTTGGGTTGGCTTTGTCTTCTTCTTTTGCCAATTCCACTAATTGTTTGGG | G20T | -2.0674277408427173 | 0.054370402335101944 | 6.211679398699622 |
| GATCCAGATCTAACTTGGGTTGGG | Del_k39_20-58 | -6.4988828780973416 | 0.41875066061070676 | 0.07390137512412327 |
| GATCCAGATCTAACTTGGGTTGGTGGCTTTGTCTTCTTCTTTTGCCAATTCCACTAATTGTTTGGG | Ins_k3_18_GTT;Ins_k3_19_TTG | -0.14787392879280103 | 0.03642335738417776 | 42.35070661181686 |
| GATCCAGATCTAACTTGGGTTGTCTTCTTCTTTTGCCAATTCCACTAATTGTTTGGG | Del_k6_20-25 | -0.13742639070768814 | 0.030672395845074987 | 42.79548661440713 |
| GATCCAGATCTAACTTGGGTTGTGGCTTTGTCTTCTTCTTTTGCCAATTCCACTAATTGTTTGGG | Ins_k2_19_TT | 0.07160462578404769 | 0.0343253757343354 | 52.7447193683764 |
| GATCCAGATCTAACTTGGGTTGTTTGGG | Del_k35_20-54 | -5.288729082754756 | 0.20055033823546448 | 0.2478652448700365 |
| GATCCAGATCTAACTTGGGTTTGCCAATTCCACTAATTGTTTGGG | Del_k18_20-37 | -0.8258133889208712 | 0.03079784596174474 | 21.499843281609895 |
| GATCCAGATCTAACTTGGGTTTGGG | Del_k38_19-56;Del_k38_20-57 | -5.772439971897373 | 0.24394555697854314 | 0.15280679442640724 |
| GATCCAGATCTAACTTGGGTTTGTCTTCTTCTTTTGCCAATTCCACTAATTGTTTGGG | Del_k5_20-24 | -0.5906923128984541 | 0.035428385399950316 | 27.19863319626407 |
| GATCCAGATCTAACTTGGGTTTGTGGCTTTGTCTTCTTCTTTTGCCAATTCCACTAATTGTTTGGG | Ins_k3_19_TTT | -0.5699352076382311 | 0.041044186333260915 | 27.76909820731754 |
| GATCCAGATCTAACTTGGGTTTTGCCAATTCCACTAATTGTTTGGG | Del_k17_20-36 | -0.6569301711991785 | 0.03340885356612106 | 25.45542449740681 |
| GATCCAGATCTAACTTGGTAAGGGTGGCTTTGTCTTCTTCTTTTGCCAATTCCACTAATTGTTTGGG | Ins_k4_17_GTAA;Ins_k4_18_TAA | -3.236843812502678 | 0.14801185940332112 | 1.9290260192493196 |
| GATCCAGATCTAACTTGGTAAGGTGGCTTTGTCTTCTTCTTTTGCCAATTCCACTAATTGTTTGGG | Ins_k3_18_TAA | -3.7256611812601084 | 0.17460510669650073 | 1.1831706819994159 |
| GATCCAGATCTAACTTGGTAATTGTTTGGG | Del_k33_19-51 | -5.2910434521631675 | 0.23858556939142878 | 0.24729225643910724 |
| GATCCAGATCTAACTTGGTACGGTGGCTTTGTCTTCTTCTTTTGCCAATTCCACTAATTGTTTGGG | Ins_k3_18_TAC | 0.18985306612509212 | 0.03008619720391327 | 59.365431811171185 |
| GATCCAGATCTAACTTGGTACTGGTGGCTTTGTCTTCTTCTTTTGCCAATTCCACTAATTGTTTGGG | Ins_k4_18_TACT | 0.5458938315149242 | 0.029491809204279427 | 84.75399351873432 |
| GATCCAGATCTAACTTGGTAGGGTGGCTTTGTCTTCTTCTTTTGCCAATTCCACTAATTGTTTGGG | Ins_k3_17_GTA;Ins_k3_18_TAG | -2.995101627274099 | 0.1304896499252343 | 2.456548724914609 |
| GATCCAGATCTAACTTGGTAGGTGGCTTTGTCTTCTTCTTTTGCCAATTCCACTAATTGTTTGGG | Ins_k2_18_TA | -5.217229575348961 | 0.7005492823563165 | 0.26623642729637836 |
| GATCCAGATCTAACTTGGTATGGTGGCTTTGTCTTCTTCTTTTGCCAATTCCACTAATTGTTTGGG | Ins_k3_18_TAT | -0.04227339271076558 | 0.03213509698427921 | 47.067636518209945 |
| GATCCAGATCTAACTTGGTCAGGTGGCTTTGTCTTCTTCTTTTGCCAATTCCACTAATTGTTTGGG | Ins_k3_18_TCA | -0.04639218454931715 | 0.03454101449542956 | 46.87417341169039 |
| GATCCAGATCTAACTTGGTCCACTAATTGTTTGGG | Del_k28_19-46 | -2.2934073881707002 | 0.05027839412919217 | 4.9552699500441815 |
| GATCCAGATCTAACTTGGTCCGGTGGCTTTGTCTTCTTCTTTTGCCAATTCCACTAATTGTTTGGG | Ins_k3_18_TCC | 0.6591434226823216 | 0.02962019043272148 | 94.91696506839382 |
| GATCCAGATCTAACTTGGTCGGGTGGCTTTGTCTTCTTCTTTTGCCAATTCCACTAATTGTTTGGG | Ins_k3_17_GTC;Ins_k3_18_TCG | 0.5652783094501137 | 0.029496909465260568 | 86.41293232102203 |
| GATCCAGATCTAACTTGGTCGGTGGCTTTGTCTTCTTCTTTTGCCAATTCCACTAATTGTTTGGG | Ins_k2_18_TC | 0.654328673402301 | 0.03185157458585341 | 94.46106208916059 |
| GATCCAGATCTAACTTGGTCTGGTGGCTTTGTCTTCTTCTTTTGCCAATTCCACTAATTGTTTGGG | Ins_k3_18_TCT | -2.2706269047410714 | 0.08532115559769922 | 5.069448984098325 |
| GATCCAGATCTAACTTGGTCTTCTTCTTTTGCCAATTCCACTAATTGTTTGGG | Del_k10_18-27;Del_k10_19-28 | -0.09942960388501942 | 0.032236370270635965 | 44.45286573744839 |
| GATCCAGATCTAACTTGGTCTTCTTTTGCCAATTCCACTAATTGTTTGGG | Del_k13_19-31 | -0.226235514331789 | 0.03099849829639663 | 39.15873526232155 |
| GATCCAGATCTAACTTGGTCTTTTGCCAATTCCACTAATTGTTTGGG | Del_k16_19-34 | -1.2090853834369613 | 0.03979725445729579 | 14.654883891835594 |
| GATCCAGATCTAACTTGGTGAGGTGGCTTTGTCTTCTTCTTTTGCCAATTCCACTAATTGTTTGGG | Ins_k3_18_TGA | 0.056399961336279336 | 0.030154994314733827 | 51.948819636239165 |
| GATCCAGATCTAACTTGGTGCCAATTCCACTAATTGTTTGGG | Del_k21_19-39 | -0.4160279535920124 | 0.030735440076832148 | 32.38939586817117 |
| GATCCAGATCTAACTTGGTGCGGTGGCTTTGTCTTCTTCTTTTGCCAATTCCACTAATTGTTTGGG | Ins_k3_18_TGC | 0.6582077253570306 | 0.025636566265086366 | 94.82819305639856 |
| GATCCAGATCTAACTTGGTGGCTTTGTCTTCTTCTTTTGCCAATTCCACTAATTGTTTGGG | Del_k2_17-18;Del_k2_18-19;Del_ | 0.20048732308450146 | 0.030386118420356587 | 60.00010773986927 |
| GATCCAGATCTAACTTGGTGGG | Del_k41_19-59 | -5.9666343106250705 | 0.2444400174775337 | 0.12583606685770887 |
| GATCCAGATCTAACTTGGTGGGGTGGCTTTGTCTTCTTCTTTTGCCAATTCCACTAATTGTTTGGG | Ins_k3_15_TGG;Ins_k3_16_GGT; | -1.1398416027858769 | 0.05294853982456875 | 15.705601497690049 |
| GATCCAGATCTAACTTGGTGGGTGGCTTTGTCTTCTTCTTTTGCCAATTCCACTAATTGTTTGGG | Ins_k2_17_GT;Ins_k2_18_TG | -1.3593294999373282 | 0.06903547300298903 | 12.610496653196728 |
| GATCCAGATCTAACTTGGTGGTGGCTTTGTCTTCTTCTTTTGCCAATTCCACTAATTGTTTGGG | Ins_k1_18_T | 0.5009804144195551 | 0.029666962088091296 | 81.03162002851587 |
| GATCCAGATCTAACTTGGTGTCTTCTTCTTTTGCCAATTCCACTAATTGTTTGGG | Del_k8_19-26 | 0.31007242805007273 | 0.027634327968982523 | 66.94902191540444 |
| GATCCAGATCTAACTTGGTGTGGCTTTGTCTTCTTCTTTTGCCAATTCCACTAATTGTTTGGG | G19T | -0.34502810297640135 | 0.026515858712980643 | 34.77264212679643 |
| GATCCAGATCTAACTTGGTGTGGTGGCTTTGTCTTCTTCTTTTGCCAATTCCACTAATTGTTTGGG | Ins_k3_18_TGT | 0.47150707282777393 | 0.027878228192989324 | 78.67819935857422 |
| GATCCAGATCTAACTTGGTGTTTGGG | Del_k37_19-55 | -4.35956830681819 | 0.10233397279440126 | 0.6276897444957948 |
| GATCCAGATCTAACTTGGTTAGGTGGCTTTGTCTTCTTCTTTTGCCAATTCCACTAATTGTTTGGG | Ins_k3_18_TTA | -1.8965155420679352 | 0.06824294516697937 | 7.369453067030788 |
| GATCCAGATCTAACTTGGTTCCACTAATTGTTTGGG | Del_k27_19-45 | -2.209954723432578 | 0.044531563514283626 | 5.386545720891075 |
| GATCCAGATCTAACTTGGTTCGGTGGCTTTGTCTTCTTCTTTTGCCAATTCCACTAATTGTTTGGG | Ins_k3_18_TTC | 0.6847556408199511 | 0.027771685504484524 | 97.37939866729519 |

| | | | | |
|---|---|---|---|---|
| **GATCCAGATCTAACTTGGTTCTTCTTTTGCCAATTCCACTAATTGTTTGGG** | Del_k12_19-30 | -0.3866891476908071 | 0.03161118955447252 | 33.35373923645483 |
| **GATCCAGATCTAACTTGGTTCTTTTGCCAATTCCACTAATTGTTTGGG** | Del_k15_19-33 | -1.3827748204597783 | 0.044516948906079025 | 12.318278477108025 |
| **GATCCAGATCTAACTTGGTTGCCAATTCCACTAATTGTTTGGG** | Del_k20_19-38 | -0.6360614614278606 | 0.03185699944005168 | 25.99222808095185 |
| **GATCCAGATCTAACTTGGTTGGG** | Del_k40_19-58 | -6.097470508511149 | 0.21874359615296812 | 0.11040371738828739 |
| **GATCCAGATCTAACTTGGTTGGGTGGCTTTGTCTTCTTCTTTTGCCAATTCCACTAATTGTTTGGG** | Ins_k3_17_GTT;Ins_k3_18_TTG | -0.35189270998142197 | 0.0401098932167545 | 34.534759025215486 |
| **GATCCAGATCTAACTTGGTTGGTGGCTTTGTCTTCTTCTTTTGCCAATTCCACTAATTGTTTGGG** | Ins_k2_18_TT | 0.1752028796471008 | 0.032998409288393725 | 58.5020569084317 |
| **GATCCAGATCTAACTTGGTTGTCTTCTTCTTTTGCCAATTCCACTAATTGTTTGGG** | Del_k7_19-25 | -0.03290466346871368 | 0.03222890042739671 | 47.510672562636 |
| **GATCCAGATCTAACTTGGTTGTTTGGG** | Del_k36_19-54 | -5.452133452480823 | 0.1533965631587401 | 0.2104989913241704 |
| **GATCCAGATCTAACTTGGTTTGCCAATTCCACTAATTGTTTGGG** | Del_k19_19-37 | -0.6221970112024274 | 0.03365620317222917 | 26.355105770493314 |
| **GATCCAGATCTAACTTGGTTTGGG** | Del_k39_18-56;Del_k39_19-57 | -5.1663512405415055 | 0.16279998912920426 | 0.28013260300942094 |
| **GATCCAGATCTAACTTGGTTTGGTGGCTTTGTCTTCTTCTTTTGCCAATTCCACTAATTGTTTGGG** | Ins_k3_18_TTT | 0.3980012688513308 | 0.02986640998994004 | 73.10233462438306 |
| **GATCCAGATCTAACTTGGTTTGTCTTCTTCTTTTGCCAATTCCACTAATTGTTTGGG** | Del_k6_19-24 | -1.0195904935063616 | 0.04599769355944533 | 17.712463446349652 |
| **GATCCAGATCTAACTTGGTTTTGCCAATTCCACTAATTGTTTGGG** | Del_k18_19-36 | -1.170021408844387 | 0.03500548849224744 | 15.238690569887979 |
| **GATCCAGATCTAACTTGTAAGGGTGGCTTTGTCTTCTTCTTTTGCCAATTCCACTAATTGTTTGGG** | Ins_k3_17_TAA | -0.7729950799541575 | 0.041590258090516065 | 22.66595354688662 |
| **GATCCAGATCTAACTTGTAATTGTTTGGG** | Del_k34_18-51 | -5.2352173413359475 | 0.17596361065069108 | 0.26149024334026794 |
| **GATCCAGATCTAACTTGTACGGGTGGCTTTGTCTTCTTCTTTTGCCAATTCCACTAATTGTTTGGG** | Ins_k3_17_TAC | 0.39341940973855927 | 0.03390241220940896 | 72.76815619020061 |
| **GATCCAGATCTAACTTGTAGGGGTGGCTTTGTCTTCTTCTTTTGCCAATTCCACTAATTGTTTGGG** | Ins_k3_16_GTA;Ins_k3_17_TAG | -1.1719070351833218 | 0.0568953256665952 | 15.209983167801237 |
| **GATCCAGATCTAACTTGTAGGGTGGCTTTGTCTTCTTCTTTTGCCAATTCCACTAATTGTTTGGG** | Ins_k2_17_TA | -2.5283951802816444 | 0.1046137141275817 | 3.9175397991454908 |
| **GATCCAGATCTAACTTGTATGGGTGGCTTTGTCTTCTTCTTTTGCCAATTCCACTAATTGTTTGGG** | Ins_k3_17_TAT | -0.07032463031731251 | 0.0305740831165152 | 45.765677219299015 |
| **GATCCAGATCTAACTTGTCAGGGTGGCTTTGTCTTCTTCTTTTGCCAATTCCACTAATTGTTTGGG** | Ins_k3_17_TCA | -0.5467451243259611 | 0.041609328471176696 | 28.420590795574643 |
| **GATCCAGATCTAACTTGTCCACTAATTGTTTGGG** | Del_k29_18-46 | -2.7855807802018053 | 0.05923160882142223 | 3.0291384967809507 |
| **GATCCAGATCTAACTTGTCCGGGTGGCTTTGTCTTCTTCTTTTGCCAATTCCACTAATTGTTTGGG** | Ins_k3_17_TCC | 0.6860645400325001 | 0.028952022855804612 | 97.50694193796346 |
| **GATCCAGATCTAACTTGTCGGGGTGGCTTTGTCTTCTTCTTTTGCCAATTCCACTAATTGTTTGGG** | Ins_k3_16_GTC;Ins_k3_17_TCG | 0.5431537357200502 | 0.027948851781921098 | 84.52207733887407 |
| **GATCCAGATCTAACTTGTCGGGTGGCTTTGTCTTCTTCTTTTGCCAATTCCACTAATTGTTTGGG** | Ins_k2_17_TC | 0.545786246645005 | 0.0279849053654969 | 84.74487576184161 |
| **GATCCAGATCTAACTTGTCTGGGTGGCTTTGTCTTCTTCTTTTGCCAATTCCACTAATTGTTTGGG** | Ins_k3_17_TCT | -0.14817078613918483 | 0.03364354076423167 | 42.33813635931281 |
| **GATCCAGATCTAACTTGTCTTCTTCTTTTGCCAATTCCACTAATTGTTTGGG** | Del_k11_15-25;Del_k11_16-26;D | -1.4749145194367967 | 0.04794326667760387 | 11.233995745161918 |
| **GATCCAGATCTAACTTGTCTTCTTTTGCCAATTCCACTAATTGTTTGGG** | Del_k14_18-31 | -1.273900044095699 | 0.039887102823791674 | 13.735160304284262 |
| **GATCCAGATCTAACTTGTCTTTTGCCAATTCCACTAATTGTTTGGG** | Del_k17_18-34 | -1.9279096508810225 | 0.045467536392281876 | 7.141689577910808 |
| **GATCCAGATCTAACTTGTGAGGGTGGCTTTGTCTTCTTCTTTTGCCAATTCCACTAATTGTTTGGG** | Ins_k3_17_TGA | 0.23800731623646143 | 0.028582956103545377 | 62.2940771242343 |
| **GATCCAGATCTAACTTGTGCCAATTCCACTAATTGTTTGGG** | Del_k22_18-39 | -0.9817100034190194 | 0.03360246354504153 | 18.39629032251359 |
| **GATCCAGATCTAACTTGTGCGGGTGGCTTTGTCTTCTTCTTTTGCCAATTCCACTAATTGTTTGGG** | Ins_k3_17_TGC | 0.39834182878686253 | 0.029186650233990908 | 73.12723459047471 |
| **GATCCAGATCTAACTTGTGGCTTTGTCTTCTTCTTTTGCCAATTCCACTAATTGTTTGGG** | Del_k3_17-19;Del_k3_18-20 | -0.08328102105163604 | 0.033828838802104876 | 45.176543984581386 |
| **GATCCAGATCTAACTTGTGGG** | Del_k42_18-59 | -6.088176155283055 | 0.23906355721532316 | 0.11143463195617409 |
| **GATCCAGATCTAACTTGTGGGGGTGGCTTTGTCTTCTTCTTTTGCCAATTCCACTAATTGTTTGGG** | Ins_k3_16_GTG;Ins_k3_17_TGG | -1.0622479515750292 | 0.05499493087111743 | 16.972783375765015 |
| **GATCCAGATCTAACTTGTGGGGTGGCTTTGTCTTCTTCTTTTGCCAATTCCACTAATTGTTTGGG** | Ins_k2_15_TG;Ins_k2_16_GT;Ins_ | -0.600950885210485 | 0.0472436610075412 | 26.921040338480136 |
| **GATCCAGATCTAACTTGTGGGTGGCTTTGTCTTCTTCTTTTGCCAATTCCACTAATTGTTTGGG** | Ins_k1_17_T | -0.18281220935286052 | 0.040651617091349235 | 40.89659572379716 |
| **GATCCAGATCTAACTTGTGGTGGCTTTGTCTTCTTCTTTTGCCAATTCCACTAATTGTTTGGG** | G18T | -0.4171364179557132 | 0.02764660429840216 | 32.353513268055316 |
| **GATCCAGATCTAACTTGTGTCTTCTTCTTTTGCCAATTCCACTAATTGTTTGGG** | Del_k9_18-26 | -0.8761971595514894 | 0.039820666007038874 | 20.443436455474593 |
| **GATCCAGATCTAACTTGTGTGGGTGGCTTTGTCTTCTTCTTTTGCCAATTCCACTAATTGTTTGGG** | Ins_k3_17_TGT | -0.2698680564284705 | 0.03396556963590082 | 37.486878998347855 |
| **GATCCAGATCTAACTTGTGTTTGGG** | Del_k38_18-55 | -5.820891713504484 | 0.20826372056467174 | 0.14557953947121174 |
| **GATCCAGATCTAACTTGTTAGGGTGGCTTTGTCTTCTTCTTTTGCCAATTCCACTAATTGTTTGGG** | Ins_k3_17_TTA | -3.8598125551009512 | 0.16818537649576878 | 1.0346326950273268 |
| **GATCCAGATCTAACTTGTTCAGGGTGGCTTTGTCTTCTTCTTTTGCCAATTCCACTAATTGTTTGGG** | Ins_k4_17_TTCA | -0.37913911143518275 | 0.03489252369368842 | 33.60651420629997 |
| **GATCCAGATCTAACTTGTTCCACTAATTGTTTGGG** | Del_k28_18-45 | -2.689408707595948 | 0.048748614998597 | 3.3349254563693034 |
| **GATCCAGATCTAACTTGTTCGGGTGGCTTTGTCTTCTTCTTTTGCCAATTCCACTAATTGTTTGGG** | Ins_k3_17_TTC | 0.44187245997427227 | 0.026550933569125707 | 76.38081062351361 |
| **GATCCAGATCTAACTTGTTCTTCTTTTGCCAATTCCACTAATTGTTTGGG** | Del_k13_18-30 | -1.6035199787325054 | 0.055076308700122416 | 9.878286406385527 |
| **GATCCAGATCTAACTTGTTCTTTTGCCAATTCCACTAATTGTTTGGG** | Del_k16_18-33 | -2.945537847410631 | 0.09134466429478574 | 2.5813723790926377 |
| **GATCCAGATCTAACTTGTTGCCAATTCCACTAATTGTTTGGG** | Del_k21_18-38 | -1.1980426247144789 | 0.04177854549210208 | 14.817611063596711 |
| **GATCCAGATCTAACTTGTTGGG** | Del_k41_18-58 | -5.783635271809991 | 0.2784549841081223 | 0.1511056168984494 |
| **GATCCAGATCTAACTTGTTGGGGTGGCTTTGTCTTCTTCTTTTGCCAATTCCACTAATTGTTTGGG** | Ins_k3_14_TTG;Ins_k3_15_TGT;I | -0.012632549265160331 | 0.03515801573290068 | 48.48364310705261 |
| **GATCCAGATCTAACTTGTTGGGTGGCTTTGTCTTCTTCTTTTGCCAATTCCACTAATTGTTTGGG** | Ins_k2_17_TT | 0.1017068189209357 | 0.033259489592746716 | 54.35658980184657 |
| **GATCCAGATCTAACTTGTTGTCTTCTTCTTTTGCCAATTCCACTAATTGTTTGGG** | Del_k8_18-25 | -1.4807446416738501 | 0.0503809756329353 | 11.168690729846201 |
| **GATCCAGATCTAACTTGTTGTTTGGG** | Del_k37_18-54 | -5.513179495314376 | 0.15850928954560942 | 0.19803322476646953 |
| **GATCCAGATCTAACTTGTTTGCCAATTCCACTAATTGTTTGGG** | Del_k20_18-37 | -1.4305062013382281 | 0.041401597524328974 | 11.744121687144418 |
| **GATCCAGATCTAACTTGTTTGGG** | Del_k40_15-54;Del_k40_16-55;D | -5.46401302507791 | 0.1597432350070036 | 0.20801314788823194 |
| **GATCCAGATCTAACTTGTTTGGGTGGCTTTGTCTTCTTCTTTTGCCAATTCCACTAATTGTTTGGG** | Ins_k3_17_TTT | -0.16085772027940698 | 0.0352044441461585 | 41.80438818491774 |
| **GATCCAGATCTAACTTGTTTGTCTTCTTCTTTTGCCAATTCCACTAATTGTTTGGG** | Del_k7_18-24 | -2.7989376108181765 | 0.10212841707373602 | 2.98894781454957 |
| **GATCCAGATCTAACTTGTTTTGCCAATTCCACTAATTGTTTGGG** | Del_k19_18-36 | -1.7260388576150392 | 0.04509298843125133 | 8.739213326313005 |
| **GATCCAGATCTAACTTTAAGGGGTGGCTTTGTCTTCTTCTTTTGCCAATTCCACTAATTGTTTGGG** | Ins_k3_16_TAA | -0.9556667836921262 | 0.05111273552427574 | 18.881682101510396 |
| **GATCCAGATCTAACTTTAATTGTTTGGG** | Del_k35_17-51 | -5.620431935087853 | 0.2788005040706585 | 0.17789302361448 |
| **GATCCAGATCTAACTTTACGGGGTGGCTTTGTCTTCTTCTTTTGCCAATTCCACTAATTGTTTGGG** | Ins_k3_16_TAC | 0.23598079589987642 | 0.03069134668561819 | 62.16796473791072 |
| **GATCCAGATCTAACTTTAGGGGGTGGCTTTGTCTTCTTCTTTTGCCAATTCCACTAATTGTTTGGG** | Ins_k3_16_TAG | -3.3536446397935435 | 0.18739476786333747 | 1.7163748049291052 |
| **GATCCAGATCTAACTTTAGGGGTGGCTTTGTCTTCTTCTTTTGCCAATTCCACTAATTGTTTGGG** | Ins_k2_16_TA | -1.8557683158783669 | 0.0923487936291073 | 7.6759396728386635 |
| **GATCCAGATCTAACTTTATGGGGTGGCTTTGTCTTCTTCTTTTGCCAATTCCACTAATTGTTTGGG** | Ins_k3_15_TTA;Ins_k3_16_TAT | -0.5911139044740308 | 0.04011124446768162 | 27.18716889843682 |
| **GATCCAGATCTAACTTTCAGGGGTGGCTTTGTCTTCTTCTTTTGCCAATTCCACTAATTGTTTGGG** | Ins_k3_16_TCA | -0.5965222807002174 | 0.04688909665702471 | 27.040527364247918 |
| **GATCCAGATCTAACTTTCCACTAATTGTTTGGG** | Del_k30_16-45;Del_k30_17-46 | -3.5912433858845243 | 0.08569502460357771 | 1.3533941859413776 |
| **GATCCAGATCTAACTTTCCGGGGTGGCTTTGTCTTCTTCTTTTGCCAATTCCACTAATTGTTTGGG** | Ins_k3_16_TCC | 0.516109722420385 | 0.03122942121968584 | 82.26689321767509 |
| **GATCCAGATCTAACTTTCGGGGGTGGCTTTGTCTTCTTCTTTTGCCAATTCCACTAATTGTTTGGG** | Ins_k3_16_TCG | -0.4214778245446033 | 0.04457569900809816 | 32.213357968085994 |
| **GATCCAGATCTAACTTTCGGGGTGGCTTTGTCTTCTTCTTTTGCCAATTCCACTAATTGTTTGGG** | Ins_k2_16_TC | 0.8241863310865707 | 0.18697190572947645 | 100 |
| **GATCCAGATCTAACTTTCTGGGGTGGCTTTGTCTTCTTCTTTTGCCAATTCCACTAATTGTTTGGG** | Ins_k3_15_TTC;Ins_k3_16_TCT | -0.485696299870177 | 0.04362256716635095 | 30.209690032843387 |
| **GATCCAGATCTAACTTTCTTCTTCTTTTGCCAATTCCACTAATTGTTTGGG** | Del_k12_17-28 | -3.004757233576384 | 0.11021233974174628 | 2.4329434028077426 |
| **GATCCAGATCTAACTTTCTTCTTTTGCCAATTCCACTAATTGTTTGGG** | Del_k15_16-30;Del_k15_17-31 | -2.5282235261146244 | 0.09068757244497229 | 3.9182123188952382 |
| **GATCCAGATCTAACTTTCTTTTGCCAATTCCACTAATTGTTTGGG** | Del_k18_16-33;Del_k18_17-34 | -3.2187255791863527 | 0.11684744365866093 | 1.964295104687697 |
| **GATCCAGATCTAACTTTGAGGGGTGGCTTTGTCTTCTTCTTTTGCCAATTCCACTAATTGTTTGGG** | Ins_k3_16_TGA | 0.1752635902408337 | 0.035185991574211796 | 58.50560871085614 |
| **GATCCAGATCTAACTTTGCCAATTCCACTAATTGTTTGGG** | Del_k23_15-37;Del_k23_16-38;D | -1.8320825285285283 | 0.045945503228150525 | 7.859920613071248 |
| **GATCCAGATCTAACTTTGCGGGGTGGCTTTGTCTTCTTCTTTTGCCAATTCCACTAATTGTTTGGG** | Ins_k3_16_TGC | 0.3658476080888974 | 0.030207057910280413 | 70.789213888725655 |

| | | | | |
|---|---|---|---|---|
| GATCCAGATCTAACTTTGGCTTTGTCTTCTTCTTTTGCCAATTCCACTAATTGTTTGGG | Del_k4_17-20 | -0.47984461596258987 | 0.036201349169928954 | 30.386985823407766 |
| GATCCAGATCTAACTTTGGG | Del_k43_15-57;Del_k43_16-58;D | -6.29448510859802 | 0.2826363117945569 | 0.09066117493146676 |
| GATCCAGATCTAACTTTGGGGGGTGGCTTTGTCTTCTTCTTTTGCCAATTCCACTAATTGTTTGGG | Ins_k3_16_TGG | -4.506103866811747 | 0.3364599438577707 | 0.5421325056724704 |
| GATCCAGATCTAACTTTGGGGGTGGCTTTGTCTTCTTCTTTTGCCAATTCCACTAATTGTTTGGG | Ins_k2_16_TG | -1.2278811104606824 | 0.07822900504177224 | 14.382007186104584 |
| GATCCAGATCTAACTTTGGGGTGGCTTTGTCTTCTTCTTTTGCCAATTCCACTAATTGTTTGGG | Ins_k1_14_T;Ins_k1_15_T;Ins_k1_ | -0.11622566686877539 | 0.04091631414355742 | 43.71246789502544 |
| GATCCAGATCTAACTTTGGGTGGCTTTGTCTTCTTCTTTTGCCAATTCCACTAATTGTTTGGG | G17T | -0.26948149674593497 | 0.02976240116261957 | 37.50137271555558 |
| GATCCAGATCTAACTTTGTCGGGGTGGCTTTGTCTTCTTCTTTTGCCAATTCCACTAATTGTTTGGG | Ins_k4_16_TGTC | 0.6934463559366049 | 0.031011830726328536 | 98.22938341716626 |
| GATCCAGATCTAACTTTGTCTTCTTCTTTTGCCAATTCCACTAATTGTTTGGG | Del_k10_14-23;Del_k10_15-24;D | -2.292866193831382 | 0.08192108549914755 | 4.9579524398996275 |
| GATCCAGATCTAACTTTGTGGGGGTGGCTTTGTCTTCTTCTTTTGCCAATTCCACTAATTGTTTGGG | Ins_k4_16_TGTG | -1.3953401336962794 | 0.0780492785144969 | 12.164463837843648 |
| GATCCAGATCTAACTTTGTGGGGTGGCTTTGTCTTCTTCTTTTGCCAATTCCACTAATTGTTTGGG | Ins_k3_15_TTG;Ins_k3_16_TGT | -0.9842074198364448 | 0.05675956495700859 | 18.350404446961406 |
| GATCCAGATCTAACTTTGTTTGGG | Del_k39_16-54;Del_k39_17-55 | -5.7685715780623426 | 0.19755561126303203 | 0.15339905610004725 |
| GATCCAGATCTAACTTTTAGGGGTGGCTTTGTCTTCTTCTTTTGCCAATTCCACTAATTGTTTGGG | Ins_k3_16_TTA | -2.6655826511039646 | 0.13442721583378087 | 3.4153377283496966 |
| GATCCAGATCTAACTTTTATGGGGTGGCTTTGTCTTCTTCTTTTGCCAATTCCACTAATTGTTTGGG | Ins_k4_15_TTTA;Ins_k4_16_TTAT | -0.9888116195538084 | 0.057369473438426806 | 18.266109723774715 |
| GATCCAGATCTAACTTTTCCACTAATTGTTTGGG | Del_k29_17-45 | -3.482905283186839 | 0.09039852326550711 | 1.508255600557269 |
| GATCCAGATCTAACTTTTCGGGGTGGCTTTGTCTTCTTCTTTTGCCAATTCCACTAATTGTTTGGG | Ins_k3_16_TTC | 0.025898227039716115 | 0.03641813919257468 | 50.38821215156919 |
| GATCCAGATCTAACTTTTCTTCTTTTGCCAATTCCACTAATTGTTTGGG | Del_k14_17-30 | -3.185143598256224 | 0.1298412818943851 | 2.031380145236063 |
| GATCCAGATCTAACTTTTCTTTTGCCAATTCCACTAATTGTTTGGG | Del_k17_17-33 | -4.184762235156849 | 0.1710776586979389 | 0.747588040771616 |
| GATCCAGATCTAACTTTTGAGGGGTGGCTTTGTCTTCTTCTTTTGCCAATTCCACTAATTGTTTGGG | Ins_k4_16_TTGA | 0.3043516849120115 | 0.03864498993467695 | 66.56711718865044 |
| GATCCAGATCTAACTTTTGCCAATTCCACTAATTGTTTGGG | Del_k22_14-35;Del_k22_15-36;D | -2.085647577749844 | 0.04978464113114539 | 6.099528402046155 |
| GATCCAGATCTAACTTTTGGG | Del_k42_16-57;Del_k42_17-58 | -5.965344760901757 | 0.25292857240637645 | 0.12599844339671676 |
| GATCCAGATCTAACTTTTGGGGGTGGCTTTGTCTTCTTCTTTTGCCAATTCCACTAATTGTTTGGG | Ins_k3_16_TTG | -1.8283215432081048 | 0.0823009710089321 | 7.889537318202612 |
| GATCCAGATCTAACTTTTGGGGTGGCTTTGTCTTCTTCTTTTGCCAATTCCACTAATTGTTTGGG | Ins_k2_14_TT;Ins_k2_15_TT;Ins_I | -0.4190492739577163 | 0.04026808277528355 | 32.291684809336424 |
| GATCCAGATCTAACTTTTGTCTTCTTCTTTTGCCAATTCCACTAATTGTTTGGG | Del_k9_16-24;Del_k9_17-25 | -3.133993439946793 | 0.11962177099659974 | 2.1379888444813746 |
| GATCCAGATCTAACTTTTGTTTGGG | Del_k38_17-54 | -5.782166779464634 | 0.24276775368738143 | 0.15132767734733427 |
| GATCCAGATCTAACTTTTTCGGGGTGGCTTTGTCTTCTTCTTTTGCCAATTCCACTAATTGTTTGGG | Ins_k4_16_TTTC | 0.25883528646214427 | 0.04191099889405775 | 63.6051423261702 |
| GATCCAGATCTAACTTTTTGCCAATTCCACTAATTGTTTGGG | Del_k21_16-36;Del_k21_17-37 | -2.3066319467488228 | 0.07388667147716826 | 4.890170099480622 |
| GATCCAGATCTAACTTTTTGGG | Del_k41_17-57 | -5.880759726605008 | 0.19666648790628685 | 0.13711974390507298 |
| GATCCAGATCTAACTTTTTGGGGTGGCTTTGTCTTCTTCTTTTGCCAATTCCACTAATTGTTTGGG | Ins_k3_14_TTT;Ins_k3_15_TTT;In | -1.0213800273743856 | 0.05643423509560386 | 17.680794737691027 |
| GATCCAGATCTAACTTTTTGTCTTCTTCTTTTGCCAATTCCACTAATTGTTTGGG | Del_k8_17-24 | -3.7815916535060166 | 0.19035930545476695 | 1.1188119700479404 |
| GATCCAGATCTAACTTTTTTGCCAATTCCACTAATTGTTTGGG | Del_k20_17-36 | -2.528328935324125 | 0.07163963223272532 | 3.917799324999111 |
| GATCCAGATCTAAG | Del_k49_14-62 | -6.165657171351324 | 0.20124540817888353 | 0.10312657759526253 |
| GATCCAGATCTAAGAACTTGGGGTGGCTTTGTCTTCTTCTTTTGCCAATTCCACTAATTGTTTGGG | Ins_k3_11_AAG;Ins_k3_12_AGA; | 0.43288879358824994 | 0.031122187771219802 | 75.69770389828378 |
| GATCCAGATCTAAGACCTTGGGGTGGCTTTGTCTTCTTCTTTTGCCAATTCCACTAATTGTTTGGG | Ins_k3_13_GAC | 0.5025828403993138 | 0.02941057533077662 | 81.16157129246515 |
| GATCCAGATCTAAGACTTGGGGTGGCTTTGTCTTCTTCTTTTGCCAATTCCACTAATTGTTTGGG | Ins_k2_12_AG;Ins_k2_13_GA | 0.48146451454060313 | 0.03581353116997442 | 79.46554641849937 |
| GATCCAGATCTAAGAGCTTGGGGTGGCTTTGTCTTCTTCTTTTGCCAATTCCACTAATTGTTTGGG | Ins_k3_13_GAG | 0.510281259050517 | 0.03064198967789523 | 81.78879827687351 |
| GATCCAGATCTAAGATCTTGGGGTGGCTTTGTCTTCTTCTTTTGCCAATTCCACTAATTGTTTGGG | Ins_k3_13_GAT | 0.06653186096114172 | 0.0381369662328497 | 52.47783530372704 |
| GATCCAGATCTAAGCACTTGGGGTGGCTTTGTCTTCTTCTTTTGCCAATTCCACTAATTGTTTGGG | Ins_k3_12_AGC;Ins_k3_13_GCA | 0.20277668968730733 | 0.034686163460131696 | 60.13762733901848 |
| GATCCAGATCTAAGCCAATTCCACTAATTGTTTGGG | Del_k27_14-40 | -2.456324322069095 | 0.05086573049684104 | 4.210303404738455 |
| GATCCAGATCTAAGCCCTTGGGGTGGCTTTGTCTTCTTCTTTTGCCAATTCCACTAATTGTTTGGG | Ins_k3_13_GCC | 0.5675681310039391 | 0.03100237396123251 | 86.61102923271251 |
| GATCCAGATCTAAGCCTTGGGGTGGCTTTGTCTTCTTCTTTTGCCAATTCCACTAATTGTTTGGG | Ins_k2_13_GC | 0.7420852982323212 | 0.03289377587358283 | 100 |
| GATCCAGATCTAAGCGCTTGGGGTGGCTTTGTCTTCTTCTTTTGCCAATTCCACTAATTGTTTGGG | Ins_k3_13_GCG | 0.47597655294321206 | 0.03269335550307034 | 79.03063702600062 |
| GATCCAGATCTAAGCTCTTGGGGTGGCTTTGTCTTCTTCTTTTGCCAATTCCACTAATTGTTTGGG | Ins_k3_13_GCT | -0.04664230146143822 | 0.03943500438035073 | 46.86245085424447 |
| GATCCAGATCTAAGCTTGGGGTGGCTTTGTCTTCTTCTTTTGCCAATTCCACTAATTGTTTGGG | Ins_k1_13_G | 0.25878831181770023 | 0.032920243201181525 | 63.60215456739963 |
| GATCCAGATCTAAGCTTTGTCTTCTTCTTTTGCCAATTCCACTAATTGTTTGGG | Del_k9_14-22 | -0.8996489283211878 | 0.04490478234639747 | 19.969579816680262 |
| GATCCAGATCTAAGG | Del_k48_14-61 | -6.054223788739201 | 0.2994421862704965 | 0.1152830634167376 |
| GATCCAGATCTAAGGACTTGGGGTGGCTTTGTCTTCTTCTTTTGCCAATTCCACTAATTGTTTGGG | Ins_k3_12_AGG;Ins_k3_13_GGA | 0.6114809889334742 | 0.031821603790826415 | 90.49911066340998 |
| GATCCAGATCTAAGGCCTTGGGGTGGCTTTGTCTTCTTCTTTTGCCAATTCCACTAATTGTTTGGG | Ins_k3_13_GGC | 0.6090830871343433 | 0.03255810756838656 | 90.28236265720464 |
| GATCCAGATCTAAGGCTTGGGGTGGCTTTGTCTTCTTCTTTTGCCAATTCCACTAATTGTTTGGG | Ins_k2_13_GG | 0.41902344445656836 | 0.03582546890972684 | 74.65537165430025 |
| GATCCAGATCTAAGGCTTTGTCTTCTTCTTTTGCCAATTCCACTAATTGTTTGGG | Del_k8_14-21 | 0.11541554791097586 | 0.028393615940349053 | 55.10688057661527 |
| GATCCAGATCTAAGGG | Del_k47_14-60 | -6.9420532636075745 | 0.6260400933323623 | 0.047444521033248935 |
| GATCCAGATCTAAGGGCTTGGGGTGGCTTTGTCTTCTTCTTTTGCCAATTCCACTAATTGTTTGGG | Ins_k3_13_GGG | -1.5378935162516854 | 0.07107209203342742 | 10.548308534121439 |
| GATCCAGATCTAAGGGGTGGCTTTGTCTTCTTCTTTTGCCAATTCCACTAATTGTTTGGG | Del_k3_14-16 | 0.15354879912125075 | 0.03143981876576464 | 57.24886597388404 |
| GATCCAGATCTAAGGGTGGCTTTGTCTTCTTCTTTTGCCAATTCCACTAATTGTTTGGG | Del_k4_14-17 | 0.25795457826240475 | 0.027235066459546733 | 63.54914941610468 |
| GATCCAGATCTAAGGTCTTGGGGTGGCTTTGTCTTCTTCTTTTGCCAATTCCACTAATTGTTTGGG | Ins_k3_13_GGT | -0.4579588375699126 | 0.042109709878621004 | 31.059359534184644 |
| GATCCAGATCTAAGGTGGCTTTGTCTTCTTCTTTTGCCAATTCCACTAATTGTTTGGG | Del_k5_14-18 | -0.04338533654741061 | 0.028958017808629125 | 47.01532903675837 4 |
| GATCCAGATCTAAGTACTTGGGGTGGCTTTGTCTTCTTCTTTTGCCAATTCCACTAATTGTTTGGG | Ins_k3_12_AGT;Ins_k3_13_GTA | -0.31045247502296913 | 0.039412739888488015 | 35.995954594883145 |
| GATCCAGATCTAAGTCCTTGGGGTGGCTTTGTCTTCTTCTTTTGCCAATTCCACTAATTGTTTGGG | Ins_k3_13_GTC | 0.4598086075365915 | 0.0334034256918963 | 77.76314795841417 |
| GATCCAGATCTAAGTCTTCTTCTTTTGCCAATTCCACTAATTGTTTGGG | Del_k14_14-27 | -0.5622252320474956 | 0.034429548402899336 | 27.984024751207844 |
| GATCCAGATCTAAGTCTTGGGGTGGCTTTGTCTTCTTCTTTTGCCAATTCCACTAATTGTTTGGG | Ins_k2_13_GT | 0.12764917277317056 | 0.037809018355092414 | 55.78517803970162 |
| GATCCAGATCTAAGTGCTTGGGGTGGCTTTGTCTTCTTCTTTTGCCAATTCCACTAATTGTTTGGG | Ins_k3_13_GTG | 0.2327628142792756 | 0.03736018206574172 | 61.96823091224435 |
| GATCCAGATCTAAGTGGCTTTGTCTTCTTCTTTTGCCAATTCCACTAATTGTTTGGG | Del_k6_14-19 | 0.00887219149619134 | 0.02938425453982568 | 49.537562802697146 |
| GATCCAGATCTAAGTTCTTGGGGTGGCTTTGTCTTCTTCTTTTGCCAATTCCACTAATTGTTTGGG | Ins_k3_13_GTT | -0.3055532722158844 | 0.04080127694403283 | 36.17273877385056 |
| GATCCAGATCTAAGTTGGGGTGGCTTTGTCTTCTTCTTTTGCCAATTCCACTAATTGTTTGGG | C14G | -0.17338367337017113 | 0.03580435943107042 | 41.284014272980116 |
| GATCCAGATCTAAGTTTGGG | Del_k43_14-56 | -6.246326705525122 | 0.2829660886898619 | 0.09513411265764851 |
| GATCCAGATCTAATAACTTGGGGTGGCTTTGTCTTCTTCTTTTGCCAATTCCACTAATTGTTTGGG | Ins_k3_10_TAA;Ins_k3_11_AAT;Ir | -0.5921292521766186 | 0.05392201003296781 | 27.159578478261114 |
| GATCCAGATCTAATAATTGTTTGGG | Del_k38_14-51 | -5.567501042634117 | 0.20290140375774743 | 0.18756271531210333 |
| GATCCAGATCTAATACCTTGGGGTGGCTTTGTCTTCTTCTTTTGCCAATTCCACTAATTGTTTGGG | Ins_k3_13_TAC | 0.1468638346823371 | 0.0374141440289623 | 56.86743568510784 |
| GATCCAGATCTAATACTTGGGGTGGCTTTGTCTTCTTCTTTTGCCAATTCCACTAATTGTTTGGG | Ins_k2_12_AT;Ins_k2_13_TA | -0.056093975386597994 | 0.0409097797825386 | 46.42160887858789 |
| GATCCAGATCTAATAGCTTGGGGTGGCTTTGTCTTCTTCTTTTGCCAATTCCACTAATTGTTTGGG | Ins_k3_13_TAG | -0.7692419289325709 | 0.05700230117276027 | 22.751182131425775 |
| GATCCAGATCTAATATCTTGGGGTGGCTTTGTCTTCTTCTTTTGCCAATTCCACTAATTGTTTGGG | Ins_k3_13_TAT | -0.39801061448537656 | 0.045558478020146884 | 32.97825550913604 |
| GATCCAGATCTAATCACTTGGGGTGGCTTTGTCTTCTTCTTTTGCCAATTCCACTAATTGTTTGGG | Ins_k3_12_ATC;Ins_k3_13_TCA | 0.014742018964177483 | 0.03450458768281135 | 49.82919482702206 |
| GATCCAGATCTAATCCACTAATTGTTTGGG | Del_k33_14-46 | -3.6697087409829026 | 0.08671721682627809 | 1.2512590617983752 |
| GATCCAGATCTAATCCCTTGGGGTGGCTTTGTCTTCTTCTTTTGCCAATTCCACTAATTGTTTGGG | Ins_k3_13_TCC | 0.34612636497179594 | 0.033679688375669675 | 69.40683847570983 |
| GATCCAGATCTAATCCTTGGGGTGGCTTTGTCTTCTTCTTTTGCCAATTCCACTAATTGTTTGGG | Ins_k2_13_TC | 0.5009650900602449 | 0.037513218403071785 | 81.03037828036959 |
| GATCCAGATCTAATCGCTTGGGGTGGCTTTGTCTTCTTCTTTTGCCAATTCCACTAATTGTTTGGG | Ins_k3_13_TCG | 0.7240444933841699 | 0.029779007740217776 | 100 |

| | | | | |
|---|---|---|---|---|
| GATCCAGATCTAATCTCTTGGGGTGGCTTTGTCTTCTTCTTTTGCCAATTCCACTAATTGTTTGGG | Ins_k3_13_TCT | -0.26623096641262334 | 0.03895823707056818 | 37.62347039871323 |
| GATCCAGATCTAATCTTCTTCTTTTGCCAATTCCACTAATTGTTTGGG | Del_k15_14-28 | -1.0815998480623077 | 0.04356843961801192 | 16.647485545573456 |
| GATCCAGATCTAATCTTCTTTTGCCAATTCCACTAATTGTTTGGG | Del_k18_14-31 | -0.9878843812600259 | 0.037371647833951166 | 18.28305461495096 |
| GATCCAGATCTAATCTTGGGGTGGCTTTGTCTTCTTCTTTTGCCAATTCCACTAATTGTTTGGG | Ins_k1_13_T | 0.18900520179151736 | 0.03687961745363655 | 59.315119310992465 |
| GATCCAGATCTAATCTTTTGCCAATTCCACTAATTGTTTGGG | Del_k21_14-34 | -2.0464078104097796 | 0.05703161999511878 | 6.3436304096951535 |
| GATCCAGATCTAATGACTTGGGGTGGCTTTGTCTTCTTCTTTTGCCAATTCCACTAATTGTTTGGG | Ins_k3_12_ATG;Ins_k3_13_TGA | 0.454121184249467 | 0.02998545375978127 | 77.3221313327373 |
| GATCCAGATCTAATGCCAATTCCACTAATTGTTTGGG | Del_k26_14-39 | -1.8197537697088964 | 0.04136676373676217 | 7.957423488348582 |
| GATCCAGATCTAATGCCTTGGGGTGGCTTTGTCTTCTTCTTTTGCCAATTCCACTAATTGTTTGGG | Ins_k3_13_TGC | 0.5455816224280958 | 0.03355733648267665 | 84.72753668206009 |
| GATCCAGATCTAATGCTTGGGGTGGCTTTGTCTTCTTCTTTTGCCAATTCCACTAATTGTTTGGG | Ins_k2_13_TG | 0.4290789113269502 | 0.034877602743777415 | 75.40985324571228 |
| GATCCAGATCTAATGGCCTTGGGGTGGCTTTGTCTTCTTCTTTTGCCAATTCCACTAATTGTTTGGG | Ins_k4_13_TGGC | 0.920289682817578 | 0.03134245424576251 | 100 |
| GATCCAGATCTAATGGCTTGGGGTGGCTTTGTCTTCTTCTTTTGCCAATTCCACTAATTGTTTGGG | Ins_k3_13_TGG | 0.3569151033101845 | 0.03494177187108893 | 70.15970463030045 |
| GATCCAGATCTAATGGCTTTGTCTTCTTCTTTTGCCAATTCCACTAATTGTTTGGG | Del_k7_14-20 | 0.03759037514998087 | 0.029617233831645478 | 50.980816247441204 |
| GATCCAGATCTAATGGG | Del_k46_14-59 | -6.229120225205919 | 0.24415474541204832 | 0.09678519985921669 |
| GATCCAGATCTAATGGGGTGGCTTTGTCTTCTTCTTTTGCCAATTCCACTAATTGTTTGGG | Del_k2_14-15 | 0.11564026444871534 | 0.03235864627594608 | 55.11926539550876 |
| GATCCAGATCTAATGTCTTCTTCTTTTGCCAATTCCACTAATTGTTTGGG | Del_k13_14-26 | -0.7301860147757946 | 0.032978736564749964 | 23.65733039110544 |
| GATCCAGATCTAATGTCTTGGGGTGGCTTTGTCTTCTTCTTTTGCCAATTCCACTAATTGTTTGGG | Ins_k3_13_TGT | 0.12406378830806564 | 0.03540839504925242 | 55.58552485955832 |
| GATCCAGATCTAATGTTTGGG | Del_k42_14-55 | -5.98361148722758 | 0.20569554205402565 | 0.12371775805779904 |
| GATCCAGATCTAATTACTTGGGGTGGCTTTGTCTTCTTCTTTTGCCAATTCCACTAATTGTTTGGG | Ins_k3_12_ATT;Ins_k3_13_TTA | -0.2904704022763038 | 0.03945600285270234 | 36.72246277444773 |
| GATCCAGATCTAATTCCACTAATTGTTTGGG | Del_k32_12-43;Del_k32_13-44;D | -3.569142306144134 | 0.08702469304951838 | 1.3836386452839027 |
| GATCCAGATCTAATTCCTTGGGGTGGCTTTGTCTTCTTCTTTTGCCAATTCCACTAATTGTTTGGG | Ins_k3_13_TTC | 0.3796689420299387 | 0.03618914260920549 | 71.77440791778346 |
| GATCCAGATCTAATTCTTCTTTTGCCAATTCCACTAATTGTTTGGG | Del_k17_14-30 | -0.8707710399438852 | 0.03662146539188754 | 20.554666487691915 |
| GATCCAGATCTAATTCTTGGGGTGGCTTTGTCTTCTTCTTTTGCCAATTCCACTAATTGTTTGGG | Ins_k2_13_TT | -0.2184926865487311 | 0.04506241496557087 | 39.46311145121976 |
| GATCCAGATCTAATTCTTTTGCCAATTCCACTAATTGTTTGGG | Del_k20_14-33 | -2.2846323232651002 | 0.0647452885591318 | 4.99894410701307 |
| GATCCAGATCTAATTGCCAATTCCACTAATTGTTTGGG | Del_k25_14-38 | -1.8870126538276708 | 0.0455234683746991 | 7.439817961257919 |
| GATCCAGATCTAATTGCTTGGGGTGGCTTTGTCTTCTTCTTTTGCCAATTCCACTAATTGTTTGGG | Ins_k3_13_TTG | 0.2190431953140235 | 0.037203260442242064 | 61.12385589248469 |
| GATCCAGATCTAATTGGG | Del_k45_14-58 | -6.528511596442642 | 0.31585121494169865 | 0.07174389165876972 |
| GATCCAGATCTAATTGGGGTGGCTTTGTCTTCTTCTTTTGCCAATTCCACTAATTGTTTGGG | Del_k1_14-14 | -0.15594203558875747 | 0.0350260379004412 | 42.01039128424044 |
| GATCCAGATCTAATTGTCTTCTTCTTTTGCCAATTCCACTAATTGTTTGGG | Del_k12_14-25 | -1.088083538340256 | 0.04354532722795915 | 16.539897565858897 |
| GATCCAGATCTAATTGTTTGGG | Del_k41_10-50;Del_k41_11-51;D | -5.4755580697322275 | 0.1601686585457129 | 0.2056254364458483 |
| GATCCAGATCTAATTTCTTGGGGTGGCTTTGTCTTCTTCTTTTGCCAATTCCACTAATTGTTTGGG | Ins_k3_13_TTT | -0.7746229913770615 | 0.05714981166494008 | 22.629085399377878 |
| GATCCAGATCTAATTTGCCAATTCCACTAATTGTTTGGG | Del_k24_14-37 | -1.7973766412550034 | 0.04371404209488673 | 8.137495003664288 |
| GATCCAGATCTAATTTGGG | Del_k44_14-57 | -5.642130283539855 | 0.25150949680945733 | 0.17407461519269296 |
| GATCCAGATCTAATTTGGGGTGGCTTTGTCTTCTTCTTTTGCCAATTCCACTAATTGTTTGGG | C14T | -0.416567802807422 | 0.03445187848175248 | 32.37191519711197 |
| GATCCAGATCTAATTTGTCTTCTTCTTTTGCCAATTCCACTAATTGTTTGGG | Del_k11_14-24 | -1.304756672905653 | 0.04849588028363247 | 13.31781166092445 |
| GATCCAGATCTAATTTTGCCAATTCCACTAATTGTTTGGG | Del_k23_14-36 | -2.0250339365001597 | 0.06040258478177104 | 6.480677765268552 |
| GATCCAGATCTACAAACTTGGGGTGGCTTTGTCTTCTTCTTTTGCCAATTCCACTAATTGTTTGGG | Ins_k3_11_ACA;Ins_k3_12_CAA | 0.49714911405027573 | 0.02866901514507167 | 80.72175751998465 |
| GATCCAGATCTACAACTTGGGGTGGCTTTGTCTTCTTCTTTTGCCAATTCCACTAATTGTTTGGG | Ins_k2_11_AC;Ins_k2_12_CA | 0.6013594677904328 | 0.032743973304229346 | 89.58774200043092 |
| GATCCAGATCTACAATTCCACTAATTGTTTGGG | Del_k30_13-42 | -1.4573262719991429 | 0.03395682862863953 | 11.433329872435817 |
| GATCCAGATCTACACACTTGGGGTGGCTTTGTCTTCTTCTTTTGCCAATTCCACTAATTGTTTGGG | Ins_k3_12_CAC | 0.23067200965357482 | 0.029648044186507898 | 61.838802800003606 |
| GATCCAGATCTACACGACTTGGGGTGGCTTTGTCTTCTTCTTTTGCCAATTCCACTAATTGTTTGGG | Ins_k4_12_CACG | 0.8807989400419667 | 0.028373997879658205 | 100 |
| GATCCAGATCTACACTAATTGTTTGGG | Del_k36_13-48 | -3.537758171424124 | 0.06511721508445364 | 1.4277515488433372 |
| GATCCAGATCTACACTTGGGGTGGCTTTGTCTTCTTCTTTTGCCAATTCCACTAATTGTTTGGG | Ins_k1_12_C | 0.5114482786438264 | 0.02753195109484937 | 81.88430312415058 |
| GATCCAGATCTACAGACTTGGGGTGGCTTTGTCTTCTTCTTTTGCCAATTCCACTAATTGTTTGGG | Ins_k3_12_CAG | 0.6554465199055881 | 0.026382744671005983 | 94.56671409747605 |
| GATCCAGATCTACATACTTGGGGTGGCTTTGTCTTCTTCTTTTGCCAATTCCACTAATTGTTTGGG | Ins_k3_12_CAT | 0.4913054093208846 | 0.028222953383892344 | 80.25141900197198 |
| GATCCAGATCTACCAACTTGGGGTGGCTTTGTCTTCTTCTTTTGCCAATTCCACTAATTGTTTGGG | Ins_k3_11_ACC;Ins_k3_12_CCA | 0.4911609581641434 | 0.031288448316637435 | 80.23982742889525 |
| GATCCAGATCTACCAATTCCACTAATTGTTTGGG | Del_k29_13-41 | -1.517716576770559 | 0.036921477991344556 | 10.763302786260018 |
| GATCCAGATCTACCACTAATTGTTTGGG | Del_k35_13-47 | -3.3620668582464686 | 0.06904757174647212 | 1.7019798252971778 |
| GATCCAGATCTACCACTTGGGGTGGCTTTGTCTTCTTCTTTTGCCAATTCCACTAATTGTTTGGG | Ins_k2_12_CC | 0.7370193713255002 | 0.033207667617195034 | 100 |
| GATCCAGATCTACCATACTTGGGGTGGCTTTGTCTTCTTCTTTTGCCAATTCCACTAATTGTTTGGG | Ins_k4_12_CCAT | 0.6330635755641789 | 0.03128607603519954 | 92.47354562438821 |
| GATCCAGATCTACCCACTTGGGGTGGCTTTGTCTTCTTCTTTTGCCAATTCCACTAATTGTTTGGG | Ins_k3_12_CCC | 0.09377620074611381 | 0.031711888964057924 | 53.92721330382886 |
| GATCCAGATCTACCGACTTGGGGTGGCTTTGTCTTCTTCTTTTGCCAATTCCACTAATTGTTTGGG | Ins_k3_12_CCG | 0.8034400187845507 | 0.028052729608625676 | 100 |
| GATCCAGATCTACCGGACTTGGGGTGGCTTTGTCTTCTTCTTTTGCCAATTCCACTAATTGTTTGGG | Ins_k4_12_CCGG | 1.0621659471107305 | 0.03006993954473548 | 100 |
| GATCCAGATCTACCTACTTGGGGTGGCTTTGTCTTCTTCTTTTGCCAATTCCACTAATTGTTTGGG | Ins_k3_12_CCT | 0.5939911912482969 | 0.029666502232999492 | 88.93006070694898 |
| GATCCAGATCTACCTTGGGGTGGCTTTGTCTTCTTCTTTTGCCAATTCCACTAATTGTTTGGG | A13C | 0.31236315843290086 | 0.02889037468571316 | 67.10255986388827 |
| GATCCAGATCTACGAACTTGGGGTGGCTTTGTCTTCTTCTTTTGCCAATTCCACTAATTGTTTGGG | Ins_k3_11_ACG;Ins_k3_12_CGA | 0.8094486751808698 | 0.03155020503420626 | 100 |
| GATCCAGATCTACGACTTGGGGTGGCTTTGTCTTCTTCTTTTGCCAATTCCACTAATTGTTTGGG | Ins_k2_12_CG | 0.8534519491689492 | 0.030093118294164486 | 100 |
| GATCCAGATCTACGCACTTGGGGTGGCTTTGTCTTCTTCTTTTGCCAATTCCACTAATTGTTTGGG | Ins_k3_12_CGC | 0.797137179422529 | 0.028278134686039066 | 100 |
| GATCCAGATCTACGGACTTGGGGTGGCTTTGTCTTCTTCTTTTGCCAATTCCACTAATTGTTTGGG | Ins_k3_12_CGG | 0.8789888213103421 | 0.029312935008375028 | 100 |
| GATCCAGATCTACGTACTTGGGGTGGCTTTGTCTTCTTCTTTTGCCAATTCCACTAATTGTTTGGG | Ins_k3_12_CGT | 0.7668370977952295 | 0.03230981341258919 | 100 |
| GATCCAGATCTACTAACTTGGGGTGGCTTTGTCTTCTTCTTTTGCCAATTCCACTAATTGTTTGGG | Ins_k3_9_CTA;Ins_k3_10_TAC;Ins | -0.042112478195245995 | 0.03882552163701055 | 47.07521099354209 |
| GATCCAGATCTACTAATTGTTTGGG | Del_k38_12-49;Del_k38_13-50 | -3.9415167981064827 | 0.09345799638257649 | 0.953460040258447 |
| GATCCAGATCTACTACTTGGGGTGGCTTTGTCTTCTTCTTTTGCCAATTCCACTAATTGTTTGGG | Ins_k2_12_CT | 0.48976560779635125 | 0.03089762128614525 | 80.12794283343977 |
| GATCCAGATCTACTCAACTTGGGGTGGCTTTGTCTTCTTCTTTTGCCAATTCCACTAATTGTTTGGG | Ins_k4_11_ACTC;Ins_k4_12_CTC | 0.7974604419552842 | 0.02965605997939862 | 100 |
| GATCCAGATCTACTCACTTGGGGTGGCTTTGTCTTCTTCTTTTGCCAATTCCACTAATTGTTTGGG | Ins_k3_12_CTC | 0.32462755957668 | 0.03181775361656334 | 67.93059990429497 |
| GATCCAGATCTACTGACTTGGGGTGGCTTTGTCTTCTTCTTTTGCCAATTCCACTAATTGTTTGGG | Ins_k3_12_CTG | 0.677730586547842 | 0.029764382068792228 | 96.6977003939166 |
| GATCCAGATCTACTTACTTGGGGTGGCTTTGTCTTCTTCTTTTGCCAATTCCACTAATTGTTTGGG | Ins_k3_12_CTT | 0.30410969134637067 | 0.03380473696960687 | 66.551010323565 |
| GATCCAGATCTACTTCTTCTTTTGCCAATTCCACTAATTGTTTGGG | Del_k17_13-29 | 0.17967689792655983 | 0.02670557568939658 | 58.76438256774604 |
| GATCCAGATCTACTTCTTTTGCCAATTCCACTAATTGTTTGGG | Del_k20_13-32 | -0.29026624908452564 | 0.031728814903781814 | 36.72996054775422 |
| GATCCAGATCTACTTGGGGTGGCTTTGTCTTCTTCTTTTGCCAATTCCACTAATTGTTTGGG | Del_k1_12-12;Del_k1_13-13 | 0.46458996588289403 | 0.027806176783718477 | 78.13585174006491 |
| GATCCAGATCTACTTTGTCTTCTTCTTTTGCCAATTCCACTAATTGTTTGGG | Del_k11_13-23 | 0.31841020948302334 | 0.02612994112064447 | 67.50956180952043 |
| GATCCAGATCTACTTTTGCCAATTCCACTAATTGTTTGGG | Del_k23_13-35 | -0.6188495315369126 | 0.029846641492641674 | 26.443476778698045 |
| GATCCAGATCTAG | Del_k50_13-62 | -6.029478143707297 | 0.18460760728899775 | 0.11817140675420357 |
| GATCCAGATCTAGAAACTTGGGGTGGCTTTGTCTTCTTCTTTTGCCAATTCCACTAATTGTTTGGG | Ins_k3_11_AGA;Ins_k3_12_GAA | 0.2952657847823704 | 0.03421184690863354 | 65.96503438407427 |
| GATCCAGATCTAGAACTTGGGGTGGCTTTGTCTTCTTCTTTTGCCAATTCCACTAATTGTTTGGG | Ins_k2_11_AG;Ins_k2_12_GA | 0.36149267540373886 | 0.03607052387910683 | 70.4816019282616 |
| GATCCAGATCTAGACACTTGGGGTGGCTTTGTCTTCTTCTTTTGCCAATTCCACTAATTGTTTGGG | Ins_k3_12_GAC | 0.021131794038765528 | 0.03332188592662648 | 50.14861158794907 |

| | | | | |
|---|---|---|---|---|
| GATCCAGATCTAGACTTGGGGTGGCTTTGTCTTCTTCTTTTGCCAATTCCACTAATTGTTTGGG | Ins_k1_12_G | 0.180837678000335 | 0.035389155972661665 | 58.83263469727072 |
| GATCCAGATCTAGAGACTTGGGGTGGCTTTGTCTTCTTCTTTTGCCAATTCCACTAATTGTTTGGG | Ins_k3_12_GAG | 0.4322664064104029 | 0.02832038845666433 | 75.6506052762946 |
| GATCCAGATCTAGATACTTGGGGTGGCTTTGTCTTCTTCTTTTGCCAATTCCACTAATTGTTTGGG | Ins_k3_12_GAT | -0.542915573112644 | 0.039169703430017744 | 28.52963757037498 |
| GATCCAGATCTAGCAACTTGGGGTGGCTTTGTCTTCTTCTTTTGCCAATTCCACTAATTGTTTGGG | Ins_k3_11_AGC;Ins_k3_12_GCA | 0.14833980476575648 | 0.033056117044470514 | 56.951432291890406 |
| GATCCAGATCTAGCACTTGGGGTGGCTTTGTCTTCTTCTTTTGCCAATTCCACTAATTGTTTGGG | Ins_k2_12_GC | -0.044842867018671795 | 0.03366510718657365 | 46.946852677383674 |
| GATCCAGATCTAGCCAATTCCACTAATTGTTTGGG | Del_k28_13-40 | -3.2783340936963814 | 0.07408659932470775 | 1.8506278152291036 |
| GATCCAGATCTAGCCACTTGGGGTGGCTTTGTCTTCTTCTTTTGCCAATTCCACTAATTGTTTGGG | Ins_k3_12_GCC | 0.2630667127784867 | 0.029925788623452156 | 63.874853027313165 |
| GATCCAGATCTAGCGAACTTGGGGTGGCTTTGTCTTCTTCTTTTGCCAATTCCACTAATTGTTTGGG | Ins_k4_11_AGCG;Ins_k4_12_GC | 0.5363075606376233 | 0.030418225694165187 | 83.94540066622183 |
| GATCCAGATCTAGCGACTTGGGGTGGCTTTGTCTTCTTCTTTTGCCAATTCCACTAATTGTTTGGG | Ins_k3_12_GCG | 0.6171745676290723 | 0.034071723851097524 | 91.01584410721257 |
| GATCCAGATCTAGCTAACTTGGGGTGGCTTTGTCTTCTTCTTTTGCCAATTCCACTAATTGTTTGGG | Ins_k4_9_CTAG;Ins_k4_10_TAGC | -1.0566454290123013 | 0.05786412449437379 | 17.06814064879209 |
| GATCCAGATCTAGCTACTTGGGGTGGCTTTGTCTTCTTCTTTTGCCAATTCCACTAATTGTTTGGG | Ins_k3_12_GCT | 0.013325503552881202 | 0.03078180762331172 | 49.75866097255697 |
| GATCCAGATCTAGCTTGGGGTGGCTTTGTCTTCTTCTTTTGCCAATTCCACTAATTGTTTGGG | A13G | -0.17895367375852733 | 0.030003646900767907 | 41.054701526357455 |
| GATCCAGATCTAGCTTTGTCTTCTTCTTTTGCCAATTCCACTAATTGTTTGGG | Del_k10_13-22 | -1.2945469802611607 | 0.04753631378432343 | 13.45447890279081 |
| GATCCAGATCTAGG | Del_k49_13-61 | -6.1470033469960486 | 0.31784045041742676 | 0.10506833697422452 |
| GATCCAGATCTAGGAACTTGGGGTGGCTTTGTCTTCTTCTTTTGCCAATTCCACTAATTGTTTGGG | Ins_k3_11_AGG;Ins_k3_12_GGA | 0.13957049831881818 | 0.03384437406382138 | 56.45419114603717 |
| GATCCAGATCTAGGACTTGGGGTGGCTTTGTCTTCTTCTTTTGCCAATTCCACTAATTGTTTGGG | Ins_k2_12_GG | 0.44993862383684413 | 0.03524285508596377 | 76.99940223466021 |
| GATCCAGATCTAGGCACTTGGGGTGGCTTTGTCTTCTTCTTTTGCCAATTCCACTAATTGTTTGGG | Ins_k3_12_GGC | -0.24399231556413103 | 0.03420403158450199 | 38.46953845787192 |
| GATCCAGATCTAGGCTTTGTCTTCTTCTTTTGCCAATTCCACTAATTGTTTGGG | Del_k9_13-21 | -0.9803532614461699 | 0.0405503035110228 | 18.421266280873173 |
| GATCCAGATCTAGGG | Del_k48_13-60 | -6.290068535737592 | 0.25254435724617097 | 0.09106247214303814 |
| GATCCAGATCTAGGGACTTGGGGTGGCTTTGTCTTCTTCTTTTGCCAATTCCACTAATTGTTTGGG | Ins_k3_12_GGG | -1.259713335382425 | 0.06517363190440217 | 13.931405769882923 |
| GATCCAGATCTAGGGGTGGCTTTGTCTTCTTCTTTTGCCAATTCCACTAATTGTTTGGG | Del_k4_13-16 | -0.059387624409742545 | 0.03768817048315563 | 46.26896390932169 |
| GATCCAGATCTAGGGTGGCTTTGTCTTCTTCTTTTGCCAATTCCACTAATTGTTTGGG | Del_k5_13-17 | -0.24802858885070767 | 0.0367468730014075 | 38.314577829548156 |
| GATCCAGATCTAGGTACTTGGGGTGGCTTTGTCTTCTTCTTTTGCCAATTCCACTAATTGTTTGGG | Ins_k3_12_GGT | -1.80883085911097 | 0.07561032954993754 | 8.04481814680268 |
| GATCCAGATCTAGGTGGCTTTGTCTTCTTCTTTTGCCAATTCCACTAATTGTTTGGG | Del_k6_13-18 | -0.2501719723718108 | 0.02929349943931821 | 38.23254294232932 |
| GATCCAGATCTAGTAACTTGGGGTGGCTTTGTCTTCTTCTTTTGCCAATTCCACTAATTGTTTGGG | Ins_k3_10_TAG;Ins_k3_11_AGT;I | -1.1336782437922355 | 0.05797791447487326 | 15.802699676024604 |
| GATCCAGATCTAGTACTTGGGGTGGCTTTGTCTTCTTCTTTTGCCAATTCCACTAATTGTTTGGG | Ins_k2_12_GT | -0.4610887979298593 | 0.04598894708706206 | 30.962296950297993 |
| GATCCAGATCTAGTCACTTGGGGTGGCTTTGTCTTCTTCTTTTGCCAATTCCACTAATTGTTTGGG | Ins_k3_12_GTC | 0.09763289885698656 | 0.03522275314943518 | 54.135595861518446 |
| GATCCAGATCTAGTCTTCTTCTTTTGCCAATTCCACTAATTGTTTGGG | Del_k15_13-27 | -1.0726973650414402 | 0.04094050893482005 | 16.79635115658612 |
| GATCCAGATCTAGTGACTTGGGGTGGCTTTGTCTTCTTCTTTTGCCAATTCCACTAATTGTTTGGG | Ins_k3_12_GTG | 0.12221076312555357 | 0.03392155288381187 | 55.48261885536473 |
| GATCCAGATCTAGTGGCTTTGTCTTCTTCTTTTGCCAATTCCACTAATTGTTTGGG | Del_k7_13-19 | -0.14119206056140438 | 0.030302567876443132 | 42.634635985845 |
| GATCCAGATCTAGTTACTTGGGGTGGCTTTGTCTTCTTCTTTTGCCAATTCCACTAATTGTTTGGG | Ins_k3_12_GTT | -1.4092385637768317 | 0.057174685450411114 | 11.996566346170555 |
| GATCCAGATCTAGTTTACTTGGGGTGGCTTTGTCTTCTTCTTTTGCCAATTCCACTAATTGTTTGGG | Ins_k4_12_GTTT | -1.7803855199555991 | 0.0798737577411739 | 8.276941489610572 |
| GATCCAGATCTAGTTTGGG | Del_k44_13-56 | -6.40259925183168 | 0.3388876345941301 | 0.081370684100187 |
| GATCCAGATCTATAAACTTGGGGTGGCTTTGTCTTCTTCTTTTGCCAATTCCACTAATTGTTTGGG | Ins_k3_11_ATA;Ins_k3_12_TAA | -0.5173725227821866 | 0.04880283176201315 | 29.26775833501942 |
| GATCCAGATCTATAACTTGGGGTGGCTTTGTCTTCTTCTTTTGCCAATTCCACTAATTGTTTGGG | Ins_k2_10_TA;Ins_k2_11_AT;Ins_ | -0.4627847898216325 | 0.04784093001393019 | 30.909829650353622 |
| GATCCAGATCTATAATTGTTTGGG | Del_k39_13-51 | -5.834073533384755 | 0.19712315860470453 | 0.14367312878956748 |
| GATCCAGATCTATACACTTGGGGTGGCTTTGTCTTCTTCTTTTGCCAATTCCACTAATTGTTTGGG | Ins_k3_12_TAC | -0.04800627654669487 | 0.035306658200080374 | 46.7985752111411 |
| GATCCAGATCTATACTTGGGGTGGCTTTGTCTTCTTCTTTTGCCAATTCCACTAATTGTTTGGG | Ins_k1_12_T | -0.06925083446315605 | 0.0366897528512601 | 45.814846607978104 |
| GATCCAGATCTATAGACTTGGGGTGGCTTTGTCTTCTTCTTTTGCCAATTCCACTAATTGTTTGGG | Ins_k3_12_TAG | -0.34793133140849747 | 0.03840205298958678 | 34.67183560649869 |
| GATCCAGATCTATATACTTGGGGTGGCTTTGTCTTCTTCTTTTGCCAATTCCACTAATTGTTTGGG | Ins_k3_12_TAT | -0.29919482079767235 | 0.04186367923276526 | 36.40347415849101 |
| GATCCAGATCTATCAACTTGGGGTGGCTTTGTCTTCTTCTTTTGCCAATTCCACTAATTGTTTGGG | Ins_k3_11_ATC;Ins_k3_12_TCA | 0.4276768828644417 | 0.028354930430117053 | 75.30420056644658 |
| GATCCAGATCTATCACTTGGGGTGGCTTTGTCTTCTTCTTTTGCCAATTCCACTAATTGTTTGGG | Ins_k2_12_TC | 0.2119829938312887 | 0.03599615734366493 | 60.6938289790825 |
| GATCCAGATCTATCCAACTTGGGGTGGCTTTGTCTTCTTCTTTTGCCAATTCCACTAATTGTTTGGG | Ins_k4_11_ATCC;Ins_k4_12_TCC | 0.4874467295503291 | 0.032278002736337734 | 79.94235115532089 |
| GATCCAGATCTATCCACTAATTGTTTGGG | Del_k34_13-46 | -3.5939721879640936 | 0.08594905117442303 | 1.3497060754227135 |
| GATCCAGATCTATCCACTTGGGGTGGCTTTGTCTTCTTCTTTTGCCAATTCCACTAATTGTTTGGG | Ins_k3_12_TCC | 0.2970452914740843 | 0.029301332829595152 | 66.08252411005265 |
| GATCCAGATCTATCGACTTGGGGTGGCTTTGTCTTCTTCTTTTGCCAATTCCACTAATTGTTTGGG | Ins_k3_12_TCG | 0.7085812294346427 | 0.029463679229067834 | 99.727380109495 |
| GATCCAGATCTATCTACTTGGGGTGGCTTTGTCTTCTTCTTTTGCCAATTCCACTAATTGTTTGGG | Ins_k3_12_TCT | 0.3547210019274767 | 0.03127717324611996 | 70.0059358791634 |
| GATCCAGATCTATCTTCTTCTTTTGCCAATTCCACTAATTGTTTGGG | Del_k16_13-28 | -0.3921488067017031 | 0.03570494334261399 | 33.17213539211987 |
| GATCCAGATCTATCTTCTTTTGCCAATTCCACTAATTGTTTGGG | Del_k19_13-31 | -0.47311864233934814 | 0.03227768507753848 | 30.59205676632729 |
| GATCCAGATCTATCTTGGGGTGGCTTTGTCTTCTTCTTTTGCCAATTCCACTAATTGTTTGGG | A13T | -0.2091615125633004 | 0.037553400524808926 | 39.83307200895034 |
| GATCCAGATCTATCTTTTGCCAATTCCACTAATTGTTTGGG | Del_k22_13-34 | -1.6727347831442918 | 0.04223466254996804 | 9.217688043803555 |
| GATCCAGATCTATGAACTTGGGGTGGCTTTGTCTTCTTCTTTTGCCAATTCCACTAATTGTTTGGG | Ins_k3_11_ATG;Ins_k3_12_TGA | 0.132139581710208 | 0.03131635302991791 | 56.03623956432012 |
| GATCCAGATCTATGACTTGGGGTGGCTTTGTCTTCTTCTTTTGCCAATTCCACTAATTGTTTGGG | Ins_k2_12_TG | 0.315042561577045 | 0.03921096232992623 | 67.28259576043185 |
| GATCCAGATCTATGCACTTGGGGTGGCTTTGTCTTCTTCTTTTGCCAATTCCACTAATTGTTTGGG | Ins_k3_12_TGC | 0.19655360308637326 | 0.02863902109762674 | 59.76454773347462 |
| GATCCAGATCTATGCCAATTCCACTAATTGTTTGGG | Del_k27_13-39 | -1.981826640305645 | 0.04445168935099528 | 6.766827695728845 |
| GATCCAGATCTATGGACTTGGGGTGGCTTTGTCTTCTTCTTTTGCCAATTCCACTAATTGTTTGGG | Ins_k3_12_TGG | -0.004315719084641012 | 0.3934755252261934 | 48.88855479019587 |
| GATCCAGATCTATGGCTTTGTCTTCTTCTTTTGCCAATTCCACTAATTGTTTGGG | Del_k8_13-20 | 0.27438144268564635 | 0.02620488530600247 | 64.60168393060987 |
| GATCCAGATCTATGGG | Del_k47_13-59 | -6.831355753998821 | 0.4784670061771124 | 0.05299823227975869 |
| GATCCAGATCTATGGGGTGGCTTTGTCTTCTTCTTTTGCCAATTCCACTAATTGTTTGGG | Del_k3_13-15 | 0.3255972498665183 | 0.03300216327273367 | 67.99650349527708 |
| GATCCAGATCTATGTACTTGGGGTGGCTTTGTCTTCTTCTTTTGCCAATTCCACTAATTGTTTGGG | Ins_k3_12_TGT | -0.06902749350795134 | 0.03200957747909232 | 45.82508008231652 |
| GATCCAGATCTATGTCTTCTTCTTTTGCCAATTCCACTAATTGTTTGGG | Del_k14_13-26 | -0.22099875333056085 | 0.0278764216143971 | 39.36433807653527 |
| GATCCAGATCTATGTTTGGG | Del_k43_13-55 | -5.6877049579867425 | 0.25623050981352086 | 0.16631928686230224 |
| GATCCAGATCTATTAACTTGGGGTGGCTTTGTCTTCTTCTTTTGCCAATTCCACTAATTGTTTGGG | Ins_k3_10_TAT;Ins_k3_11_ATT;Ins | -0.868900057427149 | 0.04771123639574111 | 20.593159908357173 |
| GATCCAGATCTATTACTTGGGGTGGCTTTGTCTTCTTCTTTTGCCAATTCCACTAATTGTTTGGG | Ins_k2_12_TT | 0.027293230554543135 | 0.04299423845774967 | 50.458552936040135 |
| GATCCAGATCTATTCACTTGGGGTGGCTTTGTCTTCTTCTTTTGCCAATTCCACTAATTGTTTGGG | Ins_k3_12_TTC | 0.14572345434761846 | 0.03466856131869104 | 56.802622142835176 |
| GATCCAGATCTATTCCACTAATTGTTTGGG | Del_k33_12-44;Del_k33_13-45 | -3.7413911479256035 | 0.08268794627569548 | 1.164705059174752 |
| GATCCAGATCTATTCTTCTTTTGCCAATTCCACTAATTGTTTGGG | Del_k18_13-30 | -0.6785469380921412 | 0.033634981515577714 | 24.91106535956822 |
| GATCCAGATCTATTCTTTTGCCAATTCCACTAATTGTTTGGG | Del_k21_13-33 | -2.030336080311833 | 0.05359233055171696 | 6.446407213742855 |
| GATCCAGATCTATTGACTTGGGGTGGCTTTGTCTTCTTCTTTTGCCAATTCCACTAATTGTTTGGG | Ins_k3_12_TTG | 0.49000426622464044 | 0.03332336178220125 | 80.14706832447749 |
| GATCCAGATCTATTGCCAATTCCACTAATTGTTTGGG | Del_k26_13-38 | -1.6753580069831955 | 0.041845911650430445 | 9.193539671927548 |
| GATCCAGATCTATTGGG | Del_k46_13-58 | -5.96031725248447 | 0.25323669620809575 | 0.12663349666165138 |
| GATCCAGATCTATTGGGGTGGCTTTGTCTTCTTCTTTTGCCAATTCCACTAATTGTTTGGG | Del_k2_13-14 | 0.1595384664530628 | 0.029176011136653298 | 57.592796623037245 |
| GATCCAGATCTATTGTCTTCTTCTTTTGCCAATTCCACTAATTGTTTGGG | Del_k13_13-25 | -0.38187096771521756 | 0.031773717207109455 | 33.51483132865734 |
| GATCCAGATCTATTGTTTGGG | Del_k42_12-53;Del_k42_13-54 | -5.753141768668795 | 0.2298187039129256 | 0.15578432912746962 |
| GATCCAGATCTATTTACTTGGGGTGGCTTTGTCTTCTTCTTTTGCCAATTCCACTAATTGTTTGGG | Ins_k3_12_TTT | -0.38005833678172296 | 0.04318413309218167 | 33.57563644061935 |

| | | | | |
|---|---|---|---|---|
| **GATCCAGATCTATTTGCCAATTCCACTAATTGTTTGGG** | Del_k25_13-37 | -1.6117657830922292 | 0.03944093886528077 | 9.797166896703787 |
| **GATCCAGATCTATTTGGG** | Del_k45_13-57 | -6.620272203400381 | 0.5137538877892653 | 0.06545364030105502 |
| **GATCCAGATCTATTTGTCTTCTTCTTTTGCCAATTCCACTAATTGTTTGGG** | Del_k12_13-24 | -0.45522235943495115 | 0.03521037828328543 | 31.14446918968088 |
| **GATCCAGATCTATTTTGCCAATTCCACTAATTGTTTGGG** | Del_k24_13-36 | -1.8452274038844152 | 0.04636982332548843 | 7.757279019710678 |
| **GATCCAGATCTCAAAACTTGGGGTGGCTTTGTCTTCTTCTTTTGCCAATTCCACTAATTGTTTGGG** | Ins_k3_11_CAA | 0.4371023247213188 | 0.03132939721254983 | 76.01733143777066 |
| **GATCCAGATCTCAAACTTGGGGTGGCTTTGTCTTCTTCTTTTGCCAATTCCACTAATTGTTTGGG** | Ins_k2_11_CA | 0.44809369282943345 | 0.035256836111981965 | 76.85747461351734 |
| **GATCCAGATCTCAACTTGGGGTGGCTTTGTCTTCTTCTTTTGCCAATTCCACTAATTGTTTGGG** | Ins_k1_11_C | 0.8934507328935124 | 0.3020949654272741 | 100 |
| **GATCCAGATCTCAATTCCACTAATTGTTTGGG** | Del_k31_12-42 | -2.5449472754338425 | 0.04742654646217819 | 3.853230006778136 |
| **GATCCAGATCTCACAACTTGGGGTGGCTTTGTCTTCTTCTTTTGCCAATTCCACTAATTGTTTGGG** | Ins_k3_11_CAC | 0.4638135310306357 | 0.03023207354185126 | 78.07520788760995 |
| **GATCCAGATCTCACTAATTGTTTGGG** | Del_k37_12-48 | -4.588354779156873 | 0.12924383070126655 | 0.49932617255354317 |
| **GATCCAGATCTCACTTGGGGTGGCTTTGTCTTCTTCTTTTGCCAATTCCACTAATTGTTTGGG** | A12C | 0.2114329090470407 | 0.03314905938680069 | 60.660451408351896 |
| **GATCCAGATCTCAGAACTTGGGGTGGCTTTGTCTTCTTCTTTTGCCAATTCCACTAATTGTTTGGG** | Ins_k3_11_CAG | 0.62895834347102 | 0.03016319196927535 | 92.09469841703677 |
| **GATCCAGATCTCATAACTTGGGGTGGCTTTGTCTTCTTCTTTTGCCAATTCCACTAATTGTTTGGG** | Ins_k3_10_TCA;Ins_k3_11_CAT | 0.03446966583984812 | 0.03859279348688345 | 50.82196792817041 |
| **GATCCAGATCTCCAAACTTGGGGTGGCTTTGTCTTCTTCTTTTGCCAATTCCACTAATTGTTTGGG** | Ins_k3_11_CCA | 0.5977598516219753 | 0.028476766424416995 | 89.26584022482835 |
| **GATCCAGATCTCCAACTTGGGGTGGCTTTGTCTTCTTCTTTTGCCAATTCCACTAATTGTTTGGG** | Ins_k2_11_CC | 0.7983660843992679 | 0.031638670928915216 | 100 |
| **GATCCAGATCTCCAATTCCACTAATTGTTTGGG** | Del_k30_12-41 | -1.5147701945737557 | 0.03539453809165986 | 10.795062354906232 |
| **GATCCAGATCTCCACTAATTGTTTGGG** | Del_k36_11-46;Del_k36_12-47 | -3.1308779957373285 | 0.05712378206803037 | 2.1446600158821085 |
| **GATCCAGATCTCCCAACTTGGGGTGGCTTTGTCTTCTTCTTTTGCCAATTCCACTAATTGTTTGGG** | Ins_k3_11_CCC | 0.46589561821875214 | 0.02980721277361143 | 78.2379366265866 |
| **GATCCAGATCTCCGAACTTGGGGTGGCTTTGTCTTCTTCTTTTGCCAATTCCACTAATTGTTTGGG** | Ins_k3_11_CCG | 0.8984002639664861 | 0.029604714552042232 | 100 |
| **GATCCAGATCTCCTAACTTGGGGTGGCTTTGTCTTCTTCTTTTGCCAATTCCACTAATTGTTTGGG** | Ins_k3_9_CTC;Ins_k3_10_TCC;In | 0.3612108583580891 | 0.034604470941072746 | 70.46174181002489 |
| **GATCCAGATCTCGAAACTTGGGGTGGCTTTGTCTTCTTCTTTTGCCAATTCCACTAATTGTTTGGG** | Ins_k3_11_CGA | 0.7401224783607612 | 0.029475798094437344 | 100 |
| **GATCCAGATCTCGAACTTGGGGTGGCTTTGTCTTCTTCTTTTGCCAATTCCACTAATTGTTTGGG** | Ins_k2_11_CG | 0.635706454056477 | 0.03303767257430567 | 92.71826520884541 |
| **GATCCAGATCTCGCAACTTGGGGTGGCTTTGTCTTCTTCTTTTGCCAATTCCACTAATTGTTTGGG** | Ins_k3_11_CGC | 0.6586904867681722 | 0.028821520110669078 | 94.87398350073491 |
| **GATCCAGATCTCGGAACTTGGGGTGGCTTTGTCTTCTTCTTTTGCCAATTCCACTAATTGTTTGGG** | Ins_k3_11_CGG | 0.7786219266370737 | 0.03455563693825025 | 100 |
| **GATCCAGATCTCGTAACTTGGGGTGGCTTTGTCTTCTTCTTTTGCCAATTCCACTAATTGTTTGGG** | Ins_k3_10_TCG;Ins_k3_11_CGT | 0.37397088912224197 | 0.03279609640451012 | 71.3665965129245 |
| **GATCCAGATCTCTAAACTTGGGGTGGCTTTGTCTTCTTCTTTTGCCAATTCCACTAATTGTTTGGG** | Ins_k3_11_CTA | -0.38718239825867573 | 0.039437770188883677 | 33.3372915423866 |
| **GATCCAGATCTCTAACTTGGGGTGGCTTTGTCTTCTTCTTTTGCCAATTCCACTAATTGTTTGGG** | Ins_k2_8_TC;Ins_k2_9_CT;Ins_k2 | -0.27330992774873863 | 0.04713906581092906 | 37.35807577385461 |
| **GATCCAGATCTCTAATTGTTTGGG** | Del_k39_12-50 | -5.703673748291879 | 0.21502008141616072 | 0.16368446251571545 |
| **GATCCAGATCTCTCAACTTGGGGTGGCTTTGTCTTCTTCTTTTGCCAATTCCACTAATTGTTTGGG** | Ins_k3_11_CTC | 0.5652715034039937 | 0.03186628654968536 | 86.4123441926207 |
| **GATCCAGATCTCTGAACTTGGGGTGGCTTTGTCTTCTTCTTTTGCCAATTCCACTAATTGTTTGGG** | Ins_k3_11_CTG | 0.5431647565389832 | 0.03380583462419166 | 84.52300884651726 |
| **GATCCAGATCTCTTAACTTGGGGTGGCTTTGTCTTCTTCTTTTGCCAATTCCACTAATTGTTTGGG** | Ins_k3_10_TCT;Ins_k3_11_CTT | -0.22698542808240024 | 0.03918783045031218 | 39.129380596401276 |
| **GATCCAGATCTCTTCTTCTTTTGCCAATTCCACTAATTGTTTGGG** | Del_k18_11-28;Del_k18_12-29 | -0.20354309340225069 | 0.030907651853567267 | 40.05750078105421 |
| **GATCCAGATCTCTTCTTTTGCCAATTCCACTAATTGTTTGGG** | Del_k21_11-31;Del_k21_12-32 | -0.30592859445357573 | 0.031597942489964274 | 36.15916488804048 |
| **GATCCAGATCTCTTGAACTTGGGGTGGCTTTGTCTTCTTCTTTTGCCAATTCCACTAATTGTTTGGG** | Ins_k4_11_CTTG | 0.698036923599946 | 0.03516911801689849 | 98.68134864304642 |
| **GATCCAGATCTCTTGGGGTGGCTTTGTCTTCTTCTTTTGCCAATTCCACTAATTGTTTGGG** | Del_k2_12-13 | 0.5180983116586728 | 0.0329903783715622 | 82.4306510457576 |
| **GATCCAGATCTCTTTGTCTTCTTCTTTTGCCAATTCCACTAATTGTTTGGG** | Del_k12_12-23 | -0.47611829621970403 | 0.032579297350425085 | 30.500428679518386 |
| **GATCCAGATCTCTTTTGCCAATTCCACTAATTGTTTGGG** | Del_k24_11-34;Del_k24_12-35 | -1.4084195897757386 | 0.04148097589322152 | 12.006395246367948 |
| **GATCCAGATCTG** | Del_k51_12-62 | -6.6632744430282695 | 0.27934986708419557 | 0.06269864714393507 |
| **GATCCAGATCTGAAAACTTGGGGTGGCTTTGTCTTCTTCTTTTGCCAATTCCACTAATTGTTTGGG** | Ins_k3_11_GAA | 0.6131847053037489 | 0.028030169771875762 | 90.65342689796286 |
| **GATCCAGATCTGAAACTTGGGGTGGCTTTGTCTTCTTCTTTTGCCAATTCCACTAATTGTTTGGG** | Ins_k2_11_GA | 0.6930212708645593 | 0.031006828201016953 | 98.18763644627506 |
| **GATCCAGATCTGAACTTGGGGTGGCTTTGTCTTCTTCTTTTGCCAATTCCACTAATTGTTTGGG** | Ins_k1_11_G | 0.570109894038485 | 0.030994671280779502 | 86.83145396024013 |
| **GATCCAGATCTGACAACTTGGGGTGGCTTTGTCTTCTTCTTTTGCCAATTCCACTAATTGTTTGGG** | Ins_k3_11_GAC | 0.5092601121232241 | 0.028589728100810192 | 81.7053225245837 |
| **GATCCAGATCTGACTTGGGGTGGCTTTGTCTTCTTCTTTTGCCAATTCCACTAATTGTTTGGG** | A12G | 0.40516282795334324 | 0.02725515079199701 | 73.62774043107989 |
| **GATCCAGATCTGAGAACTTGGGGTGGCTTTGTCTTCTTCTTTTGCCAATTCCACTAATTGTTTGGG** | Ins_k3_11_GAG | 0.6182448913206053 | 0.029677458105810753 | 91.11331267361837 |
| **GATCCAGATCTGATAACTTGGGGTGGCTTTGTCTTCTTCTTTTGCCAATTCCACTAATTGTTTGGG** | Ins_k3_10_TGA;Ins_k3_11_GAT | 0.20187702528528895 | 0.03447018865512283 | 60.08354798676141 |
| **GATCCAGATCTGCAAACTTGGGGTGGCTTTGTCTTCTTCTTTTGCCAATTCCACTAATTGTTTGGG** | Ins_k3_11_GCA | 0.7253319690514921 | 0.02690277347101435 | 100 |
| **GATCCAGATCTGCAACTTGGGGTGGCTTTGTCTTCTTCTTTTGCCAATTCCACTAATTGTTTGGG** | Ins_k2_11_GC | 0.9080258550952494 | 0.029171330178896247 | 100 |
| **GATCCAGATCTGCCAACTTGGGGTGGCTTTGTCTTCTTCTTTTGCCAATTCCACTAATTGTTTGGG** | Ins_k3_11_GCC | 0.5883818762539985 | 0.030551225170181627 | 88.432620437866575 |
| **GATCCAGATCTGCCAATTCCACTAATTGTTTGGG** | Del_k29_11-39;Del_k29_12-40 | -1.2252047158573682 | 0.032743518164322215 | 14.420550668298642 |
| **GATCCAGATCTGCGAACTTGGGGTGGCTTTGTCTTCTTCTTTTGCCAATTCCACTAATTGTTTGGG** | Ins_k3_11_GCG | 0.8039228484077876 | 0.02844405415426105 | 100 |
| **GATCCAGATCTGCTAACTTGGGGTGGCTTTGTCTTCTTCTTTTGCCAATTCCACTAATTGTTTGGG** | Ins_k3_9_CTG;Ins_k3_10_TGC;In | 0.44604979038807024 | 0.03328206896178232 | 76.70054586167856 |
| **GATCCAGATCTGCTTTGTCTTCTTCTTTTGCCAATTCCACTAATTGTTTGGG** | Del_k11_12-22 | 0.5197504166762066 | 0.02606588804726173 | 82.56694769513678 |
| **GATCCAGATCTGG** | Del_k50_12-61 | -6.215870112392481 | 0.32477569513983623 | 0.09807614839578298 |
| **GATCCAGATCTGGAAACTTGGGGTGGCTTTGTCTTCTTCTTTTGCCAATTCCACTAATTGTTTGGG** | Ins_k3_11_GGA | 0.7883390779512598 | 0.02981034712316718 | 100 |
| **GATCCAGATCTGGAACTTGGGGTGGCTTTGTCTTCTTCTTTTGCCAATTCCACTAATTGTTTGGG** | Ins_k2_11_GG | 0.7431508716731403 | 0.032969399332294626 | 100 |
| **GATCCAGATCTGGCAACTTGGGGTGGCTTTGTCTTCTTCTTTTGCCAATTCCACTAATTGTTTGGG** | Ins_k3_11_GGC | 0.5127236336324135 | 0.029840524180646765 | 81.98880130059814 |
| **GATCCAGATCTGGCTTTGTCTTCTTCTTTTGCCAATTCCACTAATTGTTTGGG** | Del_k10_11-20;Del_k10_12-21 | 0.37382717246662023 | 0.027727883253746304 | 71.35634068133513 |
| **GATCCAGATCTGGG** | Del_k49_11-59;Del_k49_12-60 | -6.305511014594702 | 0.2934099832509592 | 0.08966704400758546 |
| **GATCCAGATCTGGGAACTTGGGGTGGCTTTGTCTTCTTCTTTTGCCAATTCCACTAATTGTTTGGG** | Ins_k3_11_GGG | 0.14324686076763216 | 0.034587383384651164 | 56.66211918968047 |
| **GATCCAGATCTGGGGTGGCTTTGTCTTCTTCTTTTGCCAATTCCACTAATTGTTTGGG** | Del_k5_11-15;Del_k5_12-16 | 0.3896421114127778 | 0.030221740922215783 | 72.49380763003163 |
| **GATCCAGATCTGGGTGGCTTTGTCTTCTTCTTTTGCCAATTCCACTAATTGTTTGGG** | Del_k6_12-17 | 0.4262145864375839 | 0.029127680842198446 | 75.19416397569533 |
| **GATCCAGATCTGGTAACTTGGGGTGGCTTTGTCTTCTTCTTTTGCCAATTCCACTAATTGTTTGGG** | Ins_k3_10_TGG;Ins_k3_11_GGT | -0.40841329716612906 | 0.041800186397758676 | 32.63697139457272 |
| **GATCCAGATCTGGTGGCTTTGTCTTCTTCTTTTGCCAATTCCACTAATTGTTTGGG** | Del_k7_12-18 | 0.512474016028875 | 0.02624196995661978 | 81.96833800660595 |
| **GATCCAGATCTGGTTAACTTGGGGTGGCTTTGTCTTCTTCTTTTGCCAATTCCACTAATTGTTTGGG** | Ins_k4_10_TGGT;Ins_k4_11_GGT | 0.13406784540939742 | 0.03734955093797397 | 56.144396454901624 |
| **GATCCAGATCTGTAAACTTGGGGTGGCTTTGTCTTCTTCTTTTGCCAATTCCACTAATTGTTTGGG** | Ins_k3_11_GTA | -0.42046000209730017 | 0.04007702893668327 | 32.246162138507444 |
| **GATCCAGATCTGTAACTTGGGGTGGCTTTGTCTTCTTCTTTTGCCAATTCCACTAATTGTTTGGG** | Ins_k2_10_TG;Ins_k2_11_GT | -0.1707365618775707 | 0.04019032823933593 | 41.39344243199274 |
| **GATCCAGATCTGTCAACTTGGGGTGGCTTTGTCTTCTTCTTTTGCCAATTCCACTAATTGTTTGGG** | Ins_k3_11_GTC | 0.5097980572684635 | 0.03108540988200297 | 81.74928733044274 |
| **GATCCAGATCTGTCTTCTTCTTTTGCCAATTCCACTAATTGTTTGGG** | Del_k16_11-26;Del_k16_12-27 | -0.08547565316350503 | 0.030013897322611824 | 45.077506805286845 |
| **GATCCAGATCTGTGAACTTGGGGTGGCTTTGTCTTCTTCTTTTGCCAATTCCACTAATTGTTTGGG** | Ins_k3_11_GTG | 0.2916860330338481 | 0.031546213754755845 | 65.72931809151966 |
| **GATCCAGATCTGTGGCTTTGTCTTCTTCTTTTGCCAATTCCACTAATTGTTTGGG** | Del_k8_12-19 | 0.39259539516373587 | 0.027553538395713294 | 72.70821886692228 |
| **GATCCAGATCTGTTAACTTGGGGTGGCTTTGTCTTCTTCTTTTGCCAATTCCACTAATTGTTTGGG** | Ins_k3_10_TGT;Ins_k3_11_GTT | -0.006935649834436064 | 0.03874137743550704 | 48.76063780218205 |
| **GATCCAGATCTGTTTGGG** | Del_k45_11-55;Del_k45_12-56 | -6.156176454066048 | 0.2556542179943079 | 0.10410894091829204 |
| **GATCCAGATCTTAAAACTTGGGGTGGCTTTGTCTTCTTCTTTTGCCAATTCCACTAATTGTTTGGG** | Ins_k3_11_TAA | -0.21748507403340855 | 0.040308943036442516 | 39.50289501605453 |
| **GATCCAGATCTTAAACTTGGGGTGGCTTTGTCTTCTTCTTTTGCCAATTCCACTAATTGTTTGGG** | Ins_k2_11_TA | 0.023620067819247192 | 0.039134911006007723 | 50.273550439862305 |
| **GATCCAGATCTTAACTTGGGGTGGCTTTGTCTTCTTCTTTTGCCAATTCCACTAATTGTTTGGG** | Ins_k1_10_T;Ins_k1_11_T | -0.14191586186968153 | 0.17527646332371108 | 42.60378814574077 |

| | | | | |
|---|---|---|---|---|
| **GATCCAGATCTTAATTGTTTGGG** | Del_k40_12-51 | -5.907680117307705 | 0.28397344845880035 | 0.13347766980548428 |
| **GATCCAGATCTTACAACTTGGGGTGGCTTTGTCTTCTTCTTTTGCCAATTCCACTAATTGTTTGGG** | Ins_k3_11_TAC | 0.39109466334174237 | 0.03349579231836751 | 72.59918516478523 |
| **GATCCAGATCTTACTTGGGGTGGCTTTGTCTTCTTCTTTTGCCAATTCCACTAATTGTTTGGG** | A12T | 0.046229548740301274 | 0.036784694754080215 | 51.42315634403984 |
| **GATCCAGATCTTAGAACTTGGGGTGGCTTTGTCTTCTTCTTTTGCCAATTCCACTAATTGTTTGGG** | Ins_k3_11_TAG | 0.07046595251702847 | 0.035463675427146645 | 52.68469454727034 |
| **GATCCAGATCTTATAACTTGGGGTGGCTTTGTCTTCTTCTTTTGCCAATTCCACTAATTGTTTGGG** | Ins_k3_10_TTA;Ins_k3_11_TAT | -0.5427470885942729 | 0.04742024047767786 | 28.53444477757895 |
| **GATCCAGATCTTCAAACTTGGGGTGGCTTTGTCTTCTTCTTTTGCCAATTCCACTAATTGTTTGGG** | Ins_k3_11_TCA | 0.7037701986450158 | 0.02929895671481104 | 99.24874091039693 |
| **GATCCAGATCTTCAACTTGGGGTGGCTTTGTCTTCTTCTTTTGCCAATTCCACTAATTGTTTGGG** | Ins_k2_11_TC | 0.8941458184238731 | 0.034999513368880145 | 100 |
| **GATCCAGATCTTCCAACTTGGGGTGGCTTTGTCTTCTTCTTTTGCCAATTCCACTAATTGTTTGGG** | Ins_k3_11_TCC | 0.6884525128768559 | 0.029390104575995214 | 97.7400641013654 |
| **GATCCAGATCTTCCACTAATTGTTTGGG** | Del_k35_11-45;Del_k35_12-46 | -2.863941435846044 | 0.057018125700197944 | 2.8008350334548537 |
| **GATCCAGATCTTCGAAACTTGGGGTGGCTTTGTCTTCTTCTTTTGCCAATTCCACTAATTGTTTGGG** | Ins_k4_11_TCGA | 1.0791402842447515 | 0.029341286128606 | 100 |
| **GATCCAGATCTTCGAACTTGGGGTGGCTTTGTCTTCTTCTTTTGCCAATTCCACTAATTGTTTGGG** | Ins_k3_11_TCG | 0.8273143434190704 | 0.027979693268294553 | 100 |
| **GATCCAGATCTTCTAACTTGGGGTGGCTTTGTCTTCTTCTTTTGCCAATTCCACTAATTGTTTGGG** | Ins_k3_8_TCT;Ins_k3_9_CTT;Ins_ | 0.3478536663858629 | 0.032508581473490276 | 69.52682860568218 |
| **GATCCAGATCTTCTTCTTCTTTTGCCAATTCCACTAATTGTTTGGG** | Del_k17_12-28 | 0.32629641193002834 | 0.029306181547816317 | 68.04406069412967 |
| **GATCCAGATCTTCTTCTTTTGCCAATTCCACTAATTGTTTGGG** | Del_k20_9-28;Del_k20_10-29;De | 0.06031645047616874 | 0.02835781683930714 | 52.152675563360056 |
| **GATCCAGATCTTCTTTTGCCAATTCCACTAATTGTTTGGG** | Del_k23_9-31;Del_k23_10-32;De | -0.24429645274461453 | 0.03162125934162178 | 38.45784021993556 |
| **GATCCAGATCTTGAAACTTGGGGTGGCTTTGTCTTCTTCTTTTGCCAATTCCACTAATTGTTTGGG** | Ins_k3_11_TGA | 0.6077263725050823 | 0.0316019299370434 | 90.15995830768476 |
| **GATCCAGATCTTGAACTTGGGGTGGCTTTGTCTTCTTCTTTTGCCAATTCCACTAATTGTTTGGG** | Ins_k2_11_TG | 0.641944681738821 | 0.03177412628374986 | 93.29847070293722 |
| **GATCCAGATCTTGCAACTTGGGGTGGCTTTGTCTTCTTCTTTTGCCAATTCCACTAATTGTTTGGG** | Ins_k3_11_TGC | 0.5847232247112024 | 0.03243112226984128 | 88.10966744115466 |
| **GATCCAGATCTTGCCAATTCCACTAATTGTTTGGG** | Del_k28_11-38;Del_k28_12-39 | -1.8006290960282316 | 0.03742262197920078 | 8.111071163663397 |
| **GATCCAGATCTTGGAACTTGGGGTGGCTTTGTCTTCTTCTTTTGCCAATTCCACTAATTGTTTGGG** | Ins_k3_11_TGG | 0.7049207114733103 | 0.03190261317567229 | 99.36299357198003 |
| **GATCCAGATCTTGGCTTTGTCTTCTTCTTTTGCCAATTCCACTAATTGTTTGGG** | Del_k9_12-20 | 0.18346138904777032 | 0.027333677201127986 | 58.98719720589083 |
| **GATCCAGATCTTGGG** | Del_k48_11-58;Del_k48_12-59 | -6.53991999093924 | 0.3475695626214738 | 0.07093006012237715 |
| **GATCCAGATCTTGGGGTGGCTTTGTCTTCTTCTTTTGCCAATTCCACTAATTGTTTGGG** | Del_k4_10-13;Del_k4_11-14;Del_ | 0.41227859961242364 | 0.0312976242220706 | 74.15352709010101 |
| **GATCCAGATCTTGTAACTTGGGGTGGCTTTGTCTTCTTCTTTTGCCAATTCCACTAATTGTTTGGG** | Ins_k3_10_TTG;Ins_k3_11_TGT | -0.3268878261127749 | 0.04663738938335642 | 35.40918354559997 |
| **GATCCAGATCTTGTCTTCTTCTTTTGCCAATTCCACTAATTGTTTGGG** | Del_k15_11-25;Del_k15_12-26 | -0.14890658047546745 | 0.03046316913464172 | 42.30699565635225 |
| **GATCCAGATCTTGTTTGGG** | Del_k44_11-54;Del_k44_12-55 | -5.7394562079124185 | 0.24005498460975122 | 0.15793098058855543 |
| **GATCCAGATCTTTAAACTTGGGGTGGCTTTGTCTTCTTCTTTTGCCAATTCCACTAATTGTTTGGG** | Ins_k3_11_TTA | -0.9074379642231909 | 0.05321563104714101 | 19.814640240835676 |
| **GATCCAGATCTTTAACTTGGGGTGGCTTTGTCTTCTTCTTTTGCCAATTCCACTAATTGTTTGGG** | Ins_k2_10_TT;Ins_k2_11_TT | -0.6298339360679928 | 0.053035585235319226 | 26.154600404606175 |
| **GATCCAGATCTTTCAACTTGGGGTGGCTTTGTCTTCTTCTTTTGCCAATTCCACTAATTGTTTGGG** | Ins_k3_11_TTC | 0.47005104747149606 | 0.03392021633503597 | 78.56372526413776 |
| **GATCCAGATCTTTCCAACTTGGGGTGGCTTTGTCTTCTTCTTTTGCCAATTCCACTAATTGTTTGGG** | Ins_k4_11_TTCC | 0.48149376664649846 | 0.03457584120637675 | 79.46787098707733 |
| **GATCCAGATCTTTCCACTAATTGTTTGGG** | Del_k34_12-45 | -3.703111888187283 | 0.0893923277329682 | 1.2101534220012484 |
| **GATCCAGATCTTTCTTCTTTTGCCAATTCCACTAATTGTTTGGG** | Del_k19_12-30 | -0.33043059300998645 | 0.0344604006621854 | 35.28395901391657 |
| **GATCCAGATCTTTCTTTTGCCAATTCCACTAATTGTTTGGG** | Del_k22_12-33 | -1.8571580908792567 | 0.05126472416243224 | 7.665279253281175 |
| **GATCCAGATCTTTGAACTTGGGGTGGCTTTGTCTTCTTCTTTTGCCAATTCCACTAATTGTTTGGG** | Ins_k3_11_TTG | 0.24437381970853833 | 0.03563333882322493 | 62.691937729145096 |
| **GATCCAGATCTTTGCCAATTCCACTAATTGTTTGGG** | Del_k27_11-37;Del_k27_12-38 | -1.7998680030307983 | 0.04181419163734724 | 8.117246792943734 |
| **GATCCAGATCTTTGGG** | Del_k47_11-57;Del_k47_12-58 | -6.42254761206505 | 0.3253137788823296 | 0.07976355546610672 |
| **GATCCAGATCTTTGGGGTGGCTTTGTCTTCTTCTTTTGCCAATTCCACTAATTGTTTGGG** | Del_k3_12-14 | 0.25596723509872643 | 0.03463213288893495 | 63.42298086008923 |
| **GATCCAGATCTTTGTCTTCTTCTTTTGCCAATTCCACTAATTGTTTGGG** | Del_k14_10-23;Del_k14_11-24;D | -0.17424274205820783 | 0.029439197370079655 | 41.248563698421236 |
| **GATCCAGATCTTTGTTTGGG** | Del_k43_12-54 | -5.5059996663210535 | 0.2479323683762435 | 0.1994601859941106 |
| **GATCCAGATCTTTTAACTTGGGGTGGCTTTGTCTTCTTCTTTTGCCAATTCCACTAATTGTTTGGG** | Ins_k3_10_TTT;Ins_k3_11_TTT | -0.6137029474892032 | 0.050743881018911495 | 26.579921164532575 |
| **GATCCAGATCTTTTCAACTTGGGGTGGCTTTGTCTTCTTCTTTTGCCAATTCCACTAATTGTTTGGG** | Ins_k4_11_TTTC | 0.6469487466006514 | 0.03387543317725921 | 93.7665123806284 |
| **GATCCAGATCTTTTGCCAATTCCACTAATTGTTTGGG** | Del_k26_9-34;Del_k26_10-35;De | -1.446810179354301 | 0.03714220191616006 | 11.554198246239114 |
| **GATCCAGATCTTTTGGG** | Del_k46_12-57 | -5.762448920460485 | 0.2875939372398608 | 0.15434114710431354 |
| **GATCCAGATCTTTTGTCTTCTTCTTTTGCCAATTCCACTAATTGTTTGGG** | Del_k13_12-24 | -0.199252353335172 | 0.03544547866262872 | 40.229746370909574 |
| **GATCCAGATCTTTTTGCCAATTCCACTAATTGTTTGGG** | Del_k25_12-36 | -1.535709992904404 | 0.043160916705447284 | 10.571366176368974 |
| **GATCCAGATG** | Del_k53_10-62 | -7.002941886615758 | 0.4577912210795742 | 0.044641879789525024 |
| **GATCCAGATGAACTAACTTGGGGTGGCTTTGTCTTCTTCTTTTGCCAATTCCACTAATTGTTTGGG** | Ins_k3_9_GAA | 0.38619817486591246 | 0.031352001453595714 | 72.24457297701964 |
| **GATCCAGATGACCTAACTTGGGGTGGCTTTGTCTTCTTCTTTTGCCAATTCCACTAATTGTTTGGG** | Ins_k3_9_GAC | 0.5630777368786408 | 0.0308321008993166 | 86.22298346710737 |
| **GATCCAGATGACTAACTTGGGGTGGCTTTGTCTTCTTCTTTTGCCAATTCCACTAATTGTTTGGG** | Ins_k2_9_GA | 0.3549857559863573 | 0.03423240151319526 | 70.02447268857264 |
| **GATCCAGATGAGCCTAACTTGGGGTGGCTTTGTCTTCTTCTTTTGCCAATTCCACTAATTGTTTGGG** | Ins_k4_9_GAGC | 0.7691683868238721 | 0.03165081268145178 | 100 |
| **GATCCAGATGAGCTAACTTGGGGTGGCTTTGTCTTCTTCTTTTGCCAATTCCACTAATTGTTTGGG** | Ins_k3_9_GAG | 0.4158880479778444 | 0.02912227948227793 | 74.42166403937328 |
| **GATCCAGATGATCTAACTTGGGGTGGCTTTGTCTTCTTCTTTTGCCAATTCCACTAATTGTTTGGG** | Ins_k3_6_GAT;Ins_k3_7_ATG;Ins_ | 0.5308743757205208 | 0.033820701830676224 | 83.49054655363554 |
| **GATCCAGATGCACTAACTTGGGGTGGCTTTGTCTTCTTCTTTTGCCAATTCCACTAATTGTTTGGG** | Ins_k3_9_GCA | 0.3133111944004291 | 0.0342354242975025 | 67.16620566864158 |
| **GATCCAGATGCCAATTCCACTAATTGTTTGGG** | Del_k31_9-39;Del_k31_10-40 | -0.9913692321174058 | 0.029048185228489884 | 18.219451783957616 |
| **GATCCAGATGCCCCTAACTTGGGGTGGCTTTGTCTTCTTCTTTTGCCAATTCCACTAATTGTTTGGG** | Ins_k4_9_GCCC | 0.6342912138778531 | 0.03338717353094401 | 92.5871394037695 |
| **GATCCAGATGCCCTAACTTGGGGTGGCTTTGTCTTCTTCTTTTGCCAATTCCACTAATTGTTTGGG** | Ins_k3_9_GCC | 0.5855216896096271 | 0.03119645884177431 | 88.18004801228685 |
| **GATCCAGATGCCTAACTTGGGGTGGCTTTGTCTTCTTCTTTTGCCAATTCCACTAATTGTTTGGG** | Ins_k2_9_GC | 0.6786254470176969 | 0.03400276001638784 | 96.78427007163933 |
| **GATCCAGATGCGCTAACTTGGGGTGGCTTTGTCTTCTTCTTTTGCCAATTCCACTAATTGTTTGGG** | Ins_k3_9_GCG | 0.6472601069500004 | 0.029612250403242436 | 93.79571210026202 |
| **GATCCAGATGCTAACTTGGGGTGGCTTTGTCTTCTTCTTTTGCCAATTCCACTAATTGTTTGGG** | Ins_k1_9_G | 0.6036646719968528 | 0.031175754337011106 | 89.79449825627135 |
| **GATCCAGATGCTCTAACTTGGGGTGGCTTTGTCTTCTTCTTTTGCCAATTCCACTAATTGTTTGGG** | Ins_k3_8_TGC;Ins_k3_9_GCT | 0.5267294660057416 | 0.03408661731901902 | 83.14520198159508 |
| **GATCCAGATGCTTTGTCTTCTTCTTTTGCCAATTCCACTAATTGTTTGGG** | Del_k13_10-22 | 0.454863027039756 | 0.026373637374905345 | 77.3795134800308 |
| **GATCCAGATGG** | Del_k52_10-61 | -8.774775118536787 | 2.0686444336561456 | 0.007590057713352011 |
| **GATCCAGATGGACTAACTTGGGGTGGCTTTGTCTTCTTCTTTTGCCAATTCCACTAATTGTTTGGG** | Ins_k3_9_GGA | 0.5473863901127229 | 0.030688067101359774 | 84.88058827200061 |
| **GATCCAGATGGCCTAACTTGGGGTGGCTTTGTCTTCTTCTTTTGCCAATTCCACTAATTGTTTGGG** | Ins_k3_9_GGC | 0.6739072054731743 | 0.03225477334573197 | 96.32869411160324 |
| **GATCCAGATGGCTAACTTGGGGTGGCTTTGTCTTCTTCTTTTGCCAATTCCACTAATTGTTTGGG** | Ins_k2_9_GG | 0.5356019352693331 | 0.03337152742034804 | 83.88618755555386 |
| **GATCCAGATGGCTTTGTCTTCTTCTTTTGCCAATTCCACTAATTGTTTGGG** | Del_k12_9-20;Del_k12_10-21 | 0.3886071862374807 | 0.02679283037892545 | 72.41882077304822 |
| **GATCCAGATGGG** | Del_k51_9-59;Del_k51_10-60 | -6.103653475810027 | 0.47607400042834985 | 0.10972320078790344 |
| **GATCCAGATGGGCTAACTTGGGGTGGCTTTGTCTTCTTCTTTTGCCAATTCCACTAATTGTTTGGG** | Ins_k3_9_GGG | -0.5105764939477155 | 0.04285352572015718 | 29.467340278749575 |
| **GATCCAGATGGGGTGGCTTTGTCTTCTTCTTTTGCCAATTCCACTAATTGTTTGGG** | Del_k7_9-15;Del_k7_10-16 | 0.5399423912406545 | 0.028691303966005572 | 84.25108319309264 |
| **GATCCAGATGGGTGGCTTTGTCTTCTTCTTTTGCCAATTCCACTAATTGTTTGGG** | Del_k8_10-17 | 0.6285497346467621 | 0.02748549669349218 | 92.05707539766853 |
| **GATCCAGATGGTCTAACTTGGGGTGGCTTTGTCTTCTTCTTTTGCCAATTCCACTAATTGTTTGGG** | Ins_k3_8_TGG;Ins_k3_9_GGT | 0.3228411663746217 | 0.0318360376664637 | 67.80935746798862 |
| **GATCCAGATGGTGGCTTTGTCTTCTTCTTTTGCCAATTCCACTAATTGTTTGGG** | Del_k9_10-18 | 0.5310633253971762 | 0.027106517723368046 | 83.50632355589342 |
| **GATCCAGATGTAACTTGGGGTGGCTTTGTCTTCTTCTTTTGCCAATTCCACTAATTGTTTGGG** | C10G | 0.09534154183015975 | 0.033584092571808145 | 54.01169388957712 |
| **GATCCAGATGTACTAACTTGGGGTGGCTTTGTCTTCTTCTTTTGCCAATTCCACTAATTGTTTGGG** | Ins_k3_9_GTA | 0.07228462164648208 | 0.034817500125781296 | 52.7805977565074 |
| **GATCCAGATGTCCTAACTTGGGGTGGCTTTGTCTTCTTCTTTTGCCAATTCCACTAATTGTTTGGG** | Ins_k3_9_GTC | 0.6763262491731692 | 0.02728889783025892 | 96.56199950642458 |

| | | | | |
|---|---|---|---|---|
| **GATCCAGATGTCTAACTTGGGGTGGCTTTGTCTTCTTCTTTTGCCAATTCCACTAATTGTTTGGG** | Ins_k2_8_TG;Ins_k2_9_GT | 0.567521875253796 | 0.0332854018698956 | 86.6070230672395 |
| **GATCCAGATGTCTCTAACTTGGGGTGGCTTTGTCTTCTTCTTTTGCCAATTCCACTAATTGTTTGGG** | Ins_k4_8_TGTC;Ins_k4_9_GTCT | 0.8179337476973936 | 0.034050802780671865 | 100 |
| **GATCCAGATGTCTTCTTCTTTTGCCAATTCCACTAATTGTTTGGG** | Del_k18_9-26;Del_k18_10-27 | 0.3318409274891313 | 0.02507157458716286 | 68.42237987646998 |
| **GATCCAGATGTGCTAACTTGGGGTGGCTTTGTCTTCTTCTTTTGCCAATTCCACTAATTGTTTGGG** | Ins_k3_9_GTG | 0.5199950589270581 | 0.03040381960616226 | 82.58714953007728 |
| **GATCCAGATGTGGCTTTGTCTTCTTCTTTTGCCAATTCCACTAATTGTTTGGG** | Del_k10_10-19 | 0.5511852865335702 | 0.028135679534771193 | 85.20365409325254 |
| **GATCCAGATGTGTCTAACTTGGGGTGGCTTTGTCTTCTTCTTTTGCCAATTCCACTAATTGTTTGGG** | Ins_k4_8_TGTG;Ins_k4_9_GTGT | 0.7183725450628948 | 0.031229840558658836 | 100 |
| **GATCCAGATGTTCTAACTTGGGGTGGCTTTGTCTTCTTCTTTTGCCAATTCCACTAATTGTTTGGG** | Ins_k3_8_TGT;Ins_k3_9_GTT | 0.4679603458979926 | 0.03578361271214112 | 78.3996435428314 |
| **GATCCAGATGTTTGGG** | Del_k47_9-55;Del_k47_10-56 | -6.275490366138073 | 0.38428450476397874 | 0.09239971993573802 |
| **GATCCAGATTAAACTAACTTGGGGTGGCTTTGTCTTCTTCTTTTGCCAATTCCACTAATTGTTTGGG** | Ins_k4_9_TAAA | -1.5804896949407659 | 0.08362187709884074 | 10.108426064552978 |
| **GATCCAGATTAACTAACTTGGGGTGGCTTTGTCTTCTTCTTTTGCCAATTCCACTAATTGTTTGGG** | Ins_k3_9_TAA | -1.6043181865426919 | 0.07534939595232211 | 9.870404627092066 |
| **GATCCAGATTAACTTGGGGTGGCTTTGTCTTCTTCTTTTGCCAATTCCACTAATTGTTTGGG** | Del_k1_10-10 | -0.5466504199185023 | 0.043527496816388025 | 28.42328247824066 |
| **GATCCAGATTAATTGTTTGGG** | Del_k42_10-51 | -6.064858645992994 | 0.25365108302331224 | 0.11406354071245854 |
| **GATCCAGATTACCTAACTTGGGGTGGCTTTGTCTTCTTCTTTTGCCAATTCCACTAATTGTTTGGG** | Ins_k3_9_TAC | 0.2907613453446417 | 0.03829818507585853 | 65.66856709243972 |
| **GATCCAGATTACTAACTTGGGGTGGCTTTGTCTTCTTCTTTTGCCAATTCCACTAATTGTTTGGG** | Ins_k2_9_TA | -1.0968292499639158 | 0.061500669990833416 | 16.395875098936234 |
| **GATCCAGATTAGCTAACTTGGGGTGGCTTTGTCTTCTTCTTTTGCCAATTCCACTAATTGTTTGGG** | Ins_k3_9_TAG | -2.53486987410216 | 0.10767135248383942 | 3.8922568663658046 |
| **GATCCAGATTATCTAACTTGGGGTGGCTTTGTCTTCTTCTTTTGCCAATTCCACTAATTGTTTGGG** | Ins_k3_7_ATT;Ins_k3_8_TTA;Ins_ | -0.3853107133874168 | 0.04512692400380277 | 33.39974687673463 |
| **GATCCAGATTCAACTAACTTGGGGTGGCTTTGTCTTCTTCTTTTGCCAATTCCACTAATTGTTTGGG** | Ins_k4_9_TCAA | 0.5371611317696332 | 0.03514309724297206 | 84.01708462622265 |
| **GATCCAGATTCACTAACTTGGGGTGGCTTTGTCTTCTTCTTTTGCCAATTCCACTAATTGTTTGGG** | Ins_k3_9_TCA | -0.19544938347402652 | 0.0359989416892003 | 40.38303016596619 |
| **GATCCAGATTCCACTAATTGTTTGGG** | Del_k37_8-44;Del_k37_9-45;Del_ | -4.32578868039543 | 0.09592663592867906 | 0.6492550530313409 |
| **GATCCAGATTCCCTAACTTGGGGTGGCTTTGTCTTCTTCTTTTGCCAATTCCACTAATTGTTTGGG** | Ins_k3_9_TCC | 0.3788484275820375 | 0.0339599534499427 | 71.71554013333387 |
| **GATCCAGATTCCTAACTTGGGGTGGCTTTGTCTTCTTCTTTTGCCAATTCCACTAATTGTTTGGG** | Ins_k2_9_TC | 0.1961651335283524 | 0.03814531296453422 | 59.74133553493944 |
| **GATCCAGATTCGCTAACTTGGGGTGGCTTTGTCTTCTTCTTTTGCCAATTCCACTAATTGTTTGGG** | Ins_k3_9_TCG | 0.4261772529216463 | 0.029998822903883282 | 75.19135676557796 |
| **GATCCAGATTCTAACTTGGGGTGGCTTTGTCTTCTTCTTTTGCCAATTCCACTAATTGTTTGGG** | Ins_k1_8_T;Ins_k1_9_T | 0.22434437278044728 | 0.04101447296489501 | 61.448744685729665 |
| **GATCCAGATTCTCTAACTTGGGGTGGCTTTGTCTTCTTCTTTTGCCAATTCCACTAATTGTTTGGG** | Ins_k3_8_TTC;Ins_k3_9_TCT | -0.19596111809686317 | 0.04701235038836237 | 40.36237005795236 |
| **GATCCAGATTCTTCTTCTTTTGCCAATTCCACTAATTGTTTGGG** | Del_k19_10-28 | 0.0729760081467381 | 0.03283420173727272 | 52.817102167145904 |
| **GATCCAGATTCTTCTTTTGCCAATTCCACTAATTGTTTGGG** | Del_k22_9-30;Del_k22_10-31 | -0.2917306007296158 | 0.02939320076556869 | 36.67621433088668 |
| **GATCCAGATTCTTTTGCCAATTCCACTAATTGTTTGGG** | Del_k25_9-33;Del_k25_10-34 | -1.5347306626106527 | 0.04193302608661013 | 10.581724106601412 |
| **GATCCAGATTGACTAACTTGGGGTGGCTTTGTCTTCTTCTTTTGCCAATTCCACTAATTGTTTGGG** | Ins_k3_9_TGA | -0.0951139469837664 | 0.0402216871434393 | 44.64512361536906 |
| **GATCCAGATTGCCAATTCCACTAATTGTTTGGG** | Del_k30_9-38;Del_k30_10-39 | -2.061460364162752 | 0.04063735020853084 | 6.248857647469098 |
| **GATCCAGATTGCCTAACTTGGGGTGGCTTTGTCTTCTTCTTTTGCCAATTCCACTAATTGTTTGGG** | Ins_k3_9_TGC | 0.4882875501424859 | 0.0347649960087127 | 80.00959659707203 |
| **GATCCAGATTGCTAACTTGGGGTGGCTTTGTCTTCTTCTTTTGCCAATTCCACTAATTGTTTGGG** | Ins_k2_9_TG | -0.014953705151468544 | 0.04637960903894161 | 48.371235521698004 |
| **GATCCAGATTGGCTAACTTGGGGTGGCTTTGTCTTCTTCTTTTGCCAATTCCACTAATTGTTTGGG** | Ins_k3_9_TGG | 0.46604499918201014 | 0.039151457384299725 | 78.24962475789363 |
| **GATCCAGATTGGCTTTGTCTTCTTCTTTTGCCAATTCCACTAATTGTTTGGG** | Del_k11_10-20 | 0.22925496311358406 | 0.03218851401455182 | 61.75123639594656 |
| **GATCCAGATTGGG** | Del_k50_9-58;Del_k50_10-59 | -6.330216987152246 | 0.3549742244437076 | 0.08747887421153494 |
| **GATCCAGATTGGGGTGGCTTTGTCTTCTTCTTTTGCCAATTCCACTAATTGTTTGGG** | Del_k6_9-14;Del_k6_10-15 | 0.0897577788472288 | 0.029398276049775615 | 53.71094582694418 |
| **GATCCAGATTGTCTAACTTGGGGTGGCTTTGTCTTCTTCTTTTGCCAATTCCACTAATTGTTTGGG** | Ins_k3_8_TTG;Ins_k3_9_TGT | -0.006745445100407421 | 0.04088006448956962 | 48.769913188409504 |
| **GATCCAGATTGTCTTCTTCTTTTGCCAATTCCACTAATTGTTTGGG** | Del_k17_9-25;Del_k17_10-26 | 0.478824938415654 | 0.1314597564553761 | 79.25606764862307 |
| **GATCCAGATTGTTTGGG** | Del_k46_8-53;Del_k46_9-54;Del_ | -6.98145609508849 | 1.1470153721761733 | 0.04561142432997446 |
| **GATCCAGATTTAACTTGGGGTGGCTTTGTCTTCTTCTTTTGCCAATTCCACTAATTGTTTGGG** | C10T | -0.4374524992114188 | 0.03750230769556513 | 31.70284852214918 |
| **GATCCAGATTTACTAACTTGGGGTGGCTTTGTCTTCTTCTTTTGCCAATTCCACTAATTGTTTGGG** | Ins_k3_9_TTA | -1.2118750457661678 | 0.07090129467345656 | 14.614058685050248 |
| **GATCCAGATTTCCACTAATTGTTTGGG** | Del_k36_10-45 | -3.949021152986702 | 0.09214672033448572 | 0.9463317179352408 |
| **GATCCAGATTTCCTAACTTGGGGTGGCTTTGTCTTCTTCTTTTGCCAATTCCACTAATTGTTTGGG** | Ins_k3_9_TTC | 0.23335254875512956 | 0.03728187149443851 | 62.00478649240446 |
| **GATCCAGATTTCTAACTTGGGGTGGCTTTGTCTTCTTCTTTTGCCAATTCCACTAATTGTTTGGG** | Ins_k2_8_TT;Ins_k2_9_TT | -0.5165433463347509 | 0.050225847344373166 | 29.29203653496484 |
| **GATCCAGATTTCTTCTTTTGCCAATTCCACTAATTGTTTGGG** | Del_k21_10-30 | -0.24985079157525175 | 0.03747636149034789 | 38.24482447311635 |
| **GATCCAGATTTCTTTTGCCAATTCCACTAATTGTTTGGG** | Del_k24_10-33 | -1.706790403606516 | 0.057067102317907015 | 8.909059062014398 |
| **GATCCAGATTTGCCAATTCCACTAATTGTTTGGG** | Del_k29_9-37;Del_k29_10-38 | -1.758259871439411 | 0.04014063140559753 | 8.462115177600541 |
| **GATCCAGATTTGCTAACTTGGGGTGGCTTTGTCTTCTTCTTTTGCCAATTCCACTAATTGTTTGGG** | Ins_k3_9_TTG | -0.11975214187895805 | 0.041129712717509335 | 43.558588454831124 |
| **GATCCAGATTTGGG** | Del_k49_9-57;Del_k49_10-58 | -5.9087338290675175 | 0.3030296371102147 | 0.13333709688981163 |
| **GATCCAGATTTGGGGTGGCTTTGTCTTCTTCTTTTGCCAATTCCACTAATTGTTTGGG** | Del_k5_10-14 | 0.12415814617390086 | 0.030845016029922064 | 55.59077003851346 |
| **GATCCAGATTTGTCTTCTTCTTTTGCCAATTCCACTAATTGTTTGGG** | Del_k16_9-24;Del_k16_10-25 | 0.139334426078485 | 0.0327247250638613 | 56.440865451632284 |
| **GATCCAGATTTGTTTGGG** | Del_k45_10-54 | -6.8266904701852615 | 0.843549312507664 | 0.05324606172280744 |
| **GATCCAGATTTTCTAACTTGGGGTGGCTTTGTCTTCTTCTTTTGCCAATTCCACTAATTGTTTGGG** | Ins_k3_8_TTT;Ins_k3_9_TTT | -0.6237035397022543 | 0.06046755815162739 | 26.315430945665202 |
| **GATCCAGATTTTGCCAATTCCACTAATTGTTTGGG** | Del_k28_9-36;Del_k28_10-37 | -1.9676973822548394 | 0.04471874797445993 | 6.863116593883525 |
| **GATCCAGATTTTGGG** | Del_k48_10-57 | -6.431661226415338 | 0.4888909910367627 | 0.07903992364167925 |
| **GATCCAGATTTTGTCTTCTTCTTTTGCCAATTCCACTAATTGTTTGGG** | Del_k15_10-24 | -0.13703108230387456 | 0.031243339141645853 | 42.81240737415001 |
| **GATCCAGATTTTTGCCAATTCCACTAATTGTTTGGG** | Del_k27_10-36 | -2.0269600588954018 | 0.048268881246834636 | 6.468207200461029 |
| **GATCCAGCAAATCTAACTTGGGGTGGCTTTGTCTTCTTCTTTTGCCAATTCCACTAATTGTTTGGG** | Ins_k3_7_CAA | 0.08211291683064836 | 0.03765172139881837 | 53.30189860429636 |
| **GATCCAGCAATCTAACTTGGGGTGGCTTTGTCTTCTTCTTTTGCCAATTCCACTAATTGTTTGGG** | Ins_k2_7_CA | 0.028664038564808703 | 0.04535998914239072 | 50.52776935496322 |
| **GATCCAGCAATTCCACTAATTGTTTGGG** | Del_k35_8-42 | -3.4506265754923566 | 0.07359412336114697 | 1.557734402118485 |
| **GATCCAGCACATCTAACTTGGGGTGGCTTTGTCTTCTTCTTTTGCCAATTCCACTAATTGTTTGGG** | Ins_k3_7_CAC | 0.3416029899841956 | 0.03273082749812708 | 69.09359431309066 |
| **GATCCAGCACTAATTGTTTGGG** | Del_k41_8-48 | -5.105898464878914 | 0.1700430870630997 | 0.2975897466497967 |
| **GATCCAGCAGATCTAACTTGGGGTGGCTTTGTCTTCTTCTTTTGCCAATTCCACTAATTGTTTGGG** | Ins_k3_4_CAG;Ins_k3_5_AGC;Ins | 0.3898673413024689 | 0.03256826983708129 | 72.51013724121665 |
| **GATCCAGCATATCTAACTTGGGGTGGCTTTGTCTTCTTCTTTTGCCAATTCCACTAATTGTTTGGG** | Ins_k3_7_CAT | -0.19208247381501775 | 0.035652207716469084 | 40.519225330038815 |
| **GATCCAGCATCTAACTTGGGGTGGCTTTGTCTTCTTCTTTTGCCAATTCCACTAATTGTTTGGG** | Ins_k1_7_C | 0.34035882268033213 | 0.031516332008434636 | 69.0076837767622 |
| **GATCCAGCCAATCTAACTTGGGGTGGCTTTGTCTTCTTCTTTTGCCAATTCCACTAATTGTTTGGG** | Ins_k3_7_CCA | 0.18288077041158435 | 0.03656037686414165 | 58.95295808079565 |
| **GATCCAGCCAATTCCACTAATTGTTTGGG** | Del_k34_7-40;Del_k34_8-41 | -3.3028758992721095 | 0.07531683299170579 | 1.8057628526150056 |
| **GATCCAGCCACTAATTGTTTGGG** | Del_k40_8-47 | -5.262925819991648 | 0.17802733408196458 | 0.25434420661449864 |
| **GATCCAGCCATCTAACTTGGGGTGGCTTTGTCTTCTTCTTTTGCCAATTCCACTAATTGTTTGGG** | Ins_k2_7_CC | -0.8783479764184585 | 0.06829138785601295 | 20.399513619447934 |
| **GATCCAGCCCATCTAACTTGGGGTGGCTTTGTCTTCTTCTTTTGCCAATTCCACTAATTGTTTGGG** | Ins_k3_7_CCC | 0.01796436682446856 | 0.04233613273439836 | 49.99002080581556 |
| **GATCCAGCCGATCTAACTTGGGGTGGCTTTGTCTTCTTCTTTTGCCAATTCCACTAATTGTTTGGG** | Ins_k3_6_GCC;Ins_k3_7_CCG | 0.6352471741171237 | 0.03237432676432858 | 92.67569134703207 |
| **GATCCAGCCTATCTAACTTGGGGTGGCTTTGTCTTCTTCTTTTGCCAATTCCACTAATTGTTTGGG** | Ins_k3_7_CCT | -0.028813542695411143 | 0.040320087312810254 | 47.70544260433257 |
| **GATCCAGCGAATCTAACTTGGGGTGGCTTTGTCTTCTTCTTTTGCCAATTCCACTAATTGTTTGGG** | Ins_k3_7_CGA | 0.6381685700938668 | 0.030399254794218455 | 92.94682959713468 |
| **GATCCAGCGATCTAACTTGGGGTGGCTTTGTCTTCTTCTTTTGCCAATTCCACTAATTGTTTGGG** | Ins_k2_6_GC;Ins_k2_7_CG | 0.3679042935752147 | 0.03636488678324522 | 70.9349548578132 |
| **GATCCAGCGCATCTAACTTGGGGTGGCTTTGTCTTCTTCTTTTGCCAATTCCACTAATTGTTTGGG** | Ins_k3_7_CGC | 0.6174176375639748 | 0.032243455995468254 | 91.03797001147693 |
| **GATCCAGCGGATCTAACTTGGGGTGGCTTTGTCTTCTTCTTTTGCCAATTCCACTAATTGTTTGGG** | Ins_k3_6_GCG;Ins_k3_7_CGG | 0.7780447587431654 | 0.03050474118458226 | 100 |

| | | | | |
|---|---|---|---|---|
| GATCCAGCGTATCTAACTTGGGGTGGCTTTGTCTTCTTCTTTTGCCAATTCCACTAATTGTTTGGG | Ins_k3_7_CGT | 0.2422921116481993 | 0.032079709060256416 | 62.56156716086764 |
| GATCCAGCTAACTTGGGGTGGCTTTGTCTTCTTCTTTTGCCAATTCCACTAATTGTTTGGG | Del_k2_8-9 | 0.1898776486276862 | 0.03427836569551675 | 59.36689117999007 |
| GATCCAGCTAATCTAACTTGGGGTGGCTTTGTCTTCTTCTTTTGCCAATTCCACTAATTGTTTGGG | Ins_k3_7_CTA | -0.5007496804733718 | 0.0472973646583117 | 29.758337782641966 |
| GATCCAGCTAATTGTTTGGG | Del_k43_8-50 | -5.830226438860464 | 0.2601709829761952 | 0.1442269174522754 |
| GATCCAGCTATCTAACTTGGGGTGGCTTTGTCTTCTTCTTTTGCCAATTCCACTAATTGTTTGGG | Ins_k2_7_CT | -0.45507308130512353 | 0.049621771393747266 | 31.149118724824312 |
| GATCCAGCTCATCTAACTTGGGGTGGCTTTGTCTTCTTCTTTTGCCAATTCCACTAATTGTTTGGG | Ins_k3_7_CTC | 0.5360888464444293 | 0.03267061377551937 | 83.92704262329822 |
| GATCCAGCTCTAACTTGGGGTGGCTTTGTCTTCTTCTTTTGCCAATTCCACTAATTGTTTGGG | A8C | -0.05079973511816749 | 0.03554394359692441 | 46.66802775443374 |
| GATCCAGCTGATCTAACTTGGGGTGGCTTTGTCTTCTTCTTTTGCCAATTCCACTAATTGTTTGGG | Ins_k3_6_GCT;Ins_k3_7_CTG | 0.49598466907847927 | 0.03260017526764075 | 80.62781618070086 |
| GATCCAGCTTATCTAACTTGGGGTGGCTTTGTCTTCTTCTTTTGCCAATTCCACTAATTGTTTGGG | Ins_k3_7_CTT | -0.4660526148941059 | 0.04030020276908207 | 30.808986592511932 |
| GATCCAGCTTCTTCTTTTGCCAATTCCACTAATTGTTTGGG | Del_k22_8-29 | -0.37294964081635423 | 0.030439418292796692 | 33.81516579390475 |
| GATCCAGCTTCTTTTGCCAATTCCACTAATTGTTTGGG | Del_k25_8-32 | -1.0358137778667509 | 0.04023723049175399 | 17.42742747657352 |
| GATCCAGCTTGGGGTGGCTTTGTCTTCTTCTTTTGCCAATTCCACTAATTGTTTGGG | Del_k6_8-13 | 0.30275299662494526 | 0.032678747483325485 | 66.46078213904397 |
| GATCCAGCTTTGTCTTCTTCTTTTGCCAATTCCACTAATTGTTTGGG | Del_k16_7-22;Del_k16_8-23 | 0.08476480395444486 | 0.028228410494855246 | 53.44343681160642 |
| GATCCAGCTTTTGCCAATTCCACTAATTGTTTGGG | Del_k28_8-35 | -1.593570611042138 | 0.03840577028444024 | 9.97705966091203 |
| GATCCAGG | Del_k55_7-61;Del_k55_8-62 | -6.789730537569707 | 0.4667228921545509 | 0.05525085283536323 |
| GATCCAGGAAAATCTAACTTGGGGTGGCTTTGTCTTCTTCTTTTGCCAATTCCACTAATTGTTTGGG | Ins_k4_7_GAAA | 0.6352390203986171 | 0.03543062009388843 | 92.6749356986131 |
| GATCCAGGAAATCTAACTTGGGGTGGCTTTGTCTTCTTCTTTTGCCAATTCCACTAATTGTTTGGG | Ins_k3_7_GAA | 0.41431435725166854 | 0.03306913014031432 | 74.30463946124559 |
| GATCCAGGAATCTAACTTGGGGTGGCTTTGTCTTCTTCTTTTGCCAATTCCACTAATTGTTTGGG | Ins_k2_7_GA | 0.44581018002856254 | 0.03725496663097324 | 76.68216981794335 |
| GATCCAGGACATCTAACTTGGGGTGGCTTTGTCTTCTTCTTTTGCCAATTCCACTAATTGTTTGGG | Ins_k3_7_GAC | 0.48949463713123453 | 0.031688673414212006 | 80.10623345291124 |
| GATCCAGGAGATCTAACTTGGGGTGGCTTTGTCTTCTTCTTTTGCCAATTCCACTAATTGTTTGGG | Ins_k3_5_AGG;Ins_k3_6_GGA;In | 0.47855592841527805 | 0.034052349914574016 | 79.23474984131536 |
| GATCCAGGATAATCTAACTTGGGGTGGCTTTGTCTTCTTCTTTTGCCAATTCCACTAATTGTTTGGG | Ins_k4_7_GATA | -0.3043069167021471 | 0.048429110707569924 | 36.217850973347836 |
| GATCCAGGATATCTAACTTGGGGTGGCTTTGTCTTCTTCTTTTGCCAATTCCACTAATTGTTTGGG | Ins_k3_7_GAT | 0.32658412355040123 | 0.03398488912275153 | 68.06364057763368 |
| GATCCAGGATCTAACTTGGGGTGGCTTTGTCTTCTTCTTTTGCCAATTCCACTAATTGTTTGGG | Ins_k1_6_G;Ins_k1_7_G | 0.6457720706390608 | 0.03127424360848215 | 93.65624446703234 |
| GATCCAGGCAATCTAACTTGGGGTGGCTTTGTCTTCTTCTTTTGCCAATTCCACTAATTGTTTGGG | Ins_k3_7_GCA | -0.23375359651424954 | 0.041140950587985135 | 38.86544056074811 |
| GATCCAGGCATCTAACTTGGGGTGGCTTTGTCTTCTTCTTTTGCCAATTCCACTAATTGTTTGGG | Ins_k2_7_GC | 0.23028801356854345 | 0.04406622889205635 | 61.81506150040016 |
| GATCCAGGCCAATTCCACTAATTGTTTGGG | Del_k33_8-40 | -2.8078073105492938 | 0.06667937691288925 | 2.9625539706942843 |
| GATCCAGGCCATCTAACTTGGGGTGGCTTTGTCTTCTTCTTTTGCCAATTCCACTAATTGTTTGGG | Ins_k3_7_GCC | -0.4025031927024424 | 0.04794007159571159 | 32.830430423311704 |
| GATCCAGGCGATCTAACTTGGGGTGGCTTTGTCTTCTTCTTTTGCCAATTCCACTAATTGTTTGGG | Ins_k3_6_GGC;Ins_k3_7_GCG | 0.45073825384077537 | 0.03238141554311122 | 77.06099789055703 |
| GATCCAGGCTATCTAACTTGGGGTGGCTTTGTCTTCTTCTTTTGCCAATTCCACTAATTGTTTGGG | Ins_k3_7_GCT | -0.42693895190954845 | 0.04067642128875333 | 32.037916208297126 |
| GATCCAGGCTTTGTCTTCTTCTTTTGCCAATTCCACTAATTGTTTGGG | Del_k15_7-21;Del_k15_8-22 | -0.12745676938143363 | 0.03145728425696904 | 43.224275289228046 |
| GATCCAGGG | Del_k54_7-60;Del_k54_8-61 | -6.167197579834793 | 0.32579162989462523 | 0.10296784282984116 |
| GATCCAGGGAATCTAACTTGGGGTGGCTTTGTCTTCTTCTTTTGCCAATTCCACTAATTGTTTGGG | Ins_k3_7_GGA | -0.513422765503014 | 0.05099071171168209 | 29.383587474458075 |
| GATCCAGGGATCTAACTTGGGGTGGCTTTGTCTTCTTCTTTTGCCAATTCCACTAATTGTTTGGG | Ins_k2_6_GG;Ins_k2_7_GG | -0.17301164751138481 | 0.04604396068822756 | 41.29937585111915 |
| GATCCAGGGCATCTAACTTGGGGTGGCTTTGTCTTCTTCTTTTGCCAATTCCACTAATTGTTTGGG | Ins_k3_7_GGC | -0.7622843190268052 | 0.0448206274093672 | 22.910027934369396 |
| GATCCAGGGCTTTGTCTTCTTCTTTTGCCAATTCCACTAATTGTTTGGG | Del_k14_8-21 | -0.12627262452040822 | 0.035193654196522005 | 43.275489409167434 |
| GATCCAGGGG | Del_k53_8-60 | -6.373209997886754 | 0.5731592806213862 | 0.08379759567906746 |
| GATCCAGGGGATCTAACTTGGGGTGGCTTTGTCTTCTTCTTTTGCCAATTCCACTAATTGTTTGGG | Ins_k3_6_GGG;Ins_k3_7_GGG | -1.1635414414402432 | 0.061897375546269597 | 15.337757416379754 |
| GATCCAGGGGGTGGCTTTGTCTTCTTCTTTTGCCAATTCCACTAATTGTTTGGG | Del_k9_8-16 | -0.5150454887857094 | 0.0436133418509964 | 29.33594470890115 |
| GATCCAGGGGTGGCTTTGTCTTCTTCTTTTGCCAATTCCACTAATTGTTTGGG | Del_k10_7-16;Del_k10_8-17 | 0.2892424827536909 | 0.03419320943075063 | 65.56890127098585 |
| GATCCAGGGTATCTAACTTGGGGTGGCTTTGTCTTCTTCTTTTGCCAATTCCACTAATTGTTTGGG | Ins_k3_7_GGT | -2.8069072081656476 | 0.14554392159108692 | 2.9652217730524995 |
| GATCCAGGGTGGCTTTGTCTTCTTCTTTTGCCAATTCCACTAATTGTTTGGG | Del_k11_7-17;Del_k11_8-18 | 0.34438844254407086 | 0.02865406535543112 | 69.28631953110816 |
| GATCCAGGTAATCTAACTTGGGGTGGCTTTGTCTTCTTCTTTTGCCAATTCCACTAATTGTTTGGG | Ins_k3_7_GTA | -4.607800498770814 | 0.5538556245349385 | 0.48971021343834364 |
| GATCCAGGTATCTAACTTGGGGTGGCTTTGTCTTCTTCTTTTGCCAATTCCACTAATTGTTTGGG | Ins_k2_7_GT | -2.1962346653316702 | 0.10602568045503029 | 5.4609587493934795 |
| GATCCAGGTCATCTAACTTGGGGTGGCTTTGTCTTCTTCTTTTGCCAATTCCACTAATTGTTTGGG | Ins_k3_7_GTC | 0.2925619777131572 | 0.035275557704805456 | 65.78691856170624 |
| GATCCAGGTCTAACTTGGGGTGGCTTTGTCTTCTTCTTTTGCCAATTCCACTAATTGTTTGGG | A8G | 0.10232013808424134 | 0.03020267666069238 | 54.389937965512544 |
| GATCCAGGTCTTCTTCTTTTGCCAATTCCACTAATTGTTTGGG | Del_k20_8-27 | -0.4294738540115032 | 0.033179364415435134 | 31.95680607371642 |
| GATCCAGGTGATCTAACTTGGGGTGGCTTTGTCTTCTTCTTTTGCCAATTCCACTAATTGTTTGGG | Ins_k3_6_GGT;Ins_k3_7_GTG | -0.32966471076325093 | 0.04857422128551365 | 35.310992722717245 |
| GATCCAGGTGGCTTTGTCTTCTTCTTTTGCCAATTCCACTAATTGTTTGGG | Del_k12_7-18;Del_k12_8-19 | 0.13334718929765166 | 0.02761120126136107 | 56.103950228116844 |
| GATCCAGGTTATCTAACTTGGGGTGGCTTTGTCTTCTTCTTTTGCCAATTCCACTAATTGTTTGGG | Ins_k3_7_GTT | -1.366395381210869 | 0.06561119359101254 | 12.521706441010027 |
| GATCCAGGTTGATCTAACTTGGGGTGGCTTTGTCTTCTTCTTTTGCCAATTCCACTAATTGTTTGGG | Ins_k4_6_GGTT;Ins_k4_7_GTTG | -1.044199403172423 | 0.07047814677820204 | 17.281898627694563 |
| GATCCAGGTTTGGG | Del_k49_8-56 | -8.311641647823627 | 1.9411134090818514 | 0.012060946441534291 |
| GATCCAGTAAATCTAACTTGGGGTGGCTTTGTCTTCTTCTTTTGCCAATTCCACTAATTGTTTGGG | Ins_k3_7_TAA | -0.749311051833594 | 0.0567012034563899 | 23.20918215622462 |
| GATCCAGTAACTTGGGGTGGCTTTGTCTTCTTCTTTTGCCAATTCCACTAATTGTTTGGG | Del_k3_8-10 | 0.09761434789643797 | 0.03427571595252233 | 54.13459160353035 |
| GATCCAGTAATCTAACTTGGGGTGGCTTTGTCTTCTTCTTTTGCCAATTCCACTAATTGTTTGGG | Ins_k2_7_TA | -0.8104602205448591 | 0.05902109741791854 | 21.832480982806207 |
| GATCCAGTAATTGTTTGGG | Del_k44_8-51 | -6.172711533322532 | 0.45055818264962144 | 0.10240164536153709 |
| GATCCAGTACATCTAACTTGGGGTGGCTTTGTCTTCTTCTTTTGCCAATTCCACTAATTGTTTGGG | Ins_k3_7_TAC | -0.043953202550233494 | 0.0392035970597086 | 46.988638208910984 |
| GATCCAGTACCATCTAACTTGGGGTGGCTTTGTCTTCTTCTTTTGCCAATTCCACTAATTGTTTGGG | Ins_k4_7_TACC | 0.626676306474359 | 0.037539943253285774 | 91.88477452619703 |
| GATCCAGTAGATCTAACTTGGGGTGGCTTTGTCTTCTTCTTTTGCCAATTCCACTAATTGTTTGGG | Ins_k3_5_AGT;Ins_k3_6_GTA;Ins | -1.0467480522870412 | 0.0647510060599959 | 17.23790921266152 |
| GATCCAGTATATCTAACTTGGGGTGGCTTTGTCTTCTTCTTTTGCCAATTCCACTAATTGTTTGGG | Ins_k3_7_TAT | -0.839446229079781 | 0.04863573279101535 | 21.208728225956406 |
| GATCCAGTATCTAACTTGGGGTGGCTTTGTCTTCTTCTTTTGCCAATTCCACTAATTGTTTGGG | Ins_k1_7_T | -0.2611434436619999 | 0.047895324651396703 | 37.81536838911407 |
| GATCCAGTCAATCTAACTTGGGGTGGCTTTGTCTTCTTCTTTTGCCAATTCCACTAATTGTTTGGG | Ins_k3_7_TCA | -0.2766537552124463 | 0.046027515606757045 | 37.23336543521912 |
| GATCCAGTCATCTAACTTGGGGTGGCTTTGTCTTCTTCTTTTGCCAATTCCACTAATTGTTTGGG | Ins_k2_7_TC | 0.13363501910837217 | 0.03990847086696464 | 56.1201009417086 |
| GATCCAGTCCACTAATTGTTTGGG | Del_k39_8-46 | -5.01111954996892 | 0.1856358218433649 | 0.32717485861860046 |
| GATCCAGTCCATCTAACTTGGGGTGGCTTTGTCTTCTTCTTTTGCCAATTCCACTAATTGTTTGGG | Ins_k3_7_TCC | 0.43414358510531814 | 0.03052359299185105 | 75.79274835301402 |
| GATCCAGTCGATCTAACTTGGGGTGGCTTTGTCTTCTTCTTTTGCCAATTCCACTAATTGTTTGGG | Ins_k3_6_GTC;Ins_k3_7_TCG | 0.3885994565468175 | 0.03375678353213848 | 72.4182610001289 |
| GATCCAGTCTAACTTGGGGTGGCTTTGTCTTCTTCTTTTGCCAATTCCACTAATTGTTTGGG | Del_k1_8-8 | -0.30737433412266685 | 0.038867830761702006 | 36.10692592003379 |
| GATCCAGTCTATCTAACTTGGGGTGGCTTTGTCTTCTTCTTTTGCCAATTCCACTAATTGTTTGGG | Ins_k3_7_TCT | -0.6693549854828541 | 0.05483427435815545 | 25.141102316515756 |
| GATCCAGTCTTCTTCTTTTGCCAATTCCACTAATTGTTTGGG | Del_k21_7-27;Del_k21_8-28 | -0.5869731221462823 | 0.036597401344607934 | 27.299978445656013 |
| GATCCAGTCTTCTTTTGCCAATTCCACTAATTGTTTGGG | Del_k24_8-31 | -1.0776440093678024 | 0.03774507197910495 | 16.71347074040605 |
| GATCCAGTCTTTTGCCAATTCCACTAATTGTTTGGG | Del_k27_8-34 | -2.1370627028615914 | 0.05133963915184367 | 5.793846070481218 |
| GATCCAGTGAATCTAACTTGGGGTGGCTTTGTCTTCTTCTTTTGCCAATTCCACTAATTGTTTGGG | Ins_k3_7_TGA | 0.2161618449014706 | 0.037045092547221284 | 60.94799013220494 |
| GATCCAGTGATCTAACTTGGGGTGGCTTTGTCTTCTTCTTTTGCCAATTCCACTAATTGTTTGGG | Ins_k2_6_GT;Ins_k2_7_TG | 0.054903874632256766 | 0.03864151847450499 | 51.87115780679266 |
| GATCCAGTGCATCTAACTTGGGGTGGCTTTGTCTTCTTCTTTTGCCAATTCCACTAATTGTTTGGG | Ins_k3_7_TGC | 0.32049180189852844 | 0.03291064885083236 | 67.65023556327138 |
| GATCCAGTGCCAATTCCACTAATTGTTTGGG | Del_k32_8-39 | -2.3457442921595897 | 0.05075252614984367 | 4.702596216240263 |
| GATCCAGTGCCATCTAACTTGGGGTGGCTTTGTCTTCTTCTTTTGCCAATTCCACTAATTGTTTGGG | Ins_k4_7_TGCC | 0.35928117086905265 | 0.036810889684112225 | 70.32590377303023 |

| | | | | |
|---|---|---|---|---|
| **GATCCAGTGGATCTAACTTGGGGTGGCTTTGTCTTCTTCTTTTGCCAATTCCACTAATTGTTTGGG** | Ins_k3_6_GTG;Ins_k3_7_TGG | 0.2525012019283903 | 0.0378397007961176 | 63.203535227278145 |
| **GATCCAGTGGCTTTGTCTTCTTCTTTTGCCAATTCCACTAATTGTTTGGG** | Del_k13_7-19;Del_k13_8-20 | 0.2654089502722681 | 0.028420755343276444 | 64.02463845102338 |
| **GATCCAGTGGG** | Del_k52_8-59 | -7.406978462135406 | 0.5812222464764453 | 0.02980379849451762 |
| **GATCCAGTGGGGTGGCTTTGTCTTCTTCTTTTGCCAATTCCACTAATTGTTTGGG** | Del_k8_8-15 | -0.2943666800166518 | 0.039466632153755385 | 36.57966023999384 |
| **GATCCAGTGTATCTAACTTGGGGTGGCTTTGTCTTCTTCTTTTGCCAATTCCACTAATTGTTTGGG** | Ins_k3_7_TGT | -0.2629158672964314 | 0.04281159198898348 | 37.748402899575574 |
| **GATCCAGTGTCTTCTTCTTTTGCCAATTCCACTAATTGTTTGGG** | Del_k19_8-26 | 0.0015146968875018318 | 0.03008675752011129 | 49.1744279708542 |
| **GATCCAGTGTTTGGG** | Del_k48_8-55 | -5.957290612857109 | 0.465992956224138 | 0.12701735122248486 |
| **GATCCAGTTAATCTAACTTGGGGTGGCTTTGTCTTCTTCTTTTGCCAATTCCACTAATTGTTTGGG** | Ins_k3_7_TTA | -1.1249292520075287 | 0.05975512821869642 | 15.94156394104561 |
| **GATCCAGTTATCTAACTTGGGGTGGCTTTGTCTTCTTCTTTTGCCAATTCCACTAATTGTTTGGG** | Ins_k2_7_TT | -0.5344818238312797 | 0.05676240775946113 | 28.771266868975175 |
| **GATCCAGTTCATCTAACTTGGGGTGGCTTTGTCTTCTTCTTTTGCCAATTCCACTAATTGTTTGGG** | Ins_k3_7_TTC | 0.3607412395136844 | 0.031628706571343305 | 70.42865941692693 |
| **GATCCAGTTCCACTAATTGTTTGGG** | Del_k38_8-45 | -4.517672658164592 | 0.1124207091424368 | 0.535896827008309 |
| **GATCCAGTTCTAACTTGGGGTGGCTTTGTCTTCTTCTTTTGCCAATTCCACTAATTGTTTGGG** | A8T | -0.27919253093804425 | 0.039613459966563805 | 37.138958161025464 |
| **GATCCAGTTCTTCTTTTGCCAATTCCACTAATTGTTTGGG** | Del_k23_8-30 | -0.5905468402134603 | 0.037677150098956363 | 27.202590142270097 |
| **GATCCAGTTCTTTTGCCAATTCCACTAATTGTTTGGG** | Del_k26_8-33 | -1.7684498610476194 | 0.05152212336026988 | 8.376324159018841 |
| **GATCCAGTTGATCTAACTTGGGGTGGCTTTGTCTTCTTCTTTTGCCAATTCCACTAATTGTTTGGG** | Ins_k3_6_GTT;Ins_k3_7_TTG | 0.147174619174115 | 0.04007033287593803 | 56.885111948818 |
| **GATCCAGTTGCCAATTCCACTAATTGTTTGGG** | Del_k31_8-38 | -2.326713080148732 | 0.05428953857428008 | 4.792949359702596 |
| **GATCCAGTTGGG** | Del_k51_8-58 | -6.724785322852892 | 0.5818633173462173 | 0.05895821607713592 |
| **GATCCAGTTGGGGTGGCTTTGTCTTCTTCTTTTGCCAATTCCACTAATTGTTTGGG** | Del_k7_8-14 | 0.3880426082135392 | 0.03222197750321106 | 72.37794623783789 |
| **GATCCAGTTGTCTTCTTCTTTTGCCAATTCCACTAATTGTTTGGG** | Del_k18_8-25 | -0.18334480408646187 | 0.03046873087901466 | 40.874820211568746 |
| **GATCCAGTTGTTTGGG** | Del_k47_8-54 | -6.59226237202453 | 0.9817641058655927 | 0.06731290299047307 |
| **GATCCAGTTTATCTAACTTGGGGTGGCTTTGTCTTCTTCTTTTGCCAATTCCACTAATTGTTTGGG** | Ins_k3_7_TTT | -0.8208618027220738 | 0.058723714552956045 | 21.606565613238367 |
| **GATCCAGTTTGCCAATTCCACTAATTGTTTGGG** | Del_k30_8-37 | -2.308005288872406 | 0.04934826269108413 | 4.883458832379251 |
| **GATCCAGTTTGGG** | Del_k50_7-56;Del_k50_8-57 | -6.220670715702786 | 0.323721653416833 | 0.09760645202814425 |
| **GATCCAGTTTGTCTTCTTCTTTTGCCAATTCCACTAATTGTTTGGG** | Del_k17_8-24 | -0.15194048002086913 | 0.031360648914931046 | 42.17883499305724 |
| **GATCCAGTTTTGCCAATTCCACTAATTGTTTGGG** | Del_k29_8-36 | -2.0680661291984763 | 0.04576468064599509 | 6.207715200385437 |
| **GATCCATAACTTGGGGTGGCTTTGTCTTCTTCTTTTGCCAATTCCACTAATTGTTTGGG** | Del_k4_7-10 | -0.6723136136874674 | 0.05132773067148174 | 25.06682906974752 |
| **GATCCATAAGATCTAACTTGGGGTGGCTTTGTCTTCTTCTTTTGCCAATTCCACTAATTGTTTGGG** | Ins_k3_5_ATA;Ins_k3_6_TAA | -2.346626395666919 | 0.1286505151431872 | 4.6984498686471 |
| **GATCCATAAGGATCTAACTTGGGGTGGCTTTGTCTTCTTCTTTTGCCAATTCCACTAATTGTTTGGG** | Ins_k4_6_TAAG | -2.0563577021775625 | 0.1085026372512044 | 6.2808249456348175 |
| **GATCCATAATTGTTTGGG** | Del_k45_7-51 | -6.773034557067352 | 0.8729043448946601 | 0.05618106378057145 |
| **GATCCATACGATCTAACTTGGGGTGGCTTTGTCTTCTTCTTTTGCCAATTCCACTAATTGTTTGGG** | Ins_k3_6_TAC | -0.4488836849970596 | 0.04665847249919507 | 31.34251083809333 |
| **GATCCATAGAGATCTAACTTGGGGTGGCTTTGTCTTCTTCTTTTGCCAATTCCACTAATTGTTTGGG** | Ins_k4_5_ATAG;Ins_k4_6_TAGA | -0.19709180878908722 | 0.6595331630913003 | 40.31675849295711 |
| **GATCCATAGATCTAACTTGGGGTGGCTTTGTCTTCTTCTTTTGCCAATTCCACTAATTGTTTGGG** | Ins_k2_5_AT;Ins_k2_6_TA | -2.318533695865376 | 0.1413580272243611 | 4.832313502127724 |
| **GATCCATAGGATCTAACTTGGGGTGGCTTTGTCTTCTTCTTTTGCCAATTCCACTAATTGTTTGGG** | Ins_k3_6_TAG | -0.8997514755730402 | 0.06441708565922784 | 19.967532096145266 |
| **GATCCATAGTGATCTAACTTGGGGTGGCTTTGTCTTCTTCTTTTGCCAATTCCACTAATTGTTTGGG** | Ins_k4_6_TAGT | -3.183467865737405 | 0.19588309873336823 | 2.0347870487357724 |
| **GATCCATATCTAACTTGGGGTGGCTTTGTCTTCTTCTTTTGCCAATTCCACTAATTGTTTGGG** | G7T | -1.8021990023485426 | 0.05020951956179908 | 8.0983475318473 |
| **GATCCATATGATCTAACTTGGGGTGGCTTTGTCTTCTTCTTTTGCCAATTCCACTAATTGTTTGGG** | Ins_k3_6_TAT | -1.1031592537107477 | 0.06940037920219033 | 16.292416938852597 |
| **GATCCATCAGATCTAACTTGGGGTGGCTTTGTCTTCTTCTTTTGCCAATTCCACTAATTGTTTGGG** | Ins_k3_4_CAT;Ins_k3_5_ATC;Ins_ | 0.08644073270456065 | 0.040230459067490655 | 53.53307930009397 |
| **GATCCATCCACTAATTGTTTGGG** | Del_k40_7-46 | -5.458042163362706 | 0.203550789903767 | 0.20925888097823733 |
| **GATCCATCCGATCTAACTTGGGGTGGCTTTGTCTTCTTCTTTTGCCAATTCCACTAATTGTTTGGG** | Ins_k3_6_TCC | 0.2877410609303098 | 0.0374528452917628 | 65.47052855963017 |
| **GATCCATCGATCTAACTTGGGGTGGCTTTGTCTTCTTCTTTTGCCAATTCCACTAATTGTTTGGG** | Ins_k2_6_TC | 0.22970933753974976 | 0.04241923531587199 | 61.779300953974875 |
| **GATCCATCGGATCTAACTTGGGGTGGCTTTGTCTTCTTCTTTTGCCAATTCCACTAATTGTTTGGG** | Ins_k3_6_TCG | 0.5835376315263678 | 0.03357383838242165 | 88.00526712030187 |
| **GATCCATCTAACTTGGGGTGGCTTTGTCTTCTTCTTTTGCCAATTCCACTAATTGTTTGGG** | Del_k2_6-7;Del_k2_7-8 | -0.32526954013822074 | 0.04127157949792146 | 35.466532121384546 |
| **GATCCATCTGATCTAACTTGGGGTGGCTTTGTCTTCTTCTTTTGCCAATTCCACTAATTGTTTGGG** | Ins_k3_6_TCT | 0.2019387124148958 | 0.038395651742872454 | 60.08725448269367 |
| **GATCCATCTTCTTCTTTTGCCAATTCCACTAATTGTTTGGG** | Del_k22_7-28 | -1.1885519474245845 | 0.039704374629647585 | 14.958909677584856 |
| **GATCCATCTTCTTTTGCCAATTCCACTAATTGTTTGGG** | Del_k25_7-31 | -1.8729308015958626 | 0.04787209485999004 | 7.545325505751586 |
| **GATCCATCTTTTGCCAATTCCACTAATTGTTTGGG** | Del_k28_7-34 | -2.503073420680662 | 0.05774954675831052 | 4.018005415171017 |
| **GATCCATGAGATCTAACTTGGGGTGGCTTTGTCTTCTTCTTTTGCCAATTCCACTAATTGTTTGGG** | Ins_k3_5_ATG;Ins_k3_6_TGA | -0.7184111758917948 | 0.053302339242068265 | 23.937538107759913 |
| **GATCCATGATCTAACTTGGGGTGGCTTTGTCTTCTTCTTTTGCCAATTCCACTAATTGTTTGGG** | Ins_k1_6_T | -0.5352943134536288 | 0.05504823963750306 | 28.74790000717411 |
| **GATCCATGCCAATTCCACTAATTGTTTGGG** | Del_k33_7-39 | -4.500596215107285 | 0.13930549430325417 | 0.5451266203932287 |
| **GATCCATGCGATCTAACTTGGGGTGGCTTTGTCTTCTTCTTTTGCCAATTCCACTAATTGTTTGGG** | Ins_k3_6_TGC | -0.23552847658341086 | 0.04373936598737583 | 38.796520245657206 |
| **GATCCATGGATCTAACTTGGGGTGGCTTTGTCTTCTTCTTTTGCCAATTCCACTAATTGTTTGGG** | Ins_k2_6_TG | 0.5515918588600635 | 0.04153721662002821 | 85.23830258420446 |
| **GATCCATGGCTTTGTCTTCTTCTTTTGCCAATTCCACTAATTGTTTGGG** | Del_k14_7-20 | 0.2651605354647758 | 0.026971170164530833 | 64.00873575810178 |
| **GATCCATGGG** | Del_k53_7-59 | -6.60679767179142 | 0.7726348090129744 | 0.0663415662045392 |
| **GATCCATGGGATCTAACTTGGGGTGGCTTTGTCTTCTTCTTTTGCCAATTCCACTAATTGTTTGGG** | Ins_k3_6_TGG | -0.41977064650778373 | 0.04714660644355783 | 32.26839887424652 |
| **GATCCATGGGGTGGCTTTGTCTTCTTCTTTTGCCAATTCCACTAATTGTTTGGG** | Del_k9_7-15 | 0.0398225633541377 | 0.0344969041815788 | 51.09474212880677 |
| **GATCCATGTCTTCTTCTTTTGCCAATTCCACTAATTGTTTGGG** | Del_k20_7-26 | -0.9019074225194497 | 0.03763140969562887 | 19.924529528579075 |
| **GATCCATGTGATCTAACTTGGGGTGGCTTTGTCTTCTTCTTTTGCCAATTCCACTAATTGTTTGGG** | Ins_k3_6_TGT | -0.701287704184207 | 0.05811349910944408 | 24.35096138346285 |
| **GATCCATGTTTGGG** | Del_k49_7-55 | -6.129726869569452 | 0.38165636776392364 | 0.10689931864104439 |
| **GATCCATTAGATCTAACTTGGGGTGGCTTTGTCTTCTTCTTTTGCCAATTCCACTAATTGTTTGGG** | Ins_k3_5_ATT;Ins_k3_6_TTA | -2.942809793468427 | 0.16828994783784085 | 2.58842411657442 |
| **GATCCATTCCACTAATTGTTTGGG** | Del_k39_6-44;Del_k39_7-45 | -5.393586776111906 | 0.22131660084648003 | 0.22319091754370102 |
| **GATCCATTCGATCTAACTTGGGGTGGCTTTGTCTTCTTCTTTTGCCAATTCCACTAATTGTTTGGG** | Ins_k3_6_TTC | -0.21290947218044903 | 0.0484020340850554 | 39.68405868602716 |
| **GATCCATTCGGATCTAACTTGGGGTGGCTTTGTCTTCTTCTTTTGCCAATTCCACTAATTGTTTGGG** | Ins_k4_6_TTCG | -1.4200973068249247 | 0.09378897352487997 | 11.867003433035165 |
| **GATCCATTCTGATCTAACTTGGGGTGGCTTTGTCTTCTTCTTTTGCCAATTCCACTAATTGTTTGGG** | Ins_k4_6_TTCT | -0.8858826193085569 | 0.0733319883165354 | 20.246388166417752 |
| **GATCCATTCTTCTTTTGCCAATTCCACTAATTGTTTGGG** | Del_k24_7-30 | -2.1329276351976696 | 0.07412973501607809 | 5.817853618225874 |
| **GATCCATTCTTTTGCCAATTCCACTAATTGTTTGGG** | Del_k27_7-33 | -3.4296902291948097 | 0.10330621401722842 | 1.5906914654139188 |
| **GATCCATTGATCTAACTTGGGGTGGCTTTGTCTTCTTCTTTTGCCAATTCCACTAATTGTTTGGG** | Ins_k2_6_TT | -0.5319804848908628 | 0.605730051086242 | 28.843323640794527 |
| **GATCCATTGCCAATTCCACTAATTGTTTGGG** | Del_k32_7-38 | -3.885917290177692 | 0.11141062259913054 | 1.0079733639253257 |
| **GATCCATTGGATCTAACTTGGGGTGGCTTTGTCTTCTTCTTTTGCCAATTCCACTAATTGTTTGGG** | Ins_k3_6_TTG | -0.38023222252859323 | 0.07337063090953144 | 33.56979862357152 |
| **GATCCATTGGGGTGGCTTTGTCTTCTTCTTTTGCCAATTCCACTAATTGTTTGGG** | Del_k8_7-14 | -0.06627319531478604 | 0.044311880894426184 | 45.95146999544243 |
| **GATCCATTGTCTTCTTCTTTTGCCAATTCCACTAATTGTTTGGG** | Del_k19_7-25 | -1.1013754312215578 | 0.04661101008281036 | 16.321505655431377 |
| **GATCCATTGTTTGGG** | Del_k48_6-53;Del_k48_7-54 | -5.84290596106838 | 0.4010529337008748 | 0.14240973390472783 |
| **GATCCATTTGATCTAACTTGGGGTGGCTTTGTCTTCTTCTTTTGCCAATTCCACTAATTGTTTGGG** | Ins_k3_6_TTT | 0.12446968297865724 | 0.5981907120578852 | 55.60809130735148 |
| **GATCCATTTGCCAATTCCACTAATTGTTTGGG** | Del_k31_7-37 | -3.7209487398189753 | 0.09809142888908279 | 1.1887594626113351 |
| **GATCCATTTGGG** | Del_k51_7-57 | -6.534591002166084 | 0.48917859434713223 | 0.0713090545480747 |
| **GATCCATTTGTCTTCTTCTTTTGCCAATTCCACTAATTGTTTGGG** | Del_k18_7-24 | -1.4822478915522086 | 0.055643797643484046 | 11.15191400982558 |

| | | | | |
|---|---|---|---|---|
| GATCCATTTTGCCAATTCCACTAATTGTTTGGG | Del_k30_7-36 | -3.955581431453712 | 0.13220007316393678 | 0.9401438376447667 |
| GATCCCAAAGATCTAACTTGGGGTGGCTTTGTCTTCTTCTTTTGCCAATTCCACTAATTGTTTGGG | Ins_k3_5_CAA | -0.6432336858487537 | 0.05718920878167126 | 25.806472923078225 |
| GATCCCAAGATCTAACTTGGGGTGGCTTTGTCTTCTTCTTTTGCCAATTCCACTAATTGTTTGGG | Ins_k2_5_CA | -1.6437679399879608 | 0.08632026879779868 | 9.48860015924279 |
| GATCCCAATTCCACTAATTGTTTGGG | Del_k37_5-41;Del_k37_6-42 | -5.557008696296479 | 0.3591515044659819 | 0.1895410488118935 |
| GATCCCACAGATCTAACTTGGGGTGGCTTTGTCTTCTTCTTTTGCCAATTCCACTAATTGTTTGGG | Ins_k3_4_CCA;Ins_k3_5_CAC | -0.3041816923418356 | 0.0422838963285447 | 36.22238661454829 |
| GATCCCACTAATTGTTTGGG | Del_k43_5-47;Del_k43_6-48 | -7.029839330712699 | 0.6155867502568542 | 0.04345713209281138 |
| GATCCCAGAGATCTAACTTGGGGTGGCTTTGTCTTCTTCTTTTGCCAATTCCACTAATTGTTTGGG | Ins_k3_5_CAG | -0.9777637886841213 | 1.2074867472699011 | 18.469029462189315 |
| GATCCCAGATCTAACTTGGGGTGGCTTTGTCTTCTTCTTTTGCCAATTCCACTAATTGTTTGGG | Ins_k1_3_C;Ins_k1_4_C;Ins_k1_5 | -0.1352688417097555 | 0.03986629955249521 | 42.88791965221569 |
| GATCCCATAGATCTAACTTGGGGTGGCTTTGTCTTCTTCTTTTGCCAATTCCACTAATTGTTTGGG | Ins_k3_5_CAT | -2.31959504672011 | 0.11278032497454082 | 4.827187442816808 |
| GATCCCCAAGATCTAACTTGGGGTGGCTTTGTCTTCTTCTTTTGCCAATTCCACTAATTGTTTGGG | Ins_k3_5_CCA | -3.22706509374934 | 0.16403834921065547 | 1.9479819534819751 |
| GATCCCCAATTCCACTAATTGTTTGGG | Del_k36_6-41 | -5.402122405944204 | 0.4132845368594308 | 0.22129394991362744 |
| GATCCCCACTAATTGTTTGGG | Del_k42_6-47 | -5.558650234104401 | 0.4022712257311096 | 0.18923016524748812 |
| GATCCCCAGATCTAACTTGGGGTGGCTTTGTCTTCTTCTTTTGCCAATTCCACTAATTGTTTGGG | Ins_k2_3_CC;Ins_k2_4_CC;Ins_k | -1.6410304294246019 | 0.08567726866274518 | 9.514610888489623 |
| GATCCCCCAGATCTAACTTGGGGTGGCTTTGTCTTCTTCTTTTGCCAATTCCACTAATTGTTTGGG | Ins_k3_3_CCC;Ins_k3_4_CCC;In | -1.6073479029528328 | 0.07564341658713523 | 9.840545355621321 |
| GATCCCCGAGATCTAACTTGGGGTGGCTTTGTCTTCTTCTTTTGCCAATTCCACTAATTGTTTGGG | Ins_k3_5_CCG | -1.8447123020022556 | 0.07922949352535405 | 7.761275838030115 |
| GATCCCCTAGATCTAACTTGGGGTGGCTTTGTCTTCTTCTTTTGCCAATTCCACTAATTGTTTGGG | Ins_k3_5_CCT | -3.478403585952181 | 0.231989910402729 | 1.5150606161913618 |
| GATCCCGAAGATCTAACTTGGGGTGGCTTTGTCTTCTTCTTTTGCCAATTCCACTAATTGTTTGGG | Ins_k3_5_CGA | 0.7859669036864341 | 0.03478831297290923 | 100 |
| GATCCCGAGATCTAACTTGGGGTGGCTTTGTCTTCTTCTTTTGCCAATTCCACTAATTGTTTGGG | Ins_k2_5_CG | -0.13487076332884262 | 0.04465682856891788 | 42.90499580442958 |
| GATCCCGATAGATCTAACTTGGGGTGGCTTTGTCTTCTTCTTTTGCCAATTCCACTAATTGTTTGGG | Ins_k4_5_CGAT | -0.687572284406367 | 0.0523370844880618 | 24.68724591096027 |
| GATCCCGATCTAACTTGGGGTGGCTTTGTCTTCTTCTTTTGCCAATTCCACTAATTGTTTGGG | A6C | -0.13073399667351926 | 0.037498091504384816 | 43.082851380293576 |
| GATCCCGCAGATCTAACTTGGGGTGGCTTTGTCTTCTTCTTTTGCCAATTCCACTAATTGTTTGGG | Ins_k3_4_CCG;Ins_k3_5_CGC | 0.3664383344747224 | 0.034275513983245805 | 70.83104329885327 |
| GATCCCGGAGATCTAACTTGGGGTGGCTTTGTCTTCTTCTTTTGCCAATTCCACTAATTGTTTGGG | Ins_k3_5_CGG | 0.749360372004372 | 0.03286892081829195 | 100 |
| GATCCCGTAGATCTAACTTGGGGTGGCTTTGTCTTCTTCTTTTGCCAATTCCACTAATTGTTTGGG | Ins_k3_5_CGT | -0.9553476807291845 | 0.063655093209410707 | 18.88770826364627 |
| GATCCCTAACTTGGGGTGGCTTTGTCTTCTTCTTTTGCCAATTCCACTAATTGTTTGGG | Del_k4_6-9 | -1.4429800723972193 | 0.06639187841937537 | 11.598536918050245 |
| GATCCCTAAGATCTAACTTGGGGTGGCTTTGTCTTCTTCTTTTGCCAATTCCACTAATTGTTTGGG | Ins_k3_5_CTA | -3.2271867173787925 | 0.1799394179816279 | 1.9477450472536746 |
| GATCCCTAATTGTTTGGG | Del_k45_6-50 | -6.271876268311017 | 1.0414332544323677 | 0.09273426573926444 |
| GATCCCTAGATCTAACTTGGGGTGGCTTTGTCTTCTTCTTTTGCCAATTCCACTAATTGTTTGGG | Ins_k2_5_CT | -2.876049241691811 | 0.16181603908465036 | 2.767127540344114 |
| GATCCCTCAGATCTAACTTGGGGTGGCTTTGTCTTCTTCTTTTGCCAATTCCACTAATTGTTTGGG | Ins_k3_4_CCT;Ins_k3_5_CTC | -0.6728169549828147 | 0.05871338536813379 | 25.054215074372006 |
| GATCCCTGAGATCTAACTTGGGGTGGCTTTGTCTTCTTCTTTTGCCAATTCCACTAATTGTTTGGG | Ins_k3_5_CTG | -1.5385448049330182 | 0.07718177762743576 | 10.541440776855007 |
| GATCCCTTAGATCTAACTTGGGGTGGCTTTGTCTTCTTCTTTTGCCAATTCCACTAATTGTTTGGG | Ins_k3_5_CTT | -3.448597899433195 | 0.2267995199824347 | 1.560897748223948 |
| GATCCCTTCTTCTTTTGCCAATTCCACTAATTGTTTGGG | Del_k24_6-29 | -2.7090623488944443 | 0.09128250170529073 | 3.2700219123331777 |
| GATCCCTTCTTTTGCCAATTCCACTAATTGTTTGGG | Del_k27_6-32 | -3.8126209151471864 | 0.14476134690743744 | 1.0846291373335677 |
| GATCCCTTGGGGTGGCTTTGTCTTCTTCTTTTGCCAATTCCACTAATTGTTTGGG | Del_k8_6-13 | -0.2762103994535434 | 0.04014376980056889 | 37.24987672212476 |
| GATCCCTTTGTCTTCTTCTTTTGCCAATTCCACTAATTGTTTGGG | Del_k18_6-23 | -2.1135484333139942 | 0.07413610625248393 | 5.931698526875353 |
| GATCCCTTTTGCCAATTCCACTAATTGTTTGGG | Del_k30_6-35 | -4.4175434990147 33 | 0.14811254251666284 | 0.5923340887254606 |
| GATCCG | Del_k57_6-62 | -7.685702719720574 | 0.5262813110535717 | 0.022553981060330 26 |
| GATCCGAAAGATCTAACTTGGGGTGGCTTTGTCTTCTTCTTTTGCCAATTCCACTAATTGTTTGGG | Ins_k3_5_GAA | 0.20736558206809785 | 0.036130216839701336 | 60.41422659564006 |
| GATCCGAAGATCTAACTTGGGGTGGCTTTGTCTTCTTCTTTTGCCAATTCCACTAATTGTTTGGG | Ins_k2_5_GA | 0.7026458354186427 | 0.03282351020706371 | 99.13721198711298 |
| GATCCGACAGATCTAACTTGGGGTGGCTTTGTCTTCTTCTTTTGCCAATTCCACTAATTGTTTGGG | Ins_k3_4_CGA;Ins_k3_5_GAC | 0.4032829544688248 | 0.03332934227081809 | 73.48945961006574 |
| GATCCGAGAGATCTAACTTGGGGTGGCTTTGTCTTCTTCTTTTGCCAATTCCACTAATTGTTTGGG | Ins_k3_5_GAG | 0.23753192712729132 | 0.04020530940741359 | 62.26447023635724 |
| GATCCGAGATCTAACTTGGGGTGGCTTTGTCTTCTTCTTTTGCCAATTCCACTAATTGTTTGGG | Ins_k1_5_G | 0.4283574411441433 | 0.03458453891234524 | 75.35546690652579 |
| GATCCGATAGATCTAACTTGGGGTGGCTTTGTCTTCTTCTTTTGCCAATTCCACTAATTGTTTGGG | Ins_k3_5_GAT | -1.496244913516544 | 0.07140993120728133 | 10.99690776805438 |
| GATCCGATCTAACTTGGGGTGGCTTTGTCTTCTTCTTTTGCCAATTCCACTAATTGTTTGGG | Del_k1_6-6 | -0.4158974373110418 | 0.039293177301329724 | 32.393623487543344 |
| GATCCGCAAGATCTAACTTGGGGTGGCTTTGTCTTCTTCTTTTGCCAATTCCACTAATTGTTTGGG | Ins_k3_5_GCA | 0.5553289835811683 | 0.031542170704112055 | 85.55744471752404 |
| GATCCGCAGATCTAACTTGGGGTGGCTTTGTCTTCTTCTTTTGCCAATTCCACTAATTGTTTGGG | Ins_k2_4_CG;Ins_k2_5_GC | 0.2573531343171688 | 0.03729784965008724 | 63.51093965662953 |
| GATCCGCCAATTCCACTAATTGTTTGGG | Del_k35_6-40 | -3.6410839108733772 | 0.08363760974546433 | 1.287593695262183 |
| GATCCGCCAGATCTAACTTGGGGTGGCTTTGTCTTCTTCTTTTGCCAATTCCACTAATTGTTTGGG | Ins_k3_3_CCG;Ins_k3_4_CGC;In | 0.6947766057414217 | 0.03170529481723088 | 98.36013998545211 |
| GATCCGCGAGATCTAACTTGGGGTGGCTTTGTCTTCTTCTTTTGCCAATTCCACTAATTGTTTGGG | Ins_k3_5_GCG | 0.6344000617045467 | 0.032430926668323624 | 92.59721786117252 |
| GATCCGCTAGATCTAACTTGGGGTGGCTTTGTCTTCTTCTTTTGCCAATTCCACTAATTGTTTGGG | Ins_k3_5_GCT | -0.7713499024965599 | 0.05070625394154323 | 22.703273753480612 |
| GATCCGCTTTGTCTTCTTCTTTTGCCAATTCCACTAATTGTTTGGG | Del_k17_6-22 | -0.6462448501342336 | 0.0336249279570005 | 25.728882271204856 |
| GATCCGG | Del_k56_6-61 | -7.6557994983304 68 | 0.536438365117729 | 0.02323860293380021 |
| GATCCGGAAGATCTAACTTGGGGTGGCTTTGTCTTCTTCTTTTGCCAATTCCACTAATTGTTTGGG | Ins_k3_5_GGA | 0.9436109558390999 | 0.02943570278645304 | 100 |
| GATCCGGAGAGATCTAACTTGGGGTGGCTTTGTCTTCTTCTTTTGCCAATTCCACTAATTGTTTGGG | Ins_k4_5_GGAG | 0.718073779094603 | 0.029191011098843526 | 100 |
| GATCCGGAGATCTAACTTGGGGTGGCTTTGTCTTCTTCTTTTGCCAATTCCACTAATTGTTTGGG | Ins_k2_5_GG | 0.5087248666452795 | 0.0344220694269816 | 81.66160182187642 |
| GATCCGGATCTAACTTGGGGTGGCTTTGTCTTCTTCTTTTGCCAATTCCACTAATTGTTTGGG | A6G | 0.13394622982909377 | 0.026090008204749517 | 56.13756883672675 |
| GATCCGGCAGATCTAACTTGGGGTGGCTTTGTCTTCTTCTTTTGCCAATTCCACTAATTGTTTGGG | Ins_k3_4_CGG;Ins_k3_5_GGC | 0.5693556636239729 | 0.03108924697382563 | 86.76598772814214 |
| GATCCGGCTTTGTCTTCTTCTTTTGCCAATTCCACTAATTGTTTGGG | Del_k16_6-21 | 0.2239963591862305 | 0.028928528573870554 | 61.42736340793495 |
| GATCCGGG | Del_k55_6-60 | -6.889721336395727 | 0.40314964207849957 | 0.049993499020600914 |
| GATCCGGGAGATCTAACTTGGGGTGGCTTTGTCTTCTTCTTTTGCCAATTCCACTAATTGTTTGGG | Ins_k3_5_GGG | 0.35416214677616964 | 0.03439100895887371 | 69.96682363133382 |
| GATCCGGGCAGATCTAACTTGGGGTGGCTTTGTCTTCTTCTTTTGCCAATTCCACTAATTGTTTGGG | Ins_k4_4_CGGG;Ins_k4_5_GGG( | 0.38171824602985416 | 0.03976816663494753 | 71.92164631561063 |
| GATCCGGGGTGGCTTTGTCTTCTTCTTTTGCCAATTCCACTAATTGTTTGGG | Del_k11_6-16 | 0.39275995127682295 | 0.03517200342909521 | 72.72018443328506 |
| GATCCGGGTGGCTTTGTCTTCTTCTTTTGCCAATTCCACTAATTGTTTGGG | Del_k12_6-17 | 0.46415595989987013 | 0.027635703929662254 | 78.1019476707377 |
| GATCCGGTAGATCTAACTTGGGGTGGCTTTGTCTTCTTCTTTTGCCAATTCCACTAATTGTTTGGG | Ins_k3_5_GGT | -0.5463782082758728 | 0.0492732762921241 | 28.431020679819483 |
| GATCCGGTGGCTTTGTCTTCTTCTTTTGCCAATTCCACTAATTGTTTGGG | Del_k13_6-18 | 0.422134181137672 | 0.027806031596441794 | 74.88796644035098 |
| GATCCGTAAGATCTAACTTGGGGTGGCTTTGTCTTCTTCTTTTGCCAATTCCACTAATTGTTTGGG | Ins_k3_5_GTA | -1.9708390441673835 | 0.10282476937960944 | 6.841588836054238 |
| GATCCGTAGATCTAACTTGGGGTGGCTTTGTCTTCTTCTTTTGCCAATTCCACTAATTGTTTGGG | Ins_k2_5_GT | -0.12792727277351956 | 1.2282268299084544 | 43.20394290468704 |
| GATCCGTCAGATCTAACTTGGGGTGGCTTTGTCTTCTTCTTTTGCCAATTCCACTAATTGTTTGGG | Ins_k3_4_CGT;Ins_k3_5_GTC | 0.5549595798723659 | 0.03157580723142916 | 85.52584531695695 |
| GATCCGTCTTCTTCTTTTGCCAATTCCACTAATTGTTTGGG | Del_k22_6-27 | -0.7686870537257904 | 0.03207775822047221 | 22.7638097013542 |
| GATCCGTGAGATCTAACTTGGGGTGGCTTTGTCTTCTTCTTTTGCCAATTCCACTAATTGTTTGGG | Ins_k3_5_GTG | -0.295784382894229 | 0.04288400314331055 | 36.52783789343546 |
| GATCCGTGGCTTTGTCTTCTTCTTTTGCCAATTCCACTAATTGTTTGGG | Del_k14_6-19 | 0.3349608436900543 | 0.02750030104695937 | 68.63618532205942 |
| GATCCGTTAGATCTAACTTGGGGTGGCTTTGTCTTCTTCTTTTGCCAATTCCACTAATTGTTTGGG | Ins_k3_5_GTT | -2.6747097414191363 | 0.12762434000125253 | 3.384307455907057 |
| GATCCGTTTGGG | Del_k51_6-56 | -6.407405158595397 | 0.4345011533130593 | 0.08098056237430025 |
| GATCCTAAAGATCTAACTTGGGGTGGCTTTGTCTTCTTCTTTTGCCAATTCCACTAATTGTTTGGG | Ins_k3_5_TAA | -1.1506265127446433 | 0.06363043592721929 | 15.537128117790722 |
| GATCCTAACTTGGGGTGGCTTTGTCTTCTTCTTTTGCCAATTCCACTAATTGTTTGGG | Del_k5_5-9;Del_k5_6-10 | -0.6852329743381483 | 0.046321290024423735 | 24.745064635471437 |
| GATCCTAAGATCTAACTTGGGGTGGCTTTGTCTTCTTCTTTTGCCAATTCCACTAATTGTTTGGG | Ins_k2_5_TA | -2.141974045416428 | 0.12754416072489688 | 5.76546027097011 |

| | | | | |
|---|---|---|---|---|
| **GATCCTACAGATCTAACTTGGGGTGGCTTTGTCTTCTTCTTTTGCCAATTCCACTAATTGTTTGGG** | Ins_k3_4_CTA;Ins_k3_5_TAC | -0.21635790540584976 | 0.045055849866111254 | 39.54744654383966 |
| **GATCCTAGAGATCTAACTTGGGGTGGCTTTGTCTTCTTCTTTTGCCAATTCCACTAATTGTTTGGG** | Ins_k3_5_TAG | -2.0916241908604123 | 0.09433046116739152 | 6.063182601214066 |
| **GATCCTAGATCTAACTTGGGGTGGCTTTGTCTTCTTCTTTTGCCAATTCCACTAATTGTTTGGG** | Ins_k1_5_T | -2.0828488873548365 | 0.10909113560176273 | 6.116623003689725 |
| **GATCCTATAGATCTAACTTGGGGTGGCTTTGTCTTCTTCTTTTGCCAATTCCACTAATTGTTTGGG** | Ins_k3_5_TAT | -3.4019838651528524 | 0.1804091747817893 | 1.6353799614061064 |
| **GATCCTCAAGATCTAACTTGGGGTGGCTTTGTCTTCTTCTTTTGCCAATTCCACTAATTGTTTGGG** | Ins_k3_5_TCA | 0.07397571981883733 | 0.04051600621011794 | 52.86993044279094 |
| **GATCCTCAGATCTAACTTGGGGTGGCTTTGTCTTCTTCTTTTGCCAATTCCACTAATTGTTTGGG** | Ins_k2_4_CT;Ins_k2_5_TC | -0.42616986108207 | 0.05870456011571841 | 32.06256575343897 |
| **GATCCTCCACTAATTGTTTGGG** | Del_k41_6-46 | -5.297303191878777 | 0.25095843907651866 | 0.2457491061785398 |
| **GATCCTCCAGATCTAACTTGGGGTGGCTTTGTCTTCTTCTTTTGCCAATTCCACTAATTGTTTGGG** | Ins_k3_2_TCC;Ins_k3_3_CCT;Ins | 0.3522529150661988 | 0.03626899495268937 | 69.83336819222797 |
| **GATCCTCGAGATCTAACTTGGGGTGGCTTTGTCTTCTTCTTTTGCCAATTCCACTAATTGTTTGGG** | Ins_k3_5_TCG | 0.17025706458754197 | 0.03445566411682487 | 58.213430888159564 |
| **GATCCTCTAACTTGGGGTGGCTTTGTCTTCTTCTTTTGCCAATTCCACTAATTGTTTGGG** | Del_k3_6-8 | -0.850697365847894 | 0.05285743649723472 | 20.971443289758582 |
| **GATCCTCTAGATCTAACTTGGGGTGGCTTTGTCTTCTTCTTTTGCCAATTCCACTAATTGTTTGGG** | Ins_k3_5_TCT | -1.8490682220364856 | 0.08063160803483561 | 7.7275418656981385 |
| **GATCCTCTTCTTCTTTTGCCAATTCCACTAATTGTTTGGG** | Del_k23_6-28 | -1.7517949415644187 | 0.05879422191393923 | 8.516999378915788 |
| **GATCCTCTTCTTTTGCCAATTCCACTAATTGTTTGGG** | Del_k26_6-31 | -2.400138892068669 | 0.059483875002199656 | 4.453632889265553 |
| **GATCCTCTTTTGCCAATTCCACTAATTGTTTGGG** | Del_k29_6-34 | -3.5769038486789766 | 0.09661129562915605 | 1.3729410437336869 |
| **GATCCTGAAGATCTAACTTGGGGTGGCTTTGTCTTCTTCTTTTGCCAATTCCACTAATTGTTTGGG** | Ins_k3_5_TGA | 0.7582491534867901 | 0.03045100086068427 | 100 |
| **GATCCTGAGATCTAACTTGGGGTGGCTTTGTCTTCTTCTTTTGCCAATTCCACTAATTGTTTGGG** | Ins_k2_5_TG | 0.05522066465077047 | 0.04283822460583277 | 51.88759267489824 |
| **GATCCTGATCTAACTTGGGGTGGCTTTGTCTTCTTCTTTTGCCAATTCCACTAATTGTTTGGG** | A6T | -0.13077698605439209 | 0.03358342154459094 | 43.08099931499636 |
| **GATCCTGCAGATCTAACTTGGGGTGGCTTTGTCTTCTTCTTTTGCCAATTCCACTAATTGTTTGGG** | Ins_k3_4_CTG;Ins_k3_5_TGC | 0.5589775836171047 | 0.03532679125370272 | 85.87017978905692 |
| **GATCCTGCCAATTCCACTAATTGTTTGGG** | Del_k34_6-39 | -3.0918607015727555 | 0.06664267287256781 | 2.2299927475069867 |
| **GATCCTGGAGATCTAACTTGGGGTGGCTTTGTCTTCTTCTTTTGCCAATTCCACTAATTGTTTGGG** | Ins_k3_5_TGG | 0.5340180956100442 | 0.03257489229940821 | 83.75343044557704 |
| **GATCCTGGCTTTGTCTTCTTCTTTTGCCAATTCCACTAATTGTTTGGG** | Del_k15_6-20 | 0.10420039144955828 | 0.030985991304172653 | 54.49230103350567 |
| **GATCCTGGGGTGGCTTTGTCTTCTTCTTTTGCCAATTCCACTAATTGTTTGGG** | Del_k10_6-15 | 0.21277720907572095 | 0.03877403366217608 | 60.74205209053093 |
| **GATCCTGTAGATCTAACTTGGGGTGGCTTTGTCTTCTTCTTTTGCCAATTCCACTAATTGTTTGGG** | Ins_k3_5_TGT | -1.7560265257865366 | 0.08713054998674204 | 8.481035125273875 |
| **GATCCTGTCTTCTTCTTTTGCCAATTCCACTAATTGTTTGGG** | Del_k21_6-26 | -0.5562154134436983 | 0.035103473042438015 | 28.152710040047136 |
| **GATCCTGTTTGGG** | Del_k50_6-55 | -6.011356831508759 | 0.45060910997597087 | 0.12033234812132602 |
| **GATCCTTAAGATCTAACTTGGGGTGGCTTTGTCTTCTTCTTTTGCCAATTCCACTAATTGTTTGGG** | Ins_k3_5_TTA | -3.4164279846449155 | 0.17676197311489372 | 1.611928116189453 |
| **GATCCTTAGATCTAACTTGGGGTGGCTTTGTCTTCTTCTTTTGCCAATTCCACTAATTGTTTGGG** | Ins_k2_5_TT | -3.4662729881005943 | 0.3099428685356203 | 1.5335511310052492 |
| **GATCCTTCAGATCTAACTTGGGGTGGCTTTGTCTTCTTCTTTTGCCAATTCCACTAATTGTTTGGG** | Ins_k3_4_CTT;Ins_k3_5_TTC | -0.13710445969041418 | 0.041320585013575675 | 42.80926602683877 |
| **GATCCTTCCACTAATTGTTTGGG** | Del_k40_6-45 | -5.440794912062264 | 0.19952373596409873 | 0.21289932506994239 |
| **GATCCTTCTTCTTTTGCCAATTCCACTAATTGTTTGGG** | Del_k25_5-29;Del_k25_6-30 | -2.0457323084443693 | 0.05662137636550716 | 6.347916992139174 |
| **GATCCTTCTTTTGCCAATTCCACTAATTGTTTGGG** | Del_k28_5-32;Del_k28_6-33 | -3.1648114397817126 | 0.08390669323803572 | 2.073105231395326 |
| **GATCCTTGAGATCTAACTTGGGGTGGCTTTGTCTTCTTCTTTTGCCAATTCCACTAATTGTTTGGG** | Ins_k3_5_TTG | -2.348662926786517 | 0.1340498889557056 | 4.688891065979008 |
| **GATCCTTGCCAATTCCACTAATTGTTTGGG** | Del_k33_6-38 | -4.424614952078976 | 0.12418838206093105 | 0.5881602011368554 |
| **GATCCTTGGG** | Del_k53_6-58 | -5.912432715527715 | 0.535307105870828 | 0.1328448091270341 |
| **GATCCTTGGGGTGGCTTTGTCTTCTTCTTTTGCCAATTCCACTAATTGTTTGGG** | Del_k9_5-13;Del_k9_6-14 | -0.19426780283106326 | 0.04673699050046273 | 40.43077417385752 |
| **GATCCTTGTCTTCTTCTTTTGCCAATTCCACTAATTGTTTGGG** | Del_k20_6-25 | -1.435451863814309 | 0.04687422390711122 | 11.686182616841776 |
| **GATCCTTGTTTGGG** | Del_k49_6-54 | -6.076276886324641 | 0.35227433087775534 | 0.11276854315788767 |
| **GATCCTTTAGATCTAACTTGGGGTGGCTTTGTCTTCTTCTTTTGCCAATTCCACTAATTGTTTGGG** | Ins_k3_5_TTT | -3.8178318521879526 | 0.2812728455778157 | 1.0789919035774866 |
| **GATCCTTTGCCAATTCCACTAATTGTTTGGG** | Del_k32_6-37 | -4.002358296440595 | 0.10752104250680068 | 0.8971795572581167 |
| **GATCCTTTGTCTTCTTCTTTTGCCAATTCCACTAATTGTTTGGG** | Del_k19_5-23;Del_k19_6-24 | -1.8304426049334344 | 0.06279380694819973 | 7.8728208571564116 |
| **GATCCTTTTGCCAATTCCACTAATTGTTTGGG** | Del_k31_5-35;Del_k31_6-36 | -4.25015293311683 | 0.14864358080723827 | 0.7002667827606377 |
| **GATCG** | Del_k58_5-62 | -7.3193362354280636 | 0.6977848010256321 | 0.032533752157266706 |
| **GATCGAACAGATCTAACTTGGGGTGGCTTTGTCTTCTTCTTTTGCCAATTCCACTAATTGTTTGGG** | Ins_k3_4_GAA | -0.053535552246576135 | 0.05036370872458718 | 46.54052705358006 |
| **GATCGACAGATCTAACTTGGGGTGGCTTTGTCTTCTTCTTTTGCCAATTCCACTAATTGTTTGGG** | Ins_k2_4_GA | 0.33145441841310463 | 0.035932471752167114 | 68.39593911577053 |
| **GATCGACCAGATCTAACTTGGGGTGGCTTTGTCTTCTTCTTTTGCCAATTCCACTAATTGTTTGGG** | Ins_k3_3_CGA;Ins_k3_4_GAC | 0.6486498854781582 | 0.0339937511287745 | 93.92615799142537 |
| **GATCGAGATCTAACTTGGGGTGGCTTTGTCTTCTTCTTTTGCCAATTCCACTAATTGTTTGGG** | C5G | -0.11249531671385754 | 0.03432799277337179 | 43.875835225638056 |
| **GATCGAGCAGATCTAACTTGGGGTGGCTTTGTCTTCTTCTTTTGCCAATTCCACTAATTGTTTGGG** | Ins_k3_4_GAG | 0.08460028219400362 | 0.03198849580843584 | 53.43464492654617 |
| **GATCGATCAGATCTAACTTGGGGTGGCTTTGTCTTCTTCTTTTGCCAATTCCACTAATTGTTTGGG** | Ins_k3_4_GAT | -0.9703976361621389 | 0.051968834087631026 | 18.605577449364606 |
| **GATCGATCTAACTTGGGGTGGCTTTGTCTTCTTCTTTTGCCAATTCCACTAATTGTTTGGG** | Del_k2_5-6 | -0.5983586672478145 | 0.04642326231265927 | 26.99091607025481 |
| **GATCGCACAGATCTAACTTGGGGTGGCTTTGTCTTCTTCTTTTGCCAATTCCACTAATTGTTTGGG** | Ins_k3_4_GCA | -0.25013604547508245 | 0.04416652669525316 | 38.23391654362573 |
| **GATCGCAGATCTAACTTGGGGTGGCTTTGTCTTCTTCTTTTGCCAATTCCACTAATTGTTTGGG** | Ins_k1_4_G | 0.2679067536048456 | 0.03571997223582013 | 64.18475929825655 |
| **GATCGCCAATTCCACTAATTGTTTGGG** | Del_k36_5-40 | -3.929097738965973 | 0.08176078011779339 | 0.9653749497302326 |
| **GATCGCCAGATCTAACTTGGGGTGGCTTTGTCTTCTTCTTTTGCCAATTCCACTAATTGTTTGGG** | Ins_k2_3_CG;Ins_k2_4_GC | 0.4037571542891365 | 0.034914084459932715 | 73.52431656253476 |
| **GATCGCCCAGATCTAACTTGGGGTGGCTTTGTCTTCTTCTTTTGCCAATTCCACTAATTGTTTGGG** | Ins_k3_3_CGC;Ins_k3_4_GCC | 0.46735193941563336 | 0.037921857696413305 | 78.35195919868823 |
| **GATCGCGCAGATCTAACTTGGGGTGGCTTTGTCTTCTTCTTTTGCCAATTCCACTAATTGTTTGGG** | Ins_k3_4_GCG | 0.5590136813693094 | 0.03235585425102946 | 85.87327956547585 |
| **GATCGCTCAGATCTAACTTGGGGTGGCTTTGTCTTCTTCTTTTGCCAATTCCACTAATTGTTTGGG** | Ins_k3_4_GCT | -0.6751300218432681 | 0.04788566662675558 | 24.99632997163368 |
| **GATCGCTGCAGATCTAACTTGGGGTGGCTTTGTCTTCTTCTTTTGCCAATTCCACTAATTGTTTGGG** | Ins_k4_4_GCTG | 0.9011206756602199 | 0.03295733229090933 | 100 |
| **GATCGCTTTGTCTTCTTCTTTTGCCAATTCCACTAATTGTTTGGG** | Del_k18_5-22 | -1.4472359978065228 | 0.05045404182160392 | 11.549279302788005 |
| **GATCGGACAGATCTAACTTGGGGTGGCTTTGTCTTCTTCTTTTGCCAATTCCACTAATTGTTTGGG** | Ins_k3_4_GGA | -0.1911397240595143 | 0.045792441157412836 | 40.55744283175153 |
| **GATCGGCAGATCTAACTTGGGGTGGCTTTGTCTTCTTCTTTTGCCAATTCCACTAATTGTTTGGG** | Ins_k2_4_GG | 0.24668450106804907 | 0.23581187502075932 | 62.83696631394256 |
| **GATCGGCCAGATCTAACTTGGGGTGGCTTTGTCTTCTTCTTTTGCCAATTCCACTAATTGTTTGGG** | Ins_k3_3_CGG;Ins_k3_4_GGC | 0.8146306960697822 | 0.030333632181056346 | 100 |
| **GATCGGCTTTGTCTTCTTCTTTTGCCAATTCCACTAATTGTTTGGG** | Del_k17_5-21 | 0.00922072417297326 | 0.0281570307300265 | 49.5548312712 |
| **GATCGGG** | Del_k56_5-60 | -7.630806285924559 | 0.6559340653473373 | 0.023826729241173218 |
| **GATCGGGCAGATCTAACTTGGGGTGGCTTTGTCTTCTTCTTTTGCCAATTCCACTAATTGTTTGGG** | Ins_k3_4_GGG | -0.9409027767461415 | 0.06769720871697579 | 19.162519428033132 |
| **GATCGGGGTGGCTTTGTCTTCTTCTTTTGCCAATTCCACTAATTGTTTGGG** | Del_k12_5-16 | -0.5794217188195762 | 0.03570279629699002 | 27.506911929466906 |
| **GATCGGGTGGCTTTGTCTTCTTCTTTTGCCAATTCCACTAATTGTTTGGG** | Del_k13_5-17 | -0.02610220677345776 | 0.033320514226172056 | 47.83496359264642 |
| **GATCGGTCAGATCTAACTTGGGGTGGCTTTGTCTTCTTCTTTTGCCAATTCCACTAATTGTTTGGG** | Ins_k3_4_GGT | -1.3041871670553304 | 0.06927962009569057 | 13.325398392717855 |
| **GATCGGTGGCTTTGTCTTCTTCTTTTGCCAATTCCACTAATTGTTTGGG** | Del_k14_5-18 | 0.26315223353695794 | 0.027839419468172994 | 63.88031588678191 |
| **GATCGTACAGATCTAACTTGGGGTGGCTTTGTCTTCTTCTTTTGCCAATTCCACTAATTGTTTGGG** | Ins_k3_4_GTA | -0.6448673569271198 | 0.06048481307762552 | 25.764348053088895 |
| **GATCGTCAGATCTAACTTGGGGTGGCTTTGTCTTCTTCTTTTGCCAATTCCACTAATTGTTTGGG** | Ins_k2_4_GT | 0.4086902092317437 | 0.03513455385202513 | 73.88791213688582 |
| **GATCGTCCAGATCTAACTTGGGGTGGCTTTGTCTTCTTCTTTTGCCAATTCCACTAATTGTTTGGG** | Ins_k3_2_TCG;Ins_k3_3_CGT;Ins | 0.7657475412627619 | 0.028382178590866784 | 100 |
| **GATCGTCTTCTTCTTTTGCCAATTCCACTAATTGTTTGGG** | Del_k23_5-27 | -0.571780565716173 | 0.030638832848507352 | 27.717901530252835 |
| **GATCGTGCAGATCTAACTTGGGGTGGCTTTGTCTTCTTCTTTTGCCAATTCCACTAATTGTTTGGG** | Ins_k3_4_GTG | 0.16718277091567413 | 0.03513433811735623 | 58.03474052003408 |
| **GATCGTGGCTTTGTCTTCTTCTTTTGCCAATTCCACTAATTGTTTGGG** | Del_k15_5-19 | 0.05460051568514801 | 0.02723376283853949 | 51.85542461349821 |
| **GATCGTTCAGATCTAACTTGGGGTGGCTTTGTCTTCTTCTTTTGCCAATTCCACTAATTGTTTGGG** | Ins_k3_4_GTT | -1.1923062671528002 | 0.06821055840579823 | 14.902854438356101 |

| | | | | |
|---|---|---|---|---|
| GATCGTTTGGG | Del_k52_5-56 | -6.473877782513346 | 0.9265509299014565 | 0.07577258344689132 |
| GATCTAACAGATCTAACTTGGGGTGGCTTTGTCTTCTTCTTTTGCCAATTCCACTAATTGTTTGGG | Ins_k3_4_TAA | -2.3473682687949817 | 0.1040886251812377 | 4.69496550758273 |
| GATCTAACTTGGGGTGGCTTTGTCTTCTTCTTTTGCCAATTCCACTAATTGTTTGGG | Del_k6_1-6;Del_k6_2-7;Del_k6_3 | -2.374784434954419 | 0.1050063406456319 | 4.567996014386778 |
| GATCTAATTGTTTGGG | Del_k47_4-50;Del_k47_5-51 | -6.002423955228909 | 0.9031501358460952 | 0.12141207746401894 |
| GATCTACAGATCTAACTTGGGGTGGCTTTGTCTTCTTCTTTTGCCAATTCCACTAATTGTTTGGG | Ins_k2_4_TA | -0.05328202837782381 | 0.044339776528739355 | 46.55232768385997 |
| GATCTACCAGATCTAACTTGGGGTGGCTTTGTCTTCTTCTTTTGCCAATTCCACTAATTGTTTGGG | Ins_k3_3_CTA;Ins_k3_4_TAC | -0.12878483983535943 | 0.04256581212433774 | 43.166908508319274 |
| GATCTAGATCTAACTTGGGGTGGCTTTGTCTTCTTCTTTTGCCAATTCCACTAATTGTTTGGG | C5T | -1.952426444518042 | 0.055978613296790325 | 6.968727153841046 |
| GATCTAGCAGATCTAACTTGGGGTGGCTTTGTCTTCTTCTTTTGCCAATTCCACTAATTGTTTGGG | Ins_k3_4_TAG | -1.9721504090834927 | 0.07658390359769242 | 6.832622896578304 |
| GATCTATCAGATCTAACTTGGGGTGGCTTTGTCTTCTTCTTTTGCCAATTCCACTAATTGTTTGGG | Ins_k3_4_TAT | -2.9773032829497263 | 0.2051178271475808 | 2.500662637743725 |
| GATCTCACAGATCTAACTTGGGGTGGCTTTGTCTTCTTCTTTTGCCAATTCCACTAATTGTTTGGG | Ins_k3_4_TCA | -0.9531559569014034 | 0.05649249578517839 | 18.929150302061988 |
| GATCTCAGATCTAACTTGGGGTGGCTTTGTCTTCTTCTTTTGCCAATTCCACTAATTGTTTGGG | Ins_k1_4_T | -1.2382249246989503 | 0.07276765269854252 | 14.23400912719168 |
| GATCTCCACTAATTGTTTGGG | Del_k42_5-46 | -3.013142894242983 | 0.7243502668903578 | 2.41262686784838 |
| GATCTCCAGATCTAACTTGGGGTGGCTTTGTCTTCTTCTTTTGCCAATTCCACTAATTGTTTGGG | Ins_k2_2_TC;Ins_k2_3_CT;Ins_k2 | 0.15843789596224045 | 0.040157916315683974 | 57.52944655759073 |
| GATCTCCCAGATCTAACTTGGGGTGGCTTTGTCTTCTTCTTTTGCCAATTCCACTAATTGTTTGGG | Ins_k3_3_CTC;Ins_k3_4_TCC | -0.17318997264926156 | 0.04217531501718502 | 41.29201179084432 |
| GATCTCGCAGATCTAACTTGGGGTGGCTTTGTCTTCTTCTTTTGCCAATTCCACTAATTGTTTGGG | Ins_k3_4_TCG | 0.2826887826096317 | 0.04110636417088741 | 65.14058740663842 |
| GATCTCTAACTTGGGGTGGCTTTGTCTTCTTCTTTTGCCAATTCCACTAATTGTTTGGG | Del_k4_5-8 | -2.5769428837739214 | 0.1307323861372049 | 3.7318950130074655 |
| GATCTCTCAGATCTAACTTGGGGTGGCTTTGTCTTCTTCTTTTGCCAATTCCACTAATTGTTTGGG | Ins_k3_4_TCT | -1.9516656334243487 | 0.08936039515338508 | 6.974031056146627 |
| GATCTCTTCTTCTTTTGCCAATTCCACTAATTGTTTGGG | Del_k24_5-28 | -2.3703575920573074 | 0.06208946861518599 | 4.588262640585245 |
| GATCTCTTCTTTTGCCAATTCCACTAATTGTTTGGG | Del_k27_5-31 | -3.150523395953346 | 0.07464782487461723 | 2.1029384715922688 |
| GATCTCTTTTGCCAATTCCACTAATTGTTTGGG | Del_k30_5-34 | -4.470449900574639 | 0.14281326676335063 | 0.561810392574676 |
| GATCTGACAGATCTAACTTGGGGTGGCTTTGTCTTCTTCTTTTGCCAATTCCACTAATTGTTTGGG | Ins_k3_4_TGA | -0.5085439418600417 | 0.04944920451088304 | 29.527295092751746 |
| GATCTGCAGATCTAACTTGGGGTGGCTTTGTCTTCTTCTTTTGCCAATTCCACTAATTGTTTGGG | Ins_k2_4_TG | -0.03767434765122901 | 0.039590767243362544 | 47.28460123222899 |
| GATCTGCCAATTCCACTAATTGTTTGGG | Del_k35_5-39 | -4.282447537022928 | 0.10728112701602203 | 0.6780132135482355 |
| GATCTGCCAGATCTAACTTGGGGTGGCTTTGTCTTCTTCTTTTGCCAATTCCACTAATTGTTTGGG | Ins_k3_3_CTG;Ins_k3_4_TGC | 0.29712955608911307 | 0.036523137381203594 | 66.08809276312383 |
| GATCTGGCAGATCTAACTTGGGGTGGCTTTGTCTTCTTCTTTTGCCAATTCCACTAATTGTTTGGG | Ins_k3_4_TGG | 0.013829549898175975 | 0.03767644498485113 | 49.78374796573942 |
| GATCTGGCTTTGTCTTCTTCTTTTGCCAATTCCACTAATTGTTTGGG | Del_k16_5-20 | -0.8375707321793291 | 0.03920020090161313 | 21.24854245406691 |
| GATCTGGGGTGGCTTTGTCTTCTTCTTTTGCCAATTCCACTAATTGTTTGGG | Del_k11_5-15 | -1.3293638223587885 | 0.05636242410785846 | 12.994097455149069 |
| GATCTGTCAGATCTAACTTGGGGTGGCTTTGTCTTCTTCTTTTGCCAATTCCACTAATTGTTTGGG | Ins_k3_4_TGT | -2.016029129372045 | 0.08665002599392088 | 6.539298556953046 |
| GATCTGTCTTCTTCTTTTGCCAATTCCACTAATTGTTTGGG | Del_k22_5-26 | -2.2112433345098164 | 0.06033219321585582 | 5.379609028716147 |
| GATCTGTTTGGG | Del_k51_5-55 | -6.769411775999239 | 0.5160713202950856 | 0.05638496459582011 |
| GATCTTACAGATCTAACTTGGGGTGGCTTTGTCTTCTTCTTTTGCCAATTCCACTAATTGTTTGGG | Ins_k3_4_TTA | -1.9706879690042483 | 0.09634011218979031 | 6.842622508282884 |
| GATCTTCAGATCTAACTTGGGGTGGCTTTGTCTTCTTCTTTTGCCAATTCCACTAATTGTTTGGG | Ins_k2_4_TT | -0.10843304202395532 | 0.04141001323467964 | 44.054433432457735 |
| GATCTTCCACTAATTGTTTGGG | Del_k41_5-45 | -5.797711136604872 | 0.3518204038096337 | 0.14899357395218463 |
| GATCTTCCAGATCTAACTTGGGGTGGCTTTGTCTTCTTCTTTTGCCAATTCCACTAATTGTTTGGG | Ins_k3_2_TCT;Ins_k3_3_CTT;Ins_ | 0.18250591255810422 | 0.03685035990962242 | 58.93086324294366 |
| GATCTTCTTCTTTTGCCAATTCCACTAATTGTTTGGG | Del_k26_3-28;Del_k26_4-29;Del_ | -2.0808335982438306 | 0.055180945571837905 | 6.128962196769725 |
| GATCTTCTTTTGCCAATTCCACTAATTGTTTGGG | Del_k29_3-31;Del_k29_4-32;Del_ | -3.01739671800891 | 0.07773697406514318 | 2.402385775682957 |
| GATCTTGCAGATCTAACTTGGGGTGGCTTTGTCTTCTTCTTTTGCCAATTCCACTAATTGTTTGGG | Ins_k3_4_TTG | -0.3319952315353299 | 0.04417995408565047 | 35.22879553901954 |
| GATCTTGCCAATTCCACTAATTGTTTGGG | Del_k34_5-38 | -5.201291527944716 | 0.20529092211546268 | 0.2705137113301368 |
| GATCTTGGGGTGGCTTTGTCTTCTTCTTTTGCCAATTCCACTAATTGTTTGGG | Del_k10_4-13;Del_k10_5-14 | -1.606069530608712 | 0.07813797781266404 | 9.853133280968528 |
| GATCTTGTCTTCTTCTTTTGCCAATTCCACTAATTGTTTGGG | Del_k21_5-25 | -2.252033382816095 | 0.0645289020759601 | 5.164589654018328 |
| GATCTTGTTTGGG | Del_k50_5-54 | -6.7894225565240935 | 0.45946350770446237 | 0.055267871671395415 |
| GATCTTTCAGATCTAACTTGGGGTGGCTTTGTCTTCTTCTTTTGCCAATTCCACTAATTGTTTGGG | Ins_k3_4_TTT | -2.894421705457365 | 0.15264101632774668 | 2.716752761257979 |
| GATCTTTGCCAATTCCACTAATTGTTTGGG | Del_k33_5-37 | -4.497721847138941 | 0.1672481180444162 | 0.5466957689644163 |
| GATCTTTGTCTTCTTCTTTTGCCAATTCCACTAATTGTTTGGG | Del_k20_4-23;Del_k20_5-24 | -2.4908824514148193 | 0.07403540400190245 | 4.0672885901636775 |
| GATCTTTTGCCAATTCCACTAATTGTTTGGG | Del_k32_3-34;Del_k32_4-35;Del_ | -4.468449082291793 | 0.1259269840289254 | 0.5629355983704212 |
| GATGAAACCAGATCTAACTTGGGGTGGCTTTGTCTTCTTCTTTTGCCAATTCCACTAATTGTTTGGG | Ins_k4_3_GAAA | 0.7337217337621937 | 0.03785261032111623 | 100 |
| GATGAACCAGATCTAACTTGGGGTGGCTTTGTCTTCTTCTTTTGCCAATTCCACTAATTGTTTGGG | Ins_k3_3_GAA | 0.3766615145368615 | 0.03362743135262708 | 71.55887585109411 |
| GATGAAGCCAGATCTAACTTGGGGTGGCTTTGTCTTCTTCTTTTGCCAATTCCACTAATTGTTTGGG | Ins_k4_3_GAAG | 0.48736207828073286 | 0.034860335946801824 | 79.93558422021995 |
| GATGACCAGATCTAACTTGGGGTGGCTTTGTCTTCTTCTTTTGCCAATTCCACTAATTGTTTGGG | Ins_k2_3_GA | 0.019579683050270136 | 0.042160829564696585 | 50.07083575082516 |
| GATGACCCAGATCTAACTTGGGGTGGCTTTGTCTTCTTCTTTTGCCAATTCCACTAATTGTTTGGG | Ins_k3_3_GAC | -0.18304033596408908 | 0.03851920897073905 | 40.887267186088216 |
| GATGAGCCAGATCTAACTTGGGGTGGCTTTGTCTTCTTCTTTTGCCAATTCCACTAATTGTTTGGG | Ins_k3_3_GAG | -0.3547519674270262 | 0.04438681767748996 | 34.43615629081774 |
| GATGATCCAGATCTAACTTGGGGTGGCTTTGTCTTCTTCTTTTGCCAATTCCACTAATTGTTTGGG | Ins_k3_1_ATG;Ins_k3_2_TGA;Ins | 0.412077000140997 | 0.03143796574081622 | 74.13857928502067 |
| GATGATCTAACTTGGGGTGGCTTTGTCTTCTTCTTTTGCCAATTCCACTAATTGTTTGGG | Del_k3_4-6 | -1.7568255905226238 | 0.07682536420870698 | 8.474260936048061 |
| GATGCACCAGATCTAACTTGGGGTGGCTTTGTCTTCTTCTTTTGCCAATTCCACTAATTGTTTGGG | Ins_k3_3_GCA | 0.32006949365686577 | 0.03422113978256547 | 67.62167234290878 |
| GATGCAGATCTAACTTGGGGTGGCTTTGTCTTCTTCTTTTGCCAATTCCACTAATTGTTTGGG | C4G | -0.7565026151836056 | 0.039950673011642235 | 23.042870589313413 |
| GATGCCAATTCCACTAATTGTTTGGG | Del_k37_3-39;Del_k37_4-40 | -5.405941787320322 | 0.23784098234362172 | 0.2204503559517191 |
| GATGCCAGATCTAACTTGGGGTGGCTTTGTCTTCTTCTTTTGCCAATTCCACTAATTGTTTGGG | Ins_k1_3_G | 0.11363508101570186 | 0.038812390224114666 | 55.00885189435498 |
| GATGCCCAGATCTAACTTGGGGTGGCTTTGTCTTCTTCTTTTGCCAATTCCACTAATTGTTTGGG | Ins_k2_3_GC | -0.7457853834724595 | 0.055980601701932434 | 23.291154454097253 |
| GATGCCCCAGATCTAACTTGGGGTGGCTTTGTCTTCTTCTTTTGCCAATTCCACTAATTGTTTGGG | Ins_k3_3_GCC | -2.947116307676413 | 0.1436761563234928 | 2.577300999461872 |
| GATGCCCCCAGATCTAACTTGGGGTGGCTTTGTCTTCTTCTTTTGCCAATTCCACTAATTGTTTGGG | Ins_k4_3_GCCC | -1.531435936497847 | 0.07891762101173269 | 10.616645485934196 |
| GATGCGCCAGATCTAACTTGGGGTGGCTTTGTCTTCTTCTTTTGCCAATTCCACTAATTGTTTGGG | Ins_k3_3_GCG | 0.3264429471139426 | 0.034291609301234816 | 68.05403227365355 |
| GATGCTCCAGATCTAACTTGGGGTGGCTTTGTCTTCTTCTTTTGCCAATTCCACTAATTGTTTGGG | Ins_k3_2_TGC;Ins_k3_3_GCT | 0.47082964975628916 | 0.033732796717125646 | 78.62491897982228 |
| GATGCTTTGTCTTCTTCTTTTGCCAATTCCACTAATTGTTTGGG | Del_k19_4-22 | -2.4434830377086536 | 0.06907966708026582 | 4.264717735212772 |
| GATGG | Del_k58_4-61 | -7.1843224563936525 | 0.7565073613642793 | 0.03723658915766914 |
| GATGGACCAGATCTAACTTGGGGTGGCTTTGTCTTCTTCTTTTGCCAATTCCACTAATTGTTTGGG | Ins_k3_3_GGA | 0.29684233989737885 | 0.04887885400344766 | 66.06911391844764 |
| GATGGCCAGATCTAACTTGGGGTGGCTTTGTCTTCTTCTTTTGCCAATTCCACTAATTGTTTGGG | Ins_k2_3_GG | -0.24530790373767808 | 0.05142648026474541 | 38.41896166444531 |
| GATGGCCCAGATCTAACTTGGGGTGGCTTTGTCTTCTTCTTTTGCCAATTCCACTAATTGTTTGGG | Ins_k3_3_GGC | 0.0733914355366534 | 0.0421047156224768 | 52.83904839625881 |
| GATGGCTTTGTCTTCTTCTTTTGCCAATTCCACTAATTGTTTGGG | Del_k18_3-20;Del_k18_4-21 | -2.02986133121048 | 0.0631203956477261 | 6.449468366356768 |
| GATGGG | Del_k57_3-59;Del_k57_4-60 | -6.43865658782091 | 0.41297841514760947 | 0.07848894022128035 |
| GATGGGCCAGATCTAACTTGGGGTGGCTTTGTCTTCTTCTTTTGCCAATTCCACTAATTGTTTGGG | Ins_k3_3_GGG | -2.0796764512577464 | 0.09909071744356886 | 6.136058411793642 |
| GATGGGGTGGCTTTGTCTTCTTCTTTTGCCAATTCCACTAATTGTTTGGG | Del_k13_3-15;Del_k13_4-16 | -3.141485576384553 | 0.11428506957820289 | 2.1220305956921726 |
| GATGGGTGGCTTTGTCTTCTTCTTTTGCCAATTCCACTAATTGTTTGGG | Del_k14_4-17 | -2.4102819956612125 | 0.07975467584278466 | 4.408687557425205 |
| GATGGTCCAGATCTAACTTGGGGTGGCTTTGTCTTCTTCTTTTGCCAATTCCACTAATTGTTTGGG | Ins_k3_2_TGG;Ins_k3_3_GGT | 0.12853178030498436 | 0.03771098031771098 | 55.83443619261376 |
| GATGGTGGCTTTGTCTTCTTCTTTTGCCAATTCCACTAATTGTTTGGG | Del_k15_4-18 | -0.8246991905105373 | 0.03393666610830307 | 21.523811723136802 |
| GATGTACCAGATCTAACTTGGGGTGGCTTTGTCTTCTTCTTTTGCCAATTCCACTAATTGTTTGGG | Ins_k3_3_GTA | 0.1959396823550139 | 0.03898247356127424 | 59.7278682989054 |

| | | | | |
|---|---|---|---|---|
| GATGTCCAGATCTAACTTGGGGTGGCTTTGTCTTCTTCTTTTGCCAATTCCACTAATTGTTTGGG | Ins_k2_2_TG;Ins_k2_3_GT | -0.27890462989318515 | 0.04754726880365594 | 37.14965204520134 |
| GATGTCCCAGATCTAACTTGGGGTGGCTTTGTCTTCTTCTTTTGCCAATTCCACTAATTGTTTGGG | Ins_k3_3_GTC | -0.4216516707277006 | 0.044335230855252566 | 32.20775828551432 |
| GATGTCTTCTTCTTTTGCCAATTCCACTAATTGTTTGGG | Del_k24_3-26;Del_k24_4-27 | -2.435207692283405 | 0.0634345114136954 | 4.300156178051641 |
| GATGTGCCAGATCTAACTTGGGGTGGCTTTGTCTTCTTCTTTTGCCAATTCCACTAATTGTTTGGG | Ins_k3_3_GTG | 0.12934232888241692 | 0.0385592832552113 | 55.87971106170304 |
| GATGTGGCTTTGTCTTCTTCTTTTGCCAATTCCACTAATTGTTTGGG | Del_k16_4-19 | -1.3802146959712616 | 0.0462978538992059 | 12.349855206425854 |
| GATGTTCCAGATCTAACTTGGGGTGGCTTTGTCTTCTTCTTTTGCCAATTCCACTAATTGTTTGGG | Ins_k3_2_TGT;Ins_k3_3_GTT | 0.6886171121918736 | 0.030441417739249996 | 97.7561533730715 |
| GATGTTTGGG | Del_k53_3-55;Del_k53_4-56 | -6.540593002125693 | 0.5502948506057409 | 0.07088233945853055 |
| GATTAACCAGATCTAACTTGGGGTGGCTTTGTCTTCTTCTTTTGCCAATTCCACTAATTGTTTGGG | Ins_k3_3_TAA | -1.4428581440870412 | 0.07629755690480199 | 11.599951194275604 |
| GATTAACTTGGGGTGGCTTTGTCTTCTTCTTTTGCCAATTCCACTAATTGTTTGGG | Del_k7_4-10 | -3.0811573520519406 | 0.14313677134312633 | 2.2539893321384055 |
| GATTAATTGTTTGGG | Del_k48_4-51 | -7.3013112262754865 | 1.4713117540552787 | 0.033125490359653925 |
| GATTACCAGATCTAACTTGGGGTGGCTTTGTCTTCTTCTTTTGCCAATTCCACTAATTGTTTGGG | Ins_k2_3_TA | -0.4415115985997258 | 0.05643770446432024 | 31.574424328717285 |
| GATTACCCAGATCTAACTTGGGGTGGCTTTGTCTTCTTCTTTTGCCAATTCCACTAATTGTTTGGG | Ins_k3_3_TAC | -0.720958486500939 | 0.05790350686911337 | 23.876639359924383 |
| GATTAGCCAGATCTAACTTGGGGTGGCTTTGTCTTCTTCTTTTGCCAATTCCACTAATTGTTTGGG | Ins_k3_3_TAG | -3.645152121557619 | 0.20468439828574297 | 1.2823661334553769 |
| GATTATCCAGATCTAACTTGGGGTGGCTTTGTCTTCTTCTTTTGCCAATTCCACTAATTGTTTGGG | Ins_k3_1_ATT;Ins_k3_2_TTA;Ins_ | -0.030073889902966666 | 0.04239491426812312 | 47.645355056501515 |
| GATTCACCAGATCTAACTTGGGGTGGCTTTGTCTTCTTCTTTTGCCAATTCCACTAATTGTTTGGG | Ins_k3_3_TCA | 0.25480777453227543 | 0.03542920815495274 | 63.349487029607786 |
| GATTCAGATCTAACTTGGGGTGGCTTTGTCTTCTTCTTTTGCCAATTCCACTAATTGTTTGGG | C4T | -1.4608819734422531 | 0.048035525493561444 | 11.392748555188826 |
| GATTCCACTAATTGTTTGGG | Del_k43_2-44;Del_k43_3-45;Del_ | -3.473875846059334 | 0.06889393341381886 | 1.5219359697440145 |
| GATTCCAGATCTAACTTGGGGTGGCTTTGTCTTCTTCTTTTGCCAATTCCACTAATTGTTTGGG | Ins_k1_2_T;Ins_k1_3_T | 0.4301789365998041 | 0.04030913615705386 | 75.49285163189658 |
| GATTCCCAGATCTAACTTGGGGTGGCTTTGTCTTCTTCTTTTGCCAATTCCACTAATTGTTTGGG | Ins_k2_3_TC | -0.5325611971453938 | 0.05672077715017901 | 28.8265788317234 |
| GATTCCCCAGATCTAACTTGGGGTGGCTTTGTCTTCTTCTTTTGCCAATTCCACTAATTGTTTGGG | Ins_k3_3_TCC | -1.3573818396243647 | 0.06788217474379547 | 12.635081550797942 |
| GATTCGCCAGATCTAACTTGGGGTGGCTTTGTCTTCTTCTTTTGCCAATTCCACTAATTGTTTGGG | Ins_k3_3_TCG | 0.4780583473784472 | 0.032165085082829344 | 79.19533393945218 |
| GATTCTAACTTGGGGTGGCTTTGTCTTCTTCTTTTGCCAATTCCACTAATTGTTTGGG | Del_k5_4-8 | -2.457194817747256 | 0.11495082162016156 | 4.206639948563602 |
| GATTCTCCAGATCTAACTTGGGGTGGCTTTGTCTTCTTCTTTTGCCAATTCCACTAATTGTTTGGG | Ins_k3_2_TTC;Ins_k3_3_TCT | 0.4011077767526148 | 0.0347056158076231 | 73.3297807031044 |
| GATTCTTCTTCTTTTGCCAATTCCACTAATTGTTTGGG | Del_k25_4-28 | -2.446251694755998 | 0.058374722010223014 | 4.252926524841175 |
| GATTCTTCTTTTGCCAATTCCACTAATTGTTTGGG | Del_k28_3-30;Del_k28_4-31 | -3.3086639752830194 | 0.0784929975981482 | 1.7953411498657184 |
| GATTCTTTTGCCAATTCCACTAATTGTTTGGG | Del_k31_3-33;Del_k31_4-34 | -4.645050217116792 | 0.15079969671731286 | 0.4718042130476121 |
| GATTGACCAGATCTAACTTGGGGTGGCTTTGTCTTCTTCTTTTGCCAATTCCACTAATTGTTTGGG | Ins_k3_3_TGA | -0.10392603074375895 | 0.04284738525660432 | 44.253435376019766 |
| GATTGCCAATTCCACTAATTGTTTGGG | Del_k36_3-38;Del_k36_4-39 | -5.4142785056393485 | 0.16729852831813602 | 0.21862016293264394 |
| GATTGCCAGATCTAACTTGGGGTGGCTTTGTCTTCTTCTTTTGCCAATTCCACTAATTGTTTGGG | Ins_k2_3_TG | -0.3365425781844578 | 0.04882360592414814 | 35.06896167908118 |
| GATTGCCCAGATCTAACTTGGGGTGGCTTTGTCTTCTTCTTTTGCCAATTCCACTAATTGTTTGGG | Ins_k3_3_TGC | -0.26623626786388294 | 0.04596937446587768 | 37.6232709402474 |
| GATTGCCCCAGATCTAACTTGGGGTGGCTTTGTCTTCTTCTTTTGCCAATTCCACTAATTGTTTGGG | Ins_k4_3_TGCC | -1.8820437944396602 | 0.11700880012715976 | 7.476877365816588 |
| GATTGGCCAGATCTAACTTGGGGTGGCTTTGTCTTCTTCTTTTGCCAATTCCACTAATTGTTTGGG | Ins_k3_3_TGG | -0.3086198019457802 | 0.04610932542292654 | 36.06198389833928 |
| GATTGGCTTTGTCTTCTTCTTTTGCCAATTCCACTAATTGTTTGGG | Del_k17_4-20 | -3.349365590115105 | 0.10444859773813621 | 1.7237349940631088 |
| GATTGGG | Del_k56_3-58;Del_k56_4-59 | -6.843310536423553 | 0.9134392713296036 | 0.052368422066845895 |
| GATTGGGGTGGCTTTGTCTTCTTCTTTTGCCAATTCCACTAATTGTTTGGG | Del_k12_3-14;Del_k12_4-15 | -3.0773820657457236 | 0.14045212237328022 | 2.262514870245968 |
| GATTGTCCAGATCTAACTTGGGGTGGCTTTGTCTTCTTCTTTTGCCAATTCCACTAATTGTTTGGG | Ins_k3_2_TTG;Ins_k3_3_TGT | 0.11860431045983755 | 0.04539277498672072 | 55.282883800690875 |
| GATTGTCTTCTTCTTTTGCCAATTCCACTAATTGTTTGGG | Del_k23_3-25;Del_k23_4-26 | -2.129325728353769 | 0.05982922799099192 | 5.838846769982044 |
| GATTGTTTGGG | Del_k52_2-53;Del_k52_3-54;Del_ | -6.012516074023524 | 0.23684291434646404 | 0.12019293457015258 |
| GATTTACCAGATCTAACTTGGGGTGGCTTTGTCTTCTTCTTTTGCCAATTCCACTAATTGTTTGGG | Ins_k3_3_TTA | -0.34238387864902925 | 0.04690592958070543 | 34.864710464059144 |
| GATTTCCACTAATTGTTTGGG | Del_k42_4-45 | -6.335677703462069 | 0.5302437363309084 | 0.08700247881025104 |
| GATTTCCAGATCTAACTTGGGGTGGCTTTGTCTTCTTCTTTTGCCAATTCCACTAATTGTTTGGG | Ins_k2_2_TT;Ins_k2_3_TT | 0.023010490890925106 | 0.04452885986435605 | 50.242914181935994 |
| GATTTCCCAGATCTAACTTGGGGTGGCTTTGTCTTCTTCTTTTGCCAATTCCACTAATTGTTTGGG | Ins_k3_3_TTC | -0.5741811055735068 | 0.05684418101156998 | 27.65144340247435 |
| GATTTCTTCTTTTGCCAATTCCACTAATTGTTTGGG | Del_k27_4-30 | -3.22682787147725 | 0.10470756195989835 | 1.9484441130020689 |
| GATTTCTTTTGCCAATTCCACTAATTGTTTGGG | Del_k30_4-33 | -4.663813256525646 | 0.16328718946303986 | 0.46303426473588266 |
| GATTTGCCAATTCCACTAATTGTTTGGG | Del_k35_3-37;Del_k35_4-38 | -5.04610472576032 | 0.15059446772988014 | 0.3159264988710323 |
| GATTTGCCAGATCTAACTTGGGGTGGCTTTGTCTTCTTCTTTTGCCAATTCCACTAATTGTTTGGG | Ins_k3_3_TTG | -0.06537528615901633 | 0.04148850375249595 | 45.9927487705941 |
| GATTTGGG | Del_k55_3-57;Del_k55_4-58 | -5.953073856140313 | 0.44815663356116603 | 0.1275540833304583 |
| GATTTGGGGTGGCTTTGTCTTCTTCTTTTGCCAATTCCACTAATTGTTTGGG | Del_k11_4-14 | -2.5299061540354026 | 0.09529666839624461 | 3.911624969031526 |
| GATTTGTCTTCTTCTTTTGCCAATTCCACTAATTGTTTGGG | Del_k22_3-24;Del_k22_4-25 | -2.2650600428868835 | 0.068977212847872 | 5.097748603221133 |
| GATTTGTTTGGG | Del_k51_4-54 | -6.0333229134307915 | 0.33466316915116995 | 0.11771793720892969 |
| GATTTTCCAGATCTAACTTGGGGTGGCTTTGTCTTCTTCTTTTGCCAATTCCACTAATTGTTTGGG | Ins_k3_2_TTT;Ins_k3_3_TTT | 0.06143710566778138 | 0.04650564897547289 | 52.21115349066299 |
| GATTTTCCCAGATCTAACTTGGGGTGGCTTTGTCTTCTTCTTTTGCCAATTCCACTAATTGTTTGGG | Ins_k4_3_TTTC | -0.8021103260493966 | 0.0648518321283363 | 22.015543106696164 |
| GATTTTGCCAATTCCACTAATTGTTTGGG | Del_k34_3-36;Del_k34_4-37 | -4.50498855102556 | 0.14605996205293528 | 0.5427374919268987 |
| GATTTTGTCTTCTTCTTTTGCCAATTCCACTAATTGTTTGGG | Del_k21_4-24 | -2.409976661159094 | 0.09031021519949352 | 4.4100338873755796 |
| GATTTTTGCCAATTCCACTAATTGTTTGGG | Del_k33_4-36 | -4.932725191431037 | 0.16492784243524813 | 0.3538556714930189 |
| GCAAATCCAGATCTAACTTGGGGTGGCTTTGTCTTCTTCTTTTGCCAATTCCACTAATTGTTTGGG | Ins_k3_1_CAA | -0.07179038598110088 | 0.044690974295057986 | 45.69864505709464 |
| GCAATCCAGATCTAACTTGGGGTGGCTTTGTCTTCTTCTTTTGCCAATTCCACTAATTGTTTGGG | Ins_k2_1_CA | -0.17714137201753466 | 0.04367307868023781 | 41.1291724949597 |
| GCAATTCCACTAATTGTTTGGG | Del_k41_2-42 | -4.63576912328693 | 0.1222804570364827 | 0.4762034555306558 |
| GCACATCCAGATCTAACTTGGGGTGGCTTTGTCTTCTTCTTTTGCCAATTCCACTAATTGTTTGGG | Ins_k3_1_CAC | 0.4084878790192503 | 0.03999949924887424 | 73.87296389221422 |
| GCACTAATTGTTTGGG | Del_k47_2-48 | -7.141462200668531 | 1.1051949359642763 | 0.03886725464083141 |
| GCAGATCCAGATCTAACTTGGGGTGGCTTTGTCTTCTTCTTTTGCCAATTCCACTAATTGTTTGGG | Ins_k3_1_CAG | 0.27538751075873535 | 0.036522928432564244 | 64.66671032728621 |
| GCAGATCTAACTTGGGGTGGCTTTGTCTTCTTCTTTTGCCAATTCCACTAATTGTTTGGG | Del_k3_2-4 | -3.1906076486813384 | 0.14656293991309668 | 2.020310850821593 |
| GCAGCATCCAGATCTAACTTGGGGTGGCTTTGTCTTCTTCTTTTGCCAATTCCACTAATTGTTTGGG | Ins_k4_1_CAGC | 0.6564555833609231 | 0.03407552519597687 | 94.66218607330296 |
| GCATATCCAGATCTAACTTGGGGTGGCTTTGTCTTCTTCTTTTGCCAATTCCACTAATTGTTTGGG | Ins_k3_1_CAT | 0.044216611073418344 | 0.03993879716083367 | 51.31974884702436 |
| GCATCCAGATCTAACTTGGGGTGGCTTTGTCTTCTTCTTTTGCCAATTCCACTAATTGTTTGGG | Ins_k1_1_C | 0.2775953336552317 | 0.03594906158290718 | 64.80964069539826 |
| GCCAATCCAGATCTAACTTGGGGTGGCTTTGTCTTCTTCTTTTGCCAATTCCACTAATTGTTTGGG | Ins_k3_1_CCA | -0.5399195739421382 | 0.05312780276433316 | 28.615240510007656 |
| GCCAATTCCACTAATTGTTTGGG | Del_k40_1-40;Del_k40_2-41 | -1.3381984849784625 | 0.029164677876857663 | 12.879804600880156 |
| GCCACTAATTGTTTGGG | Del_k46_2-47 | -6.5051885548229835 | 0.5507710781659461 | 0.07343684307667896 |
| GCCAGATCTAACTTGGGGTGGCTTTGTCTTCTTCTTTTGCCAATTCCACTAATTGTTTGGG | Del_k2_2-3 | -1.9114099843135148 | 0.08748770105516908 | 7.260502566574897 |
| GCCATCCAGATCTAACTTGGGGTGGCTTTGTCTTCTTCTTTTGCCAATTCCACTAATTGTTTGGG | Ins_k2_1_CC | -2.9308973568407906 | 0.23805282777482945 | 2.6194429429174546 |
| GCCCATCCAGATCTAACTTGGGGTGGCTTTGTCTTCTTCTTTTGCCAATTCCACTAATTGTTTGGG | Ins_k3_1_CCC | 0.3505388516683725 | 0.04573598038239086 | 69.71377189895844 |
| GCCGATCCAGATCTAACTTGGGGTGGCTTTGTCTTCTTCTTTTGCCAATTCCACTAATTGTTTGGG | Ins_k3_1_CCG | 0.5920129508466667 | 0.034790404594279296 | 88.7543095642605 |
| GCCTATCCAGATCTAACTTGGGGTGGCTTTGTCTTCTTCTTTTGCCAATTCCACTAATTGTTTGGG | Ins_k3_1_CCT | -0.09269258321382856 | 0.042405939427719135 | 44.75335668312931 |
| GCCTTATCCAGATCTAACTTGGGGTGGCTTTGTCTTCTTCTTTTGCCAATTCCACTAATTGTTTGGG | Ins_k4_1_CCTT | 0.2976075918643768 | 0.040693170047077586 | 66.11969278815302 |
| GCGAATCCAGATCTAACTTGGGGTGGCTTTGTCTTCTTCTTTTGCCAATTCCACTAATTGTTTGGG | Ins_k3_1_CGA | 0.5463462734275149 | 0.031559281597545306 | 84.79234845369741 |

| | | | | |
|---|---|---|---|---|
| **GCGATCCAGATCTAACTTGGGGTGGCTTTGTCTTCTTCTTTTGCCAATTCCACTAATTGTTTGGG** | Ins_k2_1_CG | 0.42260299061161843 | 0.034343511043442444 | 74.92308285931459 |
| **GCGCATCCAGATCTAACTTGGGGTGGCTTTGTCTTCTTCTTTTGCCAATTCCACTAATTGTTTGGG** | Ins_k3_1_CGC | 0.6473366150179625 | 0.03095110381977945 | 93.80288850350082 |
| **GCGGATCCAGATCTAACTTGGGGTGGCTTTGTCTTCTTCTTTTGCCAATTCCACTAATTGTTTGGG** | Ins_k3_1_CGG | 0.6728097752627689 | 0.03443364498078933 | 96.22303807822166 |
| **GCGTATCCAGATCTAACTTGGGGTGGCTTTGTCTTCTTCTTTTGCCAATTCCACTAATTGTTTGGG** | Ins_k3_1_CGT | 0.6085075805958 | 0.03256128546122586 | 90.23041951542281 |
| **GCGTGATCCAGATCTAACTTGGGGTGGCTTTGTCTTCTTCTTTTGCCAATTCCACTAATTGTTTGGG** | Ins_k4_1_CGTG | 0.39328472827464245 | 0.03959901894781696 | 72.75835632834307 |
| **GCTAACTTGGGGTGGCTTTGTCTTCTTCTTTTGCCAATTCCACTAATTGTTTGGG** | Del_k8_2-9 | -2.8367940456530385 | 0.12461607880850147 | 2.877911878952655 |
| **GCTAATCCAGATCTAACTTGGGGTGGCTTTGTCTTCTTCTTTTGCCAATTCCACTAATTGTTTGGG** | Ins_k3_1_CTA | -0.38038617525819557 | 0.042901043703957496 | 33.56463085924714 |
| **GCTAATTGTTTGGG** | Del_k49_2-50 | -4.82997425860062 | 0.12073988002794306 | 0.3921482884742035 |
| **GCTATCCAGATCTAACTTGGGGTGGCTTTGTCTTCTTCTTTTGCCAATTCCACTAATTGTTTGGG** | Ins_k2_1_CT | -0.3732430880408315 | 0.051927862665529505 | 33.80524428314838 |
| **GCTCATCCAGATCTAACTTGGGGTGGCTTTGTCTTCTTCTTTTGCCAATTCCACTAATTGTTTGGG** | Ins_k3_1_CTC | 0.5552394496199609 | 0.03588694543152719 | 85.54978476350581 |
| **GCTCCAGATCTAACTTGGGGTGGCTTTGTCTTCTTCTTTTGCCAATTCCACTAATTGTTTGGG** | A2C | 0.007144651093307419 | 0.04195210909170718 | 49.45205853879523 |
| **GCTGATCCAGATCTAACTTGGGGTGGCTTTGTCTTCTTCTTTTGCCAATTCCACTAATTGTTTGGG** | Ins_k3_1_CTG | 0.6357000293917112 | 0.035959835199954535 | 92.71766952698732 |
| **GCTGTATCCAGATCTAACTTGGGGTGGCTTTGTCTTCTTCTTTTGCCAATTCCACTAATTGTTTGGG** | Ins_k4_1_CTGT | 0.7211387602197921 | 0.03857519399495953 | 100 |
| **GCTTATCCAGATCTAACTTGGGGTGGCTTTGTCTTCTTCTTTTGCCAATTCCACTAATTGTTTGGG** | Ins_k3_1_CTT | -0.04192782412393081 | 0.03921500048939404 | 47.08390442552418 |
| **GCTTCTTCTTTTGCCAATTCCACTAATTGTTTGGG** | Del_k28_2-29 | -3.1287488130646275 | 0.07231272107731805 | 2.1492312535998854 |
| **GCTTCTTTTGCCAATTCCACTAATTGTTTGGG** | Del_k31_2-32 | -5.151935243088632 | 0.22559325058156351 | 0.28420024299990837 |
| **GCTTGGGGTGGCTTTGTCTTCTTCTTTTGCCAATTCCACTAATTGTTTGGG** | Del_k12_2-13 | -2.848181461338493 | 0.14336133584898184 | 2.8453257878910505 |
| **GCTTTGTCTTCTTCTTTTGCCAATTCCACTAATTGTTTGGG** | Del_k22_1-22;Del_k22_2-23 | -2.1643706569188517 | 0.06961810675194394 | 5.637768763241641 |
| **GCTTTTGCCAATTCCACTAATTGTTTGGG** | Del_k34_2-35 | -5.490116219427303 | 0.21353625749735589 | 0.2026535953024854 |
| **GGAAATCCAGATCTAACTTGGGGTGGCTTTGTCTTCTTCTTTTGCCAATTCCACTAATTGTTTGGG** | Ins_k3_1_GAA | -0.5184303587623039 | 0.051598765568934835 | 29.236814216981553 |
| **GGAATCCAGATCTAACTTGGGGTGGCTTTGTCTTCTTCTTTTGCCAATTCCACTAATTGTTTGGG** | Ins_k2_1_GA | -0.11097674503233207 | 0.050372239586562145 | 43.94251444228621 |
| **GGACATCCAGATCTAACTTGGGGTGGCTTTGTCTTCTTCTTTTGCCAATTCCACTAATTGTTTGGG** | Ins_k3_1_GAC | -0.3565842768448779 | 0.04500774343839329 | 34.37311636928972 |
| **GGAGATCCAGATCTAACTTGGGGTGGCTTTGTCTTCTTCTTTTGCCAATTCCACTAATTGTTTGGG** | Ins_k3_1_GAG | 0.34244859837972175 | 0.04109838295343207 | 69.15204514629357 |
| **GGATATCCAGATCTAACTTGGGGTGGCTTTGTCTTCTTCTTTTGCCAATTCCACTAATTGTTTGGG** | Ins_k3_1_GAT | -0.21747152491944982 | 0.04220437261940725 | 39.50343024890676 |
| **GGATCCAGATCTAACTTGGGGTGGCTTTGTCTTCTTCTTTTGCCAATTCCACTAATTGTTTGGG** | Ins_k1_1_G | -0.1223998423050024 | 0.024741932336212552 | 43.44341090672889 |
| **GGATCTAACTTGGGGTGGCTTTGTCTTCTTCTTTTGCCAATTCCACTAATTGTTTGGG** | Del_k5_2-6 | -3.7486972137491272 | 0.1907356218745442 | 1.1562266569435549 |
| **GGCAATCCAGATCTAACTTGGGGTGGCTTTGTCTTCTTCTTTTGCCAATTCCACTAATTGTTTGGG** | Ins_k3_1_GCA | -0.10991983963731433 | 0.042050165835265725 | 43.988982074490934 |
| **GGCATCCAGATCTAACTTGGGGTGGCTTTGTCTTCTTCTTTTGCCAATTCCACTAATTGTTTGGG** | Ins_k2_1_GC | 0.13824134295289203 | 0.04066781812579837 | 56.37920460037077 |
| **GGCCAATTCCACTAATTGTTTGGG** | Del_k39_2-40 | -3.816536000269792 | 0.06985256867169176 | 1.0803910236358358 |
| **GGCCATCCAGATCTAACTTGGGGTGGCTTTGTCTTCTTCTTTTGCCAATTCCACTAATTGTTTGGG** | Ins_k3_1_GCC | -3.4990716722426085 | 0.22695701302219534 | 1.4840685888190142 |
| **GGCGATCCAGATCTAACTTGGGGTGGCTTTGTCTTCTTCTTTTGCCAATTCCACTAATTGTTTGGG** | Ins_k3_1_GCG | 0.5933112859056939 | 0.03290630047234157 | 88.86961723380492 |
| **GGCTATCCAGATCTAACTTGGGGTGGCTTTGTCTTCTTCTTTTGCCAATTCCACTAATTGTTTGGG** | Ins_k3_1_GCT | -0.5572658538774217 | 0.05363383117925944 | 28.12315282186857 |
| **GGCTTTGTCTTCTTCTTTTGCCAATTCCACTAATTGTTTGGG** | Del_k21_1-21;Del_k21_2-22 | -2.2957209673877474 | 0.0809602826530866 | 4.9438187921611805 |
| **GGG** | Del_k60_1-60;Del_k60_2-61 | -6.682736529907313 | 0.24650961436048957 | 0.06149019823854058 |
| **GGGAATCCAGATCTAACTTGGGGTGGCTTTGTCTTCTTCTTTTGCCAATTCCACTAATTGTTTGGG** | Ins_k3_1_GGA | -0.36664101714274344 | 0.04587331666644565 | 34.02916726699042 |
| **GGGATCCAGATCTAACTTGGGGTGGCTTTGTCTTCTTCTTTTGCCAATTCCACTAATTGTTTGGG** | Ins_k2_1_GG | -0.1275634650632339 | 0.04560794294181958 | 43.21966369172888 |
| **GGGCATCCAGATCTAACTTGGGGTGGCTTTGTCTTCTTCTTTTGCCAATTCCACTAATTGTTTGGG** | Ins_k3_1_GGC | 0.08115529996288973 | 0.0379414935356962 | 53.25088023903169 |
| **GGGCTTTGTCTTCTTCTTTTGCCAATTCCACTAATTGTTTGGG** | Del_k20_2-21 | -2.5805867683747445 | 0.07818124417472214 | 3.718321164026659 |
| **GGGG** | Del_k59_2-60 | -6.885298808045961 | 0.8511489878427904 | 0.05021508631420324 |
| **GGGGATCCAGATCTAACTTGGGGTGGCTTTGTCTTCTTCTTTTGCCAATTCCACTAATTGTTTGGG** | Ins_k3_1_GGG | -0.5034934248516229 | 0.04810523291253088 | 29.67680042063606 |
| **GGGGGTGGCTTTGTCTTCTTCTTTTGCCAATTCCACTAATTGTTTGGG** | Del_k15_2-16 | -5.997682505660476 | 2.0687664561367316 | 0.12198911361918234 |
| **GGGGTGGCTTTGTCTTCTTCTTTTGCCAATTCCACTAATTGTTTGGG** | Del_k16_1-16;Del_k16_2-17 | -3.7827921291037403 | 0.18502985141778644 | 1.1174696694402226 |
| **GGGTATCCAGATCTAACTTGGGGTGGCTTTGTCTTCTTCTTTTGCCAATTCCACTAATTGTTTGGG** | Ins_k3_1_GGT | -0.1732824287774723 | 0.04360165417366922 | 41.288194267787524 |
| **GGGTGGCTTTGTCTTCTTCTTTTGCCAATTCCACTAATTGTTTGGG** | Del_k17_1-17;Del_k17_2-18 | -2.7530156442418554 | 0.09743325504205672 | 3.1294065644641442 |
| **GGTAATCCAGATCTAACTTGGGGTGGCTTTGTCTTCTTCTTTTGCCAATTCCACTAATTGTTTGGG** | Ins_k3_1_GTA | -0.6051727252735096 | 0.05513699964618873 | 26.807623593979702 |
| **GGTATCCAGATCTAACTTGGGGTGGCTTTGTCTTCTTCTTTTGCCAATTCCACTAATTGTTTGGG** | Ins_k2_1_GT | -0.07625785931132656 | 0.048285662729948886 | 45.494942934769874 |
| **GGTCATCCAGATCTAACTTGGGGTGGCTTTGTCTTCTTCTTTTGCCAATTCCACTAATTGTTTGGG** | Ins_k3_1_GTC | 0.091010774911508 | 0.03871211770244505 | 53.778287611358934 |
| **GGTCCAGATCTAACTTGGGGTGGCTTTGTCTTCTTCTTTTGCCAATTCCACTAATTGTTTGGG** | A2G | -0.05902746481401755 | 0.029484163119207273 | 46.28563112190599 |
| **GGTCTTCTTCTTTTGCCAATTCCACTAATTGTTTGGG** | Del_k26_2-27 | -1.7910220398338643 | 0.04298994841694791 | 8.189370189479808 |
| **GGTGATCCAGATCTAACTTGGGGTGGCTTTGTCTTCTTCTTTTGCCAATTCCACTAATTGTTTGGG** | Ins_k3_1_GTG | 0.2939313797315235 | 0.03322539484243831 | 65.87706901278361 |
| **GGTGGCTTTGTCTTCTTCTTTTGCCAATTCCACTAATTGTTTGGG** | Del_k18_1-18;Del_k18_2-19 | -1.8321110243421983 | 0.06553628898110415 | 7.859696641429138 |
| **GGTTATCCAGATCTAACTTGGGGTGGCTTTGTCTTCTTCTTTTGCCAATTCCACTAATTGTTTGGG** | Ins_k3_1_GTT | -0.6730006842416324 | 0.04775107299350192 | 25.049612304850797 |
| **GGTTTGGG** | Del_k55_2-56 | -6.178259853460001 | 0.324429730882522 | 0.1018350614982658 |
| **GTAAATCCAGATCTAACTTGGGGTGGCTTTGTCTTCTTCTTTTGCCAATTCCACTAATTGTTTGGG** | Ins_k3_1_TAA | -0.17998969223836103 | 0.04396685543741853 | 41.01219012199476 |
| **GTAACTTGGGGTGGCTTTGTCTTCTTCTTTTGCCAATTCCACTAATTGTTTGGG** | Del_k9_2-10 | -1.0600497972763707 | 0.05168323325479931 | 17.01013320781042 |
| **GTAATCCAGATCTAACTTGGGGTGGCTTTGTCTTCTTCTTTTGCCAATTCCACTAATTGTTTGGG** | Ins_k2_1_TA | -0.15563512246066968 | 0.046701947139175005 | 42.02328680364255 |
| **GTAATTGTTTGGG** | Del_k50_2-51 | -4.107040687503462 | 0.07908915126191218 | 0.808009346802569 |
| **GTACATCCAGATCTAACTTGGGGTGGCTTTGTCTTCTTCTTTTGCCAATTCCACTAATTGTTTGGG** | Ins_k3_1_TAC | 0.30888454730687853 | 0.035732789315258695 | 66.86954167736351 |
| **GTAGATCCAGATCTAACTTGGGGTGGCTTTGTCTTCTTCTTTTGCCAATTCCACTAATTGTTTGGG** | Ins_k3_1_TAG | 0.3354827179087563 | 0.0331787459592353 | 68.67201412590038 |
| **GTATATCCAGATCTAACTTGGGGTGGCTTTGTCTTCTTCTTTTGCCAATTCCACTAATTGTTTGGG** | Ins_k3_1_TAT | 0.19915600073146134 | 0.03804970798124052 | 59.920281404333274 |
| **GTATCCAGATCTAACTTGGGGTGGCTTTGTCTTCTTCTTTTGCCAATTCCACTAATTGTTTGGG** | Ins_k1_1_T | 0.5617514719753902 | 0.0364987378573669 | 86.10870474895907 |
| **GTATGATCCAGATCTAACTTGGGGTGGCTTTGTCTTCTTCTTTTGCCAATTCCACTAATTGTTTGGG** | Ins_k4_1_TATG | 0.11971987638764059 | 0.042510009013534356 | 55.34458991446929 |
| **GTCAATCCAGATCTAACTTGGGGTGGCTTTGTCTTCTTCTTTTGCCAATTCCACTAATTGTTTGGG** | Ins_k3_1_TCA | 0.6159530655308894 | 0.03625913234475164 | 90.90473593591499 |
| **GTCATCCAGATCTAACTTGGGGTGGCTTTGTCTTCTTCTTTTGCCAATTCCACTAATTGTTTGGG** | Ins_k2_1_TC | 0.4893864918663262 | 0.03539465679280098 | 80.09757081149394 |
| **GTCCACTAATTGTTTGGG** | Del_k45_2-46 | -4.088678190851729 | 0.09081923816101288 | 0.8229834761807694 |
| **GTCCAGATCTAACTTGGGGTGGCTTTGTCTTCTTCTTTTGCCAATTCCACTAATTGTTTGGG** | Del_k1_2-2 | 0.28685260213543196 | 0.03745384431867485 | 65.41238652495167 |
| **GTCCATCCAGATCTAACTTGGGGTGGCTTTGTCTTCTTCTTTTGCCAATTCCACTAATTGTTTGGG** | Ins_k3_1_TCC | 0.6300131686894694 | 0.03152631312178511 | 92.19189348030243 |
| **GTCGATCCAGATCTAACTTGGGGTGGCTTTGTCTTCTTCTTTTGCCAATTCCACTAATTGTTTGGG** | Ins_k3_1_TCG | 0.8544396437724542 | 0.034502030434770285 | 100 |
| **GTCTAACTTGGGGTGGCTTTGTCTTCTTCTTTTGCCAATTCCACTAATTGTTTGGG** | Del_k7_2-8 | -0.839955503044318 | 0.06061412368921286 | 21.19792992273072 |
| **GTCTATCCAGATCTAACTTGGGGTGGCTTTGTCTTCTTCTTTTGCCAATTCCACTAATTGTTTGGG** | Ins_k3_1_TCT | 0.38699570117060966 | 0.03488296569293031 | 72.30221290596505 |
| **GTCTTCTTCTTTTGCCAATTCCACTAATTGTTTGGG** | Del_k27_1-27;Del_k27_2-28 | -1.0452018907825458 | 0.036638676593572436 | 17.264582419533177 |
| **GTCTTCTTTTGCCAATTCCACTAATTGTTTGGG** | Del_k30_2-31 | -1.8615946803238645 | 0.040824744371346595 | 7.6313468839104575 |
| **GTCTTTTGCCAATTCCACTAATTGTTTGGG** | Del_k33_2-34 | -1.2603853950701651 | 0.03390937826282967 | 13.92204617912554 |
| **GTGAATCCAGATCTAACTTGGGGTGGCTTTGTCTTCTTCTTTTGCCAATTCCACTAATTGTTTGGG** | Ins_k3_1_TGA | 0.7198810717177708 | 0.03463195096235024 | 100 |
| **GTGATCCAGATCTAACTTGGGGTGGCTTTGTCTTCTTCTTTTGCCAATTCCACTAATTGTTTGGG** | Ins_k2_1_TG | 0.6052722971943314 | 0.03755436456477006 | 89.9389702515579 |

| | | | | |
|---|---|---|---|---|
| **GTGCATCCAGATCTAACTTGGGGTGGCTTTGTCTTCTTCTTTTGCCAATTCCACTAATTGTTTGGG** | Ins_k3_1_TGC | 0.7101444956265675 | 0.03427606749876689 | 99.88340247169742 |
| **GTGCCAATTCCACTAATTGTTTGGG** | Del_k38_2-39 | -2.107645559753663 | 0.03201217622290066 | 5.966816138719194 |
| **GTGGATCCAGATCTAACTTGGGGTGGCTTTGTCTTCTTCTTTTGCCAATTCCACTAATTGTTTGGG** | Ins_k3_1_TGG | 0.7486001891701293 | 0.034603211412844524 | 100 |
| **GTGGCTTTGTCTTCTTCTTTTGCCAATTCCACTAATTGTTTGGG** | Del_k19_1-19;Del_k19_2-20 | -0.16159304786989298 | 0.028863800914913203 | 41.77365956406795 |
| **GTGGG** | Del_k58_2-59 | -6.491538851957533 | 0.27633155146718835 | 0.07444610656745618 |
| **GTGGGATCCAGATCTAACTTGGGGTGGCTTTGTCTTCTTCTTTTGCCAATTCCACTAATTGTTTGGG** | Ins_k4_1_TGGG | 0.22794490645197252 | 0.049575274696062994 | 61.67039174444289 |
| **GTGGGGTGGCTTTGTCTTCTTCTTTTGCCAATTCCACTAATTGTTTGGG** | Del_k14_2-15 | -1.2623431883458565 | 0.05145179932483288 | 13.894816354613402 |
| **GTGTATCCAGATCTAACTTGGGGTGGCTTTGTCTTCTTCTTTTGCCAATTCCACTAATTGTTTGGG** | Ins_k3_1_TGT | 0.6762244822272883 | 0.031731735534205815 | 96.55217318665241 |
| **GTGTCTTCTTCTTTTGCCAATTCCACTAATTGTTTGGG** | Del_k25_2-26 | -0.2818890700382612 | 0.030186195187824033 | 37.038946411600804 |
| **GTGTTTGGG** | Del_k54_2-55 | -6.222180377107175 | 0.25917106215776425 | 0.09745921050508488 |
| **GTTAATCCAGATCTAACTTGGGGTGGCTTTGTCTTCTTCTTTTGCCAATTCCACTAATTGTTTGGG** | Ins_k3_1_TTA | 0.290001549242769 | 0.03859957261119884 | 65.61869132125497 |
| **GTTATCCAGATCTAACTTGGGGTGGCTTTGTCTTCTTCTTTTGCCAATTCCACTAATTGTTTGGG** | Ins_k2_1_TT | 0.3848290626951606 | 0.040584333285583016 | 72.14572973207073 |
| **GTTCATCCAGATCTAACTTGGGGTGGCTTTGTCTTCTTCTTTTGCCAATTCCACTAATTGTTTGGG** | Ins_k3_1_TTC | 0.7642929127840432 | 0.03268982256694197 | 100 |
| **GTTCCACTAATTGTTTGGG** | Del_k44_2-45 | -4.7195428647044535 | 0.11927392011938096 | 0.43793541671323466 |
| **GTTCCAGATCTAACTTGGGGTGGCTTTGTCTTCTTCTTTTGCCAATTCCACTAATTGTTTGGG** | A2T | 0.4083315624058559 | 0.031131904305678863 | 73.86141722316879 |
| **GTTCTTCTTTTGCCAATTCCACTAATTGTTTGGG** | Del_k29_2-30 | -0.9915246906505978 | 0.03386441796352934 | 18.2166196348543 |
| **GTTCTTTTGCCAATTCCACTAATTGTTTGGG** | Del_k32_2-33 | -2.671981593275233 | 0.062192491775822936 | 3.393552953820915 |
| **GTTGATCCAGATCTAACTTGGGGTGGCTTTGTCTTCTTCTTTTGCCAATTCCACTAATTGTTTGGG** | Ins_k3_1_TTG | 0.6810269812847534 | 0.0333149724980522 | 97.01698013147623 |
| **GTTGCCAATTCCACTAATTGTTTGGG** | Del_k37_2-38 | -2.4273375741512555 | 0.043552319331186896 | 4.334132438380401 |
| **GTTGGG** | Del_k57_2-58 | -5.919775013882713 | 0.7006945770785349 | 0.13187299494416704 |
| **GTTGGGGTGGCTTTGTCTTCTTCTTTTGCCAATTCCACTAATTGTTTGGG** | Del_k13_2-14 | -4.238946833302373 | 0.341428666443034 | 0.7081581755113573 |
| **GTTGTCTTCTTCTTTTGCCAATTCCACTAATTGTTTGGG** | Del_k24_2-25 | -1.0568707784297855 | 0.042930556033252604 | 17.064294786587027 |
| **GTTGTTTGGG** | Del_k53_2-54 | -5.384599652126311 | 0.19504733124824433 | 0.22520580244075122 |
| **GTTTATCCAGATCTAACTTGGGGTGGCTTTGTCTTCTTCTTTTGCCAATTCCACTAATTGTTTGGG** | Ins_k3_1_TTT | 0.5429170156888599 | 0.03641751641642003 | 84.50207163805862 |
| **GTTTGCCAATTCCACTAATTGTTTGGG** | Del_k36_2-37 | -1.0286138559684783 | 0.031668628361204064 | 17.55335638845718 |
| **GTTTGGG** | Del_k56_1-56;Del_k56_2-57 | -5.153680797477356 | 0.18123723141142728 | 0.283704588740051 |
| **GTTTGTCTTCTTCTTTTGCCAATTCCACTAATTGTTTGGG** | Del_k23_2-24 | -0.7509525448663608 | 0.0439431087329715 | 23.171115696880364 |
| **GTTTTGCCAATTCCACTAATTGTTTGGG** | Del_k35_2-36 | -1.5879178591537575 | 0.036985936172345815 | 10.03361720603436 |
| **TAACTTGGGGTGGCTTTGTCTTCTTCTTTTGCCAATTCCACTAATTGTTTGGG** | Del_k10_1-10 | -4.577567844788717 | 0.6451583268458184 | 0.5047415262276582 |
| **TAATTGTTTGGG** | Del_k51_1-51 | -5.508546755803748 | 0.18969599518480112 | 0.19895278951857998 |
| **TATCCAGATCTAACTTGGGGTGGCTTTGTCTTCTTCTTTTGCCAATTCCACTAATTGTTTGGG** | G1T | -3.6909288730960137 | 0.11999743792957383 | 1.2249869142619123 |
| **TCCACTAATTGTTTGGG** | Del_k46_1-46 | -6.730125120559903 | 0.36894864325131477 | 0.058644230186754326 |
| **TCCAGATCTAACTTGGGGTGGCTTTGTCTTCTTCTTTTGCCAATTCCACTAATTGTTTGGG** | Del_k2_1-2 | 0.746554291393688 | 0.02902936099364325 | 100 |
| **TCTAACTTGGGGTGGCTTTGTCTTCTTCTTTTGCCAATTCCACTAATTGTTTGGG** | Del_k8_1-8 | -4.748878740640092 | 0.6662296232399241 | 0.4252748106024913 |
| **TCTTCTTCTTTTGCCAATTCCACTAATTGTTTGGG** | Del_k28_1-28 | -4.933817000835815 | 0.18756246985031283 | 0.3534695393726462 |
| **TCTTCTTTTGCCAATTCCACTAATTGTTTGGG** | Del_k31_1-31 | -6.982275013478602 | 0.702937442435571 | 0.04557408758575126 |
| **TCTTTTGCCAATTCCACTAATTGTTTGGG** | Del_k34_1-34 | -5.449069607774973 | 0.2338649407686553 | 0.21114491654632922 |
| **TGCCAATTCCACTAATTGTTTGGG** | Del_k39_1-39 | -3.0443291657027376 | 0.04623852712040404 | 2.3385471701579648 |
| **TGGCTTTGTCTTCTTCTTTTGCCAATTCCACTAATTGTTTGGG** | Del_k20_1-20 | -4.613227749271928 | 0.2113924034916062 | 0.4870596326261592 |
| **TGGG** | Del_k59_1-59 | -7.567750065288243 | 1.0354995907259954 | 0.025377532833885514 |
| **TGTCTTCTTCTTTTGCCAATTCCACTAATTGTTTGGG** | Del_k26_1-26 | -6.184047525766668 | 0.50670474127268506 | 0.10124737583947277 |
| **TGTTTGGG** | Del_k55_1-55 | -6.418884475958091 | 0.26092266615683 | 0.08005627603647521 |
| **TTCCACTAATTGTTTGGG** | Del_k45_1-45 | -4.804620644625075 | 0.10675334447512339 | 0.4022177743423333 |
| **TTCTTCTTTTGCCAATTCCACTAATTGTTTGGG** | Del_k30_1-30 | -5.535893650588324 | 0.37530644415498005 | 0.1935857686680014 |
| **TTCTTTTGCCAATTCCACTAATTGTTTGGG** | Del_k33_1-33 | -3.937775694159202 | 0.10739170778558974 | 0.9570337139529631 |
| **TTGCCAATTCCACTAATTGTTTGGG** | Del_k38_1-38 | -5.461904015020711 | 0.23919803201947726 | 0.20845231264782424 |
| **TTGGG** | Del_k58_1-58 | -6.734828119025026 | 0.8476255252354122 | 0.0583690739989252 |
| **TTGGGGTGGCTTTGTCTTCTTCTTTTGCCAATTCCACTAATTGTTTGGG** | Del_k14_1-14 | -5.318047148235889 | 0.6914996389939249 | 0.24070380808650085 |
| **TTGTCTTCTTCTTTTGCCAATTCCACTAATTGTTTGGG** | Del_k25_1-25 | -5.526885372604712 | 0.5537182998915945 | 0.19533752137744043 |
| **TTGTTTGGG** | Del_k54_1-54 | -6.243962250283498 | 0.33885637724043666 | 0.0953593191493932 |
| **TTTGCCAATTCCACTAATTGTTTGGG** | Del_k37_1-37 | -5.229080046284577 | 0.1595048465579299 | 0.26310002090376633 |
| **TTTGGG** | Del_k57_1-57 | -4.6158150522815244 | 0.11460383381779037 | 0.485801090589683 |
| **TTTGTCTTCTTCTTTTGCCAATTCCACTAATTGTTTGGG** | Del_k24_1-24 | -4.902753194107814 | 0.6051655818477933 | 0.3646219705210007 |
| **TTTTGCCAATTCCACTAATTGTTTGGG** | Del_k36_1-36 | -4.646410390436549 | 0.15699544647177988 | 0.4711629137828212 |
