## Supplementary material for "Deep indel mutagenesis reveals the regulatory and modulatory architecture of alternative exon splicing": Baeza Minana et al Supplementary Tables 2_4

### Baeza, Miñana et al Supplementary Table 2

##### Indel library amplification primers

FAS_i5_GC_F 5’-tgtccaatgttccaacctacag-3’

FAS_i6_GC_R 5’-ctacttcccaagttatttcaatctg-3’

##### pCMV FAS 567 Vector Backbone amplification primers

PT1 5’-GTCGACGACACTTGCTCAAC-3’

PT2 5’-AAGCTTGCATCGAATCAGTAG-3’

##### Primers used for amplifying technical replicates (Input library)

| Forward primer | Barcode | Forward primer (FAS_i5_TR_F) | Reverse primer | Barcode | Reverse Primer (PT2) | Amplicon Size (nt) |
|---|---|---|---|---|---|---|
| **FAS_TR1** | GCCGAATT | **aaaatgtccaatgttccaacc** | PT2_TR1 | GCCGAATT | aagcttgcatcgaatcagtag | 231+16 |
| **FAS_TR2** | CGGCAATT | **aaaatgtccaatgttccaacc** | PT2_TR2 | CGGCAATT | aagcttgcatcgaatcagtag | 231+16 |
| **FAS_TR3** | GAACGTTC | **aaaatgtccaatgttccaacc** | PT2_TR3 | GAACGTTC | aagcttgcatcgaatcagtag | 231+16 |

##### Primers used for amplifying biological replicates (Output library)

| Forward primer | Barcode | sequence | Reverse primer | Barcode | Reverse Primer (PT2) | Inclusion amplicon Size (nt) |
|---|---|---|---|---|---|---|
| **FAS_BR1** | CAACCATG | CAGCAACACCAAGTGCAAAG | PT2_BR1 | CAACCATG | aagcttgcatcgaatcagtag | 195 +16 |
| **FAS_BR2** | CGTACCTT | CAGCAACACCAAGTGCAAAG | PT2_BR2 | CGTACCTT | aagcttgcatcgaatcagtag | 195 +16 |
| **FAS_BR3** | ACACTGTG | CAGCAACACCAAGTGCAAAG | PT2_BR3 | ACACTGTG | aagcttgcatcgaatcagtag | 195 +16 |
| **FAS_BR4** | GACACACT | CAGCAACACCAAGTGCAAAG | PT2_BR4 | GACACACT | aagcttgcatcgaatcagtag | 195 +16 |
| **FAS_BR5** | ACGACTTG | CAGCAACACCAAGTGCAAAG | PT2_BR5 | ACGACTTG | aagcttgcatcgaatcagtag | 195 +16 |
| **FAS_BR6** | ATCGATCG | CAGCAACACCAAGTGCAAAG | PT2_BR6 | ATCGATCG | aagcttgcatcgaatcagtag | 195 +16 |
| **FAS_BR7** | AACCAACC | CAGCAACACCAAGTGCAAAG | PT2_BR7 | AACCAACC | aagcttgcatcgaatcagtag | 195 +16 |
| **FAS_BR8** | CCATCGTT | CAGCAACACCAAGTGCAAAG | PT2_BR8 | CCATCGTT | aagcttgcatcgaatcagtag | 195 +16 |
| **FAS_BR9** | GCGCATAT | CAGCAACACCAAGTGCAAAG | PT2_BR9 | GCGCATAT | aagcttgcatcgaatcagtag | 195 +16 |

### Supplementary Table 3

##### Experimental validation of individual clones and PSI determination by Hek293 transfection

| SampleID | sequenceID | sequence | PSI | SD |
|---|---|---|---|---|
| **Fas wt** | | GATCCAGATCTAACTTGGGGTGGCTTTGTCTTCTTCTTTTGCCAATTCCACTAATTGTTTGGG | 49.9 | 1.3 |
| **Indel1** | EXK513 | ATCCAGATCTAACTTGGGGTGGCTTTGTCTTCTTCTTTTGCCAATTCCACTAATTGTTTGGG | 53.3 | 0.5 |
| **Indel3** | EXK515 | GATCCAGATCTAACTTGGGGTGGCTTTGTCTTCTTCTTTTGCCAATTCCACTAATTGAGATTTGGG | 67.0 | 1.2 |
| **Indel5** | EXK517 | GATCCAGATCTAACTTGGGGTGGCTTTGTCTTCTTCTTTTGCCAATTCCCTGACTAATTGTTTGGG | 47.8 | 0.6 |
| **Indel6** | EXK518 | TTTGCCAATTCCACTAATTGTTTGGG | 1.2 | 1.4 |
| **Indel8** | EXK520 | GATCCAGATCTAACTTGGGGTGGCTGTTTGGG | 21.5 | 1.9 |
| **Indel9** | EXK521 | GATCCAGATCTAACTTGGGGTGGCTTTGTCTTCTTCTTTTGCCAATTCCAATTTCTAATTGTTTGGG | 5.1 | 0.8 |
| **Indel10** | EXK522 | GATCCAGATCTAACTTGGGGTGGCTTTGTCTTCTTCTTTTGCCAATTCCACTAATTGTTTGTACGG | 44.9 | 2.1 |
| **Indel13** | EXK525 | GATCCAGATCTAACTTGGGGTGGCTTTGTCTTCTTATCCTTTTGCCAATTCCACTAATTGTTTGGG | 52.3 | 0.3 |
| **Indel16** | EXK528 | GATCCAGATCTAACTTGGGGTGGCTTTGTCTTCTTCTTTTGCCAATTCCACTTGTTTGGG | 62.6 | 0.2 |
| **Indel18** | EXK530 | GATCCAGATCTAACTTGGGGTGGCCACTAATTGTTTGGG | 38.0 | 1.3 |
| **Indel19** | EXK531 | GATCCAGATCTAACAATTCCACTAATTGTTTGGG | 3.7 | 0.5 |
| **Indel20** | EXK532 | GATCCAGATCTAACTTGGGGTGGCTTTGTCTTCTTCTCCTTTTGCCAATTCCACTAATTGTTTGGG | 64.3 | 0.7 |
| **Indel22** | EXK534 | GATCCAGTTTGCCAATTCCACTAATTGTTTGGG | 13.1 | 1.5 |
| **Indel23** | EXK535 | GATCCAGATCTAACGGATTGGGGTGGCTTTGTCTTCTTCTTTTGCCAATTCCACTAATTGTTTGGG | 79.3 | 2.6 |
| **Indel24** | EXK536 | GATCCAGATCTAACTTGGGGTGGCTTTGTCTTCTTCTTTTGCCAATTCCACTAATTGCGTTTTGGG | 63.1 | 0.9 |

### Supplementary Table 4

##### AON Walk

| RNA oligo 2' O-Me PS | Sequence DNA FAS exon 6 | Size | Position | Sequence RNA 2' O-MePS |
|---|---|---|---|---|
| **FAON_2** | GATCCAGATCTAACTTGGGGT | 21 | 0_21 | ACCCCAAGUUAGAUCUGGAUC |
| **FAON_3** | AGATCTAACTTGGGGTGGCTT | 21 | 6_26 | AAGCCACCCCAAGUUAGAUCU |
| **FAON_4** | TAACTTGGGGTGGCTTTGTCT | 21 | 11_31 | AGACAAAGCCACCCCAAGUUA |
| **FAON_5** | TGGGGTGGCTTTGTCTTCTTC | 21 | 16_36 | GAAGAAGACAAAGCCACCCCA |
| **FAON_6** | TGGCTTTGTCTTCTTCTTTTG | 21 | 21_41 | CAAAAGAAGAAGACAAAGCCA |
| **FAON_7** | TTGTCTTCTTCTTTTGCCAAT | 21 | 26_46 | AUUGGCAAAAGAAGAAGACAA |
| **FAON_8** | TTCTTCTTTTGCCAATTCCAC | 21 | 31_51 | GUGGAAUUGGCAAAAGAAGAA |
| **FAON_9** | CTTTTGCCAATTCCACTAATT | 21 | 36_56 | AAUUAGUGGAAUUGGCAAAAG |
| **FAON_10** | GCCAATTCCACTAATTGTTTG | 21 | 41_61 | CAAACAAUUAGUGGAAUUGGC |
